# Mouse, pig, and human atherosclerotic lesions have common and distinct mesenchymal cell populations

**DOI:** 10.1101/2025.07.24.666663

**Authors:** D Sharysh, P Nogales, D Morales-Cano, A Markov, R Izquierdo-Serrano, L Carramolino, J Albarrán-Juárez, JF Bentzon

**Author notes:** **Correspondence:** Diana Sharysh Department of Clinical Medicine, Aarhus University, Palle Juul-Jensens Boulevard 11, 8200 Aarhus N, Denmark. or Jacob Fog Bentzon Department of Clinical Medicine, Aarhus University, Palle Juul-Jensens Boulevard 11, 8200 Aarhus N, Denmark.

## Abstract

The proliferation and phenotypic modulation of smooth muscle cells (SMCs) to alternative mesenchymal states is a key process by which atherosclerotic lesions grow. The underlying mechanisms can be studied in mouse and pig atherosclerosis, but it remains unclear to what extent the mesenchymal plaque cell types in these species recapitulate human disease. Here, we integrate published and new single-cell RNA sequencing data of plaque mesenchymal cells from human carotid and coronary arteries, pig aorta and coronary arteries, and mouse brachiocephalic arteries. By applying consensus across multiple integration and gene homology-matching strategies, we identify a conserved core continuum of mesenchymal plaque cells, ranging from SMCs to extracellular matrix-producing fibroblast-like cells, which is stable across species and vascular beds. Notably, several other populations differed between human and experimental lesions. Subpopulations of SMCs marked by DLX5 and RERGL expression were specific to human carotid and coronary plaques, respectively. Mesenchymal cell states with strong pro-angiogenic and inflammation-associated gene signatures were identified in pig, but not human, coronary plaque datasets, with the pro-angiogenic phenotype associated with early stages of necrotic core development. Pericytes were solely present in pig and human plaques, while chondrocyte-like cells were unique to mouse lesions. The presented interspecies maps of mesenchymal cell diversity, and their markers may inform translational research into the role of SMCs and their derived progeny in atherosclerosis.

## Introduction

During atherosclerosis and other types of arterial disease, a subset of smooth muscle cells (SMCs) loses its differentiated phenotype, proliferates, and transitions to several mesenchymal cell types not typically present in the arterial intima^1,2^. Additional mesenchymal cells may enter the lesion as pericytes accompanying neovessel ingrowth or, under certain conditions, may originate from endothelial cells undergoing endothelial-to-mesenchymal transitioning^3,4^ or from invading adventitial fibroblasts^5^. SMC-derived mesenchymal cells contribute both to stabilizing components of plaques, such as the fibrous cap and extracellular matrix, and to destabilizing elements, including the necrotic core^6^. Their relevance to clinical outcomes and potential as therapeutic targets is underscored by their expression of multiple coronary artery disease-associated risk genes identified in genome-wide association studies^7,8^.

The abundance and diversity of plaque mesenchymal cells have been extensively studied in mouse models of atherosclerosis. By using SMC lineage tracing with fluorescence reporter transgenes, including stochastically recombining versions that allow tracking of individual cells, up to 70% of all murine plaque cells have been found to originate from a small number of medial SMCs that undergo clonal expansion^2,9,10^. In human carotid and coronary atherosclerosis, single-cell RNA-sequencing (scRNA-seq) has revealed a large and heterogeneous population of putative SMC-derived mesenchymal cells^11–14^, but the extent to which these human cell types correspond to those observed in mouse models has not been systematically analyzed. Moreover, mesenchymal cell populations remain uncharacterized in large-animal models, such as pigs, whose lesions show higher morphological resemblance to human lesions than those in mice^15^.

Potential differences between experimental and human atherosclerosis may arise not only from divergent regulatory mechanisms governing SMC responses in different species but also from variation in the arterial sites studied. While clinically relevant atherosclerosis in humans primarily affects the coronary and carotid arteries, murine studies typically focus on the aortic root or brachiocephalic artery^16^, whereas porcine studies often examine the abdominal aorta as well as the coronary arteries^15^. SMCs across these vascular regions differ in their embryonic origin, which could influence the repertoire of mesenchymal phenotypes in plaques, as embryonic programs are known to play critical roles in metaplastic processes in general^17^ and SMC fate decisions in particular^18,19^.

Resolving the similarities and differences between plaque mesenchymal cells in human and experimental atherosclerosis is essential for determining which human processes can be reliably studied in animal models. Moreover, comparing SMC responses across multiple species may provide a valuable vantage point for distinguishing core regulatory mechanisms of SMC plasticity from species- or model-specific phenomena. It may also help define a meaningful set of markers that can be used across species to describe plaque mesenchymal cells in translational studies.

Here, we performed consensus integration of plaque mesenchymal scRNA-seq data from mouse, pig, and human atherosclerotic lesions. We identified a conserved spectrum of mesenchymal cell states, ranging from contractile SMCs to non-contractile, extracellular matrix–producing phenotypes, consistently observed across all species and vascular sites examined. Importantly, several other mesenchymal cell states were specific to models or human lesions. Based on these findings, we propose a set of consensus markers applicable across species and provide a comparative map to support cross-species interpretation of mesenchymal cell phenotypes in experimental and human atherosclerosis.

## Results

### Plaque mesenchymal scRNA-seq datasets from humans, pigs, and mice

Atherosclerosis develops both in large elastic arteries, e.g. carotid arteries and aorta, and in medium-sized muscular arteries, e.g. coronary arteries. To minimize confounding from intrinsic differences in arterial wall structure, we stratified our analysis accordingly. To compare plaque mesenchymal cells in elastic arteries across species, we obtained published scRNA-seq data from 6 human carotid plaque endarterectomies and 15 brachiocephalic artery lesions from high-fat-fed Apoe^-/-^ mice^13,14,20^, and generated scRNA-seq data of 6 atherosclerotic lesions from the abdominal aorta of high-fat-fed PCSK9 transgenic minipigs, as described elsewhere^21^ (**Table S1**). The murine dataset is similar to several published mouse atherosclerosis datasets from *Apoe-/-*, *Ldlr^-/-^*, and PCSK9-overexpressing mice^22,23^. Murine and porcine lesions were harvested without the underlying arterial wall, yielding samples comparable to endarterectomy specimens.

The mesenchymal supercluster in human and pig lesions was identified based on expression of canonical SMC and fibroblast markers (**Fig. S1A-B**), whereas the mouse dataset included only lineage-traced SMC-derived cells. In all datasets, the mesenchymal supercluster comprised a main population spanning SMCs and cells with low or absent expression of contractile genes, forming a phenotypic continuum, as well as a variable number of smaller adjacent clusters (**Fig. 1**). To harmonize cluster granularity and naming across species, we applied the lowest clustering resolution that separated adjacent clusters from the main population and subdivided the latter into at least three clusters, labeled as contractile, intermediate, and terminal cells. These labels reflect spatial positioning in the UMAP plot and do not imply a linear transitioning trajectory - i.e., that cells necessarily pass from contractile to intermediate to terminal states. Explorative analysis showed that higher resolution did not produce subsets with distinct gene expression patterns, indicating that the chosen resolution was sufficient to distinguish relevant cell types. Full sets of cluster marker genes are provided in **Table S2-S4**.

**Figure 1.**
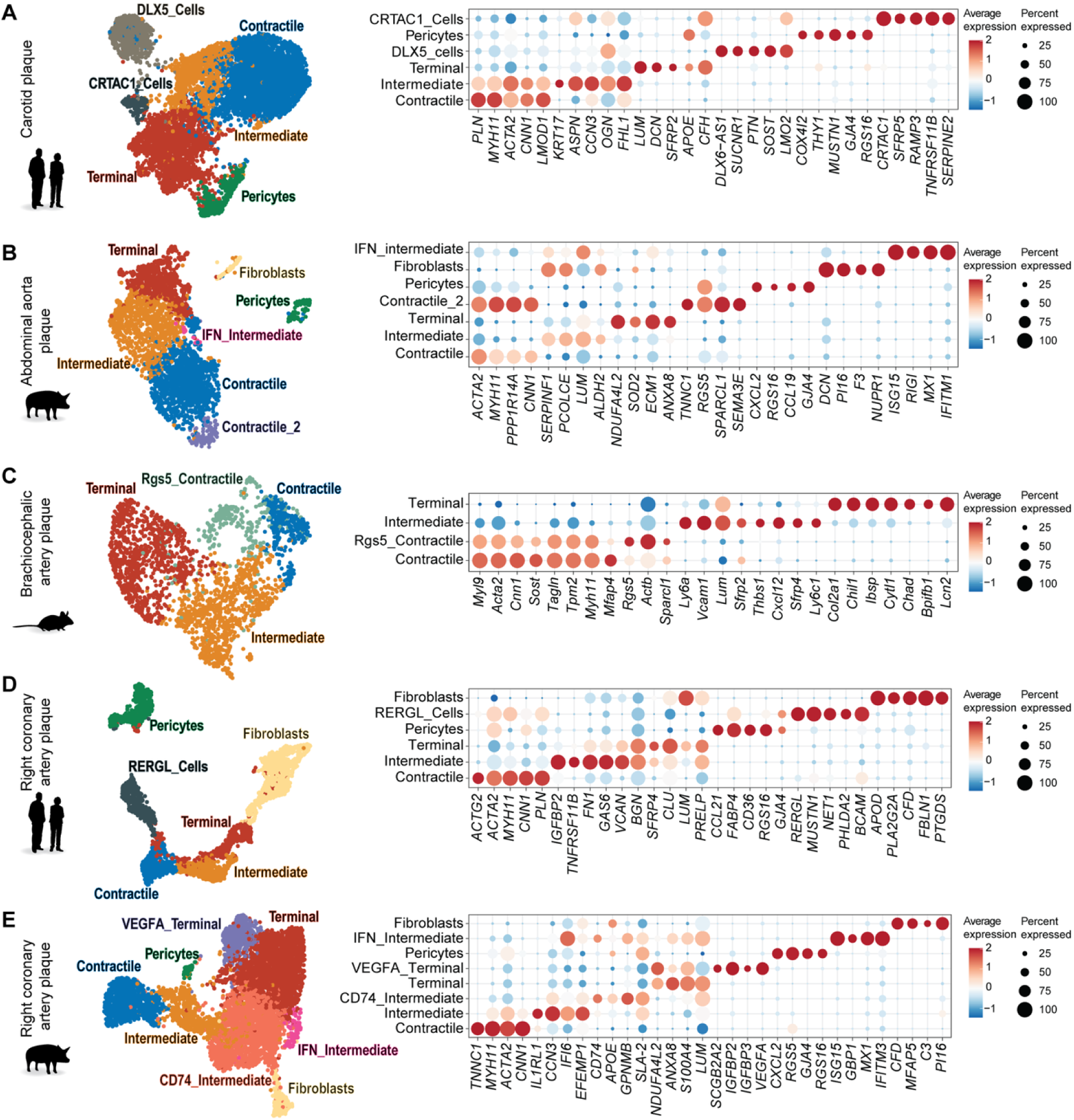
Individual analysis of plaque mesenchymal cells from human, pig, and mouse atherosclerosis. **A-E**, UMAP representations of the mesenchymal cell supercluster from human carotid plaques (from Pan et al.^13^ and Alsaigh et al.^14^, 6 patients, 8264 cells in total) (A), pig abdominal aorta plaques (from Nogales et al^21^, 6 pigs, 3153 cells) (B), mouse brachiocephalic artery plaques (Alencar et al.^20^, 15 mice, 2412 cells) (C), human right coronary artery plaques (Wirka et al.^12^, 4 patients 3609 cells) (D), and pig right coronary artery plaques (this paper, 6 pigs, 6222 cells) (E). The cells were clustered at the minimum resolution that separated the adjacent cell clusters, e.g., pericyte and fibroblasts, from the main continuous cell cluster and divided the main cluster into ≥3 cellular subclusters (contractile, intermediate, terminal). The identified cellular subtypes are labeled, and top markers, selected from the top 20 over-expressed genes in each cluster, are shown in the dot plots.

Following this strategy, the human mesenchymal supercluster in carotid plaques comprised 6 clusters (**Fig. 1A**), including the contractile, intermediate, and terminal clusters; a cluster of pericytes, identified by expression of canonical pericyte markers (e.g., *GJA4* and *RGS16*)^24,25^; and two additional SMC subclusters, DLX5_Cells and CRTAC1_Cells, with unknown functional roles, named after their top-expressed genes. The porcine mesenchymal supercluster in abdominal plaques comprised 7 clusters, including the contractile, intermediate, and terminal clusters, as well as a small secondary SMC cluster with high expression of *RGS5*, a pericyte cluster, a small fibroblast cluster, and a cluster with a type 1 interferon gene expression signature, named IFN_Intermediate (**Fig. 1B**). The murine mesenchymal supercluster in brachiocephalic plaques comprised 4 clusters, including the 3 core populations and a separate cluster of *Rgs5-*expressing SMCs (**Fig. 1C**).

To compare mesenchymal cells in coronary atherosclerosis across species, we utilized a published human scRNA-seq dataset consisting of 8 atherosclerotic right coronary artery segments from 4 heart transplant patients^12^, and generated scRNA-seq from raised atherosclerotic lesions in the right coronary arteries of 5 high-fat-fed PCSK9 transgenic minipigs. Murine coronary arteries are not routinely used for experimental atherosclerosis research, and no suitable datasets were available. Mesenchymal cell isolation (**Fig. S1C-D**), marker analysis (**Table S5-S6)**, and clustering were performed as for elastic artery atherosclerosis. The mesenchymal supercluster in human coronary plaques comprised 6 populations (**Fig. 1D**), including the contractile, intermediate, and terminal cells, as well as pericytes, fibroblasts, and a population of RERGL_cells with intermediate expression of contractile genes and high expression of *RERGL*. The mesenchymal supercluster in pig coronary lesions encompassed 8 clusters (**Fig. 1E**). In this dataset, separation of adjacent pericyte and fibroblast clusters could not be achieved without splitting the intermediate cluster into two subsets. The other identified clusters were VEGFA_Terminal cells, characterized by high *VEGFA* expression, and IFN_Intermediate cells with a type 1 interferon signature.

### Integration of plaque mesenchymal cells from mouse, pig, and human elastic arteries

The goal of scRNA-seq data integration is to generate a unified representation of cell types and states from datasets obtained under different conditions. This involves minimizing variability in gene expression caused by technical and experimental differences (batch effects) while preserving meaningful biological variation. In cross-species integration, it also requires separating trivial species-specific gene expression differences within comparable cell types from true differences in cell type and state. Numerous scRNA-seq data integration algorithms have been developed, and because their performance is highly context-dependent, it is advisable to evaluate multiple approaches on the specific datasets under study. Several metrics have been developed for this purpose to assess integration quality, including batch mixing efficiency and potential overcorrection, an undesirable effect that can mask genuine biological differences^26^.

Recently, Song et al. introduced the BENGAL pipeline, a three-step framework developed for cross-species integration^27^. In the first step, homologous genes are selected across datasets, using either one-to-one orthologs (O2O) or expanded sets that include one-to-many and many-to-many orthologs, selected based on average expression levels (HE) or homology confidence (HHC). In the second step, 9 integration algorithms - scVI^28^, scANVI^29^, scanorama^30^, harmony^31^, Seurat RPCA and CCA ^32^, LIGER^33^, LIGER UINMF^34^, fastMNN^35^ - are applied, generating a total of 27 integrated datasets. In the third step, the quality of integration is evaluated using batch-mixing and biology preservation metrics^26^.

To understand similarities and differences in elastic artery atherosclerosis across species, we applied the BENGAL pipeline to the scRNA-seq datasets from human carotid, pig abdominal aorta, and mouse brachiocephalic artery plaques. UMAP representations of the 27 integrated and 3 unintegrated homology-matched datasets are shown in **Fig. 2A**. RPCA, CCA, scVI, and scANVI integration algorithms demonstrated the best overall performance based on a composite score of batch mixing and biology conservation metrics (**Fig. 2B**). RPCA and CCA provided slightly better mixing, while scVI and scANVI achieved higher biology preservation metrics (**Fig. S1E)**.

**Figure 2.**
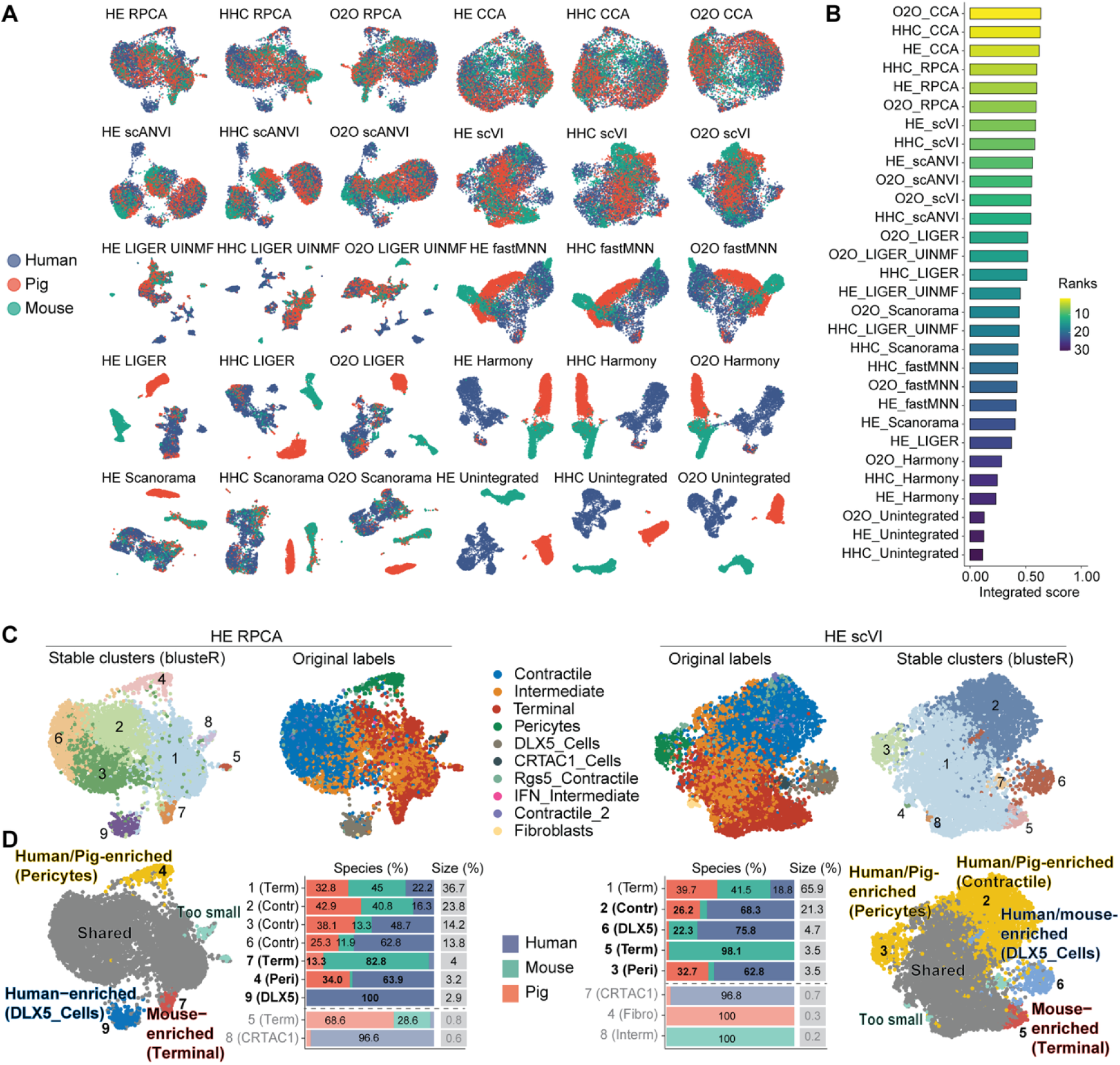
Identification of shared and species-specific mesenchymal plaque cell types in elastic mouse, pig, and human arteries. **A**, Combined datasets produced by 9 integration algorithms (or without integration) and 3 methods for gene homology matching between mouse, pig, and human datasets. **B,** Combined score for the integration performance combining 5 metrics for batch mixing and 5 metrics for biological preservation of differences between known cell types. CCA, RPCA, scANVI, and scVI outperform the others. **C,** Two examples of integrated data sets, HE RPCA and HE scVI, with stable clusters defined by the bluster R package and cell labels from the individual species datasets. **D**, Process for identifying species-enriched clusters. The contribution from the 3 species was analyzed in each stable cluster (using 1500 random cells from each data set) and defined as enriched in one species if that species contributed ≥80% of cells and in two species if those contributed ≥90% of cells. Names of clusters fulfilling these criteria are set in bold typeface. Clusters with less than 50 cells (1.1%) are labeled (light gray) but not further considered in the analysis. Cells not fulfilling the criteria to be species-enriched were assigned to the shared cluster. The enriched clusters are named after the dominant contributor from the individual species datasets.

The 12 highest-performing integrated datasets - generated using the 4 top-performing algorithms and the 3 gene homology strategies - were subsequently clustered using the bluster R package, which identifies stable clusters through permutation-based robustness testing (**Fig. S1F**)^36,37^. Cluster labels were assigned based on the dominant contributing cluster from the individual species datasets. The relative species contribution to stable clusters was then analyzed using random subsampling of 1500 cells per species, omitting small clusters with less than 50 sampled cells. A cluster was considered enriched for one species if ≥80% of its cells originated from that species, or for two species if those species together contributed ≥90% of cells. Enrichment was considered robust if reproduced in ≥6 of the 12 integrated datasets. Two representative examples showing shared and species-enriched clusters are presented in **Fig. 2C**, with additional results shown in **Fig. S2**, and a summary of findings below.

In all 12 integrated datasets, the majority of cells (median 77.1%, interquartile range (IQR) 65.0-89.8%) were assigned to shared clusters. The DLX5_Cell cluster was consistently enriched for human cells (median 95.9%, IQR 83.6-99.7%, in 8 integrations); minor mouse cell contributions were observed in scVI-based integrations, including the example in **Fig. 2C**, but this was not replicated by other algorithms. The pericyte cluster was highly enriched for human and pig cells (median 99.0%, IQR 94.5-100%, in 10 integrations). In contrast, a stable cluster at the tip of the terminal cells was dominated by mouse cells (median 92.3%, IQR 84.0-97.7%, in 6 integrations), while mouse cells were underrepresented in a cluster at the contractile end of the central axis (median 94.1%, IQR 94.9-100% human and pig cells, in 6 integrations).

### Transcriptional profiles of shared and species-specific plaque mesenchymal cells from elastic arteries

To analyze the transcriptome of the shared and species-enriched mesenchymal cell populations, we transferred cluster labels back to the individual mouse, pig, and human datasets, assigning cells to a given species-enriched cluster if they consistently belonged to that cluster in ≥4 integrated datasets. The remaining cells were defined as shared. They were re-clustered in the integrated dataset (using HE RPCA as a representative integration) into contractile, intermediate, and terminal cells before labels were transferred, which also resolved a small, shared population of proliferating cells (**Fig. 3A** and **Fig. S3A**).

**Figure 3.**
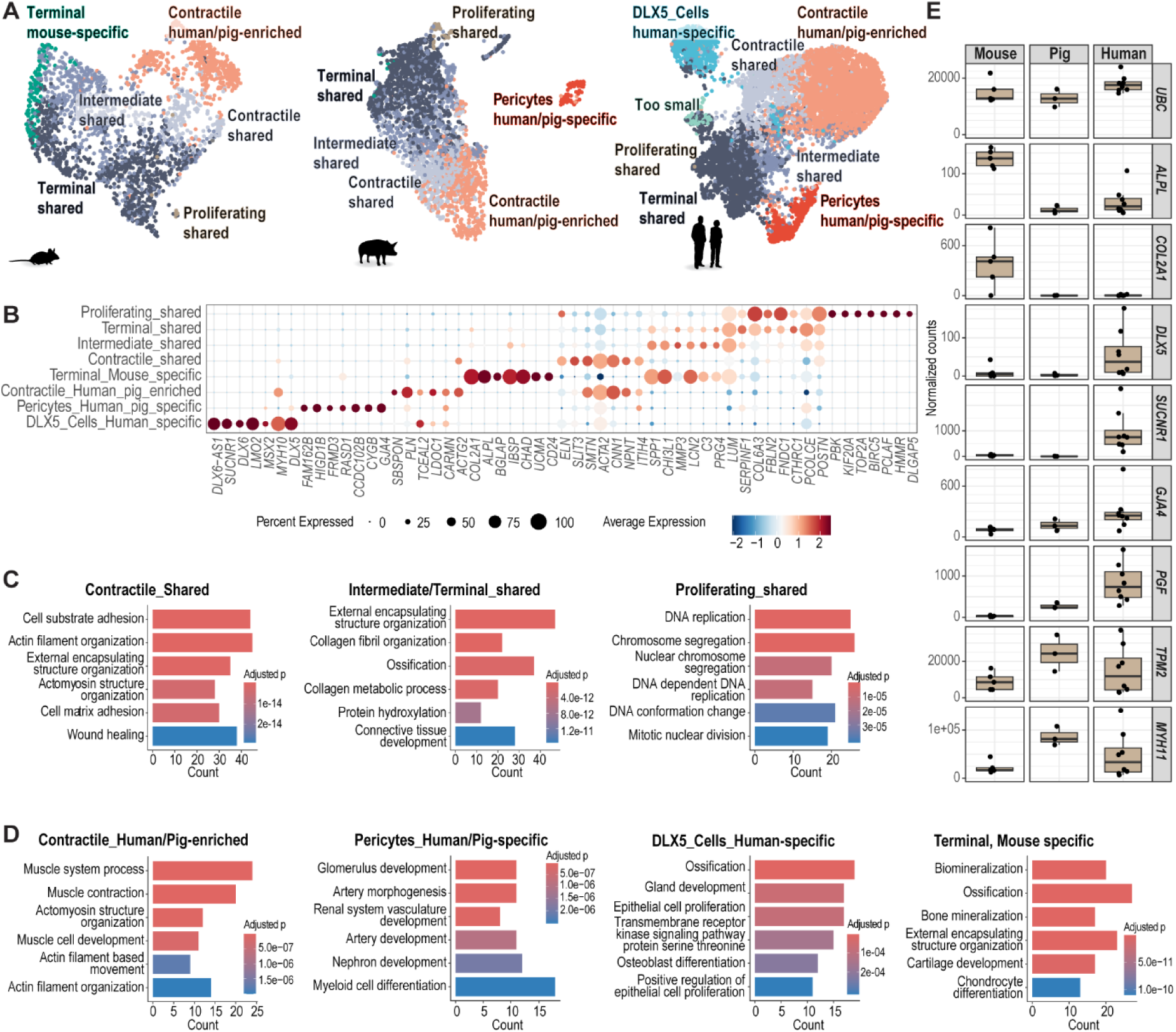
Gene expression analysis of shared and species-specific plaque mesenchymal cell types in elastic arteries. **A**, Shared and species-enriched/specific clusters in the mouse, pig, and human datasets based on consensus among integrations. **B,** Top markers for the shared and species-enriched/specific clusters. Markers were analyzed in merged data from all species. **C-D,** Enrichment analysis for shared and species-enriched/specific phenotypes. For the analysis of shared clusters, only genes expressed in ≥5% of cluster cells in all species were included. **E,** Validation of specific marker gene expression in mouse, pig, and human atherosclerosis using published RNA-seq data. Top markers for mouse-specific chondrocyte-like terminal cells (*Alpl*, *Col2a1*), human-specific DLX5_Cells (*DLX5*, *SUCNR1*), human/pig-specific pericytes (*GJA4*, *PGF*), and the human/pig-enriched contractile type of cells are shown. *UBC* is a housekeeping gene for comparison. DESeq2 normalized counts are shown.

The shared contractile cells were characterized by expression of canonical SMC markers (e.g., *MYH11*, *CNN1*, *ACTA2*, *ITGA8*), and enrichment of pathways (Gene Ontology) related to contractility (e.g., *Actin filament organization, Actomyosin structure*), but also *Cell adhesion*, and *Wound healing* (**Fig. 3B-C**). Notably, the shared intermediate cluster lacked strong marker genes that distinguished it from terminal cells, suggesting the absence of a distinct, conserved transitional cell state across species. Both intermediate and terminal cells exhibited high expression of collagen genes and *LUM*, which encodes the small leucine-rich proteoglycan extracellular matrix protein lumican, and enrichment for gene sets related to extracellular matrix organization.

Inspection of the species-enriched clusters revealed that the human/pig-enriched contractile cluster did not represent a unique cell population, but rather a group of contractile cells with elevated expression levels of contractile genes that were more abundant in humans and pigs than in mice (**Fig. 3A-B**). In contrast, DLX5_Cells, pericytes, and the terminal mouse-enriched cluster were truly species-*specific* populations, either absent or very rare in other species (**Fig. 3A**). Human-specific DLX5_Cells were marked by expression of the embryonic contractile gene *MYH10* and genes associated with neural crest cell fates (e.g. *DLX5*, *MSX2*, *PDGFRA*)^38,39^ and were enriched for gene sets related to bone development (e.g. *Ossification*, *Osteoblast differentiation*), among others (**Fig. 3B, D**). This cluster comprised, on average, 9% of mesenchymal cells in carotid plaques, but was present in only 3 out of 6 carotid plaque scRNA-seq datasets (data not shown). The human/pig-specific pericytes, comprising 6% and 3% of mesenchymal cells in human and pig plaques, respectively, expressed known pericyte markers (*GJA4*, *PGF*, *RGS16*), and their expressed genes were enriched in categories such as *Artery morphogenesis* (**Fig. 3B, D**) ^40,41^. Their species specificity likely reflects the absence - or minimal presence - of plaque neovascularization in mouse models of atherosclerosis^42^. Finally, the mouse-specific terminal cell cluster, representing 7% of mesenchymal cells in mouse plaques, exhibited a highly distinct chondrocyte-like transcriptional profile. It expressed genes such as *Col2a1, Bglap*, *Chad*, *Mia*, and *Alpl* (**Fig. 3B**), which encode proteins associated with cartilage extracellular matrix, and were enriched for gene categories such as *Cartilage development* and *Biomineralization* (**Fig. 3D**). Notably, although these chondrocyte-like mouse cells and the human *DLX5*-expressing cells are both enriched for osteochondrogenic gene expression signatures, they represent very distinct cell types.

### Validation of species-specific cell clusters in mouse, pig, and human plaque bulk RNA-seq data

To validate the observed differences in plaque mesenchymal cell types across species, we analyzed publicly available bulk RNA-seq data from atherosclerotic plaques: human carotid endarterectomies (GSE120521, N=8) ^43^, aortas from *Ldlr*^-/-^ mice fed a high-fat diet for 3 months (GSE205929, N=5)^44^, and aortas from minipigs fed a high-fat, high-fructose diet for 6 months (GSE245530 N=3). All datasets were processed uniformly to ensure comparability of RNA counts (**Fig. 3E**).

This analysis confirmed the species-specific patterns of marker gene expression observed in the scRNA-seq data: high expression of *DLX5* and *SUCNR1* (marking DLX5_SMCs) in human carotid plaques; high *Col2a1* and *Alpl* expression (marking the chondrocyte-like population) in murine aortic plaques; and elevated expression of *GJA4* and *PGF* (marking pericytes), along with *MYH11* and *TPM2* (marking contractile cells), in human and pig atherosclerotic lesions.

### Regulatory modules governing the shared and species-specific SMC populations

To identify regulatory networks underlying the gene expression signatures of shared and species-specific mesenchymal cell types, we applied the SCENIC pipeline to human and mouse datasets^46^. This technique, which is not yet available for porcine data, detects the activity of co-expressed genes that share a cis-regulatory motif recognized by the same transcription factor (regulon). Regulons were inferred separately for each dataset and then binarized - classifying each regulon as either active (on) or inactive (off) - to facilitate cross-species comparison^46^.

For the shared mesenchymal clusters, the aim was to identify regulons *active* in both species, reasoning that key regulatory mechanisms are likely to be conserved and that cross-species analysis may help distinguish those core programs from species-specific side phenomena or noise. For all regulons active in ≥30% of cells in at least one species, we calculated a relative activity index: (f_h_-f_m_)/(f_h_+f_m_), where f_h/m_ is the fraction of cells with the regulon active in human or mouse, respectively. This metric ranges from –1 (mouse-specific) over 0 (conserved) to +1 (human-specific) regulon activity (**Fig. 4A**).

**Figure 4.**
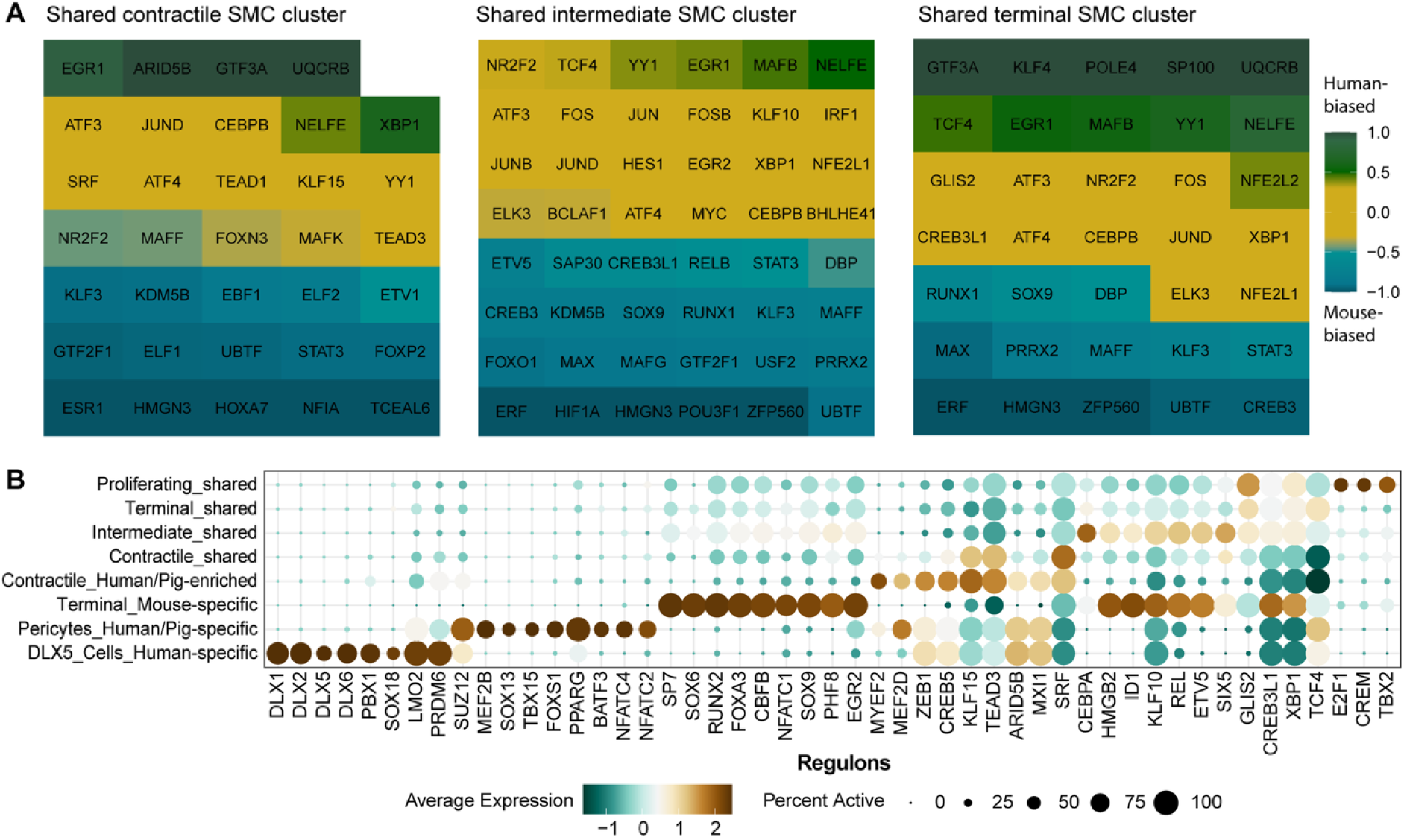
Activity of regulons across species in shared and specific plaque mesenchymal cell clusters. **A**, Plots showing the conservation of regulon activity in the shared contractile, intermediate, and terminal cell types in mouse and human lesions. Regulons detected in ≥30% of cells in the designated cluster in ≥1 species are shown with a color code indicating whether regulon activity is balanced between species (yellow), or higher in human (green) or mouse cells (blue). The color code is based on (f_h_-f_m_)/(f_h_+f_m_), where f_h/m_ is the fraction of cells with the regulon active in human or mouse, respectively. **B**, Dot plot showing differentially active regulons across shared and specific clusters.

In the shared contractile cluster, regulons with conserved high activity across species (relative activity index ≈ 0) included those important for SMC differentiation (TEAD1/3, SRF, and KLF15)^47,48^, as well as regulons associated with endoplasmic reticulum (ER) stress and inflammation (ATF3/4, YY1, JUND, CEBPB, NELFE, MAFK)^49^. In the shared intermediate cluster, regulons with conserved high activity included a broader set linked to ER stress (e.g., ATF3/4, XBP1, NFE2L1), AP1 (i.e., JUN/FOS)- and interferon-driven inflammatory signaling (JUN/B/D, FOS/B, CEBPB, IRF1),^50^ cell cycle control (MYC, EGR2), and cellular plasticity (HES1, YY1, KLF10) ^51^. Murine intermediate-type cells exhibited more activity of regulons involved in cartilage development and hypoxia (SOX9, RUNX1, HIF1A)^52,53^, while ER stress–related regulons were more active in humans. In cells of the shared terminal phenotype, ER stress and inflammatory regulons remained active in both species. Regulons related to cartilage development (SOX9, RUNX1) and cartilage-related epigenetic reprogramming (HMGN3, UBTF, ERF, MAX) were again more active in mice, while those related to cellular plasticity (KLF4, TCF4, YY1) were more active in humans.

To explore mechanisms underlying species-specific phenotypes, we identified the most differentially active regulons across clusters (**Fig. 4B**). DLX5_Cells, pericytes, and the mouse-specific terminal cells each showed distinct regulon activity. DLX5_Cells expressed regulons involved in neural crest development (DLX family regulons and PBX1) and vascular development (SOX18, LMO2, PRDM6).^54,55^ The human/pig-specific pericytes were characterized by regulons associated with mesenchymal and SMC differentiation and plasticity (TBX15, MEF2B, SUZ12)^56–58^, mechanotransduction (SOX13)^59^, and inflammation/lipid metabolism (BATF3, NFATC2/4, and PPARG)^50^. In contrast, the mouse-specific terminal cells displayed activity of regulons important for cartilage development (SOX6/9, SP7, RUNX2)^60^.

Together, these findings suggest a conserved regulatory network in modulated SMCs, characterized by ER stress and inflammatory signaling, particularly AP1-driven transcription. This conserved core is complemented by species-specific transcriptional programs, including a chondrocyte-related gene expression module in mouse lesions that is already detectable in shared mesenchymal cell types and becomes prominent in the mouse-specific chondrocyte-like cells.

### Location of shared and species-specific populations in atherosclerotic lesions of elastic arteries

To localize the shared and species-specific/enriched cell types in plaques from the respective species, we stained atherosclerotic lesions for RNA or protein encoded by top marker genes. The main continuum from shared contractile to shared terminal cells was visualized by double-staining for ACTA2 and LUM proteins in human, pig, and mouse plaques (**Fig. 5A-C**). Neither marker is specific to any individual subcluster, but their expression patterns delineate the gradual transition from ACTA2+ LUM-cells to ACTA2-LUM+ cells across the contractile, intermediate, and terminal clusters (**Fig. S3B**). Notably, the two markers together label nearly the entire population of mesenchymal plaque cells in all analyzed species (**Fig. S3C**), providing a practical tool for studies in human and pig lesions, where lineage tracing is currently not feasible. Consistent with the presence of a human/pig-enriched cluster of cells with strong contractile gene expression in the integrated scRNA-seq analysis, ACTA2+ plaque cells (with or without LUM expression) were more abundant in pig and human plaques than in those from mice (**Fig. S3D).**

**Figure 5.**
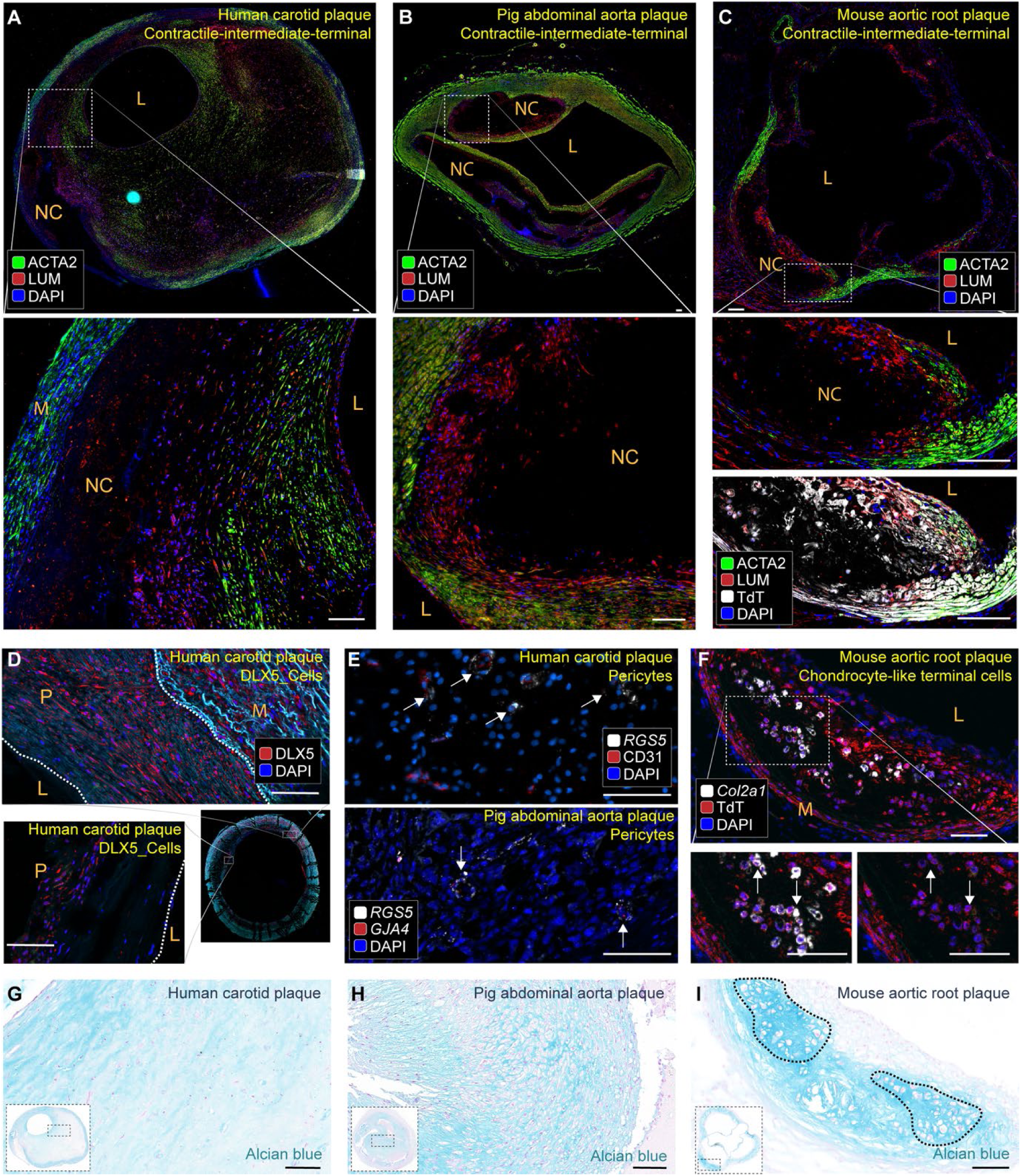
Mapping of shared and species-specific mesenchymal populations in atherosclerosis of elastic arteries. **A-C**, Representative ACTA2-LUM staining of human carotid plaque (N=6) (A), porcine abdominal aorta plaque from PCSK9 transgenic minipigs (N=6) (B), and mouse aortic root plaques from rAAV-PCSK9-induced mice (N=8) (C). Findings in Apoe-/- mice (N=8) were similar (not shown). Overview (top) and magnified views of the boxed regions are shown (bottom). ACTA2+ and ACTA2+ LUM+ cells predominate in the luminal part, while LUM+ cells localize to the plaque interior. **D**, Representative examples of DLX5+ cells in human carotid plaque (N=9) showing their localization in media and intima/plaque (N=9). The white dotted lines indicate the lumen and internal elastic lamina. **E**, Representative staining of pericytes (marked by arrows), identified as *RGS5+* cells wrapped around CD31+ endothelial cells in neovessels of human carotid plaques (N=4), and as *GJA4*+*RGS5*+ cells forming neovessel structures in porcine plaques (N=6). **F**, Mouse-specific chondrocyte-like cells mapped as *Col2a1+*tdTomato+ cells (marked by arrows) in SMC lineage-traced mice and found in the plaque interior. Representative of stained sections from aortic roots in rAAV-PCSK9-induced mice (N=8). Findings in Apoe-/- mice (N=8) were similar (not shown). **G-I,** Representative examples of Alcian blue-stained sections from human carotid (N=9), porcine abdominal aorta (N=6), and mouse aortic roots from rAAV-PCSK9-induced mice showing areas of chondroid metaplasia (marked by dotted lines) in murine but not porcine or human atherosclerosis. Findings in Apoe-/- mice (N=8) were similar (not shown). Scale bars, 100 μm. L, Lumen. NC, Necrotic core. P, plaque/intima. M, media.

The human-specific DLX5_Cells were identified by DLX5 antibody staining in human carotid endarterectomy samples. Positive staining was observed in 6/9 plaques, consistent with the variable presence of this cluster in scRNA-seq datasets. DLX5+ cells were primarily located in the media in a patchy manner (**Fig. 5D**); however, some samples also showed DLX5+ cells within the plaque. Pericytes were detected in neovessels of human plaques using CD31 immunostaining combined with RNAscope detection of *RGS5* mRNA, and in porcine plaques by double detection of *GJA4* and *RGS5* mRNA (**Fig. 5E**). *RGS5* expression was also seen in the fibrous cap and media, marking cells with the morphology of contractile SMCs (data not shown). Mouse-specific chondrocyte-like cells were identified by detection of *Col2a1* mRNA in aortic root plaques from Apoe^-/-^ and rAAV-PCSK9-treated mice carrying a tdTomato transgene for SMC-lineage tracing (**Fig. 5F**). These cells localized to areas with morphological features of chondroid metaplasia, which were not observed in human or pig plaques (**Fig. 5G-I**).

### Integration of plaque mesenchymal cells from pig and human coronary arteries

The pipeline used to identify shared and species-enriched mesenchymal cell populations was also applied to the human and pig coronary scRNA-seq datasets. A predefined limitation of this analysis was that, although PCSK9 transgenic pigs develop coronary atherosclerosis that morphologically resembles human disease^61^, lesion development is slower than in the abdominal aorta^62^. Consequently, the porcine lesions examined (after 15 months of high-fat diet) were likely less advanced than those from explanted human hearts, some of which were obtained from heart transplants due to advanced ischemic heart disease^12^.

CCA, RPCA, scANVI, and scVI integration algorithms successfully integrated the data based on visual assessment (**Fig. S4A**) and metrics (**Fig. S4B**). Clustering resolution (**Fig. S4C**) and naming of the stable clusters after their dominant contributor were performed as for elastic arteries. Since only two species were examined, the threshold for robust species-enrichment was set at ≥90% cell contribution in ≥6 integrated datasets. Contributions were calculated using random subsampling of 3600 cells per species, omitting small clusters with less than 100 sampled cells. Two representative examples showing shared and species-enriched clusters are presented in **Fig. 6A**, with additional results shown in **Fig. S5,** and a summary of findings below.

**Figure 6.**
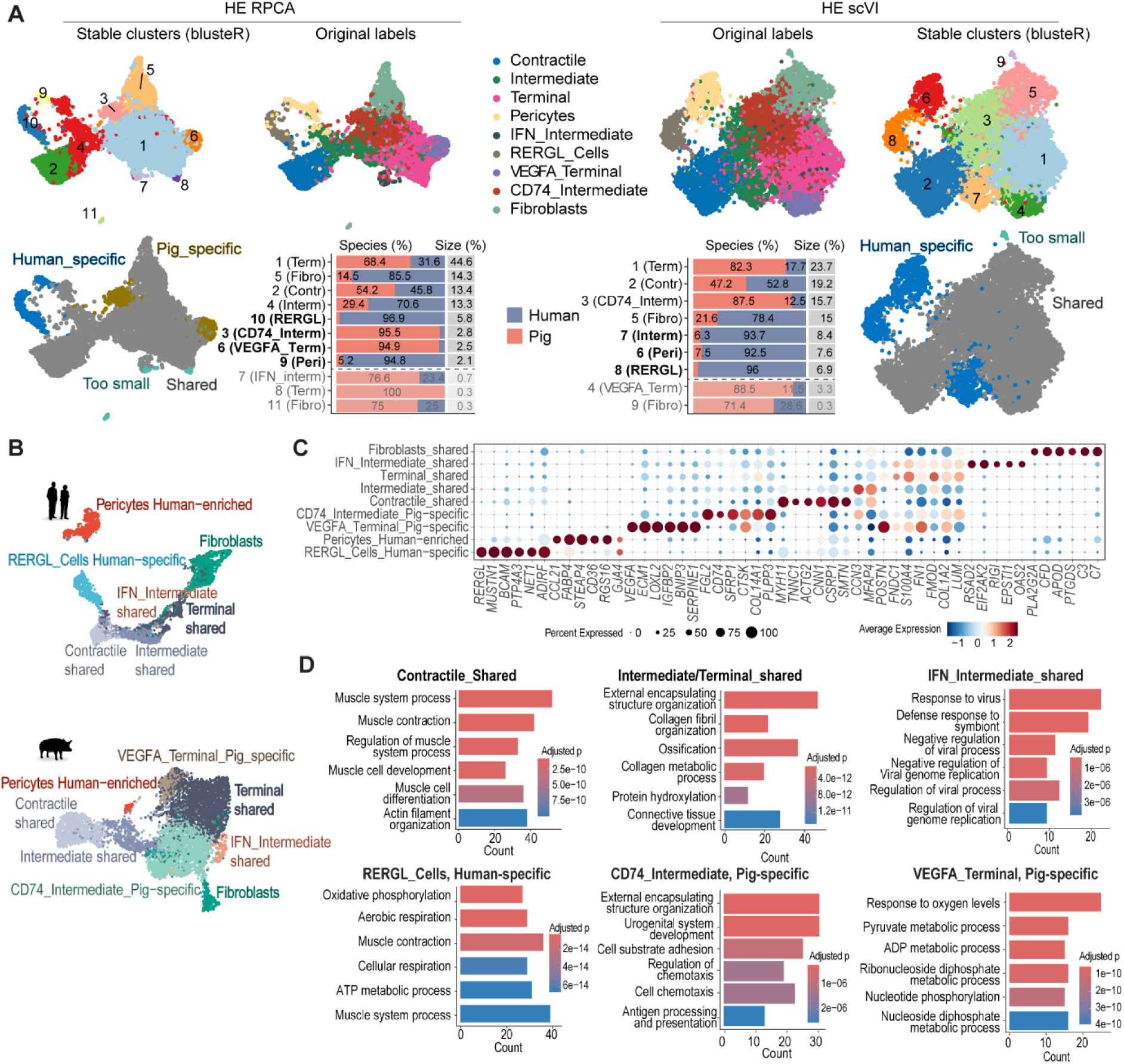
Identification of shared and species-specific mesenchymal plaque cell types in coronary arteries. **A**, Two examples of integrated datasets, HE RPCA and HE scVI, with stable clusters defined by *bluster* and cell labels from the individual species datasets shown in the top row. The bottom row illustrates the decision process for identifying species-enriched clusters. Stable clusters were defined as enriched (highlighted in bold) in one species if that species contributed ≥90% of cells (using 3600 random cells from each dataset and only considering clusters with ≥100 cells). The enriched clusters are named after the dominant contributor from the individual species datasets. **B,** Shared and species-enriched/specific cell types labeled in the pig and human datasets based on consensus among integrations. **C,** Top markers for the shared and species-enriched/specific clusters. Markers were analyzed in merged data from all species. **D,** Enrichment analysis for shared and species-enriched/specific mesenchymal cell phenotypes.

In all 12 integrated datasets, most cells (median 78.6%, IQR 71.0-89.7%) belonged to shared clusters. RERGL_Cells (median 97.7%, IQR 96.7-98.9%, in 9 integrations) and a pericyte population (94.4, IQR 92.2-95.1% in 6 integrations) were enriched for human cells. In contrast, two non-contractile clusters were enriched for pig cells: CD74_Intermediate (median 96.5%, IQR 95.4-99.6%, in 8 integrations) and VEGFA_Terminal cells (median 95.4%, IQR 95.0-96.3%, in 7 integrations).

To analyze transcriptional profiles, cluster labels were transferred back to the individual datasets. Cells belonging to species-enriched clusters (in ≥4 integrated data sets) were identified first. The remaining cells were considered shared and re-clustered (using HE RPCA as a representative integration) to separate them into contractile, intermediate, and terminal cells. This also resolved shared clusters of IFN_SMCs and fibroblasts (**Fig. 6B** and **Fig. S6A**).

The markers and enriched gene sets of shared contractile, intermediate, and terminal cells largely recapitulated those observed in plaques of elastic arteries. The main axis again extended from cells expressing canonical contractile markers (e.g., *MYH11, ACTG2, CNN1*) and gene sets related to contractility to intermediate and terminal cells characterized by expression of extracellular matrix genes (*LUM, COL14A1, CLU*) and gene sets related to *Collagen organization* and *Connective tissue development*. The population of IFN-SMCs shared markers with intermediate and terminal cell clusters but was distinguished from them by the strong induction of type 1 interferon response genes and enrichment for gene sets, such as *Response to virus*.

Analysis of the species-enriched clusters from the integrated datasets revealed that pericytes were also present in porcine coronary atherosclerosis (0.8% of all mesenchymal cells) but were merely less abundant than in the human dataset, where they constituted 15.2% of all mesenchymal cells. Other enriched populations were species-specific. The human-specific RERGL+ cells, which comprised 13.0% of mesenchymal cells, matched cell types variably described in the literature as contractile pericytes - due to the co-expression of pericyte markers (e.g*., KCNAB1, COX4I2, GJA4, MUSTN1*)^63^ - or as a medial SMC subpopulation due to co-expression of contractile genes.^64,65^ Gene set enrichment analysis indicated increased expression of *Oxidative phosphorylation* genes, suggesting that these may represent a more metabolically active subset of contractile cells compared to canonical contractile SMCs. The pig-specific CD74_Intermediate cluster, which comprised 28.3% of pig mesenchymal cells, shared markers with other intermediate-type cells but exhibited elevated expression of CD74 as well as several other genes involved in antigen presentation, suggesting an inflammation-associated cell state. The pig-specific VEGFA_Terminal population, constituting 6.7% of pig mesenchymal cells, expressed genes (e.g., *VEGFA, IGFPB2, EGLN3*) and gene sets (e.g., *Response to oxygen levels*) related with hypoxia and its metabolic responses (**Fig. 6C,D**).

### Location of shared and species-specific populations in atherosclerotic lesions of coronary arteries

The main mesenchymal phenotypic spectrum - from contractile to intermediate and terminal cells - was mapped by ACTA2/LUM immunostaining in sections of human and porcine coronary atherosclerosis (**Fig. 7A,B**). ACTA2/LUM staining revealed an architecture consistent with that observed in human carotid and porcine abdominal aorta lesions: ACTA2+ and ACTA2+/LUM+ cells were located near the lumen and in the media, while ACTA2-LUM+ cells were restricted to the plaque interior.

**Figure 7.**
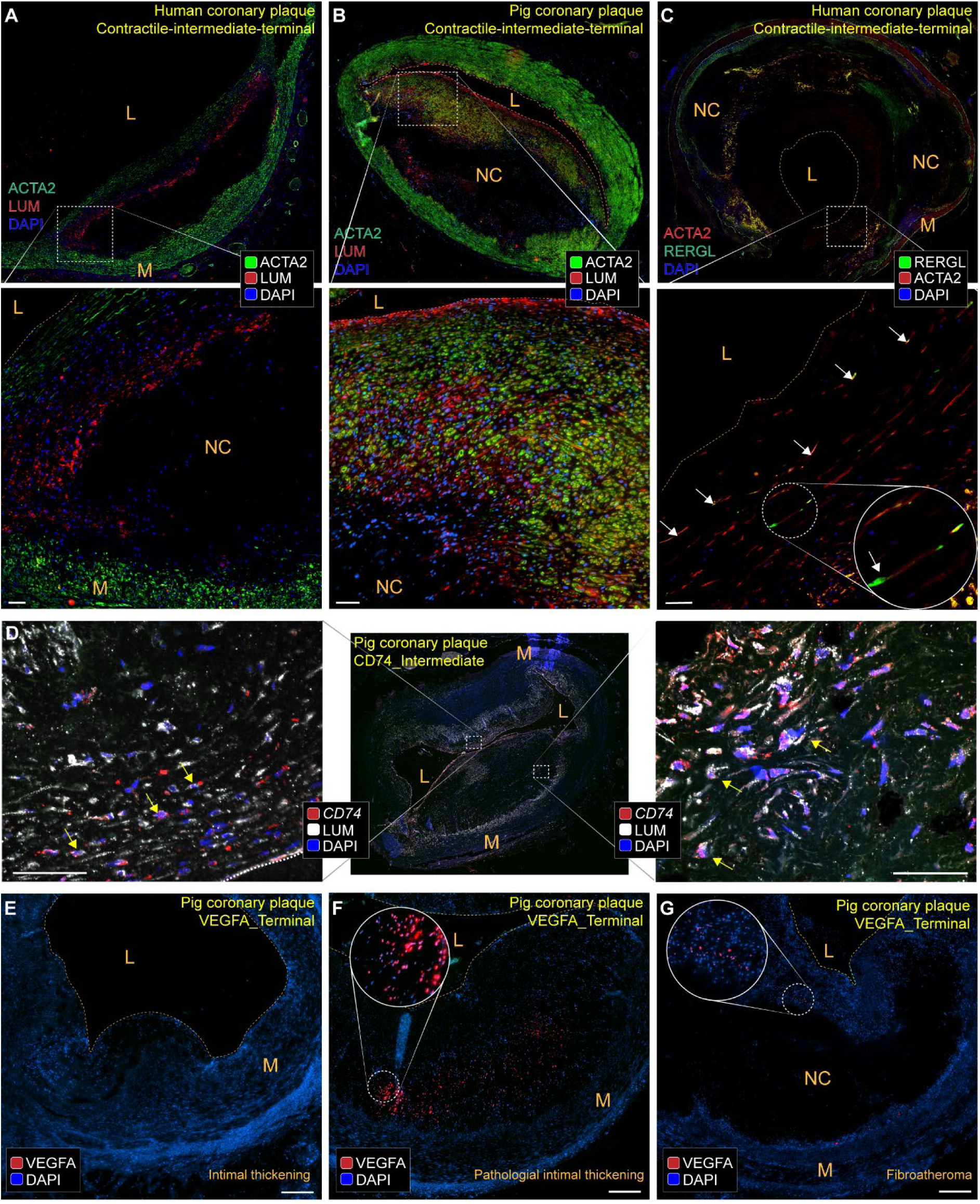
Mapping of shared and species-specific mesenchymal populations in coronary atherosclerosis. **A-B**, Representative ACTA2-LUM staining of human (N=5) (A) and porcine (N=6) (B) coronary lesions. Overview (top) and magnified views of the boxed regions (bottom) are shown. **C**, Representative examples of combined RERGL and ACTA2 immunostaining showing RERGL+ACTA2+ (marked by white arrows) in the fibrous cap region of a human coronary plaque (N=7). **D,** Representative images of combined CD74 mRNA RNAscope and LUM immunstaining in porcine coronary plaque (N=5). Cells with co-expression are both in the fibrous cap (left) and in the interior region of advanced lesions (right). Overview (middle) and magnified views of the boxed regions (sides) are shown. The dotted line indicates lumen border. **E-G,** RNAscope for VEGFA in porcine LAD in intimal thickening (E), pathological intimal thickening (F), and large fibroatheroma (G), respectively (N=4). VEGFA signal is most abundant in early stages of plaque development. Scale bars, 50 μm. L, Lumen. NC, Necrotic core. P, plaque/intima. M, media.

The human-specific RERGL_Cells population was localized by combined RERGL and ACTA2 immunostaining and identified in 6 out of 8 human coronary fibroatheromas. They were found in discrete regions of the fibrous cap, exhibiting a spindle-shaped morphology and layered organization (**Fig. 7C**). The CD74_Intermediate pig cells were detected by combining RNAscope for *CD74* mRNA with LUM immunostaining to distinguish them from CD74-expressing immune cells. They appeared in plaque shoulder regions but also deeper inside advanced plaques, (**Fig. 7D)**. VEGFA_Terminal cells, detected by RNAScope for *VEGFA* mRNA, were absent in adaptive intimal thickening and small lesions (**Fig.7E**), abundantly expressed in the core of pathological intimal thickenings (**Fig. 7F**), and restricted to the necrotic border zone in fibroatheromas (**Fig. 7G**). This spatial and temporal pattern suggests that *VEGFA* expression marks regions of the plaque destined for necrosis.

### Overview of nomenclature, subpopulations, and their markers across species

Single-cell RNA-seq studies of mouse and human atherosclerotic plaques have introduced over 15 different terms to describe mesenchymal plaque cell types, many of which carry implied meanings about derivation (e.g., fibromyocytes), fates (e.g., transitional), or function (e.g., fibrochondrocytes)^12,13,22,23,66^. While these terms are supported in the specific contexts in which they were coined, they are not necessarily translatable across species. In the current study, we have therefore mainly relied on generic terms based on the relative positioning of mesenchymal cells in UMAP plots. The identified shared and specific populations are summarized in **Figure 8**, alongside key terms from prior mouse and human studies to contextualize the findings within the existing literature. These terms were positioned on the map based on reported marker gene expression (see **Table S9** and Chen et al.^67^).

**Figure 8.**
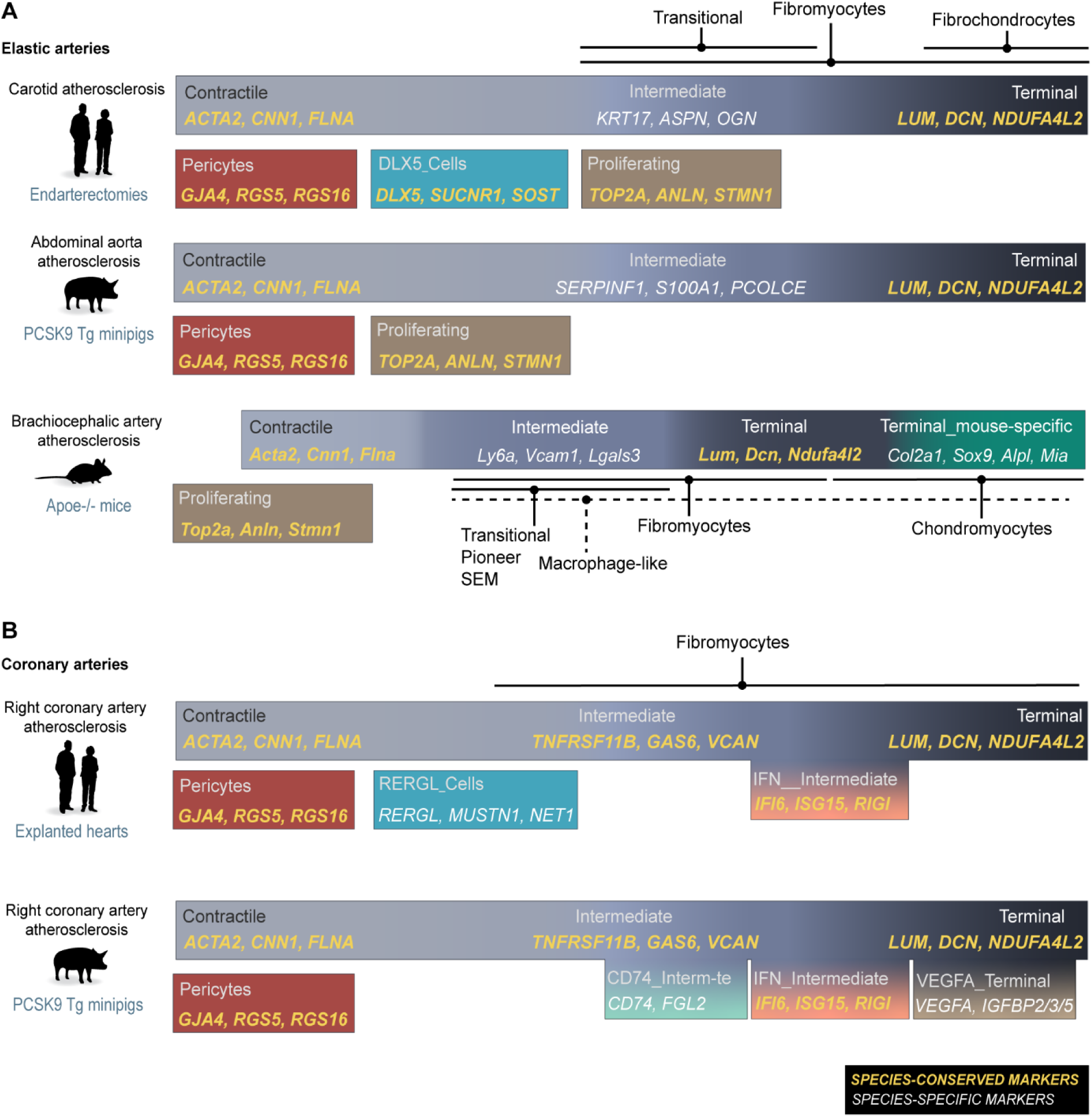
Summary of shared and specific mesenchymal cell types and their markers. The schematic provides an overview of the mesenchymal cell subtypes/states identified in the analysis, along with their correspondence to terminology used in the literature. Most mesenchymal cells form a continuous population in UMAP space, while some cell types/states (e.g., pericytes, DLX5_Cells) are sufficiently distinct to form separate clusters. Cell states that appear to be driven by specific signaling activity in intermediate/terminal cells are shown as side populations. Conserved markers are shown in yellow; markers that only work within a specific species are shown in white.

To identify conserved markers for mesenchymal cell populations that may aid translational workflows from experimental to human atherosclerosis, we overlapped marker gene lists for each shared mesenchymal cell phenotype (**Table S7** and **S8**). Such conserved markers (e.g. ACTA2 for contractile cells and LUM for terminal cells) were identified for most cell states in both elastic and coronary atherosclerosis. Notably, markers often used in murine studies to identify intermediate-type cells - also referred to as pioneer, transitional or SEM (stem, endothelial, and macrophage-like) cells - such as *LGALS3, VCAM1*, and *LY6A*^13,20,66^, were not suitable for identifying intermediate-type cells in human or porcine lesions. This was due either to gene absence (*LY6A*) or lack of specificity (*LGALS3, VCAM1*) (**Fig. S6**). Indeed, no conserved markers were identified that could define an intermediate-type cell state across murine brachiocephalic, porcine abdominal, and human carotid atherosclerosis. In contrast, intermediate-type cell markers such as TNFRSF11B and GAS4 were conserved across human and porcine coronary lesions.

## Discussion

Translational atherosclerosis research benefits from a thorough understanding of the similarities and differences in cell types and states between human disease and the models used to investigate it. Previous scRNA-seq studies comparing mouse and human lesions have addressed this for plaque macrophages^68^ and adaptive immune cells^69^. In the present study, we extend this comparative approach to mesenchymal cell populations. Furthermore, we incorporate porcine atherosclerosis, which despite its close anatomical and pathological resemblance to human atherosclerosis, has not previously been analysed for mesenchymal cell diversity. We identify a conserved core of mesenchymal cell diversity present across species and vascular beds, as well as mesenchymal cell populations unique to specific contexts. Some, such as DLX5 or RERGL-expressing SMCs, are found only in human carotid or coronary atherosclerosis and not in experimental models. Conversely, *Col2a1*-expressing chondrocyte-like cells are present in mouse lesions but absent from advanced human carotid or coronary plaques. Recognizing these similarities and differences is essential for accurately interpreting murine and porcine studies of plaque mesenchymal cells.

A key feature of our approach is the use of consensus across multiple integration and gene homology strategies. While many studies rely on a single integration method to infer overlap in cell types between mouse and human plaques^12,13^, our analysis shows that different methods yield differing results. Using the BENGAL pipeline^27^, we performed 9 data integrations and applied 3 gene homology-matching approaches, basing our conclusions on agreement among at least half of the 12 best-performing method combinations. In addition, because cluster number is subject to analyst choice, we used cluster stability metrics to select the most robust resolution.

Some of the observed differences in plaque mesenchymal cells between pig and human coronary atherosclerosis may reflect differences in lesion stage or disease activity. The *VEGFA*-expressing cells that were found in porcine lesions appeared to transiently populate the regions of pathological intimal thickenings destined to convert into necrotic core(s). This is consistent with *VEGFA* being downstream of hypoxia and with hypoxia acting as a causal driver of necrotic core formation in mice^70^. It is understandable that similar widespread hypoxic areas will be less prevalent in advanced human lesions, where hypoxic areas may have already resolved through neovessel formation or necrosis. Similarly, the increased prevalence of *CD74-*expressing cells may reflect the high inflammatory activity of accelerated porcine atherosclerosis compared with coronary atherosclerotic lesions from heart transplant patients, who are likely under medical therapy to lower atherosclerotic disease activity. Indeed, CD74+ mesenchymal cells have also been described in humans^71^, and CD74 was recently implicated causally in murine atherogenesis^72^. More scRNA-seq data at different stages and levels of disease activity are needed to fully clarify this.

Our cross-species comparison of atherosclerosis in major elastic arteries also involved differences in anatomical location. This could be seen as a limitation if the goal was to isolate the pure effect of species biology. SMCs vary in embryonic origin across the arterial tree^73^, and because the transition of SMCs in atherosclerosis involves the reactivation of developmental pathways^74^, this may theoretically influence the resulting phenotypes. For example, human DLX5_Cells, bearing a neural crest signature, were found in the neural crest-derived carotid but not in the second heart field- and epicardial-derived coronary artery^75^. However, embryonic origin alone cannot explain the absence of DLX5_Cells in the neural crest-derived murine brachiocephalic artery, nor the presence of chondrocyte-like cells in mouse brachiocephalic but not human carotid plaques. Importantly, translational research must contend with the fact that clinically important arterial sites for human atherosclerosis cannot always be studied in experimental models. For example, pigs lack a proper carotid bifurcation^76^, the predilection site of human carotid atherosclerosis, and mouse coronary arteries are small and mostly intra-myocardial^77^. Therefore, using porcine and murine models to understand human carotid or coronary atherosclerosis, respectively, necessarily involves extrapolating across vascular sites. Arguably, the most salient finding of our analysis is that the majority of mesenchymal cells belonged to shared clusters: cell types present in all analyzed plaques, irrespective of species, disease stage, or anatomical location. This supports the utility of animal models for studying key aspects of mesenchymal cell function in human atherosclerosis. To this end, we provide conserved markers and regulon analysis that may facilitate the flow of information between models and human disease. Nevertheless, two important exceptions warrant mention. First, murine chondrocyte-like cells, often referred to as chondromyocytes^78^, were not detected to any significant extent in porcine or human atherosclerosis. While other mesenchymal phenotypes (e.g., DLX5_Cells) exhibit ossification-related signatures, they are molecularly distinct. Chondroid metaplasia has previously been reported as rare in human coronary atherosclerosis, consistent with our findings, but not uncommon in peripheral artery disease (e.g., femoral plaques)^79,80^. Whether murine chondrocyte-like cells correspond to subtypes of mesenchymal cells in peripheral atherosclerosis remains an open question for future scRNA-seq studies. Second, markers such as VCAM1 and LGALS3, commonly used to define transitional SMC-derived states in mice, cannot be applied equivalently in human or pig lesions. Although comparable contractile and matrix-producing (fibromyocyte-like) states are clearly present across species, our cross-species analysis did not support the existence of a distinct, conserved intermediate cell type. That said, analysis of more data at higher cluster resolution could potentially uncover small, shared populations with such transitional characteristics.

## Methods

### Deposited scRNA-seq datasets

Murine plaque scRNA-seq data count matrices (GEO GSE150644, 15 *Apoe-/-* mice with lineage tracing of SMC-derived cells, Alencar et al.^20^) were downloaded from the Gene Expression Omnibus (GEO). Only sorted SMC-derived (eGFP+, tdTomato+, or eYFP+) cells were used. Doublets were removed with scDblFinder (v1.12.0)^81^, low-quality cells were removed (genes <800, UMIs >25000, percent of mitochondrial counts >15% and percent of hemoglobin counts >1%), and datasets were integrated (Seurat RPCA algorithm) and clustered at moderate resolution. The final dataset comprised 2412 cells (482±327 cells per sample) with 13243 expressed genes. Human scRNA-seq raw data of carotid endarterectomies (GSE155512, 3 patients, Pan et al.^13^ and GSE159677, 3 patients, Alsaigh et al.^14^) and atherosclerotic coronary artery segments (GSE131778, 4 patients, 8 samples, Wirka et al.^12^) were downloaded (GEO database) and processed (CellRanger v6.1.0) using the human reference genome Ensembl hg38 2020. Data were cleaned of doublets (scDblFinder) and low-quality cells (number of UMI >25000 or <1200, number of detected genes > 4000 or <1000, cell with high rate of mitochondrial genes expressed >10%, percent of hemoglobin counts > 1%, library complexity <80%), and ribosomal, mitochondrial, and lowly expressed (less than 5 counts in total) genes were removed. Datasets were integrated (Seurat RPCA algorithm) separately for carotid endarterectomy samples and coronary plaques, and the mesenchymal cell population was identified based on expression of marker genes (*ACTA2, LUM*), isolated, and re-clustered with moderate resolution. The final datasets consisted of 8264 mesenchymal cells (1204±570 cells per sample) with 18924 expressed genes for carotid endarterectomy samples, and 3609 mesenchymal cells (516±234 cells per individual library) with 16242 expressed genes for the coronary artery samples.

### Porcine plaque scRNA-seq datasets

Animal procedures were approved by the Comunidad de Madrid (license nos. PROEX265/16 and 043.6/21). PCSK9 transgenic minipigs (2 males and 5 females) were fed high-fat, high-cholesterol diet from 3 to 18 months of age to induce severe hypercholesterolemia and atherosclerosis. The group of pigs served as the control group of a recently reported atherosclerosis regression study,^21^ and readers are referred to this work for details on diet composition, plasma biochemistry, and lesion histopathology. Immediately after sedation (by 5 mg/kg Azaperone, 6 mg/kg Tiletamine and Zolazepam, and 0.1 mg/kg Medetomidine) and euthanization (by 50 mg/kg pentobarbital), raised lesions were harvested from the opened right coronary artery. Aorta plaques from the distal abdominal aorta were dissected free of underlying arterial media as previously reported^21^.

Plaques were minced, washed in HBSS, and digested 60 minutes at 37°C in an enzyme solution containing 1.5 mg/ml (8 U/ml) elastase (LS002279, Worthington Biochemical), 2 mg/ml liberase (05401119001, Sigma Aldrich), and 300 µg/ml DNaseI (Sigma Aldrich, DN25). The reaction was stopped with 0.5% bovine serum albumin, and cells were sorted for viability in a FACSAria Cell Sorter after staining with DAPI (Sigma-Aldrich, No 1.24653.0100) (final concentration 1 µg/ml) and Draq5 (Thermo Scientific 65-0880-96; final concentration, 5 µM). Viable cells (Draq5 positive and DAPI negative) were processed for scRNA-seq using the Chromium Next GEM Chip G and Chromium Next GEM Single Cell 3’ Kit v3.1 (10x Genomics). Mean library size was calculated using a 2100 Bioanalyzer (Agilent), and the concentration was determined with a Qubit® fluorometer (Thermofisher). Libraries were sequenced in paired-end reads using a HiSeq 4000 system (Illumina) and processed with RTA v1.18.66.3. FASTQ files were aligned to Sus scrofa reference genome (v11.1) using CellRanger software (v6.0.0). Count matrices containing all cell types from the atherosclerotic plaques were cleaned from low quality cells (number of detected molecules ≤1200, number of detected genes ≤900, cells with high (>10%) content of mitochondrial counts, and percent of hemoglobin counts >1%) and doublets (scDblFinder R package). Ribosomal, mitochondrial, and lowly expressed (less than 5 counts in total) genes were removed. The mesenchymal cell supercluster was identified by expression of *ACTA2* and *LUM* and isolated yielding 3153 cells (526±299 cells per pig) with 17481 expressed genes in aortic plaques and 6222 cells (1244±542 cells per pig) with 19475 expressed genes in right coronary arteries. SMCs were re-clustered at moderate resolution to identify and characterize the main SMC phenotypes. Data from the abdominal aorta have been previously reported^21^.

### Cross-species scRNA-seq integration and metrics using the BENGAL pipeline

Datasets were prepared for processing in the BENGAL pipeline as recommended^27^. Gene names were converted to ENSEMBL ids using the gprofiler R package (v0.2.2, function gconvert()), and data were stored in an AnnData object. The cluster names in **Figure 1** were used as labels, and the species from which the data were derived as batch and species keys.

Homologous genes were matched by the BENGAL pipeline in 3 different ways: one-to-one orthologs (O2O) based on biomaRt database information, or expanded sets that include one-to-many and many-to-many orthologs, selected based on higher average expression levels (HE), calculated in the individual species datasets, or higher homology confidence (HHC) in biomaRt.

Metrics evaluating batch (species)-mixing (Principal component regression (PCR), batch average silhouette width (bASW), graph connectivity (GC), k-nearest neighbors batch effect test (kBET), integration local inverse Simpson’s Index (iLISI)) and biology conservation metrics (adjusted Rand index (ARI), cell type average silhouette width (cASW), cell type local inverse Simpson’s Index (cLISI), isolated label F1 score (iF1), Normalized Mutual Information (NMI)) were calculated in R using the following packages: kBET v0.99.6, lisi v1.0.0, FNN v1.1.3.2, cluster v2.1.3, bluster v1.8.0, aricode v1.0.3 with some examples of code from https://github.com/MillerLab-CPHG/Human_athero_scRNA_meta.git ^11^. PCR, bASW, GC, cASW, and iF1 scores were subtracted from 1 to make score directionality aligned with the other metrics, with the lowest value indicating batch removal failure and the highest value complete mixing. Raw metrics were min-max scaled, so the metrics have equal weight when averaged. Scripts for metrics plotting were obtained from Song et al^27^.

### Clustering and species-specificity

Integrated datasets from top-performing algorithms (RPCA, CCA, scVI, scANVI), each with the 3 different gene homology match approaches, were gathered and imported as Seurat objects, preserving the original UMAP structure from the BENGAL pipeline. Each of the 12 integrated datasets was clustered using assessment of cluster stability with the bluster R package to find the optimal resolution^36^. Briefly, we performed 20 iterations using bootstrapping at different resolutions (ranging from 0.1 to 1), assessed the stability of clusters based on the average adjusted Rand index (ARI), and chose the resolution with maximum average ARI and at least 7 clusters for elastic arteries and 6 clusters for coronary arteries. The rationale for the minimum number of clusters was that integrated datasets should have at least the number of mesenchymal cell types observed in the individual species datasets. The final resolution varied for each dataset, resulting in a different number of clusters (8-11) in each integrated dataset.

Clusters adopted their names from the major contributor of the individual species dataset to indicate cluster properties. To find species-specific and shared clusters, we calculated the contribution of each species to a cluster and set cutoffs as follows. In elastic arteries, a stable cluster was species-specific if one species contributed ≥80% or two species contributed ≥90% (equivalent to one species contributing <10%) of cells in ≥6 integrated datasets. In coronary arteries, the cutoff was defined at 90% in ≥6 integrated datasets, since only two species were analyzed. The proportions were calculated on randomly sampled cells from each species (1500 for elastic and 3600 for coronary arteries). Integrated clusters with ≥50 cells (elastic arteries) or ≥100 cells (coronary arteries) were considered for subsequent analysis. Cells in the individual datasets were categorized as being of a particular species-specific type if they belonged to that species-specific cluster in ≥4 integrated datasets. Cells not meeting that criterion were considered of a shared type. Those cells were separated and re-clustered in the HE_RPCA dataset, and labels were transferred to the individual species datasets. Markers were calculated in merged datasets with the transferred consensus labels using random subsampling of 2500 cells from each individual dataset and Seurat (v4.4.0)^32^ and R (v4.4.3). Over-representation analysis was performed using the EnrichR R package (v3.2), the GO database, and marker genes identified by the FindAllMarkers() function with arguments min.pct=0.3, only.pos.=TRUE. The enriched categories were ordered by adjusted p-value, only significantly enriched categories are shown (p-value adjusted < 0.05). Conserved markers for shared phenotypes were identified by overlapping lists of gene markers (log2FC > 0.6) from the individual datasets.

### Analysis of bulk RNA sequencing data

Published bulk RNA-seq datasets of human carotid plaques (GSE120521), mouse atherosclerotic aortas (GSE205929), and pig atherosclerotic aortas (GSE245530) were downloaded as FASTQ files and processed with the “nf-core/rnaseq” pipeline (v3.14.0)^82^. This included alignment with the appropriate reference genome (hg38, mm10, or Sscrofa11.1) using STAR (v2.6.1d)^83^ and quantification with Salmon software (v1.10.1)^84^. To compare expression of genes of interest, matrices were normalized with the median of ratios method implemented in DESeq2^85^.

### Transcription factor activity inference

SCENIC was used to infer regulon activity scores in human and mouse mesenchymal scRNA-seq datasets^46^. Activities were binarized to mitigate the context-dependency of AUC scores in individual datasets, and the binarized matrices from both species were merged, preserving all inferred regulons. For regulons that were not found in a species, all cells were assigned an activity of 0. To analyze differences in regulon activity associated with shared mesenchymal cell types, we calculated for each shared cluster (excluding proliferating cells because of its small size) and regulon active ≥30% of cluster cells in ≥1 species the metric (f_h_-f_m_)/(f_h_+f_m_), where f_h/m_ is the fraction of cells with the regulon active in human or mouse, respectively. To identify regulon activity associated with species-specific clusters, we added the merged matrix to the integrated HE_RPCA dataset as an Assay and used the function FindMarkers to find mesenchymal type-specific regulons on this AUC Assay.

### Histological material

Paraffin sections (5 µm) of formalin-fixed human carotid plaques (N=11, all females) from endarterectomies were obtained from a previously described material^86^. Ethics board approval was not required (Danish Act no. 593 of 14 June 2011 on Research Ethics Review of Health Research Projects) because the samples were anonymized and removed during standard surgical procedures. Paraffin sections (5 µm) of formalin-fixed human proximal left anterior descending (LAD) coronary artery (N=6) were obtained from a repository of arterial segments collected during forensic autopsies at the Institute of Forensic Medicine, University of Aarhus, Denmark, between 1996 and 1999^87^. The samples were obtained from individuals aged 20-80 years, irrespective of the cause of death, with authorization from the Regional Research Ethics Committee and the Danish Data Protection Agency, which complies with the Declaration of Helsinki. The sample collection is anonymized and is not linked to any clinical information except for sex and age group (<45 or ≥45 years). Samples from individuals aged ≥45 years were decalcified in 10% formic acid for 24 hours before paraffin embedding. The present study was exempted from separate ethical approval.

Cryosections (5 µm) of fixed murine aortic roots were obtained from Myh11-CreERT2 x tdTomato C57BL/6J mice injected with AAV-PCSK9 at 8 weeks of age and 8 Myh11-CreERT2 x tdTomato Apoe-/- C57BL/6J mice, both fed on a high-fat diet from 8 to 26 weeks of age. Animal procedures were approved by the Comunidad de Madrid (license nos. PROEX020.8/21 and PROEX50/16).

Formalin-fixed, paraffin-embedded (4 µm) sections of the proximal LAD and abdominal aorta were obtained from the same pigs used for scRNA-seq (N=6).

### Staining procedures

Immunostaining was performed with antibodies against DLX5 (dilution 1:2300, ab109737, Abcam), RERGL (dilution 1:200, hpa041740, Merck), ACTA2 (1:200, M0851, Dako), LUM (1:200, ab198974, Abcam). Sections were dewaxed, rehydrated, pretreated in 0.05 % Tween-20 Tris-EDTA (pH 9.0) for 20 minutes at 95 °C, permeabilized in PBS containing 0.3 % Triton X-100 for 10 minutes at room temperature, and blocked for 1 hour in PBS with 0.1 % Tween 20 and 10% normal donkey or goat serum (16210-072, Life Technologies). Primary antibodies were applied overnight at 4 °C, and sections were then washed and incubated for 1 hour at room temperature with fluorophore-conjugated secondary antibodies (1:500, Alexa Fluor 594 Donkey Anti-Rabbit, A21207, Alexa Fluor 488 Donkey Anti-Mouse A21202, or Alexa Fluor 568–conjugated Goat Anti-rabbit, A-11011, Invitrogen). Nuclei were counterstained with DAPI (Sigma-Aldrich 1.24653) before mounting in SlowFade Antifade medium (Invitrogen S36937). Control slides were stained with isotype-matched antibodies at identical concentrations to confirm specificity.

RNAscope was performed using reagents (323110) and probes from ACD Bio for porcine *VEGFA* mRNA (429211-C1), porcine *CD74* mRNA (1103941-C1) and *GJA4* mRNA (580041-C3), mouse *Col2a1* mRNA (407221-C2), human and pig *RGS5* mRNA (533421-C1 and 1801301-C2, respectively), and Opal 570, Opal 650, and/or Opal 690 (FP1488001KT, FP1496001KT, FP1497001KT, respectively) from Akoya Biosciences. RNAscope for *RGS5* mRNA was combined with anti-CD31 (ab28364, Abcam), *CD74* mRNA with anti-LUM (ab198974, Abcam), and *Col2a1* mRNA with anti-RFP (1:100, Ab62341) (to detect endogenous tdTomato).

Stained sections were analyzed in NIS-Elements (Nikon) and images were processed in Fiji (ImageJ)^88^ and QuPath software (v0.5.1)^89^. Quantification of the fraction of ACTA2+ cells in plaque was performed by training a cell classifier in QuPath.

Alcian Blue staining of mouse aortic root, pig abdominal aorta plaques, and human carotid endarterectomies was carried out by the histology unit at CNIC using a standard protocol, and sections were scanned in a NanoZoomer 2.0-RS (Hamamatsu) digital slide scanner with 20× magnification.

## Acknowledgment

We thank Dorte Wilhardt Qualmann, Lisa Maria Røge, Ainoa Caballero Molano, Leticia González-Cintado, and Vanessa Cumbicus for excellent technical assistance. We also extend our thanks to members of the Animal Facility and Histopathology Unit, Microscopy Unit, Genomics Unit, Bioinformatics Unit, and Cellomics Unit at CNIC, as well as the Confocal Microscopy Core at the Department of Clinical Medicine, Aarhus University, for their excellent technical support. We thank Carlos Torroja and Fátima Sánchez Cabo for their help with bioinformatics analysis. Furthermore, we thank Yuyao Song from the European Bioinformatics Institute, Cambridge, for helping implement the BENGAL pipeline in the study.

