## Supplementary Figures 1-7 for "Mouse, pig, and human atherosclerotic lesions have common and distinct mesenchymal cell populations"

**Supplementary material**


| | |
|---|---|
| Supplementary Figure 1 | 2 |
| Supplementary Figure 2 | 3 |
| Supplementary Figure 3 | 4 |
| Supplementary Figure 4 | 5 |
| Supplementary Figure 5 | 6 |
| Supplementary Figure 6 | 7 |
| Supplementary Figure 7 | 8 |

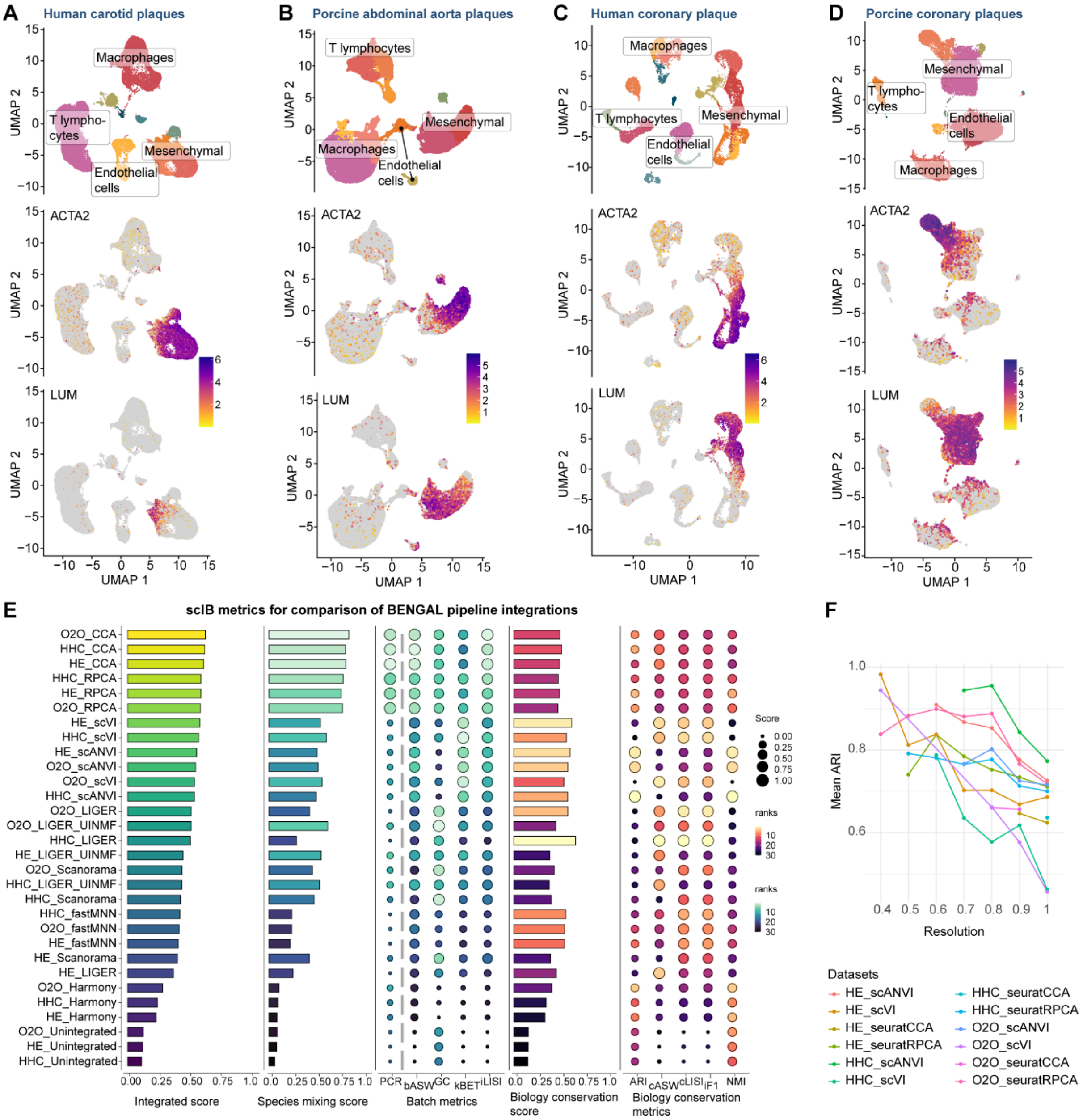


**Supplementary Figure 1. Individual species scRNA-seq data sets, integration metrics, and stable clusters in integrated data. A-D,** UMAP representations of atherosclerotic plaque scRNAseq data from human carotid (from Pan et al. (ref. 13) and Alsaigh et al. (ref. 14)), pig abdominal aorta (from Nogales et al. (ref. 21)), human coronary artery (from Wirka et al. (ref. 12)) and pig coronary artery (this paper). Plots show identified superclusters (top) and expression of markers for SMCs (ACTA2) and other mesenchymal cells (LUM) (below). **E,** ScIB metrics (batch mixing and biology conservation) and their ranks assessing 'goodness' for integration of elastic artery plaque scRNAseq datasets. **F**, Mean diagonal Adjusted Rand index (ARI) from bootstrapStability() function with 20 iterations for different clustering resolutions of the integrated datasets of elastic arteries.

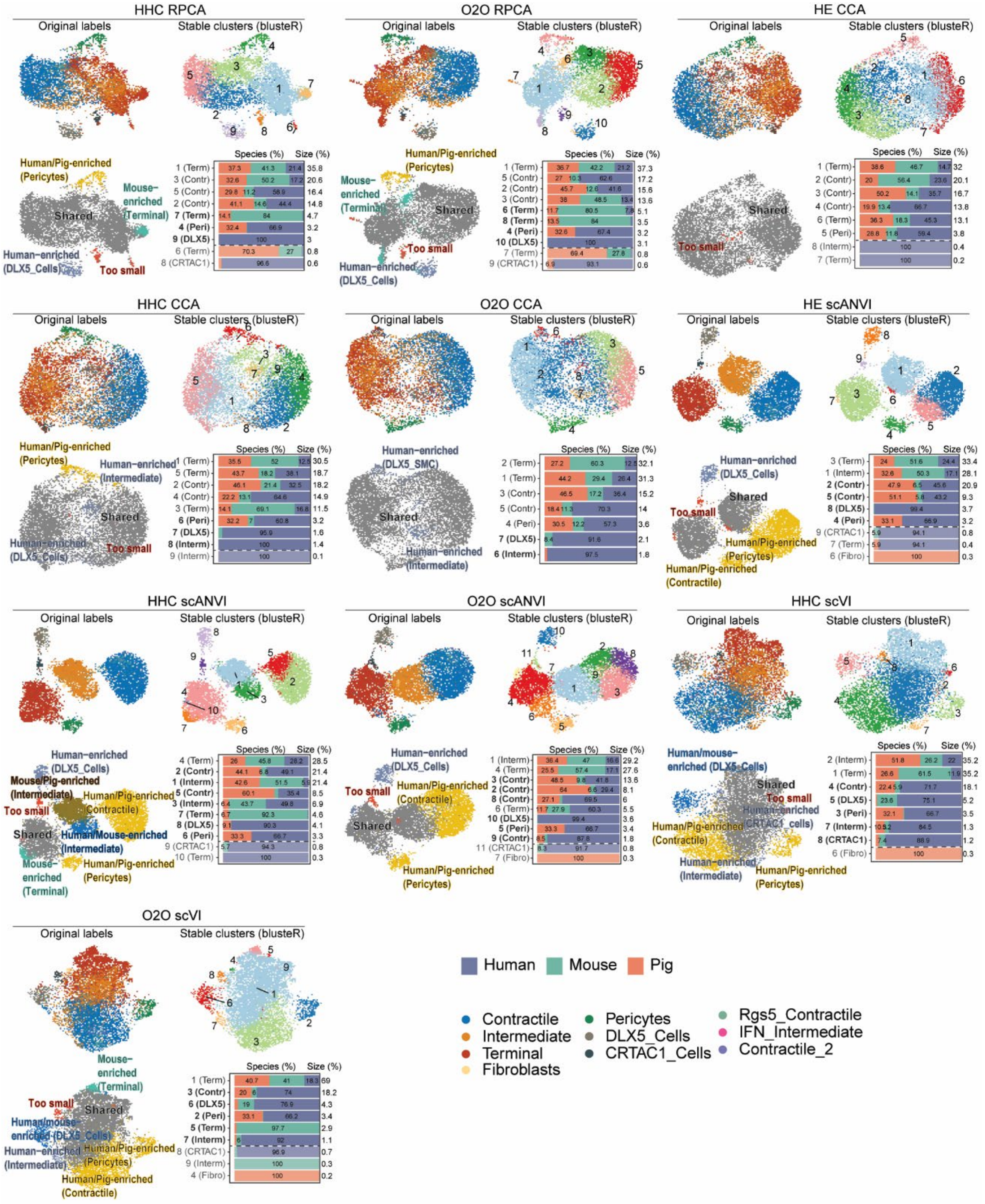


**Supplementary Figure 2. Identification of shared and species-specific mesenchymal plaque cell types in elastic artery atherosclerosis (additional data).** The ten additional integrations (to the two shown in Figure 2) produced by RPCA, CCA, scANVI, and scVI combined with the 3 gene homology match approaches. Each set of 4 plots consists of integrated dataset with cell type labels from the individual species data sets (top left); stable clusters defined using the *bluster* R package (top right); clusters fulfilling the criteria for being species-enriched (bottom left); and percentage contribution from each species to stable clusters (bottom right). Fractions were calculated using a random subsample of 1500 cells from each species dataset, and one or two species were considered overrepresented in a cluster if contributing ≥80% or ≥90% of cells, respectively. Only clusters with more than 50 cells (1.1%) were analyzed for species contributions.

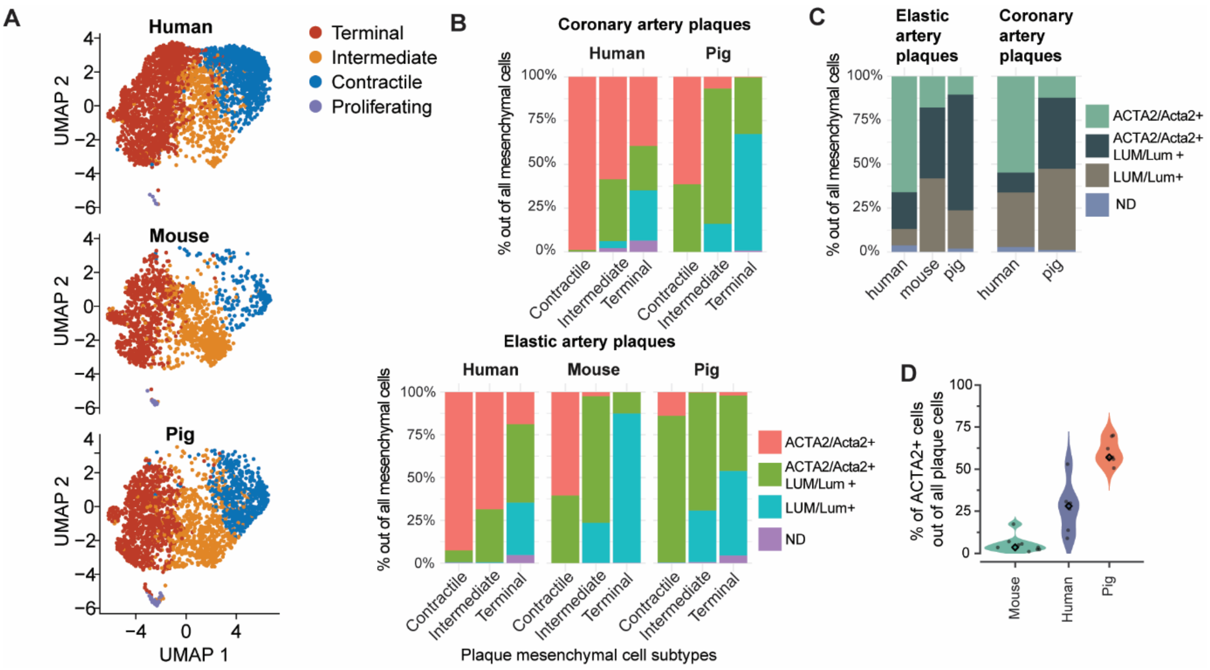


**Supplementary Figure 3. Reclustering of shared cell types and their markers. A**, UMAP representation of re-clustered species-shared SMC phenotypes of mouse, human and pig elastic arteries (using HE RPCA as a representative integration). **B-C**, Distribution of *ACTA2/Acta2*-expressing (ACTA2/Acta2+), *ACTA2/Acta2*- and *LUM/Lum*-expressing (*ACTA2/Acta2+ LUM/Lum* +), *LUM/Lum-expressing (LUM/Lum+)*, and cells without detectable *ACTA2/Acta2 and LUM/Lum* expression in contractile, intermediate, and terminal cell clusters (B) and in all plaque mesenchymal cells (C). **D**, Quantification of the percentage of ACTA2+ cells out of all plaque cells in human carotid (N=6), pig abdominal aorta (N=7), and mouse aortic root plaques (N=9). Quantification was performed in ACTA2- and DAPI-stained sections and the percentage calculated as the number of DAPI-stained nuclei in ACTA2+ cells out of all DAPI-stained nuclei within the plaque area.

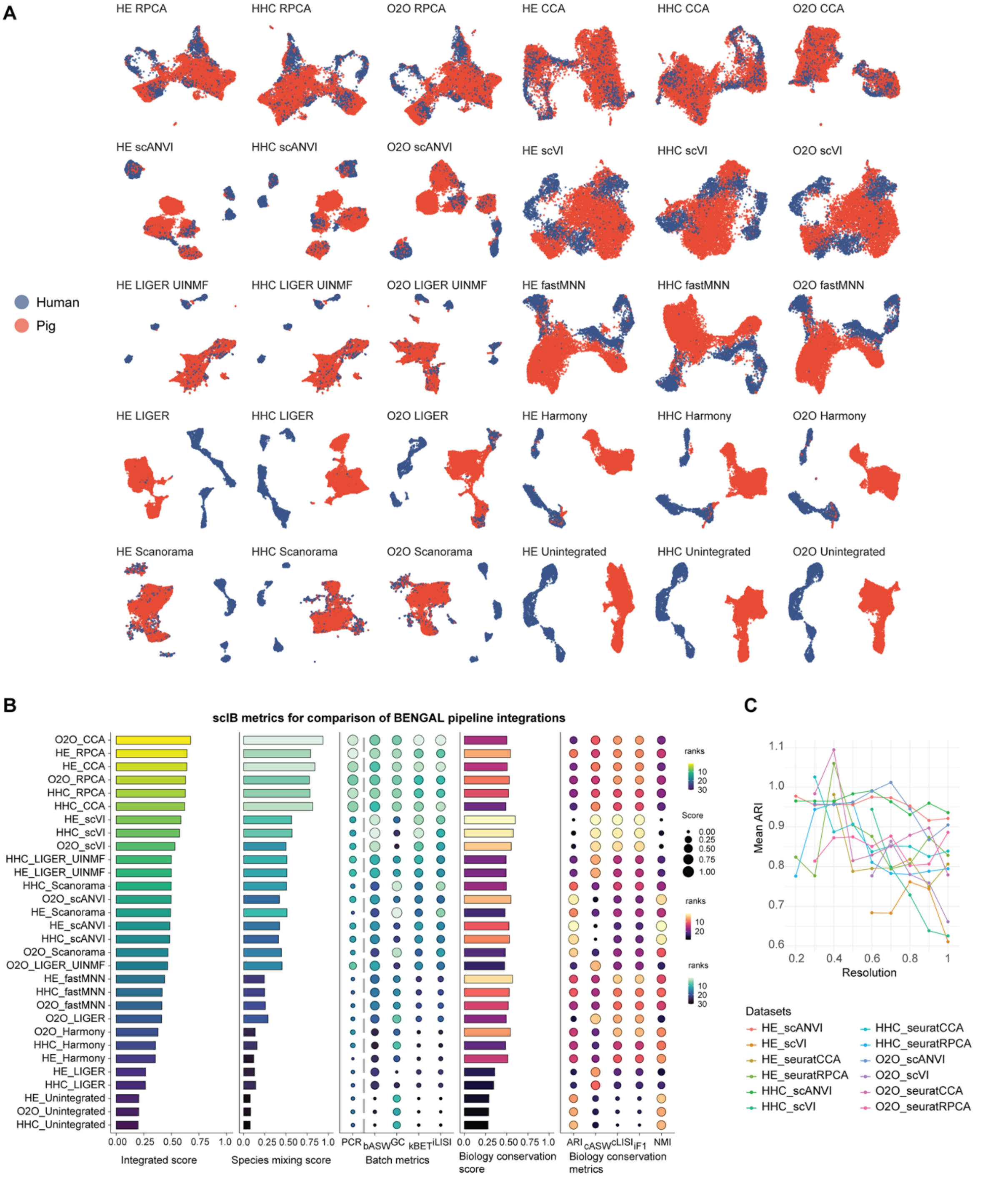


**Supplementary Figure 4. Coronary plaque scRNA-seq data integration and performance metrics. A,** Combined datasets produced by 9 integration algorithms (or without integration) and 3 methods for gene homology matching between pig and human coronary plaque datasets. **B,** scIB metrics (batch mixing and biology conservation) and their ranks assessing 'goodness' for integration of coronary artery plaque scRNAseq datasets. **C,** Mean diagonal Adjusted Rand index (ARI) produced by the bootstrapStability() function with 20 iterations for different clustering resolutions of the integrated datasets of coronary arteries.

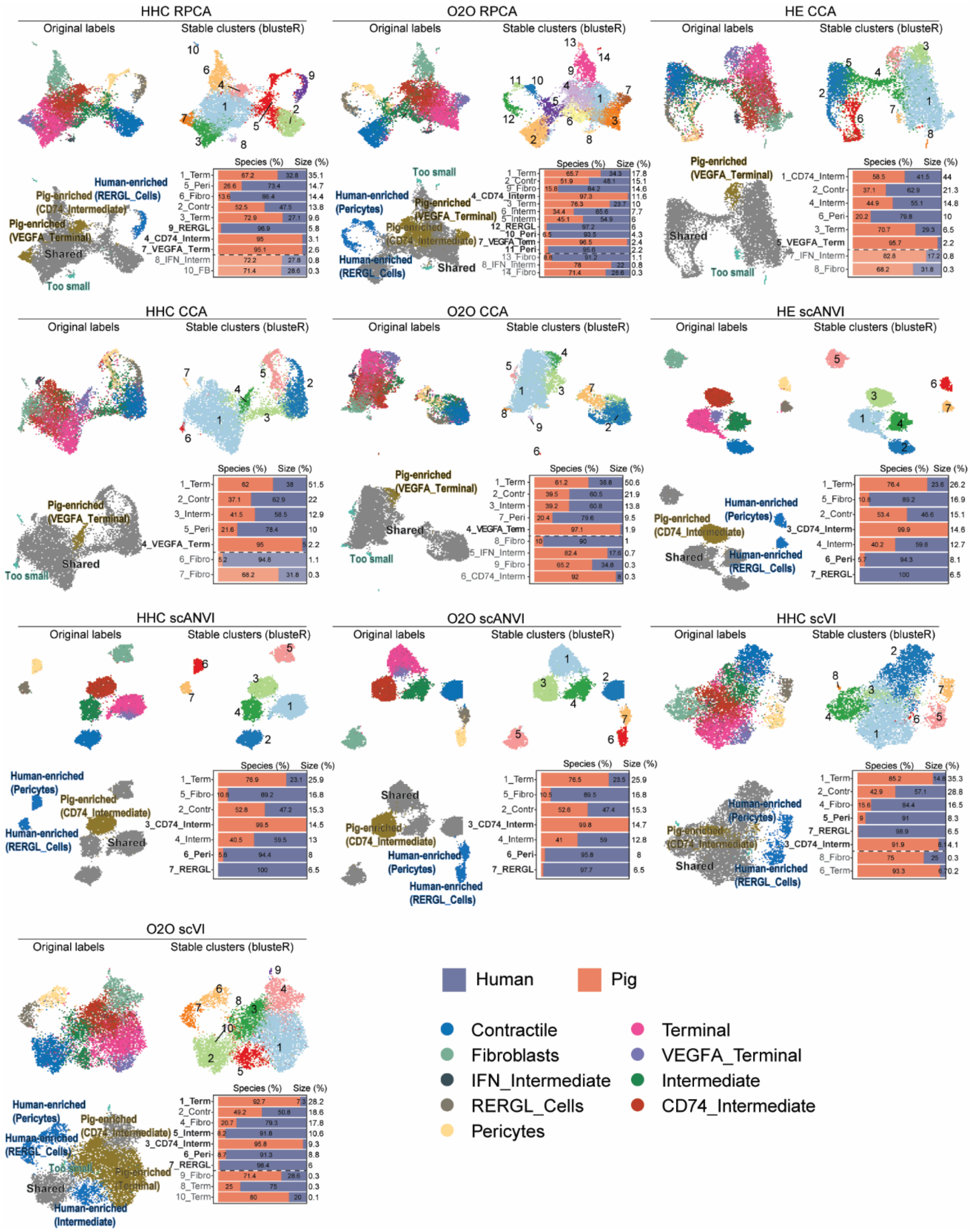


**Supplementary Figure 5. Identification of shared and species-specific mesenchymal plaque cell types in coronary atherosclerosis (additional data).** The ten additional integrations (to the two shown in Figure 6) produced by RPCA, CCA, scANVI, and scVI combined with the 3 gene homology match approaches. Each set of 4 plots consists of integrated dataset with cell type labels from the individual species data sets (top left); stable clusters defined using the *bluster* R package (top right); clusters fulfilling the criteria for being species-enriched (bottom left); and percentage contribution from each species to stable clusters (bottom right).. Fractions were calculated using a random subsample of 3200 cells from each species dataset, and one species were considered overrepresented in a cluster if contributing ≥90% of cells. Only clusters with more than 100 cells were analyzed for species contributions.

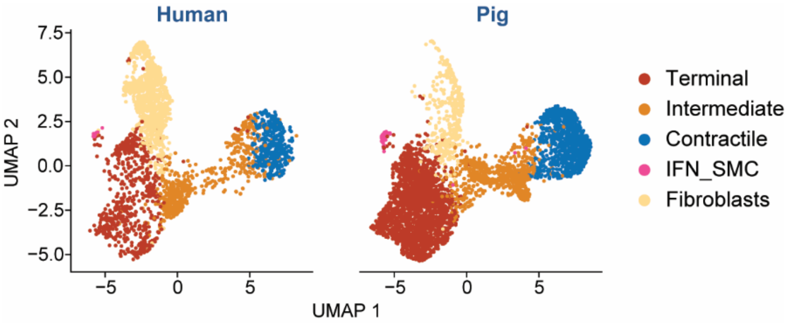


**Supplementary Figure 6.** UMAP representation of re-clustered species-shared SMC phenotypes of human and pig coronary arteries.

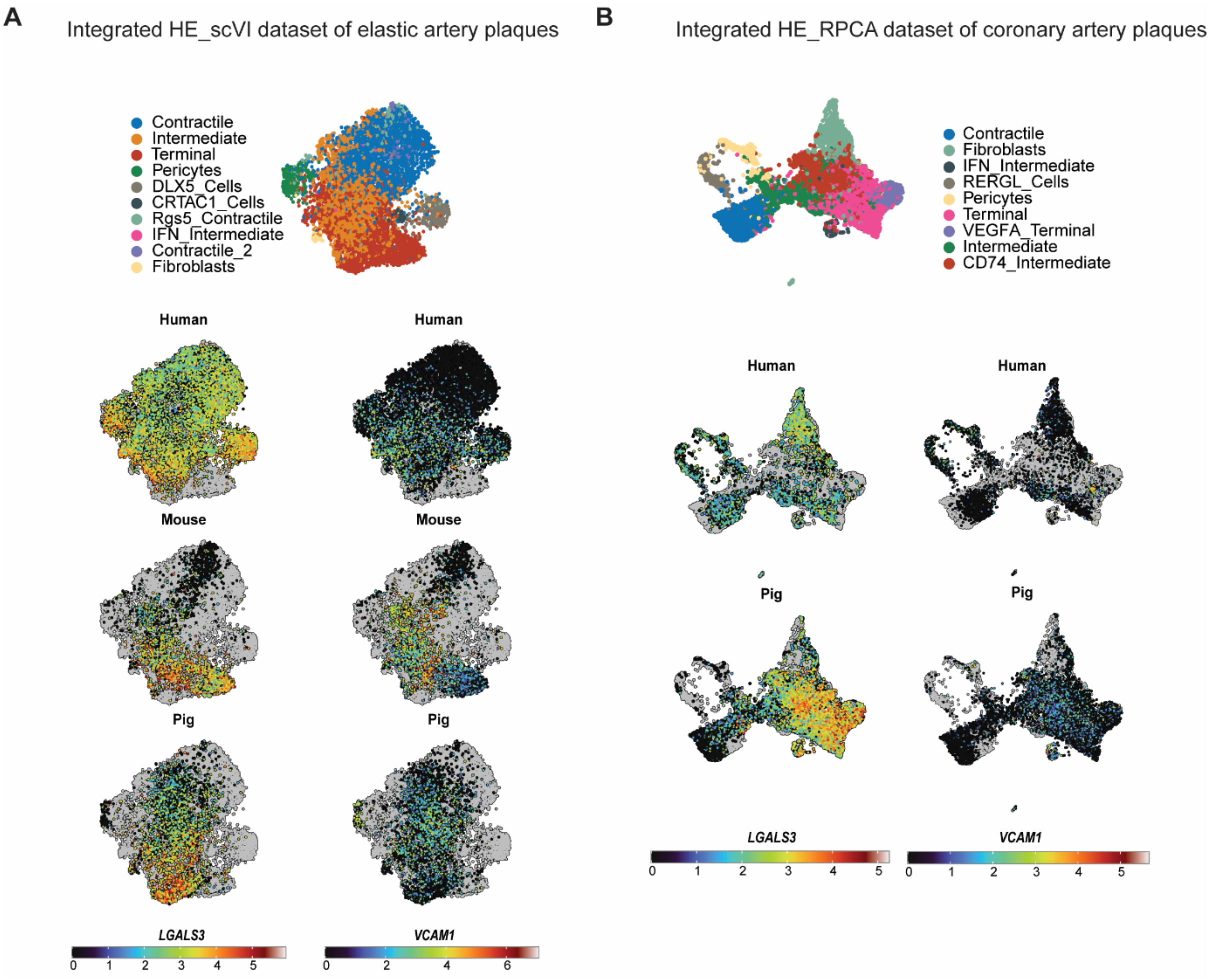


**Supplementary Figure 7.** Commonly used “transitional cell” markers visualized using integrated datasets of elastic artery plaques (mice, humans, pigs) (A) and coronary artery plaques (humans, pigs) (B).
