## Supplementary Tables 1-9 for "Mouse, pig, and human atherosclerotic lesions have common and distinct mesenchymal cell populations"

**Table S1. Datasets.**

List of publicly available and newly generated scRNA-seq and bulk RNA-seq datasets used for analysis in this paper .

| Reference | Species | Tissue | Type | Number of samples | Number of individuals | Number of males | GSE/E-MTAB | GSM |
|---|---|---|---|---|---|---|---|---|
| Wirka R. et al, 2019 (ref. 12) | Human | Right coronary artery segments with atherosclerosis in explanted hearts | scRNA-seq | 8 | 4 | 3 | GSE131778 | GSM3819856, GSM3819857, GSM3819858, GSM3819859, GSM3819860, GSM3819861, GSM3819862, GSM3819863 |
| This paper | PCSK9-transgenic pigs fed on high-fat diet for 15 months | Raised leions isolated from the right coronary artery | scRNA-seq | 5 | 5 | 1 | | |
| Alsaigh T. et al, 2022 (ref. 14) | Human | Carotid plaques from endarterectomies | scRNA-seq | 3 | 3 | 2 | GSE159677 | GSM4837523, GSM4837525, GSM4837527 |
| Pan H. et al., 2020 (ref. 13) | Human | Carotid plaques from endarterectomies | scRNA-seq | 3 | 3 | 2 | GSE155512 | GSM4705589, GSM4705590, GSM4705591 |
| Nogales et al. (ref. 21) | PCSK9-transgenic pigs fed on high-fat diet for 15 months | Raised lesions isolated from the distal abdominal aorta | scRNA-seq | 6 | 6 | 2 | E-MTAB-13898 | |
| Alencar GF. et al., 2020 (ref. 20) | Apoe-/- mice fed on high-fat diet for 18 weeks | Atherosclerotic plaques microdissected from the brachiocephalic artery | scRNA-seq | 8 | 15 | 15 | GSE150644 | GSM4555585, GSM4555586, GSM4555592, GSM4555593, GSM4555594, GSM4555595, GSM4555596, GSM4555597 |
| Mahmoud A.D., et al (ref. 43) | Human | Stable and unstable carotid plaques from endarterectomies | RNA-seq | 8 | 4 | | GSE120521 | GSM3402504, GSM3402505, GSM3402506, GSM3402507, GSM3402508, GSM3402509, GSM3402510, GSM3402511 |
| Cai Z., et al (ref. 45) | Wuzhishan miniature pigs fed on high-fat diet for 6 months | Atherosclerotic plaques isolated from the abdominal aorta | RNA-seq | 3 | 5 | 5 | GSE245530 | GSM7844585, GSM7844586, GSM7844587 |
| Örd T., et al. (ref. 50) | LDLR-/- ApoB100/100 mice fed high-fat diet for 3 months | Atherosclerotic plaques isolated from the aorta | RNA-seq | 5 | 5 | 4 | GSE205929 | GSM6235805, GSM6235806, GSM6235807, GSM6235808, GSM6235809 |

**Table S2. Marker genes for mesenchymal cell clusters in human carotid plaque scRNA-seq data. Corresponds to Fig. 1A.**
The listed genes have significantly higher expression in the indicated cluster than all other clusters combined.
Analysis was performed using the Wilcoxon test in Seurat v4.4.0 with P values adjusted for multiple testing by the Bonferroni method.

| cluster | gene | p_val | avg_log2F( | pct.1 | pct.2 | p_val_adj |
|---|---|---|---|---|---|---|
| Contractile | PLN | 0 | 1.846231 | 0.945 | 0.525 | 0 |
| Contractile | MYH11 | 0 | 1.845066 | 0.972 | 0.657 | 0 |
| Contractile | C11orf96 | 0 | 1.677769 | 0.976 | 0.821 | 0 |
| Contractile | CNN1 | 0 | 1.4203 | 0.829 | 0.353 | 0 |
| Contractile | C12orf75 | 0 | 1.389136 | 0.947 | 0.596 | 0 |
| Contractile | LMOD1 | 0 | 1.332413 | 0.932 | 0.619 | 0 |
| Contractile | CSRP2 | 0 | 1.332293 | 0.799 | 0.365 | 0 |
| Contractile | TPM2 | 0 | 1.279295 | 0.999 | 0.968 | 0 |
| Contractile | FLNA | 0 | 1.276862 | 0.957 | 0.77 | 0 |
| Contractile | PPP1R12B | 0 | 1.21016 | 0.825 | 0.446 | 0 |
| Contractile | FILIP1L | 0 | 1.196784 | 0.91 | 0.666 | 0 |
| Contractile | MCAM | 0 | 1.192363 | 0.788 | 0.438 | 0 |
| Contractile | SORBS2 | 0 | 1.173899 | 0.713 | 0.319 | 0 |
| Contractile | KCNMA1 | 0 | 1.159951 | 0.633 | 0.227 | 0 |
| Contractile | SUSD5 | 0 | 1.122489 | 0.612 | 0.232 | 0 |
| Contractile | MYLK | 0 | 1.119975 | 0.869 | 0.596 | 0 |
| Contractile | LPP | 0 | 1.113786 | 0.905 | 0.699 | 0 |
| Contractile | MYL9 | 0 | 1.108889 | 0.998 | 0.969 | 0 |
| Contractile | RERGL | 2.9095E-183 | 1.092689 | 0.303 | 0.065 | 5.5059E-179 |
| Contractile | MFAP4 | 0 | 1.089423 | 0.951 | 0.63 | 0 |
| Contractile | ITGA8 | 0 | 1.088678 | 0.938 | 0.647 | 0 |
| Contractile | LDOC1 | 2.0126E-303 | 1.069072 | 0.634 | 0.269 | 3.8086E-299 |
| Contractile | TNS1 | 0 | 1.068711 | 0.779 | 0.46 | 0 |
| Contractile | FILIP1 | 1.0422E-288 | 1.067537 | 0.694 | 0.346 | 1.9723E-284 |
| Contractile | SYNPO2 | 0 | 1.05931 | 0.775 | 0.43 | 0 |
| Contractile | DSTN | 0 | 1.041364 | 0.999 | 0.991 | 0 |
| Contractile | ACTC1 | 1.8349E-140 | 1.036683 | 0.372 | 0.133 | 3.4723E-136 |
| Contractile | HSPB7 | 7.6882E-266 | 1.025249 | 0.611 | 0.259 | 1.4549E-261 |
| Contractile | GRIA2 | 2.7856E-281 | 1.0188 | 0.419 | 0.091 | 5.2715E-277 |
| Contractile | MTRNR2L1 | 2.81939E-72 | 1.010402 | 0.836 | 0.795 | 5.33542E-68 |
| Contractile | CARMN | 1.6187E-231 | 0.993892 | 0.69 | 0.367 | 3.0631E-227 |
| Contractile | SLMAP | 1.3256E-270 | 0.9868 | 0.64 | 0.278 | 2.5085E-266 |
| Contractile | CSRP1 | 0 | 0.976665 | 0.948 | 0.812 | 0 |
| Contractile | PRUNE2 | 3.2057E-241 | 0.964278 | 0.578 | 0.252 | 6.0664E-237 |
| Contractile | PPP1R12A | 5.888E-261 | 0.954149 | 0.823 | 0.614 | 1.1142E-256 |
| Contractile | SVIL | 1.8487E-262 | 0.952493 | 0.873 | 0.672 | 3.4984E-258 |
| Contractile | FIBIN | 1.5612E-239 | 0.948577 | 0.664 | 0.333 | 2.9544E-235 |
| Contractile | SMTN | 1.5764E-228 | 0.946359 | 0.5 | 0.192 | 2.9832E-224 |
| Contractile | PALLD | 0 | 0.945169 | 0.904 | 0.742 | 0 |
| Contractile | SBSPON | 5.5333E-281 | 0.931352 | 0.4 | 0.079 | 1.0471E-276 |
| Contractile | ACTG2 | 3.607E-258 | 0.931196 | 0.559 | 0.189 | 6.8259E-254 |
| Contractile | PCDH7 | 7.7719E-261 | 0.925361 | 0.779 | 0.486 | 1.4707E-256 |
| Contractile | SCRG1 | 1.6692E-250 | 0.921978 | 0.621 | 0.269 | 3.1589E-246 |
| Contractile | SORBS1 | 1.3462E-236 | 0.921399 | 0.613 | 0.291 | 2.5476E-232 |
| Contractile | CALD1 | 0 | 0.897286 | 0.999 | 0.994 | 0 |
| Contractile | TCEAL2 | 4.3791E-262 | 0.887764 | 0.564 | 0.193 | 8.287E-258 |
| Contractile | TAGLN | 0 | 0.887382 | 0.999 | 0.973 | 0 |
| Contractile | RTN1 | 6.0284E-196 | 0.886592 | 0.393 | 0.114 | 1.1408E-191 |
| Contractile | TTLL7 | 7.5917E-231 | 0.873886 | 0.708 | 0.418 | 1.4366E-226 |
| Contractile | NEXN | 5.194E-277 | 0.857073 | 0.897 | 0.633 | 9.829E-273 |
| Contractile | NPNT | 4.4645E-232 | 0.852808 | 0.475 | 0.157 | 8.4487E-228 |
| Contractile | MAP3K20 | 1.933E-194 | 0.847078 | 0.594 | 0.313 | 3.658E-190 |
| Contractile | RAMP1 | 0 | 0.846094 | 0.93 | 0.543 | 0 |

| | | | | | | |
|---|---|---|---|---|---|---|
| Contractile | DES | 8.04443E-86 | 0.843079 | 0.152 | 0.033 | 1.52233E-81 |
| Contractile | MYOCD | 5.1945E-197 | 0.841996 | 0.397 | 0.121 | 9.8301E-193 |
| Contractile | NET1 | 5.6744E-170 | 0.808075 | 0.594 | 0.324 | 1.0738E-165 |
| Contractile | ADRA2C | 1.2202E-242 | 0.800733 | 0.294 | 0.035 | 2.3091E-238 |
| Contractile | HSPB6 | 6.1867E-169 | 0.800412 | 0.582 | 0.313 | 1.1708E-164 |
| Contractile | MYH10 | 5.3116E-228 | 0.795699 | 0.922 | 0.679 | 1.0052E-223 |
| Contractile | ACTA2 | 0 | 0.794951 | 0.995 | 0.828 | 0 |
| Contractile | TLN1 | 2.4114E-265 | 0.790986 | 0.921 | 0.801 | 4.5633E-261 |
| Contractile | SPEG | 1.0189E-201 | 0.78105 | 0.494 | 0.198 | 1.9283E-197 |
| Contractile | PART1 | 1.1123E-241 | 0.777523 | 0.276 | 0.027 | 2.105E-237 |
| Contractile | DACT3 | 3.8489E-192 | 0.771512 | 0.501 | 0.211 | 7.2837E-188 |
| Contractile | RGS5 | 3.0906E-291 | 0.75514 | 0.958 | 0.657 | 5.8487E-287 |
| Contractile | CPED1 | 6.7525E-166 | 0.75035 | 0.529 | 0.266 | 1.2779E-161 |
| Contractile | PPP1R14A | 4.193E-253 | 0.750146 | 0.962 | 0.768 | 7.9348E-249 |
| Contractile | ALDH1B1 | 1.4378E-171 | 0.710991 | 0.483 | 0.201 | 2.7209E-167 |
| Contractile | CRISPLD1 | 1.3708E-139 | 0.710406 | 0.534 | 0.282 | 2.5941E-135 |
| Contractile | CAMK2G | 8.559E-171 | 0.707784 | 0.457 | 0.189 | 1.6197E-166 |
| Contractile | CPE | 2.0879E-271 | 0.704417 | 0.958 | 0.648 | 3.9511E-267 |
| Contractile | SCX | 1.4375E-122 | 0.703207 | 0.343 | 0.133 | 2.7203E-118 |
| Contractile | FHL5 | 1.2598E-156 | 0.699968 | 0.386 | 0.131 | 2.3841E-152 |
| Contractile | MXI1 | 3.886E-157 | 0.697104 | 0.625 | 0.369 | 7.3538E-153 |
| Contractile | PRKG1 | 1.4585E-164 | 0.697034 | 0.68 | 0.428 | 2.7601E-160 |
| Contractile | FBXO32 | 3.7764E-105 | 0.689432 | 0.551 | 0.344 | 7.1465E-101 |
| Contractile | ACTN1 | 6.9421E-182 | 0.688 | 0.865 | 0.727 | 1.3137E-177 |
| Contractile | SPINT2 | 5.0304E-177 | 0.684214 | 0.816 | 0.558 | 9.5196E-173 |
| Contractile | MTRNR2L8 | 1.60505E-18 | 0.68229 | 0.477 | 0.441 | 3.0374E-14 |
| Contractile | NCS1 | 6.4118E-162 | 0.681167 | 0.413 | 0.163 | 1.2134E-157 |
| Contractile | A2M | 1.9634E-188 | 0.680835 | 0.956 | 0.841 | 3.7156E-184 |
| Contractile | ADSSL1 | 1.0094E-169 | 0.680217 | 0.282 | 0.061 | 1.9101E-165 |
| Contractile | RASSF3 | 3.7449E-131 | 0.679422 | 0.438 | 0.209 | 7.0868E-127 |
| Contractile | NIBAN1 | 7.126E-154 | 0.676972 | 0.742 | 0.522 | 1.3485E-149 |
| Contractile | SLC25A4 | 1.4991E-184 | 0.674675 | 0.82 | 0.592 | 2.8368E-180 |
| Contractile | HSPB8 | 1.2987E-104 | 0.666921 | 0.438 | 0.236 | 2.4576E-100 |
| Contractile | FOXN3 | 1.7565E-133 | 0.66595 | 0.477 | 0.242 | 3.324E-129 |
| Contractile | DMD | 9.7067E-129 | 0.665584 | 0.563 | 0.326 | 1.8369E-124 |
| Contractile | ST6GAL2 | 1.7069E-155 | 0.662333 | 0.331 | 0.106 | 3.2301E-151 |
| Contractile | PDLIM5 | 1.3364E-159 | 0.6573 | 0.762 | 0.558 | 2.529E-155 |
| Contractile | ZBTB16 | 3.4863E-111 | 0.650865 | 0.582 | 0.371 | 6.5974E-107 |
| Contractile | MRPS6 | 5.5335E-130 | 0.648022 | 0.614 | 0.393 | 1.0472E-125 |
| Contractile | MRAP2 | 2.0525E-139 | 0.64709 | 0.427 | 0.189 | 3.8842E-135 |
| Contractile | ANGPTL1 | 1.1225E-119 | 0.643173 | 0.307 | 0.11 | 2.1243E-115 |
| Contractile | HIST1H4C | 4.66426E-94 | 0.642617 | 0.716 | 0.605 | 8.82665E-90 |
| Contractile | RNF115 | 1.1273E-100 | 0.640778 | 0.565 | 0.392 | 2.13336E-96 |
| Contractile | SMOC2 | 3.6321E-154 | 0.637001 | 0.782 | 0.552 | 6.8733E-150 |
| Contractile | AKAP1 | 4.4744E-137 | 0.635465 | 0.363 | 0.138 | 8.4674E-133 |
| Contractile | MRVI1 | 3.2473E-128 | 0.634174 | 0.586 | 0.35 | 6.1451E-124 |
| Contractile | ZNF385D | 1.1638E-130 | 0.629569 | 0.451 | 0.213 | 2.2025E-126 |
| Contractile | SLC22A3 | 4.0088E-155 | 0.629436 | 0.335 | 0.103 | 7.5863E-151 |
| Contractile | MYL6 | 0 | 0.627423 | 0.997 | 0.994 | 0 |
| Contractile | CFL2 | 2.1054E-123 | 0.623089 | 0.638 | 0.43 | 3.9842E-119 |
| Contractile | RBPMS | 1.7539E-155 | 0.622699 | 0.764 | 0.56 | 3.319E-151 |
| Contractile | GASK1B | 5.9631E-121 | 0.615801 | 0.46 | 0.237 | 1.1285E-116 |
| Contractile | BTC | 1.6524E-177 | 0.613802 | 0.28 | 0.057 | 3.127E-173 |
| Contractile | HACD1 | 4.3321E-131 | 0.600554 | 0.464 | 0.222 | 8.198E-127 |
| Contractile | AOC3 | 4.9105E-115 | 0.599397 | 0.53 | 0.314 | 9.2926E-111 |
| Contractile | SNHG14 | 8.02484E-86 | 0.598001 | 0.479 | 0.295 | 1.51862E-81 |
| Contractile | ACTN4 | 5.0693E-148 | 0.597461 | 0.857 | 0.707 | 9.5931E-144 |
| Contractile | RAP1A | 2.2418E-112 | 0.594328 | 0.588 | 0.394 | 4.2424E-108 |

| | | | | | | |
|---|---|---|---|---|---|---|
| Contractile | SYNE2 | 2.5671E-106 | 0.593958 | 0.489 | 0.275 | 4.858E-102 |
| Contractile | SYNM | 3.1852E-105 | 0.592231 | 0.512 | 0.305 | 6.0277E-101 |
| Contractile | GSTP1 | 4.5856E-273 | 0.590167 | 0.988 | 0.974 | 8.6777E-269 |
| Contractile | FABP3 | 2.53E-158 | 0.588429 | 0.289 | 0.072 | 4.7878E-154 |
| Contractile | LDB3 | 3.1975E-153 | 0.581226 | 0.312 | 0.089 | 6.0509E-149 |
| Contractile | MRPL33 | 1.3768E-118 | 0.578687 | 0.787 | 0.681 | 2.6055E-114 |
| Contractile | TPM1 | 4.1137E-222 | 0.577931 | 0.99 | 0.953 | 7.7847E-218 |
| Contractile | MAP1B | 1.5002E-159 | 0.574047 | 0.948 | 0.878 | 2.8389E-155 |
| Contractile | DUSP26 | 8.9329E-136 | 0.571645 | 0.378 | 0.148 | 1.6905E-131 |
| Contractile | NPR1 | 2.0011E-130 | 0.570013 | 0.353 | 0.135 | 3.7869E-126 |
| Contractile | VCL | 5.753E-108 | 0.56896 | 0.724 | 0.561 | 1.0887E-103 |
| Contractile | KALRN | 3.05E-101 | 0.568115 | 0.382 | 0.189 | 5.77175E-97 |
| Contractile | NFIA | 2.9793E-126 | 0.565834 | 0.828 | 0.683 | 5.638E-122 |
| Contractile | MEF2A | 6.0381E-100 | 0.561064 | 0.62 | 0.434 | 1.14266E-95 |
| Contractile | MYEF2 | 7.5957E-125 | 0.56028 | 0.353 | 0.138 | 1.4374E-120 |
| Contractile | TWSG1 | 1.01361E-95 | 0.558503 | 0.504 | 0.321 | 1.91816E-91 |
| Contractile | PLS3 | 6.1571E-164 | 0.558374 | 0.93 | 0.853 | 1.1652E-159 |
| Contractile | GULP1 | 9.7541E-110 | 0.5578 | 0.533 | 0.316 | 1.8459E-105 |
| Contractile | RRAD | 5.87366E-88 | 0.557632 | 0.465 | 0.259 | 1.11153E-83 |
| Contractile | PLEKHO1 | 9.34537E-94 | 0.556051 | 0.502 | 0.31 | 1.76852E-89 |
| Contractile | MGST3 | 1.6627E-219 | 0.548967 | 0.974 | 0.951 | 3.1464E-215 |
| Contractile | TES | 1.27699E-92 | 0.54532 | 0.5 | 0.304 | 2.41658E-88 |
| Contractile | GMDS | 1.38909E-78 | 0.544793 | 0.642 | 0.492 | 2.62872E-74 |
| Contractile | KCNA5 | 4.0379E-167 | 0.543652 | 0.227 | 0.033 | 7.6413E-163 |
| Contractile | ADH5 | 7.7387E-109 | 0.541294 | 0.82 | 0.709 | 1.4645E-104 |
| Contractile | STK38L | 5.28733E-59 | 0.540689 | 0.541 | 0.403 | 1.00058E-54 |
| Contractile | ROCK1 | 1.37382E-84 | 0.54063 | 0.685 | 0.55 | 2.59981E-80 |
| Contractile | RHOB | 1.06863E-67 | 0.539153 | 0.732 | 0.598 | 2.02228E-63 |
| Contractile | BEX4 | 5.6313E-111 | 0.539013 | 0.564 | 0.334 | 1.0657E-106 |
| Contractile | COX7C | 9.1515E-235 | 0.538289 | 0.976 | 0.953 | 1.7318E-230 |
| Contractile | AC092164. | 2.9264E-99 | 0.537539 | 0.335 | 0.146 | 5.53799E-95 |
| Contractile | CCDC3 | 7.93805E-95 | 0.537529 | 0.626 | 0.429 | 1.5022E-90 |
| Contractile | NDUFA4 | 1.3893E-189 | 0.53593 | 0.971 | 0.92 | 2.6292E-185 |
| Contractile | PDLIM7 | 8.4689E-114 | 0.53358 | 0.783 | 0.605 | 1.6026E-109 |
| Contractile | SH3BGR | 6.50319E-82 | 0.530764 | 0.443 | 0.263 | 1.23066E-77 |
| Contractile | TBL1X | 7.6372E-104 | 0.530484 | 0.327 | 0.138 | 1.4453E-99 |
| Contractile | MXRA7 | 1.9016E-124 | 0.529451 | 0.822 | 0.711 | 3.5985E-120 |
| Contractile | PDE4DIP | 3.3041E-78 | 0.52238 | 0.51 | 0.341 | 6.25268E-74 |
| Contractile | ADI1 | 7.2889E-109 | 0.518454 | 0.826 | 0.724 | 1.3794E-104 |
| Contractile | ATP1A2 | 1.4257E-110 | 0.509648 | 0.236 | 0.065 | 2.698E-106 |
| Contractile | APLP2 | 3.966E-99 | 0.5088 | 0.836 | 0.721 | 7.50529E-95 |
| Contractile | SYNE1 | 1.73218E-73 | 0.507287 | 0.61 | 0.459 | 3.27799E-69 |
| Contractile | FERMT2 | 1.7274E-108 | 0.506883 | 0.781 | 0.646 | 3.2689E-104 |
| Contractile | ARID5A | 7.62748E-62 | 0.505616 | 0.456 | 0.306 | 1.44342E-57 |
| Contractile | SH3BGRL | 2.45E-141 | 0.503568 | 0.906 | 0.842 | 4.6363E-137 |
| Contractile | MSRB3 | 3.878E-103 | 0.50339 | 0.766 | 0.633 | 7.3387E-99 |
| Contractile | PPP1R3C | 3.69745E-72 | 0.5005 | 0.382 | 0.216 | 6.99706E-68 |
| Contractile | SIVA1 | 9.4955E-109 | 0.497806 | 0.833 | 0.739 | 1.7969E-104 |
| Contractile | TMEM47 | 3.10507E-88 | 0.497794 | 0.718 | 0.549 | 5.87603E-84 |
| Contractile | PPP1R1A | 1.112E-141 | 0.495968 | 0.195 | 0.028 | 2.1043E-137 |
| Contractile | ZEB1 | 5.29132E-78 | 0.494073 | 0.519 | 0.349 | 1.00133E-73 |
| Contractile | MEF2C | 2.6619E-120 | 0.492535 | 0.842 | 0.704 | 5.0375E-116 |
| Contractile | CCN5 | 5.83662E-59 | 0.491391 | 0.444 | 0.28 | 1.10452E-54 |
| Contractile | LDHB | 2.535E-105 | 0.490464 | 0.819 | 0.694 | 4.7972E-101 |
| Contractile | PHACTR1 | 2.3644E-118 | 0.488864 | 0.222 | 0.054 | 4.4743E-114 |
| Contractile | CNTN4 | 3.9347E-113 | 0.48349 | 0.234 | 0.064 | 7.446E-109 |
| Contractile | LBH | 5.37599E-88 | 0.480811 | 0.611 | 0.421 | 1.01735E-83 |
| Contractile | SPARCL1 | 1.3131E-143 | 0.480188 | 0.989 | 0.884 | 2.4849E-139 |

| | | | | | | |
|---|---|---|---|---|---|---|
| Contractile | INSR | 3.6076E-80 | 0.477508 | 0.303 | 0.139 | 6.82702E-76 |
| Contractile | SMDT1 | 8.29526E-84 | 0.474865 | 0.731 | 0.62 | 1.5698E-79 |
| Contractile | PPP1CB | 1.0341E-114 | 0.474061 | 0.895 | 0.799 | 1.9568E-110 |
| Contractile | FXYD1 | 1.086E-120 | 0.471715 | 0.919 | 0.821 | 2.0552E-116 |
| Contractile | KANK2 | 1.56699E-68 | 0.470867 | 0.637 | 0.497 | 2.96538E-64 |
| Contractile | TGFB1I1 | 5.795E-110 | 0.466186 | 0.854 | 0.756 | 1.0967E-105 |
| Contractile | CCNI | 4.9376E-144 | 0.466172 | 0.965 | 0.934 | 9.3438E-140 |
| Contractile | APP | 1.85376E-67 | 0.465483 | 0.67 | 0.561 | 3.50806E-63 |
| Contractile | PHGDH | 1.19964E-70 | 0.46238 | 0.335 | 0.172 | 2.2702E-66 |
| Contractile | LIMCH1 | 3.40523E-71 | 0.462001 | 0.681 | 0.544 | 6.44406E-67 |
| Contractile | PRDM16-D | 6.41609E-92 | 0.459438 | 0.303 | 0.127 | 1.21418E-87 |
| Contractile | SGCA | 7.54783E-76 | 0.459388 | 0.658 | 0.489 | 1.42835E-71 |
| Contractile | KANK1 | 6.51385E-78 | 0.455111 | 0.385 | 0.208 | 1.23268E-73 |
| Contractile | MFGE8 | 7.1052E-144 | 0.455086 | 0.97 | 0.92 | 1.3446E-139 |
| Contractile | LINC02269 | 9.28659E-98 | 0.454106 | 0.199 | 0.052 | 1.7574E-93 |
| Contractile | MAP4 | 1.92696E-72 | 0.454017 | 0.755 | 0.639 | 3.64658E-68 |
| Contractile | PPP2R2B | 2.8541E-111 | 0.453408 | 0.252 | 0.076 | 5.4012E-107 |
| Contractile | IQGAP1 | 2.17194E-70 | 0.45079 | 0.681 | 0.56 | 4.11017E-66 |
| Contractile | PITPNB | 6.18339E-64 | 0.450612 | 0.523 | 0.377 | 1.17014E-59 |
| Contractile | SELENOM | 3.3808E-158 | 0.450078 | 0.985 | 0.977 | 6.3978E-154 |
| Contractile | KTN1 | 2.9623E-104 | 0.447643 | 0.88 | 0.811 | 5.6058E-100 |
| Contractile | PDLIM3 | 2.68426E-93 | 0.445373 | 0.905 | 0.828 | 5.0797E-89 |
| Contractile | MPP7 | 2.3189E-103 | 0.442345 | 0.256 | 0.084 | 4.3882E-99 |
| Contractile | PDE4D | 4.51703E-95 | 0.440813 | 0.172 | 0.039 | 8.54802E-91 |
| Contractile | TSPAN2 | 2.67189E-75 | 0.439658 | 0.474 | 0.292 | 5.05628E-71 |
| Contractile | ITGA7 | 2.06257E-75 | 0.439311 | 0.344 | 0.173 | 3.90322E-71 |
| Contractile | TC2N | 6.52281E-80 | 0.438982 | 0.255 | 0.101 | 1.23438E-75 |
| Contractile | MACF1 | 5.59073E-47 | 0.437993 | 0.638 | 0.543 | 1.05799E-42 |
| Contractile | ACADL | 1.9049E-112 | 0.436632 | 0.179 | 0.034 | 3.6049E-108 |
| Contractile | JAK2 | 1.74522E-75 | 0.436182 | 0.296 | 0.138 | 3.30266E-71 |
| Contractile | PGRMC1 | 2.25426E-77 | 0.435767 | 0.739 | 0.624 | 4.26596E-73 |
| Contractile | SUN1 | 1.63417E-62 | 0.435643 | 0.39 | 0.236 | 3.09251E-58 |
| Contractile | WDR1 | 4.19542E-68 | 0.432428 | 0.601 | 0.449 | 7.93941E-64 |
| Contractile | LIMS2 | 1.46374E-71 | 0.430714 | 0.483 | 0.309 | 2.76999E-67 |
| Contractile | ACTB | 8.4572E-207 | 0.429565 | 0.999 | 0.997 | 1.6004E-202 |
| Contractile | KCNAB1 | 2.37634E-68 | 0.428559 | 0.238 | 0.101 | 4.49698E-64 |
| Contractile | CCDC6 | 6.91868E-59 | 0.428054 | 0.444 | 0.297 | 1.30929E-54 |
| Contractile | RIMKLB | 9.02964E-80 | 0.426145 | 0.321 | 0.154 | 1.70877E-75 |
| Contractile | RSL24D1 | 2.27421E-77 | 0.425062 | 0.813 | 0.732 | 4.30372E-73 |
| Contractile | LINC00632 | 3.00442E-43 | 0.42443 | 0.306 | 0.189 | 5.68557E-39 |
| Contractile | RCAN2 | 7.89534E-70 | 0.424357 | 0.713 | 0.551 | 1.49412E-65 |
| Contractile | DST | 3.43872E-51 | 0.424242 | 0.753 | 0.682 | 6.50744E-47 |
| Contractile | LPAR1 | 8.82388E-97 | 0.420718 | 0.228 | 0.071 | 1.66983E-92 |
| Contractile | SNHG6 | 3.06605E-68 | 0.419527 | 0.785 | 0.693 | 5.80219E-64 |
| Contractile | SORT1 | 2.72853E-59 | 0.418129 | 0.466 | 0.314 | 5.16348E-55 |
| Contractile | CRNDE | 9.36581E-59 | 0.416911 | 0.565 | 0.43 | 1.77239E-54 |
| Contractile | NTRK3 | 1.60878E-64 | 0.416102 | 0.319 | 0.164 | 3.04446E-60 |
| Contractile | ALDH2 | 2.77449E-59 | 0.41557 | 0.689 | 0.569 | 5.25045E-55 |
| Contractile | SELENOW | 2.0089E-131 | 0.413957 | 0.975 | 0.963 | 3.8015E-127 |
| Contractile | MZT2B | 5.8098E-102 | 0.413366 | 0.911 | 0.873 | 1.09945E-97 |
| Contractile | CKMT2 | 4.13586E-97 | 0.4125 | 0.215 | 0.062 | 7.8267E-93 |
| Contractile | PDK3 | 7.15865E-57 | 0.412434 | 0.377 | 0.235 | 1.3547E-52 |
| Contractile | MAOB | 6.98163E-69 | 0.41231 | 0.271 | 0.126 | 1.3212E-64 |
| Contractile | GYPC | 3.02269E-70 | 0.411509 | 0.738 | 0.602 | 5.72013E-66 |
| Contractile | MYLIP | 8.54298E-45 | 0.411325 | 0.417 | 0.289 | 1.61667E-40 |
| Contractile | PAM | 9.91099E-57 | 0.408144 | 0.571 | 0.435 | 1.87556E-52 |
| Contractile | ADCY5 | 7.6676E-102 | 0.407593 | 0.189 | 0.044 | 1.45102E-97 |
| Contractile | COPRS | 1.09124E-57 | 0.407381 | 0.744 | 0.66 | 2.06507E-53 |

| | | | | | | |
|---|---|---|---|---|---|---|
| Contractile | FOXC1 | 2.93865E-56 | 0.406665 | 0.741 | 0.635 | 5.5611E-52 |
| Contractile | FBLIM1 | 1.37464E-76 | 0.406429 | 0.822 | 0.705 | 2.60136E-72 |
| Contractile | ENPP1 | 2.14469E-37 | 0.403719 | 0.298 | 0.186 | 4.0586E-33 |
| Contractile | RYR2 | 1.3146E-115 | 0.402075 | 0.172 | 0.028 | 2.4878E-111 |
| Contractile | CLIC4 | 5.65589E-57 | 0.401798 | 0.73 | 0.633 | 1.07032E-52 |
| Contractile | CRIM1 | 9.05413E-55 | 0.401098 | 0.668 | 0.533 | 1.7134E-50 |
| Contractile | COX6C | 1.6613E-118 | 0.40102 | 0.93 | 0.914 | 3.1438E-114 |
| Contractile | BAG2 | 1.49918E-42 | 0.400679 | 0.431 | 0.308 | 2.83705E-38 |
| Contractile | TCAP | 8.82078E-90 | 0.400184 | 0.108 | 0.009 | 1.66924E-85 |
| Contractile | TMEM106 | 1.38667E-63 | 0.399276 | 0.574 | 0.417 | 2.62414E-59 |
| Contractile | C5orf24 | 4.69372E-45 | 0.398483 | 0.551 | 0.448 | 8.88239E-41 |
| Contractile | CAV2 | 6.80585E-80 | 0.398428 | 0.82 | 0.704 | 1.28794E-75 |
| Contractile | KCNMB1 | 1.90817E-55 | 0.397457 | 0.562 | 0.418 | 3.61102E-51 |
| Contractile | HADH | 3.14798E-53 | 0.397373 | 0.325 | 0.19 | 5.95724E-49 |
| Contractile | DAPK3 | 3.32771E-59 | 0.39689 | 0.398 | 0.247 | 6.29737E-55 |
| Contractile | DANCR | 7.6314E-45 | 0.396827 | 0.423 | 0.299 | 1.44417E-40 |
| Contractile | ROCK2 | 1.13849E-48 | 0.396431 | 0.617 | 0.511 | 2.15448E-44 |
| Contractile | CALM3 | 1.90158E-52 | 0.395795 | 0.734 | 0.64 | 3.59855E-48 |
| Contractile | ST13 | 5.08128E-76 | 0.395456 | 0.853 | 0.781 | 9.61581E-72 |
| Contractile | SNRPN | 1.7071E-65 | 0.395067 | 0.798 | 0.717 | 3.23051E-61 |
| Contractile | EFHD1 | 1.24594E-65 | 0.394331 | 0.634 | 0.471 | 2.35782E-61 |
| Contractile | RASL12 | 3.92648E-61 | 0.393115 | 0.385 | 0.227 | 7.43047E-57 |
| Contractile | SOD1 | 8.7123E-119 | 0.393008 | 0.949 | 0.924 | 1.6487E-114 |
| Contractile | EMP2 | 4.59656E-76 | 0.392689 | 0.842 | 0.76 | 8.69853E-72 |
| Contractile | SUN2 | 6.99628E-49 | 0.391573 | 0.526 | 0.397 | 1.32398E-44 |
| Contractile | SESTD1 | 2.42502E-49 | 0.390478 | 0.381 | 0.249 | 4.5891E-45 |
| Contractile | RERG | 9.67612E-47 | 0.390228 | 0.436 | 0.302 | 1.83111E-42 |
| Contractile | LINC00844 | 8.63897E-74 | 0.389926 | 0.16 | 0.044 | 1.63484E-69 |
| Contractile | RBMS3 | 9.20916E-40 | 0.388608 | 0.501 | 0.388 | 1.74274E-35 |
| Contractile | SLC20A2 | 3.46729E-60 | 0.387622 | 0.288 | 0.148 | 6.56151E-56 |
| Contractile | PGM5 | 3.21373E-64 | 0.386937 | 0.292 | 0.146 | 6.08166E-60 |
| Contractile | PLCG2 | 2.27756E-21 | 0.385547 | 0.612 | 0.588 | 4.31006E-17 |
| Contractile | PNISR | 3.34467E-56 | 0.385355 | 0.772 | 0.678 | 6.32945E-52 |
| Contractile | AOPEP | 3.32044E-73 | 0.385026 | 0.821 | 0.729 | 6.28361E-69 |
| Contractile | ZC3H13 | 2.10357E-42 | 0.385009 | 0.573 | 0.466 | 3.98079E-38 |
| Contractile | SLC5A3 | 4.92839E-59 | 0.38473 | 0.203 | 0.084 | 9.32648E-55 |
| Contractile | PPM1K | 2.31915E-45 | 0.383996 | 0.345 | 0.221 | 4.38876E-41 |
| Contractile | CRTAP | 1.2403E-60 | 0.382781 | 0.732 | 0.631 | 2.34714E-56 |
| Contractile | AL627171. | 1.47226E-47 | 0.382695 | 0.282 | 0.157 | 2.78611E-43 |
| Contractile | SAMD4A | 4.30514E-50 | 0.382193 | 0.267 | 0.143 | 8.14705E-46 |
| Contractile | RBP7 | 1.91558E-93 | 0.381481 | 0.154 | 0.029 | 3.62505E-89 |
| Contractile | ENAH | 4.83343E-42 | 0.380951 | 0.616 | 0.515 | 9.14678E-38 |
| Contractile | HSPB1 | 6.07E-123 | 0.380834 | 0.994 | 0.975 | 1.1487E-118 |
| Contractile | NBEAL1 | 4.95366E-40 | 0.380252 | 0.503 | 0.397 | 9.3743E-36 |
| Contractile | FAU | 6.1605E-180 | 0.379085 | 0.995 | 0.997 | 1.1658E-175 |
| Contractile | MYH9 | 4.44453E-64 | 0.379044 | 0.842 | 0.754 | 8.41082E-60 |
| Contractile | BRD7 | 3.89718E-47 | 0.375861 | 0.589 | 0.475 | 7.37502E-43 |
| Contractile | IRS2 | 1.27458E-65 | 0.375854 | 0.247 | 0.11 | 2.41201E-61 |
| Contractile | PTGER3 | 5.40441E-68 | 0.374641 | 0.295 | 0.141 | 1.02273E-63 |
| Contractile | TMEM159 | 4.32519E-35 | 0.371062 | 0.457 | 0.351 | 8.18499E-31 |
| Contractile | UBE2H | 1.22627E-41 | 0.370164 | 0.569 | 0.472 | 2.32059E-37 |
| Contractile | PJA2 | 3.38784E-52 | 0.3688 | 0.699 | 0.606 | 6.41114E-48 |
| Contractile | TRIM36 | 3.26479E-71 | 0.367821 | 0.212 | 0.078 | 6.17828E-67 |
| Contractile | KIF1B | 3.56927E-47 | 0.365703 | 0.274 | 0.152 | 6.75448E-43 |
| Contractile | NPM1 | 5.45926E-88 | 0.365374 | 0.948 | 0.919 | 1.03311E-83 |
| Contractile | THRB | 2.24429E-49 | 0.365095 | 0.32 | 0.188 | 4.24709E-45 |
| Contractile | LINC00702 | 4.73993E-38 | 0.365051 | 0.386 | 0.267 | 8.96984E-34 |
| Contractile | SERBP1 | 3.33423E-65 | 0.364561 | 0.831 | 0.757 | 6.3097E-61 |

| | | | | | | |
|---|---|---|---|---|---|---|
| Contractile | SLC4A7 | 8.51765E-43 | 0.363648 | 0.294 | 0.177 | 1.61188E-38 |
| Contractile | UNC13C | 1.3123E-103 | 0.363206 | 0.132 | 0.014 | 2.4834E-99 |
| Contractile | CAV1 | 4.02942E-86 | 0.362562 | 0.957 | 0.896 | 7.62528E-82 |
| Contractile | HNRNPA1 | 2.53118E-74 | 0.362456 | 0.908 | 0.864 | 4.79E-70 |
| Contractile | ALCAM | 7.47846E-47 | 0.362398 | 0.529 | 0.387 | 1.41522E-42 |
| Contractile | HRCT1 | 3.60666E-34 | 0.361944 | 0.282 | 0.18 | 6.82524E-30 |
| Contractile | GUK1 | 6.61039E-89 | 0.361085 | 0.939 | 0.917 | 1.25095E-84 |
| Contractile | YAP1 | 1.21553E-37 | 0.361019 | 0.458 | 0.347 | 2.30027E-33 |
| Contractile | EIF4B | 1.73846E-50 | 0.360921 | 0.737 | 0.632 | 3.28985E-46 |
| Contractile | PRR34-AS1 | 3.15243E-41 | 0.359668 | 0.329 | 0.209 | 5.96566E-37 |
| Contractile | AKAP6 | 1.90772E-44 | 0.359571 | 0.26 | 0.144 | 3.61017E-40 |
| Contractile | BEX5 | 1.70156E-68 | 0.357822 | 0.219 | 0.085 | 3.22003E-64 |
| Contractile | SORL1 | 1.1482E-103 | 0.356934 | 0.158 | 0.026 | 2.1729E-99 |
| Contractile | PGM1 | 7.20026E-50 | 0.356696 | 0.326 | 0.195 | 1.36258E-45 |
| Contractile | PCDH11X | 4.09502E-89 | 0.35648 | 0.12 | 0.015 | 7.74941E-85 |
| Contractile | UQCRB | 5.53188E-92 | 0.354802 | 0.957 | 0.929 | 1.04685E-87 |
| Contractile | ATRX | 1.78144E-41 | 0.354012 | 0.679 | 0.588 | 3.37119E-37 |
| Contractile | SNTA1 | 9.32197E-46 | 0.353798 | 0.462 | 0.333 | 1.76409E-41 |
| Contractile | MRGPRF | 1.13681E-39 | 0.353742 | 0.426 | 0.305 | 2.15129E-35 |
| Contractile | CD44 | 6.71534E-30 | 0.353293 | 0.555 | 0.472 | 1.27081E-25 |
| Contractile | LGI2 | 8.01146E-70 | 0.351772 | 0.17 | 0.052 | 1.51609E-65 |
| Contractile | HINT1 | 3.2378E-102 | 0.351663 | 0.935 | 0.905 | 6.12718E-98 |
| Contractile | TXNRD1 | 1.19149E-21 | 0.3505 | 0.363 | 0.287 | 2.25477E-17 |
| Contractile | COX20 | 2.74985E-40 | 0.350094 | 0.63 | 0.544 | 5.20382E-36 |
| Contractile | SEMA3D | 3.19331E-77 | 0.34973 | 0.138 | 0.029 | 6.04302E-73 |
| Contractile | PDZRN4 | 9.55259E-71 | 0.349491 | 0.155 | 0.043 | 1.80773E-66 |
| Contractile | PBXIP1 | 2.14719E-39 | 0.348296 | 0.419 | 0.301 | 4.06333E-35 |
| Contractile | SNHG8 | 1.60553E-39 | 0.347803 | 0.686 | 0.609 | 3.03831E-35 |
| Contractile | ANAPC16 | 3.77356E-64 | 0.347471 | 0.836 | 0.779 | 7.14109E-60 |
| Contractile | CCDC88A | 2.60423E-34 | 0.34722 | 0.346 | 0.241 | 4.92824E-30 |
| Contractile | EPB41L4A- | 1.84958E-36 | 0.346952 | 0.506 | 0.391 | 3.50015E-32 |
| Contractile | CASQ2 | 5.02778E-84 | 0.345641 | 0.145 | 0.028 | 9.51456E-80 |
| Contractile | NDUFB7 | 6.12172E-74 | 0.345306 | 0.883 | 0.825 | 1.15847E-69 |
| Contractile | KLF15 | 1.76997E-74 | 0.345014 | 0.19 | 0.062 | 3.3495E-70 |
| Contractile | EIF5B | 4.41259E-46 | 0.344706 | 0.73 | 0.657 | 8.35038E-42 |
| Contractile | MPP6 | 2.83014E-77 | 0.344636 | 0.171 | 0.048 | 5.35575E-73 |
| Contractile | SMIM10 | 9.81696E-38 | 0.343938 | 0.491 | 0.374 | 1.85776E-33 |
| Contractile | PDZRN3 | 1.0545E-72 | 0.343889 | 0.205 | 0.072 | 1.99554E-68 |
| Contractile | NCKAP1 | 3.371E-39 | 0.342845 | 0.464 | 0.349 | 6.37928E-35 |
| Contractile | CSDE1 | 1.93937E-50 | 0.341815 | 0.776 | 0.705 | 3.67006E-46 |
| Contractile | VEZT | 1.88321E-41 | 0.341518 | 0.324 | 0.208 | 3.56379E-37 |
| Contractile | LGR6 | 3.30732E-74 | 0.341505 | 0.181 | 0.056 | 6.25877E-70 |
| Contractile | AP003548. | 3.20947E-68 | 0.341094 | 0.162 | 0.049 | 6.07359E-64 |
| Contractile | TNRC6A | 2.06936E-36 | 0.340219 | 0.443 | 0.326 | 3.91606E-32 |
| Contractile | ITM2C | 2.83957E-23 | 0.339124 | 0.469 | 0.389 | 5.37361E-19 |
| Contractile | EIF3A | 2.01756E-30 | 0.339064 | 0.539 | 0.456 | 3.81803E-26 |
| Contractile | ATL3 | 1.05473E-33 | 0.338668 | 0.482 | 0.382 | 1.99597E-29 |
| Contractile | RALY | 8.51119E-40 | 0.337615 | 0.56 | 0.459 | 1.61066E-35 |
| Contractile | CAVIN1 | 1.5583E-72 | 0.337592 | 0.913 | 0.857 | 2.94892E-68 |
| Contractile | PDE8B | 3.11098E-66 | 0.33656 | 0.179 | 0.06 | 5.88722E-62 |
| Contractile | RNF150 | 1.38621E-60 | 0.33639 | 0.161 | 0.054 | 2.62326E-56 |
| Contractile | MZT2A | 2.65623E-38 | 0.335307 | 0.546 | 0.435 | 5.02664E-34 |
| Contractile | ZFYVE21 | 1.74068E-40 | 0.335219 | 0.601 | 0.503 | 3.29407E-36 |
| Contractile | COX14 | 8.01148E-47 | 0.333598 | 0.668 | 0.569 | 1.51609E-42 |
| Contractile | FRK | 5.1654E-69 | 0.332986 | 0.184 | 0.062 | 9.775E-65 |
| Contractile | HSPA2 | 1.61354E-33 | 0.332172 | 0.327 | 0.22 | 3.05347E-29 |
| Contractile | GLS | 2.19276E-32 | 0.331476 | 0.592 | 0.509 | 4.14958E-28 |
| Contractile | ITIH5 | 1.12762E-41 | 0.331357 | 0.328 | 0.199 | 2.13392E-37 |

| | | | | | | |
|---|---|---|---|---|---|---|
| Contractile | ATP2A2 | 3.24797E-33 | 0.331265 | 0.381 | 0.274 | 6.14645E-29 |
| Contractile | MYOM1 | 1.39088E-78 | 0.330927 | 0.175 | 0.049 | 2.6321E-74 |
| Contractile | PRMT2 | 3.47254E-37 | 0.330631 | 0.646 | 0.567 | 6.57144E-33 |
| Contractile | CAPZA2 | 3.85747E-43 | 0.32915 | 0.683 | 0.599 | 7.29987E-39 |
| Contractile | KCTD10 | 3.16216E-39 | 0.328583 | 0.304 | 0.193 | 5.98407E-35 |
| Contractile | FRYL | 1.18114E-37 | 0.328525 | 0.31 | 0.196 | 2.23519E-33 |
| Contractile | SELENOK | 1.23977E-50 | 0.328221 | 0.757 | 0.703 | 2.34615E-46 |
| Contractile | TPH1 | 4.68567E-39 | 0.327975 | 0.261 | 0.148 | 8.86715E-35 |
| Contractile | KIAA0232 | 3.68731E-39 | 0.32672 | 0.343 | 0.227 | 6.97786E-35 |
| Contractile | PMEPA1 | 1.77043E-55 | 0.32579 | 0.782 | 0.688 | 3.35037E-51 |
| Contractile | GSG1L | 8.75644E-77 | 0.325442 | 0.178 | 0.052 | 1.65707E-72 |
| Contractile | C14orf132 | 2.31607E-60 | 0.325243 | 0.209 | 0.086 | 4.38294E-56 |
| Contractile | CAP1 | 1.33252E-42 | 0.325104 | 0.733 | 0.662 | 2.52166E-38 |
| Contractile | PTPN11 | 7.33408E-33 | 0.324843 | 0.444 | 0.341 | 1.3879E-28 |
| Contractile | ILK | 2.07922E-60 | 0.324723 | 0.829 | 0.741 | 3.93471E-56 |
| Contractile | CCDC107 | 5.40382E-50 | 0.324376 | 0.779 | 0.7 | 1.02262E-45 |
| Contractile | PLAAT3 | 4.7122E-31 | 0.323934 | 0.529 | 0.431 | 8.91736E-27 |
| Contractile | CFDP1 | 4.44143E-31 | 0.323561 | 0.56 | 0.464 | 8.40496E-27 |
| Contractile | CAMTA1 | 1.74073E-49 | 0.322502 | 0.753 | 0.686 | 3.29416E-45 |
| Contractile | PERP | 1.2561E-42 | 0.321792 | 0.304 | 0.182 | 2.37705E-38 |
| Contractile | PSIP1 | 1.38529E-42 | 0.32083 | 0.732 | 0.631 | 2.62152E-38 |
| Contractile | FTL | 7.81526E-72 | 0.319416 | 0.997 | 1 | 1.47896E-67 |
| Contractile | EIF3L | 5.56532E-31 | 0.31824 | 0.643 | 0.567 | 1.05318E-26 |
| Contractile | LMCD1 | 5.45832E-35 | 0.317111 | 0.685 | 0.59 | 1.03293E-30 |
| Contractile | WFS1 | 2.53319E-42 | 0.316001 | 0.241 | 0.131 | 4.79381E-38 |
| Contractile | SQSTM1 | 1.50698E-45 | 0.314814 | 0.859 | 0.821 | 2.85181E-41 |
| Contractile | TBC1D1 | 4.03316E-32 | 0.314087 | 0.382 | 0.277 | 7.63235E-28 |
| Contractile | KMT2A | 1.24311E-27 | 0.31387 | 0.575 | 0.496 | 2.35246E-23 |
| Contractile | MAOA | 9.01802E-36 | 0.313208 | 0.258 | 0.154 | 1.70657E-31 |
| Contractile | GLUL | 2.06818E-16 | 0.312973 | 0.374 | 0.311 | 3.91382E-12 |
| Contractile | PFN2 | 3.68098E-40 | 0.312711 | 0.597 | 0.475 | 6.96589E-36 |
| Contractile | CHMP3 | 8.29103E-34 | 0.312585 | 0.634 | 0.562 | 1.56899E-29 |
| Contractile | KIF1C | 2.00886E-42 | 0.311937 | 0.288 | 0.169 | 3.80156E-38 |
| Contractile | PTPRD | 1.7574E-40 | 0.311287 | 0.18 | 0.083 | 3.3257E-36 |
| Contractile | NQO1 | 7.4975E-13 | 0.311084 | 0.213 | 0.16 | 1.41883E-08 |
| Contractile | BCAM | 7.79987E-36 | 0.310132 | 0.438 | 0.311 | 1.47605E-31 |
| Contractile | ARPP19 | 8.52053E-34 | 0.310129 | 0.459 | 0.356 | 1.61243E-29 |
| Contractile | NXPH3 | 7.26849E-36 | 0.309073 | 0.258 | 0.154 | 1.37549E-31 |
| Contractile | LITAF | 8.14545E-35 | 0.307588 | 0.585 | 0.491 | 1.54144E-30 |
| Contractile | MAF1 | 5.90488E-38 | 0.307459 | 0.629 | 0.535 | 1.11744E-33 |
| Contractile | SAMMSON | 8.9066E-69 | 0.306232 | 0.121 | 0.025 | 1.68549E-64 |
| Contractile | ERBB4 | 1.53885E-44 | 0.305094 | 0.179 | 0.08 | 2.91212E-40 |
| Contractile | MSN | 1.13168E-29 | 0.304365 | 0.625 | 0.561 | 2.14159E-25 |
| Contractile | ARHGAP10 | 1.17251E-28 | 0.304194 | 0.494 | 0.397 | 2.21886E-24 |
| Contractile | NGRN | 2.15012E-33 | 0.304087 | 0.672 | 0.601 | 4.06889E-29 |
| Contractile | MAN2A1 | 2.35361E-38 | 0.303337 | 0.236 | 0.133 | 4.45398E-34 |
| Contractile | CCNB1IP1 | 4.25083E-32 | 0.30263 | 0.367 | 0.261 | 8.04427E-28 |
| Contractile | YWHAH | 1.07452E-30 | 0.301732 | 0.661 | 0.593 | 2.03342E-26 |
| Contractile | WSB1 | 3.72772E-22 | 0.301024 | 0.601 | 0.541 | 7.05433E-18 |
| Contractile | CDS2 | 2.53368E-34 | 0.300842 | 0.338 | 0.229 | 4.79474E-30 |
| Contractile | FLNC | 3.08676E-52 | 0.300602 | 0.161 | 0.06 | 5.84139E-48 |
| Contractile | PKD1 | 2.94464E-28 | 0.300438 | 0.34 | 0.24 | 5.57243E-24 |
| Contractile | NFIX | 7.44869E-29 | 0.300057 | 0.577 | 0.496 | 1.40959E-24 |
| Contractile | S1PR3 | 8.88749E-46 | 0.299815 | 0.203 | 0.095 | 1.68187E-41 |
| Contractile | H2AFZ | 2.98873E-30 | 0.299343 | 0.772 | 0.741 | 5.65587E-26 |
| Contractile | KLHL42 | 1.60566E-33 | 0.298049 | 0.329 | 0.22 | 3.03855E-29 |
| Contractile | SLC41A2 | 3.08667E-41 | 0.297348 | 0.201 | 0.101 | 5.84121E-37 |
| Contractile | GYG1 | 1.25778E-25 | 0.297013 | 0.424 | 0.339 | 2.38023E-21 |

| | | | | | | |
|---|---|---|---|---|---|---|
| Contractile | CREBRF | 8.23458E-32 | 0.296555 | 0.46 | 0.351 | 1.55831E-27 |
| Contractile | IGF1R | 1.57121E-29 | 0.295816 | 0.288 | 0.192 | 2.97335E-25 |
| Contractile | HEBP2 | 6.57523E-39 | 0.295694 | 0.699 | 0.627 | 1.2443E-34 |
| Contractile | TKT | 5.78562E-17 | 0.295367 | 0.479 | 0.421 | 1.09487E-12 |
| Contractile | SCARA3 | 1.41517E-43 | 0.294942 | 0.409 | 0.267 | 2.67807E-39 |
| Contractile | EHBP1L1 | 3.372E-34 | 0.29482 | 0.369 | 0.256 | 6.38117E-30 |
| Contractile | RBM27 | 3.46356E-25 | 0.2947 | 0.415 | 0.327 | 6.55445E-21 |
| Contractile | PACSIN3 | 7.97633E-52 | 0.294532 | 0.211 | 0.095 | 1.50944E-47 |
| Contractile | PDGFC | 8.79921E-35 | 0.294255 | 0.279 | 0.174 | 1.66516E-30 |
| Contractile | RHOQ | 1.90117E-30 | 0.293527 | 0.533 | 0.441 | 3.59778E-26 |
| Contractile | ASH1L | 3.21751E-22 | 0.292941 | 0.432 | 0.349 | 6.08882E-18 |
| Contractile | SHROOM3 | 7.27497E-38 | 0.292859 | 0.273 | 0.163 | 1.37671E-33 |
| Contractile | ANP32B | 4.01333E-34 | 0.292286 | 0.734 | 0.653 | 7.59482E-30 |
| Contractile | LINC01088 | 1.82604E-27 | 0.292189 | 0.197 | 0.116 | 3.45559E-23 |
| Contractile | DTNA | 1.29266E-52 | 0.292078 | 0.18 | 0.072 | 2.44624E-48 |
| Contractile | PRNP | 8.10052E-29 | 0.291515 | 0.562 | 0.478 | 1.53294E-24 |
| Contractile | SPOP | 4.28616E-21 | 0.290266 | 0.493 | 0.426 | 8.11113E-17 |
| Contractile | ALPK3 | 1.69337E-41 | 0.290174 | 0.176 | 0.081 | 3.20454E-37 |
| Contractile | CLASP2 | 3.04503E-39 | 0.289849 | 0.207 | 0.107 | 5.76242E-35 |
| Contractile | CD151 | 2.12591E-58 | 0.289619 | 0.957 | 0.928 | 4.02308E-54 |
| Contractile | METAP2 | 6.9227E-29 | 0.288605 | 0.609 | 0.537 | 1.31005E-24 |
| Contractile | HAND2 | 1.36502E-39 | 0.288435 | 0.285 | 0.166 | 2.58316E-35 |
| Contractile | NEDD9 | 3.16364E-24 | 0.288291 | 0.276 | 0.189 | 5.98688E-20 |
| Contractile | CALCOCO1 | 3.78354E-30 | 0.288196 | 0.34 | 0.238 | 7.15997E-26 |
| Contractile | ARHGEF10 | 5.81424E-37 | 0.288031 | 0.22 | 0.12 | 1.10029E-32 |
| Contractile | PIK3R1 | 4.64801E-19 | 0.287956 | 0.414 | 0.338 | 8.79589E-15 |
| Contractile | EIF4H | 6.30378E-30 | 0.287863 | 0.585 | 0.501 | 1.19293E-25 |
| Contractile | DUSP3 | 5.82367E-31 | 0.287801 | 0.276 | 0.178 | 1.10207E-26 |
| Contractile | NME4 | 3.37987E-18 | 0.287532 | 0.584 | 0.542 | 6.39607E-14 |
| Contractile | MTURN | 1.96524E-27 | 0.286975 | 0.348 | 0.253 | 3.71902E-23 |
| Contractile | LUC7L3 | 1.12478E-29 | 0.286887 | 0.672 | 0.597 | 2.12854E-25 |
| Contractile | LAMA5 | 6.94036E-38 | 0.28612 | 0.248 | 0.14 | 1.31339E-33 |
| Contractile | FAT3 | 1.80513E-52 | 0.286117 | 0.132 | 0.04 | 3.41602E-48 |
| Contractile | UBC | 1.3227E-77 | 0.285623 | 0.988 | 0.988 | 2.50307E-73 |
| Contractile | SDK1 | 1.92307E-71 | 0.28543 | 0.146 | 0.037 | 3.63921E-67 |
| Contractile | PXMP2 | 6.67576E-31 | 0.285323 | 0.276 | 0.178 | 1.26332E-26 |
| Contractile | SOD3 | 1.40407E-74 | 0.284599 | 0.981 | 0.957 | 2.65706E-70 |
| Contractile | MANBAL | 3.89899E-27 | 0.284044 | 0.377 | 0.285 | 7.37845E-23 |
| Contractile | TSR2 | 1.39811E-26 | 0.283312 | 0.463 | 0.369 | 2.64578E-22 |
| Contractile | DNAJB6 | 4.86115E-34 | 0.283303 | 0.626 | 0.53 | 9.19923E-30 |
| Contractile | JPH2 | 1.21632E-61 | 0.282735 | 0.157 | 0.05 | 2.30177E-57 |
| Contractile | GLIPR1 | 2.12184E-28 | 0.28273 | 0.658 | 0.579 | 4.01537E-24 |
| Contractile | SGCD | 9.1925E-22 | 0.281129 | 0.292 | 0.213 | 1.73959E-17 |
| Contractile | PEBP1 | 1.9153E-64 | 0.280976 | 0.961 | 0.941 | 3.62451E-60 |
| Contractile | CAB39 | 4.55292E-30 | 0.280574 | 0.265 | 0.17 | 8.61595E-26 |
| Contractile | TOMM20 | 9.33101E-40 | 0.278854 | 0.841 | 0.805 | 1.7658E-35 |
| Contractile | NOVA1 | 3.51949E-39 | 0.278177 | 0.218 | 0.114 | 6.66028E-35 |
| Contractile | GCNT2 | 1.87971E-51 | 0.277541 | 0.137 | 0.045 | 3.55717E-47 |
| Contractile | PYGB | 8.97597E-34 | 0.277481 | 0.266 | 0.164 | 1.69861E-29 |
| Contractile | PCBD1 | 9.33872E-29 | 0.277467 | 0.56 | 0.47 | 1.76726E-24 |
| Contractile | RBBP7 | 7.76586E-23 | 0.277204 | 0.531 | 0.457 | 1.46961E-18 |
| Contractile | LMO4 | 1.8873E-16 | 0.277098 | 0.453 | 0.39 | 3.57153E-12 |
| Contractile | GOLGA4 | 2.67293E-16 | 0.275973 | 0.54 | 0.488 | 5.05825E-12 |
| Contractile | NDUFB1 | 3.12331E-35 | 0.275144 | 0.771 | 0.732 | 5.91055E-31 |
| Contractile | DHRS7 | 2.37748E-25 | 0.274348 | 0.654 | 0.587 | 4.49914E-21 |
| Contractile | HEY2 | 4.00605E-24 | 0.274242 | 0.319 | 0.228 | 7.58105E-20 |
| Contractile | FBXL22 | 5.152E-37 | 0.274168 | 0.194 | 0.099 | 9.74964E-33 |
| Contractile | PRKAR1A | 1.1489E-18 | 0.274155 | 0.51 | 0.451 | 2.17417E-14 |

| | | | | | | |
|---|---|---|---|---|---|---|
| Contractile | GLDN | 7.32361E-33 | 0.273675 | 0.283 | 0.178 | 1.38592E-28 |
| Contractile | RSF1 | 3.95568E-22 | 0.273203 | 0.6 | 0.532 | 7.48574E-18 |
| Contractile | RBMX | 5.37674E-26 | 0.273144 | 0.588 | 0.519 | 1.01749E-21 |
| Contractile | PREX2 | 3.67166E-50 | 0.272974 | 0.15 | 0.053 | 6.94825E-46 |
| Contractile | ITGB1 | 2.37365E-40 | 0.272893 | 0.964 | 0.954 | 4.4919E-36 |
| Contractile | LINC02762 | 1.14757E-35 | 0.272547 | 0.401 | 0.275 | 2.17166E-31 |
| Contractile | AK4 | 1.38426E-41 | 0.271745 | 0.186 | 0.088 | 2.61957E-37 |
| Contractile | ITPR1 | 4.98623E-32 | 0.271532 | 0.28 | 0.177 | 9.43594E-28 |
| Contractile | SZRD1 | 5.57687E-22 | 0.271376 | 0.379 | 0.298 | 1.05537E-17 |
| Contractile | CENPV | 3.20706E-38 | 0.271082 | 0.212 | 0.112 | 6.06903E-34 |
| Contractile | PRKAR2A | 7.50389E-23 | 0.270402 | 0.295 | 0.213 | 1.42004E-18 |
| Contractile | GADD45G | 1.80424E-12 | 0.270305 | 0.171 | 0.119 | 3.41434E-08 |
| Contractile | PPIP5K2 | 3.14957E-24 | 0.269734 | 0.329 | 0.242 | 5.96024E-20 |
| Contractile | RAD23A | 4.72104E-27 | 0.269132 | 0.644 | 0.584 | 8.93409E-23 |
| Contractile | MIB1 | 1.09101E-30 | 0.269036 | 0.253 | 0.157 | 2.06463E-26 |
| Contractile | INPP5A | 2.07357E-36 | 0.268308 | 0.197 | 0.103 | 3.92403E-32 |
| Contractile | YPEL5 | 1.95603E-24 | 0.268186 | 0.515 | 0.435 | 3.7016E-20 |
| Contractile | NTF3 | 5.56146E-38 | 0.267909 | 0.182 | 0.09 | 1.05245E-33 |
| Contractile | SMAD9 | 3.05079E-27 | 0.267792 | 0.262 | 0.172 | 5.77332E-23 |
| Contractile | CNBP | 2.08727E-37 | 0.267604 | 0.887 | 0.852 | 3.94994E-33 |
| Contractile | MAGI2 | 1.42378E-27 | 0.266557 | 0.217 | 0.131 | 2.69436E-23 |
| Contractile | C22orf39 | 1.83138E-25 | 0.265279 | 0.355 | 0.264 | 3.4657E-21 |
| Contractile | RAB11B | 4.32189E-22 | 0.264925 | 0.469 | 0.397 | 8.17874E-18 |
| Contractile | CCDC146 | 1.63178E-43 | 0.264535 | 0.16 | 0.066 | 3.08798E-39 |
| Contractile | SYNC | 1.39973E-29 | 0.264242 | 0.281 | 0.183 | 2.64885E-25 |
| Contractile | SLURP2 | 7.70686E-66 | 0.264039 | 0.124 | 0.028 | 1.45845E-61 |
| Contractile | ANKRD12 | 5.49514E-27 | 0.263927 | 0.691 | 0.619 | 1.0399E-22 |
| Contractile | PKD2 | 3.05754E-19 | 0.263543 | 0.42 | 0.341 | 5.78609E-15 |
| Contractile | YTHDC1 | 1.49219E-25 | 0.26333 | 0.468 | 0.374 | 2.82382E-21 |
| Contractile | PPT1 | 6.65042E-20 | 0.262952 | 0.393 | 0.317 | 1.25853E-15 |
| Contractile | FMC1 | 5.60497E-24 | 0.262924 | 0.4 | 0.311 | 1.06068E-19 |
| Contractile | NT5DC3 | 6.30417E-39 | 0.262369 | 0.174 | 0.082 | 1.193E-34 |
| Contractile | CCDC115 | 1.46774E-18 | 0.261725 | 0.401 | 0.333 | 2.77755E-14 |
| Contractile | NINJ1 | 7.70856E-30 | 0.261444 | 0.45 | 0.343 | 1.45877E-25 |
| Contractile | PFN1 | 1.61469E-49 | 0.26139 | 0.944 | 0.921 | 3.05565E-45 |
| Contractile | ZBTB10 | 6.2583E-25 | 0.261355 | 0.259 | 0.172 | 1.18432E-20 |
| Contractile | HCFC1R1 | 1.70523E-33 | 0.26119 | 0.869 | 0.805 | 3.22697E-29 |
| Contractile | APBB2 | 1.59235E-29 | 0.261188 | 0.256 | 0.161 | 3.01336E-25 |
| Contractile | SOX5 | 1.158E-30 | 0.261105 | 0.174 | 0.091 | 2.1914E-26 |
| Contractile | BEX2 | 6.02845E-37 | 0.261034 | 0.158 | 0.07 | 1.14082E-32 |
| Contractile | PHIP | 1.61017E-14 | 0.2609 | 0.367 | 0.306 | 3.04709E-10 |
| Contractile | SON | 9.80454E-43 | 0.26054 | 0.881 | 0.848 | 1.85541E-38 |
| Contractile | FNBP1 | 1.51886E-18 | 0.260223 | 0.321 | 0.248 | 2.87429E-14 |
| Contractile | GAPDH | 2.19261E-85 | 0.259021 | 0.998 | 0.998 | 4.14929E-81 |
| Contractile | PKN2 | 4.48032E-22 | 0.258845 | 0.425 | 0.342 | 8.47855E-18 |
| Contractile | ATP5ME | 1.01924E-37 | 0.258555 | 0.847 | 0.828 | 1.92881E-33 |
| Contractile | IGFBPL1 | 2.07683E-54 | 0.25816 | 0.128 | 0.038 | 3.93018E-50 |
| Contractile | NUPR2 | 2.90003E-75 | 0.257739 | 0.105 | 0.014 | 5.48802E-71 |
| Contractile | GKAP1 | 2.92888E-35 | 0.257472 | 0.171 | 0.083 | 5.54262E-31 |
| Contractile | MRPL20 | 2.7155E-27 | 0.257185 | 0.706 | 0.653 | 5.1388E-23 |
| Contractile | IL13RA2 | 2.3478E-09 | 0.256647 | 0.128 | 0.09 | 4.44297E-05 |
| Contractile | ZMAT2 | 4.11597E-20 | 0.256195 | 0.441 | 0.372 | 7.78907E-16 |
| Contractile | DYNLL2 | 2.11275E-29 | 0.256183 | 0.255 | 0.16 | 3.99817E-25 |
| Contractile | ZNF292 | 2.96231E-18 | 0.255316 | 0.32 | 0.245 | 5.60588E-14 |
| Contractile | ZYX | 1.22357E-28 | 0.255174 | 0.579 | 0.48 | 2.31548E-24 |
| Contractile | KCTD9 | 1.51472E-24 | 0.255016 | 0.266 | 0.181 | 2.86645E-20 |
| Contractile | PPFIBP1 | 5.70221E-21 | 0.254987 | 0.58 | 0.511 | 1.07909E-16 |
| Contractile | PFKP | 2.58251E-22 | 0.254418 | 0.261 | 0.184 | 4.88713E-18 |

| | | | | | | |
|---|---|---|---|---|---|---|
| Contractile | EFR3A | 1.42146E-19 | 0.254024 | 0.322 | 0.247 | 2.68996E-15 |
| Contractile | CHD2 | 3.8357E-12 | 0.253945 | 0.328 | 0.274 | 7.25868E-08 |
| Contractile | RTF1 | 1.55256E-22 | 0.253732 | 0.617 | 0.558 | 2.93806E-18 |
| Contractile | ETS2 | 4.64728E-21 | 0.25366 | 0.421 | 0.333 | 8.7945E-17 |
| Contractile | PID1 | 1.59207E-15 | 0.253576 | 0.35 | 0.285 | 3.01283E-11 |
| Contractile | NSRP1 | 3.28946E-18 | 0.253389 | 0.565 | 0.506 | 6.22497E-14 |
| Contractile | RHOU | 4.49664E-30 | 0.252567 | 0.184 | 0.101 | 8.50944E-26 |
| Contractile | SLK | 2.01181E-21 | 0.252426 | 0.381 | 0.299 | 3.80714E-17 |
| Contractile | FXR1 | 4.01369E-23 | 0.252387 | 0.542 | 0.457 | 7.59551E-19 |
| Contractile | CARMIL1 | 4.40797E-24 | 0.252346 | 0.256 | 0.172 | 8.34164E-20 |
| Contractile | ADARB1 | 1.36515E-24 | 0.252328 | 0.225 | 0.144 | 2.58341E-20 |
| Contractile | AMIGO2 | 1.11536E-27 | 0.252093 | 0.219 | 0.132 | 2.11072E-23 |
| Contractile | C1QTNF7 | 9.35386E-30 | 0.251493 | 0.277 | 0.175 | 1.77012E-25 |
| Contractile | ESYT2 | 1.03767E-16 | 0.251387 | 0.394 | 0.329 | 1.96368E-12 |
| Contractile | CES1 | 6.97335E-27 | 0.250802 | 0.374 | 0.266 | 1.31964E-22 |
| Contractile | CWF19L2 | 3.83489E-24 | 0.250548 | 0.398 | 0.308 | 7.25715E-20 |
| Terminal | LUM | 0 | 3.669083 | 0.785 | 0.157 | 0 |
| Terminal | DCN | 0 | 3.389492 | 0.589 | 0.101 | 0 |
| Terminal | SFRP2 | 0 | 3.307987 | 0.431 | 0.051 | 0 |
| Terminal | APOE | 1.3677E-194 | 2.996978 | 0.452 | 0.163 | 2.5883E-190 |
| Terminal | CCL19 | 2.9294E-134 | 2.91959 | 0.175 | 0.025 | 5.5436E-130 |
| Terminal | CFH | 0 | 2.881024 | 0.92 | 0.288 | 0 |
| Terminal | PTGDS | 1.2437E-139 | 2.578267 | 0.175 | 0.023 | 2.3536E-135 |
| Terminal | COL3A1 | 0 | 2.468253 | 0.863 | 0.518 | 0 |
| Terminal | COL1A1 | 0 | 2.293387 | 0.772 | 0.393 | 0 |
| Terminal | SLPI | 1.01344E-87 | 2.280345 | 0.13 | 0.022 | 1.91783E-83 |
| Terminal | POSTN | 1.3324E-248 | 2.216939 | 0.477 | 0.141 | 2.5215E-244 |
| Terminal | TIMP1 | 0 | 2.207223 | 1 | 0.966 | 0 |
| Terminal | COL1A2 | 0 | 2.034456 | 0.968 | 0.75 | 0 |
| Terminal | CCL2 | 2.8654E-211 | 1.837853 | 0.507 | 0.18 | 5.4225E-207 |
| Terminal | CCDC80 | 0 | 1.803953 | 0.859 | 0.433 | 0 |
| Terminal | CRTAC1 | 0 | 1.675458 | 0.403 | 0.052 | 0 |
| Terminal | SERPINF1 | 8.17E-245 | 1.664877 | 0.341 | 0.06 | 1.5461E-240 |
| Terminal | CCL21 | 2.4547E-112 | 1.644678 | 0.168 | 0.029 | 4.6453E-108 |
| Terminal | C1R | 0 | 1.605304 | 0.901 | 0.599 | 0 |
| Terminal | VCAM1 | 0 | 1.598911 | 0.587 | 0.161 | 0 |
| Terminal | C7 | 2.026E-238 | 1.595109 | 0.257 | 0.026 | 3.834E-234 |
| Terminal | COL6A3 | 2.8253E-307 | 1.59172 | 0.604 | 0.206 | 5.3465E-303 |
| Terminal | PCOLCE | 0 | 1.493518 | 0.834 | 0.444 | 0 |
| Terminal | TNC | 0 | 1.487921 | 0.44 | 0.089 | 0 |
| Terminal | COL15A1 | 0 | 1.454843 | 0.586 | 0.152 | 0 |
| Terminal | SFRP4 | 4.0073E-117 | 1.434931 | 0.457 | 0.224 | 7.5835E-113 |
| Terminal | S100A10 | 3.8691E-251 | 1.421326 | 0.953 | 0.816 | 7.3219E-247 |
| Terminal | MARCKS | 0 | 1.406602 | 0.774 | 0.305 | 0 |
| Terminal | FN1 | 5.9927E-265 | 1.374084 | 0.984 | 0.886 | 1.1341E-260 |
| Terminal | TNFSF10 | 0 | 1.362815 | 0.392 | 0.053 | 0 |
| Terminal | IFI27 | 5.0812E-116 | 1.358129 | 0.313 | 0.108 | 9.6156E-112 |
| Terminal | PCOLCE2 | 2.5271E-192 | 1.331292 | 0.495 | 0.192 | 4.7823E-188 |
| Terminal | MGP | 0 | 1.322907 | 0.999 | 0.997 | 0 |
| Terminal | GSN | 2.82E-105 | 1.318278 | 0.978 | 0.964 | 5.3366E-101 |
| Terminal | SERPINA3 | 1.5587E-273 | 1.317686 | 0.289 | 0.029 | 2.9498E-269 |
| Terminal | AGT | 6.9141E-270 | 1.309947 | 0.516 | 0.146 | 1.3084E-265 |
| Terminal | DPT | 4.5136E-185 | 1.303068 | 0.552 | 0.235 | 8.5415E-181 |
| Terminal | TFPI2 | 1.21671E-57 | 1.29858 | 0.289 | 0.15 | 2.3025E-53 |
| Terminal | VCAN | 0 | 1.276243 | 0.972 | 0.725 | 0 |
| Terminal | SPRY1 | 0 | 1.273783 | 0.645 | 0.217 | 0 |
| Terminal | FGF7 | 1.1918E-236 | 1.241388 | 0.364 | 0.073 | 2.2554E-232 |
| Terminal | CXCL12 | 1.9423E-190 | 1.21003 | 0.682 | 0.354 | 3.6756E-186 |

| | | | | | | |
|---|---|---|---|---|---|---|
| Terminal | CTSK | 0 | 1.197409 | 0.677 | 0.247 | 0 |
| Terminal | MT2A | 4.136E-203 | 1.188434 | 0.904 | 0.708 | 7.8269E-199 |
| Terminal | PRSS23 | 3.9557E-246 | 1.179811 | 0.814 | 0.48 | 7.4857E-242 |
| Terminal | CTSC | 2.1495E-199 | 1.176422 | 0.502 | 0.18 | 4.0677E-195 |
| Terminal | C1S | 1.1526E-278 | 1.171041 | 0.808 | 0.439 | 2.1812E-274 |
| Terminal | AEBP1 | 0 | 1.166456 | 0.972 | 0.848 | 0 |
| Terminal | GGT5 | 2.7229E-306 | 1.165222 | 0.446 | 0.087 | 5.1529E-302 |
| Terminal | ECRG4 | 5.43E-103 | 1.163786 | 0.673 | 0.428 | 1.02758E-98 |
| Terminal | LY6E | 6.7679E-250 | 1.148264 | 0.732 | 0.357 | 1.2808E-245 |
| Terminal | CCL5 | 3.7781E-75 | 1.140485 | 0.143 | 0.033 | 7.14968E-71 |
| Terminal | FAP | 0 | 1.13993 | 0.544 | 0.124 | 0 |
| Terminal | LTBP2 | 8.1245E-246 | 1.138393 | 0.809 | 0.453 | 1.5375E-241 |
| Terminal | FBLN1 | 1.2171E-136 | 1.13534 | 0.435 | 0.179 | 2.3033E-132 |
| Terminal | APOC1 | 5.9293E-106 | 1.116065 | 0.182 | 0.039 | 1.1221E-101 |
| Terminal | THY1 | 3.6812E-306 | 1.097658 | 0.464 | 0.093 | 6.9662E-302 |
| Terminal | HLA-B | 1.5762E-247 | 1.079278 | 0.937 | 0.711 | 2.9828E-243 |
| Terminal | RGS16 | 3.377E-177 | 1.07812 | 0.397 | 0.116 | 6.3906E-173 |
| Terminal | SULF1 | 1.0098E-173 | 1.075435 | 0.745 | 0.453 | 1.9109E-169 |
| Terminal | HLA-DRB1 | 4.23307E-91 | 1.067422 | 0.328 | 0.138 | 8.01067E-87 |
| Terminal | TFPI | 7.1587E-261 | 1.052463 | 0.747 | 0.367 | 1.3547E-256 |
| Terminal | TYMP | 2.2403E-304 | 1.041775 | 0.486 | 0.108 | 4.2395E-300 |
| Terminal | STEAP4 | 4.1807E-239 | 1.039529 | 0.353 | 0.061 | 7.9116E-235 |
| Terminal | FBLN2 | 6.8739E-230 | 1.039403 | 0.347 | 0.068 | 1.3008E-225 |
| Terminal | BGN | 0 | 1.024256 | 0.993 | 0.977 | 0 |
| Terminal | TMSB10 | 0 | 1.011933 | 0.995 | 0.95 | 0 |
| Terminal | THBS2 | 1.8504E-174 | 1.010686 | 0.754 | 0.446 | 3.5016E-170 |
| Terminal | GAP43 | 3.1335E-231 | 1.009495 | 0.329 | 0.059 | 5.9298E-227 |
| Terminal | NPC2 | 7.27E-259 | 1.007265 | 0.9 | 0.641 | 1.3758E-254 |
| Terminal | NNMT | 3.0257E-220 | 0.998157 | 0.931 | 0.749 | 5.7258E-216 |
| Terminal | TNFAIP6 | 8.7704E-255 | 0.99664 | 0.269 | 0.026 | 1.6597E-250 |
| Terminal | F2R | 8.8006E-260 | 0.986382 | 0.532 | 0.161 | 1.6654E-255 |
| Terminal | PLAC9 | 1.594E-283 | 0.985898 | 0.923 | 0.538 | 3.0165E-279 |
| Terminal | RCN3 | 4.0493E-271 | 0.974584 | 0.663 | 0.271 | 7.663E-267 |
| Terminal | CD74 | 1.36661E-45 | 0.961223 | 0.389 | 0.237 | 2.58618E-41 |
| Terminal | CST3 | 0 | 0.957325 | 1 | 0.993 | 0 |
| Terminal | FXYD5 | 9.4029E-240 | 0.94452 | 0.777 | 0.403 | 1.7794E-235 |
| Terminal | MMP2 | 7.2804E-193 | 0.940732 | 0.588 | 0.259 | 1.3777E-188 |
| Terminal | HLA-A | 2.273E-207 | 0.930679 | 0.967 | 0.858 | 4.3015E-203 |
| Terminal | IL32 | 3.0696E-190 | 0.914805 | 0.374 | 0.097 | 5.8089E-186 |
| Terminal | COL6A2 | 6.007E-219 | 0.913497 | 0.965 | 0.869 | 1.1368E-214 |
| Terminal | CP | 1.07624E-87 | 0.910414 | 0.408 | 0.201 | 2.03667E-83 |
| Terminal | SH3BGRL3 | 5.4531E-164 | 0.908328 | 0.909 | 0.778 | 1.0319E-159 |
| Terminal | SOD2 | 1.91387E-36 | 0.906944 | 0.519 | 0.392 | 3.6218E-32 |
| Terminal | ISLR | 9.0492E-177 | 0.906113 | 0.58 | 0.27 | 1.7125E-172 |
| Terminal | PMP22 | 5.4656E-227 | 0.90578 | 0.821 | 0.504 | 1.0343E-222 |
| Terminal | IGFBP6 | 1.10528E-54 | 0.903285 | 0.846 | 0.768 | 2.09163E-50 |
| Terminal | PLTP | 2.2405E-239 | 0.901705 | 0.459 | 0.124 | 4.24E-235 |
| Terminal | KDELR3 | 1.3829E-274 | 0.876601 | 0.46 | 0.114 | 2.617E-270 |
| Terminal | TNFSF13B | 5.3773E-264 | 0.871101 | 0.258 | 0.02 | 1.0176E-259 |
| Terminal | CLEC11A | 7.6102E-238 | 0.870547 | 0.38 | 0.081 | 1.4402E-233 |
| Terminal | CYTL1 | 1.36433E-30 | 0.868136 | 0.193 | 0.1 | 2.58187E-26 |
| Terminal | ITM2B | 4.1638E-295 | 0.863309 | 0.995 | 0.984 | 7.8796E-291 |
| Terminal | CDH11 | 1.9439E-175 | 0.853408 | 0.571 | 0.256 | 3.6786E-171 |
| Terminal | SERPINE2 | 3.6833E-241 | 0.845401 | 0.357 | 0.064 | 6.9702E-237 |
| Terminal | SOCS3 | 1.1517E-165 | 0.830876 | 0.47 | 0.172 | 2.1795E-161 |
| Terminal | SPARC | 4.4644E-146 | 0.82868 | 0.95 | 0.834 | 8.4483E-142 |
| Terminal | NDUFA4L2 | 1.1419E-127 | 0.828404 | 0.597 | 0.307 | 2.1609E-123 |
| Terminal | COL5A2 | 5.4198E-140 | 0.82203 | 0.529 | 0.258 | 1.0256E-135 |

| | | | | | | |
|---|---|---|---|---|---|---|
| Terminal | G0S2 | 3.3295E-117 | 0.817134 | 0.221 | 0.052 | 6.3007E-113 |
| Terminal | TGM2 | 1.29462E-76 | 0.816074 | 0.56 | 0.37 | 2.44993E-72 |
| Terminal | HLA-C | 6.2261E-205 | 0.815045 | 0.976 | 0.893 | 1.1782E-200 |
| Terminal | RRBP1 | 4.1699E-189 | 0.809137 | 0.751 | 0.438 | 7.8912E-185 |
| Terminal | COL6A1 | 6.8686E-116 | 0.796477 | 0.85 | 0.691 | 1.2998E-111 |
| Terminal | IGFBP4 | 6.4866E-100 | 0.789736 | 0.513 | 0.282 | 1.22752E-95 |
| Terminal | STEAP1 | 1.6452E-209 | 0.787392 | 0.358 | 0.082 | 3.1134E-205 |
| Terminal | HSPA1A | 3.44457E-07 | 0.772833 | 0.721 | 0.731 | 0.006518495 |
| Terminal | HLA-E | 2.6701E-159 | 0.768248 | 0.875 | 0.667 | 5.0529E-155 |
| Terminal | LGALS3BP | 1.489E-152 | 0.766447 | 0.732 | 0.454 | 2.8178E-148 |
| Terminal | IER3 | 2.49733E-82 | 0.761101 | 0.444 | 0.244 | 4.72594E-78 |
| Terminal | FOS | 1.04271E-81 | 0.758011 | 0.818 | 0.67 | 1.97323E-77 |
| Terminal | PPIC | 1.0172E-154 | 0.757656 | 0.789 | 0.561 | 1.9249E-150 |
| Terminal | CD9 | 1.6472E-125 | 0.754911 | 0.979 | 0.966 | 3.1172E-121 |
| Terminal | ECM1 | 3.5251E-245 | 0.753509 | 0.388 | 0.083 | 6.6709E-241 |
| Terminal | HSPA1B | 1.93144E-33 | 0.750561 | 0.516 | 0.413 | 3.65506E-29 |
| Terminal | RGS2 | 1.5928E-121 | 0.750242 | 0.357 | 0.13 | 3.0143E-117 |
| Terminal | NUCB2 | 2.2032E-143 | 0.738231 | 0.785 | 0.546 | 4.1694E-139 |
| Terminal | CRABP2 | 4.3667E-174 | 0.738115 | 0.215 | 0.028 | 8.2635E-170 |
| Terminal | HTRA1 | 1.4613E-124 | 0.737793 | 0.895 | 0.767 | 2.7655E-120 |
| Terminal | PPIB | 1.7752E-245 | 0.737553 | 0.974 | 0.899 | 3.3594E-241 |
| Terminal | FOSB | 5.0055E-108 | 0.735295 | 0.547 | 0.286 | 9.4724E-104 |
| Terminal | LRRC17 | 3.322E-133 | 0.735163 | 0.554 | 0.273 | 6.2866E-129 |
| Terminal | SERPINE1 | 3.29621E-76 | 0.733835 | 0.484 | 0.27 | 6.23775E-72 |
| Terminal | EMP3 | 5.2816E-119 | 0.73324 | 0.875 | 0.723 | 9.9949E-115 |
| Terminal | CEBPD | 8.445E-99 | 0.72175 | 0.75 | 0.541 | 1.59812E-94 |
| Terminal | CFB | 1.1844E-160 | 0.720802 | 0.488 | 0.192 | 2.2413E-156 |
| Terminal | EMP1 | 3.145E-214 | 0.716159 | 0.37 | 0.086 | 5.9517E-210 |
| Terminal | FBN1 | 2.9777E-114 | 0.714903 | 0.559 | 0.303 | 5.635E-110 |
| Terminal | B2M | 5.6273E-203 | 0.704068 | 0.998 | 0.997 | 1.0649E-198 |
| Terminal | SERPINH1 | 6.2408E-124 | 0.700256 | 0.598 | 0.339 | 1.181E-119 |
| Terminal | MMP23B | 6.089E-117 | 0.699297 | 0.367 | 0.141 | 1.1523E-112 |
| Terminal | CYP1B1 | 3.59E-155 | 0.697216 | 0.192 | 0.025 | 6.7937E-151 |
| Terminal | MXRA8 | 5.7192E-115 | 0.693183 | 0.752 | 0.523 | 1.0823E-110 |
| Terminal | DIO2 | 9.2453E-102 | 0.690822 | 0.221 | 0.06 | 1.74957E-97 |
| Terminal | PDGFRL | 1.538E-127 | 0.68596 | 0.379 | 0.139 | 2.9105E-123 |
| Terminal | ISG15 | 6.52132E-43 | 0.684229 | 0.4 | 0.247 | 1.23409E-38 |
| Terminal | MXRA5 | 1.5276E-247 | 0.676868 | 0.239 | 0.017 | 2.8909E-243 |
| Terminal | TM4SF1 | 1.27228E-97 | 0.674947 | 0.808 | 0.622 | 2.40767E-93 |
| Terminal | MDK | 4.874E-133 | 0.674569 | 0.327 | 0.103 | 9.2236E-129 |
| Terminal | SOX4 | 1.26084E-81 | 0.674379 | 0.482 | 0.266 | 2.38602E-77 |
| Terminal | TNXB | 6.4394E-112 | 0.6725 | 0.251 | 0.072 | 1.2186E-107 |
| Terminal | FAM3C | 4.5047E-142 | 0.672433 | 0.632 | 0.351 | 8.5246E-138 |
| Terminal | TIMP3 | 1.66331E-24 | 0.657711 | 0.358 | 0.541 | 3.14764E-20 |
| Terminal | PROCR | 1.5253E-117 | 0.657635 | 0.338 | 0.119 | 2.8865E-113 |
| Terminal | IFI16 | 1.3502E-126 | 0.656697 | 0.663 | 0.384 | 2.5552E-122 |
| Terminal | FKBP10 | 6.9823E-149 | 0.654273 | 0.583 | 0.284 | 1.3213E-144 |
| Terminal | TNFRSF11I | 2.61146E-28 | 0.648744 | 0.478 | 0.354 | 4.94193E-24 |
| Terminal | ANKH | 2.92789E-68 | 0.645738 | 0.503 | 0.317 | 5.54073E-64 |
| Terminal | LAMB1 | 7.3938E-154 | 0.645314 | 0.329 | 0.092 | 1.3992E-149 |
| Terminal | IRF1 | 3.44392E-88 | 0.644277 | 0.316 | 0.125 | 6.51727E-84 |
| Terminal | FHL2 | 6.6176E-98 | 0.643447 | 0.479 | 0.244 | 1.25232E-93 |
| Terminal | IL34 | 4.8818E-165 | 0.639875 | 0.447 | 0.158 | 9.2383E-161 |
| Terminal | HSP90B1 | 3.2135E-159 | 0.635082 | 0.971 | 0.893 | 6.0813E-155 |
| Terminal | BST2 | 1.08802E-78 | 0.633423 | 0.621 | 0.382 | 2.05898E-74 |
| Terminal | TSPAN4 | 5.9085E-135 | 0.633046 | 0.589 | 0.299 | 1.1181E-130 |
| Terminal | ODF3B | 2.928E-191 | 0.632531 | 0.358 | 0.086 | 5.5409E-187 |
| Terminal | IGFBP3 | 7.64684E-26 | 0.631851 | 0.379 | 0.274 | 1.44709E-21 |

| | | | | | | |
|---|---|---|---|---|---|---|
| Terminal | HLA-DRB5 | 4.93358E-57 | 0.627563 | 0.157 | 0.051 | 9.33631E-53 |
| Terminal | PLAAT4 | 1.68428E-95 | 0.625674 | 0.465 | 0.229 | 3.18733E-91 |
| Terminal | COL5A1 | 2.4456E-111 | 0.624887 | 0.358 | 0.141 | 4.628E-107 |
| Terminal | IFI27L2 | 3.481E-121 | 0.62415 | 0.708 | 0.435 | 6.5874E-117 |
| Terminal | AKR1C2 | 1.5865E-122 | 0.619372 | 0.452 | 0.194 | 3.0023E-118 |
| Terminal | UACA | 1.06035E-93 | 0.618871 | 0.73 | 0.515 | 2.00662E-89 |
| Terminal | MT1E | 1.36857E-89 | 0.618778 | 0.689 | 0.456 | 2.58989E-85 |
| Terminal | MMP11 | 6.052E-101 | 0.617999 | 0.156 | 0.028 | 1.14528E-96 |
| Terminal | CTSZ | 1.57E-135 | 0.617526 | 0.871 | 0.682 | 2.9711E-131 |
| Terminal | DAB2 | 2.4585E-147 | 0.615778 | 0.468 | 0.186 | 4.6524E-143 |
| Terminal | KDELR2 | 6.5345E-121 | 0.614221 | 0.755 | 0.536 | 1.2366E-116 |
| Terminal | NBL1 | 2.07427E-92 | 0.613921 | 0.752 | 0.543 | 3.92535E-88 |
| Terminal | ANGPT2 | 2.24794E-62 | 0.61276 | 0.44 | 0.253 | 4.254E-58 |
| Terminal | PRDX4 | 3.802E-116 | 0.609749 | 0.769 | 0.566 | 7.1949E-112 |
| Terminal | FAM114A1 | 2.9251E-118 | 0.607217 | 0.705 | 0.458 | 5.5354E-114 |
| Terminal | DUSP6 | 1.72015E-95 | 0.607211 | 0.322 | 0.128 | 3.25522E-91 |
| Terminal | COL12A1 | 1.08614E-56 | 0.607059 | 0.385 | 0.218 | 2.0554E-52 |
| Terminal | IFITM2 | 1.0035E-124 | 0.607049 | 0.921 | 0.746 | 1.899E-120 |
| Terminal | SMOC1 | 6.73E-150 | 0.606586 | 0.168 | 0.018 | 1.2736E-145 |
| Terminal | JUNB | 2.3922E-126 | 0.601851 | 0.882 | 0.639 | 4.527E-122 |
| Terminal | DPP7 | 4.9413E-117 | 0.601243 | 0.689 | 0.431 | 9.3509E-113 |
| Terminal | IGKC | 2.69283E-73 | 0.599925 | 0.418 | 0.221 | 5.0959E-69 |
| Terminal | PRRX1 | 5.8126E-104 | 0.597837 | 0.745 | 0.498 | 1.1E-99 |
| Terminal | CCDC102B | 6.5843E-126 | 0.590904 | 0.332 | 0.106 | 1.246E-121 |
| Terminal | LRRC32 | 8.498E-100 | 0.588041 | 0.358 | 0.149 | 1.60816E-95 |
| Terminal | UNC5B | 5.4211E-177 | 0.587709 | 0.331 | 0.08 | 1.0259E-172 |
| Terminal | PHLDA2 | 1.0293E-145 | 0.586642 | 0.433 | 0.159 | 1.9478E-141 |
| Terminal | MRC2 | 1.1555E-119 | 0.58448 | 0.461 | 0.206 | 2.1867E-115 |
| Terminal | OMD | 2.42167E-82 | 0.584187 | 0.64 | 0.386 | 4.58278E-78 |
| Terminal | ARPC1B | 1.73363E-97 | 0.583257 | 0.811 | 0.625 | 3.28071E-93 |
| Terminal | CD59 | 7.4898E-128 | 0.576519 | 0.916 | 0.822 | 1.4174E-123 |
| Terminal | EGR1 | 1.37039E-87 | 0.575807 | 0.663 | 0.415 | 2.59333E-83 |
| Terminal | LAP3 | 1.65976E-67 | 0.574694 | 0.435 | 0.25 | 3.14094E-63 |
| Terminal | ZFP36 | 4.11324E-62 | 0.574089 | 0.721 | 0.548 | 7.78389E-58 |
| Terminal | NFKBIA | 2.46119E-56 | 0.570892 | 0.71 | 0.54 | 4.65756E-52 |
| Terminal | PDGFRA | 2.8113E-106 | 0.568638 | 0.506 | 0.252 | 5.3202E-102 |
| Terminal | ABI3BP | 5.63996E-59 | 0.568063 | 0.377 | 0.207 | 1.06731E-54 |
| Terminal | TPM3 | 1.7491E-112 | 0.565302 | 0.537 | 0.276 | 3.3099E-108 |
| Terminal | S100A13 | 6.27693E-93 | 0.564072 | 0.823 | 0.638 | 1.18785E-88 |
| Terminal | S100A16 | 1.76424E-92 | 0.562599 | 0.646 | 0.414 | 3.33864E-88 |
| Terminal | FKBP11 | 9.6932E-164 | 0.562564 | 0.337 | 0.091 | 1.8343E-159 |
| Terminal | GPM6B | 5.2111E-112 | 0.560693 | 0.37 | 0.144 | 9.8615E-108 |
| Terminal | PDIA3 | 1.1487E-120 | 0.559672 | 0.897 | 0.758 | 2.1739E-116 |
| Terminal | ANXA1 | 3.8645E-142 | 0.559471 | 0.979 | 0.932 | 7.3131E-138 |
| Terminal | GADD45B | 3.39245E-40 | 0.556823 | 0.655 | 0.524 | 6.41987E-36 |
| Terminal | GPX8 | 6.5384E-114 | 0.555029 | 0.547 | 0.285 | 1.2373E-109 |
| Terminal | CERCAM | 7.3094E-115 | 0.554988 | 0.438 | 0.193 | 1.3832E-110 |
| Terminal | KLF2 | 3.4789E-109 | 0.554317 | 0.677 | 0.418 | 6.5835E-105 |
| Terminal | CISD1 | 2.38788E-87 | 0.546758 | 0.529 | 0.303 | 4.51882E-83 |
| Terminal | NTRK2 | 1.07644E-85 | 0.546211 | 0.267 | 0.096 | 2.03706E-81 |
| Terminal | TAGLN2 | 3.9724E-131 | 0.54607 | 0.939 | 0.797 | 7.5174E-127 |
| Terminal | COL14A1 | 6.72236E-73 | 0.544178 | 0.941 | 0.861 | 1.27214E-68 |
| Terminal | LAMA2 | 2.4162E-101 | 0.54359 | 0.262 | 0.084 | 4.57236E-97 |
| Terminal | PLPP1 | 6.03478E-66 | 0.543323 | 0.645 | 0.45 | 1.14202E-61 |
| Terminal | CALU | 3.9159E-100 | 0.542177 | 0.761 | 0.552 | 7.41039E-96 |
| Terminal | HLA-DRA | 5.16428E-22 | 0.540451 | 0.298 | 0.196 | 9.77288E-18 |
| Terminal | CALCRL | 6.01005E-94 | 0.539837 | 0.38 | 0.166 | 1.13734E-89 |
| Terminal | DDIT4 | 1.84265E-72 | 0.5384 | 0.291 | 0.121 | 3.48703E-68 |

| | | | | | | |
|---|---|---|---|---|---|---|
| Terminal | FCGRT | 1.52386E-94 | 0.536061 | 0.682 | 0.451 | 2.88376E-90 |
| Terminal | LAMA4 | 3.8029E-120 | 0.535405 | 0.405 | 0.161 | 7.1966E-116 |
| Terminal | CYBA | 1.6237E-104 | 0.534318 | 0.676 | 0.409 | 3.0728E-100 |
| Terminal | P4HB | 4.18667E-94 | 0.533518 | 0.808 | 0.636 | 7.92286E-90 |
| Terminal | ARHGAP29 | 2.26065E-72 | 0.533306 | 0.555 | 0.344 | 4.27806E-68 |
| Terminal | TGFBR2 | 7.32915E-73 | 0.532903 | 0.539 | 0.327 | 1.38697E-68 |
| Terminal | HLA-DPA1 | 1.34668E-49 | 0.532591 | 0.262 | 0.125 | 2.54846E-45 |
| Terminal | LOXL1 | 4.74629E-59 | 0.5324 | 0.467 | 0.289 | 8.98188E-55 |
| Terminal | TSC22D3 | 3.13783E-58 | 0.528612 | 0.727 | 0.57 | 5.93804E-54 |
| Terminal | ITM2A | 1.89662E-56 | 0.526536 | 0.241 | 0.107 | 3.58916E-52 |
| Terminal | CCN1 | 2.05869E-41 | 0.526424 | 0.819 | 0.734 | 3.89586E-37 |
| Terminal | CD55 | 6.05267E-58 | 0.526379 | 0.438 | 0.254 | 1.14541E-53 |
| Terminal | MYDGF | 1.98846E-85 | 0.523832 | 0.772 | 0.598 | 3.76296E-81 |
| Terminal | PSMB9 | 1.42593E-85 | 0.523343 | 0.416 | 0.198 | 2.69843E-81 |
| Terminal | CHPF | 1.20463E-88 | 0.522905 | 0.457 | 0.239 | 2.27964E-84 |
| Terminal | MEG3 | 4.76949E-77 | 0.522866 | 0.225 | 0.077 | 9.02579E-73 |
| Terminal | IKBIP | 4.2184E-105 | 0.522465 | 0.455 | 0.211 | 7.9829E-101 |
| Terminal | DEPP1 | 1.76301E-25 | 0.521209 | 0.361 | 0.247 | 3.33632E-21 |
| Terminal | PLAUR | 6.5155E-109 | 0.520809 | 0.256 | 0.075 | 1.233E-104 |
| Terminal | PTHLH | 1.1978E-119 | 0.520494 | 0.196 | 0.039 | 2.2667E-115 |
| Terminal | PSME2 | 3.63127E-48 | 0.518031 | 0.62 | 0.47 | 6.87182E-44 |
| Terminal | SSR4 | 6.818E-116 | 0.517881 | 0.93 | 0.826 | 1.2902E-111 |
| Terminal | PPP3CA | 5.1106E-107 | 0.513017 | 0.415 | 0.179 | 9.6714E-103 |
| Terminal | IFI6 | 1.63943E-31 | 0.512841 | 0.507 | 0.369 | 3.10246E-27 |
| Terminal | COLEC12 | 3.2625E-131 | 0.512111 | 0.272 | 0.072 | 6.1739E-127 |
| Terminal | GPX3 | 5.74094E-60 | 0.507723 | 0.522 | 0.328 | 1.08642E-55 |
| Terminal | PDGFD | 2.03012E-72 | 0.506946 | 0.449 | 0.246 | 3.8418E-68 |
| Terminal | TCF4 | 3.29919E-69 | 0.505718 | 0.733 | 0.535 | 6.24339E-65 |
| Terminal | FMO2 | 1.31471E-05 | 0.496767 | 0.453 | 0.44 | 0.248794953 |
| Terminal | CTSA | 2.28867E-85 | 0.496707 | 0.516 | 0.291 | 4.33108E-81 |
| Terminal | OS9 | 3.11266E-88 | 0.49528 | 0.642 | 0.41 | 5.89041E-84 |
| Terminal | ARHGDIB | 4.10399E-83 | 0.491601 | 0.674 | 0.425 | 7.76638E-79 |
| Terminal | CD109 | 1.30666E-89 | 0.49082 | 0.324 | 0.133 | 2.47272E-85 |
| Terminal | ANXA2 | 1.4634E-109 | 0.490124 | 0.973 | 0.922 | 2.7694E-105 |
| Terminal | IFNGR2 | 8.8346E-110 | 0.488901 | 0.412 | 0.174 | 1.6719E-105 |
| Terminal | APOD | 1.8266E-105 | 0.487602 | 0.103 | 0.007 | 3.4567E-101 |
| Terminal | CDKN2A | 9.08886E-98 | 0.487054 | 0.208 | 0.054 | 1.71998E-93 |
| Terminal | IFIT3 | 1.71753E-59 | 0.485398 | 0.203 | 0.075 | 3.25026E-55 |
| Terminal | BICC1 | 1.8216E-111 | 0.485053 | 0.373 | 0.143 | 3.4471E-107 |
| Terminal | EFEMP1 | 7.25718E-62 | 0.484497 | 0.874 | 0.743 | 1.37335E-57 |
| Terminal | MT1M | 2.24363E-60 | 0.482722 | 0.528 | 0.332 | 4.24584E-56 |
| Terminal | TPPP3 | 1.98204E-49 | 0.48185 | 0.208 | 0.087 | 3.75081E-45 |
| Terminal | SPTBN1 | 5.64398E-63 | 0.479787 | 0.753 | 0.581 | 1.06807E-58 |
| Terminal | CTHRC1 | 2.49465E-16 | 0.479731 | 0.433 | 0.342 | 4.72087E-12 |
| Terminal | LIMA1 | 1.23649E-81 | 0.478359 | 0.691 | 0.453 | 2.33994E-77 |
| Terminal | IGLC2 | 5.22693E-70 | 0.477386 | 0.291 | 0.123 | 9.89145E-66 |
| Terminal | BZW1 | 1.94755E-69 | 0.477113 | 0.712 | 0.529 | 3.68555E-65 |
| Terminal | TEX41 | 4.7993E-119 | 0.476582 | 0.306 | 0.095 | 9.0821E-115 |
| Terminal | KCNT2 | 1.3325E-118 | 0.472248 | 0.22 | 0.05 | 2.5217E-114 |
| Terminal | HES1 | 2.218E-53 | 0.470883 | 0.468 | 0.288 | 4.19735E-49 |
| Terminal | FGL2 | 2.98223E-13 | 0.46842 | 0.643 | 0.563 | 5.64357E-09 |
| Terminal | PHLDA1 | 9.11067E-71 | 0.467445 | 0.394 | 0.2 | 1.7241E-66 |
| Terminal | NEAT1 | 6.26024E-39 | 0.467187 | 0.866 | 0.768 | 1.18469E-34 |
| Terminal | TENT5A | 1.30942E-64 | 0.4634 | 0.483 | 0.286 | 2.47795E-60 |
| Terminal | PDGFRB | 8.59617E-57 | 0.460784 | 0.598 | 0.4 | 1.62674E-52 |
| Terminal | CTSS | 3.1767E-110 | 0.459947 | 0.197 | 0.043 | 6.0115E-106 |
| Terminal | LSP1 | 5.5148E-136 | 0.456409 | 0.212 | 0.039 | 1.0436E-131 |
| Terminal | CDC42EP5 | 9.25171E-77 | 0.453531 | 0.557 | 0.33 | 1.75079E-72 |

| | | | | | | |
|---|---|---|---|---|---|---|
| Terminal | ARL4C | 2.4418E-103 | 0.452124 | 0.217 | 0.057 | 4.6209E-99 |
| Terminal | ARF4 | 5.00554E-69 | 0.451886 | 0.786 | 0.625 | 9.47248E-65 |
| Terminal | C1GALT1 | 5.54245E-80 | 0.449444 | 0.289 | 0.116 | 1.04885E-75 |
| Terminal | ARHGAP18 | 7.1788E-114 | 0.449404 | 0.305 | 0.099 | 1.3585E-109 |
| Terminal | DNAJB1 | 4.56655E-24 | 0.447752 | 0.407 | 0.304 | 8.64175E-20 |
| Terminal | OLFML3 | 6.13821E-58 | 0.446272 | 0.342 | 0.179 | 1.16159E-53 |
| Terminal | C1QTNF1 | 3.36668E-41 | 0.444269 | 0.582 | 0.426 | 6.3711E-37 |
| Terminal | S100A11 | 3.1075E-105 | 0.444264 | 0.992 | 0.964 | 5.8806E-101 |
| Terminal | OSTC | 1.07443E-67 | 0.444231 | 0.76 | 0.597 | 2.03325E-63 |
| Terminal | TMOD3 | 1.77084E-74 | 0.443743 | 0.55 | 0.326 | 3.35113E-70 |
| Terminal | P4HA1 | 6.51644E-77 | 0.443004 | 0.419 | 0.215 | 1.23317E-72 |
| Terminal | JUN | 7.84167E-15 | 0.442244 | 0.808 | 0.797 | 1.48396E-10 |
| Terminal | PLSCR1 | 5.25192E-68 | 0.441386 | 0.431 | 0.232 | 9.93874E-64 |
| Terminal | CD68 | 7.8158E-144 | 0.440788 | 0.243 | 0.05 | 1.4791E-139 |
| Terminal | DAD1 | 7.0211E-105 | 0.440003 | 0.929 | 0.831 | 1.3287E-100 |
| Terminal | IFITM3 | 2.75804E-93 | 0.439367 | 0.991 | 0.977 | 5.21931E-89 |
| Terminal | PDIA6 | 3.07882E-58 | 0.438222 | 0.743 | 0.597 | 5.82636E-54 |
| Terminal | CD63 | 4.5463E-189 | 0.436345 | 0.997 | 0.993 | 8.6034E-185 |
| Terminal | COPZ2 | 4.11231E-45 | 0.436108 | 0.695 | 0.561 | 7.78214E-41 |
| Terminal | STAT1 | 1.28078E-57 | 0.434093 | 0.342 | 0.176 | 2.42374E-53 |
| Terminal | A4GALT | 2.97656E-74 | 0.432312 | 0.376 | 0.183 | 5.63285E-70 |
| Terminal | ANKRD28 | 9.88554E-78 | 0.431709 | 0.283 | 0.113 | 1.87074E-73 |
| Terminal | DKK3 | 1.02918E-49 | 0.429892 | 0.908 | 0.818 | 1.94762E-45 |
| Terminal | COLEC11 | 3.83364E-78 | 0.429217 | 0.112 | 0.018 | 7.25479E-74 |
| Terminal | SLC20A1 | 2.77451E-76 | 0.428346 | 0.208 | 0.068 | 5.25048E-72 |
| Terminal | VMP1 | 2.88575E-57 | 0.428095 | 0.555 | 0.357 | 5.461E-53 |
| Terminal | C4orf48 | 2.9249E-128 | 0.426702 | 0.214 | 0.043 | 5.535E-124 |
| Terminal | CMTM3 | 4.1898E-111 | 0.426667 | 0.286 | 0.09 | 7.9288E-107 |
| Terminal | LTC4S | 3.1384E-106 | 0.425767 | 0.326 | 0.114 | 5.9391E-102 |
| Terminal | THBS1 | 3.78377E-50 | 0.425313 | 0.208 | 0.087 | 7.1604E-46 |
| Terminal | SH3KBP1 | 5.3359E-101 | 0.425214 | 0.281 | 0.093 | 1.00976E-96 |
| Terminal | LMAN1 | 3.08813E-58 | 0.424159 | 0.667 | 0.49 | 5.84397E-54 |
| Terminal | CSF1 | 6.8153E-107 | 0.423508 | 0.249 | 0.071 | 1.2897E-102 |
| Terminal | ANGPTL2 | 2.88548E-72 | 0.421358 | 0.247 | 0.096 | 5.46048E-68 |
| Terminal | HAPLN1 | 2.89376E-32 | 0.41868 | 0.192 | 0.097 | 5.47614E-28 |
| Terminal | FST | 2.23928E-63 | 0.415386 | 0.206 | 0.075 | 4.23761E-59 |
| Terminal | BACE2 | 3.10887E-62 | 0.4152 | 0.467 | 0.277 | 5.88323E-58 |
| Terminal | TGFB1 | 1.32126E-44 | 0.414828 | 0.523 | 0.361 | 2.50035E-40 |
| Terminal | NME2 | 1.0839E-112 | 0.412609 | 0.974 | 0.945 | 2.0511E-108 |
| Terminal | CCPG1 | 5.02264E-67 | 0.412505 | 0.809 | 0.652 | 9.50485E-63 |
| Terminal | CAPG | 1.00997E-39 | 0.412127 | 0.287 | 0.161 | 1.91126E-35 |
| Terminal | KLF4 | 6.45683E-82 | 0.412104 | 0.284 | 0.109 | 1.22189E-77 |
| Terminal | PLD3 | 5.69256E-59 | 0.411896 | 0.63 | 0.439 | 1.07726E-54 |
| Terminal | CYB5A | 1.28322E-40 | 0.408792 | 0.538 | 0.389 | 2.42836E-36 |
| Terminal | RND3 | 6.28103E-29 | 0.408012 | 0.395 | 0.274 | 1.18862E-24 |
| Terminal | F3 | 3.12949E-44 | 0.407586 | 0.214 | 0.1 | 5.92225E-40 |
| Terminal | THBD | 1.08297E-31 | 0.406384 | 0.229 | 0.125 | 2.0494E-27 |
| Terminal | LRP1 | 3.0726E-37 | 0.403025 | 0.724 | 0.579 | 5.81459E-33 |
| Terminal | LMNA | 4.76066E-72 | 0.40286 | 0.957 | 0.894 | 9.00908E-68 |
| Terminal | NPDC1 | 3.61557E-62 | 0.402488 | 0.581 | 0.375 | 6.8421E-58 |
| Terminal | COL4A1 | 5.96273E-22 | 0.401188 | 0.621 | 0.512 | 1.12839E-17 |
| Terminal | EVA1B | 2.47656E-56 | 0.4005 | 0.713 | 0.537 | 4.68665E-52 |
| Terminal | NCOA7 | 5.46542E-53 | 0.400349 | 0.41 | 0.236 | 1.03428E-48 |
| Terminal | PLAGL1 | 2.13502E-49 | 0.397625 | 0.356 | 0.199 | 4.04032E-45 |
| Terminal | SRPX | 1.39357E-50 | 0.396974 | 0.221 | 0.098 | 2.63718E-46 |
| Terminal | VAMP5 | 5.14844E-44 | 0.396572 | 0.676 | 0.523 | 9.7429E-40 |
| Terminal | SELENOP | 2.88836E-15 | 0.395535 | 0.412 | 0.322 | 5.46594E-11 |
| Terminal | CHST1 | 2.3206E-110 | 0.395018 | 0.19 | 0.04 | 4.3916E-106 |

| | | | | | | |
|---|---|---|---|---|---|---|
| Terminal | MYC | 3.8541E-78 | 0.39423 | 0.207 | 0.064 | 7.2935E-74 |
| Terminal | SDF4 | 7.40095E-62 | 0.394049 | 0.637 | 0.44 | 1.40056E-57 |
| Terminal | RUNX1 | 1.10922E-91 | 0.393011 | 0.239 | 0.075 | 2.09909E-87 |
| Terminal | SERPING1 | 3.20615E-59 | 0.392902 | 0.938 | 0.875 | 6.06731E-55 |
| Terminal | GRN | 1.06814E-49 | 0.392478 | 0.555 | 0.382 | 2.02134E-45 |
| Terminal | SPATS2L | 5.84736E-45 | 0.391184 | 0.595 | 0.431 | 1.10655E-40 |
| Terminal | IL33 | 2.88631E-45 | 0.391116 | 0.395 | 0.234 | 5.46204E-41 |
| Terminal | TNFAIP2 | 2.07346E-90 | 0.390595 | 0.154 | 0.031 | 3.92382E-86 |
| Terminal | MFAP2 | 1.48697E-50 | 0.390275 | 0.288 | 0.143 | 2.81393E-46 |
| Terminal | GALNT1 | 1.97497E-39 | 0.386544 | 0.335 | 0.203 | 3.73743E-35 |
| Terminal | ODF2L | 1.69543E-79 | 0.386392 | 0.31 | 0.128 | 3.20843E-75 |
| Terminal | EPHA3 | 5.27075E-90 | 0.383564 | 0.187 | 0.047 | 9.97437E-86 |
| Terminal | CD200 | 6.39467E-43 | 0.383244 | 0.385 | 0.232 | 1.21013E-38 |
| Terminal | P4HA2 | 1.0891E-63 | 0.382515 | 0.395 | 0.211 | 2.06102E-59 |
| Terminal | PPA1 | 1.29231E-43 | 0.38198 | 0.678 | 0.52 | 2.44556E-39 |
| Terminal | TMEM119 | 9.2856E-139 | 0.381766 | 0.121 | 0.005 | 1.7572E-134 |
| Terminal | TWIST1 | 1.4006E-64 | 0.381754 | 0.493 | 0.282 | 2.65049E-60 |
| Terminal | PTTG1IP | 2.18015E-38 | 0.380796 | 0.722 | 0.579 | 4.12572E-34 |
| Terminal | CALR | 3.62778E-52 | 0.37976 | 0.89 | 0.8 | 6.86522E-48 |
| Terminal | APCDD1 | 7.64604E-51 | 0.379461 | 0.121 | 0.035 | 1.44694E-46 |
| Terminal | SLC2A3 | 1.01006E-82 | 0.379214 | 0.207 | 0.062 | 1.91144E-78 |
| Terminal | EFEMP2 | 1.11362E-48 | 0.377586 | 0.703 | 0.544 | 2.10741E-44 |
| Terminal | IL1R1 | 1.79462E-58 | 0.377558 | 0.41 | 0.226 | 3.39613E-54 |
| Terminal | TPBG | 5.41056E-79 | 0.376336 | 0.305 | 0.125 | 1.02389E-74 |
| Terminal | OLFML2B | 1.61858E-45 | 0.375512 | 0.359 | 0.205 | 3.063E-41 |
| Terminal | ADD3 | 1.5404E-63 | 0.374077 | 0.392 | 0.206 | 2.91506E-59 |
| Terminal | CRLF1 | 2.00952E-40 | 0.373939 | 0.132 | 0.047 | 3.80282E-36 |
| Terminal | TMED9 | 2.85892E-50 | 0.373331 | 0.747 | 0.598 | 5.41022E-46 |
| Terminal | SELENOF | 7.83008E-54 | 0.372789 | 0.67 | 0.486 | 1.48176E-49 |
| Terminal | GOLM1 | 5.5414E-121 | 0.368089 | 0.181 | 0.031 | 1.0487E-116 |
| Terminal | HM13 | 9.91294E-51 | 0.367814 | 0.556 | 0.376 | 1.87592E-46 |
| Terminal | GUSB | 3.07301E-67 | 0.366093 | 0.351 | 0.17 | 5.81536E-63 |
| Terminal | SMC4 | 1.20566E-47 | 0.365005 | 0.298 | 0.156 | 2.28158E-43 |
| Terminal | TMEM258 | 5.35166E-56 | 0.364951 | 0.845 | 0.719 | 1.01275E-51 |
| Terminal | SSR2 | 1.64418E-56 | 0.362626 | 0.838 | 0.717 | 3.11145E-52 |
| Terminal | TUBA1B | 6.26062E-38 | 0.362385 | 0.936 | 0.895 | 1.18476E-33 |
| Terminal | ABCA6 | 8.6673E-109 | 0.361901 | 0.153 | 0.024 | 1.6402E-104 |
| Terminal | CLU | 5.33563E-42 | 0.361694 | 0.798 | 0.905 | 1.00971E-37 |
| Terminal | ANXA5 | 1.77518E-54 | 0.36126 | 0.939 | 0.885 | 3.35934E-50 |
| Terminal | ARL6IP5 | 2.22785E-57 | 0.361038 | 0.946 | 0.897 | 4.21599E-53 |
| Terminal | SLIT2 | 8.48836E-66 | 0.359909 | 0.209 | 0.075 | 1.60634E-61 |
| Terminal | IFI44L | 1.07139E-65 | 0.359552 | 0.173 | 0.053 | 2.02749E-61 |
| Terminal | CTSF | 2.73296E-41 | 0.359471 | 0.639 | 0.484 | 5.17185E-37 |
| Terminal | CYP7B1 | 2.0855E-123 | 0.359283 | 0.178 | 0.028 | 3.9466E-119 |
| Terminal | HSPA5 | 3.68107E-43 | 0.358104 | 0.793 | 0.656 | 6.96605E-39 |
| Terminal | HEXA | 5.57462E-51 | 0.357381 | 0.462 | 0.288 | 1.05494E-46 |
| Terminal | CEMIP | 2.3237E-117 | 0.354114 | 0.106 | 0.005 | 4.3974E-113 |
| Terminal | PYCR1 | 1.2491E-163 | 0.353519 | 0.181 | 0.018 | 2.3638E-159 |
| Terminal | CCN2 | 4.28216E-34 | 0.353236 | 0.86 | 0.807 | 8.10356E-30 |
| Terminal | PIGT | 1.8035E-45 | 0.350605 | 0.545 | 0.376 | 3.41295E-41 |
| Terminal | PRELP | 1.17908E-05 | 0.350385 | 0.782 | 0.739 | 0.223129573 |
| Terminal | SLC29A1 | 3.1584E-29 | 0.349066 | 0.359 | 0.235 | 5.97695E-25 |
| Terminal | UGCG | 2.63716E-41 | 0.348728 | 0.272 | 0.143 | 4.99057E-37 |
| Terminal | SRM | 4.33845E-42 | 0.348685 | 0.588 | 0.427 | 8.21009E-38 |
| Terminal | REXO2 | 8.67508E-40 | 0.34831 | 0.738 | 0.607 | 1.64167E-35 |
| Terminal | ATRAID | 3.33663E-51 | 0.346651 | 0.801 | 0.667 | 6.31424E-47 |
| Terminal | IL6ST | 1.33242E-41 | 0.345212 | 0.676 | 0.513 | 2.52147E-37 |
| Terminal | CYTOR | 3.69498E-57 | 0.344356 | 0.495 | 0.298 | 6.99237E-53 |

| | | | | | | |
|---|---|---|---|---|---|---|
| Terminal | ENG | 6.71806E-60 | 0.344216 | 0.63 | 0.417 | 1.27132E-55 |
| Terminal | KCNQ1OT1 | 4.64459E-72 | 0.343587 | 0.168 | 0.047 | 8.78942E-68 |
| Terminal | RAB31 | 9.95213E-36 | 0.343255 | 0.517 | 0.367 | 1.88334E-31 |
| Terminal | MEDAG | 2.3264E-33 | 0.342844 | 0.274 | 0.158 | 4.40247E-29 |
| Terminal | PHLDB1 | 8.70661E-43 | 0.342625 | 0.281 | 0.149 | 1.64764E-38 |
| Terminal | TNFRSF1A | 7.35684E-44 | 0.340635 | 0.57 | 0.397 | 1.39221E-39 |
| Terminal | IER5L | 1.75071E-44 | 0.338454 | 0.456 | 0.284 | 3.31304E-40 |
| Terminal | SLC39A7 | 7.96983E-49 | 0.337292 | 0.431 | 0.261 | 1.50821E-44 |
| Terminal | PYCARD | 8.69449E-59 | 0.337243 | 0.26 | 0.113 | 1.64534E-54 |
| Terminal | IFIT1 | 7.1696E-23 | 0.337133 | 0.195 | 0.109 | 1.35678E-18 |
| Terminal | GPC1 | 1.81269E-41 | 0.336788 | 0.343 | 0.201 | 3.43034E-37 |
| Terminal | CFI | 2.12255E-55 | 0.335489 | 0.192 | 0.071 | 4.01672E-51 |
| Terminal | YIF1A | 2.41636E-40 | 0.335126 | 0.636 | 0.491 | 4.57271E-36 |
| Terminal | HEXB | 1.19416E-51 | 0.334571 | 0.455 | 0.276 | 2.25983E-47 |
| Terminal | EPB41L2 | 1.01147E-40 | 0.334143 | 0.611 | 0.449 | 1.9141E-36 |
| Terminal | BHLHE40 | 1.97367E-60 | 0.333876 | 0.214 | 0.081 | 3.73498E-56 |
| Terminal | CIB1 | 1.11408E-47 | 0.332741 | 0.707 | 0.539 | 2.10828E-43 |
| Terminal | CPQ | 3.83654E-39 | 0.332678 | 0.691 | 0.535 | 7.26026E-35 |
| Terminal | ITPR2 | 1.15407E-62 | 0.332567 | 0.248 | 0.101 | 2.18397E-58 |
| Terminal | SELENOS | 1.41071E-42 | 0.33213 | 0.77 | 0.634 | 2.66963E-38 |
| Terminal | SRGN | 9.63439E-29 | 0.331684 | 0.436 | 0.3 | 1.82321E-24 |
| Terminal | CD81 | 1.19585E-77 | 0.331279 | 0.995 | 0.983 | 2.26303E-73 |
| Terminal | GLT8D1 | 3.92084E-56 | 0.330357 | 0.361 | 0.19 | 7.4198E-52 |
| Terminal | VKORC1 | 1.28688E-46 | 0.328932 | 0.815 | 0.704 | 2.4353E-42 |
| Terminal | AXL | 4.10245E-43 | 0.326794 | 0.471 | 0.302 | 7.76348E-39 |
| Terminal | LDB2 | 6.197E-103 | 0.326428 | 0.151 | 0.025 | 1.17272E-98 |
| Terminal | HDLBP | 2.05315E-42 | 0.32582 | 0.638 | 0.469 | 3.88537E-38 |
| Terminal | GLT8D2 | 4.74375E-42 | 0.325275 | 0.523 | 0.356 | 8.97708E-38 |
| Terminal | SPOCK1 | 1.56241E-88 | 0.324029 | 0.149 | 0.03 | 2.95671E-84 |
| Terminal | CMKLR1 | 5.74909E-71 | 0.32376 | 0.229 | 0.083 | 1.08796E-66 |
| Terminal | TMEM45A | 1.12202E-57 | 0.322589 | 0.374 | 0.198 | 2.12331E-53 |
| Terminal | MMP14 | 1.18227E-25 | 0.322543 | 0.271 | 0.17 | 2.23732E-21 |
| Terminal | TCTN1 | 1.27743E-68 | 0.32238 | 0.269 | 0.109 | 2.41741E-64 |
| Terminal | NME1 | 6.3159E-45 | 0.321761 | 0.462 | 0.293 | 1.19522E-40 |
| Terminal | PODNL1 | 1.11249E-95 | 0.321368 | 0.2 | 0.051 | 2.10527E-91 |
| Terminal | IFIT2 | 6.20037E-21 | 0.321302 | 0.16 | 0.085 | 1.17336E-16 |
| Terminal | HAPLN3 | 2.43255E-47 | 0.321046 | 0.275 | 0.137 | 4.60336E-43 |
| Terminal | SPHK1 | 3.03191E-85 | 0.319919 | 0.149 | 0.031 | 5.73758E-81 |
| Terminal | TAPBP | 4.01122E-42 | 0.319066 | 0.433 | 0.268 | 7.59083E-38 |
| Terminal | ITGA10 | 2.5958E-30 | 0.318711 | 0.455 | 0.319 | 4.91229E-26 |
| Terminal | XAF1 | 1.96693E-66 | 0.318407 | 0.219 | 0.079 | 3.72222E-62 |
| Terminal | XBP1 | 6.08265E-40 | 0.317642 | 0.577 | 0.406 | 1.15108E-35 |
| Terminal | COL8A1 | 1.16224E-09 | 0.317494 | 0.661 | 0.579 | 2.19941E-05 |
| Terminal | SP100 | 2.20357E-35 | 0.316869 | 0.593 | 0.441 | 4.17004E-31 |
| Terminal | AC020916. | 4.99877E-37 | 0.316463 | 0.307 | 0.175 | 9.45967E-33 |
| Terminal | MCL1 | 4.71283E-30 | 0.316259 | 0.516 | 0.376 | 8.91856E-26 |
| Terminal | EDIL3 | 9.0031E-36 | 0.316175 | 0.849 | 0.731 | 1.70375E-31 |
| Terminal | RCN1 | 1.72411E-37 | 0.315893 | 0.59 | 0.429 | 3.26271E-33 |
| Terminal | PXK | 1.00596E-59 | 0.314919 | 0.264 | 0.115 | 1.90369E-55 |
| Terminal | TMEM167 | 2.36338E-39 | 0.31339 | 0.552 | 0.388 | 4.47245E-35 |
| Terminal | APOL3 | 4.43602E-81 | 0.31328 | 0.18 | 0.048 | 8.39472E-77 |
| Terminal | KLF6 | 8.04548E-10 | 0.312838 | 0.736 | 0.682 | 1.52253E-05 |
| Terminal | PSAP | 9.27697E-48 | 0.311923 | 0.908 | 0.812 | 1.75557E-43 |
| Terminal | TMED3 | 1.09052E-36 | 0.311003 | 0.481 | 0.329 | 2.0637E-32 |
| Terminal | TMEM204 | 1.89124E-39 | 0.310788 | 0.42 | 0.265 | 3.57898E-35 |
| Terminal | ACAN | 2.49588E-07 | 0.309472 | 0.103 | 0.069 | 0.0047232 |
| Terminal | ELK3 | 3.97789E-67 | 0.309067 | 0.261 | 0.105 | 7.52777E-63 |
| Terminal | YIPF2 | 9.08992E-49 | 0.308676 | 0.427 | 0.253 | 1.72018E-44 |

| | | | | | | |
|---|---|---|---|---|---|---|
| Terminal | ABRACL | 2.01004E-50 | 0.30817 | 0.262 | 0.124 | 3.80381E-46 |
| Terminal | PPP1R15A | 4.22273E-24 | 0.307812 | 0.605 | 0.488 | 7.99109E-20 |
| Terminal | APOL1 | 2.64246E-65 | 0.307359 | 0.189 | 0.063 | 5.00059E-61 |
| Terminal | AGTRAP | 3.68604E-53 | 0.307135 | 0.343 | 0.18 | 6.97546E-49 |
| Terminal | LAMB2 | 6.57842E-12 | 0.3071 | 0.479 | 0.405 | 1.2449E-07 |
| Terminal | ERLEC1 | 5.35398E-37 | 0.306548 | 0.684 | 0.535 | 1.01319E-32 |
| Terminal | KRTCAP2 | 9.69456E-51 | 0.306016 | 0.909 | 0.82 | 1.8346E-46 |
| Terminal | IFNGR1 | 8.24026E-37 | 0.305835 | 0.471 | 0.314 | 1.55939E-32 |
| Terminal | SERTAD4 | 6.90391E-63 | 0.305401 | 0.211 | 0.077 | 1.3065E-58 |
| Terminal | CHSY1 | 1.26079E-50 | 0.305255 | 0.264 | 0.125 | 2.38592E-46 |
| Terminal | BIRC3 | 7.83991E-50 | 0.305133 | 0.136 | 0.043 | 1.48363E-45 |
| Terminal | LGALS3 | 3.49151E-40 | 0.304599 | 0.936 | 0.883 | 6.60734E-36 |
| Terminal | DAAM1 | 3.00597E-20 | 0.304104 | 0.546 | 0.437 | 5.68849E-16 |
| Terminal | KCNN4 | 5.353E-124 | 0.303531 | 0.111 | 0.005 | 1.013E-119 |
| Terminal | TMED10 | 2.83112E-44 | 0.303428 | 0.751 | 0.599 | 5.35762E-40 |
| Terminal | FOXS1 | 4.60288E-24 | 0.302771 | 0.252 | 0.155 | 8.7105E-20 |
| Terminal | ASAP1 | 2.17213E-51 | 0.302758 | 0.31 | 0.158 | 4.11054E-47 |
| Terminal | EVL | 1.62576E-37 | 0.302702 | 0.277 | 0.154 | 3.07659E-33 |
| Terminal | MPZL1 | 1.94958E-47 | 0.302241 | 0.373 | 0.212 | 3.68938E-43 |
| Terminal | BTG1 | 1.61052E-23 | 0.302034 | 0.774 | 0.692 | 3.04776E-19 |
| Terminal | IFT57 | 5.54219E-40 | 0.301982 | 0.493 | 0.33 | 1.0488E-35 |
| Terminal | NUDT4 | 1.2628E-23 | 0.299128 | 0.453 | 0.332 | 2.38972E-19 |
| Terminal | TIMP2 | 2.23713E-34 | 0.299061 | 0.899 | 0.803 | 4.23354E-30 |
| Terminal | CASP4 | 3.15913E-50 | 0.298782 | 0.383 | 0.212 | 5.97834E-46 |
| Terminal | RARRES1 | 4.84035E-48 | 0.298763 | 0.152 | 0.053 | 9.15988E-44 |
| Terminal | JTB | 3.54267E-37 | 0.298488 | 0.71 | 0.573 | 6.70414E-33 |
| Terminal | TRAM1 | 1.12107E-38 | 0.298295 | 0.662 | 0.499 | 2.12152E-34 |
| Terminal | IFI35 | 1.91798E-49 | 0.29813 | 0.31 | 0.159 | 3.62959E-45 |
| Terminal | GDI2 | 9.79828E-36 | 0.297421 | 0.629 | 0.477 | 1.85423E-31 |
| Terminal | ART4 | 1.56358E-47 | 0.297049 | 0.219 | 0.097 | 2.95892E-43 |
| Terminal | COL16A1 | 1.2775E-24 | 0.296421 | 0.342 | 0.233 | 2.41755E-20 |
| Terminal | DAP | 1.91522E-34 | 0.296287 | 0.507 | 0.354 | 3.62435E-30 |
| Terminal | NR2F2 | 3.09838E-13 | 0.295998 | 0.691 | 0.619 | 5.86337E-09 |
| Terminal | S100A3 | 3.85402E-74 | 0.295923 | 0.187 | 0.055 | 7.29335E-70 |
| Terminal | SLC38A2 | 8.84663E-23 | 0.295865 | 0.647 | 0.519 | 1.67414E-18 |
| Terminal | LOX | 9.55598E-56 | 0.295807 | 0.223 | 0.092 | 1.80837E-51 |
| Terminal | B4GALT1 | 4.04538E-41 | 0.295531 | 0.316 | 0.175 | 7.65548E-37 |
| Terminal | PDIA4 | 2.69799E-48 | 0.294926 | 0.343 | 0.186 | 5.10568E-44 |
| Terminal | SEMA5A | 1.47188E-39 | 0.294356 | 0.273 | 0.146 | 2.78538E-35 |
| Terminal | PSMB8 | 4.90381E-39 | 0.293723 | 0.487 | 0.318 | 9.27996E-35 |
| Terminal | ALDH1A2 | 1.27499E-60 | 0.292859 | 0.152 | 0.045 | 2.4128E-56 |
| Terminal | ETHE1 | 1.14971E-35 | 0.292743 | 0.428 | 0.279 | 2.17571E-31 |
| Terminal | LPAR6 | 6.39643E-60 | 0.291872 | 0.216 | 0.082 | 1.21046E-55 |
| Terminal | NFKBIZ | 1.19998E-44 | 0.29155 | 0.274 | 0.137 | 2.27085E-40 |
| Terminal | H3F3B | 2.67855E-74 | 0.291021 | 0.995 | 0.983 | 5.06889E-70 |
| Terminal | STK17B | 1.60363E-50 | 0.290988 | 0.22 | 0.093 | 3.03471E-46 |
| Terminal | ZFP36L2 | 6.49138E-21 | 0.29009 | 0.711 | 0.6 | 1.22843E-16 |
| Terminal | DUXAP8 | 1.30086E-65 | 0.289667 | 0.173 | 0.053 | 2.46175E-61 |
| Terminal | HGF | 3.65924E-36 | 0.289533 | 0.284 | 0.156 | 6.92474E-32 |
| Terminal | DNPH1 | 1.21911E-48 | 0.289393 | 0.404 | 0.229 | 2.30704E-44 |
| Terminal | SEC61B | 3.36058E-40 | 0.288395 | 0.902 | 0.847 | 6.35957E-36 |
| Terminal | DECR1 | 1.55487E-36 | 0.288392 | 0.428 | 0.278 | 2.94243E-32 |
| Terminal | CD82 | 1.30951E-41 | 0.288391 | 0.296 | 0.16 | 2.47813E-37 |
| Terminal | CD99 | 1.97968E-39 | 0.288208 | 0.947 | 0.921 | 3.74635E-35 |
| Terminal | PLXDC2 | 1.34357E-27 | 0.288055 | 0.553 | 0.413 | 2.54258E-23 |
| Terminal | LTBP4 | 3.17367E-16 | 0.287602 | 0.721 | 0.633 | 6.00585E-12 |
| Terminal | ANK3 | 2.75527E-32 | 0.287167 | 0.211 | 0.11 | 5.21407E-28 |
| Terminal | APOL6 | 6.35301E-57 | 0.28677 | 0.197 | 0.074 | 1.20224E-52 |

| | | | | | | |
|---|---|---|---|---|---|---|
| Terminal | AHR | 1.23474E-36 | 0.286447 | 0.358 | 0.216 | 2.33663E-32 |
| Terminal | ANGPTL4 | 8.83843E-50 | 0.285477 | 0.132 | 0.04 | 1.67258E-45 |
| Terminal | MAP4K4 | 9.67433E-32 | 0.28501 | 0.346 | 0.217 | 1.83077E-27 |
| Terminal | CTSB | 1.23921E-24 | 0.284939 | 0.556 | 0.421 | 2.34509E-20 |
| Terminal | PHACTR2 | 1.89933E-23 | 0.284698 | 0.522 | 0.398 | 3.59429E-19 |
| Terminal | EMILIN1 | 1.18673E-14 | 0.284557 | 0.548 | 0.459 | 2.24577E-10 |
| Terminal | TSPAN15 | 7.8285E-101 | 0.284223 | 0.125 | 0.015 | 1.48146E-96 |
| Terminal | BCAT1 | 2.0707E-99 | 0.28364 | 0.141 | 0.022 | 3.91855E-95 |
| Terminal | P3H4 | 8.21843E-65 | 0.282961 | 0.225 | 0.084 | 1.55526E-60 |
| Terminal | WARS | 1.75147E-29 | 0.281647 | 0.152 | 0.071 | 3.31448E-25 |
| Terminal | CHCHD10 | 7.18624E-34 | 0.281267 | 0.746 | 0.612 | 1.35992E-29 |
| Terminal | GBP4 | 7.60701E-35 | 0.280279 | 0.103 | 0.034 | 1.43955E-30 |
| Terminal | CASC4 | 7.27578E-34 | 0.280119 | 0.518 | 0.366 | 1.37687E-29 |
| Terminal | TMEM9B | 2.43854E-46 | 0.280059 | 0.287 | 0.147 | 4.61469E-42 |
| Terminal | TNFAIP8L3 | 2.69563E-50 | 0.279652 | 0.252 | 0.117 | 5.10122E-46 |
| Terminal | INTS6 | 1.82925E-25 | 0.279582 | 0.268 | 0.164 | 3.46168E-21 |
| Terminal | TEX9 | 8.1238E-70 | 0.279016 | 0.166 | 0.046 | 1.53735E-65 |
| Terminal | EIF4A1 | 4.09191E-32 | 0.27891 | 0.891 | 0.811 | 7.74354E-28 |
| Terminal | OSMR | 1.27355E-40 | 0.277473 | 0.267 | 0.139 | 2.41007E-36 |
| Terminal | FTH1 | 1.0116E-22 | 0.277215 | 1 | 0.999 | 1.91435E-18 |
| Terminal | ARHGAP22 | 5.24337E-87 | 0.276709 | 0.159 | 0.035 | 9.92255E-83 |
| Terminal | RECQL | 5.44574E-35 | 0.276499 | 0.434 | 0.286 | 1.03055E-30 |
| Terminal | GNG11 | 3.4528E-14 | 0.276346 | 0.625 | 0.539 | 6.53408E-10 |
| Terminal | C5orf15 | 5.61688E-41 | 0.275923 | 0.324 | 0.182 | 1.06294E-36 |
| Terminal | PRAF2 | 5.8924E-40 | 0.275547 | 0.396 | 0.239 | 1.11508E-35 |
| Terminal | CRELD2 | 3.5012E-39 | 0.274492 | 0.396 | 0.244 | 6.62567E-35 |
| Terminal | NUCB1 | 1.01723E-32 | 0.274467 | 0.71 | 0.568 | 1.925E-28 |
| Terminal | OSBPL8 | 1.8958E-28 | 0.274452 | 0.56 | 0.416 | 3.58762E-24 |
| Terminal | GALK1 | 8.33996E-47 | 0.274055 | 0.281 | 0.141 | 1.57825E-42 |
| Terminal | MSC | 6.8326E-35 | 0.273738 | 0.318 | 0.186 | 1.293E-30 |
| Terminal | C1GALT1C | 3.22865E-43 | 0.273403 | 0.377 | 0.219 | 6.1099E-39 |
| Terminal | CHID1 | 5.47177E-31 | 0.272705 | 0.474 | 0.331 | 1.03548E-26 |
| Terminal | GPX7 | 1.00255E-49 | 0.272342 | 0.21 | 0.089 | 1.89723E-45 |
| Terminal | PDGFA | 5.44871E-16 | 0.271223 | 0.567 | 0.471 | 1.03111E-11 |
| Terminal | RPN2 | 4.22272E-25 | 0.270717 | 0.619 | 0.496 | 7.99107E-21 |
| Terminal | LGMN | 1.63743E-31 | 0.268918 | 0.424 | 0.285 | 3.09867E-27 |
| Terminal | FNDC1 | 1.49749E-69 | 0.265075 | 0.12 | 0.025 | 2.83386E-65 |
| Terminal | NIBAN2 | 1.17914E-40 | 0.264677 | 0.25 | 0.127 | 2.2314E-36 |
| Terminal | PLAT | 4.33954E-51 | 0.264564 | 0.166 | 0.06 | 8.21214E-47 |
| Terminal | NT5E | 5.369E-38 | 0.26452 | 0.31 | 0.176 | 1.01603E-33 |
| Terminal | MIR4435-2 | 3.52958E-50 | 0.264392 | 0.256 | 0.119 | 6.67937E-46 |
| Terminal | IL15RA | 1.26792E-56 | 0.263972 | 0.173 | 0.059 | 2.39941E-52 |
| Terminal | SEC31A | 9.48195E-28 | 0.263807 | 0.637 | 0.497 | 1.79436E-23 |
| Terminal | MAFB | 1.40159E-19 | 0.263772 | 0.238 | 0.149 | 2.65237E-15 |
| Terminal | ATP1A1 | 4.04719E-25 | 0.263603 | 0.451 | 0.328 | 7.6589E-21 |
| Terminal | BEND6 | 3.78251E-81 | 0.26293 | 0.142 | 0.029 | 7.15803E-77 |
| Terminal | CCND1 | 5.61351E-35 | 0.261464 | 0.561 | 0.4 | 1.0623E-30 |
| Terminal | DPYSL2 | 4.53914E-26 | 0.260849 | 0.552 | 0.415 | 8.58987E-22 |
| Terminal | AKR1A1 | 1.32288E-28 | 0.260704 | 0.605 | 0.465 | 2.50342E-24 |
| Terminal | PSMB10 | 4.86325E-29 | 0.260086 | 0.342 | 0.216 | 9.20321E-25 |
| Terminal | UBXN4 | 1.40048E-32 | 0.260081 | 0.778 | 0.655 | 2.65027E-28 |
| Terminal | SERTAD1 | 1.04481E-25 | 0.259704 | 0.469 | 0.339 | 1.97719E-21 |
| Terminal | MAP1A | 9.12247E-38 | 0.259621 | 0.274 | 0.146 | 1.72634E-33 |
| Terminal | EEA1 | 4.46707E-25 | 0.25949 | 0.661 | 0.527 | 8.45348E-21 |
| Terminal | MX1 | 4.52082E-15 | 0.258795 | 0.234 | 0.156 | 8.5552E-11 |
| Terminal | ANPEP | 6.6369E-108 | 0.258477 | 0.114 | 0.01 | 1.256E-103 |
| Terminal | PLGRKT | 2.57161E-42 | 0.258029 | 0.267 | 0.136 | 4.86651E-38 |
| Terminal | LDHA | 1.44055E-27 | 0.257923 | 0.934 | 0.879 | 2.7261E-23 |

| | | | | | | |
|---|---|---|---|---|---|---|
| Terminal | FRMD4B | 1.21387E-33 | 0.257658 | 0.257 | 0.14 | 2.29713E-29 |
| Terminal | CHIC2 | 2.31604E-41 | 0.257236 | 0.283 | 0.15 | 4.38288E-37 |
| Terminal | HIF1A | 7.81348E-25 | 0.256585 | 0.41 | 0.285 | 1.47862E-20 |
| Terminal | TMEM179 | 2.25753E-33 | 0.256385 | 0.548 | 0.387 | 4.27215E-29 |
| Terminal | ITGBL1 | 1.4149E-19 | 0.255158 | 0.696 | 0.576 | 2.67755E-15 |
| Terminal | GAA | 4.12884E-35 | 0.254834 | 0.277 | 0.153 | 7.81341E-31 |
| Terminal | LAMP2 | 4.19635E-30 | 0.2548 | 0.451 | 0.313 | 7.94118E-26 |
| Terminal | LTBP1 | 2.54571E-15 | 0.254787 | 0.882 | 0.819 | 4.8175E-11 |
| Terminal | AKR1C1 | 6.04641E-33 | 0.253471 | 0.281 | 0.162 | 1.14422E-28 |
| Terminal | RBMS1 | 8.84437E-25 | 0.253397 | 0.511 | 0.384 | 1.67371E-20 |
| Terminal | CCDC151 | 1.02659E-49 | 0.253111 | 0.231 | 0.101 | 1.94271E-45 |
| Terminal | ECE1 | 1.02176E-31 | 0.253024 | 0.305 | 0.181 | 1.93358E-27 |
| Terminal | SAMD9L | 5.14722E-50 | 0.253002 | 0.203 | 0.083 | 9.74061E-46 |
| Terminal | CKAP4 | 2.13138E-32 | 0.252736 | 0.338 | 0.208 | 4.03343E-28 |
| Terminal | RASSF8-AS | 2.75544E-45 | 0.252454 | 0.212 | 0.094 | 5.21439E-41 |
| Terminal | GALNT13 | 1.0154E-123 | 0.25228 | 0.101 | 0.003 | 1.9215E-119 |
| Terminal | CNN3 | 1.40312E-30 | 0.251959 | 0.795 | 0.668 | 2.65526E-26 |
| Terminal | SLC14A1 | 4.69127E-29 | 0.251853 | 0.205 | 0.109 | 8.87775E-25 |
| Terminal | HLA-DPB1 | 1.40269E-16 | 0.251133 | 0.323 | 0.23 | 2.65445E-12 |
| Terminal | TRIM22 | 2.02792E-27 | 0.250549 | 0.282 | 0.169 | 3.83763E-23 |
| Terminal | MSX1 | 1.3923E-35 | 0.250225 | 0.226 | 0.115 | 2.63479E-31 |
| Intermediate | KRT17 | 1.3334E-199 | 1.86015 | 0.476 | 0.131 | 2.5232E-195 |
| Intermediate | ASPN | 1.5841E-287 | 1.565559 | 0.937 | 0.553 | 2.9978E-283 |
| Intermediate | KRT18 | 1.17943E-56 | 1.342346 | 0.217 | 0.074 | 2.23196E-52 |
| Intermediate | CCN3 | 4.208E-177 | 1.240326 | 0.924 | 0.7 | 7.9631E-173 |
| Intermediate | OGN | 4.5741E-225 | 1.193635 | 0.982 | 0.849 | 8.656E-221 |
| Intermediate | FHL1 | 1.2053E-220 | 1.126446 | 0.984 | 0.879 | 2.2809E-216 |
| Intermediate | MYOZ2 | 3.5068E-152 | 1.089745 | 0.4 | 0.111 | 6.6363E-148 |
| Intermediate | KRT7 | 8.0158E-148 | 1.064425 | 0.308 | 0.067 | 1.5169E-143 |
| Intermediate | TNFRSF12A | 2.35428E-71 | 1.048039 | 0.708 | 0.528 | 4.45524E-67 |
| Intermediate | PPP1R14A | 5.6556E-199 | 1.026653 | 0.988 | 0.829 | 1.0703E-194 |
| Intermediate | ADH1B | 8.1208E-122 | 0.995398 | 0.748 | 0.393 | 1.5368E-117 |
| Intermediate | CRYAB | 1.56636E-82 | 0.993603 | 0.96 | 0.905 | 2.96417E-78 |
| Intermediate | BDNF | 7.9182E-246 | 0.970547 | 0.28 | 0.026 | 1.4984E-241 |
| Intermediate | CYFIP2 | 2.165E-132 | 0.93945 | 0.523 | 0.205 | 4.0971E-128 |
| Intermediate | CCDC144N | 1.3363E-268 | 0.93827 | 0.325 | 0.034 | 2.5288E-264 |
| Intermediate | ACKR4 | 6.435E-162 | 0.914496 | 0.382 | 0.095 | 1.2178E-157 |
| Intermediate | EFEMP1 | 1.0238E-118 | 0.891081 | 0.939 | 0.752 | 1.9374E-114 |
| Intermediate | INHBA | 4.50822E-91 | 0.870711 | 0.709 | 0.444 | 8.53136E-87 |
| Intermediate | FSTL3 | 1.3775E-123 | 0.845238 | 0.543 | 0.227 | 2.6068E-119 |
| Intermediate | ACTA2 | 2.2589E-150 | 0.842806 | 0.996 | 0.884 | 4.2748E-146 |
| Intermediate | SERPINE1 | 2.0587E-41 | 0.83369 | 0.493 | 0.3 | 3.89588E-37 |
| Intermediate | TCEAL4 | 6.034E-116 | 0.828297 | 0.94 | 0.826 | 1.1419E-111 |
| Intermediate | CCN2 | 1.02363E-36 | 0.821259 | 0.896 | 0.809 | 1.93713E-32 |
| Intermediate | WFDC1 | 5.2012E-106 | 0.817656 | 0.48 | 0.199 | 9.8427E-102 |
| Intermediate | AKAP12 | 3.59113E-78 | 0.811495 | 0.539 | 0.283 | 6.79586E-74 |
| Intermediate | CREB5 | 3.4317E-110 | 0.806912 | 0.337 | 0.101 | 6.4942E-106 |
| Intermediate | NEXN | 2.7805E-114 | 0.804231 | 0.924 | 0.717 | 5.2618E-110 |
| Intermediate | MFAP4 | 1.2138E-117 | 0.792474 | 0.953 | 0.737 | 2.297E-113 |
| Intermediate | FBLN5 | 4.911E-103 | 0.786617 | 0.806 | 0.566 | 9.2935E-99 |
| Intermediate | PDK4 | 2.31799E-25 | 0.771469 | 0.248 | 0.137 | 4.38656E-21 |
| Intermediate | FSTL1 | 5.84523E-69 | 0.759093 | 0.75 | 0.549 | 1.10615E-64 |
| Intermediate | SUGCT | 7.93487E-93 | 0.756997 | 0.739 | 0.46 | 1.50159E-88 |
| Intermediate | CRLF1 | 7.3565E-88 | 0.750066 | 0.207 | 0.047 | 1.39214E-83 |
| Intermediate | CRIM1 | 3.52067E-82 | 0.729314 | 0.806 | 0.556 | 6.66251E-78 |
| Intermediate | TGFBR1 | 4.78545E-83 | 0.722428 | 0.468 | 0.221 | 9.05599E-79 |
| Intermediate | PXDC1 | 3.12087E-80 | 0.714855 | 0.697 | 0.456 | 5.90594E-76 |
| Intermediate | MMP23B | 6.0975E-115 | 0.711619 | 0.454 | 0.16 | 1.1539E-110 |

| | | | | | | |
|---|---|---|---|---|---|---|
| Intermediate | RAB32 | 1.92982E-57 | 0.708402 | 0.538 | 0.34 | 3.652E-53 |
| Intermediate | NPY1R | 9.2045E-136 | 0.689168 | 0.301 | 0.067 | 1.7419E-131 |
| Intermediate | TNFRSF11I | 6.1429E-92 | 0.686302 | 0.645 | 0.346 | 1.16248E-87 |
| Intermediate | MYH10 | 2.76762E-96 | 0.684193 | 0.949 | 0.756 | 5.23744E-92 |
| Intermediate | PRSS23 | 4.42846E-78 | 0.676314 | 0.791 | 0.533 | 8.38041E-74 |
| Intermediate | CNN1 | 2.66221E-65 | 0.660883 | 0.771 | 0.521 | 5.03796E-61 |
| Intermediate | ACTG2 | 5.50953E-21 | 0.660122 | 0.462 | 0.328 | 1.04262E-16 |
| Intermediate | MBNL1-AS | 9.60231E-39 | 0.654014 | 0.536 | 0.356 | 1.81714E-34 |
| Intermediate | GEM | 6.92856E-83 | 0.64232 | 0.859 | 0.639 | 1.31116E-78 |
| Intermediate | ITGBL1 | 1.08915E-80 | 0.638851 | 0.819 | 0.574 | 2.0611E-76 |
| Intermediate | WTIP | 1.02749E-59 | 0.636284 | 0.598 | 0.366 | 1.94442E-55 |
| Intermediate | ELN | 2.89302E-35 | 0.636209 | 0.593 | 0.411 | 5.47476E-31 |
| Intermediate | BAG2 | 7.86963E-45 | 0.619684 | 0.524 | 0.335 | 1.48925E-40 |
| Intermediate | DNAJB4 | 4.22783E-52 | 0.618956 | 0.59 | 0.388 | 8.00074E-48 |
| Intermediate | FILIP1L | 3.78205E-55 | 0.613106 | 0.907 | 0.748 | 7.15715E-51 |
| Intermediate | NDUFA4 | 2.4457E-117 | 0.612611 | 0.978 | 0.936 | 4.6283E-113 |
| Intermediate | KRT8 | 7.67059E-09 | 0.610096 | 0.378 | 0.313 | 0.000145158 |
| Intermediate | NPR3 | 2.062E-124 | 0.606508 | 0.295 | 0.069 | 3.9021E-120 |
| Intermediate | MEGF6 | 1.43233E-82 | 0.605396 | 0.4 | 0.163 | 2.71053E-78 |
| Intermediate | CDH2 | 2.8575E-92 | 0.596814 | 0.387 | 0.144 | 5.40753E-88 |
| Intermediate | CCN1 | 3.65663E-21 | 0.594861 | 0.827 | 0.745 | 6.91981E-17 |
| Intermediate | TMEM47 | 1.16834E-64 | 0.593822 | 0.793 | 0.594 | 2.21096E-60 |
| Intermediate | PDLIM7 | 8.82687E-67 | 0.593044 | 0.829 | 0.657 | 1.6704E-62 |
| Intermediate | MYL12A | 5.26004E-67 | 0.580947 | 0.957 | 0.942 | 9.95411E-63 |
| Intermediate | TPM2 | 2.61126E-75 | 0.580085 | 0.997 | 0.979 | 4.94154E-71 |
| Intermediate | TCEAL3 | 1.7514E-50 | 0.57827 | 0.786 | 0.637 | 3.31434E-46 |
| Intermediate | COL8A1 | 4.15189E-68 | 0.572528 | 0.804 | 0.568 | 7.85704E-64 |
| Intermediate | OMD | 5.5173E-52 | 0.560091 | 0.65 | 0.422 | 1.04409E-47 |
| Intermediate | ACTN4 | 2.40381E-71 | 0.556495 | 0.899 | 0.751 | 4.54897E-67 |
| Intermediate | MAP3K7CL | 1.80065E-83 | 0.554385 | 0.354 | 0.13 | 3.40754E-79 |
| Intermediate | TAGLN | 3.23776E-87 | 0.538749 | 0.998 | 0.982 | 6.12713E-83 |
| Intermediate | TPH1 | 9.57811E-57 | 0.538138 | 0.369 | 0.169 | 1.81256E-52 |
| Intermediate | C5orf46 | 8.09594E-60 | 0.537668 | 0.308 | 0.121 | 1.53208E-55 |
| Intermediate | GLIPR1 | 5.33466E-52 | 0.514073 | 0.78 | 0.586 | 1.00953E-47 |
| Intermediate | PDLIM1 | 6.22143E-37 | 0.513718 | 0.804 | 0.701 | 1.17734E-32 |
| Intermediate | KRT16 | 2.77108E-82 | 0.512732 | 0.112 | 0.013 | 5.24399E-78 |
| Intermediate | TCEAL1 | 3.73461E-39 | 0.503475 | 0.54 | 0.356 | 7.06738E-35 |
| Intermediate | CD151 | 1.4422E-87 | 0.50197 | 0.979 | 0.934 | 2.72921E-83 |
| Intermediate | USP53 | 3.34576E-37 | 0.50191 | 0.479 | 0.305 | 6.33151E-33 |
| Intermediate | TPM1 | 3.46624E-80 | 0.500509 | 0.995 | 0.965 | 6.55952E-76 |
| Intermediate | FN1 | 5.60253E-61 | 0.500114 | 0.989 | 0.899 | 1.06022E-56 |
| Intermediate | TMSB4X | 3.08513E-71 | 0.499205 | 1 | 0.996 | 5.8383E-67 |
| Intermediate | SLMAP | 5.78331E-36 | 0.498102 | 0.588 | 0.407 | 1.09443E-31 |
| Intermediate | VCAN | 3.57977E-48 | 0.495797 | 0.938 | 0.767 | 6.77436E-44 |
| Intermediate | GAS6 | 2.89569E-54 | 0.494088 | 0.932 | 0.858 | 5.4798E-50 |
| Intermediate | PKD2 | 8.80122E-52 | 0.490328 | 0.562 | 0.345 | 1.66554E-47 |
| Intermediate | MORF4L2 | 8.10897E-40 | 0.487527 | 0.765 | 0.618 | 1.53454E-35 |
| Intermediate | ILK | 7.25361E-57 | 0.479041 | 0.872 | 0.763 | 1.37267E-52 |
| Intermediate | PCGF5 | 2.13047E-48 | 0.47885 | 0.619 | 0.407 | 4.0317E-44 |
| Intermediate | OXTR | 9.4839E-145 | 0.474885 | 0.138 | 0.008 | 1.7947E-140 |
| Intermediate | FGF1 | 1.89718E-73 | 0.473936 | 0.302 | 0.107 | 3.59021E-69 |
| Intermediate | GJA1 | 4.30884E-39 | 0.469366 | 0.575 | 0.38 | 8.15404E-35 |
| Intermediate | HSPB1 | 1.1946E-47 | 0.466617 | 0.99 | 0.982 | 2.26066E-43 |
| Intermediate | LINC02762 | 6.17493E-40 | 0.462436 | 0.494 | 0.302 | 1.16854E-35 |
| Intermediate | PPP1R12B | 2.76974E-46 | 0.459772 | 0.809 | 0.575 | 5.24146E-42 |
| Intermediate | ABCA8 | 1.58563E-43 | 0.457104 | 0.173 | 0.059 | 3.00064E-39 |
| Intermediate | FLNA | 7.11288E-50 | 0.453283 | 0.958 | 0.832 | 1.34604E-45 |
| Intermediate | MTHFD1L | 5.07461E-59 | 0.450968 | 0.33 | 0.14 | 9.60318E-55 |

| | | | | | | |
|---|---|---|---|---|---|---|
| Intermediate | MICAL2 | 4.42395E-47 | 0.448425 | 0.603 | 0.392 | 8.37188E-43 |
| Intermediate | JAG1 | 2.08119E-39 | 0.445145 | 0.666 | 0.474 | 3.93845E-35 |
| Intermediate | ALDH1B1 | 8.10738E-27 | 0.442756 | 0.453 | 0.3 | 1.53424E-22 |
| Intermediate | CSRP1 | 2.32691E-48 | 0.431178 | 0.946 | 0.858 | 4.40345E-44 |
| Intermediate | SDC2 | 2.08304E-41 | 0.430235 | 0.702 | 0.519 | 3.94195E-37 |
| Intermediate | RAB23 | 1.65427E-46 | 0.42307 | 0.386 | 0.198 | 3.13054E-42 |
| Intermediate | PDLIM3 | 2.62256E-41 | 0.421565 | 0.927 | 0.85 | 4.96293E-37 |
| Intermediate | C12orf75 | 3.34898E-53 | 0.420503 | 0.924 | 0.718 | 6.3376E-49 |
| Intermediate | ITGA1 | 1.00613E-39 | 0.419221 | 0.653 | 0.462 | 1.90401E-35 |
| Intermediate | SLC25A4 | 5.81718E-44 | 0.41664 | 0.83 | 0.666 | 1.10084E-39 |
| Intermediate | ARID5B | 2.87681E-31 | 0.410157 | 0.839 | 0.75 | 5.44407E-27 |
| Intermediate | MYH9 | 4.23689E-41 | 0.409599 | 0.888 | 0.776 | 8.0179E-37 |
| Intermediate | ENAH | 1.29608E-21 | 0.40184 | 0.661 | 0.542 | 2.45271E-17 |
| Intermediate | INMT | 3.00814E-36 | 0.401547 | 0.504 | 0.308 | 5.69261E-32 |
| Intermediate | HES4 | 1.76447E-33 | 0.400784 | 0.614 | 0.426 | 3.33909E-29 |
| Intermediate | AHR | 4.66033E-40 | 0.400574 | 0.411 | 0.229 | 8.8192E-36 |
| Intermediate | KIF5B | 5.02314E-34 | 0.399482 | 0.828 | 0.733 | 9.5058E-30 |
| Intermediate | CCDC107 | 5.26214E-39 | 0.397759 | 0.837 | 0.717 | 9.95808E-35 |
| Intermediate | GALNT2 | 3.38346E-39 | 0.395251 | 0.423 | 0.241 | 6.40286E-35 |
| Intermediate | NIBAN1 | 1.72365E-35 | 0.390936 | 0.779 | 0.59 | 3.26184E-31 |
| Intermediate | AC099066. | 4.41204E-56 | 0.389933 | 0.146 | 0.036 | 8.34934E-52 |
| Intermediate | DIPK2A | 4.31968E-35 | 0.389052 | 0.336 | 0.18 | 8.17457E-31 |
| Intermediate | MGARP | 4.02489E-91 | 0.385122 | 0.199 | 0.042 | 7.6167E-87 |
| Intermediate | AOPEP | 7.5854E-34 | 0.38486 | 0.865 | 0.753 | 1.43546E-29 |
| Intermediate | PTK2 | 7.79266E-31 | 0.381602 | 0.589 | 0.417 | 1.47468E-26 |
| Intermediate | ITGA5 | 8.95104E-24 | 0.379512 | 0.185 | 0.09 | 1.69389E-19 |
| Intermediate | LTBP4 | 1.18122E-32 | 0.377228 | 0.804 | 0.632 | 2.23535E-28 |
| Intermediate | CAVIN1 | 1.08431E-38 | 0.373809 | 0.938 | 0.872 | 2.05194E-34 |
| Intermediate | MYL9 | 6.38326E-54 | 0.370047 | 0.997 | 0.979 | 1.20797E-49 |
| Intermediate | CFL2 | 1.72071E-29 | 0.369865 | 0.661 | 0.496 | 3.25627E-25 |
| Intermediate | CITED2 | 1.78176E-26 | 0.366047 | 0.474 | 0.315 | 3.3718E-22 |
| Intermediate | ZYX | 3.37128E-31 | 0.365943 | 0.665 | 0.5 | 6.37981E-27 |
| Intermediate | SACS | 4.11817E-36 | 0.365272 | 0.314 | 0.16 | 7.79323E-32 |
| Intermediate | SCUBE3 | 2.18931E-44 | 0.362697 | 0.304 | 0.137 | 4.14306E-40 |
| Intermediate | THUMPD2 | 2.35665E-34 | 0.361818 | 0.448 | 0.273 | 4.45972E-30 |
| Intermediate | COL18A1 | 3.57252E-30 | 0.360652 | 0.786 | 0.642 | 6.76064E-26 |
| Intermediate | ACTN1 | 2.77398E-30 | 0.358798 | 0.858 | 0.774 | 5.24948E-26 |
| Intermediate | ESAM | 1.17142E-25 | 0.358703 | 0.493 | 0.332 | 2.2168E-21 |
| Intermediate | RGS5 | 9.28621E-43 | 0.358146 | 0.953 | 0.758 | 1.75732E-38 |
| Intermediate | VGLL3 | 5.6441E-55 | 0.357448 | 0.293 | 0.117 | 1.06809E-50 |
| Intermediate | FHL5 | 2.20627E-23 | 0.354413 | 0.359 | 0.221 | 4.17515E-19 |
| Intermediate | ANTXR1 | 1.54462E-21 | 0.353708 | 0.665 | 0.542 | 2.92304E-17 |
| Intermediate | TPM4 | 6.07415E-37 | 0.352315 | 0.966 | 0.908 | 1.14947E-32 |
| Intermediate | PPP1R14B | 2.26608E-25 | 0.348357 | 0.793 | 0.659 | 4.28832E-21 |
| Intermediate | AVEN | 2.80803E-36 | 0.347946 | 0.377 | 0.208 | 5.31392E-32 |
| Intermediate | GPC6 | 6.16115E-12 | 0.347179 | 0.282 | 0.195 | 1.16594E-07 |
| Intermediate | CITED4 | 5.45027E-29 | 0.347141 | 0.34 | 0.196 | 1.03141E-24 |
| Intermediate | SULF1 | 4.52927E-34 | 0.345592 | 0.712 | 0.501 | 8.5712E-30 |
| Intermediate | ART4 | 2.93024E-37 | 0.345295 | 0.248 | 0.111 | 5.54518E-33 |
| Intermediate | SPARC | 1.00514E-38 | 0.341795 | 0.961 | 0.849 | 1.90213E-34 |
| Intermediate | SMS | 7.49961E-26 | 0.341502 | 0.518 | 0.358 | 1.41923E-21 |
| Intermediate | CYB5R3 | 1.02115E-34 | 0.339779 | 0.935 | 0.866 | 1.93242E-30 |
| Intermediate | MAP1B | 2.11758E-30 | 0.33914 | 0.953 | 0.901 | 4.00731E-26 |
| Intermediate | CTPS1 | 8.84001E-42 | 0.333495 | 0.33 | 0.162 | 1.67288E-37 |
| Intermediate | COMT | 6.03022E-27 | 0.332754 | 0.736 | 0.609 | 1.14116E-22 |
| Intermediate | PPDPF | 9.22003E-20 | 0.331776 | 0.899 | 0.871 | 1.7448E-15 |
| Intermediate | MSRB3 | 9.99975E-26 | 0.331039 | 0.799 | 0.672 | 1.89235E-21 |
| Intermediate | NEGR1 | 4.84152E-42 | 0.330826 | 0.208 | 0.08 | 9.16209E-38 |

| | | | | | | |
|---|---|---|---|---|---|---|
| Intermediate | LTBP1 | 1.35379E-29 | 0.330029 | 0.931 | 0.82 | 2.56192E-25 |
| Intermediate | RTN1 | 5.31488E-09 | 0.327835 | 0.297 | 0.223 | 0.000100579 |
| Intermediate | CCT8 | 2.9276E-20 | 0.326663 | 0.669 | 0.564 | 5.54019E-16 |
| Intermediate | ARHGEF9 | 1.88594E-19 | 0.32442 | 0.424 | 0.297 | 3.56895E-15 |
| Intermediate | SUB1 | 1.78244E-21 | 0.324347 | 0.824 | 0.789 | 3.37309E-17 |
| Intermediate | CPXM2 | 9.2765E-26 | 0.323311 | 0.509 | 0.347 | 1.75549E-21 |
| Intermediate | BAMBI | 3.87936E-27 | 0.322729 | 0.219 | 0.108 | 7.3413E-23 |
| Intermediate | SYNPO2 | 7.72182E-26 | 0.322005 | 0.737 | 0.551 | 1.46128E-21 |
| Intermediate | DSTN | 2.36171E-44 | 0.321052 | 0.999 | 0.993 | 4.46929E-40 |
| Intermediate | HSP90AA1 | 8.61541E-15 | 0.320079 | 0.968 | 0.961 | 1.63038E-10 |
| Intermediate | HSPG2 | 1.81466E-17 | 0.318903 | 0.571 | 0.434 | 3.43405E-13 |
| Intermediate | LINC00702 | 1.26513E-23 | 0.318634 | 0.448 | 0.297 | 2.39414E-19 |
| Intermediate | EFHD1 | 3.51036E-27 | 0.317855 | 0.692 | 0.516 | 6.643E-23 |
| Intermediate | SBDS | 6.55189E-34 | 0.317611 | 0.926 | 0.873 | 1.23988E-29 |
| Intermediate | F2R | 1.72942E-28 | 0.315844 | 0.396 | 0.237 | 3.27276E-24 |
| Intermediate | ANXA1 | 1.8365E-32 | 0.315342 | 0.971 | 0.94 | 3.47539E-28 |
| Intermediate | HSPH1 | 7.78727E-16 | 0.314262 | 0.319 | 0.216 | 1.47366E-11 |
| Intermediate | TINAGL1 | 3.90193E-21 | 0.312741 | 0.816 | 0.71 | 7.38401E-17 |
| Intermediate | LASP1 | 3.87267E-25 | 0.310721 | 0.3 | 0.171 | 7.32864E-21 |
| Intermediate | IGFBP7 | 8.96906E-29 | 0.309456 | 1 | 1 | 1.69731E-24 |
| Intermediate | HSPA4 | 4.30547E-18 | 0.309269 | 0.395 | 0.277 | 8.14767E-14 |
| Intermediate | LMCD1 | 8.61493E-28 | 0.308499 | 0.764 | 0.609 | 1.63029E-23 |
| Intermediate | QSOX1 | 1.24316E-24 | 0.308362 | 0.356 | 0.218 | 2.35256E-20 |
| Intermediate | ITGB1 | 8.56035E-34 | 0.306915 | 0.969 | 0.957 | 1.61996E-29 |
| Intermediate | ADGRL3 | 3.36541E-29 | 0.303969 | 0.175 | 0.074 | 6.36869E-25 |
| Intermediate | SLIT3 | 7.68784E-32 | 0.303347 | 0.443 | 0.264 | 1.45485E-27 |
| Intermediate | HACD1 | 7.10233E-26 | 0.302932 | 0.468 | 0.302 | 1.34405E-21 |
| Intermediate | WEE1 | 2.02867E-32 | 0.302002 | 0.288 | 0.147 | 3.83905E-28 |
| Intermediate | RTN4 | 3.40532E-26 | 0.301074 | 0.853 | 0.767 | 6.44422E-22 |
| Intermediate | MAP2 | 2.21115E-34 | 0.300961 | 0.27 | 0.131 | 4.18438E-30 |
| Intermediate | CNN3 | 1.27422E-17 | 0.30033 | 0.778 | 0.689 | 2.41134E-13 |
| Intermediate | HSPA8 | 7.35303E-20 | 0.299785 | 0.88 | 0.843 | 1.39149E-15 |
| Intermediate | MIR100HG | 3.7788E-18 | 0.29867 | 0.646 | 0.52 | 7.151E-14 |
| Intermediate | TC2N | 2.88125E-23 | 0.297764 | 0.269 | 0.15 | 5.45247E-19 |
| Intermediate | NEDD9 | 2.16501E-13 | 0.297547 | 0.309 | 0.213 | 4.09707E-09 |
| Intermediate | HMCN1 | 2.50473E-23 | 0.296935 | 0.576 | 0.414 | 4.73994E-19 |
| Intermediate | CES1 | 1.81011E-14 | 0.294502 | 0.408 | 0.297 | 3.42545E-10 |
| Intermediate | PODN | 1.31321E-38 | 0.293668 | 0.17 | 0.06 | 2.48512E-34 |
| Intermediate | DACT1 | 2.20773E-29 | 0.29333 | 0.233 | 0.113 | 4.1779E-25 |
| Intermediate | SCG2 | 5.01802E-27 | 0.292043 | 0.138 | 0.053 | 9.4961E-23 |
| Intermediate | MATN2 | 3.97287E-28 | 0.29104 | 0.186 | 0.083 | 7.51827E-24 |
| Intermediate | PLPP1 | 9.97468E-18 | 0.290932 | 0.619 | 0.483 | 1.88761E-13 |
| Intermediate | CCND2 | 4.72073E-16 | 0.289643 | 0.546 | 0.42 | 8.9335E-12 |
| Intermediate | PKIG | 3.69332E-19 | 0.289463 | 0.674 | 0.552 | 6.98924E-15 |
| Intermediate | TP53BP2 | 3.31705E-31 | 0.288297 | 0.342 | 0.186 | 6.27719E-27 |
| Intermediate | MYRFL | 4.37332E-38 | 0.288187 | 0.163 | 0.056 | 8.27606E-34 |
| Intermediate | CDH13 | 3.70641E-23 | 0.284956 | 0.67 | 0.491 | 7.01401E-19 |
| Intermediate | AC092807. | 2.32808E-28 | 0.284529 | 0.236 | 0.116 | 4.40566E-24 |
| Intermediate | FST | 9.40578E-22 | 0.282559 | 0.193 | 0.097 | 1.77995E-17 |
| Intermediate | RNF141 | 1.85189E-28 | 0.28228 | 0.345 | 0.196 | 3.50451E-24 |
| Intermediate | SAMD4A | 1.89346E-23 | 0.281167 | 0.305 | 0.179 | 3.58318E-19 |
| Intermediate | TRIM36 | 6.64853E-20 | 0.281113 | 0.221 | 0.121 | 1.25817E-15 |
| Intermediate | TGFB2 | 2.24656E-34 | 0.280215 | 0.192 | 0.078 | 4.25139E-30 |
| Intermediate | CTNNAL1 | 1.2043E-21 | 0.280044 | 0.412 | 0.271 | 2.27903E-17 |
| Intermediate | YAP1 | 5.25455E-17 | 0.279883 | 0.503 | 0.377 | 9.94371E-13 |
| Intermediate | ACTB | 2.59838E-39 | 0.279697 | 1 | 0.998 | 4.91718E-35 |
| Intermediate | C1orf198 | 2.73219E-18 | 0.278603 | 0.441 | 0.31 | 5.1704E-14 |
| Intermediate | OSR1 | 1.86377E-22 | 0.27812 | 0.307 | 0.179 | 3.527E-18 |

| | | | | | | |
|---|---|---|---|---|---|---|
| Intermediate | TSPAN2 | 4.44796E-23 | 0.277971 | 0.513 | 0.347 | 8.41732E-19 |
| Intermediate | AC018742. | 6.59779E-22 | 0.277644 | 0.157 | 0.072 | 1.24856E-17 |
| Intermediate | ANK3 | 6.86347E-25 | 0.277603 | 0.234 | 0.121 | 1.29884E-20 |
| Intermediate | ADAMTS8 | 4.18213E-46 | 0.276448 | 0.149 | 0.043 | 7.91425E-42 |
| Intermediate | PLCG2 | 0.006919222 | 0.275732 | 0.619 | 0.594 | 1 |
| Intermediate | LIMS2 | 3.3426E-21 | 0.275201 | 0.518 | 0.362 | 6.32554E-17 |
| Intermediate | SYTL2 | 1.51865E-28 | 0.274778 | 0.217 | 0.102 | 2.8739E-24 |
| Intermediate | SHROOM3 | 2.83582E-23 | 0.274592 | 0.324 | 0.192 | 5.36651E-19 |
| Intermediate | FXYD1 | 7.95615E-26 | 0.274469 | 0.94 | 0.851 | 1.50562E-21 |
| Intermediate | PRELP | 1.12731E-25 | 0.274441 | 0.86 | 0.733 | 2.13332E-21 |
| Intermediate | DIAPH2 | 9.62946E-18 | 0.272516 | 0.409 | 0.281 | 1.82228E-13 |
| Intermediate | EXT1 | 4.315E-21 | 0.271815 | 0.383 | 0.252 | 8.16571E-17 |
| Intermediate | ISYNA1 | 2.81381E-17 | 0.271399 | 0.45 | 0.324 | 5.32485E-13 |
| Intermediate | MYOZ1 | 1.53321E-23 | 0.271189 | 0.272 | 0.151 | 2.90145E-19 |
| Intermediate | PLS3 | 7.88499E-28 | 0.268719 | 0.958 | 0.875 | 1.49216E-23 |
| Intermediate | HSPA5 | 0.000101989 | 0.267579 | 0.73 | 0.687 | 1 |
| Intermediate | RCN1 | 1.66155E-13 | 0.265605 | 0.57 | 0.456 | 3.14432E-09 |
| Intermediate | ANGPT1 | 4.56985E-07 | 0.264497 | 0.499 | 0.417 | 0.008647983 |
| Intermediate | RBP4 | 2.54595E-39 | 0.263502 | 0.1 | 0.024 | 4.81796E-35 |
| Intermediate | LIMA1 | 6.37451E-12 | 0.263328 | 0.612 | 0.501 | 1.20631E-07 |
| Intermediate | B3GALNT2 | 7.11124E-21 | 0.262791 | 0.313 | 0.19 | 1.34573E-16 |
| Intermediate | ITGA8 | 2.25344E-29 | 0.26225 | 0.919 | 0.747 | 4.26442E-25 |
| Intermediate | CALD1 | 5.33627E-28 | 0.261546 | 0.998 | 0.996 | 1.00984E-23 |
| Intermediate | WDR1 | 4.06526E-19 | 0.261468 | 0.638 | 0.494 | 7.69309E-15 |
| Intermediate | LINC0067C | 4.5742E-36 | 0.258548 | 0.157 | 0.056 | 8.65622E-32 |
| Intermediate | CDK5RAP2 | 4.80666E-15 | 0.258178 | 0.366 | 0.255 | 9.09613E-11 |
| Intermediate | DNAJA1 | 6.58974E-12 | 0.257516 | 0.697 | 0.624 | 1.24704E-07 |
| Intermediate | MAFK | 8.34677E-26 | 0.255938 | 0.232 | 0.118 | 1.57954E-21 |
| Intermediate | ECM2 | 2.95116E-16 | 0.255529 | 0.53 | 0.403 | 5.58477E-12 |
| Intermediate | ODC1 | 5.13684E-15 | 0.255148 | 0.402 | 0.29 | 9.72095E-11 |
| Intermediate | DNAJB6 | 4.33112E-18 | 0.255045 | 0.673 | 0.554 | 8.19621E-14 |
| Intermediate | EVL | 1.55391E-23 | 0.254757 | 0.295 | 0.169 | 2.94063E-19 |
| Intermediate | UGP2 | 4.01362E-15 | 0.253974 | 0.67 | 0.556 | 7.59537E-11 |
| Intermediate | KCNK6 | 6.2868E-23 | 0.25393 | 0.234 | 0.125 | 1.18971E-18 |
| Intermediate | S100A4 | 3.79595E-31 | 0.251576 | 0.997 | 0.982 | 7.18345E-27 |
| Intermediate | PXDN | 1.04474E-22 | 0.251115 | 0.267 | 0.149 | 1.97707E-18 |
| Intermediate | PFN2 | 6.69811E-15 | 0.250524 | 0.627 | 0.511 | 1.26755E-10 |
| Intermediate | LINC01252 | 2.79451E-52 | 0.250442 | 0.129 | 0.03 | 5.28832E-48 |
| DLX5_Cells | DLX6-AS1 | 0 | 2.955431 | 0.738 | 0.015 | 0 |
| DLX5_Cells | SUCNR1 | 0 | 2.927668 | 0.62 | 0.032 | 0 |
| DLX5_Cells | SOST | 0 | 2.746167 | 0.712 | 0.127 | 0 |
| DLX5_Cells | LMO2 | 0 | 2.637359 | 0.827 | 0.113 | 0 |
| DLX5_Cells | DLX5 | 0 | 2.267233 | 0.736 | 0.057 | 0 |
| DLX5_Cells | ATP1B1 | 0 | 2.064726 | 0.755 | 0.202 | 0 |
| DLX5_Cells | TSPAN13 | 0 | 1.326556 | 0.33 | 0.005 | 0 |
| DLX5_Cells | DLX6 | 0 | 1.193458 | 0.417 | 0.021 | 0 |
| DLX5_Cells | RSPO2 | 0 | 1.178262 | 0.268 | 0.008 | 0 |
| DLX5_Cells | SAMD11 | 5.8255E-308 | 1.304371 | 0.446 | 0.05 | 1.1024E-303 |
| DLX5_Cells | FRZB | 5.8465E-302 | 2.093755 | 0.993 | 0.793 | 1.1064E-297 |
| DLX5_Cells | PHYHIPL | 2.2406E-293 | 0.800866 | 0.289 | 0.015 | 4.2401E-289 |
| DLX5_Cells | GREM2 | 1.0635E-291 | 1.240841 | 0.224 | 0.005 | 2.0126E-287 |
| DLX5_Cells | LINC02544 | 3.2321E-274 | 1.108845 | 0.38 | 0.038 | 6.1164E-270 |
| DLX5_Cells | ENPP2 | 9.3852E-258 | 1.666412 | 0.588 | 0.128 | 1.7761E-253 |
| DLX5_Cells | PDE5A | 7.7677E-250 | 1.977944 | 0.942 | 0.634 | 1.47E-245 |
| DLX5_Cells | MSX2 | 2.4331E-216 | 0.996526 | 0.31 | 0.033 | 4.6043E-212 |
| DLX5_Cells | NDNF | 8.4324E-211 | 0.874919 | 0.147 | 0.002 | 1.5958E-206 |
| DLX5_Cells | MECOM | 3.6164E-208 | 1.721799 | 0.497 | 0.109 | 6.8437E-204 |
| DLX5_Cells | CAVIN2 | 4.2211E-204 | 1.455222 | 0.394 | 0.063 | 7.9879E-200 |

| | | | | | | |
|---|---|---|---|---|---|---|
| DLX5_Cells | TMEM233 | 3.818E-203 | 0.462281 | 0.13 | 0.001 | 7.2252E-199 |
| DLX5_Cells | THSD4 | 2.2882E-192 | 1.348104 | 0.527 | 0.134 | 4.3301E-188 |
| DLX5_Cells | BST2 | 3.5195E-183 | 1.315843 | 0.879 | 0.402 | 6.6602E-179 |
| DLX5_Cells | CPNE4 | 9.0416E-177 | 0.549275 | 0.197 | 0.013 | 1.711E-172 |
| DLX5_Cells | PTN | 2.0664E-170 | 2.823152 | 0.626 | 0.234 | 3.9104E-166 |
| DLX5_Cells | DLX3 | 1.6479E-163 | 0.484629 | 0.156 | 0.007 | 3.1185E-159 |
| DLX5_Cells | LMO7 | 1.2712E-158 | 1.501191 | 0.776 | 0.388 | 2.4057E-154 |
| DLX5_Cells | RGS7BP | 2.7494E-158 | 1.033765 | 0.433 | 0.097 | 5.2029E-154 |
| DLX5_Cells | PLK2 | 2.1668E-152 | 1.421839 | 0.598 | 0.209 | 4.1004E-148 |
| DLX5_Cells | ID1 | 4.5462E-152 | 1.469125 | 0.681 | 0.267 | 8.6032E-148 |
| DLX5_Cells | GPC3 | 1.7932E-148 | 1.071271 | 0.302 | 0.049 | 3.3935E-144 |
| DLX5_Cells | CCDC68 | 2.0709E-147 | 1.166578 | 0.317 | 0.054 | 3.919E-143 |
| DLX5_Cells | AC012085. | 8.0552E-145 | 0.433354 | 0.135 | 0.006 | 1.5244E-140 |
| DLX5_Cells | RAMP1 | 7.5995E-144 | 1.188156 | 0.961 | 0.683 | 1.4381E-139 |
| DLX5_Cells | NBL1 | 6.6101E-142 | 1.377975 | 0.863 | 0.572 | 1.2509E-137 |
| DLX5_Cells | ECRG4 | 6.7901E-136 | 0.964811 | 0.852 | 0.457 | 1.285E-131 |
| DLX5_Cells | AK1 | 3.6453E-135 | 1.377435 | 0.822 | 0.51 | 6.8984E-131 |
| DLX5_Cells | HMCN1 | 1.2046E-125 | 1.304453 | 0.753 | 0.405 | 2.2797E-121 |
| DLX5_Cells | FGL2 | 1.3597E-125 | 1.534191 | 0.834 | 0.56 | 2.573E-121 |
| DLX5_Cells | PTGIS | 9.9534E-125 | 1.123749 | 0.788 | 0.447 | 1.8836E-120 |
| DLX5_Cells | LMO3 | 4.5521E-123 | 0.925482 | 0.501 | 0.163 | 8.6145E-119 |
| DLX5_Cells | PRDM6 | 1.7711E-117 | 1.136142 | 0.61 | 0.262 | 3.3517E-113 |
| DLX5_Cells | LSAMP | 1.0367E-116 | 0.335819 | 0.114 | 0.006 | 1.9618E-112 |
| DLX5_Cells | MAB21L2 | 4.6758E-116 | 0.700255 | 0.303 | 0.063 | 8.8485E-112 |
| DLX5_Cells | CDO1 | 6.0285E-114 | 1.159134 | 0.548 | 0.212 | 1.1408E-109 |
| DLX5_Cells | KRT19 | 7.6956E-114 | 0.730808 | 0.151 | 0.013 | 1.4563E-109 |
| DLX5_Cells | ID3 | 8.7541E-113 | 1.085303 | 0.963 | 0.785 | 1.6566E-108 |
| DLX5_Cells | TMEFF2 | 1.8523E-111 | 0.453085 | 0.113 | 0.006 | 3.5053E-107 |
| DLX5_Cells | CLIC3 | 9.4933E-110 | 0.771213 | 0.344 | 0.084 | 1.7965E-105 |
| DLX5_Cells | NPTX2 | 2.5165E-109 | 0.822375 | 0.401 | 0.114 | 4.7622E-105 |
| DLX5_Cells | BEX2 | 1.6925E-106 | 0.863289 | 0.341 | 0.084 | 3.2028E-102 |
| DLX5_Cells | KAZALD1 | 9.5698E-104 | 0.727466 | 0.336 | 0.086 | 1.811E-99 |
| DLX5_Cells | S100A4 | 2.0828E-103 | 0.653001 | 0.992 | 0.984 | 3.9414E-99 |
| DLX5_Cells | OGN | 3.8855E-99 | 0.952919 | 0.97 | 0.857 | 7.35289E-95 |
| DLX5_Cells | BARX1 | 7.6788E-99 | 0.547577 | 0.167 | 0.021 | 1.45313E-94 |
| DLX5_Cells | LGALS3 | 4.04463E-92 | 0.725308 | 0.961 | 0.891 | 7.65406E-88 |
| DLX5_Cells | NCAM1 | 5.02587E-92 | 0.412179 | 0.144 | 0.016 | 9.51095E-88 |
| DLX5_Cells | RAMP2 | 2.48666E-91 | 0.538868 | 0.159 | 0.02 | 4.70576E-87 |
| DLX5_Cells | TAGLN2 | 2.12642E-88 | 0.742177 | 0.948 | 0.823 | 4.02403E-84 |
| DLX5_Cells | ACTC1 | 2.4283E-88 | 1.613037 | 0.516 | 0.207 | 4.59532E-84 |
| DLX5_Cells | IL11RA | 7.00494E-84 | 0.868139 | 0.554 | 0.263 | 1.32562E-79 |
| DLX5_Cells | EFEMP1 | 2.19701E-83 | 0.788716 | 0.946 | 0.762 | 4.15762E-79 |
| DLX5_Cells | GDF10 | 8.64206E-83 | 0.746768 | 0.215 | 0.043 | 1.63542E-78 |
| DLX5_Cells | RGS17 | 6.2511E-81 | 0.816062 | 0.379 | 0.132 | 1.18296E-76 |
| DLX5_Cells | TPD52L1 | 1.98343E-80 | 0.869201 | 0.512 | 0.236 | 3.75344E-76 |
| DLX5_Cells | DAPL1 | 2.1481E-80 | 0.923872 | 0.245 | 0.058 | 4.06507E-76 |
| DLX5_Cells | SOD3 | 3.93357E-80 | 0.692965 | 0.996 | 0.965 | 7.44389E-76 |
| DLX5_Cells | CD34 | 1.28433E-79 | 0.606516 | 0.318 | 0.093 | 2.43047E-75 |
| DLX5_Cells | BEX3 | 5.41119E-78 | 0.719784 | 0.894 | 0.764 | 1.02401E-73 |
| DLX5_Cells | CXXC5 | 1.83166E-76 | 0.809536 | 0.519 | 0.236 | 3.46624E-72 |
| DLX5_Cells | PRRX2 | 2.68959E-76 | 0.818965 | 0.662 | 0.382 | 5.08978E-72 |
| DLX5_Cells | IFITM1 | 2.25256E-75 | 0.597955 | 0.321 | 0.095 | 4.26274E-71 |
| DLX5_Cells | CDKN1C | 2.28982E-75 | 0.926643 | 0.628 | 0.348 | 4.33325E-71 |
| DLX5_Cells | IER2 | 1.22545E-73 | 1.075059 | 0.822 | 0.612 | 2.31905E-69 |
| DLX5_Cells | SLC16A9 | 5.03472E-73 | 0.415969 | 0.159 | 0.026 | 9.5277E-69 |
| DLX5_Cells | DLX2 | 9.13023E-73 | 0.410665 | 0.16 | 0.027 | 1.7278E-68 |
| DLX5_Cells | SYNGR1 | 9.76548E-73 | 0.655027 | 0.427 | 0.17 | 1.84802E-68 |
| DLX5_Cells | GNG2 | 1.24262E-72 | 0.320492 | 0.121 | 0.015 | 2.35154E-68 |

| | | | | | | |
|---|---|---|---|---|---|---|
| DLX5_Cells | BMPR1B | 2.15746E-72 | 0.576071 | 0.255 | 0.067 | 4.08277E-68 |
| DLX5_Cells | INMT | 4.54712E-71 | 0.887889 | 0.584 | 0.311 | 8.60497E-67 |
| DLX5_Cells | S100A13 | 6.11877E-71 | 0.762281 | 0.855 | 0.671 | 1.15792E-66 |
| DLX5_Cells | PAMR1 | 1.06291E-70 | 0.38661 | 0.132 | 0.018 | 2.01145E-66 |
| DLX5_Cells | KCNK17 | 1.46215E-70 | 0.733952 | 0.326 | 0.106 | 2.76698E-66 |
| DLX5_Cells | SEM1 | 1.01527E-69 | 0.61247 | 0.914 | 0.826 | 1.92129E-65 |
| DLX5_Cells | CYTL1 | 1.75416E-67 | 0.825646 | 0.323 | 0.105 | 3.31957E-63 |
| DLX5_Cells | SRP14 | 7.64263E-66 | 0.383513 | 0.993 | 0.985 | 1.44629E-61 |
| DLX5_Cells | PGF | 4.3286E-65 | 0.736284 | 0.531 | 0.257 | 8.19144E-61 |
| DLX5_Cells | LMCD1 | 8.10319E-65 | 0.860695 | 0.821 | 0.612 | 1.53345E-60 |
| DLX5_Cells | GABARAPL | 3.92814E-63 | 0.578094 | 0.943 | 0.849 | 7.43361E-59 |
| DLX5_Cells | VAT1L | 3.78928E-62 | 0.411381 | 0.16 | 0.032 | 7.17083E-58 |
| DLX5_Cells | PDIA5 | 5.92363E-61 | 0.588603 | 0.288 | 0.097 | 1.12099E-56 |
| DLX5_Cells | OMD | 3.88692E-60 | 0.90944 | 0.689 | 0.43 | 7.35562E-56 |
| DLX5_Cells | GEM | 7.73119E-60 | 1.024355 | 0.81 | 0.655 | 1.46305E-55 |
| DLX5_Cells | DMKN | 1.30821E-59 | 0.673277 | 0.505 | 0.251 | 2.47566E-55 |
| DLX5_Cells | MBNL1-AS | 1.8742E-58 | 0.882732 | 0.618 | 0.358 | 3.54673E-54 |
| DLX5_Cells | JUNB | 5.5738E-58 | 0.800254 | 0.875 | 0.687 | 1.05479E-53 |
| DLX5_Cells | HAND2 | 4.4757E-56 | 0.721138 | 0.425 | 0.197 | 8.46981E-52 |
| DLX5_Cells | ASS1 | 1.64585E-55 | 0.632609 | 0.33 | 0.128 | 3.11461E-51 |
| DLX5_Cells | ENG | 3.43479E-55 | 0.833606 | 0.685 | 0.452 | 6.49999E-51 |
| DLX5_Cells | IGFBP5 | 4.23094E-55 | 0.895488 | 0.933 | 0.793 | 8.00663E-51 |
| DLX5_Cells | CXCL12 | 2.13401E-54 | 0.625554 | 0.683 | 0.417 | 4.0384E-50 |
| DLX5_Cells | DYNC1I1 | 4.64114E-54 | 0.517371 | 0.28 | 0.097 | 8.78289E-50 |
| DLX5_Cells | SLC9A3R2 | 4.182E-53 | 0.632052 | 0.598 | 0.364 | 7.91401E-49 |
| DLX5_Cells | DOK5 | 4.23298E-52 | 0.603118 | 0.302 | 0.115 | 8.0105E-48 |
| DLX5_Cells | MT1F | 4.5382E-52 | 0.493569 | 0.227 | 0.068 | 8.58809E-48 |
| DLX5_Cells | TIMP3 | 1.23529E-51 | 0.501957 | 0.736 | 0.469 | 2.33766E-47 |
| DLX5_Cells | OSR1 | 8.25844E-51 | 0.831733 | 0.387 | 0.178 | 1.56283E-46 |
| DLX5_Cells | FOXD2 | 3.83681E-49 | 0.523146 | 0.276 | 0.1 | 7.26079E-45 |
| DLX5_Cells | ARID5B | 9.91555E-49 | 0.772479 | 0.863 | 0.752 | 1.87642E-44 |
| DLX5_Cells | DPT | 1.22265E-48 | 0.931955 | 0.542 | 0.297 | 2.31375E-44 |
| DLX5_Cells | DLX1 | 2.74676E-47 | 0.329839 | 0.136 | 0.029 | 5.19798E-43 |
| DLX5_Cells | TMEM38A | 2.33164E-46 | 0.381102 | 0.167 | 0.044 | 4.41239E-42 |
| DLX5_Cells | CHMP4C | 3.31116E-46 | 0.382903 | 0.215 | 0.067 | 6.26603E-42 |
| DLX5_Cells | LNP1 | 1.5274E-45 | 0.652023 | 0.314 | 0.133 | 2.89045E-41 |
| DLX5_Cells | BGN | 1.81581E-45 | 0.311285 | 1 | 0.98 | 3.43624E-41 |
| DLX5_Cells | KAZN | 4.83196E-45 | 0.435047 | 0.207 | 0.065 | 9.144E-41 |
| DLX5_Cells | CTSH | 1.28372E-44 | 0.430936 | 0.231 | 0.078 | 2.42932E-40 |
| DLX5_Cells | BICC1 | 2.12864E-44 | 0.541506 | 0.391 | 0.186 | 4.02823E-40 |
| DLX5_Cells | MICOS10 | 2.22269E-44 | 0.500047 | 0.887 | 0.805 | 4.20621E-40 |
| DLX5_Cells | NUPR1 | 3.82972E-44 | 0.533527 | 0.904 | 0.826 | 7.24736E-40 |
| DLX5_Cells | S100A11 | 4.72194E-44 | 0.392993 | 0.988 | 0.97 | 8.9358E-40 |
| DLX5_Cells | SKP1 | 4.78925E-44 | 0.388449 | 0.981 | 0.963 | 9.06318E-40 |
| DLX5_Cells | SMPX | 1.61966E-43 | 0.387782 | 0.145 | 0.036 | 3.06505E-39 |
| DLX5_Cells | SPON1 | 3.04674E-43 | 0.419294 | 0.166 | 0.045 | 5.76566E-39 |
| DLX5_Cells | FOS | 6.49621E-43 | 0.985784 | 0.864 | 0.694 | 1.22934E-38 |
| DLX5_Cells | FTH1 | 1.38454E-42 | 0.356089 | 1 | 0.999 | 2.6201E-38 |
| DLX5_Cells | NACA | 1.88264E-42 | 0.370064 | 0.993 | 0.992 | 3.56271E-38 |
| DLX5_Cells | IGFBP2 | 2.28692E-42 | 0.371188 | 0.973 | 0.885 | 4.32777E-38 |
| DLX5_Cells | JUP | 2.32154E-42 | 0.332537 | 0.196 | 0.06 | 4.39328E-38 |
| DLX5_Cells | RGS10 | 3.25899E-42 | 0.505318 | 0.409 | 0.205 | 6.16732E-38 |
| DLX5_Cells | S100A10 | 3.90882E-42 | 0.28844 | 0.973 | 0.84 | 7.39704E-38 |
| DLX5_Cells | CYSTM1 | 5.13576E-42 | 0.567135 | 0.739 | 0.583 | 9.71891E-38 |
| DLX5_Cells | NDUFS5 | 7.69992E-42 | 0.428924 | 0.929 | 0.891 | 1.45713E-37 |
| DLX5_Cells | GMDS | 9.92926E-42 | 0.671668 | 0.723 | 0.539 | 1.87901E-37 |
| DLX5_Cells | NPC2 | 1.18502E-41 | 0.661666 | 0.838 | 0.697 | 2.24254E-37 |
| DLX5_Cells | SUSD3 | 1.49975E-41 | 0.324402 | 0.143 | 0.035 | 2.83812E-37 |

| | | | | | | |
|---|---|---|---|---|---|---|
| DLX5_Cells | CRIP2 | 2.53706E-41 | 0.485398 | 0.927 | 0.847 | 4.80114E-37 |
| DLX5_Cells | HIGD1A | 4.65065E-41 | 0.664417 | 0.606 | 0.424 | 8.80089E-37 |
| DLX5_Cells | SNCG | 5.59019E-41 | 0.83803 | 0.632 | 0.438 | 1.05789E-36 |
| DLX5_Cells | CLU | 2.44728E-40 | 0.678093 | 0.982 | 0.866 | 4.63122E-36 |
| DLX5_Cells | IFITM3 | 2.66745E-40 | 0.436087 | 0.996 | 0.979 | 5.04789E-36 |
| DLX5_Cells | NRP1 | 3.24837E-40 | 0.545322 | 0.447 | 0.24 | 6.14722E-36 |
| DLX5_Cells | IMP3 | 3.53273E-40 | 0.582692 | 0.56 | 0.352 | 6.68533E-36 |
| DLX5_Cells | CCN2 | 1.39725E-39 | 0.830106 | 0.908 | 0.813 | 2.64416E-35 |
| DLX5_Cells | PLAC9 | 2.37917E-39 | 0.377795 | 0.885 | 0.616 | 4.50234E-35 |
| DLX5_Cells | ID4 | 1.44502E-38 | 0.653422 | 0.883 | 0.751 | 2.73456E-34 |
| DLX5_Cells | LAPTM4B | 1.63979E-38 | 0.503124 | 0.436 | 0.236 | 3.10313E-34 |
| DLX5_Cells | AP3S1 | 1.75215E-38 | 0.635323 | 0.681 | 0.51 | 3.31577E-34 |
| DLX5_Cells | SSR3 | 3.96271E-38 | 0.5843 | 0.755 | 0.607 | 7.49904E-34 |
| DLX5_Cells | MYL12B | 4.01407E-38 | 0.417318 | 0.978 | 0.958 | 7.59622E-34 |
| DLX5_Cells | IL1R1 | 1.45701E-37 | 0.612449 | 0.448 | 0.258 | 2.75724E-33 |
| DLX5_Cells | NUDT14 | 5.18431E-37 | 0.370365 | 0.14 | 0.038 | 9.81079E-33 |
| DLX5_Cells | PEBP1 | 6.17786E-37 | 0.35457 | 0.976 | 0.947 | 1.1691E-32 |
| DLX5_Cells | CD81 | 2.16151E-36 | 0.308629 | 0.993 | 0.986 | 4.09045E-32 |
| DLX5_Cells | FZD8 | 2.31363E-36 | 0.50709 | 0.207 | 0.075 | 4.37831E-32 |
| DLX5_Cells | TMEM98 | 3.80308E-36 | 0.499238 | 0.586 | 0.387 | 7.19696E-32 |
| DLX5_Cells | IFITM2 | 9.71426E-36 | 0.429074 | 0.931 | 0.779 | 1.83833E-31 |
| DLX5_Cells | AKR1C2 | 1.89645E-35 | 0.626033 | 0.439 | 0.245 | 3.58883E-31 |
| DLX5_Cells | SLIT3 | 7.13785E-35 | 0.678946 | 0.457 | 0.272 | 1.35077E-30 |
| DLX5_Cells | ATP6V0E1 | 9.61237E-35 | 0.412014 | 0.894 | 0.822 | 1.81904E-30 |
| DLX5_Cells | VEGFB | 1.12635E-34 | 0.510403 | 0.841 | 0.745 | 2.1315E-30 |
| DLX5_Cells | VIM | 1.42437E-34 | 0.388136 | 1 | 1 | 2.69547E-30 |
| DLX5_Cells | NAV1 | 1.92837E-34 | 0.473146 | 0.265 | 0.115 | 3.64924E-30 |
| DLX5_Cells | MT1M | 4.53772E-34 | 0.429475 | 0.582 | 0.365 | 8.58717E-30 |
| DLX5_Cells | PRDX5 | 6.8578E-34 | 0.431243 | 0.844 | 0.756 | 1.29777E-29 |
| DLX5_Cells | NDUFC2 | 1.37185E-33 | 0.396728 | 0.902 | 0.86 | 2.5961E-29 |
| DLX5_Cells | SELENBP1 | 1.38456E-33 | 0.482805 | 0.469 | 0.277 | 2.62014E-29 |
| DLX5_Cells | SOCS2 | 1.54792E-33 | 0.528454 | 0.308 | 0.15 | 2.92928E-29 |
| DLX5_Cells | BTG2 | 2.80028E-33 | 0.773609 | 0.518 | 0.329 | 5.29926E-29 |
| DLX5_Cells | CRYAB | 3.47368E-33 | 0.453396 | 0.952 | 0.908 | 6.57359E-29 |
| DLX5_Cells | GABARAP | 7.3747E-33 | 0.381003 | 0.973 | 0.946 | 1.39559E-28 |
| DLX5_Cells | KCNE4 | 1.46289E-32 | 0.571537 | 0.367 | 0.191 | 2.76838E-28 |
| DLX5_Cells | TUBA4A | 1.62869E-32 | 0.506199 | 0.401 | 0.222 | 3.08212E-28 |
| DLX5_Cells | H3F3B | 2.14479E-32 | 0.345175 | 0.992 | 0.986 | 4.05881E-28 |
| DLX5_Cells | SH3BGRL2 | 4.13095E-32 | 0.277639 | 0.103 | 0.024 | 7.81741E-28 |
| DLX5_Cells | NRGN | 4.22294E-32 | 0.886693 | 0.552 | 0.369 | 7.9915E-28 |
| DLX5_Cells | FLRT2 | 7.21468E-32 | 0.493801 | 0.182 | 0.066 | 1.36531E-27 |
| DLX5_Cells | EEF1D | 8.96061E-32 | 0.315673 | 0.98 | 0.959 | 1.69571E-27 |
| DLX5_Cells | TLE2 | 1.21427E-31 | 0.336771 | 0.254 | 0.108 | 2.29788E-27 |
| DLX5_Cells | CASP4 | 2.36014E-31 | 0.553281 | 0.417 | 0.242 | 4.46632E-27 |
| DLX5_Cells | FNDC5 | 2.54978E-31 | 0.321423 | 0.144 | 0.044 | 4.82521E-27 |
| DLX5_Cells | CSTB | 4.5286E-31 | 0.465019 | 0.879 | 0.803 | 8.56992E-27 |
| DLX5_Cells | IGFBP6 | 5.24451E-31 | 0.938754 | 0.855 | 0.782 | 9.92471E-27 |
| DLX5_Cells | SMAD6 | 7.82113E-31 | 0.319527 | 0.164 | 0.056 | 1.48007E-26 |
| DLX5_Cells | VASN | 8.45499E-31 | 0.560876 | 0.614 | 0.443 | 1.60002E-26 |
| DLX5_Cells | ATOH8 | 9.8672E-31 | 0.463469 | 0.276 | 0.129 | 1.86727E-26 |
| DLX5_Cells | NFKBIA | 2.17605E-30 | 0.757572 | 0.724 | 0.571 | 4.11795E-26 |
| DLX5_Cells | NPB | 2.31156E-30 | 0.264593 | 0.121 | 0.034 | 4.3744E-26 |
| DLX5_Cells | AP001528. | 2.72183E-30 | 0.41434 | 0.302 | 0.147 | 5.15079E-26 |
| DLX5_Cells | PDGFRA | 3.21289E-30 | 0.52522 | 0.482 | 0.304 | 6.08008E-26 |
| DLX5_Cells | BTF3 | 4.05379E-30 | 0.305883 | 0.989 | 0.974 | 7.6714E-26 |
| DLX5_Cells | SEC14L1 | 5.08162E-30 | 0.470491 | 0.283 | 0.135 | 9.61646E-26 |
| DLX5_Cells | MYL6B | 5.10918E-30 | 0.478414 | 0.523 | 0.349 | 9.66861E-26 |
| DLX5_Cells | EEF1A1 | 6.27563E-30 | 0.250373 | 1 | 0.999 | 1.1876E-25 |

| | | | | | | |
|---|---|---|---|---|---|---|
| DLX5_Cells | HSPB2 | 6.9293E-30 | 0.543551 | 0.61 | 0.455 | 1.3113E-25 |
| DLX5_Cells | ACTG1 | 7.90313E-30 | 0.269346 | 0.999 | 0.998 | 1.49559E-25 |
| DLX5_Cells | AC009041. | 1.06163E-29 | 0.359654 | 0.166 | 0.059 | 2.00902E-25 |
| DLX5_Cells | SFRP1 | 1.10753E-29 | 0.618554 | 0.215 | 0.087 | 2.09589E-25 |
| DLX5_Cells | IER5L | 1.63154E-29 | 0.586172 | 0.493 | 0.314 | 3.08753E-25 |
| DLX5_Cells | TLE5 | 2.38057E-29 | 0.439249 | 0.787 | 0.653 | 4.505E-25 |
| DLX5_Cells | AIF1L | 2.53747E-29 | 0.263098 | 0.129 | 0.038 | 4.80191E-25 |
| DLX5_Cells | BEX4 | 3.45598E-29 | 0.518909 | 0.603 | 0.415 | 6.54009E-25 |
| DLX5_Cells | UBB | 4.50067E-29 | 0.329447 | 0.974 | 0.953 | 8.51708E-25 |
| DLX5_Cells | MID1 | 4.55211E-29 | 0.370811 | 0.194 | 0.075 | 8.61441E-25 |
| DLX5_Cells | CD74 | 5.78097E-29 | 0.411053 | 0.444 | 0.261 | 1.09399E-24 |
| DLX5_Cells | ZNF503 | 9.09635E-29 | 0.560065 | 0.508 | 0.332 | 1.72139E-24 |
| DLX5_Cells | NPR3 | 9.82362E-29 | 0.495952 | 0.212 | 0.089 | 1.85902E-24 |
| DLX5_Cells | ID2 | 1.10052E-28 | 0.564109 | 0.484 | 0.311 | 2.08262E-24 |
| DLX5_Cells | VAMP2 | 1.29615E-28 | 0.454529 | 0.81 | 0.705 | 2.45283E-24 |
| DLX5_Cells | PDP1 | 1.43193E-28 | 0.387853 | 0.261 | 0.122 | 2.70978E-24 |
| DLX5_Cells | CDKN2C | 1.45457E-28 | 0.521206 | 0.433 | 0.265 | 2.75263E-24 |
| DLX5_Cells | PFN2 | 2.27457E-28 | 0.50077 | 0.66 | 0.514 | 4.30439E-24 |
| DLX5_Cells | ETFRF1 | 2.44435E-28 | 0.525127 | 0.477 | 0.318 | 4.62569E-24 |
| DLX5_Cells | MT1X | 2.84065E-28 | 0.748561 | 0.715 | 0.571 | 5.37564E-24 |
| DLX5_Cells | CAPG | 2.92578E-28 | 0.380422 | 0.341 | 0.18 | 5.53675E-24 |
| DLX5_Cells | ADIRF | 6.23575E-28 | 0.322753 | 0.999 | 0.993 | 1.18005E-23 |
| DLX5_Cells | LXN | 6.89112E-28 | 0.441147 | 0.197 | 0.08 | 1.30408E-23 |
| DLX5_Cells | NCALD | 7.37477E-28 | 0.472499 | 0.349 | 0.19 | 1.3956E-23 |
| DLX5_Cells | TRABD2B | 1.84667E-27 | 0.331552 | 0.2 | 0.082 | 3.49464E-23 |
| DLX5_Cells | RNASET2 | 4.12794E-27 | 0.709998 | 0.379 | 0.23 | 7.81171E-23 |
| DLX5_Cells | ZFPM2 | 6.12517E-27 | 0.33834 | 0.215 | 0.092 | 1.15913E-22 |
| DLX5_Cells | ECH1 | 6.80622E-27 | 0.477404 | 0.671 | 0.535 | 1.28801E-22 |
| DLX5_Cells | MGST2 | 1.10244E-26 | 0.473851 | 0.452 | 0.292 | 2.08627E-22 |
| DLX5_Cells | LMO7-AS1 | 1.20118E-26 | 0.292791 | 0.133 | 0.043 | 2.2731E-22 |
| DLX5_Cells | H3F3A | 1.34675E-26 | 0.282831 | 0.988 | 0.976 | 2.54858E-22 |
| DLX5_Cells | PTPRF | 1.86826E-26 | 0.309733 | 0.185 | 0.074 | 3.5355E-22 |
| DLX5_Cells | SEPTIN4 | 1.8857E-26 | 0.368258 | 0.512 | 0.331 | 3.56849E-22 |
| DLX5_Cells | DBNDD2 | 2.09725E-26 | 0.434016 | 0.86 | 0.806 | 3.96884E-22 |
| DLX5_Cells | TSPO | 2.19402E-26 | 0.384112 | 0.92 | 0.883 | 4.15196E-22 |
| DLX5_Cells | SLC14A1 | 2.73036E-26 | 0.563995 | 0.255 | 0.122 | 5.16693E-22 |
| DLX5_Cells | SNHG29 | 4.15563E-26 | 0.325934 | 0.965 | 0.945 | 7.86411E-22 |
| DLX5_Cells | FOSB | 5.53546E-26 | 0.490917 | 0.533 | 0.338 | 1.04753E-21 |
| DLX5_Cells | MIR100HG | 2.93766E-25 | 0.501541 | 0.656 | 0.526 | 5.55923E-21 |
| DLX5_Cells | DCLK2 | 4.14106E-25 | 0.354525 | 0.173 | 0.069 | 7.83653E-21 |
| DLX5_Cells | OCIAD2 | 4.7443E-25 | 0.462427 | 0.382 | 0.228 | 8.97811E-21 |
| DLX5_Cells | MAPK10 | 6.94531E-25 | 0.315806 | 0.217 | 0.098 | 1.31433E-20 |
| DLX5_Cells | HRH1 | 7.96393E-25 | 0.405845 | 0.235 | 0.112 | 1.50709E-20 |
| DLX5_Cells | EMC2 | 9.06551E-25 | 0.458682 | 0.549 | 0.4 | 1.71556E-20 |
| DLX5_Cells | SEC11A | 1.56003E-24 | 0.400776 | 0.859 | 0.806 | 2.9522E-20 |
| DLX5_Cells | ANKMY2 | 3.3819E-24 | 0.346241 | 0.204 | 0.09 | 6.3999E-20 |
| DLX5_Cells | DNPH1 | 6.62185E-24 | 0.403578 | 0.417 | 0.262 | 1.25312E-19 |
| DLX5_Cells | COPRS | 7.12158E-24 | 0.359953 | 0.799 | 0.686 | 1.34769E-19 |
| DLX5_Cells | PLCD3 | 7.42746E-24 | 0.335769 | 0.249 | 0.12 | 1.40557E-19 |
| DLX5_Cells | RFTN1 | 1.0703E-23 | 0.439729 | 0.341 | 0.199 | 2.02544E-19 |
| DLX5_Cells | TMEM14A | 1.80592E-23 | 0.471362 | 0.461 | 0.314 | 3.41752E-19 |
| DLX5_Cells | CLTA | 2.08505E-23 | 0.354772 | 0.855 | 0.79 | 3.94574E-19 |
| DLX5_Cells | ADAMTSL3 | 2.64952E-23 | 0.380897 | 0.285 | 0.152 | 5.01394E-19 |
| DLX5_Cells | G0S2 | 4.84604E-23 | 0.5898 | 0.2 | 0.087 | 9.17064E-19 |
| DLX5_Cells | PRTFDC1 | 7.01056E-23 | 0.363187 | 0.211 | 0.098 | 1.32668E-18 |
| DLX5_Cells | FAM118A | 7.99327E-23 | 0.439758 | 0.303 | 0.168 | 1.51265E-18 |
| DLX5_Cells | YWHAE | 1.06023E-22 | 0.40378 | 0.749 | 0.653 | 2.00638E-18 |
| DLX5_Cells | HOXA3 | 1.14483E-22 | 0.333982 | 0.196 | 0.087 | 2.16648E-18 |

| | | | | | | |
|---|---|---|---|---|---|---|
| DLX5_Cells | CYP4X1 | 3.04959E-22 | 0.345704 | 0.242 | 0.12 | 5.77105E-18 |
| DLX5_Cells | KLF2 | 4.21041E-22 | 1.122466 | 0.592 | 0.477 | 7.96778E-18 |
| DLX5_Cells | NAT14 | 4.65158E-22 | 0.324483 | 0.19 | 0.086 | 8.80264E-18 |
| DLX5_Cells | KRT10 | 6.22773E-22 | 0.353274 | 0.8 | 0.695 | 1.17854E-17 |
| DLX5_Cells | GATA2 | 6.22809E-22 | 0.252086 | 0.12 | 0.041 | 1.1786E-17 |
| DLX5_Cells | SNAI2 | 6.58672E-22 | 0.31244 | 0.155 | 0.062 | 1.24647E-17 |
| DLX5_Cells | CYBA | 8.7606E-22 | 0.380595 | 0.632 | 0.464 | 1.65786E-17 |
| DLX5_Cells | LINC01503 | 1.26748E-21 | 0.273875 | 0.129 | 0.047 | 2.39858E-17 |
| DLX5_Cells | CLMP | 2.21904E-21 | 0.324578 | 0.266 | 0.139 | 4.19931E-17 |
| DLX5_Cells | SCUBE3 | 2.42521E-21 | 0.343747 | 0.279 | 0.149 | 4.58947E-17 |
| DLX5_Cells | PRDX1 | 2.8946E-21 | 0.266447 | 0.963 | 0.941 | 5.47775E-17 |
| DLX5_Cells | MTHFS | 3.564E-21 | 0.361153 | 0.277 | 0.152 | 6.74452E-17 |
| DLX5_Cells | GNAS | 3.86915E-21 | 0.256551 | 0.985 | 0.98 | 7.32198E-17 |
| DLX5_Cells | MYH10 | 5.96596E-21 | 0.321339 | 0.918 | 0.769 | 1.129E-16 |
| DLX5_Cells | RGS3 | 6.55522E-21 | 0.45061 | 0.522 | 0.383 | 1.24051E-16 |
| DLX5_Cells | NDUFAF3 | 7.05752E-21 | 0.37776 | 0.735 | 0.631 | 1.33557E-16 |
| DLX5_Cells | VDAC2 | 8.85752E-21 | 0.36099 | 0.822 | 0.748 | 1.6762E-16 |
| DLX5_Cells | C1QTNF1 | 9.67886E-21 | 0.426381 | 0.601 | 0.454 | 1.83163E-16 |
| DLX5_Cells | CCDC151 | 1.52111E-20 | 0.388323 | 0.242 | 0.125 | 2.87855E-16 |
| DLX5_Cells | HSP90AB1 | 1.66572E-20 | 0.298583 | 0.993 | 0.979 | 3.15221E-16 |
| DLX5_Cells | CARD16 | 1.8313E-20 | 0.329114 | 0.189 | 0.087 | 3.46555E-16 |
| DLX5_Cells | PIR | 2.39082E-20 | 0.360309 | 0.238 | 0.122 | 4.52438E-16 |
| DLX5_Cells | GNG5 | 2.45502E-20 | 0.340235 | 0.895 | 0.834 | 4.64587E-16 |
| DLX5_Cells | COX4I1 | 2.94898E-20 | 0.288566 | 0.95 | 0.93 | 5.58066E-16 |
| DLX5_Cells | CITED2 | 4.08619E-20 | 0.523368 | 0.466 | 0.324 | 7.7327E-16 |
| DLX5_Cells | ZNF428 | 8.3088E-20 | 0.433893 | 0.643 | 0.545 | 1.57236E-15 |
| DLX5_Cells | H1F0 | 9.07022E-20 | 0.460742 | 0.345 | 0.213 | 1.71645E-15 |
| DLX5_Cells | HLA-DPB1 | 1.31581E-19 | 0.34642 | 0.383 | 0.242 | 2.49004E-15 |
| DLX5_Cells | ARL4D | 1.37638E-19 | 0.29115 | 0.204 | 0.099 | 2.60467E-15 |
| DLX5_Cells | MT1E | 2.38665E-19 | 0.286179 | 0.664 | 0.503 | 4.51649E-15 |
| DLX5_Cells | RAB29 | 3.38037E-19 | 0.321149 | 0.235 | 0.122 | 6.39702E-15 |
| DLX5_Cells | VSTM4 | 3.75429E-19 | 0.348537 | 0.243 | 0.129 | 7.10462E-15 |
| DLX5_Cells | F10 | 4.23318E-19 | 0.33108 | 0.285 | 0.16 | 8.01086E-15 |
| DLX5_Cells | PNKD | 4.52542E-19 | 0.356553 | 0.663 | 0.538 | 8.5639E-15 |
| DLX5_Cells | COMT | 4.69429E-19 | 0.362459 | 0.712 | 0.618 | 8.88347E-15 |
| DLX5_Cells | ATP13A3 | 4.78653E-19 | 0.43728 | 0.327 | 0.203 | 9.05804E-15 |
| DLX5_Cells | STARD10 | 6.82082E-19 | 0.348955 | 0.307 | 0.182 | 1.29077E-14 |
| DLX5_Cells | ATP5IF1 | 1.03203E-18 | 0.340694 | 0.84 | 0.79 | 1.95301E-14 |
| DLX5_Cells | CKB | 1.33998E-18 | 0.376941 | 0.326 | 0.194 | 2.53578E-14 |
| DLX5_Cells | RNASEK | 1.59479E-18 | 0.260083 | 0.946 | 0.914 | 3.01797E-14 |
| DLX5_Cells | PLAAT3 | 1.69864E-18 | 0.427159 | 0.575 | 0.463 | 3.21451E-14 |
| DLX5_Cells | ELN | 2.45064E-18 | 0.581295 | 0.549 | 0.425 | 4.6376E-14 |
| DLX5_Cells | LINC00623 | 3.8297E-18 | 0.406557 | 0.317 | 0.194 | 7.24733E-14 |
| DLX5_Cells | MDFIC | 4.02711E-18 | 0.378623 | 0.284 | 0.166 | 7.62089E-14 |
| DLX5_Cells | PRDX2 | 4.54678E-18 | 0.293363 | 0.837 | 0.787 | 8.60432E-14 |
| DLX5_Cells | GCHFR | 4.79534E-18 | 0.286049 | 0.168 | 0.077 | 9.0747E-14 |
| DLX5_Cells | SPINT2 | 7.54371E-18 | 0.329453 | 0.814 | 0.654 | 1.42757E-13 |
| DLX5_Cells | HMGN1 | 7.83005E-18 | 0.368741 | 0.647 | 0.544 | 1.48176E-13 |
| DLX5_Cells | MTCH1 | 8.26385E-18 | 0.391308 | 0.738 | 0.674 | 1.56385E-13 |
| DLX5_Cells | NDUFA11 | 8.9392E-18 | 0.271753 | 0.872 | 0.821 | 1.69166E-13 |
| DLX5_Cells | PLD5 | 1.24804E-17 | 0.30545 | 0.155 | 0.07 | 2.36179E-13 |
| DLX5_Cells | ARL5A | 1.25273E-17 | 0.428287 | 0.656 | 0.568 | 2.37067E-13 |
| DLX5_Cells | CCSER2 | 1.32941E-17 | 0.54222 | 0.601 | 0.524 | 2.51578E-13 |
| DLX5_Cells | BZW2 | 1.40603E-17 | 0.366349 | 0.213 | 0.114 | 2.66076E-13 |
| DLX5_Cells | RHOB | 1.53363E-17 | 0.399732 | 0.731 | 0.647 | 2.90224E-13 |
| DLX5_Cells | ITM2C | 1.71608E-17 | 0.447593 | 0.538 | 0.412 | 3.2475E-13 |
| DLX5_Cells | CLIC1 | 1.83737E-17 | 0.26426 | 0.933 | 0.9 | 3.47704E-13 |
| DLX5_Cells | STMN1 | 2.63649E-17 | 0.424845 | 0.379 | 0.252 | 4.9893E-13 |

| | | | | | | |
|---|---|---|---|---|---|---|
| DLX5_Cells | PDLIM4 | 3.29089E-17 | 0.369408 | 0.511 | 0.382 | 6.22769E-13 |
| DLX5_Cells | ROM1 | 4.79879E-17 | 0.269297 | 0.213 | 0.111 | 9.08123E-13 |
| DLX5_Cells | GOLIM4 | 5.48793E-17 | 0.478373 | 0.622 | 0.525 | 1.03854E-12 |
| DLX5_Cells | KCTD12 | 6.77624E-17 | 0.633018 | 0.272 | 0.166 | 1.28234E-12 |
| DLX5_Cells | FZD1 | 6.84787E-17 | 0.480626 | 0.359 | 0.244 | 1.29589E-12 |
| DLX5_Cells | PREX2 | 7.08376E-17 | 0.333255 | 0.178 | 0.086 | 1.34053E-12 |
| DLX5_Cells | CHCHD10 | 1.35105E-16 | 0.304511 | 0.746 | 0.638 | 2.55672E-12 |
| DLX5_Cells | GADD45B | 1.68436E-16 | 0.435282 | 0.67 | 0.548 | 3.18748E-12 |
| DLX5_Cells | H1FX | 2.39674E-16 | 0.37381 | 0.654 | 0.524 | 4.53559E-12 |
| DLX5_Cells | VAMP5 | 6.42576E-16 | 0.31339 | 0.667 | 0.554 | 1.21601E-11 |
| DLX5_Cells | EPB41L2 | 7.00505E-16 | 0.409005 | 0.601 | 0.481 | 1.32564E-11 |
| DLX5_Cells | ATP5MC2 | 8.18171E-16 | 0.306845 | 0.933 | 0.908 | 1.54831E-11 |
| DLX5_Cells | EEF1B2 | 1.29699E-15 | 0.259215 | 0.963 | 0.944 | 2.45443E-11 |
| DLX5_Cells | TBCA | 1.31728E-15 | 0.286579 | 0.859 | 0.795 | 2.49281E-11 |
| DLX5_Cells | NFIA | 1.43054E-15 | 0.325978 | 0.817 | 0.738 | 2.70715E-11 |
| DLX5_Cells | HDDC2 | 1.60476E-15 | 0.332795 | 0.558 | 0.438 | 3.03684E-11 |
| DLX5_Cells | EIF3E | 1.76456E-15 | 0.341053 | 0.838 | 0.815 | 3.33925E-11 |
| DLX5_Cells | NDUFB8 | 3.01003E-15 | 0.274878 | 0.87 | 0.839 | 5.69617E-11 |
| DLX5_Cells | XIST | 3.34032E-15 | 0.300824 | 0.136 | 0.06 | 6.32122E-11 |
| DLX5_Cells | HSPA2 | 3.5872E-15 | 0.359196 | 0.376 | 0.255 | 6.78842E-11 |
| DLX5_Cells | SEMA5A | 3.94132E-15 | 0.379022 | 0.279 | 0.17 | 7.45856E-11 |
| DLX5_Cells | SH3BGRL3 | 4.51121E-15 | 0.35762 | 0.84 | 0.81 | 8.53701E-11 |
| DLX5_Cells | TIPARP | 5.24879E-15 | 0.253342 | 0.168 | 0.084 | 9.93281E-11 |
| DLX5_Cells | LGALS3BP | 7.09583E-15 | 0.285641 | 0.658 | 0.515 | 1.34282E-10 |
| DLX5_Cells | ZFYVE21 | 1.02079E-14 | 0.351011 | 0.645 | 0.535 | 1.93174E-10 |
| DLX5_Cells | KRAS | 1.2566E-14 | 0.364171 | 0.391 | 0.281 | 2.37799E-10 |
| DLX5_Cells | DNAJC15 | 1.37468E-14 | 0.341192 | 0.588 | 0.469 | 2.60145E-10 |
| DLX5_Cells | PSMB9 | 1.43986E-14 | 0.295647 | 0.367 | 0.245 | 2.72479E-10 |
| DLX5_Cells | LINC01578 | 3.84537E-14 | 0.330653 | 0.754 | 0.678 | 7.27698E-10 |
| DLX5_Cells | TMBIM1 | 4.23793E-14 | 0.344274 | 0.556 | 0.447 | 8.01985E-10 |
| DLX5_Cells | BEND5 | 5.15348E-14 | 0.253289 | 0.215 | 0.12 | 9.75245E-10 |
| DLX5_Cells | MSRB2 | 5.30905E-14 | 0.34672 | 0.622 | 0.537 | 1.00469E-09 |
| DLX5_Cells | HSPA1B | 5.6211E-14 | 0.317244 | 0.56 | 0.429 | 1.06374E-09 |
| DLX5_Cells | TSPAN4 | 7.65913E-14 | 0.29392 | 0.488 | 0.365 | 1.44941E-09 |
| DLX5_Cells | LACTB2 | 8.42715E-14 | 0.304836 | 0.197 | 0.11 | 1.59475E-09 |
| DLX5_Cells | AAMDC | 8.53537E-14 | 0.320267 | 0.592 | 0.495 | 1.61523E-09 |
| DLX5_Cells | PXMP2 | 9.81551E-14 | 0.308843 | 0.319 | 0.21 | 1.85749E-09 |
| DLX5_Cells | THYN1 | 1.15286E-13 | 0.33111 | 0.47 | 0.36 | 2.18167E-09 |
| DLX5_Cells | IGFBP3 | 1.44508E-13 | 1.515106 | 0.39 | 0.293 | 2.73467E-09 |
| DLX5_Cells | RAB34 | 1.62612E-13 | 0.283293 | 0.76 | 0.701 | 3.07727E-09 |
| DLX5_Cells | SYNPO | 3.13411E-13 | 0.446138 | 0.567 | 0.474 | 5.931E-09 |
| DLX5_Cells | TMEM237 | 5.19396E-13 | 0.309287 | 0.321 | 0.218 | 9.82905E-09 |
| DLX5_Cells | HEBP1 | 5.8437E-13 | 0.299042 | 0.622 | 0.54 | 1.10586E-08 |
| DLX5_Cells | ITGA10 | 6.38284E-13 | 0.306696 | 0.457 | 0.345 | 1.20789E-08 |
| DLX5_Cells | EFHD1 | 7.61192E-13 | 0.304362 | 0.649 | 0.529 | 1.44048E-08 |
| DLX5_Cells | PEX2 | 9.43496E-13 | 0.305032 | 0.378 | 0.273 | 1.78547E-08 |
| DLX5_Cells | HSPB11 | 1.03759E-12 | 0.343892 | 0.427 | 0.327 | 1.96354E-08 |
| DLX5_Cells | ANKRD35 | 1.3254E-12 | 0.309065 | 0.33 | 0.228 | 2.50818E-08 |
| DLX5_Cells | ACKR4 | 1.45265E-12 | 0.371185 | 0.215 | 0.127 | 2.74899E-08 |
| DLX5_Cells | C4orf3 | 3.43426E-12 | 0.273644 | 0.908 | 0.847 | 6.499E-08 |
| DLX5_Cells | PSIP1 | 4.13966E-12 | 0.304 | 0.74 | 0.668 | 7.83389E-08 |
| DLX5_Cells | AKR7A2 | 4.35311E-12 | 0.326942 | 0.531 | 0.44 | 8.23783E-08 |
| DLX5_Cells | BLOC1S1 | 6.11653E-12 | 0.256351 | 0.761 | 0.7 | 1.15749E-07 |
| DLX5_Cells | TCEAL3 | 8.78676E-12 | 0.25847 | 0.736 | 0.65 | 1.66281E-07 |
| DLX5_Cells | GLRX | 9.44096E-12 | 0.383522 | 0.311 | 0.212 | 1.78661E-07 |
| DLX5_Cells | CUX1 | 1.00257E-11 | 0.344915 | 0.47 | 0.371 | 1.89727E-07 |
| DLX5_Cells | ACAA2 | 1.07858E-11 | 0.284245 | 0.28 | 0.185 | 2.0411E-07 |
| DLX5_Cells | VCAM1 | 1.13126E-11 | 0.265157 | 0.387 | 0.263 | 2.1408E-07 |

| | | | | | | |
|---|---|---|---|---|---|---|
| DLX5_Cells | CCS | 1.20001E-11 | 0.289642 | 0.429 | 0.323 | 2.2709E-07 |
| DLX5_Cells | SNX2 | 1.36411E-11 | 0.302432 | 0.52 | 0.428 | 2.58145E-07 |
| DLX5_Cells | SEC62 | 1.38806E-11 | 0.270093 | 0.914 | 0.878 | 2.62676E-07 |
| DLX5_Cells | DPY30 | 1.39599E-11 | 0.3248 | 0.501 | 0.417 | 2.64176E-07 |
| DLX5_Cells | RRAD | 1.41485E-11 | 0.327926 | 0.446 | 0.337 | 2.67746E-07 |
| DLX5_Cells | NEO1 | 1.41984E-11 | 0.336087 | 0.283 | 0.191 | 2.68691E-07 |
| DLX5_Cells | AKR1C3 | 1.72684E-11 | 0.396895 | 0.497 | 0.393 | 3.26788E-07 |
| DLX5_Cells | SERTAD3 | 3.05423E-11 | 0.287669 | 0.302 | 0.204 | 5.77982E-07 |
| DLX5_Cells | ETFB | 3.2293E-11 | 0.287155 | 0.602 | 0.523 | 6.11113E-07 |
| DLX5_Cells | MIR497HG | 3.87663E-11 | 0.289074 | 0.357 | 0.259 | 7.33613E-07 |
| DLX5_Cells | TOB1 | 4.25324E-11 | 0.315013 | 0.393 | 0.291 | 8.04883E-07 |
| DLX5_Cells | SSBP4 | 4.3466E-11 | 0.300053 | 0.418 | 0.32 | 8.22551E-07 |
| DLX5_Cells | KLF13 | 5.44098E-11 | 0.313089 | 0.405 | 0.306 | 1.02965E-06 |
| DLX5_Cells | FZD4 | 6.98788E-11 | 0.269276 | 0.201 | 0.122 | 1.32239E-06 |
| DLX5_Cells | ARHGAP21 | 7.23728E-11 | 0.30902 | 0.395 | 0.299 | 1.36958E-06 |
| DLX5_Cells | PSMA2 | 7.86936E-11 | 0.295281 | 0.765 | 0.71 | 1.4892E-06 |
| DLX5_Cells | BLVRB | 8.93817E-11 | 0.29884 | 0.658 | 0.575 | 1.69146E-06 |
| DLX5_Cells | OCIAD1 | 1.0115E-10 | 0.273113 | 0.7 | 0.632 | 1.91417E-06 |
| DLX5_Cells | ISYNA1 | 1.0588E-10 | 0.27719 | 0.439 | 0.332 | 2.00368E-06 |
| DLX5_Cells | CES1 | 1.06252E-10 | 0.290507 | 0.413 | 0.302 | 2.01071E-06 |
| DLX5_Cells | LRP1 | 1.14024E-10 | 0.411659 | 0.666 | 0.613 | 2.15779E-06 |
| DLX5_Cells | HES1 | 1.32946E-10 | 0.346486 | 0.423 | 0.327 | 2.51586E-06 |
| DLX5_Cells | RPA2 | 1.36897E-10 | 0.280642 | 0.329 | 0.234 | 2.59063E-06 |
| DLX5_Cells | GATA6 | 2.18764E-10 | 0.251551 | 0.291 | 0.197 | 4.13989E-06 |
| DLX5_Cells | EML4 | 2.60143E-10 | 0.276722 | 0.253 | 0.167 | 4.92295E-06 |
| DLX5_Cells | C1GALT1 | 2.88947E-10 | 0.339184 | 0.234 | 0.154 | 5.46804E-06 |
| DLX5_Cells | PKIG | 3.64418E-10 | 0.262473 | 0.636 | 0.562 | 6.89624E-06 |
| DLX5_Cells | CITED4 | 3.65098E-10 | 0.275633 | 0.298 | 0.208 | 6.90912E-06 |
| DLX5_Cells | MRPS18C | 3.88486E-10 | 0.293556 | 0.598 | 0.535 | 7.3517E-06 |
| DLX5_Cells | PLEKHA4 | 4.23036E-10 | 0.305064 | 0.312 | 0.222 | 8.00553E-06 |
| DLX5_Cells | RARRES1 | 4.27022E-10 | 0.48629 | 0.137 | 0.074 | 8.08096E-06 |
| DLX5_Cells | PLIN3 | 4.52879E-10 | 0.274932 | 0.48 | 0.394 | 8.57029E-06 |
| DLX5_Cells | CTSF | 5.7669E-10 | 0.255938 | 0.615 | 0.516 | 1.09133E-05 |
| DLX5_Cells | PRMT1 | 6.13109E-10 | 0.250812 | 0.602 | 0.525 | 1.16025E-05 |
| DLX5_Cells | SKIL | 6.91958E-10 | 0.334616 | 0.423 | 0.334 | 1.30946E-05 |
| DLX5_Cells | ST5 | 7.32194E-10 | 0.267706 | 0.207 | 0.128 | 1.3856E-05 |
| DLX5_Cells | TCEAL2 | 8.12755E-10 | 0.44709 | 0.427 | 0.344 | 1.53806E-05 |
| DLX5_Cells | MEIS2 | 8.19093E-10 | 0.348943 | 0.28 | 0.194 | 1.55005E-05 |
| DLX5_Cells | SMAD7 | 8.40236E-10 | 0.272863 | 0.303 | 0.211 | 1.59006E-05 |
| DLX5_Cells | IRF1 | 8.78144E-10 | 0.319389 | 0.255 | 0.168 | 1.6618E-05 |
| DLX5_Cells | MSX1 | 8.98702E-10 | 0.271317 | 0.216 | 0.137 | 1.7007E-05 |
| DLX5_Cells | WWTR1 | 9.47991E-10 | 0.281549 | 0.534 | 0.451 | 1.79398E-05 |
| DLX5_Cells | CNRIP1 | 9.60962E-10 | 0.266979 | 0.253 | 0.17 | 1.81852E-05 |
| DLX5_Cells | KLF10 | 9.71714E-10 | 0.259101 | 0.274 | 0.188 | 1.83887E-05 |
| DLX5_Cells | PCBP1 | 1.00179E-09 | 0.291471 | 0.732 | 0.707 | 1.89578E-05 |
| DLX5_Cells | DEXI | 1.23213E-09 | 0.263204 | 0.435 | 0.349 | 2.33167E-05 |
| DLX5_Cells | IGLC1 | 1.27316E-09 | 0.261703 | 0.121 | 0.062 | 2.40933E-05 |
| DLX5_Cells | SCRG1 | 1.32301E-09 | 0.598923 | 0.497 | 0.411 | 2.50367E-05 |
| DLX5_Cells | PDGFD | 1.35138E-09 | 0.356117 | 0.38 | 0.292 | 2.55735E-05 |
| DLX5_Cells | ZFP36 | 1.71463E-09 | 0.413709 | 0.674 | 0.586 | 3.24476E-05 |
| DLX5_Cells | BHLHE41 | 2.02319E-09 | 0.295716 | 0.224 | 0.147 | 3.82868E-05 |
| DLX5_Cells | GJA1 | 2.14451E-09 | 0.396143 | 0.481 | 0.4 | 4.05826E-05 |
| DLX5_Cells | TCEA3 | 2.26669E-09 | 0.278147 | 0.412 | 0.322 | 4.28949E-05 |
| DLX5_Cells | HYI | 2.35634E-09 | 0.268509 | 0.561 | 0.483 | 4.45914E-05 |
| DLX5_Cells | FKBP8 | 3.06866E-09 | 0.250892 | 0.754 | 0.702 | 5.80713E-05 |
| DLX5_Cells | DUSP1 | 3.10537E-09 | 0.263883 | 0.908 | 0.862 | 5.8766E-05 |
| DLX5_Cells | FAM92A | 3.39062E-09 | 0.268536 | 0.291 | 0.205 | 6.41641E-05 |
| DLX5_Cells | ACADVL | 3.43309E-09 | 0.317083 | 0.477 | 0.4 | 6.49678E-05 |

| | | | | | | |
|---|---|---|---|---|---|---|
| DLX5_Cells | PLAAT4 | 4.39406E-09 | 0.290702 | 0.376 | 0.283 | 8.31532E-05 |
| DLX5_Cells | PCBP2 | 4.43753E-09 | 0.25177 | 0.746 | 0.702 | 8.39758E-05 |
| DLX5_Cells | MYCBP2 | 4.91794E-09 | 0.394912 | 0.477 | 0.404 | 9.30671E-05 |
| DLX5_Cells | CAT | 4.97513E-09 | 0.268029 | 0.364 | 0.276 | 9.41494E-05 |
| DLX5_Cells | HNRNPH1 | 5.30731E-09 | 0.280299 | 0.632 | 0.558 | 0.000100436 |
| DLX5_Cells | SMAD9 | 5.8778E-09 | 0.267276 | 0.285 | 0.203 | 0.000111232 |
| DLX5_Cells | LARP6 | 7.9086E-09 | 0.275839 | 0.428 | 0.344 | 0.000149662 |
| DLX5_Cells | CARHSP1 | 9.17204E-09 | 0.260658 | 0.457 | 0.369 | 0.000173572 |
| DLX5_Cells | SERPINB6 | 9.17565E-09 | 0.269857 | 0.613 | 0.549 | 0.00017364 |
| DLX5_Cells | LINC02762 | 1.19996E-08 | 0.281262 | 0.408 | 0.321 | 0.00022708 |
| DLX5_Cells | OXR1 | 1.40406E-08 | 0.26272 | 0.439 | 0.356 | 0.000265704 |
| DLX5_Cells | GSTM3 | 1.60263E-08 | 0.264894 | 0.469 | 0.389 | 0.000303282 |
| DLX5_Cells | NORAD | 1.64121E-08 | 0.289653 | 0.726 | 0.679 | 0.000310582 |
| DLX5_Cells | FOXC2 | 2.01303E-08 | 0.269391 | 0.458 | 0.375 | 0.000380947 |
| DLX5_Cells | UROD | 2.91139E-08 | 0.274208 | 0.361 | 0.283 | 0.000550951 |
| DLX5_Cells | LMO4 | 4.17743E-08 | 0.303408 | 0.482 | 0.41 | 0.000790538 |
| DLX5_Cells | BDH2 | 4.31945E-08 | 0.253091 | 0.515 | 0.447 | 0.000817412 |
| DLX5_Cells | IGSF8 | 4.37097E-08 | 0.265976 | 0.296 | 0.222 | 0.000827163 |
| DLX5_Cells | PBX1 | 5.04509E-08 | 0.327746 | 0.333 | 0.262 | 0.000954732 |
| DLX5_Cells | TECR | 8.35412E-08 | 0.274174 | 0.48 | 0.404 | 0.001580934 |
| DLX5_Cells | FAM200B | 8.94294E-08 | 0.2824 | 0.433 | 0.359 | 0.001692361 |
| DLX5_Cells | HIP1 | 9.43794E-08 | 0.250781 | 0.3 | 0.225 | 0.001786036 |
| DLX5_Cells | FAM162A | 1.22224E-07 | 0.269184 | 0.615 | 0.566 | 0.002312963 |
| DLX5_Cells | SNHG18 | 1.36953E-07 | 0.250635 | 0.374 | 0.295 | 0.002591708 |
| DLX5_Cells | LITAF | 1.61703E-07 | 0.280275 | 0.596 | 0.525 | 0.003060063 |
| DLX5_Cells | KLF4 | 2.48646E-07 | 0.330326 | 0.216 | 0.15 | 0.004705376 |
| DLX5_Cells | PAXX | 3.92119E-07 | 0.252995 | 0.405 | 0.342 | 0.007420452 |
| DLX5_Cells | ZSWIM7 | 4.60275E-07 | 0.290093 | 0.299 | 0.232 | 0.008710246 |
| DLX5_Cells | PPP1R15A | 9.30065E-07 | 0.303907 | 0.576 | 0.513 | 0.017600558 |
| DLX5_Cells | CKLF | 1.70375E-06 | 0.25178 | 0.34 | 0.273 | 0.032241676 |
| DLX5_Cells | ING2 | 3.23227E-06 | 0.259052 | 0.209 | 0.15 | 0.061167403 |
| DLX5_Cells | GNB4 | 3.31845E-06 | 0.260772 | 0.323 | 0.262 | 0.062798262 |
| DLX5_Cells | GLDN | 5.01557E-06 | 0.258261 | 0.279 | 0.217 | 0.094914611 |
| DLX5_Cells | EXT1 | 5.8973E-06 | 0.273778 | 0.326 | 0.265 | 0.111600441 |
| DLX5_Cells | ZFAND5 | 6.2662E-06 | 0.460453 | 0.474 | 0.431 | 0.118581518 |
| DLX5_Cells | TRAM1 | 1.95141E-05 | 0.262175 | 0.58 | 0.538 | 0.369284813 |
| DLX5_Cells | BZW1 | 2.64941E-05 | 0.278852 | 0.61 | 0.574 | 0.50137369 |
| DLX5_Cells | CP | 4.86052E-05 | 0.536671 | 0.311 | 0.25 | 0.919805048 |
| DLX5_Cells | CD55 | 6.733E-05 | 0.297742 | 0.355 | 0.297 | 1 |
| DLX5_Cells | CCND1 | 7.75242E-05 | 0.550241 | 0.467 | 0.44 | 1 |
| DLX5_Cells | IER3 | 0.000123785 | 0.262412 | 0.353 | 0.291 | 1 |
| DLX5_Cells | ITGBL1 | 0.000315986 | 0.350998 | 0.622 | 0.606 | 1 |
| DLX5_Cells | NR4A1 | 0.000363097 | 0.353714 | 0.326 | 0.277 | 1 |
| Pericytes | COX4I2 | 0 | 2.390993 | 0.51 | 0.005 | 0 |
| Pericytes | HIGD1B | 0 | 1.651261 | 0.359 | 0 | 0 |
| Pericytes | NRARP | 0 | 1.347918 | 0.385 | 0.023 | 0 |
| Pericytes | FAM162B | 0 | 1.341263 | 0.318 | 0.007 | 0 |
| Pericytes | OLFML2A | 0 | 1.133349 | 0.291 | 0.009 | 0 |
| Pericytes | CPM | 0 | 0.996108 | 0.258 | 0.006 | 0 |
| Pericytes | PLXDC1 | 5.7849E-296 | 1.423064 | 0.367 | 0.023 | 1.0947E-291 |
| Pericytes | CCDC102B | 1.0796E-280 | 2.211468 | 0.709 | 0.132 | 2.0431E-276 |
| Pericytes | OR51E1 | 4.4786E-271 | 0.683767 | 0.172 | 0.001 | 8.4753E-267 |
| Pericytes | ABCC9 | 1.2106E-257 | 2.047448 | 0.514 | 0.069 | 2.2909E-253 |
| Pericytes | SLCO1C1 | 1.276E-257 | 0.757834 | 0.207 | 0.004 | 2.4147E-253 |
| Pericytes | CD36 | 5.1205E-251 | 1.906957 | 0.211 | 0.005 | 9.6901E-247 |
| Pericytes | LINGO1 | 1.2181E-240 | 0.783407 | 0.236 | 0.009 | 2.3052E-236 |
| Pericytes | NEURL1B | 3.555E-233 | 0.901296 | 0.248 | 0.011 | 6.7274E-229 |
| Pericytes | ENPEP | 1.2465E-229 | 0.651442 | 0.174 | 0.003 | 2.3589E-225 |

| | | | | | | |
|---|---|---|---|---|---|---|
| Pericytes | ADAP2 | 3.1131E-229 | 0.649369 | 0.178 | 0.003 | 5.8912E-225 |
| Pericytes | EDNRB | 3.5332E-207 | 0.837563 | 0.24 | 0.013 | 6.6862E-203 |
| Pericytes | TMEM74B | 7.1105E-204 | 0.440199 | 0.127 | 0.001 | 1.3456E-199 |
| Pericytes | NCKAP5 | 3.6144E-194 | 0.642055 | 0.15 | 0.003 | 6.8399E-190 |
| Pericytes | HOPX | 2.1464E-185 | 1.227469 | 0.311 | 0.031 | 4.0618E-181 |
| Pericytes | RGS16 | 2.1531E-185 | 2.15751 | 0.648 | 0.162 | 4.0745E-181 |
| Pericytes | EBF2 | 5.0216E-184 | 0.568992 | 0.152 | 0.003 | 9.5028E-180 |
| Pericytes | IGFBP4 | 1.7227E-182 | 1.999499 | 0.816 | 0.313 | 3.26E-178 |
| Pericytes | STEAP4 | 1.2304E-180 | 2.145542 | 0.549 | 0.113 | 2.3283E-176 |
| Pericytes | AC079822. | 6.3075E-176 | 0.676753 | 0.131 | 0.002 | 1.1936E-171 |
| Pericytes | PGF | 1.3429E-174 | 1.809392 | 0.744 | 0.252 | 2.5414E-170 |
| Pericytes | CRISPLD2 | 5.483E-169 | 0.975882 | 0.297 | 0.03 | 1.0376E-164 |
| Pericytes | NDUFA4L2 | 1.4175E-167 | 2.089741 | 0.861 | 0.353 | 2.6825E-163 |
| Pericytes | PTP4A3 | 2.631E-167 | 1.775622 | 0.623 | 0.175 | 4.9789E-163 |
| Pericytes | GUCY1A2 | 3.5345E-166 | 0.701695 | 0.199 | 0.011 | 6.6887E-162 |
| Pericytes | AC100803. | 2.8197E-163 | 0.700347 | 0.158 | 0.006 | 5.3361E-159 |
| Pericytes | TESC | 3.7142E-161 | 0.627106 | 0.113 | 0.001 | 7.0287E-157 |
| Pericytes | APOE | 3.6221E-160 | 1.796369 | 0.713 | 0.21 | 6.8545E-156 |
| Pericytes | HLA-B | 1.7978E-159 | 1.642461 | 0.986 | 0.758 | 3.4021E-155 |
| Pericytes | CYGB | 2.8853E-159 | 1.262827 | 0.287 | 0.031 | 5.4602E-155 |
| Pericytes | GMFG | 2.2604E-149 | 0.874978 | 0.232 | 0.02 | 4.2776E-145 |
| Pericytes | MUSTN1 | 1.0091E-147 | 2.270815 | 0.736 | 0.284 | 1.9096E-143 |
| Pericytes | TPPP3 | 1.7326E-147 | 1.394014 | 0.473 | 0.097 | 3.2788E-143 |
| Pericytes | ADGRF5 | 3.0515E-145 | 0.906705 | 0.268 | 0.028 | 5.7746E-141 |
| Pericytes | B2M | 6.1302E-145 | 1.124435 | 1 | 0.997 | 1.1601E-140 |
| Pericytes | CDH6 | 1.5597E-141 | 0.977912 | 0.311 | 0.043 | 2.9516E-137 |
| Pericytes | GUCY1B1 | 9.577E-141 | 1.468652 | 0.668 | 0.233 | 1.8124E-136 |
| Pericytes | GJA4 | 5.381E-140 | 1.512108 | 0.545 | 0.148 | 1.0183E-135 |
| Pericytes | FABP5 | 7.632E-133 | 1.46027 | 0.316 | 0.047 | 1.4443E-128 |
| Pericytes | KLHL23 | 1.4201E-132 | 1.154518 | 0.363 | 0.064 | 2.6873E-128 |
| Pericytes | SEPTIN4 | 7.3651E-132 | 1.595074 | 0.723 | 0.324 | 1.3938E-127 |
| Pericytes | LHFPL6 | 7.8032E-131 | 1.544044 | 0.934 | 0.759 | 1.4767E-126 |
| Pericytes | LPL | 1.7823E-129 | 0.680399 | 0.137 | 0.006 | 3.3727E-125 |
| Pericytes | THY1 | 1.7206E-126 | 2.333702 | 0.559 | 0.168 | 3.256E-122 |
| Pericytes | MT2A | 5.4982E-124 | 1.763107 | 0.947 | 0.748 | 1.0405E-119 |
| Pericytes | PHLDA1 | 6.6787E-122 | 1.571588 | 0.625 | 0.228 | 1.2639E-117 |
| Pericytes | LGI4 | 1.0454E-121 | 0.992864 | 0.305 | 0.048 | 1.9784E-117 |
| Pericytes | TBX2-AS1 | 4.7801E-120 | 1.285855 | 0.465 | 0.12 | 9.0459E-116 |
| Pericytes | IFITM2 | 3.4732E-118 | 1.28694 | 0.965 | 0.781 | 6.5727E-114 |
| Pericytes | TBX2 | 1.2376E-116 | 1.548925 | 0.596 | 0.22 | 2.3421E-112 |
| Pericytes | SERPINF1 | 5.2423E-115 | 1.26879 | 0.471 | 0.113 | 9.9205E-111 |
| Pericytes | SEPTIN7 | 5.6509E-115 | 1.005918 | 0.975 | 0.952 | 1.0694E-110 |
| Pericytes | PLAC9 | 8.341E-115 | 1.391073 | 0.939 | 0.621 | 1.5784E-110 |
| Pericytes | FXYD6 | 1.6124E-113 | 0.950444 | 0.266 | 0.038 | 3.0513E-109 |
| Pericytes | GJC1 | 2.0416E-113 | 0.994036 | 0.303 | 0.051 | 3.8636E-109 |
| Pericytes | HLA-A | 2.6359E-113 | 1.172761 | 0.986 | 0.881 | 4.9882E-109 |
| Pericytes | TDO2 | 3.2622E-111 | 0.944449 | 0.199 | 0.02 | 6.1733E-107 |
| Pericytes | CAMK2N1 | 1.2077E-109 | 1.300526 | 0.496 | 0.147 | 2.2854E-105 |
| Pericytes | ARHGAP29 | 2.6144E-109 | 1.493314 | 0.742 | 0.378 | 4.9474E-105 |
| Pericytes | ADAMTS4 | 4.4056E-108 | 0.514756 | 0.141 | 0.009 | 8.3371E-104 |
| Pericytes | RCSD1 | 1.2541E-107 | 0.75898 | 0.279 | 0.044 | 2.3733E-103 |
| Pericytes | ATP1B2 | 2.6E-107 | 0.720862 | 0.207 | 0.023 | 4.9203E-103 |
| Pericytes | CLMN | 6.7819E-107 | 0.73106 | 0.242 | 0.033 | 1.2834E-102 |
| Pericytes | PTMA | 2.2993E-106 | 0.698042 | 1 | 0.998 | 4.3512E-102 |
| Pericytes | KCNJ8 | 2.239E-105 | 1.055026 | 0.291 | 0.05 | 4.2371E-101 |
| Pericytes | IGKC | 2.6076E-104 | 1.588884 | 0.641 | 0.25 | 4.9347E-100 |
| Pericytes | BTG1 | 5.6054E-103 | 1.323388 | 0.906 | 0.702 | 1.06077E-98 |
| Pericytes | GPR4 | 3.7395E-101 | 0.445031 | 0.137 | 0.009 | 7.07665E-97 |

| | | | | | | |
|---|---|---|---|---|---|---|
| Pericytes | ANKRD29 | 3.975E-100 | 0.877789 | 0.295 | 0.054 | 7.52222E-96 |
| Pericytes | AGT | 4.80007E-98 | 1.298516 | 0.609 | 0.221 | 9.08365E-94 |
| Pericytes | APOLD1 | 8.37562E-98 | 0.72754 | 0.182 | 0.02 | 1.585E-93 |
| Pericytes | RGS5 | 2.1465E-97 | 1.51884 | 0.975 | 0.773 | 4.06204E-93 |
| Pericytes | HES4 | 2.93779E-97 | 1.401928 | 0.762 | 0.433 | 5.55947E-93 |
| Pericytes | NOTCH3 | 5.65551E-95 | 1.419184 | 0.725 | 0.39 | 1.07025E-90 |
| Pericytes | IFITM3 | 1.49231E-94 | 0.86272 | 1 | 0.98 | 2.82405E-90 |
| Pericytes | NR2F2 | 1.61361E-94 | 1.334459 | 0.879 | 0.623 | 3.0536E-90 |
| Pericytes | TMEM176 | 2.88995E-92 | 0.67445 | 0.125 | 0.008 | 5.46895E-88 |
| Pericytes | HLA-C | 1.78319E-88 | 0.899491 | 0.973 | 0.911 | 3.37451E-84 |
| Pericytes | TUSC3 | 1.43131E-87 | 0.526117 | 0.176 | 0.021 | 2.7086E-83 |
| Pericytes | EPS8 | 1.11214E-84 | 1.102292 | 0.818 | 0.584 | 2.10461E-80 |
| Pericytes | ANGPT2 | 5.90349E-84 | 1.279839 | 0.633 | 0.282 | 1.11718E-79 |
| Pericytes | H3F3B | 6.44038E-83 | 0.688561 | 0.996 | 0.986 | 1.21878E-78 |
| Pericytes | IGLC2 | 3.2961E-81 | 1.11301 | 0.463 | 0.149 | 6.23753E-77 |
| Pericytes | SLC7A2 | 6.55929E-81 | 0.979946 | 0.408 | 0.119 | 1.24128E-76 |
| Pericytes | ARHGAP42 | 1.11045E-80 | 0.957418 | 0.369 | 0.101 | 2.10142E-76 |
| Pericytes | EPHX1 | 2.50834E-77 | 1.367219 | 0.625 | 0.331 | 4.74677E-73 |
| Pericytes | GPR20 | 4.63773E-77 | 0.432191 | 0.139 | 0.014 | 8.77645E-73 |
| Pericytes | MYC | 1.48649E-76 | 1.187469 | 0.334 | 0.087 | 2.81303E-72 |
| Pericytes | C20orf27 | 2.63033E-75 | 1.391648 | 0.449 | 0.17 | 4.97764E-71 |
| Pericytes | ADGRA2 | 3.42468E-75 | 0.760559 | 0.275 | 0.06 | 6.48086E-71 |
| Pericytes | CD248 | 1.01811E-74 | 0.779706 | 0.215 | 0.037 | 1.92666E-70 |
| Pericytes | ISYNA1 | 6.53863E-74 | 1.253952 | 0.633 | 0.323 | 1.23737E-69 |
| Pericytes | NT5DC2 | 1.71769E-73 | 1.0587 | 0.49 | 0.193 | 3.25056E-69 |
| Pericytes | PHLDA2 | 2.19693E-72 | 1.363218 | 0.512 | 0.213 | 4.15747E-68 |
| Pericytes | OAZ2 | 5.91545E-72 | 1.126453 | 0.564 | 0.274 | 1.11944E-67 |
| Pericytes | IFITM1 | 7.36812E-72 | 0.968333 | 0.352 | 0.101 | 1.39434E-67 |
| Pericytes | KCNE4 | 2.90063E-71 | 1.314498 | 0.482 | 0.189 | 5.48915E-67 |
| Pericytes | ARPC5 | 3.48216E-71 | 0.915543 | 0.902 | 0.823 | 6.58964E-67 |
| Pericytes | FRMD3 | 6.35539E-70 | 0.665761 | 0.219 | 0.041 | 1.20269E-65 |
| Pericytes | PDGFRB | 6.44336E-70 | 1.207072 | 0.713 | 0.436 | 1.21934E-65 |
| Pericytes | EDNRA | 6.64205E-70 | 0.924155 | 0.371 | 0.114 | 1.25694E-65 |
| Pericytes | SERPINI1 | 6.91117E-70 | 1.066439 | 0.389 | 0.128 | 1.30787E-65 |
| Pericytes | ZFP36 | 1.46536E-68 | 1.506519 | 0.799 | 0.581 | 2.77305E-64 |
| Pericytes | SOCS3 | 8.03481E-67 | 1.484676 | 0.531 | 0.234 | 1.52051E-62 |
| Pericytes | FABP4 | 9.10766E-66 | 1.393429 | 0.186 | 0.031 | 1.72353E-61 |
| Pericytes | CHST2 | 1.88184E-65 | 0.348132 | 0.102 | 0.009 | 3.5612E-61 |
| Pericytes | COL18A1 | 6.06874E-64 | 1.196393 | 0.84 | 0.651 | 1.14845E-59 |
| Pericytes | AC100793. | 1.90956E-63 | 0.768662 | 0.291 | 0.077 | 3.61366E-59 |
| Pericytes | SCIN | 3.48282E-63 | 0.60435 | 0.172 | 0.028 | 6.5909E-59 |
| Pericytes | COL5A3 | 5.06617E-63 | 0.457921 | 0.15 | 0.021 | 9.58721E-59 |
| Pericytes | ZFP36L1 | 3.1085E-62 | 1.084685 | 0.762 | 0.533 | 5.88253E-58 |
| Pericytes | TINAGL1 | 3.3728E-61 | 0.935212 | 0.861 | 0.716 | 6.38268E-57 |
| Pericytes | TYMP | 4.29565E-60 | 0.902463 | 0.477 | 0.191 | 8.12909E-56 |
| Pericytes | SLC2A3 | 5.926E-60 | 0.874275 | 0.305 | 0.087 | 1.12144E-55 |
| Pericytes | PSME1 | 4.7472E-59 | 0.879302 | 0.842 | 0.706 | 8.98361E-55 |
| Pericytes | TRPC6 | 8.45319E-59 | 0.446962 | 0.143 | 0.02 | 1.59968E-54 |
| Pericytes | NRIP2 | 3.67423E-58 | 0.406525 | 0.133 | 0.018 | 6.95312E-54 |
| Pericytes | PPARG | 1.10206E-57 | 0.441822 | 0.145 | 0.022 | 2.08554E-53 |
| Pericytes | JUNB | 1.24538E-57 | 1.366019 | 0.879 | 0.692 | 2.35675E-53 |
| Pericytes | CYTOR | 7.77428E-57 | 1.146865 | 0.59 | 0.335 | 1.47121E-52 |
| Pericytes | TFPI | 2.14089E-56 | 1.145099 | 0.703 | 0.453 | 4.05141E-52 |
| Pericytes | RBPMS2 | 2.20253E-56 | 0.42851 | 0.129 | 0.017 | 4.16806E-52 |
| Pericytes | CABP1 | 6.22494E-56 | 0.358109 | 0.107 | 0.012 | 1.17801E-51 |
| Pericytes | UGCG | 1.74472E-55 | 0.87549 | 0.414 | 0.163 | 3.30171E-51 |
| Pericytes | IGHA1 | 3.11036E-55 | 0.752558 | 0.309 | 0.094 | 5.88604E-51 |
| Pericytes | LZTS1 | 3.69402E-55 | 0.413892 | 0.133 | 0.019 | 6.99055E-51 |

| | | | | | | |
|---|---|---|---|---|---|---|
| Pericytes | LAMA4 | 1.45999E-54 | 1.067288 | 0.467 | 0.211 | 2.76289E-50 |
| Pericytes | JUN | 1.76917E-53 | 1.423906 | 0.885 | 0.795 | 3.34798E-49 |
| Pericytes | SFTA1P | 2.4106E-53 | 0.53019 | 0.178 | 0.035 | 4.56181E-49 |
| Pericytes | OAF | 5.05517E-53 | 0.408813 | 0.123 | 0.017 | 9.56641E-49 |
| Pericytes | TGFBI | 9.39231E-53 | 0.938325 | 0.418 | 0.173 | 1.7774E-48 |
| Pericytes | FOS | 1.49274E-52 | 1.154665 | 0.9 | 0.697 | 2.82485E-48 |
| Pericytes | PCOLCE | 7.04884E-52 | 0.84033 | 0.793 | 0.532 | 1.33392E-47 |
| Pericytes | SPARCL1 | 1.01969E-51 | 1.030092 | 0.988 | 0.925 | 1.92967E-47 |
| Pericytes | ATP5MC2 | 9.00852E-51 | 0.654506 | 0.932 | 0.909 | 1.70477E-46 |
| Pericytes | ETS1 | 1.55974E-50 | 0.99353 | 0.459 | 0.219 | 2.95165E-46 |
| Pericytes | FAM13C | 3.5427E-49 | 0.93214 | 0.42 | 0.178 | 6.7042E-45 |
| Pericytes | SNRK | 3.80626E-48 | 0.730434 | 0.254 | 0.075 | 7.20296E-44 |
| Pericytes | IFI27 | 4.75107E-48 | 1.030783 | 0.391 | 0.148 | 8.99093E-44 |
| Pericytes | CHCHD10 | 8.52889E-48 | 0.879333 | 0.764 | 0.64 | 1.61401E-43 |
| Pericytes | IFI16 | 2.13092E-47 | 0.954803 | 0.678 | 0.444 | 4.03256E-43 |
| Pericytes | EPAS1 | 2.60087E-47 | 0.936228 | 0.734 | 0.544 | 4.92189E-43 |
| Pericytes | MYO1B | 3.17667E-47 | 1.134596 | 0.461 | 0.236 | 6.01153E-43 |
| Pericytes | EIF4A1 | 7.78787E-47 | 0.801304 | 0.852 | 0.831 | 1.47378E-42 |
| Pericytes | B3GNT2 | 3.02888E-46 | 0.736557 | 0.365 | 0.146 | 5.73186E-42 |
| Pericytes | KIAA0040 | 3.07528E-46 | 1.028362 | 0.408 | 0.186 | 5.81966E-42 |
| Pericytes | STOM | 7.1554E-46 | 0.889779 | 0.643 | 0.445 | 1.35409E-41 |
| Pericytes | UACA | 7.89753E-46 | 1.142801 | 0.717 | 0.563 | 1.49453E-41 |
| Pericytes | PNRC1 | 4.31492E-45 | 0.855535 | 0.904 | 0.874 | 8.16555E-41 |
| Pericytes | SLC25A5 | 1.87403E-44 | 0.904641 | 0.74 | 0.615 | 3.5464E-40 |
| Pericytes | H3F3A | 3.4192E-44 | 0.460935 | 0.982 | 0.977 | 6.47049E-40 |
| Pericytes | PTK2 | 5.00285E-44 | 0.825448 | 0.639 | 0.428 | 9.46739E-40 |
| Pericytes | GGT5 | 5.03449E-44 | 0.844624 | 0.408 | 0.168 | 9.52727E-40 |
| Pericytes | HIC1 | 2.47572E-43 | 0.555706 | 0.178 | 0.042 | 4.68506E-39 |
| Pericytes | RACK1 | 1.93829E-42 | 0.593726 | 0.98 | 0.984 | 3.66802E-38 |
| Pericytes | GUCY1A1 | 7.08989E-42 | 0.666403 | 0.789 | 0.574 | 1.34169E-37 |
| Pericytes | AC025164. | 4.96469E-41 | 0.410295 | 0.125 | 0.022 | 9.39518E-37 |
| Pericytes | SLCO3A1 | 1.03213E-40 | 0.863463 | 0.426 | 0.213 | 1.95321E-36 |
| Pericytes | SLC12A2 | 1.77127E-40 | 0.757721 | 0.221 | 0.066 | 3.35195E-36 |
| Pericytes | ARHGDIB | 4.58486E-40 | 0.919509 | 0.664 | 0.48 | 8.67639E-36 |
| Pericytes | CD81 | 1.25666E-39 | 0.457247 | 0.992 | 0.986 | 2.3781E-35 |
| Pericytes | ZEB2 | 2.83308E-39 | 0.911882 | 0.59 | 0.398 | 5.36133E-35 |
| Pericytes | ADCY3 | 3.71465E-39 | 0.522874 | 0.205 | 0.058 | 7.0296E-35 |
| Pericytes | DDX5 | 7.08832E-39 | 0.636737 | 0.918 | 0.881 | 1.34139E-34 |
| Pericytes | DLC1 | 1.21502E-38 | 0.867113 | 0.539 | 0.331 | 2.2993E-34 |
| Pericytes | COL4A1 | 1.28037E-38 | 1.111052 | 0.684 | 0.532 | 2.42296E-34 |
| Pericytes | ARHGAP15 | 4.25682E-37 | 0.419408 | 0.115 | 0.021 | 8.0556E-33 |
| Pericytes | PODXL2 | 4.48587E-37 | 0.460396 | 0.148 | 0.034 | 8.48905E-33 |
| Pericytes | BST2 | 4.94988E-37 | 0.662863 | 0.697 | 0.429 | 9.36716E-33 |
| Pericytes | CYSTM1 | 1.04789E-36 | 0.782973 | 0.715 | 0.589 | 1.98303E-32 |
| Pericytes | NDRG2 | 1.64235E-36 | 0.918951 | 0.482 | 0.284 | 3.10798E-32 |
| Pericytes | HOXB-AS1 | 1.70397E-36 | 0.644716 | 0.277 | 0.102 | 3.22459E-32 |
| Pericytes | FOSB | 1.84107E-36 | 1.034248 | 0.578 | 0.341 | 3.48403E-32 |
| Pericytes | RASD1 | 1.99877E-36 | 0.598094 | 0.133 | 0.028 | 3.78247E-32 |
| Pericytes | CHN1 | 5.03647E-36 | 0.775 | 0.273 | 0.103 | 9.53102E-32 |
| Pericytes | REM1 | 5.61492E-36 | 0.749877 | 0.291 | 0.116 | 1.06257E-31 |
| Pericytes | MAP3K7CL | 2.13774E-35 | 0.964781 | 0.336 | 0.15 | 4.04547E-31 |
| Pericytes | GPSM3 | 6.80541E-35 | 0.363254 | 0.119 | 0.024 | 1.28786E-30 |
| Pericytes | MARCKSL1 | 8.19052E-35 | 0.601987 | 0.18 | 0.05 | 1.54997E-30 |
| Pericytes | EEF1D | 1.02379E-34 | 0.506659 | 0.971 | 0.961 | 1.93743E-30 |
| Pericytes | INPP4B | 2.40207E-34 | 0.830538 | 0.381 | 0.189 | 4.54568E-30 |
| Pericytes | HLA-F | 2.61455E-34 | 0.762341 | 0.461 | 0.261 | 4.94777E-30 |
| Pericytes | GPRC5C | 2.92473E-34 | 0.868239 | 0.467 | 0.269 | 5.53475E-30 |
| Pericytes | MIR4435-2 | 8.57115E-34 | 0.670756 | 0.326 | 0.145 | 1.622E-29 |

| | | | | | | |
|---|---|---|---|---|---|---|
| Pericytes | MEF2B | 1.98669E-33 | 0.605679 | 0.152 | 0.038 | 3.75961E-29 |
| Pericytes | SH3KBP1 | 5.3815E-33 | 0.658469 | 0.309 | 0.132 | 1.0184E-28 |
| Pericytes | ARHGEF17 | 5.43679E-33 | 0.815033 | 0.445 | 0.254 | 1.02886E-28 |
| Pericytes | CCR10 | 1.37556E-32 | 0.468235 | 0.148 | 0.037 | 2.60311E-28 |
| Pericytes | EGR1 | 1.54214E-32 | 0.957647 | 0.668 | 0.469 | 2.91835E-28 |
| Pericytes | RNASEK | 2.52986E-31 | 0.484233 | 0.928 | 0.916 | 4.78751E-27 |
| Pericytes | TAGLN2 | 5.60994E-31 | 0.527673 | 0.891 | 0.831 | 1.06162E-26 |
| Pericytes | CSRP2 | 1.46076E-30 | 0.838558 | 0.746 | 0.538 | 2.76435E-26 |
| Pericytes | NACA | 1.58198E-30 | 0.350221 | 0.994 | 0.992 | 2.99374E-26 |
| Pericytes | MT1E | 2.34097E-30 | 1.094931 | 0.66 | 0.509 | 4.43006E-26 |
| Pericytes | JAK1 | 3.23231E-30 | 0.74128 | 0.631 | 0.499 | 6.11683E-26 |
| Pericytes | PPP1R15A | 3.57509E-30 | 0.912821 | 0.639 | 0.511 | 6.76549E-26 |
| Pericytes | NME2 | 2.37608E-29 | 0.448747 | 0.965 | 0.952 | 4.49649E-25 |
| Pericytes | CEBPD | 3.30433E-28 | 1.253858 | 0.715 | 0.589 | 6.25312E-24 |
| Pericytes | DUXAP8 | 3.58718E-28 | 0.540005 | 0.213 | 0.077 | 6.78838E-24 |
| Pericytes | SPARC | 6.30102E-28 | 0.560185 | 0.91 | 0.862 | 1.19241E-23 |
| Pericytes | NRP1 | 9.22113E-28 | 0.770877 | 0.43 | 0.247 | 1.74501E-23 |
| Pericytes | SYTL2 | 1.63507E-27 | 0.629033 | 0.26 | 0.109 | 3.09421E-23 |
| Pericytes | IGFBP7 | 2.23354E-27 | 0.297217 | 1 | 1 | 4.22675E-23 |
| Pericytes | HGF | 2.90039E-27 | 0.820007 | 0.348 | 0.18 | 5.4887E-23 |
| Pericytes | ANGPTL4 | 4.87992E-27 | 0.543064 | 0.176 | 0.057 | 9.23477E-23 |
| Pericytes | LBH | 8.76117E-27 | 1.043907 | 0.621 | 0.494 | 1.65796E-22 |
| Pericytes | CALM2 | 3.48928E-26 | 0.361523 | 0.994 | 0.985 | 6.60311E-22 |
| Pericytes | MT1A | 4.20166E-26 | 0.842628 | 0.193 | 0.069 | 7.95122E-22 |
| Pericytes | MGLL | 5.51071E-26 | 1.003631 | 0.498 | 0.363 | 1.04285E-21 |
| Pericytes | CYBA | 5.59391E-26 | 0.757692 | 0.621 | 0.47 | 1.05859E-21 |
| Pericytes | PLEKHH3 | 6.20687E-26 | 0.680751 | 0.369 | 0.204 | 1.17459E-21 |
| Pericytes | LGALSL | 6.46452E-26 | 0.638291 | 0.34 | 0.178 | 1.22335E-21 |
| Pericytes | KRT18 | 2.61756E-25 | 0.80299 | 0.225 | 0.085 | 4.95348E-21 |
| Pericytes | CCNL1 | 5.03602E-25 | 0.712074 | 0.467 | 0.305 | 9.53016E-21 |
| Pericytes | UBA2 | 6.42122E-25 | 0.689019 | 0.59 | 0.467 | 1.21515E-20 |
| Pericytes | MMP11 | 8.13139E-25 | 0.422168 | 0.168 | 0.055 | 1.53878E-20 |
| Pericytes | PFDN5 | 8.66601E-25 | 0.354217 | 0.984 | 0.968 | 1.63996E-20 |
| Pericytes | COL4A2 | 1.07618E-24 | 0.745695 | 0.768 | 0.682 | 2.03656E-20 |
| Pericytes | IER5L | 1.22872E-24 | 0.880914 | 0.475 | 0.321 | 2.32523E-20 |
| Pericytes | RELL1 | 1.26138E-24 | 0.567256 | 0.244 | 0.105 | 2.38703E-20 |
| Pericytes | INAFM1 | 1.6178E-24 | 0.839886 | 0.525 | 0.382 | 3.06153E-20 |
| Pericytes | FOXS1 | 3.4922E-24 | 0.676521 | 0.332 | 0.171 | 6.60863E-20 |
| Pericytes | RERGL | 4.08303E-24 | 1.243084 | 0.318 | 0.157 | 7.72672E-20 |
| Pericytes | ACYP1 | 6.29623E-24 | 0.634031 | 0.277 | 0.135 | 1.1915E-19 |
| Pericytes | ATF3 | 6.34009E-24 | 0.988338 | 0.566 | 0.408 | 1.1998E-19 |
| Pericytes | IER2 | 6.88781E-24 | 0.793773 | 0.734 | 0.624 | 1.30345E-19 |
| Pericytes | SUMO2 | 6.9903E-24 | 0.349838 | 0.957 | 0.951 | 1.32284E-19 |
| Pericytes | BATF3 | 1.11124E-23 | 0.412307 | 0.174 | 0.061 | 2.10291E-19 |
| Pericytes | ZNF385A | 1.36949E-23 | 0.338442 | 0.129 | 0.037 | 2.59163E-19 |
| Pericytes | POPDC2 | 1.5794E-23 | 0.386302 | 0.133 | 0.039 | 2.98886E-19 |
| Pericytes | LAMA3 | 3.25216E-23 | 0.301364 | 0.1 | 0.024 | 6.15439E-19 |
| Pericytes | FARP1 | 5.43534E-23 | 0.662883 | 0.289 | 0.146 | 1.02858E-18 |
| Pericytes | IRF1 | 1.19977E-22 | 0.539484 | 0.33 | 0.166 | 2.27045E-18 |
| Pericytes | CLIC1 | 3.22585E-22 | 0.418739 | 0.928 | 0.901 | 6.10461E-18 |
| Pericytes | DUSP5 | 8.98191E-22 | 0.308804 | 0.121 | 0.035 | 1.69974E-17 |
| Pericytes | CTSC | 1.05E-21 | 0.361961 | 0.441 | 0.254 | 1.98702E-17 |
| Pericytes | JAG1 | 1.14636E-21 | 0.736178 | 0.596 | 0.494 | 2.16938E-17 |
| Pericytes | EVA1B | 2.04342E-21 | 0.610695 | 0.666 | 0.578 | 3.86697E-17 |
| Pericytes | RASL11A | 2.13536E-21 | 0.611848 | 0.404 | 0.25 | 4.04096E-17 |
| Pericytes | MEF2C | 2.36661E-21 | 0.547876 | 0.807 | 0.76 | 4.47857E-17 |
| Pericytes | MIF | 2.57552E-21 | 0.379979 | 0.951 | 0.954 | 4.87392E-17 |
| Pericytes | HMGB1 | 3.89635E-21 | 0.34167 | 0.98 | 0.98 | 7.37345E-17 |

| | | | | | | |
|---|---|---|---|---|---|---|
| Pericytes | PPP1R10 | 6.43598E-21 | 0.677396 | 0.434 | 0.288 | 1.21794E-16 |
| Pericytes | KITLG | 1.33594E-20 | 0.458569 | 0.201 | 0.081 | 2.52813E-16 |
| Pericytes | MAPRE2 | 1.5663E-20 | 0.592024 | 0.369 | 0.222 | 2.96406E-16 |
| Pericytes | PDGFA | 2.33901E-20 | 0.584949 | 0.617 | 0.489 | 4.42635E-16 |
| Pericytes | ABTB1 | 2.78847E-20 | 0.60703 | 0.389 | 0.241 | 5.27689E-16 |
| Pericytes | SMARCB1 | 2.79907E-20 | 0.645227 | 0.414 | 0.284 | 5.29695E-16 |
| Pericytes | NTRK2 | 3.13035E-20 | 0.558876 | 0.273 | 0.133 | 5.92388E-16 |
| Pericytes | LIMD2 | 5.29721E-20 | 0.461323 | 0.168 | 0.064 | 1.00244E-15 |
| Pericytes | PRXL2A | 5.44891E-20 | 0.733223 | 0.359 | 0.22 | 1.03115E-15 |
| Pericytes | NFIL3 | 5.68538E-20 | 0.494688 | 0.201 | 0.085 | 1.0759E-15 |
| Pericytes | NES | 6.15125E-20 | 0.488012 | 0.209 | 0.091 | 1.16406E-15 |
| Pericytes | MCAM | 6.18583E-20 | 0.519477 | 0.756 | 0.577 | 1.17061E-15 |
| Pericytes | SMAP2 | 6.71728E-20 | 0.422309 | 0.17 | 0.065 | 1.27118E-15 |
| Pericytes | KNOP1 | 7.78322E-20 | 0.588214 | 0.266 | 0.134 | 1.4729E-15 |
| Pericytes | SELENOP | 9.77938E-20 | 0.568938 | 0.498 | 0.336 | 1.85065E-15 |
| Pericytes | TPI1 | 1.41585E-19 | 0.45692 | 0.836 | 0.831 | 2.67935E-15 |
| Pericytes | PPM1L | 1.69544E-19 | 0.500276 | 0.186 | 0.076 | 3.20846E-15 |
| Pericytes | BTF3 | 1.84357E-19 | 0.309269 | 0.98 | 0.975 | 3.48877E-15 |
| Pericytes | CAVIN3 | 2.23581E-19 | 0.467966 | 0.92 | 0.921 | 4.23105E-15 |
| Pericytes | SOX4 | 2.32894E-19 | 0.712571 | 0.467 | 0.314 | 4.40729E-15 |
| Pericytes | LY6E | 2.39632E-19 | 0.542547 | 0.6 | 0.447 | 4.5348E-15 |
| Pericytes | ADAMTS1 | 2.84962E-19 | 0.900714 | 0.541 | 0.415 | 5.39262E-15 |
| Pericytes | TMEM50A | 3.2472E-19 | 0.467681 | 0.754 | 0.715 | 6.14499E-15 |
| Pericytes | HES1 | 3.69728E-19 | 0.872329 | 0.465 | 0.328 | 6.99674E-15 |
| Pericytes | ESAM | 4.23058E-19 | 0.584759 | 0.486 | 0.346 | 8.00594E-15 |
| Pericytes | LINC01197 | 4.93123E-19 | 0.604448 | 0.277 | 0.146 | 9.33186E-15 |
| Pericytes | ODF3B | 6.34379E-19 | 0.528373 | 0.287 | 0.15 | 1.2005E-14 |
| Pericytes | LINC00989 | 6.54502E-19 | 0.32793 | 0.127 | 0.042 | 1.23858E-14 |
| Pericytes | COX4I1 | 7.76419E-19 | 0.369087 | 0.936 | 0.931 | 1.4693E-14 |
| Pericytes | NCK2 | 8.06019E-19 | 0.534005 | 0.309 | 0.175 | 1.52531E-14 |
| Pericytes | DLX5 | 8.07089E-19 | 0.332688 | 0.246 | 0.11 | 1.52734E-14 |
| Pericytes | PMEPA1 | 1.00573E-18 | 0.70499 | 0.77 | 0.725 | 1.90324E-14 |
| Pericytes | GBP4 | 1.28734E-18 | 0.354927 | 0.137 | 0.047 | 2.43617E-14 |
| Pericytes | CRIP1 | 2.90247E-18 | 0.419293 | 0.98 | 0.914 | 5.49263E-14 |
| Pericytes | CSRNP3 | 4.50867E-18 | 0.384108 | 0.131 | 0.045 | 8.53221E-14 |
| Pericytes | CFI | 5.64191E-18 | 0.429678 | 0.215 | 0.096 | 1.06767E-13 |
| Pericytes | RAB13 | 6.14374E-18 | 0.538133 | 0.732 | 0.712 | 1.16264E-13 |
| Pericytes | PSMB8 | 8.15049E-18 | 0.623692 | 0.473 | 0.356 | 1.5424E-13 |
| Pericytes | HRH2 | 1.17303E-17 | 0.410423 | 0.15 | 0.057 | 2.21984E-13 |
| Pericytes | THBS1 | 1.35907E-17 | 0.739837 | 0.232 | 0.112 | 2.5719E-13 |
| Pericytes | EHD2 | 1.36712E-17 | 0.555862 | 0.461 | 0.338 | 2.58713E-13 |
| Pericytes | ENTPD1 | 1.39803E-17 | 0.529512 | 0.223 | 0.109 | 2.64563E-13 |
| Pericytes | BTG2 | 8.34632E-17 | 0.859097 | 0.467 | 0.338 | 1.57946E-12 |
| Pericytes | ATP5F1B | 1.04895E-16 | 0.463495 | 0.684 | 0.651 | 1.98503E-12 |
| Pericytes | TNFAIP3 | 2.061E-16 | 0.367549 | 0.137 | 0.052 | 3.90023E-12 |
| Pericytes | CLEC11A | 2.18862E-16 | 0.457078 | 0.285 | 0.153 | 4.14174E-12 |
| Pericytes | GNG11 | 2.73135E-16 | 0.595663 | 0.629 | 0.557 | 5.16881E-12 |
| Pericytes | EPB41L1 | 3.26149E-16 | 0.486725 | 0.217 | 0.108 | 6.17204E-12 |
| Pericytes | ATP5F1D | 4.8016E-16 | 0.409399 | 0.82 | 0.823 | 9.08654E-12 |
| Pericytes | MAF | 5.06274E-16 | 0.469996 | 0.215 | 0.105 | 9.58073E-12 |
| Pericytes | A1BG | 5.61806E-16 | 0.532294 | 0.322 | 0.195 | 1.06316E-11 |
| Pericytes | MEIS3 | 6.70277E-16 | 0.254393 | 0.102 | 0.033 | 1.26843E-11 |
| Pericytes | SEM1 | 8.8025E-16 | 0.390372 | 0.85 | 0.833 | 1.66579E-11 |
| Pericytes | SSBP4 | 8.89232E-16 | 0.562341 | 0.439 | 0.322 | 1.68278E-11 |
| Pericytes | PEAK1 | 1.23516E-15 | 0.541041 | 0.246 | 0.133 | 2.33743E-11 |
| Pericytes | NR4A1 | 1.6554E-15 | 0.832058 | 0.4 | 0.274 | 3.13267E-11 |
| Pericytes | MDK | 2.26435E-15 | 0.508581 | 0.279 | 0.155 | 4.28505E-11 |
| Pericytes | PPP1CA | 2.90739E-15 | 0.571384 | 0.361 | 0.244 | 5.50194E-11 |

| | | | | | | |
|---|---|---|---|---|---|---|
| Pericytes | LOXL2 | 3.27262E-15 | 0.288292 | 0.121 | 0.044 | 6.1931E-11 |
| Pericytes | FAM241A | 3.29465E-15 | 0.439864 | 0.211 | 0.105 | 6.2348E-11 |
| Pericytes | ASAP1 | 3.70738E-15 | 0.561621 | 0.309 | 0.191 | 7.01585E-11 |
| Pericytes | C5orf38 | 4.05391E-15 | 0.408589 | 0.189 | 0.088 | 7.67162E-11 |
| Pericytes | SLC25A6 | 4.28042E-15 | 0.407287 | 0.867 | 0.861 | 8.10027E-11 |
| Pericytes | MOCS1 | 4.31728E-15 | 0.369627 | 0.148 | 0.061 | 8.17002E-11 |
| Pericytes | DBNDD2 | 5.48299E-15 | 0.393423 | 0.828 | 0.81 | 1.0376E-10 |
| Pericytes | PTEN | 6.45686E-15 | 0.529116 | 0.535 | 0.444 | 1.2219E-10 |
| Pericytes | EBF1 | 7.63838E-15 | 0.537364 | 0.449 | 0.33 | 1.44549E-10 |
| Pericytes | TUBA1A | 8.06691E-15 | 0.510199 | 0.848 | 0.831 | 1.52658E-10 |
| Pericytes | PSMB9 | 1.1512E-14 | 0.484916 | 0.379 | 0.248 | 2.17853E-10 |
| Pericytes | C12orf57 | 1.24847E-14 | 0.326328 | 0.951 | 0.946 | 2.3626E-10 |
| Pericytes | UBXN1 | 1.28811E-14 | 0.408822 | 0.766 | 0.744 | 2.43761E-10 |
| Pericytes | CD63 | 1.3333E-14 | 0.270507 | 0.996 | 0.994 | 2.52314E-10 |
| Pericytes | LDHB | 1.35291E-14 | 0.418048 | 0.758 | 0.747 | 2.56025E-10 |
| Pericytes | EIF3K | 1.41861E-14 | 0.338094 | 0.9 | 0.898 | 2.68457E-10 |
| Pericytes | TCF4 | 1.61656E-14 | 0.541203 | 0.654 | 0.583 | 3.05917E-10 |
| Pericytes | FHOD1 | 1.61771E-14 | 0.287044 | 0.1 | 0.034 | 3.06135E-10 |
| Pericytes | ARHGAP6 | 1.82828E-14 | 0.539299 | 0.262 | 0.15 | 3.45983E-10 |
| Pericytes | HOXB2 | 2.01205E-14 | 0.519307 | 0.328 | 0.209 | 3.8076E-10 |
| Pericytes | C4orf48 | 2.36331E-14 | 0.456067 | 0.178 | 0.083 | 4.47234E-10 |
| Pericytes | JUND | 2.52456E-14 | 0.503466 | 0.926 | 0.918 | 4.77747E-10 |
| Pericytes | ARID5B | 3.13157E-14 | 0.441064 | 0.803 | 0.76 | 5.92619E-10 |
| Pericytes | CCL4 | 3.69457E-14 | 0.416125 | 0.236 | 0.125 | 6.99161E-10 |
| Pericytes | FZD10-AS1 | 6.23888E-14 | 0.254801 | 0.115 | 0.042 | 1.18065E-09 |
| Pericytes | IQCK | 6.63306E-14 | 0.308528 | 0.121 | 0.047 | 1.25524E-09 |
| Pericytes | ARPC3 | 8.34565E-14 | 0.332572 | 0.844 | 0.839 | 1.57933E-09 |
| Pericytes | COX6A1 | 8.49168E-14 | 0.334805 | 0.906 | 0.907 | 1.60697E-09 |
| Pericytes | NRGN | 1.86625E-13 | 0.330908 | 0.514 | 0.377 | 3.5317E-09 |
| Pericytes | CPE | 2.1091E-13 | 0.719841 | 0.826 | 0.777 | 3.99127E-09 |
| Pericytes | IGHG1 | 2.64612E-13 | 0.268448 | 0.107 | 0.039 | 5.00751E-09 |
| Pericytes | BCAM | 2.94969E-13 | 0.692497 | 0.465 | 0.359 | 5.582E-09 |
| Pericytes | HMGN2 | 3.07635E-13 | 0.41136 | 0.787 | 0.77 | 5.82169E-09 |
| Pericytes | MRPL32 | 3.84544E-13 | 0.507526 | 0.502 | 0.421 | 7.2771E-09 |
| Pericytes | VEGFA | 4.50136E-13 | 0.526333 | 0.16 | 0.073 | 8.51837E-09 |
| Pericytes | HIGD2A | 5.63476E-13 | 0.60402 | 0.5 | 0.431 | 1.06632E-08 |
| Pericytes | VMP1 | 5.80984E-13 | 0.510488 | 0.502 | 0.404 | 1.09945E-08 |
| Pericytes | CCT5 | 6.70416E-13 | 0.461272 | 0.533 | 0.476 | 1.2687E-08 |
| Pericytes | SIX1 | 8.17492E-13 | 0.293471 | 0.143 | 0.062 | 1.54702E-08 |
| Pericytes | TMEM256 | 9.04875E-13 | 0.592383 | 0.52 | 0.449 | 1.71239E-08 |
| Pericytes | PHLDB2 | 9.5625E-13 | 0.553065 | 0.387 | 0.282 | 1.80961E-08 |
| Pericytes | PLEKHA4 | 9.78598E-13 | 0.470218 | 0.334 | 0.224 | 1.8519E-08 |
| Pericytes | YBX3 | 1.04585E-12 | 0.407811 | 0.846 | 0.851 | 1.97917E-08 |
| Pericytes | TCIM | 1.06262E-12 | 0.495093 | 0.125 | 0.051 | 2.0109E-08 |
| Pericytes | TIMP3 | 1.1448E-12 | 0.58509 | 0.592 | 0.486 | 2.16643E-08 |
| Pericytes | CCL19 | 1.73218E-12 | 1.186757 | 0.139 | 0.06 | 3.27798E-08 |
| Pericytes | GADD45B | 2.01356E-12 | 1.002312 | 0.617 | 0.555 | 3.81045E-08 |
| Pericytes | UTRN | 2.29951E-12 | 0.530726 | 0.482 | 0.389 | 4.3516E-08 |
| Pericytes | RAP1B | 2.44574E-12 | 0.478549 | 0.627 | 0.611 | 4.62832E-08 |
| Pericytes | LYSMD2 | 2.49709E-12 | 0.429973 | 0.305 | 0.2 | 4.72549E-08 |
| Pericytes | EPHA3 | 3.52826E-12 | 0.310613 | 0.166 | 0.078 | 6.67687E-08 |
| Pericytes | SLIT3 | 4.69659E-12 | 0.547937 | 0.387 | 0.282 | 8.88782E-08 |
| Pericytes | GALNT16 | 5.36118E-12 | 0.28495 | 0.117 | 0.048 | 1.01455E-07 |
| Pericytes | NCL | 5.75969E-12 | 0.376829 | 0.818 | 0.818 | 1.08996E-07 |
| Pericytes | BCL6 | 5.98754E-12 | 0.430282 | 0.191 | 0.101 | 1.13308E-07 |
| Pericytes | C1orf54 | 6.31702E-12 | 0.503271 | 0.463 | 0.393 | 1.19543E-07 |
| Pericytes | NOP53 | 6.64412E-12 | 0.365219 | 0.793 | 0.772 | 1.25733E-07 |
| Pericytes | GSTK1 | 6.74417E-12 | 0.524784 | 0.529 | 0.466 | 1.27627E-07 |

| | | | | | | |
|---|---|---|---|---|---|---|
| Pericytes | HNRNPA2I | 6.77503E-12 | 0.325191 | 0.877 | 0.895 | 1.28211E-07 |
| Pericytes | POLD4 | 7.4227E-12 | 0.507246 | 0.561 | 0.517 | 1.40467E-07 |
| Pericytes | KCNS3 | 7.61268E-12 | 0.405125 | 0.166 | 0.083 | 1.44062E-07 |
| Pericytes | SERTAD1 | 8.22841E-12 | 0.671904 | 0.449 | 0.369 | 1.55714E-07 |
| Pericytes | HOXB4 | 8.87231E-12 | 0.28418 | 0.119 | 0.049 | 1.679E-07 |
| Pericytes | PLTP | 8.92219E-12 | 0.492269 | 0.318 | 0.206 | 1.68844E-07 |
| Pericytes | IGLC3 | 9.73227E-12 | 0.384131 | 0.158 | 0.076 | 1.84173E-07 |
| Pericytes | IGHG3 | 9.92118E-12 | 0.309076 | 0.1 | 0.039 | 1.87748E-07 |
| Pericytes | ALKBH7 | 1.00796E-11 | 0.445503 | 0.635 | 0.606 | 1.90746E-07 |
| Pericytes | S100A3 | 1.33813E-11 | 0.385799 | 0.168 | 0.085 | 2.53227E-07 |
| Pericytes | RAP2A | 1.38933E-11 | 0.430848 | 0.277 | 0.175 | 2.62916E-07 |
| Pericytes | CCL21 | 1.78902E-11 | 0.902604 | 0.137 | 0.061 | 3.38555E-07 |
| Pericytes | TSPAN12 | 2.16589E-11 | 0.308758 | 0.133 | 0.06 | 4.09874E-07 |
| Pericytes | NR3C1 | 2.49694E-11 | 0.456049 | 0.557 | 0.494 | 4.7252E-07 |
| Pericytes | PSMB10 | 2.5234E-11 | 0.446268 | 0.342 | 0.243 | 4.77527E-07 |
| Pericytes | PSMB3 | 2.7951E-11 | 0.378247 | 0.682 | 0.66 | 5.28945E-07 |
| Pericytes | RABGAP1 | 2.89579E-11 | 0.519543 | 0.467 | 0.377 | 5.48E-07 |
| Pericytes | PLCE1 | 3.13518E-11 | 0.385183 | 0.154 | 0.076 | 5.93301E-07 |
| Pericytes | H1FX | 3.13622E-11 | 0.57644 | 0.588 | 0.533 | 5.93498E-07 |
| Pericytes | METRN | 3.26572E-11 | 0.441784 | 0.408 | 0.32 | 6.18005E-07 |
| Pericytes | LGALS3 | 3.79886E-11 | 0.334407 | 0.891 | 0.898 | 7.18895E-07 |
| Pericytes | ANKRD28 | 4.11076E-11 | 0.410895 | 0.252 | 0.152 | 7.7792E-07 |
| Pericytes | TMEM38B | 4.71045E-11 | 0.33124 | 0.184 | 0.099 | 8.91405E-07 |
| Pericytes | PLEKHG2 | 4.72175E-11 | 0.264302 | 0.129 | 0.058 | 8.93544E-07 |
| Pericytes | WDR54 | 4.83909E-11 | 0.394694 | 0.191 | 0.105 | 9.15749E-07 |
| Pericytes | CYP7B1 | 6.50753E-11 | 0.312792 | 0.137 | 0.064 | 1.23149E-06 |
| Pericytes | BAZ1A | 6.69087E-11 | 0.36527 | 0.189 | 0.103 | 1.26618E-06 |
| Pericytes | PSMA7 | 6.77571E-11 | 0.286515 | 0.887 | 0.905 | 1.28223E-06 |
| Pericytes | YWHAB | 6.96992E-11 | 0.289966 | 0.852 | 0.865 | 1.31899E-06 |
| Pericytes | NID1 | 7.56853E-11 | 0.313258 | 0.154 | 0.076 | 1.43227E-06 |
| Pericytes | FAT1 | 7.67296E-11 | 0.48859 | 0.334 | 0.238 | 1.45203E-06 |
| Pericytes | AXL | 9.42764E-11 | 0.468122 | 0.43 | 0.341 | 1.78409E-06 |
| Pericytes | NFASC | 1.16024E-10 | 0.291764 | 0.127 | 0.058 | 2.19564E-06 |
| Pericytes | RNF152 | 1.21674E-10 | 0.389507 | 0.197 | 0.109 | 2.30257E-06 |
| Pericytes | HVCN1 | 1.26225E-10 | 0.279617 | 0.109 | 0.046 | 2.38869E-06 |
| Pericytes | ATP5PD | 1.29483E-10 | 0.340455 | 0.766 | 0.771 | 2.45034E-06 |
| Pericytes | ATP5MC3 | 1.34561E-10 | 0.370445 | 0.785 | 0.807 | 2.54643E-06 |
| Pericytes | MAML2 | 1.72782E-10 | 0.47556 | 0.32 | 0.225 | 3.26972E-06 |
| Pericytes | HNRNPAB | 1.91714E-10 | 0.470937 | 0.48 | 0.414 | 3.628E-06 |
| Pericytes | DDIT4 | 2.23505E-10 | 0.38221 | 0.26 | 0.16 | 4.22961E-06 |
| Pericytes | NFATC4 | 2.25109E-10 | 0.269716 | 0.111 | 0.047 | 4.25996E-06 |
| Pericytes | SRGN | 2.25527E-10 | 0.410986 | 0.43 | 0.33 | 4.26786E-06 |
| Pericytes | SPRY2 | 2.28559E-10 | 0.429292 | 0.191 | 0.108 | 4.32526E-06 |
| Pericytes | YWHAZ | 2.85159E-10 | 0.370231 | 0.689 | 0.678 | 5.39635E-06 |
| Pericytes | TLE1 | 3.97591E-10 | 0.42184 | 0.285 | 0.191 | 7.52401E-06 |
| Pericytes | PKIG | 4.00285E-10 | 0.465809 | 0.6 | 0.566 | 7.57499E-06 |
| Pericytes | IER3 | 4.34656E-10 | 0.549741 | 0.393 | 0.291 | 8.22544E-06 |
| Pericytes | LGALS3BP | 4.42208E-10 | 0.35856 | 0.592 | 0.524 | 8.36835E-06 |
| Pericytes | H2AFZ | 4.81687E-10 | 0.307026 | 0.77 | 0.753 | 9.11544E-06 |
| Pericytes | CSRNP1 | 4.9249E-10 | 0.413473 | 0.197 | 0.114 | 9.31988E-06 |
| Pericytes | SMC6 | 6.49409E-10 | 0.386328 | 0.193 | 0.11 | 1.22894E-05 |
| Pericytes | PSME2 | 6.64487E-10 | 0.4369 | 0.564 | 0.506 | 1.25748E-05 |
| Pericytes | ATP6V1G1 | 7.4744E-10 | 0.325561 | 0.795 | 0.815 | 1.41446E-05 |
| Pericytes | PDE1A | 7.6164E-10 | 0.483923 | 0.404 | 0.318 | 1.44133E-05 |
| Pericytes | PLAAT4 | 8.16924E-10 | 0.34035 | 0.391 | 0.285 | 1.54595E-05 |
| Pericytes | ODF2L | 9.70011E-10 | 0.454756 | 0.26 | 0.171 | 1.83565E-05 |
| Pericytes | ADIRF | 1.01334E-09 | 0.336475 | 0.994 | 0.993 | 1.91764E-05 |
| Pericytes | MT1F | 1.08958E-09 | 0.335269 | 0.152 | 0.078 | 2.06192E-05 |

| | | | | | | |
|---|---|---|---|---|---|---|
| Pericytes | GPX3 | 1.14503E-09 | 0.428269 | 0.459 | 0.374 | 2.16685E-05 |
| Pericytes | EGFLAM | 1.50585E-09 | 0.357591 | 0.15 | 0.078 | 2.84967E-05 |
| Pericytes | GLRX5 | 1.6037E-09 | 0.442038 | 0.51 | 0.467 | 3.03484E-05 |
| Pericytes | TIPARP | 1.73479E-09 | 0.343447 | 0.162 | 0.087 | 3.28292E-05 |
| Pericytes | ATP2B1-AS | 1.86334E-09 | 0.256153 | 0.109 | 0.048 | 3.52619E-05 |
| Pericytes | IFIT3 | 1.86452E-09 | 0.329441 | 0.186 | 0.104 | 3.52842E-05 |
| Pericytes | ZMIZ1 | 3.28261E-09 | 0.281813 | 0.156 | 0.083 | 6.21202E-05 |
| Pericytes | TSC22D1 | 3.77212E-09 | 0.445011 | 0.74 | 0.71 | 7.13836E-05 |
| Pericytes | FUS | 3.99193E-09 | 0.39149 | 0.625 | 0.613 | 7.55432E-05 |
| Pericytes | NR2F2-AS1 | 4.22504E-09 | 0.418318 | 0.236 | 0.149 | 7.99546E-05 |
| Pericytes | OSTF1 | 4.36775E-09 | 0.438257 | 0.305 | 0.226 | 8.26553E-05 |
| Pericytes | ARPC1A | 4.59187E-09 | 0.443263 | 0.58 | 0.562 | 8.68966E-05 |
| Pericytes | ASXL1 | 4.8194E-09 | 0.42773 | 0.279 | 0.196 | 9.12024E-05 |
| Pericytes | CASQ2 | 4.99671E-09 | 0.363008 | 0.143 | 0.074 | 9.45577E-05 |
| Pericytes | GLRX | 5.20628E-09 | 0.408945 | 0.305 | 0.216 | 9.85237E-05 |
| Pericytes | ARGLU1 | 6.50646E-09 | 0.394938 | 0.691 | 0.672 | 0.000123128 |
| Pericytes | SPATS2L | 6.69525E-09 | 0.489857 | 0.52 | 0.472 | 0.000126701 |
| Pericytes | SERPINB6 | 7.33065E-09 | 0.363101 | 0.59 | 0.552 | 0.000138725 |
| Pericytes | RASSF4 | 7.40176E-09 | 0.296234 | 0.119 | 0.057 | 0.000140071 |
| Pericytes | ZNHIT1 | 7.40744E-09 | 0.376676 | 0.639 | 0.643 | 0.000140178 |
| Pericytes | DEK | 8.82942E-09 | 0.39834 | 0.709 | 0.723 | 0.000167088 |
| Pericytes | AC023157. | 1.00084E-08 | 0.368322 | 0.133 | 0.068 | 0.000189399 |
| Pericytes | FAS | 1.20997E-08 | 0.442135 | 0.266 | 0.184 | 0.000228975 |
| Pericytes | KLF9 | 1.40721E-08 | 0.434766 | 0.449 | 0.385 | 0.0002663 |
| Pericytes | SMOC2 | 1.44719E-08 | 0.356547 | 0.707 | 0.646 | 0.000273867 |
| Pericytes | IMPDH2 | 2.05585E-08 | 0.46513 | 0.379 | 0.31 | 0.00038905 |
| Pericytes | GCC2 | 2.08245E-08 | 0.390486 | 0.48 | 0.425 | 0.000394083 |
| Pericytes | HNRNPA3 | 2.22952E-08 | 0.322693 | 0.734 | 0.722 | 0.000421915 |
| Pericytes | FLYWCH2 | 2.5437E-08 | 0.423955 | 0.328 | 0.251 | 0.00048137 |
| Pericytes | ARPC2 | 3.23508E-08 | 0.309725 | 0.834 | 0.866 | 0.000612207 |
| Pericytes | SIGIRR | 3.85676E-08 | 0.404705 | 0.289 | 0.21 | 0.000729853 |
| Pericytes | PRXL2C | 3.94411E-08 | 0.446757 | 0.367 | 0.297 | 0.000746384 |
| Pericytes | CD40 | 4.83732E-08 | 0.409502 | 0.256 | 0.18 | 0.000915415 |
| Pericytes | DDX3X | 4.99585E-08 | 0.404066 | 0.512 | 0.466 | 0.000945415 |
| Pericytes | SERPINH1 | 6.00635E-08 | 0.431643 | 0.465 | 0.404 | 0.001136642 |
| Pericytes | LAP3 | 6.68601E-08 | 0.42002 | 0.367 | 0.295 | 0.001265261 |
| Pericytes | PGP | 6.69576E-08 | 0.427686 | 0.287 | 0.215 | 0.001267105 |
| Pericytes | PLA2G5 | 7.65337E-08 | 0.357962 | 0.115 | 0.058 | 0.001448323 |
| Pericytes | PRMT1 | 7.65572E-08 | 0.415251 | 0.551 | 0.531 | 0.001448768 |
| Pericytes | IGLC1 | 8.98077E-08 | 0.480476 | 0.123 | 0.064 | 0.001699522 |
| Pericytes | MEOX2 | 9.04099E-08 | 0.350875 | 0.145 | 0.082 | 0.001710916 |
| Pericytes | ARHGEF7 | 9.17046E-08 | 0.40357 | 0.225 | 0.152 | 0.001735417 |
| Pericytes | COX7B | 9.83257E-08 | 0.364231 | 0.748 | 0.766 | 0.001860715 |
| Pericytes | PLAU | 1.05187E-07 | 0.341907 | 0.15 | 0.086 | 0.001990562 |
| Pericytes | POMP | 1.09039E-07 | 0.27837 | 0.758 | 0.763 | 0.002063448 |
| Pericytes | CYC1 | 1.15736E-07 | 0.371504 | 0.539 | 0.514 | 0.002190185 |
| Pericytes | VASP | 1.26774E-07 | 0.399541 | 0.389 | 0.33 | 0.002399077 |
| Pericytes | IFIT2 | 1.55925E-07 | 0.457635 | 0.168 | 0.101 | 0.002950726 |
| Pericytes | CNN3 | 1.70775E-07 | 0.359352 | 0.693 | 0.702 | 0.003231744 |
| Pericytes | NDUFB2 | 1.84641E-07 | 0.299941 | 0.783 | 0.808 | 0.003494144 |
| Pericytes | ARHGAP10 | 1.87055E-07 | 0.412868 | 0.486 | 0.435 | 0.003539828 |
| Pericytes | UQCRH | 1.91733E-07 | 0.287891 | 0.809 | 0.844 | 0.003628353 |
| Pericytes | RBM39 | 2.08963E-07 | 0.298707 | 0.691 | 0.709 | 0.003954414 |
| Pericytes | PRRX1 | 2.37724E-07 | 0.411992 | 0.611 | 0.561 | 0.004498692 |
| Pericytes | TLE4 | 3.20158E-07 | 0.392854 | 0.258 | 0.184 | 0.006058669 |
| Pericytes | ANKRD50 | 3.54404E-07 | 0.366238 | 0.174 | 0.108 | 0.006706739 |
| Pericytes | TRA2B | 3.76811E-07 | 0.445656 | 0.443 | 0.39 | 0.007130765 |
| Pericytes | TSPAN14 | 3.82626E-07 | 0.264624 | 0.109 | 0.056 | 0.007240808 |

| | | | | | | |
|---|---|---|---|---|---|---|
| Pericytes | NREP | 4.25369E-07 | 0.381838 | 0.189 | 0.121 | 0.00804968 |
| Pericytes | CCDC151 | 4.29405E-07 | 0.387139 | 0.201 | 0.131 | 0.008126063 |
| Pericytes | NME3 | 4.46818E-07 | 0.367565 | 0.719 | 0.732 | 0.008455575 |
| Pericytes | MT1M | 4.59083E-07 | 0.837386 | 0.432 | 0.381 | 0.008687696 |
| Pericytes | CSF1 | 4.74521E-07 | 0.345839 | 0.18 | 0.114 | 0.008979828 |
| Pericytes | C1QBP | 5.08239E-07 | 0.402649 | 0.543 | 0.549 | 0.009617906 |
| Pericytes | TBCA | 5.91273E-07 | 0.271157 | 0.781 | 0.802 | 0.011189242 |
| Pericytes | PIM3 | 6.18904E-07 | 0.333633 | 0.244 | 0.173 | 0.011712137 |
| Pericytes | RFXANK | 6.9452E-07 | 0.350409 | 0.217 | 0.149 | 0.013143091 |
| Pericytes | LRRC32 | 7.11027E-07 | 0.397994 | 0.273 | 0.2 | 0.013455483 |
| Pericytes | ISG15 | 7.24354E-07 | 0.568013 | 0.35 | 0.284 | 0.013707677 |
| Pericytes | STAT3 | 7.62069E-07 | 0.376569 | 0.523 | 0.493 | 0.014421394 |
| Pericytes | RASGRP2 | 8.37854E-07 | 0.353301 | 0.268 | 0.196 | 0.015855545 |
| Pericytes | MICOS10 | 8.97873E-07 | 0.252584 | 0.797 | 0.814 | 0.01699135 |
| Pericytes | HNRNPU | 9.63267E-07 | 0.354169 | 0.656 | 0.653 | 0.018228864 |
| Pericytes | SNRPD2 | 1.06665E-06 | 0.257938 | 0.738 | 0.741 | 0.020185226 |
| Pericytes | FAM110B | 1.14221E-06 | 0.25262 | 0.123 | 0.068 | 0.021615144 |
| Pericytes | PSMA4 | 1.17424E-06 | 0.344281 | 0.531 | 0.507 | 0.022221384 |
| Pericytes | CYCS | 1.25377E-06 | 0.381232 | 0.52 | 0.517 | 0.023726322 |
| Pericytes | C19orf53 | 1.28694E-06 | 0.268819 | 0.775 | 0.815 | 0.024354104 |
| Pericytes | UBE2V1 | 1.42166E-06 | 0.369133 | 0.541 | 0.52 | 0.026903493 |
| Pericytes | ATP2B1 | 1.47073E-06 | 0.395548 | 0.227 | 0.161 | 0.027832146 |
| Pericytes | COMMD7 | 1.47923E-06 | 0.323077 | 0.377 | 0.322 | 0.027992883 |
| Pericytes | RBX1 | 1.50808E-06 | 0.273184 | 0.697 | 0.717 | 0.02853899 |
| Pericytes | IFI27L2 | 1.5139E-06 | 0.329838 | 0.541 | 0.505 | 0.028648983 |
| Pericytes | NDUFA7 | 1.66478E-06 | 0.401026 | 0.465 | 0.427 | 0.031504345 |
| Pericytes | CHSY1 | 1.84748E-06 | 0.327542 | 0.225 | 0.158 | 0.034961774 |
| Pericytes | SORBS3 | 1.85838E-06 | 0.383101 | 0.412 | 0.357 | 0.035168055 |
| Pericytes | BAX | 1.93753E-06 | 0.42726 | 0.477 | 0.444 | 0.036665906 |
| Pericytes | CFAP36 | 1.97499E-06 | 0.31905 | 0.502 | 0.464 | 0.03737478 |
| Pericytes | PABPC1 | 1.99991E-06 | 0.337316 | 0.664 | 0.677 | 0.037846244 |
| Pericytes | RBMS1 | 2.05695E-06 | 0.373784 | 0.449 | 0.416 | 0.038925694 |
| Pericytes | SAMHD1 | 2.10435E-06 | 0.407587 | 0.246 | 0.179 | 0.039822795 |
| Pericytes | MRPL34 | 2.14868E-06 | 0.287028 | 0.592 | 0.578 | 0.040661526 |
| Pericytes | SSR2 | 2.51454E-06 | 0.262618 | 0.74 | 0.75 | 0.047585162 |
| Pericytes | SLC38A2 | 2.70675E-06 | 0.377246 | 0.574 | 0.552 | 0.05122254 |
| Pericytes | RANBP1 | 2.80746E-06 | 0.345312 | 0.545 | 0.542 | 0.053128419 |
| Pericytes | C2orf27A | 2.96098E-06 | 0.261215 | 0.127 | 0.074 | 0.056033622 |
| Pericytes | GAS5 | 3.18508E-06 | 0.355966 | 0.666 | 0.664 | 0.060274381 |
| Pericytes | SMG7 | 3.75118E-06 | 0.274881 | 0.127 | 0.074 | 0.070987417 |
| Pericytes | DHRS4L2 | 3.76464E-06 | 0.375693 | 0.355 | 0.307 | 0.071242138 |
| Pericytes | CMTM6 | 4.17721E-06 | 0.333361 | 0.305 | 0.242 | 0.079049607 |
| Pericytes | MIR497HG | 4.48528E-06 | 0.354775 | 0.326 | 0.264 | 0.0848795 |
| Pericytes | SNRPB | 4.62865E-06 | 0.372629 | 0.545 | 0.55 | 0.087592495 |
| Pericytes | FOXN2 | 4.93982E-06 | 0.278661 | 0.158 | 0.1 | 0.093481131 |
| Pericytes | SPTSSA | 5.11717E-06 | 0.340222 | 0.359 | 0.3 | 0.096837414 |
| Pericytes | AP2M1 | 5.47581E-06 | 0.288338 | 0.617 | 0.625 | 0.103624225 |
| Pericytes | TECR | 5.52057E-06 | 0.347861 | 0.445 | 0.409 | 0.104471212 |
| Pericytes | MAP1A | 5.66095E-06 | 0.323889 | 0.24 | 0.176 | 0.107127851 |
| Pericytes | BCAS3 | 7.03907E-06 | 0.296309 | 0.143 | 0.089 | 0.133207319 |
| Pericytes | ATP5PB | 7.36971E-06 | 0.383928 | 0.484 | 0.466 | 0.1394643 |
| Pericytes | LAPTM4B | 7.46102E-06 | 0.371059 | 0.309 | 0.25 | 0.141192318 |
| Pericytes | ARHGAP17 | 7.5803E-06 | 0.290102 | 0.148 | 0.093 | 0.143449575 |
| Pericytes | SRRM2 | 7.58106E-06 | 0.2985 | 0.744 | 0.772 | 0.143464029 |
| Pericytes | MIDN | 7.6448E-06 | 0.35341 | 0.262 | 0.199 | 0.144670134 |
| Pericytes | ITM2C | 7.94387E-06 | 0.253082 | 0.496 | 0.418 | 0.150329883 |
| Pericytes | XRCC5 | 9.38585E-06 | 0.330344 | 0.631 | 0.652 | 0.17761785 |
| Pericytes | DIO2 | 9.98199E-06 | 0.364635 | 0.158 | 0.099 | 0.188899142 |

| | | | | | | |
|---|---|---|---|---|---|---|
| Pericytes | GPM6B | 1.02439E-05 | 0.330395 | 0.266 | 0.2 | 0.193856505 |
| Pericytes | SLFN5 | 1.05075E-05 | 0.299662 | 0.164 | 0.107 | 0.198844035 |
| Pericytes | SEMA5A | 1.07061E-05 | 0.330577 | 0.24 | 0.176 | 0.202601831 |
| Pericytes | MAP4K4 | 1.10747E-05 | 0.378163 | 0.307 | 0.247 | 0.209576976 |
| Pericytes | C1QTNF1 | 1.12662E-05 | 0.290615 | 0.514 | 0.464 | 0.213200946 |
| Pericytes | TMEM35B | 1.17873E-05 | 0.363137 | 0.543 | 0.526 | 0.223062099 |
| Pericytes | DAAM2 | 1.19046E-05 | 0.285874 | 0.174 | 0.116 | 0.225282564 |
| Pericytes | SERTAD3 | 1.20107E-05 | 0.346832 | 0.268 | 0.209 | 0.227290338 |
| Pericytes | FAM133B | 1.27537E-05 | 0.400706 | 0.445 | 0.422 | 0.241351518 |
| Pericytes | SMIM12 | 1.35476E-05 | 0.354127 | 0.34 | 0.291 | 0.256374618 |
| Pericytes | DNMT1 | 1.43758E-05 | 0.328941 | 0.176 | 0.12 | 0.272046712 |
| Pericytes | IFI35 | 1.47125E-05 | 0.349045 | 0.256 | 0.196 | 0.278419222 |
| Pericytes | PTPN2 | 1.54789E-05 | 0.282016 | 0.232 | 0.17 | 0.292922079 |
| Pericytes | TNC | 1.57318E-05 | 0.269012 | 0.25 | 0.177 | 0.297708781 |
| Pericytes | TUBB4B | 1.80702E-05 | 0.300709 | 0.574 | 0.566 | 0.341960656 |
| Pericytes | TKT | 1.88272E-05 | 0.284388 | 0.477 | 0.444 | 0.356285738 |
| Pericytes | IER5 | 2.03335E-05 | 0.328706 | 0.262 | 0.2 | 0.384790244 |
| Pericytes | ANO1 | 2.07394E-05 | 0.420257 | 0.291 | 0.234 | 0.392472558 |
| Pericytes | MCL1 | 2.20262E-05 | 0.372682 | 0.445 | 0.411 | 0.416823119 |
| Pericytes | OSBPL8 | 2.3333E-05 | 0.345812 | 0.475 | 0.453 | 0.441553746 |
| Pericytes | GABPB1-A | 2.55911E-05 | 0.280228 | 0.158 | 0.104 | 0.484285648 |
| Pericytes | PACSIN2 | 2.83271E-05 | 0.414397 | 0.32 | 0.268 | 0.536061618 |
| Pericytes | APRT | 2.88106E-05 | 0.359744 | 0.598 | 0.622 | 0.545211853 |
| Pericytes | PLCL1 | 3.12837E-05 | 0.357824 | 0.219 | 0.161 | 0.592012721 |
| Pericytes | H1F0 | 3.18461E-05 | 0.402543 | 0.279 | 0.221 | 0.60265637 |
| Pericytes | MRPL24 | 3.23876E-05 | 0.355888 | 0.299 | 0.249 | 0.612902848 |
| Pericytes | RUVBL1 | 3.26355E-05 | 0.310717 | 0.193 | 0.136 | 0.617593395 |
| Pericytes | STARD13 | 3.37142E-05 | 0.293813 | 0.145 | 0.096 | 0.638007212 |
| Pericytes | APEX1 | 3.39297E-05 | 0.352364 | 0.455 | 0.436 | 0.642085434 |
| Pericytes | KLHDC8B | 3.694E-05 | 0.263879 | 0.195 | 0.139 | 0.699052491 |
| Pericytes | LINC01615 | 3.83975E-05 | 0.297051 | 0.186 | 0.131 | 0.726634255 |
| Pericytes | IRF9 | 3.98693E-05 | 0.317343 | 0.369 | 0.321 | 0.754487386 |
| Pericytes | MGAT5 | 4.05191E-05 | 0.25971 | 0.137 | 0.087 | 0.766783033 |
| Pericytes | APOL6 | 4.17292E-05 | 0.289783 | 0.156 | 0.103 | 0.789684303 |
| Pericytes | CSNK2B | 4.39702E-05 | 0.295544 | 0.59 | 0.612 | 0.832092766 |
| Pericytes | ITGA7 | 4.51145E-05 | 0.373763 | 0.299 | 0.242 | 0.853747326 |
| Pericytes | MRTFB | 5.22497E-05 | 0.282097 | 0.135 | 0.086 | 0.988772646 |
| Pericytes | RDH5 | 5.28906E-05 | 0.300516 | 0.139 | 0.09 | 1 |
| Pericytes | SYNGR1 | 5.74872E-05 | 0.35493 | 0.244 | 0.19 | 1 |
| Pericytes | RSU1 | 6.95951E-05 | 0.318084 | 0.498 | 0.493 | 1 |
| Pericytes | TMOD3 | 7.3905E-05 | 0.317087 | 0.416 | 0.383 | 1 |
| Pericytes | PTGIR | 7.63835E-05 | 0.350587 | 0.332 | 0.284 | 1 |
| Pericytes | RAPGEF5 | 8.23155E-05 | 0.279234 | 0.219 | 0.163 | 1 |
| Pericytes | PTPRK | 9.43368E-05 | 0.27309 | 0.141 | 0.094 | 1 |
| Pericytes | TAPBP | 9.62895E-05 | 0.3136 | 0.352 | 0.309 | 1 |
| Pericytes | TRIB2 | 9.98595E-05 | 0.327183 | 0.174 | 0.124 | 1 |
| Pericytes | INTS6 | 0.00011168 | 0.295148 | 0.244 | 0.188 | 1 |
| Pericytes | DEPP1 | 0.000120638 | 0.610818 | 0.33 | 0.274 | 1 |
| Pericytes | ANTKMT | 0.00012274 | 0.318183 | 0.346 | 0.304 | 1 |
| Pericytes | NDUFS6 | 0.000126217 | 0.263348 | 0.674 | 0.706 | 1 |
| Pericytes | TUBA1C | 0.00013676 | 0.425865 | 0.457 | 0.458 | 1 |
| Pericytes | CSNK1E | 0.000139638 | 0.347452 | 0.305 | 0.264 | 1 |
| Pericytes | MTUS1 | 0.000145284 | 0.389465 | 0.301 | 0.25 | 1 |
| Pericytes | CDKN1A | 0.000149372 | 0.417932 | 0.314 | 0.26 | 1 |
| Pericytes | SFPQ | 0.000155068 | 0.311926 | 0.527 | 0.532 | 1 |
| Pericytes | ECE1 | 0.000163394 | 0.365068 | 0.26 | 0.211 | 1 |
| Pericytes | SNX6 | 0.000172467 | 0.29431 | 0.541 | 0.559 | 1 |
| Pericytes | TMX1 | 0.000173212 | 0.303824 | 0.275 | 0.225 | 1 |

| | | | | | | |
|---|---|---|---|---|---|---|
| Pericytes | LYRM2 | 0.000174587 | 0.317579 | 0.291 | 0.243 | 1 |
| Pericytes | GSTM3 | 0.000186441 | 0.399205 | 0.418 | 0.395 | 1 |
| Pericytes | ADGRL2 | 0.000200291 | 0.272327 | 0.156 | 0.109 | 1 |
| Pericytes | ARL3 | 0.000212702 | 0.300224 | 0.51 | 0.52 | 1 |
| Pericytes | GPATCH4 | 0.000224541 | 0.29805 | 0.186 | 0.139 | 1 |
| Pericytes | EIF4A3 | 0.000240386 | 0.329277 | 0.328 | 0.294 | 1 |
| Pericytes | RNASEH2C | 0.000256496 | 0.262207 | 0.639 | 0.666 | 1 |
| Pericytes | PELO | 0.000283237 | 0.388771 | 0.307 | 0.273 | 1 |
| Pericytes | TNFRSF1A | 0.000302328 | 0.342114 | 0.455 | 0.442 | 1 |
| Pericytes | JPT1 | 0.000321921 | 0.275924 | 0.164 | 0.119 | 1 |
| Pericytes | BHLHE40 | 0.00036727 | 0.264938 | 0.16 | 0.114 | 1 |
| Pericytes | GSDMD | 0.000423876 | 0.261619 | 0.309 | 0.267 | 1 |
| Pericytes | FADS3 | 0.000467842 | 0.323548 | 0.461 | 0.461 | 1 |
| Pericytes | SP100 | 0.000482606 | 0.258148 | 0.49 | 0.481 | 1 |
| Pericytes | FDPS | 0.000492582 | 0.278993 | 0.424 | 0.405 | 1 |
| Pericytes | XRN2 | 0.00053014 | 0.303288 | 0.449 | 0.44 | 1 |
| Pericytes | NECTIN2 | 0.000569896 | 0.338693 | 0.242 | 0.202 | 1 |
| Pericytes | PTPN12 | 0.000654399 | 0.394384 | 0.285 | 0.252 | 1 |
| Pericytes | IER3IP1 | 0.000684862 | 0.258293 | 0.459 | 0.457 | 1 |
| Pericytes | COX17 | 0.000701917 | 0.31078 | 0.609 | 0.622 | 1 |
| Pericytes | TMEM107 | 0.000716424 | 0.267866 | 0.174 | 0.13 | 1 |
| Pericytes | AGAP2-AS1 | 0.000743602 | 0.264347 | 0.115 | 0.078 | 1 |
| Pericytes | PUF60 | 0.0007591 | 0.264288 | 0.434 | 0.426 | 1 |
| Pericytes | DLEU2 | 0.000794286 | 0.254676 | 0.158 | 0.115 | 1 |
| Pericytes | TNS2 | 0.000827149 | 0.378129 | 0.309 | 0.276 | 1 |
| Pericytes | TMEM165 | 0.000871355 | 0.32886 | 0.469 | 0.476 | 1 |
| Pericytes | IRF2BPL | 0.000995927 | 0.282544 | 0.189 | 0.143 | 1 |
| Pericytes | ABRACL | 0.001117457 | 0.25927 | 0.201 | 0.158 | 1 |
| Pericytes | NAB1 | 0.001131173 | 0.345557 | 0.205 | 0.166 | 1 |
| Pericytes | A4GALT | 0.00121396 | 0.303753 | 0.268 | 0.232 | 1 |
| Pericytes | TXNL4A | 0.001323006 | 0.263982 | 0.453 | 0.456 | 1 |
| Pericytes | SRA1 | 0.001344521 | 0.343006 | 0.32 | 0.297 | 1 |
| Pericytes | HDGFL3 | 0.001347422 | 0.295777 | 0.406 | 0.392 | 1 |
| Pericytes | NOL3 | 0.001348812 | 0.303036 | 0.32 | 0.292 | 1 |
| Pericytes | BACH1 | 0.001354809 | 0.345365 | 0.223 | 0.186 | 1 |
| Pericytes | SPRY1 | 0.001420484 | 0.250036 | 0.373 | 0.327 | 1 |
| Pericytes | MAT2A | 0.00142468 | 0.282982 | 0.266 | 0.229 | 1 |
| Pericytes | TWF1 | 0.001448044 | 0.253141 | 0.234 | 0.194 | 1 |
| Pericytes | UBE2D1 | 0.001459476 | 0.302941 | 0.219 | 0.181 | 1 |
| Pericytes | SELENBP1 | 0.001465759 | 0.356947 | 0.324 | 0.292 | 1 |
| Pericytes | SNHG9 | 0.001468935 | 0.298687 | 0.252 | 0.213 | 1 |
| Pericytes | DNPH1 | 0.001493108 | 0.28235 | 0.307 | 0.274 | 1 |
| Pericytes | SNHG17 | 0.001530704 | 0.270718 | 0.205 | 0.165 | 1 |
| Pericytes | RSBN1L | 0.001678652 | 0.336293 | 0.451 | 0.453 | 1 |
| Pericytes | NTHL1 | 0.00169161 | 0.264766 | 0.242 | 0.203 | 1 |
| Pericytes | COBLL1 | 0.00206773 | 0.297466 | 0.344 | 0.32 | 1 |
| Pericytes | POLR2G | 0.002118466 | 0.295687 | 0.326 | 0.302 | 1 |
| Pericytes | ING1 | 0.002165995 | 0.302396 | 0.197 | 0.157 | 1 |
| Pericytes | LGMN | 0.00221489 | 0.293117 | 0.342 | 0.321 | 1 |
| Pericytes | DNAJC15 | 0.002249142 | 0.264772 | 0.471 | 0.48 | 1 |
| Pericytes | DMAC2L | 0.002288575 | 0.255712 | 0.166 | 0.129 | 1 |
| Pericytes | BBIP1 | 0.002588465 | 0.253058 | 0.254 | 0.218 | 1 |
| Pericytes | HNRNPC | 0.002764371 | 0.272355 | 0.508 | 0.527 | 1 |
| Pericytes | MIEN1 | 0.002851316 | 0.324218 | 0.346 | 0.333 | 1 |
| Pericytes | SPECC1 | 0.002881655 | 0.278042 | 0.361 | 0.342 | 1 |
| Pericytes | PSMC4 | 0.003007375 | 0.273169 | 0.406 | 0.403 | 1 |
| Pericytes | MAP2K2 | 0.003424801 | 0.257124 | 0.477 | 0.492 | 1 |
| Pericytes | MAP7D3 | 0.003684224 | 0.289855 | 0.389 | 0.379 | 1 |

| | | | | | | |
|---|---|---|---|---|---|---|
| Pericytes | PGLS | 0.003797953 | 0.261978 | 0.506 | 0.525 | 1 |
| Pericytes | DNAJB4 | 0.00389386 | 0.27834 | 0.436 | 0.414 | 1 |
| Pericytes | SYPL1 | 0.003991862 | 0.257402 | 0.508 | 0.538 | 1 |
| Pericytes | OLFML2B | 0.004011612 | 0.326945 | 0.281 | 0.243 | 1 |
| Pericytes | ZNF24 | 0.004086255 | 0.286858 | 0.34 | 0.319 | 1 |
| Pericytes | OSMR | 0.004172579 | 0.310816 | 0.205 | 0.171 | 1 |
| Pericytes | FHL3 | 0.00461075 | 0.29804 | 0.289 | 0.259 | 1 |
| Pericytes | ID4 | 0.004980283 | 0.457783 | 0.736 | 0.764 | 1 |
| Pericytes | HIF1A | 0.005323433 | 0.275464 | 0.34 | 0.317 | 1 |
| Pericytes | ANXA6 | 0.005491572 | 0.259259 | 0.457 | 0.462 | 1 |
| Pericytes | HSDL2 | 0.005622305 | 0.266813 | 0.318 | 0.292 | 1 |
| Pericytes | GBP1 | 0.005848629 | 0.379286 | 0.266 | 0.24 | 1 |
| Pericytes | BOLA3 | 0.0060819 | 0.305128 | 0.328 | 0.315 | 1 |
| Pericytes | IL34 | 0.006327184 | 0.297324 | 0.264 | 0.233 | 1 |
| Pericytes | SAT2 | 0.00700557 | 0.27752 | 0.418 | 0.423 | 1 |
| Pericytes | AC020916. | 0.007599276 | 0.266815 | 0.242 | 0.208 | 1 |
| Pericytes | ZNF22 | 0.007613622 | 0.309825 | 0.314 | 0.297 | 1 |
| Pericytes | CITED2 | 0.007625917 | 0.350273 | 0.359 | 0.336 | 1 |
| Pericytes | MED11 | 0.00764757 | 0.251258 | 0.207 | 0.174 | 1 |
| Pericytes | TIMM13 | 0.007771826 | 0.289827 | 0.385 | 0.386 | 1 |
| Pericytes | ECI1 | 0.008358806 | 0.273363 | 0.379 | 0.381 | 1 |
| Pericytes | SEPTIN9 | 0.008433739 | 0.25124 | 0.17 | 0.14 | 1 |
| Pericytes | TMEM173 | 0.008968554 | 0.27336 | 0.342 | 0.332 | 1 |
| Pericytes | CISD3 | 0.00960985 | 0.297332 | 0.359 | 0.355 | 1 |
| CRTAC1_Cells | SFRP5 | 0 | 2.720916 | 0.638 | 0.013 | 0 |
| CRTAC1_Cells | RAMP3 | 0 | 2.698095 | 0.765 | 0.019 | 0 |
| CRTAC1_Cells | EGFL7 | 0 | 2.110457 | 0.582 | 0.022 | 0 |
| CRTAC1_Cells | SGCG | 0 | 1.966432 | 0.765 | 0.03 | 0 |
| CRTAC1_Cells | PLVAP | 0 | 1.741994 | 0.393 | 0.008 | 0 |
| CRTAC1_Cells | SOX18 | 0 | 1.573352 | 0.403 | 0.01 | 0 |
| CRTAC1_Cells | BMP6 | 0 | 1.47205 | 0.51 | 0.01 | 0 |
| CRTAC1_Cells | PALMD | 0 | 1.228384 | 0.551 | 0.025 | 0 |
| CRTAC1_Cells | FLT1 | 0 | 1.047538 | 0.378 | 0.005 | 0 |
| CRTAC1_Cells | BMX | 1.7004E-286 | 0.84419 | 0.281 | 0.005 | 3.2179E-282 |
| CRTAC1_Cells | CRTAC1 | 1.8109E-256 | 3.190227 | 0.949 | 0.126 | 3.427E-252 |
| CRTAC1_Cells | TM6SF1 | 4.339E-226 | 0.587865 | 0.306 | 0.01 | 8.2111E-222 |
| CRTAC1_Cells | LTC4S | 3.0994E-193 | 2.416159 | 0.883 | 0.153 | 5.8653E-189 |
| CRTAC1_Cells | ECSCR | 2.1104E-192 | 0.714965 | 0.204 | 0.004 | 3.9936E-188 |
| CRTAC1_Cells | PLLP | 3.2157E-177 | 0.446177 | 0.24 | 0.007 | 6.0854E-173 |
| CRTAC1_Cells | SERPINE2 | 9.193E-169 | 2.595347 | 0.781 | 0.126 | 1.7397E-164 |
| CRTAC1_Cells | RAMP2 | 5.0832E-167 | 1.275977 | 0.378 | 0.024 | 9.6195E-163 |
| CRTAC1_Cells | HHIP-AS1 | 1.1318E-159 | 0.424048 | 0.214 | 0.007 | 2.1419E-155 |
| CRTAC1_Cells | GATA2 | 3.1621E-145 | 0.747216 | 0.439 | 0.039 | 5.984E-141 |
| CRTAC1_Cells | MASP1 | 3.1586E-141 | 0.753692 | 0.413 | 0.035 | 5.9772E-137 |
| CRTAC1_Cells | HHIP | 1.6284E-130 | 0.305302 | 0.117 | 0.002 | 3.0816E-126 |
| CRTAC1_Cells | PLAT | 3.1373E-129 | 1.78194 | 0.561 | 0.076 | 5.937E-125 |
| CRTAC1_Cells | PROCR | 2.8674E-128 | 1.725741 | 0.791 | 0.162 | 5.4262E-124 |
| CRTAC1_Cells | PNP | 3.3683E-126 | 0.869518 | 0.449 | 0.048 | 6.3741E-122 |
| CRTAC1_Cells | SHISA3 | 2.1613E-125 | 0.669494 | 0.357 | 0.03 | 4.09E-121 |
| CRTAC1_Cells | LMO2 | 1.0331E-122 | 1.503527 | 0.816 | 0.161 | 1.955E-118 |
| CRTAC1_Cells | ACVRL1 | 2.4914E-122 | 0.79348 | 0.383 | 0.035 | 4.7147E-118 |
| CRTAC1_Cells | LINC01133 | 7.0455E-121 | 0.584236 | 0.296 | 0.021 | 1.3333E-116 |
| CRTAC1_Cells | LYL1 | 5.9892E-112 | 0.525527 | 0.337 | 0.029 | 1.1334E-107 |
| CRTAC1_Cells | KCNT2 | 4.5667E-111 | 1.077233 | 0.561 | 0.084 | 8.6421E-107 |
| CRTAC1_Cells | TCN2 | 6.329E-109 | 0.705287 | 0.423 | 0.049 | 1.1977E-104 |
| CRTAC1_Cells | KCNK15 | 4.6527E-108 | 1.226519 | 0.638 | 0.114 | 8.8049E-104 |
| CRTAC1_Cells | PTGS1 | 1.1867E-105 | 0.528559 | 0.224 | 0.013 | 2.2457E-101 |
| CRTAC1_Cells | MARCKSL1 | 9.3675E-102 | 0.786858 | 0.413 | 0.049 | 1.77271E-97 |

| | | | | | | |
|---|---|---|---|---|---|---|
| CRTAC1_Cells | HHEX | 1.2583E-100 | 0.387338 | 0.209 | 0.012 | 2.38112E-96 |
| CRTAC1_Cells | EMCN | 5.1144E-100 | 1.246317 | 0.204 | 0.012 | 9.67853E-96 |
| CRTAC1_Cells | ENG | 1.76204E-95 | 1.908618 | 0.974 | 0.461 | 3.33448E-91 |
| CRTAC1_Cells | ANXA3 | 8.08357E-95 | 0.950555 | 0.393 | 0.049 | 1.52974E-90 |
| CRTAC1_Cells | ART4 | 2.23434E-94 | 1.073445 | 0.617 | 0.118 | 4.22826E-90 |
| CRTAC1_Cells | HTR2B | 7.91462E-94 | 0.708716 | 0.398 | 0.049 | 1.49776E-89 |
| CRTAC1_Cells | TNXB | 2.1267E-91 | 1.093444 | 0.587 | 0.108 | 4.02456E-87 |
| CRTAC1_Cells | EDN1 | 4.79072E-91 | 0.852928 | 0.296 | 0.028 | 9.06597E-87 |
| CRTAC1_Cells | MALL | 1.426E-90 | 0.572072 | 0.347 | 0.039 | 2.69856E-86 |
| CRTAC1_Cells | MGP | 1.79768E-90 | 2.003437 | 1 | 0.997 | 3.40192E-86 |
| CRTAC1_Cells | PLCXD3 | 4.57331E-89 | 0.638714 | 0.347 | 0.039 | 8.65453E-85 |
| CRTAC1_Cells | FAM167B | 5.66139E-87 | 0.371858 | 0.214 | 0.015 | 1.07136E-82 |
| CRTAC1_Cells | CD59 | 7.61456E-86 | 1.729525 | 0.995 | 0.843 | 1.44098E-81 |
| CRTAC1_Cells | SMOC1 | 6.65811E-83 | 0.635103 | 0.383 | 0.05 | 1.25998E-78 |
| CRTAC1_Cells | ANKRD1 | 7.72925E-83 | 1.1154 | 0.189 | 0.012 | 1.46268E-78 |
| CRTAC1_Cells | MMRN2 | 1.60361E-82 | 0.522112 | 0.265 | 0.025 | 3.03468E-78 |
| CRTAC1_Cells | SCN1B | 1.59735E-81 | 0.827414 | 0.449 | 0.073 | 3.02282E-77 |
| CRTAC1_Cells | FBLN2 | 8.72998E-80 | 1.091962 | 0.612 | 0.131 | 1.65206E-75 |
| CRTAC1_Cells | CTSC | 2.85994E-79 | 1.761011 | 0.806 | 0.252 | 5.41214E-75 |
| CRTAC1_Cells | TNFRSF11I | 4.4895E-76 | 2.684824 | 0.883 | 0.375 | 8.49594E-72 |
| CRTAC1_Cells | ARPC1B | 4.60524E-76 | 1.495095 | 0.974 | 0.667 | 8.71496E-72 |
| CRTAC1_Cells | PCDH1 | 2.58179E-75 | 0.342335 | 0.209 | 0.017 | 4.88578E-71 |
| CRTAC1_Cells | PTGS2 | 5.56045E-74 | 0.798357 | 0.311 | 0.039 | 1.05226E-69 |
| CRTAC1_Cells | ITPR2 | 6.30547E-74 | 0.840229 | 0.597 | 0.129 | 1.19325E-69 |
| CRTAC1_Cells | ASS1 | 6.6152E-74 | 1.023034 | 0.597 | 0.135 | 1.25186E-69 |
| CRTAC1_Cells | PODXL | 1.08776E-73 | 0.589684 | 0.25 | 0.025 | 2.05847E-69 |
| CRTAC1_Cells | CDC42EP2 | 1.50329E-71 | 0.40007 | 0.245 | 0.024 | 2.84482E-67 |
| CRTAC1_Cells | ADCY4 | 1.97553E-71 | 0.399113 | 0.173 | 0.012 | 3.73849E-67 |
| CRTAC1_Cells | DOK5 | 4.78002E-71 | 0.788967 | 0.561 | 0.121 | 9.0457E-67 |
| CRTAC1_Cells | EFEMP1 | 4.55072E-70 | 1.564434 | 0.995 | 0.773 | 8.61178E-66 |
| CRTAC1_Cells | KLF4 | 1.52629E-69 | 1.02823 | 0.607 | 0.145 | 2.88836E-65 |
| CRTAC1_Cells | FAP | 4.31804E-68 | 0.989891 | 0.765 | 0.222 | 8.17146E-64 |
| CRTAC1_Cells | GRASP | 1.27966E-65 | 0.703755 | 0.291 | 0.038 | 2.42163E-61 |
| CRTAC1_Cells | TM4SF1 | 1.5123E-63 | 1.573086 | 0.98 | 0.664 | 2.86187E-59 |
| CRTAC1_Cells | RAB38 | 1.7831E-62 | 0.478338 | 0.311 | 0.044 | 3.37435E-58 |
| CRTAC1_Cells | FHL2 | 3.09853E-62 | 1.233861 | 0.806 | 0.294 | 5.86366E-58 |
| CRTAC1_Cells | CD68 | 1.34548E-61 | 0.703712 | 0.459 | 0.092 | 2.54619E-57 |
| CRTAC1_Cells | FXYD5 | 1.51615E-61 | 1.281995 | 0.959 | 0.491 | 2.86916E-57 |
| CRTAC1_Cells | IER3 | 5.2981E-61 | 1.703692 | 0.796 | 0.285 | 1.00261E-56 |
| CRTAC1_Cells | SH3BGRL3 | 7.82238E-60 | 1.393382 | 0.995 | 0.808 | 1.48031E-55 |
| CRTAC1_Cells | OMD | 1.52492E-59 | 1.317613 | 0.929 | 0.442 | 2.88576E-55 |
| CRTAC1_Cells | CFH | 8.04322E-59 | 1.31274 | 0.969 | 0.443 | 1.5221E-54 |
| CRTAC1_Cells | HEG1 | 1.14794E-58 | 0.867033 | 0.546 | 0.134 | 2.17237E-54 |
| CRTAC1_Cells | TXNDC5 | 1.33836E-57 | 1.196234 | 0.709 | 0.248 | 2.53271E-53 |
| CRTAC1_Cells | PLA2G4A | 7.43026E-57 | 0.356104 | 0.255 | 0.033 | 1.4061E-52 |
| CRTAC1_Cells | CAVIN2 | 1.9808E-56 | 1.128299 | 0.418 | 0.085 | 3.74847E-52 |
| CRTAC1_Cells | PIR | 1.9803E-55 | 0.900292 | 0.51 | 0.124 | 3.74751E-51 |
| CRTAC1_Cells | SERPINE1 | 1.18242E-54 | 1.520999 | 0.796 | 0.315 | 2.23761E-50 |
| CRTAC1_Cells | ITM2B | 1.65701E-54 | 0.85341 | 0.995 | 0.987 | 3.13573E-50 |
| CRTAC1_Cells | MCUB | 1.12442E-53 | 0.491607 | 0.337 | 0.058 | 2.12785E-49 |
| CRTAC1_Cells | CD34 | 1.42538E-52 | 0.632841 | 0.464 | 0.105 | 2.6974E-48 |
| CRTAC1_Cells | KLF2 | 2.84433E-51 | 1.687425 | 0.903 | 0.477 | 5.38261E-47 |
| CRTAC1_Cells | IFITM2 | 1.15548E-50 | 1.13421 | 1 | 0.787 | 2.18662E-46 |
| CRTAC1_Cells | HTRA1 | 2.1993E-50 | 1.13521 | 0.98 | 0.796 | 4.16195E-46 |
| CRTAC1_Cells | PRR15 | 5.63558E-50 | 0.280815 | 0.173 | 0.017 | 1.06648E-45 |
| CRTAC1_Cells | PECAM1 | 1.32307E-49 | 0.475939 | 0.163 | 0.016 | 2.50378E-45 |
| CRTAC1_Cells | STAB1 | 1.55697E-49 | 0.288283 | 0.173 | 0.018 | 2.94642E-45 |
| CRTAC1_Cells | HLA-E | 3.48248E-49 | 1.226171 | 0.959 | 0.716 | 6.59025E-45 |

| | | | | | | |
|---|---|---|---|---|---|---|
| CRTAC1_Cells | TMEM255 | 8.04933E-49 | 0.469844 | 0.245 | 0.035 | 1.52326E-44 |
| CRTAC1_Cells | SRGN | 8.38217E-49 | 1.30573 | 0.765 | 0.325 | 1.58624E-44 |
| CRTAC1_Cells | MMP28 | 4.46665E-48 | 0.425727 | 0.281 | 0.046 | 8.45269E-44 |
| CRTAC1_Cells | SLC9A3R2 | 4.57733E-48 | 1.466344 | 0.796 | 0.375 | 8.66214E-44 |
| CRTAC1_Cells | ID1 | 4.11327E-47 | 1.51511 | 0.74 | 0.293 | 7.78396E-43 |
| CRTAC1_Cells | IL32 | 4.26107E-47 | 0.782729 | 0.561 | 0.161 | 8.06365E-43 |
| CRTAC1_Cells | QPCT | 1.02187E-46 | 0.311011 | 0.204 | 0.026 | 1.93379E-42 |
| CRTAC1_Cells | AKR1C1 | 3.94023E-46 | 1.045648 | 0.582 | 0.184 | 7.45649E-42 |
| CRTAC1_Cells | TMSB10 | 4.051E-46 | 0.935569 | 0.995 | 0.961 | 7.66611E-42 |
| CRTAC1_Cells | SMAD6 | 1.83022E-45 | 0.470721 | 0.316 | 0.059 | 3.46352E-41 |
| CRTAC1_Cells | PCOLCE2 | 2.12215E-45 | 1.271361 | 0.689 | 0.262 | 4.01596E-41 |
| CRTAC1_Cells | CNTNAP3E | 3.60389E-45 | 0.362072 | 0.25 | 0.038 | 6.82E-41 |
| CRTAC1_Cells | TPM3 | 4.65726E-45 | 0.888396 | 0.801 | 0.334 | 8.8134E-41 |
| CRTAC1_Cells | ABLIM1 | 5.13292E-45 | 0.50858 | 0.388 | 0.084 | 9.71353E-41 |
| CRTAC1_Cells | NPDC1 | 5.98504E-45 | 1.03734 | 0.832 | 0.42 | 1.13261E-40 |
| CRTAC1_Cells | HYAL2 | 2.62373E-44 | 0.572502 | 0.423 | 0.102 | 4.96514E-40 |
| CRTAC1_Cells | IFI27 | 1.0357E-43 | 1.407735 | 0.526 | 0.153 | 1.95997E-39 |
| CRTAC1_Cells | PLAUR | 2.7824E-43 | 0.571484 | 0.454 | 0.115 | 5.26542E-39 |
| CRTAC1_Cells | CARD16 | 5.48867E-43 | 0.445657 | 0.393 | 0.089 | 1.03867E-38 |
| CRTAC1_Cells | SERPINA3 | 2.63764E-42 | 1.266743 | 0.383 | 0.091 | 4.99147E-38 |
| CRTAC1_Cells | FRMD4B | 3.31636E-42 | 0.549217 | 0.561 | 0.162 | 6.27588E-38 |
| CRTAC1_Cells | IFITM3 | 4.43693E-42 | 0.939048 | 1 | 0.98 | 8.39644E-38 |
| CRTAC1_Cells | FSCN1 | 1.1679E-41 | 0.384824 | 0.316 | 0.063 | 2.21013E-37 |
| CRTAC1_Cells | HRH1 | 2.3622E-41 | 0.54007 | 0.444 | 0.115 | 4.47022E-37 |
| CRTAC1_Cells | KDELR3 | 3.3041E-41 | 0.632921 | 0.612 | 0.196 | 6.25269E-37 |
| CRTAC1_Cells | ARHGAP18 | 5.05616E-41 | 0.62808 | 0.505 | 0.145 | 9.56827E-37 |
| CRTAC1_Cells | EMP1 | 5.53462E-41 | 0.545339 | 0.531 | 0.152 | 1.04737E-36 |
| CRTAC1_Cells | MYL12B | 1.45368E-40 | 0.808108 | 0.995 | 0.959 | 2.75094E-36 |
| CRTAC1_Cells | PI16 | 2.09369E-40 | 0.442497 | 0.107 | 0.008 | 3.96209E-36 |
| CRTAC1_Cells | CLIC2 | 3.437E-40 | 0.290226 | 0.209 | 0.03 | 6.50418E-36 |
| CRTAC1_Cells | CD14 | 4.12879E-40 | 0.447979 | 0.311 | 0.063 | 7.81333E-36 |
| CRTAC1_Cells | ABCA8 | 6.27916E-40 | 0.380688 | 0.332 | 0.069 | 1.18827E-35 |
| CRTAC1_Cells | SNAI1 | 4.01788E-39 | 0.394818 | 0.245 | 0.042 | 7.60343E-35 |
| CRTAC1_Cells | TMEM204 | 4.09967E-39 | 0.996213 | 0.704 | 0.296 | 7.75821E-35 |
| CRTAC1_Cells | CUBN | 1.15734E-38 | 0.259587 | 0.168 | 0.021 | 2.19015E-34 |
| CRTAC1_Cells | LEPR | 1.6246E-38 | 0.48975 | 0.26 | 0.048 | 3.0744E-34 |
| CRTAC1_Cells | FERMT3 | 5.84949E-38 | 0.269012 | 0.204 | 0.031 | 1.10696E-33 |
| CRTAC1_Cells | LXN | 6.0675E-38 | 0.527215 | 0.357 | 0.084 | 1.14821E-33 |
| CRTAC1_Cells | TRAF1 | 6.24131E-38 | 0.451531 | 0.281 | 0.056 | 1.18111E-33 |
| CRTAC1_Cells | SERPINB1 | 8.11969E-38 | 0.803872 | 0.827 | 0.365 | 1.53657E-33 |
| CRTAC1_Cells | CYTL1 | 1.04029E-37 | 1.293807 | 0.423 | 0.118 | 1.96865E-33 |
| CRTAC1_Cells | SULT1A1 | 2.50931E-37 | 0.270148 | 0.194 | 0.028 | 4.74862E-33 |
| CRTAC1_Cells | GATA2-AS1 | 5.65976E-37 | 0.359379 | 0.214 | 0.035 | 1.07105E-32 |
| CRTAC1_Cells | S1PR1 | 6.71534E-37 | 0.261973 | 0.153 | 0.018 | 1.27081E-32 |
| CRTAC1_Cells | IGFBP4 | 2.4791E-36 | 1.225676 | 0.765 | 0.333 | 4.69144E-32 |
| CRTAC1_Cells | JAM2 | 3.47678E-36 | 0.361088 | 0.296 | 0.063 | 6.57946E-32 |
| CRTAC1_Cells | TIMP1 | 8.15606E-36 | 0.767156 | 1 | 0.975 | 1.54345E-31 |
| CRTAC1_Cells | CDH11 | 1.74553E-35 | 0.783635 | 0.76 | 0.329 | 3.30324E-31 |
| CRTAC1_Cells | ARHGDIB | 2.18773E-35 | 0.75091 | 0.929 | 0.481 | 4.14005E-31 |
| CRTAC1_Cells | ERG | 2.49861E-35 | 0.646718 | 0.658 | 0.249 | 4.72836E-31 |
| CRTAC1_Cells | THBD | 3.47844E-35 | 0.651236 | 0.474 | 0.145 | 6.5826E-31 |
| CRTAC1_Cells | DHRS3 | 5.57687E-35 | 0.653375 | 0.449 | 0.135 | 1.05537E-30 |
| CRTAC1_Cells | LPCAT2 | 6.90392E-35 | 0.424069 | 0.398 | 0.105 | 1.3065E-30 |
| CRTAC1_Cells | CMTM8 | 7.65084E-35 | 0.297328 | 0.158 | 0.021 | 1.44785E-30 |
| CRTAC1_Cells | ECE1 | 2.18155E-34 | 0.614804 | 0.592 | 0.204 | 4.12836E-30 |
| CRTAC1_Cells | CTSK | 4.5333E-34 | 0.608044 | 0.791 | 0.35 | 8.57883E-30 |
| CRTAC1_Cells | DUSP5 | 5.31681E-34 | 0.439912 | 0.209 | 0.036 | 1.00615E-29 |
| CRTAC1_Cells | CFI | 1.08884E-33 | 0.597006 | 0.367 | 0.097 | 2.06053E-29 |

| | | | | | | |
|---|---|---|---|---|---|---|
| CRTAC1_Cells | IL1R1 | 2.63529E-33 | 0.691674 | 0.663 | 0.265 | 4.98702E-29 |
| CRTAC1_Cells | PTTG1IP | 2.83459E-33 | 0.854621 | 0.939 | 0.609 | 5.36417E-29 |
| CRTAC1_Cells | NAMPT | 4.76979E-33 | 0.903491 | 0.592 | 0.228 | 9.02634E-29 |
| CRTAC1_Cells | CCN2 | 8.54232E-33 | 1.406804 | 0.974 | 0.817 | 1.61655E-28 |
| CRTAC1_Cells | F2R | 1.74702E-32 | 0.580601 | 0.663 | 0.25 | 3.30607E-28 |
| CRTAC1_Cells | YBX3 | 1.82925E-32 | 0.849838 | 0.985 | 0.848 | 3.46168E-28 |
| CRTAC1_Cells | JUNB | 2.89948E-32 | 0.867701 | 0.974 | 0.697 | 5.48698E-28 |
| CRTAC1_Cells | TGFB2 | 3.05308E-32 | 0.333959 | 0.347 | 0.087 | 5.77764E-28 |
| CRTAC1_Cells | MTUS1 | 3.68881E-32 | 0.697789 | 0.622 | 0.244 | 6.9807E-28 |
| CRTAC1_Cells | SRPX2 | 5.92532E-32 | 0.267729 | 0.235 | 0.045 | 1.12131E-27 |
| CRTAC1_Cells | PLSCR1 | 5.96605E-32 | 0.827493 | 0.663 | 0.276 | 1.12901E-27 |
| CRTAC1_Cells | CCL2 | 7.43901E-32 | 1.635489 | 0.643 | 0.258 | 1.40776E-27 |
| CRTAC1_Cells | LAPTM5 | 2.76964E-31 | 0.267575 | 0.163 | 0.024 | 5.24127E-27 |
| CRTAC1_Cells | VCAM1 | 3.2875E-31 | 0.807674 | 0.648 | 0.265 | 6.22126E-27 |
| CRTAC1_Cells | COL8A1 | 4.14089E-31 | 1.053512 | 0.878 | 0.594 | 7.83622E-27 |
| CRTAC1_Cells | ICAM1 | 4.78699E-31 | 0.385386 | 0.209 | 0.038 | 9.05891E-27 |
| CRTAC1_Cells | FGL2 | 4.82142E-31 | 0.991254 | 0.893 | 0.576 | 9.12405E-27 |
| CRTAC1_Cells | FKBP11 | 1.91902E-30 | 0.585669 | 0.454 | 0.149 | 3.63156E-26 |
| CRTAC1_Cells | IL18BP | 4.47743E-30 | 0.281492 | 0.204 | 0.037 | 8.47308E-26 |
| CRTAC1_Cells | RAB11A | 4.8484E-30 | 0.59937 | 0.724 | 0.321 | 9.17511E-26 |
| CRTAC1_Cells | NCOA7 | 1.80355E-29 | 0.718665 | 0.643 | 0.274 | 3.41304E-25 |
| CRTAC1_Cells | PAPSS2 | 2.24276E-29 | 0.304024 | 0.24 | 0.05 | 4.2442E-25 |
| CRTAC1_Cells | SOCS3 | 2.34006E-29 | 0.574803 | 0.617 | 0.242 | 4.42833E-25 |
| CRTAC1_Cells | ITGA5 | 3.4972E-29 | 0.424345 | 0.352 | 0.097 | 6.61809E-25 |
| CRTAC1_Cells | CCDC71L | 4.56035E-29 | 0.555658 | 0.571 | 0.212 | 8.63002E-25 |
| CRTAC1_Cells | NUCB2 | 6.54569E-29 | 0.757521 | 0.898 | 0.602 | 1.23871E-24 |
| CRTAC1_Cells | FKBP10 | 8.62647E-29 | 0.601466 | 0.765 | 0.353 | 1.63247E-24 |
| CRTAC1_Cells | ATP1B3 | 9.94659E-29 | 0.768556 | 0.776 | 0.384 | 1.88229E-24 |
| CRTAC1_Cells | SLC16A3 | 9.98764E-29 | 0.38083 | 0.321 | 0.085 | 1.89006E-24 |
| CRTAC1_Cells | C1R | 4.5665E-28 | 0.75916 | 0.959 | 0.672 | 8.64165E-24 |
| CRTAC1_Cells | SUSD6 | 9.52169E-28 | 0.309584 | 0.367 | 0.103 | 1.80188E-23 |
| CRTAC1_Cells | NT5E | 9.6497E-28 | 0.585212 | 0.536 | 0.204 | 1.82611E-23 |
| CRTAC1_Cells | GSN | 9.88465E-28 | 0.784144 | 0.995 | 0.967 | 1.87057E-23 |
| CRTAC1_Cells | TSPAN6 | 1.63654E-27 | 0.34174 | 0.311 | 0.082 | 3.09699E-23 |
| CRTAC1_Cells | BCO2 | 1.97826E-27 | 0.286661 | 0.128 | 0.017 | 3.74366E-23 |
| CRTAC1_Cells | MMP2 | 3.41584E-27 | 0.661782 | 0.74 | 0.337 | 6.46413E-23 |
| CRTAC1_Cells | NANS | 5.85055E-27 | 0.538088 | 0.622 | 0.256 | 1.10716E-22 |
| CRTAC1_Cells | C4orf48 | 9.44157E-27 | 0.337199 | 0.311 | 0.083 | 1.78672E-22 |
| CRTAC1_Cells | ABCA6 | 1.42101E-26 | 0.285404 | 0.24 | 0.054 | 2.68913E-22 |
| CRTAC1_Cells | NAV1 | 1.43058E-26 | 0.404358 | 0.393 | 0.122 | 2.70723E-22 |
| CRTAC1_Cells | GAS6 | 1.88105E-26 | 0.73619 | 0.98 | 0.865 | 3.5597E-22 |
| CRTAC1_Cells | KRT18 | 3.42738E-26 | 0.596801 | 0.316 | 0.088 | 6.48598E-22 |
| CRTAC1_Cells | GAP43 | 3.51322E-26 | 0.685096 | 0.388 | 0.124 | 6.64842E-22 |
| CRTAC1_Cells | HOMER3 | 6.4261E-26 | 0.457231 | 0.449 | 0.158 | 1.21608E-21 |
| CRTAC1_Cells | TCF4 | 6.8132E-26 | 0.638847 | 0.944 | 0.578 | 1.28933E-21 |
| CRTAC1_Cells | SLC20A1 | 7.4852E-26 | 0.378748 | 0.342 | 0.099 | 1.4165E-21 |
| CRTAC1_Cells | ABCA9 | 1.22476E-25 | 0.274977 | 0.143 | 0.023 | 2.31773E-21 |
| CRTAC1_Cells | PAPPA | 1.74158E-25 | 0.379508 | 0.245 | 0.059 | 3.29576E-21 |
| CRTAC1_Cells | ODF3B | 4.47787E-25 | 0.427773 | 0.444 | 0.151 | 8.47391E-21 |
| CRTAC1_Cells | BACE2 | 5.37841E-25 | 0.578469 | 0.699 | 0.318 | 1.01781E-20 |
| CRTAC1_Cells | CLIC3 | 7.06936E-25 | 1.011495 | 0.321 | 0.102 | 1.33781E-20 |
| CRTAC1_Cells | HSPA8 | 7.82186E-25 | 0.669591 | 0.974 | 0.845 | 1.48021E-20 |
| CRTAC1_Cells | HAPLN3 | 9.19342E-25 | 0.572233 | 0.449 | 0.167 | 1.73976E-20 |
| CRTAC1_Cells | NFKBIA | 1.4776E-24 | 1.061959 | 0.852 | 0.578 | 2.7962E-20 |
| CRTAC1_Cells | ID3 | 1.86701E-24 | 0.728898 | 0.969 | 0.797 | 3.53313E-20 |
| CRTAC1_Cells | APOL1 | 2.29374E-24 | 0.301731 | 0.321 | 0.091 | 4.34067E-20 |
| CRTAC1_Cells | MEG3 | 2.46262E-24 | 0.5527 | 0.352 | 0.111 | 4.66026E-20 |
| CRTAC1_Cells | B2M | 2.94007E-24 | 0.512072 | 1 | 0.997 | 5.56378E-20 |

| | | | | | | |
|---|---|---|---|---|---|---|
| CRTAC1_Cells | RARRES1 | 2.97986E-24 | 0.436643 | 0.276 | 0.074 | 5.63908E-20 |
| CRTAC1_Cells | BST1 | 3.08012E-24 | 0.283335 | 0.286 | 0.076 | 5.82882E-20 |
| CRTAC1_Cells | NFKBIZ | 4.15566E-24 | 0.561922 | 0.454 | 0.166 | 7.86417E-20 |
| CRTAC1_Cells | IRF7 | 4.41319E-24 | 0.409508 | 0.276 | 0.074 | 8.35152E-20 |
| CRTAC1_Cells | CASP4 | 6.89047E-24 | 0.417682 | 0.612 | 0.249 | 1.30395E-19 |
| CRTAC1_Cells | GADD45B | 7.2944E-24 | 0.900656 | 0.867 | 0.551 | 1.38039E-19 |
| CRTAC1_Cells | GDF7 | 9.31055E-24 | 0.58731 | 0.342 | 0.108 | 1.76193E-19 |
| CRTAC1_Cells | BMP4 | 1.17767E-23 | 0.417089 | 0.311 | 0.09 | 2.22862E-19 |
| CRTAC1_Cells | CCDC68 | 1.28735E-23 | 0.285227 | 0.276 | 0.073 | 2.43618E-19 |
| CRTAC1_Cells | EDEM2 | 2.67009E-23 | 0.375891 | 0.342 | 0.107 | 5.05288E-19 |
| CRTAC1_Cells | ITGA10 | 4.94364E-23 | 0.570737 | 0.724 | 0.346 | 9.35535E-19 |
| CRTAC1_Cells | SIGIRR | 5.16672E-23 | 0.394185 | 0.531 | 0.207 | 9.7775E-19 |
| CRTAC1_Cells | IL15RA | 6.63869E-23 | 0.334494 | 0.296 | 0.084 | 1.2563E-18 |
| CRTAC1_Cells | IGFBP3 | 8.00333E-23 | 1.362798 | 0.587 | 0.295 | 1.51455E-18 |
| CRTAC1_Cells | STEAP1 | 8.35674E-23 | 0.590807 | 0.413 | 0.149 | 1.58143E-18 |
| CRTAC1_Cells | PLPP3 | 8.6415E-23 | 0.689949 | 0.597 | 0.269 | 1.63532E-18 |
| CRTAC1_Cells | SGK1 | 1.48451E-22 | 0.324489 | 0.255 | 0.067 | 2.80928E-18 |
| CRTAC1_Cells | GLRX | 2.56748E-22 | 0.543501 | 0.52 | 0.214 | 4.85869E-18 |
| CRTAC1_Cells | IL13RA1 | 2.70103E-22 | 0.522555 | 0.617 | 0.275 | 5.11143E-18 |
| CRTAC1_Cells | MATN2 | 3.02345E-22 | 0.342512 | 0.306 | 0.092 | 5.72157E-18 |
| CRTAC1_Cells | IL33 | 4.87352E-22 | 0.440826 | 0.622 | 0.268 | 9.22264E-18 |
| CRTAC1_Cells | ST6GAL1 | 7.46427E-22 | 0.307146 | 0.235 | 0.061 | 1.41254E-17 |
| CRTAC1_Cells | ODF2L | 7.82344E-22 | 0.378804 | 0.464 | 0.169 | 1.48051E-17 |
| CRTAC1_Cells | NNMT | 7.93336E-22 | 0.657004 | 0.969 | 0.793 | 1.50131E-17 |
| CRTAC1_Cells | RGS3 | 9.44046E-22 | 0.876758 | 0.694 | 0.388 | 1.78651E-17 |
| CRTAC1_Cells | CPNE8 | 1.15925E-21 | 0.431712 | 0.459 | 0.173 | 2.19377E-17 |
| CRTAC1_Cells | CD58 | 1.3387E-21 | 0.53442 | 0.551 | 0.239 | 2.53336E-17 |
| CRTAC1_Cells | AKR1C2 | 1.46492E-21 | 0.671157 | 0.561 | 0.255 | 2.77221E-17 |
| CRTAC1_Cells | C1S | 1.54133E-21 | 0.528746 | 0.898 | 0.528 | 2.91681E-17 |
| CRTAC1_Cells | MYOZ2 | 1.80694E-21 | 0.572752 | 0.398 | 0.145 | 3.41946E-17 |
| CRTAC1_Cells | MYC | 2.82088E-21 | 0.341254 | 0.311 | 0.096 | 5.33823E-17 |
| CRTAC1_Cells | IFNGR2 | 3.03666E-21 | 0.504244 | 0.536 | 0.23 | 5.74657E-17 |
| CRTAC1_Cells | NPC2 | 3.12259E-21 | 0.57542 | 0.964 | 0.704 | 5.90919E-17 |
| CRTAC1_Cells | PTGIS | 3.15788E-21 | 0.692767 | 0.791 | 0.47 | 5.97596E-17 |
| CRTAC1_Cells | HSP90B1 | 4.08206E-21 | 0.608589 | 0.985 | 0.912 | 7.7249E-17 |
| CRTAC1_Cells | SERTAD1 | 5.28047E-21 | 0.57031 | 0.724 | 0.365 | 9.99277E-17 |
| CRTAC1_Cells | ERCC6 | 5.96981E-21 | 0.326255 | 0.372 | 0.128 | 1.12973E-16 |
| CRTAC1_Cells | GMPPB | 6.83769E-21 | 0.25424 | 0.291 | 0.086 | 1.29396E-16 |
| CRTAC1_Cells | MELTF | 7.19944E-21 | 0.325612 | 0.112 | 0.018 | 1.36242E-16 |
| CRTAC1_Cells | P2RY6 | 8.25536E-21 | 0.319516 | 0.194 | 0.046 | 1.56224E-16 |
| CRTAC1_Cells | KIF9 | 8.80307E-21 | 0.337099 | 0.337 | 0.111 | 1.66589E-16 |
| CRTAC1_Cells | BGN | 8.88176E-21 | 0.501821 | 1 | 0.981 | 1.68078E-16 |
| CRTAC1_Cells | GCLM | 1.07002E-20 | 0.293792 | 0.347 | 0.114 | 2.0249E-16 |
| CRTAC1_Cells | CISD1 | 1.21606E-20 | 0.454934 | 0.724 | 0.354 | 2.30128E-16 |
| CRTAC1_Cells | VAMP5 | 1.27605E-20 | 0.569049 | 0.862 | 0.556 | 2.41481E-16 |
| CRTAC1_Cells | FCGRT | 1.7385E-20 | 0.535518 | 0.837 | 0.505 | 3.28994E-16 |
| CRTAC1_Cells | IFITM1 | 1.92548E-20 | 0.388293 | 0.337 | 0.11 | 3.64377E-16 |
| CRTAC1_Cells | RAB29 | 2.05038E-20 | 0.335763 | 0.367 | 0.126 | 3.88015E-16 |
| CRTAC1_Cells | PRCP | 2.24407E-20 | 0.338943 | 0.444 | 0.162 | 4.24669E-16 |
| CRTAC1_Cells | S100A11 | 2.34911E-20 | 0.509877 | 1 | 0.971 | 4.44546E-16 |
| CRTAC1_Cells | ELK3 | 2.45321E-20 | 0.375234 | 0.393 | 0.141 | 4.64246E-16 |
| CRTAC1_Cells | KRT16 | 2.73327E-20 | 0.532703 | 0.133 | 0.024 | 5.17245E-16 |
| CRTAC1_Cells | APOL3 | 2.80591E-20 | 0.25607 | 0.27 | 0.078 | 5.3099E-16 |
| CRTAC1_Cells | TNFSF10 | 2.85654E-20 | 0.46645 | 0.383 | 0.137 | 5.40572E-16 |
| CRTAC1_Cells | ST6GALNA | 3.572E-20 | 0.328704 | 0.372 | 0.129 | 6.75965E-16 |
| CRTAC1_Cells | CDH13 | 3.98536E-20 | 0.58486 | 0.852 | 0.508 | 7.54189E-16 |
| CRTAC1_Cells | RNASE1 | 4.25054E-20 | 0.660257 | 0.158 | 0.033 | 8.04372E-16 |
| CRTAC1_Cells | KAZALD1 | 7.03422E-20 | 0.268363 | 0.321 | 0.103 | 1.33116E-15 |

| | | | | | | |
|---|---|---|---|---|---|---|
| CRTAC1_Cells | ADAM15 | 7.40565E-20 | 0.295295 | 0.276 | 0.082 | 1.40145E-15 |
| CRTAC1_Cells | CD200 | 8.48769E-20 | 0.629129 | 0.561 | 0.266 | 1.60621E-15 |
| CRTAC1_Cells | FAS | 9.98784E-20 | 0.331704 | 0.469 | 0.182 | 1.8901E-15 |
| CRTAC1_Cells | RILPL2 | 1.14253E-19 | 0.60713 | 0.765 | 0.465 | 2.16211E-15 |
| CRTAC1_Cells | PLPP5 | 1.20695E-19 | 0.426401 | 0.546 | 0.241 | 2.28403E-15 |
| CRTAC1_Cells | SLC41A3 | 1.32841E-19 | 0.413801 | 0.577 | 0.255 | 2.51388E-15 |
| CRTAC1_Cells | TAGLN2 | 1.5454E-19 | 0.549284 | 0.99 | 0.831 | 2.92452E-15 |
| CRTAC1_Cells | WWTR1 | 2.06648E-19 | 0.6425 | 0.765 | 0.451 | 3.9106E-15 |
| CRTAC1_Cells | AFAP1 | 2.62977E-19 | 0.314642 | 0.321 | 0.107 | 4.97658E-15 |
| CRTAC1_Cells | SP100 | 3.05214E-19 | 0.532534 | 0.811 | 0.473 | 5.77587E-15 |
| CRTAC1_Cells | S100A13 | 3.3013E-19 | 0.535232 | 0.923 | 0.681 | 6.24739E-15 |
| CRTAC1_Cells | SSR4 | 3.8782E-19 | 0.447256 | 0.985 | 0.85 | 7.3391E-15 |
| CRTAC1_Cells | AC020916. | 5.64063E-19 | 0.373497 | 0.495 | 0.203 | 1.06743E-14 |
| CRTAC1_Cells | IER2 | 6.22489E-19 | 0.558445 | 0.908 | 0.624 | 1.178E-14 |
| CRTAC1_Cells | SAT1 | 6.93938E-19 | 0.753932 | 0.842 | 0.561 | 1.31321E-14 |
| CRTAC1_Cells | DNAJC1 | 7.03381E-19 | 0.442594 | 0.643 | 0.311 | 1.33108E-14 |
| CRTAC1_Cells | FLNB | 7.06639E-19 | 0.285101 | 0.286 | 0.089 | 1.33724E-14 |
| CRTAC1_Cells | THBS1 | 7.80375E-19 | 0.389031 | 0.332 | 0.114 | 1.47678E-14 |
| CRTAC1_Cells | NFE2L2 | 7.88462E-19 | 0.565235 | 0.888 | 0.626 | 1.49209E-14 |
| CRTAC1_Cells | IKBIP | 7.93534E-19 | 0.422639 | 0.587 | 0.268 | 1.50168E-14 |
| CRTAC1_Cells | MIEN1 | 9.3727E-19 | 0.421599 | 0.653 | 0.326 | 1.77369E-14 |
| CRTAC1_Cells | BAMBI | 9.76651E-19 | 0.385495 | 0.337 | 0.118 | 1.84821E-14 |
| CRTAC1_Cells | NUDT5 | 1.14585E-18 | 0.319078 | 0.469 | 0.185 | 2.1684E-14 |
| CRTAC1_Cells | ATP1A1 | 1.41774E-18 | 0.502213 | 0.684 | 0.353 | 2.68293E-14 |
| CRTAC1_Cells | MSX1 | 1.49319E-18 | 0.471578 | 0.367 | 0.139 | 2.82571E-14 |
| CRTAC1_Cells | CFB | 1.71421E-18 | 0.47299 | 0.566 | 0.263 | 3.24397E-14 |
| CRTAC1_Cells | SULF2 | 1.84188E-18 | 0.345748 | 0.301 | 0.1 | 3.48558E-14 |
| CRTAC1_Cells | BTG2 | 1.90106E-18 | 0.532617 | 0.658 | 0.338 | 3.59757E-14 |
| CRTAC1_Cells | PRSS23 | 1.91821E-18 | 0.568542 | 0.862 | 0.561 | 3.63002E-14 |
| CRTAC1_Cells | BCL3 | 2.55436E-18 | 0.330575 | 0.245 | 0.073 | 4.83387E-14 |
| CRTAC1_Cells | TNFRSF6B | 2.59253E-18 | 0.365774 | 0.143 | 0.03 | 4.9061E-14 |
| CRTAC1_Cells | CTSZ | 3.91005E-18 | 0.519721 | 0.939 | 0.727 | 7.39938E-14 |
| CRTAC1_Cells | ABRACL | 4.10338E-18 | 0.345936 | 0.398 | 0.155 | 7.76523E-14 |
| CRTAC1_Cells | ROM1 | 4.11964E-18 | 0.253668 | 0.337 | 0.115 | 7.79601E-14 |
| CRTAC1_Cells | VIM | 7.48334E-18 | 0.503733 | 1 | 1 | 1.41615E-13 |
| CRTAC1_Cells | MIR22HG | 7.55139E-18 | 0.479838 | 0.561 | 0.266 | 1.42903E-13 |
| CRTAC1_Cells | PHACTR2 | 7.59422E-18 | 0.558014 | 0.735 | 0.424 | 1.43713E-13 |
| CRTAC1_Cells | PIM3 | 8.02247E-18 | 0.438569 | 0.418 | 0.171 | 1.51817E-13 |
| CRTAC1_Cells | BZW1 | 1.14389E-17 | 0.614429 | 0.852 | 0.57 | 2.1647E-13 |
| CRTAC1_Cells | RELB | 1.23122E-17 | 0.293503 | 0.25 | 0.076 | 2.32996E-13 |
| CRTAC1_Cells | CXCL2 | 1.26486E-17 | 1.041357 | 0.143 | 0.032 | 2.39361E-13 |
| CRTAC1_Cells | LTBP2 | 1.7402E-17 | 0.597656 | 0.862 | 0.54 | 3.29315E-13 |
| CRTAC1_Cells | PLOD1 | 2.03861E-17 | 0.297959 | 0.367 | 0.137 | 3.85786E-13 |
| CRTAC1_Cells | MEGF6 | 2.14152E-17 | 0.384931 | 0.449 | 0.19 | 4.05262E-13 |
| CRTAC1_Cells | CCDC80 | 2.58786E-17 | 0.711712 | 0.821 | 0.539 | 4.89727E-13 |
| CRTAC1_Cells | GRAMD1A | 2.76349E-17 | 0.360198 | 0.352 | 0.132 | 5.22963E-13 |
| CRTAC1_Cells | GDF15 | 2.99712E-17 | 0.388863 | 0.158 | 0.037 | 5.67175E-13 |
| CRTAC1_Cells | NR4A1 | 3.41642E-17 | 0.401048 | 0.577 | 0.274 | 6.46524E-13 |
| CRTAC1_Cells | GLT8D1 | 3.43836E-17 | 0.335882 | 0.526 | 0.228 | 6.50675E-13 |
| CRTAC1_Cells | MARCH2 | 3.85394E-17 | 0.474652 | 0.648 | 0.347 | 7.29319E-13 |
| CRTAC1_Cells | ORMDL2 | 4.26199E-17 | 0.313217 | 0.515 | 0.222 | 8.06539E-13 |
| CRTAC1_Cells | SLC25A37 | 4.43479E-17 | 0.412825 | 0.536 | 0.245 | 8.3924E-13 |
| CRTAC1_Cells | PTGDS | 5.83612E-17 | 0.729396 | 0.209 | 0.06 | 1.10443E-12 |
| CRTAC1_Cells | PCGF2 | 6.6056E-17 | 0.331331 | 0.378 | 0.146 | 1.25004E-12 |
| CRTAC1_Cells | TMEM37 | 7.0635E-17 | 0.282782 | 0.153 | 0.036 | 1.3367E-12 |
| CRTAC1_Cells | NME1 | 8.35685E-17 | 0.497726 | 0.638 | 0.33 | 1.58145E-12 |
| CRTAC1_Cells | H3F3B | 9.08133E-17 | 0.400959 | 0.995 | 0.986 | 1.71855E-12 |
| CRTAC1_Cells | CLMP | 9.38941E-17 | 0.412364 | 0.362 | 0.145 | 1.77685E-12 |

| | | | | | | |
|---|---|---|---|---|---|---|
| CRTAC1_Cells | PCDH10 | 1.00453E-16 | 0.859913 | 0.5 | 0.245 | 1.90098E-12 |
| CRTAC1_Cells | MTRF1L | 1.17692E-16 | 0.296843 | 0.464 | 0.196 | 2.2272E-12 |
| CRTAC1_Cells | PKN1 | 1.30872E-16 | 0.393689 | 0.5 | 0.222 | 2.47661E-12 |
| CRTAC1_Cells | MYDGF | 1.62983E-16 | 0.527448 | 0.888 | 0.638 | 3.08429E-12 |
| CRTAC1_Cells | CD99 | 1.67974E-16 | 0.445013 | 0.985 | 0.927 | 3.17875E-12 |
| CRTAC1_Cells | IFI44L | 1.86391E-16 | 0.364254 | 0.25 | 0.081 | 3.52726E-12 |
| CRTAC1_Cells | MGAT4B | 1.87305E-16 | 0.364374 | 0.464 | 0.202 | 3.54455E-12 |
| CRTAC1_Cells | SLC35B1 | 2.0753E-16 | 0.262204 | 0.449 | 0.183 | 3.92729E-12 |
| CRTAC1_Cells | SRPRB | 2.52558E-16 | 0.323822 | 0.429 | 0.175 | 4.7794E-12 |
| CRTAC1_Cells | KDELR2 | 2.82699E-16 | 0.531581 | 0.857 | 0.587 | 5.34979E-12 |
| CRTAC1_Cells | FUCA2 | 2.86315E-16 | 0.278039 | 0.464 | 0.194 | 5.41822E-12 |
| CRTAC1_Cells | IFNAR2 | 3.09251E-16 | 0.330075 | 0.398 | 0.16 | 5.85226E-12 |
| CRTAC1_Cells | LY96 | 3.73062E-16 | 0.284588 | 0.311 | 0.113 | 7.05982E-12 |
| CRTAC1_Cells | TSPAN4 | 5.21464E-16 | 0.42732 | 0.689 | 0.368 | 9.86818E-12 |
| CRTAC1_Cells | C2orf27A | 5.41621E-16 | 0.268941 | 0.235 | 0.073 | 1.02496E-11 |
| CRTAC1_Cells | PLTP | 6.3278E-16 | 0.420712 | 0.469 | 0.206 | 1.19747E-11 |
| CRTAC1_Cells | HLA-B | 6.49424E-16 | 0.384057 | 0.98 | 0.766 | 1.22897E-11 |
| CRTAC1_Cells | PTPRE | 6.80934E-16 | 0.340509 | 0.199 | 0.057 | 1.2886E-11 |
| CRTAC1_Cells | HDGFL3 | 7.00247E-16 | 0.465921 | 0.699 | 0.386 | 1.32515E-11 |
| CRTAC1_Cells | MAP3K8 | 7.14323E-16 | 0.378093 | 0.434 | 0.187 | 1.35178E-11 |
| CRTAC1_Cells | CSRNP1 | 7.44407E-16 | 0.283974 | 0.316 | 0.114 | 1.40872E-11 |
| CRTAC1_Cells | RGS10 | 9.01371E-16 | 0.312668 | 0.49 | 0.217 | 1.70575E-11 |
| CRTAC1_Cells | CDC42EP5 | 1.05024E-15 | 0.43579 | 0.694 | 0.383 | 1.98747E-11 |
| CRTAC1_Cells | SHC1 | 1.29777E-15 | 0.34171 | 0.408 | 0.168 | 2.45589E-11 |
| CRTAC1_Cells | CCN1 | 1.40045E-15 | 0.785223 | 0.898 | 0.753 | 2.65021E-11 |
| CRTAC1_Cells | DNAJB9 | 1.53322E-15 | 0.475891 | 0.48 | 0.224 | 2.90147E-11 |
| CRTAC1_Cells | HM13 | 1.99481E-15 | 0.468992 | 0.714 | 0.417 | 3.77498E-11 |
| CRTAC1_Cells | GPX8 | 2.09679E-15 | 0.439638 | 0.653 | 0.347 | 3.96796E-11 |
| CRTAC1_Cells | VCAN | 2.20431E-15 | 0.594507 | 0.964 | 0.786 | 4.17143E-11 |
| CRTAC1_Cells | MRPL17 | 2.51559E-15 | 0.353398 | 0.602 | 0.291 | 4.76051E-11 |
| CRTAC1_Cells | ST3GAL1 | 2.89338E-15 | 0.262911 | 0.281 | 0.097 | 5.47543E-11 |
| CRTAC1_Cells | BSCL2 | 3.1556E-15 | 0.252661 | 0.485 | 0.207 | 5.97167E-11 |
| CRTAC1_Cells | BAG3 | 3.21482E-15 | 0.408754 | 0.464 | 0.208 | 6.08373E-11 |
| CRTAC1_Cells | SELENOS | 3.71918E-15 | 0.488535 | 0.913 | 0.664 | 7.03818E-11 |
| CRTAC1_Cells | CEBPD | 3.84681E-15 | 0.484701 | 0.888 | 0.589 | 7.2797E-11 |
| CRTAC1_Cells | MFAP2 | 3.88018E-15 | 0.379807 | 0.413 | 0.176 | 7.34285E-11 |
| CRTAC1_Cells | EPSTI1 | 4.33963E-15 | 0.319829 | 0.158 | 0.041 | 8.21232E-11 |
| CRTAC1_Cells | TMEM173 | 4.38744E-15 | 0.468197 | 0.612 | 0.326 | 8.3028E-11 |
| CRTAC1_Cells | FLOT1 | 4.6209E-15 | 0.412673 | 0.724 | 0.41 | 8.74458E-11 |
| CRTAC1_Cells | TSPAN9 | 5.15831E-15 | 0.337939 | 0.418 | 0.182 | 9.76159E-11 |
| CRTAC1_Cells | MEDAG | 6.11324E-15 | 0.313437 | 0.423 | 0.183 | 1.15687E-10 |
| CRTAC1_Cells | TNIP2 | 7.90041E-15 | 0.292038 | 0.423 | 0.182 | 1.49507E-10 |
| CRTAC1_Cells | PLK3 | 9.16716E-15 | 0.323607 | 0.367 | 0.148 | 1.73479E-10 |
| CRTAC1_Cells | BST2 | 1.17179E-14 | 0.621005 | 0.75 | 0.438 | 2.2175E-10 |
| CRTAC1_Cells | SULF1 | 1.20181E-14 | 0.577352 | 0.786 | 0.524 | 2.27431E-10 |
| CRTAC1_Cells | SPSB1 | 1.30894E-14 | 0.355982 | 0.342 | 0.136 | 2.47705E-10 |
| CRTAC1_Cells | IGFBP6 | 1.39447E-14 | 0.787969 | 0.903 | 0.786 | 2.63889E-10 |
| CRTAC1_Cells | AK1 | 1.6692E-14 | 0.386057 | 0.821 | 0.531 | 3.15879E-10 |
| CRTAC1_Cells | S100A16 | 1.66934E-14 | 0.451237 | 0.776 | 0.468 | 3.15906E-10 |
| CRTAC1_Cells | AGTRAP | 1.70384E-14 | 0.322879 | 0.474 | 0.217 | 3.22435E-10 |
| CRTAC1_Cells | CD9 | 2.20285E-14 | 0.474303 | 0.995 | 0.969 | 4.16867E-10 |
| CRTAC1_Cells | SH3BP5 | 2.25517E-14 | 0.418964 | 0.464 | 0.211 | 4.26768E-10 |
| CRTAC1_Cells | EMP3 | 2.46439E-14 | 0.612181 | 0.929 | 0.759 | 4.66362E-10 |
| CRTAC1_Cells | TOR1AIP2 | 2.53442E-14 | 0.262297 | 0.51 | 0.23 | 4.79613E-10 |
| CRTAC1_Cells | RHOJ | 2.92537E-14 | 0.306269 | 0.265 | 0.095 | 5.53597E-10 |
| CRTAC1_Cells | SEC13 | 2.9555E-14 | 0.337125 | 0.515 | 0.241 | 5.59299E-10 |
| CRTAC1_Cells | NDUFA4L2 | 3.40667E-14 | 0.260338 | 0.699 | 0.376 | 6.44678E-10 |
| CRTAC1_Cells | RCN3 | 3.9462E-14 | 0.351272 | 0.694 | 0.367 | 7.46779E-10 |

| | | | | | | |
|---|---|---|---|---|---|---|
| CRTAC1_Cells | B4GALT1 | 3.96277E-14 | 0.422011 | 0.444 | 0.207 | 7.49915E-10 |
| CRTAC1_Cells | LAGE3 | 3.96336E-14 | 0.26159 | 0.362 | 0.147 | 7.50026E-10 |
| CRTAC1_Cells | PPA1 | 4.28492E-14 | 0.444208 | 0.847 | 0.555 | 8.10878E-10 |
| CRTAC1_Cells | SSR3 | 4.89942E-14 | 0.452519 | 0.878 | 0.614 | 9.27167E-10 |
| CRTAC1_Cells | LIMA1 | 4.94728E-14 | 0.407658 | 0.816 | 0.509 | 9.36223E-10 |
| CRTAC1_Cells | KIFC3 | 5.56626E-14 | 0.291277 | 0.255 | 0.089 | 1.05336E-09 |
| CRTAC1_Cells | SEC61B | 5.61124E-14 | 0.422235 | 0.98 | 0.859 | 1.06187E-09 |
| CRTAC1_Cells | SNCG | 6.15833E-14 | 0.498369 | 0.75 | 0.448 | 1.1654E-09 |
| CRTAC1_Cells | FTH1 | 6.57927E-14 | 0.555022 | 1 | 0.999 | 1.24506E-09 |
| CRTAC1_Cells | SHMT2 | 8.17965E-14 | 0.303041 | 0.316 | 0.123 | 1.54792E-09 |
| CRTAC1_Cells | IL6ST | 8.42601E-14 | 0.529786 | 0.821 | 0.55 | 1.59454E-09 |
| CRTAC1_Cells | CALU | 8.58021E-14 | 0.475984 | 0.867 | 0.601 | 1.62372E-09 |
| CRTAC1_Cells | MAPK3 | 9.01184E-14 | 0.342044 | 0.526 | 0.251 | 1.7054E-09 |
| CRTAC1_Cells | ASPN | 9.81738E-14 | 0.711655 | 0.837 | 0.601 | 1.85784E-09 |
| CRTAC1_Cells | LUZP1 | 9.83203E-14 | 0.422183 | 0.556 | 0.29 | 1.86061E-09 |
| CRTAC1_Cells | TLE5 | 1.13942E-13 | 0.419465 | 0.918 | 0.659 | 2.15624E-09 |
| CRTAC1_Cells | FAM174C | 1.20537E-13 | 0.303151 | 0.515 | 0.244 | 2.28104E-09 |
| CRTAC1_Cells | ETS2 | 1.41289E-13 | 0.428566 | 0.663 | 0.364 | 2.67375E-09 |
| CRTAC1_Cells | MTHFD1L | 1.41706E-13 | 0.296674 | 0.378 | 0.162 | 2.68165E-09 |
| CRTAC1_Cells | NES | 1.47573E-13 | 0.323534 | 0.26 | 0.094 | 2.79267E-09 |
| CRTAC1_Cells | PPIC | 1.58817E-13 | 0.436222 | 0.862 | 0.616 | 3.00546E-09 |
| CRTAC1_Cells | FKBP1A | 1.62568E-13 | 0.588862 | 0.857 | 0.671 | 3.07644E-09 |
| CRTAC1_Cells | CIB1 | 2.01371E-13 | 0.425525 | 0.852 | 0.577 | 3.81075E-09 |
| CRTAC1_Cells | SOD2 | 2.6183E-13 | 0.747702 | 0.684 | 0.419 | 4.95488E-09 |
| CRTAC1_Cells | GNL3 | 2.7422E-13 | 0.359739 | 0.429 | 0.198 | 5.18935E-09 |
| CRTAC1_Cells | GNG5 | 3.51321E-13 | 0.426325 | 0.939 | 0.837 | 6.64839E-09 |
| CRTAC1_Cells | DDIT4 | 4.05566E-13 | 0.384324 | 0.372 | 0.161 | 7.67493E-09 |
| CRTAC1_Cells | IFIT3 | 4.5033E-13 | 0.678684 | 0.27 | 0.105 | 8.52205E-09 |
| CRTAC1_Cells | IFT22 | 5.00856E-13 | 0.251941 | 0.485 | 0.225 | 9.47819E-09 |
| CRTAC1_Cells | GABARAPL | 5.35793E-13 | 0.425051 | 0.974 | 0.854 | 1.01394E-08 |
| CRTAC1_Cells | EXT1 | 5.88392E-13 | 0.366477 | 0.52 | 0.264 | 1.11347E-08 |
| CRTAC1_Cells | TAF10 | 5.9899E-13 | 0.374297 | 0.673 | 0.391 | 1.13353E-08 |
| CRTAC1_Cells | SPTBN1 | 6.10932E-13 | 0.413755 | 0.888 | 0.62 | 1.15613E-08 |
| CRTAC1_Cells | CMTM6 | 6.45709E-13 | 0.292644 | 0.49 | 0.24 | 1.22194E-08 |
| CRTAC1_Cells | OAZ1 | 8.04334E-13 | 0.353655 | 0.974 | 0.946 | 1.52212E-08 |
| CRTAC1_Cells | CFLAR | 8.17394E-13 | 0.312998 | 0.551 | 0.279 | 1.54684E-08 |
| CRTAC1_Cells | OCIAD2 | 9.20357E-13 | 0.459166 | 0.459 | 0.236 | 1.74168E-08 |
| CRTAC1_Cells | SMC4 | 1.04988E-12 | 0.276272 | 0.413 | 0.188 | 1.98679E-08 |
| CRTAC1_Cells | MDK | 1.24648E-12 | 0.447377 | 0.362 | 0.157 | 2.35884E-08 |
| CRTAC1_Cells | BHLHE40 | 1.29564E-12 | 0.270176 | 0.286 | 0.112 | 2.45188E-08 |
| CRTAC1_Cells | TMEM120 | 1.41541E-12 | 0.30708 | 0.571 | 0.291 | 2.67853E-08 |
| CRTAC1_Cells | GPR108 | 1.49467E-12 | 0.261775 | 0.474 | 0.224 | 2.82852E-08 |
| CRTAC1_Cells | TGFBR2 | 1.54513E-12 | 0.426057 | 0.648 | 0.377 | 2.924E-08 |
| CRTAC1_Cells | OAS1 | 1.63114E-12 | 0.303564 | 0.143 | 0.039 | 3.08677E-08 |
| CRTAC1_Cells | ETHE1 | 1.87201E-12 | 0.303637 | 0.602 | 0.311 | 3.54259E-08 |
| CRTAC1_Cells | ARRDC2 | 1.92633E-12 | 0.301968 | 0.265 | 0.102 | 3.64538E-08 |
| CRTAC1_Cells | DEPP1 | 2.38139E-12 | 0.61127 | 0.51 | 0.271 | 4.50654E-08 |
| CRTAC1_Cells | PTHLH | 2.4043E-12 | 0.327242 | 0.219 | 0.077 | 4.5499E-08 |
| CRTAC1_Cells | PLOD2 | 2.49391E-12 | 0.382723 | 0.439 | 0.209 | 4.71948E-08 |
| CRTAC1_Cells | POLR2L | 2.6484E-12 | 0.441845 | 0.949 | 0.774 | 5.01182E-08 |
| CRTAC1_Cells | CKAP4 | 2.8662E-12 | 0.331389 | 0.469 | 0.237 | 5.424E-08 |
| CRTAC1_Cells | RPN2 | 3.05601E-12 | 0.367466 | 0.827 | 0.521 | 5.7832E-08 |
| CRTAC1_Cells | SKIL | 3.09175E-12 | 0.4058 | 0.607 | 0.335 | 5.85082E-08 |
| CRTAC1_Cells | EIF4A1 | 3.57544E-12 | 0.464571 | 0.964 | 0.829 | 6.76615E-08 |
| CRTAC1_Cells | CDK2AP1 | 3.74006E-12 | 0.418498 | 0.755 | 0.481 | 7.07769E-08 |
| CRTAC1_Cells | MESD | 4.28467E-12 | 0.412873 | 0.679 | 0.415 | 8.10831E-08 |
| CRTAC1_Cells | CCDC85B | 5.04433E-12 | 0.478538 | 0.918 | 0.768 | 9.5459E-08 |
| CRTAC1_Cells | FAM92A | 6.24335E-12 | 0.300809 | 0.434 | 0.208 | 1.18149E-07 |

| | | | | | | |
|---|---|---|---|---|---|---|
| CRTAC1_Cells | PSMB8 | 6.69881E-12 | 0.325648 | 0.653 | 0.356 | 1.26768E-07 |
| CRTAC1_Cells | TMBIM1 | 1.05218E-11 | 0.383363 | 0.755 | 0.449 | 1.99114E-07 |
| CRTAC1_Cells | MFAP5 | 1.23802E-11 | 0.472886 | 0.138 | 0.039 | 2.34283E-07 |
| CRTAC1_Cells | FAM114A1 | 1.26069E-11 | 0.385362 | 0.791 | 0.517 | 2.38574E-07 |
| CRTAC1_Cells | DPP7 | 1.32473E-11 | 0.314513 | 0.796 | 0.492 | 2.50691E-07 |
| CRTAC1_Cells | DAP | 1.3513E-11 | 0.349844 | 0.679 | 0.387 | 2.5572E-07 |
| CRTAC1_Cells | ARL2BP | 1.44658E-11 | 0.326103 | 0.628 | 0.361 | 2.73752E-07 |
| CRTAC1_Cells | EIF1AY | 1.66739E-11 | 0.251085 | 0.515 | 0.254 | 3.15537E-07 |
| CRTAC1_Cells | FAM177A1 | 1.692E-11 | 0.419764 | 0.847 | 0.583 | 3.20195E-07 |
| CRTAC1_Cells | ZFP36 | 1.79631E-11 | 0.341654 | 0.816 | 0.588 | 3.39933E-07 |
| CRTAC1_Cells | TMEM167 | 1.81806E-11 | 0.346238 | 0.704 | 0.425 | 3.44049E-07 |
| CRTAC1_Cells | SPHK1 | 1.83821E-11 | 0.270951 | 0.179 | 0.059 | 3.47863E-07 |
| CRTAC1_Cells | YIPF3 | 2.00213E-11 | 0.355589 | 0.73 | 0.451 | 3.78883E-07 |
| CRTAC1_Cells | DDX21 | 2.11221E-11 | 0.391755 | 0.704 | 0.428 | 3.99715E-07 |
| CRTAC1_Cells | PDIA3 | 2.32672E-11 | 0.419808 | 0.939 | 0.791 | 4.40308E-07 |
| CRTAC1_Cells | XBP1 | 2.33592E-11 | 0.335039 | 0.735 | 0.444 | 4.42049E-07 |
| CRTAC1_Cells | CTSB | 2.38789E-11 | 0.380961 | 0.719 | 0.45 | 4.51884E-07 |
| CRTAC1_Cells | PPIB | 2.64541E-11 | 0.346671 | 0.98 | 0.917 | 5.00617E-07 |
| CRTAC1_Cells | MT1E | 2.98847E-11 | 0.490446 | 0.781 | 0.511 | 5.65537E-07 |
| CRTAC1_Cells | CST6 | 3.02186E-11 | 0.278173 | 0.398 | 0.191 | 5.71856E-07 |
| CRTAC1_Cells | SURF4 | 3.19009E-11 | 0.305669 | 0.571 | 0.298 | 6.03692E-07 |
| CRTAC1_Cells | PAPSS1 | 3.23858E-11 | 0.265983 | 0.434 | 0.213 | 6.1287E-07 |
| CRTAC1_Cells | INSIG2 | 3.46803E-11 | 0.268148 | 0.48 | 0.238 | 6.5629E-07 |
| CRTAC1_Cells | IRF1 | 3.4941E-11 | 0.380746 | 0.367 | 0.171 | 6.61223E-07 |
| CRTAC1_Cells | FAM107B | 4.0439E-11 | 0.306431 | 0.515 | 0.267 | 7.65269E-07 |
| CRTAC1_Cells | PLPP1 | 5.04839E-11 | 0.375425 | 0.765 | 0.495 | 9.55357E-07 |
| CRTAC1_Cells | IFI6 | 5.49746E-11 | 0.775446 | 0.617 | 0.4 | 1.04034E-06 |
| CRTAC1_Cells | GSDMD | 6.22875E-11 | 0.275791 | 0.51 | 0.264 | 1.17873E-06 |
| CRTAC1_Cells | ARPC3 | 6.40023E-11 | 0.334008 | 0.954 | 0.836 | 1.21118E-06 |
| CRTAC1_Cells | TMBIM4 | 6.54332E-11 | 0.367814 | 0.959 | 0.781 | 1.23826E-06 |
| CRTAC1_Cells | LDHA | 7.20002E-11 | 0.366859 | 0.974 | 0.892 | 1.36253E-06 |
| CRTAC1_Cells | CTNNAL1 | 7.49262E-11 | 0.383425 | 0.531 | 0.285 | 1.4179E-06 |
| CRTAC1_Cells | LINC02185 | 7.62089E-11 | 0.280047 | 0.133 | 0.039 | 1.44218E-06 |
| CRTAC1_Cells | CCNY | 7.79645E-11 | 0.265707 | 0.454 | 0.225 | 1.4754E-06 |
| CRTAC1_Cells | TM9SF1 | 9.62132E-11 | 0.29829 | 0.515 | 0.268 | 1.82074E-06 |
| CRTAC1_Cells | KLF3 | 1.0291E-10 | 0.331213 | 0.551 | 0.301 | 1.94746E-06 |
| CRTAC1_Cells | SRM | 1.15242E-10 | 0.357723 | 0.724 | 0.463 | 2.18084E-06 |
| CRTAC1_Cells | FOS | 1.15366E-10 | 0.321616 | 0.913 | 0.704 | 2.18318E-06 |
| CRTAC1_Cells | PGM2L1 | 1.17688E-10 | 0.339586 | 0.398 | 0.195 | 2.22713E-06 |
| CRTAC1_Cells | GPCPD1 | 1.33889E-10 | 0.290066 | 0.321 | 0.148 | 2.53371E-06 |
| CRTAC1_Cells | JTB | 1.44282E-10 | 0.34557 | 0.852 | 0.603 | 2.73039E-06 |
| CRTAC1_Cells | PPP1CA | 1.62612E-10 | 0.295363 | 0.464 | 0.246 | 3.07726E-06 |
| CRTAC1_Cells | HIF1A | 1.6648E-10 | 0.488516 | 0.546 | 0.313 | 3.15047E-06 |
| CRTAC1_Cells | P4HB | 1.84033E-10 | 0.418229 | 0.872 | 0.677 | 3.48265E-06 |
| CRTAC1_Cells | CRIM1 | 1.90028E-10 | 0.415073 | 0.888 | 0.583 | 3.59609E-06 |
| CRTAC1_Cells | MT1M | 1.98846E-10 | 0.439665 | 0.643 | 0.378 | 3.76296E-06 |
| CRTAC1_Cells | SERPINB6 | 2.40512E-10 | 0.308138 | 0.832 | 0.548 | 4.55145E-06 |
| CRTAC1_Cells | XAF1 | 2.63277E-10 | 0.267568 | 0.265 | 0.112 | 4.98226E-06 |
| CRTAC1_Cells | YIPF5 | 2.72072E-10 | 0.310384 | 0.587 | 0.337 | 5.1487E-06 |
| CRTAC1_Cells | OGN | 2.84371E-10 | 0.468103 | 0.934 | 0.866 | 5.38144E-06 |
| CRTAC1_Cells | CHST12 | 2.98097E-10 | 0.251679 | 0.378 | 0.178 | 5.64119E-06 |
| CRTAC1_Cells | LY6E | 2.98166E-10 | 0.444548 | 0.724 | 0.45 | 5.64248E-06 |
| CRTAC1_Cells | BLOC1S2 | 3.68542E-10 | 0.251676 | 0.408 | 0.201 | 6.97428E-06 |
| CRTAC1_Cells | CCPG1 | 3.78358E-10 | 0.388504 | 0.878 | 0.689 | 7.16004E-06 |
| CRTAC1_Cells | POLR2H | 3.79516E-10 | 0.293802 | 0.469 | 0.243 | 7.18197E-06 |
| CRTAC1_Cells | AUP1 | 4.3804E-10 | 0.29378 | 0.536 | 0.286 | 8.28946E-06 |
| CRTAC1_Cells | ACKR4 | 4.4375E-10 | 0.277557 | 0.296 | 0.131 | 8.39752E-06 |
| CRTAC1_Cells | SMIM14 | 5.76679E-10 | 0.331781 | 0.776 | 0.492 | 1.09131E-05 |

| | | | | | | |
|---|---|---|---|---|---|---|
| CRTAC1_Cells | DUXAP8 | 9.30184E-10 | 0.347208 | 0.209 | 0.082 | 1.76028E-05 |
| CRTAC1_Cells | PLAAT3 | 9.34429E-10 | 0.390999 | 0.714 | 0.467 | 1.76831E-05 |
| CRTAC1_Cells | POLD4 | 9.58965E-10 | 0.320079 | 0.801 | 0.513 | 1.81475E-05 |
| CRTAC1_Cells | C1GALT1C | 9.96072E-10 | 0.264227 | 0.474 | 0.256 | 1.88497E-05 |
| CRTAC1_Cells | RAB13 | 1.16251E-09 | 0.358477 | 0.878 | 0.709 | 2.19994E-05 |
| CRTAC1_Cells | UFD1 | 1.31267E-09 | 0.272273 | 0.714 | 0.431 | 2.4841E-05 |
| CRTAC1_Cells | EPRS | 1.31993E-09 | 0.342797 | 0.566 | 0.326 | 2.49783E-05 |
| CRTAC1_Cells | CNIH1 | 1.35936E-09 | 0.268774 | 0.633 | 0.355 | 2.57245E-05 |
| CRTAC1_Cells | CYB5R3 | 1.4995E-09 | 0.351895 | 0.974 | 0.873 | 2.83766E-05 |
| CRTAC1_Cells | CPXM2 | 1.50792E-09 | 0.2541 | 0.633 | 0.363 | 2.85358E-05 |
| CRTAC1_Cells | ATP6V0E1 | 1.87042E-09 | 0.308692 | 0.949 | 0.825 | 3.53958E-05 |
| CRTAC1_Cells | POMP | 2.08043E-09 | 0.306907 | 0.939 | 0.759 | 3.93701E-05 |
| CRTAC1_Cells | LMAN1 | 2.15476E-09 | 0.294484 | 0.801 | 0.53 | 4.07767E-05 |
| CRTAC1_Cells | ARF4 | 2.20611E-09 | 0.354861 | 0.888 | 0.662 | 4.17484E-05 |
| CRTAC1_Cells | TXNDC12 | 2.37693E-09 | 0.275194 | 0.541 | 0.307 | 4.49809E-05 |
| CRTAC1_Cells | SLC39A7 | 2.4121E-09 | 0.262872 | 0.546 | 0.3 | 4.56466E-05 |
| CRTAC1_Cells | FADS3 | 2.51568E-09 | 0.333704 | 0.719 | 0.454 | 4.76067E-05 |
| CRTAC1_Cells | KLF10 | 2.55969E-09 | 0.339116 | 0.378 | 0.192 | 4.84396E-05 |
| CRTAC1_Cells | RCAN1 | 2.72609E-09 | 0.323199 | 0.362 | 0.184 | 5.15885E-05 |
| CRTAC1_Cells | FAM89B | 3.0201E-09 | 0.365218 | 0.816 | 0.574 | 5.71523E-05 |
| CRTAC1_Cells | HIGD1A | 3.09835E-09 | 0.418548 | 0.653 | 0.435 | 5.86331E-05 |
| CRTAC1_Cells | AAMDC | 3.33024E-09 | 0.346161 | 0.75 | 0.497 | 6.30215E-05 |
| CRTAC1_Cells | REXO2 | 3.36304E-09 | 0.340864 | 0.862 | 0.637 | 6.36421E-05 |
| CRTAC1_Cells | TFG | 4.01792E-09 | 0.276127 | 0.684 | 0.409 | 7.60352E-05 |
| CRTAC1_Cells | COPZ2 | 4.14933E-09 | 0.369861 | 0.832 | 0.591 | 7.8522E-05 |
| CRTAC1_Cells | ARPC2 | 4.55362E-09 | 0.345892 | 0.959 | 0.862 | 8.61727E-05 |
| CRTAC1_Cells | GNAQ | 4.77413E-09 | 0.25684 | 0.485 | 0.259 | 9.03456E-05 |
| CRTAC1_Cells | HSPG2 | 4.88853E-09 | 0.410745 | 0.694 | 0.447 | 9.25106E-05 |
| CRTAC1_Cells | HRCT1 | 5.76295E-09 | 0.376819 | 0.403 | 0.219 | 0.000109058 |
| CRTAC1_Cells | RND3 | 6.2784E-09 | 0.30594 | 0.52 | 0.3 | 0.000118812 |
| CRTAC1_Cells | NPLOC4 | 6.29236E-09 | 0.251673 | 0.27 | 0.122 | 0.000119077 |
| CRTAC1_Cells | OS9 | 6.87473E-09 | 0.265956 | 0.73 | 0.465 | 0.000130097 |
| CRTAC1_Cells | PDIA6 | 7.65916E-09 | 0.315232 | 0.832 | 0.631 | 0.000144942 |
| CRTAC1_Cells | NDUFAF3 | 8.9131E-09 | 0.309973 | 0.878 | 0.634 | 0.000168671 |
| CRTAC1_Cells | SOX4 | 9.87981E-09 | 0.327252 | 0.546 | 0.318 | 0.000186965 |
| CRTAC1_Cells | PSME2 | 1.00636E-08 | 0.255923 | 0.786 | 0.503 | 0.000190444 |
| CRTAC1_Cells | ARPC5L | 1.08604E-08 | 0.324304 | 0.49 | 0.279 | 0.000205523 |
| CRTAC1_Cells | NEAT1 | 1.11393E-08 | 0.499621 | 0.908 | 0.792 | 0.0002108 |
| CRTAC1_Cells | SERP1 | 1.24534E-08 | 0.331845 | 0.827 | 0.601 | 0.000235669 |
| CRTAC1_Cells | TMED10 | 1.32295E-08 | 0.34616 | 0.827 | 0.635 | 0.000250355 |
| CRTAC1_Cells | ABI3BP | 1.76822E-08 | 0.304493 | 0.449 | 0.247 | 0.000334618 |
| CRTAC1_Cells | PPP1R14B | 2.09172E-08 | 0.314022 | 0.888 | 0.673 | 0.000395838 |
| CRTAC1_Cells | TFPI | 2.54016E-08 | 0.254015 | 0.724 | 0.461 | 0.0004807 |
| CRTAC1_Cells | IFIT1 | 2.59663E-08 | 0.55929 | 0.27 | 0.129 | 0.000491386 |
| CRTAC1_Cells | DIXDC1 | 2.82546E-08 | 0.329555 | 0.643 | 0.403 | 0.00053469 |
| CRTAC1_Cells | SELENOF | 3.34387E-08 | 0.256226 | 0.781 | 0.529 | 0.000632793 |
| CRTAC1_Cells | GSTO1 | 3.54766E-08 | 0.39419 | 0.893 | 0.722 | 0.000671359 |
| CRTAC1_Cells | ANGPTL4 | 3.65063E-08 | 0.360656 | 0.163 | 0.062 | 0.000690845 |
| CRTAC1_Cells | METRNL | 4.17346E-08 | 0.251646 | 0.551 | 0.324 | 0.000789786 |
| CRTAC1_Cells | SNHG7 | 4.32619E-08 | 0.271419 | 0.724 | 0.455 | 0.000818687 |
| CRTAC1_Cells | FABP5 | 4.66321E-08 | 0.295264 | 0.158 | 0.06 | 0.000882467 |
| CRTAC1_Cells | HSPA1B | 4.97042E-08 | 0.321189 | 0.663 | 0.435 | 0.000940603 |
| CRTAC1_Cells | DNAJC3 | 5.22141E-08 | 0.282659 | 0.735 | 0.497 | 0.0009881 |
| CRTAC1_Cells | RHOC | 5.23924E-08 | 0.338835 | 0.944 | 0.826 | 0.000991475 |
| CRTAC1_Cells | MT2A | 5.88793E-08 | 0.819163 | 0.888 | 0.757 | 0.001114231 |
| CRTAC1_Cells | LRRC17 | 6.14024E-08 | 0.250847 | 0.571 | 0.342 | 0.001161978 |
| CRTAC1_Cells | SDF2L1 | 6.56515E-08 | 0.259177 | 0.577 | 0.347 | 0.001242388 |
| CRTAC1_Cells | KDELR1 | 7.95446E-08 | 0.293713 | 0.908 | 0.721 | 0.001505302 |

| | | | | | | |
|---|---|---|---|---|---|---|
| CRTAC1_Cells | ISG15 | 8.41791E-08 | 0.443056 | 0.474 | 0.283 | 0.001593005 |
| CRTAC1_Cells | SERPINH1 | 1.21603E-07 | 0.30556 | 0.617 | 0.402 | 0.002301221 |
| CRTAC1_Cells | SLC25A6 | 1.22528E-07 | 0.285271 | 0.959 | 0.859 | 0.002318719 |
| CRTAC1_Cells | PRELP | 1.26464E-07 | 0.284 | 0.918 | 0.747 | 0.002393213 |
| CRTAC1_Cells | UGP2 | 1.28345E-07 | 0.378982 | 0.76 | 0.567 | 0.002428807 |
| CRTAC1_Cells | OSTC | 2.78233E-07 | 0.279179 | 0.867 | 0.634 | 0.005265277 |
| CRTAC1_Cells | MYL12A | 3.49076E-07 | 0.336925 | 0.995 | 0.943 | 0.00660591 |
| CRTAC1_Cells | LMAN2 | 5.15161E-07 | 0.269494 | 0.76 | 0.515 | 0.009748909 |
| CRTAC1_Cells | DNAJB1 | 1.48087E-06 | 0.353064 | 0.505 | 0.327 | 0.028024042 |
| CRTAC1_Cells | C9orf16 | 1.69634E-06 | 0.26022 | 0.893 | 0.722 | 0.03210147 |
| CRTAC1_Cells | TSPO | 2.47405E-06 | 0.278888 | 0.944 | 0.885 | 0.046818986 |
| CRTAC1_Cells | BMPR2 | 2.72366E-06 | 0.313241 | 0.531 | 0.336 | 0.05154245 |
| CRTAC1_Cells | MX1 | 2.76146E-06 | 0.47946 | 0.311 | 0.174 | 0.052257805 |
| CRTAC1_Cells | GPX3 | 2.84337E-06 | 0.317611 | 0.592 | 0.374 | 0.053807907 |
| CRTAC1_Cells | CCND1 | 3.54707E-06 | 0.68682 | 0.597 | 0.439 | 0.067124709 |
| CRTAC1_Cells | HSPA5 | 4.71214E-06 | 0.402803 | 0.857 | 0.689 | 0.089172568 |
| CRTAC1_Cells | SLC25A5 | 5.08921E-06 | 0.284039 | 0.816 | 0.618 | 0.096308269 |
| CRTAC1_Cells | SNTB2 | 6.81562E-06 | 0.35799 | 0.367 | 0.224 | 0.128978818 |
| CRTAC1_Cells | TGFB1 | 8.80653E-06 | 0.274042 | 0.587 | 0.4 | 0.166654787 |
| CRTAC1_Cells | TFPI2 | 1.05306E-05 | 0.843526 | 0.296 | 0.184 | 0.199281192 |
| CRTAC1_Cells | ARID5B | 2.71704E-05 | 0.327922 | 0.842 | 0.76 | 0.51417191 |
| CRTAC1_Cells | HERPUD1 | 5.90767E-05 | 0.290087 | 0.76 | 0.584 | 1 |
| CRTAC1_Cells | INHBA | 7.82795E-05 | 0.319 | 0.643 | 0.477 | 1 |
| CRTAC1_Cells | CLU | 0.000140721 | 0.545061 | 0.929 | 0.875 | 1 |
| CRTAC1_Cells | AAK1 | 0.000153561 | 0.311485 | 0.474 | 0.326 | 1 |
| CRTAC1_Cells | INMT | 0.000166394 | 0.349213 | 0.48 | 0.332 | 1 |
| CRTAC1_Cells | SOST | 0.000279885 | 0.635834 | 0.27 | 0.177 | 1 |
| CRTAC1_Cells | ARL5A | 0.002081298 | 0.292288 | 0.699 | 0.573 | 1 |

**Table S3. Marker genes for mesenchymal cell clusters in pig abdominal aorta plaque scRNA-seq data. Corresponds to Fig. 1B.**
The listed genes have significantly higher expression in the indicated cluster than all other clusters combined.
Analysis was performed using the Wilcoxon test in Seurat v4.4.0 with P values adjusted for multiple testing by the Bonferroni method.

| cluster | gene | p_val | avg_log2F( | pct.1 | pct.2 | p_val_adj |
|---|---|---|---|---|---|---|
| IFN_Interm | ISG15 | 1.62E-46 | 5.167669 | 1 | 0.095 | 2.82E-42 |
| Pericytes | CXCL2 | 2.8E-112 | 4.24042 | 0.6 | 0.044 | 4.9E-108 |
| Fibroblasts | DCN | 3.86E-43 | 4.167784 | 0.983 | 0.426 | 6.74E-39 |
| Fibroblasts | PI16 | 1.1E-127 | 4.013529 | 0.85 | 0.059 | 1.9E-123 |
| Pericytes | RASD1 | 3.8E-151 | 3.960995 | 0.7 | 0.044 | 6.6E-147 |
| Pericytes | CCL19 | 1.19E-24 | 3.664457 | 0.5 | 0.155 | 2.08E-20 |
| Fibroblasts | F3 | 5.4E-153 | 3.577797 | 0.65 | 0.024 | 9.4E-149 |
| IFN_Interm | RIGI | 1.27E-61 | 3.514733 | 0.789 | 0.039 | 2.22E-57 |
| Pericytes | GJA4 | 0 | 3.418973 | 0.7 | 0.003 | 0 |
| Terminal | MT1X | 8.59E-62 | 3.378345 | 0.258 | 0.052 | 1.5E-57 |
| IFN_Interm | MX1 | 1.45E-23 | 3.376867 | 0.895 | 0.159 | 2.54E-19 |
| IFN_Interm | ENSSSCG0( | 6.89E-20 | 3.272129 | 1 | 0.3 | 1.2E-15 |
| Terminal | SAA3 | 1.67E-31 | 3.243956 | 0.185 | 0.053 | 2.92E-27 |
| Fibroblasts | NUPR1 | 3.96E-41 | 3.204886 | 0.817 | 0.209 | 6.93E-37 |
| Fibroblasts | OGN | 6.08E-97 | 3.116458 | 0.717 | 0.056 | 1.06E-92 |
| IFN_Interm | PLAC8 | 1.18E-75 | 3.106195 | 0.947 | 0.045 | 2.06E-71 |
| Contractile | TNNC1 | 6.7E-263 | 3.094443 | 0.714 | 0.018 | 1.2E-258 |
| IFN_Interm | ISG12(A) | 2.41E-13 | 3.088045 | 1 | 0.485 | 4.21E-09 |
| IFN_Interm | IFIT1 | 5.04E-66 | 2.999487 | 0.632 | 0.022 | 8.81E-62 |
| Fibroblasts | FBLN1 | 4.6E-39 | 2.930535 | 0.917 | 0.343 | 8.04E-35 |
| Pericytes | ATF3 | 2.04E-36 | 2.926003 | 0.556 | 0.128 | 3.57E-32 |
| Fibroblasts | ITGBL1 | 5.09E-77 | 2.925724 | 0.817 | 0.1 | 8.91E-73 |
| Pericytes | C7 | 7.1E-186 | 2.884784 | 0.644 | 0.024 | 1.2E-181 |
| Fibroblasts | PTX3 | 2.62E-08 | 2.881889 | 0.1 | 0.013 | 0.000459 |
| Pericytes | BTG2 | 3.23E-26 | 2.856698 | 0.633 | 0.236 | 5.65E-22 |
| Fibroblasts | APOD | 0.000778 | 2.760409 | 0.3 | 0.172 | 1 |
| Pericytes | HIGD1B | 0 | 2.756228 | 0.667 | 0.001 | 0 |
| Pericytes | COL5A3 | 2.65E-60 | 2.666672 | 0.689 | 0.133 | 4.63E-56 |
| IFN_Interm | IFI6 | 8.24E-12 | 2.622318 | 1 | 0.734 | 1.44E-07 |
| Contractile | ACTA2 | 0 | 2.617769 | 0.995 | 0.665 | 0 |
| IFN_Interm | MX2 | 6.57E-93 | 2.605486 | 0.684 | 0.017 | 1.15E-88 |
| Fibroblasts | HSD11B1 | 5.98E-38 | 2.593198 | 0.817 | 0.225 | 1.04E-33 |
| Fibroblasts | ALDH1A3 | 2.41E-70 | 2.55381 | 0.417 | 0.023 | 4.22E-66 |
| Contractile | RGS5 | 7.12E-52 | 2.472963 | 1 | 0.462 | 1.24E-47 |
| Pericytes | FABP4 | 7.52E-22 | 2.458146 | 0.733 | 0.419 | 1.31E-17 |
| Fibroblasts | MFAP5 | 7.4E-218 | 2.457254 | 0.35 | 0.001 | 1.3E-213 |
| Contractile | MYH11 | 0 | 2.434086 | 0.964 | 0.317 | 0 |
| Pericytes | RGS16 | 1.7E-89 | 2.422276 | 0.367 | 0.018 | 2.97E-85 |
| IFN_Interm | ENSSSCG0( | 3.58E-28 | 2.408959 | 0.632 | 0.057 | 6.27E-24 |
| Pericytes | GJC1 | 4.7E-54 | 2.407248 | 0.7 | 0.159 | 8.22E-50 |
| Pericytes | PGF | 1.19E-52 | 2.398014 | 0.644 | 0.131 | 2.09E-48 |
| IFN_Interm | EPSTI1 | 1.28E-19 | 2.395449 | 0.632 | 0.083 | 2.23E-15 |
| Terminal | SCGB2A2 | 7.99E-29 | 2.395034 | 0.102 | 0.015 | 1.4E-24 |
| Fibroblasts | CFD | 4.35E-21 | 2.359924 | 0.283 | 0.039 | 7.61E-17 |
| Pericytes | TINAGL1 | 2.38E-53 | 2.356875 | 0.778 | 0.207 | 4.17E-49 |
| Terminal | NDUFA4L2 | 5.6E-231 | 2.355381 | 0.898 | 0.34 | 9.8E-227 |
| Pericytes | FHL5 | 3.2E-99 | 2.347121 | 0.778 | 0.099 | 5.65E-95 |
| Pericytes | RGS5 | 7.41E-41 | 2.339562 | 0.967 | 0.463 | 1.3E-36 |
| Pericytes | CYGB | 2.2E-257 | 2.332006 | 0.544 | 0.007 | 3.8E-253 |
| Terminal | SOD2 | 5E-99 | 2.313432 | 0.661 | 0.274 | 8.66E-95 |
| Pericytes | ADAMTS1 | 8.1E-24 | 2.295859 | 0.489 | 0.141 | 1.42E-19 |
| IFN_Interm | LGALS9 | 6.77E-43 | 2.282502 | 0.632 | 0.035 | 1.18E-38 |
| Contractile | SPARCL1 | 1.76E-46 | 2.282266 | 1 | 0.53 | 3.08E-42 |
| Fibroblasts | TXNIP | 7.77E-20 | 2.279826 | 0.9 | 0.534 | 1.36E-15 |

| | | | | | | |
|---|---|---|---|---|---|---|
| Pericytes | FRMD3 | 7.6E-306 | 2.271047 | 0.589 | 0.005 | 1.3E-301 |
| Pericytes | GUCY1B1 | 4.75E-67 | 2.265132 | 0.744 | 0.143 | 8.31E-63 |
| Fibroblasts | PDPN | 3.8E-179 | 2.246606 | 0.55 | 0.011 | 6.7E-175 |
| Pericytes | ZFP36 | 1.17E-23 | 2.24284 | 0.744 | 0.416 | 2.04E-19 |
| Terminal | ECM1 | 3E-199 | 2.216447 | 0.94 | 0.658 | 5.2E-195 |
| Fibroblasts | MYOC | 5.2E-137 | 2.207542 | 0.2 | 0 | 9.1E-133 |
| Pericytes | ITIH5 | 1.42E-59 | 2.202086 | 0.667 | 0.114 | 2.49E-55 |
| Pericytes | SOCS3 | 7.15E-32 | 2.195399 | 0.656 | 0.221 | 1.25E-27 |
| Terminal | ANXA8 | 1E-185 | 2.190909 | 0.743 | 0.222 | 1.8E-181 |
| Contractile | SEMA3E | 2.26E-65 | 2.189396 | 0.813 | 0.169 | 3.96E-61 |
| Pericytes | FAM162B | 1.2E-294 | 2.188912 | 0.456 | 0.001 | 2.2E-290 |
| Contractile | PPP1R14A | 4.6E-308 | 2.186129 | 0.901 | 0.282 | 8E-304 |
| Contractile | CNN1 | 2E-275 | 2.183986 | 0.784 | 0.178 | 3.6E-271 |
| Pericytes | SEPTIN4 | 1.24E-58 | 2.183946 | 0.689 | 0.137 | 2.16E-54 |
| Fibroblasts | CRISPLD2 | 2.02E-41 | 2.149387 | 0.767 | 0.169 | 3.53E-37 |
| Pericytes | MYO1B | 1.71E-74 | 2.129729 | 0.5 | 0.049 | 2.99E-70 |
| Pericytes | PTP4A3 | 3.8E-149 | 2.126252 | 0.578 | 0.027 | 6.6E-145 |
| IFN_Interm | ENSSSCG0( | 4.1E-11 | 2.122927 | 0.947 | 0.465 | 7.16E-07 |
| Pericytes | APOD | 5.25E-52 | 2.117459 | 0.733 | 0.158 | 9.17E-48 |
| Fibroblasts | ANGPTL4 | 2.67E-30 | 2.112366 | 0.733 | 0.195 | 4.67E-26 |
| Fibroblasts | OMD | 5.38E-18 | 2.109293 | 0.617 | 0.226 | 9.4E-14 |
| Pericytes | FOS | 3.79E-18 | 2.104658 | 0.8 | 0.532 | 6.63E-14 |
| IFN_Interm | HERC5 | 3.34E-53 | 2.097028 | 0.632 | 0.027 | 5.83E-49 |
| Pericytes | NR4A1 | 4.37E-30 | 2.082414 | 0.567 | 0.162 | 7.64E-26 |
| Pericytes | NR2F2 | 4.67E-41 | 2.080893 | 0.822 | 0.323 | 8.16E-37 |
| IFN_Interm | OAS2 | 1.08E-52 | 2.069021 | 0.684 | 0.033 | 1.89E-48 |
| IFN_Interm | RSAD2 | 6.6E-190 | 2.066686 | 0.579 | 0.004 | 1.2E-185 |
| Pericytes | JUN | 1.79E-16 | 2.062373 | 0.789 | 0.582 | 3.13E-12 |
| Pericytes | ARPC1A | 1.03E-35 | 2.055559 | 0.722 | 0.264 | 1.81E-31 |
| Fibroblasts | LCN2 | 1.17E-12 | 2.054822 | 0.433 | 0.132 | 2.05E-08 |
| IFN_Interm | LGALS3BP | 1.32E-07 | 2.035303 | 0.737 | 0.29 | 0.002305 |
| Pericytes | FOSB | 2.62E-09 | 2.021567 | 0.456 | 0.236 | 4.58E-05 |
| Pericytes | CCL2 | 3.77E-13 | 2.020364 | 0.5 | 0.21 | 6.59E-09 |
| Contractile | DES | 1.39E-39 | 1.999399 | 0.407 | 0.058 | 2.43E-35 |
| Pericytes | AVPR1A | 2.7E-249 | 1.982326 | 0.389 | 0.001 | 4.7E-245 |
| Contractile | MYL9 | 1.2E-282 | 1.966679 | 0.96 | 0.536 | 2.1E-278 |
| Pericytes | EPS8 | 1.23E-33 | 1.941797 | 0.589 | 0.165 | 2.15E-29 |
| Pericytes | SLC2A3 | 2.72E-30 | 1.937272 | 0.744 | 0.311 | 4.75E-26 |
| Fibroblasts | MEDAG | 1.02E-20 | 1.932808 | 0.383 | 0.074 | 1.78E-16 |
| Fibroblasts | BTG2 | 1.73E-17 | 1.909541 | 0.667 | 0.24 | 3.03E-13 |
| Pericytes | NRP1 | 3.71E-28 | 1.901547 | 0.733 | 0.333 | 6.49E-24 |
| Intermedia | SERPINF1 | 1.1E-190 | 1.900568 | 0.867 | 0.392 | 1.9E-186 |
| Terminal | S100A10 | 7E-143 | 1.892978 | 0.914 | 0.717 | 1.2E-138 |
| Fibroblasts | SPTBN1 | 5.72E-17 | 1.892155 | 0.817 | 0.525 | 1E-12 |
| Pericytes | ADAMTS4 | 1.99E-52 | 1.889604 | 0.333 | 0.029 | 3.48E-48 |
| IFN_Interm | B2M.1 | 1.44E-10 | 1.885543 | 1 | 0.859 | 2.52E-06 |
| Fibroblasts | GALNT15 | 5.22E-16 | 1.876106 | 0.533 | 0.17 | 9.13E-12 |
| Pericytes | BTG3 | 4.68E-34 | 1.875272 | 0.633 | 0.199 | 8.18E-30 |
| IFN_Interm | S100A1 | 0.000343 | 1.873935 | 0.526 | 0.216 | 1 |
| Fibroblasts | PCOLCE2 | 8.11E-26 | 1.871988 | 0.6 | 0.15 | 1.42E-21 |
| Fibroblasts | C3 | 9.68E-09 | 1.87178 | 0.2 | 0.046 | 0.000169 |
| Fibroblasts | FOS | 9.53E-11 | 1.866815 | 0.767 | 0.535 | 1.67E-06 |
| Pericytes | ITIH3 | 7.2E-192 | 1.866549 | 0.422 | 0.006 | 1.3E-187 |
| IFN_Interm | ZNFX1 | 8.32E-09 | 1.864933 | 0.526 | 0.12 | 0.000145 |
| Pericytes | LOC10052: | 2.09E-26 | 1.853721 | 0.944 | 0.887 | 3.65E-22 |
| IFN_Interm | IFIT5 | 2.05E-12 | 1.852201 | 0.474 | 0.072 | 3.58E-08 |
| Pericytes | GUCY1A1 | 1.3E-49 | 1.851952 | 0.622 | 0.126 | 2.27E-45 |
| Pericytes | NOTCH3 | 9.96E-32 | 1.851283 | 0.844 | 0.432 | 1.74E-27 |
| Pericytes | KLHL23 | 7.5E-57 | 1.848568 | 0.444 | 0.051 | 1.31E-52 |

| | | | | | | |
|---|---|---|---|---|---|---|
| Fibroblasts | LAMA2 | 4.1E-239 | 1.847693 | 0.367 | 0 | 7.1E-235 |
| Fibroblasts | LEPR | 1.5E-185 | 1.84417 | 0.4 | 0.004 | 2.7E-181 |
| Pericytes | ENSSSCG0( | 2.18E-24 | 1.840572 | 0.289 | 0.047 | 3.81E-20 |
| Pericytes | APOLD1 | 3.1E-118 | 1.837133 | 0.311 | 0.007 | 5.4E-114 |
| Contractile | NEXN | 2.3E-279 | 1.833696 | 0.965 | 0.6 | 4E-275 |
| IFN_Interm | ENSSSCG0( | 7.25E-12 | 1.830564 | 0.421 | 0.057 | 1.27E-07 |
| Contractile | MYH11 | 8.31E-36 | 1.828375 | 1 | 0.564 | 1.45E-31 |
| IFN_Interm | PARP14 | 2.36E-40 | 1.82194 | 0.526 | 0.025 | 4.13E-36 |
| IFN_Interm | SAMD9 | 1.56E-12 | 1.819754 | 0.526 | 0.082 | 2.72E-08 |
| Pericytes | MYC | 2.25E-53 | 1.818014 | 0.489 | 0.067 | 3.94E-49 |
| Terminal | BNIP3 | 2.3E-145 | 1.813915 | 0.841 | 0.498 | 4.1E-141 |
| Fibroblasts | PTGFR | 4.09E-95 | 1.804061 | 0.4 | 0.014 | 7.15E-91 |
| Pericytes | PDGFRB | 1.02E-21 | 1.803671 | 0.767 | 0.476 | 1.79E-17 |
| Pericytes | ENSSSCG0( | 3.5E-36 | 1.802623 | 0.611 | 0.168 | 6.11E-32 |
| Fibroblasts | PDK4 | 1.12E-34 | 1.800881 | 0.333 | 0.032 | 1.96E-30 |
| Pericytes | CYCS | 4.47E-07 | 1.795218 | 0.533 | 0.405 | 0.007817 |
| Contractile | TGM2 | 1.15E-07 | 1.793755 | 0.637 | 0.416 | 0.002007 |
| Fibroblasts | FIBIN | 3.47E-10 | 1.787459 | 0.517 | 0.226 | 6.06E-06 |
| Contractile | FBXO32 | 8.11E-34 | 1.785046 | 0.879 | 0.403 | 1.42E-29 |
| Contractile | CPE | 5.75E-36 | 1.780685 | 0.956 | 0.625 | 1E-31 |
| Pericytes | RHOB | 2.33E-22 | 1.776162 | 0.844 | 0.557 | 4.08E-18 |
| Pericytes | ENSSSCG0( | 1.31E-37 | 1.773081 | 0.467 | 0.084 | 2.28E-33 |
| Pericytes | ENSSSCG0( | 4.33E-20 | 1.769833 | 0.622 | 0.272 | 7.58E-16 |
| Pericytes | EBF1 | 1.65E-27 | 1.769244 | 0.578 | 0.193 | 2.88E-23 |
| Pericytes | PDGFA | 1.1E-26 | 1.768161 | 0.689 | 0.286 | 1.92E-22 |
| Pericytes | NFKBIA | 2.59E-14 | 1.767545 | 0.622 | 0.358 | 4.53E-10 |
| IFN_Interm | IFI44L | 2.63E-15 | 1.76695 | 0.684 | 0.118 | 4.59E-11 |
| Fibroblasts | ZFP36 | 2.17E-17 | 1.762485 | 0.8 | 0.418 | 3.79E-13 |
| Contractile | FHL1 | 4.11E-34 | 1.755949 | 0.978 | 0.698 | 7.19E-30 |
| Contractile | ADIRF | 3.38E-38 | 1.752642 | 0.989 | 0.676 | 5.91E-34 |
| Fibroblasts | SAA3 | 1.77E-05 | 1.751831 | 0.233 | 0.081 | 0.308897 |
| Contractile | PPP1R14A | 4E-36 | 1.739691 | 0.989 | 0.517 | 6.99E-32 |
| Fibroblasts | ENSSSCG0( | 3.56E-55 | 1.738866 | 0.25 | 0.01 | 6.22E-51 |
| IFN_Interm | UBE2L6 | 7.14E-19 | 1.738862 | 0.632 | 0.084 | 1.25E-14 |
| Fibroblasts | CHRDL1 | 5.4E-167 | 1.738666 | 0.417 | 0.005 | 9.4E-163 |
| Contractile | ACTG2 | 5.06E-88 | 1.738456 | 0.285 | 0.038 | 8.85E-84 |
| Pericytes | ADAP2 | 1E-182 | 1.737896 | 0.4 | 0.006 | 1.8E-178 |
| Contractile | FLNA | 6.8E-251 | 1.736277 | 0.975 | 0.75 | 1.2E-246 |
| IFN_Interm | EIF2AK2 | 2.63E-06 | 1.735756 | 0.632 | 0.254 | 0.045971 |
| Pericytes | CAMK2N1 | 2.85E-20 | 1.735226 | 0.6 | 0.269 | 4.99E-16 |
| Fibroblasts | TGFBR3 | 5.11E-72 | 1.731465 | 0.55 | 0.042 | 8.94E-68 |
| Pericytes | EDNRB | 1.3E-191 | 1.731176 | 0.356 | 0.003 | 2.2E-187 |
| Pericytes | DEPP1 | 3.37E-37 | 1.728767 | 0.256 | 0.024 | 5.89E-33 |
| Pericytes | SLC7A2 | 1.73E-28 | 1.720514 | 0.678 | 0.249 | 3.03E-24 |
| Contractile | IL1RL1 | 5.05E-95 | 1.720426 | 0.393 | 0.09 | 8.82E-91 |
| Fibroblasts | IGFBP6 | 2.52E-09 | 1.716327 | 0.883 | 0.648 | 4.41E-05 |
| Contractile | ITIH4 | 2.3E-178 | 1.714472 | 0.891 | 0.524 | 4E-174 |
| Fibroblasts | PLSCR4 | 1.77E-23 | 1.713109 | 0.5 | 0.112 | 3.09E-19 |
| Fibroblasts | IGF2 | 3.99E-24 | 1.712885 | 0.417 | 0.07 | 6.98E-20 |
| Fibroblasts | CXCL12 | 3.99E-43 | 1.707257 | 0.45 | 0.047 | 6.97E-39 |
| Fibroblasts | ENSSSCG0( | 1.65E-37 | 1.70268 | 0.333 | 0.03 | 2.89E-33 |
| IFN_Interm | STAT1 | 2.19E-09 | 1.696684 | 0.684 | 0.2 | 3.83E-05 |
| Pericytes | SPRY2 | 3.15E-45 | 1.696375 | 0.478 | 0.076 | 5.51E-41 |
| Fibroblasts | TFPI2 | 5.21E-09 | 1.690564 | 0.717 | 0.496 | 9.11E-05 |
| Contractile | CSRP1 | 2.9E-283 | 1.690371 | 0.983 | 0.726 | 5E-279 |
| Pericytes | PHLDA2 | 1.13E-18 | 1.688779 | 0.844 | 0.582 | 1.97E-14 |
| IFN_Interm | GBP1 | 8.4E-35 | 1.685763 | 0.474 | 0.024 | 1.47E-30 |
| Contractile | SMTN | 4.51E-34 | 1.685657 | 0.912 | 0.427 | 7.89E-30 |
| Pericytes | GTF2E2 | 4.68E-16 | 1.681869 | 0.656 | 0.404 | 8.19E-12 |

| | | | | | | |
|---|---|---|---|---|---|---|
| Fibroblasts | FGL2 | 2.64E-15 | 1.677276 | 0.833 | 0.469 | 4.62E-11 |
| Contractile | FIBIN | 1.96E-34 | 1.675307 | 0.714 | 0.218 | 3.43E-30 |
| Fibroblasts | SLPI | 3.44E-53 | 1.668253 | 0.283 | 0.014 | 6.02E-49 |
| Pericytes | RCSD1 | 6.63E-77 | 1.654753 | 0.478 | 0.041 | 1.16E-72 |
| Pericytes | ANO3 | 1.8E-110 | 1.653846 | 0.389 | 0.015 | 3.2E-106 |
| Terminal | COL6A3 | 7.7E-136 | 1.653038 | 0.905 | 0.663 | 1.3E-131 |
| Pericytes | IFRD1 | 5.58E-18 | 1.64874 | 0.456 | 0.158 | 9.75E-14 |
| Fibroblasts | SVEP1 | 3.37E-24 | 1.639027 | 0.517 | 0.116 | 5.89E-20 |
| Pericytes | JUNB | 1.77E-15 | 1.63021 | 0.8 | 0.612 | 3.1E-11 |
| Contractile | SYNPO2 | 5.52E-27 | 1.629096 | 0.934 | 0.608 | 9.66E-23 |
| IFN_Interm | IRF7 | 1.08E-23 | 1.624164 | 0.474 | 0.036 | 1.89E-19 |
| Pericytes | EPHX3 | 2.64E-77 | 1.622538 | 0.333 | 0.018 | 4.62E-73 |
| Terminal | COMP | 6.26E-18 | 1.622096 | 0.214 | 0.099 | 1.09E-13 |
| Contractile | TPM2 | 1.5E-272 | 1.60947 | 0.994 | 0.823 | 2.6E-268 |
| Pericytes | ENSSSCG0( | 3.78E-18 | 1.60933 | 0.689 | 0.378 | 6.61E-14 |
| Contractile | CCN3 | 1.3E-219 | 1.599796 | 0.976 | 0.667 | 2.3E-215 |
| Pericytes | RELL1 | 4.16E-16 | 1.594903 | 0.389 | 0.129 | 7.26E-12 |
| Contractile | A2M | 4.5E-219 | 1.594726 | 0.974 | 0.674 | 7.9E-215 |
| Fibroblasts | JUN | 2.7E-10 | 1.591472 | 0.783 | 0.584 | 4.72E-06 |
| Terminal | GAPDH | 2.4E-168 | 1.590124 | 0.977 | 0.897 | 4.3E-164 |
| Pericytes | ABCC9 | 1.27E-58 | 1.588144 | 0.333 | 0.025 | 2.22E-54 |
| IFN_Interm | SETX | 0.000198 | 1.587705 | 0.474 | 0.188 | 1 |
| Terminal | LCN2 | 8.91E-90 | 1.587627 | 0.361 | 0.071 | 1.56E-85 |
| Fibroblasts | TNXB | 1.28E-56 | 1.581355 | 0.333 | 0.018 | 2.23E-52 |
| Pericytes | STOM | 1.14E-26 | 1.57964 | 0.833 | 0.468 | 1.99E-22 |
| Pericytes | TRA2B | 4.15E-17 | 1.574924 | 0.7 | 0.442 | 7.26E-13 |
| IFN_Interm | SP140 | 2.21E-08 | 1.562687 | 0.579 | 0.154 | 0.000387 |
| Fibroblasts | GFPT2 | 2.4E-106 | 1.557345 | 0.333 | 0.007 | 4.2E-102 |
| Pericytes | CXCL14 | 3.45E-18 | 1.557297 | 0.278 | 0.057 | 6.03E-14 |
| Contractile | MYLK | 2.7E-199 | 1.550068 | 0.912 | 0.502 | 4.7E-195 |
| Pericytes | DES | 8.54E-31 | 1.549761 | 0.367 | 0.059 | 1.49E-26 |
| Fibroblasts | CLEC3B | 4.08E-37 | 1.540905 | 0.3 | 0.024 | 7.13E-33 |
| IFN_Interm | IFITM1 | 3.54E-15 | 1.538253 | 0.526 | 0.071 | 6.19E-11 |
| Fibroblasts | GADD45G | 1.82E-07 | 1.534004 | 0.433 | 0.199 | 0.003185 |
| Fibroblasts | MMP2 | 1.67E-18 | 1.532209 | 0.867 | 0.533 | 2.91E-14 |
| Fibroblasts | CDKN1C | 3.45E-34 | 1.521125 | 0.367 | 0.039 | 6.04E-30 |
| Fibroblasts | SERPINF1 | 2.43E-21 | 1.520916 | 0.933 | 0.519 | 4.24E-17 |
| Terminal | SPP1 | 9.43E-25 | 1.520111 | 0.281 | 0.127 | 1.65E-20 |
| Pericytes | GADD45A | 2.63E-12 | 1.516259 | 0.567 | 0.324 | 4.6E-08 |
| Pericytes | PPP1R15A | 1.09E-11 | 1.51447 | 0.544 | 0.303 | 1.91E-07 |
| Terminal | LGALS3 | 1.2E-139 | 1.509919 | 0.904 | 0.659 | 2.1E-135 |
| Pericytes | GNG2 | 4.95E-45 | 1.50832 | 0.3 | 0.027 | 8.66E-41 |
| Contractile | LMOD1 | 1.59E-30 | 1.506521 | 0.967 | 0.547 | 2.79E-26 |
| Contractile | GLDN | 7.66E-39 | 1.505923 | 0.495 | 0.087 | 1.34E-34 |
| Terminal | IGFBP3 | 1.89E-45 | 1.489654 | 0.184 | 0.034 | 3.31E-41 |
| Pericytes | ZFP36L1 | 1.25E-19 | 1.488499 | 0.9 | 0.724 | 2.19E-15 |
| Fibroblasts | CEBPD | 2.38E-15 | 1.479595 | 0.933 | 0.745 | 4.17E-11 |
| Fibroblasts | EGR1 | 7.95E-08 | 1.478331 | 0.717 | 0.477 | 0.001389 |
| Contractile | MYH10 | 4E-175 | 1.477939 | 0.922 | 0.554 | 7E-171 |
| IFN_Interm | HLA-DRA | 6.74E-05 | 1.475809 | 0.368 | 0.101 | 1 |
| Pericytes | EDNRA | 2.8E-33 | 1.472033 | 0.411 | 0.073 | 4.89E-29 |
| Contractile | MYL9 | 3.31E-28 | 1.471072 | 0.989 | 0.698 | 5.79E-24 |
| Pericytes | KCNE4 | 3.02E-50 | 1.470226 | 0.344 | 0.033 | 5.27E-46 |
| Fibroblasts | MATN2 | 9.48E-20 | 1.469713 | 0.383 | 0.074 | 1.66E-15 |
| Terminal | HAPLN1 | 3.33E-83 | 1.469149 | 0.222 | 0.02 | 5.83E-79 |
| Pericytes | CDKN1C | 4.09E-17 | 1.465398 | 0.222 | 0.04 | 7.14E-13 |
| Pericytes | RASL11A | 3.93E-11 | 1.451992 | 0.6 | 0.361 | 6.86E-07 |
| Pericytes | ITIH4 | 2.21E-13 | 1.450429 | 0.833 | 0.667 | 3.87E-09 |
| Contractile | CNN1 | 9.28E-29 | 1.447326 | 0.923 | 0.407 | 1.62E-24 |

| | | | | | | |
|---|---|---|---|---|---|---|
| Fibroblasts | ENSSSCG0( | 2.2E-216 | 1.44599 | 0.333 | 0 | 3.9E-212 |
| Terminal | ENO1 | 1.7E-201 | 1.445563 | 0.988 | 0.895 | 2.9E-197 |
| Pericytes | PEAR1 | 6.9E-30 | 1.44515 | 0.533 | 0.14 | 1.21E-25 |
| Contractile | TAGLN | 4.5E-274 | 1.44383 | 0.999 | 0.933 | 7.9E-270 |
| Fibroblasts | FGF7 | 6.67E-06 | 1.443369 | 0.283 | 0.111 | 0.116554 |
| Pericytes | STUM | 1.4E-203 | 1.441644 | 0.311 | 0 | 2.4E-199 |
| Fibroblasts | MATN4 | 1.5E-67 | 1.437827 | 0.35 | 0.016 | 2.61E-63 |
| Terminal | TIMP3 | 3.13E-59 | 1.437109 | 0.704 | 0.447 | 5.48E-55 |
| Fibroblasts | SOCS3 | 0.000173 | 1.434341 | 0.4 | 0.23 | 1 |
| IFN_Interm | HERC6 | 6.27E-12 | 1.429052 | 0.579 | 0.108 | 1.1E-07 |
| IFN_Interm | GHSR | 1.14E-05 | 1.421595 | 0.368 | 0.092 | 0.198721 |
| Pericytes | LRRC10B | 1.03E-81 | 1.420859 | 0.333 | 0.016 | 1.79E-77 |
| Terminal | SERPINE2 | 3.67E-87 | 1.41881 | 0.775 | 0.422 | 6.41E-83 |
| Terminal | EDIL3 | 3.5E-112 | 1.417067 | 0.727 | 0.334 | 6.2E-108 |
| IFN_Interm | PSMB8 | 9.19E-09 | 1.412961 | 0.579 | 0.144 | 0.000161 |
| IFN_Interm | ENSSSCG0( | 5.55E-07 | 1.406777 | 0.579 | 0.182 | 0.009708 |
| Fibroblasts | RSPO3 | 3.79E-74 | 1.406111 | 0.283 | 0.008 | 6.63E-70 |
| Terminal | LOXL2 | 1.5E-105 | 1.399673 | 0.634 | 0.249 | 2.7E-101 |
| Fibroblasts | AQP3 | 3.03E-09 | 1.397066 | 0.45 | 0.18 | 5.3E-05 |
| Pericytes | EGR1 | 1.29E-07 | 1.396566 | 0.622 | 0.477 | 0.002258 |
| Pericytes | OLFML2A | 8.3E-32 | 1.393634 | 0.378 | 0.065 | 1.45E-27 |
| Pericytes | P3H2 | 6.78E-66 | 1.391562 | 0.411 | 0.036 | 1.19E-61 |
| Pericytes | IER2 | 0.002006 | 1.385313 | 0.389 | 0.313 | 1 |
| Pericytes | PRXL2A | 1.66E-29 | 1.37352 | 0.356 | 0.062 | 2.91E-25 |
| IFN_Interm | LOC11025! | 4.46E-15 | 1.373155 | 0.421 | 0.045 | 7.8E-11 |
| Intermedia | S100A1 | 6.16E-44 | 1.371113 | 0.373 | 0.156 | 1.08E-39 |
| Terminal | DDIT4 | 1.66E-80 | 1.367119 | 0.77 | 0.432 | 2.9E-76 |
| Contractile | ITGA8 | 1.9E-191 | 1.364335 | 0.93 | 0.558 | 3.4E-187 |
| Contractile | SORBS1 | 7.09E-27 | 1.357237 | 0.681 | 0.241 | 1.24E-22 |
| IFN_Interm | DTX3L | 1.97E-17 | 1.355405 | 0.368 | 0.03 | 3.44E-13 |
| Fibroblasts | CILP2 | 1.57E-06 | 1.354782 | 0.2 | 0.056 | 0.027527 |
| Fibroblasts | GNG11 | 1.44E-06 | 1.35362 | 0.683 | 0.656 | 0.025231 |
| Pericytes | NEURL1B | 2.68E-93 | 1.349417 | 0.289 | 0.009 | 4.68E-89 |
| Fibroblasts | FAM180B | 1.98E-66 | 1.345469 | 0.367 | 0.018 | 3.46E-62 |
| Pericytes | ENSSSCG0( | 4.84E-34 | 1.344264 | 1 | 0.996 | 8.45E-30 |
| Fibroblasts | DHRS3 | 3.24E-10 | 1.341355 | 0.55 | 0.271 | 5.67E-06 |
| Pericytes | PLAU | 9.42E-23 | 1.34016 | 0.311 | 0.06 | 1.65E-18 |
| Intermedia | PCOLCE | 1.7E-141 | 1.340112 | 0.867 | 0.504 | 2.9E-137 |
| Terminal | RGCC | 3.11E-47 | 1.337757 | 0.476 | 0.226 | 5.44E-43 |
| Pericytes | IFITM1 | 1.51E-18 | 1.333606 | 0.3 | 0.067 | 2.64E-14 |
| IFN_Interm | CMPK2 | 2.27E-50 | 1.331176 | 0.526 | 0.019 | 3.96E-46 |
| Contractile | CSRP1 | 3.53E-29 | 1.328652 | 1 | 0.824 | 6.17E-25 |
| IFN_Interm | SLA-DQB1 | 0.000847 | 1.327128 | 0.526 | 0.242 | 1 |
| IFN_Interm | SMPDL3B | 0.007744 | 1.326915 | 0.474 | 0.261 | 1 |
| Pericytes | RALGPS2 | 1.84E-51 | 1.326734 | 0.244 | 0.015 | 3.21E-47 |
| Pericytes | GPR4 | 2.6E-227 | 1.326536 | 0.367 | 0.001 | 4.6E-223 |
| Fibroblasts | CYP1B1 | 9.18E-05 | 1.322032 | 0.267 | 0.119 | 1 |
| Contractile | RAMP1 | 4.3E-37 | 1.321457 | 0.626 | 0.152 | 7.52E-33 |
| Pericytes | CD59 | 9.35E-22 | 1.319598 | 0.656 | 0.272 | 1.63E-17 |
| Pericytes | ITGA7 | 1.01E-14 | 1.318718 | 0.633 | 0.348 | 1.76E-10 |
| Terminal | IGFBP5 | 4.89E-38 | 1.317673 | 0.687 | 0.471 | 8.56E-34 |
| Pericytes | GPER1 | 1.04E-12 | 1.314782 | 0.467 | 0.213 | 1.82E-08 |
| IFN_Interm | IFIH1 | 3.16E-09 | 1.314509 | 0.526 | 0.11 | 5.52E-05 |
| Contractile | CRIM1 | 3.33E-16 | 1.314257 | 0.747 | 0.439 | 5.83E-12 |
| Contractile | SORBS2 | 2.05E-19 | 1.313591 | 0.813 | 0.479 | 3.58E-15 |
| Fibroblasts | ADGRD1 | 2.39E-56 | 1.312492 | 0.25 | 0.009 | 4.17E-52 |
| Terminal | MAL | 3.9E-123 | 1.308018 | 0.281 | 0.016 | 6.8E-119 |
| Fibroblasts | HTRA3 | 1.09E-32 | 1.306074 | 0.317 | 0.031 | 1.9E-28 |
| Fibroblasts | ABCA9 | 3.8E-99 | 1.303384 | 0.267 | 0.004 | 6.63E-95 |

| | | | | | | |
|---|---|---|---|---|---|---|
| Terminal | ANXA2 | 1.3E-108 | 1.303181 | 0.938 | 0.895 | 2.3E-104 |
| Contractile | DSTN | 1.2E-209 | 1.300748 | 0.975 | 0.818 | 2.1E-205 |
| Contractile | EFEMP1 | 6.6E-165 | 1.298873 | 0.858 | 0.42 | 1.2E-160 |
| Fibroblasts | PMP22 | 2.74E-08 | 1.298273 | 0.7 | 0.502 | 0.000479 |
| Fibroblasts | FOSB | 6.99E-08 | 1.298157 | 0.517 | 0.237 | 0.001222 |
| Pericytes | MCAM | 1.17E-14 | 1.297553 | 0.556 | 0.243 | 2.05E-10 |
| Contractile | SMTN | 1.6E-160 | 1.2964 | 0.718 | 0.255 | 2.8E-156 |
| Pericytes | EGFLAM | 1.6E-141 | 1.294701 | 0.278 | 0.003 | 2.7E-137 |
| Contractile | LMOD1 | 4.7E-187 | 1.29437 | 0.843 | 0.369 | 8.3E-183 |
| Contractile | PDLIM7 | 1.3E-160 | 1.290184 | 0.78 | 0.359 | 2.2E-156 |
| Contractile | CALD1 | 5.2E-239 | 1.287981 | 0.997 | 0.943 | 9.1E-235 |
| Contractile | RGS5 | 2.2E-195 | 1.287586 | 0.81 | 0.255 | 3.8E-191 |
| Fibroblasts | PROCR | 8.52E-14 | 1.284409 | 0.233 | 0.04 | 1.49E-09 |
| Pericytes | LOC10051 | 1.03E-11 | 1.283067 | 0.622 | 0.375 | 1.8E-07 |
| Contractile | ANGPTL7 | 8.48E-60 | 1.28252 | 0.352 | 0.028 | 1.48E-55 |
| Fibroblasts | YBX3 | 4.21E-07 | 1.282462 | 0.933 | 0.797 | 0.007352 |
| Pericytes | RBM24 | 3.08E-28 | 1.280261 | 0.3 | 0.046 | 5.39E-24 |
| Terminal | SMPDL3B | 2.9E-107 | 1.280085 | 0.564 | 0.172 | 5E-103 |
| Terminal | CCL2 | 7.85E-46 | 1.279668 | 0.403 | 0.163 | 1.37E-41 |
| Terminal | CAPG | 9.42E-94 | 1.277641 | 0.83 | 0.604 | 1.65E-89 |
| Pericytes | SPP1 | 0.00337 | 1.275472 | 0.256 | 0.16 | 1 |
| Fibroblasts | RNASE4 | 5.93E-10 | 1.274273 | 0.817 | 0.615 | 1.04E-05 |
| Pericytes | RASL12 | 3.68E-36 | 1.272658 | 0.367 | 0.053 | 6.43E-32 |
| Pericytes | SNX10 | 1.51E-19 | 1.261304 | 0.456 | 0.145 | 2.64E-15 |
| Contractile | NET1 | 1.95E-20 | 1.258185 | 0.857 | 0.523 | 3.41E-16 |
| IFN_Interm | ENSSSCG0( | 1E-05 | 1.258064 | 0.947 | 0.677 | 0.175132 |
| Contractile | ITIH5 | 3.19E-19 | 1.257034 | 0.429 | 0.121 | 5.57E-15 |
| IFN_Interm | DDX60 | 3.58E-44 | 1.25508 | 0.421 | 0.014 | 6.25E-40 |
| Contractile | PDZRN4 | 5.64E-28 | 1.251686 | 0.593 | 0.173 | 9.86E-24 |
| Intermedia | COL11A1 | 1.46E-47 | 1.250679 | 0.182 | 0.032 | 2.56E-43 |
| Pericytes | PCP4L1 | 1.36E-33 | 1.249968 | 0.311 | 0.041 | 2.37E-29 |
| Pericytes | SOX18 | 1.59E-43 | 1.249835 | 0.256 | 0.02 | 2.77E-39 |
| Pericytes | MARCKS | 1.8E-06 | 1.249461 | 0.522 | 0.367 | 0.031412 |
| Terminal | LDHA | 1.1E-129 | 1.246701 | 0.927 | 0.71 | 1.9E-125 |
| Contractile | EFHD1 | 1.7E-154 | 1.246246 | 0.641 | 0.193 | 3E-150 |
| Fibroblasts | JUNB | 1.08E-07 | 1.2458 | 0.8 | 0.614 | 0.00189 |
| Pericytes | DLX5 | 6.58E-36 | 1.24445 | 0.344 | 0.046 | 1.15E-31 |
| Terminal | RCAN1 | 2.26E-75 | 1.244366 | 0.357 | 0.088 | 3.94E-71 |
| Contractile | FRZB | 4.52E-28 | 1.243036 | 0.769 | 0.281 | 7.9E-24 |
| Pericytes | CHN1 | 5.72E-45 | 1.236304 | 0.267 | 0.021 | 1E-40 |
| Contractile | ADIRF | 3E-166 | 1.231705 | 0.919 | 0.528 | 5.3E-162 |
| IFN_Interm | IFIT3 | 3.61E-87 | 1.231383 | 0.316 | 0.003 | 6.32E-83 |
| Pericytes | SYNE2 | 2.1E-10 | 1.230496 | 0.611 | 0.4 | 3.67E-06 |
| Pericytes | SEPTIN11 | 1.1E-12 | 1.228547 | 0.667 | 0.472 | 1.92E-08 |
| Fibroblasts | EPHA3 | 2.9E-74 | 1.227804 | 0.3 | 0.01 | 5.06E-70 |
| Contractile | TPM2 | 1.76E-28 | 1.226246 | 1 | 0.888 | 3.08E-24 |
| Fibroblasts | COL14A1 | 4.76E-10 | 1.223098 | 0.883 | 0.665 | 8.32E-06 |
| Pericytes | VCAM1 | 1.39E-11 | 1.223042 | 0.7 | 0.462 | 2.42E-07 |
| Pericytes | ANGPT1 | 1.38E-18 | 1.221067 | 0.256 | 0.05 | 2.41E-14 |
| Intermedia | LUM | 5.4E-146 | 1.220847 | 0.991 | 0.852 | 9.4E-142 |
| Pericytes | ENSSSCG0( | 8.53E-31 | 1.219344 | 0.278 | 0.036 | 1.49E-26 |
| Intermedia | ALDH2 | 4.28E-93 | 1.219207 | 0.594 | 0.256 | 7.48E-89 |
| Pericytes | NRIP2 | 1E-141 | 1.219075 | 0.278 | 0.003 | 1.8E-137 |
| Contractile | DSTN | 1.65E-26 | 1.218486 | 0.989 | 0.878 | 2.88E-22 |
| Contractile | VCL | 4.3E-169 | 1.213747 | 0.934 | 0.66 | 7.5E-165 |
| Fibroblasts | GLT8D2 | 1.59E-09 | 1.210131 | 0.517 | 0.241 | 2.77E-05 |
| Terminal | EMP1 | 5.19E-92 | 1.209416 | 0.877 | 0.672 | 9.07E-88 |
| Fibroblasts | CYP27A1 | 2.76E-79 | 1.209374 | 0.3 | 0.009 | 4.83E-75 |
| IFN_Interm | PHF11 | 3.16E-06 | 1.208895 | 0.632 | 0.22 | 0.055199 |

| | | | | | | |
|---|---|---|---|---|---|---|
| Pericytes | HGF | 2.68E-22 | 1.207397 | 0.378 | 0.089 | 4.68E-18 |
| Intermedia | SFRP2 | 1.04E-21 | 1.205863 | 0.116 | 0.031 | 1.81E-17 |
| Pericytes | PARM1 | 3.17E-69 | 1.205837 | 0.256 | 0.01 | 5.54E-65 |
| IFN_Interm | PML | 0.000652 | 1.205257 | 0.263 | 0.072 | 1 |
| Pericytes | ANGPT2 | 5.78E-18 | 1.204585 | 0.411 | 0.124 | 1.01E-13 |
| IFN_Interm | GNG11 | 0.000168 | 1.203083 | 0.789 | 0.656 | 1 |
| Fibroblasts | SLA-6 | 1.27E-07 | 1.201639 | 0.583 | 0.34 | 0.002213 |
| Pericytes | ISYNA1 | 1.74E-40 | 1.200771 | 0.233 | 0.018 | 3.05E-36 |
| Pericytes | PDE1C | 3.55E-20 | 1.195951 | 0.322 | 0.071 | 6.21E-16 |
| Fibroblasts | C7 | 0.0003 | 1.194714 | 0.133 | 0.04 | 1 |
| Pericytes | CEBPB | 2.79E-09 | 1.194587 | 0.656 | 0.512 | 4.88E-05 |
| Pericytes | LOC10012 | 6.97E-27 | 1.191585 | 0.278 | 0.041 | 1.22E-22 |
| Fibroblasts | PLPP3 | 7.68E-10 | 1.189696 | 0.867 | 0.672 | 1.34E-05 |
| Pericytes | ENSSSCG0 | 1.3E-23 | 1.187862 | 0.222 | 0.03 | 2.28E-19 |
| Terminal | VEGFA | 1.38E-77 | 1.183131 | 0.486 | 0.163 | 2.41E-73 |
| Terminal | EGLN3 | 1.38E-85 | 1.182483 | 0.471 | 0.139 | 2.41E-81 |
| Pericytes | DLC1 | 7.07E-12 | 1.18125 | 0.467 | 0.221 | 1.24E-07 |
| Terminal | PTGES | 8.6E-96 | 1.177728 | 0.512 | 0.156 | 1.5E-91 |
| Pericytes | ETS1 | 4.86E-14 | 1.175935 | 0.567 | 0.298 | 8.49E-10 |
| IFN_Interm | LOC10051 | 1.94E-06 | 1.175935 | 0.526 | 0.155 | 0.033831 |
| Fibroblasts | MGP | 2.26E-14 | 1.172006 | 1 | 0.992 | 3.95E-10 |
| IFN_Interm | ENSSSCG0 | 3.87E-13 | 1.169664 | 0.368 | 0.04 | 6.76E-09 |
| Contractile | TNS1 | 7.89E-20 | 1.168099 | 0.736 | 0.365 | 1.38E-15 |
| Pericytes | ENSSSCG0 | 4.18E-37 | 1.166734 | 1 | 0.996 | 7.31E-33 |
| Contractile | SLIT3 | 6.8E-161 | 1.166064 | 0.897 | 0.501 | 1.2E-156 |
| Contractile | PDLIM7 | 3.65E-16 | 1.165051 | 0.835 | 0.519 | 6.39E-12 |
| Contractile | RND3 | 4.7E-11 | 1.162017 | 0.571 | 0.296 | 8.22E-07 |
| Pericytes | ETS2 | 9.98E-08 | 1.162016 | 0.4 | 0.221 | 0.001745 |
| Pericytes | AIG1 | 2.56E-26 | 1.161922 | 0.389 | 0.082 | 4.47E-22 |
| Intermedia | SFRP1 | 6.54E-61 | 1.159185 | 0.632 | 0.391 | 1.14E-56 |
| Contractile | KANK1 | 2.93E-20 | 1.156886 | 0.857 | 0.535 | 5.12E-16 |
| Pericytes | PLEKHA5 | 1.27E-20 | 1.156547 | 0.267 | 0.049 | 2.23E-16 |
| Intermedia | ABI3BP | 3.07E-70 | 1.15453 | 0.648 | 0.339 | 5.38E-66 |
| Fibroblasts | GLUL | 2.45E-09 | 1.153667 | 0.567 | 0.263 | 4.27E-05 |
| Pericytes | ADAMTS1 | 1.86E-34 | 1.153638 | 0.289 | 0.034 | 3.24E-30 |
| IFN_Interm | SCGB2A2 | 2.96E-05 | 1.15314 | 0.211 | 0.034 | 0.518084 |
| IFN_Interm | ENSSSCG0 | 0.00013 | 1.152429 | 0.895 | 0.76 | 1 |
| IFN_Interm | ENSSSCG0 | 0.000224 | 1.151797 | 0.947 | 0.805 | 1 |
| Contractile | MYLK | 1.86E-28 | 1.151171 | 1 | 0.657 | 3.26E-24 |
| IFN_Interm | SLC25A28 | 5.4E-07 | 1.150469 | 0.474 | 0.123 | 0.009438 |
| Contractile | LOC10051 | 1.35E-14 | 1.14686 | 0.714 | 0.372 | 2.36E-10 |
| Pericytes | COL4A1 | 3.28E-06 | 1.142688 | 0.933 | 0.927 | 0.057401 |
| Pericytes | PRPF4B | 3.29E-09 | 1.142453 | 0.589 | 0.405 | 5.76E-05 |
| Fibroblasts | C4B | 1.24E-23 | 1.142273 | 0.133 | 0.007 | 2.17E-19 |
| Pericytes | PIM3 | 5.45E-19 | 1.138397 | 0.311 | 0.071 | 9.53E-15 |
| Pericytes | SHOX2 | 1.64E-31 | 1.137796 | 0.3 | 0.04 | 2.87E-27 |
| Terminal | PCOLCE2 | 3.85E-90 | 1.133388 | 0.394 | 0.087 | 6.72E-86 |
| Contractile | RHOB | 1.06E-15 | 1.13313 | 0.857 | 0.556 | 1.86E-11 |
| Fibroblasts | PDGFRL | 5.52E-08 | 1.131614 | 0.683 | 0.435 | 0.000965 |
| Contractile | HSPB7 | 9.5E-107 | 1.131426 | 0.528 | 0.174 | 1.7E-102 |
| Fibroblasts | LTBP4 | 1.48E-07 | 1.131021 | 0.65 | 0.425 | 0.002585 |
| Terminal | SOX9 | 1.8E-100 | 1.130998 | 0.291 | 0.032 | 3.16E-96 |
| Pericytes | RERGL | 1.6E-99 | 1.129967 | 0.211 | 0.003 | 2.79E-95 |
| Terminal | SLC39A14 | 9.92E-63 | 1.12789 | 0.42 | 0.142 | 1.73E-58 |
| Terminal | SLC16A3 | 9.1E-113 | 1.123636 | 0.558 | 0.156 | 1.6E-108 |
| Pericytes | COX4I2 | 5.51E-15 | 1.120938 | 0.878 | 0.738 | 9.64E-11 |
| Pericytes | ENSSSCG0 | 1.03E-52 | 1.117647 | 0.167 | 0.005 | 1.8E-48 |
| Contractile | FGL2 | 1.08E-17 | 1.114779 | 0.824 | 0.466 | 1.88E-13 |
| Contractile | ACTA2 | 2.71E-22 | 1.112705 | 1 | 0.792 | 4.74E-18 |

| | | | | | | |
|---|---|---|---|---|---|---|
| Pericytes | SIK1 | 1.6E-18 | 1.110171 | 0.333 | 0.083 | 2.79E-14 |
| Contractile | MCAM | 2.1E-137 | 1.10896 | 0.491 | 0.092 | 3.6E-133 |
| Pericytes | TRPC6 | 1.74E-13 | 1.107899 | 0.278 | 0.076 | 3.05E-09 |
| Pericytes | JAG1 | 1.98E-15 | 1.105169 | 0.711 | 0.423 | 3.47E-11 |
| Pericytes | LDHB | 5.99E-13 | 1.104224 | 0.733 | 0.556 | 1.05E-08 |
| Fibroblasts | ENSSSCG0( | 4.94E-11 | 1.103965 | 0.917 | 0.713 | 8.63E-07 |
| Pericytes | EPAS1 | 1.99E-12 | 1.101672 | 0.778 | 0.645 | 3.48E-08 |
| Pericytes | SLC8A1 | 4.69E-25 | 1.100827 | 0.389 | 0.084 | 8.21E-21 |
| Pericytes | CCNL1 | 5.92E-10 | 1.100548 | 0.467 | 0.241 | 1.03E-05 |
| Fibroblasts | ALDH1A1 | 1.81E-16 | 1.100395 | 0.267 | 0.044 | 3.16E-12 |
| Fibroblasts | FGF18 | 1.92E-37 | 1.100065 | 0.233 | 0.014 | 3.36E-33 |
| Pericytes | TNFRSF21 | 1.5E-86 | 1.099736 | 0.244 | 0.006 | 2.63E-82 |
| Contractile | MYL6 | 1.4E-194 | 1.099318 | 0.991 | 0.941 | 2.5E-190 |
| Contractile | C6H19ORF | 6.55E-15 | 1.099111 | 0.692 | 0.394 | 1.15E-10 |
| Contractile | LMCD1 | 5.98E-93 | 1.096339 | 0.552 | 0.209 | 1.05E-88 |
| Contractile | ADGRE5 | 1.3E-117 | 1.096056 | 0.659 | 0.269 | 2.3E-113 |
| IFN_Interm | RARRES1 | 0.00096 | 1.095347 | 0.789 | 0.705 | 1 |
| Pericytes | CISD1 | 1.28E-07 | 1.093307 | 0.422 | 0.245 | 0.002229 |
| Fibroblasts | FOXO1 | 8.03E-16 | 1.08962 | 0.483 | 0.142 | 1.4E-11 |
| Contractile | ITGA7 | 3.9E-134 | 1.087739 | 0.612 | 0.184 | 6.7E-130 |
| Contractile | FLNA | 1.3E-20 | 1.083953 | 0.989 | 0.836 | 2.28E-16 |
| Terminal | SEMA3C | 6.1E-70 | 1.083494 | 0.48 | 0.169 | 1.07E-65 |
| Contractile | CAMK2G | 4.47E-17 | 1.083254 | 0.626 | 0.291 | 7.82E-13 |
| Pericytes | NCKAP5 | 1.43E-80 | 1.082955 | 0.256 | 0.008 | 2.5E-76 |
| Contractile | LOC10051 | 2.3E-100 | 1.082498 | 0.604 | 0.233 | 3.97E-96 |
| Pericytes | LDLR | 2.45E-71 | 1.081413 | 0.289 | 0.014 | 4.29E-67 |
| Contractile | PCGF5 | 3.7E-113 | 1.08103 | 0.861 | 0.538 | 6.5E-109 |
| Fibroblasts | PDGFRA | 0.000429 | 1.080697 | 0.367 | 0.209 | 1 |
| IFN_Interm | ITGA11 | 0.009029 | 1.079919 | 0.579 | 0.344 | 1 |
| Contractile | PGM5 | 3.17E-30 | 1.078892 | 0.396 | 0.071 | 5.54E-26 |
| Fibroblasts | FMO2 | 4.73E-84 | 1.077837 | 0.15 | 0.001 | 8.27E-80 |
| Pericytes | LRRC8A | 4.41E-15 | 1.076555 | 0.356 | 0.109 | 7.71E-11 |
| Fibroblasts | ABI3BP | 6.65E-13 | 1.074999 | 0.783 | 0.42 | 1.16E-08 |
| Contractile | TAGLN | 9.7E-27 | 1.0742 | 1 | 0.959 | 1.7E-22 |
| Intermedia | MMP2 | 9.33E-68 | 1.072183 | 0.715 | 0.469 | 1.63E-63 |
| Pericytes | TBX2 | 7.33E-32 | 1.071843 | 0.3 | 0.04 | 1.28E-27 |
| Pericytes | LAMA4 | 3.15E-06 | 1.071678 | 0.578 | 0.47 | 0.055016 |
| Pericytes | IGFBP4 | 4.16E-27 | 1.070652 | 0.322 | 0.054 | 7.28E-23 |
| Contractile | SCUBE1 | 2.37E-44 | 1.070529 | 0.308 | 0.029 | 4.14E-40 |
| Pericytes | ARHGAP29 | 3.5E-08 | 1.070373 | 0.433 | 0.239 | 0.000612 |
| Contractile | IFI6 | 4.33E-89 | 1.070051 | 0.89 | 0.633 | 7.56E-85 |
| Terminal | ACKR3 | 1.33E-83 | 1.068914 | 0.384 | 0.089 | 2.32E-79 |
| Contractile | SPARCL1 | 2.1E-113 | 1.066156 | 0.766 | 0.395 | 3.7E-109 |
| Fibroblasts | SAT1 | 1.69E-12 | 1.065957 | 0.767 | 0.4 | 2.96E-08 |
| Fibroblasts | NR4A1 | 4.37E-07 | 1.065417 | 0.4 | 0.169 | 0.00764 |
| IFN_Interm | CAMK2D | 0.00085 | 1.064729 | 0.632 | 0.323 | 1 |
| Terminal | PGAM1 | 6.59E-87 | 1.064256 | 0.77 | 0.467 | 1.15E-82 |
| Fibroblasts | CD302 | 1.21E-11 | 1.063414 | 0.683 | 0.358 | 2.11E-07 |
| Contractile | FILIP1L | 3.57E-89 | 1.057373 | 0.669 | 0.329 | 6.24E-85 |
| Fibroblasts | FBLN5 | 3.16E-05 | 1.055836 | 0.533 | 0.376 | 0.552536 |
| Contractile | ATP1A2 | 2.45E-47 | 1.055226 | 0.352 | 0.036 | 4.28E-43 |
| Fibroblasts | KLF2 | 0.000267 | 1.053413 | 0.633 | 0.447 | 1 |
| Contractile | NUF2 | 3.25E-21 | 1.051342 | 0.385 | 0.092 | 5.69E-17 |
| Pericytes | KCNJ8 | 1.53E-90 | 1.049774 | 0.233 | 0.005 | 2.68E-86 |
| Pericytes | FARP1 | 2.79E-08 | 1.048928 | 0.522 | 0.346 | 0.000487 |
| Contractile | TTLL7 | 5.1E-121 | 1.048285 | 0.785 | 0.45 | 8.9E-117 |
| Pericytes | CDC42EP4 | 5.02E-28 | 1.045776 | 0.333 | 0.056 | 8.77E-24 |
| Fibroblasts | TGFBR2 | 3.77E-09 | 1.045111 | 0.55 | 0.273 | 6.59E-05 |
| Terminal | UAP1 | 9.21E-55 | 1.044475 | 0.508 | 0.231 | 1.61E-50 |

| | | | | | | |
|---|---|---|---|---|---|---|
| Contractile | LPP | 7.17E-17 | 1.043516 | 0.791 | 0.454 | 1.25E-12 |
| Fibroblasts | MT1X | 3.84E-28 | 1.043257 | 0.517 | 0.091 | 6.71E-24 |
| Fibroblasts | ABCA6 | 3.7E-103 | 1.041384 | 0.15 | 0 | 6.5E-99 |
| Terminal | FN1 | 7.53E-72 | 1.040858 | 0.999 | 0.995 | 1.32E-67 |
| Terminal | CP | 1.2E-24 | 1.040855 | 0.247 | 0.108 | 2.1E-20 |
| IFN_Interm | GPX3 | 0.001437 | 1.040536 | 0.368 | 0.134 | 1 |
| IFN_Interm | PLPP1 | 0.00027 | 1.037958 | 0.737 | 0.469 | 1 |
| IFN_Interm | CTSS | 0.000458 | 1.03695 | 0.474 | 0.203 | 1 |
| Fibroblasts | PCOLCE | 1.16E-10 | 1.035544 | 0.867 | 0.603 | 2.03E-06 |
| Fibroblasts | GPNMB | 9.91E-11 | 1.034988 | 0.317 | 0.082 | 1.73E-06 |
| Contractile | FABP4 | 5.5E-161 | 1.034851 | 0.713 | 0.237 | 9.5E-157 |
| Terminal | CD9 | 1.16E-93 | 1.034746 | 0.962 | 0.909 | 2.02E-89 |
| Pericytes | NAALAD2 | 6.89E-11 | 1.034015 | 0.278 | 0.089 | 1.2E-06 |
| Pericytes | DBNDD2 | 5.02E-10 | 1.031922 | 0.433 | 0.218 | 8.78E-06 |
| Fibroblasts | IER2 | 0.000321 | 1.031463 | 0.483 | 0.312 | 1 |
| Terminal | ENSSSCG0( | 3.53E-36 | 1.031459 | 0.475 | 0.258 | 6.17E-32 |
| Contractile | PALLD | 1.4E-104 | 1.030168 | 0.681 | 0.319 | 2.4E-100 |
| Contractile | S100A2 | 6.98E-50 | 1.029782 | 0.285 | 0.088 | 1.22E-45 |
| Pericytes | KCNA5 | 5.05E-08 | 1.028779 | 0.322 | 0.143 | 0.000884 |
| Pericytes | ECE1 | 3.93E-35 | 1.027697 | 0.267 | 0.028 | 6.87E-31 |
| Pericytes | TOP1 | 8.67E-08 | 1.023611 | 0.611 | 0.482 | 0.001515 |
| Pericytes | CRISPLD2 | 3.68E-12 | 1.022899 | 0.433 | 0.173 | 6.43E-08 |
| Contractile | COL4A5 | 7.54E-09 | 1.022723 | 0.604 | 0.374 | 0.000132 |
| Fibroblasts | CD34 | 8.73E-64 | 1.021974 | 0.333 | 0.016 | 1.53E-59 |
| IFN_Interm | PSMB9 | 1.31E-05 | 1.021892 | 0.263 | 0.05 | 0.22973 |
| Terminal | RNH1 | 7.86E-85 | 1.021289 | 0.801 | 0.545 | 1.37E-80 |
| Intermedia | CTHRC1 | 2.7E-36 | 1.020153 | 0.385 | 0.191 | 4.72E-32 |
| Pericytes | BAALC | 1.1E-95 | 1.019087 | 0.222 | 0.004 | 1.93E-91 |
| Fibroblasts | CD59 | 3.65E-06 | 1.018413 | 0.5 | 0.279 | 0.063804 |
| IFN_Interm | IFIT2 | 1.3E-145 | 1.017127 | 0.211 | 0 | 2.2E-141 |
| Pericytes | TAGLN2 | 1.46E-16 | 1.016706 | 0.856 | 0.67 | 2.56E-12 |
| Pericytes | WTIP | 4.21E-12 | 1.014807 | 0.333 | 0.115 | 7.36E-08 |
| Fibroblasts | PIK3R1 | 1.63E-08 | 1.01472 | 0.683 | 0.441 | 0.000284 |
| Contractile | IGFBP7 | 8.9E-221 | 1.01407 | 1 | 0.998 | 1.6E-216 |
| Pericytes | TPPP3 | 8.95E-10 | 1.013505 | 0.322 | 0.122 | 1.56E-05 |
| Pericytes | REM1 | 4.74E-18 | 1.012695 | 0.289 | 0.064 | 8.28E-14 |
| Fibroblasts | SELENBP1 | 7.08E-18 | 1.008353 | 0.233 | 0.031 | 1.24E-13 |
| Pericytes | OAZ2 | 3.72E-08 | 1.008032 | 0.4 | 0.209 | 0.00065 |
| Pericytes | HEYL | 6.93E-07 | 1.006451 | 0.378 | 0.213 | 0.012122 |
| Contractile | LPP | 3.3E-114 | 1.005286 | 0.689 | 0.312 | 5.7E-110 |
| Pericytes | BCAM | 5.85E-15 | 1.003626 | 0.389 | 0.13 | 1.02E-10 |
| Contractile | FRZB | 2.1E-100 | 1.001475 | 0.504 | 0.155 | 3.63E-96 |
| Terminal | CRLF1 | 1.25E-65 | 0.999711 | 0.298 | 0.065 | 2.19E-61 |
| Contractile | CCDC3 | 2.24E-12 | 0.999589 | 0.659 | 0.391 | 3.91E-08 |
| Contractile | ZBTB16 | 2.18E-12 | 0.999208 | 0.571 | 0.293 | 3.81E-08 |
| Contractile | PKDCC | 2.01E-97 | 0.999112 | 0.75 | 0.438 | 3.52E-93 |
| Pericytes | REV3L | 1.32E-09 | 0.998828 | 0.378 | 0.166 | 2.3E-05 |
| Fibroblasts | THBS4 | 4.76E-07 | 0.997586 | 0.117 | 0.021 | 0.008329 |
| Pericytes | ITM2B | 4.56E-22 | 0.995753 | 0.978 | 0.951 | 7.98E-18 |
| Pericytes | ENSSSCG0( | 2.27E-14 | 0.994055 | 0.911 | 0.802 | 3.97E-10 |
| Pericytes | ENSSSCG0( | 1.39E-12 | 0.993475 | 0.289 | 0.084 | 2.42E-08 |
| Contractile | CD200 | 1.35E-10 | 0.992593 | 0.593 | 0.321 | 2.36E-06 |
| Pericytes | NR4A3 | 4.5E-13 | 0.99056 | 0.156 | 0.026 | 7.86E-09 |
| Fibroblasts | ADM | 0.00387 | 0.990181 | 0.217 | 0.107 | 1 |
| Terminal | TSPO | 5.8E-111 | 0.990071 | 0.952 | 0.868 | 1E-106 |
| IFN_Interm | FUNDC1 | 0.000179 | 0.989201 | 0.474 | 0.174 | 1 |
| Fibroblasts | ZFP36L2 | 0.001875 | 0.989102 | 0.783 | 0.751 | 1 |
| Terminal | EMP3 | 4.2E-79 | 0.988899 | 0.882 | 0.724 | 7.33E-75 |
| Contractile | BTC | 5.49E-85 | 0.988872 | 0.286 | 0.01 | 9.6E-81 |

| | | | | | | |
|---|---|---|---|---|---|---|
| Contractile | ITIH4 | 0.00014 | 0.987891 | 0.736 | 0.669 | 1 |
| Pericytes | ENSSSCG0( | 4.03E-22 | 0.982187 | 0.211 | 0.028 | 7.04E-18 |
| Contractile | C7 | 3.21E-12 | 0.982039 | 0.187 | 0.038 | 5.62E-08 |
| Fibroblasts | ALDH2 | 2.85E-12 | 0.981429 | 0.733 | 0.345 | 4.99E-08 |
| Fibroblasts | CD55 | 7.23E-06 | 0.981092 | 0.3 | 0.122 | 0.126429 |
| Terminal | COL6A1 | 6.21E-79 | 0.979707 | 0.966 | 0.922 | 1.09E-74 |
| IFN_Interm | TIMP1 | 0.003036 | 0.979211 | 0.947 | 0.933 | 1 |
| Pericytes | PLEKHA2 | 4.6E-11 | 0.978927 | 0.267 | 0.084 | 8.04E-07 |
| Pericytes | AJUBA | 1.03E-22 | 0.978456 | 0.256 | 0.041 | 1.8E-18 |
| Fibroblasts | OSR1 | 3.48E-06 | 0.978262 | 0.117 | 0.024 | 0.060895 |
| Terminal | TPI1 | 5.23E-72 | 0.976855 | 0.85 | 0.667 | 9.14E-68 |
| Contractile | LIMCH1 | 1.7E-99 | 0.975096 | 0.763 | 0.432 | 2.96E-95 |
| IFN_Interm | UBA7 | 3.42E-12 | 0.974712 | 0.263 | 0.022 | 5.97E-08 |
| Contractile | CD74 | 3.69E-40 | 0.974195 | 0.455 | 0.241 | 6.45E-36 |
| IFN_Interm | CD74 | 6.4E-05 | 0.972638 | 0.737 | 0.325 | 1 |
| Fibroblasts | ACKR4 | 2.86E-41 | 0.971194 | 0.167 | 0.005 | 5.01E-37 |
| IFN_Interm | SNX10 | 0.000265 | 0.970879 | 0.421 | 0.152 | 1 |
| Pericytes | SEPTIN8 | 3.06E-05 | 0.970674 | 0.467 | 0.341 | 0.535538 |
| Pericytes | TULP3 | 6.79E-15 | 0.970005 | 0.378 | 0.124 | 1.19E-10 |
| Terminal | MYBL1 | 1.01E-49 | 0.969993 | 0.432 | 0.179 | 1.76E-45 |
| Contractile | FILIP1 | 4.6E-100 | 0.969616 | 0.589 | 0.217 | 8.02E-96 |
| Pericytes | HNRNPA2E | 5.75E-16 | 0.966986 | 0.967 | 0.869 | 1E-11 |
| Contractile | RASSF3 | 4.72E-12 | 0.966183 | 0.637 | 0.347 | 8.26E-08 |
| Terminal | CDA | 1.1E-103 | 0.966004 | 0.209 | 0.005 | 1.9E-99 |
| Contractile | PAWR | 1.4E-133 | 0.964827 | 0.925 | 0.692 | 2.4E-129 |
| Pericytes | ARHGAP1 | 1.22E-09 | 0.964471 | 0.411 | 0.198 | 2.14E-05 |
| Contractile | PRKG1 | 1.88E-91 | 0.964186 | 0.699 | 0.35 | 3.28E-87 |
| Terminal | NDRG1 | 1.42E-49 | 0.96378 | 0.561 | 0.289 | 2.48E-45 |
| Fibroblasts | NPM1 | 9.8E-07 | 0.961599 | 0.867 | 0.76 | 0.017127 |
| Pericytes | ANKRD24 | 5E-121 | 0.961169 | 0.189 | 0 | 8.8E-117 |
| Intermedia | COL14A1 | 1.34E-63 | 0.960691 | 0.854 | 0.596 | 2.34E-59 |
| Pericytes | RFTN1 | 1.54E-09 | 0.95935 | 0.267 | 0.092 | 2.69E-05 |
| Contractile | NEXN | 1.46E-19 | 0.957989 | 1 | 0.739 | 2.56E-15 |
| Terminal | P4HA1 | 5.98E-71 | 0.957688 | 0.77 | 0.501 | 1.05E-66 |
| IFN_Interm | CTHRC1 | 0.000575 | 0.956918 | 0.579 | 0.244 | 1 |
| Pericytes | CSNK1E | 5.36E-08 | 0.954825 | 0.389 | 0.206 | 0.000937 |
| Pericytes | ENSSSCG0( | 2.74E-37 | 0.954444 | 0.256 | 0.024 | 4.8E-33 |
| Pericytes | NREP | 1.01E-06 | 0.953964 | 0.389 | 0.209 | 0.017608 |
| Pericytes | RCAN2 | 4.83E-08 | 0.953898 | 0.7 | 0.546 | 0.000844 |
| Terminal | PKM | 1.6E-101 | 0.953816 | 0.967 | 0.861 | 2.81E-97 |
| Pericytes | ARHGDIB | 6.85E-05 | 0.953266 | 0.489 | 0.399 | 1 |
| Contractile | AIF1L | 1.2E-114 | 0.952579 | 0.538 | 0.153 | 2.1E-110 |
| Pericytes | PLCL1 | 5.44E-13 | 0.952011 | 0.189 | 0.039 | 9.51E-09 |
| Contractile | PTGER3 | 3.72E-72 | 0.951227 | 0.341 | 0.02 | 6.5E-68 |
| Contractile | KCNE1 | 2.53E-86 | 0.950935 | 0.456 | 0.144 | 4.43E-82 |
| Fibroblasts | LPAR1 | 5.04E-24 | 0.950908 | 0.267 | 0.03 | 8.81E-20 |
| Contractile | MGLL | 1.92E-95 | 0.949568 | 0.663 | 0.32 | 3.36E-91 |
| Pericytes | IL13RA1 | 0.000193 | 0.949412 | 0.444 | 0.365 | 1 |
| Pericytes | TMEM150 | 7.49E-15 | 0.949178 | 0.278 | 0.069 | 1.31E-10 |
| Terminal | ALDOA | 1.74E-93 | 0.94833 | 0.929 | 0.8 | 3.05E-89 |
| Fibroblasts | DKK2 | 5.1E-95 | 0.945858 | 0.167 | 0.001 | 8.91E-91 |
| Contractile | TNC | 6.07E-66 | 0.945287 | 0.782 | 0.545 | 1.06E-61 |
| Pericytes | ENSSSCG0( | 1.4E-114 | 0.945271 | 0.189 | 0.001 | 2.4E-110 |
| Terminal | TXN | 2.5E-89 | 0.944698 | 0.981 | 0.917 | 4.36E-85 |
| IFN_Interm | IFI44 | 4.98E-13 | 0.943954 | 0.368 | 0.04 | 8.7E-09 |
| Pericytes | MAFB | 1.7E-05 | 0.943421 | 0.344 | 0.193 | 0.296628 |
| Pericytes | FILIP1 | 1.03E-08 | 0.942416 | 0.567 | 0.361 | 0.000181 |
| Contractile | HSPB1 | 2.4E-124 | 0.941584 | 0.978 | 0.909 | 4.2E-120 |
| Contractile | SYNPO2 | 1.49E-89 | 0.939924 | 0.795 | 0.499 | 2.61E-85 |

| | | | | | | |
|---|---|---|---|---|---|---|
| Contractile | KANK1 | 4.89E-94 | 0.93907 | 0.748 | 0.408 | 8.55E-90 |
| Fibroblasts | FBXO21 | 9.97E-11 | 0.938719 | 0.433 | 0.147 | 1.74E-06 |
| Contractile | FHL1 | 8.4E-116 | 0.936417 | 0.882 | 0.587 | 1.5E-111 |
| Fibroblasts | LTBP2 | 0.002443 | 0.934878 | 0.75 | 0.734 | 1 |
| Fibroblasts | TSC22D3 | 6.74E-07 | 0.934019 | 0.6 | 0.392 | 0.011781 |
| Pericytes | IER5 | 3.01E-06 | 0.93313 | 0.278 | 0.132 | 0.052664 |
| Fibroblasts | BEX3 | 5.89E-15 | 0.929778 | 0.417 | 0.105 | 1.03E-10 |
| Fibroblasts | FKBP5 | 0.000146 | 0.929383 | 0.55 | 0.403 | 1 |
| Contractile | ENSSSCG0( | 8.3E-91 | 0.929381 | 0.898 | 0.669 | 1.45E-86 |
| Fibroblasts | CCDC80 | 3.4E-08 | 0.929178 | 0.883 | 0.776 | 0.000595 |
| Pericytes | COTL1 | 9.77E-31 | 0.928816 | 0.267 | 0.032 | 1.71E-26 |
| Contractile | PEAR1 | 1.04E-18 | 0.928772 | 0.462 | 0.142 | 1.82E-14 |
| Contractile | CNTN1 | 2E-112 | 0.925681 | 0.176 | 0 | 3.5E-108 |
| Contractile | TES | 3.93E-16 | 0.923029 | 0.374 | 0.111 | 6.87E-12 |
| Fibroblasts | SPON1 | 4.96E-11 | 0.922893 | 0.517 | 0.194 | 8.67E-07 |
| Pericytes | SOCS1 | 2E-18 | 0.922749 | 0.211 | 0.034 | 3.49E-14 |
| Contractile | NDUFB1 | 1.83E-76 | 0.922178 | 0.676 | 0.393 | 3.2E-72 |
| Pericytes | LOC10051! | 2.01E-09 | 0.921123 | 0.356 | 0.151 | 3.52E-05 |
| IFN_Interm | EHD4 | 2.31E-05 | 0.919103 | 0.421 | 0.119 | 0.404416 |
| Fibroblasts | LIFR | 1.53E-10 | 0.916585 | 0.217 | 0.044 | 2.67E-06 |
| Pericytes | CAMK1 | 1.08E-06 | 0.916209 | 0.333 | 0.171 | 0.018888 |
| Pericytes | ATP2B1 | 1.94E-09 | 0.914714 | 0.367 | 0.159 | 3.39E-05 |
| Fibroblasts | MRTO4 | 1.43E-06 | 0.914496 | 0.3 | 0.113 | 0.024919 |
| Intermedia | CLU | 5.97E-86 | 0.91406 | 0.994 | 0.902 | 1.04E-81 |
| Pericytes | PPP1R12A | 4.09E-12 | 0.91386 | 0.833 | 0.679 | 7.16E-08 |
| Terminal | NT5E | 6E-68 | 0.913607 | 0.49 | 0.179 | 1.05E-63 |
| Fibroblasts | SFRP2 | 0.005859 | 0.913069 | 0.133 | 0.054 | 1 |
| Contractile | ENSSSCG0( | 8.64E-11 | 0.913054 | 0.604 | 0.32 | 1.51E-06 |
| Contractile | VCAN | 1.44E-88 | 0.912063 | 0.913 | 0.681 | 2.52E-84 |
| Contractile | STOM | 1.09E-10 | 0.908719 | 0.725 | 0.471 | 1.91E-06 |
| Terminal | TNFRSF12A | 3.55E-40 | 0.907053 | 0.628 | 0.422 | 6.21E-36 |
| IFN_Interm | ENSSSCG0( | 0.000188 | 0.906662 | 0.368 | 0.106 | 1 |
| Contractile | SCHIP1 | 1.93E-09 | 0.906649 | 0.451 | 0.222 | 3.37E-05 |
| Contractile | FILIP1L | 5.85E-14 | 0.905997 | 0.78 | 0.456 | 1.02E-09 |
| Terminal | IER3 | 1.16E-45 | 0.905931 | 0.342 | 0.12 | 2.03E-41 |
| Contractile | COL4A6 | 2.12E-10 | 0.905735 | 0.484 | 0.222 | 3.7E-06 |
| Terminal | SLC20A1 | 8.54E-53 | 0.905084 | 0.396 | 0.144 | 1.49E-48 |
| Pericytes | CHD4 | 3.63E-06 | 0.905041 | 0.611 | 0.509 | 0.06349 |
| Pericytes | RND3 | 0.001136 | 0.903999 | 0.4 | 0.301 | 1 |
| Pericytes | EIF1B | 1.19E-06 | 0.903749 | 0.556 | 0.447 | 0.02082 |
| Fibroblasts | TNFSF13 | 9.44E-06 | 0.903715 | 0.2 | 0.064 | 0.165096 |
| Contractile | MAP1B | 5.85E-97 | 0.902969 | 0.93 | 0.744 | 1.02E-92 |
| Pericytes | COL4A2 | 3.74E-05 | 0.902966 | 0.878 | 0.845 | 0.653844 |
| IFN_Interm | TAP1 | 4.92E-05 | 0.902698 | 0.316 | 0.077 | 0.859734 |
| Fibroblasts | C1QTNF4 | 1.47E-09 | 0.902695 | 0.317 | 0.093 | 2.57E-05 |
| Pericytes | SPTBN1 | 3.58E-10 | 0.902555 | 0.7 | 0.525 | 6.27E-06 |
| Fibroblasts | IL6 | 4.66E-05 | 0.901627 | 0.117 | 0.028 | 0.815194 |
| IFN_Interm | CXCL16 | 0.005478 | 0.898856 | 0.526 | 0.299 | 1 |
| Pericytes | LBH | 0.000268 | 0.898311 | 0.311 | 0.195 | 1 |
| Fibroblasts | ENSSSCG0( | 6.36E-36 | 0.897563 | 0.217 | 0.012 | 1.11E-31 |
| Contractile | GJA5 | 5.44E-78 | 0.89753 | 0.509 | 0.198 | 9.51E-74 |
| Terminal | YBX3 | 2.72E-85 | 0.897443 | 0.93 | 0.76 | 4.76E-81 |
| Intermedia | PDGFD | 8.36E-61 | 0.896208 | 0.605 | 0.323 | 1.46E-56 |
| Pericytes | ARL5B | 8.89E-08 | 0.896022 | 0.222 | 0.078 | 0.001553 |
| Contractile | PPP1R12B | 1.45E-14 | 0.895961 | 0.341 | 0.099 | 2.53E-10 |
| Pericytes | MSX2 | 6.59E-63 | 0.895269 | 0.222 | 0.008 | 1.15E-58 |
| Fibroblasts | H2AC6 | 0.000892 | 0.895173 | 0.183 | 0.073 | 1 |
| Pericytes | ENSSSCG0( | 1.96E-06 | 0.891894 | 0.422 | 0.252 | 0.034265 |
| IFN_Interm | IRF9 | 0.006015 | 0.89129 | 0.421 | 0.185 | 1 |

| | | | | | | |
|---|---|---|---|---|---|---|
| Terminal | POSTN | 2.89E-58 | 0.890951 | 0.962 | 0.909 | 5.05E-54 |
| Terminal | TCEA3 | 1.34E-51 | 0.890742 | 0.599 | 0.325 | 2.34E-47 |
| Fibroblasts | RUVBL1 | 9.77E-05 | 0.890167 | 0.283 | 0.127 | 1 |
| Pericytes | PLEKHH3 | 1.03E-09 | 0.889521 | 0.289 | 0.103 | 1.8E-05 |
| Fibroblasts | ADD3 | 2.59E-07 | 0.886724 | 0.367 | 0.145 | 0.004533 |
| Terminal | TPM3 | 7.43E-60 | 0.886513 | 0.674 | 0.394 | 1.3E-55 |
| Contractile | LSAMP | 6.63E-94 | 0.886162 | 0.375 | 0.078 | 1.16E-89 |
| Intermedia | COL1A1 | 2.17E-90 | 0.885402 | 1 | 0.975 | 3.8E-86 |
| Contractile | KCNAB1 | 1.64E-95 | 0.885243 | 0.447 | 0.118 | 2.87E-91 |
| Pericytes | UTRN | 1.62E-06 | 0.88464 | 0.711 | 0.588 | 0.028265 |
| Contractile | PLN | 1.31E-13 | 0.882732 | 0.549 | 0.226 | 2.28E-09 |
| Pericytes | INPP4B | 3.82E-34 | 0.880968 | 0.156 | 0.009 | 6.69E-30 |
| Contractile | TPM1 | 2.4E-134 | 0.880592 | 0.99 | 0.905 | 4.2E-130 |
| Contractile | EFHD1 | 5.28E-15 | 0.880383 | 0.703 | 0.363 | 9.24E-11 |
| Terminal | MIF | 1.2E-92 | 0.880241 | 0.951 | 0.818 | 2.09E-88 |
| Pericytes | GLUL | 0.000308 | 0.879577 | 0.378 | 0.266 | 1 |
| Contractile | SYTL2 | 3.52E-80 | 0.875474 | 0.536 | 0.209 | 6.16E-76 |
| Terminal | P4HA2 | 1.73E-58 | 0.873996 | 0.738 | 0.509 | 3.02E-54 |
| Pericytes | SLC22A18 | 1.16E-17 | 0.873529 | 0.256 | 0.051 | 2.02E-13 |
| Contractile | ACTN1 | 4.8E-102 | 0.87321 | 0.928 | 0.713 | 8.39E-98 |
| Pericytes | ARHGEF7 | 1.94E-08 | 0.871831 | 0.367 | 0.171 | 0.000339 |
| Contractile | ACTN4 | 1.1E-85 | 0.87179 | 0.855 | 0.603 | 1.93E-81 |
| Pericytes | EIF4A1 | 6.33E-07 | 0.870616 | 0.611 | 0.477 | 0.011073 |
| IFN_Interm | SELL | 4.14E-17 | 0.870467 | 0.105 | 0.002 | 7.24E-13 |
| Terminal | ENSSSCG0( | 2.84E-61 | 0.869837 | 0.579 | 0.276 | 4.97E-57 |
| Intermedia | MFAP2 | 1.06E-52 | 0.869686 | 0.628 | 0.391 | 1.84E-48 |
| Intermedia | SPARC | 2.3E-118 | 0.869393 | 0.996 | 0.969 | 4.1E-114 |
| Terminal | ANXA1 | 1.57E-56 | 0.869079 | 0.859 | 0.734 | 2.74E-52 |
| Contractile | IRAG1 | 2.65E-11 | 0.868835 | 0.648 | 0.362 | 4.63E-07 |
| IFN_Interm | LOC106510 | 5.59E-08 | 0.86723 | 0.316 | 0.048 | 0.000977 |
| Fibroblasts | IGF1 | 4.42E-18 | 0.866769 | 0.117 | 0.007 | 7.73E-14 |
| Contractile | MYL6 | 4.74E-20 | 0.865842 | 1 | 0.96 | 8.28E-16 |
| Fibroblasts | GHSR | 5.99E-05 | 0.864673 | 0.233 | 0.091 | 1 |
| Contractile | KCNAB1 | 1.2E-11 | 0.864261 | 0.516 | 0.242 | 2.1E-07 |
| Pericytes | BZW2 | 5.73E-10 | 0.862666 | 0.3 | 0.108 | 1E-05 |
| Terminal | SH3BGRL3 | 8.29E-53 | 0.861944 | 0.83 | 0.7 | 1.45E-48 |
| Contractile | COL18A1 | 5.1E-124 | 0.861606 | 0.976 | 0.786 | 8.8E-120 |
| Terminal | UPP1 | 4.25E-98 | 0.861398 | 0.258 | 0.022 | 7.43E-94 |
| IFN_Interm | ADD3 | 0.000475 | 0.860332 | 0.421 | 0.147 | 1 |
| Pericytes | B2M.1 | 1.11E-14 | 0.860295 | 0.944 | 0.857 | 1.94E-10 |
| Fibroblasts | GNG2 | 9.33E-33 | 0.859931 | 0.317 | 0.03 | 1.63E-28 |
| Terminal | CD44 | 1.26E-57 | 0.859418 | 0.623 | 0.32 | 2.2E-53 |
| Contractile | KCNMA1 | 1.19E-14 | 0.859366 | 0.516 | 0.206 | 2.09E-10 |
| Intermedia | CDO1 | 7.38E-58 | 0.85909 | 0.709 | 0.44 | 1.29E-53 |
| Contractile | IGF2 | 3.32E-23 | 0.85818 | 0.133 | 0.038 | 5.8E-19 |
| Intermedia | THBS1 | 1.14E-28 | 0.857967 | 0.921 | 0.866 | 1.99E-24 |
| IFN_Interm | GM2A | 0.003186 | 0.857736 | 0.316 | 0.11 | 1 |
| Pericytes | RBPMS | 3.12E-06 | 0.857145 | 0.511 | 0.35 | 0.054509 |
| Intermedia | IGFBP6 | 9.43E-39 | 0.857043 | 0.773 | 0.605 | 1.65E-34 |
| Pericytes | BRD3 | 8.51E-07 | 0.856636 | 0.444 | 0.275 | 0.014871 |
| Pericytes | TLR5 | 9.5E-100 | 0.856257 | 0.167 | 0.001 | 1.66E-95 |
| Fibroblasts | PLAC9 | 0.009469 | 0.855564 | 0.6 | 0.58 | 1 |
| Fibroblasts | MPZL1 | 1.31E-05 | 0.855522 | 0.483 | 0.283 | 0.228415 |
| Contractile | CAV2 | 4.39E-09 | 0.853274 | 0.538 | 0.303 | 7.68E-05 |
| Terminal | PSAT1 | 2.74E-90 | 0.852521 | 0.219 | 0.015 | 4.79E-86 |
| Contractile | IRAG1 | 8.05E-81 | 0.852519 | 0.567 | 0.238 | 1.41E-76 |
| Contractile | NIBAN1 | 8.95E-15 | 0.852148 | 0.934 | 0.732 | 1.57E-10 |
| Pericytes | TM4SF1 | 6.34E-08 | 0.85199 | 0.767 | 0.747 | 0.001108 |
| Terminal | BHLHE41 | 6.67E-65 | 0.85129 | 0.403 | 0.12 | 1.17E-60 |

| | | | | | | |
|---|---|---|---|---|---|---|
| Fibroblasts | ANG | 1.02E-07 | 0.849685 | 0.55 | 0.29 | 0.001783 |
| Contractile | MFAP4 | 1.7E-123 | 0.848405 | 0.988 | 0.834 | 3.1E-119 |
| Contractile | NFIB | 7.1E-10 | 0.847595 | 0.615 | 0.387 | 1.24E-05 |
| Terminal | DUT | 1.5E-23 | 0.847386 | 0.336 | 0.18 | 2.63E-19 |
| Contractile | CD93 | 1.4E-108 | 0.846595 | 0.346 | 0.046 | 2.4E-104 |
| Contractile | PALM2AKA | 8.19E-72 | 0.84635 | 0.751 | 0.477 | 1.43E-67 |
| Terminal | ITGA6 | 1.3E-102 | 0.846012 | 0.226 | 0.01 | 2.27E-98 |
| Pericytes | ENSSSCG00 | 8.1E-100 | 0.845494 | 0.167 | 0.001 | 1.42E-95 |
| Contractile | ITGA7 | 1.23E-09 | 0.845136 | 0.626 | 0.348 | 2.16E-05 |
| Contractile | PDLIM3 | 7.15E-11 | 0.844524 | 0.396 | 0.155 | 1.25E-06 |
| Terminal | LTC4S | 4.9E-33 | 0.844152 | 0.752 | 0.626 | 8.56E-29 |
| Pericytes | ATP5MC2 | 1.26E-05 | 0.843233 | 0.578 | 0.494 | 0.219643 |
| IFN_Interm | PARP9 | 5.6E-08 | 0.843148 | 0.263 | 0.034 | 0.000978 |
| Terminal | EZR | 2.65E-46 | 0.842945 | 0.594 | 0.34 | 4.63E-42 |
| Contractile | PLCL1 | 4.83E-34 | 0.842166 | 0.297 | 0.035 | 8.45E-30 |
| Contractile | SH3BGR | 1.72E-76 | 0.84095 | 0.496 | 0.199 | 3.01E-72 |
| Fibroblasts | SEMA3C | 0.000676 | 0.840676 | 0.4 | 0.238 | 1 |
| Contractile | CPED1 | 2.1E-18 | 0.84053 | 0.44 | 0.129 | 3.67E-14 |
| Pericytes | RAP2A | 6.72E-08 | 0.84028 | 0.256 | 0.096 | 0.001174 |
| Contractile | RASSF3 | 8.34E-70 | 0.839534 | 0.534 | 0.236 | 1.46E-65 |
| Intermedia | ITGA11 | 3.66E-54 | 0.837858 | 0.528 | 0.272 | 6.41E-50 |
| IFN_Interm | SIPA1L1 | 0.009114 | 0.83776 | 0.421 | 0.209 | 1 |
| Contractile | SORBS1 | 1.33E-72 | 0.837 | 0.425 | 0.139 | 2.32E-68 |
| Pericytes | EFHD2 | 9.11E-08 | 0.835934 | 0.367 | 0.185 | 0.001593 |
| Pericytes | SMARCA5 | 1.67E-06 | 0.834886 | 0.578 | 0.449 | 0.029172 |
| Pericytes | MXI1 | 1.14E-05 | 0.833945 | 0.367 | 0.214 | 0.199703 |
| Terminal | TMEM45A | 3.76E-66 | 0.833426 | 0.525 | 0.21 | 6.57E-62 |
| Fibroblasts | PIK3IP1 | 4.35E-06 | 0.833057 | 0.317 | 0.13 | 0.076003 |
| Fibroblasts | CCN1 | 0.005272 | 0.831743 | 0.633 | 0.553 | 1 |
| Pericytes | ITPK1 | 1.32E-12 | 0.831435 | 0.211 | 0.048 | 2.31E-08 |
| Fibroblasts | BICC1 | 1.1E-05 | 0.831125 | 0.333 | 0.148 | 0.19252 |
| Pericytes | SEMA6D | 5.96E-26 | 0.830528 | 0.167 | 0.015 | 1.04E-21 |
| Terminal | SHISA4 | 4.69E-40 | 0.830352 | 0.642 | 0.426 | 8.2E-36 |
| Pericytes | BAMBI | 0.000344 | 0.829677 | 0.278 | 0.167 | 1 |
| Terminal | SSR3 | 2.86E-60 | 0.828936 | 0.698 | 0.414 | 5E-56 |
| Terminal | HOPX | 5.32E-64 | 0.828848 | 0.479 | 0.184 | 9.3E-60 |
| Pericytes | NEK4 | 2.33E-12 | 0.828503 | 0.2 | 0.044 | 4.08E-08 |
| Contractile | NIBAN1 | 9.72E-79 | 0.827797 | 0.866 | 0.652 | 1.7E-74 |
| Pericytes | NFATC4 | 1.99E-10 | 0.827514 | 0.233 | 0.067 | 3.48E-06 |
| Pericytes | TBC1D1 | 1.04E-09 | 0.826591 | 0.278 | 0.097 | 1.81E-05 |
| Terminal | PLP2 | 3.75E-55 | 0.826155 | 0.827 | 0.669 | 6.56E-51 |
| Contractile | NET1 | 1.25E-81 | 0.82464 | 0.737 | 0.394 | 2.18E-77 |
| Pericytes | MAN1A1 | 4.49E-08 | 0.824431 | 0.3 | 0.125 | 0.000784 |
| Terminal | RFLNB | 3.06E-45 | 0.82389 | 0.413 | 0.171 | 5.35E-41 |
| IFN_Interm | AKAP7 | 0.001163 | 0.82388 | 0.211 | 0.05 | 1 |
| IFN_Interm | HSD17B12 | 0.005158 | 0.822944 | 0.421 | 0.199 | 1 |
| Contractile | LIMS2 | 4.33E-73 | 0.821935 | 0.644 | 0.353 | 7.56E-69 |
| Fibroblasts | PCDH18 | 3.66E-07 | 0.82033 | 0.267 | 0.087 | 0.006403 |
| Fibroblasts | PRELP | 0.000418 | 0.820137 | 0.8 | 0.741 | 1 |
| Pericytes | HSP90AB1 | 3.86E-09 | 0.82007 | 0.867 | 0.785 | 6.75E-05 |
| Pericytes | ARGLU1 | 1.03E-07 | 0.820045 | 0.678 | 0.552 | 0.001798 |
| Terminal | TIMP1 | 0.001643 | 0.819993 | 0.947 | 0.929 | 1 |
| Pericytes | RASA3 | 5.18E-05 | 0.819673 | 0.433 | 0.307 | 0.904647 |
| IFN_Interm | CASP5 | 0.000774 | 0.819644 | 0.474 | 0.193 | 1 |
| Fibroblasts | VIPR2 | 1E-44 | 0.818827 | 0.267 | 0.015 | 1.75E-40 |
| Contractile | GAS6 | 2.99E-82 | 0.81797 | 0.767 | 0.456 | 5.22E-78 |
| Contractile | C6H19ORF | 4.79E-65 | 0.816985 | 0.573 | 0.288 | 8.38E-61 |
| Terminal | C4ORF3 | 3.82E-55 | 0.816884 | 0.819 | 0.641 | 6.68E-51 |
| Contractile | PI15 | 7.88E-26 | 0.814698 | 0.143 | 0.01 | 1.38E-21 |

| | | | | | | |
|---|---|---|---|---|---|---|
| Fibroblasts | ABLIM1 | 5.44E-16 | 0.813349 | 0.317 | 0.06 | 9.5E-12 |
| Pericytes | PPA1 | 9.88E-05 | 0.813261 | 0.267 | 0.145 | 1 |
| Terminal | NBL1 | 4.31E-38 | 0.813059 | 0.723 | 0.493 | 7.53E-34 |
| Contractile | PCDH7 | 6.72E-70 | 0.813036 | 0.615 | 0.309 | 1.17E-65 |
| Fibroblasts | IER3 | 0.002873 | 0.811574 | 0.3 | 0.168 | 1 |
| IFN_Interm | DZIP3 | 1.38E-05 | 0.811493 | 0.158 | 0.019 | 0.241493 |
| Fibroblasts | TAGLN2 | 3.17E-05 | 0.811141 | 0.783 | 0.673 | 0.553533 |
| Contractile | CAVIN3 | 1.3E-117 | 0.81111 | 0.987 | 0.928 | 2.2E-113 |
| Contractile | LOC10015: | 5.25E-17 | 0.811035 | 0.297 | 0.069 | 9.19E-13 |
| Fibroblasts | GPC1 | 6.62E-14 | 0.810851 | 0.167 | 0.02 | 1.16E-09 |
| Pericytes | PELI1 | 1.37E-06 | 0.810604 | 0.244 | 0.101 | 0.023961 |
| Contractile | PPP1CB | 4.38E-08 | 0.809654 | 0.692 | 0.489 | 0.000766 |
| IFN_Interm | ST3GAL6 | 0.000535 | 0.808992 | 0.368 | 0.119 | 1 |
| IFN_Interm | SDC2 | 0.005529 | 0.808771 | 0.947 | 0.827 | 1 |
| Pericytes | MAT2A | 0.000724 | 0.807684 | 0.467 | 0.394 | 1 |
| Terminal | IL18 | 7.64E-61 | 0.807267 | 0.376 | 0.116 | 1.34E-56 |
| Pericytes | FGF7 | 3.07E-07 | 0.806998 | 0.278 | 0.11 | 0.00537 |
| Fibroblasts | QKI | 0.000165 | 0.806988 | 0.617 | 0.451 | 1 |
| Terminal | ECRG4 | 1.55E-24 | 0.806902 | 0.103 | 0.02 | 2.71E-20 |
| Pericytes | ENSSSCG0( | 0.002466 | 0.806195 | 0.344 | 0.252 | 1 |
| Contractile | FAM13C | 3.89E-76 | 0.805334 | 0.567 | 0.245 | 6.79E-72 |
| Terminal | PGK1 | 1.09E-50 | 0.804724 | 0.694 | 0.446 | 1.9E-46 |
| Intermedia | PDGFRL | 6.25E-55 | 0.804595 | 0.615 | 0.37 | 1.09E-50 |
| Contractile | ADARB1 | 6.99E-70 | 0.804384 | 0.594 | 0.286 | 1.22E-65 |
| Contractile | MSRB3 | 1.26E-61 | 0.804156 | 0.654 | 0.399 | 2.21E-57 |
| Contractile | ZYX | 1.31E-65 | 0.804126 | 0.568 | 0.292 | 2.3E-61 |
| Terminal | ANXA5 | 1.75E-89 | 0.803329 | 0.975 | 0.953 | 3.06E-85 |
| Pericytes | ETFB | 2.35E-05 | 0.802562 | 0.344 | 0.21 | 0.410819 |
| Contractile | ENSSSCG0( | 1.5E-67 | 0.802446 | 0.551 | 0.252 | 2.63E-63 |
| Pericytes | GGTA1 | 0.00121 | 0.801795 | 0.456 | 0.388 | 1 |
| Intermedia | ENSSSCG0( | 1.39E-47 | 0.801035 | 0.687 | 0.489 | 2.43E-43 |
| Pericytes | LRRC32 | 4.69E-05 | 0.800996 | 0.367 | 0.229 | 0.819213 |
| Pericytes | PSME1 | 3.44E-05 | 0.799979 | 0.5 | 0.369 | 0.601548 |
| Pericytes | TBX3 | 2.24E-09 | 0.799531 | 0.167 | 0.04 | 3.92E-05 |
| Contractile | FOXO3 | 1.73E-05 | 0.798668 | 0.637 | 0.472 | 0.303097 |
| Pericytes | ESAM | 2.35E-05 | 0.798519 | 0.3 | 0.156 | 0.410796 |
| Fibroblasts | ENSSSCG0( | 2.67E-05 | 0.798468 | 0.583 | 0.382 | 0.467099 |
| IFN_Interm | MAMDC2 | 0.003536 | 0.797704 | 0.263 | 0.083 | 1 |
| Fibroblasts | NFIX | 6.92E-05 | 0.796936 | 0.583 | 0.41 | 1 |
| Contractile | CAP1 | 1.55E-78 | 0.796777 | 0.766 | 0.514 | 2.7E-74 |
| Pericytes | MDK | 3.41E-10 | 0.796748 | 0.278 | 0.092 | 5.96E-06 |
| Contractile | ISG12(A) | 7.91E-64 | 0.796101 | 0.672 | 0.364 | 1.38E-59 |
| Contractile | RCAN2 | 2.19E-63 | 0.795856 | 0.702 | 0.448 | 3.83E-59 |
| Terminal | S100A4 | 5.51E-37 | 0.794657 | 0.955 | 0.86 | 9.63E-33 |
| Contractile | PHLDB2 | 6.35E-12 | 0.794478 | 0.648 | 0.323 | 1.11E-07 |
| IFN_Interm | GPX8 | 0.004421 | 0.794312 | 0.684 | 0.444 | 1 |
| Pericytes | BTBD10 | 5.31E-05 | 0.794175 | 0.244 | 0.122 | 0.928458 |
| Pericytes | F3 | 3.74E-10 | 0.793255 | 0.156 | 0.032 | 6.55E-06 |
| Pericytes | ODC1 | 1.91E-06 | 0.792752 | 0.233 | 0.094 | 0.033315 |
| IFN_Interm | ENSSSCG0( | 2.37E-07 | 0.79199 | 0.158 | 0.014 | 0.004151 |
| Intermedia | MARCKS | 1.92E-35 | 0.791184 | 0.521 | 0.312 | 3.36E-31 |
| Contractile | PLN | 4.4E-75 | 0.789344 | 0.405 | 0.122 | 7.69E-71 |
| Pericytes | HSBP1 | 3.03E-10 | 0.789216 | 0.822 | 0.717 | 5.31E-06 |
| Pericytes | TGFBR2 | 0.002854 | 0.787195 | 0.356 | 0.276 | 1 |
| Pericytes | DOCK11 | 9.85E-12 | 0.786821 | 0.167 | 0.033 | 1.72E-07 |
| IFN_Interm | SERINC3 | 0.005546 | 0.785532 | 0.684 | 0.465 | 1 |
| Contractile | TMSB4X | 3E-117 | 0.785014 | 0.999 | 0.994 | 5.3E-113 |
| Pericytes | ENSSSCG0( | 6.46E-06 | 0.784692 | 0.189 | 0.07 | 0.113011 |
| Fibroblasts | LVRN | 5.2E-102 | 0.784032 | 0.2 | 0.001 | 9.15E-98 |

| | | | | | | |
|---|---|---|---|---|---|---|
| Contractile | ROCK1 | 4.73E-12 | 0.783049 | 0.824 | 0.605 | 8.28E-08 |
| Terminal | COL14A1 | 4.83E-52 | 0.782412 | 0.831 | 0.62 | 8.44E-48 |
| Contractile | LRRFIP1 | 2.12E-12 | 0.78233 | 0.901 | 0.693 | 3.71E-08 |
| Contractile | ENSSSCG0( | 2.8E-78 | 0.781634 | 0.91 | 0.735 | 4.89E-74 |
| Contractile | ITGB1 | 3.4E-145 | 0.781585 | 0.998 | 0.971 | 6E-141 |
| Pericytes | ID3 | 0.003558 | 0.781285 | 0.744 | 0.721 | 1 |
| Terminal | CD59 | 2.64E-48 | 0.781276 | 0.492 | 0.22 | 4.62E-44 |
| Pericytes | ARHGEF17 | 2.25E-07 | 0.780808 | 0.467 | 0.272 | 0.003931 |
| Contractile | KCTD10 | 3.38E-10 | 0.780471 | 0.692 | 0.406 | 5.91E-06 |
| Contractile | WIF1 | 1.03E-84 | 0.779731 | 0.852 | 0.501 | 1.79E-80 |
| Pericytes | ADGRF5 | 3.77E-77 | 0.779644 | 0.156 | 0.002 | 6.6E-73 |
| Terminal | GPI | 7.05E-51 | 0.779502 | 0.571 | 0.288 | 1.23E-46 |
| Pericytes | DOCK6 | 4.59E-15 | 0.779171 | 0.233 | 0.049 | 8.02E-11 |
| Contractile | LYNX1 | 3.43E-68 | 0.778585 | 0.496 | 0.214 | 6E-64 |
| Pericytes | AIF1 | 1.7E-12 | 0.778215 | 0.2 | 0.043 | 2.96E-08 |
| Fibroblasts | PHACTR2 | 4.54E-09 | 0.777143 | 0.283 | 0.078 | 7.93E-05 |
| Contractile | DMD | 2.94E-68 | 0.776381 | 0.408 | 0.135 | 5.14E-64 |
| Contractile | IL17B | 1.7E-64 | 0.776343 | 0.68 | 0.402 | 2.97E-60 |
| Fibroblasts | ENSSSCG0( | 0.003189 | 0.775367 | 0.633 | 0.544 | 1 |
| Pericytes | ARHGAP42 | 3.11E-07 | 0.775155 | 0.3 | 0.132 | 0.005435 |
| Fibroblasts | KLF9 | 5.59E-06 | 0.774702 | 0.433 | 0.212 | 0.097725 |
| Fibroblasts | KLF5 | 5.03E-06 | 0.774304 | 0.133 | 0.03 | 0.087909 |
| Pericytes | LAMC1 | 0.00054 | 0.774093 | 0.689 | 0.635 | 1 |
| Pericytes | SH2B3 | 7.59E-15 | 0.773678 | 0.211 | 0.042 | 1.33E-10 |
| Pericytes | SPC24 | 1.79E-05 | 0.773511 | 0.278 | 0.135 | 0.312056 |
| Pericytes | CNN3 | 0.000103 | 0.772909 | 0.7 | 0.673 | 1 |
| Contractile | TLN1 | 5.59E-76 | 0.772199 | 0.83 | 0.6 | 9.77E-72 |
| IFN_Interm | USP18 | 0.000162 | 0.77208 | 0.211 | 0.041 | 1 |
| Fibroblasts | CYBRD1 | 1.87E-05 | 0.771796 | 0.7 | 0.559 | 0.326357 |
| IFN_Interm | XAF1 | 1.69E-09 | 0.771466 | 0.211 | 0.019 | 2.95E-05 |
| IFN_Interm | CCND2 | 0.007023 | 0.769881 | 0.632 | 0.38 | 1 |
| Intermedia | PLXDC2 | 5.24E-61 | 0.769753 | 0.847 | 0.656 | 9.16E-57 |
| Contractile | HSPG2 | 1.14E-85 | 0.768757 | 0.916 | 0.715 | 1.99E-81 |
| Pericytes | ENPEP | 9.58E-57 | 0.768646 | 0.122 | 0.002 | 1.67E-52 |
| Fibroblasts | CITED2 | 0.000343 | 0.76853 | 0.25 | 0.112 | 1 |
| Intermedia | CLEC11A | 6.91E-58 | 0.768321 | 0.769 | 0.524 | 1.21E-53 |
| Pericytes | PDE3A | 2.42E-05 | 0.767933 | 0.489 | 0.353 | 0.423349 |
| Pericytes | MAP3K20 | 4.07E-07 | 0.76695 | 0.356 | 0.176 | 0.007113 |
| Contractile | ENSSSCG0( | 8.67E-19 | 0.765672 | 0.341 | 0.08 | 1.52E-14 |
| Terminal | CGREF1 | 4.05E-38 | 0.764118 | 0.488 | 0.256 | 7.08E-34 |
| Pericytes | DEPTOR | 6.56E-28 | 0.763653 | 0.167 | 0.013 | 1.15E-23 |
| Terminal | LMNA | 1.45E-39 | 0.762973 | 0.892 | 0.846 | 2.54E-35 |
| Fibroblasts | ADA | 3.02E-06 | 0.762176 | 0.183 | 0.05 | 0.05274 |
| Pericytes | IFNAR2 | 1.11E-08 | 0.761904 | 0.2 | 0.059 | 0.000194 |
| Intermedia | FNDC1 | 5.31E-49 | 0.761693 | 0.839 | 0.66 | 9.28E-45 |
| Terminal | IGFBP6 | 3.26E-42 | 0.761213 | 0.794 | 0.61 | 5.71E-38 |
| Contractile | DPYSL3 | 2.86E-44 | 0.760682 | 0.56 | 0.332 | 5.01E-40 |
| Pericytes | CCDC9B | 1.37E-07 | 0.759968 | 0.256 | 0.099 | 0.002397 |
| Contractile | VLDLR | 5.62E-65 | 0.759177 | 0.726 | 0.455 | 9.82E-61 |
| Contractile | FGL2 | 2.39E-68 | 0.758859 | 0.667 | 0.348 | 4.18E-64 |
| Pericytes | KANK2 | 2.77E-07 | 0.758531 | 0.633 | 0.481 | 0.004845 |
| Contractile | ENSSSCG0( | 5.8E-14 | 0.758386 | 0.319 | 0.09 | 1.01E-09 |
| Contractile | RRAGD | 1.09E-11 | 0.75827 | 0.275 | 0.08 | 1.9E-07 |
| IFN_Interm | PLEKHA4 | 0.005054 | 0.75697 | 0.158 | 0.038 | 1 |
| Pericytes | TP53I11 | 3.2E-06 | 0.756112 | 0.244 | 0.105 | 0.055947 |
| Pericytes | MEF2C | 0.000148 | 0.755314 | 0.544 | 0.464 | 1 |
| Contractile | MCAM | 6.62E-13 | 0.755099 | 0.571 | 0.243 | 1.16E-08 |
| Contractile | WDR1 | 1.85E-70 | 0.754753 | 0.717 | 0.45 | 3.24E-66 |
| Fibroblasts | FOLR2 | 0.000232 | 0.754569 | 0.6 | 0.482 | 1 |

| | | | | | | |
|---|---|---|---|---|---|---|
| Fibroblasts | RCAN3 | 3.07E-05 | 0.754188 | 0.167 | 0.05 | 0.536832 |
| Contractile | FBXO32 | 4.84E-60 | 0.753902 | 0.59 | 0.3 | 8.45E-56 |
| Intermedia | MXRA8 | 1.11E-44 | 0.752902 | 0.793 | 0.651 | 1.95E-40 |
| IFN_Interm | ABTB2 | 7.43E-05 | 0.752758 | 0.158 | 0.023 | 1 |
| Fibroblasts | PA2G4 | 0.006923 | 0.752039 | 0.583 | 0.48 | 1 |
| Pericytes | TXNIP | 4.24E-06 | 0.751899 | 0.678 | 0.537 | 0.074183 |
| Pericytes | TNFRSF1A | 0.001587 | 0.751886 | 0.344 | 0.251 | 1 |
| Intermedia | COL1A2 | 1.64E-66 | 0.751697 | 0.996 | 0.977 | 2.86E-62 |
| Contractile | MSRB3 | 6.15E-09 | 0.751601 | 0.725 | 0.494 | 0.000107 |
| Pericytes | TESC | 1.02E-79 | 0.751459 | 0.2 | 0.004 | 1.79E-75 |
| Contractile | FBLIM1 | 1.28E-07 | 0.750767 | 0.604 | 0.388 | 0.002237 |
| Pericytes | REL | 4.27E-05 | 0.750696 | 0.244 | 0.118 | 0.746469 |
| Contractile | RASL12 | 3.76E-23 | 0.750074 | 0.308 | 0.055 | 6.58E-19 |
| Pericytes | UACA | 0.004797 | 0.749201 | 0.422 | 0.359 | 1 |
| IFN_Interm | ZBP1 | 3.92E-19 | 0.74832 | 0.105 | 0.002 | 6.85E-15 |
| Contractile | WFDC1 | 7.63E-80 | 0.747856 | 0.89 | 0.669 | 1.33E-75 |
| Intermedia | OLFML3 | 1.07E-46 | 0.747732 | 0.491 | 0.255 | 1.87E-42 |
| Pericytes | ENSSSCG0( | 2.03E-18 | 0.74735 | 0.156 | 0.019 | 3.54E-14 |
| Contractile | PDLIM5 | 4.25E-67 | 0.746939 | 0.672 | 0.382 | 7.43E-63 |
| Fibroblasts | GOLIM4 | 0.000434 | 0.74677 | 0.733 | 0.643 | 1 |
| Contractile | ITGA1 | 1.71E-70 | 0.746286 | 0.842 | 0.589 | 2.98E-66 |
| Pericytes | NDRG2 | 0.000109 | 0.74615 | 0.311 | 0.18 | 1 |
| Contractile | CFL2 | 6.44E-56 | 0.746064 | 0.59 | 0.332 | 1.13E-51 |
| Pericytes | ENSSSCG0( | 4.35E-07 | 0.745656 | 0.211 | 0.074 | 0.007608 |
| Fibroblasts | ARRDC3 | 2.03E-07 | 0.745397 | 0.35 | 0.13 | 0.003553 |
| Fibroblasts | ENSSSCG0( | 0.000559 | 0.744669 | 0.75 | 0.673 | 1 |
| Fibroblasts | DDX39B | 0.0014 | 0.744551 | 0.4 | 0.261 | 1 |
| Pericytes | CTSC | 0.000402 | 0.744437 | 0.456 | 0.324 | 1 |
| Fibroblasts | SFRP1 | 0.002347 | 0.744179 | 0.617 | 0.456 | 1 |
| Contractile | ENSSSCG0( | 2.87E-26 | 0.743819 | 0.22 | 0.025 | 5.02E-22 |
| Fibroblasts | TCF4 | 0.000444 | 0.743092 | 0.433 | 0.272 | 1 |
| Pericytes | SIX1 | 1.39E-05 | 0.742974 | 0.256 | 0.12 | 0.242844 |
| Fibroblasts | CCPG1 | 3.62E-06 | 0.742425 | 0.817 | 0.633 | 0.063317 |
| Pericytes | PDE1A | 1.09E-06 | 0.74123 | 0.333 | 0.169 | 0.018993 |
| Contractile | ISG12(A) | 7.79E-07 | 0.74095 | 0.67 | 0.482 | 0.013623 |
| Contractile | ENSSSCG0( | 8.09E-06 | 0.740307 | 0.385 | 0.21 | 0.14142 |
| Contractile | SLIT3 | 1.62E-08 | 0.739154 | 0.835 | 0.655 | 0.000283 |
| Fibroblasts | SPOCK1 | 4.55E-08 | 0.73915 | 0.233 | 0.062 | 0.000796 |
| Contractile | AKAP12 | 1.05E-08 | 0.73878 | 0.758 | 0.521 | 0.000184 |
| Contractile | FAM13C | 9.03E-11 | 0.738527 | 0.67 | 0.365 | 1.58E-06 |
| Pericytes | JAK1 | 5.14E-06 | 0.738508 | 0.7 | 0.638 | 0.089934 |
| IFN_Interm | ENPP4 | 1.36E-11 | 0.737957 | 0.263 | 0.023 | 2.38E-07 |
| Terminal | TAGLN2 | 2.45E-47 | 0.737566 | 0.797 | 0.638 | 4.29E-43 |
| Contractile | RGS7BP | 1.3E-114 | 0.737055 | 0.242 | 0.003 | 2.2E-110 |
| Fibroblasts | DNAJB9 | 1.39E-06 | 0.737021 | 0.433 | 0.203 | 0.024262 |
| Pericytes | PNISR | 0.000129 | 0.736905 | 0.622 | 0.577 | 1 |
| Fibroblasts | SPRY1 | 0.000393 | 0.73682 | 0.367 | 0.199 | 1 |
| Pericytes | BTG1 | 3.93E-08 | 0.736793 | 0.3 | 0.125 | 0.000688 |
| Contractile | L1CAM | 4.56E-64 | 0.736139 | 0.348 | 0.103 | 7.97E-60 |
| Pericytes | PSPC1 | 4.96E-05 | 0.735658 | 0.356 | 0.21 | 0.867664 |
| Contractile | SH3BGRL | 6.78E-10 | 0.735552 | 0.736 | 0.518 | 1.19E-05 |
| IFN_Interm | ETV1 | 0.002546 | 0.735432 | 0.421 | 0.175 | 1 |
| Pericytes | MSI2 | 3.7E-06 | 0.735416 | 0.256 | 0.115 | 0.06461 |
| Pericytes | SEMA5A | 1.57E-60 | 0.734776 | 0.156 | 0.003 | 2.74E-56 |
| Intermedia | POSTN | 1.73E-67 | 0.734593 | 0.988 | 0.895 | 3.02E-63 |
| Pericytes | OSMR | 0.000243 | 0.734243 | 0.356 | 0.234 | 1 |
| Pericytes | ENSSSCG0( | 2.11E-15 | 0.733808 | 0.111 | 0.011 | 3.7E-11 |
| Contractile | NDRG4 | 3.77E-15 | 0.733214 | 0.165 | 0.025 | 6.6E-11 |
| Contractile | LRCH1 | 1.15E-08 | 0.73305 | 0.407 | 0.187 | 0.000201 |

| | | | | | | |
|---|---|---|---|---|---|---|
| IFN_Interm | PLEKHA7 | 7.62E-26 | 0.732643 | 0.158 | 0.003 | 1.33E-21 |
| Pericytes | ENSSSCG0( | 3.2E-63 | 0.732459 | 0.156 | 0.003 | 5.59E-59 |
| Pericytes | SEPTIN7 | 9.68E-14 | 0.731866 | 0.978 | 0.933 | 1.69E-09 |
| Terminal | NMB | 8.43E-50 | 0.731501 | 0.209 | 0.039 | 1.47E-45 |
| Pericytes | PTK2 | 6E-07 | 0.731343 | 0.322 | 0.152 | 0.010488 |
| Contractile | AKAP6 | 4.87E-10 | 0.730711 | 0.319 | 0.113 | 8.51E-06 |
| Pericytes | CACNA2D1 | 0.002889 | 0.730576 | 0.389 | 0.306 | 1 |
| Pericytes | ENSSSCG0( | 4.82E-16 | 0.730409 | 0.189 | 0.031 | 8.43E-12 |
| Terminal | KLF4 | 7.7E-52 | 0.730195 | 0.451 | 0.177 | 1.35E-47 |
| IFN_Interm | TRAF5 | 0.006183 | 0.729313 | 0.211 | 0.064 | 1 |
| Contractile | PLXDC1 | 1.9E-07 | 0.729267 | 0.604 | 0.392 | 0.003313 |
| Contractile | PDZRN4 | 3.07E-80 | 0.729137 | 0.348 | 0.075 | 5.36E-76 |
| Contractile | KCNA5 | 2.13E-10 | 0.728599 | 0.374 | 0.142 | 3.72E-06 |
| Pericytes | AHI1 | 0.001798 | 0.728375 | 0.367 | 0.279 | 1 |
| Intermedia | TFPI2 | 9.9E-49 | 0.728328 | 0.664 | 0.434 | 1.73E-44 |
| Contractile | CDKN1B | 2.44E-08 | 0.728018 | 0.484 | 0.261 | 0.000427 |
| Contractile | RBPMS | 1.09E-66 | 0.727896 | 0.53 | 0.237 | 1.91E-62 |
| Pericytes | LOC10052( | 2.35E-05 | 0.726841 | 0.444 | 0.299 | 0.410438 |
| Terminal | PTPRR | 1.92E-65 | 0.726435 | 0.326 | 0.079 | 3.36E-61 |
| IFN_Interm | PSMF1 | 2.72E-05 | 0.725987 | 0.368 | 0.093 | 0.47472 |
| Intermedia | CDH11 | 5.25E-40 | 0.725381 | 0.738 | 0.556 | 9.18E-36 |
| Terminal | VDAC1 | 9.68E-42 | 0.724868 | 0.676 | 0.467 | 1.69E-37 |
| Pericytes | CD4 | 4.49E-80 | 0.724804 | 0.167 | 0.002 | 7.85E-76 |
| IFN_Interm | AP1S3 | 1.11E-13 | 0.724481 | 0.158 | 0.007 | 1.94E-09 |
| Pericytes | VPS36 | 0.000783 | 0.724343 | 0.5 | 0.416 | 1 |
| Pericytes | TTYH2 | 2.78E-12 | 0.723519 | 0.211 | 0.049 | 4.86E-08 |
| Pericytes | PLK2 | 6.18E-05 | 0.723453 | 0.144 | 0.052 | 1 |
| Intermedia | PRELP | 1.78E-38 | 0.723211 | 0.842 | 0.702 | 3.11E-34 |
| Pericytes | ADCY3 | 4.91E-14 | 0.722354 | 0.2 | 0.04 | 8.59E-10 |
| Pericytes | MAP3K7CL | 5.38E-07 | 0.722163 | 0.178 | 0.056 | 0.009397 |
| Pericytes | HOXA9 | 9.47E-05 | 0.721994 | 0.311 | 0.178 | 1 |
| Terminal | HOMER3 | 1.14E-45 | 0.72179 | 0.392 | 0.156 | 1.99E-41 |
| Contractile | EPAS1 | 4.56E-08 | 0.721712 | 0.791 | 0.645 | 0.000797 |
| Terminal | KDELR3 | 4.32E-34 | 0.721688 | 0.667 | 0.497 | 7.56E-30 |
| Intermedia | COL12A1 | 5.57E-34 | 0.721355 | 0.855 | 0.724 | 9.73E-30 |
| Pericytes | NCK2 | 3.19E-07 | 0.721025 | 0.256 | 0.102 | 0.005571 |
| Terminal | USP53 | 6.68E-24 | 0.72096 | 0.565 | 0.389 | 1.17E-19 |
| Contractile | TM4SF1 | 3.1E-09 | 0.720735 | 0.868 | 0.744 | 5.43E-05 |
| Fibroblasts | MCFD2 | 0.000981 | 0.720653 | 0.417 | 0.28 | 1 |
| Contractile | SMOC2 | 3.47E-56 | 0.720584 | 0.572 | 0.295 | 6.07E-52 |
| Fibroblasts | SLIT2 | 0.000177 | 0.720502 | 0.283 | 0.128 | 1 |
| Terminal | NME2 | 1.16E-66 | 0.720372 | 0.945 | 0.827 | 2.03E-62 |
| Contractile | TNS1 | 1.56E-61 | 0.720345 | 0.544 | 0.264 | 2.73E-57 |
| Terminal | YBX1 | 3.88E-58 | 0.719372 | 0.956 | 0.877 | 6.79E-54 |
| Contractile | VCL | 4.99E-13 | 0.718777 | 0.934 | 0.766 | 8.72E-09 |
| Fibroblasts | EML4 | 7.04E-05 | 0.717726 | 0.35 | 0.168 | 1 |
| Pericytes | NFE2L3 | 2.82E-06 | 0.717449 | 0.133 | 0.038 | 0.049358 |
| Pericytes | ENSSSCG0( | 1.66E-15 | 0.71703 | 0.2 | 0.036 | 2.91E-11 |
| Pericytes | ENSSSCG0( | 3.43E-17 | 0.716764 | 0.144 | 0.017 | 6E-13 |
| Contractile | LMCD1 | 7.9E-09 | 0.715876 | 0.593 | 0.339 | 0.000138 |
| Contractile | COL4A1 | 2.4E-108 | 0.71549 | 0.993 | 0.883 | 4.2E-104 |
| Pericytes | NAB1 | 3.39E-05 | 0.714793 | 0.356 | 0.21 | 0.593049 |
| Terminal | H2AZ2 | 1.63E-23 | 0.714373 | 0.723 | 0.577 | 2.84E-19 |
| Terminal | FAM167B | 1.38E-29 | 0.713423 | 0.398 | 0.205 | 2.41E-25 |
| Pericytes | ZFHX3 | 0.000331 | 0.71334 | 0.378 | 0.253 | 1 |
| Pericytes | EBF2 | 1.55E-12 | 0.713115 | 0.211 | 0.048 | 2.71E-08 |
| Contractile | SPINT2 | 2.87E-62 | 0.712778 | 0.557 | 0.262 | 5.01E-58 |
| Terminal | DPP4 | 1.19E-98 | 0.712453 | 0.228 | 0.012 | 2.09E-94 |
| Contractile | ANGPTL1 | 2.47E-19 | 0.711793 | 0.198 | 0.028 | 4.33E-15 |

| | | | | | | |
|---|---|---|---|---|---|---|
| Fibroblasts | SNED1 | 8.6E-14 | 0.711095 | 0.133 | 0.013 | 1.5E-09 |
| Fibroblasts | BEX1 | 1.65E-22 | 0.710838 | 0.15 | 0.01 | 2.89E-18 |
| Terminal | NPC2 | 8.64E-39 | 0.71077 | 0.709 | 0.51 | 1.51E-34 |
| Pericytes | CD248 | 0.000229 | 0.710531 | 0.244 | 0.128 | 1 |
| Fibroblasts | SPOCK2 | 3.1E-53 | 0.709793 | 0.217 | 0.007 | 5.42E-49 |
| Pericytes | FILIP1L | 1.32E-07 | 0.709715 | 0.644 | 0.46 | 0.0023 |
| Fibroblasts | SLC39A14 | 0.006508 | 0.709138 | 0.317 | 0.204 | 1 |
| Contractile | PDK1 | 4.67E-07 | 0.709073 | 0.516 | 0.307 | 0.008162 |
| Pericytes | PEMT | 1.33E-05 | 0.708778 | 0.2 | 0.081 | 0.232239 |
| Pericytes | PDE4B | 0.000703 | 0.707901 | 0.167 | 0.077 | 1 |
| IFN_Interm | MAGI1 | 0.003815 | 0.707326 | 0.211 | 0.059 | 1 |
| Contractile | ACTG2 | 4.41E-11 | 0.706372 | 0.374 | 0.13 | 7.72E-07 |
| Fibroblasts | PIGK | 6.44E-05 | 0.706363 | 0.367 | 0.184 | 1 |
| Pericytes | IRAG1 | 0.00478 | 0.706355 | 0.456 | 0.368 | 1 |
| IFN_Interm | TRIM8 | 0.008339 | 0.706138 | 0.263 | 0.095 | 1 |
| Contractile | SVIL | 1.33E-07 | 0.705562 | 0.527 | 0.296 | 0.002317 |
| Contractile | DMD | 9.64E-11 | 0.705548 | 0.516 | 0.236 | 1.69E-06 |
| IFN_Interm | CDC42SE1 | 0.005266 | 0.705418 | 0.316 | 0.119 | 1 |
| Pericytes | BCL3 | 6.05E-05 | 0.705279 | 0.2 | 0.089 | 1 |
| Pericytes | JARID2 | 7.26E-05 | 0.704846 | 0.122 | 0.04 | 1 |
| Pericytes | H3-3A | 4.06E-13 | 0.704585 | 0.989 | 0.928 | 7.1E-09 |
| IFN_Interm | PLXDC2 | 0.005839 | 0.70437 | 0.947 | 0.709 | 1 |
| Contractile | ENSSSCG0( | 1.48E-07 | 0.70417 | 0.637 | 0.415 | 0.002592 |
| Terminal | OSTC | 2.31E-35 | 0.703379 | 0.584 | 0.374 | 4.04E-31 |
| Pericytes | PHLDB2 | 0.000527 | 0.703267 | 0.456 | 0.329 | 1 |
| Contractile | ITGA1 | 1.07E-11 | 0.703043 | 0.901 | 0.684 | 1.87E-07 |
| Pericytes | ENSSSCG0( | 5.75E-09 | 0.702546 | 0.2 | 0.057 | 0.000101 |
| IFN_Interm | BNC2 | 3.76E-06 | 0.7019 | 0.263 | 0.044 | 0.065724 |
| Pericytes | RAP1A | 0.00054 | 0.701805 | 0.544 | 0.485 | 1 |
| Contractile | PALLD | 7.47E-09 | 0.70158 | 0.714 | 0.457 | 0.000131 |
| Pericytes | PSMB8 | 0.00015 | 0.701542 | 0.267 | 0.143 | 1 |
| Terminal | VMP1 | 3.93E-44 | 0.701399 | 0.857 | 0.663 | 6.86E-40 |
| Pericytes | SUB1 | 0.000753 | 0.70128 | 0.533 | 0.497 | 1 |
| Terminal | PHGDH | 2.22E-76 | 0.700996 | 0.207 | 0.019 | 3.88E-72 |
| Intermedia | FOLR2 | 9.63E-38 | 0.700801 | 0.613 | 0.433 | 1.68E-33 |
| Terminal | IVNS1ABP | 8.36E-37 | 0.70075 | 0.362 | 0.15 | 1.46E-32 |
| Pericytes | ABCC4 | 1.7E-69 | 0.700632 | 0.167 | 0.003 | 2.97E-65 |
| Contractile | LOC100520 | 3.78E-08 | 0.700447 | 0.527 | 0.297 | 0.000661 |
| Terminal | LMAN1 | 1.07E-42 | 0.700189 | 0.636 | 0.396 | 1.86E-38 |
| Fibroblasts | NFIA | 0.000122 | 0.698182 | 0.55 | 0.332 | 1 |
| Pericytes | CCND1 | 0.001869 | 0.697718 | 0.589 | 0.599 | 1 |
| Fibroblasts | ENSSSCG0( | 2.38E-09 | 0.697703 | 1 | 0.996 | 4.16E-05 |
| Contractile | ITGAV | 4.55E-60 | 0.697639 | 0.8 | 0.596 | 7.96E-56 |
| Pericytes | PRMT1 | 0.000464 | 0.696526 | 0.422 | 0.328 | 1 |
| Contractile | TGFB3 | 2.26E-24 | 0.696379 | 0.344 | 0.197 | 3.95E-20 |
| Pericytes | THSD7A | 5.64E-08 | 0.695888 | 0.2 | 0.061 | 0.000986 |
| Pericytes | PRR13 | 0.001991 | 0.695851 | 0.411 | 0.332 | 1 |
| Pericytes | ATP8B1 | 1.6E-05 | 0.69558 | 0.267 | 0.13 | 0.279023 |
| Pericytes | ENSSSCG0( | 3.27E-14 | 0.695572 | 0.189 | 0.035 | 5.72E-10 |
| Fibroblasts | TLE1 | 8.89E-06 | 0.695348 | 0.283 | 0.111 | 0.155429 |
| Terminal | CDO1 | 4.04E-21 | 0.695201 | 0.653 | 0.476 | 7.06E-17 |
| Fibroblasts | CTSV | 8.44E-06 | 0.694609 | 0.817 | 0.725 | 0.147477 |
| Terminal | IL33 | 1.19E-66 | 0.694424 | 0.185 | 0.017 | 2.09E-62 |
| Terminal | PYCR1 | 1.56E-38 | 0.693455 | 0.433 | 0.207 | 2.72E-34 |
| Pericytes | ENSSSCG0( | 6.13E-15 | 0.693191 | 0.122 | 0.014 | 1.07E-10 |
| Contractile | GASK1B | 1.8E-09 | 0.693054 | 0.736 | 0.473 | 3.15E-05 |
| Contractile | PDE4DIP | 8.31E-06 | 0.693027 | 0.396 | 0.219 | 0.145271 |
| Terminal | PRDX1 | 7.18E-32 | 0.692252 | 0.738 | 0.569 | 1.26E-27 |
| Pericytes | CHD3 | 7.46E-06 | 0.692092 | 0.367 | 0.203 | 0.130479 |

| | | | | | | |
|---|---|---|---|---|---|---|
| Pericytes | FAM76A | 0.000172 | 0.691992 | 0.356 | 0.233 | 1 |
| Contractile | ADCY5 | 3.43E-10 | 0.691814 | 0.352 | 0.132 | 5.99E-06 |
| Pericytes | ARPC1B | 3.87E-05 | 0.6914 | 0.633 | 0.534 | 0.675783 |
| Contractile | NPR1 | 3.17E-08 | 0.690971 | 0.385 | 0.172 | 0.000555 |
| Contractile | FBLN5 | 6.4E-07 | 0.689759 | 0.571 | 0.373 | 0.011194 |
| Fibroblasts | NFIB | 0.000184 | 0.689552 | 0.567 | 0.39 | 1 |
| Terminal | ANPEP | 1.36E-21 | 0.689252 | 0.56 | 0.392 | 2.38E-17 |
| Pericytes | DGKZ | 2.67E-05 | 0.689083 | 0.167 | 0.063 | 0.466399 |
| Terminal | GJA1 | 5.02E-35 | 0.688454 | 0.543 | 0.308 | 8.77E-31 |
| Terminal | PPP1R14B | 8.34E-30 | 0.688141 | 0.624 | 0.44 | 1.46E-25 |
| Pericytes | DDX5 | 2.79E-10 | 0.6878 | 0.867 | 0.83 | 4.87E-06 |
| Pericytes | FBL | 0.000181 | 0.686771 | 0.344 | 0.225 | 1 |
| Contractile | FOXP2 | 4.8E-14 | 0.686042 | 0.286 | 0.074 | 8.4E-10 |
| Contractile | MTUS1 | 3.28E-08 | 0.685185 | 0.286 | 0.107 | 0.000573 |
| Contractile | CCDC50 | 1.25E-05 | 0.684936 | 0.604 | 0.431 | 0.217889 |
| Fibroblasts | CRYL1 | 0.000434 | 0.684316 | 0.383 | 0.224 | 1 |
| Pericytes | SMARCC1 | 5.18E-05 | 0.683912 | 0.256 | 0.13 | 0.905612 |
| Terminal | PAPSS1 | 5.08E-39 | 0.683761 | 0.612 | 0.377 | 8.87E-35 |
| Contractile | HMG20B | 1.24E-47 | 0.683051 | 0.495 | 0.262 | 2.17E-43 |
| Pericytes | LDB2 | 8.55E-31 | 0.68186 | 0.144 | 0.008 | 1.49E-26 |
| IFN_Interm | RNASEL | 4.3E-13 | 0.681672 | 0.211 | 0.013 | 7.52E-09 |
| Fibroblasts | LARP7 | 0.008414 | 0.681533 | 0.383 | 0.263 | 1 |
| Pericytes | SNTB1 | 1.97E-67 | 0.680493 | 0.178 | 0.004 | 3.44E-63 |
| Terminal | ARF4 | 1.47E-34 | 0.679771 | 0.771 | 0.599 | 2.56E-30 |
| Fibroblasts | PPA1 | 0.00366 | 0.679581 | 0.267 | 0.146 | 1 |
| Pericytes | PTPN1 | 0.001626 | 0.679203 | 0.267 | 0.165 | 1 |
| Intermedia | OLFML2B | 6.59E-50 | 0.67885 | 0.27 | 0.075 | 1.15E-45 |
| Pericytes | GSTO1.1 | 0.000195 | 0.678841 | 0.4 | 0.285 | 1 |
| Contractile | MYL12A | 4.82E-86 | 0.678758 | 0.95 | 0.882 | 8.42E-82 |
| Contractile | THRA | 3.35E-05 | 0.67866 | 0.407 | 0.254 | 0.585327 |
| Pericytes | ENSSSCG0( | 8.21E-07 | 0.678605 | 0.478 | 0.299 | 0.014346 |
| Fibroblasts | ASAH1 | 0.00069 | 0.678535 | 0.383 | 0.236 | 1 |
| IFN_Interm | AATF | 1.97E-05 | 0.678531 | 0.316 | 0.069 | 0.343793 |
| Pericytes | ARRDC2 | 0.003788 | 0.678052 | 0.233 | 0.144 | 1 |
| Intermedia | CTSH | 1.15E-34 | 0.677656 | 0.697 | 0.519 | 2E-30 |
| Terminal | TYMS | 1.19E-65 | 0.677482 | 0.209 | 0.026 | 2.08E-61 |
| Contractile | MAP3K20 | 5.82E-10 | 0.676905 | 0.418 | 0.174 | 1.02E-05 |
| Contractile | HSPB7 | 6.19E-06 | 0.676758 | 0.505 | 0.311 | 0.108281 |
| Pericytes | ENSSSCG0( | 8.17E-10 | 0.676573 | 0.189 | 0.047 | 1.43E-05 |
| Contractile | TLN1 | 1.79E-08 | 0.676467 | 0.846 | 0.687 | 0.000314 |
| Contractile | YPEL3 | 1.03E-05 | 0.676342 | 0.593 | 0.411 | 0.18088 |
| Pericytes | CREM | 0.000177 | 0.676097 | 0.233 | 0.119 | 1 |
| Fibroblasts | IGFBP4 | 5.58E-05 | 0.674842 | 0.183 | 0.059 | 0.975842 |
| Pericytes | UBB | 9.35E-09 | 0.674092 | 0.978 | 0.96 | 0.000163 |
| Fibroblasts | TIMP3 | 2.03E-07 | 0.673647 | 0.8 | 0.5 | 0.003548 |
| Terminal | POLR2L | 2.08E-26 | 0.673351 | 0.634 | 0.46 | 3.63E-22 |
| Contractile | WWP2 | 2.89E-05 | 0.672833 | 0.429 | 0.266 | 0.504698 |
| Pericytes | ENSSSCG0( | 3.35E-63 | 0.672434 | 0.156 | 0.003 | 5.86E-59 |
| Contractile | MYH9 | 1.46E-69 | 0.672356 | 0.953 | 0.835 | 2.54E-65 |
| Pericytes | FXYD6 | 0.001858 | 0.672016 | 0.4 | 0.301 | 1 |
| Intermedia | FAP | 2.56E-49 | 0.671607 | 0.861 | 0.719 | 4.47E-45 |
| Fibroblasts | PID1 | 5.77E-39 | 0.671533 | 0.217 | 0.011 | 1.01E-34 |
| Contractile | SLA-DQB1 | 7.53E-38 | 0.671484 | 0.362 | 0.164 | 1.32E-33 |
| Contractile | PHLDB2 | 3.66E-57 | 0.671113 | 0.496 | 0.223 | 6.4E-53 |
| Contractile | SLC22A3 | 4.49E-12 | 0.670602 | 0.231 | 0.058 | 7.85E-08 |
| Pericytes | UGCG | 4.49E-08 | 0.67053 | 0.222 | 0.074 | 0.000784 |
| Terminal | PRSS35 | 5.08E-41 | 0.670465 | 0.316 | 0.106 | 8.88E-37 |
| Fibroblasts | CREG1 | 0.000143 | 0.670296 | 0.5 | 0.315 | 1 |
| Terminal | NDFIP1 | 8.84E-42 | 0.670181 | 0.737 | 0.527 | 1.55E-37 |

| | | | | | | |
|---|---|---|---|---|---|---|
| Terminal | LOXL3 | 3.52E-31 | 0.669586 | 0.281 | 0.11 | 6.15E-27 |
| Fibroblasts | CAT | 0.005536 | 0.669516 | 0.333 | 0.203 | 1 |
| Contractile | ENSSSCG0( | 5.37E-68 | 0.66946 | 0.908 | 0.757 | 9.39E-64 |
| IFN_Interm | MORC3 | 0.004427 | 0.669318 | 0.316 | 0.116 | 1 |
| Pericytes | ENSSSCG0( | 0.001632 | 0.66878 | 0.356 | 0.252 | 1 |
| Contractile | KCNMB1 | 7.36E-14 | 0.66805 | 0.363 | 0.111 | 1.29E-09 |
| Fibroblasts | FBLN2 | 0.009222 | 0.667151 | 0.433 | 0.303 | 1 |
| Pericytes | MYLIP | 0.000626 | 0.666791 | 0.322 | 0.207 | 1 |
| Pericytes | CPM | 5.09E-24 | 0.666724 | 0.133 | 0.01 | 8.89E-20 |
| Contractile | ROCK2 | 3.56E-06 | 0.666652 | 0.736 | 0.576 | 0.062167 |
| Terminal | FTL | 9.19E-39 | 0.666016 | 0.993 | 0.974 | 1.61E-34 |
| Contractile | FBLN5 | 2.3E-53 | 0.665942 | 0.537 | 0.273 | 4.01E-49 |
| Fibroblasts | ECRG4 | 1.49E-12 | 0.665854 | 0.217 | 0.036 | 2.6E-08 |
| Intermedia | F5 | 2.68E-21 | 0.665189 | 0.57 | 0.435 | 4.68E-17 |
| Pericytes | GALNT16 | 3.87E-15 | 0.664832 | 0.167 | 0.026 | 6.77E-11 |
| Contractile | COL13A1 | 1.5E-66 | 0.664464 | 0.302 | 0.069 | 2.62E-62 |
| Pericytes | TMEM131 | 0.000114 | 0.663378 | 0.222 | 0.105 | 1 |
| Terminal | GARS1 | 2.14E-35 | 0.663337 | 0.473 | 0.252 | 3.74E-31 |
| Fibroblasts | ZPR1 | 0.000957 | 0.663262 | 0.25 | 0.122 | 1 |
| Pericytes | KLHL5 | 1.08E-07 | 0.663056 | 0.167 | 0.047 | 0.001881 |
| Terminal | FNDC1 | 2.17E-31 | 0.662348 | 0.801 | 0.684 | 3.79E-27 |
| IFN_Interm | TULP3 | 0.001756 | 0.662079 | 0.368 | 0.13 | 1 |
| Fibroblasts | CTSH | 0.000377 | 0.662048 | 0.75 | 0.566 | 1 |
| IFN_Interm | AP3M1 | 0.006516 | 0.660801 | 0.316 | 0.122 | 1 |
| Pericytes | ARHGAP6 | 1.81E-06 | 0.660714 | 0.2 | 0.073 | 0.031584 |
| Contractile | HOXA10 | 1.68E-06 | 0.660562 | 0.374 | 0.187 | 0.029302 |
| Contractile | LRRFIP1 | 8.31E-66 | 0.659894 | 0.828 | 0.612 | 1.45E-61 |
| Contractile | CALML4 | 2.02E-43 | 0.659617 | 0.517 | 0.288 | 3.53E-39 |
| Fibroblasts | CILK1 | 4.85E-06 | 0.659229 | 0.4 | 0.184 | 0.084696 |
| Pericytes | PRDX2 | 1.53E-05 | 0.659162 | 0.722 | 0.65 | 0.268038 |
| Terminal | ANKH | 2.78E-26 | 0.659062 | 0.342 | 0.164 | 4.85E-22 |
| Intermedia | COL5A1 | 5.32E-44 | 0.658875 | 0.867 | 0.718 | 9.3E-40 |
| Pericytes | CEP85L | 5.23E-07 | 0.65872 | 0.256 | 0.104 | 0.009145 |
| IFN_Interm | CNN3 | 0.004155 | 0.658622 | 0.895 | 0.673 | 1 |
| Pericytes | ENSSSCG0( | 9.97E-05 | 0.658455 | 0.322 | 0.19 | 1 |
| Fibroblasts | TVP23B | 0.000372 | 0.657933 | 0.583 | 0.434 | 1 |
| Terminal | CLIC1.1 | 3.76E-24 | 0.657124 | 0.749 | 0.646 | 6.57E-20 |
| Contractile | ATP2A2 | 0.002757 | 0.656932 | 0.462 | 0.363 | 1 |
| Contractile | AEBP1 | 3.07E-12 | 0.656846 | 1 | 0.873 | 5.37E-08 |
| Pericytes | S1PR5 | 1.69E-23 | 0.656419 | 0.144 | 0.012 | 2.95E-19 |
| Contractile | COL4A2 | 5.46E-81 | 0.655873 | 0.95 | 0.776 | 9.54E-77 |
| Pericytes | H1-3 | 2.68E-05 | 0.655651 | 0.178 | 0.069 | 0.468268 |
| Contractile | IL17B | 0.000252 | 0.655225 | 0.637 | 0.51 | 1 |
| Pericytes | PPP2R2A | 6.67E-05 | 0.654928 | 0.389 | 0.26 | 1 |
| Fibroblasts | CPXM2 | 0.002361 | 0.654787 | 0.817 | 0.756 | 1 |
| Fibroblasts | BCLAF1 | 0.001207 | 0.653902 | 0.533 | 0.376 | 1 |
| Fibroblasts | APOA1 | 0.002888 | 0.652212 | 0.083 | 0.255 | 1 |
| Contractile | ID4 | 1.01E-53 | 0.650801 | 0.396 | 0.154 | 1.76E-49 |
| Contractile | ANXA6 | 6.52E-08 | 0.650769 | 0.648 | 0.418 | 0.001139 |
| Contractile | NOTCH3 | 2.65E-66 | 0.650623 | 0.631 | 0.318 | 4.62E-62 |
| Pericytes | AGTR1 | 1.17E-14 | 0.650582 | 0.133 | 0.017 | 2.05E-10 |
| Terminal | SELENOM | 2.32E-41 | 0.64983 | 0.841 | 0.693 | 4.06E-37 |
| IFN_Interm | TRMT1 | 0.009864 | 0.649747 | 0.158 | 0.042 | 1 |
| Terminal | EEF1G | 2.97E-39 | 0.649726 | 0.826 | 0.694 | 5.19E-35 |
| Pericytes | PDCD4 | 8.36E-05 | 0.649494 | 0.444 | 0.306 | 1 |
| Contractile | ADGRE5 | 6.49E-06 | 0.649257 | 0.626 | 0.42 | 0.113477 |
| Pericytes | B2M | 2.98E-08 | 0.649108 | 0.944 | 0.861 | 0.000521 |
| Contractile | BMPR1A | 5.86E-08 | 0.648279 | 0.516 | 0.294 | 0.001024 |
| Contractile | B2M | 4.29E-80 | 0.647849 | 0.939 | 0.813 | 7.49E-76 |

| | | | | | | |
|---|---|---|---|---|---|---|
| IFN_Interm | PDIA4 | 0.00466 | 0.647387 | 0.737 | 0.411 | 1 |
| IFN_Interm | FAM76A | 0.007658 | 0.647158 | 0.474 | 0.235 | 1 |
| Contractile | GYG2 | 1.86E-07 | 0.647081 | 0.374 | 0.179 | 0.003245 |
| Pericytes | ENSSSCG0( | 3.92E-05 | 0.646989 | 0.333 | 0.192 | 0.68575 |
| IFN_Interm | INTS6 | 0.005691 | 0.646906 | 0.263 | 0.09 | 1 |
| Pericytes | TRPV2 | 2.09E-48 | 0.646019 | 0.133 | 0.003 | 3.65E-44 |
| Contractile | ILK | 1.27E-47 | 0.645797 | 0.706 | 0.508 | 2.21E-43 |
| Terminal | METRNL | 4.86E-40 | 0.645201 | 0.472 | 0.226 | 8.5E-36 |
| Contractile | RASL11A | 2.47E-58 | 0.6451 | 0.536 | 0.255 | 4.32E-54 |
| Pericytes | AIMP1 | 2.64E-05 | 0.645038 | 0.633 | 0.578 | 0.461192 |
| Pericytes | CDH6 | 1.39E-79 | 0.644612 | 0.144 | 0.001 | 2.43E-75 |
| Intermedia | ENSSSCG0( | 9.18E-35 | 0.644566 | 0.781 | 0.633 | 1.6E-30 |
| Pericytes | ENSSSCG0( | 2.77E-16 | 0.6444 | 0.167 | 0.024 | 4.84E-12 |
| Pericytes | MED13 | 0.005038 | 0.644172 | 0.278 | 0.195 | 1 |
| Pericytes | SOX13 | 1.07E-17 | 0.643238 | 0.156 | 0.019 | 1.87E-13 |
| Pericytes | MOB3B | 9.96E-19 | 0.643139 | 0.144 | 0.016 | 1.74E-14 |
| Contractile | ITGA5 | 0.00114 | 0.642793 | 0.549 | 0.413 | 1 |
| Contractile | DYRK3 | 1.2E-54 | 0.64266 | 0.426 | 0.174 | 2.09E-50 |
| Pericytes | EHD2 | 0.001074 | 0.641621 | 0.211 | 0.114 | 1 |
| Intermedia | TPST1 | 2.82E-28 | 0.641541 | 0.434 | 0.259 | 4.94E-24 |
| Contractile | KCNMB1 | 1.04E-74 | 0.641455 | 0.248 | 0.032 | 1.83E-70 |
| Pericytes | TIPARP | 0.000135 | 0.641225 | 0.156 | 0.061 | 1 |
| Intermedia | CHI3L1 | 7.85E-09 | 0.640988 | 0.532 | 0.439 | 0.000137 |
| Fibroblasts | LOX | 0.000292 | 0.64079 | 0.883 | 0.823 | 1 |
| Terminal | TPD52L2 | 4.01E-33 | 0.640763 | 0.561 | 0.353 | 7.01E-29 |
| Fibroblasts | FZD2 | 8.93E-16 | 0.639264 | 0.217 | 0.03 | 1.56E-11 |
| Contractile | CEP70 | 7.95E-12 | 0.639187 | 0.33 | 0.106 | 1.39E-07 |
| Pericytes | ENSSSCG0( | 1.46E-07 | 0.639136 | 0.122 | 0.028 | 0.002544 |
| Pericytes | ROCK1 | 2.26E-05 | 0.638952 | 0.689 | 0.609 | 0.394413 |
| IFN_Interm | ENSSSCG0( | 2.09E-15 | 0.638908 | 0.158 | 0.006 | 3.65E-11 |
| Contractile | GLDN | 2.13E-37 | 0.638908 | 0.182 | 0.043 | 3.72E-33 |
| Terminal | PRDX4 | 2.36E-38 | 0.638904 | 0.776 | 0.62 | 4.13E-34 |
| Fibroblasts | FBLN7 | 5.86E-05 | 0.638694 | 0.233 | 0.088 | 1 |
| Contractile | TMEM70 | 3.29E-52 | 0.638333 | 0.511 | 0.257 | 5.75E-48 |
| Fibroblasts | HNRNPDL | 0.000251 | 0.638154 | 0.683 | 0.6 | 1 |
| Fibroblasts | UST | 1.62E-09 | 0.637999 | 0.133 | 0.02 | 2.84E-05 |
| Terminal | RACK1 | 8.61E-47 | 0.637832 | 0.926 | 0.832 | 1.51E-42 |
| Pericytes | ADAMTS9 | 8.84E-07 | 0.637564 | 0.244 | 0.097 | 0.015451 |
| Pericytes | ACYP1 | 0.005069 | 0.636945 | 0.344 | 0.264 | 1 |
| IFN_Interm | CD302 | 0.009471 | 0.636351 | 0.632 | 0.362 | 1 |
| Pericytes | CTNND1 | 0.0003 | 0.635786 | 0.322 | 0.203 | 1 |
| Pericytes | ENSSSCG0( | 0.000335 | 0.63571 | 0.222 | 0.112 | 1 |
| Fibroblasts | PTGS2 | 0.000357 | 0.635672 | 0.1 | 0.026 | 1 |
| Terminal | EMILIN2 | 9.71E-51 | 0.63549 | 0.273 | 0.071 | 1.7E-46 |
| Pericytes | RBPMS2 | 5.37E-08 | 0.635279 | 0.189 | 0.056 | 0.000939 |
| IFN_Interm | LACTB | 0.006517 | 0.634322 | 0.263 | 0.092 | 1 |
| Terminal | VEGFC | 7.22E-64 | 0.634246 | 0.25 | 0.044 | 1.26E-59 |
| Contractile | LYNX1 | 1.2E-06 | 0.633703 | 0.538 | 0.321 | 0.020978 |
| Contractile | FERMT2 | 7.51E-10 | 0.633684 | 0.901 | 0.736 | 1.31E-05 |
| Fibroblasts | MESD | 2.16E-05 | 0.633674 | 0.65 | 0.444 | 0.377372 |
| Fibroblasts | PTPRS | 5.72E-10 | 0.633571 | 0.233 | 0.051 | 1E-05 |
| Terminal | ADM | 2.69E-32 | 0.633125 | 0.229 | 0.073 | 4.71E-28 |
| Pericytes | PNRC1 | 0.001828 | 0.633061 | 0.589 | 0.55 | 1 |
| Terminal | ENSSSCG0( | 5.02E-31 | 0.632651 | 0.575 | 0.358 | 8.78E-27 |
| IFN_Interm | RTP4 | 4.98E-10 | 0.631677 | 0.316 | 0.037 | 8.71E-06 |
| IFN_Interm | CES1 | 5.45E-13 | 0.631675 | 0.105 | 0.003 | 9.53E-09 |
| Contractile | HMCN1 | 2.45E-44 | 0.631675 | 0.577 | 0.335 | 4.29E-40 |
| Contractile | MACF1 | 3.54E-35 | 0.631078 | 0.565 | 0.358 | 6.18E-31 |
| Contractile | ITGAV | 1.66E-05 | 0.631056 | 0.802 | 0.674 | 0.290223 |

| | | | | | | |
|---|---|---|---|---|---|---|
| Pericytes | SLC12A2 | 0.005031 | 0.631038 | 0.133 | 0.065 | 1 |
| Terminal | SSR4 | 1.47E-28 | 0.630691 | 0.739 | 0.599 | 2.56E-24 |
| Terminal | CD109 | 7.82E-23 | 0.630589 | 0.608 | 0.453 | 1.37E-18 |
| Fibroblasts | ANGPTL5 | 2.76E-17 | 0.630505 | 0.15 | 0.013 | 4.83E-13 |
| Fibroblasts | EBF1 | 0.001563 | 0.63033 | 0.35 | 0.201 | 1 |
| Pericytes | AFAP1L2 | 6.42E-16 | 0.629454 | 0.133 | 0.016 | 1.12E-11 |
| Contractile | KCTD10 | 5.19E-48 | 0.626994 | 0.563 | 0.315 | 9.07E-44 |
| Contractile | RCAN2 | 5.46E-06 | 0.626834 | 0.736 | 0.544 | 0.095489 |
| Contractile | SETBP1 | 1.79E-08 | 0.626634 | 0.242 | 0.08 | 0.000314 |
| Fibroblasts | NENF | 0.009852 | 0.626522 | 0.4 | 0.302 | 1 |
| Pericytes | MARCHF8 | 3.59E-08 | 0.626124 | 0.233 | 0.08 | 0.000628 |
| Contractile | PENK | 1.57E-66 | 0.625904 | 0.726 | 0.407 | 2.74E-62 |
| Intermedia | GPX3 | 1.2E-20 | 0.625122 | 0.222 | 0.101 | 2.1E-16 |
| Terminal | RCN1 | 2.43E-29 | 0.625084 | 0.676 | 0.52 | 4.26E-25 |
| Contractile | ARPC1A | 1.43E-06 | 0.625037 | 0.473 | 0.272 | 0.025029 |
| Fibroblasts | TENM3 | 4.06E-61 | 0.623092 | 0.15 | 0.002 | 7.1E-57 |
| Pericytes | PABPC1 | 9.55E-07 | 0.622832 | 0.767 | 0.688 | 0.016702 |
| Contractile | SLC7A2 | 0.000529 | 0.622727 | 0.407 | 0.257 | 1 |
| Terminal | ADAMTSL4 | 1.36E-41 | 0.622254 | 0.339 | 0.126 | 2.37E-37 |
| Contractile | C1QTNF7 | 5.35E-06 | 0.621562 | 0.571 | 0.362 | 0.09351 |
| Terminal | ERO1A | 3.4E-23 | 0.621524 | 0.517 | 0.333 | 5.95E-19 |
| Contractile | RBPMS | 7.25E-07 | 0.621089 | 0.571 | 0.348 | 0.012681 |
| Fibroblasts | VGLL3 | 0.003176 | 0.620661 | 0.267 | 0.143 | 1 |
| Contractile | DNAJB4 | 3.01E-05 | 0.620521 | 0.593 | 0.418 | 0.525953 |
| IFN_Interm | PARP12 | 2.69E-08 | 0.620289 | 0.211 | 0.021 | 0.00047 |
| Terminal | CKS2 | 4.96E-52 | 0.620239 | 0.229 | 0.047 | 8.68E-48 |
| Contractile | EPAS1 | 9.51E-58 | 0.61939 | 0.796 | 0.55 | 1.66E-53 |
| Terminal | TSPAN6 | 5.34E-35 | 0.618512 | 0.524 | 0.285 | 9.33E-31 |
| Fibroblasts | SAMD12 | 1.92E-08 | 0.618129 | 0.3 | 0.09 | 0.000335 |
| Intermedia | VMP1 | 1.04E-33 | 0.617681 | 0.819 | 0.663 | 1.82E-29 |
| Contractile | ADGRL2 | 0.00084 | 0.617298 | 0.396 | 0.264 | 1 |
| Fibroblasts | CLMP | 6E-14 | 0.616993 | 0.2 | 0.029 | 1.05E-09 |
| Terminal | EIF1 | 1.23E-61 | 0.616723 | 0.996 | 0.962 | 2.15E-57 |
| Pericytes | PAIP2 | 0.001193 | 0.616278 | 0.478 | 0.384 | 1 |
| Fibroblasts | MTR | 3.07E-09 | 0.615937 | 0.167 | 0.03 | 5.37E-05 |
| Contractile | SLC8A1 | 3.65E-09 | 0.615899 | 0.264 | 0.087 | 6.37E-05 |
| Contractile | LIMS2 | 7.79E-07 | 0.615772 | 0.67 | 0.464 | 0.013616 |
| Pericytes | ENSSSCG0( | 0.007865 | 0.615121 | 0.233 | 0.152 | 1 |
| Pericytes | UBE2E3 | 0.000166 | 0.614533 | 0.256 | 0.137 | 1 |
| Contractile | C1ORF198 | 2.44E-38 | 0.613837 | 0.671 | 0.462 | 4.27E-34 |
| Terminal | CTSV | 1.1E-30 | 0.613545 | 0.835 | 0.694 | 1.93E-26 |
| Contractile | AIF1L | 4.64E-07 | 0.613445 | 0.527 | 0.301 | 0.00811 |
| Fibroblasts | EIF4EBP2 | 0.007662 | 0.613146 | 0.433 | 0.317 | 1 |
| Contractile | ITGA8 | 1.89E-09 | 0.613109 | 0.934 | 0.701 | 3.3E-05 |
| Contractile | ENSSSCG0( | 1.74E-43 | 0.613054 | 0.426 | 0.201 | 3.05E-39 |
| Contractile | ENSSSCG0( | 3.37E-45 | 0.612746 | 0.602 | 0.362 | 5.89E-41 |
| Pericytes | C5ORF24 | 0.003827 | 0.612544 | 0.489 | 0.438 | 1 |
| Terminal | SELENOF | 2.46E-33 | 0.612312 | 0.705 | 0.543 | 4.3E-29 |
| Contractile | MYOM1 | 2.19E-07 | 0.611724 | 0.264 | 0.1 | 0.003823 |
| Pericytes | STAT3 | 0.000761 | 0.611685 | 0.467 | 0.364 | 1 |
| Fibroblasts | ENSSSCG0( | 0.000142 | 0.611524 | 0.367 | 0.182 | 1 |
| IFN_Interm | SUMF2 | 4.72E-06 | 0.610702 | 0.263 | 0.045 | 0.082468 |
| Terminal | PAPSS2 | 1.9E-41 | 0.610534 | 0.299 | 0.101 | 3.32E-37 |
| Pericytes | MAPRE2 | 0.000297 | 0.610095 | 0.411 | 0.297 | 1 |
| Pericytes | DYNC1LI1 | 0.007627 | 0.609737 | 0.322 | 0.247 | 1 |
| Contractile | MRAP2 | 6.9E-12 | 0.609456 | 0.319 | 0.1 | 1.21E-07 |
| Pericytes | MEOX2 | 1.06E-10 | 0.608945 | 0.222 | 0.059 | 1.85E-06 |
| Fibroblasts | F2RL2 | 2.99E-14 | 0.608814 | 0.183 | 0.024 | 5.23E-10 |
| Terminal | RBP4 | 5.9E-57 | 0.608323 | 0.114 | 0.002 | 1.03E-52 |

| | | | | | | |
|---|---|---|---|---|---|---|
| Pericytes | BCAS3 | 3.25E-07 | 0.60813 | 0.211 | 0.073 | 0.005684 |
| Contractile | NPNT | 1.27E-46 | 0.60764 | 0.431 | 0.192 | 2.23E-42 |
| Pericytes | SPRY1 | 0.005679 | 0.607206 | 0.289 | 0.199 | 1 |
| Contractile | YAP1 | 3.47E-05 | 0.607088 | 0.505 | 0.351 | 0.607007 |
| Pericytes | LIFR | 2.67E-08 | 0.607081 | 0.167 | 0.044 | 0.000466 |
| Fibroblasts | SLC35A1 | 0.000554 | 0.606856 | 0.25 | 0.115 | 1 |
| Pericytes | PNPLA2 | 0.007788 | 0.606841 | 0.311 | 0.235 | 1 |
| Terminal | ENSSSCG0( | 7.54E-32 | 0.606603 | 0.421 | 0.208 | 1.32E-27 |
| Pericytes | TAF1D | 0.000181 | 0.606124 | 0.556 | 0.485 | 1 |
| Fibroblasts | CELF2 | 1.89E-20 | 0.606024 | 0.217 | 0.023 | 3.31E-16 |
| Pericytes | CACNA1H | 7.57E-12 | 0.605778 | 0.167 | 0.032 | 1.32E-07 |
| Terminal | AP3S1 | 1.03E-32 | 0.605305 | 0.617 | 0.405 | 1.8E-28 |
| Terminal | CYP7A1 | 9.29E-51 | 0.60507 | 0.121 | 0.007 | 1.62E-46 |
| Contractile | MYOZ2 | 7.15E-06 | 0.604821 | 0.264 | 0.116 | 0.124951 |
| Terminal | TRPS1 | 6.71E-27 | 0.604819 | 0.494 | 0.293 | 1.17E-22 |
| Intermedia | MRC2 | 3.82E-22 | 0.604582 | 0.497 | 0.359 | 6.68E-18 |
| Contractile | GASK1B | 1.65E-46 | 0.604579 | 0.637 | 0.375 | 2.89E-42 |
| Pericytes | CNN2 | 0.004336 | 0.604577 | 0.333 | 0.252 | 1 |
| Pericytes | HNRNPM | 0.00011 | 0.604233 | 0.656 | 0.574 | 1 |
| Terminal | CIDEC | 9.91E-56 | 0.604126 | 0.204 | 0.033 | 1.73E-51 |
| IFN_Interm | AGRN | 0.007928 | 0.604069 | 0.316 | 0.116 | 1 |
| Terminal | KCNK12 | 1.49E-84 | 0.603926 | 0.178 | 0.006 | 2.6E-80 |
| Intermedia | CCN5 | 1.67E-22 | 0.603858 | 0.377 | 0.23 | 2.92E-18 |
| Intermedia | COL3A1 | 4.55E-34 | 0.603291 | 0.99 | 0.965 | 7.96E-30 |
| IFN_Interm | SLC22A18 | 0.002912 | 0.603286 | 0.211 | 0.056 | 1 |
| Fibroblasts | SLC39A1 | 0.000836 | 0.603031 | 0.333 | 0.187 | 1 |
| Fibroblasts | CLSTN1 | 0.004863 | 0.60298 | 0.45 | 0.335 | 1 |
| Fibroblasts | MOSMO | 0.001138 | 0.602817 | 0.333 | 0.185 | 1 |
| Terminal | GALNT15 | 1.61E-20 | 0.602585 | 0.292 | 0.143 | 2.82E-16 |
| Contractile | CALD1 | 2.38E-12 | 0.602584 | 1 | 0.964 | 4.17E-08 |
| Pericytes | NR1H3 | 2.59E-06 | 0.602139 | 0.167 | 0.055 | 0.045339 |
| Pericytes | ENSSSCG0( | 0.003768 | 0.602062 | 0.522 | 0.466 | 1 |
| Contractile | FERMT2 | 1.29E-52 | 0.60192 | 0.841 | 0.674 | 2.26E-48 |
| Intermedia | PRR15 | 3.51E-36 | 0.601783 | 0.21 | 0.062 | 6.14E-32 |
| Contractile | SMARCD3 | 2.54E-52 | 0.601777 | 0.403 | 0.166 | 4.44E-48 |
| Fibroblasts | CEP126 | 0.006068 | 0.601527 | 0.267 | 0.153 | 1 |
| Terminal | ATF4 | 2.23E-26 | 0.601028 | 0.85 | 0.758 | 3.89E-22 |
| IFN_Interm | NAPB | 0.000203 | 0.600893 | 0.105 | 0.012 | 1 |
| Fibroblasts | IL17D | 0.00042 | 0.600792 | 0.25 | 0.118 | 1 |
| Contractile | TTLL7 | 8.88E-08 | 0.600415 | 0.802 | 0.578 | 0.001552 |
| Intermedia | MAFB | 2.11E-21 | 0.600243 | 0.3 | 0.156 | 3.69E-17 |
| Contractile | CFL2 | 2.99E-06 | 0.600235 | 0.626 | 0.43 | 0.052309 |
| Fibroblasts | ZNF462 | 2.98E-09 | 0.60012 | 0.133 | 0.02 | 5.21E-05 |
| Fibroblasts | GPM6B | 2.34E-07 | 0.599755 | 0.183 | 0.043 | 0.004091 |
| Fibroblasts | APPL1 | 0.008304 | 0.598926 | 0.5 | 0.377 | 1 |
| Fibroblasts | FOLH1 | 7.33E-09 | 0.598668 | 0.2 | 0.044 | 0.000128 |
| Pericytes | CH25H | 2.43E-26 | 0.597541 | 0.122 | 0.007 | 4.25E-22 |
| Contractile | CRIM1 | 1.63E-34 | 0.597402 | 0.572 | 0.365 | 2.85E-30 |
| Terminal | ENSSSCG0( | 8.92E-52 | 0.596543 | 0.978 | 0.942 | 1.56E-47 |
| Pericytes | GTF2H5 | 0.006639 | 0.596497 | 0.333 | 0.259 | 1 |
| Pericytes | ZNF217 | 3.25E-07 | 0.59612 | 0.167 | 0.049 | 0.005677 |
| Pericytes | PITRM1 | 5.3E-05 | 0.596093 | 0.178 | 0.072 | 0.926429 |
| Fibroblasts | CCDC146 | 8.23E-16 | 0.595874 | 0.2 | 0.026 | 1.44E-11 |
| Contractile | ADCY5 | 2.06E-63 | 0.595726 | 0.266 | 0.052 | 3.61E-59 |
| IFN_Interm | CEP20 | 0.004386 | 0.595676 | 0.211 | 0.059 | 1 |
| Contractile | PEA15 | 6.5E-40 | 0.595603 | 0.508 | 0.29 | 1.14E-35 |
| IFN_Interm | SLC49A3 | 0.008933 | 0.595228 | 0.211 | 0.066 | 1 |
| Fibroblasts | NID1 | 0.000443 | 0.595221 | 0.15 | 0.051 | 1 |
| Terminal | PNP | 1.28E-44 | 0.593609 | 0.196 | 0.04 | 2.23E-40 |

| | | | | | | |
|---|---|---|---|---|---|---|
| IFN_Interm | PDIA3 | 0.004355 | 0.593516 | 0.895 | 0.835 | 1 |
| Fibroblasts | ANKRD46 | 1.3E-06 | 0.592417 | 0.317 | 0.119 | 0.022739 |
| Contractile | ATOSA | 3.74E-06 | 0.592371 | 0.286 | 0.127 | 0.065464 |
| Pericytes | PAK1 | 1.26E-07 | 0.59224 | 0.211 | 0.07 | 0.0022 |
| Pericytes | ELL2 | 0.007009 | 0.592128 | 0.233 | 0.152 | 1 |
| Fibroblasts | GSN | 0.001587 | 0.592042 | 0.95 | 0.966 | 1 |
| Terminal | PLOD1 | 3.88E-36 | 0.590987 | 0.383 | 0.163 | 6.78E-32 |
| Intermedia | SULF2 | 7.3E-26 | 0.590696 | 0.502 | 0.329 | 1.28E-21 |
| Pericytes | SUMO2 | 1.99E-07 | 0.590269 | 0.889 | 0.869 | 0.003485 |
| Pericytes | KDM5B | 0.004354 | 0.589415 | 0.244 | 0.156 | 1 |
| IFN_Interm | C1R | 0.003806 | 0.589403 | 0.947 | 0.785 | 1 |
| Pericytes | EIF3D | 0.001186 | 0.589054 | 0.456 | 0.375 | 1 |
| Contractile | ENSSSCG0( | 5.94E-06 | 0.588967 | 0.648 | 0.453 | 0.103849 |
| Fibroblasts | NCEH1 | 9.07E-10 | 0.588537 | 0.233 | 0.052 | 1.59E-05 |
| Intermedia | FBLN7 | 1.58E-24 | 0.587777 | 0.172 | 0.059 | 2.76E-20 |
| Contractile | ENSSSCG0( | 1.58E-05 | 0.587342 | 0.451 | 0.26 | 0.276766 |
| Fibroblasts | DAPK1 | 6.71E-51 | 0.587165 | 0.117 | 0.001 | 1.17E-46 |
| Contractile | ACTB | 3.23E-57 | 0.587044 | 0.861 | 0.659 | 5.64E-53 |
| Fibroblasts | ENSSSCG0( | 7.81E-08 | 0.586994 | 0.117 | 0.018 | 0.001366 |
| Pericytes | UBE2S | 0.001723 | 0.586972 | 0.222 | 0.128 | 1 |
| Contractile | BCAM | 2.33E-08 | 0.586902 | 0.33 | 0.132 | 0.000408 |
| IFN_Interm | PEX1 | 0.006443 | 0.586848 | 0.158 | 0.039 | 1 |
| Fibroblasts | ENSSSCG0( | 6.5E-08 | 0.586702 | 0.15 | 0.029 | 0.001137 |
| Terminal | PDGFRA | 3.48E-37 | 0.586538 | 0.38 | 0.161 | 6.09E-33 |
| Pericytes | RBM39 | 0.00068 | 0.586189 | 0.5 | 0.398 | 1 |
| Pericytes | STK39 | 2.48E-05 | 0.585856 | 0.156 | 0.056 | 0.433806 |
| Contractile | PIAS1 | 1.52E-07 | 0.585756 | 0.44 | 0.217 | 0.002657 |
| Intermedia | RNASE4 | 1.03E-39 | 0.585249 | 0.751 | 0.566 | 1.79E-35 |
| IFN_Interm | HES6 | 0.007656 | 0.58522 | 0.263 | 0.093 | 1 |
| Pericytes | UQCRC1 | 0.000509 | 0.585119 | 0.4 | 0.293 | 1 |
| Contractile | ACTR2 | 1.3E-05 | 0.584355 | 0.67 | 0.517 | 0.227473 |
| Fibroblasts | ATP6AP2 | 0.009358 | 0.584346 | 0.517 | 0.417 | 1 |
| IFN_Interm | STK17A | 0.004156 | 0.58398 | 0.579 | 0.292 | 1 |
| Terminal | EIF4EBP1 | 8.25E-23 | 0.583974 | 0.444 | 0.269 | 1.44E-18 |
| Intermedia | GHSR | 2.51E-28 | 0.583533 | 0.183 | 0.058 | 4.39E-24 |
| Fibroblasts | GRIA4 | 1.16E-34 | 0.583473 | 0.15 | 0.005 | 2.03E-30 |
| Pericytes | CCT7 | 0.000213 | 0.582894 | 0.5 | 0.407 | 1 |
| Pericytes | GPR20 | 1.26E-05 | 0.58243 | 0.111 | 0.03 | 0.220508 |
| Fibroblasts | PLAAT3 | 2.63E-06 | 0.582404 | 0.283 | 0.102 | 0.045939 |
| IFN_Interm | CACNA1G | 2.03E-05 | 0.582369 | 0.105 | 0.009 | 0.355386 |
| Contractile | ZC2HC1A | 6.03E-05 | 0.582342 | 0.527 | 0.378 | 1 |
| Pericytes | POLD4 | 0.003304 | 0.581473 | 0.533 | 0.454 | 1 |
| Contractile | PDE5A | 8.08E-05 | 0.580693 | 0.363 | 0.207 | 1 |
| Pericytes | PDGFB | 1.79E-29 | 0.580621 | 0.133 | 0.008 | 3.12E-25 |
| Contractile | VASP | 6.01E-46 | 0.580149 | 0.405 | 0.181 | 1.05E-41 |
| Terminal | RCN3 | 8.3E-26 | 0.579899 | 0.641 | 0.472 | 1.45E-21 |
| Contractile | FBLIM1 | 1.18E-42 | 0.579893 | 0.534 | 0.301 | 2.07E-38 |
| Contractile | PDLIM3 | 1.27E-48 | 0.57946 | 0.279 | 0.083 | 2.21E-44 |
| Contractile | ACTN4 | 3.78E-08 | 0.578985 | 0.901 | 0.699 | 0.000662 |
| Pericytes | PHACTR2 | 0.00446 | 0.578776 | 0.156 | 0.08 | 1 |
| Pericytes | GPM6B | 6.34E-10 | 0.578582 | 0.178 | 0.042 | 1.11E-05 |
| Terminal | SDCBP | 1.13E-30 | 0.578529 | 0.8 | 0.637 | 1.98E-26 |
| Pericytes | ENSSSCG0( | 9.59E-61 | 0.578397 | 0.122 | 0.001 | 1.68E-56 |
| Fibroblasts | SERPING1 | 4.07E-06 | 0.578194 | 0.9 | 0.817 | 0.071202 |
| Contractile | GNAI1 | 0.000201 | 0.578076 | 0.352 | 0.21 | 1 |
| Pericytes | PTP4A1 | 0.00054 | 0.57763 | 0.278 | 0.167 | 1 |
| Pericytes | SLC25A5 | 0.001276 | 0.577576 | 0.267 | 0.163 | 1 |
| Terminal | ARF6 | 7.63E-26 | 0.577513 | 0.466 | 0.273 | 1.33E-21 |
| Terminal | ANKRD28 | 1.21E-38 | 0.577218 | 0.343 | 0.132 | 2.11E-34 |

| | | | | | | |
|---|---|---|---|---|---|---|
| Pericytes | FBLIM1 | 0.000383 | 0.576862 | 0.5 | 0.391 | 1 |
| Pericytes | HIPK3 | 8.97E-05 | 0.576703 | 0.367 | 0.22 | 1 |
| Fibroblasts | IGSF10 | 5.22E-48 | 0.576384 | 0.15 | 0.003 | 9.12E-44 |
| Intermedia | COL5A2 | 1.19E-39 | 0.576159 | 0.918 | 0.817 | 2.08E-35 |
| Contractile | SH3BGR | 8.72E-06 | 0.575969 | 0.505 | 0.313 | 0.152445 |
| Intermedia | ASPN | 1.75E-16 | 0.575806 | 0.569 | 0.445 | 3.06E-12 |
| Pericytes | SLC2A4 | 2.43E-17 | 0.575737 | 0.144 | 0.017 | 4.24E-13 |
| Contractile | ENSSSCG00 | 2.2E-17 | 0.575553 | 0.165 | 0.022 | 3.84E-13 |
| Fibroblasts | ATF6 | 0.009089 | 0.575338 | 0.35 | 0.236 | 1 |
| Pericytes | PARP1 | 3.16E-05 | 0.575247 | 0.3 | 0.157 | 0.552359 |
| Terminal | YWHAE | 3.83E-27 | 0.574908 | 0.8 | 0.643 | 6.7E-23 |
| Pericytes | RASGEF1B | 0.000374 | 0.574446 | 0.167 | 0.073 | 1 |
| Contractile | SBSPON | 5.04E-12 | 0.574212 | 0.176 | 0.035 | 8.82E-08 |
| Contractile | SVIL | 3.74E-38 | 0.574133 | 0.431 | 0.216 | 6.53E-34 |
| Pericytes | SDC1 | 0.001018 | 0.573927 | 0.156 | 0.071 | 1 |
| Contractile | ID3 | 1.57E-51 | 0.57344 | 0.854 | 0.632 | 2.74E-47 |
| Terminal | MCUB | 1.39E-34 | 0.573425 | 0.359 | 0.155 | 2.42E-30 |
| Terminal | AQP1 | 4.27E-09 | 0.573123 | 0.583 | 0.521 | 7.47E-05 |
| Contractile | INPP4A | 3.41E-07 | 0.573072 | 0.341 | 0.153 | 0.005962 |
| Fibroblasts | NCL | 0.003172 | 0.573025 | 0.667 | 0.599 | 1 |
| Fibroblasts | ENSSSCG00 | 0.000379 | 0.572764 | 0.167 | 0.059 | 1 |
| Contractile | RNF19A | 1.63E-06 | 0.572663 | 0.703 | 0.508 | 0.02846 |
| Contractile | TINAGL1 | 7.92E-73 | 0.572628 | 0.393 | 0.109 | 1.39E-68 |
| Terminal | NME1 | 5.25E-35 | 0.572254 | 0.377 | 0.168 | 9.18E-31 |
| Contractile | GNB4 | 2.82E-05 | 0.570951 | 0.538 | 0.381 | 0.492643 |
| Contractile | TMEM70 | 5.94E-05 | 0.570908 | 0.516 | 0.354 | 1 |
| Contractile | PPP1R12A | 4.9E-06 | 0.570372 | 0.78 | 0.681 | 0.085576 |
| Fibroblasts | SRPK2 | 0.000853 | 0.570211 | 0.3 | 0.153 | 1 |
| Contractile | COL8A1 | 4.83E-08 | 0.569368 | 0.934 | 0.863 | 0.000844 |
| Pericytes | TGFB3 | 0.002827 | 0.569186 | 0.367 | 0.253 | 1 |
| Contractile | MSANTD2 | 3.23E-07 | 0.569047 | 0.253 | 0.097 | 0.005651 |
| Intermedia | FBN1 | 5.19E-20 | 0.568817 | 0.732 | 0.626 | 9.08E-16 |
| Pericytes | ELK3 | 0.007084 | 0.568289 | 0.278 | 0.193 | 1 |
| Terminal | VIM | 1.85E-51 | 0.567133 | 0.999 | 0.999 | 3.24E-47 |
| Pericytes | NACA | 1.95E-09 | 0.566886 | 0.978 | 0.955 | 3.41E-05 |
| Pericytes | ARID1A | 5.87E-06 | 0.56677 | 0.2 | 0.077 | 0.102612 |
| Terminal | PLAUR | 2.05E-36 | 0.566504 | 0.274 | 0.093 | 3.58E-32 |
| Pericytes | ATP11A | 0.004758 | 0.56589 | 0.189 | 0.107 | 1 |
| Contractile | UBE2H | 0.000516 | 0.565701 | 0.582 | 0.469 | 1 |
| Pericytes | ENSSSCG00 | 8.3E-12 | 0.565277 | 0.144 | 0.025 | 1.45E-07 |
| Fibroblasts | RUFY3 | 0.000239 | 0.56504 | 0.233 | 0.098 | 1 |
| Fibroblasts | ENSSSCG00 | 0.000493 | 0.564957 | 0.267 | 0.127 | 1 |
| Fibroblasts | AFF1 | 0.001631 | 0.564907 | 0.35 | 0.203 | 1 |
| Contractile | MXI1 | 0.000109 | 0.56404 | 0.363 | 0.215 | 1 |
| Contractile | COL15A1 | 1.28E-45 | 0.56375 | 0.919 | 0.769 | 2.23E-41 |
| Contractile | RAB27B | 4.59E-05 | 0.563638 | 0.165 | 0.062 | 0.802148 |
| Terminal | BBLN | 5.93E-24 | 0.563283 | 0.436 | 0.257 | 1.04E-19 |
| Pericytes | EIF4A2 | 0.000172 | 0.562748 | 0.767 | 0.73 | 1 |
| Fibroblasts | HTATSF1 | 0.002205 | 0.56242 | 0.4 | 0.246 | 1 |
| Terminal | RER1 | 2.17E-29 | 0.562201 | 0.569 | 0.354 | 3.8E-25 |
| Pericytes | LGALSL | 0.000736 | 0.561539 | 0.511 | 0.433 | 1 |
| IFN_Interm | CENPB | 0.000446 | 0.560793 | 0.211 | 0.045 | 1 |
| Contractile | FZD7 | 2.48E-06 | 0.560746 | 0.33 | 0.157 | 0.043365 |
| Intermedia | BGN | 4.88E-82 | 0.560514 | 0.998 | 0.989 | 8.54E-78 |
| Contractile | DTNA | 2.97E-09 | 0.560353 | 0.198 | 0.053 | 5.19E-05 |
| Pericytes | MYOM1 | 0.003561 | 0.560216 | 0.189 | 0.103 | 1 |
| Contractile | RHOB | 4.97E-55 | 0.56018 | 0.724 | 0.459 | 8.68E-51 |
| Contractile | ENSSSCG00 | 1.77E-06 | 0.560047 | 0.363 | 0.179 | 0.031009 |
| Pericytes | PPP1R2 | 0.005301 | 0.55992 | 0.433 | 0.373 | 1 |

| | | | | | | |
|---|---|---|---|---|---|---|
| Pericytes | NCL | 0.000243 | 0.559789 | 0.678 | 0.598 | 1 |
| Fibroblasts | NR2F2 | 0.002185 | 0.559761 | 0.483 | 0.335 | 1 |
| Fibroblasts | CLDN11 | 1.38E-34 | 0.559585 | 0.15 | 0.005 | 2.41E-30 |
| Pericytes | APBB2 | 0.004173 | 0.559301 | 0.311 | 0.217 | 1 |
| Intermedia | OMD | 3.53E-28 | 0.559186 | 0.362 | 0.182 | 6.16E-24 |
| Intermedia | FBLN1 | 2.3E-27 | 0.55888 | 0.488 | 0.301 | 4.01E-23 |
| Terminal | SULF1 | 2.9E-35 | 0.55849 | 0.413 | 0.184 | 5.06E-31 |
| Terminal | IMPA2 | 4.83E-41 | 0.558317 | 0.283 | 0.091 | 8.45E-37 |
| Contractile | PDE3A | 7.82E-37 | 0.558064 | 0.492 | 0.267 | 1.37E-32 |
| Intermedia | ENSSSCG0( | 4.8E-23 | 0.557778 | 0.396 | 0.247 | 8.39E-19 |
| Contractile | DHX40 | 5.05E-06 | 0.557674 | 0.407 | 0.221 | 0.088353 |
| Pericytes | KCTD20 | 0.002363 | 0.55734 | 0.278 | 0.178 | 1 |
| Pericytes | ATP1A2 | 1.28E-07 | 0.557317 | 0.156 | 0.042 | 0.00223 |
| Pericytes | AGAP1 | 0.000117 | 0.556789 | 0.222 | 0.106 | 1 |
| Pericytes | ENSSSCG0( | 2.06E-08 | 0.556636 | 0.111 | 0.021 | 0.000361 |
| Pericytes | MAP3K5 | 3.19E-23 | 0.556187 | 0.122 | 0.008 | 5.57E-19 |
| Terminal | FAM180A | 7.16E-14 | 0.556169 | 0.407 | 0.274 | 1.25E-09 |
| Pericytes | KITLG | 0.001962 | 0.556108 | 0.189 | 0.1 | 1 |
| Terminal | HSD3B7 | 4.29E-39 | 0.555961 | 0.347 | 0.135 | 7.5E-35 |
| Contractile | MYOM1 | 8.49E-55 | 0.555922 | 0.21 | 0.034 | 1.48E-50 |
| Pericytes | SRSF1 | 0.002693 | 0.555595 | 0.322 | 0.223 | 1 |
| Contractile | PNRC1 | 1.25E-06 | 0.555422 | 0.736 | 0.545 | 0.021879 |
| Pericytes | INPP5B | 0.006483 | 0.555338 | 0.133 | 0.066 | 1 |
| Fibroblasts | NR2F1 | 0.000102 | 0.555239 | 0.133 | 0.037 | 1 |
| Contractile | MAP2 | 5.92E-36 | 0.554959 | 0.413 | 0.21 | 1.03E-31 |
| Contractile | KANK2 | 1.27E-06 | 0.554568 | 0.67 | 0.479 | 0.022265 |
| Fibroblasts | CHCHD4 | 0.000436 | 0.55419 | 0.183 | 0.069 | 1 |
| IFN_Interm | TAPBPL | 0.004059 | 0.554184 | 0.211 | 0.059 | 1 |
| Contractile | ESAM | 2.19E-06 | 0.554179 | 0.33 | 0.155 | 0.038334 |
| Contractile | RNF149 | 5.04E-05 | 0.553239 | 0.374 | 0.21 | 0.881519 |
| Contractile | SMIM13 | 3.45E-05 | 0.55306 | 0.363 | 0.201 | 0.603341 |
| Intermedia | MGP | 2.01E-35 | 0.552986 | 0.999 | 0.989 | 3.51E-31 |
| IFN_Interm | TENT2 | 0.009956 | 0.552796 | 0.263 | 0.094 | 1 |
| Contractile | HSPB8 | 0.00102 | 0.552648 | 0.341 | 0.225 | 1 |
| Terminal | PLOD2 | 2.62E-24 | 0.552404 | 0.733 | 0.568 | 4.59E-20 |
| Fibroblasts | DNAJC3 | 0.007079 | 0.55177 | 0.517 | 0.378 | 1 |
| Pericytes | SOCS4 | 0.001141 | 0.551479 | 0.267 | 0.161 | 1 |
| Fibroblasts | CRISPLD1 | 4.92E-11 | 0.551161 | 0.283 | 0.065 | 8.61E-07 |
| Contractile | RBPMS2 | 7.9E-09 | 0.551094 | 0.198 | 0.056 | 0.000138 |
| Contractile | HDAC5 | 7.83E-05 | 0.550889 | 0.319 | 0.173 | 1 |
| Intermedia | CGNL1 | 5.78E-23 | 0.550554 | 0.25 | 0.114 | 1.01E-18 |
| Fibroblasts | PRNP | 0.006294 | 0.550493 | 0.367 | 0.232 | 1 |
| Fibroblasts | ENSSSCG0( | 7.13E-05 | 0.550429 | 0.25 | 0.101 | 1 |
| Contractile | SLMAP | 0.00024 | 0.550251 | 0.385 | 0.247 | 1 |
| IFN_Interm | ATP6V1A | 0.002976 | 0.5496 | 0.263 | 0.079 | 1 |
| Contractile | ADAMTSL5 | 1.95E-43 | 0.5495 | 0.299 | 0.105 | 3.42E-39 |
| Terminal | NGEF | 3.57E-66 | 0.549401 | 0.241 | 0.038 | 6.24E-62 |
| Contractile | NOTCH3 | 1.41E-05 | 0.548967 | 0.659 | 0.437 | 0.247203 |
| Intermedia | ITM2C | 1.17E-26 | 0.54864 | 0.713 | 0.594 | 2.05E-22 |
| Pericytes | NDRG4 | 3.18E-05 | 0.547891 | 0.1 | 0.027 | 0.555488 |
| Terminal | TUBB2A | 2.66E-16 | 0.547071 | 0.399 | 0.257 | 4.64E-12 |
| IFN_Interm | ETV7 | 2.31E-37 | 0.547003 | 0.105 | 0.001 | 4.03E-33 |
| Pericytes | NFIL3 | 0.000669 | 0.546727 | 0.2 | 0.1 | 1 |
| Fibroblasts | ABCA8 | 2.47E-69 | 0.54668 | 0.1 | 0 | 4.31E-65 |
| Terminal | UGDH | 1.8E-23 | 0.546458 | 0.395 | 0.217 | 3.15E-19 |
| Pericytes | MGAT3 | 3.35E-76 | 0.545535 | 0.111 | 0 | 5.86E-72 |
| Intermedia | CD302 | 3.26E-23 | 0.545412 | 0.473 | 0.32 | 5.71E-19 |
| IFN_Interm | PIGK | 0.007664 | 0.545379 | 0.421 | 0.186 | 1 |
| Contractile | NFIA | 4.48E-05 | 0.545335 | 0.505 | 0.331 | 0.783705 |

| | | | | | | |
|---|---|---|---|---|---|---|
| Fibroblasts | IFNAR1 | 0.005702 | 0.545295 | 0.267 | 0.149 | 1 |
| Pericytes | ENSSSCG0( | 0.004119 | 0.545221 | 0.556 | 0.515 | 1 |
| Fibroblasts | GGNBP2 | 0.00358 | 0.545203 | 0.467 | 0.32 | 1 |
| Pericytes | GRK5 | 0.00021 | 0.544785 | 0.211 | 0.102 | 1 |
| Pericytes | CGGBP1 | 0.002822 | 0.544399 | 0.533 | 0.495 | 1 |
| IFN_Interm | ENSSSCG0( | 0.003956 | 0.544154 | 0.105 | 0.018 | 1 |
| Contractile | TGFB1I1 | 3.48E-39 | 0.54374 | 0.474 | 0.257 | 6.08E-35 |
| Terminal | PTPN18 | 5.68E-35 | 0.543524 | 0.399 | 0.178 | 9.93E-31 |
| Fibroblasts | SH3BP5 | 2.28E-09 | 0.5434 | 0.25 | 0.061 | 3.99E-05 |
| Contractile | GPC6 | 2.6E-06 | 0.543054 | 0.308 | 0.137 | 0.045389 |
| Pericytes | PNN | 1.81E-05 | 0.543028 | 0.678 | 0.596 | 0.315592 |
| IFN_Interm | ACAD9 | 0.002384 | 0.543021 | 0.211 | 0.054 | 1 |
| Terminal | CTSC | 3.69E-15 | 0.542659 | 0.442 | 0.293 | 6.46E-11 |
| IFN_Interm | GJA1 | 0.005727 | 0.542547 | 0.684 | 0.36 | 1 |
| Pericytes | COBLL1 | 0.00602 | 0.541811 | 0.156 | 0.083 | 1 |
| Fibroblasts | OSR2 | 2.85E-12 | 0.541764 | 0.117 | 0.012 | 4.99E-08 |
| Terminal | PHLDA2 | 4.97E-17 | 0.54162 | 0.687 | 0.561 | 8.69E-13 |
| Contractile | CAMK2G | 4.87E-44 | 0.541484 | 0.44 | 0.207 | 8.52E-40 |
| Contractile | SORT1 | 9.98E-11 | 0.541264 | 0.231 | 0.062 | 1.74E-06 |
| Pericytes | CALCRL | 1.25E-14 | 0.541038 | 0.178 | 0.029 | 2.19E-10 |
| Pericytes | CORO7 | 0.00666 | 0.540969 | 0.1 | 0.044 | 1 |
| Terminal | PPIA | 7.27E-29 | 0.540872 | 0.908 | 0.851 | 1.27E-24 |
| Contractile | NARS2 | 0.000198 | 0.540782 | 0.297 | 0.157 | 1 |
| Contractile | ACADL | 3.25E-05 | 0.540643 | 0.297 | 0.151 | 0.568832 |
| Fibroblasts | MXI1 | 0.004306 | 0.540398 | 0.35 | 0.216 | 1 |
| Intermedia | VCAM1 | 5.52E-16 | 0.540344 | 0.562 | 0.432 | 9.65E-12 |
| Terminal | KDELR1 | 2.29E-27 | 0.539688 | 0.702 | 0.544 | 4E-23 |
| Contractile | SAMD11 | 1.02E-53 | 0.539451 | 0.214 | 0.039 | 1.78E-49 |
| Terminal | SRPX2 | 6.3E-68 | 0.538435 | 0.203 | 0.023 | 1.1E-63 |
| Fibroblasts | ENSSSCG0( | 0.004897 | 0.537934 | 0.467 | 0.325 | 1 |
| Fibroblasts | FAM180A | 0.007348 | 0.537873 | 0.417 | 0.303 | 1 |
| Contractile | DSG2 | 7.05E-44 | 0.537615 | 0.223 | 0.057 | 1.23E-39 |
| Pericytes | PRDM1 | 1.52E-31 | 0.537486 | 0.1 | 0.003 | 2.66E-27 |
| Pericytes | HS6ST1 | 0.001414 | 0.537362 | 0.156 | 0.072 | 1 |
| IFN_Interm | NIPAL2 | 0.006873 | 0.537241 | 0.158 | 0.04 | 1 |
| Contractile | DIAPH2 | 3.16E-05 | 0.537097 | 0.659 | 0.5 | 0.552637 |
| Terminal | MRPL17 | 8.55E-28 | 0.537049 | 0.7 | 0.537 | 1.49E-23 |
| Pericytes | EMID1 | 6.54E-39 | 0.536919 | 0.122 | 0.004 | 1.14E-34 |
| Pericytes | MCM7 | 0.007384 | 0.536616 | 0.122 | 0.059 | 1 |
| Contractile | PDE3A | 2.55E-06 | 0.536548 | 0.571 | 0.351 | 0.044505 |
| Pericytes | KLHL15 | 2.36E-05 | 0.536388 | 0.178 | 0.068 | 0.413338 |
| Pericytes | PRKAR1A | 0.000838 | 0.536374 | 0.578 | 0.513 | 1 |
| Pericytes | HDGFL3 | 0.000323 | 0.535642 | 0.411 | 0.288 | 1 |
| Contractile | UBC | 6.87E-10 | 0.534759 | 0.967 | 0.919 | 1.2E-05 |
| Pericytes | ENSSSCG0( | 2.12E-06 | 0.534423 | 0.144 | 0.042 | 0.037088 |
| Pericytes | ENSSSCG0( | 2.56E-05 | 0.534419 | 0.1 | 0.027 | 0.44718 |
| Intermedia | PLPP3 | 2.28E-20 | 0.534099 | 0.746 | 0.648 | 3.98E-16 |
| Pericytes | GPC1 | 1.05E-12 | 0.53403 | 0.133 | 0.02 | 1.84E-08 |
| Pericytes | ENSSSCG0( | 3.48E-06 | 0.533134 | 0.8 | 0.714 | 0.060879 |
| Pericytes | FAM241A | 0.000215 | 0.532988 | 0.144 | 0.057 | 1 |
| Contractile | DCTN6 | 3.35E-05 | 0.532922 | 0.473 | 0.296 | 0.585221 |
| Terminal | FKBP1A | 1.17E-20 | 0.532921 | 0.628 | 0.479 | 2.04E-16 |
| Contractile | CYS1 | 0.000247 | 0.532817 | 0.495 | 0.333 | 1 |
| IFN_Interm | IFT52 | 0.004131 | 0.532683 | 0.211 | 0.058 | 1 |
| Terminal | SPON2 | 4.51E-33 | 0.53204 | 0.162 | 0.036 | 7.89E-29 |
| Pericytes | PSIP1 | 0.000914 | 0.531923 | 0.4 | 0.289 | 1 |
| Pericytes | ENSSSCG0( | 0.004392 | 0.531873 | 0.178 | 0.098 | 1 |
| Fibroblasts | HEG1 | 0.00579 | 0.531581 | 0.283 | 0.162 | 1 |
| Contractile | HACD1 | 5.09E-06 | 0.531213 | 0.385 | 0.205 | 0.088949 |

| | | | | | | |
|---|---|---|---|---|---|---|
| Pericytes | PTMS | 5.41E-09 | 0.531101 | 0.944 | 0.929 | 9.45E-05 |
| Contractile | AKT3 | 8.4E-06 | 0.531086 | 0.582 | 0.389 | 0.146897 |
| Pericytes | ENSSSCG0( | 2.35E-06 | 0.53104 | 0.189 | 0.066 | 0.041111 |
| Contractile | THRB | 0.001371 | 0.530859 | 0.363 | 0.242 | 1 |
| Pericytes | NR2F1 | 1.85E-05 | 0.530597 | 0.122 | 0.037 | 0.324106 |
| Pericytes | CCT2 | 0.0007 | 0.530043 | 0.544 | 0.465 | 1 |
| IFN_Interm | SLC25A37 | 0.006185 | 0.529787 | 0.158 | 0.039 | 1 |
| Intermedia | C1QTNF5 | 8.05E-27 | 0.529668 | 0.785 | 0.681 | 1.41E-22 |
| Fibroblasts | VSTM4 | 0.000746 | 0.529195 | 0.217 | 0.095 | 1 |
| Fibroblasts | ESR1 | 3.01E-45 | 0.529106 | 0.133 | 0.003 | 5.26E-41 |
| Fibroblasts | DPYSL2 | 0.000363 | 0.528939 | 0.317 | 0.157 | 1 |
| Pericytes | IRF2BP2 | 0.005301 | 0.528884 | 0.378 | 0.294 | 1 |
| Intermedia | CYBRD1 | 3.41E-24 | 0.528829 | 0.66 | 0.523 | 5.96E-20 |
| Intermedia | FMOD | 3.13E-28 | 0.528044 | 0.945 | 0.887 | 5.47E-24 |
| Fibroblasts | GABARAPL | 0.006092 | 0.527777 | 0.35 | 0.218 | 1 |
| Fibroblasts | RRP15 | 0.002137 | 0.527757 | 0.35 | 0.208 | 1 |
| Pericytes | H1-0 | 1.48E-05 | 0.527538 | 0.189 | 0.073 | 0.258653 |
| Pericytes | ENSSSCG0( | 0.000706 | 0.527229 | 0.1 | 0.034 | 1 |
| Pericytes | MYSM1 | 0.006739 | 0.526818 | 0.222 | 0.139 | 1 |
| Fibroblasts | RALBP1 | 0.006351 | 0.52643 | 0.6 | 0.488 | 1 |
| Contractile | PRKG1 | 1.02E-05 | 0.526359 | 0.703 | 0.484 | 0.178624 |
| Terminal | HSPA5 | 2.75E-11 | 0.526295 | 0.733 | 0.675 | 4.81E-07 |
| Pericytes | INKA2 | 1.23E-06 | 0.526246 | 0.133 | 0.036 | 0.021467 |
| Contractile | CLIC4 | 7.3E-53 | 0.525446 | 0.938 | 0.837 | 1.28E-48 |
| Fibroblasts | MLLT3 | 3.79E-07 | 0.525345 | 0.217 | 0.059 | 0.006624 |
| IFN_Interm | MIS18BP1 | 0.001878 | 0.524858 | 0.263 | 0.075 | 1 |
| Terminal | ATP1B1 | 6.07E-23 | 0.524623 | 0.405 | 0.224 | 1.06E-18 |
| Fibroblasts | PTCH1 | 2.56E-06 | 0.524622 | 0.133 | 0.029 | 0.044799 |
| IFN_Interm | SOST | 0.007308 | 0.524421 | 0.158 | 0.039 | 1 |
| Contractile | STAT5B | 0.003573 | 0.52404 | 0.308 | 0.197 | 1 |
| Contractile | HCFC1R1 | 0.001058 | 0.52379 | 0.385 | 0.261 | 1 |
| Contractile | NEDD1 | 1.08E-32 | 0.523294 | 0.299 | 0.131 | 1.9E-28 |
| Fibroblasts | QSOX1 | 0.002093 | 0.523232 | 0.5 | 0.342 | 1 |
| Fibroblasts | NPM3 | 8.51E-07 | 0.523229 | 0.15 | 0.033 | 0.014873 |
| Contractile | RRAS | 5.55E-42 | 0.52319 | 0.841 | 0.686 | 9.7E-38 |
| Pericytes | CBL | 0.004282 | 0.52294 | 0.156 | 0.08 | 1 |
| Pericytes | IRAK2 | 8.42E-07 | 0.522939 | 0.122 | 0.03 | 0.014717 |
| Pericytes | CHCHD2 | 5.76E-05 | 0.522896 | 0.633 | 0.526 | 1 |
| Pericytes | ENSSSCG0( | 6.01E-07 | 0.522558 | 0.1 | 0.021 | 0.010504 |
| Contractile | KSR1 | 4.45E-08 | 0.522487 | 0.297 | 0.115 | 0.000779 |
| Fibroblasts | RNF144A | 2.01E-07 | 0.522362 | 0.167 | 0.037 | 0.003521 |
| Pericytes | BCAT2 | 0.001924 | 0.52211 | 0.222 | 0.128 | 1 |
| Contractile | KDSR | 0.000168 | 0.522011 | 0.484 | 0.327 | 1 |
| Pericytes | TTC33 | 9.77E-05 | 0.521997 | 0.178 | 0.074 | 1 |
| Pericytes | GAB1 | 0.000503 | 0.521768 | 0.2 | 0.098 | 1 |
| Pericytes | SPRY4 | 0.003791 | 0.521659 | 0.2 | 0.113 | 1 |
| Fibroblasts | ENSSSCG0( | 0.002453 | 0.521509 | 0.333 | 0.191 | 1 |
| Fibroblasts | TFAM | 0.002428 | 0.521193 | 0.317 | 0.178 | 1 |
| Contractile | AKAP6 | 1.26E-51 | 0.520168 | 0.227 | 0.047 | 2.2E-47 |
| Contractile | SGCD | 2.48E-32 | 0.519637 | 0.42 | 0.231 | 4.33E-28 |
| Intermedia | KLF2 | 1.16E-08 | 0.519572 | 0.506 | 0.429 | 0.000203 |
| Contractile | SEPTIN9 | 3.93E-35 | 0.519488 | 0.529 | 0.323 | 6.87E-31 |
| Fibroblasts | MT1A | 3.91E-07 | 0.519205 | 0.133 | 0.025 | 0.006836 |
| Pericytes | CLK4 | 0.002169 | 0.518654 | 0.267 | 0.169 | 1 |
| Pericytes | KIRREL1 | 0.000273 | 0.518631 | 0.133 | 0.051 | 1 |
| Fibroblasts | CLK1 | 0.000414 | 0.518602 | 0.6 | 0.426 | 1 |
| Contractile | PARP8 | 0.000565 | 0.518133 | 0.253 | 0.141 | 1 |
| Pericytes | OR51E1 | 2.28E-10 | 0.517903 | 0.122 | 0.021 | 3.98E-06 |
| Terminal | LOC11025( | 2.75E-08 | 0.517461 | 0.119 | 0.06 | 0.000481 |

| | | | | | | |
|---|---|---|---|---|---|---|
| Contractile | AEBP1 | 5.66E-60 | 0.517414 | 0.967 | 0.817 | 9.89E-56 |
| Fibroblasts | RERG | 2.49E-48 | 0.517259 | 0.133 | 0.002 | 4.35E-44 |
| Fibroblasts | ENSSSCG0( | 0.007218 | 0.517222 | 0.183 | 0.088 | 1 |
| Pericytes | ART4 | 2.28E-10 | 0.516501 | 0.122 | 0.021 | 3.99E-06 |
| Contractile | ITPR1 | 8.76E-34 | 0.516133 | 0.323 | 0.144 | 1.53E-29 |
| Pericytes | NME3 | 0.004121 | 0.515941 | 0.256 | 0.164 | 1 |
| Fibroblasts | LONRF1 | 2.32E-08 | 0.51573 | 0.2 | 0.045 | 0.000405 |
| Fibroblasts | SPON2 | 0.001227 | 0.515409 | 0.167 | 0.063 | 1 |
| Terminal | ENSSSCG0( | 1.17E-44 | 0.515283 | 0.999 | 0.993 | 2.05E-40 |
| Contractile | ESAM | 7.39E-52 | 0.514966 | 0.284 | 0.077 | 1.29E-47 |
| Pericytes | DUSP5 | 2.81E-10 | 0.514228 | 0.122 | 0.021 | 4.92E-06 |
| Pericytes | SOS2 | 0.004221 | 0.514201 | 0.222 | 0.133 | 1 |
| Contractile | UTRN | 4.77E-29 | 0.513803 | 0.695 | 0.522 | 8.34E-25 |
| Fibroblasts | BOK | 0.004312 | 0.513594 | 0.233 | 0.121 | 1 |
| Pericytes | ZNF703 | 0.002383 | 0.513552 | 0.189 | 0.101 | 1 |
| Contractile | CAP2 | 0.000187 | 0.513491 | 0.264 | 0.137 | 1 |
| Pericytes | ENSSSCG0( | 2.95E-11 | 0.51323 | 0.111 | 0.016 | 5.16E-07 |
| Pericytes | MICU3 | 0.009321 | 0.513118 | 0.144 | 0.077 | 1 |
| Intermedia | RARRES1 | 4.63E-24 | 0.512987 | 0.794 | 0.67 | 8.09E-20 |
| Fibroblasts | MICU2 | 0.00449 | 0.5116 | 0.433 | 0.294 | 1 |
| Intermedia | CTSK | 9.81E-36 | 0.511447 | 0.895 | 0.815 | 1.71E-31 |
| Contractile | LTBP1 | 1.99E-06 | 0.511411 | 0.901 | 0.764 | 0.034874 |
| Contractile | USP47 | 0.002198 | 0.510583 | 0.341 | 0.231 | 1 |
| Pericytes | ENSSSCG0( | 0.000259 | 0.510578 | 0.156 | 0.064 | 1 |
| Contractile | FGFBP3 | 6.32E-12 | 0.510404 | 0.121 | 0.018 | 1.11E-07 |
| Terminal | CCDC85B | 2.54E-22 | 0.510161 | 0.414 | 0.24 | 4.44E-18 |
| Contractile | TMEM106 | 0.000537 | 0.509826 | 0.527 | 0.391 | 1 |
| Intermedia | THBS2 | 3.25E-23 | 0.509461 | 0.806 | 0.701 | 5.69E-19 |
| Contractile | RNF11 | 7.5E-05 | 0.508877 | 0.484 | 0.325 | 1 |
| Pericytes | STK3 | 0.0021 | 0.50882 | 0.222 | 0.128 | 1 |
| Fibroblasts | SLC40A1 | 0.001061 | 0.508782 | 0.25 | 0.119 | 1 |
| Pericytes | SRRM2 | 0.004183 | 0.508538 | 0.689 | 0.669 | 1 |
| Contractile | PDK4 | 1.6E-06 | 0.508474 | 0.132 | 0.035 | 0.027926 |
| Contractile | DACT3 | 2.88E-06 | 0.508327 | 0.308 | 0.144 | 0.050318 |
| Intermedia | ITGB5 | 9.7E-18 | 0.508229 | 0.647 | 0.545 | 1.7E-13 |
| Contractile | PAWR | 7.39E-09 | 0.508133 | 0.956 | 0.781 | 0.000129 |
| Fibroblasts | HNRNPA2E | 0.006424 | 0.508077 | 0.9 | 0.871 | 1 |
| Terminal | TSKU | 2.95E-23 | 0.508054 | 0.465 | 0.278 | 5.15E-19 |
| Fibroblasts | NONO | 0.008437 | 0.50804 | 0.417 | 0.296 | 1 |
| IFN_Interm | UNC93B1 | 6.26E-05 | 0.508021 | 0.158 | 0.022 | 1 |
| Terminal | HINT1 | 5.55E-29 | 0.507804 | 0.827 | 0.672 | 9.7E-25 |
| IFN_Interm | CEP44 | 0.0002 | 0.507672 | 0.211 | 0.041 | 1 |
| Contractile | ZBTB10 | 0.00065 | 0.506963 | 0.297 | 0.174 | 1 |
| Fibroblasts | NVL | 0.004501 | 0.506963 | 0.333 | 0.2 | 1 |
| Pericytes | CTNNA3 | 0.000362 | 0.5068 | 0.178 | 0.079 | 1 |
| Fibroblasts | RRS1 | 0.000361 | 0.506783 | 0.183 | 0.068 | 1 |
| Contractile | IGFBP2 | 7.06E-36 | 0.506565 | 0.487 | 0.257 | 1.23E-31 |
| Contractile | MPRIP | 3.27E-29 | 0.506321 | 0.411 | 0.225 | 5.71E-25 |
| Contractile | CD47 | 0.000331 | 0.505808 | 0.626 | 0.507 | 1 |
| Intermedia | SERTAD4 | 1.48E-20 | 0.505658 | 0.276 | 0.145 | 2.58E-16 |
| Intermedia | TTR | 1.87E-15 | 0.505341 | 0.101 | 0.033 | 3.26E-11 |
| Contractile | SPOCK1 | 1.75E-13 | 0.505065 | 0.253 | 0.06 | 3.06E-09 |
| Contractile | PPFIBP1 | 9.67E-06 | 0.504748 | 0.736 | 0.559 | 0.169043 |
| Fibroblasts | PTGFRN | 7.6E-05 | 0.504491 | 0.233 | 0.09 | 1 |
| Fibroblasts | ENSSSCG0( | 0.000289 | 0.504455 | 0.683 | 0.549 | 1 |
| Contractile | FHL5 | 3.41E-05 | 0.503835 | 0.253 | 0.115 | 0.595663 |
| Fibroblasts | PPP3CA | 1.69E-05 | 0.503438 | 0.233 | 0.082 | 0.294883 |
| Contractile | KIF5B | 2.1E-29 | 0.502725 | 0.686 | 0.499 | 3.67E-25 |
| Pericytes | ENSSSCG0( | 5.1E-09 | 0.502668 | 0.156 | 0.036 | 8.91E-05 |

| | | | | | | |
|---|---|---|---|---|---|---|
| Contractile | MARF1 | 0.000418 | 0.50266 | 0.297 | 0.169 | 1 |
| Terminal | CLU | 1.61E-33 | 0.502346 | 0.977 | 0.914 | 2.82E-29 |
| Terminal | EIF4EBP3 | 8.23E-35 | 0.502151 | 0.251 | 0.083 | 1.44E-30 |
| Contractile | ENSSSCG0( | 2.54E-45 | 0.502043 | 0.421 | 0.188 | 4.44E-41 |
| Terminal | HK2 | 1.71E-53 | 0.501791 | 0.152 | 0.015 | 2.99E-49 |
| Terminal | TUBB | 4.43E-13 | 0.501209 | 0.658 | 0.546 | 7.74E-09 |
| Pericytes | IL17B | 0.003281 | 0.501096 | 0.589 | 0.512 | 1 |
| Terminal | RRM2 | 4.2E-49 | 0.500715 | 0.103 | 0.003 | 7.35E-45 |
| Contractile | ENSSSCG0( | 4.34E-12 | 0.50065 | 0.187 | 0.039 | 7.59E-08 |
| Fibroblasts | COLEC12 | 2.06E-27 | 0.500217 | 0.183 | 0.012 | 3.6E-23 |
| Terminal | BZW1 | 3.59E-25 | 0.499927 | 0.674 | 0.501 | 6.27E-21 |
| Terminal | COX7A2L | 4.28E-22 | 0.499756 | 0.62 | 0.447 | 7.49E-18 |
| Pericytes | TRIB2 | 0.001129 | 0.499736 | 0.178 | 0.086 | 1 |
| Fibroblasts | HSD3B1 | 2.47E-69 | 0.499612 | 0.1 | 0 | 4.31E-65 |
| Intermedia | PRSS35 | 3.91E-19 | 0.499567 | 0.243 | 0.12 | 6.84E-15 |
| Terminal | CCDC80 | 7.92E-12 | 0.499276 | 0.807 | 0.77 | 1.39E-07 |
| Fibroblasts | ENSSSCG0( | 0.000912 | 0.499018 | 0.567 | 0.405 | 1 |
| Contractile | ALDH1B1 | 1.31E-08 | 0.498913 | 0.165 | 0.042 | 0.000228 |
| Contractile | ENSSSCG0( | 0.000201 | 0.498588 | 0.462 | 0.314 | 1 |
| Pericytes | ENSSSCG0( | 0.009663 | 0.49839 | 0.178 | 0.104 | 1 |
| Contractile | GYG1 | 3.78E-32 | 0.498387 | 0.555 | 0.355 | 6.6E-28 |
| Pericytes | SRRM1 | 0.004912 | 0.498237 | 0.544 | 0.504 | 1 |
| Pericytes | NPM1 | 0.000175 | 0.49812 | 0.789 | 0.762 | 1 |
| Terminal | MYDGF | 3.3E-20 | 0.497615 | 0.484 | 0.315 | 5.77E-16 |
| Contractile | CSRP2 | 0.000221 | 0.497489 | 0.78 | 0.684 | 1 |
| Contractile | ENSSSCG0( | 3.24E-07 | 0.497482 | 0.769 | 0.545 | 0.005666 |
| Pericytes | SNRK | 0.000417 | 0.496956 | 0.167 | 0.073 | 1 |
| Fibroblasts | RAMP2 | 0.002377 | 0.496555 | 0.117 | 0.04 | 1 |
| Pericytes | MAP1LC3A | 0.002748 | 0.496529 | 0.589 | 0.583 | 1 |
| Contractile | GPR20 | 4.05E-08 | 0.496173 | 0.132 | 0.03 | 0.000709 |
| Contractile | CAVIN3 | 5.15E-08 | 0.495637 | 0.989 | 0.951 | 0.000901 |
| Terminal | CLEC11A | 3.8E-18 | 0.495515 | 0.711 | 0.559 | 6.64E-14 |
| Contractile | FLNC | 0.001531 | 0.495307 | 0.143 | 0.065 | 1 |
| Pericytes | ENSSSCG0( | 0.009717 | 0.495298 | 0.178 | 0.103 | 1 |
| Pericytes | TRAK1 | 0.000452 | 0.495161 | 0.167 | 0.074 | 1 |
| Pericytes | ABI2 | 0.00375 | 0.495143 | 0.144 | 0.07 | 1 |
| Pericytes | ENSSSCG0( | 3.15E-07 | 0.495048 | 0.133 | 0.034 | 0.005504 |
| Contractile | ARHGEF17 | 4.89E-33 | 0.494872 | 0.395 | 0.198 | 8.54E-29 |
| Pericytes | TPM1 | 0.002188 | 0.494774 | 0.978 | 0.938 | 1 |
| Terminal | SLC1A5 | 9.94E-23 | 0.494729 | 0.385 | 0.208 | 1.74E-18 |
| Contractile | ATP1B3 | 1.13E-29 | 0.494111 | 0.769 | 0.625 | 1.98E-25 |
| Terminal | PRDX6 | 1.93E-23 | 0.493965 | 0.388 | 0.211 | 3.38E-19 |
| Contractile | PNRC2 | 0.000888 | 0.493914 | 0.648 | 0.519 | 1 |
| Contractile | CXCL12 | 2.97E-07 | 0.493232 | 0.176 | 0.051 | 0.005188 |
| Pericytes | FHOD1 | 0.005821 | 0.493136 | 0.144 | 0.073 | 1 |
| Intermedia | SLC40A1 | 2.87E-24 | 0.4928 | 0.214 | 0.085 | 5.02E-20 |
| Terminal | ENSSSCG0( | 8.44E-29 | 0.492799 | 0.955 | 0.898 | 1.48E-24 |
| Contractile | AKAP12 | 8.28E-37 | 0.492658 | 0.664 | 0.436 | 1.45E-32 |
| Terminal | GDF15 | 4.36E-42 | 0.492482 | 0.117 | 0.011 | 7.62E-38 |
| Terminal | PRSS23 | 3.2E-05 | 0.492057 | 0.657 | 0.625 | 0.559987 |
| Fibroblasts | EBF2 | 0.000417 | 0.491901 | 0.15 | 0.05 | 1 |
| Pericytes | TYRO3 | 0.000221 | 0.491179 | 0.133 | 0.05 | 1 |
| Contractile | PBXIP1 | 0.000151 | 0.491136 | 0.495 | 0.332 | 1 |
| Pericytes | PLA2R1 | 0.005017 | 0.491029 | 0.133 | 0.064 | 1 |
| Pericytes | HMGB1 | 8.39E-07 | 0.49099 | 0.989 | 0.952 | 0.014661 |
| Intermedia | GNG11 | 3.34E-17 | 0.490936 | 0.72 | 0.631 | 5.83E-13 |
| Terminal | LOX | 8.23E-18 | 0.490912 | 0.875 | 0.809 | 1.44E-13 |
| Fibroblasts | ENSSSCG0( | 0.004554 | 0.490635 | 0.167 | 0.073 | 1 |
| Contractile | ACTB | 7.4E-07 | 0.49049 | 0.912 | 0.735 | 0.012942 |

| | | | | | | |
|---|---|---|---|---|---|---|
| Terminal | FAM111A | 1.57E-18 | 0.490405 | 0.654 | 0.498 | 2.74E-14 |
| Fibroblasts | LDHB | 0.001934 | 0.489844 | 0.667 | 0.559 | 1 |
| Pericytes | CACNA1D | 1.61E-14 | 0.489742 | 0.1 | 0.01 | 2.82E-10 |
| Contractile | SMC5 | 4.04E-05 | 0.489413 | 0.451 | 0.273 | 0.705364 |
| Pericytes | SLC25A3 | 0.000179 | 0.488746 | 0.8 | 0.779 | 1 |
| Contractile | SGCA | 0.000118 | 0.488601 | 0.363 | 0.211 | 1 |
| Terminal | P4HA3 | 3.05E-28 | 0.488294 | 0.32 | 0.144 | 5.33E-24 |
| Fibroblasts | ENSSSCG0( | 5.68E-06 | 0.487828 | 0.25 | 0.084 | 0.099321 |
| Fibroblasts | RGL1 | 5.49E-19 | 0.487751 | 0.167 | 0.015 | 9.59E-15 |
| Contractile | CAP2 | 6.76E-38 | 0.487619 | 0.238 | 0.076 | 1.18E-33 |
| Pericytes | ETNK2 | 1.93E-41 | 0.487329 | 0.111 | 0.003 | 3.37E-37 |
| Terminal | CEMIP | 7.11E-31 | 0.48724 | 0.204 | 0.061 | 1.24E-26 |
| Contractile | ARID5B | 5.1E-35 | 0.486868 | 0.582 | 0.353 | 8.91E-31 |
| Contractile | TBC1D1 | 0.000314 | 0.48684 | 0.209 | 0.099 | 1 |
| Contractile | PLCB1 | 4.38E-11 | 0.486816 | 0.143 | 0.026 | 7.65E-07 |
| Contractile | LRRK2 | 0.006792 | 0.486491 | 0.418 | 0.326 | 1 |
| Fibroblasts | ARHGEF10 | 2.71E-07 | 0.486401 | 0.167 | 0.038 | 0.004733 |
| Pericytes | HNRNPA1 | 0.002791 | 0.485831 | 0.6 | 0.572 | 1 |
| Pericytes | CALD1 | 2.52E-08 | 0.485805 | 0.989 | 0.964 | 0.00044 |
| Pericytes | SON | 5.6E-06 | 0.485775 | 0.822 | 0.828 | 0.0979 |
| Fibroblasts | RABGAP1L | 0.004395 | 0.485769 | 0.217 | 0.107 | 1 |
| Pericytes | SCD5 | 1.9E-08 | 0.485695 | 0.144 | 0.034 | 0.000333 |
| Intermedia | COL16A1 | 3.69E-20 | 0.48567 | 0.637 | 0.517 | 6.45E-16 |
| IFN_Interm | RIPK2 | 0.005458 | 0.485363 | 0.211 | 0.06 | 1 |
| Fibroblasts | ENSSSCG0( | 0.002663 | 0.484861 | 0.45 | 0.302 | 1 |
| Intermedia | FOS | 8.55E-12 | 0.484629 | 0.625 | 0.505 | 1.5E-07 |
| Fibroblasts | MAN1A2 | 0.009309 | 0.484261 | 0.317 | 0.199 | 1 |
| Fibroblasts | TIMM8A | 0.000156 | 0.483877 | 0.183 | 0.064 | 1 |
| Contractile | KCTD1 | 1.02E-15 | 0.483819 | 0.176 | 0.027 | 1.79E-11 |
| Pericytes | ZNF423 | 8.33E-07 | 0.483547 | 0.111 | 0.026 | 0.014564 |
| Contractile | HIP1 | 0.000145 | 0.483409 | 0.44 | 0.269 | 1 |
| Fibroblasts | PIM3 | 0.005886 | 0.483257 | 0.167 | 0.076 | 1 |
| Pericytes | SYDE1 | 0.003091 | 0.482812 | 0.2 | 0.113 | 1 |
| Contractile | SMG1 | 0.003028 | 0.482616 | 0.363 | 0.247 | 1 |
| Intermedia | SFRP4 | 3.44E-18 | 0.482472 | 0.365 | 0.227 | 6.02E-14 |
| Contractile | TNC | 5.02E-05 | 0.482399 | 0.857 | 0.634 | 0.877815 |
| Pericytes | PTGER4 | 6.98E-11 | 0.482307 | 0.111 | 0.016 | 1.22E-06 |
| Fibroblasts | KIAA0408 | 1.28E-06 | 0.482272 | 0.15 | 0.034 | 0.022451 |
| Pericytes | RO60 | 0.006149 | 0.481845 | 0.289 | 0.199 | 1 |
| Fibroblasts | LAPTM4B | 0.000393 | 0.481793 | 0.417 | 0.231 | 1 |
| Pericytes | C21ORF91 | 0.001143 | 0.481284 | 0.289 | 0.178 | 1 |
| Contractile | HSPB6 | 1.1E-27 | 0.481274 | 0.67 | 0.504 | 1.92E-23 |
| Fibroblasts | TMED4 | 0.003256 | 0.481054 | 0.567 | 0.423 | 1 |
| Contractile | ANKRD10 | 0.007651 | 0.480658 | 0.308 | 0.203 | 1 |
| Intermedia | NREP | 0.000586 | 0.480602 | 0.246 | 0.201 | 1 |
| Terminal | ANLN | 5.94E-09 | 0.48059 | 0.123 | 0.06 | 0.000104 |
| Pericytes | EPHA5 | 1.02E-53 | 0.480075 | 0.111 | 0.001 | 1.79E-49 |
| Fibroblasts | FTL | 1.99E-06 | 0.480049 | 0.983 | 0.978 | 0.034786 |
| Terminal | C16ORF74 | 2.51E-53 | 0.479441 | 0.187 | 0.027 | 4.39E-49 |
| Terminal | FAM162A | 5.68E-14 | 0.479036 | 0.532 | 0.395 | 9.93E-10 |
| Pericytes | KIAA0040 | 0.007654 | 0.478892 | 0.144 | 0.075 | 1 |
| Pericytes | FADS2 | 3.69E-07 | 0.477399 | 0.167 | 0.049 | 0.006456 |
| Contractile | GJC1 | 1.66E-09 | 0.477269 | 0.407 | 0.168 | 2.9E-05 |
| Pericytes | ZRANB2 | 0.001492 | 0.477022 | 0.644 | 0.611 | 1 |
| Pericytes | KDM7A | 0.000378 | 0.476735 | 0.222 | 0.111 | 1 |
| Pericytes | PSMB9 | 0.000214 | 0.47666 | 0.133 | 0.049 | 1 |
| Contractile | LNPEP | 0.000969 | 0.47658 | 0.319 | 0.192 | 1 |
| Intermedia | LOXL1 | 3.31E-14 | 0.47654 | 0.576 | 0.471 | 5.79E-10 |
| Pericytes | DAB2IP | 1.69E-05 | 0.476462 | 0.111 | 0.031 | 0.295751 |

| | | | | | | |
|---|---|---|---|---|---|---|
| Fibroblasts | ATP6V0B | 0.004309 | 0.476372 | 0.517 | 0.375 | 1 |
| Pericytes | ENSSSCG0( | 0.001502 | 0.476241 | 0.156 | 0.073 | 1 |
| Terminal | HK1 | 3.85E-23 | 0.475969 | 0.757 | 0.611 | 6.73E-19 |
| Pericytes | FTH1 | 2.79E-09 | 0.475699 | 1 | 0.998 | 4.87E-05 |
| Intermedia | TRPS1 | 7.58E-16 | 0.475593 | 0.437 | 0.301 | 1.33E-11 |
| Contractile | ARHGEF10 | 0.000819 | 0.475389 | 0.385 | 0.253 | 1 |
| Pericytes | IL10RB | 6.35E-05 | 0.475022 | 0.156 | 0.058 | 1 |
| Contractile | PRKAR2B | 0.000218 | 0.474725 | 0.462 | 0.312 | 1 |
| Contractile | ASPN | 2.86E-05 | 0.474597 | 0.637 | 0.475 | 0.500599 |
| Contractile | MRPS6 | 0.001507 | 0.474524 | 0.385 | 0.255 | 1 |
| IFN_Intern | ITPKC | 7.54E-05 | 0.474072 | 0.105 | 0.011 | 1 |
| Terminal | UGP2 | 1.69E-15 | 0.473642 | 0.444 | 0.298 | 2.96E-11 |
| Fibroblasts | ANGPTL1 | 9.53E-06 | 0.473424 | 0.133 | 0.031 | 0.166665 |
| Contractile | B2M.1 | 1.33E-53 | 0.473356 | 0.931 | 0.812 | 2.32E-49 |
| IFN_Intern | PITRM1 | 0.002005 | 0.473205 | 0.263 | 0.074 | 1 |
| IFN_Intern | PLSCR1 | 0.000526 | 0.47251 | 0.316 | 0.088 | 1 |
| Terminal | EEF1D | 5.65E-20 | 0.472443 | 0.76 | 0.657 | 9.87E-16 |
| Pericytes | PIP5K1B | 1.12E-31 | 0.472303 | 0.1 | 0.003 | 1.95E-27 |
| IFN_Intern | ERAP2 | 0.007473 | 0.472165 | 0.158 | 0.039 | 1 |
| Pericytes | GNB1 | 0.004968 | 0.472114 | 0.533 | 0.507 | 1 |
| Pericytes | EPHX4 | 4.43E-08 | 0.472017 | 0.133 | 0.031 | 0.000774 |
| Terminal | VDAC3 | 1.25E-19 | 0.471812 | 0.457 | 0.29 | 2.19E-15 |
| Fibroblasts | SMIM15 | 0.002681 | 0.471522 | 0.417 | 0.271 | 1 |
| Contractile | SYCP2 | 1.15E-21 | 0.471472 | 0.449 | 0.287 | 2E-17 |
| Contractile | RARRES2 | 3.03E-45 | 0.471436 | 0.954 | 0.864 | 5.29E-41 |
| Contractile | LOC10052( | 9.87E-30 | 0.471179 | 0.415 | 0.228 | 1.73E-25 |
| Contractile | SOX13 | 9.04E-23 | 0.471045 | 0.176 | 0.019 | 1.58E-18 |
| Fibroblasts | SEMA3B | 3.85E-06 | 0.471019 | 0.133 | 0.03 | 0.067375 |
| Pericytes | CSRNP1 | 2.18E-05 | 0.470995 | 0.156 | 0.054 | 0.380837 |
| Contractile | PLCXD3 | 4.68E-06 | 0.470918 | 0.11 | 0.027 | 0.081736 |
| IFN_Intern | CASTOR1 | 0.007913 | 0.470839 | 0.421 | 0.183 | 1 |
| Contractile | VIPR2 | 4.15E-18 | 0.470676 | 0.143 | 0.016 | 7.26E-14 |
| Pericytes | SET | 0.006942 | 0.470495 | 0.689 | 0.676 | 1 |
| Contractile | CRTC3 | 7.71E-05 | 0.470318 | 0.352 | 0.195 | 1 |
| Contractile | CCDC3 | 3.53E-34 | 0.470304 | 0.533 | 0.308 | 6.17E-30 |
| Fibroblasts | ENSSSCG0( | 0.009152 | 0.469861 | 0.467 | 0.348 | 1 |
| Pericytes | C1S | 0.00031 | 0.469819 | 0.789 | 0.69 | 1 |
| Intermedia | CTSF | 2.37E-22 | 0.469653 | 0.669 | 0.535 | 4.15E-18 |
| Contractile | PPP1R3B | 1.99E-05 | 0.469129 | 0.264 | 0.121 | 0.347532 |
| Terminal | RBPJ | 3.26E-21 | 0.468864 | 0.58 | 0.388 | 5.71E-17 |
| Contractile | HDAC9 | 1.4E-27 | 0.468456 | 0.264 | 0.115 | 2.45E-23 |
| IFN_Intern | LOC11025! | 5.16E-21 | 0.468429 | 0.158 | 0.004 | 9.03E-17 |
| Fibroblasts | OS9 | 0.00239 | 0.46758 | 0.383 | 0.23 | 1 |
| Terminal | DDIT3 | 1.93E-17 | 0.467542 | 0.265 | 0.133 | 3.37E-13 |
| Fibroblasts | LGR4 | 3.57E-05 | 0.467431 | 0.15 | 0.042 | 0.62458 |
| Fibroblasts | RECK | 0.000808 | 0.467185 | 0.283 | 0.138 | 1 |
| Terminal | ENSSSCG0( | 1.32E-22 | 0.467089 | 0.198 | 0.073 | 2.31E-18 |
| Intermedia | FBLN2 | 2.01E-16 | 0.466941 | 0.398 | 0.269 | 3.52E-12 |
| Intermedia | JUN | 0.003572 | 0.466839 | 0.592 | 0.586 | 1 |
| Contractile | KLF9 | 0.000453 | 0.46681 | 0.352 | 0.212 | 1 |
| Contractile | DMPK | 3.32E-36 | 0.466768 | 0.324 | 0.137 | 5.8E-32 |
| Pericytes | VPS72 | 0.003573 | 0.466487 | 0.156 | 0.079 | 1 |
| Terminal | PSMB2 | 5.89E-18 | 0.466472 | 0.527 | 0.367 | 1.03E-13 |
| Pericytes | ADGRA2 | 6.64E-05 | 0.466236 | 0.178 | 0.072 | 1 |
| Fibroblasts | DENND4C | 0.003287 | 0.466174 | 0.283 | 0.155 | 1 |
| Contractile | FAM107B | 9.96E-38 | 0.465998 | 0.334 | 0.137 | 1.74E-33 |
| IFN_Intern | TOPAZ1 | 0.006743 | 0.465931 | 0.105 | 0.019 | 1 |
| Fibroblasts | KIAA2026 | 0.002418 | 0.465881 | 0.367 | 0.22 | 1 |
| Contractile | UACA | 0.000642 | 0.46586 | 0.505 | 0.357 | 1 |

| | | | | | | |
|---|---|---|---|---|---|---|
| Intermedia | MMP14 | 1.61E-09 | 0.465306 | 0.456 | 0.371 | 2.81E-05 |
| Terminal | CD55 | 1.5E-25 | 0.464954 | 0.237 | 0.091 | 2.63E-21 |
| Pericytes | FGF13 | 9.56E-69 | 0.464827 | 0.1 | 0 | 1.67E-64 |
| Terminal | TMSB10 | 4.19E-33 | 0.464763 | 0.993 | 0.995 | 7.32E-29 |
| Pericytes | ENSSSCG0( | 0.0012 | 0.464312 | 0.156 | 0.072 | 1 |
| Terminal | PRELID1 | 7.63E-19 | 0.463751 | 0.472 | 0.307 | 1.33E-14 |
| Contractile | HSPB6 | 2E-05 | 0.463687 | 0.747 | 0.565 | 0.349373 |
| Terminal | TMEM258 | 3.17E-15 | 0.463657 | 0.654 | 0.536 | 5.55E-11 |
| Terminal | NCAM1 | 1.12E-27 | 0.463655 | 0.355 | 0.164 | 1.95E-23 |
| Pericytes | PRRX1 | 0.002133 | 0.462935 | 0.633 | 0.606 | 1 |
| Terminal | RHOD | 2.7E-20 | 0.462525 | 0.385 | 0.216 | 4.71E-16 |
| Pericytes | MFSD6 | 1.26E-10 | 0.46251 | 0.122 | 0.02 | 2.2E-06 |
| Pericytes | SMC3 | 0.006646 | 0.462318 | 0.522 | 0.483 | 1 |
| Terminal | SLC3A2 | 7.65E-21 | 0.46212 | 0.281 | 0.133 | 1.34E-16 |
| Pericytes | MTUS1 | 0.008555 | 0.461904 | 0.189 | 0.11 | 1 |
| Pericytes | PA2G4 | 0.007444 | 0.460908 | 0.5 | 0.482 | 1 |
| Terminal | TMEM208 | 7.32E-18 | 0.460807 | 0.487 | 0.318 | 1.28E-13 |
| Terminal | SELENOS | 5.37E-19 | 0.460741 | 0.68 | 0.537 | 9.39E-15 |
| Fibroblasts | NTRK2 | 1.92E-21 | 0.460019 | 0.1 | 0.004 | 3.36E-17 |
| Contractile | ILK | 2.08E-06 | 0.459735 | 0.758 | 0.583 | 0.036437 |
| Pericytes | PIP4P2 | 1.53E-05 | 0.459663 | 0.133 | 0.042 | 0.267939 |
| Contractile | COL4A5 | 1.62E-25 | 0.458763 | 0.491 | 0.307 | 2.83E-21 |
| Fibroblasts | UBL3 | 0.004744 | 0.45873 | 0.333 | 0.201 | 1 |
| Contractile | NARS2 | 1.93E-19 | 0.458082 | 0.231 | 0.114 | 3.37E-15 |
| Fibroblasts | ODC1 | 0.004863 | 0.458028 | 0.2 | 0.096 | 1 |
| Fibroblasts | NR4A2 | 0.000687 | 0.45798 | 0.167 | 0.062 | 1 |
| Contractile | LAMB2 | 5.03E-28 | 0.457794 | 0.667 | 0.472 | 8.8E-24 |
| Contractile | FLOT2 | 0.000218 | 0.457526 | 0.297 | 0.161 | 1 |
| Intermedia | AQP3 | 1.77E-14 | 0.457498 | 0.265 | 0.153 | 3.09E-10 |
| Terminal | CTSK | 1.42E-13 | 0.457474 | 0.879 | 0.825 | 2.48E-09 |
| Terminal | ENSSSCG0( | 1.8E-30 | 0.457276 | 0.967 | 0.938 | 3.14E-26 |
| Fibroblasts | DBN1 | 7.5E-06 | 0.457193 | 0.167 | 0.045 | 0.131025 |
| Fibroblasts | AFDN | 3.33E-18 | 0.457182 | 0.15 | 0.012 | 5.82E-14 |
| Fibroblasts | RASL11B | 0.000441 | 0.457114 | 0.2 | 0.08 | 1 |
| Contractile | KIF13B | 8E-05 | 0.457041 | 0.176 | 0.071 | 1 |
| Contractile | UNC45A | 3.53E-34 | 0.45693 | 0.313 | 0.133 | 6.17E-30 |
| Contractile | COL13A1 | 0.000503 | 0.456594 | 0.297 | 0.159 | 1 |
| Fibroblasts | CDC40 | 0.00289 | 0.456479 | 0.3 | 0.165 | 1 |
| Contractile | ENSSSCG0( | 3.06E-25 | 0.456401 | 0.767 | 0.619 | 5.36E-21 |
| Contractile | CRTC3 | 5.94E-27 | 0.456151 | 0.292 | 0.137 | 1.04E-22 |
| Contractile | ZC2HC1A | 9.05E-28 | 0.455591 | 0.495 | 0.306 | 1.58E-23 |
| Terminal | ENSSSCG0( | 3.61E-48 | 0.454832 | 0.213 | 0.043 | 6.31E-44 |
| Terminal | KDELR2 | 6.36E-19 | 0.453942 | 0.646 | 0.49 | 1.11E-14 |
| Pericytes | ETV1 | 0.00557 | 0.453883 | 0.267 | 0.173 | 1 |
| Contractile | ENSSSCG0( | 6.1E-07 | 0.453565 | 0.923 | 0.815 | 0.010655 |
| Contractile | DYNLL2 | 0.004975 | 0.453462 | 0.407 | 0.299 | 1 |
| Terminal | PDGFC | 1.67E-26 | 0.453405 | 0.28 | 0.118 | 2.92E-22 |
| Contractile | PLS3 | 3.16E-05 | 0.453319 | 0.912 | 0.802 | 0.551738 |
| Contractile | ITPRID2 | 0.001764 | 0.453305 | 0.418 | 0.286 | 1 |
| Pericytes | PGM5 | 0.001142 | 0.452896 | 0.167 | 0.078 | 1 |
| Pericytes | SMUG1 | 0.002032 | 0.452842 | 0.133 | 0.059 | 1 |
| Contractile | GJA5 | 0.000254 | 0.452757 | 0.484 | 0.318 | 1 |
| Terminal | NRP2 | 6.46E-23 | 0.452598 | 0.361 | 0.181 | 1.13E-18 |
| Pericytes | MLLT6 | 0.002827 | 0.452375 | 0.189 | 0.102 | 1 |
| Fibroblasts | FBP1 | 0.002 | 0.452142 | 0.117 | 0.039 | 1 |
| Contractile | DUSP3 | 3.18E-05 | 0.45211 | 0.385 | 0.214 | 0.556371 |
| Contractile | DAAM1 | 0.005321 | 0.451863 | 0.429 | 0.337 | 1 |
| Terminal | HERPUD1 | 5.42E-14 | 0.451652 | 0.561 | 0.42 | 9.48E-10 |
| Fibroblasts | FRMD6 | 0.002108 | 0.451335 | 0.3 | 0.161 | 1 |

| | | | | | | |
|---|---|---|---|---|---|---|
| Pericytes | RACK1 | 0.000567 | 0.450801 | 0.856 | 0.853 | 1 |
| Terminal | CALR | 9.82E-14 | 0.450584 | 0.802 | 0.716 | 1.72E-09 |
| Terminal | CDK2AP1 | 4.47E-18 | 0.450378 | 0.461 | 0.301 | 7.82E-14 |
| Terminal | RTN4 | 8.47E-17 | 0.450229 | 0.735 | 0.616 | 1.48E-12 |
| Terminal | VDAC2 | 1.01E-16 | 0.450019 | 0.593 | 0.455 | 1.77E-12 |
| Pericytes | CDCA7L | 0.00196 | 0.449984 | 0.144 | 0.067 | 1 |
| Terminal | GNPTAB | 1.04E-19 | 0.449922 | 0.307 | 0.159 | 1.81E-15 |
| Pericytes | EEF1G | 0.005495 | 0.449691 | 0.733 | 0.724 | 1 |
| Contractile | CA4 | 6.9E-08 | 0.449556 | 0.513 | 0.434 | 0.001206 |
| Pericytes | SERBP1 | 5.39E-06 | 0.449287 | 0.8 | 0.712 | 0.094276 |
| Contractile | CRYAB | 2.96E-37 | 0.449159 | 0.979 | 0.914 | 5.17E-33 |
| IFN_Interm | DHX58 | 4.62E-08 | 0.448834 | 0.158 | 0.013 | 0.000808 |
| Intermedia | CILP2 | 1.75E-18 | 0.448134 | 0.117 | 0.035 | 3.05E-14 |
| Contractile | DUSP3 | 1.19E-25 | 0.447919 | 0.309 | 0.157 | 2.08E-21 |
| Contractile | ENSSSCG0( | 0.000111 | 0.447772 | 0.451 | 0.287 | 1 |
| Terminal | SLC16A1.1 | 2.94E-36 | 0.447659 | 0.258 | 0.082 | 5.13E-32 |
| Pericytes | CDH24 | 5.82E-21 | 0.447411 | 0.1 | 0.006 | 1.02E-16 |
| Fibroblasts | ENSSSCG0( | 1.62E-06 | 0.447288 | 0.133 | 0.028 | 0.028333 |
| Contractile | ITGB1 | 2.58E-08 | 0.447241 | 1 | 0.981 | 0.000451 |
| Terminal | ENPP1 | 1.34E-15 | 0.44708 | 0.244 | 0.125 | 2.34E-11 |
| Contractile | TPM1 | 8.89E-07 | 0.447058 | 1 | 0.937 | 0.015533 |
| Contractile | ITGA9 | 5.47E-05 | 0.446967 | 0.132 | 0.044 | 0.956145 |
| Terminal | UBA52 | 1.16E-25 | 0.446733 | 0.918 | 0.88 | 2.02E-21 |
| Terminal | HACD4 | 6.4E-29 | 0.446566 | 0.226 | 0.078 | 1.12E-24 |
| Contractile | SMAD6 | 7E-27 | 0.446328 | 0.253 | 0.109 | 1.22E-22 |
| Contractile | CDK19 | 2.26E-05 | 0.446099 | 0.187 | 0.072 | 0.394328 |
| Terminal | SEC61B | 4.04E-24 | 0.446091 | 0.866 | 0.789 | 7.05E-20 |
| Terminal | CA5B | 6.35E-31 | 0.445694 | 0.226 | 0.074 | 1.11E-26 |
| Contractile | HEY2 | 0.000373 | 0.445601 | 0.429 | 0.273 | 1 |
| Pericytes | LINGO1 | 9.56E-09 | 0.445523 | 0.133 | 0.028 | 0.000167 |
| Intermedia | RUNX1 | 5.8E-15 | 0.445325 | 0.407 | 0.281 | 1.01E-10 |
| Pericytes | NAP1L1 | 0.000535 | 0.445183 | 0.833 | 0.826 | 1 |
| Contractile | ENSSSCG0( | 0.002046 | 0.445069 | 0.275 | 0.167 | 1 |
| Terminal | ARL6IP5 | 1.36E-17 | 0.444961 | 0.683 | 0.571 | 2.38E-13 |
| Intermedia | COL8A2 | 4.79E-47 | 0.444854 | 0.144 | 0.017 | 8.37E-43 |
| Intermedia | SDCBP | 1.35E-19 | 0.444812 | 0.746 | 0.646 | 2.36E-15 |
| Contractile | EP300 | 0.003873 | 0.444539 | 0.374 | 0.261 | 1 |
| Contractile | LRP5 | 0.000352 | 0.444022 | 0.242 | 0.122 | 1 |
| Terminal | FKBP11 | 3.47E-24 | 0.443656 | 0.292 | 0.131 | 6.06E-20 |
| Terminal | REEP5 | 7.18E-24 | 0.443543 | 0.771 | 0.653 | 1.25E-19 |
| IFN_Interm | NTAQ1 | 0.001283 | 0.443184 | 0.211 | 0.049 | 1 |
| Pericytes | TNFAIP1 | 0.000915 | 0.443102 | 0.156 | 0.07 | 1 |
| Contractile | CCND2 | 0.000547 | 0.443074 | 0.527 | 0.378 | 1 |
| Pericytes | STARD8 | 2.59E-13 | 0.442926 | 0.111 | 0.013 | 4.53E-09 |
| Terminal | TGFBI | 5.11E-14 | 0.442811 | 0.475 | 0.33 | 8.94E-10 |
| Contractile | SPINT2 | 0.002401 | 0.442705 | 0.516 | 0.377 | 1 |
| Terminal | CSTB | 8.58E-19 | 0.442622 | 0.857 | 0.774 | 1.5E-14 |
| Contractile | CDC42EP3 | 9.27E-29 | 0.442586 | 0.684 | 0.49 | 1.62E-24 |
| Pericytes | SHISAL1 | 1.17E-23 | 0.442539 | 0.1 | 0.005 | 2.05E-19 |
| Contractile | C5ORF24 | 0.000118 | 0.442513 | 0.615 | 0.434 | 1 |
| Intermedia | ADAMTS2 | 1.3E-12 | 0.442159 | 0.408 | 0.303 | 2.27E-08 |
| Contractile | SUSD5 | 0.000382 | 0.441907 | 0.253 | 0.132 | 1 |
| Contractile | ACSF2 | 0.000263 | 0.441358 | 0.297 | 0.16 | 1 |
| Contractile | LGALSL | 8.13E-05 | 0.441325 | 0.593 | 0.431 | 1 |
| Terminal | MGMT | 7.33E-15 | 0.441054 | 0.395 | 0.254 | 1.28E-10 |
| Pericytes | FOSL2 | 0.000291 | 0.440793 | 0.133 | 0.051 | 1 |
| Pericytes | OAZ1 | 0.001179 | 0.440755 | 0.822 | 0.768 | 1 |
| Contractile | CAPZA2 | 0.00012 | 0.440639 | 0.791 | 0.706 | 1 |
| Pericytes | TRIQK | 0.001516 | 0.440454 | 0.156 | 0.073 | 1 |

| | | | | | | |
|---|---|---|---|---|---|---|
| Contractile | SLK | 0.000472 | 0.440311 | 0.527 | 0.387 | 1 |
| Fibroblasts | PICK1 | 1.33E-05 | 0.440176 | 0.133 | 0.032 | 0.231891 |
| Intermedia | KIAA1217 | 1.53E-15 | 0.439972 | 0.244 | 0.135 | 2.68E-11 |
| Contractile | ARHGEF25 | 1.2E-25 | 0.439916 | 0.448 | 0.269 | 2.1E-21 |
| IFN_Interm | ERAP1 | 0.009376 | 0.439889 | 0.158 | 0.041 | 1 |
| Pericytes | ENSSSCG0( | 0.001452 | 0.439331 | 0.667 | 0.598 | 1 |
| Intermedia | SCARA3 | 1.22E-13 | 0.439159 | 0.457 | 0.343 | 2.14E-09 |
| Intermedia | GUSB | 1.04E-16 | 0.438952 | 0.373 | 0.242 | 1.81E-12 |
| Terminal | FXYD6 | 2.37E-18 | 0.43882 | 0.435 | 0.264 | 4.14E-14 |
| Terminal | VKORC1 | 6.52E-20 | 0.438785 | 0.734 | 0.604 | 1.14E-15 |
| Contractile | ETS2 | 0.001081 | 0.438688 | 0.352 | 0.222 | 1 |
| Fibroblasts | ENSSSCG0( | 0.001523 | 0.438639 | 0.917 | 0.932 | 1 |
| Contractile | MGLL | 0.006571 | 0.438602 | 0.56 | 0.455 | 1 |
| Intermedia | FSTL1 | 5.7E-29 | 0.438396 | 0.886 | 0.806 | 9.96E-25 |
| Terminal | ALPK3 | 3.71E-42 | 0.438119 | 0.213 | 0.049 | 6.48E-38 |
| Contractile | MAP3K20 | 1.11E-27 | 0.437965 | 0.272 | 0.12 | 1.93E-23 |
| Contractile | DDAH2 | 4.85E-28 | 0.437757 | 0.667 | 0.494 | 8.47E-24 |
| Terminal | POLE4 | 2.56E-16 | 0.437361 | 0.567 | 0.399 | 4.47E-12 |
| Terminal | BTF3 | 4.61E-25 | 0.437182 | 0.907 | 0.784 | 8.07E-21 |
| Pericytes | RXYLT1 | 0.005179 | 0.437177 | 0.156 | 0.08 | 1 |
| Fibroblasts | SEMA3D | 2.44E-05 | 0.437138 | 0.117 | 0.027 | 0.426868 |
| Terminal | NPM1 | 1.6E-16 | 0.437076 | 0.846 | 0.737 | 2.79E-12 |
| Contractile | ARL14EP | 0.001994 | 0.436928 | 0.396 | 0.275 | 1 |
| Pericytes | ENSSSCG0( | 0.000472 | 0.436862 | 0.611 | 0.549 | 1 |
| Contractile | EVA1C | 1.55E-05 | 0.436249 | 0.176 | 0.063 | 0.271377 |
| Contractile | PPP1R12A | 2.36E-26 | 0.436157 | 0.766 | 0.628 | 4.13E-22 |
| Contractile | PYGB | 1.15E-40 | 0.436029 | 0.204 | 0.05 | 2.02E-36 |
| Contractile | RSF1 | 0.000525 | 0.436007 | 0.473 | 0.323 | 1 |
| Intermedia | CST3 | 2.69E-34 | 0.435968 | 0.993 | 0.989 | 4.7E-30 |
| Contractile | BBX | 0.002528 | 0.435715 | 0.429 | 0.317 | 1 |
| Fibroblasts | XPNPEP1 | 0.001704 | 0.435561 | 0.15 | 0.057 | 1 |
| Contractile | AKT3 | 4.88E-27 | 0.435474 | 0.507 | 0.319 | 8.53E-23 |
| Contractile | ENSSSCG0( | 1.82E-32 | 0.435437 | 0.239 | 0.086 | 3.18E-28 |
| Terminal | RHOC | 1.87E-14 | 0.435394 | 0.701 | 0.585 | 3.26E-10 |
| Intermedia | GLIS2 | 9.49E-15 | 0.434528 | 0.319 | 0.203 | 1.66E-10 |
| Contractile | MGAT5 | 0.006722 | 0.434507 | 0.242 | 0.149 | 1 |
| Terminal | NAMPT | 3.34E-25 | 0.434491 | 0.353 | 0.171 | 5.84E-21 |
| Contractile | OGDH | 0.000612 | 0.434249 | 0.352 | 0.212 | 1 |
| Pericytes | RGMA | 1.33E-07 | 0.433792 | 0.122 | 0.027 | 0.002319 |
| Terminal | ANKRD37 | 7.6E-26 | 0.433721 | 0.254 | 0.101 | 1.33E-21 |
| Intermedia | FGD4 | 2.64E-14 | 0.433615 | 0.301 | 0.189 | 4.62E-10 |
| Contractile | CLIP1 | 2.91E-18 | 0.433608 | 0.59 | 0.461 | 5.09E-14 |
| Contractile | LY75 | 6.13E-35 | 0.43337 | 0.121 | 0.005 | 1.07E-30 |
| Terminal | DAB2 | 4.78E-15 | 0.4333 | 0.599 | 0.45 | 8.36E-11 |
| Fibroblasts | CD9 | 0.006575 | 0.432307 | 0.933 | 0.921 | 1 |
| Contractile | ADAMTSL5 | 3.4E-05 | 0.432168 | 0.341 | 0.178 | 0.594216 |
| Fibroblasts | SOCS1 | 9.55E-11 | 0.432064 | 0.2 | 0.036 | 1.67E-06 |
| Contractile | EHBP1L1 | 0.001763 | 0.432024 | 0.385 | 0.254 | 1 |
| Pericytes | NTN5 | 2.86E-08 | 0.431957 | 0.156 | 0.039 | 0.000501 |
| Terminal | NUCB2 | 2.7E-16 | 0.43189 | 0.567 | 0.42 | 4.72E-12 |
| Contractile | UBE2H | 2.26E-24 | 0.43183 | 0.579 | 0.401 | 3.95E-20 |
| Intermedia | LAMC3 | 5.85E-12 | 0.431654 | 0.324 | 0.215 | 1.02E-07 |
| Pericytes | ADAMTS7 | 1.91E-05 | 0.431391 | 0.1 | 0.026 | 0.334216 |
| Pericytes | FAM171B | 1.01E-05 | 0.431059 | 0.133 | 0.04 | 0.175945 |
| Pericytes | TNFSF13 | 0.000323 | 0.43101 | 0.156 | 0.064 | 1 |
| Terminal | APLN | 1.43E-38 | 0.430928 | 0.187 | 0.041 | 2.5E-34 |
| Contractile | KIF1C | 0.000831 | 0.430702 | 0.418 | 0.281 | 1 |
| Pericytes | TAGLN | 0.000313 | 0.430637 | 0.967 | 0.96 | 1 |
| Contractile | HUWE1 | 0.001489 | 0.430276 | 0.44 | 0.298 | 1 |

| | | | | | | |
|---|---|---|---|---|---|---|
| Contractile | NPR1 | 3.4E-24 | 0.430045 | 0.26 | 0.122 | 5.94E-20 |
| Pericytes | ENSSSCG0( | 3.09E-05 | 0.429673 | 0.944 | 0.95 | 0.539966 |
| Contractile | CTNNA3 | 1.45E-43 | 0.42961 | 0.165 | 0.026 | 2.53E-39 |
| Terminal | BCL2A1 | 4.37E-30 | 0.429421 | 0.213 | 0.068 | 7.64E-26 |
| Terminal | SARS1 | 1.3E-08 | 0.429237 | 0.461 | 0.362 | 0.000227 |
| Fibroblasts | KLF11 | 3.27E-10 | 0.428741 | 0.133 | 0.018 | 5.72E-06 |
| Terminal | CDKN1A | 8.19E-13 | 0.428648 | 0.221 | 0.116 | 1.43E-08 |
| Pericytes | SCN1B | 0.001745 | 0.428486 | 0.156 | 0.073 | 1 |
| Contractile | MAPRE2 | 0.001812 | 0.428295 | 0.44 | 0.296 | 1 |
| Contractile | MMP28 | 7.57E-38 | 0.428223 | 0.227 | 0.068 | 1.32E-33 |
| Contractile | ARHGEF17 | 0.000789 | 0.428056 | 0.418 | 0.273 | 1 |
| Contractile | PPP1R12B | 1.99E-33 | 0.427975 | 0.186 | 0.051 | 3.47E-29 |
| Contractile | MEF2C | 0.000627 | 0.427763 | 0.604 | 0.462 | 1 |
| Intermedia | DAB2 | 4.18E-17 | 0.427763 | 0.577 | 0.448 | 7.31E-13 |
| Contractile | TPM4 | 1.14E-35 | 0.427747 | 0.987 | 0.964 | 2E-31 |
| Pericytes | MGST1 | 2.41E-06 | 0.427678 | 0.122 | 0.032 | 0.042123 |
| Contractile | ENSSSCG0( | 2.12E-25 | 0.427673 | 0.611 | 0.437 | 3.7E-21 |
| Terminal | MRPL28 | 2.94E-20 | 0.42742 | 0.505 | 0.327 | 5.13E-16 |
| Contractile | GNG12 | 7.82E-06 | 0.427131 | 0.78 | 0.604 | 0.136661 |
| Intermedia | GSN | 6.9E-11 | 0.427011 | 0.968 | 0.965 | 1.21E-06 |
| Pericytes | ARHGAP24 | 0.002458 | 0.426979 | 0.189 | 0.1 | 1 |
| Terminal | GPSM2 | 6.68E-33 | 0.426418 | 0.17 | 0.04 | 1.17E-28 |
| Contractile | GSN | 1.01E-48 | 0.426384 | 0.996 | 0.945 | 1.77E-44 |
| Terminal | TPPP3 | 8.14E-09 | 0.426256 | 0.189 | 0.109 | 0.000142 |
| Intermedia | CCND3 | 4.88E-17 | 0.426236 | 0.658 | 0.561 | 8.52E-13 |
| Fibroblasts | EIF1 | 0.000224 | 0.426085 | 0.983 | 0.969 | 1 |
| Fibroblasts | LOC10052: | 0.001716 | 0.425933 | 0.917 | 0.888 | 1 |
| IFN_Interm | KIAA0319L | 0.001554 | 0.425845 | 0.158 | 0.031 | 1 |
| Contractile | KCNA5 | 2.44E-35 | 0.425798 | 0.246 | 0.083 | 4.26E-31 |
| Contractile | ENSSSCG0( | 2.18E-06 | 0.425619 | 0.385 | 0.188 | 0.0381 |
| IFN_Interm | SLA-DMA | 6.7E-06 | 0.425583 | 0.158 | 0.018 | 0.117039 |
| Fibroblasts | SHOX2 | 0.000153 | 0.425039 | 0.15 | 0.046 | 1 |
| Terminal | CHCHD3 | 2.4E-25 | 0.424925 | 0.298 | 0.13 | 4.19E-21 |
| Fibroblasts | CPED1 | 0.002043 | 0.424748 | 0.267 | 0.135 | 1 |
| Fibroblasts | PCSK6 | 1.99E-18 | 0.424726 | 0.1 | 0.005 | 3.48E-14 |
| IFN_Interm | CDHR4 | 0.000732 | 0.424497 | 0.105 | 0.014 | 1 |
| Terminal | SNRPD1 | 1.19E-18 | 0.424487 | 0.45 | 0.281 | 2.08E-14 |
| Terminal | PSMA7 | 1.1E-13 | 0.424408 | 0.672 | 0.564 | 1.92E-09 |
| Contractile | C1ORF54 | 7.76E-32 | 0.42438 | 0.41 | 0.214 | 1.36E-27 |
| Fibroblasts | HMGN4 | 0.00115 | 0.424037 | 0.233 | 0.106 | 1 |
| Intermedia | CYP1B1 | 8.35E-11 | 0.423893 | 0.182 | 0.098 | 1.46E-06 |
| Contractile | SMOC2 | 0.003047 | 0.423804 | 0.56 | 0.402 | 1 |
| Terminal | SSR1 | 4.55E-17 | 0.423383 | 0.525 | 0.359 | 7.95E-13 |
| Terminal | BCAT1 | 3.82E-37 | 0.42331 | 0.288 | 0.095 | 6.68E-33 |
| Terminal | ARRDC3 | 5.64E-21 | 0.422827 | 0.24 | 0.103 | 9.86E-17 |
| Contractile | MAP7D1 | 3.28E-24 | 0.422651 | 0.621 | 0.448 | 5.74E-20 |
| Terminal | DNAJA1 | 7.15E-15 | 0.422565 | 0.587 | 0.436 | 1.25E-10 |
| Terminal | TUSC3 | 4.98E-26 | 0.422244 | 0.289 | 0.124 | 8.7E-22 |
| Contractile | CYS1 | 2.49E-26 | 0.421915 | 0.445 | 0.266 | 4.34E-22 |
| Intermedia | SERPINE2 | 1.04E-40 | 0.421851 | 0.688 | 0.43 | 1.82E-36 |
| Terminal | SEC61A1 | 2.37E-14 | 0.421434 | 0.422 | 0.276 | 4.15E-10 |
| Pericytes | HEY1 | 0.000329 | 0.420774 | 0.111 | 0.038 | 1 |
| Contractile | ENSSSCG0( | 1.16E-05 | 0.420443 | 0.187 | 0.07 | 0.203009 |
| Contractile | VGLL4 | 0.00249 | 0.420408 | 0.264 | 0.155 | 1 |
| Fibroblasts | SORT1 | 0.001212 | 0.420395 | 0.167 | 0.065 | 1 |
| Contractile | SOD3 | 3.44E-29 | 0.420001 | 0.89 | 0.758 | 6.02E-25 |
| Contractile | TMEM131 | 1.03E-06 | 0.419978 | 0.264 | 0.104 | 0.017933 |
| Fibroblasts | LAPTM4A | 0.000462 | 0.419977 | 0.833 | 0.765 | 1 |
| Contractile | SORBS2 | 3.5E-24 | 0.419889 | 0.597 | 0.416 | 6.12E-20 |

| | | | | | | |
|---|---|---|---|---|---|---|
| Terminal | SPCS1 | 4.32E-17 | 0.419855 | 0.717 | 0.595 | 7.56E-13 |
| Terminal | CYP1B1 | 7.04E-13 | 0.419829 | 0.198 | 0.099 | 1.23E-08 |
| Pericytes | KAT7 | 0.002056 | 0.419771 | 0.189 | 0.099 | 1 |
| Pericytes | IRF1 | 0.003801 | 0.419492 | 0.111 | 0.048 | 1 |
| Terminal | GPX7 | 3.01E-25 | 0.41937 | 0.353 | 0.171 | 5.26E-21 |
| Contractile | CBX3 | 0.005144 | 0.419266 | 0.549 | 0.46 | 1 |
| Terminal | ARHGDIA | 7.36E-17 | 0.419032 | 0.661 | 0.539 | 1.29E-12 |
| Contractile | ACAA2 | 0.001255 | 0.418636 | 0.198 | 0.101 | 1 |
| Contractile | ISLR | 3.83E-25 | 0.418619 | 0.591 | 0.409 | 6.69E-21 |
| Intermedia | LIMA1 | 7.87E-12 | 0.418512 | 0.489 | 0.389 | 1.38E-07 |
| Contractile | MED11 | 0.001529 | 0.418348 | 0.264 | 0.155 | 1 |
| Contractile | CLPX | 0.000606 | 0.418207 | 0.264 | 0.143 | 1 |
| Terminal | SNRPG | 8.58E-14 | 0.417658 | 0.623 | 0.486 | 1.5E-09 |
| Contractile | ROCK1 | 3.63E-21 | 0.417347 | 0.7 | 0.552 | 6.34E-17 |
| Contractile | LRP1B | 7.83E-09 | 0.417184 | 0.143 | 0.031 | 0.000137 |
| Terminal | WDR83OS | 1.68E-14 | 0.416992 | 0.535 | 0.395 | 2.94E-10 |
| Pericytes | ENSSSCG0( | 7.3E-21 | 0.416878 | 0.1 | 0.006 | 1.28E-16 |
| Intermedia | GLT8D2 | 3.96E-16 | 0.416876 | 0.336 | 0.21 | 6.92E-12 |
| Intermedia | ECM2 | 3.47E-23 | 0.416791 | 0.873 | 0.733 | 6.07E-19 |
| Pericytes | EIF2B4 | 0.004086 | 0.416776 | 0.211 | 0.122 | 1 |
| Fibroblasts | SCN9A | 2.47E-69 | 0.416764 | 0.1 | 0 | 4.31E-65 |
| Pericytes | ENSSSCG0( | 2.55E-05 | 0.416468 | 0.889 | 0.912 | 0.446355 |
| Contractile | CLOCK | 0.001398 | 0.416405 | 0.231 | 0.123 | 1 |
| Intermedia | HTRA1 | 3.21E-36 | 0.416295 | 0.982 | 0.929 | 5.62E-32 |
| Terminal | SSR2 | 1.33E-13 | 0.416221 | 0.551 | 0.41 | 2.33E-09 |
| Contractile | HECTD1 | 0.001673 | 0.416134 | 0.549 | 0.418 | 1 |
| Contractile | PFKFB3 | 4.93E-08 | 0.415925 | 0.187 | 0.054 | 0.000862 |
| Pericytes | ENSSSCG0( | 5.36E-06 | 0.415875 | 0.967 | 0.957 | 0.093669 |
| Pericytes | PLEKHH2 | 0.002083 | 0.415794 | 0.111 | 0.045 | 1 |
| Contractile | ENSSSCG0( | 2.33E-28 | 0.415515 | 0.251 | 0.102 | 4.07E-24 |
| IFN_Interm | PALD1 | 4.87E-17 | 0.415351 | 0.105 | 0.002 | 8.52E-13 |
| Contractile | LOC10015: | 0.005494 | 0.415266 | 0.473 | 0.368 | 1 |
| Intermedia | COL8A1 | 2.69E-21 | 0.415056 | 0.939 | 0.836 | 4.7E-17 |
| Contractile | ENSSSCG0( | 0.001632 | 0.414885 | 0.253 | 0.145 | 1 |
| Fibroblasts | GALNT10 | 0.000147 | 0.414773 | 0.167 | 0.054 | 1 |
| Terminal | ENSSSCG0( | 1.18E-17 | 0.414655 | 0.403 | 0.241 | 2.07E-13 |
| Terminal | SEC61G | 2.14E-18 | 0.414616 | 0.837 | 0.757 | 3.75E-14 |
| Terminal | EIF2S2 | 1.29E-14 | 0.414593 | 0.691 | 0.557 | 2.26E-10 |
| Contractile | TSPAN2 | 3.09E-06 | 0.414505 | 0.231 | 0.088 | 0.054086 |
| Contractile | NR2F2 | 0.003143 | 0.414415 | 0.462 | 0.334 | 1 |
| Terminal | RFNG | 3.1E-19 | 0.414271 | 0.442 | 0.271 | 5.41E-15 |
| Intermedia | NBL1 | 3.55E-19 | 0.414145 | 0.642 | 0.508 | 6.2E-15 |
| Contractile | PROS1 | 5.96E-26 | 0.413976 | 0.668 | 0.49 | 1.04E-21 |
| Contractile | DYNC1LI2 | 0.000825 | 0.413895 | 0.637 | 0.5 | 1 |
| Contractile | PTGES3L | 7.92E-06 | 0.413718 | 0.22 | 0.087 | 0.13852 |
| IFN_Interm | MRRF | 0.001782 | 0.413688 | 0.158 | 0.031 | 1 |
| Intermedia | COX4I2 | 5.09E-20 | 0.413595 | 0.81 | 0.714 | 8.91E-16 |
| Contractile | FILIP1 | 3.66E-05 | 0.413483 | 0.571 | 0.361 | 0.640665 |
| Intermedia | TENT5A | 7.45E-16 | 0.41338 | 0.625 | 0.497 | 1.3E-11 |
| Terminal | LRRC59 | 6.18E-20 | 0.41333 | 0.296 | 0.148 | 1.08E-15 |
| Terminal | EIF4A1 | 3.31E-14 | 0.413211 | 0.591 | 0.447 | 5.78E-10 |
| Contractile | GUCY1A1 | 0.000928 | 0.412906 | 0.253 | 0.137 | 1 |
| Contractile | MAP2 | 0.009937 | 0.412742 | 0.396 | 0.288 | 1 |
| Contractile | ITGA3 | 0.000111 | 0.412542 | 0.22 | 0.101 | 1 |
| Pericytes | YWHAZ | 0.006077 | 0.412405 | 0.767 | 0.762 | 1 |
| Terminal | UBE2D3 | 3.23E-16 | 0.412388 | 0.626 | 0.461 | 5.65E-12 |
| Terminal | CNIH1 | 5.63E-12 | 0.411985 | 0.545 | 0.429 | 9.84E-08 |
| Fibroblasts | KAZALD1 | 2.37E-29 | 0.411893 | 0.1 | 0.003 | 4.14E-25 |
| Contractile | LRRK2 | 2.23E-24 | 0.411857 | 0.432 | 0.259 | 3.89E-20 |

| | | | | | | |
|---|---|---|---|---|---|---|
| Terminal | MAFG | 8.3E-26 | 0.411419 | 0.222 | 0.081 | 1.45E-21 |
| Terminal | NDUFB4 | 2.32E-17 | 0.411381 | 0.689 | 0.551 | 4.05E-13 |
| IFN_Interm | ENSSSCG00 | 3.13E-05 | 0.41133 | 0.105 | 0.01 | 0.54789 |
| Terminal | BTG3 | 1.26E-24 | 0.411291 | 0.348 | 0.17 | 2.19E-20 |
| Contractile | EXOC7 | 0.001138 | 0.410972 | 0.209 | 0.108 | 1 |
| Contractile | TST | 7.39E-09 | 0.410859 | 0.231 | 0.071 | 0.000129 |
| Fibroblasts | ENDOG | 0.005719 | 0.410043 | 0.117 | 0.044 | 1 |
| Contractile | ANKRD12 | 0.009856 | 0.40999 | 0.714 | 0.629 | 1 |
| Contractile | PRDM6 | 0.000184 | 0.409705 | 0.154 | 0.06 | 1 |
| Terminal | HMGB2 | 2.44E-05 | 0.409694 | 0.412 | 0.328 | 0.426094 |
| Terminal | SLC2A5 | 1.07E-33 | 0.409681 | 0.141 | 0.027 | 1.87E-29 |
| Terminal | ENSSSCG00 | 3.25E-28 | 0.409381 | 0.995 | 0.985 | 5.69E-24 |
| Contractile | UCHL1 | 3.94E-16 | 0.409371 | 0.363 | 0.234 | 6.88E-12 |
| Contractile | GNAI2 | 0.000713 | 0.40929 | 0.824 | 0.711 | 1 |
| Contractile | DDX6 | 0.002886 | 0.409004 | 0.396 | 0.273 | 1 |
| Contractile | ARHGEF10 | 1.88E-19 | 0.408967 | 0.342 | 0.2 | 3.29E-15 |
| Intermedia | SERPING1 | 1.96E-26 | 0.408944 | 0.901 | 0.786 | 3.42E-22 |
| Pericytes | ENSSSCG00 | 0.00163 | 0.408873 | 0.1 | 0.037 | 1 |
| Terminal | PROCR | 2.49E-22 | 0.408799 | 0.108 | 0.024 | 4.36E-18 |
| Pericytes | MRGPRF | 0.007647 | 0.408561 | 0.222 | 0.136 | 1 |
| Contractile | ENSSSCG00 | 0.000449 | 0.408465 | 0.538 | 0.367 | 1 |
| Contractile | PDLIM2 | 3.64E-23 | 0.408379 | 0.442 | 0.273 | 6.36E-19 |
| Pericytes | SLC25A25 | 2.69E-07 | 0.408301 | 0.111 | 0.024 | 0.004698 |
| Contractile | SRL | 7.48E-21 | 0.408257 | 0.132 | 0.011 | 1.31E-16 |
| Fibroblasts | PEG10 | 4.18E-06 | 0.407706 | 0.1 | 0.018 | 0.073116 |
| Terminal | GALNT1 | 6.91E-09 | 0.407593 | 0.521 | 0.415 | 0.000121 |
| Intermedia | ID1 | 2E-05 | 0.40722 | 0.322 | 0.265 | 0.350316 |
| Terminal | ENSSSCG00 | 7.04E-12 | 0.407213 | 0.542 | 0.415 | 1.23E-07 |
| Contractile | SERPINB9 | 3.61E-25 | 0.407186 | 0.279 | 0.13 | 6.31E-21 |
| Intermedia | ENSSSCG00 | 1.06E-16 | 0.407121 | 0.166 | 0.072 | 1.85E-12 |
| Terminal | SRPRA | 4E-15 | 0.406927 | 0.631 | 0.493 | 6.99E-11 |
| Contractile | RBM41 | 0.004326 | 0.406892 | 0.242 | 0.144 | 1 |
| Fibroblasts | ESM1 | 1.21E-07 | 0.406881 | 0.1 | 0.014 | 0.002123 |
| Contractile | MAP1A | 1.81E-20 | 0.406309 | 0.503 | 0.341 | 3.16E-16 |
| Fibroblasts | ENSSSCG00 | 0.000721 | 0.406126 | 0.167 | 0.062 | 1 |
| Terminal | HSPE1 | 1.37E-15 | 0.405682 | 0.69 | 0.566 | 2.39E-11 |
| Contractile | KATNAL1 | 0.00473 | 0.405293 | 0.22 | 0.124 | 1 |
| Contractile | RABEP1 | 0.007174 | 0.404933 | 0.451 | 0.349 | 1 |
| Contractile | JAZF1 | 6.9E-05 | 0.404128 | 0.319 | 0.169 | 1 |
| Contractile | PALM2AKA | 1.5E-05 | 0.403886 | 0.78 | 0.581 | 0.262669 |
| Contractile | PHIP | 0.000368 | 0.403777 | 0.297 | 0.164 | 1 |
| IFN_Interm | ENSSSCG00 | 8.11E-05 | 0.403659 | 0.105 | 0.011 | 1 |
| Pericytes | KLC4 | 0.000946 | 0.40358 | 0.122 | 0.049 | 1 |
| Terminal | ENSSSCG00 | 1.66E-13 | 0.403437 | 0.661 | 0.54 | 2.89E-09 |
| Terminal | PHLDA1 | 1.16E-23 | 0.403043 | 0.199 | 0.07 | 2.03E-19 |
| Intermedia | SPON1 | 8.34E-10 | 0.403014 | 0.263 | 0.175 | 1.46E-05 |
| Intermedia | SDF4 | 6.11E-22 | 0.402996 | 0.781 | 0.695 | 1.07E-17 |
| Intermedia | C4ORF48 | 8.73E-17 | 0.402851 | 0.312 | 0.183 | 1.53E-12 |
| Fibroblasts | FAM102A | 0.001831 | 0.402788 | 0.15 | 0.057 | 1 |
| Terminal | ENSSSCG00 | 1.34E-31 | 0.402777 | 0.996 | 0.985 | 2.34E-27 |
| Contractile | RIMKLB | 6.3E-29 | 0.40265 | 0.35 | 0.17 | 1.1E-24 |
| Contractile | PDLIM1 | 1.64E-25 | 0.402328 | 0.429 | 0.251 | 2.87E-21 |
| IFN_Interm | LOC106508 | 2.52E-06 | 0.40226 | 0.105 | 0.008 | 0.044041 |
| Intermedia | PCSK1N | 6.25E-14 | 0.402019 | 0.749 | 0.678 | 1.09E-09 |
| Terminal | DNASE1L1 | 1.97E-12 | 0.401989 | 0.471 | 0.338 | 3.44E-08 |
| Fibroblasts | PWWP3B | 2.63E-14 | 0.401868 | 0.1 | 0.007 | 4.6E-10 |
| Contractile | ISCU | 7.52E-28 | 0.401568 | 0.76 | 0.607 | 1.31E-23 |
| Terminal | PPA1 | 8.19E-26 | 0.40149 | 0.272 | 0.111 | 1.43E-21 |
| Contractile | INPP4A | 2.56E-22 | 0.401471 | 0.234 | 0.107 | 4.48E-18 |

| | | | | | | |
|---|---|---|---|---|---|---|
| Pericytes | TRMT6 | 0.00566 | 0.401355 | 0.156 | 0.08 | 1 |
| Contractile | MAP4 | 2.13E-23 | 0.401222 | 0.676 | 0.501 | 3.72E-19 |
| Contractile | C1R | 5.13E-36 | 0.401197 | 0.903 | 0.708 | 8.96E-32 |
| Contractile | PRKCD | 3.46E-32 | 0.400983 | 0.251 | 0.094 | 6.04E-28 |
| Contractile | MEGF6 | 7.77E-27 | 0.400894 | 0.212 | 0.078 | 1.36E-22 |
| Contractile | ANK2 | 0.000112 | 0.400315 | 0.198 | 0.085 | 1 |
| Contractile | METTL24 | 4.28E-08 | 0.400301 | 0.176 | 0.048 | 0.000748 |
| Contractile | ZYX | 0.003257 | 0.400292 | 0.527 | 0.399 | 1 |
| Contractile | USF2 | 0.005001 | 0.400101 | 0.396 | 0.288 | 1 |
| Pericytes | ENSSSCG0( | 2.53E-12 | 0.399656 | 0.1 | 0.012 | 4.43E-08 |
| Terminal | TMED9 | 1.91E-17 | 0.399575 | 0.369 | 0.214 | 3.34E-13 |
| Intermedia | FLRT2 | 5.75E-17 | 0.398893 | 0.154 | 0.062 | 1.01E-12 |
| Contractile | TRIM24 | 0.000836 | 0.398863 | 0.176 | 0.081 | 1 |
| Intermedia | ENSSSCG0( | 6.31E-12 | 0.398598 | 0.464 | 0.355 | 1.1E-07 |
| Contractile | CDK18 | 5.25E-06 | 0.397988 | 0.11 | 0.028 | 0.091698 |
| Contractile | ATP2B4 | 0.001193 | 0.397631 | 0.659 | 0.541 | 1 |
| Contractile | PDLIM5 | 0.003901 | 0.39748 | 0.615 | 0.495 | 1 |
| Contractile | APP | 0.000439 | 0.397053 | 0.901 | 0.852 | 1 |
| Contractile | SELENOW | 0.001115 | 0.397032 | 0.593 | 0.48 | 1 |
| Terminal | LAMTOR5 | 3.35E-14 | 0.396804 | 0.613 | 0.485 | 5.85E-10 |
| Contractile | COL8A1 | 1.43E-31 | 0.396608 | 0.938 | 0.817 | 2.5E-27 |
| Terminal | CREB3L1 | 3.9E-18 | 0.396479 | 0.364 | 0.203 | 6.81E-14 |
| Contractile | IST1 | 0.005508 | 0.39577 | 0.231 | 0.139 | 1 |
| Contractile | TNK2 | 0.004873 | 0.395624 | 0.176 | 0.094 | 1 |
| Terminal | TXNDC17 | 9.47E-17 | 0.39559 | 0.678 | 0.532 | 1.65E-12 |
| Contractile | MEIS1 | 0.000646 | 0.395528 | 0.22 | 0.108 | 1 |
| Contractile | SAMD4A | 6.06E-16 | 0.395112 | 0.521 | 0.386 | 1.06E-11 |
| Terminal | AGFG1 | 2.19E-23 | 0.394851 | 0.344 | 0.168 | 3.83E-19 |
| Contractile | DIAPH2 | 1.06E-20 | 0.394783 | 0.606 | 0.436 | 1.86E-16 |
| Contractile | KCNG1 | 6.65E-33 | 0.394762 | 0.109 | 0.013 | 1.16E-28 |
| Contractile | KLHL21 | 0.000235 | 0.394652 | 0.22 | 0.103 | 1 |
| Contractile | APLP2 | 9.19E-28 | 0.394372 | 0.715 | 0.534 | 1.61E-23 |
| Terminal | IARS1 | 3.74E-15 | 0.394363 | 0.326 | 0.187 | 6.54E-11 |
| Terminal | ENSSSCG0( | 2.46E-12 | 0.394226 | 0.383 | 0.258 | 4.31E-08 |
| Terminal | DSEL | 1.91E-20 | 0.394199 | 0.255 | 0.116 | 3.33E-16 |
| IFN_Interm | ZCWPW1 | 1.77E-10 | 0.394194 | 0.105 | 0.004 | 3.1E-06 |
| Contractile | DAPK3 | 1.45E-24 | 0.394084 | 0.297 | 0.146 | 2.54E-20 |
| Pericytes | NGF | 0.000772 | 0.393695 | 0.1 | 0.035 | 1 |
| Fibroblasts | ENSSSCG0( | 0.008323 | 0.393479 | 0.133 | 0.055 | 1 |
| Contractile | RTL6 | 5.28E-06 | 0.393453 | 0.154 | 0.049 | 0.09227 |
| Contractile | NDUFB8 | 0.002649 | 0.392945 | 0.44 | 0.307 | 1 |
| Contractile | KANK2 | 5.75E-22 | 0.392888 | 0.589 | 0.415 | 1E-17 |
| Contractile | SENP6 | 0.001167 | 0.392698 | 0.484 | 0.334 | 1 |
| Contractile | LTBP1 | 2.08E-31 | 0.39268 | 0.874 | 0.697 | 3.64E-27 |
| IFN_Interm | CD274 | 0.000454 | 0.392663 | 0.105 | 0.013 | 1 |
| Contractile | CCND1 | 6.41E-27 | 0.392555 | 0.717 | 0.519 | 1.12E-22 |
| Contractile | TEAD3 | 0.004893 | 0.392544 | 0.275 | 0.169 | 1 |
| IFN_Interm | NMI | 0.000129 | 0.392373 | 0.263 | 0.056 | 1 |
| Intermedia | C1QTNF4 | 3.08E-16 | 0.392373 | 0.165 | 0.071 | 5.39E-12 |
| Contractile | TPP1 | 3.82E-22 | 0.392048 | 0.533 | 0.366 | 6.68E-18 |
| Intermedia | HOXA5 | 4.79E-09 | 0.392047 | 0.35 | 0.26 | 8.38E-05 |
| Contractile | ACTN1 | 0.000139 | 0.391877 | 0.901 | 0.796 | 1 |
| Contractile | HSD17B4 | 3.04E-23 | 0.391817 | 0.331 | 0.175 | 5.32E-19 |
| Contractile | MECOM | 2.65E-06 | 0.391774 | 0.121 | 0.031 | 0.046406 |
| Terminal | BHLHE40 | 8.14E-14 | 0.391722 | 0.764 | 0.646 | 1.42E-09 |
| Contractile | DACT3 | 3.15E-28 | 0.391701 | 0.233 | 0.092 | 5.5E-24 |
| Contractile | WDTC1 | 0.000618 | 0.391641 | 0.209 | 0.1 | 1 |
| Fibroblasts | RPA1 | 0.001678 | 0.391496 | 0.183 | 0.076 | 1 |
| Terminal | MEF2A | 1.3E-09 | 0.391402 | 0.646 | 0.56 | 2.27E-05 |

| | | | | | | |
|---|---|---|---|---|---|---|
| Pericytes | TTC5 | 0.000228 | 0.391131 | 0.111 | 0.037 | 1 |
| Contractile | MAP1B | 4.15E-06 | 0.39069 | 0.934 | 0.815 | 0.072509 |
| Contractile | PHF3 | 0.000309 | 0.390628 | 0.516 | 0.338 | 1 |
| Fibroblasts | FTH1 | 9.09E-05 | 0.390448 | 1 | 0.998 | 1 |
| Pericytes | FAM110B | 5.71E-12 | 0.390444 | 0.111 | 0.015 | 9.97E-08 |
| Contractile | CASP8AP2 | 4.92E-05 | 0.39028 | 0.242 | 0.109 | 0.859627 |
| Terminal | ARPC1B | 2.85E-16 | 0.390257 | 0.645 | 0.505 | 4.99E-12 |
| Contractile | ECM2 | 8.74E-29 | 0.390054 | 0.869 | 0.708 | 1.53E-24 |
| Intermedia | GALNT1 | 8.06E-10 | 0.389922 | 0.495 | 0.418 | 1.41E-05 |
| Contractile | ACCSL | 7.52E-20 | 0.389764 | 0.126 | 0.04 | 1.32E-15 |
| Contractile | RYR2 | 0.000771 | 0.389743 | 0.198 | 0.095 | 1 |
| Contractile | ENSSSCG0( | 0.000401 | 0.389622 | 0.242 | 0.123 | 1 |
| Intermedia | ANTXR1 | 1.25E-11 | 0.389609 | 0.637 | 0.53 | 2.19E-07 |
| Contractile | WIF1 | 3.58E-05 | 0.389563 | 0.835 | 0.636 | 0.626101 |
| Fibroblasts | DNASE2 | 0.009029 | 0.389469 | 0.183 | 0.09 | 1 |
| Contractile | SLC20A2 | 4.77E-05 | 0.389389 | 0.165 | 0.061 | 0.834496 |
| Contractile | LIMCH1 | 0.000244 | 0.389261 | 0.725 | 0.56 | 1 |
| Contractile | INPP5A | 1.38E-24 | 0.389187 | 0.26 | 0.12 | 2.41E-20 |
| Contractile | LAMA5 | 5.07E-25 | 0.388847 | 0.622 | 0.421 | 8.86E-21 |
| Contractile | RRAD | 0.002034 | 0.388343 | 0.176 | 0.085 | 1 |
| Fibroblasts | CEP68 | 0.004444 | 0.388184 | 0.133 | 0.052 | 1 |
| Contractile | APPBP2 | 0.001068 | 0.388184 | 0.242 | 0.132 | 1 |
| Terminal | TMEM167. | 4.34E-12 | 0.388132 | 0.517 | 0.384 | 7.59E-08 |
| Contractile | ASB2 | 4.24E-06 | 0.388011 | 0.11 | 0.027 | 0.074048 |
| Fibroblasts | YPEL3 | 0.002163 | 0.387995 | 0.6 | 0.412 | 1 |
| Fibroblasts | ENSSSCG0( | 2.12E-05 | 0.387864 | 0.117 | 0.026 | 0.370997 |
| Terminal | PCNA.1 | 3.13E-08 | 0.387833 | 0.182 | 0.107 | 0.000547 |
| Fibroblasts | ENSSSCG0( | 0.004759 | 0.387407 | 0.117 | 0.043 | 1 |
| Pericytes | FADS1 | 0.000502 | 0.387255 | 0.1 | 0.033 | 1 |
| Terminal | BIRC3 | 1.47E-09 | 0.386978 | 0.273 | 0.174 | 2.57E-05 |
| Fibroblasts | CERS5 | 0.003051 | 0.38645 | 0.233 | 0.114 | 1 |
| Terminal | STX8 | 8.53E-17 | 0.386405 | 0.376 | 0.222 | 1.49E-12 |
| Contractile | CFLAR | 0.000653 | 0.386367 | 0.275 | 0.152 | 1 |
| Pericytes | TSPAN14 | 0.000406 | 0.386276 | 0.133 | 0.052 | 1 |
| Contractile | CD47 | 5.66E-19 | 0.385874 | 0.597 | 0.452 | 9.89E-15 |
| Terminal | P4HB | 1.52E-14 | 0.38573 | 0.678 | 0.557 | 2.65E-10 |
| Contractile | MAP7D1 | 0.001404 | 0.385629 | 0.626 | 0.514 | 1 |
| Intermedia | LTBP2 | 2.68E-17 | 0.385607 | 0.817 | 0.701 | 4.68E-13 |
| Terminal | PGLS | 6.7E-14 | 0.385484 | 0.401 | 0.26 | 1.17E-09 |
| Terminal | ATP5PD | 6.74E-16 | 0.385438 | 0.671 | 0.547 | 1.18E-11 |
| Intermedia | DUSP1 | 0.000217 | 0.385428 | 0.499 | 0.459 | 1 |
| IFN_Interm | AGK | 0.009176 | 0.385052 | 0.158 | 0.04 | 1 |
| Fibroblasts | ENSSSCG0( | 2.22E-20 | 0.384705 | 0.1 | 0.005 | 3.88E-16 |
| Contractile | PCMTD1 | 0.001449 | 0.384647 | 0.582 | 0.445 | 1 |
| Terminal | EIF3M | 5.54E-10 | 0.384646 | 0.532 | 0.421 | 9.68E-06 |
| Fibroblasts | DUSP4 | 1.56E-07 | 0.384599 | 0.117 | 0.019 | 0.002725 |
| Terminal | RNFT1 | 2.37E-19 | 0.384089 | 0.278 | 0.136 | 4.14E-15 |
| Terminal | ENSSSCG0( | 2.25E-22 | 0.383859 | 0.993 | 0.979 | 3.94E-18 |
| Contractile | PRPF8 | 0.002024 | 0.383456 | 0.308 | 0.185 | 1 |
| Pericytes | ENSSSCG0( | 0.000139 | 0.383199 | 0.111 | 0.036 | 1 |
| IFN_Interm | OAS1 | 5.23E-17 | 0.383167 | 0.105 | 0.002 | 9.13E-13 |
| Fibroblasts | PAIP2B | 0.001904 | 0.38311 | 0.117 | 0.038 | 1 |
| Terminal | SERP1 | 4.17E-14 | 0.383036 | 0.54 | 0.387 | 7.28E-10 |
| Pericytes | ACOX2 | 0.004202 | 0.382981 | 0.144 | 0.071 | 1 |
| Terminal | ATAD2 | 2.23E-09 | 0.382973 | 0.128 | 0.061 | 3.9E-05 |
| Contractile | NES | 8.63E-27 | 0.382688 | 0.163 | 0.049 | 1.51E-22 |
| Intermedia | P2RY1 | 5.79E-26 | 0.38266 | 0.11 | 0.023 | 1.01E-21 |
| Contractile | DYRK2 | 0.002211 | 0.382415 | 0.308 | 0.192 | 1 |
| Intermedia | RRBP1 | 5.67E-20 | 0.382185 | 0.91 | 0.828 | 9.9E-16 |

| | | | | | | |
|---|---|---|---|---|---|---|
| Contractile | DEK | 0.000643 | 0.382113 | 0.857 | 0.78 | 1 |
| Contractile | PDE5A | 1.48E-23 | 0.381905 | 0.3 | 0.152 | 2.59E-19 |
| Contractile | DYRK3 | 0.002442 | 0.381535 | 0.396 | 0.272 | 1 |
| Contractile | TFDP2 | 0.000844 | 0.381121 | 0.286 | 0.157 | 1 |
| Terminal | UQCC2 | 7.16E-14 | 0.381019 | 0.458 | 0.31 | 1.25E-09 |
| Terminal | PSPH | 4.3E-27 | 0.381008 | 0.169 | 0.048 | 7.51E-23 |
| Fibroblasts | IQCD | 7.38E-08 | 0.380948 | 0.1 | 0.014 | 0.001291 |
| Terminal | SSPN | 7.47E-13 | 0.380749 | 0.749 | 0.686 | 1.31E-08 |
| Pericytes | TMEM106 | 0.005428 | 0.380669 | 0.489 | 0.392 | 1 |
| Fibroblasts | EVL | 0.002018 | 0.380629 | 0.167 | 0.067 | 1 |
| Terminal | GLIPR2 | 3.51E-14 | 0.380624 | 0.477 | 0.324 | 6.14E-10 |
| Contractile | HNRNPUL2 | 0.004637 | 0.380385 | 0.473 | 0.348 | 1 |
| Pericytes | UBE2D3 | 0.004231 | 0.380219 | 0.567 | 0.497 | 1 |
| Terminal | UBE2E1 | 2.97E-15 | 0.380088 | 0.362 | 0.218 | 5.2E-11 |
| Contractile | PNPLA2 | 0.000838 | 0.38008 | 0.374 | 0.234 | 1 |
| Pericytes | MYO6 | 0.008536 | 0.380068 | 0.178 | 0.1 | 1 |
| Intermedia | CPXM2 | 4.9E-18 | 0.379977 | 0.819 | 0.732 | 8.56E-14 |
| Terminal | MAMDC2 | 2.12E-29 | 0.379891 | 0.187 | 0.053 | 3.71E-25 |
| Fibroblasts | NAALADL2 | 1.31E-05 | 0.379634 | 0.117 | 0.025 | 0.229595 |
| Contractile | SCHIP1 | 6.19E-19 | 0.379525 | 0.309 | 0.175 | 1.08E-14 |
| Intermedia | NFIX | 1.34E-09 | 0.3795 | 0.475 | 0.389 | 2.35E-05 |
| Contractile | ZNF292 | 0.000427 | 0.379198 | 0.429 | 0.28 | 1 |
| Intermedia | TSC22D1 | 7.28E-14 | 0.37912 | 0.594 | 0.477 | 1.27E-09 |
| Terminal | NUDC | 1.37E-08 | 0.378833 | 0.472 | 0.372 | 0.000239 |
| Contractile | TEAD3 | 3.11E-24 | 0.378351 | 0.256 | 0.116 | 5.43E-20 |
| Contractile | RAB18 | 0.002085 | 0.378048 | 0.626 | 0.526 | 1 |
| Contractile | ATP2A2 | 1.07E-17 | 0.377982 | 0.454 | 0.308 | 1.87E-13 |
| Pericytes | NPY1R | 0.000716 | 0.377967 | 0.122 | 0.047 | 1 |
| Contractile | HTR1B | 9.82E-25 | 0.377951 | 0.21 | 0.083 | 1.72E-20 |
| IFN_Interm | CLN3 | 0.000162 | 0.377932 | 0.263 | 0.057 | 1 |
| Contractile | ZCCHC24 | 7.05E-22 | 0.377545 | 0.498 | 0.321 | 1.23E-17 |
| Terminal | NANS | 1.41E-19 | 0.377426 | 0.313 | 0.159 | 2.46E-15 |
| Contractile | ROCK2 | 2.12E-19 | 0.376418 | 0.672 | 0.519 | 3.7E-15 |
| Contractile | KLF15 | 0.00147 | 0.376268 | 0.154 | 0.07 | 1 |
| Terminal | ANXA7 | 3.36E-13 | 0.376135 | 0.475 | 0.334 | 5.88E-09 |
| Contractile | ESF1 | 0.003653 | 0.376074 | 0.462 | 0.328 | 1 |
| Terminal | DGCR6L | 3.17E-13 | 0.375811 | 0.379 | 0.24 | 5.54E-09 |
| Contractile | DNAJB4 | 8.74E-20 | 0.375591 | 0.515 | 0.362 | 1.53E-15 |
| Contractile | ACSS3 | 1.54E-11 | 0.375411 | 0.143 | 0.024 | 2.69E-07 |
| Fibroblasts | EPHA7 | 0.009931 | 0.375128 | 0.2 | 0.101 | 1 |
| Terminal | ENSSSCG0( | 8.43E-09 | 0.375072 | 0.499 | 0.394 | 0.000147 |
| Intermedia | PTGIS | 4.03E-12 | 0.37498 | 0.748 | 0.679 | 7.04E-08 |
| Terminal | ENSSSCG0( | 1.05E-08 | 0.374945 | 0.298 | 0.196 | 0.000184 |
| Fibroblasts | MARCHF9 | 0.002619 | 0.374915 | 0.1 | 0.032 | 1 |
| Terminal | PFKL | 4.3E-24 | 0.374834 | 0.252 | 0.102 | 7.52E-20 |
| Contractile | ITFG1 | 7.14E-19 | 0.374748 | 0.552 | 0.396 | 1.25E-14 |
| Pericytes | GNG11 | 0.00548 | 0.374631 | 0.689 | 0.656 | 1 |
| Terminal | YKT6 | 1.45E-12 | 0.374631 | 0.483 | 0.351 | 2.53E-08 |
| Terminal | UBE2M | 1.13E-12 | 0.37453 | 0.484 | 0.343 | 1.97E-08 |
| Contractile | PRRC2B | 0.001697 | 0.374529 | 0.374 | 0.244 | 1 |
| Contractile | SGTB | 1.21E-11 | 0.374399 | 0.121 | 0.018 | 2.12E-07 |
| Terminal | LOC10052: | 2.22E-23 | 0.374392 | 0.938 | 0.874 | 3.88E-19 |
| Terminal | ITGA5 | 5.23E-10 | 0.374199 | 0.512 | 0.389 | 9.13E-06 |
| Terminal | TMED3 | 1.09E-13 | 0.374097 | 0.462 | 0.318 | 1.91E-09 |
| Contractile | ENSSSCG0( | 3.59E-05 | 0.374005 | 0.209 | 0.085 | 0.62832 |
| Terminal | CTSD | 6.19E-08 | 0.373948 | 0.927 | 0.913 | 0.001082 |
| Fibroblasts | UXS1 | 0.003257 | 0.373897 | 0.217 | 0.104 | 1 |
| Terminal | TUBB3 | 5.78E-34 | 0.373815 | 0.129 | 0.021 | 1.01E-29 |
| Intermedia | S100A14 | 2.08E-12 | 0.373719 | 0.203 | 0.112 | 3.64E-08 |

| | | | | | | |
|---|---|---|---|---|---|---|
| Contractile | CD93 | 0.001233 | 0.373506 | 0.286 | 0.163 | 1 |
| Contractile | ENSSSCG0( | 0.009897 | 0.373469 | 0.363 | 0.258 | 1 |
| Terminal | GADD45G | 5.12E-11 | 0.37345 | 0.287 | 0.178 | 8.95E-07 |
| Fibroblasts | ORAI1 | 0.001675 | 0.373302 | 0.117 | 0.039 | 1 |
| Contractile | NAV2 | 2.35E-22 | 0.373189 | 0.213 | 0.09 | 4.11E-18 |
| Contractile | GALM | 0.00079 | 0.373124 | 0.253 | 0.137 | 1 |
| Contractile | MTMR7 | 1.3E-06 | 0.373099 | 0.121 | 0.03 | 0.022702 |
| Terminal | SNRPD2 | 3.44E-16 | 0.372789 | 0.789 | 0.695 | 6.02E-12 |
| Contractile | PPFIBP2 | 0.003592 | 0.372708 | 0.176 | 0.091 | 1 |
| Terminal | ENSSSCG0( | 3.87E-15 | 0.372692 | 0.842 | 0.738 | 6.76E-11 |
| Contractile | TBL1XR1 | 0.00989 | 0.372202 | 0.582 | 0.498 | 1 |
| Terminal | HSP90AB1 | 2.5E-13 | 0.372089 | 0.834 | 0.773 | 4.37E-09 |
| Contractile | SESN1 | 0.006364 | 0.372015 | 0.33 | 0.217 | 1 |
| Fibroblasts | LPL | 0.001101 | 0.371949 | 0.1 | 0.028 | 1 |
| Contractile | NCS1 | 0.003469 | 0.371612 | 0.231 | 0.135 | 1 |
| Contractile | FAIM | 4.8E-07 | 0.371608 | 0.187 | 0.059 | 0.008386 |
| Contractile | CCDC85B | 0.002768 | 0.371532 | 0.407 | 0.276 | 1 |
| Terminal | COL1A2 | 1.82E-12 | 0.37153 | 0.982 | 0.983 | 3.18E-08 |
| Contractile | VWA1 | 0.000427 | 0.371527 | 0.132 | 0.051 | 1 |
| Pericytes | EEF1B2 | 0.001528 | 0.371299 | 0.9 | 0.876 | 1 |
| Fibroblasts | ENSSSCG0( | 0.001671 | 0.371258 | 0.133 | 0.046 | 1 |
| Contractile | RBBP4 | 0.008613 | 0.37123 | 0.462 | 0.354 | 1 |
| Contractile | LRCH1 | 4.9E-22 | 0.371108 | 0.276 | 0.137 | 8.57E-18 |
| Contractile | MXRA7 | 0.00512 | 0.370833 | 0.78 | 0.743 | 1 |
| Contractile | EIF4B | 0.003089 | 0.370459 | 0.505 | 0.39 | 1 |
| Terminal | GALNS | 8.88E-24 | 0.370454 | 0.23 | 0.089 | 1.55E-19 |
| Terminal | COPZ2 | 5.54E-11 | 0.370438 | 0.66 | 0.568 | 9.69E-07 |
| Intermedia | GXYLT2 | 5.65E-07 | 0.370036 | 0.337 | 0.269 | 0.009881 |
| Intermedia | ENSSSCG0( | 2.84E-09 | 0.369892 | 0.243 | 0.163 | 4.97E-05 |
| Terminal | ENSSSCG0( | 3.78E-28 | 0.369772 | 0.999 | 0.986 | 6.61E-24 |
| Contractile | TMEM47 | 7.12E-19 | 0.369612 | 0.68 | 0.552 | 1.24E-14 |
| Contractile | APOA1 | 1.77E-31 | 0.369528 | 0.361 | 0.178 | 3.1E-27 |
| Terminal | CALM3 | 1.71E-12 | 0.369478 | 0.339 | 0.212 | 2.99E-08 |
| Terminal | MRPL23 | 7.59E-12 | 0.369252 | 0.598 | 0.474 | 1.33E-07 |
| Contractile | ADA | 7.91E-08 | 0.369151 | 0.176 | 0.049 | 0.001383 |
| Terminal | SNF8 | 1.33E-12 | 0.368906 | 0.56 | 0.428 | 2.32E-08 |
| Intermedia | ITGA10 | 7.17E-14 | 0.368573 | 0.289 | 0.175 | 1.25E-09 |
| IFN_Interm | RPTOR | 0.004701 | 0.368394 | 0.158 | 0.036 | 1 |
| Contractile | COL4A2 | 0.000218 | 0.36827 | 0.945 | 0.843 | 1 |
| Contractile | HRH1 | 5.13E-05 | 0.368268 | 0.121 | 0.038 | 0.896809 |
| Contractile | RNF103 | 0.00269 | 0.36794 | 0.209 | 0.112 | 1 |
| Terminal | AP1S2 | 9.41E-15 | 0.367926 | 0.377 | 0.231 | 1.65E-10 |
| Pericytes | SIPA1 | 6.99E-06 | 0.367739 | 0.111 | 0.029 | 0.122213 |
| Contractile | RAP1A | 1.6E-20 | 0.367629 | 0.576 | 0.426 | 2.8E-16 |
| Contractile | PNN | 0.001029 | 0.367511 | 0.736 | 0.595 | 1 |
| Fibroblasts | CDON | 0.000938 | 0.367461 | 0.133 | 0.045 | 1 |
| Fibroblasts | CDC42EP1 | 0.001735 | 0.367411 | 0.283 | 0.142 | 1 |
| Terminal | ATP5MK | 7.65E-11 | 0.367287 | 0.711 | 0.612 | 1.34E-06 |
| Terminal | MEDAG | 1.33E-26 | 0.367277 | 0.174 | 0.051 | 2.33E-22 |
| Contractile | EBF1 | 0.000728 | 0.36722 | 0.341 | 0.2 | 1 |
| Contractile | ENSSSCG0( | 0.000568 | 0.36721 | 0.758 | 0.656 | 1 |
| Intermedia | FGF7 | 6.28E-10 | 0.366649 | 0.169 | 0.093 | 1.1E-05 |
| Terminal | SYPL1 | 4.46E-13 | 0.366169 | 0.443 | 0.3 | 7.8E-09 |
| Contractile | MCCC2 | 0.000424 | 0.365905 | 0.176 | 0.077 | 1 |
| Contractile | RASL11A | 0.001151 | 0.3659 | 0.538 | 0.363 | 1 |
| Intermedia | ENSSSCG0( | 7.2E-07 | 0.365763 | 0.194 | 0.129 | 0.012581 |
| Contractile | KIF13A | 0.003739 | 0.36576 | 0.33 | 0.208 | 1 |
| Contractile | DYNLL2 | 6.61E-21 | 0.365176 | 0.395 | 0.24 | 1.16E-16 |
| Fibroblasts | ERO1B | 0.000246 | 0.364674 | 0.15 | 0.048 | 1 |

| | | | | | | |
|---|---|---|---|---|---|---|
| Terminal | ZNF511 | 4.53E-21 | 0.364451 | 0.309 | 0.151 | 7.92E-17 |
| Contractile | DES | 2.26E-13 | 0.364284 | 0.109 | 0.041 | 3.95E-09 |
| Intermedia | ZFP36L2 | 4.98E-15 | 0.364252 | 0.799 | 0.733 | 8.7E-11 |
| Contractile | KAT6B | 0.001632 | 0.36423 | 0.341 | 0.216 | 1 |
| Contractile | TADA3 | 0.007092 | 0.364071 | 0.198 | 0.114 | 1 |
| Fibroblasts | PLTP | 0.000231 | 0.364034 | 0.15 | 0.047 | 1 |
| Contractile | OAT | 0.001757 | 0.363757 | 0.648 | 0.524 | 1 |
| Contractile | KRR1 | 0.002739 | 0.363413 | 0.253 | 0.148 | 1 |
| Contractile | MAP4K3 | 0.001386 | 0.363399 | 0.275 | 0.156 | 1 |
| Contractile | TOB2 | 0.001825 | 0.362812 | 0.571 | 0.437 | 1 |
| Terminal | CALU | 2.19E-14 | 0.36278 | 0.793 | 0.72 | 3.83E-10 |
| Contractile | PTPRS | 0.004403 | 0.362705 | 0.121 | 0.053 | 1 |
| Terminal | P3H4 | 2.13E-13 | 0.362414 | 0.396 | 0.255 | 3.73E-09 |
| Fibroblasts | ATAD3A | 0.002024 | 0.362402 | 0.1 | 0.03 | 1 |
| Intermedia | MPZL1 | 2.8E-11 | 0.36236 | 0.357 | 0.258 | 4.9E-07 |
| Contractile | DYRK2 | 2.2E-18 | 0.36188 | 0.27 | 0.145 | 3.85E-14 |
| Contractile | PCDH7 | 0.003453 | 0.361828 | 0.56 | 0.428 | 1 |
| Intermedia | TIMP3 | 1.7E-17 | 0.361772 | 0.621 | 0.461 | 2.97E-13 |
| Contractile | FADS3 | 2.93E-22 | 0.361643 | 0.518 | 0.34 | 5.13E-18 |
| Terminal | SPPL2A | 5.12E-09 | 0.361553 | 0.531 | 0.42 | 8.94E-05 |
| Contractile | FBXO30 | 1.29E-05 | 0.361257 | 0.385 | 0.196 | 0.225973 |
| Fibroblasts | ASNSD1 | 0.007615 | 0.361213 | 0.167 | 0.076 | 1 |
| Contractile | VSIR | 0.00085 | 0.36091 | 0.242 | 0.128 | 1 |
| IFN_Interm | ENSSSCG0( | 4.43E-06 | 0.360848 | 0.105 | 0.008 | 0.077481 |
| Contractile | EIF4H | 0.001845 | 0.360149 | 0.582 | 0.462 | 1 |
| Contractile | CBX6 | 1.33E-05 | 0.36007 | 0.231 | 0.095 | 0.232344 |
| Contractile | A4GALT | 0.004331 | 0.359938 | 0.538 | 0.431 | 1 |
| Terminal | ATP5MGL | 1.26E-14 | 0.359855 | 0.808 | 0.719 | 2.21E-10 |
| Terminal | MGAT4A | 1.05E-15 | 0.359817 | 0.178 | 0.077 | 1.84E-11 |
| Terminal | ATP5MC3 | 2.05E-13 | 0.359587 | 0.754 | 0.66 | 3.59E-09 |
| Fibroblasts | DBP | 0.001038 | 0.359556 | 0.2 | 0.083 | 1 |
| Pericytes | UBA7 | 1.04E-06 | 0.359467 | 0.1 | 0.022 | 0.01812 |
| Contractile | COX6C | 0.00146 | 0.359414 | 0.835 | 0.781 | 1 |
| Terminal | FUOM | 2.5E-21 | 0.359405 | 0.221 | 0.09 | 4.37E-17 |
| Intermedia | MMP16 | 5.97E-16 | 0.359251 | 0.178 | 0.082 | 1.04E-11 |
| Pericytes | ENSSSCG0( | 0.000219 | 0.359229 | 0.122 | 0.042 | 1 |
| Contractile | FN3K | 1.63E-07 | 0.358828 | 0.154 | 0.04 | 0.002852 |
| Contractile | GALNT2 | 0.001563 | 0.358718 | 0.418 | 0.283 | 1 |
| Contractile | ATOH8 | 8.31E-05 | 0.358663 | 0.154 | 0.057 | 1 |
| Contractile | RAP1A | 0.004533 | 0.358536 | 0.593 | 0.483 | 1 |
| Contractile | ESCO1 | 0.007014 | 0.358507 | 0.231 | 0.138 | 1 |
| Fibroblasts | ACADSB | 0.002751 | 0.357812 | 0.167 | 0.068 | 1 |
| Terminal | COPE | 6.71E-14 | 0.357714 | 0.686 | 0.569 | 1.17E-09 |
| Contractile | DENND2B | 1.54E-20 | 0.357644 | 0.282 | 0.146 | 2.69E-16 |
| Pericytes | ENSSSCG0( | 0.000324 | 0.357631 | 0.1 | 0.032 | 1 |
| Terminal | CIB1 | 1.52E-14 | 0.357453 | 0.353 | 0.212 | 2.66E-10 |
| Intermedia | ANG | 2.42E-10 | 0.357368 | 0.367 | 0.266 | 4.22E-06 |
| Contractile | PRKACA | 0.006867 | 0.357152 | 0.319 | 0.208 | 1 |
| Intermedia | ABCA1 | 2.17E-08 | 0.356885 | 0.446 | 0.355 | 0.000379 |
| Contractile | PCMTD2 | 4.05E-05 | 0.356786 | 0.198 | 0.08 | 0.708185 |
| Pericytes | ENSSSCG0( | 4.65E-05 | 0.356699 | 0.122 | 0.038 | 0.812346 |
| Contractile | PLPPR4 | 1.42E-38 | 0.356694 | 0.11 | 0.003 | 2.49E-34 |
| Contractile | CDHR1 | 1.85E-08 | 0.356584 | 0.143 | 0.032 | 0.000323 |
| Terminal | CLTB | 4.06E-09 | 0.356542 | 0.502 | 0.393 | 7.1E-05 |
| Terminal | LHFPL2 | 2.84E-21 | 0.356183 | 0.177 | 0.064 | 4.96E-17 |
| Terminal | CHIC2 | 1.05E-12 | 0.35584 | 0.362 | 0.227 | 1.83E-08 |
| Intermedia | NUAK1 | 1.04E-10 | 0.355577 | 0.239 | 0.15 | 1.81E-06 |
| Terminal | GPX4 | 8.43E-15 | 0.355558 | 0.739 | 0.616 | 1.47E-10 |
| Terminal | ZMIZ1 | 8.06E-11 | 0.35547 | 0.274 | 0.166 | 1.41E-06 |

| | | | | | | |
|---|---|---|---|---|---|---|
| Terminal | ASNS | 4.34E-18 | 0.355333 | 0.196 | 0.083 | 7.58E-14 |
| Contractile | CCDC146 | 7.59E-08 | 0.354954 | 0.121 | 0.026 | 0.001326 |
| Fibroblasts | ENSSSCG0( | 0.001691 | 0.354595 | 0.133 | 0.047 | 1 |
| Fibroblasts | FAN1 | 0.001207 | 0.354324 | 0.167 | 0.064 | 1 |
| Contractile | CSRP2 | 1.21E-18 | 0.354306 | 0.766 | 0.633 | 2.11E-14 |
| Terminal | RAN | 4.35E-11 | 0.354289 | 0.702 | 0.593 | 7.6E-07 |
| Contractile | POLR2B | 1.39E-19 | 0.354024 | 0.509 | 0.35 | 2.43E-15 |
| Contractile | MAP4 | 0.001992 | 0.354007 | 0.692 | 0.568 | 1 |
| Terminal | ENSSSCG0( | 2.54E-16 | 0.353675 | 0.905 | 0.869 | 4.45E-12 |
| Contractile | LIN7A | 1.35E-08 | 0.353304 | 0.121 | 0.024 | 0.000237 |
| Fibroblasts | FAM20B | 0.000511 | 0.353264 | 0.167 | 0.058 | 1 |
| Contractile | DSG2 | 0.00102 | 0.3532 | 0.231 | 0.121 | 1 |
| Contractile | MBNL1 | 0.003044 | 0.353052 | 0.901 | 0.818 | 1 |
| Contractile | ENSSSCG0( | 4.64E-26 | 0.353009 | 0.179 | 0.059 | 8.11E-22 |
| IFN_Interm | C1QTNF3 | 0.002755 | 0.35289 | 0.211 | 0.053 | 1 |
| Terminal | COX5A | 1.63E-10 | 0.352587 | 0.573 | 0.448 | 2.85E-06 |
| Fibroblasts | KIAA1755 | 2.41E-21 | 0.35225 | 0.1 | 0.004 | 4.21E-17 |
| Contractile | ENSSSCG0( | 0.001563 | 0.352009 | 0.286 | 0.166 | 1 |
| Contractile | CAB39 | 0.003774 | 0.351912 | 0.33 | 0.214 | 1 |
| Contractile | DBI | 0.009266 | 0.35179 | 0.637 | 0.554 | 1 |
| Terminal | TSPAN2 | 0.001989 | 0.351699 | 0.119 | 0.084 | 1 |
| Terminal | ZNHIT1 | 2.28E-11 | 0.351399 | 0.538 | 0.401 | 3.98E-07 |
| Fibroblasts | GK5 | 0.000387 | 0.351091 | 0.167 | 0.058 | 1 |
| Contractile | ENSSSCG0( | 2.29E-13 | 0.351087 | 0.333 | 0.213 | 4.01E-09 |
| Contractile | LAMB1 | 7.73E-19 | 0.350915 | 0.749 | 0.608 | 1.35E-14 |
| Fibroblasts | PTPRU | 0.000572 | 0.350903 | 0.117 | 0.034 | 1 |
| Terminal | TCP1 | 2.1E-10 | 0.350868 | 0.446 | 0.325 | 3.68E-06 |
| Contractile | MESP2 | 4.06E-21 | 0.350862 | 0.218 | 0.099 | 7.1E-17 |
| Contractile | RBBP7 | 0.002645 | 0.35085 | 0.429 | 0.295 | 1 |
| Contractile | KLHDC3 | 0.000209 | 0.350472 | 0.22 | 0.103 | 1 |
| Pericytes | PHYHD1 | 1.44E-07 | 0.350334 | 0.1 | 0.019 | 0.002522 |
| Terminal | ENSSSCG0( | 3.87E-25 | 0.350322 | 0.117 | 0.025 | 6.77E-21 |
| Fibroblasts | ENSSSCG0( | 0.006248 | 0.350269 | 0.2 | 0.096 | 1 |
| Intermedia | EVA1B | 2.49E-10 | 0.350152 | 0.469 | 0.379 | 4.35E-06 |
| Terminal | PERP | 1.62E-44 | 0.349692 | 0.103 | 0.005 | 2.84E-40 |
| Terminal | CCDC141 | 5.26E-17 | 0.349212 | 0.329 | 0.18 | 9.2E-13 |
| Fibroblasts | MSMO1 | 0.009373 | 0.349205 | 0.167 | 0.08 | 1 |
| Terminal | ARL1 | 9.11E-12 | 0.349109 | 0.554 | 0.425 | 1.59E-07 |
| Terminal | SEC62 | 6.99E-18 | 0.349064 | 0.889 | 0.836 | 1.22E-13 |
| Pericytes | ENSSSCG0( | 0.003227 | 0.348709 | 0.1 | 0.04 | 1 |
| Contractile | GATA5 | 1.59E-24 | 0.348703 | 0.18 | 0.064 | 2.78E-20 |
| Fibroblasts | C1QTNF3 | 0.004468 | 0.34858 | 0.133 | 0.052 | 1 |
| Contractile | SMARCC2 | 0.005203 | 0.348519 | 0.308 | 0.194 | 1 |
| Contractile | ENSSSCG0( | 1.85E-16 | 0.348354 | 0.564 | 0.419 | 3.23E-12 |
| Contractile | WFS1 | 0.000826 | 0.348329 | 0.198 | 0.096 | 1 |
| Contractile | PRKAR2B | 1.14E-20 | 0.348225 | 0.41 | 0.252 | 2E-16 |
| Contractile | CAPZA2 | 7.61E-20 | 0.348169 | 0.768 | 0.669 | 1.33E-15 |
| Terminal | ENSSSCG0( | 2.29E-10 | 0.348028 | 0.521 | 0.41 | 4.01E-06 |
| Pericytes | FAM120B | 0.007072 | 0.347878 | 0.111 | 0.05 | 1 |
| Terminal | GEM | 1.55E-06 | 0.347853 | 0.54 | 0.45 | 0.027111 |
| Contractile | DCAF8 | 0.001347 | 0.347791 | 0.286 | 0.167 | 1 |
| Fibroblasts | ENSSSCG0( | 0.006262 | 0.347717 | 0.833 | 0.761 | 1 |
| Contractile | DSC3 | 1.25E-22 | 0.347532 | 0.15 | 0.049 | 2.18E-18 |
| Terminal | AMDHD2 | 6.93E-13 | 0.347469 | 0.615 | 0.47 | 1.21E-08 |
| Terminal | GFUS | 2.92E-18 | 0.347359 | 0.257 | 0.124 | 5.11E-14 |
| Terminal | EIF3F | 1.45E-12 | 0.347316 | 0.669 | 0.543 | 2.53E-08 |
| Contractile | HSPG2 | 0.004123 | 0.34705 | 0.857 | 0.794 | 1 |
| Contractile | TAX1BP1 | 0.002058 | 0.346889 | 0.769 | 0.676 | 1 |
| Terminal | CHCHD5 | 3.08E-19 | 0.346663 | 0.298 | 0.148 | 5.39E-15 |

| | | | | | | |
|---|---|---|---|---|---|---|
| Contractile | PKN2 | 0.00166 | 0.346645 | 0.451 | 0.313 | 1 |
| Terminal | S100A6 | 2.71E-09 | 0.346578 | 0.993 | 0.99 | 4.74E-05 |
| Terminal | ALYREF | 1.56E-09 | 0.346206 | 0.353 | 0.251 | 2.73E-05 |
| Fibroblasts | ENSSSCG0( | 0.000603 | 0.346189 | 0.15 | 0.051 | 1 |
| Fibroblasts | RGMA | 6.76E-05 | 0.346112 | 0.117 | 0.028 | 1 |
| Contractile | TERF1 | 0.005483 | 0.345941 | 0.33 | 0.22 | 1 |
| Contractile | MMP23B | 2.17E-19 | 0.345677 | 0.629 | 0.454 | 3.79E-15 |
| Contractile | IRS2 | 0.002861 | 0.345673 | 0.143 | 0.065 | 1 |
| Terminal | CCDC167 | 2.84E-22 | 0.345222 | 0.173 | 0.059 | 4.96E-18 |
| Contractile | GULP1 | 1.16E-14 | 0.345221 | 0.47 | 0.337 | 2.03E-10 |
| Contractile | CPQ | 4.65E-20 | 0.344872 | 0.445 | 0.287 | 8.13E-16 |
| Contractile | PTMS | 8.66E-39 | 0.344627 | 0.964 | 0.907 | 1.51E-34 |
| IFN_Interm | WDR36 | 0.005107 | 0.344481 | 0.211 | 0.058 | 1 |
| Contractile | IFI44L | 0.006459 | 0.344452 | 0.209 | 0.119 | 1 |
| Fibroblasts | TMEM243 | 0.00592 | 0.34432 | 0.2 | 0.097 | 1 |
| Contractile | FZD7 | 8.52E-17 | 0.344306 | 0.228 | 0.117 | 1.49E-12 |
| Contractile | ZNF664 | 0.004108 | 0.344176 | 0.308 | 0.195 | 1 |
| Contractile | SLMAP | 3.86E-16 | 0.344137 | 0.327 | 0.2 | 6.74E-12 |
| Contractile | CPED1 | 3.2E-19 | 0.34378 | 0.206 | 0.092 | 5.59E-15 |
| Contractile | PLS3 | 1.35E-18 | 0.343566 | 0.872 | 0.76 | 2.36E-14 |
| Contractile | ARHGAP29 | 0.003789 | 0.343558 | 0.363 | 0.241 | 1 |
| Contractile | LMO4 | 1.27E-13 | 0.343557 | 0.751 | 0.664 | 2.23E-09 |
| Terminal | ANGPTL4 | 1.16E-06 | 0.343191 | 0.269 | 0.186 | 0.020207 |
| Contractile | ZZZ3 | 0.008879 | 0.342974 | 0.264 | 0.167 | 1 |
| Terminal | PSMD8 | 1.61E-08 | 0.342971 | 0.488 | 0.389 | 0.000281 |
| Contractile | ABCC9 | 2.02E-10 | 0.34287 | 0.154 | 0.03 | 3.53E-06 |
| Intermedia | CRISPLD2 | 2.42E-15 | 0.342738 | 0.264 | 0.147 | 4.24E-11 |
| Terminal | CD63 | 2.01E-25 | 0.342529 | 0.975 | 0.955 | 3.52E-21 |
| Contractile | PUM2 | 0.003021 | 0.342479 | 0.396 | 0.268 | 1 |
| Pericytes | ENSSSCG0( | 0.006602 | 0.342338 | 0.144 | 0.073 | 1 |
| Fibroblasts | ENSSSCG0( | 2.36E-14 | 0.342237 | 0.117 | 0.01 | 4.12E-10 |
| Terminal | COX6B | 2.33E-11 | 0.342119 | 0.632 | 0.518 | 4.07E-07 |
| Fibroblasts | SLC47A1 | 0.003134 | 0.342115 | 0.167 | 0.069 | 1 |
| Terminal | ATP6V0B | 1.68E-09 | 0.342066 | 0.465 | 0.351 | 2.94E-05 |
| Intermedia | HEXA | 1.36E-10 | 0.342024 | 0.516 | 0.418 | 2.38E-06 |
| Contractile | WASF3 | 3.05E-22 | 0.341816 | 0.398 | 0.226 | 5.34E-18 |
| Contractile | BCAM | 9.49E-27 | 0.341699 | 0.219 | 0.083 | 1.66E-22 |
| Terminal | ATOX1 | 6.9E-12 | 0.341305 | 0.595 | 0.465 | 1.21E-07 |
| Fibroblasts | GCK | 0.00559 | 0.341303 | 0.1 | 0.034 | 1 |
| Intermedia | PLEKHG5 | 7.57E-10 | 0.340895 | 0.311 | 0.221 | 1.32E-05 |
| Contractile | SERPINE1 | 1.94E-33 | 0.340827 | 0.876 | 0.652 | 3.39E-29 |
| Contractile | PLEKHG3 | 0.001164 | 0.340813 | 0.165 | 0.076 | 1 |
| Contractile | COX17 | 1.36E-16 | 0.340793 | 0.336 | 0.204 | 2.38E-12 |
| Contractile | MRPS22 | 0.001197 | 0.340544 | 0.209 | 0.106 | 1 |
| Fibroblasts | NKD1 | 0.001353 | 0.340534 | 0.167 | 0.065 | 1 |
| Terminal | SPCS2 | 1.33E-11 | 0.34031 | 0.525 | 0.384 | 2.32E-07 |
| Terminal | YIF1A | 2.65E-12 | 0.340274 | 0.405 | 0.272 | 4.63E-08 |
| Intermedia | COMP | 1.55E-11 | 0.339987 | 0.19 | 0.1 | 2.71E-07 |
| Terminal | ENSSSCG0( | 5.43E-15 | 0.339502 | 0.289 | 0.157 | 9.5E-11 |
| Terminal | COX7A2 | 9.34E-13 | 0.339118 | 0.767 | 0.675 | 1.63E-08 |
| Contractile | SFR1 | 0.002138 | 0.338892 | 0.22 | 0.118 | 1 |
| Contractile | HNRNPA1 | 0.001024 | 0.338805 | 0.736 | 0.568 | 1 |
| Terminal | ENSSSCG0( | 1.79E-25 | 0.338802 | 0.99 | 0.98 | 3.13E-21 |
| Terminal | LGMN | 1.48E-09 | 0.338783 | 0.439 | 0.317 | 2.59E-05 |
| Pericytes | TMEM94 | 0.00033 | 0.338742 | 0.1 | 0.032 | 1 |
| Contractile | SLC6A17 | 0.000273 | 0.338739 | 0.11 | 0.037 | 1 |
| Contractile | BLTP3B | 0.008347 | 0.33862 | 0.451 | 0.342 | 1 |
| Terminal | ANAPC11 | 1.25E-11 | 0.338571 | 0.495 | 0.357 | 2.18E-07 |
| Fibroblasts | S1PR2 | 0.002037 | 0.338317 | 0.117 | 0.039 | 1 |

| | | | | | | |
|---|---|---|---|---|---|---|
| Fibroblasts | PI15 | 1.84E-11 | 0.338107 | 0.117 | 0.012 | 3.22E-07 |
| Contractile | EEF2 | 0.001809 | 0.338084 | 0.89 | 0.834 | 1 |
| Terminal | EFCAB7 | 1.73E-15 | 0.337819 | 0.348 | 0.202 | 3.03E-11 |
| Fibroblasts | CEMIP2 | 2.58E-08 | 0.337724 | 0.133 | 0.022 | 0.000452 |
| Intermedia | ISM1 | 4.55E-09 | 0.337498 | 0.361 | 0.273 | 7.95E-05 |
| Contractile | ARIH1 | 0.004135 | 0.337356 | 0.264 | 0.161 | 1 |
| Terminal | SAR1A | 1.06E-10 | 0.337182 | 0.498 | 0.377 | 1.84E-06 |
| Terminal | SNRPE | 1.03E-11 | 0.337134 | 0.641 | 0.51 | 1.81E-07 |
| Fibroblasts | ENSSSCG0( | 0.000189 | 0.337084 | 0.25 | 0.102 | 1 |
| IFN_Interm | BLTP2 | 0.005197 | 0.337045 | 0.158 | 0.036 | 1 |
| Contractile | NUPR1 | 3.54E-05 | 0.337044 | 0.396 | 0.216 | 0.618103 |
| Terminal | ENSSSCG0( | 5.24E-10 | 0.336958 | 0.28 | 0.177 | 9.15E-06 |
| Contractile | ACYP2 | 8.75E-27 | 0.336931 | 0.215 | 0.081 | 1.53E-22 |
| Contractile | AFTPH | 0.003977 | 0.336901 | 0.209 | 0.116 | 1 |
| Terminal | ACTG1 | 2.01E-08 | 0.336835 | 0.966 | 0.989 | 0.000352 |
| Terminal | ACP1 | 3.19E-11 | 0.336548 | 0.524 | 0.388 | 5.57E-07 |
| Fibroblasts | PGAP4 | 0.000184 | 0.336265 | 0.1 | 0.024 | 1 |
| Terminal | FTH1 | 1.54E-12 | 0.335925 | 1 | 0.998 | 2.7E-08 |
| Terminal | ATF3 | 8.7E-08 | 0.335864 | 0.2 | 0.123 | 0.001521 |
| Terminal | RHOA | 3.3E-09 | 0.335706 | 0.776 | 0.733 | 5.77E-05 |
| Terminal | FOSL1 | 2.24E-49 | 0.335591 | 0.118 | 0.007 | 3.91E-45 |
| Intermedia | SAT1 | 2.19E-07 | 0.335441 | 0.465 | 0.384 | 0.003828 |
| Contractile | LAMB2 | 0.005328 | 0.335397 | 0.637 | 0.548 | 1 |
| Terminal | NDUFS4 | 4.71E-10 | 0.335334 | 0.554 | 0.434 | 8.23E-06 |
| Fibroblasts | RIOX2 | 0.006669 | 0.335289 | 0.1 | 0.035 | 1 |
| Contractile | PAIP2 | 0.001052 | 0.335109 | 0.527 | 0.382 | 1 |
| Intermedia | SESN3 | 1.23E-08 | 0.334518 | 0.439 | 0.35 | 0.000215 |
| Intermedia | CHPF | 1.2E-06 | 0.334464 | 0.363 | 0.297 | 0.020993 |
| Contractile | FIS1 | 0.001029 | 0.334442 | 0.747 | 0.648 | 1 |
| Terminal | ERP29 | 1.31E-14 | 0.334355 | 0.333 | 0.196 | 2.29E-10 |
| Intermedia | HSD11B1 | 2.06E-12 | 0.334186 | 0.315 | 0.204 | 3.6E-08 |
| Contractile | LIN7C | 0.007116 | 0.334097 | 0.462 | 0.356 | 1 |
| Contractile | TACC1 | 3.07E-17 | 0.333668 | 0.679 | 0.548 | 5.37E-13 |
| Terminal | MORF4L2 | 8.83E-11 | 0.333574 | 0.531 | 0.405 | 1.54E-06 |
| Contractile | ZNF207 | 0.005804 | 0.333435 | 0.352 | 0.242 | 1 |
| Fibroblasts | LOC10216: | 8E-06 | 0.333222 | 0.117 | 0.024 | 0.139893 |
| Contractile | MAP3K7CL | 3.71E-20 | 0.333206 | 0.107 | 0.028 | 6.48E-16 |
| Fibroblasts | ENSSSCG0( | 8.01E-12 | 0.333094 | 0.133 | 0.016 | 1.4E-07 |
| Terminal | HSPD1 | 1.84E-10 | 0.333004 | 0.599 | 0.496 | 3.21E-06 |
| IFN_Interm | SLFN11 | 9.09E-09 | 0.332915 | 0.158 | 0.011 | 0.000159 |
| Contractile | SYNC | 0.005141 | 0.332679 | 0.198 | 0.111 | 1 |
| Pericytes | PRR16 | 5.8E-06 | 0.332415 | 0.111 | 0.028 | 0.10142 |
| Fibroblasts | TMEM94 | 1.71E-05 | 0.332237 | 0.133 | 0.032 | 0.299538 |
| Contractile | FBH1 | 0.003211 | 0.332132 | 0.198 | 0.105 | 1 |
| Contractile | ETV6 | 0.004593 | 0.33207 | 0.253 | 0.153 | 1 |
| Contractile | CALM1 | 1.7E-25 | 0.332003 | 0.946 | 0.896 | 2.97E-21 |
| Intermedia | NFKBIZ | 1.78E-08 | 0.331858 | 0.499 | 0.402 | 0.000312 |
| Terminal | TALDO1 | 5.32E-12 | 0.331782 | 0.776 | 0.71 | 9.31E-08 |
| IFN_Interm | MED26 | 0.000999 | 0.33126 | 0.158 | 0.029 | 1 |
| Terminal | CD46 | 4.43E-12 | 0.331194 | 0.627 | 0.483 | 7.74E-08 |
| Terminal | FAU | 1.43E-23 | 0.331073 | 0.989 | 0.972 | 2.5E-19 |
| Terminal | WARS1 | 7.71E-18 | 0.331018 | 0.188 | 0.078 | 1.35E-13 |
| Terminal | PTPRM | 6.88E-21 | 0.330828 | 0.247 | 0.107 | 1.2E-16 |
| Fibroblasts | TBC1D4 | 0.00074 | 0.330762 | 0.15 | 0.052 | 1 |
| Contractile | DAAM1 | 1.66E-16 | 0.330711 | 0.427 | 0.28 | 2.9E-12 |
| Terminal | PTGES3 | 1.1E-10 | 0.330365 | 0.606 | 0.485 | 1.93E-06 |
| Contractile | PHKB | 0.000858 | 0.33029 | 0.176 | 0.08 | 1 |
| Terminal | ARPC5L | 3.72E-11 | 0.330207 | 0.266 | 0.159 | 6.5E-07 |
| Pericytes | COX6C | 0.007867 | 0.329848 | 0.8 | 0.782 | 1 |

| | | | | | | |
|---|---|---|---|---|---|---|
| Contractile | CACNA1H | 3.25E-08 | 0.329844 | 0.143 | 0.033 | 0.000569 |
| Pericytes | GPX1 | 0.004263 | 0.329766 | 0.867 | 0.878 | 1 |
| Terminal | FGF2 | 1.33E-12 | 0.3295 | 0.254 | 0.141 | 2.32E-08 |
| Fibroblasts | CORO2B | 0.003149 | 0.329455 | 0.117 | 0.04 | 1 |
| Terminal | ACTR3 | 2.07E-10 | 0.329191 | 0.679 | 0.575 | 3.62E-06 |
| Contractile | EIF3A | 0.00163 | 0.328984 | 0.714 | 0.587 | 1 |
| Terminal | ZDHHC12 | 5.83E-19 | 0.328829 | 0.221 | 0.096 | 1.02E-14 |
| Terminal | TPT1 | 1.36E-14 | 0.328827 | 0.974 | 0.944 | 2.39E-10 |
| Contractile | HIP1R | 3.55E-05 | 0.328739 | 0.11 | 0.031 | 0.620673 |
| Intermedia | RARRES2 | 3.38E-14 | 0.328503 | 0.936 | 0.886 | 5.9E-10 |
| Terminal | CTSH | 3.31E-07 | 0.328352 | 0.632 | 0.55 | 0.005784 |
| Terminal | GSPT1 | 6.03E-10 | 0.328086 | 0.562 | 0.434 | 1.05E-05 |
| Intermedia | PPIC | 6.38E-10 | 0.328035 | 0.513 | 0.427 | 1.12E-05 |
| Terminal | ABL2 | 6.58E-18 | 0.327946 | 0.166 | 0.063 | 1.15E-13 |
| Terminal | MANF | 4.98E-08 | 0.327782 | 0.554 | 0.447 | 0.000871 |
| Terminal | AK4 | 1.53E-15 | 0.327683 | 0.303 | 0.164 | 2.67E-11 |
| Contractile | PPDPF | 0.001215 | 0.327672 | 0.901 | 0.841 | 1 |
| Terminal | PTP4A2 | 1.93E-06 | 0.32722 | 0.514 | 0.419 | 0.033657 |
| Terminal | METTL26 | 3.61E-15 | 0.327179 | 0.233 | 0.116 | 6.31E-11 |
| Contractile | YPEL2 | 5.4E-17 | 0.327171 | 0.363 | 0.226 | 9.44E-13 |
| Contractile | SLC25A4 | 8.14E-18 | 0.327151 | 0.384 | 0.245 | 1.42E-13 |
| Terminal | NDUFB2 | 2.03E-09 | 0.327131 | 0.538 | 0.433 | 3.55E-05 |
| Terminal | TIMM9 | 2.74E-18 | 0.32673 | 0.265 | 0.127 | 4.79E-14 |
| Contractile | DST | 2.97E-16 | 0.326551 | 0.72 | 0.57 | 5.18E-12 |
| Terminal | CRACR2B | 5.32E-35 | 0.326472 | 0.123 | 0.018 | 9.3E-31 |
| Terminal | ELK3 | 3.59E-15 | 0.325927 | 0.3 | 0.163 | 6.28E-11 |
| Terminal | SURF4 | 6.12E-11 | 0.325678 | 0.418 | 0.287 | 1.07E-06 |
| Fibroblasts | RPIA | 0.008542 | 0.325646 | 0.183 | 0.087 | 1 |
| Contractile | GOT2 | 0.003881 | 0.325625 | 0.374 | 0.25 | 1 |
| Intermedia | C1S | 1.73E-11 | 0.325376 | 0.76 | 0.666 | 3.02E-07 |
| Terminal | RABAC1 | 2.33E-14 | 0.325176 | 0.867 | 0.804 | 4.07E-10 |
| Terminal | UBALD2 | 2.28E-08 | 0.325148 | 0.235 | 0.146 | 0.000398 |
| IFN_Interm | SLA-DMB | 4.95E-06 | 0.325104 | 0.158 | 0.017 | 0.086541 |
| Contractile | FGF1 | 8.67E-17 | 0.324647 | 0.181 | 0.083 | 1.52E-12 |
| Terminal | SERPINH1 | 1.95E-07 | 0.324619 | 0.781 | 0.75 | 0.003403 |
| Terminal | IMPDH2 | 9.58E-12 | 0.324584 | 0.335 | 0.209 | 1.68E-07 |
| Pericytes | NUCKS1 | 0.004793 | 0.324311 | 0.844 | 0.845 | 1 |
| Terminal | YWHAZ | 5.54E-12 | 0.32422 | 0.804 | 0.75 | 9.69E-08 |
| Terminal | SLC38A5 | 4.09E-32 | 0.323956 | 0.166 | 0.039 | 7.15E-28 |
| Contractile | PEAR1 | 1.42E-29 | 0.323819 | 0.243 | 0.089 | 2.49E-25 |
| Terminal | TOMM22 | 6.22E-09 | 0.323798 | 0.361 | 0.25 | 0.000109 |
| Fibroblasts | LOC10052 | 0.005822 | 0.323626 | 0.2 | 0.095 | 1 |
| Contractile | USP12 | 0.000649 | 0.323154 | 0.132 | 0.052 | 1 |
| Terminal | STMP1 | 1.71E-13 | 0.323132 | 0.38 | 0.24 | 2.99E-09 |
| Contractile | ARHGEF6 | 0.000324 | 0.323131 | 0.286 | 0.152 | 1 |
| Contractile | MGST2 | 0.000632 | 0.323068 | 0.242 | 0.124 | 1 |
| Terminal | HSPA9 | 1.77E-08 | 0.322697 | 0.353 | 0.246 | 0.000309 |
| Fibroblasts | DEPTOR | 9.96E-12 | 0.32257 | 0.133 | 0.016 | 1.74E-07 |
| Terminal | TFG | 1.06E-10 | 0.32256 | 0.546 | 0.426 | 1.84E-06 |
| Terminal | TMEM147 | 3.91E-11 | 0.322465 | 0.49 | 0.359 | 6.83E-07 |
| Contractile | CAPNS1 | 2.44E-17 | 0.322357 | 0.732 | 0.599 | 4.26E-13 |
| Terminal | SFT2D1 | 3.6E-10 | 0.322331 | 0.432 | 0.305 | 6.3E-06 |
| Terminal | C9H1ORF2 | 4.58E-11 | 0.322251 | 0.524 | 0.393 | 8E-07 |
| Terminal | SHISA5 | 6.65E-13 | 0.322182 | 0.384 | 0.244 | 1.16E-08 |
| Contractile | ZBTB16 | 3.35E-15 | 0.321884 | 0.381 | 0.247 | 5.85E-11 |
| Intermedia | AHR | 4.29E-09 | 0.32179 | 0.225 | 0.144 | 7.51E-05 |
| Contractile | SH3BGRL | 8.6E-15 | 0.321764 | 0.6 | 0.473 | 1.5E-10 |
| Contractile | ANXA6 | 1.66E-17 | 0.321723 | 0.519 | 0.362 | 2.9E-13 |
| Contractile | STK17A | 2.06E-17 | 0.321485 | 0.376 | 0.239 | 3.61E-13 |

| | | | | | | |
|---|---|---|---|---|---|---|
| Terminal | LGALS1 | 0.002948 | 0.32148 | 0.956 | 0.968 | 1 |
| Terminal | NUSAP1 | 8.97E-13 | 0.321307 | 0.11 | 0.04 | 1.57E-08 |
| Fibroblasts | CHAD | 1.25E-08 | 0.321245 | 0.117 | 0.016 | 0.000218 |
| Pericytes | RBM19 | 0.004916 | 0.321114 | 0.1 | 0.041 | 1 |
| Contractile | SPOPL | 0.005237 | 0.321094 | 0.198 | 0.108 | 1 |
| Terminal | CRYL1 | 3.06E-14 | 0.321057 | 0.333 | 0.196 | 5.35E-10 |
| Terminal | PRRX2 | 4.01E-13 | 0.321001 | 0.808 | 0.737 | 7E-09 |
| Contractile | AFF3 | 3.72E-06 | 0.320893 | 0.143 | 0.042 | 0.064946 |
| Terminal | CISD2 | 2.48E-09 | 0.320612 | 0.38 | 0.262 | 4.34E-05 |
| Contractile | ENSSSCG00 | 1.17E-14 | 0.320548 | 0.53 | 0.384 | 2.04E-10 |
| Terminal | CDC42EP5 | 2.22E-07 | 0.320466 | 0.396 | 0.297 | 0.003886 |
| Terminal | CMPK1 | 4.86E-09 | 0.320423 | 0.556 | 0.448 | 8.49E-05 |
| Intermedia | THSD7A | 1.8E-15 | 0.320304 | 0.12 | 0.043 | 3.15E-11 |
| Contractile | CBR1 | 0.007246 | 0.320295 | 0.231 | 0.137 | 1 |
| Terminal | ENSSSCG00 | 2.29E-08 | 0.320229 | 0.311 | 0.212 | 0.000401 |
| Contractile | ATF1 | 0.003473 | 0.320185 | 0.253 | 0.149 | 1 |
| Terminal | TNIP1 | 9.47E-12 | 0.320149 | 0.289 | 0.172 | 1.66E-07 |
| Contractile | EIF4A2 | 0.000672 | 0.320116 | 0.813 | 0.729 | 1 |
| Pericytes | CGRRF1 | 0.007347 | 0.319814 | 0.122 | 0.058 | 1 |
| Contractile | ENSSSCG00 | 1.11E-05 | 0.319671 | 0.132 | 0.04 | 0.193503 |
| Terminal | ARHGDIB | 3.67E-08 | 0.319656 | 0.483 | 0.377 | 0.000642 |
| Intermedia | LTBP3 | 9.72E-10 | 0.319344 | 0.667 | 0.582 | 1.7E-05 |
| Contractile | ZBTB44 | 0.002158 | 0.319143 | 0.231 | 0.125 | 1 |
| Terminal | DARS1 | 1.81E-10 | 0.318652 | 0.398 | 0.274 | 3.16E-06 |
| Terminal | HDLBP | 1.7E-12 | 0.318079 | 0.613 | 0.472 | 2.97E-08 |
| Terminal | SPTSSA | 1.71E-09 | 0.317988 | 0.342 | 0.233 | 2.99E-05 |
| Contractile | PPP1R12C | 0.000307 | 0.317955 | 0.154 | 0.062 | 1 |
| Terminal | PHB2 | 4.38E-13 | 0.317834 | 0.353 | 0.218 | 7.66E-09 |
| Terminal | ATP5MC1 | 7.14E-10 | 0.317811 | 0.597 | 0.49 | 1.25E-05 |
| Contractile | GNG12 | 3.34E-13 | 0.317794 | 0.673 | 0.566 | 5.84E-09 |
| Contractile | BCLAF3 | 0.000675 | 0.31778 | 0.154 | 0.066 | 1 |
| IFN_Interm | NSDHL | 0.002332 | 0.317712 | 0.158 | 0.033 | 1 |
| Contractile | DCAF11 | 0.008849 | 0.317703 | 0.187 | 0.104 | 1 |
| Contractile | SMARCD3 | 0.004469 | 0.317656 | 0.385 | 0.258 | 1 |
| Contractile | MFGE8 | 0.000124 | 0.317645 | 0.945 | 0.873 | 1 |
| Pericytes | AOX1 | 0.008638 | 0.317624 | 0.1 | 0.044 | 1 |
| Terminal | GNB2 | 6.96E-11 | 0.31749 | 0.413 | 0.285 | 1.22E-06 |
| Intermedia | TCF4 | 7.03E-10 | 0.317484 | 0.342 | 0.248 | 1.23E-05 |
| Contractile | ZMYND11 | 0.006231 | 0.317479 | 0.495 | 0.381 | 1 |
| Fibroblasts | ITGA4 | 0.000529 | 0.317459 | 0.1 | 0.026 | 1 |
| Terminal | GMDS | 3.37E-14 | 0.317406 | 0.289 | 0.162 | 5.9E-10 |
| Terminal | SMG6 | 1.16E-16 | 0.31721 | 0.188 | 0.08 | 2.03E-12 |
| Terminal | SFXN3 | 6.59E-19 | 0.317061 | 0.251 | 0.116 | 1.15E-14 |
| Terminal | EPRS1 | 2.18E-11 | 0.316968 | 0.472 | 0.333 | 3.81E-07 |
| IFN_Interm | APBA3 | 0.00639 | 0.316789 | 0.105 | 0.019 | 1 |
| Terminal | HNRNPF | 4.81E-10 | 0.316459 | 0.654 | 0.542 | 8.41E-06 |
| Terminal | BEND6 | 2.01E-28 | 0.316241 | 0.145 | 0.034 | 3.52E-24 |
| Contractile | ANO6 | 0.005203 | 0.316207 | 0.473 | 0.356 | 1 |
| Terminal | NDUFV2 | 1.02E-08 | 0.316138 | 0.513 | 0.399 | 0.000178 |
| Contractile | VIM | 9.85E-06 | 0.316046 | 1 | 0.999 | 0.172268 |
| Intermedia | NT5E | 7.83E-12 | 0.315869 | 0.332 | 0.219 | 1.37E-07 |
| Contractile | ITGB5 | 0.005261 | 0.315768 | 0.681 | 0.571 | 1 |
| Terminal | PRICKLE4 | 3.01E-09 | 0.315457 | 0.546 | 0.423 | 5.26E-05 |
| Fibroblasts | PTGER4 | 3.75E-06 | 0.315411 | 0.1 | 0.017 | 0.065475 |
| Terminal | CHSY1 | 1.69E-09 | 0.315408 | 0.37 | 0.251 | 2.96E-05 |
| Terminal | ANAPC16 | 6.17E-07 | 0.315397 | 0.534 | 0.435 | 0.010789 |
| Contractile | CYLD | 0.005842 | 0.315354 | 0.319 | 0.207 | 1 |
| Contractile | LRRC32 | 1.73E-15 | 0.315045 | 0.308 | 0.182 | 3.03E-11 |
| Contractile | ABLIM1 | 0.000493 | 0.31497 | 0.154 | 0.062 | 1 |

| | | | | | | |
|---|---|---|---|---|---|---|
| Terminal | CYCS | 2.05E-12 | 0.314694 | 0.509 | 0.378 | 3.58E-08 |
| Terminal | SRM | 3.09E-15 | 0.314683 | 0.162 | 0.068 | 5.4E-11 |
| Fibroblasts | OAZ1 | 0.009694 | 0.314451 | 0.8 | 0.769 | 1 |
| Terminal | RALA | 1.08E-10 | 0.314242 | 0.401 | 0.276 | 1.89E-06 |
| Contractile | PSTPIP1 | 0.009343 | 0.314214 | 0.253 | 0.156 | 1 |
| Terminal | OAZ1 | 7.36E-10 | 0.313972 | 0.808 | 0.757 | 1.29E-05 |
| Contractile | CXCL16 | 3.76E-15 | 0.313865 | 0.38 | 0.248 | 6.58E-11 |
| Intermedia | SCARF2 | 1.09E-08 | 0.313728 | 0.343 | 0.26 | 0.00019 |
| Fibroblasts | SOBP | 0.00262 | 0.313645 | 0.117 | 0.04 | 1 |
| Terminal | PSMB6 | 5.76E-08 | 0.313388 | 0.641 | 0.555 | 0.001006 |
| Contractile | PDK3 | 0.00985 | 0.313036 | 0.308 | 0.203 | 1 |
| Terminal | GLRX.2 | 1.11E-10 | 0.313025 | 0.195 | 0.104 | 1.94E-06 |
| Terminal | C4ORF48 | 9.23E-16 | 0.313015 | 0.333 | 0.185 | 1.61E-11 |
| Contractile | PEX5 | 0.000299 | 0.312805 | 0.11 | 0.037 | 1 |
| Terminal | RAP1B | 9.99E-09 | 0.312713 | 0.477 | 0.373 | 0.000175 |
| Terminal | RPN1 | 2.6E-13 | 0.312662 | 0.58 | 0.428 | 4.55E-09 |
| Contractile | RSU1 | 1.94E-16 | 0.312294 | 0.65 | 0.502 | 3.39E-12 |
| Terminal | WLS | 6.13E-24 | 0.312176 | 0.224 | 0.083 | 1.07E-19 |
| Contractile | PDE4DIP | 6.23E-18 | 0.312161 | 0.304 | 0.171 | 1.09E-13 |
| Contractile | HACD1 | 8.56E-16 | 0.311884 | 0.279 | 0.163 | 1.5E-11 |
| Contractile | NPTN | 1.25E-13 | 0.311883 | 0.589 | 0.461 | 2.18E-09 |
| Fibroblasts | PHOSPHO2 | 0.00307 | 0.311791 | 0.1 | 0.032 | 1 |
| Terminal | GOLT1B | 5.1E-14 | 0.311758 | 0.277 | 0.153 | 8.92E-10 |
| Intermedia | PODNL1 | 3.79E-17 | 0.311699 | 0.157 | 0.063 | 6.63E-13 |
| Terminal | ADSS2 | 9.22E-13 | 0.311698 | 0.244 | 0.132 | 1.61E-08 |
| Terminal | CMC1 | 4.45E-12 | 0.311581 | 0.399 | 0.262 | 7.79E-08 |
| Terminal | OLA1 | 3.01E-11 | 0.311539 | 0.313 | 0.195 | 5.26E-07 |
| Intermedia | MAN2B1 | 4.24E-07 | 0.311425 | 0.243 | 0.172 | 0.007406 |
| Pericytes | CHST7 | 0.006805 | 0.311107 | 0.144 | 0.073 | 1 |
| Terminal | SLC39A7 | 8.64E-13 | 0.310999 | 0.369 | 0.229 | 1.51E-08 |
| Contractile | CCND2 | 9.19E-13 | 0.310951 | 0.463 | 0.328 | 1.61E-08 |
| Contractile | EIF3L | 0.004353 | 0.310871 | 0.473 | 0.348 | 1 |
| Contractile | C1QTNF1 | 3.52E-15 | 0.31086 | 0.764 | 0.629 | 6.15E-11 |
| Intermedia | RAB31 | 2.51E-08 | 0.310457 | 0.421 | 0.341 | 0.000438 |
| Pericytes | HIBCH | 0.005917 | 0.310356 | 0.1 | 0.042 | 1 |
| Intermedia | S100A13 | 2.5E-08 | 0.310186 | 0.6 | 0.543 | 0.000437 |
| Contractile | ENSSSCG0 | 5.54E-20 | 0.31011 | 0.156 | 0.057 | 9.69E-16 |
| Intermedia | MEOX1 | 1.16E-20 | 0.310099 | 0.104 | 0.026 | 2.02E-16 |
| Intermedia | SDC2 | 5.23E-13 | 0.310069 | 0.868 | 0.812 | 9.14E-09 |
| Terminal | COPS8 | 2.16E-11 | 0.309817 | 0.302 | 0.189 | 3.78E-07 |
| Terminal | DYNLRB1 | 4.72E-11 | 0.309798 | 0.75 | 0.637 | 8.26E-07 |
| Contractile | YWHAQ | 0.006204 | 0.309714 | 0.736 | 0.611 | 1 |
| Contractile | H1-10 | 0.002983 | 0.309488 | 0.736 | 0.623 | 1 |
| Pericytes | PITX1 | 3.01E-06 | 0.309426 | 0.111 | 0.027 | 0.052642 |
| Pericytes | SHC4 | 1.58E-07 | 0.309303 | 0.1 | 0.019 | 0.002763 |
| Fibroblasts | HGF | 0.006737 | 0.30911 | 0.2 | 0.095 | 1 |
| Contractile | STAU2 | 3.05E-18 | 0.308914 | 0.21 | 0.1 | 5.33E-14 |
| Contractile | PHF21A | 0.007118 | 0.308507 | 0.165 | 0.086 | 1 |
| Terminal | PCMT1 | 6.18E-10 | 0.308407 | 0.476 | 0.349 | 1.08E-05 |
| Terminal | MICOS13 | 3E-07 | 0.308377 | 0.494 | 0.392 | 0.005238 |
| Contractile | PTGES3L | 1.41E-26 | 0.308222 | 0.158 | 0.046 | 2.46E-22 |
| Contractile | MYO1C | 1.66E-14 | 0.308174 | 0.49 | 0.357 | 2.9E-10 |
| Contractile | DPM3 | 0.007921 | 0.30803 | 0.297 | 0.195 | 1 |
| Terminal | ARL8B | 1.69E-12 | 0.307633 | 0.487 | 0.335 | 2.95E-08 |
| Terminal | ENSSSCG0 | 1.58E-10 | 0.307548 | 0.772 | 0.674 | 2.76E-06 |
| Terminal | HMGN1 | 2.04E-06 | 0.307504 | 0.492 | 0.403 | 0.035575 |
| Terminal | FBL | 1.61E-12 | 0.307226 | 0.328 | 0.198 | 2.82E-08 |
| Terminal | SLC25A24 | 1.03E-09 | 0.307142 | 0.37 | 0.255 | 1.8E-05 |
| Terminal | CTHRC1 | 1.48E-24 | 0.307043 | 0.401 | 0.2 | 2.59E-20 |

| | | | | | | |
|---|---|---|---|---|---|---|
| Contractile | CPXM2 | 6.2E-22 | 0.306857 | 0.86 | 0.688 | 1.08E-17 |
| Pericytes | HMBOX1 | 0.003813 | 0.30666 | 0.133 | 0.061 | 1 |
| Contractile | ESYT2 | 1.63E-12 | 0.306655 | 0.457 | 0.334 | 2.86E-08 |
| Contractile | ARID4A | 0.009812 | 0.306517 | 0.374 | 0.258 | 1 |
| Terminal | CCDC47 | 9.38E-13 | 0.306284 | 0.343 | 0.21 | 1.64E-08 |
| Terminal | DRAP1 | 1.55E-08 | 0.306089 | 0.606 | 0.509 | 0.00027 |
| Terminal | ADAMTS6 | 2.54E-30 | 0.306043 | 0.115 | 0.019 | 4.44E-26 |
| Intermedia | PDGFRA | 2.25E-10 | 0.306035 | 0.282 | 0.184 | 3.94E-06 |
| Terminal | FAM114A1 | 1.73E-07 | 0.305989 | 0.649 | 0.567 | 0.003017 |
| Intermedia | BHLHE41 | 7.9E-09 | 0.305978 | 0.246 | 0.161 | 0.000138 |
| Fibroblasts | DCLK1 | 0.003383 | 0.305807 | 0.15 | 0.06 | 1 |
| Contractile | ROR1 | 1.53E-19 | 0.305365 | 0.19 | 0.081 | 2.68E-15 |
| Terminal | ATP5MC2 | 2.01E-10 | 0.305317 | 0.598 | 0.466 | 3.52E-06 |
| Contractile | KIF1C | 9.43E-18 | 0.305106 | 0.369 | 0.228 | 1.65E-13 |
| Terminal | TAF10 | 1.09E-10 | 0.305011 | 0.422 | 0.292 | 1.91E-06 |
| Terminal | THYN1 | 6.09E-12 | 0.304369 | 0.391 | 0.254 | 1.06E-07 |
| Intermedia | PTPN13 | 2.65E-06 | 0.304309 | 0.395 | 0.326 | 0.046359 |
| Terminal | RTCB | 7.49E-13 | 0.304227 | 0.285 | 0.165 | 1.31E-08 |
| Contractile | CAV3 | 2.95E-05 | 0.30373 | 0.165 | 0.06 | 0.514928 |
| Terminal | C1QBP | 2.61E-08 | 0.303538 | 0.383 | 0.283 | 0.000456 |
| Terminal | RAB2A | 4.39E-13 | 0.303469 | 0.849 | 0.779 | 7.68E-09 |
| Terminal | OSBPL3 | 2.55E-20 | 0.303224 | 0.198 | 0.076 | 4.45E-16 |
| Terminal | TJP2 | 8.94E-14 | 0.302766 | 0.355 | 0.213 | 1.56E-09 |
| Contractile | TMEM51 | 6E-19 | 0.30255 | 0.196 | 0.086 | 1.05E-14 |
| Terminal | RAC1 | 4.11E-13 | 0.3025 | 0.774 | 0.694 | 7.18E-09 |
| Contractile | AGAP1 | 0.004769 | 0.30238 | 0.198 | 0.106 | 1 |
| Intermedia | TSC22D3 | 1.12E-08 | 0.302351 | 0.46 | 0.371 | 0.000196 |
| Contractile | FCHO2 | 0.003115 | 0.302293 | 0.407 | 0.274 | 1 |
| Fibroblasts | CD320 | 0.002919 | 0.302177 | 0.117 | 0.04 | 1 |
| Terminal | MED10 | 2.78E-14 | 0.302127 | 0.274 | 0.148 | 4.86E-10 |
| Intermedia | ATRAID | 2.69E-08 | 0.302124 | 0.536 | 0.461 | 0.000471 |
| Terminal | ENO2 | 5.8E-29 | 0.301992 | 0.121 | 0.022 | 1.01E-24 |
| Terminal | GPRC5A | 3.4E-25 | 0.301928 | 0.104 | 0.019 | 5.94E-21 |
| Terminal | NR1D1 | 2.86E-25 | 0.301839 | 0.166 | 0.048 | 5.01E-21 |
| Terminal | AVPI1 | 1.47E-18 | 0.301754 | 0.162 | 0.058 | 2.57E-14 |
| Terminal | DAD1 | 1.81E-14 | 0.301751 | 0.27 | 0.144 | 3.16E-10 |
| Terminal | PLIN2 | 2.75E-08 | 0.301713 | 0.357 | 0.245 | 0.000481 |
| Terminal | XYLT1 | 4.19E-14 | 0.301514 | 0.193 | 0.091 | 7.33E-10 |
| Pericytes | NTHL1 | 0.006477 | 0.301378 | 0.1 | 0.042 | 1 |
| Terminal | ALCAM | 3.54E-11 | 0.30132 | 0.347 | 0.221 | 6.19E-07 |
| Intermedia | ATP8B4 | 2.79E-10 | 0.301204 | 0.234 | 0.144 | 4.88E-06 |
| Intermedia | DCN | 1.19E-15 | 0.301004 | 0.53 | 0.399 | 2.09E-11 |
| Contractile | TP53I11 | 0.000577 | 0.300903 | 0.22 | 0.106 | 1 |
| Contractile | RAB11FIP2 | 0.001282 | 0.300794 | 0.264 | 0.143 | 1 |
| Intermedia | TIMP2 | 3.04E-20 | 0.300716 | 0.949 | 0.9 | 5.31E-16 |
| Contractile | DGKH | 0.005872 | 0.300641 | 0.154 | 0.078 | 1 |
| Contractile | MRPL37 | 0.007373 | 0.3006 | 0.22 | 0.131 | 1 |
| Intermedia | DPT | 1.04E-09 | 0.300508 | 0.144 | 0.075 | 1.83E-05 |
| Contractile | SYNC | 1.21E-19 | 0.300333 | 0.176 | 0.072 | 2.11E-15 |
| Intermedia | NRP2 | 3.78E-06 | 0.300306 | 0.269 | 0.204 | 0.066134 |
| Terminal | H1-3 | 1.71E-15 | 0.30011 | 0.14 | 0.052 | 3E-11 |
| Contractile | ADGRL2 | 5.74E-14 | 0.300032 | 0.341 | 0.219 | 1E-09 |
| Terminal | GPX1 | 1.67E-11 | 0.299594 | 0.915 | 0.866 | 2.92E-07 |
| Intermedia | RCN2 | 2.95E-10 | 0.299397 | 0.641 | 0.576 | 5.15E-06 |
| Terminal | DERL1 | 1.17E-10 | 0.299303 | 0.477 | 0.342 | 2.05E-06 |
| Contractile | CTSA | 6.58E-14 | 0.299299 | 0.676 | 0.532 | 1.15E-09 |
| Terminal | FJX1 | 3.66E-14 | 0.299272 | 0.233 | 0.119 | 6.4E-10 |
| Terminal | ACOT13 | 4.62E-08 | 0.29925 | 0.343 | 0.239 | 0.000807 |
| Contractile | PLEKHO1 | 0.001044 | 0.299132 | 0.209 | 0.105 | 1 |

| | | | | | | |
|---|---|---|---|---|---|---|
| Contractile | PDCL3 | 0.001333 | 0.299072 | 0.187 | 0.09 | 1 |
| Contractile | TMEM64 | 0.00463 | 0.299061 | 0.165 | 0.084 | 1 |
| Terminal | KIT | 1.58E-14 | 0.29892 | 0.133 | 0.05 | 2.76E-10 |
| Terminal | VPS13C | 5.36E-08 | 0.298856 | 0.45 | 0.339 | 0.000937 |
| Contractile | PACSIN3 | 0.000683 | 0.298815 | 0.308 | 0.172 | 1 |
| Terminal | ENSSSCG00 | 2.86E-05 | 0.298795 | 0.669 | 0.58 | 0.499572 |
| Intermedia | LMO7 | 6.47E-09 | 0.298524 | 0.241 | 0.16 | 0.000113 |
| Contractile | HEYL | 0.002617 | 0.29831 | 0.341 | 0.214 | 1 |
| Contractile | MYO9B | 1.71E-12 | 0.298243 | 0.267 | 0.163 | 2.99E-08 |
| Pericytes | DDHD1 | 0.006052 | 0.298184 | 0.122 | 0.056 | 1 |
| Terminal | SERBP1 | 2.16E-09 | 0.29818 | 0.764 | 0.7 | 3.78E-05 |
| Terminal | SLC2A3 | 1.68E-13 | 0.298167 | 0.444 | 0.288 | 2.94E-09 |
| Contractile | FAM210B | 2.04E-14 | 0.298047 | 0.414 | 0.285 | 3.57E-10 |
| Contractile | TRAK2 | 1.91E-14 | 0.298008 | 0.28 | 0.166 | 3.34E-10 |
| Contractile | COL17A1 | 1.71E-09 | 0.297894 | 0.11 | 0.018 | 2.99E-05 |
| Contractile | RETREG1 | 0.00224 | 0.297888 | 0.209 | 0.109 | 1 |
| Terminal | NDUFB3 | 2.8E-09 | 0.297873 | 0.562 | 0.446 | 4.89E-05 |
| IFN_Interm | CBX7 | 0.001475 | 0.297856 | 0.105 | 0.015 | 1 |
| Terminal | KLF6 | 8.25E-08 | 0.297679 | 0.645 | 0.538 | 0.001442 |
| Contractile | PLAC9 | 6.7E-16 | 0.297595 | 0.665 | 0.524 | 1.17E-11 |
| Fibroblasts | LPCAT3 | 0.00202 | 0.297422 | 0.117 | 0.038 | 1 |
| Terminal | LIMS1 | 1.73E-10 | 0.297285 | 0.587 | 0.459 | 3.03E-06 |
| Terminal | IFNGR2 | 1.95E-11 | 0.297226 | 0.354 | 0.225 | 3.42E-07 |
| Terminal | COPB1 | 1.3E-08 | 0.296907 | 0.538 | 0.425 | 0.000227 |
| Terminal | CCNYL1 | 2.48E-08 | 0.296669 | 0.281 | 0.185 | 0.000434 |
| Terminal | PHB1 | 6.25E-08 | 0.296329 | 0.337 | 0.243 | 0.001093 |
| Terminal | TEX264 | 5.66E-12 | 0.296235 | 0.211 | 0.112 | 9.9E-08 |
| Contractile | ENAH | 1.93E-14 | 0.295771 | 0.47 | 0.334 | 3.37E-10 |
| Terminal | KRTCAP2 | 6E-11 | 0.29576 | 0.763 | 0.682 | 1.05E-06 |
| Intermedia | SESN1 | 1.8E-08 | 0.295731 | 0.282 | 0.195 | 0.000314 |
| Terminal | GNPNAT1 | 5.24E-16 | 0.29569 | 0.204 | 0.093 | 9.16E-12 |
| Terminal | ARPC2 | 2.25E-09 | 0.295687 | 0.849 | 0.811 | 3.93E-05 |
| Contractile | NKD1 | 0.002202 | 0.295663 | 0.143 | 0.064 | 1 |
| Contractile | MSN | 7.09E-17 | 0.295465 | 0.842 | 0.742 | 1.24E-12 |
| Contractile | DDAH1 | 1.19E-13 | 0.295344 | 0.223 | 0.125 | 2.07E-09 |
| Contractile | RNF149 | 8.88E-14 | 0.295312 | 0.281 | 0.171 | 1.55E-09 |
| Intermedia | CREB3L1 | 2.42E-09 | 0.295257 | 0.305 | 0.214 | 4.24E-05 |
| Terminal | BRI3 | 3.25E-07 | 0.29515 | 0.61 | 0.507 | 0.005688 |
| Terminal | ENSSSCG00 | 3.28E-10 | 0.294952 | 0.752 | 0.672 | 5.74E-06 |
| Contractile | PAIP2B | 0.000462 | 0.294931 | 0.11 | 0.038 | 1 |
| Pericytes | EEF2 | 0.004595 | 0.294849 | 0.856 | 0.835 | 1 |
| Terminal | TOMM20 | 5.17E-08 | 0.294574 | 0.373 | 0.269 | 0.000903 |
| Intermedia | ENSSSCG00 | 7.3E-19 | 0.294533 | 0.954 | 0.922 | 1.28E-14 |
| Terminal | GLS | 1.67E-08 | 0.294491 | 0.538 | 0.432 | 0.000292 |
| Terminal | LOC10062 | 5.92E-13 | 0.294278 | 0.169 | 0.078 | 1.04E-08 |
| Terminal | VAPA | 1.32E-08 | 0.294106 | 0.615 | 0.514 | 0.00023 |
| Fibroblasts | NABP1 | 0.003233 | 0.294106 | 0.1 | 0.032 | 1 |
| Contractile | EIF3A | 9.66E-16 | 0.294095 | 0.676 | 0.534 | 1.69E-11 |
| Contractile | PGM5 | 1.56E-22 | 0.294047 | 0.139 | 0.041 | 2.73E-18 |
| Contractile | TRAPPC5 | 0.009617 | 0.293612 | 0.56 | 0.435 | 1 |
| Terminal | CANX | 1.41E-09 | 0.293516 | 0.77 | 0.696 | 2.46E-05 |
| Terminal | CARD19 | 2.1E-10 | 0.293497 | 0.347 | 0.229 | 3.68E-06 |
| Terminal | TRAM1 | 4.98E-08 | 0.293202 | 0.468 | 0.363 | 0.000871 |
| Contractile | DAAM2 | 9.15E-16 | 0.293131 | 0.187 | 0.089 | 1.6E-11 |
| Terminal | LXN | 2.92E-07 | 0.293108 | 0.374 | 0.277 | 0.005105 |
| Contractile | STK40 | 0.009897 | 0.293092 | 0.22 | 0.133 | 1 |
| Terminal | CHCHD2 | 2.06E-08 | 0.293033 | 0.61 | 0.504 | 0.00036 |
| Contractile | BMPR1A | 2.52E-14 | 0.292998 | 0.378 | 0.249 | 4.4E-10 |
| Terminal | PSMA2 | 2.64E-08 | 0.292558 | 0.553 | 0.447 | 0.000462 |

| | | | | | | |
|---|---|---|---|---|---|---|
| Terminal | SRSF3 | 8.53E-06 | 0.292154 | 0.549 | 0.464 | 0.149169 |
| Terminal | LUM | 2.15E-10 | 0.292108 | 0.94 | 0.877 | 3.76E-06 |
| Terminal | TIMM8B | 5.93E-08 | 0.292107 | 0.543 | 0.429 | 0.001036 |
| Pericytes | PGM2 | 0.006235 | 0.291904 | 0.111 | 0.049 | 1 |
| Terminal | RTRAF | 1.78E-10 | 0.291578 | 0.66 | 0.547 | 3.12E-06 |
| Terminal | PSMB7 | 9.86E-11 | 0.291532 | 0.41 | 0.282 | 1.72E-06 |
| Intermedia | SH3D19 | 3.86E-05 | 0.291387 | 0.276 | 0.221 | 0.674278 |
| Fibroblasts | TMEM204 | 3.13E-08 | 0.291351 | 0.133 | 0.022 | 0.000547 |
| Fibroblasts | PIM1 | 0.003695 | 0.291332 | 0.117 | 0.041 | 1 |
| Terminal | ATP6V1F | 3.06E-09 | 0.29132 | 0.45 | 0.329 | 5.35E-05 |
| Terminal | PDLIM4 | 1.82E-14 | 0.291141 | 0.218 | 0.107 | 3.18E-10 |
| Pericytes | EIF4G2 | 0.005376 | 0.291013 | 0.856 | 0.87 | 1 |
| Fibroblasts | HOXB4 | 0.000143 | 0.290967 | 0.117 | 0.03 | 1 |
| Contractile | TNFRSF11I | 5.17E-19 | 0.290924 | 0.548 | 0.364 | 9.03E-15 |
| Contractile | SRPX | 2.61E-12 | 0.290713 | 0.6 | 0.476 | 4.57E-08 |
| Terminal | ADAM12 | 2.78E-13 | 0.290629 | 0.25 | 0.132 | 4.86E-09 |
| Terminal | CCT7 | 1.54E-08 | 0.290323 | 0.497 | 0.384 | 0.000269 |
| Contractile | LIN7C | 1.95E-12 | 0.290254 | 0.433 | 0.309 | 3.42E-08 |
| Contractile | SLC2A13 | 4.95E-11 | 0.290238 | 0.4 | 0.291 | 8.66E-07 |
| Contractile | INHBA | 4.92E-11 | 0.289884 | 0.399 | 0.283 | 8.61E-07 |
| Intermedia | PLD3 | 1.71E-06 | 0.289877 | 0.382 | 0.317 | 0.02986 |
| Terminal | SLC38A1 | 1.14E-30 | 0.289701 | 0.115 | 0.019 | 1.99E-26 |
| Pericytes | ENSSSCG0( | 0.001384 | 0.289643 | 0.867 | 0.758 | 1 |
| Terminal | SLC50A1 | 1.24E-10 | 0.289621 | 0.262 | 0.156 | 2.17E-06 |
| Terminal | HSPA8 | 2.61E-09 | 0.289561 | 0.783 | 0.694 | 4.57E-05 |
| Fibroblasts | HRH1 | 0.002088 | 0.289501 | 0.117 | 0.038 | 1 |
| Contractile | ENTPD3 | 5.66E-05 | 0.289174 | 0.132 | 0.043 | 0.988627 |
| Contractile | MEF2D | 1.41E-13 | 0.288652 | 0.294 | 0.181 | 2.47E-09 |
| Terminal | DTD1 | 4.5E-13 | 0.288651 | 0.27 | 0.151 | 7.86E-09 |
| Contractile | CYLD | 4.24E-13 | 0.288343 | 0.274 | 0.167 | 7.42E-09 |
| Terminal | C6ORF62 | 9.79E-10 | 0.288295 | 0.357 | 0.241 | 1.71E-05 |
| Intermedia | SELENOM | 1.75E-11 | 0.288177 | 0.798 | 0.699 | 3.06E-07 |
| Terminal | CHRAC1 | 1.74E-10 | 0.288059 | 0.403 | 0.27 | 3.04E-06 |
| Intermedia | LAMB1 | 5.44E-10 | 0.287942 | 0.737 | 0.636 | 9.5E-06 |
| Pericytes | ENSSSCG0( | 0.0002 | 0.287924 | 0.978 | 0.988 | 1 |
| Contractile | MFGE8 | 3.1E-21 | 0.287909 | 0.933 | 0.837 | 5.42E-17 |
| Contractile | MRPL47 | 0.006154 | 0.28768 | 0.231 | 0.136 | 1 |
| Contractile | PPM1K | 0.005283 | 0.287621 | 0.253 | 0.147 | 1 |
| Contractile | PTPN11 | 7.55E-11 | 0.287285 | 0.455 | 0.341 | 1.32E-06 |
| Terminal | SUPT4H1 | 1.23E-10 | 0.286955 | 0.407 | 0.274 | 2.14E-06 |
| Terminal | HAPLN3 | 6.49E-11 | 0.286864 | 0.147 | 0.069 | 1.13E-06 |
| Terminal | ARFGAP1 | 9.84E-10 | 0.286773 | 0.42 | 0.295 | 1.72E-05 |
| Intermedia | SRPX | 6.61E-07 | 0.286759 | 0.574 | 0.506 | 0.011559 |
| Contractile | UBE2B | 0.009706 | 0.286605 | 0.681 | 0.611 | 1 |
| Fibroblasts | ENSSSCG0( | 7.05E-06 | 0.286467 | 0.117 | 0.024 | 0.123263 |
| Terminal | GNG10 | 1.93E-09 | 0.286433 | 0.796 | 0.72 | 3.37E-05 |
| Pericytes | PITPNC1 | 0.000923 | 0.286137 | 0.133 | 0.054 | 1 |
| Contractile | NTF3 | 7.28E-29 | 0.285994 | 0.113 | 0.019 | 1.27E-24 |
| Terminal | NDUFA2 | 1.02E-08 | 0.285952 | 0.565 | 0.45 | 0.000179 |
| Terminal | COPS9 | 1.35E-08 | 0.285768 | 0.742 | 0.685 | 0.000236 |
| Fibroblasts | MTF1 | 0.001744 | 0.285398 | 0.1 | 0.029 | 1 |
| Intermedia | ASAH1 | 1.19E-07 | 0.285362 | 0.294 | 0.217 | 0.002072 |
| Terminal | ARCN1 | 5.92E-08 | 0.285306 | 0.588 | 0.481 | 0.001035 |
| Terminal | ETV4 | 7.28E-23 | 0.285254 | 0.121 | 0.029 | 1.27E-18 |
| Contractile | ENSSSCG0( | 3.7E-19 | 0.285109 | 0.193 | 0.082 | 6.47E-15 |
| Terminal | UQCRQ | 1.54E-08 | 0.284955 | 0.42 | 0.311 | 0.000269 |
| Terminal | EIF2S3B | 2.65E-10 | 0.284759 | 0.486 | 0.344 | 4.63E-06 |
| IFN_Interm | ENSSSCG0( | 1.15E-07 | 0.284685 | 0.105 | 0.006 | 0.002004 |
| Contractile | ENSSSCG0( | 2.59E-17 | 0.284674 | 0.293 | 0.163 | 4.53E-13 |

| | | | | | | |
|---|---|---|---|---|---|---|
| Intermedia | NUCB1 | 3.23E-08 | 0.28436 | 0.624 | 0.559 | 0.000565 |
| Contractile | GIT2 | 0.003087 | 0.284083 | 0.209 | 0.113 | 1 |
| Terminal | SNRPF | 1.38E-08 | 0.283997 | 0.406 | 0.294 | 0.000241 |
| Terminal | SMIM15 | 6.85E-09 | 0.283866 | 0.361 | 0.247 | 0.00012 |
| Contractile | CD81 | 1.08E-23 | 0.283681 | 0.972 | 0.932 | 1.88E-19 |
| Terminal | EEF1B2 | 5.08E-07 | 0.283475 | 0.908 | 0.867 | 0.008887 |
| Contractile | GRIK5 | 0.000189 | 0.283464 | 0.132 | 0.047 | 1 |
| Terminal | CTNNBIP1 | 3.77E-18 | 0.283372 | 0.236 | 0.106 | 6.59E-14 |
| Terminal | FKBP10 | 2.71E-09 | 0.28337 | 0.516 | 0.399 | 4.74E-05 |
| Intermedia | LURAP1L | 3.5E-10 | 0.283343 | 0.139 | 0.071 | 6.12E-06 |
| Fibroblasts | PPP1R15B | 0.009797 | 0.282875 | 0.233 | 0.123 | 1 |
| Terminal | ATP5MF | 1.2E-10 | 0.282757 | 0.808 | 0.725 | 2.1E-06 |
| Terminal | C17ORF49 | 9.8E-14 | 0.282623 | 0.237 | 0.123 | 1.71E-09 |
| Intermedia | ALDH1A2 | 1.43E-09 | 0.28262 | 0.138 | 0.071 | 2.51E-05 |
| Contractile | ACTR2 | 5.78E-11 | 0.282566 | 0.59 | 0.475 | 1.01E-06 |
| Contractile | LGR4 | 3.37E-05 | 0.28253 | 0.132 | 0.041 | 0.589233 |
| Terminal | EIF3G | 7.39E-08 | 0.282363 | 0.652 | 0.551 | 0.001291 |
| Contractile | BPNT2 | 1.12E-12 | 0.281898 | 0.488 | 0.364 | 1.96E-08 |
| Terminal | RAB5IF | 4.76E-10 | 0.281666 | 0.513 | 0.382 | 8.33E-06 |
| Intermedia | GPR155 | 8.48E-07 | 0.281579 | 0.229 | 0.162 | 0.014822 |
| Terminal | CENPH | 6.61E-17 | 0.281457 | 0.152 | 0.057 | 1.16E-12 |
| Terminal | RANBP1 | 0.000202 | 0.281109 | 0.524 | 0.456 | 1 |
| Contractile | TBC1D10A | 0.005298 | 0.281085 | 0.11 | 0.047 | 1 |
| Terminal | SLC17A9 | 5.64E-21 | 0.281035 | 0.176 | 0.062 | 9.86E-17 |
| IFN_Interm | ENSSSCG0( | 4.48E-07 | 0.280812 | 0.105 | 0.007 | 0.007839 |
| Terminal | VBP1 | 2.07E-07 | 0.280702 | 0.388 | 0.285 | 0.003614 |
| Intermedia | GPX4 | 3.36E-08 | 0.28066 | 0.677 | 0.631 | 0.000587 |
| Contractile | EXT1 | 5.23E-11 | 0.280644 | 0.385 | 0.273 | 9.14E-07 |
| Terminal | SNX5 | 7.91E-11 | 0.280331 | 0.267 | 0.158 | 1.38E-06 |
| Terminal | CCT2 | 8.43E-09 | 0.280312 | 0.565 | 0.438 | 0.000147 |
| Terminal | CAPZA1 | 2.57E-12 | 0.28022 | 0.436 | 0.284 | 4.49E-08 |
| Terminal | UQCRFS1 | 4.08E-06 | 0.28014 | 0.374 | 0.287 | 0.071335 |
| Terminal | DPM2 | 2.97E-14 | 0.279911 | 0.266 | 0.143 | 5.2E-10 |
| Terminal | OMD | 0.000484 | 0.279793 | 0.278 | 0.22 | 1 |
| IFN_Interm | TMEM140 | 0.001493 | 0.279777 | 0.105 | 0.015 | 1 |
| Contractile | FLNC | 5.63E-31 | 0.279693 | 0.13 | 0.024 | 9.84E-27 |
| Terminal | CCT3 | 1.24E-06 | 0.279624 | 0.484 | 0.388 | 0.021606 |
| Terminal | MRPS21 | 1.58E-08 | 0.279583 | 0.41 | 0.295 | 0.000277 |
| Contractile | NMRAL1 | 0.005084 | 0.279565 | 0.176 | 0.092 | 1 |
| Terminal | LTBP2 | 5.42E-10 | 0.279498 | 0.789 | 0.718 | 9.48E-06 |
| Contractile | TFEB | 0.006948 | 0.279383 | 0.165 | 0.086 | 1 |
| Terminal | YWHAQ | 3.97E-07 | 0.279273 | 0.668 | 0.599 | 0.006933 |
| Pericytes | TGFBR3 | 0.007712 | 0.278971 | 0.111 | 0.05 | 1 |
| Terminal | NARS1 | 5.71E-09 | 0.278943 | 0.523 | 0.403 | 9.98E-05 |
| Contractile | CSPG4 | 6.93E-15 | 0.278913 | 0.525 | 0.38 | 1.21E-10 |
| Contractile | MORF4L1 | 0.003631 | 0.278857 | 0.912 | 0.833 | 1 |
| Terminal | TMED2 | 6.34E-09 | 0.278654 | 0.719 | 0.615 | 0.000111 |
| Contractile | THRB | 1.37E-10 | 0.278609 | 0.305 | 0.205 | 2.4E-06 |
| Terminal | ABI3BP | 5.6E-06 | 0.27844 | 0.494 | 0.406 | 0.097962 |
| Contractile | DYNC1LI2 | 3.88E-10 | 0.278006 | 0.566 | 0.463 | 6.79E-06 |
| Terminal | COX4I1 | 4.73E-13 | 0.277837 | 0.94 | 0.903 | 8.27E-09 |
| Fibroblasts | NOP2 | 0.006159 | 0.277772 | 0.117 | 0.043 | 1 |
| Contractile | SYDE2 | 0.000885 | 0.27776 | 0.143 | 0.059 | 1 |
| Terminal | GHITM | 2.51E-07 | 0.277651 | 0.462 | 0.354 | 0.004383 |
| Intermedia | SOX6 | 4.01E-07 | 0.277607 | 0.179 | 0.116 | 0.00701 |
| IFN_Interm | MTM1 | 0.002129 | 0.277607 | 0.158 | 0.032 | 1 |
| Contractile | DTNA | 8.82E-23 | 0.277353 | 0.107 | 0.024 | 1.54E-18 |
| Terminal | POMP | 3.75E-07 | 0.277255 | 0.63 | 0.531 | 0.006553 |
| Contractile | RSL1D1 | 3.94E-13 | 0.2771 | 0.399 | 0.272 | 6.89E-09 |

| | | | | | | |
|---|---|---|---|---|---|---|
| Terminal | ROBO1 | 5.04E-12 | 0.277005 | 0.303 | 0.178 | 8.81E-08 |
| Intermedia | DLX5 | 1.08E-13 | 0.276989 | 0.103 | 0.036 | 1.88E-09 |
| Contractile | TP53BP2 | 0.009033 | 0.276924 | 0.231 | 0.138 | 1 |
| Terminal | PVR | 1.17E-23 | 0.27684 | 0.1 | 0.019 | 2.04E-19 |
| Contractile | NCS1 | 6.81E-16 | 0.276706 | 0.198 | 0.098 | 1.19E-11 |
| Contractile | ENSSSCG0( | 1.25E-14 | 0.276643 | 0.203 | 0.104 | 2.18E-10 |
| Terminal | COX7C | 8.8E-11 | 0.2766 | 0.82 | 0.763 | 1.54E-06 |
| Contractile | ARHGEF7 | 8.73E-19 | 0.27635 | 0.253 | 0.126 | 1.53E-14 |
| Fibroblasts | ENSSSCG0( | 0.003608 | 0.27633 | 1 | 0.987 | 1 |
| Intermedia | C1QTNF7 | 5.46E-06 | 0.276164 | 0.415 | 0.35 | 0.095392 |
| Fibroblasts | EPB41 | 0.004242 | 0.276132 | 0.1 | 0.033 | 1 |
| Terminal | COX6A2 | 2.34E-11 | 0.276089 | 0.233 | 0.131 | 4.1E-07 |
| Terminal | PSMA6 | 1.48E-07 | 0.275868 | 0.446 | 0.339 | 0.002579 |
| Terminal | FAM216A | 2.96E-20 | 0.275715 | 0.125 | 0.035 | 5.17E-16 |
| Terminal | DPM1 | 3.46E-09 | 0.275684 | 0.469 | 0.343 | 6.05E-05 |
| Contractile | PAXBP1 | 0.002192 | 0.27556 | 0.242 | 0.131 | 1 |
| Intermedia | YPEL3 | 6.93E-09 | 0.275368 | 0.477 | 0.391 | 0.000121 |
| Terminal | ATP2B1 | 1.34E-10 | 0.275104 | 0.244 | 0.141 | 2.35E-06 |
| Contractile | YPEL5 | 0.006016 | 0.274722 | 0.418 | 0.297 | 1 |
| Terminal | PSMD2 | 9.76E-08 | 0.274597 | 0.34 | 0.238 | 0.001705 |
| Terminal | CEP170 | 4.54E-10 | 0.274571 | 0.346 | 0.221 | 7.94E-06 |
| Terminal | NID1 | 1.71E-20 | 0.274534 | 0.121 | 0.033 | 3E-16 |
| Contractile | ENDOU | 1.55E-06 | 0.274422 | 0.121 | 0.03 | 0.027126 |
| Contractile | PGRMC2 | 0.006828 | 0.274145 | 0.593 | 0.463 | 1 |
| Contractile | PIAS1 | 3.95E-12 | 0.274048 | 0.286 | 0.181 | 6.91E-08 |
| Contractile | CALM1 | 0.001288 | 0.274002 | 0.934 | 0.915 | 1 |
| Terminal | LAT2 | 2.12E-13 | 0.273932 | 0.239 | 0.125 | 3.7E-09 |
| Terminal | PSME2 | 5.4E-08 | 0.273874 | 0.243 | 0.154 | 0.000944 |
| Contractile | CACNA2D1 | 1.57E-13 | 0.273772 | 0.384 | 0.257 | 2.74E-09 |
| Terminal | TIMM10B | 4.33E-09 | 0.273618 | 0.311 | 0.2 | 7.58E-05 |
| Pericytes | CLIC2 | 0.000102 | 0.273548 | 0.1 | 0.029 | 1 |
| Terminal | SLC29A1 | 3.1E-14 | 0.273423 | 0.214 | 0.104 | 5.42E-10 |
| Terminal | PSENEN | 4.87E-09 | 0.273275 | 0.527 | 0.396 | 8.52E-05 |
| Contractile | PBXIP1 | 6.58E-13 | 0.273056 | 0.411 | 0.287 | 1.15E-08 |
| Terminal | ST8SIA1 | 7.85E-21 | 0.27285 | 0.139 | 0.041 | 1.37E-16 |
| Contractile | DYSF | 0.000121 | 0.272792 | 0.121 | 0.04 | 1 |
| Contractile | NEDD4 | 0.004709 | 0.272655 | 0.176 | 0.09 | 1 |
| Contractile | CASKIN2 | 1.93E-06 | 0.272565 | 0.143 | 0.04 | 0.03376 |
| IFN_Interm | XKR8 | 0.001277 | 0.272523 | 0.105 | 0.015 | 1 |
| Terminal | CASP3 | 1.23E-07 | 0.272489 | 0.309 | 0.212 | 0.002151 |
| Terminal | EMC6 | 3.48E-07 | 0.272432 | 0.45 | 0.352 | 0.006084 |
| Terminal | NDUFV3 | 1.14E-08 | 0.271664 | 0.543 | 0.422 | 0.0002 |
| Contractile | RNF19A | 3.02E-12 | 0.271524 | 0.594 | 0.459 | 5.28E-08 |
| Fibroblasts | DPY19L4 | 0.00798 | 0.271381 | 0.15 | 0.065 | 1 |
| Contractile | ADARB1 | 0.007278 | 0.271205 | 0.527 | 0.407 | 1 |
| Contractile | GPRC5B | 0.000351 | 0.271144 | 0.121 | 0.043 | 1 |
| Terminal | BAD | 1.36E-08 | 0.270849 | 0.357 | 0.249 | 0.000238 |
| Contractile | C3ORF70 | 1.76E-17 | 0.270764 | 0.125 | 0.042 | 3.08E-13 |
| Contractile | PLEKHB2 | 8.82E-13 | 0.270585 | 0.239 | 0.14 | 1.54E-08 |
| Terminal | TIPARP | 3.75E-23 | 0.270576 | 0.143 | 0.04 | 6.56E-19 |
| Terminal | GPX8 | 1.13E-08 | 0.270534 | 0.532 | 0.42 | 0.000198 |
| Fibroblasts | RAB3IL1 | 2.53E-07 | 0.270521 | 0.117 | 0.02 | 0.004429 |
| Terminal | GON7 | 3.59E-07 | 0.270385 | 0.381 | 0.278 | 0.00628 |
| Terminal | IDH2 | 1.97E-06 | 0.270359 | 0.432 | 0.339 | 0.034404 |
| Contractile | ENSSSCG0( | 0.008363 | 0.270212 | 0.132 | 0.064 | 1 |
| Terminal | PSMG4 | 5.84E-08 | 0.270055 | 0.414 | 0.308 | 0.001021 |
| Intermedia | PENK | 1.83E-05 | 0.270034 | 0.592 | 0.513 | 0.319404 |
| Intermedia | RNF130 | 1.48E-06 | 0.269863 | 0.476 | 0.407 | 0.02595 |
| Contractile | PLXDC1 | 1.17E-12 | 0.269841 | 0.481 | 0.342 | 2.05E-08 |

| | | | | | | |
|---|---|---|---|---|---|---|
| Intermedia | RASL11B | 1.28E-11 | 0.269665 | 0.134 | 0.061 | 2.24E-07 |
| Terminal | GNG5 | 5.98E-10 | 0.269551 | 0.366 | 0.244 | 1.05E-05 |
| Contractile | SSC5D | 7.92E-12 | 0.269351 | 0.339 | 0.23 | 1.38E-07 |
| Terminal | NFATC2 | 1.94E-17 | 0.269218 | 0.106 | 0.029 | 3.39E-13 |
| Fibroblasts | C5ORF49 | 2.55E-07 | 0.269149 | 0.117 | 0.019 | 0.004457 |
| Contractile | CD151 | 1.18E-11 | 0.269096 | 0.746 | 0.633 | 2.07E-07 |
| Terminal | SMDT1 | 5.71E-06 | 0.26889 | 0.49 | 0.402 | 0.099764 |
| Terminal | UBE2L3 | 2.9E-06 | 0.268817 | 0.528 | 0.434 | 0.050694 |
| Intermedia | STK17B | 0.000187 | 0.268625 | 0.182 | 0.136 | 1 |
| Terminal | SLBP | 1.94E-10 | 0.268479 | 0.241 | 0.139 | 3.39E-06 |
| Terminal | ETV1 | 1.69E-09 | 0.268352 | 0.254 | 0.153 | 2.95E-05 |
| Intermedia | EDIL3 | 8.04E-16 | 0.268231 | 0.548 | 0.376 | 1.41E-11 |
| Terminal | GPNMB | 3.51E-08 | 0.267975 | 0.137 | 0.072 | 0.000614 |
| Terminal | SRP72 | 2.81E-11 | 0.26783 | 0.359 | 0.229 | 4.91E-07 |
| Terminal | FSCN1 | 6.2E-14 | 0.267807 | 0.336 | 0.197 | 1.08E-09 |
| Contractile | TINAGL1 | 4.23E-05 | 0.267744 | 0.407 | 0.218 | 0.739889 |
| Terminal | SPCS3 | 3.8E-06 | 0.267712 | 0.546 | 0.461 | 0.066368 |
| Terminal | TMEM230 | 6.68E-13 | 0.26768 | 0.302 | 0.174 | 1.17E-08 |
| Terminal | ADIPOR1 | 1.72E-10 | 0.267394 | 0.224 | 0.127 | 3E-06 |
| Intermedia | TMEM223 | 3.4E-05 | 0.267136 | 0.498 | 0.461 | 0.595122 |
| Contractile | NFIA | 5.63E-14 | 0.267097 | 0.418 | 0.282 | 9.84E-10 |
| Contractile | ENSSSCG0( | 0.00596 | 0.267045 | 0.165 | 0.085 | 1 |
| Contractile | ZEB1 | 3.9E-10 | 0.266983 | 0.325 | 0.225 | 6.83E-06 |
| Terminal | PGM3 | 6.57E-13 | 0.26659 | 0.277 | 0.158 | 1.15E-08 |
| Contractile | SGCA | 1.15E-11 | 0.266482 | 0.277 | 0.174 | 2.01E-07 |
| Terminal | CLTA | 2.3E-09 | 0.26637 | 0.723 | 0.64 | 4.02E-05 |
| Terminal | YIPF3 | 6.9E-08 | 0.266288 | 0.488 | 0.376 | 0.001206 |
| Intermedia | NOX4 | 9.63E-05 | 0.26627 | 0.423 | 0.363 | 1 |
| IFN_Interm | MAPK3 | 0.004355 | 0.266209 | 0.158 | 0.035 | 1 |
| Terminal | MRPS25 | 5.55E-08 | 0.266179 | 0.277 | 0.184 | 0.000969 |
| Intermedia | COL6A1 | 1.47E-11 | 0.266116 | 0.954 | 0.923 | 2.57E-07 |
| Intermedia | C1QTNF1 | 3.59E-08 | 0.266064 | 0.72 | 0.669 | 0.000628 |
| Terminal | RASSF1 | 6.23E-10 | 0.265943 | 0.277 | 0.169 | 1.09E-05 |
| Contractile | GPC6 | 9.44E-14 | 0.265899 | 0.2 | 0.103 | 1.65E-09 |
| Contractile | SLC14A1 | 3.79E-18 | 0.265883 | 0.13 | 0.045 | 6.62E-14 |
| Intermedia | HOXA10 | 6.08E-06 | 0.2658 | 0.237 | 0.175 | 0.106287 |
| Fibroblasts | ENSSSCG0( | 0.000275 | 0.265729 | 0.1 | 0.025 | 1 |
| Contractile | CEP95 | 0.004098 | 0.265691 | 0.154 | 0.075 | 1 |
| Fibroblasts | N6AMT1 | 0.004433 | 0.265621 | 0.133 | 0.052 | 1 |
| IFN_Interm | SHFL | 0.008092 | 0.265543 | 0.263 | 0.085 | 1 |
| Terminal | BACH1 | 5.81E-11 | 0.265543 | 0.305 | 0.187 | 1.02E-06 |
| Intermedia | CXCL12 | 6.71E-13 | 0.265455 | 0.101 | 0.036 | 1.17E-08 |
| IFN_Interm | ZNF782 | 4.59E-05 | 0.265391 | 0.105 | 0.01 | 0.802725 |
| Intermedia | CTSB | 7.07E-10 | 0.265388 | 0.8 | 0.73 | 1.24E-05 |
| Terminal | XPOT | 1.87E-11 | 0.265281 | 0.306 | 0.184 | 3.26E-07 |
| Intermedia | VKORC1 | 5.34E-07 | 0.265267 | 0.66 | 0.624 | 0.009328 |
| Terminal | IAH1 | 5.16E-10 | 0.265267 | 0.429 | 0.3 | 9.03E-06 |
| Terminal | PSMD7 | 5.6E-06 | 0.265251 | 0.465 | 0.378 | 0.09786 |
| Contractile | S100A11 | 1.9E-24 | 0.264611 | 0.968 | 0.939 | 3.32E-20 |
| Contractile | UBC | 4.41E-16 | 0.264551 | 0.945 | 0.903 | 7.7E-12 |
| Intermedia | EFEMP2 | 2.4E-07 | 0.264427 | 0.685 | 0.634 | 0.004203 |
| Terminal | MAP2K2 | 4.03E-09 | 0.264334 | 0.355 | 0.24 | 7.04E-05 |
| Terminal | RUVBL1 | 4.8E-13 | 0.26382 | 0.21 | 0.106 | 8.38E-09 |
| Terminal | CIAO2B | 8.57E-09 | 0.263805 | 0.326 | 0.216 | 0.00015 |
| Contractile | ENSSSCG0( | 5.49E-22 | 0.263762 | 0.109 | 0.026 | 9.6E-18 |
| Contractile | BLOC1S1 | 2.38E-11 | 0.263612 | 0.571 | 0.463 | 4.16E-07 |
| Terminal | MRPS6 | 1.07E-06 | 0.26359 | 0.331 | 0.238 | 0.018791 |
| Terminal | DNAJB9 | 5.03E-07 | 0.263277 | 0.274 | 0.187 | 0.008794 |
| Fibroblasts | ITM2A | 0.000144 | 0.263115 | 0.133 | 0.037 | 1 |

| | | | | | | |
|---|---|---|---|---|---|---|
| Contractile | A4GALT | 1.11E-09 | 0.263043 | 0.496 | 0.392 | 1.95E-05 |
| Contractile | TMED5 | 2.46E-13 | 0.262859 | 0.382 | 0.255 | 4.31E-09 |
| Terminal | SLC1A4 | 1.67E-20 | 0.262694 | 0.126 | 0.035 | 2.91E-16 |
| Contractile | HEY2 | 9E-13 | 0.262647 | 0.348 | 0.23 | 1.57E-08 |
| Contractile | ENSSSCG0( | 1.89E-22 | 0.26251 | 0.15 | 0.048 | 3.3E-18 |
| Terminal | EIF3H | 1.52E-06 | 0.262505 | 0.709 | 0.629 | 0.026559 |
| Contractile | ADAM19 | 1.57E-13 | 0.262446 | 0.197 | 0.103 | 2.74E-09 |
| Contractile | PKIA | 1.11E-11 | 0.262383 | 0.183 | 0.101 | 1.94E-07 |
| Contractile | AGRN | 7.09E-15 | 0.262374 | 0.171 | 0.08 | 1.24E-10 |
| Contractile | IL11RA | 5.92E-13 | 0.262268 | 0.2 | 0.107 | 1.03E-08 |
| Terminal | PALMD | 1.67E-11 | 0.261871 | 0.18 | 0.089 | 2.91E-07 |
| Intermedia | ENSSSCG0( | 1.61E-08 | 0.261759 | 0.9 | 0.869 | 0.000282 |
| IFN_Interm | ERP27 | 0.001956 | 0.261572 | 0.105 | 0.016 | 1 |
| Terminal | TUBB6 | 4.16E-05 | 0.261415 | 0.409 | 0.33 | 0.727586 |
| Terminal | EBPL | 1.15E-16 | 0.261333 | 0.169 | 0.067 | 2.01E-12 |
| Terminal | PPP1R3C | 1.38E-06 | 0.260998 | 0.32 | 0.233 | 0.024206 |
| Contractile | HIP1 | 6.95E-12 | 0.260822 | 0.343 | 0.227 | 1.22E-07 |
| Terminal | BET1 | 1.99E-07 | 0.260641 | 0.305 | 0.21 | 0.003482 |
| Contractile | GYG2 | 4.62E-11 | 0.260468 | 0.241 | 0.146 | 8.07E-07 |
| Contractile | SASH1 | 4E-10 | 0.260462 | 0.427 | 0.32 | 6.99E-06 |
| Terminal | PDGFA | 4.94E-11 | 0.260269 | 0.399 | 0.267 | 8.63E-07 |
| Contractile | CTNNA1 | 1E-09 | 0.260248 | 0.573 | 0.465 | 1.75E-05 |
| Terminal | SELENOT | 1.4E-05 | 0.260246 | 0.449 | 0.358 | 0.245305 |
| Terminal | RNF13 | 1.04E-09 | 0.260151 | 0.385 | 0.259 | 1.82E-05 |
| Intermedia | B4GALT1 | 4.29E-06 | 0.260134 | 0.356 | 0.284 | 0.07497 |
| Terminal | PSMD12 | 3.31E-07 | 0.260131 | 0.394 | 0.292 | 0.005789 |
| Terminal | MAGOH | 3.25E-06 | 0.260094 | 0.329 | 0.241 | 0.056744 |
| Terminal | PI4K2B | 9.32E-11 | 0.260014 | 0.305 | 0.187 | 1.63E-06 |
| Terminal | FOLR3 | 1.55E-05 | 0.26 | 0.346 | 0.262 | 0.271673 |
| Fibroblasts | MBD1 | 0.003971 | 0.259906 | 0.15 | 0.06 | 1 |
| Terminal | TMX1 | 6.65E-10 | 0.259875 | 0.314 | 0.2 | 1.16E-05 |
| Terminal | IL17D | 7.34E-14 | 0.259834 | 0.202 | 0.097 | 1.28E-09 |
| Contractile | A2M | 0.002239 | 0.25951 | 0.967 | 0.79 | 1 |
| Contractile | TBC1D9 | 0.002769 | 0.259267 | 0.121 | 0.051 | 1 |
| Contractile | STOM | 9.07E-17 | 0.258987 | 0.576 | 0.413 | 1.59E-12 |
| IFN_Interm | CGAS | 2.08E-07 | 0.258818 | 0.158 | 0.014 | 0.003645 |
| Contractile | SIK2 | 0.00192 | 0.258796 | 0.143 | 0.063 | 1 |
| Intermedia | DUSP6 | 0.005213 | 0.258463 | 0.122 | 0.091 | 1 |
| Terminal | TOMM5.1 | 1.47E-08 | 0.25837 | 0.281 | 0.184 | 0.000257 |
| Contractile | HIBADH | 0.002219 | 0.25824 | 0.176 | 0.087 | 1 |
| Contractile | BLTP3B | 1.39E-09 | 0.258208 | 0.41 | 0.302 | 2.43E-05 |
| Contractile | CUX1 | 1.74E-10 | 0.257965 | 0.675 | 0.566 | 3.04E-06 |
| Contractile | ANKRD27 | 0.003021 | 0.25785 | 0.143 | 0.065 | 1 |
| Terminal | NDUFB5 | 3.75E-09 | 0.257817 | 0.519 | 0.392 | 6.56E-05 |
| Terminal | MRPS24 | 2.41E-08 | 0.257777 | 0.329 | 0.225 | 0.00042 |
| Terminal | DOK1 | 1.39E-10 | 0.257353 | 0.154 | 0.075 | 2.43E-06 |
| Terminal | NFATC1 | 3.16E-11 | 0.25721 | 0.177 | 0.088 | 5.52E-07 |
| Terminal | MRPL13 | 1.68E-07 | 0.257053 | 0.269 | 0.179 | 0.002929 |
| Intermedia | RFLNB | 1.62E-09 | 0.257029 | 0.297 | 0.199 | 2.83E-05 |
| Terminal | DPCD | 3.24E-10 | 0.257009 | 0.192 | 0.105 | 5.66E-06 |
| Contractile | ENSSSCG0( | 4.01E-18 | 0.256962 | 0.137 | 0.049 | 7.01E-14 |
| Contractile | ZNF362 | 0.001122 | 0.256857 | 0.187 | 0.088 | 1 |
| Intermedia | CNPY2 | 1.04E-07 | 0.256509 | 0.449 | 0.368 | 0.001814 |
| Contractile | GALT | 0.004821 | 0.256415 | 0.176 | 0.091 | 1 |
| Contractile | JCAD | 1.3E-11 | 0.256395 | 0.228 | 0.135 | 2.27E-07 |
| Terminal | ALG5 | 2.09E-12 | 0.256119 | 0.252 | 0.139 | 3.66E-08 |
| Terminal | SSBP1 | 2.17E-11 | 0.256018 | 0.292 | 0.174 | 3.79E-07 |
| Contractile | FBLN1 | 0.00382 | 0.256016 | 0.484 | 0.35 | 1 |
| Terminal | TTLL3 | 1.15E-08 | 0.255883 | 0.706 | 0.62 | 0.000201 |

| | | | | | | |
|---|---|---|---|---|---|---|
| Fibroblasts | ENSSSCG0( | 0.005872 | 0.255865 | 0.1 | 0.034 | 1 |
| Contractile | PBX2 | 0.000995 | 0.255846 | 0.198 | 0.094 | 1 |
| Contractile | STK38L | 1.89E-10 | 0.255564 | 0.532 | 0.411 | 3.31E-06 |
| Contractile | PPM1L | 4.63E-05 | 0.255478 | 0.143 | 0.048 | 0.809027 |
| Contractile | ARID4A | 1.75E-11 | 0.255412 | 0.328 | 0.216 | 3.05E-07 |
| Terminal | ALDH1L2 | 6.51E-10 | 0.255286 | 0.32 | 0.204 | 1.14E-05 |
| Intermedia | FOXS1 | 0.000492 | 0.255084 | 0.492 | 0.454 | 1 |
| Terminal | SDF2 | 7.35E-08 | 0.254888 | 0.39 | 0.279 | 0.001285 |
| Contractile | ENSSSCG0( | 0.000821 | 0.254795 | 0.132 | 0.052 | 1 |
| Contractile | TNS2 | 5.09E-11 | 0.254779 | 0.399 | 0.285 | 8.9E-07 |
| Contractile | KCNH2 | 0.004495 | 0.254709 | 0.121 | 0.053 | 1 |
| Terminal | TRAPPC2L | 3.1E-06 | 0.254696 | 0.417 | 0.325 | 0.054201 |
| Contractile | NPY1R | 2.54E-05 | 0.254685 | 0.143 | 0.046 | 0.444463 |
| Fibroblasts | ENSSSCG0( | 0.008582 | 0.254574 | 0.117 | 0.045 | 1 |
| Contractile | DMPK | 0.004875 | 0.254427 | 0.33 | 0.209 | 1 |
| Contractile | FIBIN | 1.39E-17 | 0.254412 | 0.315 | 0.176 | 2.42E-13 |
| Contractile | PRKAR1A | 0.004494 | 0.254335 | 0.637 | 0.511 | 1 |
| Terminal | THOC7 | 4.31E-09 | 0.254168 | 0.535 | 0.417 | 7.54E-05 |
| Terminal | DNMT1 | 5.95E-06 | 0.254098 | 0.224 | 0.153 | 0.104012 |
| Terminal | AHR | 4.71E-10 | 0.253994 | 0.244 | 0.144 | 8.23E-06 |
| Terminal | MGAT4B | 1.24E-11 | 0.253928 | 0.294 | 0.172 | 2.16E-07 |
| Terminal | MPC1 | 2.41E-06 | 0.253883 | 0.525 | 0.431 | 0.042128 |
| Contractile | COL4A6 | 2.42E-09 | 0.253658 | 0.283 | 0.193 | 4.24E-05 |
| Terminal | SAT1 | 1.2E-06 | 0.253601 | 0.486 | 0.383 | 0.02106 |
| Terminal | VAT1 | 5.08E-08 | 0.253435 | 0.368 | 0.259 | 0.000888 |
| Terminal | SET | 2.84E-07 | 0.253411 | 0.734 | 0.659 | 0.004973 |
| Terminal | SEC11A | 1.79E-07 | 0.253193 | 0.56 | 0.458 | 0.003124 |
| Terminal | SIGIRR | 5.22E-18 | 0.252943 | 0.151 | 0.053 | 9.12E-14 |
| Contractile | TOB2 | 5.74E-09 | 0.252887 | 0.501 | 0.4 | 0.0001 |
| Terminal | GLUD1 | 8.4E-08 | 0.252546 | 0.505 | 0.391 | 0.001468 |
| Terminal | BAX | 9.78E-10 | 0.252439 | 0.21 | 0.119 | 1.71E-05 |
| Terminal | FKBP7 | 7.2E-06 | 0.252379 | 0.425 | 0.339 | 0.125845 |
| Contractile | BCL7A | 0.000186 | 0.252277 | 0.11 | 0.035 | 1 |
| Terminal | ANP32B | 1.7E-06 | 0.25212 | 0.706 | 0.62 | 0.029671 |
| Contractile | NF2 | 0.005392 | 0.251944 | 0.143 | 0.069 | 1 |
| Contractile | SYT11 | 3.94E-10 | 0.251816 | 0.317 | 0.219 | 6.89E-06 |
| Terminal | PRDX5 | 1.31E-08 | 0.251771 | 0.207 | 0.123 | 0.000228 |
| Contractile | DPY19L3 | 0.001561 | 0.25171 | 0.121 | 0.048 | 1 |
| Terminal | ATP6V0C | 1.96E-13 | 0.251689 | 0.236 | 0.121 | 3.43E-09 |
| Terminal | BUD23 | 1.2E-06 | 0.251453 | 0.331 | 0.243 | 0.021047 |
| Terminal | RWDD1 | 5.54E-07 | 0.25134 | 0.632 | 0.532 | 0.009691 |
| Contractile | CCDC9B | 2.68E-13 | 0.251248 | 0.152 | 0.071 | 4.69E-09 |
| Terminal | PXDC1 | 6.71E-08 | 0.251167 | 0.432 | 0.318 | 0.001173 |
| Contractile | PHACTR2 | 0.003175 | 0.251006 | 0.165 | 0.079 | 1 |
| Intermedia | FTL | 4.61E-15 | 0.250959 | 0.993 | 0.972 | 8.06E-11 |
| IFN_Interm | CIITA | 3.62E-11 | 0.250884 | 0.105 | 0.004 | 6.32E-07 |
| Terminal | ERH | 2.26E-05 | 0.250824 | 0.631 | 0.565 | 0.395104 |
| Terminal | APH1A | 2.33E-08 | 0.25074 | 0.213 | 0.128 | 0.000407 |
| Terminal | TM9SF1 | 1.74E-08 | 0.250694 | 0.343 | 0.236 | 0.000304 |
| Terminal | SMIM7 | 1.74E-05 | 0.250617 | 0.494 | 0.402 | 0.304442 |
| Terminal | PTPN2 | 3.02E-08 | 0.250493 | 0.254 | 0.163 | 0.000528 |
| Terminal | EEF2 | 8.47E-07 | 0.250265 | 0.864 | 0.827 | 0.014808 |
| Terminal | ELOVL1 | 1.51E-12 | 0.250179 | 0.2 | 0.1 | 2.64E-08 |
| IFN_Interm | DEGS2 | 2.31E-37 | 0.250178 | 0.105 | 0.001 | 4.03E-33 |
| Intermedia | ENSSSCG0( | 2.03E-07 | 0.25008 | 0.444 | 0.364 | 0.003541 |

**Table S4. Marker genes for mesenchymal cell clusters in individual mouse BCT plaque scRNA-seq data. Corresponds to Fig. 1C.**
The listed genes have significantly higher expression in the indicated cluster than all other clusters combined.
Analysis was performed using the Wilcoxon test in Seurat v4.4.0 with P values adjusted for multiple testing by the Bonferroni method.

| cluster | gene | p_val | avg_log2F( | pct.1 | pct.2 | p_val_adj |
|---|---|---|---|---|---|---|
| Terminal | Col2a1 | 2.2091E-252 | 5.026369 | 0.838 | 0.218 | 2.9255E-248 |
| Terminal | Chil1 | 2.538E-178 | 4.884945 | 0.854 | 0.378 | 3.3611E-174 |
| Terminal | Ibsp | 5.7471E-188 | 4.782929 | 0.851 | 0.333 | 7.6108E-184 |
| Terminal | Cytl1 | 3.6742E-179 | 4.704297 | 0.858 | 0.373 | 4.8657E-175 |
| Terminal | Chad | 1.8624E-191 | 4.343887 | 0.661 | 0.119 | 2.4663E-187 |
| Terminal | 3110079O | 1.6684E-119 | 3.559042 | 0.446 | 0.064 | 2.2095E-115 |
| Terminal | Bpifb1 | 1.0764E-154 | 3.557819 | 0.563 | 0.095 | 1.4255E-150 |
| Terminal | Lcn2 | 3.2367E-185 | 2.800563 | 0.909 | 0.433 | 4.2864E-181 |
| Terminal | Comp | 5.1509E-113 | 2.768993 | 0.683 | 0.245 | 6.8214E-109 |
| Terminal | Cst3 | 2.5943E-236 | 2.738546 | 1 | 0.997 | 3.4356E-232 |
| Terminal | Mia | 2.2595E-145 | 2.709511 | 0.642 | 0.171 | 2.9922E-141 |
| Terminal | Saa3 | 5.06392E-63 | 2.596204 | 0.552 | 0.24 | 6.70615E-59 |
| Contractile | Myl9 | 2.845E-163 | 2.557338 | 0.997 | 0.441 | 3.7677E-159 |
| Terminal | Hp | 1.2789E-137 | 2.540424 | 0.89 | 0.431 | 1.6937E-133 |
| Contractile | Acta2 | 2.1463E-145 | 2.52276 | 1 | 0.675 | 2.8424E-141 |
| Contractile | Cnn1 | 4.2763E-196 | 2.49834 | 0.979 | 0.236 | 5.6632E-192 |
| Intermediate | Ly6a | 9.6969E-175 | 2.497939 | 0.764 | 0.258 | 1.2842E-170 |
| Contractile | Sost | 4.2739E-197 | 2.489861 | 0.821 | 0.112 | 5.6599E-193 |
| Rgs5_Contractile | Rgs5 | 8.07586E-47 | 2.471818 | 0.573 | 0.238 | 1.06949E-42 |
| Terminal | Gpx3 | 3.4092E-114 | 2.445138 | 0.539 | 0.131 | 4.5148E-110 |
| Contractile | Tagln | 5.4806E-156 | 2.420635 | 0.997 | 0.416 | 7.258E-152 |
| Terminal | Mmp13 | 1.07991E-42 | 2.411611 | 0.387 | 0.157 | 1.43012E-38 |
| Terminal | Omd | 2.2065E-115 | 2.389784 | 0.475 | 0.085 | 2.922E-111 |
| Terminal | Alpl | 9.26429E-86 | 2.386146 | 0.324 | 0.04 | 1.22687E-81 |
| Contractile | Tpm2 | 4.1549E-140 | 2.363616 | 0.988 | 0.545 | 5.5023E-136 |
| Intermediate | Vcam1 | 6.4874E-189 | 2.362737 | 0.916 | 0.485 | 8.5913E-185 |
| Terminal | Fmod | 6.2126E-182 | 2.333158 | 0.761 | 0.201 | 8.2274E-178 |
| Contractile | Myh11 | 5.9599E-178 | 2.278796 | 0.976 | 0.25 | 7.8927E-174 |
| Contractile | Mfap4 | 1.7044E-134 | 2.238815 | 0.756 | 0.161 | 2.2571E-130 |
| Terminal | Pth1r | 3.3833E-123 | 2.233149 | 0.68 | 0.241 | 4.4805E-119 |
| Contractile | Csrp2 | 3.2115E-135 | 2.23211 | 0.976 | 0.57 | 4.253E-131 |
| Terminal | Wif1 | 7.342E-117 | 2.225074 | 0.633 | 0.217 | 9.723E-113 |
| Contractile | Csrp1 | 1.2377E-144 | 2.211573 | 0.991 | 0.58 | 1.639E-140 |
| Terminal | Hapln1 | 1.1765E-220 | 2.187905 | 0.607 | 0.034 | 1.5581E-216 |
| Terminal | Pcolce2 | 3.9617E-111 | 2.182463 | 0.662 | 0.242 | 5.2464E-107 |
| Contractile | Ppp1r14a | 4.0856E-163 | 2.165704 | 0.926 | 0.247 | 5.4105E-159 |
| Terminal | Acan | 6.2887E-217 | 2.157018 | 0.835 | 0.211 | 8.3281E-213 |
| Rgs5_Contractile | Actb | 6.20065E-79 | 2.137981 | 0.983 | 0.98 | 8.21152E-75 |
| Terminal | 1500015O | 4.7534E-228 | 2.135409 | 0.987 | 0.474 | 6.2949E-224 |
| Contractile | Sh3bgr | 1.4019E-207 | 2.122515 | 0.841 | 0.115 | 1.8566E-203 |
| Terminal | Saa2 | 2.54972E-33 | 2.117435 | 0.224 | 0.064 | 3.3766E-29 |
| Contractile | Fbxl22 | 1.6189E-172 | 2.102129 | 0.729 | 0.099 | 2.1439E-168 |
| Contractile | Dstn | 7.6146E-139 | 2.101711 | 1 | 0.876 | 1.0084E-134 |
| Contractile | Tmsb4x | 1.373E-135 | 2.093245 | 0.997 | 0.894 | 1.8183E-131 |
| Contractile | Itga8 | 8.4555E-176 | 2.071713 | 0.974 | 0.246 | 1.1198E-171 |
| Terminal | Scrg1 | 2.7515E-156 | 2.053136 | 0.455 | 0.024 | 3.6438E-152 |
| Rgs5_Contractile | Myh11 | 1.6468E-103 | 2.045273 | 0.861 | 0.283 | 2.1809E-99 |
| Rgs5_Contractile | Tagln | 3.18411E-90 | 2.039425 | 0.903 | 0.443 | 4.21672E-86 |
| Intermediate | Wisp2 | 1.64587E-98 | 2.037281 | 0.726 | 0.374 | 2.17962E-94 |
| Rgs5_Contractile | Tpm2 | 4.83061E-85 | 1.991878 | 0.92 | 0.565 | 6.39718E-81 |
| Rgs5_Contractile | Sparcl1 | 8.58456E-24 | 1.981714 | 0.385 | 0.175 | 1.13685E-19 |
| Terminal | Col11a1 | 2.731E-133 | 1.974962 | 0.375 | 0.011 | 3.6167E-129 |
| Terminal | Sparc | 1.3197E-139 | 1.966525 | 1 | 0.991 | 1.7477E-135 |
| Rgs5_Contractile | Myl9 | 1.07042E-83 | 1.957323 | 0.903 | 0.467 | 1.41756E-79 |

| | | | | | | |
|---|---|---|---|---|---|---|
| Contractile | Myl6 | 8.4917E-139 | 1.950951 | 1 | 0.978 | 1.1246E-134 |
| Terminal | Saa1 | 1.59035E-52 | 1.929992 | 0.233 | 0.035 | 2.10609E-48 |
| Terminal | Col11a2 | 5.1461E-211 | 1.928266 | 0.588 | 0.034 | 6.815E-207 |
| Contractile | Lmod1 | 5.2045E-152 | 1.927469 | 0.941 | 0.294 | 6.8923E-148 |
| Intermediate | Sfrp2 | 6.79877E-79 | 1.916322 | 0.565 | 0.219 | 9.00361E-75 |
| Rgs5_Contractile | Acta2 | 1.0749E-76 | 1.89403 | 0.938 | 0.692 | 1.42349E-72 |
| Contractile | Fblim1 | 3.0943E-146 | 1.875485 | 0.935 | 0.313 | 4.0977E-142 |
| Terminal | Spp1 | 3.7007E-180 | 1.869854 | 1 | 0.756 | 4.9009E-176 |
| Contractile | Flna | 1.5339E-119 | 1.85875 | 0.971 | 0.54 | 2.0313E-115 |
| Intermediate | Thbs1 | 1.84266E-72 | 1.855644 | 0.512 | 0.186 | 2.44024E-68 |
| Contractile | Actg2 | 1.326E-115 | 1.840234 | 0.674 | 0.139 | 1.756E-111 |
| Contractile | Tpm1 | 1.0968E-121 | 1.832631 | 0.976 | 0.645 | 1.4524E-117 |
| Terminal | Col27a1 | 3.6345E-196 | 1.832042 | 0.724 | 0.148 | 4.8131E-192 |
| Rgs5_Contractile | Pam | 6.46772E-37 | 1.829055 | 0.823 | 0.876 | 8.5652E-33 |
| Rgs5_Contractile | Mylk | 6.10186E-69 | 1.824358 | 0.785 | 0.389 | 8.08069E-65 |
| Rgs5_Contractile | Itga8 | 3.20262E-76 | 1.800578 | 0.771 | 0.291 | 4.24123E-72 |
| Rgs5_Contractile | Ppp1r14a | 1.59362E-68 | 1.792137 | 0.733 | 0.29 | 2.11044E-64 |
| Contractile | Mylk | 2.417E-131 | 1.78131 | 0.938 | 0.354 | 3.2008E-127 |
| Terminal | Col9a3 | 1.543E-171 | 1.777531 | 0.493 | 0.026 | 2.0435E-167 |
| Contractile | Tns1 | 4.0814E-109 | 1.770168 | 0.876 | 0.386 | 5.4051E-105 |
| Intermediate | Cxcl12 | 2.47398E-97 | 1.767728 | 0.713 | 0.319 | 3.27629E-93 |
| Contractile | Ramp1 | 1.7461E-115 | 1.763916 | 0.853 | 0.298 | 2.3123E-111 |
| Rgs5_Contractile | Ogn | 1.30664E-52 | 1.760402 | 0.684 | 0.334 | 1.73038E-48 |
| Contractile | Fmo2 | 1.5584E-107 | 1.755161 | 0.756 | 0.209 | 2.0638E-103 |
| Rgs5_Contractile | Tpm1 | 5.55795E-68 | 1.751362 | 0.885 | 0.665 | 7.3604E-64 |
| Contractile | Actn1 | 4.5441E-116 | 1.749076 | 0.953 | 0.597 | 6.0178E-112 |
| Contractile | Gstm1 | 6.9323E-119 | 1.737714 | 0.956 | 0.556 | 9.1805E-115 |
| Terminal | Crispld1 | 1.0946E-146 | 1.737585 | 0.765 | 0.265 | 1.4495E-142 |
| Contractile | Npnt | 7.7061E-141 | 1.734433 | 0.809 | 0.191 | 1.0205E-136 |
| Rgs5_Contractile | Flna | 2.06359E-69 | 1.723049 | 0.868 | 0.564 | 2.73281E-65 |
| Contractile | Tcap | 2.4873E-139 | 1.718777 | 0.629 | 0.086 | 3.2939E-135 |
| Intermediate | Sfrp4 | 4.94623E-74 | 1.715562 | 0.577 | 0.242 | 6.5503E-70 |
| Contractile | Synpo2 | 1.3688E-141 | 1.715345 | 0.75 | 0.143 | 1.8127E-137 |
| Contractile | Itih4 | 6.7874E-106 | 1.704742 | 0.671 | 0.145 | 8.9886E-102 |
| Rgs5_Contractile | Dstn | 1.4589E-74 | 1.701433 | 0.951 | 0.886 | 1.93202E-70 |
| Contractile | Efhd1 | 1.6355E-165 | 1.698968 | 0.744 | 0.107 | 2.1658E-161 |
| Contractile | Msrb1 | 4.66366E-97 | 1.698327 | 0.806 | 0.347 | 6.17608E-93 |
| Rgs5_Contractile | Tns1 | 2.43211E-51 | 1.678182 | 0.729 | 0.418 | 3.22085E-47 |
| Contractile | St5 | 7.4232E-150 | 1.675949 | 0.759 | 0.13 | 9.8306E-146 |
| Rgs5_Contractile | Csrp2 | 4.62364E-66 | 1.673217 | 0.892 | 0.591 | 6.12309E-62 |
| Contractile | Pdlim3 | 1.6113E-143 | 1.669607 | 0.785 | 0.157 | 2.1338E-139 |
| Terminal | Slpi | 9.54529E-37 | 1.668271 | 0.235 | 0.062 | 1.26408E-32 |
| Contractile | Map1b | 8.9394E-118 | 1.660406 | 0.9 | 0.352 | 1.1838E-113 |
| Terminal | Fzd9 | 4.7371E-189 | 1.656507 | 0.643 | 0.093 | 6.2734E-185 |
| Rgs5_Contractile | Cnn1 | 5.01453E-79 | 1.655995 | 0.809 | 0.278 | 6.64075E-75 |
| Contractile | Ogn | 1.65366E-94 | 1.654589 | 0.815 | 0.304 | 2.18994E-90 |
| Contractile | Nov | 7.1363E-100 | 1.636192 | 0.803 | 0.267 | 9.45056E-96 |
| Contractile | Fhl1 | 1.4903E-101 | 1.634002 | 0.953 | 0.63 | 1.97365E-97 |
| Contractile | Cald1 | 2.735E-120 | 1.631671 | 0.994 | 0.749 | 3.6219E-116 |
| Intermediate | Ly6c1 | 3.56882E-93 | 1.630272 | 0.559 | 0.193 | 4.72618E-89 |
| Terminal | Lect1 | 1.43696E-89 | 1.62497 | 0.282 | 0.015 | 1.90297E-85 |
| Contractile | Lpp | 3.803E-110 | 1.605401 | 0.865 | 0.325 | 5.0364E-106 |
| Rgs5_Contractile | Hspb1 | 1.03298E-50 | 1.599468 | 0.865 | 0.754 | 1.36798E-46 |
| Contractile | Fgl2 | 2.1762E-119 | 1.595757 | 0.806 | 0.22 | 2.882E-115 |
| Rgs5_Contractile | Sost | 8.15867E-66 | 1.595612 | 0.59 | 0.161 | 1.08045E-61 |
| Contractile | 1190005I0 | 7.024E-137 | 1.593294 | 0.694 | 0.124 | 9.3019E-133 |
| Terminal | Rarres1 | 1.6133E-116 | 1.591582 | 0.835 | 0.4 | 2.1365E-112 |
| Rgs5_Contractile | Rbpms | 6.98763E-49 | 1.584771 | 0.656 | 0.336 | 9.25371E-45 |
| Rgs5_Contractile | Csrp1 | 3.47803E-67 | 1.582748 | 0.892 | 0.604 | 4.60596E-63 |

| | | | | | | |
|---|---|---|---|---|---|---|
| Contractile | Slmap | 2.46E-113 | 1.580564 | 0.85 | 0.307 | 3.2578E-109 |
| Rgs5_Contractile | Cped1 | 3.6296E-43 | 1.57062 | 0.618 | 0.309 | 4.80668E-39 |
| Intermediate | Eln | 1.12376E-23 | 1.565821 | 0.699 | 0.52 | 1.4882E-19 |
| Rgs5_Contractile | Myl6 | 2.82809E-78 | 1.561131 | 0.986 | 0.981 | 3.74525E-74 |
| Rgs5_Contractile | Tinagl1 | 1.22875E-39 | 1.55899 | 0.514 | 0.219 | 1.62724E-35 |
| Contractile | Cped1 | 7.65129E-97 | 1.545866 | 0.785 | 0.274 | 1.01326E-92 |
| Terminal | Srgn | 4.96007E-91 | 1.534869 | 0.519 | 0.154 | 6.56862E-87 |
| Contractile | Sgcg | 7.3545E-145 | 1.528371 | 0.621 | 0.081 | 9.7395E-141 |
| Intermediate | Timp1 | 1.28934E-84 | 1.525377 | 0.905 | 0.652 | 1.70748E-80 |
| Terminal | Mmp3 | 5.53546E-56 | 1.525067 | 0.764 | 0.451 | 7.33061E-52 |
| Intermediate | Ltbp2 | 4.8046E-124 | 1.524757 | 0.774 | 0.34 | 6.3627E-120 |
| Rgs5_Contractile | Lpp | 3.25215E-41 | 1.517907 | 0.653 | 0.367 | 4.30682E-37 |
| Rgs5_Contractile | Filip1l | 8.88546E-31 | 1.515523 | 0.552 | 0.315 | 1.1767E-26 |
| Terminal | Dcn | 1.328E-174 | 1.513292 | 0.998 | 0.701 | 1.7586E-170 |
| Contractile | Ckb | 7.84606E-66 | 1.504859 | 0.75 | 0.378 | 1.03905E-61 |
| Terminal | Sox9 | 2.067E-153 | 1.504166 | 0.729 | 0.212 | 2.7373E-149 |
| Rgs5_Contractile | Ncam1 | 3.21256E-29 | 1.502852 | 0.604 | 0.422 | 4.2544E-25 |
| Intermediate | Col3a1 | 1.5612E-126 | 1.498444 | 0.993 | 0.916 | 2.0675E-122 |
| Contractile | E030013I1 | 3.065E-127 | 1.488674 | 0.538 | 0.062 | 4.059E-123 |
| Terminal | Fap | 1.5645E-159 | 1.488609 | 0.676 | 0.156 | 2.0718E-155 |
| Contractile | Hspb6 | 4.38099E-82 | 1.477294 | 0.688 | 0.248 | 5.80175E-78 |
| Contractile | Spint2 | 1.5158E-105 | 1.475688 | 0.738 | 0.198 | 2.0074E-101 |
| Contractile | Rarres2 | 5.93543E-95 | 1.472868 | 0.935 | 0.52 | 7.86029E-91 |
| Contractile | Rbpms | 9.893E-99 | 1.469034 | 0.818 | 0.301 | 1.31013E-94 |
| Contractile | Vim | 1.6826E-97 | 1.464298 | 1 | 0.961 | 2.22827E-93 |
| Contractile | Pbxip1 | 1.23411E-83 | 1.462483 | 0.865 | 0.499 | 1.63434E-79 |
| Rgs5_Contractile | Nfia | 1.67157E-27 | 1.461893 | 0.622 | 0.49 | 2.21366E-23 |
| Rgs5_Contractile | Ptrf | 3.06775E-42 | 1.461622 | 0.747 | 0.568 | 4.06263E-38 |
| Rgs5_Contractile | Slmap | 2.06553E-36 | 1.45003 | 0.618 | 0.352 | 2.73538E-32 |
| Intermediate | Crlf1 | 1.0549E-102 | 1.443481 | 0.582 | 0.179 | 1.39705E-98 |
| Rgs5_Contractile | Ccdc3 | 6.59719E-40 | 1.437752 | 0.455 | 0.16 | 8.73666E-36 |
| Contractile | Hspb1 | 3.97981E-88 | 1.435743 | 0.959 | 0.736 | 5.27046E-84 |
| Terminal | Ccdc80 | 1.9897E-121 | 1.433199 | 0.931 | 0.675 | 2.6349E-117 |
| Rgs5_Contractile | Rbp4 | 1.50076E-20 | 1.429536 | 0.49 | 0.29 | 1.98745E-16 |
| Terminal | Gdf10 | 1.2366E-144 | 1.42581 | 0.763 | 0.226 | 1.6376E-140 |
| Rgs5_Contractile | Map1b | 2.96434E-34 | 1.423535 | 0.646 | 0.4 | 3.92567E-30 |
| Rgs5_Contractile | Lmod1 | 3.00052E-53 | 1.422503 | 0.726 | 0.339 | 3.9736E-49 |
| Contractile | Rbp4 | 9.52086E-75 | 1.422116 | 0.709 | 0.25 | 1.26085E-70 |
| Contractile | Dmpk | 5.4725E-102 | 1.418666 | 0.668 | 0.167 | 7.24723E-98 |
| Terminal | Phlda1 | 1.2661E-125 | 1.416222 | 0.802 | 0.361 | 1.6767E-121 |
| Terminal | Enpp2 | 7.0554E-104 | 1.413751 | 0.707 | 0.279 | 9.3435E-100 |
| Contractile | Bcam | 3.48908E-88 | 1.411613 | 0.7 | 0.227 | 4.62059E-84 |
| Contractile | Rock1 | 2.42451E-84 | 1.410972 | 0.826 | 0.421 | 3.21078E-80 |
| Intermediate | Nupr1 | 5.0341E-119 | 1.405032 | 0.993 | 0.97 | 6.6667E-115 |
| Intermediate | Fbln2 | 3.3362E-130 | 1.401766 | 0.469 | 0.057 | 4.4181E-126 |
| Terminal | A230065H | 3.1283E-129 | 1.390486 | 0.471 | 0.062 | 4.1428E-125 |
| Contractile | Cpe | 2.5847E-112 | 1.390107 | 0.994 | 0.932 | 3.4229E-108 |
| Contractile | Cbr2 | 1.22195E-80 | 1.387144 | 0.729 | 0.283 | 1.61823E-76 |
| Rgs5_Contractile | Adcy5 | 6.8405E-31 | 1.386755 | 0.396 | 0.154 | 9.05888E-27 |
| Rgs5_Contractile | Ppp1r12b | 8.76936E-36 | 1.38197 | 0.441 | 0.172 | 1.16133E-31 |
| Rgs5_Contractile | Dusp3 | 8.02303E-26 | 1.37799 | 0.479 | 0.268 | 1.06249E-21 |
| Contractile | Lims2 | 3.46655E-89 | 1.377881 | 0.706 | 0.224 | 4.59075E-85 |
| Contractile | Ccdc3 | 6.73211E-98 | 1.374401 | 0.615 | 0.126 | 8.91533E-94 |
| Rgs5_Contractile | Ramp1 | 3.12967E-40 | 1.370066 | 0.649 | 0.339 | 4.14462E-36 |
| Contractile | 2210407C: | 7.27086E-66 | 1.366816 | 0.244 | 0.021 | 9.6288E-62 |
| Contractile | Ltbp1 | 2.39434E-74 | 1.364452 | 0.8 | 0.385 | 3.17083E-70 |
| Contractile | Plcb4 | 6.8521E-112 | 1.3625 | 0.591 | 0.102 | 9.0742E-108 |
| Contractile | Nexn | 2.55454E-92 | 1.361119 | 0.794 | 0.277 | 3.38297E-88 |
| Intermediate | Nbl1 | 5.0629E-120 | 1.355909 | 0.905 | 0.562 | 6.7048E-116 |

| | | | | | | |
|---|---|---|---|---|---|---|
| Contractile | Mcam | 6.62635E-84 | 1.354359 | 0.594 | 0.147 | 8.77528E-80 |
| Rgs5_Contractile | Synpo2 | 7.55114E-39 | 1.353881 | 0.497 | 0.193 | 9.99998E-35 |
| Rgs5_Contractile | Fblim1 | 2.14415E-37 | 1.351657 | 0.663 | 0.365 | 2.83949E-33 |
| Intermediate | Fstl1 | 2.4441E-129 | 1.349048 | 0.936 | 0.635 | 3.2367E-125 |
| Rgs5_Contractile | Tmsb4x | 7.1072E-61 | 1.348769 | 0.976 | 0.9 | 9.41207E-57 |
| Rgs5_Contractile | Actn1 | 1.41053E-46 | 1.347131 | 0.816 | 0.624 | 1.86796E-42 |
| Contractile | Pdlim7 | 1.81265E-77 | 1.345295 | 0.782 | 0.365 | 2.4005E-73 |
| Rgs5_Contractile | Gstm1 | 1.28827E-38 | 1.344708 | 0.764 | 0.592 | 1.70606E-34 |
| Contractile | Filip1l | 3.67005E-79 | 1.342144 | 0.738 | 0.278 | 4.86025E-75 |
| Terminal | Steap4 | 4.84244E-66 | 1.338649 | 0.455 | 0.147 | 6.41285E-62 |
| Rgs5_Contractile | Mustn1 | 2.38921E-28 | 1.331646 | 0.767 | 0.66 | 3.16403E-24 |
| Terminal | Clu | 1.81845E-63 | 1.324878 | 0.999 | 0.972 | 2.40817E-59 |
| Intermediate | Cfh | 2.82573E-82 | 1.321269 | 0.848 | 0.554 | 3.74212E-78 |
| Intermediate | Aqp1 | 1.1678E-106 | 1.318565 | 0.688 | 0.265 | 1.5465E-102 |
| Intermediate | Ifitm3 | 3.569E-165 | 1.316423 | 0.98 | 0.781 | 4.7265E-161 |
| Rgs5_Contractile | Rock1 | 1.31594E-19 | 1.313677 | 0.573 | 0.466 | 1.7427E-15 |
| Contractile | Smtn | 8.83137E-82 | 1.30608 | 0.853 | 0.461 | 1.16954E-77 |
| Contractile | Stbd1 | 2.73398E-89 | 1.302212 | 0.574 | 0.128 | 3.62061E-85 |
| Contractile | Mfap5 | 1.39827E-48 | 1.301684 | 0.594 | 0.229 | 1.85173E-44 |
| Rgs5_Contractile | Enah | 3.70061E-23 | 1.298643 | 0.493 | 0.298 | 4.90072E-19 |
| Rgs5_Contractile | Npnt | 1.07918E-26 | 1.294873 | 0.497 | 0.248 | 1.42916E-22 |
| Contractile | Pcp4l1 | 2.0479E-126 | 1.294677 | 0.532 | 0.06 | 2.712E-122 |
| Rgs5_Contractile | Pdlim3 | 1.50372E-37 | 1.292452 | 0.51 | 0.21 | 1.99138E-33 |
| Terminal | S100b | 6.1954E-113 | 1.291734 | 0.608 | 0.179 | 8.2046E-109 |
| Rgs5_Contractile | St5 | 3.36066E-36 | 1.2898 | 0.476 | 0.184 | 4.45053E-32 |
| Rgs5_Contractile | Cald1 | 1.71951E-57 | 1.287169 | 0.931 | 0.764 | 2.27715E-53 |
| Contractile | Pls3 | 3.13813E-87 | 1.28193 | 0.962 | 0.783 | 4.15582E-83 |
| Contractile | Lrrfip1 | 1.42043E-73 | 1.278969 | 0.821 | 0.469 | 1.88107E-69 |
| Intermediate | Serping1 | 7.37532E-70 | 1.277576 | 0.661 | 0.316 | 9.76714E-66 |
| Contractile | Lmcd1 | 1.855E-56 | 1.27753 | 0.821 | 0.517 | 2.45658E-52 |
| Terminal | Bglap | 7.24315E-39 | 1.275971 | 0.129 | 0.007 | 9.59211E-35 |
| Intermediate | Fbln5 | 3.39565E-96 | 1.273477 | 0.902 | 0.661 | 4.49686E-92 |
| Contractile | Npr1 | 2.471E-107 | 1.266015 | 0.524 | 0.077 | 3.2723E-103 |
| Contractile | Wfdc1 | 2.55944E-76 | 1.261253 | 0.735 | 0.311 | 3.38946E-72 |
| Intermediate | S100a4 | 2.9391E-107 | 1.256652 | 0.852 | 0.534 | 3.8922E-103 |
| Intermediate | Tmsb10 | 1.46613E-89 | 1.255271 | 0.885 | 0.73 | 1.94159E-85 |
| Contractile | Ppp1cb | 2.65498E-79 | 1.254938 | 0.882 | 0.534 | 3.51599E-75 |
| Rgs5_Contractile | Ppp1cb | 5.67376E-36 | 1.254645 | 0.715 | 0.565 | 7.51376E-32 |
| Rgs5_Contractile | Ppp1r12a | 9.43401E-17 | 1.254328 | 0.514 | 0.399 | 1.24935E-12 |
| Terminal | Dhx58os | 6.6116E-118 | 1.253151 | 0.438 | 0.055 | 8.7557E-114 |
| Contractile | Ndrg2 | 6.83906E-77 | 1.251546 | 0.762 | 0.323 | 9.05696E-73 |
| Contractile | Zyx | 4.70362E-73 | 1.250818 | 0.788 | 0.386 | 6.22901E-69 |
| Intermediate | Sdc1 | 2.6816E-101 | 1.250555 | 0.738 | 0.371 | 3.55127E-97 |
| Contractile | Pkp4 | 3.72274E-80 | 1.246895 | 0.576 | 0.148 | 4.93002E-76 |
| Rgs5_Contractile | Fmo2 | 1.46434E-29 | 1.243176 | 0.524 | 0.254 | 1.93922E-25 |
| Rgs5_Contractile | Ckb | 5.41806E-16 | 1.240059 | 0.538 | 0.416 | 7.17513E-12 |
| Rgs5_Contractile | Fhl1 | 5.54697E-43 | 1.239951 | 0.819 | 0.656 | 7.34586E-39 |
| Intermediate | Mgst1 | 1.3176E-100 | 1.233561 | 0.576 | 0.163 | 1.74487E-96 |
| Rgs5_Contractile | Tcap | 1.47844E-29 | 1.228634 | 0.378 | 0.133 | 1.95789E-25 |
| Contractile | Cav1 | 2.5509E-65 | 1.228301 | 0.706 | 0.293 | 3.37816E-61 |
| Intermediate | Mmp14 | 1.33929E-98 | 1.226666 | 0.688 | 0.296 | 1.77363E-94 |
| Contractile | Ccdc107 | 4.77331E-57 | 1.222796 | 0.706 | 0.378 | 6.3213E-53 |
| Contractile | Kcnmb1 | 6.1668E-103 | 1.219398 | 0.585 | 0.107 | 8.1668E-99 |
| Terminal | Frzb | 5.72043E-88 | 1.218395 | 0.939 | 0.614 | 7.57557E-84 |
| Rgs5_Contractile | Notch2 | 2.52711E-20 | 1.218258 | 0.378 | 0.193 | 3.34665E-16 |
| Contractile | Sh3bgrl | 2.80971E-74 | 1.210767 | 0.9 | 0.703 | 3.72089E-70 |
| Contractile | Optc | 1.3451E-127 | 1.202623 | 0.497 | 0.048 | 1.7814E-123 |
| Rgs5_Contractile | Vwc2 | 3.32797E-30 | 1.201269 | 0.33 | 0.104 | 4.40723E-26 |
| Intermediate | Sncg | 1.16291E-90 | 1.200849 | 0.782 | 0.412 | 1.54004E-86 |

| | | | | | | |
|---|---|---|---|---|---|---|
| Contractile | Ncam1 | 5.47516E-72 | 1.198649 | 0.806 | 0.384 | 7.25075E-68 |
| Contractile | Rasl12 | 5.54539E-81 | 1.194876 | 0.597 | 0.157 | 7.34375E-77 |
| Rgs5_Contractile | Smtn | 3.58935E-23 | 1.193609 | 0.622 | 0.502 | 4.75338E-19 |
| Contractile | Col18a1 | 6.77837E-69 | 1.190142 | 0.859 | 0.48 | 8.97659E-65 |
| Terminal | Tsc22d1 | 1.06143E-83 | 1.18638 | 0.953 | 0.773 | 1.40566E-79 |
| Contractile | Ntf3 | 8.854E-110 | 1.182877 | 0.509 | 0.066 | 1.1725E-105 |
| Terminal | Wfdc21 | 3.64498E-18 | 1.1813 | 0.111 | 0.027 | 4.82705E-14 |
| Rgs5_Contractile | Gucy1a3 | 9.57899E-26 | 1.18093 | 0.378 | 0.156 | 1.26855E-21 |
| Terminal | Cp | 7.87422E-98 | 1.177058 | 0.784 | 0.346 | 1.04278E-93 |
| Terminal | Serpine2 | 1.5005E-161 | 1.176271 | 0.998 | 0.957 | 1.9871E-157 |
| Intermediate | Cthrc1 | 4.26094E-27 | 1.175774 | 0.302 | 0.136 | 5.64276E-23 |
| Contractile | Slc22a1 | 8.71375E-84 | 1.174757 | 0.329 | 0.031 | 1.15396E-79 |
| Terminal | Cox4i2 | 1.53723E-96 | 1.162761 | 0.592 | 0.196 | 2.03575E-92 |
| Contractile | Rcan2 | 7.68243E-65 | 1.162472 | 0.656 | 0.249 | 1.01738E-60 |
| Rgs5_Contractile | Nexn | 2.2658E-25 | 1.161115 | 0.545 | 0.323 | 3.0006E-21 |
| Intermediate | Tpm4 | 2.3985E-108 | 1.160465 | 0.87 | 0.566 | 3.1763E-104 |
| Intermediate | Adamts2 | 2.6733E-110 | 1.158599 | 0.613 | 0.181 | 3.5403E-106 |
| Contractile | Pdlim1 | 1.09916E-56 | 1.157789 | 0.768 | 0.429 | 1.45562E-52 |
| Rgs5_Contractile | Dmd | 7.21782E-21 | 1.157046 | 0.389 | 0.193 | 9.55856E-17 |
| Rgs5_Contractile | Pcdh7 | 3.97241E-19 | 1.156923 | 0.615 | 0.54 | 5.26067E-15 |
| Rgs5_Contractile | Nox4 | 7.7402E-15 | 1.156058 | 0.441 | 0.299 | 1.02504E-10 |
| Intermediate | Mmp2 | 1.77165E-78 | 1.151416 | 0.827 | 0.534 | 2.3462E-74 |
| Rgs5_Contractile | Bcam | 1.67807E-14 | 1.149744 | 0.424 | 0.276 | 2.22226E-10 |
| Contractile | Tgfb1i1 | 2.35406E-59 | 1.149095 | 0.732 | 0.355 | 3.11749E-55 |
| Contractile | Dmd | 1.75473E-85 | 1.148394 | 0.615 | 0.151 | 2.32379E-81 |
| Intermediate | Ccl2 | 9.32603E-24 | 1.148267 | 0.135 | 0.029 | 1.23505E-19 |
| Contractile | Gstm2 | 1.52688E-66 | 1.146912 | 0.706 | 0.287 | 2.02204E-62 |
| Rgs5_Contractile | Pbxip1 | 4.28352E-27 | 1.145536 | 0.677 | 0.533 | 5.67267E-23 |
| Rgs5_Contractile | Spint2 | 5.85791E-30 | 1.14378 | 0.51 | 0.242 | 7.75763E-26 |
| Rgs5_Contractile | Lims2 | 1.06099E-18 | 1.141776 | 0.455 | 0.27 | 1.40507E-14 |
| Contractile | Itgb1 | 4.15057E-81 | 1.139851 | 0.968 | 0.749 | 5.49659E-77 |
| Rgs5_Contractile | Zyx | 1.11578E-19 | 1.136331 | 0.569 | 0.426 | 1.47763E-15 |
| Intermediate | Serpinf1 | 7.57503E-36 | 1.13517 | 0.329 | 0.133 | 1.00316E-31 |
| Rgs5_Contractile | Mir143hg | 1.86796E-25 | 1.134594 | 0.295 | 0.099 | 2.47374E-21 |
| Rgs5_Contractile | Mcam | 6.28475E-25 | 1.132457 | 0.413 | 0.182 | 8.32289E-21 |
| Rgs5_Contractile | Tmem47 | 1.42404E-18 | 1.130644 | 0.354 | 0.179 | 1.88585E-14 |
| Contractile | Dnajb4 | 3.75489E-54 | 1.130107 | 0.715 | 0.356 | 4.9726E-50 |
| Contractile | Osr1 | 3.85969E-37 | 1.127279 | 0.538 | 0.237 | 5.11139E-33 |
| Intermediate | Dpt | 3.07638E-61 | 1.127064 | 0.357 | 0.092 | 4.07405E-57 |
| Terminal | Nt5e | 1.3487E-107 | 1.124363 | 0.431 | 0.066 | 1.786E-103 |
| Contractile | Pcdh7 | 8.37941E-63 | 1.124125 | 0.829 | 0.503 | 1.10969E-58 |
| Contractile | Sparcl1 | 8.47514E-48 | 1.123988 | 0.479 | 0.154 | 1.12236E-43 |
| Rgs5_Contractile | Sh3bgr | 1.54288E-30 | 1.122976 | 0.465 | 0.184 | 2.04323E-26 |
| Terminal | Tnfrsf11b | 9.46161E-77 | 1.12279 | 0.828 | 0.447 | 1.253E-72 |
| Terminal | Fxyd5 | 6.0565E-123 | 1.121213 | 0.947 | 0.558 | 8.0207E-119 |
| Intermediate | Pcolce | 6.73795E-89 | 1.11966 | 0.859 | 0.591 | 8.92307E-85 |
| Contractile | Rrad | 6.06262E-41 | 1.115169 | 0.547 | 0.238 | 8.02872E-37 |
| Terminal | Trpv4 | 3.5218E-147 | 1.112631 | 0.501 | 0.056 | 4.6639E-143 |
| Contractile | Ppp1r12a | 3.42682E-64 | 1.112024 | 0.747 | 0.358 | 4.53814E-60 |
| Contractile | Ptrf | 4.55178E-62 | 1.111239 | 0.847 | 0.547 | 6.02793E-58 |
| Terminal | Wfdc12 | 3.08589E-29 | 1.110587 | 0.14 | 0.024 | 4.08664E-25 |
| Terminal | Id1 | 7.1078E-71 | 1.10972 | 0.733 | 0.376 | 9.41286E-67 |
| Rgs5_Contractile | Actg2 | 7.28796E-19 | 1.105766 | 0.396 | 0.189 | 9.65144E-15 |
| Terminal | Rgcc | 3.61E-115 | 1.105318 | 0.54 | 0.12 | 4.7807E-111 |
| Contractile | Atp2a2 | 2.28315E-50 | 1.104469 | 0.738 | 0.424 | 3.02358E-46 |
| Contractile | Fermt2 | 3.89012E-63 | 1.10384 | 0.841 | 0.527 | 5.15169E-59 |
| Rgs5_Contractile | E030013I1 | 3.87778E-28 | 1.101117 | 0.323 | 0.103 | 5.13534E-24 |
| Rgs5_Contractile | Col18a1 | 1.8584E-21 | 1.100956 | 0.667 | 0.515 | 2.46108E-17 |
| Contractile | Cfl2 | 7.82776E-49 | 1.098763 | 0.709 | 0.419 | 1.03663E-44 |

| | | | | | | |
|---|---|---|---|---|---|---|
| Intermediate | Timp2 | 4.2377E-141 | 1.097803 | 0.978 | 0.863 | 5.612E-137 |
| Contractile | Gucy1b3 | 4.65288E-82 | 1.096825 | 0.485 | 0.091 | 6.16181E-78 |
| Contractile | Ptgis | 3.86255E-64 | 1.095126 | 0.938 | 0.803 | 5.11517E-60 |
| Contractile | Kctd10 | 6.96309E-57 | 1.094316 | 0.729 | 0.37 | 9.22122E-53 |
| Contractile | Perp | 2.17425E-96 | 1.092866 | 0.482 | 0.069 | 2.87936E-92 |
| Contractile | Hcfc1r1 | 1.98907E-60 | 1.088415 | 0.879 | 0.711 | 2.63413E-56 |
| Rgs5_Contractile | Des | 2.4457E-12 | 1.087728 | 0.167 | 0.058 | 3.23885E-08 |
| Intermediate | S100a10 | 3.49188E-68 | 1.080562 | 0.867 | 0.659 | 4.6243E-64 |
| Contractile | Vwc2 | 2.91006E-81 | 1.078607 | 0.465 | 0.076 | 3.8538E-77 |
| Intermediate | Col1a1 | 4.5417E-65 | 1.077855 | 0.979 | 0.81 | 6.01457E-61 |
| Contractile | Aldh2 | 1.79543E-61 | 1.076598 | 0.874 | 0.629 | 2.37769E-57 |
| Rgs5_Contractile | Pdlim7 | 2.75033E-16 | 1.076024 | 0.531 | 0.41 | 3.64226E-12 |
| Contractile | Enah | 7.74713E-47 | 1.07259 | 0.609 | 0.274 | 1.02595E-42 |
| Contractile | Myh10 | 1.30769E-52 | 1.07177 | 0.579 | 0.22 | 1.73177E-48 |
| Intermediate | Lum | 5.16961E-95 | 1.071485 | 0.98 | 0.752 | 6.84612E-91 |
| Rgs5_Contractile | Pcp4l1 | 3.16783E-22 | 1.071269 | 0.292 | 0.104 | 4.19516E-18 |
| Contractile | Limch1 | 1.19871E-57 | 1.068195 | 0.55 | 0.175 | 1.58746E-53 |
| Rgs5_Contractile | Igfbp5 | 7.85653E-13 | 1.067728 | 0.226 | 0.094 | 1.04044E-08 |
| Contractile | Art3 | 7.63545E-51 | 1.067576 | 0.329 | 0.067 | 1.01116E-46 |
| Contractile | Rbp1 | 3.58959E-62 | 1.067507 | 0.932 | 0.621 | 4.7537E-58 |
| Contractile | Nox4 | 4.83479E-57 | 1.065142 | 0.65 | 0.261 | 6.40271E-53 |
| Terminal | Lbp | 1.22167E-96 | 1.06178 | 0.48 | 0.106 | 1.61786E-92 |
| Contractile | Itga9 | 1.0258E-84 | 1.061194 | 0.512 | 0.097 | 1.35847E-80 |
| Intermediate | Col14a1 | 4.46892E-67 | 1.060635 | 0.566 | 0.232 | 5.91819E-63 |
| Intermediate | Col8a1 | 7.49917E-85 | 1.059366 | 0.614 | 0.236 | 9.93116E-81 |
| Contractile | Mrvi1 | 2.56858E-73 | 1.056475 | 0.479 | 0.101 | 3.40157E-69 |
| Rgs5_Contractile | Dnajb4 | 8.9661E-13 | 1.052495 | 0.493 | 0.395 | 1.18738E-08 |
| Intermediate | Ctsl | 1.0945E-106 | 1.051816 | 0.932 | 0.71 | 1.4495E-102 |
| Rgs5_Contractile | Fbxl22 | 2.74025E-24 | 1.047395 | 0.396 | 0.16 | 3.62891E-20 |
| Rgs5_Contractile | Perp | 5.17278E-27 | 1.046766 | 0.312 | 0.103 | 6.85031E-23 |
| Rgs5_Contractile | Limch1 | 8.4854E-17 | 1.045572 | 0.378 | 0.207 | 1.12372E-12 |
| Rgs5_Contractile | Dst | 6.67259E-17 | 1.045465 | 0.597 | 0.549 | 8.83651E-13 |
| Rgs5_Contractile | Dmpk | 3.86563E-19 | 1.043616 | 0.41 | 0.214 | 5.11926E-15 |
| Contractile | Oxct1 | 5.45036E-52 | 1.040563 | 0.797 | 0.532 | 7.21791E-48 |
| Terminal | Cdkn1a | 3.39888E-56 | 1.039922 | 0.713 | 0.394 | 4.50113E-52 |
| Intermediate | Fbn1 | 2.25673E-67 | 1.038048 | 0.66 | 0.35 | 2.98859E-63 |
| Rgs5_Contractile | Thsd4 | 9.3203E-17 | 1.033773 | 0.285 | 0.126 | 1.23429E-12 |
| Terminal | Bglap2 | 2.57042E-41 | 1.033413 | 0.14 | 0.009 | 3.404E-37 |
| Contractile | Aoc3 | 1.54644E-75 | 1.030967 | 0.479 | 0.095 | 2.04795E-71 |
| Contractile | Adcy5 | 2.93124E-67 | 1.028859 | 0.518 | 0.128 | 3.88185E-63 |
| Contractile | Hspb7 | 3.15458E-52 | 1.027727 | 0.6 | 0.224 | 4.17762E-48 |
| Intermediate | Ecm1 | 3.18681E-89 | 1.02303 | 0.935 | 0.702 | 4.22029E-85 |
| Intermediate | Anxa2 | 6.51549E-92 | 1.019475 | 0.924 | 0.771 | 8.62846E-88 |
| Intermediate | Phlda3 | 1.3071E-79 | 1.018497 | 0.725 | 0.372 | 1.73099E-75 |
| Contractile | Dsp | 6.30798E-96 | 1.018167 | 0.397 | 0.043 | 8.35366E-92 |
| Rgs5_Contractile | Kcnmb1 | 2.07885E-17 | 1.017933 | 0.326 | 0.154 | 2.75303E-13 |
| Contractile | Vcl | 4.77473E-54 | 1.017842 | 0.665 | 0.295 | 6.32318E-50 |
| Terminal | Col10a1 | 1.00573E-24 | 1.016269 | 0.134 | 0.026 | 1.33188E-20 |
| Contractile | Celf2 | 1.95161E-53 | 1.014606 | 0.568 | 0.205 | 2.58452E-49 |
| Intermediate | Fndc1 | 2.51373E-70 | 1.013518 | 0.334 | 0.064 | 3.32893E-66 |
| Terminal | Prg4 | 4.7746E-103 | 1.013204 | 0.808 | 0.379 | 6.323E-99 |
| Contractile | Rassf3 | 1.62586E-57 | 1.012305 | 0.474 | 0.135 | 2.15312E-53 |
| Terminal | Ucma | 1.30596E-81 | 1.008553 | 0.245 | 0.008 | 1.72949E-77 |
| Rgs5_Contractile | Itga9 | 2.18888E-16 | 1.007206 | 0.299 | 0.136 | 2.89873E-12 |
| Rgs5_Contractile | Rrad | 0.000176002 | 1.002903 | 0.33 | 0.275 | 1 |
| Contractile | Des | 2.91078E-18 | 1.002065 | 0.182 | 0.053 | 3.85475E-14 |
| Contractile | Ilk | 4.59927E-44 | 1.001448 | 0.741 | 0.49 | 6.09081E-40 |
| Contractile | Hspb2 | 5.59907E-44 | 1.000194 | 0.688 | 0.398 | 7.41485E-40 |
| Contractile | Ccdc42 | 5.57321E-57 | 0.998418 | 0.294 | 0.044 | 7.3806E-53 |

| | | | | | | |
|---|---|---|---|---|---|---|
| Rgs5_Contractile | Rasl12 | 8.12144E-19 | 0.995137 | 0.382 | 0.197 | 1.07552E-14 |
| Rgs5_Contractile | 1190005I0 | 1.64892E-16 | 0.994023 | 0.358 | 0.184 | 2.18366E-12 |
| Intermediate | Ndufa4l2 | 9.99144E-09 | 0.990874 | 0.229 | 0.146 | 0.000132317 |
| Rgs5_Contractile | Gm13861 | 3.69192E-16 | 0.990873 | 0.17 | 0.05 | 4.88921E-12 |
| Terminal | Xylt1 | 3.5906E-118 | 0.990712 | 0.475 | 0.074 | 4.755E-114 |
| Terminal | Lgals3 | 2.0252E-105 | 0.987738 | 0.947 | 0.543 | 2.6819E-101 |
| Contractile | Asb2 | 1.81582E-94 | 0.986754 | 0.35 | 0.03 | 2.40469E-90 |
| Rgs5_Contractile | Ndrg2 | 6.19057E-17 | 0.984777 | 0.514 | 0.367 | 8.19817E-13 |
| Rgs5_Contractile | Nr2f2 | 3.09538E-10 | 0.984446 | 0.274 | 0.157 | 4.09921E-06 |
| Rgs5_Contractile | Wfdc1 | 5.8474E-12 | 0.979756 | 0.465 | 0.358 | 7.74372E-08 |
| Contractile | Net1 | 2.87014E-46 | 0.977521 | 0.444 | 0.139 | 3.80092E-42 |
| Contractile | Pfn1 | 1.32138E-69 | 0.976039 | 0.968 | 0.912 | 1.74991E-65 |
| Rgs5_Contractile | Itgb1 | 1.43073E-27 | 0.972169 | 0.809 | 0.776 | 1.89472E-23 |
| Contractile | Ppp1r12b | 1.99411E-48 | 0.972053 | 0.491 | 0.157 | 2.64079E-44 |
| Rgs5_Contractile | Vim | 2.45652E-42 | 0.970106 | 0.986 | 0.964 | 3.25317E-38 |
| Terminal | Serpina3n | 2.30072E-61 | 0.969258 | 0.678 | 0.329 | 3.04684E-57 |
| Intermediate | Igfbp2 | 1.0553E-14 | 0.969055 | 0.105 | 0.03 | 1.39754E-10 |
| Contractile | Tspan2 | 7.13399E-47 | 0.967089 | 0.518 | 0.194 | 9.44754E-43 |
| Contractile | Scube3 | 1.03976E-77 | 0.966008 | 0.485 | 0.097 | 1.37695E-73 |
| Contractile | Sorl1 | 1.82914E-58 | 0.963462 | 0.444 | 0.111 | 2.42232E-54 |
| Contractile | Mylk4 | 3.25239E-82 | 0.962289 | 0.344 | 0.036 | 4.30715E-78 |
| Rgs5_Contractile | Cav1 | 1.12654E-07 | 0.961932 | 0.417 | 0.343 | 0.001491871 |
| Rgs5_Contractile | Stbd1 | 1.99342E-15 | 0.960216 | 0.337 | 0.171 | 2.63989E-11 |
| Rgs5_Contractile | Ltbp1 | 4.69145E-13 | 0.959806 | 0.542 | 0.43 | 6.21289E-09 |
| Contractile | Tinagl1 | 7.45963E-59 | 0.956318 | 0.606 | 0.197 | 9.87879E-55 |
| Rgs5_Contractile | Sh3bgrl | 1.24127E-23 | 0.956057 | 0.75 | 0.728 | 1.64381E-19 |
| Rgs5_Contractile | Nov | 7.14399E-18 | 0.956037 | 0.524 | 0.318 | 9.46079E-14 |
| Contractile | Gm13889 | 5.86886E-39 | 0.955303 | 0.685 | 0.369 | 7.77214E-35 |
| Rgs5_Contractile | Lrrc58 | 4.61488E-07 | 0.954688 | 0.542 | 0.618 | 0.006111486 |
| Terminal | Eps8l2 | 1.8334E-122 | 0.953149 | 0.464 | 0.059 | 2.4279E-118 |
| Contractile | Kcnab1 | 1.78809E-78 | 0.95179 | 0.447 | 0.076 | 2.36797E-74 |
| Rgs5_Contractile | Efhd1 | 2.46382E-18 | 0.951312 | 0.368 | 0.174 | 3.26283E-14 |
| Rgs5_Contractile | Myom1 | 4.74665E-19 | 0.951021 | 0.267 | 0.102 | 6.28599E-15 |
| Contractile | Gucy1a3 | 8.98179E-61 | 0.948529 | 0.503 | 0.13 | 1.18946E-56 |
| Intermediate | Col6a3 | 6.347E-50 | 0.947342 | 0.755 | 0.524 | 8.40534E-46 |
| Contractile | Map1lc3a | 4.74662E-62 | 0.94622 | 0.909 | 0.722 | 6.28595E-58 |
| Intermediate | Mgp | 8.22978E-73 | 0.944905 | 1 | 1 | 1.08987E-68 |
| Intermediate | Cxcl1 | 2.44586E-19 | 0.943702 | 0.3 | 0.151 | 3.23905E-15 |
| Contractile | Tmem110 | 4.48105E-43 | 0.941504 | 0.521 | 0.208 | 5.93426E-39 |
| Rgs5_Contractile | Cpe | 3.18582E-38 | 0.939982 | 0.951 | 0.939 | 4.21898E-34 |
| Rgs5_Contractile | Plcb4 | 8.04018E-15 | 0.938282 | 0.312 | 0.152 | 1.06476E-10 |
| Rgs5_Contractile | Pdlim1 | 1.08193E-14 | 0.936549 | 0.569 | 0.464 | 1.4328E-10 |
| Rgs5_Contractile | Npr1 | 6.52266E-16 | 0.93369 | 0.281 | 0.121 | 8.63796E-12 |
| Contractile | Fam129a | 1.28936E-48 | 0.933299 | 0.579 | 0.236 | 1.7075E-44 |
| Contractile | Ttc28 | 4.49997E-43 | 0.930019 | 0.506 | 0.194 | 5.95931E-39 |
| Rgs5_Contractile | Msrb1 | 7.84248E-09 | 0.925974 | 0.476 | 0.403 | 0.000103858 |
| Intermediate | Anxa1 | 2.09356E-66 | 0.925621 | 0.933 | 0.794 | 2.7725E-62 |
| Rgs5_Contractile | Fermt2 | 1.27911E-13 | 0.923512 | 0.601 | 0.567 | 1.69392E-09 |
| Contractile | Tppp3 | 1.0604E-44 | 0.921928 | 0.485 | 0.158 | 1.40429E-40 |
| Terminal | Fibin | 8.62439E-51 | 0.921846 | 0.67 | 0.346 | 1.14213E-46 |
| Contractile | Jph2 | 3.8454E-51 | 0.919637 | 0.471 | 0.14 | 5.09246E-47 |
| Terminal | Fth1 | 1.86539E-35 | 0.916706 | 1 | 0.999 | 2.47033E-31 |
| Rgs5_Contractile | Net1 | 3.72862E-11 | 0.915663 | 0.295 | 0.166 | 4.93782E-07 |
| Terminal | Sdc4 | 1.19696E-52 | 0.913012 | 0.883 | 0.666 | 1.58514E-48 |
| Contractile | Tes | 3.81751E-65 | 0.912581 | 0.391 | 0.072 | 5.05553E-61 |
| Intermediate | Gas6 | 3.91669E-33 | 0.911823 | 0.839 | 0.694 | 5.18688E-29 |
| Contractile | Foxp2 | 2.26748E-65 | 0.911751 | 0.359 | 0.058 | 3.00282E-61 |
| Contractile | Rsu1 | 5.14547E-43 | 0.911596 | 0.794 | 0.562 | 6.81414E-39 |
| Rgs5_Contractile | Rgs4 | 2.71254E-10 | 0.910472 | 0.233 | 0.116 | 3.59222E-06 |

| | | | | | | |
|---|---|---|---|---|---|---|
| Rgs5_Contractile | Hcfc1r1 | 1.91048E-19 | 0.908293 | 0.726 | 0.736 | 2.53005E-15 |
| Rgs5_Contractile | Oxct1 | 1.20458E-09 | 0.905732 | 0.559 | 0.571 | 1.59522E-05 |
| Contractile | Hacd1 | 3.85668E-31 | 0.905135 | 0.609 | 0.374 | 5.1074E-27 |
| Contractile | Anxa11 | 1.72415E-32 | 0.904995 | 0.544 | 0.284 | 2.28329E-28 |
| Rgs5_Contractile | Aoc3 | 3.23256E-13 | 0.903744 | 0.274 | 0.132 | 4.28088E-09 |
| Intermediate | Itgbl1 | 1.83883E-63 | 0.903604 | 0.597 | 0.252 | 2.43516E-59 |
| Terminal | Ivns1abp | 4.4913E-99 | 0.902974 | 0.493 | 0.11 | 5.9478E-95 |
| Terminal | Mfi2 | 1.933E-109 | 0.902172 | 0.339 | 0.018 | 2.5599E-105 |
| Contractile | Atp1a2 | 1.03153E-43 | 0.901223 | 0.565 | 0.228 | 1.36605E-39 |
| Rgs5_Contractile | Rbpms2 | 1.22069E-11 | 0.900624 | 0.288 | 0.159 | 1.61657E-07 |
| Rgs5_Contractile | Fam129a | 6.00856E-06 | 0.899958 | 0.34 | 0.277 | 0.079571356 |
| Rgs5_Contractile | Mrvi1 | 2.32937E-09 | 0.899946 | 0.253 | 0.141 | 3.08478E-05 |
| Terminal | Znhit6 | 1.45105E-78 | 0.898136 | 0.51 | 0.16 | 1.92163E-74 |
| Contractile | Smarcd3 | 4.32756E-40 | 0.897077 | 0.55 | 0.243 | 5.73099E-36 |
| Rgs5_Contractile | Atp2a2 | 2.82493E-09 | 0.894041 | 0.5 | 0.464 | 3.74106E-05 |
| Rgs5_Contractile | Timp4 | 3.41727E-13 | 0.891054 | 0.16 | 0.052 | 4.52549E-09 |
| Rgs5_Contractile | Actn4 | 3.39352E-10 | 0.890409 | 0.594 | 0.627 | 4.49404E-06 |
| Rgs5_Contractile | Abcc9 | 3.90857E-10 | 0.890278 | 0.326 | 0.206 | 5.17611E-06 |
| Intermediate | Lgals1 | 1.61533E-92 | 0.889898 | 0.996 | 0.92 | 2.13918E-88 |
| Rgs5_Contractile | Mbnl1 | 1.5096E-14 | 0.885552 | 0.684 | 0.742 | 1.99917E-10 |
| Terminal | Pdgfrl | 1.59008E-72 | 0.885127 | 0.583 | 0.225 | 2.10574E-68 |
| Contractile | Nkd1 | 1.24899E-52 | 0.884315 | 0.403 | 0.099 | 1.65404E-48 |
| Rgs5_Contractile | Aes | 1.26143E-12 | 0.883915 | 0.556 | 0.51 | 1.67051E-08 |
| Contractile | Cav2 | 1.11012E-29 | 0.883846 | 0.556 | 0.307 | 1.47013E-25 |
| Rgs5_Contractile | Npy1r | 1.88432E-18 | 0.882748 | 0.201 | 0.061 | 2.4954E-14 |
| Intermediate | Ahnak2 | 1.22571E-68 | 0.882744 | 0.696 | 0.356 | 1.6232E-64 |
| Contractile | Ift43 | 6.6231E-37 | 0.881891 | 0.638 | 0.371 | 8.77097E-33 |
| Contractile | Bnip2 | 7.66904E-35 | 0.881202 | 0.759 | 0.557 | 1.01561E-30 |
| Contractile | Cnn2 | 1.33535E-43 | 0.88103 | 0.759 | 0.401 | 1.76841E-39 |
| Rgs5_Contractile | Adamts1 | 0.002680633 | 0.880021 | 0.274 | 0.241 | 1 |
| Contractile | Thsd4 | 1.31487E-56 | 0.878164 | 0.424 | 0.099 | 1.74129E-52 |
| Terminal | Pdzrn4 | 1.42792E-97 | 0.877953 | 0.358 | 0.036 | 1.89099E-93 |
| Rgs5_Contractile | Ccdc107 | 6.96347E-08 | 0.876957 | 0.462 | 0.419 | 0.000922173 |
| Rgs5_Contractile | Svil | 2.96804E-07 | 0.875957 | 0.274 | 0.184 | 0.003930573 |
| Rgs5_Contractile | Txnip | 1.21089E-05 | 0.872961 | 0.354 | 0.31 | 0.160358425 |
| Contractile | Mustn1 | 8.72709E-33 | 0.872122 | 0.868 | 0.64 | 1.15573E-28 |
| Contractile | Lgr6 | 9.83663E-67 | 0.870648 | 0.356 | 0.056 | 1.30266E-62 |
| Intermediate | Lpl | 1.49757E-27 | 0.870058 | 0.387 | 0.194 | 1.98323E-23 |
| Contractile | Rgs4 | 1.29825E-52 | 0.869573 | 0.388 | 0.088 | 1.71927E-48 |
| Rgs5_Contractile | Cnn2 | 8.0018E-17 | 0.869483 | 0.576 | 0.434 | 1.05968E-12 |
| Rgs5_Contractile | Rarres2 | 3.2545E-21 | 0.868836 | 0.708 | 0.561 | 4.30994E-17 |
| Terminal | C3 | 2.47424E-60 | 0.86843 | 0.798 | 0.449 | 3.27663E-56 |
| Intermediate | Crip1 | 1.40451E-58 | 0.86634 | 0.98 | 0.918 | 1.85999E-54 |
| Rgs5_Contractile | Ptgis | 1.65523E-21 | 0.865611 | 0.806 | 0.824 | 2.19202E-17 |
| Rgs5_Contractile | Optc | 4.65266E-13 | 0.864881 | 0.226 | 0.096 | 6.16152E-09 |
| Rgs5_Contractile | Lrrfip1 | 8.51947E-10 | 0.864183 | 0.552 | 0.514 | 1.12823E-05 |
| Rgs5_Contractile | Esr1 | 7.50087E-14 | 0.863865 | 0.208 | 0.081 | 9.9334E-10 |
| Contractile | Il11ra1 | 9.0054E-40 | 0.863858 | 0.815 | 0.625 | 1.19259E-35 |
| Terminal | Tubb2a | 2.56107E-54 | 0.86321 | 0.68 | 0.365 | 3.39163E-50 |
| Rgs5_Contractile | Hspb7 | 5.85297E-10 | 0.859902 | 0.382 | 0.263 | 7.75109E-06 |
| Contractile | Rgs5 | 2.34796E-50 | 0.858367 | 0.618 | 0.222 | 3.1094E-46 |
| Rgs5_Contractile | Tgfb1i1 | 5.61008E-09 | 0.857518 | 0.462 | 0.401 | 7.42943E-05 |
| Rgs5_Contractile | Pls3 | 2.33971E-27 | 0.857434 | 0.826 | 0.806 | 3.09848E-23 |
| Terminal | Bhlhe40 | 1.66519E-57 | 0.857143 | 0.574 | 0.26 | 2.20521E-53 |
| Terminal | Trf | 1.37117E-23 | 0.856935 | 0.301 | 0.138 | 1.81584E-19 |
| Contractile | Pdgfd | 1.1201E-55 | 0.854524 | 0.426 | 0.103 | 1.48334E-51 |
| Terminal | Pltp | 6.60906E-44 | 0.852982 | 0.509 | 0.231 | 8.75237E-40 |
| Rgs5_Contractile | Hsph1 | 1.2555E-05 | 0.852143 | 0.26 | 0.189 | 0.166265422 |
| Rgs5_Contractile | Tln1 | 1.5286E-07 | 0.851885 | 0.497 | 0.511 | 0.002024326 |

| | | | | | | |
|---|---|---|---|---|---|---|
| Rgs5_Contractile | Myh10 | 5.50165E-08 | 0.851792 | 0.351 | 0.26 | 0.000728583 |
| Terminal | Smoc2 | 5.67538E-50 | 0.849612 | 0.522 | 0.223 | 7.51591E-46 |
| Contractile | Fbxo32 | 2.01055E-45 | 0.849066 | 0.353 | 0.088 | 2.66257E-41 |
| Rgs5_Contractile | Pde3a | 7.81595E-06 | 0.848802 | 0.417 | 0.391 | 0.103506669 |
| Contractile | Svil | 7.7347E-41 | 0.848286 | 0.45 | 0.153 | 1.02431E-36 |
| Rgs5_Contractile | Gstm2 | 1.39174E-10 | 0.845465 | 0.438 | 0.333 | 1.84308E-06 |
| Rgs5_Contractile | Smarcd3 | 9.13198E-07 | 0.842307 | 0.351 | 0.278 | 0.012093476 |
| Rgs5_Contractile | Scube3 | 1.34901E-07 | 0.842192 | 0.236 | 0.14 | 0.00178649 |
| Rgs5_Contractile | Sun2 | 1.19598E-05 | 0.841097 | 0.24 | 0.165 | 0.158383648 |
| Terminal | Fosb | 1.68956E-46 | 0.840964 | 0.519 | 0.237 | 2.23749E-42 |
| Rgs5_Contractile | Abhd2 | 0.00077678 | 0.839403 | 0.274 | 0.23 | 1 |
| Intermediate | Ctsk | 1.52058E-25 | 0.837925 | 0.529 | 0.34 | 2.0137E-21 |
| Intermediate | Dkk3 | 7.71986E-59 | 0.836484 | 0.751 | 0.439 | 1.02234E-54 |
| Rgs5_Contractile | Tppp3 | 2.23716E-14 | 0.836155 | 0.351 | 0.184 | 2.96267E-10 |
| Rgs5_Contractile | Inpp4a | 1.74337E-06 | 0.833653 | 0.219 | 0.137 | 0.023087442 |
| Rgs5_Contractile | Tacc1 | 1.31288E-07 | 0.833411 | 0.344 | 0.265 | 0.001738653 |
| Rgs5_Contractile | Thra | 2.51291E-05 | 0.832212 | 0.448 | 0.47 | 0.332784249 |
| Contractile | Emc8 | 2.35284E-35 | 0.831417 | 0.559 | 0.282 | 3.11586E-31 |
| Intermediate | Slco2a1 | 1.91385E-67 | 0.830885 | 0.342 | 0.07 | 2.53451E-63 |
| Contractile | Dusp3 | 3.47783E-35 | 0.830607 | 0.547 | 0.252 | 4.60569E-31 |
| Contractile | Chd3 | 4.45225E-31 | 0.830549 | 0.488 | 0.231 | 5.89612E-27 |
| Contractile | 2310022B( | 1.50473E-31 | 0.82935 | 0.568 | 0.306 | 1.99271E-27 |
| Contractile | Oat | 2.956E-35 | 0.828903 | 0.859 | 0.742 | 3.91463E-31 |
| Rgs5_Contractile | Dusp1 | 6.4419E-08 | 0.827368 | 0.597 | 0.589 | 0.000853101 |
| Intermediate | Aebp1 | 7.19742E-84 | 0.822954 | 0.975 | 0.867 | 9.53155E-80 |
| Contractile | Tmem47 | 4.76919E-47 | 0.822518 | 0.482 | 0.153 | 6.31583E-43 |
| Intermediate | Igfbp4 | 4.37478E-44 | 0.822005 | 0.544 | 0.256 | 5.79352E-40 |
| Rgs5_Contractile | Ptms | 4.13502E-23 | 0.821954 | 0.799 | 0.81 | 5.476E-19 |
| Intermediate | Col5a1 | 1.11191E-42 | 0.8217 | 0.6 | 0.356 | 1.47251E-38 |
| Contractile | Bri3 | 6.89752E-45 | 0.8215 | 0.882 | 0.767 | 9.13438E-41 |
| Contractile | Esr1 | 7.93627E-56 | 0.820668 | 0.329 | 0.057 | 1.051E-51 |
| Contractile | Sod3 | 3.12646E-39 | 0.819615 | 0.953 | 0.806 | 4.14036E-35 |
| Contractile | Abcc9 | 6.4544E-40 | 0.819248 | 0.488 | 0.176 | 8.54756E-36 |
| Intermediate | Gda | 2.65233E-72 | 0.818274 | 0.381 | 0.089 | 3.51249E-68 |
| Intermediate | Htra1 | 2.12434E-78 | 0.817586 | 0.936 | 0.789 | 2.81327E-74 |
| Rgs5_Contractile | Kctd10 | 9.525E-07 | 0.817457 | 0.448 | 0.417 | 0.012613963 |
| Contractile | Nudt4 | 1.27854E-35 | 0.816543 | 0.829 | 0.656 | 1.69317E-31 |
| Terminal | Col9a1 | 7.1455E-80 | 0.815547 | 0.249 | 0.011 | 9.46279E-76 |
| Rgs5_Contractile | Jph2 | 1.44923E-08 | 0.81482 | 0.281 | 0.174 | 0.000191921 |
| Rgs5_Contractile | Sema5a | 2.00888E-08 | 0.814212 | 0.194 | 0.099 | 0.000266036 |
| Intermediate | Rnase4 | 5.89155E-65 | 0.813827 | 0.612 | 0.281 | 7.80218E-61 |
| Rgs5_Contractile | Cap1 | 2.58846E-07 | 0.813588 | 0.26 | 0.169 | 0.003427901 |
| Contractile | Postn | 8.69767E-34 | 0.813456 | 0.794 | 0.508 | 1.15183E-29 |
| Contractile | Pde5a | 6.14848E-33 | 0.813037 | 0.435 | 0.174 | 8.14243E-29 |
| Intermediate | Cx3cl1 | 2.90524E-57 | 0.812541 | 0.435 | 0.155 | 3.8474E-53 |
| Terminal | Dab2 | 1.01339E-80 | 0.812315 | 0.493 | 0.138 | 1.34204E-76 |
| Rgs5_Contractile | Samd4 | 2.29336E-05 | 0.811813 | 0.361 | 0.32 | 0.303709165 |
| Rgs5_Contractile | Ldb3 | 2.5027E-13 | 0.808537 | 0.191 | 0.072 | 3.31433E-09 |
| Contractile | Kif5b | 5.553E-30 | 0.807563 | 0.753 | 0.569 | 7.35384E-26 |
| Intermediate | Thbs2 | 4.44071E-34 | 0.805968 | 0.385 | 0.181 | 5.88083E-30 |
| Rgs5_Contractile | Mfap4 | 6.23597E-11 | 0.805667 | 0.382 | 0.226 | 8.25829E-07 |
| Intermediate | C1s1 | 6.88787E-47 | 0.80478 | 0.546 | 0.277 | 9.1216E-43 |
| Rgs5_Contractile | Map1lc3a | 9.6148E-13 | 0.803923 | 0.684 | 0.757 | 1.27329E-08 |
| Rgs5_Contractile | Pkp4 | 2.93029E-09 | 0.803771 | 0.309 | 0.194 | 3.88059E-05 |
| Contractile | Nptn | 3.18807E-30 | 0.803083 | 0.753 | 0.587 | 4.22196E-26 |
| Rgs5_Contractile | Aldh2 | 2.68514E-12 | 0.803002 | 0.646 | 0.666 | 3.55594E-08 |
| Contractile | Suclg2 | 1.44904E-31 | 0.802782 | 0.385 | 0.147 | 1.91897E-27 |
| Rgs5_Contractile | Tes | 3.88628E-06 | 0.802069 | 0.188 | 0.108 | 0.05146604 |
| Rgs5_Contractile | Plekho1 | 8.1445E-12 | 0.801368 | 0.205 | 0.088 | 1.07858E-07 |

| | | | | | | |
|---|---|---|---|---|---|---|
| Rgs5_Contractile | Fbln7 | 0.007538784 | 0.800894 | 0.16 | 0.118 | 1 |
| Rgs5_Contractile | Vcl | 5.56918E-06 | 0.800134 | 0.392 | 0.341 | 0.073752652 |
| Contractile | Npy1r | 4.86269E-51 | 0.799975 | 0.282 | 0.044 | 6.43966E-47 |
| Rgs5_Contractile | Rassf3 | 4.8115E-07 | 0.799951 | 0.264 | 0.172 | 0.006371874 |
| Intermediate | Sdc2 | 1.77708E-44 | 0.799785 | 0.756 | 0.524 | 2.35338E-40 |
| Terminal | Ier2 | 2.71451E-36 | 0.796696 | 0.721 | 0.493 | 3.59483E-32 |
| Rgs5_Contractile | Lmcd1 | 5.64718E-09 | 0.795055 | 0.58 | 0.557 | 7.47856E-05 |
| Contractile | Fbxo30 | 9.28837E-45 | 0.794404 | 0.418 | 0.125 | 1.23006E-40 |
| Rgs5_Contractile | Kcnab1 | 9.66188E-13 | 0.792345 | 0.247 | 0.113 | 1.27952E-08 |
| Intermediate | Plxdc2 | 1.80952E-59 | 0.79207 | 0.681 | 0.358 | 2.39634E-55 |
| Rgs5_Contractile | Mylk4 | 8.99231E-13 | 0.791187 | 0.181 | 0.066 | 1.19085E-08 |
| Rgs5_Contractile | Arid5b | 4.36895E-05 | 0.790749 | 0.431 | 0.417 | 0.578580538 |
| Contractile | Rasgrp2 | 1.39312E-53 | 0.790685 | 0.335 | 0.063 | 1.84491E-49 |
| Rgs5_Contractile | Cfl2 | 9.56372E-07 | 0.789168 | 0.472 | 0.458 | 0.012665236 |
| Rgs5_Contractile | Ccni | 0.000755178 | 0.789158 | 0.389 | 0.406 | 1 |
| Rgs5_Contractile | Ntrk3 | 3.65243E-06 | 0.789142 | 0.274 | 0.196 | 0.048369097 |
| Intermediate | S100a11 | 3.24444E-80 | 0.788649 | 0.973 | 0.925 | 4.29661E-76 |
| Intermediate | Emp3 | 6.85822E-81 | 0.788295 | 0.959 | 0.847 | 9.08234E-77 |
| Contractile | Rasl11b | 4.11825E-31 | 0.787994 | 0.429 | 0.167 | 5.45379E-27 |
| Contractile | Anxa6 | 7.57068E-32 | 0.787468 | 0.679 | 0.445 | 1.00259E-27 |
| Intermediate | Ftl1 | 3.27886E-75 | 0.786449 | 0.993 | 0.95 | 4.34219E-71 |
| Terminal | Ier3 | 3.97359E-45 | 0.785215 | 0.628 | 0.342 | 5.26223E-41 |
| Intermediate | Npc2 | 5.04833E-63 | 0.785003 | 0.889 | 0.72 | 6.6855E-59 |
| Rgs5_Contractile | Inpp5a | 3.28646E-06 | 0.784413 | 0.25 | 0.169 | 0.043522549 |
| Intermediate | Loxl1 | 1.4936E-51 | 0.783941 | 0.623 | 0.311 | 1.97798E-47 |
| Contractile | Cryab | 4.48933E-38 | 0.782102 | 0.882 | 0.744 | 5.94522E-34 |
| Terminal | Scara3 | 3.09618E-54 | 0.780985 | 0.815 | 0.515 | 4.10028E-50 |
| Rgs5_Contractile | Anxa11 | 3.67824E-05 | 0.780657 | 0.358 | 0.316 | 0.487109499 |
| Rgs5_Contractile | Rcan2 | 5.13008E-07 | 0.779068 | 0.375 | 0.297 | 0.006793761 |
| Intermediate | Tuba1a | 2.7966E-47 | 0.777482 | 0.773 | 0.543 | 3.70354E-43 |
| Contractile | Tagln2 | 1.5482E-37 | 0.777179 | 0.9 | 0.726 | 2.05028E-33 |
| Intermediate | Aspn | 5.60923E-21 | 0.777119 | 0.418 | 0.239 | 7.42831E-17 |
| Rgs5_Contractile | Gabra3 | 2.5903E-10 | 0.774711 | 0.174 | 0.072 | 3.43033E-06 |
| Rgs5_Contractile | Celf2 | 0.000144548 | 0.773893 | 0.302 | 0.25 | 1 |
| Terminal | Junb | 6.5552E-39 | 0.773748 | 0.829 | 0.634 | 8.68105E-35 |
| Contractile | Cnn3 | 3.16253E-34 | 0.773422 | 0.838 | 0.682 | 4.18814E-30 |
| Contractile | Tgm2 | 9.79199E-28 | 0.771714 | 0.624 | 0.386 | 1.29675E-23 |
| Rgs5_Contractile | Klhdc8b | 2.42096E-05 | 0.770346 | 0.177 | 0.108 | 0.320608223 |
| Contractile | Serinc3 | 7.34683E-36 | 0.770264 | 0.832 | 0.682 | 9.72941E-32 |
| Contractile | Ltbp4 | 6.6914E-33 | 0.769306 | 0.694 | 0.404 | 8.86141E-29 |
| Terminal | Srpx2 | 3.15526E-65 | 0.768385 | 0.602 | 0.247 | 4.17851E-61 |
| Rgs5_Contractile | Anxa6 | 0.005882619 | 0.768284 | 0.427 | 0.485 | 1 |
| Rgs5_Contractile | Ltbp4 | 6.5094E-08 | 0.768101 | 0.49 | 0.439 | 0.00086204 |
| Terminal | Klf4 | 3.19012E-46 | 0.767913 | 0.674 | 0.38 | 4.22468E-42 |
| Intermediate | Dclk1 | 3.79085E-56 | 0.767909 | 0.371 | 0.107 | 5.02022E-52 |
| Terminal | Fn1 | 5.0612E-67 | 0.766847 | 0.998 | 0.917 | 6.70254E-63 |
| Contractile | Fmo3 | 2.73545E-52 | 0.765969 | 0.203 | 0.019 | 3.62255E-48 |
| Rgs5_Contractile | Fry | 4.21826E-05 | 0.765656 | 0.309 | 0.252 | 0.55862436 |
| Contractile | Samd4 | 2.47834E-28 | 0.764973 | 0.535 | 0.29 | 3.28206E-24 |
| Rgs5_Contractile | Alkbh5 | 0.005851704 | 0.764362 | 0.295 | 0.284 | 1 |
| Rgs5_Contractile | Cryab | 2.88598E-17 | 0.764317 | 0.757 | 0.765 | 3.8219E-13 |
| Intermediate | Mrc2 | 4.10706E-73 | 0.763703 | 0.417 | 0.103 | 5.43898E-69 |
| Rgs5_Contractile | Ntf3 | 1.15617E-12 | 0.763302 | 0.247 | 0.113 | 1.53111E-08 |
| Contractile | Ptprz1 | 4.27946E-77 | 0.763111 | 0.321 | 0.032 | 5.66729E-73 |
| Terminal | Edil3 | 6.70012E-94 | 0.761806 | 0.3 | 0.016 | 8.87297E-90 |
| Intermediate | Bok | 9.82852E-52 | 0.761243 | 0.353 | 0.105 | 1.30159E-47 |
| Intermediate | Gm12840 | 4.29905E-08 | 0.761041 | 0.292 | 0.198 | 0.000569323 |
| Contractile | Ifi27 | 2.85526E-36 | 0.76089 | 0.879 | 0.741 | 3.78122E-32 |
| Rgs5_Contractile | Cox6a1 | 9.69722E-26 | 0.76017 | 0.844 | 0.9 | 1.2842E-21 |

| | | | | | | |
|---|---|---|---|---|---|---|
| Contractile | Kank1 | 9.12851E-27 | 0.7601 | 0.547 | 0.296 | 1.20889E-22 |
| Rgs5_Contractile | Gucy1b3 | 9.39649E-09 | 0.758847 | 0.243 | 0.134 | 0.000124438 |
| Intermediate | Lox | 6.47916E-37 | 0.758843 | 0.632 | 0.389 | 8.58035E-33 |
| Terminal | Sdk2 | 8.20881E-94 | 0.7565 | 0.419 | 0.076 | 1.08709E-89 |
| Rgs5_Contractile | Tgm2 | 2.95118E-05 | 0.753566 | 0.431 | 0.418 | 0.390824724 |
| Rgs5_Contractile | Zbtb38 | 0.00145596 | 0.753424 | 0.389 | 0.405 | 1 |
| Intermediate | Tgfbi | 1.0145E-30 | 0.752883 | 0.422 | 0.208 | 1.3435E-26 |
| Rgs5_Contractile | Filip1 | 6.50408E-05 | 0.75212 | 0.181 | 0.115 | 0.861334667 |
| Contractile | Hsp90aa1 | 1.18651E-42 | 0.751792 | 0.947 | 0.797 | 1.5713E-38 |
| Contractile | Fstl3 | 1.37751E-34 | 0.750093 | 0.426 | 0.155 | 1.82423E-30 |
| Rgs5_Contractile | Rasgrp2 | 2.81774E-08 | 0.748853 | 0.184 | 0.09 | 0.000373153 |
| Rgs5_Contractile | Itpr1 | 0.001118725 | 0.74879 | 0.257 | 0.215 | 1 |
| Intermediate | Bmp1 | 1.30852E-60 | 0.74759 | 0.44 | 0.149 | 1.73287E-56 |
| Rgs5_Contractile | Zfhx3 | 0.00101538 | 0.74727 | 0.431 | 0.461 | 1 |
| Intermediate | Angptl2 | 2.17924E-48 | 0.746747 | 0.499 | 0.225 | 2.88597E-44 |
| Intermediate | Ccl19 | 1.30674E-22 | 0.74606 | 0.183 | 0.06 | 1.73051E-18 |
| Rgs5_Contractile | Rhob | 0.001127524 | 0.746046 | 0.365 | 0.362 | 1 |
| Rgs5_Contractile | Rbp1 | 9.68821E-13 | 0.74536 | 0.722 | 0.657 | 1.28301E-08 |
| Terminal | Scin | 6.1579E-112 | 0.742895 | 0.331 | 0.012 | 8.1549E-108 |
| Contractile | Mmd | 1.21749E-51 | 0.74169 | 0.324 | 0.061 | 1.61232E-47 |
| Contractile | Cds2 | 5.88839E-35 | 0.74161 | 0.379 | 0.127 | 7.79799E-31 |
| Contractile | Fxyd1 | 4.46763E-49 | 0.740714 | 0.971 | 0.868 | 5.91649E-45 |
| Contractile | Rnaseh2c | 3.77173E-18 | 0.740675 | 0.462 | 0.286 | 4.9949E-14 |
| Rgs5_Contractile | Fam13c | 7.34169E-06 | 0.740406 | 0.194 | 0.119 | 0.097225987 |
| Contractile | Sema3c | 4.00473E-41 | 0.739489 | 0.332 | 0.081 | 5.30346E-37 |
| Contractile | Ndufa6 | 3.62355E-32 | 0.738747 | 0.856 | 0.791 | 4.79867E-28 |
| Rgs5_Contractile | Crip1 | 3.26344E-18 | 0.73868 | 0.958 | 0.939 | 4.32178E-14 |
| Rgs5_Contractile | Cbr2 | 5.39085E-06 | 0.738165 | 0.403 | 0.338 | 0.071390979 |
| Contractile | Rbpms2 | 1.99321E-32 | 0.73745 | 0.388 | 0.139 | 2.63961E-28 |
| Contractile | Susd5 | 3.63409E-26 | 0.737373 | 0.676 | 0.444 | 4.81262E-22 |
| Rgs5_Contractile | Nudt4 | 7.68288E-09 | 0.737352 | 0.625 | 0.688 | 0.000101744 |
| Contractile | Mir143hg | 1.7809E-45 | 0.735891 | 0.356 | 0.084 | 2.35845E-41 |
| Intermediate | Capg | 3.13576E-46 | 0.734943 | 0.71 | 0.468 | 4.15269E-42 |
| Contractile | Gabra3 | 7.00531E-62 | 0.734483 | 0.321 | 0.045 | 9.27713E-58 |
| Rgs5_Contractile | Rgs17 | 6.13496E-25 | 0.73446 | 0.132 | 0.019 | 8.12453E-21 |
| Contractile | Mbnl1 | 3.08083E-33 | 0.734435 | 0.856 | 0.715 | 4.07995E-29 |
| Intermediate | Ackr3 | 1.20142E-39 | 0.732459 | 0.372 | 0.144 | 1.59104E-35 |
| Rgs5_Contractile | Kif5b | 7.326E-08 | 0.731905 | 0.562 | 0.599 | 0.000970183 |
| Rgs5_Contractile | Tprgl | 5.11035E-06 | 0.731265 | 0.531 | 0.6 | 0.067676407 |
| Intermediate | Gja1 | 2.11922E-35 | 0.73013 | 0.6 | 0.369 | 2.80649E-31 |
| Terminal | Smpd3 | 4.10022E-50 | 0.727998 | 0.166 | 0.009 | 5.42992E-46 |
| Rgs5_Contractile | Prss23 | 1.82284E-05 | 0.726492 | 0.365 | 0.285 | 0.241398842 |
| Intermediate | Gm42418 | 8.49735E-26 | 0.725478 | 0.792 | 0.655 | 1.1253E-21 |
| Contractile | H3f3a | 4.66097E-47 | 0.723274 | 0.95 | 0.883 | 6.17252E-43 |
| Rgs5_Contractile | Pdgfd | 3.1229E-06 | 0.720598 | 0.222 | 0.139 | 0.041356515 |
| Contractile | Serpinb6b | 2.92749E-16 | 0.720571 | 0.356 | 0.182 | 3.87688E-12 |
| Contractile | Thra | 1.03864E-23 | 0.719914 | 0.629 | 0.441 | 1.37548E-19 |
| Intermediate | Ctsh | 8.42389E-46 | 0.719355 | 0.773 | 0.557 | 1.11558E-41 |
| Contractile | Speg | 8.77166E-31 | 0.719296 | 0.415 | 0.165 | 1.16163E-26 |
| Contractile | Coro1c | 1.10213E-27 | 0.719256 | 0.479 | 0.231 | 1.45956E-23 |
| Intermediate | Ankrd1 | 2.79567E-16 | 0.71886 | 0.132 | 0.04 | 3.70231E-12 |
| Rgs5_Contractile | Htr1b | 4.06008E-11 | 0.718796 | 0.16 | 0.059 | 5.37676E-07 |
| Contractile | Dapk2 | 3.18798E-46 | 0.718613 | 0.247 | 0.037 | 4.22184E-42 |
| Intermediate | Selp | 3.86193E-64 | 0.718571 | 0.221 | 0.016 | 5.11436E-60 |
| Rgs5_Contractile | Gm13889 | 1.26143E-06 | 0.717598 | 0.469 | 0.406 | 0.01670516 |
| Contractile | Wdr1 | 5.41039E-28 | 0.717415 | 0.665 | 0.434 | 7.16498E-24 |
| Contractile | Acadm | 1.0245E-21 | 0.717229 | 0.491 | 0.283 | 1.35675E-17 |
| Rgs5_Contractile | Gstt1 | 0.000411023 | 0.715201 | 0.309 | 0.267 | 1 |
| Contractile | Uqcrb | 4.65778E-45 | 0.715012 | 0.941 | 0.897 | 6.1683E-41 |

| | | | | | | |
|---|---|---|---|---|---|---|
| Rgs5_Contractile | Vasp | 0.008486763 | 0.714762 | 0.378 | 0.415 | 1 |
| Terminal | Trps1 | 1.32263E-61 | 0.712379 | 0.617 | 0.273 | 1.75156E-57 |
| Contractile | Plekho1 | 1.54337E-44 | 0.712143 | 0.315 | 0.067 | 2.04388E-40 |
| Contractile | Sepw1 | 3.45982E-45 | 0.712094 | 0.953 | 0.855 | 4.58184E-41 |
| Rgs5_Contractile | Lbh | 0.008201898 | 0.711991 | 0.333 | 0.334 | 1 |
| Contractile | Chpt1 | 2.88359E-21 | 0.710332 | 0.524 | 0.334 | 3.81874E-17 |
| Terminal | Vit | 9.58103E-64 | 0.709793 | 0.329 | 0.067 | 1.26882E-59 |
| Intermediate | Prss23 | 2.72474E-39 | 0.707899 | 0.459 | 0.198 | 3.60837E-35 |
| Contractile | Tacc2 | 5.74425E-23 | 0.707882 | 0.321 | 0.128 | 7.60711E-19 |
| Contractile | Dynll1 | 5.0515E-51 | 0.707665 | 0.985 | 0.964 | 6.6897E-47 |
| Rgs5_Contractile | Notch3 | 1.21079E-08 | 0.70729 | 0.142 | 0.058 | 0.000160345 |
| Rgs5_Contractile | Rbms3 | 9.7563E-05 | 0.707012 | 0.288 | 0.23 | 1 |
| Contractile | Sypl | 1.56222E-17 | 0.707005 | 0.582 | 0.437 | 2.06885E-13 |
| Rgs5_Contractile | Ilk | 1.38097E-05 | 0.70534 | 0.5 | 0.529 | 0.18288172 |
| Terminal | Fam180a | 6.01831E-42 | 0.705058 | 0.216 | 0.041 | 7.97005E-38 |
| Intermediate | Tm4sf1 | 1.04358E-45 | 0.705032 | 0.985 | 0.954 | 1.38201E-41 |
| Contractile | Ldhb | 2.14855E-22 | 0.704434 | 0.594 | 0.397 | 2.84533E-18 |
| Intermediate | Tnc | 1.72534E-32 | 0.703932 | 0.591 | 0.36 | 2.28487E-28 |
| Rgs5_Contractile | Slc39a1 | 4.83709E-05 | 0.703028 | 0.462 | 0.451 | 0.640575808 |
| Contractile | Lgals1 | 3.83286E-47 | 0.702016 | 0.997 | 0.94 | 5.07586E-43 |
| Contractile | Cox4i1 | 6.75381E-56 | 0.701919 | 0.971 | 0.939 | 8.94407E-52 |
| Contractile | Pdcl3 | 5.50422E-24 | 0.701717 | 0.721 | 0.556 | 7.28924E-20 |
| Intermediate | Efemp1 | 7.80052E-21 | 0.7017 | 0.217 | 0.088 | 1.03302E-16 |
| Rgs5_Contractile | Slc38a2 | 0.000660455 | 0.700879 | 0.465 | 0.512 | 1 |
| Contractile | Dst | 1.34423E-30 | 0.700754 | 0.735 | 0.526 | 1.78016E-26 |
| Intermediate | Postn | 5.83598E-24 | 0.699622 | 0.682 | 0.47 | 7.72858E-20 |
| Rgs5_Contractile | Tob2 | 0.000268594 | 0.699528 | 0.278 | 0.227 | 1 |
| Terminal | Col6a1 | 1.35895E-34 | 0.699343 | 0.891 | 0.681 | 1.79966E-30 |
| Intermediate | Angptl4 | 2.19282E-43 | 0.698852 | 0.202 | 0.032 | 2.90395E-39 |
| Terminal | Maged2 | 2.92102E-50 | 0.698442 | 0.666 | 0.367 | 3.86831E-46 |
| Contractile | Tceal6 | 1.22832E-60 | 0.698279 | 0.212 | 0.015 | 1.62666E-56 |
| Intermediate | Tmem119 | 2.2277E-57 | 0.697366 | 0.313 | 0.07 | 2.95014E-53 |
| Rgs5_Contractile | Plin4 | 0.000671305 | 0.69734 | 0.212 | 0.158 | 1 |
| Terminal | Srpx | 4.31453E-35 | 0.697303 | 0.257 | 0.078 | 5.71373E-31 |
| Rgs5_Contractile | Cox8a | 8.25313E-20 | 0.695887 | 0.875 | 0.927 | 1.09296E-15 |
| Contractile | Arl4a | 9.16244E-19 | 0.695569 | 0.541 | 0.342 | 1.21338E-14 |
| Contractile | Inpp5a | 6.69119E-27 | 0.694904 | 0.371 | 0.148 | 8.86115E-23 |
| Rgs5_Contractile | Pgm5 | 3.20696E-11 | 0.693892 | 0.146 | 0.05 | 4.24698E-07 |
| Intermediate | C4b | 9.15913E-20 | 0.692863 | 0.425 | 0.251 | 1.21294E-15 |
| Rgs5_Contractile | Ehd2 | 0.00241932 | 0.690733 | 0.361 | 0.363 | 1 |
| Rgs5_Contractile | Rasl11b | 5.82518E-07 | 0.690632 | 0.288 | 0.193 | 0.007714287 |
| Rgs5_Contractile | Speg | 0.000819392 | 0.690349 | 0.243 | 0.194 | 1 |
| Contractile | Ldb3 | 2.36079E-50 | 0.689844 | 0.297 | 0.051 | 3.12639E-46 |
| Terminal | Plod2 | 5.54912E-50 | 0.689204 | 0.728 | 0.42 | 7.3487E-46 |
| Intermediate | Sulf1 | 1.96387E-42 | 0.689049 | 0.65 | 0.391 | 2.60075E-38 |
| Contractile | Sepp1 | 2.57585E-25 | 0.688708 | 0.718 | 0.525 | 3.4112E-21 |
| Terminal | Col9a2 | 1.85052E-62 | 0.688571 | 0.19 | 0.006 | 2.45064E-58 |
| Terminal | Ppa1 | 1.25087E-40 | 0.688062 | 0.598 | 0.323 | 1.65652E-36 |
| Rgs5_Contractile | Jag1 | 0.001390298 | 0.687164 | 0.188 | 0.137 | 1 |
| Rgs5_Contractile | Entpd1 | 4.04996E-09 | 0.68712 | 0.174 | 0.077 | 5.36337E-05 |
| Contractile | Dtna | 4.53261E-52 | 0.686951 | 0.259 | 0.036 | 6.00254E-48 |
| Intermediate | Marcks | 7.10148E-49 | 0.686428 | 0.403 | 0.141 | 9.40449E-45 |
| Intermediate | Tspo | 1.2075E-66 | 0.685948 | 0.953 | 0.899 | 1.59909E-62 |
| Terminal | Cpne8 | 3.33196E-61 | 0.685802 | 0.535 | 0.216 | 4.41251E-57 |
| Contractile | Bik | 7.6752E-49 | 0.685197 | 0.271 | 0.044 | 1.01643E-44 |
| Rgs5_Contractile | Clasp2 | 5.05118E-06 | 0.684898 | 0.163 | 0.09 | 0.066892716 |
| Rgs5_Contractile | Ncoa3 | 0.001850685 | 0.683781 | 0.215 | 0.171 | 1 |
| Contractile | Zc2hc1a | 9.27951E-22 | 0.68169 | 0.471 | 0.254 | 1.22889E-17 |
| Contractile | Dek | 2.75688E-22 | 0.680321 | 0.638 | 0.468 | 3.65093E-18 |

| | | | | | | |
|---|---|---|---|---|---|---|
| Contractile | Ndufa11 | 7.91345E-31 | 0.678876 | 0.882 | 0.777 | 1.04798E-26 |
| Contractile | Lhfp | 4.1985E-32 | 0.678645 | 0.903 | 0.788 | 5.56007E-28 |
| Intermediate | Pdgfrb | 4.02621E-38 | 0.678546 | 0.556 | 0.312 | 5.3319E-34 |
| Rgs5_Contractile | Gpi1 | 0.00554681 | 0.678426 | 0.448 | 0.526 | 1 |
| Contractile | Tnfrsf12a | 7.53927E-17 | 0.677627 | 0.665 | 0.499 | 9.98426E-13 |
| Contractile | Otud1 | 5.71868E-27 | 0.677482 | 0.329 | 0.117 | 7.57325E-23 |
| Rgs5_Contractile | Penk | 4.62731E-06 | 0.677236 | 0.028 | 0.119 | 0.061279485 |
| Intermediate | F2r | 8.64994E-55 | 0.676433 | 0.39 | 0.114 | 1.14551E-50 |
| Intermediate | Cxcl2 | 7.21604E-16 | 0.675099 | 0.102 | 0.026 | 9.5562E-12 |
| Intermediate | Lgmn | 2.7522E-44 | 0.675067 | 0.771 | 0.538 | 3.64474E-40 |
| Intermediate | Sulf2 | 1.16202E-39 | 0.674314 | 0.539 | 0.277 | 1.53886E-35 |
| Contractile | Slc25a4 | 2.47914E-48 | 0.669174 | 0.974 | 0.924 | 3.28313E-44 |
| Contractile | Mtus2 | 5.59747E-20 | 0.66857 | 0.453 | 0.248 | 7.41273E-16 |
| Contractile | Tmem123 | 2.35653E-27 | 0.668171 | 0.494 | 0.242 | 3.12075E-23 |
| Contractile | Fam198b | 1.73226E-23 | 0.66769 | 0.494 | 0.261 | 2.29403E-19 |
| Contractile | Cox6a1 | 8.78652E-45 | 0.667224 | 0.962 | 0.882 | 1.1636E-40 |
| Intermediate | Ltc4s | 9.22531E-30 | 0.666968 | 0.121 | 0.014 | 1.22171E-25 |
| Contractile | Csde1 | 9.33449E-25 | 0.666814 | 0.7 | 0.523 | 1.23617E-20 |
| Intermediate | Scn1b | 5.55518E-51 | 0.666659 | 0.709 | 0.433 | 7.35673E-47 |
| Contractile | Rras | 5.33653E-19 | 0.666338 | 0.603 | 0.443 | 7.06717E-15 |
| Intermediate | Calu | 2.28998E-54 | 0.665641 | 0.783 | 0.513 | 3.03261E-50 |
| Contractile | Tln1 | 1.80946E-26 | 0.66561 | 0.697 | 0.478 | 2.39626E-22 |
| Rgs5_Contractile | Ndufv3 | 2.65312E-09 | 0.665473 | 0.67 | 0.783 | 3.51352E-05 |
| Contractile | Fam19a1 | 1.24122E-65 | 0.665306 | 0.224 | 0.015 | 1.64375E-61 |
| Terminal | Nt5dc2 | 6.32951E-24 | 0.665183 | 0.209 | 0.072 | 8.38217E-20 |
| Rgs5_Contractile | Tspan2 | 0.006173632 | 0.664717 | 0.264 | 0.236 | 1 |
| Contractile | Actb | 4.49152E-53 | 0.664532 | 1 | 0.977 | 5.94811E-49 |
| Contractile | Macf1 | 4.867E-15 | 0.663339 | 0.491 | 0.332 | 6.44537E-11 |
| Rgs5_Contractile | Phldb2 | 0.005008227 | 0.663286 | 0.24 | 0.203 | 1 |
| Contractile | Ptms | 3.02705E-33 | 0.663006 | 0.894 | 0.795 | 4.00873E-29 |
| Terminal | Gpm6b | 2.28318E-47 | 0.662727 | 0.475 | 0.194 | 3.02362E-43 |
| Rgs5_Contractile | Ptprz1 | 2.91219E-09 | 0.662408 | 0.153 | 0.062 | 3.85661E-05 |
| Contractile | Ndufv3 | 4.05408E-26 | 0.661572 | 0.838 | 0.759 | 5.36881E-22 |
| Intermediate | Ngf | 1.10336E-47 | 0.661159 | 0.167 | 0.011 | 1.46118E-43 |
| Contractile | Ndufb3 | 4.34435E-26 | 0.659078 | 0.788 | 0.648 | 5.75323E-22 |
| Terminal | Tob1 | 5.86201E-49 | 0.658391 | 0.663 | 0.357 | 7.76307E-45 |
| Contractile | Sdpr | 1.91042E-47 | 0.656654 | 0.279 | 0.048 | 2.52997E-43 |
| Contractile | Gpi1 | 1.41583E-20 | 0.656611 | 0.653 | 0.494 | 1.87498E-16 |
| Contractile | Mob2 | 1.48576E-18 | 0.656545 | 0.468 | 0.274 | 1.96759E-14 |
| Rgs5_Contractile | Csde1 | 0.002370231 | 0.655453 | 0.469 | 0.558 | 1 |
| Contractile | Cox7a2 | 6.97372E-36 | 0.654894 | 0.912 | 0.838 | 9.2353E-32 |
| Contractile | Rock2 | 1.52785E-17 | 0.654517 | 0.544 | 0.375 | 2.02333E-13 |
| Rgs5_Contractile | Hmgcs2 | 7.30965E-06 | 0.652971 | 0.163 | 0.089 | 0.096801679 |
| Intermediate | CR974586. | 2.4177E-33 | 0.651257 | 0.126 | 0.012 | 3.20175E-29 |
| Contractile | Vcan | 1.34076E-10 | 0.649931 | 0.418 | 0.285 | 1.77557E-06 |
| Rgs5_Contractile | Slc22a1 | 3.82074E-12 | 0.649767 | 0.17 | 0.06 | 5.0598E-08 |
| Contractile | Napepld | 7.44092E-33 | 0.649197 | 0.315 | 0.092 | 9.85401E-29 |
| Contractile | Arid5b | 8.38242E-23 | 0.649002 | 0.612 | 0.387 | 1.11008E-18 |
| Rgs5_Contractile | Pkdcc | 0.000104295 | 0.647932 | 0.16 | 0.098 | 1 |
| Contractile | Actn4 | 1.14716E-26 | 0.647589 | 0.771 | 0.599 | 1.51918E-22 |
| Rgs5_Contractile | 2900026A( | 0.00438234 | 0.647296 | 0.198 | 0.157 | 1 |
| Intermediate | Slurp1 | 5.33221E-14 | 0.647002 | 0.208 | 0.099 | 7.06144E-10 |
| Contractile | Eml2 | 4.36438E-27 | 0.646389 | 0.368 | 0.145 | 5.77974E-23 |
| Contractile | Vasp | 8.6674E-24 | 0.645965 | 0.606 | 0.378 | 1.14782E-19 |
| Terminal | Klf2 | 2.41351E-21 | 0.64589 | 0.713 | 0.529 | 3.19621E-17 |
| Contractile | Utrn | 6.95394E-18 | 0.644037 | 0.453 | 0.264 | 9.2091E-14 |
| Contractile | Ndufa4 | 1.73354E-37 | 0.644022 | 0.935 | 0.854 | 2.29573E-33 |
| Rgs5_Contractile | Prkar1a | 8.65103E-06 | 0.643752 | 0.566 | 0.669 | 0.114565524 |
| Contractile | Asph | 3.85157E-17 | 0.643594 | 0.559 | 0.384 | 5.10064E-13 |

| | | | | | | |
|---|---|---|---|---|---|---|
| Contractile | 4930523C( | 2.08783E-23 | 0.643533 | 0.794 | 0.653 | 2.76492E-19 |
| Intermediate | Col1a2 | 3.80354E-26 | 0.643377 | 0.979 | 0.902 | 5.03703E-22 |
| Contractile | Mgll | 6.96007E-25 | 0.642892 | 0.432 | 0.2 | 9.21722E-21 |
| Contractile | Hsph1 | 7.28148E-27 | 0.64274 | 0.4 | 0.164 | 9.64287E-23 |
| Contractile | Ppp1r2 | 1.16137E-18 | 0.64267 | 0.571 | 0.396 | 1.538E-14 |
| Intermediate | Chpf | 2.71338E-38 | 0.642303 | 0.678 | 0.455 | 3.59334E-34 |
| Contractile | Bex4 | 1.83829E-56 | 0.641725 | 0.206 | 0.016 | 2.43445E-52 |
| Contractile | Fam13c | 3.11461E-37 | 0.640876 | 0.338 | 0.093 | 4.12467E-33 |
| Contractile | Serpini1 | 6.64558E-22 | 0.64032 | 0.329 | 0.135 | 8.80074E-18 |
| Contractile | Pfkp | 1.80319E-20 | 0.640271 | 0.424 | 0.221 | 2.38797E-16 |
| Contractile | Tob2 | 1.0672E-14 | 0.639083 | 0.374 | 0.21 | 1.4133E-10 |
| Contractile | Tmem38b | 6.42767E-14 | 0.637798 | 0.465 | 0.315 | 8.51216E-10 |
| Contractile | Ptges3 | 4.20461E-20 | 0.637591 | 0.629 | 0.468 | 5.56817E-16 |
| Contractile | Ndufa5 | 1.52265E-24 | 0.637228 | 0.806 | 0.681 | 2.01644E-20 |
| Rgs5_Contractile | Unc45a | 0.00889376 | 0.636863 | 0.194 | 0.156 | 1 |
| Contractile | Eif4g2 | 5.7208E-29 | 0.636791 | 0.859 | 0.727 | 7.57605E-25 |
| Contractile | Myom1 | 7.1384E-38 | 0.636645 | 0.335 | 0.086 | 9.45339E-34 |
| Contractile | Ndufb9 | 2.27292E-27 | 0.635566 | 0.832 | 0.752 | 3.01003E-23 |
| Intermediate | Ctsb | 5.43421E-72 | 0.634621 | 0.97 | 0.912 | 7.19653E-68 |
| Terminal | Gadd45a | 6.37512E-28 | 0.634429 | 0.389 | 0.185 | 8.44257E-24 |
| Contractile | Pde3a | 1.73826E-19 | 0.633841 | 0.574 | 0.364 | 2.30198E-15 |
| Contractile | Ngfrap1 | 1.71912E-16 | 0.633378 | 0.432 | 0.261 | 2.27663E-12 |
| Intermediate | Tcn2 | 9.51458E-51 | 0.633286 | 0.398 | 0.135 | 1.26002E-46 |
| Intermediate | Mdk | 5.82683E-14 | 0.632721 | 0.385 | 0.256 | 7.71647E-10 |
| Rgs5_Contractile | Sorl1 | 0.000347277 | 0.632644 | 0.212 | 0.151 | 1 |
| Contractile | Cox17 | 3.28333E-18 | 0.63172 | 0.712 | 0.623 | 4.34811E-14 |
| Intermediate | Rtn4 | 2.10489E-44 | 0.631189 | 0.83 | 0.625 | 2.78751E-40 |
| Intermediate | Pamr1 | 1.12833E-42 | 0.631005 | 0.331 | 0.103 | 1.49425E-38 |
| Rgs5_Contractile | Gm12840 | 0.000180883 | 0.63088 | 0.299 | 0.224 | 1 |
| Terminal | Cdc42ep3 | 7.82791E-41 | 0.630694 | 0.825 | 0.575 | 1.03665E-36 |
| Intermediate | Igfbp7 | 6.05289E-67 | 0.630016 | 1 | 0.996 | 8.01584E-63 |
| Contractile | Dusp1 | 1.00554E-20 | 0.629891 | 0.762 | 0.562 | 1.33164E-16 |
| Rgs5_Contractile | Ptges3 | 0.005288855 | 0.628261 | 0.431 | 0.499 | 1 |
| Intermediate | Txndc5 | 4.53697E-50 | 0.627568 | 0.723 | 0.449 | 6.00831E-46 |
| Intermediate | Gm10116 | 3.64215E-38 | 0.626991 | 0.709 | 0.47 | 4.8233E-34 |
| Intermediate | Cd34 | 8.40688E-28 | 0.626319 | 0.314 | 0.136 | 1.11332E-23 |
| Terminal | Atp2b1 | 6.99621E-45 | 0.626283 | 0.746 | 0.438 | 9.26509E-41 |
| Intermediate | Ppp1r14b | 1.54278E-37 | 0.625477 | 0.559 | 0.312 | 2.04311E-33 |
| Rgs5_Contractile | Atp5k | 1.92351E-11 | 0.625386 | 0.712 | 0.794 | 2.54731E-07 |
| Rgs5_Contractile | Ctgf | 1.65003E-10 | 0.624934 | 0.688 | 0.623 | 2.18514E-06 |
| Contractile | Hoxa7 | 1.62849E-35 | 0.623784 | 0.229 | 0.044 | 2.1566E-31 |
| Contractile | Hipk3 | 6.33643E-19 | 0.623783 | 0.391 | 0.198 | 8.39133E-15 |
| Rgs5_Contractile | Atp1b2 | 1.34488E-06 | 0.621192 | 0.115 | 0.048 | 0.017810246 |
| Contractile | Aes | 1.18E-19 | 0.620992 | 0.668 | 0.491 | 1.56267E-15 |
| Rgs5_Contractile | Arglu1 | 0.003692035 | 0.620925 | 0.375 | 0.396 | 1 |
| Contractile | Tbl1x | 4.34489E-16 | 0.620738 | 0.429 | 0.258 | 5.75393E-12 |
| Rgs5_Contractile | Rock2 | 0.007714744 | 0.620665 | 0.375 | 0.403 | 1 |
| Contractile | Cd200 | 4.76462E-31 | 0.620549 | 0.956 | 0.849 | 6.30979E-27 |
| Intermediate | Bgn | 2.07456E-65 | 0.620427 | 0.997 | 0.995 | 2.74734E-61 |
| Intermediate | Plpp3 | 5.16541E-23 | 0.61882 | 0.668 | 0.468 | 6.84055E-19 |
| Rgs5_Contractile | Mmd | 2.23545E-08 | 0.618209 | 0.181 | 0.087 | 0.00029604 |
| Contractile | Hspa1a | 1.04355E-11 | 0.618194 | 0.55 | 0.393 | 1.38198E-07 |
| Contractile | Gm11627 | 2.7166E-41 | 0.618044 | 0.262 | 0.05 | 3.5976E-37 |
| Rgs5_Contractile | Ank2 | 0.00058146 | 0.617912 | 0.153 | 0.098 | 1 |
| Intermediate | Fzd1 | 3.13201E-33 | 0.617737 | 0.453 | 0.229 | 4.14772E-29 |
| Contractile | Adam33 | 4.22078E-56 | 0.61743 | 0.241 | 0.026 | 5.58958E-52 |
| Rgs5_Contractile | Ndufa3 | 1.10112E-06 | 0.61714 | 0.611 | 0.727 | 0.014582137 |
| Rgs5_Contractile | Marf1 | 0.003193102 | 0.61617 | 0.205 | 0.161 | 1 |
| Contractile | Phactr1 | 4.62712E-22 | 0.615871 | 0.291 | 0.107 | 6.1277E-18 |

| | | | | | | |
|---|---|---|---|---|---|---|
| Rgs5_Contractile | Art3 | 3.03825E-05 | 0.615233 | 0.167 | 0.096 | 0.402355245 |
| Rgs5_Contractile | Rnd3 | 0.00703574 | 0.614775 | 0.184 | 0.142 | 1 |
| Rgs5_Contractile | Dapk2 | 1.02321E-09 | 0.614235 | 0.146 | 0.056 | 1.35504E-05 |
| Contractile | Gamt | 2.11298E-14 | 0.613994 | 0.347 | 0.192 | 2.79821E-10 |
| Contractile | S100a6 | 2.17354E-38 | 0.613705 | 0.988 | 0.968 | 2.87842E-34 |
| Rgs5_Contractile | Atp5h | 1.04371E-15 | 0.613178 | 0.806 | 0.879 | 1.38218E-11 |
| Terminal | Cgref1 | 4.75509E-47 | 0.612819 | 0.488 | 0.206 | 6.29717E-43 |
| Contractile | Ehbp1l1 | 2.42484E-15 | 0.612544 | 0.394 | 0.229 | 3.21121E-11 |
| Contractile | Itpr1 | 1.11191E-20 | 0.612255 | 0.394 | 0.192 | 1.47251E-16 |
| Contractile | Rasl11a | 6.26241E-20 | 0.61145 | 0.45 | 0.231 | 8.29331E-16 |
| Contractile | Unc45a | 7.22572E-21 | 0.611249 | 0.324 | 0.134 | 9.56902E-17 |
| Intermediate | Mt2 | 8.46366E-05 | 0.610749 | 0.808 | 0.752 | 1 |
| Contractile | Uchl1 | 4.44739E-23 | 0.61049 | 0.403 | 0.177 | 5.88968E-19 |
| Contractile | B230219D | 2.74373E-11 | 0.610063 | 0.529 | 0.421 | 3.63352E-07 |
| Contractile | Heyl | 1.83397E-39 | 0.608644 | 0.294 | 0.065 | 2.42873E-35 |
| Intermediate | AA467197 | 1.91019E-40 | 0.60858 | 0.286 | 0.084 | 2.52966E-36 |
| Contractile | Fbxo2 | 9.22428E-32 | 0.608389 | 0.215 | 0.045 | 1.22157E-27 |
| Intermediate | Col15a1 | 2.84522E-27 | 0.607919 | 0.499 | 0.292 | 3.76792E-23 |
| Intermediate | Col4a1 | 1.24301E-20 | 0.607273 | 0.572 | 0.375 | 1.64612E-16 |
| Contractile | 2200002D | 6.72595E-13 | 0.606936 | 0.374 | 0.219 | 8.90717E-09 |
| Contractile | Atp5h | 1.23308E-32 | 0.606512 | 0.9 | 0.865 | 1.63297E-28 |
| Intermediate | Col5a2 | 1.41998E-39 | 0.606508 | 0.888 | 0.681 | 1.88048E-35 |
| Intermediate | Myl12a | 1.85071E-62 | 0.60619 | 0.964 | 0.932 | 2.45089E-58 |
| Contractile | Ank2 | 7.51445E-34 | 0.6057 | 0.288 | 0.075 | 9.95138E-30 |
| Contractile | Ccni | 4.95569E-18 | 0.605644 | 0.556 | 0.379 | 6.56282E-14 |
| Intermediate | Scarf2 | 7.14734E-29 | 0.605638 | 0.522 | 0.313 | 9.46522E-25 |
| Rgs5_Contractile | Bri3 | 3.22977E-11 | 0.604991 | 0.719 | 0.792 | 4.27718E-07 |
| Rgs5_Contractile | Hbb-bs | 3.54951E-05 | 0.604514 | 0.16 | 0.093 | 0.470062009 |
| Intermediate | Cdh13 | 3.41039E-44 | 0.604468 | 0.648 | 0.339 | 4.51638E-40 |
| Intermediate | Tulp4 | 8.09496E-40 | 0.604259 | 0.51 | 0.256 | 1.07202E-35 |
| Intermediate | Angpt2 | 3.23531E-36 | 0.603915 | 0.334 | 0.119 | 4.28452E-32 |
| Terminal | Btg2 | 4.40401E-24 | 0.603906 | 0.688 | 0.495 | 5.83223E-20 |
| Rgs5_Contractile | Hey2 | 1.06116E-05 | 0.602391 | 0.167 | 0.095 | 0.140529063 |
| Rgs5_Contractile | Foxp2 | 9.7164E-07 | 0.602152 | 0.174 | 0.09 | 0.012867433 |
| Contractile | Zbtb20 | 1.34322E-23 | 0.601924 | 0.871 | 0.824 | 1.77883E-19 |
| Contractile | Nap1l1 | 3.47387E-15 | 0.601324 | 0.629 | 0.513 | 4.60044E-11 |
| Intermediate | Cd248 | 3.71548E-66 | 0.601017 | 0.3 | 0.047 | 4.92041E-62 |
| Rgs5_Contractile | Rgs7bp | 1.04779E-08 | 0.600575 | 0.142 | 0.058 | 0.000138759 |
| Contractile | Zak | 3.3022E-15 | 0.600536 | 0.485 | 0.317 | 4.37311E-11 |
| Intermediate | Pde1a | 5.80499E-37 | 0.600337 | 0.481 | 0.239 | 7.68755E-33 |
| Intermediate | Cd44 | 1.19567E-28 | 0.600265 | 0.412 | 0.212 | 1.58342E-24 |
| Contractile | Capns1 | 9.33408E-31 | 0.599994 | 0.929 | 0.839 | 1.23611E-26 |
| Rgs5_Contractile | Irak2 | 0.000431318 | 0.599967 | 0.201 | 0.143 | 1 |
| Terminal | Tsc22d3 | 1.43449E-52 | 0.599813 | 0.516 | 0.202 | 1.8997E-48 |
| Contractile | Lbh | 1.45286E-15 | 0.599689 | 0.485 | 0.309 | 1.92402E-11 |
| Intermediate | Mmp23 | 4.85637E-38 | 0.599577 | 0.61 | 0.355 | 6.43129E-34 |
| Rgs5_Contractile | Rsu1 | 0.002481368 | 0.59857 | 0.51 | 0.606 | 1 |
| Contractile | Vldlr | 3.72993E-18 | 0.598468 | 0.288 | 0.122 | 4.93955E-14 |
| Contractile | Tmem109 | 1.51335E-13 | 0.598067 | 0.509 | 0.373 | 2.00413E-09 |
| Terminal | Ptgs2 | 2.7502E-31 | 0.596943 | 0.325 | 0.129 | 3.64209E-27 |
| Contractile | Pnpla2 | 3.68027E-31 | 0.596691 | 0.303 | 0.09 | 4.87378E-27 |
| Contractile | Pgm5 | 5.46097E-53 | 0.595504 | 0.25 | 0.031 | 7.23197E-49 |
| Intermediate | Cxcl16 | 6.67565E-46 | 0.595444 | 0.265 | 0.062 | 8.84057E-42 |
| Intermediate | Vat1 | 1.01511E-41 | 0.594232 | 0.471 | 0.213 | 1.34431E-37 |
| Contractile | Ndufa2 | 8.097E-36 | 0.593825 | 0.921 | 0.866 | 1.07229E-31 |
| Rgs5_Contractile | Eif4g2 | 2.45858E-07 | 0.592587 | 0.649 | 0.758 | 0.003255896 |
| Rgs5_Contractile | Npm1 | 7.27012E-07 | 0.592444 | 0.674 | 0.791 | 0.009627826 |
| Intermediate | Crabp2 | 1.89158E-32 | 0.592363 | 0.172 | 0.034 | 2.50502E-28 |
| Intermediate | Sod3 | 1.25522E-42 | 0.59122 | 0.916 | 0.773 | 1.66229E-38 |

| | | | | | | |
|---|---|---|---|---|---|---|
| Rgs5_Contractile | Hacd1 | 0.00185435 | 0.590092 | 0.396 | 0.408 | 1 |
| Contractile | Ppia | 1.33744E-50 | 0.58936 | 1 | 0.989 | 1.77118E-46 |
| Rgs5_Contractile | Serinc3 | 1.27512E-05 | 0.58922 | 0.597 | 0.718 | 0.168864757 |
| Terminal | Cdkn1c | 1.62764E-14 | 0.589034 | 0.222 | 0.106 | 2.15549E-10 |
| Contractile | Ndufb10 | 9.28299E-22 | 0.58886 | 0.806 | 0.721 | 1.22935E-17 |
| Contractile | Cd59a | 2.99745E-08 | 0.588546 | 0.35 | 0.245 | 0.000396953 |
| Intermediate | C1ra | 1.27764E-29 | 0.588514 | 0.606 | 0.399 | 1.69198E-25 |
| Contractile | Mmp17 | 8.62382E-37 | 0.588507 | 0.218 | 0.038 | 1.14205E-32 |
| Rgs5_Contractile | Anln | 1.66642E-06 | 0.587822 | 0.135 | 0.064 | 0.022068449 |
| Rgs5_Contractile | Adam33 | 9.21122E-08 | 0.587113 | 0.122 | 0.048 | 0.001219842 |
| Intermediate | Loxl2 | 1.83527E-53 | 0.586745 | 0.275 | 0.051 | 2.43044E-49 |
| Intermediate | Epdr1 | 6.44964E-38 | 0.586525 | 0.724 | 0.501 | 8.54126E-34 |
| Terminal | Tspan3 | 8.12475E-52 | 0.585593 | 0.908 | 0.696 | 1.07596E-47 |
| Contractile | Degs1 | 7.97636E-17 | 0.585446 | 0.644 | 0.499 | 1.05631E-12 |
| Rgs5_Contractile | Ccdc42 | 1.61507E-06 | 0.584331 | 0.146 | 0.07 | 0.021388395 |
| Contractile | Mical2 | 4.4481E-22 | 0.583142 | 0.362 | 0.154 | 5.89062E-18 |
| Contractile | Rbms3 | 1.62228E-18 | 0.582171 | 0.406 | 0.209 | 2.14839E-14 |
| Rgs5_Contractile | Lama5 | 0.004250592 | 0.58217 | 0.16 | 0.115 | 1 |
| Contractile | Ndufa3 | 4.1438E-21 | 0.581633 | 0.806 | 0.698 | 5.48764E-17 |
| Rgs5_Contractile | Heyl | 0.000263268 | 0.580856 | 0.149 | 0.09 | 1 |
| Contractile | Map7d1 | 1.22337E-13 | 0.579831 | 0.524 | 0.382 | 1.62011E-09 |
| Contractile | Capzb | 8.53457E-22 | 0.579791 | 0.844 | 0.759 | 1.13023E-17 |
| Terminal | Fgfrl1 | 1.54507E-46 | 0.579487 | 0.404 | 0.147 | 2.04614E-42 |
| Contractile | Bmyc | 2.16662E-16 | 0.578829 | 0.476 | 0.301 | 2.86925E-12 |
| Terminal | Fam20c | 1.93039E-48 | 0.578692 | 0.461 | 0.181 | 2.55641E-44 |
| Intermediate | 1110008P1 | 1.20789E-27 | 0.578629 | 0.72 | 0.536 | 1.5996E-23 |
| Rgs5_Contractile | Col4a1 | 0.000828033 | 0.578374 | 0.448 | 0.448 | 1 |
| Terminal | Itga10 | 2.88807E-59 | 0.577945 | 0.375 | 0.1 | 3.82467E-55 |
| Contractile | Camk2g | 9.8248E-16 | 0.57791 | 0.362 | 0.193 | 1.3011E-11 |
| Rgs5_Contractile | Gm9843 | 3.51616E-10 | 0.577906 | 0.75 | 0.869 | 4.65645E-06 |
| Contractile | Slc25a3 | 1.99784E-25 | 0.577043 | 0.865 | 0.769 | 2.64573E-21 |
| Contractile | Sema3d | 1.60312E-36 | 0.576921 | 0.256 | 0.054 | 2.12301E-32 |
| Intermediate | Mt1 | 3.26409E-06 | 0.576237 | 0.95 | 0.89 | 0.043226388 |
| Intermediate | Tmem176l | 1.51381E-31 | 0.575993 | 0.785 | 0.607 | 2.00474E-27 |
| Contractile | Pnck | 2.59399E-37 | 0.575589 | 0.224 | 0.04 | 3.43522E-33 |
| Intermediate | Manf | 1.32646E-28 | 0.575445 | 0.771 | 0.589 | 1.75663E-24 |
| Rgs5_Contractile | Cds2 | 0.001889233 | 0.5743 | 0.205 | 0.157 | 1 |
| Intermediate | Tubb5 | 4.73031E-33 | 0.5742 | 0.775 | 0.596 | 6.26435E-29 |
| Intermediate | Anxa5 | 5.72809E-46 | 0.574144 | 0.935 | 0.861 | 7.58571E-42 |
| Contractile | Itgb3 | 4.81267E-28 | 0.573194 | 0.253 | 0.068 | 6.37341E-24 |
| Rgs5_Contractile | Uba52 | 0.000368528 | 0.572946 | 0.517 | 0.564 | 1 |
| Rgs5_Contractile | Wdr89 | 4.66398E-11 | 0.572926 | 0.764 | 0.878 | 6.17651E-07 |
| Terminal | Vkorc1 | 2.88792E-38 | 0.572189 | 0.89 | 0.644 | 3.82447E-34 |
| Intermediate | Wisp1 | 3.30265E-43 | 0.571982 | 0.258 | 0.062 | 4.3737E-39 |
| Contractile | Ywhah | 7.08021E-20 | 0.571861 | 0.712 | 0.569 | 9.37632E-16 |
| Contractile | Entpd1 | 1.01123E-33 | 0.571826 | 0.262 | 0.06 | 1.33918E-29 |
| Contractile | Ndufa12 | 1.44111E-18 | 0.571575 | 0.776 | 0.666 | 1.90846E-14 |
| Contractile | Idh3g | 1.24061E-11 | 0.571286 | 0.441 | 0.315 | 1.64294E-07 |
| Contractile | Ethe1 | 3.86397E-16 | 0.571241 | 0.376 | 0.204 | 5.11706E-12 |
| Rgs5_Contractile | Rhobtb1 | 2.18716E-07 | 0.570661 | 0.118 | 0.048 | 0.00289645 |
| Contractile | Dnajb5 | 1.19076E-33 | 0.570093 | 0.221 | 0.043 | 1.57692E-29 |
| Contractile | Atp5b | 4.76253E-27 | 0.569937 | 0.9 | 0.834 | 6.30702E-23 |
| Contractile | Ptp4a3 | 1.42928E-24 | 0.569569 | 0.482 | 0.243 | 1.89279E-20 |
| Contractile | Atp5c1 | 2.15558E-20 | 0.568815 | 0.803 | 0.728 | 2.85464E-16 |
| Rgs5_Contractile | Rhoa | 8.2503E-10 | 0.568187 | 0.715 | 0.813 | 1.09259E-05 |
| Contractile | Tprgl | 2.5312E-19 | 0.567699 | 0.715 | 0.572 | 3.35206E-15 |
| Terminal | Mdfi | 1.35947E-67 | 0.56716 | 0.298 | 0.047 | 1.80034E-63 |
| Rgs5_Contractile | Lgr6 | 5.93041E-05 | 0.567057 | 0.156 | 0.09 | 0.785364035 |
| Rgs5_Contractile | Ndufa11 | 1.82682E-05 | 0.566923 | 0.656 | 0.81 | 0.241925185 |

| | | | | | | |
|---|---|---|---|---|---|---|
| Contractile | Psmb6 | 1.05952E-25 | 0.566858 | 0.859 | 0.753 | 1.40313E-21 |
| Rgs5_Contractile | Fxyd1 | 2.0884E-10 | 0.56671 | 0.802 | 0.894 | 2.76566E-06 |
| Rgs5_Contractile | Sepp1 | 0.000171686 | 0.566343 | 0.521 | 0.556 | 1 |
| Terminal | Fam19a5 | 1.06459E-69 | 0.56602 | 0.345 | 0.063 | 1.40983E-65 |
| Contractile | Hspb8 | 8.74826E-13 | 0.565877 | 0.432 | 0.284 | 1.15853E-08 |
| Contractile | Ak3 | 4.94335E-15 | 0.56564 | 0.326 | 0.17 | 6.54648E-11 |
| Contractile | Crispld2 | 1.39171E-18 | 0.565311 | 0.394 | 0.186 | 1.84304E-14 |
| Intermediate | Oaf | 2.5329E-33 | 0.565059 | 0.496 | 0.271 | 3.35432E-29 |
| Contractile | Camk2n1 | 2.40216E-19 | 0.564787 | 0.45 | 0.247 | 3.18118E-15 |
| Rgs5_Contractile | H3f3a | 1.95964E-12 | 0.564269 | 0.83 | 0.901 | 2.59515E-08 |
| Intermediate | Kcnn4 | 1.16482E-56 | 0.564155 | 0.235 | 0.029 | 1.54257E-52 |
| Terminal | Clint1 | 2.98432E-49 | 0.562812 | 0.507 | 0.219 | 3.95213E-45 |
| Rgs5_Contractile | Nav2 | 0.000257229 | 0.562284 | 0.149 | 0.091 | 1 |
| Contractile | Gnai1 | 2.55925E-12 | 0.562075 | 0.385 | 0.239 | 3.38921E-08 |
| Contractile | Syngr2 | 8.60103E-14 | 0.561871 | 0.406 | 0.244 | 1.13903E-09 |
| Rgs5_Contractile | Hspa8 | 2.15988E-14 | 0.561331 | 0.948 | 0.974 | 2.86033E-10 |
| Rgs5_Contractile | Gm9493 | 8.74446E-05 | 0.56064 | 0.701 | 0.842 | 1 |
| Contractile | Acat1 | 6.86169E-12 | 0.560408 | 0.438 | 0.306 | 9.08693E-08 |
| Intermediate | Mxra8 | 7.30575E-35 | 0.559755 | 0.904 | 0.769 | 9.67501E-31 |
| Terminal | H3f3b | 1.59137E-33 | 0.558278 | 0.985 | 0.949 | 2.10746E-29 |
| Contractile | Atp5k | 1.58081E-20 | 0.557066 | 0.844 | 0.774 | 2.09347E-16 |
| Contractile | Naca | 7.59292E-31 | 0.55537 | 0.941 | 0.905 | 1.00553E-26 |
| Contractile | Ehd4 | 9.12818E-17 | 0.555346 | 0.321 | 0.153 | 1.20885E-12 |
| Contractile | Msrb3 | 2.12961E-19 | 0.554457 | 0.347 | 0.161 | 2.82024E-15 |
| Terminal | Klf10 | 2.20054E-40 | 0.554178 | 0.422 | 0.173 | 2.91417E-36 |
| Intermediate | Vmp1 | 3.08121E-26 | 0.554057 | 0.607 | 0.399 | 4.08045E-22 |
| Contractile | Uqcr10 | 1.92071E-18 | 0.554013 | 0.871 | 0.817 | 2.5436E-14 |
| Contractile | Itga7 | 4.82224E-19 | 0.552947 | 0.288 | 0.116 | 6.38609E-15 |
| Rgs5_Contractile | Prdm16 | 4.18999E-08 | 0.552757 | 0.132 | 0.054 | 0.000554881 |
| Contractile | Plin4 | 1.34939E-20 | 0.552229 | 0.332 | 0.137 | 1.78699E-16 |
| Contractile | Acsl1 | 4.19177E-19 | 0.55083 | 0.276 | 0.108 | 5.55116E-15 |
| Contractile | Cox8a | 1.0742E-28 | 0.550819 | 0.944 | 0.917 | 1.42257E-24 |
| Intermediate | Acp5 | 2.53842E-34 | 0.550812 | 0.291 | 0.103 | 3.36163E-30 |
| Contractile | Gm26722 | 1.7265E-40 | 0.549777 | 0.262 | 0.051 | 2.28641E-36 |
| Intermediate | Tagln2 | 5.40799E-37 | 0.549374 | 0.872 | 0.679 | 7.1618E-33 |
| Contractile | Psmc3 | 1.36163E-13 | 0.549096 | 0.653 | 0.568 | 1.8032E-09 |
| Terminal | Slc29a1 | 6.99788E-34 | 0.548904 | 0.866 | 0.599 | 9.26729E-30 |
| Contractile | Ndufb8 | 3.9765E-15 | 0.548224 | 0.703 | 0.625 | 5.26608E-11 |
| Rgs5_Contractile | Bicd1 | 0.007380024 | 0.548093 | 0.135 | 0.095 | 1 |
| Rgs5_Contractile | Sepw1 | 5.21265E-11 | 0.547565 | 0.795 | 0.879 | 6.90311E-07 |
| Intermediate | Cd63 | 1.11949E-68 | 0.547415 | 0.994 | 0.969 | 1.48254E-64 |
| Contractile | Mfhas1 | 4.1992E-21 | 0.546959 | 0.259 | 0.09 | 5.56099E-17 |
| Intermediate | H2-D1 | 1.02591E-23 | 0.54681 | 0.902 | 0.779 | 1.35861E-19 |
| Contractile | Map1a | 5.70888E-17 | 0.546673 | 0.315 | 0.148 | 7.56027E-13 |
| Terminal | Prelp | 8.04853E-35 | 0.546202 | 0.855 | 0.606 | 1.06587E-30 |
| Rgs5_Contractile | Maml2 | 0.001373116 | 0.546167 | 0.149 | 0.1 | 1 |
| Intermediate | Uqcc2 | 3.25826E-31 | 0.545993 | 0.817 | 0.712 | 4.31491E-27 |
| Contractile | Mrpl23 | 4.12574E-13 | 0.545645 | 0.656 | 0.552 | 5.46372E-09 |
| Contractile | Bmpr1a | 4.48273E-15 | 0.545307 | 0.497 | 0.333 | 5.93648E-11 |
| Contractile | Nbeal1 | 6.35686E-15 | 0.545229 | 0.529 | 0.361 | 8.41838E-11 |
| Intermediate | Prdx1 | 6.23037E-60 | 0.544727 | 0.972 | 0.946 | 8.25087E-56 |
| Intermediate | Fat1 | 3.4235E-28 | 0.544313 | 0.628 | 0.419 | 4.53374E-24 |
| Rgs5_Contractile | Dbndd2 | 0.000235117 | 0.543742 | 0.156 | 0.097 | 1 |
| Contractile | Sod1 | 1.12737E-13 | 0.543479 | 0.494 | 0.347 | 1.49298E-09 |
| Rgs5_Contractile | Arrdc3 | 0.006498089 | 0.543196 | 0.181 | 0.14 | 1 |
| Intermediate | Ctsz | 1.28194E-38 | 0.542905 | 0.809 | 0.628 | 1.69767E-34 |
| Contractile | Enpp5 | 7.11986E-26 | 0.542573 | 0.241 | 0.066 | 9.42883E-22 |
| Contractile | Iscu | 6.10274E-18 | 0.542324 | 0.776 | 0.68 | 8.08185E-14 |
| Intermediate | Ttc9 | 3.76841E-44 | 0.542117 | 0.299 | 0.084 | 4.99051E-40 |

| | | | | | | |
|---|---|---|---|---|---|---|
| Rgs5_Contractile | Atp5b | 1.56415E-08 | 0.541799 | 0.743 | 0.857 | 0.00020714 |
| Contractile | Kank2 | 2.43069E-13 | 0.541256 | 0.374 | 0.22 | 3.21896E-09 |
| Contractile | Abracl | 8.43422E-11 | 0.541191 | 0.521 | 0.406 | 1.11694E-06 |
| Contractile | Aplp2 | 5.56736E-25 | 0.541044 | 0.906 | 0.854 | 7.37285E-21 |
| Contractile | Myeov2 | 4.77927E-20 | 0.540923 | 0.803 | 0.696 | 6.32918E-16 |
| Terminal | Tspan13 | 2.2301E-52 | 0.540558 | 0.18 | 0.012 | 2.95333E-48 |
| Contractile | Gstt1 | 3.31027E-19 | 0.540164 | 0.459 | 0.241 | 4.38379E-15 |
| Contractile | Arpc2 | 5.57178E-30 | 0.539988 | 0.926 | 0.876 | 7.37871E-26 |
| Contractile | Pdlim4 | 7.6478E-14 | 0.539888 | 0.491 | 0.331 | 1.0128E-09 |
| Contractile | Clic4 | 2.32065E-18 | 0.539836 | 0.841 | 0.754 | 3.07324E-14 |
| Rgs5_Contractile | S100a6 | 2.63494E-13 | 0.539292 | 0.951 | 0.974 | 3.48944E-09 |
| Intermediate | Clic1 | 3.23559E-32 | 0.538345 | 0.813 | 0.685 | 4.28489E-28 |
| Terminal | Gas1 | 3.97295E-08 | 0.538292 | 0.479 | 0.371 | 0.000526138 |
| Contractile | Ifi27l2a | 4.38935E-08 | 0.53795 | 0.115 | 0.043 | 0.000581282 |
| Terminal | Nfkbiz | 3.50286E-34 | 0.536566 | 0.471 | 0.231 | 4.63884E-30 |
| Intermediate | Col12a1 | 9.18553E-30 | 0.535776 | 0.327 | 0.136 | 1.21644E-25 |
| Intermediate | Prelid1 | 7.20309E-38 | 0.535749 | 0.696 | 0.469 | 9.53906E-34 |
| Intermediate | Ctsc | 6.30096E-35 | 0.53559 | 0.281 | 0.087 | 8.34436E-31 |
| Intermediate | Hsp90b1 | 2.08667E-32 | 0.5354 | 0.908 | 0.802 | 2.76338E-28 |
| Contractile | Atp5j2 | 1.78429E-25 | 0.535158 | 0.912 | 0.863 | 2.36293E-21 |
| Terminal | Col1a2 | 3.61753E-52 | 0.535049 | 0.997 | 0.892 | 4.79069E-48 |
| Contractile | Axl | 5.01025E-13 | 0.535027 | 0.565 | 0.41 | 6.63508E-09 |
| Intermediate | Mtch1 | 5.30451E-32 | 0.534693 | 0.679 | 0.47 | 7.02476E-28 |
| Contractile | Eif4a2 | 1.1582E-18 | 0.534314 | 0.824 | 0.752 | 1.53381E-14 |
| Contractile | Pdlim5 | 4.78736E-15 | 0.534309 | 0.518 | 0.347 | 6.3399E-11 |
| Rgs5_Contractile | Tnxb | 0.003442762 | 0.534277 | 0.163 | 0.116 | 1 |
| Intermediate | Smoc1 | 7.68819E-34 | 0.533103 | 0.683 | 0.406 | 1.01815E-29 |
| Intermediate | Fkbp11 | 1.17396E-27 | 0.532689 | 0.432 | 0.233 | 1.55467E-23 |
| Contractile | 1110001J0 | 9.77277E-13 | 0.532553 | 0.535 | 0.403 | 1.29421E-08 |
| Contractile | Chd6 | 1.09996E-14 | 0.532475 | 0.344 | 0.183 | 1.45667E-10 |
| Contractile | Mlf2 | 4.93213E-16 | 0.532397 | 0.703 | 0.61 | 6.53163E-12 |
| Intermediate | Psap | 3.12057E-55 | 0.532369 | 0.959 | 0.904 | 4.13257E-51 |
| Contractile | Aktip | 1.39339E-11 | 0.531313 | 0.415 | 0.282 | 1.84526E-07 |
| Contractile | Lamb2 | 9.52272E-11 | 0.530943 | 0.579 | 0.459 | 1.26109E-06 |
| Intermediate | Pmepa1 | 2.581E-37 | 0.53086 | 0.745 | 0.453 | 3.41801E-33 |
| Contractile | Id2 | 5.75303E-15 | 0.530825 | 0.665 | 0.498 | 7.61873E-11 |
| Contractile | Fam96b | 4.09279E-11 | 0.530185 | 0.503 | 0.391 | 5.42008E-07 |
| Rgs5_Contractile | Nkd1 | 7.01925E-05 | 0.529823 | 0.205 | 0.134 | 0.929559521 |
| Terminal | Sec11c | 2.43525E-38 | 0.529192 | 0.74 | 0.465 | 3.225E-34 |
| Contractile | Pink1 | 1.57997E-11 | 0.528976 | 0.538 | 0.431 | 2.09235E-07 |
| Rgs5_Contractile | Ptprd | 0.004096478 | 0.528567 | 0.139 | 0.095 | 1 |
| Contractile | Pygm | 1.03467E-22 | 0.528407 | 0.232 | 0.069 | 1.37021E-18 |
| Intermediate | Olfml2b | 1.46645E-29 | 0.528296 | 0.635 | 0.404 | 1.94202E-25 |
| Terminal | Cd24a | 1.11969E-54 | 0.528185 | 0.178 | 0.009 | 1.48281E-50 |
| Contractile | Anln | 2.03151E-32 | 0.527906 | 0.226 | 0.047 | 2.69033E-28 |
| Contractile | Ndufb7 | 7.8631E-18 | 0.527412 | 0.771 | 0.691 | 1.04131E-13 |
| Intermediate | Cd47 | 3.86191E-34 | 0.527358 | 0.781 | 0.607 | 5.11433E-30 |
| Terminal | Raph1 | 2.30627E-40 | 0.527332 | 0.524 | 0.246 | 3.05419E-36 |
| Contractile | Psma7 | 2.43429E-17 | 0.527032 | 0.8 | 0.728 | 3.22373E-13 |
| Contractile | Zfhx3 | 8.23674E-12 | 0.526981 | 0.556 | 0.442 | 1.09079E-07 |
| Contractile | Psmb7 | 9.54405E-11 | 0.526932 | 0.568 | 0.475 | 1.26392E-06 |
| Intermediate | Sox4 | 3.2935E-12 | 0.526428 | 0.39 | 0.268 | 4.36159E-08 |
| Contractile | Cox7c | 2.71124E-30 | 0.526086 | 0.938 | 0.93 | 3.5905E-26 |
| Intermediate | Calm1 | 3.74995E-38 | 0.525476 | 0.946 | 0.916 | 4.96606E-34 |
| Contractile | Ctgf | 3.2237E-21 | 0.525187 | 0.794 | 0.604 | 4.26914E-17 |
| Contractile | Eif5b | 1.22749E-10 | 0.523245 | 0.665 | 0.605 | 1.62556E-06 |
| Intermediate | Adam9 | 1.60264E-36 | 0.523145 | 0.468 | 0.219 | 2.12237E-32 |
| Rgs5_Contractile | Eif4a2 | 0.000528627 | 0.522187 | 0.628 | 0.781 | 1 |
| Rgs5_Contractile | Plekhg3 | 9.27326E-05 | 0.521822 | 0.135 | 0.076 | 1 |

| | | | | | | |
|---|---|---|---|---|---|---|
| Intermediate | Lmna | 1.69282E-35 | 0.521711 | 0.894 | 0.768 | 2.2418E-31 |
| Contractile | Hmgn5 | 3.29963E-07 | 0.521677 | 0.324 | 0.229 | 0.004369698 |
| Rgs5_Contractile | Pmepa1 | 0.001092488 | 0.521545 | 0.528 | 0.565 | 1 |
| Intermediate | Cygb | 1.09935E-31 | 0.521533 | 0.157 | 0.027 | 1.45586E-27 |
| Terminal | Chadl | 3.53056E-45 | 0.521139 | 0.252 | 0.055 | 4.67552E-41 |
| Rgs5_Contractile | Crispld2 | 0.003168099 | 0.521058 | 0.26 | 0.209 | 1 |
| Contractile | Prkcd | 2.21543E-16 | 0.521028 | 0.288 | 0.129 | 2.9339E-12 |
| Intermediate | Actg1 | 7.96944E-28 | 0.52097 | 0.974 | 0.964 | 1.05539E-23 |
| Intermediate | Pld3 | 1.5365E-29 | 0.52059 | 0.562 | 0.352 | 2.03478E-25 |
| Contractile | Myadm | 6.95162E-10 | 0.520195 | 0.465 | 0.343 | 9.20604E-06 |
| Intermediate | Col4a2 | 2.73471E-21 | 0.520165 | 0.603 | 0.412 | 3.62158E-17 |
| Contractile | Uqcr11 | 2.37574E-25 | 0.519817 | 0.894 | 0.839 | 3.14619E-21 |
| Contractile | Fkbp4 | 4.5132E-11 | 0.519664 | 0.479 | 0.358 | 5.97683E-07 |
| Intermediate | Thbd | 2.16346E-31 | 0.519537 | 0.266 | 0.09 | 2.86507E-27 |
| Contractile | Maf1 | 3.47993E-10 | 0.519502 | 0.435 | 0.311 | 4.60847E-06 |
| Terminal | Papss2 | 6.30831E-40 | 0.519217 | 0.27 | 0.074 | 8.3541E-36 |
| Intermediate | Nnmt | 1.72517E-24 | 0.518609 | 0.503 | 0.311 | 2.28464E-20 |
| Contractile | Tspan7 | 5.28274E-34 | 0.518325 | 0.176 | 0.026 | 6.99594E-30 |
| Intermediate | Pdpn | 6.68063E-36 | 0.517639 | 0.174 | 0.03 | 8.84716E-32 |
| Terminal | Nr4a1 | 6.59016E-28 | 0.517537 | 0.467 | 0.246 | 8.72735E-24 |
| Terminal | Slc38a3 | 6.65184E-75 | 0.517359 | 0.218 | 0.005 | 8.80904E-71 |
| Terminal | Tgfb1 | 2.08613E-36 | 0.516901 | 0.681 | 0.396 | 2.76266E-32 |
| Intermediate | Nme1 | 4.31705E-33 | 0.516805 | 0.784 | 0.597 | 5.71707E-29 |
| Rgs5_Contractile | Fgl2 | 4.1365E-05 | 0.516455 | 0.378 | 0.292 | 0.547796593 |
| Contractile | Ndufs5 | 8.12565E-21 | 0.516092 | 0.829 | 0.751 | 1.07608E-16 |
| Intermediate | Uap1l1 | 1.13336E-34 | 0.515849 | 0.36 | 0.148 | 1.50091E-30 |
| Contractile | Dynlrb1 | 1.30887E-20 | 0.515669 | 0.876 | 0.8 | 1.73334E-16 |
| Intermediate | Fgf7 | 1.1499E-29 | 0.514755 | 0.24 | 0.079 | 1.52281E-25 |
| Rgs5_Contractile | Fau | 1.26958E-16 | 0.514356 | 0.913 | 0.954 | 1.6813E-12 |
| Contractile | Pid1 | 2.96717E-37 | 0.514065 | 0.209 | 0.033 | 3.92943E-33 |
| Rgs5_Contractile | Cox6c | 1.1465E-17 | 0.513789 | 0.931 | 0.96 | 1.51831E-13 |
| Intermediate | Medag | 6.66627E-34 | 0.513297 | 0.248 | 0.074 | 8.82815E-30 |
| Contractile | Sorbs2 | 5.72999E-12 | 0.51246 | 0.3 | 0.163 | 7.58823E-08 |
| Intermediate | Myh9 | 8.36264E-32 | 0.512293 | 0.435 | 0.207 | 1.10746E-27 |
| Contractile | Lrrc17 | 2.88888E-08 | 0.512235 | 0.335 | 0.217 | 0.000382574 |
| Contractile | 1810037I1 | 2.30382E-16 | 0.51222 | 0.794 | 0.745 | 3.05094E-12 |
| Rgs5_Contractile | Prkcdbp | 1.60387E-06 | 0.511876 | 0.726 | 0.838 | 0.021240096 |
| Intermediate | Serpinb6a | 1.53787E-23 | 0.511656 | 0.682 | 0.52 | 2.0366E-19 |
| Contractile | Evi5 | 1.49189E-13 | 0.511266 | 0.362 | 0.208 | 1.97571E-09 |
| Intermediate | Ccdc109b | 2.1464E-39 | 0.5105 | 0.406 | 0.169 | 2.84248E-35 |
| Intermediate | Mfap2 | 1.43574E-30 | 0.510246 | 0.284 | 0.101 | 1.90135E-26 |
| Terminal | Rasd1 | 4.84058E-55 | 0.510212 | 0.3 | 0.061 | 6.41038E-51 |
| Contractile | Rbbp7 | 1.69118E-11 | 0.510091 | 0.6 | 0.5 | 2.23963E-07 |
| Intermediate | Mpp6 | 6.36219E-39 | 0.510029 | 0.376 | 0.147 | 8.42545E-35 |
| Contractile | Irak2 | 7.42447E-16 | 0.509997 | 0.288 | 0.127 | 9.83223E-12 |
| Intermediate | Rbp1 | 3.33089E-41 | 0.5088 | 0.84 | 0.562 | 4.4111E-37 |
| Intermediate | Fkbp10 | 3.58194E-28 | 0.508724 | 0.655 | 0.458 | 4.74356E-24 |
| Intermediate | Ppic | 1.11794E-33 | 0.507638 | 0.851 | 0.655 | 1.48049E-29 |
| Rgs5_Contractile | Gjc1 | 0.008924772 | 0.507324 | 0.146 | 0.107 | 1 |
| Terminal | Anxa8 | 2.10265E-42 | 0.507213 | 0.643 | 0.321 | 2.78454E-38 |
| Terminal | Ddit4l | 9.2629E-23 | 0.50659 | 0.374 | 0.19 | 1.22669E-18 |
| Intermediate | Bdh2 | 4.81215E-39 | 0.506524 | 0.22 | 0.048 | 6.37273E-35 |
| Intermediate | Fbln1 | 3.2361E-34 | 0.50616 | 0.196 | 0.042 | 4.28557E-30 |
| Intermediate | Itgb5 | 2.65256E-31 | 0.506005 | 0.842 | 0.65 | 3.51278E-27 |
| Terminal | Wwp2 | 1.39994E-23 | 0.505703 | 0.687 | 0.473 | 1.85394E-19 |
| Terminal | Serpinh1 | 1.06027E-25 | 0.505153 | 0.985 | 0.934 | 1.40411E-21 |
| Contractile | Osbpl3 | 6.58166E-17 | 0.505075 | 0.268 | 0.111 | 8.71609E-13 |
| Intermediate | Atp1b1 | 7.39028E-24 | 0.504715 | 0.199 | 0.065 | 9.78695E-20 |
| Contractile | Filip1 | 1.10204E-16 | 0.504573 | 0.253 | 0.101 | 1.45943E-12 |

| | | | | | | |
|---|---|---|---|---|---|---|
| Terminal | Ldlrad3 | 2.83664E-50 | 0.504141 | 0.374 | 0.112 | 3.75656E-46 |
| Intermediate | Pqlc3 | 4.10976E-29 | 0.503924 | 0.377 | 0.178 | 5.44256E-25 |
| Contractile | Rab3a | 1.25043E-14 | 0.503803 | 0.271 | 0.126 | 1.65594E-10 |
| Intermediate | Thy1 | 8.72215E-39 | 0.503508 | 0.182 | 0.03 | 1.15507E-34 |
| Terminal | Tuba1c | 7.35018E-19 | 0.503493 | 0.496 | 0.315 | 9.73384E-15 |
| Contractile | Pmepa1 | 4.32859E-16 | 0.503352 | 0.709 | 0.537 | 5.73235E-12 |
| Contractile | Acot9 | 3.23764E-11 | 0.502984 | 0.4 | 0.259 | 4.2876E-07 |
| Intermediate | Cd81 | 1.34893E-44 | 0.502878 | 0.98 | 0.957 | 1.78639E-40 |
| Intermediate | Gpx7 | 4.38245E-19 | 0.501441 | 0.543 | 0.385 | 5.80368E-15 |
| Intermediate | Ly6e | 6.18516E-27 | 0.50144 | 0.534 | 0.316 | 8.19101E-23 |
| Contractile | Efr3a | 1.87062E-07 | 0.500783 | 0.362 | 0.27 | 0.002477257 |
| Contractile | Hmgcs2 | 3.6914E-31 | 0.500699 | 0.274 | 0.069 | 4.88853E-27 |
| Contractile | Dazap2 | 1.18399E-14 | 0.500542 | 0.515 | 0.349 | 1.56796E-10 |
| Contractile | Vapb | 3.10463E-08 | 0.500308 | 0.432 | 0.341 | 0.000411147 |
| Contractile | Prkar1a | 3.23753E-13 | 0.499606 | 0.726 | 0.645 | 4.28747E-09 |
| Terminal | Ppp3ca | 3.2881E-42 | 0.499224 | 0.687 | 0.364 | 4.35443E-38 |
| Terminal | Ier5 | 1.24271E-33 | 0.499104 | 0.591 | 0.337 | 1.64573E-29 |
| Terminal | Ptpn2 | 2.52086E-34 | 0.49881 | 0.49 | 0.241 | 3.33837E-30 |
| Contractile | Slc38a2 | 7.67669E-14 | 0.497835 | 0.624 | 0.487 | 1.01662E-09 |
| Rgs5_Contractile | Cox4i1 | 1.98796E-12 | 0.49728 | 0.889 | 0.951 | 2.63266E-08 |
| Contractile | Ehd2 | 7.81382E-13 | 0.49642 | 0.5 | 0.34 | 1.03478E-08 |
| Contractile | Uqcrh | 4.72068E-24 | 0.496291 | 0.918 | 0.879 | 6.2516E-20 |
| Contractile | Bcar3 | 2.81232E-14 | 0.496144 | 0.203 | 0.078 | 3.72435E-10 |
| Intermediate | Bst2 | 6.51315E-26 | 0.496053 | 0.398 | 0.194 | 8.62536E-22 |
| Contractile | Fxyd7 | 6.52711E-19 | 0.495593 | 0.209 | 0.066 | 8.64385E-15 |
| Contractile | Gnas | 1.03653E-25 | 0.494772 | 0.974 | 0.941 | 1.37267E-21 |
| Terminal | Matn4 | 1.34832E-43 | 0.494455 | 0.167 | 0.016 | 1.78558E-39 |
| Contractile | Ndufs6 | 4.12075E-12 | 0.494396 | 0.694 | 0.63 | 5.45711E-08 |
| Contractile | Pcmtd1 | 5.16332E-11 | 0.493824 | 0.462 | 0.333 | 6.83779E-07 |
| Intermediate | Pi15 | 1.10479E-15 | 0.493578 | 0.54 | 0.374 | 1.46307E-11 |
| Contractile | Paqr6 | 1.80148E-25 | 0.493109 | 0.191 | 0.043 | 2.3857E-21 |
| Contractile | Klf9 | 1.19707E-10 | 0.492752 | 0.556 | 0.44 | 1.58528E-06 |
| Rgs5_Contractile | Mertk | 4.41084E-07 | 0.492539 | 0.122 | 0.051 | 0.005841282 |
| Contractile | Esyt2 | 6.86331E-18 | 0.492106 | 0.515 | 0.314 | 9.08908E-14 |
| Terminal | Dnajb1 | 9.58181E-17 | 0.4919 | 0.431 | 0.267 | 1.26892E-12 |
| Intermediate | Prr13 | 1.63688E-21 | 0.491221 | 0.549 | 0.374 | 2.16772E-17 |
| Terminal | Rnaset2a | 3.83799E-38 | 0.491105 | 0.617 | 0.331 | 5.08265E-34 |
| Contractile | Sun2 | 1.87098E-18 | 0.491086 | 0.329 | 0.149 | 2.47774E-14 |
| Contractile | Snta1 | 5.78301E-10 | 0.490738 | 0.321 | 0.199 | 7.65843E-06 |
| Terminal | Hbegf | 1.26474E-29 | 0.490601 | 0.515 | 0.287 | 1.67489E-25 |
| Intermediate | Wls | 4.73837E-29 | 0.490485 | 0.718 | 0.513 | 6.27502E-25 |
| Rgs5_Contractile | Slc25a23 | 0.002869223 | 0.490376 | 0.111 | 0.069 | 1 |
| Contractile | Park7 | 2.01628E-20 | 0.490002 | 0.891 | 0.805 | 2.67016E-16 |
| Contractile | Tcp1 | 7.6129E-11 | 0.4899 | 0.568 | 0.469 | 1.00818E-06 |
| Terminal | Fabp5 | 8.2136E-40 | 0.489549 | 0.271 | 0.072 | 1.08773E-35 |
| Contractile | Ndufc1 | 2.86777E-16 | 0.489474 | 0.829 | 0.762 | 3.79779E-12 |
| Contractile | Oaz2 | 4.50218E-11 | 0.489232 | 0.609 | 0.532 | 5.96224E-07 |
| Contractile | Uba2 | 5.42628E-11 | 0.489187 | 0.456 | 0.324 | 7.18602E-07 |
| Intermediate | Sh3bgrl3 | 8.07971E-29 | 0.488813 | 0.822 | 0.692 | 1.07E-24 |
| Contractile | Sptssa | 3.11999E-12 | 0.4883 | 0.641 | 0.555 | 4.1318E-08 |
| Contractile | Ybx1 | 2.83592E-29 | 0.488224 | 0.962 | 0.931 | 3.7556E-25 |
| Contractile | Ndufb2 | 1.83667E-15 | 0.487886 | 0.824 | 0.752 | 2.4323E-11 |
| Terminal | Fgfr1 | 8.59658E-27 | 0.487815 | 0.711 | 0.474 | 1.13845E-22 |
| Contractile | Mdh1 | 6.07882E-12 | 0.487706 | 0.768 | 0.684 | 8.05018E-08 |
| Contractile | Msra | 1.57634E-15 | 0.487543 | 0.2 | 0.073 | 2.08755E-11 |
| Intermediate | Plec | 7.79435E-30 | 0.487399 | 0.51 | 0.281 | 1.03221E-25 |
| Contractile | Cd151 | 6.49231E-15 | 0.487319 | 0.718 | 0.619 | 8.59777E-11 |
| Contractile | Scp2 | 3.30596E-13 | 0.48729 | 0.768 | 0.722 | 4.37808E-09 |
| Terminal | Hexa | 3.24099E-35 | 0.48688 | 0.894 | 0.65 | 4.29204E-31 |

| | | | | | | |
|---|---|---|---|---|---|---|
| Rgs5_Contractile | Naca | 1.7262E-08 | 0.486417 | 0.812 | 0.923 | 0.000228601 |
| Contractile | Dctn6 | 3.951E-11 | 0.486337 | 0.506 | 0.379 | 5.23231E-07 |
| Contractile | Ppp2r1a | 4.28948E-11 | 0.486215 | 0.503 | 0.38 | 5.68056E-07 |
| Contractile | Hnrnpa3 | 9.73349E-12 | 0.485609 | 0.721 | 0.673 | 1.28901E-07 |
| Contractile | Igf1r | 9.05681E-08 | 0.485156 | 0.324 | 0.22 | 0.001199394 |
| Contractile | Epn2 | 1.50373E-12 | 0.484979 | 0.303 | 0.158 | 1.99139E-08 |
| Contractile | 2010107E( | 4.90055E-17 | 0.484634 | 0.844 | 0.749 | 6.4898E-13 |
| Intermediate | Sri | 6.63992E-28 | 0.484415 | 0.683 | 0.48 | 8.79324E-24 |
| Contractile | Sntg2 | 4.78995E-32 | 0.484133 | 0.168 | 0.025 | 6.34333E-28 |
| Contractile | Smim12 | 3.69301E-12 | 0.483341 | 0.356 | 0.211 | 4.89065E-08 |
| Terminal | Golph3 | 1.66762E-34 | 0.481135 | 0.488 | 0.248 | 2.20843E-30 |
| Rgs5_Contractile | Itih4 | 1.84179E-09 | 0.480879 | 0.351 | 0.201 | 2.43908E-05 |
| Intermediate | Asah1 | 7.3959E-32 | 0.480647 | 0.578 | 0.33 | 9.79439E-28 |
| Contractile | Plat | 3.50285E-14 | 0.480509 | 0.324 | 0.164 | 4.63883E-10 |
| Contractile | Auts2 | 1.39616E-10 | 0.480412 | 0.503 | 0.38 | 1.84893E-06 |
| Contractile | Eln | 1.10828E-65 | 0.480273 | 0.95 | 0.527 | 1.46769E-61 |
| Rgs5_Contractile | Pfn1 | 3.3467E-14 | 0.479932 | 0.878 | 0.926 | 4.43203E-10 |
| Intermediate | Arhgdib | 3.19059E-28 | 0.479911 | 0.421 | 0.218 | 4.22529E-24 |
| Intermediate | Plvap | 6.79479E-22 | 0.47973 | 0.417 | 0.234 | 8.99834E-18 |
| Contractile | Rnf187 | 3.15484E-12 | 0.478713 | 0.659 | 0.567 | 4.17796E-08 |
| Contractile | Psmc6 | 4.35675E-07 | 0.478588 | 0.497 | 0.42 | 0.005769643 |
| Intermediate | Arpc3 | 5.50881E-30 | 0.478507 | 0.771 | 0.602 | 7.29531E-26 |
| Rgs5_Contractile | Hsp90ab1 | 2.58334E-14 | 0.478091 | 0.948 | 0.974 | 3.42112E-10 |
| Contractile | Mob4 | 8.85492E-12 | 0.477828 | 0.497 | 0.366 | 1.17266E-07 |
| Intermediate | Ltbp3 | 9.65513E-27 | 0.47781 | 0.673 | 0.481 | 1.27863E-22 |
| Contractile | Snrpb | 9.00051E-14 | 0.477754 | 0.8 | 0.73 | 1.19194E-09 |
| Contractile | Fry | 2.70503E-12 | 0.477512 | 0.391 | 0.237 | 3.58227E-08 |
| Intermediate | Ptgfrn | 1.68527E-34 | 0.477468 | 0.315 | 0.115 | 2.23181E-30 |
| Terminal | Ppp1r1b | 7.17431E-51 | 0.477067 | 0.244 | 0.039 | 9.50094E-47 |
| Rgs5_Contractile | Sdpr | 0.004318733 | 0.476756 | 0.118 | 0.075 | 1 |
| Contractile | Pxdc1 | 3.62457E-08 | 0.47599 | 0.371 | 0.256 | 0.000480002 |
| Contractile | D8Ertd738 | 3.39974E-18 | 0.475745 | 0.838 | 0.793 | 4.50227E-14 |
| Contractile | Cdc37 | 2.09463E-11 | 0.475564 | 0.659 | 0.577 | 2.77392E-07 |
| Contractile | Ubc | 1.4973E-18 | 0.474996 | 0.959 | 0.923 | 1.98288E-14 |
| Rgs5_Contractile | Zbtb20 | 1.22812E-06 | 0.4748 | 0.708 | 0.847 | 0.016263955 |
| Contractile | Fau | 5.04649E-27 | 0.474755 | 0.971 | 0.945 | 6.68307E-23 |
| Contractile | Rftn2 | 1.17009E-07 | 0.473126 | 0.285 | 0.184 | 0.001549549 |
| Intermediate | Rcn1 | 5.05247E-27 | 0.472961 | 0.636 | 0.434 | 6.69099E-23 |
| Contractile | Atp2b4 | 1.22684E-08 | 0.472893 | 0.376 | 0.262 | 0.00016247 |
| Intermediate | Matn2 | 8.33898E-20 | 0.472482 | 0.453 | 0.282 | 1.10433E-15 |
| Intermediate | Serpinb1a | 3.52456E-32 | 0.472453 | 0.228 | 0.065 | 4.66757E-28 |
| Contractile | Mcur1 | 1.9784E-15 | 0.47198 | 0.276 | 0.124 | 2.61999E-11 |
| Contractile | Cap2 | 1.44066E-33 | 0.471576 | 0.209 | 0.038 | 1.90786E-29 |
| Rgs5_Contractile | Hmgb1 | 2.77824E-11 | 0.471123 | 0.809 | 0.902 | 3.67922E-07 |
| Intermediate | Hgf | 2.63554E-39 | 0.470561 | 0.224 | 0.049 | 3.49025E-35 |
| Intermediate | Fads3 | 2.62707E-39 | 0.470514 | 0.45 | 0.196 | 3.47903E-35 |
| Contractile | Akr1b3 | 3.82136E-10 | 0.470463 | 0.497 | 0.38 | 5.06063E-06 |
| Contractile | Cltc | 7.77625E-08 | 0.470298 | 0.521 | 0.437 | 0.001029809 |
| Contractile | Atp6v1f | 2.55691E-18 | 0.46991 | 0.838 | 0.761 | 3.38612E-14 |
| Rgs5_Contractile | Parm1 | 6.71756E-06 | 0.46982 | 0.115 | 0.052 | 0.088960648 |
| Intermediate | Hspa5 | 4.08506E-19 | 0.469713 | 0.928 | 0.836 | 5.40984E-15 |
| Contractile | Rbfox2 | 1.05982E-06 | 0.469295 | 0.374 | 0.291 | 0.014035235 |
| Intermediate | Glipr1 | 1.36086E-20 | 0.469106 | 0.48 | 0.304 | 1.80219E-16 |
| Intermediate | Tmod3 | 2.17114E-29 | 0.468633 | 0.397 | 0.189 | 2.87524E-25 |
| Terminal | Mafb | 7.78496E-13 | 0.46857 | 0.208 | 0.102 | 1.03096E-08 |
| Contractile | Hspa8 | 2.49303E-24 | 0.468348 | 0.994 | 0.967 | 3.30152E-20 |
| Intermediate | Gm17455 | 2.28055E-15 | 0.46824 | 0.321 | 0.188 | 3.02014E-11 |
| Contractile | Dact3 | 1.61612E-16 | 0.467813 | 0.282 | 0.122 | 2.14023E-12 |
| Contractile | Unc5c | 2.37601E-27 | 0.467783 | 0.206 | 0.046 | 3.14654E-23 |

| | | | | | | |
|---|---|---|---|---|---|---|
| Intermediate | Tpm3 | 9.49533E-27 | 0.467391 | 0.659 | 0.47 | 1.25747E-22 |
| Rgs5_Contractile | Uqcrb | 1.32168E-11 | 0.466502 | 0.826 | 0.914 | 1.7503E-07 |
| Terminal | Cxcl14 | 5.62542E-16 | 0.466136 | 0.283 | 0.143 | 7.44975E-12 |
| Contractile | Polr2c | 3.63684E-07 | 0.466106 | 0.497 | 0.421 | 0.004816272 |
| Intermediate | Cyb5r3 | 7.90784E-37 | 0.465203 | 0.904 | 0.804 | 1.04724E-32 |
| Intermediate | Mxra7 | 3.46908E-28 | 0.464817 | 0.425 | 0.218 | 4.59411E-24 |
| Contractile | Atp5g3 | 8.02974E-12 | 0.464595 | 0.659 | 0.567 | 1.06338E-07 |
| Rgs5_Contractile | Cox7a2 | 2.94364E-06 | 0.464576 | 0.726 | 0.865 | 0.038982666 |
| Terminal | Cpxm2 | 1.64295E-49 | 0.464304 | 0.213 | 0.028 | 2.17576E-45 |
| Contractile | Vdac2 | 2.50625E-11 | 0.463995 | 0.653 | 0.561 | 3.31903E-07 |
| Contractile | Gadd45g | 4.37273E-05 | 0.463991 | 0.524 | 0.475 | 0.579080896 |
| Contractile | Mif4gd | 8.30591E-09 | 0.463733 | 0.274 | 0.164 | 0.000109995 |
| Intermediate | Calr | 1.13906E-31 | 0.463233 | 0.903 | 0.796 | 1.50846E-27 |
| Contractile | Rhob | 3.36185E-11 | 0.463183 | 0.488 | 0.341 | 4.4521E-07 |
| Contractile | Aldoa | 3.3689E-23 | 0.463073 | 0.944 | 0.888 | 4.46143E-19 |
| Contractile | Hmgb1 | 1.91011E-21 | 0.462821 | 0.95 | 0.881 | 2.52956E-17 |
| Contractile | Sfrp2 | 8.19691E-05 | 0.46262 | 0.441 | 0.332 | 1 |
| Rgs5_Contractile | Cep170b | 0.002856408 | 0.46258 | 0.132 | 0.087 | 1 |
| Rgs5_Contractile | Clic4 | 5.71039E-07 | 0.462331 | 0.674 | 0.779 | 0.007562264 |
| Intermediate | Ano1 | 1.02736E-30 | 0.462315 | 0.277 | 0.096 | 1.36053E-26 |
| Rgs5_Contractile | Gnas | 4.70227E-13 | 0.462149 | 0.892 | 0.952 | 6.22721E-09 |
| Rgs5_Contractile | Sema3c | 0.000786776 | 0.462129 | 0.167 | 0.109 | 1 |
| Intermediate | Arf5 | 4.31178E-30 | 0.461907 | 0.812 | 0.661 | 5.71009E-26 |
| Intermediate | Lrp1 | 1.98274E-30 | 0.461904 | 0.849 | 0.668 | 2.62574E-26 |
| Rgs5_Contractile | Ubc | 1.21723E-07 | 0.461399 | 0.865 | 0.936 | 0.001611977 |
| Contractile | Cyb5a | 1.34626E-12 | 0.461307 | 0.768 | 0.702 | 1.78286E-08 |
| Contractile | Ncald | 1.51272E-33 | 0.461184 | 0.188 | 0.03 | 2.00329E-29 |
| Terminal | Cilp2 | 1.03727E-22 | 0.46038 | 0.174 | 0.053 | 1.37365E-18 |
| Contractile | Bicd1 | 1.53414E-22 | 0.460327 | 0.244 | 0.076 | 2.03166E-18 |
| Contractile | Ndufs7 | 5.32622E-13 | 0.459992 | 0.7 | 0.627 | 7.05352E-09 |
| Contractile | Tbxa2r | 4.18125E-25 | 0.459326 | 0.206 | 0.05 | 5.53723E-21 |
| Contractile | Cystm1 | 2.43816E-16 | 0.459213 | 0.329 | 0.154 | 3.22886E-12 |
| Terminal | Serpina3m | 6.37441E-40 | 0.459016 | 0.185 | 0.028 | 8.44163E-36 |
| Rgs5_Contractile | Tbxa2r | 0.000385586 | 0.458808 | 0.118 | 0.066 | 1 |
| Rgs5_Contractile | B3galt1 | 8.67272E-05 | 0.458696 | 0.104 | 0.051 | 1 |
| Contractile | Laptm4b | 4.5916E-08 | 0.458659 | 0.276 | 0.173 | 0.000608066 |
| Contractile | Wdfy1 | 3.07223E-17 | 0.458375 | 0.274 | 0.11 | 4.06855E-13 |
| Intermediate | Nsg1 | 4.98542E-32 | 0.458294 | 0.202 | 0.051 | 6.60219E-28 |
| Intermediate | Pdgfra | 6.10078E-29 | 0.457888 | 0.314 | 0.128 | 8.07926E-25 |
| Intermediate | S100a6 | 2.18366E-32 | 0.457805 | 0.982 | 0.964 | 2.89183E-28 |
| Terminal | Sox6 | 3.70758E-48 | 0.457586 | 0.298 | 0.073 | 4.90995E-44 |
| Intermediate | P2rx4 | 5.0555E-24 | 0.457556 | 0.433 | 0.244 | 6.69499E-20 |
| Contractile | Rheb | 6.46086E-15 | 0.457254 | 0.818 | 0.774 | 8.55611E-11 |
| Contractile | Fkbp1a | 8.10665E-19 | 0.457105 | 0.909 | 0.828 | 1.07356E-14 |
| Contractile | Smim5 | 1.38878E-11 | 0.456494 | 0.259 | 0.128 | 1.83917E-07 |
| Intermediate | Lamp2 | 5.18628E-24 | 0.456455 | 0.786 | 0.624 | 6.86819E-20 |
| Intermediate | Tmem140 | 2.17156E-26 | 0.456289 | 0.397 | 0.201 | 2.8758E-22 |
| Intermediate | Plaur | 2.97209E-21 | 0.456129 | 0.286 | 0.132 | 3.93594E-17 |
| Contractile | Idh2 | 5.63285E-08 | 0.455916 | 0.471 | 0.382 | 0.000745958 |
| Terminal | Emp1 | 2.45925E-30 | 0.455862 | 0.663 | 0.377 | 3.25679E-26 |
| Contractile | Kif1c | 8.11133E-12 | 0.455797 | 0.421 | 0.271 | 1.07418E-07 |
| Contractile | Ppp1r12c | 5.16853E-10 | 0.455658 | 0.271 | 0.15 | 6.84468E-06 |
| Intermediate | Pdia6 | 2.85217E-22 | 0.455455 | 0.818 | 0.665 | 3.77713E-18 |
| Terminal | Prnp | 3.6952E-26 | 0.455121 | 0.639 | 0.406 | 4.89356E-22 |
| Contractile | Ntn1 | 1.98301E-15 | 0.454838 | 0.394 | 0.211 | 2.6261E-11 |
| Contractile | Smim14 | 5.35274E-19 | 0.454649 | 0.9 | 0.838 | 7.08863E-15 |
| Intermediate | Eif5a | 6.1885E-34 | 0.454607 | 0.914 | 0.837 | 8.19543E-30 |
| Contractile | Mapre1 | 1.23649E-09 | 0.454564 | 0.55 | 0.471 | 1.63748E-05 |
| Contractile | Hmgn1 | 3.49314E-15 | 0.454483 | 0.868 | 0.804 | 4.62597E-11 |

| | | | | | | |
|---|---|---|---|---|---|---|
| Contractile | Cdh13 | 2.05268E-13 | 0.454434 | 0.606 | 0.429 | 2.71836E-09 |
| Contractile | Pttg1 | 3.84219E-07 | 0.454079 | 0.532 | 0.462 | 0.005088212 |
| Rgs5_Contractile | Ddx5 | 7.19175E-09 | 0.454053 | 0.844 | 0.931 | 9.52403E-05 |
| Intermediate | Pdia3 | 1.21528E-27 | 0.453985 | 0.905 | 0.804 | 1.60939E-23 |
| Contractile | Chchd2 | 1.605E-25 | 0.453877 | 0.965 | 0.936 | 2.1255E-21 |
| Intermediate | Map4k4 | 4.09747E-19 | 0.453835 | 0.408 | 0.237 | 5.42628E-15 |
| Contractile | Cnbp | 2.14684E-12 | 0.45345 | 0.838 | 0.784 | 2.84307E-08 |
| Terminal | Smim3 | 4.68057E-28 | 0.45306 | 0.448 | 0.23 | 6.19848E-24 |
| Contractile | Tmcc3 | 9.47258E-25 | 0.453016 | 0.2 | 0.048 | 1.25445E-20 |
| Intermediate | Sep15 | 4.12603E-41 | 0.452886 | 0.938 | 0.877 | 5.4641E-37 |
| Contractile | Cisd3 | 1.82973E-08 | 0.452367 | 0.321 | 0.213 | 0.000242311 |
| Intermediate | Bdnf | 1.32897E-28 | 0.452173 | 0.311 | 0.125 | 1.75995E-24 |
| Contractile | Lrrc58 | 2.36408E-10 | 0.451498 | 0.668 | 0.599 | 3.13075E-06 |
| Intermediate | Ccnd1 | 1.66647E-38 | 0.451457 | 0.171 | 0.024 | 2.20691E-34 |
| Contractile | Flnc | 5.65483E-20 | 0.451297 | 0.244 | 0.084 | 7.48869E-16 |
| Contractile | Nav2 | 8.23827E-23 | 0.451249 | 0.244 | 0.074 | 1.09099E-18 |
| Intermediate | Cyp1b1 | 4.21333E-27 | 0.450852 | 0.247 | 0.083 | 5.57971E-23 |
| Contractile | Dpysl3 | 9.76581E-11 | 0.450644 | 0.603 | 0.481 | 1.29329E-06 |
| Contractile | Asap2 | 9.41981E-10 | 0.45054 | 0.344 | 0.215 | 1.24747E-05 |
| Contractile | Map1lc3b | 1.0839E-20 | 0.450466 | 0.953 | 0.885 | 1.43541E-16 |
| Contractile | Polr2f | 3.71897E-11 | 0.449708 | 0.691 | 0.611 | 4.92504E-07 |
| Intermediate | Kmt2e | 2.92618E-21 | 0.449479 | 0.635 | 0.45 | 3.87514E-17 |
| Contractile | Ndufa13 | 8.99142E-22 | 0.449465 | 0.938 | 0.901 | 1.19073E-17 |
| Contractile | Pcgf5 | 3.79327E-15 | 0.449273 | 0.265 | 0.115 | 5.02342E-11 |
| Terminal | Gm8730 | 1.66227E-33 | 0.449016 | 0.969 | 0.85 | 2.20135E-29 |
| Contractile | Mtus1 | 1.71862E-15 | 0.448427 | 0.253 | 0.106 | 2.27597E-11 |
| Contractile | Sdhc | 3.38313E-12 | 0.448227 | 0.5 | 0.353 | 4.48028E-08 |
| Contractile | Ap1s2 | 5.71527E-10 | 0.448183 | 0.241 | 0.127 | 7.56873E-06 |
| Terminal | Anxa4 | 8.33135E-28 | 0.447972 | 0.627 | 0.388 | 1.10332E-23 |
| Rgs5_Contractile | Ndufa6 | 0.00021728 | 0.447716 | 0.656 | 0.82 | 1 |
| Contractile | Fabp4 | 4.11294E-13 | 0.447488 | 0.65 | 0.49 | 5.44677E-09 |
| Terminal | Cd14 | 5.20146E-37 | 0.447355 | 0.298 | 0.094 | 6.8883E-33 |
| Contractile | Eef2 | 3.86809E-21 | 0.447194 | 0.938 | 0.88 | 5.12251E-17 |
| Contractile | Cox5b | 1.7061E-14 | 0.446951 | 0.844 | 0.804 | 2.25938E-10 |
| Contractile | Itga3 | 1.32463E-19 | 0.44692 | 0.185 | 0.051 | 1.7542E-15 |
| Terminal | Tmem176a | 2.44866E-32 | 0.4469 | 0.806 | 0.522 | 3.24277E-28 |
| Intermediate | Baiap2 | 7.84493E-35 | 0.446657 | 0.251 | 0.071 | 1.0389E-30 |
| Intermediate | Ctgf | 0.004027657 | 0.446444 | 0.662 | 0.612 | 1 |
| Contractile | Slc24a3 | 2.52136E-31 | 0.446367 | 0.191 | 0.034 | 3.33903E-27 |
| Contractile | Klf15 | 1.81316E-11 | 0.446018 | 0.279 | 0.147 | 2.40116E-07 |
| Contractile | Usmg5 | 9.17181E-15 | 0.445786 | 0.85 | 0.788 | 1.21462E-10 |
| Contractile | Cox6c | 2.48148E-30 | 0.445638 | 0.982 | 0.952 | 3.28622E-26 |
| Rgs5_Contractile | Fam126b | 0.004000411 | 0.445567 | 0.135 | 0.092 | 1 |
| Contractile | Hspa2 | 1.15841E-07 | 0.44541 | 0.344 | 0.237 | 0.001534083 |
| Contractile | Hmgn3 | 1.41458E-09 | 0.445045 | 0.621 | 0.534 | 1.87333E-05 |
| Contractile | Arhgef17 | 1.18872E-10 | 0.444492 | 0.365 | 0.227 | 1.57422E-06 |
| Intermediate | Adamtsl5 | 8.274E-37 | 0.444459 | 0.407 | 0.161 | 1.09573E-32 |
| Intermediate | Slc6a6 | 2.46148E-15 | 0.444423 | 0.522 | 0.366 | 3.25974E-11 |
| Contractile | Rgs7bp | 1.30775E-30 | 0.444396 | 0.215 | 0.044 | 1.73185E-26 |
| Terminal | Bmp2 | 2.22359E-24 | 0.44435 | 0.242 | 0.086 | 2.9447E-20 |
| Rgs5_Contractile | Rpgrip1 | 0.001526539 | 0.443909 | 0.115 | 0.068 | 1 |
| Contractile | Rtn3 | 7.53855E-08 | 0.443499 | 0.474 | 0.384 | 0.00099833 |
| Contractile | Mtss1 | 1.75882E-12 | 0.443099 | 0.194 | 0.079 | 2.3292E-08 |
| Rgs5_Contractile | Pygm | 0.003490318 | 0.442641 | 0.132 | 0.087 | 1 |
| Contractile | Tmed4 | 8.69837E-05 | 0.442598 | 0.394 | 0.337 | 1 |
| Contractile | Tax1bp3 | 7.98078E-09 | 0.442462 | 0.521 | 0.421 | 0.000105689 |
| Terminal | Col6a2 | 1.28132E-10 | 0.442268 | 0.764 | 0.611 | 1.69685E-06 |
| Intermediate | Glmp | 4.36182E-25 | 0.44213 | 0.651 | 0.445 | 5.77635E-21 |
| Intermediate | Pdgfa | 1.92479E-25 | 0.441885 | 0.757 | 0.586 | 2.549E-21 |

| | | | | | | |
|---|---|---|---|---|---|---|
| Contractile | Msn | 1.52968E-11 | 0.441837 | 0.735 | 0.658 | 2.02575E-07 |
| Contractile | Snap23 | 3.38984E-08 | 0.441679 | 0.385 | 0.278 | 0.000448917 |
| Contractile | Spag7 | 1.70931E-08 | 0.44166 | 0.344 | 0.235 | 0.000226364 |
| Intermediate | Mfap5 | 5.34487E-19 | 0.441578 | 0.396 | 0.212 | 7.07821E-15 |
| Contractile | Josd2 | 1.4277E-05 | 0.441125 | 0.412 | 0.349 | 0.189070544 |
| Contractile | Eif3f | 7.61119E-14 | 0.441018 | 0.794 | 0.739 | 1.00795E-09 |
| Rgs5_Contractile | Cd200 | 1.79038E-05 | 0.440698 | 0.795 | 0.873 | 0.237100449 |
| Contractile | Ech1 | 9.67145E-10 | 0.440541 | 0.506 | 0.404 | 1.28079E-05 |
| Intermediate | Ass1 | 1.1097E-16 | 0.439634 | 0.315 | 0.174 | 1.46957E-12 |
| Rgs5_Contractile | Atp5d | 0.005830058 | 0.439372 | 0.601 | 0.77 | 1 |
| Contractile | Kcnj8 | 4.51344E-18 | 0.439355 | 0.271 | 0.107 | 5.97715E-14 |
| Contractile | Raly | 1.49983E-09 | 0.439294 | 0.574 | 0.464 | 1.98623E-05 |
| Contractile | Cfl1 | 1.51471E-19 | 0.439043 | 0.935 | 0.858 | 2.00593E-15 |
| Intermediate | Bax | 1.49148E-19 | 0.438735 | 0.673 | 0.509 | 1.97516E-15 |
| Contractile | Prkcdbp | 4.95291E-16 | 0.43858 | 0.871 | 0.817 | 6.55914E-12 |
| Rgs5_Contractile | Atp5j2 | 3.37594E-06 | 0.438497 | 0.747 | 0.887 | 0.044707572 |
| Rgs5_Contractile | Eef2 | 1.86848E-08 | 0.438487 | 0.826 | 0.897 | 0.000247443 |
| Contractile | Anapc7 | 1.70909E-10 | 0.43718 | 0.321 | 0.189 | 2.26335E-06 |
| Intermediate | Alcam | 8.83526E-20 | 0.437013 | 0.481 | 0.3 | 1.17005E-15 |
| Terminal | G0s2 | 1.87994E-19 | 0.436748 | 0.104 | 0.02 | 2.48961E-15 |
| Rgs5_Contractile | Lgals1 | 9.21509E-15 | 0.436704 | 0.965 | 0.946 | 1.22035E-10 |
| Contractile | Pygb | 2.5216E-10 | 0.436559 | 0.244 | 0.127 | 3.33935E-06 |
| Intermediate | Rftn1 | 1.04083E-52 | 0.436413 | 0.196 | 0.017 | 1.37837E-48 |
| Contractile | Col4a1 | 5.47134E-20 | 0.436161 | 0.644 | 0.416 | 7.2457E-16 |
| Contractile | Atp5d | 5.64271E-15 | 0.436066 | 0.809 | 0.74 | 7.47265E-11 |
| Contractile | Pigyl | 2.15678E-08 | 0.436005 | 0.485 | 0.393 | 0.000285622 |
| Rgs5_Contractile | Asb2 | 1.41139E-06 | 0.435614 | 0.142 | 0.066 | 0.018691044 |
| Contractile | Ube2h | 1.10815E-06 | 0.435477 | 0.306 | 0.214 | 0.014675258 |
| Contractile | Pam | 4.86733E-30 | 0.43518 | 0.929 | 0.86 | 6.4458E-26 |
| Contractile | Bdh1 | 2.01222E-33 | 0.435115 | 0.138 | 0.014 | 2.66478E-29 |
| Contractile | Eef1d | 7.38346E-13 | 0.434853 | 0.8 | 0.75 | 9.77792E-09 |
| Intermediate | Gm26771 | 2.54574E-31 | 0.434817 | 0.174 | 0.035 | 3.37132E-27 |
| Contractile | Pecam1 | 1.30227E-09 | 0.434805 | 0.206 | 0.097 | 1.7246E-05 |
| Contractile | Sept7 | 1.32794E-12 | 0.434477 | 0.791 | 0.737 | 1.75859E-08 |
| Intermediate | Il4ra | 3.06473E-39 | 0.433658 | 0.217 | 0.046 | 4.05862E-35 |
| Contractile | Uqcrc2 | 2.94363E-06 | 0.433236 | 0.444 | 0.384 | 0.038982444 |
| Contractile | Eif3e | 2.60721E-10 | 0.433108 | 0.709 | 0.679 | 3.45273E-06 |
| Terminal | Bhlhe41 | 5.48639E-26 | 0.432492 | 0.229 | 0.076 | 7.26563E-22 |
| Contractile | Sept6 | 6.80021E-13 | 0.432404 | 0.218 | 0.093 | 9.00552E-09 |
| Contractile | Nfia | 2.35832E-13 | 0.432244 | 0.626 | 0.486 | 3.12312E-09 |
| Contractile | Snx17 | 1.11252E-06 | 0.431969 | 0.447 | 0.377 | 0.014733062 |
| Contractile | Capza2 | 5.21739E-10 | 0.430925 | 0.662 | 0.579 | 6.90939E-06 |
| Contractile | Setd3 | 6.94313E-09 | 0.430868 | 0.368 | 0.256 | 9.19479E-05 |
| Contractile | Anxa3 | 1.13837E-09 | 0.430768 | 0.55 | 0.437 | 1.50755E-05 |
| Terminal | Islr | 7.68626E-23 | 0.430003 | 0.766 | 0.537 | 1.01789E-18 |
| Intermediate | Inhba | 1.36431E-10 | 0.429521 | 0.347 | 0.218 | 1.80675E-06 |
| Intermediate | Myl12b | 3.33014E-23 | 0.429271 | 0.545 | 0.347 | 4.41011E-19 |
| Contractile | Slc25a23 | 2.93084E-22 | 0.428721 | 0.2 | 0.053 | 3.88131E-18 |
| Intermediate | Serpinb6b | 1.88318E-26 | 0.428524 | 0.327 | 0.136 | 2.49389E-22 |
| Contractile | Tomm22 | 5.35161E-07 | 0.428523 | 0.556 | 0.49 | 0.007087132 |
| Contractile | Adck3 | 2.77126E-12 | 0.428468 | 0.215 | 0.095 | 3.66998E-08 |
| Contractile | Foxn3 | 1.30976E-12 | 0.428356 | 0.335 | 0.186 | 1.73451E-08 |
| Contractile | Ywhaq | 2.3621E-14 | 0.427568 | 0.859 | 0.791 | 3.12813E-10 |
| Contractile | Glo1 | 8.08733E-09 | 0.427481 | 0.565 | 0.478 | 0.0001071 |
| Contractile | Il17b | 1.78681E-21 | 0.426698 | 0.159 | 0.036 | 2.36627E-17 |
| Intermediate | Prdx5 | 3.74937E-21 | 0.426652 | 0.834 | 0.752 | 4.96528E-17 |
| Terminal | Mpzl1 | 4.41148E-37 | 0.426626 | 0.436 | 0.189 | 5.84213E-33 |
| Contractile | Dpm1 | 0.00014461 | 0.426477 | 0.379 | 0.328 | 1 |
| Contractile | Cdkl5 | 6.00861E-22 | 0.426459 | 0.168 | 0.039 | 7.9572E-18 |

| | | | | | | |
|---|---|---|---|---|---|---|
| Contractile | Grb14 | 1.41973E-12 | 0.425947 | 0.312 | 0.165 | 1.88015E-08 |
| Intermediate | Ace | 5.63968E-15 | 0.425661 | 0.396 | 0.244 | 7.46863E-11 |
| Contractile | Cdk2ap2 | 3.58842E-08 | 0.42522 | 0.571 | 0.501 | 0.000475214 |
| Intermediate | Tubb3 | 6.88743E-38 | 0.425164 | 0.2 | 0.038 | 9.12102E-34 |
| Contractile | Gpsm2 | 1.40325E-12 | 0.424648 | 0.232 | 0.104 | 1.85832E-08 |
| Contractile | Gm9843 | 4.51883E-14 | 0.424628 | 0.844 | 0.856 | 5.98429E-10 |
| Contractile | Atpif1 | 3.29787E-16 | 0.424626 | 0.841 | 0.783 | 4.36737E-12 |
| Contractile | Atp5a1 | 7.09528E-12 | 0.424007 | 0.797 | 0.729 | 9.39628E-08 |
| Contractile | Nacc2 | 4.55984E-10 | 0.423862 | 0.391 | 0.262 | 6.03859E-06 |
| Rgs5_Contractile | 2010107E( | 0.000182729 | 0.423614 | 0.639 | 0.779 | 1 |
| Contractile | Gabarapl1 | 2.53035E-09 | 0.423537 | 0.5 | 0.388 | 3.35094E-05 |
| Contractile | Anp32e | 3.9123E-08 | 0.423505 | 0.388 | 0.284 | 0.000518106 |
| Contractile | Sept2 | 7.60442E-07 | 0.423422 | 0.482 | 0.403 | 0.010070535 |
| Contractile | Slc4a3 | 1.69937E-13 | 0.423422 | 0.191 | 0.073 | 2.25047E-09 |
| Contractile | Atp6v1a | 7.53938E-07 | 0.423197 | 0.409 | 0.318 | 0.009984401 |
| Contractile | Coq2 | 1.07949E-05 | 0.422494 | 0.312 | 0.235 | 0.142956879 |
| Contractile | Dynlt3 | 3.09796E-08 | 0.421635 | 0.529 | 0.434 | 0.000410262 |
| Terminal | Ssr4 | 9.90992E-26 | 0.421471 | 0.885 | 0.725 | 1.31237E-21 |
| Intermediate | Lman1 | 3.35842E-19 | 0.421254 | 0.654 | 0.484 | 4.44756E-15 |
| Terminal | Fgfr3 | 3.31848E-59 | 0.421202 | 0.19 | 0.009 | 4.39467E-55 |
| Intermediate | Arhgdia | 8.05327E-27 | 0.420917 | 0.798 | 0.65 | 1.06649E-22 |
| Rgs5_Contractile | Itgb5 | 0.003606117 | 0.420753 | 0.604 | 0.737 | 1 |
| Rgs5_Contractile | Figf | 0.00530662 | 0.420558 | 0.104 | 0.065 | 1 |
| Contractile | Cers4 | 3.08519E-14 | 0.420285 | 0.232 | 0.096 | 4.08572E-10 |
| Terminal | Ehd3 | 1.60684E-34 | 0.419974 | 0.284 | 0.094 | 2.12794E-30 |
| Contractile | Ndufa7 | 6.79E-14 | 0.419851 | 0.874 | 0.809 | 8.99199E-10 |
| Intermediate | Prdx2 | 5.87313E-28 | 0.419546 | 0.834 | 0.727 | 7.77778E-24 |
| Terminal | Metrnl | 1.72076E-26 | 0.419523 | 0.596 | 0.342 | 2.2788E-22 |
| Contractile | Hsp90ab1 | 5.72702E-26 | 0.419508 | 0.976 | 0.97 | 7.58429E-22 |
| Intermediate | Arf4 | 2.77645E-25 | 0.418979 | 0.853 | 0.729 | 3.67686E-21 |
| Intermediate | Mgst3 | 1.13847E-22 | 0.418899 | 0.419 | 0.226 | 1.50767E-18 |
| Terminal | Itm2b | 4.39704E-22 | 0.418825 | 0.997 | 0.986 | 5.823E-18 |
| Intermediate | Sigmar1 | 1.60574E-27 | 0.418353 | 0.414 | 0.208 | 2.12648E-23 |
| Contractile | Crtc3 | 6.88723E-09 | 0.417993 | 0.294 | 0.178 | 9.12075E-05 |
| Contractile | Cap1 | 2.16898E-12 | 0.417392 | 0.312 | 0.159 | 2.87238E-08 |
| Contractile | Smc6 | 4.39751E-05 | 0.417149 | 0.432 | 0.377 | 0.582361706 |
| Rgs5_Contractile | Unc5c | 0.009916293 | 0.417126 | 0.101 | 0.065 | 1 |
| Contractile | Ndufs3 | 1.499E-08 | 0.416932 | 0.597 | 0.529 | 0.000198512 |
| Intermediate | Loxl3 | 2.83408E-19 | 0.416717 | 0.44 | 0.268 | 3.75318E-15 |
| Contractile | Inpp4a | 7.01371E-13 | 0.416199 | 0.268 | 0.127 | 9.28825E-09 |
| Contractile | B3galnt2 | 6.54448E-16 | 0.416169 | 0.238 | 0.093 | 8.66686E-12 |
| Terminal | Adgrg6 | 9.53119E-48 | 0.41588 | 0.211 | 0.031 | 1.26222E-43 |
| Rgs5_Contractile | S100a11 | 5.7961E-15 | 0.41577 | 0.934 | 0.944 | 7.67577E-11 |
| Contractile | Prss23 | 2.57153E-11 | 0.415541 | 0.456 | 0.268 | 3.40548E-07 |
| Intermediate | Myof | 7.31252E-25 | 0.415435 | 0.365 | 0.173 | 9.68397E-21 |
| Contractile | Gyg | 4.33747E-09 | 0.415409 | 0.321 | 0.201 | 5.74411E-05 |
| Contractile | Snhg18 | 1.95386E-11 | 0.41534 | 0.856 | 0.821 | 2.58749E-07 |
| Contractile | Tpd52l1 | 3.30845E-18 | 0.415299 | 0.182 | 0.054 | 4.38138E-14 |
| Contractile | S100a11 | 6.91359E-24 | 0.414817 | 0.979 | 0.937 | 9.15566E-20 |
| Contractile | Jam3 | 6.4844E-07 | 0.414594 | 0.403 | 0.314 | 0.008587296 |
| Intermediate | Shisa5 | 1.40551E-22 | 0.414442 | 0.758 | 0.587 | 1.86132E-18 |
| Intermediate | Eif4ebp1 | 6.69888E-24 | 0.413482 | 0.553 | 0.336 | 8.87132E-20 |
| Contractile | Aldh7a1 | 5.60056E-06 | 0.413186 | 0.347 | 0.263 | 0.074168232 |
| Terminal | Slc35a2 | 3.75682E-31 | 0.413015 | 0.462 | 0.235 | 4.97516E-27 |
| Contractile | Tnxb | 6.62585E-17 | 0.412997 | 0.259 | 0.099 | 8.77462E-13 |
| Contractile | Parm1 | 6.83986E-22 | 0.412943 | 0.174 | 0.041 | 9.05803E-18 |
| Rgs5_Contractile | Tmcc3 | 0.000496224 | 0.412681 | 0.115 | 0.063 | 1 |
| Intermediate | Atp6v0b | 3.87242E-23 | 0.412549 | 0.812 | 0.667 | 5.12825E-19 |
| Terminal | Haus8 | 1.58409E-23 | 0.412314 | 0.213 | 0.073 | 2.09781E-19 |

| | | | | | | |
|---|---|---|---|---|---|---|
| Terminal | Sp7 | 6.18883E-53 | 0.412121 | 0.178 | 0.011 | 8.19587E-49 |
| Terminal | Hes1 | 3.78613E-16 | 0.412026 | 0.585 | 0.405 | 5.01397E-12 |
| Contractile | Rab18 | 1.85866E-09 | 0.411866 | 0.682 | 0.625 | 2.46143E-05 |
| Intermediate | Cst6 | 6.57891E-07 | 0.41177 | 0.3 | 0.212 | 0.008712457 |
| Contractile | Xk | 3.69975E-36 | 0.411603 | 0.162 | 0.019 | 4.89958E-32 |
| Contractile | Fryl | 2.49714E-15 | 0.411363 | 0.25 | 0.104 | 3.30696E-11 |
| Contractile | Mpc1 | 1.4347E-10 | 0.411136 | 0.765 | 0.682 | 1.89998E-06 |
| Terminal | 1700025G | 4.3935E-48 | 0.410674 | 0.307 | 0.077 | 5.81831E-44 |
| Terminal | Runx1 | 5.74878E-26 | 0.410625 | 0.522 | 0.294 | 7.61311E-22 |
| Contractile | Paip2 | 1.60546E-09 | 0.410573 | 0.738 | 0.708 | 2.12611E-05 |
| Contractile | Med28 | 4.53358E-11 | 0.410549 | 0.712 | 0.634 | 6.00381E-07 |
| Terminal | Nrn1 | 1.0459E-41 | 0.41054 | 0.38 | 0.13 | 1.38509E-37 |
| Terminal | Fxyd6 | 7.29275E-34 | 0.410132 | 0.209 | 0.049 | 9.65779E-30 |
| Contractile | Wtip | 3.98561E-15 | 0.410058 | 0.268 | 0.118 | 5.27814E-11 |
| Intermediate | Litaf | 1.45975E-22 | 0.409963 | 0.461 | 0.272 | 1.93314E-18 |
| Contractile | Nmt2 | 5.04946E-11 | 0.409904 | 0.297 | 0.163 | 6.687E-07 |
| Contractile | Gm26632 | 5.38767E-24 | 0.409732 | 0.159 | 0.031 | 7.13489E-20 |
| Contractile | 0610011F( | 1.8356E-05 | 0.40971 | 0.321 | 0.244 | 0.243088312 |
| Contractile | Hnrnpl | 3.61132E-07 | 0.409018 | 0.424 | 0.338 | 0.004782478 |
| Contractile | Comt | 1.85221E-07 | 0.408751 | 0.55 | 0.463 | 0.002452879 |
| Terminal | Hist1h1c | 1.1105E-12 | 0.408297 | 0.507 | 0.353 | 1.47064E-08 |
| Contractile | Hibadh | 1.12414E-05 | 0.408266 | 0.406 | 0.337 | 0.148869992 |
| Terminal | S100a1 | 4.03653E-23 | 0.408101 | 0.869 | 0.714 | 5.34558E-19 |
| Intermediate | Canx | 1.83084E-23 | 0.407643 | 0.744 | 0.569 | 2.42458E-19 |
| Contractile | Smco4 | 6.07318E-06 | 0.407475 | 0.279 | 0.198 | 0.080427133 |
| Terminal | Egr2 | 4.51805E-39 | 0.407416 | 0.196 | 0.034 | 5.98325E-35 |
| Contractile | 2900026A( | 7.99524E-16 | 0.407161 | 0.306 | 0.138 | 1.05881E-11 |
| Intermediate | Ifngr1 | 2.94054E-23 | 0.407156 | 0.471 | 0.271 | 3.89415E-19 |
| Contractile | Psma1 | 1.76637E-08 | 0.407016 | 0.606 | 0.537 | 0.00023392 |
| Contractile | Lamtor5 | 9.52365E-08 | 0.406829 | 0.665 | 0.61 | 0.001261218 |
| Contractile | Myo1c | 2.05756E-07 | 0.406769 | 0.421 | 0.315 | 0.002724828 |
| Contractile | Galnt16 | 3.37122E-33 | 0.406167 | 0.174 | 0.025 | 4.4645E-29 |
| Contractile | Hbb-bs | 1.97155E-17 | 0.406123 | 0.226 | 0.08 | 2.61092E-13 |
| Terminal | Tfrc | 3.97834E-35 | 0.405906 | 0.194 | 0.04 | 5.26852E-31 |
| Terminal | Ncmap | 1.6016E-61 | 0.405869 | 0.178 | 0.003 | 2.121E-57 |
| Contractile | Glul | 2.31527E-11 | 0.405537 | 0.294 | 0.153 | 3.06611E-07 |
| Rgs5_Contractile | Dtna | 0.001518889 | 0.405478 | 0.108 | 0.062 | 1 |
| Contractile | Nes | 1.56677E-09 | 0.40546 | 0.235 | 0.12 | 2.07487E-05 |
| Contractile | Htr1b | 8.3397E-18 | 0.405404 | 0.182 | 0.053 | 1.10443E-13 |
| Contractile | Higd1a | 3.64801E-17 | 0.405007 | 0.865 | 0.79 | 4.83106E-13 |
| Contractile | Fra10ac1 | 9.42659E-09 | 0.405003 | 0.253 | 0.144 | 0.000124836 |
| Contractile | Myocd | 1.63166E-24 | 0.404974 | 0.153 | 0.028 | 2.16081E-20 |
| Terminal | Rab11fip4 | 9.47318E-53 | 0.404963 | 0.178 | 0.011 | 1.25453E-48 |
| Intermediate | Tuba1b | 1.96284E-20 | 0.40483 | 0.757 | 0.603 | 2.59939E-16 |
| Terminal | Tmie | 5.71009E-32 | 0.404696 | 0.197 | 0.047 | 7.56187E-28 |
| Contractile | Amotl2 | 1.34367E-06 | 0.403954 | 0.206 | 0.119 | 0.017794168 |
| Terminal | Arhgap31 | 4.64618E-39 | 0.40385 | 0.321 | 0.104 | 6.15294E-35 |
| Intermediate | Heg1 | 9.93589E-28 | 0.403848 | 0.195 | 0.052 | 1.31581E-23 |
| Contractile | Ppef1 | 8.51157E-38 | 0.403793 | 0.129 | 0.009 | 1.12719E-33 |
| Terminal | Cnpy2 | 3.1126E-27 | 0.403689 | 0.855 | 0.612 | 4.12201E-23 |
| Intermediate | Npdc1 | 2.20848E-22 | 0.403598 | 0.793 | 0.644 | 2.9247E-18 |
| Intermediate | Slc16a3 | 3.4193E-21 | 0.403408 | 0.275 | 0.123 | 4.52818E-17 |
| Contractile | Fabp3 | 6.05192E-14 | 0.403359 | 0.203 | 0.078 | 8.01456E-10 |
| Terminal | Col5a2 | 1.30979E-22 | 0.403181 | 0.906 | 0.671 | 1.73455E-18 |
| Contractile | Elf1 | 5.37455E-06 | 0.403027 | 0.262 | 0.177 | 0.071175197 |
| Terminal | Ngef | 7.59841E-36 | 0.402332 | 0.318 | 0.109 | 1.00626E-31 |
| Contractile | Arhgef25 | 8.10164E-06 | 0.402251 | 0.318 | 0.231 | 0.107290013 |
| Terminal | Vimp | 1.07763E-26 | 0.401996 | 0.862 | 0.658 | 1.42711E-22 |
| Terminal | Kcna6 | 1.22392E-61 | 0.40197 | 0.188 | 0.006 | 1.62083E-57 |

| | | | | | | |
|---|---|---|---|---|---|---|
| Contractile | Bcl2 | 4.64474E-09 | 0.40192 | 0.3 | 0.176 | 6.15103E-05 |
| Terminal | Avpr1a | 1.20107E-53 | 0.40183 | 0.179 | 0.01 | 1.59058E-49 |
| Contractile | Vbp1 | 1.26542E-06 | 0.401243 | 0.35 | 0.261 | 0.016757933 |
| Contractile | Kcnh2 | 9.44774E-09 | 0.401138 | 0.244 | 0.136 | 0.000125116 |
| Rgs5_Contractile | Hsp90aa1 | 0.008317911 | 0.401108 | 0.663 | 0.839 | 1 |
| Intermediate | Glis2 | 1.63486E-32 | 0.400677 | 0.281 | 0.094 | 2.16504E-28 |
| Contractile | Ecm2 | 4.90094E-06 | 0.400633 | 0.456 | 0.364 | 0.064903089 |
| Contractile | Cpq | 2.84132E-06 | 0.400424 | 0.762 | 0.735 | 0.037627585 |
| Contractile | Sema3f | 4.57016E-18 | 0.400392 | 0.235 | 0.083 | 6.05227E-14 |
| Contractile | Syf2 | 2.29109E-06 | 0.400353 | 0.512 | 0.451 | 0.030340896 |
| Contractile | Rab11b | 3.26779E-09 | 0.400232 | 0.629 | 0.545 | 4.32754E-05 |
| Contractile | Snrnp27 | 8.93329E-09 | 0.400206 | 0.638 | 0.575 | 0.000118304 |
| Intermediate | Vps29 | 2.31616E-25 | 0.399969 | 0.685 | 0.482 | 3.06729E-21 |
| Terminal | Hmgb2 | 5.38736E-32 | 0.399676 | 0.401 | 0.175 | 7.13448E-28 |
| Contractile | Prkaa2 | 1.79274E-27 | 0.399477 | 0.15 | 0.024 | 2.37413E-23 |
| Contractile | Cox6b1 | 9.03269E-18 | 0.399317 | 0.903 | 0.873 | 1.1962E-13 |
| Contractile | Diaph2 | 2.07283E-13 | 0.399294 | 0.238 | 0.103 | 2.74505E-09 |
| Terminal | Trib1 | 7.48599E-37 | 0.399224 | 0.309 | 0.102 | 9.9137E-33 |
| Terminal | Nucb2 | 5.16101E-28 | 0.399024 | 0.634 | 0.378 | 6.83473E-24 |
| Terminal | Phex | 2.25316E-31 | 0.398884 | 0.144 | 0.021 | 2.98386E-27 |
| Contractile | Vapa | 1.57015E-08 | 0.398699 | 0.671 | 0.607 | 0.000207936 |
| Contractile | Stk11 | 1.70439E-05 | 0.398696 | 0.397 | 0.326 | 0.225711743 |
| Intermediate | Ppp1ca | 1.06994E-25 | 0.398668 | 0.794 | 0.648 | 1.41692E-21 |
| Intermediate | Emp1 | 4.57324E-18 | 0.39828 | 0.595 | 0.416 | 6.05634E-14 |
| Contractile | Tmpo | 8.4864E-10 | 0.398201 | 0.218 | 0.107 | 1.12385E-05 |
| Contractile | Mrfap1 | 5.49904E-11 | 0.398143 | 0.824 | 0.787 | 7.28238E-07 |
| Terminal | Tmed10 | 8.85115E-32 | 0.397413 | 0.936 | 0.758 | 1.17216E-27 |
| Contractile | Nedd4 | 5.16104E-12 | 0.397214 | 0.806 | 0.767 | 6.83476E-08 |
| Terminal | Cemip | 3.59163E-39 | 0.39707 | 0.293 | 0.087 | 4.7564E-35 |
| Rgs5_Contractile | Cox7c | 2.18247E-07 | 0.39695 | 0.844 | 0.943 | 0.002890241 |
| Contractile | Rilpl1 | 2.63369E-08 | 0.39655 | 0.297 | 0.188 | 0.00034878 |
| Contractile | Trp53inp2 | 1.80082E-08 | 0.396135 | 0.306 | 0.193 | 0.000238482 |
| Intermediate | Id3 | 5.7091E-10 | 0.395868 | 0.922 | 0.872 | 7.56056E-06 |
| Terminal | St3gal6 | 5.20946E-56 | 0.395623 | 0.248 | 0.034 | 6.89889E-52 |
| Contractile | Ndufs8 | 2.54259E-07 | 0.39515 | 0.568 | 0.507 | 0.003367145 |
| Contractile | March2 | 2.73768E-08 | 0.395017 | 0.441 | 0.333 | 0.000362551 |
| Intermediate | Grn | 2.00872E-23 | 0.395014 | 0.694 | 0.497 | 2.66015E-19 |
| Contractile | Lsm3 | 7.50003E-07 | 0.394879 | 0.379 | 0.293 | 0.009932295 |
| Contractile | Mrps24 | 3.64485E-08 | 0.394746 | 0.612 | 0.536 | 0.000482688 |
| Terminal | Ddah1 | 9.52496E-40 | 0.394562 | 0.28 | 0.078 | 1.26139E-35 |
| Rgs5_Contractile | Uqcr11 | 0.000147029 | 0.394433 | 0.722 | 0.864 | 1 |
| Intermediate | Dctn3 | 5.3845E-22 | 0.394361 | 0.609 | 0.414 | 7.1307E-18 |
| Contractile | Fam188a | 8.12919E-08 | 0.393952 | 0.229 | 0.132 | 0.001076548 |
| Contractile | Elavl1 | 3.16294E-05 | 0.393877 | 0.409 | 0.349 | 0.418867707 |
| Contractile | Rtn2 | 3.31956E-14 | 0.393858 | 0.188 | 0.068 | 4.39609E-10 |
| Contractile | Pgrmc1 | 1.23781E-07 | 0.393858 | 0.659 | 0.602 | 0.001639233 |
| Contractile | Ebf1 | 2.77507E-10 | 0.392752 | 0.541 | 0.402 | 3.67503E-06 |
| Contractile | Tub | 9.26373E-34 | 0.392607 | 0.124 | 0.01 | 1.2268E-29 |
| Terminal | Lmo4 | 1.19515E-21 | 0.392348 | 0.822 | 0.59 | 1.58273E-17 |
| Contractile | 0610009L1 | 1.59042E-09 | 0.392137 | 0.203 | 0.098 | 2.10619E-05 |
| Contractile | Nrtn | 3.37514E-10 | 0.392123 | 0.388 | 0.241 | 4.4697E-06 |
| Contractile | Ndufab1 | 6.23483E-09 | 0.391989 | 0.671 | 0.637 | 8.25679E-05 |
| Contractile | Gm13861 | 3.05078E-20 | 0.391945 | 0.179 | 0.046 | 4.04015E-16 |
| Terminal | F13a1 | 1.25719E-23 | 0.391895 | 0.162 | 0.042 | 1.66489E-19 |
| Intermediate | Rnh1 | 7.01898E-19 | 0.391815 | 0.641 | 0.481 | 9.29523E-15 |
| Intermediate | Anxa8 | 2.20647E-11 | 0.391372 | 0.523 | 0.391 | 2.92203E-07 |
| Contractile | Mrpl34 | 7.10468E-06 | 0.391326 | 0.432 | 0.366 | 0.094087319 |
| Intermediate | Zfp36l2 | 1.27656E-16 | 0.391243 | 0.433 | 0.266 | 1.69055E-12 |
| Contractile | Zfp467 | 1.14211E-13 | 0.390565 | 0.209 | 0.083 | 1.5125E-09 |

| | | | | | | |
|---|---|---|---|---|---|---|
| Intermediate | B2m | 2.07753E-17 | 0.390468 | 0.978 | 0.948 | 2.75128E-13 |
| Contractile | Psmb3 | 8.1328E-11 | 0.390241 | 0.774 | 0.738 | 1.07703E-06 |
| Contractile | Gtf2a2 | 1.84944E-07 | 0.390124 | 0.576 | 0.516 | 0.002449208 |
| Contractile | Rnd3 | 7.7766E-09 | 0.390042 | 0.244 | 0.131 | 0.000102985 |
| Contractile | Chmp3 | 4.91763E-09 | 0.389723 | 0.556 | 0.468 | 6.51242E-05 |
| Contractile | Hip1 | 5.08554E-09 | 0.389559 | 0.297 | 0.177 | 6.73478E-05 |
| Terminal | Hand2 | 6.66792E-28 | 0.389547 | 0.364 | 0.161 | 8.83032E-24 |
| Intermediate | Cd82 | 6.13212E-22 | 0.389356 | 0.484 | 0.285 | 8.12077E-18 |
| Contractile | Gm12840 | 2.88686E-27 | 0.389219 | 0.474 | 0.194 | 3.82307E-23 |
| Intermediate | Htra3 | 8.0063E-14 | 0.389153 | 0.657 | 0.511 | 1.06027E-09 |
| Contractile | Aco2 | 7.0294E-06 | 0.388999 | 0.406 | 0.331 | 0.093090339 |
| Terminal | Rcan1 | 1.18694E-07 | 0.388975 | 0.284 | 0.189 | 0.001571864 |
| Contractile | Eif4ebp1 | 4.55771E-06 | 0.388917 | 0.485 | 0.405 | 0.060357788 |
| Contractile | Map7d2 | 1.4123E-25 | 0.388669 | 0.126 | 0.017 | 1.87031E-21 |
| Contractile | Ubxn1 | 8.34144E-07 | 0.388626 | 0.585 | 0.526 | 0.011046572 |
| Contractile | Vamp2 | 2.14914E-07 | 0.388542 | 0.403 | 0.299 | 0.002846103 |
| Contractile | Ywhaz | 1.37852E-07 | 0.38853 | 0.526 | 0.451 | 0.001825574 |
| Contractile | Spr | 3.93642E-06 | 0.388363 | 0.471 | 0.403 | 0.052130041 |
| Rgs5_Contractile | Itga3 | 0.001040111 | 0.388342 | 0.111 | 0.064 | 1 |
| Contractile | Adamts1 | 1.21656E-07 | 0.388182 | 0.347 | 0.228 | 0.001611093 |
| Contractile | Syne1 | 4.64153E-08 | 0.388105 | 0.397 | 0.285 | 0.000614678 |
| Contractile | Tgfbr1 | 2.7084E-07 | 0.388029 | 0.321 | 0.217 | 0.00358674 |
| Terminal | Rnaset2b | 6.18907E-28 | 0.387785 | 0.549 | 0.313 | 8.19619E-24 |
| Intermediate | Gusb | 6.43341E-23 | 0.387503 | 0.522 | 0.318 | 8.51976E-19 |
| Rgs5_Contractile | Uqcr10 | 0.001258709 | 0.387297 | 0.66 | 0.847 | 1 |
| Intermediate | Prrx2 | 5.05702E-21 | 0.387139 | 0.312 | 0.153 | 6.69701E-17 |
| Contractile | Calm3 | 2.73853E-08 | 0.387018 | 0.676 | 0.597 | 0.000362663 |
| Contractile | Adgrl2 | 1.44117E-08 | 0.387008 | 0.274 | 0.16 | 0.000190854 |
| Contractile | Peli2 | 4.61018E-16 | 0.386187 | 0.165 | 0.049 | 6.10525E-12 |
| Contractile | Cldnd1 | 9.94248E-05 | 0.385872 | 0.462 | 0.418 | 1 |
| Contractile | Ktn1 | 1.85413E-05 | 0.384902 | 0.447 | 0.372 | 0.245542602 |
| Intermediate | Mydgf | 1.02618E-17 | 0.384794 | 0.63 | 0.481 | 1.35897E-13 |
| Contractile | Hpcal1 | 1.15493E-05 | 0.384488 | 0.226 | 0.144 | 0.152947962 |
| Intermediate | Slco3a1 | 3.29786E-28 | 0.384369 | 0.313 | 0.126 | 4.36736E-24 |
| Terminal | Serinc5 | 7.72611E-41 | 0.384303 | 0.264 | 0.066 | 1.02317E-36 |
| Intermediate | Tnfrsf12a | 7.16721E-18 | 0.384261 | 0.635 | 0.455 | 9.49154E-14 |
| Rgs5_Contractile | Lhfp | 0.000689447 | 0.383925 | 0.681 | 0.821 | 1 |
| Terminal | Creb3l1 | 4.03991E-22 | 0.383544 | 0.475 | 0.274 | 5.35006E-18 |
| Contractile | Sumo2 | 2.83758E-14 | 0.383276 | 0.882 | 0.848 | 3.75781E-10 |
| Terminal | Gm26532 | 1.47559E-32 | 0.383191 | 0.275 | 0.088 | 1.95412E-28 |
| Intermediate | Parp3 | 1.60763E-22 | 0.382986 | 0.338 | 0.164 | 2.12899E-18 |
| Contractile | Cyc1 | 1.11921E-05 | 0.382541 | 0.526 | 0.491 | 0.14821635 |
| Intermediate | Atf5 | 1.20399E-18 | 0.382454 | 0.415 | 0.241 | 1.59445E-14 |
| Contractile | Dnajc8 | 1.6777E-05 | 0.38234 | 0.574 | 0.542 | 0.222178385 |
| Contractile | Tmem43 | 5.25784E-05 | 0.382133 | 0.374 | 0.303 | 0.696295457 |
| Contractile | Mrpl30 | 2.12516E-05 | 0.382056 | 0.491 | 0.45 | 0.281434636 |
| Contractile | Wnt16 | 7.09506E-27 | 0.382032 | 0.188 | 0.038 | 9.39598E-23 |
| Contractile | Klf13 | 3.27099E-06 | 0.381911 | 0.515 | 0.448 | 0.0433177 |
| Contractile | Klhdc8b | 6.61724E-17 | 0.381735 | 0.247 | 0.095 | 8.7632E-13 |
| Contractile | Psma2 | 7.79447E-08 | 0.381694 | 0.744 | 0.713 | 0.001032221 |
| Terminal | Tpbg | 2.1647E-28 | 0.381597 | 0.239 | 0.08 | 2.86671E-24 |
| Intermediate | Tspan17 | 2.60852E-25 | 0.381539 | 0.306 | 0.134 | 3.45447E-21 |
| Contractile | Prdm6 | 5.45476E-09 | 0.381476 | 0.268 | 0.153 | 7.22374E-05 |
| Intermediate | Cfb | 1.47294E-05 | 0.381328 | 0.116 | 0.067 | 0.195060848 |
| Contractile | Sema5a | 1.94169E-14 | 0.38079 | 0.229 | 0.091 | 2.57138E-10 |
| Contractile | Fgf1 | 6.5857E-19 | 0.38071 | 0.191 | 0.056 | 8.72144E-15 |
| Contractile | Eif3i | 5.78763E-08 | 0.380615 | 0.659 | 0.601 | 0.000766455 |
| Contractile | Xpr1 | 1.47398E-08 | 0.379012 | 0.312 | 0.194 | 0.000195199 |
| Intermediate | Flrt2 | 4.4003E-31 | 0.37868 | 0.251 | 0.08 | 5.82732E-27 |

| | | | | | | |
|---|---|---|---|---|---|---|
| Intermediate | Bicc1 | 1.80698E-22 | 0.37844 | 0.257 | 0.107 | 2.39299E-18 |
| Contractile | Grcc10 | 4.29964E-06 | 0.378171 | 0.568 | 0.52 | 0.056940148 |
| Contractile | Psmd4 | 8.03428E-07 | 0.378131 | 0.55 | 0.485 | 0.01063979 |
| Contractile | Gltscr2 | 3.87511E-07 | 0.377991 | 0.621 | 0.569 | 0.005131807 |
| Rgs5_Contractile | Gm11627 | 0.000327297 | 0.377839 | 0.128 | 0.073 | 1 |
| Contractile | Dnaja2 | 9.99216E-06 | 0.377088 | 0.453 | 0.391 | 0.132326231 |
| Rgs5_Contractile | Dynll1 | 3.07616E-08 | 0.37693 | 0.924 | 0.973 | 0.000407376 |
| Contractile | Tmtc2 | 3.07964E-12 | 0.376905 | 0.126 | 0.039 | 4.07837E-08 |
| Contractile | Phf5a | 2.82425E-05 | 0.376806 | 0.379 | 0.311 | 0.374015136 |
| Contractile | Emc10 | 1.15356E-10 | 0.376745 | 0.741 | 0.687 | 1.52767E-06 |
| Contractile | Hnrnpk | 3.02191E-14 | 0.376733 | 0.885 | 0.833 | 4.00192E-10 |
| Intermediate | Hs6st2 | 3.22159E-35 | 0.376509 | 0.154 | 0.02 | 4.26635E-31 |
| Terminal | Ramp2 | 1.61357E-38 | 0.376418 | 0.249 | 0.063 | 2.13685E-34 |
| Contractile | Calm1 | 2.31502E-18 | 0.376326 | 0.965 | 0.921 | 3.06578E-14 |
| Contractile | Lzts2 | 3.81425E-06 | 0.375734 | 0.409 | 0.325 | 0.050512146 |
| Intermediate | Ctsd | 1.1956E-18 | 0.375572 | 0.927 | 0.857 | 1.58334E-14 |
| Intermediate | P3h3 | 3.56116E-17 | 0.375561 | 0.715 | 0.563 | 4.71605E-13 |
| Intermediate | Glipr2 | 2.58241E-32 | 0.375443 | 0.26 | 0.082 | 3.41989E-28 |
| Contractile | Chmp4c | 1.97273E-21 | 0.375403 | 0.147 | 0.03 | 2.61248E-17 |
| Intermediate | Abca1 | 1.72933E-20 | 0.37417 | 0.256 | 0.111 | 2.29016E-16 |
| Intermediate | Dbn1 | 9.68962E-33 | 0.374142 | 0.247 | 0.072 | 1.2832E-28 |
| Contractile | St3gal5 | 1.51239E-13 | 0.374045 | 0.218 | 0.089 | 2.00286E-09 |
| Rgs5_Contractile | Peli2 | 0.001962135 | 0.373712 | 0.104 | 0.06 | 1 |
| Intermediate | Gm9844 | 5.68247E-27 | 0.373707 | 0.251 | 0.086 | 7.5253E-23 |
| Contractile | Adm | 5.39E-18 | 0.373684 | 0.162 | 0.042 | 7.13798E-14 |
| Terminal | Clec11a | 7.30513E-23 | 0.373533 | 0.363 | 0.175 | 9.67418E-19 |
| Intermediate | Dap | 3.5776E-18 | 0.373529 | 0.819 | 0.698 | 4.73781E-14 |
| Terminal | 1190002N | 2.07671E-34 | 0.373464 | 0.281 | 0.088 | 2.75019E-30 |
| Intermediate | Rab11a | 1.35304E-20 | 0.373333 | 0.596 | 0.401 | 1.79183E-16 |
| Contractile | Pkig | 2.08687E-05 | 0.373113 | 0.547 | 0.505 | 0.276364087 |
| Contractile | Gne | 7.12826E-14 | 0.373074 | 0.203 | 0.079 | 9.43996E-10 |
| Intermediate | Itpripl2 | 1.19128E-17 | 0.372847 | 0.44 | 0.273 | 1.57761E-13 |
| Contractile | Pmvk | 3.84936E-07 | 0.372654 | 0.215 | 0.124 | 0.005097702 |
| Intermediate | Slc48a1 | 3.46138E-19 | 0.37243 | 0.706 | 0.551 | 4.5839E-15 |
| Contractile | Foxj2 | 3.49246E-09 | 0.37211 | 0.165 | 0.073 | 4.62506E-05 |
| Intermediate | Axl | 1.69204E-32 | 0.371883 | 0.609 | 0.328 | 2.24076E-28 |
| Terminal | Igfbp6 | 1.08245E-14 | 0.371876 | 0.24 | 0.118 | 1.43348E-10 |
| Contractile | Fzd2 | 1.01564E-06 | 0.371629 | 0.288 | 0.192 | 0.013450186 |
| Contractile | Dhrs3 | 5.43971E-05 | 0.37149 | 0.385 | 0.321 | 0.720380406 |
| Intermediate | AI413582 | 4.00916E-22 | 0.371451 | 0.332 | 0.164 | 5.30933E-18 |
| Contractile | Rhobtb1 | 8.27701E-23 | 0.371307 | 0.171 | 0.037 | 1.09612E-18 |
| Terminal | Egr1 | 2.02178E-20 | 0.371172 | 0.542 | 0.341 | 2.67745E-16 |
| Intermediate | Mast4 | 5.38479E-14 | 0.371006 | 0.374 | 0.228 | 7.13107E-10 |
| Contractile | Dbndd2 | 7.88471E-15 | 0.370894 | 0.221 | 0.085 | 1.04417E-10 |
| Contractile | Larp1b | 3.36281E-06 | 0.370502 | 0.203 | 0.123 | 0.044533656 |
| Contractile | Ube2b | 1.12425E-09 | 0.370463 | 0.706 | 0.648 | 1.48885E-05 |
| Intermediate | Il33 | 2.27151E-17 | 0.370139 | 0.101 | 0.022 | 3.00816E-13 |
| Intermediate | Rab31 | 9.82733E-21 | 0.370007 | 0.351 | 0.185 | 1.30143E-16 |
| Intermediate | B4galt1 | 7.9918E-23 | 0.36985 | 0.299 | 0.134 | 1.05835E-18 |
| Contractile | Pfdn1 | 2.5544E-07 | 0.369747 | 0.644 | 0.59 | 0.00338279 |
| Terminal | Ninj1 | 3.39292E-14 | 0.36951 | 0.748 | 0.574 | 4.49325E-10 |
| Contractile | Bloc1s1 | 4.71313E-05 | 0.369314 | 0.441 | 0.383 | 0.624160262 |
| Rgs5_Contractile | Ndufa4 | 0.001457895 | 0.369168 | 0.743 | 0.882 | 1 |
| Contractile | Capn2 | 3.66929E-05 | 0.369067 | 0.474 | 0.419 | 0.485923732 |
| Contractile | Mrpl24 | 0.000282506 | 0.369026 | 0.459 | 0.425 | 1 |
| Intermediate | P4hb | 6.18655E-20 | 0.369013 | 0.765 | 0.633 | 8.19285E-16 |
| Terminal | Stk32b | 3.75731E-32 | 0.368959 | 0.217 | 0.057 | 4.97581E-28 |
| Contractile | Pcnp | 5.96206E-06 | 0.368866 | 0.518 | 0.455 | 0.078955535 |
| Terminal | Ssr2 | 1.44369E-21 | 0.36881 | 0.826 | 0.601 | 1.91188E-17 |

| | | | | | | |
|---|---|---|---|---|---|---|
| Intermediate | Fn1 | 8.57393E-24 | 0.368548 | 0.982 | 0.926 | 1.13545E-19 |
| Intermediate | Sdc3 | 6.67985E-13 | 0.368485 | 0.504 | 0.366 | 8.84613E-09 |
| Contractile | Tomm7 | 1.05452E-10 | 0.368125 | 0.856 | 0.812 | 1.3965E-06 |
| Contractile | Echs1 | 0.002531607 | 0.368033 | 0.359 | 0.327 | 1 |
| Contractile | Ahsa1 | 0.000187077 | 0.36761 | 0.485 | 0.461 | 1 |
| Contractile | Ids | 4.96408E-05 | 0.367593 | 0.265 | 0.192 | 0.657393468 |
| Intermediate | H2afj | 9.98947E-20 | 0.367536 | 0.785 | 0.634 | 1.32291E-15 |
| Contractile | Sdhb | 1.02728E-07 | 0.367322 | 0.615 | 0.553 | 0.00136043 |
| Intermediate | Sec11a | 3.32074E-19 | 0.367246 | 0.724 | 0.558 | 4.39765E-15 |
| Contractile | B3galt1 | 2.27793E-18 | 0.367233 | 0.159 | 0.041 | 3.01666E-14 |
| Contractile | Eef2k | 1.3204E-09 | 0.367114 | 0.212 | 0.104 | 1.74861E-05 |
| Contractile | Fam32a | 1.84162E-08 | 0.366432 | 0.432 | 0.316 | 0.000243885 |
| Contractile | Atp1b2 | 1.07736E-23 | 0.366373 | 0.174 | 0.037 | 1.42675E-19 |
| Contractile | Atp5l | 9.02622E-18 | 0.36583 | 0.947 | 0.931 | 1.19534E-13 |
| Contractile | Aimp1 | 2.65927E-05 | 0.365779 | 0.421 | 0.359 | 0.352167471 |
| Contractile | Slit3 | 2.06427E-15 | 0.365127 | 0.165 | 0.05 | 2.73371E-11 |
| Terminal | Emb | 1.34975E-25 | 0.365013 | 0.574 | 0.327 | 1.78748E-21 |
| Contractile | Creld1 | 0.002294109 | 0.364968 | 0.226 | 0.177 | 1 |
| Intermediate | Tmem167 | 2.26168E-18 | 0.36469 | 0.772 | 0.641 | 2.99515E-14 |
| Intermediate | Cstb | 1.48802E-13 | 0.364582 | 0.794 | 0.688 | 1.97059E-09 |
| Intermediate | Ap1s1 | 1.56293E-22 | 0.364566 | 0.473 | 0.281 | 2.06979E-18 |
| Intermediate | Slc7a2 | 1.03085E-12 | 0.364553 | 0.449 | 0.307 | 1.36515E-08 |
| Terminal | Btg1 | 3.5018E-14 | 0.364428 | 0.833 | 0.698 | 4.63744E-10 |
| Contractile | Chchd10 | 2.25397E-21 | 0.36437 | 0.153 | 0.032 | 2.98494E-17 |
| Contractile | Pdcd5 | 1.21286E-07 | 0.364266 | 0.659 | 0.605 | 0.001606185 |
| Intermediate | Por | 8.27611E-18 | 0.364225 | 0.381 | 0.217 | 1.09601E-13 |
| Intermediate | Atox1 | 1.74641E-23 | 0.364224 | 0.89 | 0.814 | 2.31278E-19 |
| Intermediate | Cxcl14 | 4.8473E-05 | 0.36414 | 0.236 | 0.171 | 0.641928485 |
| Intermediate | Nid1 | 9.8684E-32 | 0.364023 | 0.225 | 0.062 | 1.30687E-27 |
| Intermediate | Ddah2 | 1.41226E-28 | 0.363947 | 0.436 | 0.208 | 1.87026E-24 |
| Terminal | Tnfaip2 | 1.10615E-21 | 0.363937 | 0.382 | 0.196 | 1.46487E-17 |
| Intermediate | Crispld2 | 0.000269405 | 0.363836 | 0.254 | 0.192 | 1 |
| Intermediate | Crip2 | 1.07357E-22 | 0.363747 | 0.894 | 0.805 | 1.42173E-18 |
| Contractile | Gm10250 | 4.8609E-06 | 0.363635 | 0.591 | 0.549 | 0.064372951 |
| Contractile | Ndufv2 | 1.02271E-07 | 0.363589 | 0.624 | 0.576 | 0.001354375 |
| Intermediate | B4galnt1 | 4.9239E-35 | 0.3634 | 0.224 | 0.055 | 6.52073E-31 |
| Contractile | Ccdc141 | 7.14526E-15 | 0.363387 | 0.188 | 0.066 | 9.46246E-11 |
| Rgs5_Contractile | Rsrp1 | 0.000199287 | 0.363144 | 0.726 | 0.858 | 1 |
| Rgs5_Contractile | Ndufa2 | 0.000454399 | 0.363019 | 0.726 | 0.894 | 1 |
| Intermediate | Tpi1 | 5.74864E-14 | 0.363016 | 0.767 | 0.651 | 7.61293E-10 |
| Contractile | Siva1 | 5.0972E-07 | 0.362984 | 0.459 | 0.373 | 0.006750216 |
| Terminal | Nfkbia | 3.33869E-20 | 0.362883 | 0.77 | 0.581 | 4.42143E-16 |
| Terminal | Ppic | 4.14232E-20 | 0.362617 | 0.887 | 0.635 | 5.48568E-16 |
| Contractile | Cltb | 5.52988E-08 | 0.362573 | 0.629 | 0.543 | 0.000732322 |
| Contractile | Morf4l1 | 9.81172E-17 | 0.362305 | 0.959 | 0.936 | 1.29937E-12 |
| Intermediate | Ece1 | 2.6097E-27 | 0.362265 | 0.235 | 0.075 | 3.45603E-23 |
| Contractile | 6330403A( | 2.67197E-27 | 0.361899 | 0.121 | 0.014 | 3.53849E-23 |
| Contractile | Gng12 | 3.23208E-06 | 0.36181 | 0.479 | 0.402 | 0.042802445 |
| Contractile | Toporsos | 1.91259E-06 | 0.361761 | 0.529 | 0.463 | 0.02532847 |
| Contractile | Dusp8 | 3.55568E-14 | 0.361518 | 0.174 | 0.059 | 4.70878E-10 |
| Intermediate | Sar1a | 1.21752E-14 | 0.361427 | 0.701 | 0.574 | 1.61236E-10 |
| Contractile | Atp13a3 | 4.92159E-07 | 0.361273 | 0.229 | 0.136 | 0.006517664 |
| Contractile | Pbx1 | 8.72374E-06 | 0.361212 | 0.435 | 0.352 | 0.115528541 |
| Terminal | Evi2a | 1.74608E-23 | 0.361109 | 0.206 | 0.07 | 2.31234E-19 |
| Intermediate | Olfml3 | 2.76413E-08 | 0.360857 | 0.341 | 0.241 | 0.000366054 |
| Intermediate | H2-T23 | 7.92467E-28 | 0.360843 | 0.229 | 0.073 | 1.04946E-23 |
| Intermediate | Tnfrsf1a | 2.83004E-15 | 0.360738 | 0.574 | 0.414 | 3.74782E-11 |
| Contractile | Nfil3 | 3.34895E-08 | 0.36072 | 0.256 | 0.147 | 0.000443501 |
| Intermediate | Ephx1 | 9.77111E-18 | 0.360715 | 0.359 | 0.196 | 1.29399E-13 |

| | | | | | | |
|---|---|---|---|---|---|---|
| Contractile | Cpne3 | 2.05129E-06 | 0.360696 | 0.385 | 0.297 | 0.02716519 |
| Contractile | Mtdh | 7.26386E-08 | 0.360678 | 0.671 | 0.628 | 0.000961953 |
| Contractile | Eef1a1 | 8.44494E-19 | 0.360377 | 0.985 | 0.984 | 1.11836E-14 |
| Contractile | Camk2d | 1.1807E-06 | 0.360294 | 0.568 | 0.508 | 0.015635993 |
| Terminal | Slc16a2 | 5.66988E-29 | 0.360155 | 0.376 | 0.171 | 7.50862E-25 |
| Intermediate | Sec61b | 2.36301E-19 | 0.359708 | 0.888 | 0.788 | 3.12934E-15 |
| Intermediate | Rbms1 | 2.04415E-16 | 0.358834 | 0.558 | 0.391 | 2.70707E-12 |
| Intermediate | Pmp22 | 1.62406E-15 | 0.358331 | 0.672 | 0.53 | 2.15074E-11 |
| Contractile | Ahnak | 6.51778E-11 | 0.358166 | 0.806 | 0.743 | 8.63149E-07 |
| Contractile | Uqcrc1 | 5.35097E-07 | 0.357808 | 0.615 | 0.556 | 0.007086296 |
| Contractile | Ndufaf3 | 0.000558965 | 0.357718 | 0.324 | 0.273 | 1 |
| Intermediate | B4galt5 | 1.67533E-23 | 0.35744 | 0.214 | 0.074 | 2.21864E-19 |
| Rgs5_Contractile | Dnajb5 | 0.009471775 | 0.357077 | 0.101 | 0.064 | 1 |
| Intermediate | Cdk4 | 3.69547E-20 | 0.356909 | 0.563 | 0.38 | 4.89391E-16 |
| Contractile | Ccnd3 | 0.0001101 | 0.356599 | 0.412 | 0.356 | 1 |
| Intermediate | Cdh11 | 5.075E-23 | 0.356389 | 0.206 | 0.07 | 6.72083E-19 |
| Contractile | Abhd17b | 4.38626E-06 | 0.356293 | 0.2 | 0.121 | 0.058087227 |
| Contractile | Csnk2b | 2.27005E-05 | 0.356252 | 0.438 | 0.376 | 0.300622921 |
| Contractile | Naa50 | 4.03168E-06 | 0.355962 | 0.397 | 0.316 | 0.05339159 |
| Intermediate | Il3ra | 1.46468E-33 | 0.355896 | 0.194 | 0.042 | 1.93968E-29 |
| Terminal | Tlr2 | 2.31375E-18 | 0.355749 | 0.265 | 0.125 | 3.0641E-14 |
| Intermediate | Lnx1 | 7.09657E-24 | 0.355741 | 0.29 | 0.125 | 9.39799E-20 |
| Terminal | Pnrc1 | 2.15327E-14 | 0.355172 | 0.657 | 0.486 | 2.85157E-10 |
| Contractile | Hmg20b | 2.81054E-05 | 0.354906 | 0.45 | 0.39 | 0.372199384 |
| Terminal | Pdcd4 | 7.66855E-29 | 0.354655 | 0.398 | 0.183 | 1.01555E-24 |
| Rgs5_Contractile | Ybx1 | 1.49408E-05 | 0.354451 | 0.882 | 0.942 | 0.19786154 |
| Contractile | Zfp367 | 2.63569E-11 | 0.354204 | 0.171 | 0.067 | 3.49045E-07 |
| Contractile | Cav3 | 1.27146E-07 | 0.35416 | 0.226 | 0.129 | 0.001683798 |
| Contractile | Hnrnpr | 0.000155771 | 0.35368 | 0.421 | 0.37 | 1 |
| Contractile | Spop | 2.08089E-05 | 0.353484 | 0.535 | 0.494 | 0.275572465 |
| Contractile | Mafk | 1.37718E-07 | 0.353468 | 0.229 | 0.131 | 0.001823798 |
| Terminal | Enpp1 | 3.51419E-28 | 0.353327 | 0.472 | 0.235 | 4.65384E-24 |
| Contractile | Tead3 | 1.14501E-11 | 0.353094 | 0.206 | 0.088 | 1.51634E-07 |
| Terminal | Abi3bp | 9.28626E-34 | 0.353062 | 0.676 | 0.376 | 1.22978E-29 |
| Contractile | Psip1 | 0.000132138 | 0.352982 | 0.335 | 0.266 | 1 |
| Contractile | Psmc4 | 4.55476E-05 | 0.352838 | 0.491 | 0.439 | 0.603186863 |
| Contractile | 1500011K: | 0.000362093 | 0.352751 | 0.468 | 0.448 | 1 |
| Rgs5_Contractile | Aldoa | 1.47296E-06 | 0.352695 | 0.819 | 0.906 | 0.019506355 |
| Intermediate | Cmtm3 | 1.8326E-25 | 0.352692 | 0.224 | 0.076 | 2.42691E-21 |
| Contractile | Zbtb7a | 0.000619531 | 0.352667 | 0.359 | 0.31 | 1 |
| Contractile | Clpp | 7.56444E-05 | 0.352662 | 0.368 | 0.302 | 1 |
| Contractile | Marveld1 | 3.91593E-06 | 0.352398 | 0.462 | 0.378 | 0.051858708 |
| Contractile | Fez1 | 0.000210597 | 0.35174 | 0.274 | 0.208 | 1 |
| Contractile | Nenf | 3.74227E-15 | 0.351715 | 0.924 | 0.886 | 4.95589E-11 |
| Intermediate | Il13ra1 | 1.96177E-17 | 0.351625 | 0.381 | 0.222 | 2.59797E-13 |
| Terminal | Stxbp6 | 9.49744E-28 | 0.351605 | 0.272 | 0.102 | 1.25775E-23 |
| Contractile | Skp1a | 1.23707E-06 | 0.3515 | 0.618 | 0.577 | 0.016382476 |
| Contractile | Srf | 5.79615E-10 | 0.351284 | 0.215 | 0.104 | 7.67584E-06 |
| Contractile | Kif13a | 1.13097E-10 | 0.351221 | 0.191 | 0.084 | 1.49775E-06 |
| Rgs5_Contractile | Impdh2 | 0.001429428 | 0.351089 | 0.25 | 0.446 | 1 |
| Contractile | Bambi | 0.00103202 | 0.350935 | 0.309 | 0.248 | 1 |
| Contractile | Ndufa10 | 0.000195698 | 0.350576 | 0.444 | 0.401 | 1 |
| Contractile | 1110059E: | 8.68399E-06 | 0.350564 | 0.229 | 0.146 | 0.115002085 |
| Contractile | Ano6 | 2.89207E-06 | 0.350522 | 0.344 | 0.247 | 0.038299717 |
| Contractile | B230118H | 0.000791588 | 0.350471 | 0.306 | 0.25 | 1 |
| Terminal | Acot1 | 1.59003E-47 | 0.350182 | 0.182 | 0.018 | 2.10568E-43 |
| Contractile | Pja2 | 8.90242E-06 | 0.35016 | 0.362 | 0.276 | 0.117894723 |
| Contractile | Fkbp8 | 0.000275623 | 0.350051 | 0.559 | 0.55 | 1 |
| Terminal | Isg20 | 2.90645E-29 | 0.349907 | 0.249 | 0.085 | 3.84901E-25 |

| | | | | | | |
|---|---|---|---|---|---|---|
| Intermediate | Asap1 | 1.57438E-18 | 0.349906 | 0.357 | 0.194 | 2.08495E-14 |
| Contractile | Sucla2 | 0.000384315 | 0.349721 | 0.341 | 0.283 | 1 |
| Contractile | Zfp106 | 0.000399238 | 0.349616 | 0.294 | 0.232 | 1 |
| Contractile | Ppp2r5a | 1.11808E-06 | 0.349535 | 0.226 | 0.136 | 0.014806776 |
| Contractile | Akirin2 | 9.41251E-06 | 0.349487 | 0.362 | 0.278 | 0.124649832 |
| Intermediate | Uap1 | 2.05711E-06 | 0.349461 | 0.446 | 0.353 | 0.027242248 |
| Contractile | Mbnl2 | 1.13984E-06 | 0.34921 | 0.606 | 0.56 | 0.015094963 |
| Intermediate | Epas1 | 5.73662E-11 | 0.349195 | 0.331 | 0.212 | 7.597E-07 |
| Contractile | Nab1 | 0.002383651 | 0.349148 | 0.3 | 0.256 | 1 |
| Intermediate | Ufm1 | 2.17181E-17 | 0.348955 | 0.534 | 0.364 | 2.87613E-13 |
| Terminal | Sertad1 | 8.22478E-22 | 0.348733 | 0.519 | 0.306 | 1.08921E-17 |
| Contractile | Prmt1 | 3.18888E-05 | 0.348606 | 0.494 | 0.442 | 0.422303134 |
| Intermediate | Fst | 7.27903E-17 | 0.348533 | 0.27 | 0.136 | 9.63962E-13 |
| Contractile | Pmpcb | 9.11668E-05 | 0.34777 | 0.282 | 0.214 | 1 |
| Contractile | Trio | 5.08086E-09 | 0.347392 | 0.203 | 0.1 | 6.72858E-05 |
| Contractile | Prkar2a | 5.09984E-09 | 0.347214 | 0.265 | 0.15 | 6.75371E-05 |
| Contractile | Lama4 | 1.56425E-07 | 0.347035 | 0.253 | 0.151 | 0.00207153 |
| Contractile | Sh3rf2 | 9.93568E-35 | 0.347025 | 0.109 | 0.006 | 1.31578E-30 |
| Intermediate | Nrp1 | 1.63361E-20 | 0.34691 | 0.38 | 0.202 | 2.16339E-16 |
| Intermediate | Pdia5 | 6.09141E-23 | 0.346771 | 0.256 | 0.105 | 8.06686E-19 |
| Contractile | Pfdn6 | 0.000778047 | 0.346722 | 0.388 | 0.345 | 1 |
| Contractile | Zfand6 | 0.000136399 | 0.346705 | 0.447 | 0.399 | 1 |
| Intermediate | Psma3 | 1.62183E-22 | 0.346575 | 0.836 | 0.725 | 2.14779E-18 |
| Rgs5_Contractile | Gm11808 | 0.003343541 | 0.346498 | 0.681 | 0.847 | 1 |
| Intermediate | Prkcsh | 2.98724E-15 | 0.346467 | 0.469 | 0.315 | 3.956E-11 |
| Contractile | Stk17b | 1.52144E-07 | 0.346338 | 0.218 | 0.122 | 0.002014839 |
| Contractile | Mpc2 | 7.10555E-07 | 0.346338 | 0.665 | 0.63 | 0.009409882 |
| Contractile | Eed | 5.07176E-06 | 0.346298 | 0.226 | 0.144 | 0.067165316 |
| Contractile | Pawr | 1.39442E-12 | 0.34625 | 0.244 | 0.108 | 1.84662E-08 |
| Contractile | Flii | 0.000168474 | 0.346134 | 0.282 | 0.216 | 1 |
| Contractile | Cct7 | 1.07394E-05 | 0.345422 | 0.568 | 0.529 | 0.142221407 |
| Intermediate | Trappc4 | 2.14238E-16 | 0.345252 | 0.555 | 0.385 | 2.83715E-12 |
| Intermediate | Hexa | 1.19885E-19 | 0.345244 | 0.862 | 0.669 | 1.58764E-15 |
| Intermediate | Rhoc | 8.62736E-21 | 0.345227 | 0.856 | 0.754 | 1.14252E-16 |
| Contractile | Cript | 3.14544E-07 | 0.345064 | 0.506 | 0.42 | 0.004165509 |
| Contractile | N6amt1 | 0.000110668 | 0.344847 | 0.332 | 0.263 | 1 |
| Intermediate | Gpnmb | 1.70079E-12 | 0.344433 | 0.112 | 0.039 | 2.25235E-08 |
| Intermediate | Tapbp | 1.23343E-15 | 0.34433 | 0.527 | 0.368 | 1.63343E-11 |
| Contractile | Faap20 | 8.2018E-06 | 0.344319 | 0.306 | 0.219 | 0.108616425 |
| Contractile | Ankrd9 | 2.25442E-09 | 0.344156 | 0.191 | 0.09 | 2.98552E-05 |
| Contractile | Camta1 | 0.000629809 | 0.344089 | 0.356 | 0.303 | 1 |
| Intermediate | Rbx1 | 6.11113E-20 | 0.343613 | 0.868 | 0.771 | 8.09297E-16 |
| Contractile | Tnfsf12 | 9.08625E-07 | 0.343518 | 0.297 | 0.198 | 0.012032917 |
| Intermediate | Scpep1 | 2.61523E-18 | 0.343513 | 0.496 | 0.316 | 3.46334E-14 |
| Contractile | Hnrnpa2b1 | 1.22264E-09 | 0.343496 | 0.824 | 0.781 | 1.61914E-05 |
| Contractile | Gadd45gip | 0.000381533 | 0.343481 | 0.412 | 0.37 | 1 |
| Contractile | Mycbp | 0.000295916 | 0.343463 | 0.247 | 0.185 | 1 |
| Contractile | Pdlim2 | 1.78104E-06 | 0.343433 | 0.453 | 0.359 | 0.023586371 |
| Contractile | Rexo2 | 4.66525E-08 | 0.343248 | 0.806 | 0.772 | 0.00061782 |
| Intermediate | Lima1 | 5.22934E-24 | 0.343226 | 0.262 | 0.106 | 6.92521E-20 |
| Intermediate | Plin2 | 2.44065E-27 | 0.343171 | 0.305 | 0.123 | 3.23215E-23 |
| Rgs5_Contractile | Mmp17 | 0.002464156 | 0.343089 | 0.101 | 0.058 | 1 |
| Contractile | Fez2 | 1.23877E-05 | 0.343028 | 0.347 | 0.268 | 0.164050358 |
| Contractile | Tesk1 | 0.000243528 | 0.342848 | 0.162 | 0.103 | 1 |
| Contractile | Armcx1 | 2.03022E-11 | 0.342666 | 0.174 | 0.069 | 2.68862E-07 |
| Intermediate | Plp2 | 1.9706E-18 | 0.34258 | 0.581 | 0.396 | 2.60967E-14 |
| Contractile | Nol7 | 5.09717E-05 | 0.342566 | 0.474 | 0.424 | 0.675018309 |
| Terminal | Chst11 | 5.38806E-26 | 0.342321 | 0.284 | 0.115 | 7.1354E-22 |
| Contractile | Smim11 | 9.16392E-08 | 0.34195 | 0.621 | 0.567 | 0.001213577 |

| | | | | | | |
|---|---|---|---|---|---|---|
| Intermediate | Mif | 9.31305E-12 | 0.341714 | 0.813 | 0.707 | 1.23333E-07 |
| Intermediate | Plac9b | 6.67107E-20 | 0.341298 | 0.326 | 0.164 | 8.8345E-16 |
| Intermediate | Eno1 | 6.28542E-10 | 0.34095 | 0.762 | 0.681 | 8.32378E-06 |
| Contractile | Vdac1 | 1.84458E-06 | 0.340856 | 0.521 | 0.446 | 0.024427785 |
| Contractile | Ccdc69 | 5.88778E-30 | 0.340744 | 0.106 | 0.008 | 7.79719E-26 |
| Contractile | Pin4 | 0.001177443 | 0.340714 | 0.391 | 0.353 | 1 |
| Contractile | Ppm1b | 5.04822E-06 | 0.340499 | 0.25 | 0.165 | 0.066853554 |
| Intermediate | Sphk1 | 5.24913E-23 | 0.34033 | 0.257 | 0.103 | 6.95143E-19 |
| Intermediate | Psma4 | 1.95396E-15 | 0.340262 | 0.709 | 0.561 | 2.58763E-11 |
| Contractile | Pdcd10 | 0.000788194 | 0.340214 | 0.456 | 0.428 | 1 |
| Intermediate | Tmbim6 | 1.66021E-20 | 0.34007 | 0.821 | 0.694 | 2.19862E-16 |
| Intermediate | Necap2 | 2.41603E-18 | 0.339759 | 0.398 | 0.231 | 3.19954E-14 |
| Contractile | Prdm16 | 2.40612E-11 | 0.339619 | 0.144 | 0.05 | 3.18642E-07 |
| Terminal | Cdo1 | 1.18389E-11 | 0.339523 | 0.635 | 0.478 | 1.56782E-07 |
| Contractile | Cbx5 | 0.000234887 | 0.339274 | 0.338 | 0.278 | 1 |
| Intermediate | Tmem50a | 1.38415E-17 | 0.339139 | 0.784 | 0.686 | 1.83304E-13 |
| Contractile | Pip5k1b | 7.51905E-36 | 0.339127 | 0.15 | 0.015 | 9.95748E-32 |
| Rgs5_Contractile | Txn1 | 0.000448842 | 0.339111 | 0.691 | 0.847 | 1 |
| Contractile | Itpkb | 0.001541561 | 0.339105 | 0.159 | 0.107 | 1 |
| Contractile | Mertk | 1.81257E-14 | 0.339041 | 0.15 | 0.044 | 2.40038E-10 |
| Contractile | Aip | 1.68825E-05 | 0.33831 | 0.362 | 0.289 | 0.223574392 |
| Contractile | Meis1 | 5.69167E-07 | 0.338304 | 0.185 | 0.1 | 0.007537478 |
| Terminal | Ldha | 4.00882E-18 | 0.338291 | 0.964 | 0.801 | 5.30888E-14 |
| Contractile | Vezf1 | 0.000320042 | 0.338238 | 0.321 | 0.261 | 1 |
| Contractile | Gstm7 | 3.44352E-14 | 0.338171 | 0.165 | 0.054 | 4.56025E-10 |
| Contractile | Zfp672 | 7.3724E-07 | 0.337602 | 0.203 | 0.116 | 0.009763263 |
| Contractile | Gnb4 | 8.17498E-08 | 0.337574 | 0.456 | 0.343 | 0.001082612 |
| Contractile | Hint2 | 0.00880098 | 0.337546 | 0.4 | 0.389 | 1 |
| Intermediate | Eif6 | 6.80266E-19 | 0.337514 | 0.54 | 0.362 | 9.00876E-15 |
| Intermediate | Pla2g15 | 7.92134E-26 | 0.336955 | 0.276 | 0.107 | 1.04902E-21 |
| Contractile | Spats2l | 7.39274E-08 | 0.336787 | 0.256 | 0.15 | 0.00097902 |
| Terminal | Capn6 | 1.2282E-38 | 0.336557 | 0.165 | 0.022 | 1.62651E-34 |
| Terminal | Bambi | 5.55757E-29 | 0.33648 | 0.397 | 0.175 | 7.35989E-25 |
| Intermediate | Ube2l6 | 6.70297E-27 | 0.336402 | 0.263 | 0.095 | 8.87674E-23 |
| Intermediate | Tmem150a | 4.19148E-19 | 0.33618 | 0.311 | 0.159 | 5.55078E-15 |
| Contractile | Eva1b | 2.75442E-06 | 0.33595 | 0.674 | 0.64 | 0.036476787 |
| Rgs5_Contractile | Cfl1 | 0.000293158 | 0.335868 | 0.747 | 0.886 | 1 |
| Contractile | Fam208a | 3.46514E-05 | 0.335844 | 0.162 | 0.095 | 0.458888512 |
| Contractile | Arhgef12 | 0.000991924 | 0.335838 | 0.312 | 0.256 | 1 |
| Contractile | Son | 1.55955E-10 | 0.334888 | 0.856 | 0.822 | 2.06532E-06 |
| Contractile | Tnrc6a | 6.24881E-05 | 0.334776 | 0.388 | 0.319 | 0.82752935 |
| Contractile | Tcf7l2 | 5.82686E-07 | 0.334583 | 0.297 | 0.197 | 0.007716509 |
| Contractile | Gstp1 | 8.59045E-05 | 0.334268 | 0.397 | 0.332 | 1 |
| Terminal | Fam101a | 4.48238E-39 | 0.334002 | 0.17 | 0.023 | 5.93601E-35 |
| Contractile | Tacc1 | 6.23654E-09 | 0.33379 | 0.388 | 0.256 | 8.25905E-05 |
| Terminal | Gsn | 1.31035E-10 | 0.333284 | 0.955 | 0.866 | 1.7353E-06 |
| Contractile | Manbal | 0.000225188 | 0.333249 | 0.441 | 0.395 | 1 |
| Contractile | Creb3l2 | 0.001480352 | 0.333177 | 0.359 | 0.318 | 1 |
| Intermediate | Spg21 | 1.46451E-17 | 0.333116 | 0.461 | 0.296 | 1.93945E-13 |
| Contractile | Npm1 | 3.43873E-10 | 0.333034 | 0.815 | 0.771 | 4.55391E-06 |
| Intermediate | Cdc42 | 2.3964E-25 | 0.333022 | 0.841 | 0.697 | 3.17355E-21 |
| Contractile | Hspa4 | 0.004317545 | 0.333005 | 0.335 | 0.295 | 1 |
| Intermediate | Arl6ip5 | 1.07163E-17 | 0.333 | 0.828 | 0.725 | 1.41916E-13 |
| Contractile | Blvrb | 0.000524153 | 0.332994 | 0.429 | 0.385 | 1 |
| Intermediate | Dnajb11 | 6.3036E-15 | 0.332965 | 0.577 | 0.415 | 8.34786E-11 |
| Intermediate | Pdlim2 | 2.13472E-16 | 0.332939 | 0.482 | 0.308 | 2.82701E-12 |
| Contractile | Dguok | 0.001598404 | 0.332721 | 0.332 | 0.288 | 1 |
| Intermediate | Akr1b8 | 1.66008E-27 | 0.332707 | 0.268 | 0.097 | 2.19844E-23 |
| Contractile | Cetn3 | 0.000173361 | 0.332273 | 0.588 | 0.578 | 1 |

| | | | | | | |
|---|---|---|---|---|---|---|
| Contractile | Hagh | 1.70765E-05 | 0.332272 | 0.324 | 0.248 | 0.226143998 |
| Intermediate | Lxn | 2.46826E-12 | 0.331985 | 0.708 | 0.579 | 3.26872E-08 |
| Contractile | Col4a5 | 5.10365E-06 | 0.331867 | 0.365 | 0.269 | 0.067587699 |
| Terminal | Emilin1 | 4.27562E-21 | 0.331786 | 0.783 | 0.56 | 5.6622E-17 |
| Contractile | Yipf4 | 1.12646E-05 | 0.33144 | 0.635 | 0.612 | 0.149177037 |
| Contractile | Tecr | 0.000127911 | 0.331343 | 0.641 | 0.644 | 1 |
| Contractile | Mrps6 | 0.000424064 | 0.331063 | 0.274 | 0.21 | 1 |
| Intermediate | Rab34 | 1.24346E-17 | 0.330975 | 0.564 | 0.385 | 1.64672E-13 |
| Intermediate | Arpc5 | 1.23373E-16 | 0.330876 | 0.64 | 0.469 | 1.63382E-12 |
| Contractile | Snrk | 2.0243E-08 | 0.330799 | 0.274 | 0.16 | 0.000268078 |
| Contractile | Snrpd2 | 6.18568E-06 | 0.330698 | 0.732 | 0.716 | 0.08191697 |
| Contractile | Arpp19 | 1.33319E-05 | 0.330609 | 0.694 | 0.675 | 0.176553729 |
| Rgs5_Contractile | Ppia | 5.54168E-12 | 0.330591 | 0.983 | 0.992 | 7.33884E-08 |
| Terminal | Vamp5 | 2.78795E-22 | 0.330543 | 0.507 | 0.28 | 3.69208E-18 |
| Contractile | Lats2 | 1.85947E-05 | 0.33041 | 0.279 | 0.196 | 0.24624985 |
| Intermediate | Qsox1 | 1.61842E-17 | 0.329981 | 0.609 | 0.424 | 2.14328E-13 |
| Contractile | Fyttd1 | 0.001946316 | 0.329692 | 0.285 | 0.236 | 1 |
| Contractile | Rhoa | 1.08333E-10 | 0.329292 | 0.871 | 0.79 | 1.43465E-06 |
| Contractile | Ppp1r18 | 0.000965546 | 0.328992 | 0.318 | 0.267 | 1 |
| Contractile | Itpk1 | 5.29732E-12 | 0.328722 | 0.165 | 0.061 | 7.01523E-08 |
| Contractile | Tm6sf1 | 1.79408E-25 | 0.328716 | 0.138 | 0.021 | 2.3759E-21 |
| Contractile | Ccl27a | 4.12364E-05 | 0.328582 | 0.209 | 0.137 | 0.54609336 |
| Contractile | Cpne2 | 3.44566E-05 | 0.328519 | 0.238 | 0.162 | 0.456309225 |
| Contractile | Acbd4 | 5.67023E-06 | 0.328321 | 0.221 | 0.138 | 0.075090866 |
| Contractile | Ndufb11 | 1.38356E-08 | 0.328209 | 0.779 | 0.752 | 0.000183225 |
| Contractile | Atp5f1 | 2.20656E-06 | 0.328133 | 0.709 | 0.705 | 0.029221421 |
| Intermediate | Lix1l | 6.62933E-16 | 0.327912 | 0.484 | 0.323 | 8.77922E-12 |
| Contractile | Pdha1 | 0.00093148 | 0.327747 | 0.338 | 0.29 | 1 |
| Intermediate | Ahnak | 2.76031E-21 | 0.327537 | 0.841 | 0.699 | 3.65548E-17 |
| Terminal | Vgll4 | 1.34978E-25 | 0.327473 | 0.421 | 0.208 | 1.78751E-21 |
| Intermediate | Aig1 | 2.31599E-15 | 0.327254 | 0.547 | 0.391 | 3.06706E-11 |
| Intermediate | Mfsd1 | 2.3738E-18 | 0.327229 | 0.272 | 0.129 | 3.14363E-14 |
| Contractile | Smarca2 | 0.000163471 | 0.3271 | 0.362 | 0.296 | 1 |
| Intermediate | Mlec | 4.85625E-16 | 0.327038 | 0.425 | 0.267 | 6.43113E-12 |
| Contractile | Yaf2 | 3.86729E-05 | 0.326515 | 0.385 | 0.318 | 0.512145452 |
| Terminal | Cst6 | 4.59282E-18 | 0.326506 | 0.348 | 0.184 | 6.08227E-14 |
| Terminal | Ank | 1.3742E-15 | 0.325759 | 0.58 | 0.385 | 1.81986E-11 |
| Contractile | Rabac1 | 6.4689E-09 | 0.325745 | 0.906 | 0.895 | 8.56676E-05 |
| Intermediate | Rin2 | 6.67321E-14 | 0.325739 | 0.455 | 0.315 | 8.83733E-10 |
| Contractile | Ddx5 | 1.56794E-12 | 0.325693 | 0.932 | 0.919 | 2.07643E-08 |
| Intermediate | Derl2 | 3.05036E-17 | 0.325549 | 0.498 | 0.317 | 4.0396E-13 |
| Contractile | Rbms2 | 0.000109994 | 0.325075 | 0.324 | 0.257 | 1 |
| Contractile | Gk | 3.63367E-14 | 0.324884 | 0.156 | 0.049 | 4.81207E-10 |
| Intermediate | Arpc4 | 9.54891E-16 | 0.324672 | 0.591 | 0.447 | 1.26456E-11 |
| Contractile | Hnrnpu | 0.000391039 | 0.324185 | 0.465 | 0.423 | 1 |
| Contractile | Csnk1a1 | 1.03274E-07 | 0.324145 | 0.682 | 0.641 | 0.001367659 |
| Contractile | Cyb5r3 | 4.29211E-09 | 0.324004 | 0.888 | 0.833 | 5.68405E-05 |
| Contractile | Ssfa2 | 3.97953E-06 | 0.323825 | 0.191 | 0.111 | 0.052700881 |
| Contractile | Mea1 | 0.000647575 | 0.323803 | 0.394 | 0.351 | 1 |
| Intermediate | H13 | 1.8158E-16 | 0.323477 | 0.673 | 0.533 | 2.40466E-12 |
| Contractile | Synpo | 7.43492E-07 | 0.323145 | 0.456 | 0.345 | 0.009846062 |
| Contractile | Ddx3x | 9.88963E-06 | 0.323092 | 0.553 | 0.5 | 0.130968423 |
| Contractile | Polr2m | 4.64809E-05 | 0.322986 | 0.506 | 0.468 | 0.615546439 |
| Intermediate | Elk3 | 4.01413E-19 | 0.322925 | 0.301 | 0.15 | 5.31592E-15 |
| Intermediate | Pdgfc | 1.38372E-14 | 0.322813 | 0.526 | 0.366 | 1.83246E-10 |
| Intermediate | Itga11 | 4.65301E-30 | 0.322782 | 0.124 | 0.014 | 6.16198E-26 |
| Contractile | Eif1ax | 1.39219E-05 | 0.322675 | 0.576 | 0.532 | 0.184367699 |
| Contractile | Impa1 | 0.000293533 | 0.322624 | 0.35 | 0.286 | 1 |
| Intermediate | Cdkn2c | 2.24852E-11 | 0.322618 | 0.385 | 0.26 | 2.97772E-07 |

| | | | | | | |
|---|---|---|---|---|---|---|
| Intermediate | Fam20a | 3.34966E-34 | 0.322468 | 0.187 | 0.037 | 4.43595E-30 |
| Intermediate | Adamts5 | 1.30773E-11 | 0.322318 | 0.258 | 0.146 | 1.73183E-07 |
| Intermediate | Pgls | 1.6476E-14 | 0.32231 | 0.642 | 0.495 | 2.18192E-10 |
| Terminal | Fuca1 | 8.38329E-20 | 0.322228 | 0.822 | 0.575 | 1.1102E-15 |
| Terminal | Slc44a1 | 8.13255E-25 | 0.322179 | 0.439 | 0.229 | 1.07699E-20 |
| Intermediate | Tcf4 | 1.22399E-12 | 0.321994 | 0.647 | 0.493 | 1.62093E-08 |
| Contractile | Fdx1 | 0.000916085 | 0.32199 | 0.344 | 0.292 | 1 |
| Contractile | Tpd52l2 | 0.007522786 | 0.321734 | 0.435 | 0.421 | 1 |
| Contractile | Ddr2 | 0.000120977 | 0.32168 | 0.344 | 0.276 | 1 |
| Contractile | Ttll7 | 1.76676E-05 | 0.321525 | 0.247 | 0.164 | 0.233972409 |
| Contractile | Npepl1 | 0.001607365 | 0.321109 | 0.224 | 0.172 | 1 |
| Intermediate | Sh3pxd2b | 1.30174E-19 | 0.320954 | 0.251 | 0.108 | 1.72389E-15 |
| Contractile | Gde1 | 4.68707E-05 | 0.320901 | 0.485 | 0.425 | 0.620708286 |
| Intermediate | Hn1 | 1.01315E-12 | 0.320892 | 0.353 | 0.225 | 1.34172E-08 |
| Contractile | Churc1 | 5.23395E-05 | 0.320871 | 0.547 | 0.506 | 0.693131561 |
| Rgs5_Contractile | Cox5b | 0.006251514 | 0.320844 | 0.677 | 0.827 | 1 |
| Rgs5_Contractile | Pin4 | 0.009149971 | 0.320834 | 0.226 | 0.376 | 1 |
| Intermediate | Txndc17 | 6.22995E-16 | 0.320621 | 0.771 | 0.638 | 8.25033E-12 |
| Contractile | Mrpl20 | 3.31635E-06 | 0.320539 | 0.635 | 0.595 | 0.043918401 |
| Intermediate | Zeb2 | 2.30671E-20 | 0.320535 | 0.342 | 0.174 | 3.05477E-16 |
| Intermediate | Tmem208 | 1.87394E-13 | 0.320356 | 0.611 | 0.48 | 2.48166E-09 |
| Contractile | Gjc1 | 5.96922E-10 | 0.320303 | 0.206 | 0.096 | 7.90504E-06 |
| Contractile | Dohh | 7.46021E-05 | 0.320216 | 0.341 | 0.27 | 0.987955659 |
| Contractile | Rcn2 | 3.99712E-07 | 0.319977 | 0.753 | 0.708 | 0.005293388 |
| Intermediate | Ccl9 | 9.0948E-11 | 0.319729 | 0.192 | 0.099 | 1.20442E-06 |
| Terminal | Slc39a14 | 2.05898E-14 | 0.319675 | 0.388 | 0.233 | 2.7267E-10 |
| Contractile | Ifrd1 | 1.03561E-05 | 0.319518 | 0.388 | 0.295 | 0.137146249 |
| Intermediate | Naglu | 8.95505E-13 | 0.319384 | 0.364 | 0.233 | 1.18592E-08 |
| Contractile | Ube2d2a | 6.77904E-06 | 0.319077 | 0.612 | 0.567 | 0.089774804 |
| Intermediate | Cib1 | 7.31856E-21 | 0.318819 | 0.437 | 0.255 | 9.69197E-17 |
| Contractile | Kalrn | 9.8008E-11 | 0.318526 | 0.215 | 0.098 | 1.29792E-06 |
| Intermediate | Leprot | 1.83792E-15 | 0.318359 | 0.623 | 0.466 | 2.43396E-11 |
| Contractile | Ube3a | 0.002729269 | 0.318191 | 0.397 | 0.358 | 1 |
| Contractile | Sarnp | 0.000209722 | 0.318113 | 0.603 | 0.582 | 1 |
| Contractile | Pomp | 1.33837E-08 | 0.317974 | 0.765 | 0.732 | 0.000177241 |
| Contractile | Abhd14b | 0.000262897 | 0.317869 | 0.259 | 0.192 | 1 |
| Intermediate | Fam129b | 8.85373E-17 | 0.317651 | 0.465 | 0.298 | 1.1725E-12 |
| Contractile | Grpel1 | 1.68218E-05 | 0.317583 | 0.456 | 0.401 | 0.222770481 |
| Contractile | Timp4 | 6.41549E-18 | 0.31747 | 0.174 | 0.047 | 8.49604E-14 |
| Terminal | Lrrc16a | 3.88806E-27 | 0.317185 | 0.255 | 0.089 | 5.14896E-23 |
| Contractile | Gm561 | 0.003090401 | 0.317068 | 0.347 | 0.307 | 1 |
| Intermediate | Trappc3 | 8.19839E-15 | 0.316777 | 0.44 | 0.283 | 1.08571E-10 |
| Contractile | Rab5b | 9.48532E-07 | 0.316755 | 0.229 | 0.139 | 0.012561412 |
| Contractile | H2-Ke6 | 3.03099E-06 | 0.316741 | 0.341 | 0.248 | 0.040139417 |
| Contractile | Atp5j | 5.86258E-14 | 0.316526 | 0.932 | 0.908 | 7.76382E-10 |
| Intermediate | Jtb | 1.11717E-12 | 0.316471 | 0.53 | 0.379 | 1.47947E-08 |
| Contractile | Dag1 | 2.65134E-05 | 0.316465 | 0.482 | 0.414 | 0.351116561 |
| Intermediate | Cd302 | 1.3358E-12 | 0.316406 | 0.794 | 0.669 | 1.76901E-08 |
| Contractile | Minos1 | 8.70847E-11 | 0.31622 | 0.876 | 0.844 | 1.15326E-06 |
| Contractile | Ndufs2 | 0.00050477 | 0.316181 | 0.524 | 0.512 | 1 |
| Intermediate | Rpn1 | 1.97148E-11 | 0.316168 | 0.594 | 0.46 | 2.61082E-07 |
| Intermediate | Eif4e2 | 3.21933E-16 | 0.316121 | 0.399 | 0.243 | 4.26335E-12 |
| Contractile | Suclg1 | 0.000257556 | 0.315874 | 0.424 | 0.371 | 1 |
| Intermediate | Ap2m1 | 8.63632E-15 | 0.315755 | 0.755 | 0.623 | 1.14371E-10 |
| Contractile | Fas | 0.004930628 | 0.315735 | 0.4 | 0.369 | 1 |
| Intermediate | Tspan4 | 3.52016E-11 | 0.315716 | 0.687 | 0.555 | 4.66175E-07 |
| Contractile | C1qtnf2 | 0.000283472 | 0.315475 | 0.397 | 0.322 | 1 |
| Contractile | Uqcrfs1 | 0.00051286 | 0.315446 | 0.438 | 0.392 | 1 |
| Intermediate | Tmem176a | 5.02385E-12 | 0.315381 | 0.727 | 0.568 | 6.65308E-08 |

| | | | | | | |
|---|---|---|---|---|---|---|
| Terminal | Eps8 | 3.267E-34 | 0.315204 | 0.271 | 0.083 | 4.32649E-30 |
| Contractile | Tpp1 | 0.005459268 | 0.315195 | 0.3 | 0.255 | 1 |
| Contractile | Pin1 | 0.000116502 | 0.315177 | 0.479 | 0.429 | 1 |
| Contractile | Psmd13 | 0.005230148 | 0.315033 | 0.403 | 0.384 | 1 |
| Contractile | Nat6 | 1.3696E-07 | 0.314779 | 0.176 | 0.09 | 0.001813755 |
| Contractile | Bpnt1 | 0.000500667 | 0.314601 | 0.182 | 0.125 | 1 |
| Contractile | Stk39 | 0.000705609 | 0.314509 | 0.179 | 0.123 | 1 |
| Intermediate | Eif1a | 7.04727E-15 | 0.314387 | 0.517 | 0.35 | 9.33269E-11 |
| Contractile | Hnrnph1 | 5.14816E-05 | 0.314318 | 0.485 | 0.426 | 0.681770866 |
| Contractile | Ginm1 | 2.89878E-05 | 0.3143 | 0.538 | 0.498 | 0.383884782 |
| Terminal | Neat1 | 4.12212E-16 | 0.314203 | 0.746 | 0.562 | 5.45892E-12 |
| Contractile | Cuedc2 | 0.002849517 | 0.314182 | 0.453 | 0.441 | 1 |
| Contractile | Alkbh7 | 4.66182E-05 | 0.313973 | 0.197 | 0.127 | 0.617365151 |
| Contractile | Sgf29 | 0.000137234 | 0.313953 | 0.165 | 0.102 | 1 |
| Contractile | Eif2a | 0.001538963 | 0.31382 | 0.324 | 0.275 | 1 |
| Intermediate | Fkbp1a | 4.49649E-21 | 0.313783 | 0.877 | 0.818 | 5.9547E-17 |
| Contractile | Ptpn11 | 0.001739756 | 0.313768 | 0.294 | 0.242 | 1 |
| Rgs5_Contractile | Cuedc1 | 0.004648358 | 0.313519 | 0.153 | 0.266 | 1 |
| Contractile | Ppil1 | 4.20796E-05 | 0.313497 | 0.244 | 0.169 | 0.557260261 |
| Contractile | Hey2 | 1.1146E-09 | 0.313442 | 0.194 | 0.088 | 1.47607E-05 |
| Intermediate | Lamb1 | 1.98556E-29 | 0.313387 | 0.132 | 0.018 | 2.62948E-25 |
| Intermediate | Igsf8 | 5.93207E-14 | 0.313314 | 0.403 | 0.258 | 7.85584E-10 |
| Rgs5_Contractile | Slc25a4 | 1.45142E-05 | 0.313282 | 0.851 | 0.942 | 0.192211355 |
| Intermediate | Nradd | 2.52074E-17 | 0.313163 | 0.252 | 0.12 | 3.33822E-13 |
| Contractile | Msantd4 | 0.000162849 | 0.313143 | 0.182 | 0.119 | 1 |
| Rgs5_Contractile | Hnrnpl | 0.005110383 | 0.313063 | 0.219 | 0.368 | 1 |
| Intermediate | Tgfb2 | 3.12593E-07 | 0.312969 | 0.36 | 0.265 | 0.004139668 |
| Intermediate | Cacna2d1 | 9.57305E-16 | 0.312807 | 0.534 | 0.368 | 1.26776E-11 |
| Intermediate | Rab7 | 3.48252E-18 | 0.312337 | 0.685 | 0.501 | 4.6119E-14 |
| Contractile | Prkaca | 0.003402527 | 0.312288 | 0.321 | 0.28 | 1 |
| Contractile | Gapdh | 1.79609E-07 | 0.31223 | 0.876 | 0.852 | 0.00237856 |
| Intermediate | Shc1 | 3.88765E-14 | 0.312164 | 0.43 | 0.28 | 5.14841E-10 |
| Intermediate | Cttn | 3.52185E-17 | 0.312031 | 0.468 | 0.298 | 4.66398E-13 |
| Intermediate | Ddost | 3.70004E-14 | 0.31195 | 0.723 | 0.586 | 4.89996E-10 |
| Contractile | Fgf13 | 1.63779E-21 | 0.311841 | 0.124 | 0.02 | 2.16892E-17 |
| Contractile | Ccdc85b | 0.006899824 | 0.311606 | 0.212 | 0.17 | 1 |
| Intermediate | Emp2 | 2.0986E-11 | 0.31158 | 0.632 | 0.51 | 2.77918E-07 |
| Intermediate | Kif26b | 1.82223E-21 | 0.311386 | 0.14 | 0.035 | 2.41318E-17 |
| Contractile | Mpp7 | 1.99743E-12 | 0.311242 | 0.118 | 0.033 | 2.6452E-08 |
| Intermediate | Atp6ap2 | 3.0158E-19 | 0.311194 | 0.556 | 0.353 | 3.99383E-15 |
| Terminal | Tspan4 | 7.56961E-20 | 0.311169 | 0.758 | 0.513 | 1.00244E-15 |
| Intermediate | Tmed1 | 8.10238E-16 | 0.311032 | 0.528 | 0.36 | 1.073E-11 |
| Terminal | Cebpb | 6.36956E-05 | 0.310976 | 0.785 | 0.731 | 0.843521156 |
| Terminal | Sgk1 | 4.81353E-16 | 0.310883 | 0.511 | 0.317 | 6.37456E-12 |
| Intermediate | Edf1 | 2.88641E-17 | 0.310873 | 0.87 | 0.821 | 3.82248E-13 |
| Contractile | Acyp2 | 0.000262389 | 0.310783 | 0.312 | 0.247 | 1 |
| Contractile | Sh3pxd2a | 0.00122071 | 0.310706 | 0.35 | 0.3 | 1 |
| Contractile | Lypla1 | 5.32537E-05 | 0.310634 | 0.271 | 0.198 | 0.705239188 |
| Contractile | Zbtb16 | 1.81331E-17 | 0.310523 | 0.129 | 0.028 | 2.40136E-13 |
| Contractile | Wdr89 | 1.93198E-05 | 0.310318 | 0.824 | 0.871 | 0.255851501 |
| Contractile | Srsf3 | 3.09035E-05 | 0.310098 | 0.632 | 0.597 | 0.409254928 |
| Contractile | Hdac5 | 1.8188E-05 | 0.310094 | 0.235 | 0.153 | 0.240863366 |
| Intermediate | Tmem258 | 7.99994E-12 | 0.310022 | 0.779 | 0.667 | 1.05943E-07 |
| Terminal | Sdf4 | 2.49883E-16 | 0.309828 | 0.821 | 0.633 | 3.3092E-12 |
| Terminal | Ppib | 3.04607E-16 | 0.309804 | 0.969 | 0.854 | 4.03391E-12 |
| Contractile | Arpc5l | 0.003165206 | 0.309769 | 0.479 | 0.471 | 1 |
| Intermediate | Eva1b | 8.42749E-18 | 0.309405 | 0.745 | 0.586 | 1.11605E-13 |
| Contractile | Afap1 | 2.60986E-06 | 0.309238 | 0.268 | 0.173 | 0.034562389 |
| Contractile | Sgca | 2.26062E-06 | 0.309184 | 0.144 | 0.072 | 0.029937398 |

| | | | | | | |
|---|---|---|---|---|---|---|
| Contractile | Nfib | 1.63458E-05 | 0.309 | 0.615 | 0.577 | 0.216468075 |
| Contractile | Por | 0.00049278 | 0.308897 | 0.332 | 0.269 | 1 |
| Contractile | Ncoa3 | 5.48407E-06 | 0.308867 | 0.256 | 0.163 | 0.072625474 |
| Contractile | Fbxw2 | 0.000482811 | 0.308764 | 0.371 | 0.32 | 1 |
| Rgs5_Contractile | 2510002D | 0.007692883 | 0.308683 | 0.122 | 0.212 | 1 |
| Contractile | Ryr2 | 1.98191E-18 | 0.308661 | 0.106 | 0.018 | 2.62464E-14 |
| Contractile | Paf1 | 4.35253E-06 | 0.308594 | 0.221 | 0.137 | 0.057640574 |
| Contractile | Mgat4a | 1.19194E-09 | 0.308563 | 0.176 | 0.078 | 1.57848E-05 |
| Rgs5_Contractile | Nrp1 | 0.005993449 | 0.308452 | 0.167 | 0.282 | 1 |
| Contractile | Endod1 | 8.28591E-05 | 0.308436 | 0.35 | 0.276 | 1 |
| Contractile | Efhd2 | 2.89781E-05 | 0.308158 | 0.247 | 0.168 | 0.38375762 |
| Contractile | Inafm2 | 9.16195E-08 | 0.307473 | 0.162 | 0.077 | 0.001213317 |
| Contractile | Btf3 | 1.8321E-10 | 0.307433 | 0.909 | 0.853 | 2.42626E-06 |
| Intermediate | Yif1b | 3.39155E-13 | 0.307426 | 0.566 | 0.429 | 4.49143E-09 |
| Intermediate | Ecscr | 1.15538E-27 | 0.307322 | 0.109 | 0.011 | 1.53007E-23 |
| Contractile | Neurl2 | 4.61544E-13 | 0.307255 | 0.121 | 0.033 | 6.11222E-09 |
| Contractile | Arhgef2 | 0.000730898 | 0.307191 | 0.315 | 0.253 | 1 |
| Intermediate | Tax1bp3 | 2.26018E-19 | 0.307091 | 0.562 | 0.36 | 2.99316E-15 |
| Contractile | Tax1bp1 | 0.003319002 | 0.306715 | 0.474 | 0.455 | 1 |
| Contractile | Syt11 | 4.18321E-06 | 0.306695 | 0.194 | 0.112 | 0.055398265 |
| Contractile | Chmp2a | 6.94846E-06 | 0.306642 | 0.682 | 0.679 | 0.09201849 |
| Terminal | Sesn3 | 8.10547E-25 | 0.30663 | 0.363 | 0.168 | 1.07341E-20 |
| Contractile | Emc9 | 0.000201699 | 0.306469 | 0.191 | 0.127 | 1 |
| Contractile | 9530077C | 2.87939E-13 | 0.306464 | 0.126 | 0.036 | 3.81318E-09 |
| Intermediate | Scx | 2.02099E-14 | 0.306457 | 0.263 | 0.136 | 2.6764E-10 |
| Contractile | Pfdn5 | 1.61356E-10 | 0.30612 | 0.935 | 0.907 | 2.13683E-06 |
| Intermediate | Ak1 | 1.37042E-14 | 0.305748 | 0.537 | 0.376 | 1.81484E-10 |
| Intermediate | H2afz | 7.22836E-08 | 0.305706 | 0.716 | 0.633 | 0.000957252 |
| Contractile | Ube2v2 | 0.000871813 | 0.305527 | 0.435 | 0.396 | 1 |
| Contractile | Svip | 1.19425E-27 | 0.30535 | 0.124 | 0.014 | 1.58154E-23 |
| Intermediate | Clec2d | 2.8778E-25 | 0.305158 | 0.163 | 0.04 | 3.81108E-21 |
| Intermediate | Sqstm1 | 5.93311E-14 | 0.305136 | 0.622 | 0.458 | 7.85722E-10 |
| Intermediate | Lrig1 | 1.38091E-24 | 0.304752 | 0.197 | 0.06 | 1.82874E-20 |
| Contractile | Smdt1 | 3.80479E-06 | 0.304591 | 0.829 | 0.806 | 0.050386868 |
| Contractile | Atp5e | 4.43758E-10 | 0.304561 | 0.912 | 0.854 | 5.87669E-06 |
| Intermediate | Osmr | 4.92389E-13 | 0.304437 | 0.377 | 0.236 | 6.5207E-09 |
| Intermediate | Abcg1 | 2.32828E-23 | 0.304417 | 0.188 | 0.059 | 3.08334E-19 |
| Contractile | Ppig | 0.001781554 | 0.304313 | 0.459 | 0.432 | 1 |
| Contractile | Mical3 | 1.56383E-06 | 0.304216 | 0.185 | 0.102 | 0.020709758 |
| Contractile | Rab9 | 0.000327784 | 0.304119 | 0.306 | 0.243 | 1 |
| Intermediate | Hexb | 3.03023E-15 | 0.303988 | 0.53 | 0.362 | 4.01293E-11 |
| Terminal | Sobp | 4.37849E-39 | 0.303923 | 0.222 | 0.045 | 5.79844E-35 |
| Rgs5_Contractile | Smim11 | 0.008839447 | 0.303802 | 0.358 | 0.604 | 1 |
| Intermediate | Pitpna | 2.3721E-11 | 0.303733 | 0.527 | 0.396 | 3.14137E-07 |
| Intermediate | Mecom | 3.81565E-17 | 0.303699 | 0.306 | 0.158 | 5.05306E-13 |
| Contractile | Ndufa9 | 0.001216721 | 0.303361 | 0.374 | 0.327 | 1 |
| Intermediate | Lgals9 | 5.01051E-25 | 0.303337 | 0.176 | 0.048 | 6.63541E-21 |
| Intermediate | Surf4 | 1.70906E-12 | 0.303324 | 0.556 | 0.412 | 2.26331E-08 |
| Terminal | Ly6e | 9.51853E-12 | 0.302805 | 0.5 | 0.336 | 1.26054E-07 |
| Intermediate | Atp6ap1 | 5.81517E-16 | 0.302717 | 0.688 | 0.53 | 7.70103E-12 |
| Contractile | Tmem65 | 0.00343874 | 0.302522 | 0.279 | 0.23 | 1 |
| Intermediate | Rap1b | 4.02018E-15 | 0.302307 | 0.636 | 0.476 | 5.32393E-11 |
| Contractile | Tmem261 | 2.9983E-05 | 0.302153 | 0.374 | 0.3 | 0.397064898 |
| Intermediate | Arhgap6 | 9.57738E-22 | 0.301956 | 0.163 | 0.047 | 1.26833E-17 |
| Intermediate | Tln2 | 3.44387E-21 | 0.301904 | 0.232 | 0.093 | 4.56072E-17 |
| Contractile | Uqcrq | 1.24194E-08 | 0.301756 | 0.841 | 0.797 | 0.00016447 |
| Rgs5_Contractile | Sec24d | 0.00499781 | 0.301755 | 0.198 | 0.338 | 1 |
| Contractile | Zfp771 | 0.00318083 | 0.301385 | 0.344 | 0.308 | 1 |
| Intermediate | Ldha | 2.31114E-16 | 0.300898 | 0.915 | 0.829 | 3.06064E-12 |

| | | | | | | |
|---|---|---|---|---|---|---|
| Intermediate | Aspscr1 | 5.40471E-16 | 0.30087 | 0.322 | 0.177 | 7.15746E-12 |
| Contractile | Gm19705 | 4.18734E-07 | 0.300846 | 0.168 | 0.085 | 0.005545297 |
| Contractile | Btc | 1.20284E-21 | 0.300724 | 0.1 | 0.013 | 1.59292E-17 |
| Contractile | Fyco1 | 7.82636E-07 | 0.300248 | 0.221 | 0.127 | 0.010364447 |
| Contractile | Sphkap | 5.3405E-35 | 0.300073 | 0.103 | 0.005 | 7.07242E-31 |
| Contractile | Pnn | 0.005059526 | 0.300001 | 0.341 | 0.306 | 1 |
| Contractile | Hiat1 | 0.004889721 | 0.299989 | 0.206 | 0.159 | 1 |
| Intermediate | Ifngr2 | 5.3695E-18 | 0.299657 | 0.454 | 0.273 | 7.11082E-14 |
| Intermediate | Vcp | 8.19054E-14 | 0.299496 | 0.71 | 0.574 | 1.08467E-09 |
| Contractile | Unc119 | 5.43666E-08 | 0.299416 | 0.182 | 0.091 | 0.000719976 |
| Contractile | Fam46b | 6.15132E-24 | 0.299068 | 0.124 | 0.018 | 8.14619E-20 |
| Rgs5_Contractile | Eef1a1 | 1.76301E-07 | 0.298633 | 0.965 | 0.987 | 0.002334755 |
| Contractile | Dnajc4 | 0.003268591 | 0.298492 | 0.262 | 0.216 | 1 |
| Contractile | Cisd1 | 0.001237249 | 0.298477 | 0.479 | 0.469 | 1 |
| Contractile | P2rx5 | 2.49597E-17 | 0.297972 | 0.109 | 0.021 | 3.30541E-13 |
| Intermediate | Napa | 4.81781E-13 | 0.297816 | 0.583 | 0.441 | 6.38023E-09 |
| Contractile | Vstm4 | 7.2721E-12 | 0.297706 | 0.118 | 0.034 | 9.63044E-08 |
| Intermediate | Gabarapl1 | 5.59338E-11 | 0.297414 | 0.493 | 0.351 | 7.40731E-07 |
| Intermediate | Snx7 | 6.74318E-15 | 0.297374 | 0.378 | 0.233 | 8.93E-11 |
| Intermediate | Erh | 6.02611E-14 | 0.297293 | 0.763 | 0.661 | 7.98038E-10 |
| Contractile | Mtpn | 0.000820376 | 0.297275 | 0.406 | 0.355 | 1 |
| Contractile | Plekhg3 | 1.87667E-08 | 0.297127 | 0.159 | 0.071 | 0.000248527 |
| Intermediate | Eif4a1 | 7.78716E-17 | 0.296978 | 0.893 | 0.802 | 1.03125E-12 |
| Intermediate | P4ha3 | 2.0734E-29 | 0.296942 | 0.163 | 0.034 | 2.7458E-25 |
| Intermediate | Igfbp6 | 0.00036351 | 0.296934 | 0.198 | 0.143 | 1 |
| Contractile | Large | 1.53503E-06 | 0.296888 | 0.215 | 0.126 | 0.020328425 |
| Contractile | Hsbp1 | 1.96255E-06 | 0.29675 | 0.8 | 0.763 | 0.025990102 |
| Rgs5_Contractile | Rsbn1l | 0.006964976 | 0.296707 | 0.142 | 0.244 | 1 |
| Contractile | Setd6 | 7.45977E-05 | 0.296608 | 0.135 | 0.077 | 0.987897999 |
| Intermediate | Sepp1 | 7.72185E-06 | 0.296593 | 0.611 | 0.518 | 0.10226041 |
| Terminal | Grin2a | 1.82242E-19 | 0.296548 | 0.345 | 0.176 | 2.41343E-15 |
| Contractile | Sacs | 1.77229E-13 | 0.296457 | 0.115 | 0.029 | 2.34704E-09 |
| Intermediate | Tomm5 | 3.0182E-14 | 0.296404 | 0.603 | 0.453 | 3.997E-10 |
| Rgs5_Contractile | Lims1 | 0.00477721 | 0.29635 | 0.205 | 0.35 | 1 |
| Terminal | Ccdc12 | 1.07435E-16 | 0.296347 | 0.667 | 0.46 | 1.42277E-12 |
| Contractile | Cab39 | 0.002585177 | 0.296258 | 0.294 | 0.247 | 1 |
| Intermediate | Pttg1ip | 1.79917E-13 | 0.29583 | 0.549 | 0.403 | 2.38263E-09 |
| Contractile | Ewsr1 | 0.00042598 | 0.295609 | 0.474 | 0.43 | 1 |
| Intermediate | Tma7 | 1.37201E-13 | 0.295422 | 0.823 | 0.723 | 1.81695E-09 |
| Contractile | Cdc42 | 4.22106E-05 | 0.295333 | 0.768 | 0.748 | 0.55899505 |
| Contractile | Mettl24 | 1.95615E-10 | 0.295283 | 0.141 | 0.052 | 2.59053E-06 |
| Terminal | P3h4 | 2.66412E-13 | 0.295033 | 0.636 | 0.447 | 3.52809E-09 |
| Contractile | Cct4 | 4.06959E-05 | 0.294951 | 0.556 | 0.514 | 0.53893571 |
| Terminal | Lrg1 | 5.4142E-33 | 0.294946 | 0.481 | 0.221 | 7.17002E-29 |
| Contractile | Aff4 | 0.000921059 | 0.294854 | 0.329 | 0.27 | 1 |
| Contractile | Crip2 | 3.83602E-07 | 0.294774 | 0.865 | 0.833 | 0.00508004 |
| Contractile | Tbca | 3.75229E-08 | 0.294649 | 0.776 | 0.726 | 0.000496915 |
| Contractile | Sirt3 | 0.000304404 | 0.294554 | 0.171 | 0.111 | 1 |
| Contractile | Got2 | 0.002211331 | 0.294504 | 0.247 | 0.196 | 1 |
| Intermediate | Minos1 | 1.49318E-16 | 0.294413 | 0.894 | 0.821 | 1.97742E-12 |
| Contractile | Gm9493 | 1.23033E-05 | 0.293888 | 0.803 | 0.829 | 0.162932134 |
| Contractile | 2810474O | 0.001720438 | 0.293801 | 0.321 | 0.268 | 1 |
| Intermediate | Pfkl | 4.92713E-10 | 0.293017 | 0.366 | 0.244 | 6.52499E-06 |
| Contractile | Uchl5 | 0.001155548 | 0.292955 | 0.212 | 0.156 | 1 |
| Intermediate | Sra1 | 2.5909E-09 | 0.292598 | 0.634 | 0.522 | 3.43113E-05 |
| Rgs5_Contractile | Lsm3 | 0.002454778 | 0.292536 | 0.184 | 0.322 | 1 |
| Contractile | Flot2 | 0.004262536 | 0.292346 | 0.268 | 0.222 | 1 |
| Terminal | Tmbim4 | 4.22489E-20 | 0.292174 | 0.66 | 0.418 | 5.59502E-16 |
| Contractile | Tceal3 | 1.90805E-13 | 0.292103 | 0.153 | 0.049 | 2.52683E-09 |

| | | | | | | |
|---|---|---|---|---|---|---|
| Rgs5_Contractile | Ogt | 0.002624887 | 0.291568 | 0.212 | 0.367 | 1 |
| Intermediate | Ablim1 | 3.86735E-17 | 0.291539 | 0.161 | 0.056 | 5.12153E-13 |
| Contractile | Trabd2b | 0.00080864 | 0.291479 | 0.235 | 0.172 | 1 |
| Contractile | Atp8b1 | 8.80475E-08 | 0.29143 | 0.2 | 0.103 | 0.001166014 |
| Intermediate | Runx1 | 4.93673E-10 | 0.291351 | 0.462 | 0.329 | 6.53771E-06 |
| Contractile | Chmp5 | 0.000157778 | 0.291277 | 0.485 | 0.439 | 1 |
| Contractile | 2210013O | 0.00023029 | 0.291216 | 0.485 | 0.445 | 1 |
| Terminal | Icam1 | 1.59872E-22 | 0.291115 | 0.367 | 0.181 | 2.11719E-18 |
| Intermediate | Hspg2 | 3.46742E-11 | 0.290633 | 0.701 | 0.569 | 4.59191E-07 |
| Intermediate | Inhbb | 1.57529E-16 | 0.290585 | 0.16 | 0.058 | 2.08616E-12 |
| Intermediate | Lman2 | 1.07433E-12 | 0.290549 | 0.572 | 0.437 | 1.42274E-08 |
| Contractile | Vasn | 0.001220016 | 0.290224 | 0.506 | 0.494 | 1 |
| Contractile | Psmb5 | 3.28904E-05 | 0.290025 | 0.665 | 0.632 | 0.435567553 |
| Intermediate | Psmd8 | 1.25316E-10 | 0.290021 | 0.638 | 0.52 | 1.65956E-06 |
| Intermediate | Pik3r1 | 1.07276E-10 | 0.289966 | 0.445 | 0.311 | 1.42065E-06 |
| Contractile | Itgb5 | 1.65851E-08 | 0.289726 | 0.774 | 0.713 | 0.000219637 |
| Contractile | Mn1 | 9.69596E-08 | 0.289688 | 0.168 | 0.08 | 0.001284036 |
| Contractile | Pfdn4 | 0.001310334 | 0.289531 | 0.379 | 0.341 | 1 |
| Contractile | AI838599 | 6.26201E-09 | 0.289484 | 0.168 | 0.074 | 8.29278E-05 |
| Intermediate | Glrx5 | 6.13053E-11 | 0.289372 | 0.47 | 0.343 | 8.11867E-07 |
| Contractile | Nae1 | 0.000422829 | 0.289209 | 0.291 | 0.229 | 1 |
| Contractile | Lamtor4 | 0.000423336 | 0.289082 | 0.565 | 0.545 | 1 |
| Intermediate | Ctsa | 4.07813E-11 | 0.288904 | 0.632 | 0.507 | 5.40067E-07 |
| Contractile | Vegfb | 3.95456E-05 | 0.288849 | 0.168 | 0.1 | 0.523702831 |
| Terminal | Col16a1 | 3.08103E-13 | 0.288694 | 0.29 | 0.162 | 4.08021E-09 |
| Rgs5_Contractile | Csnk2a1 | 0.006627761 | 0.288659 | 0.194 | 0.331 | 1 |
| Contractile | Crip1 | 2.55915E-17 | 0.288496 | 0.956 | 0.939 | 3.38908E-13 |
| Contractile | Pcsk5 | 5.22433E-12 | 0.288481 | 0.129 | 0.04 | 6.91858E-08 |
| Contractile | Ehd1 | 0.003180629 | 0.288362 | 0.244 | 0.191 | 1 |
| Contractile | Arid4b | 0.004113917 | 0.288361 | 0.3 | 0.257 | 1 |
| Intermediate | Kcnj15 | 8.63036E-15 | 0.288356 | 0.188 | 0.08 | 1.14292E-10 |
| Rgs5_Contractile | Sike1 | 0.009466625 | 0.288338 | 0.108 | 0.187 | 1 |
| Intermediate | Il6st | 4.45523E-12 | 0.288157 | 0.52 | 0.375 | 5.90006E-08 |
| Intermediate | Ap2s1 | 2.45193E-15 | 0.288132 | 0.605 | 0.437 | 3.24709E-11 |
| Intermediate | Sat1 | 6.40047E-12 | 0.288065 | 0.751 | 0.616 | 8.47614E-08 |
| Contractile | Irf2bp2 | 7.25057E-05 | 0.28805 | 0.274 | 0.195 | 0.960192777 |
| Contractile | Gkap1 | 0.002118337 | 0.287999 | 0.197 | 0.144 | 1 |
| Contractile | Use1 | 3.1828E-06 | 0.287996 | 0.676 | 0.628 | 0.042149781 |
| Contractile | Ccdc186 | 0.000670269 | 0.287967 | 0.221 | 0.161 | 1 |
| Intermediate | Capns1 | 3.75057E-23 | 0.287841 | 0.905 | 0.821 | 4.96688E-19 |
| Intermediate | B3gnt9 | 1.37581E-11 | 0.287687 | 0.456 | 0.324 | 1.82198E-07 |
| Intermediate | Ptpn1 | 1.01154E-11 | 0.2873 | 0.427 | 0.294 | 1.33958E-07 |
| Contractile | Raf1 | 0.000674445 | 0.287239 | 0.318 | 0.261 | 1 |
| Intermediate | Tubb6 | 2.40877E-11 | 0.287229 | 0.347 | 0.219 | 3.18994E-07 |
| Contractile | Sat1 | 7.24367E-05 | 0.286884 | 0.718 | 0.657 | 0.959278792 |
| Contractile | Anapc11 | 5.73053E-05 | 0.286847 | 0.715 | 0.683 | 0.758894629 |
| Intermediate | Unc5b | 3.35757E-17 | 0.286847 | 0.209 | 0.089 | 4.44644E-13 |
| Contractile | Tonsl | 6.59428E-05 | 0.286775 | 0.615 | 0.603 | 0.873280935 |
| Intermediate | Mocs2 | 5.6032E-15 | 0.286686 | 0.396 | 0.248 | 7.42031E-11 |
| Intermediate | Cyba | 8.90473E-15 | 0.286568 | 0.776 | 0.658 | 1.17925E-10 |
| Intermediate | Fam3c | 1.54112E-12 | 0.286291 | 0.545 | 0.395 | 2.0409E-08 |
| Rgs5_Contractile | Pa2g4 | 0.002191585 | 0.286261 | 0.198 | 0.351 | 1 |
| Intermediate | Ifitm2 | 2.73862E-18 | 0.286234 | 0.955 | 0.94 | 3.62676E-14 |
| Terminal | Ap3s1 | 1.45468E-18 | 0.286177 | 0.689 | 0.465 | 1.92643E-14 |
| Terminal | Dbp | 3.62041E-12 | 0.28616 | 0.306 | 0.177 | 4.79451E-08 |
| Intermediate | Cdkn2a | 4.39549E-08 | 0.286142 | 0.201 | 0.121 | 0.000582095 |
| Rgs5_Contractile | Fkbp4 | 0.004717304 | 0.286078 | 0.229 | 0.395 | 1 |
| Intermediate | Glud1 | 9.41588E-17 | 0.285835 | 0.526 | 0.344 | 1.24694E-12 |
| Contractile | Dnajb6 | 0.000743315 | 0.285825 | 0.532 | 0.509 | 1 |

| | | | | | | |
|---|---|---|---|---|---|---|
| Intermediate | Rhbdf1 | 5.33303E-16 | 0.285714 | 0.254 | 0.124 | 7.06254E-12 |
| Contractile | Bmpr2 | 0.001562928 | 0.28555 | 0.276 | 0.221 | 1 |
| Contractile | Gm20186 | 1.09047E-12 | 0.285434 | 0.141 | 0.044 | 1.44411E-08 |
| Contractile | Chchd7 | 0.001432704 | 0.285372 | 0.262 | 0.209 | 1 |
| Contractile | Tspan9 | 8.39904E-10 | 0.285316 | 0.176 | 0.076 | 1.11228E-05 |
| Contractile | Snrpn | 0.000100932 | 0.285311 | 0.232 | 0.161 | 1 |
| Intermediate | Sptbn1 | 4.46252E-14 | 0.285263 | 0.603 | 0.433 | 5.90972E-10 |
| Intermediate | Kdelr1 | 8.06489E-16 | 0.285178 | 0.659 | 0.507 | 1.06803E-11 |
| Intermediate | Slc25a5 | 4.53394E-13 | 0.285023 | 0.824 | 0.75 | 6.0043E-09 |
| Contractile | Bag2 | 2.47212E-05 | 0.28501 | 0.168 | 0.098 | 0.327382445 |
| Contractile | Sgcb | 0.001989235 | 0.284988 | 0.359 | 0.307 | 1 |
| Contractile | Ogdh | 0.001547971 | 0.284935 | 0.335 | 0.287 | 1 |
| Contractile | Fam118a | 3.3251E-07 | 0.284911 | 0.124 | 0.054 | 0.004403433 |
| Contractile | Wbp2 | 0.004756406 | 0.284854 | 0.385 | 0.349 | 1 |
| Contractile | Rsbn1l | 0.000949745 | 0.284664 | 0.282 | 0.224 | 1 |
| Intermediate | Bscl2 | 1.13787E-12 | 0.28443 | 0.435 | 0.292 | 1.50689E-08 |
| Contractile | Fuom | 0.000640065 | 0.284302 | 0.203 | 0.144 | 1 |
| Intermediate | Pxdn | 1.23923E-19 | 0.284248 | 0.18 | 0.061 | 1.64111E-15 |
| Contractile | Psmc2 | 0.002666681 | 0.284159 | 0.453 | 0.436 | 1 |
| Intermediate | Meox2 | 1.8173E-19 | 0.284029 | 0.102 | 0.019 | 2.40665E-15 |
| Contractile | Strn3 | 0.003607115 | 0.284008 | 0.353 | 0.315 | 1 |
| Contractile | Gm6133 | 1.59161E-05 | 0.28399 | 0.579 | 0.547 | 0.210777139 |
| Contractile | Map4 | 0.003432725 | 0.283723 | 0.338 | 0.295 | 1 |
| Terminal | Tubb2b | 4.12314E-18 | 0.283696 | 0.162 | 0.055 | 5.46027E-14 |
| Contractile | G3bp2 | 0.005711101 | 0.283659 | 0.344 | 0.306 | 1 |
| Contractile | Rpain | 0.001080627 | 0.283592 | 0.218 | 0.159 | 1 |
| Contractile | Eci1 | 0.005504637 | 0.283535 | 0.297 | 0.253 | 1 |
| Intermediate | Arl1 | 3.69831E-12 | 0.283501 | 0.761 | 0.642 | 4.89767E-08 |
| Intermediate | Flot1 | 3.52024E-12 | 0.283403 | 0.474 | 0.331 | 4.66185E-08 |
| Contractile | Cops6 | 0.000162672 | 0.283308 | 0.582 | 0.557 | 1 |
| Terminal | Arpc1b | 2.14439E-17 | 0.283228 | 0.98 | 0.872 | 2.83981E-13 |
| Intermediate | Ptprs | 9.26001E-12 | 0.283179 | 0.475 | 0.332 | 1.2263E-07 |
| Contractile | Cct8 | 0.000665091 | 0.283022 | 0.524 | 0.505 | 1 |
| Intermediate | Eif2s1 | 9.33369E-13 | 0.283002 | 0.395 | 0.256 | 1.23606E-08 |
| Contractile | Ppie | 0.005576627 | 0.282962 | 0.288 | 0.247 | 1 |
| Contractile | Fzd7 | 8.64009E-06 | 0.28296 | 0.188 | 0.11 | 0.114420665 |
| Terminal | Tnfrsf23 | 3.36151E-24 | 0.282847 | 0.201 | 0.064 | 4.45164E-20 |
| Intermediate | Gabarap | 4.79681E-18 | 0.282793 | 0.934 | 0.905 | 6.35241E-14 |
| Contractile | Nr1h2 | 0.001124701 | 0.282658 | 0.329 | 0.275 | 1 |
| Contractile | Smpdl3a | 0.008808431 | 0.282619 | 0.479 | 0.47 | 1 |
| Intermediate | Rnf145 | 2.06345E-16 | 0.282576 | 0.223 | 0.1 | 2.73263E-12 |
| Contractile | Nmt1 | 0.006710785 | 0.282298 | 0.379 | 0.352 | 1 |
| Contractile | Oard1 | 0.003982524 | 0.282022 | 0.25 | 0.203 | 1 |
| Intermediate | 9530068E( | 8.08528E-12 | 0.281756 | 0.685 | 0.556 | 1.07073E-07 |
| Contractile | Pdzrn3 | 0.005882816 | 0.281685 | 0.209 | 0.163 | 1 |
| Intermediate | Kazald1 | 9.58226E-10 | 0.281648 | 0.461 | 0.34 | 1.26898E-05 |
| Intermediate | Tmem9b | 2.06618E-10 | 0.281643 | 0.44 | 0.312 | 2.73625E-06 |
| Intermediate | Dnajc3 | 1.3379E-12 | 0.281374 | 0.704 | 0.553 | 1.77178E-08 |
| Contractile | Psmc5 | 0.000249664 | 0.281373 | 0.512 | 0.471 | 1 |
| Terminal | Ccnl1 | 1.81451E-19 | 0.281372 | 0.554 | 0.352 | 2.40296E-15 |
| Contractile | Aimp2 | 0.000436535 | 0.280985 | 0.247 | 0.183 | 1 |
| Rgs5_Contractile | Sf3b2 | 0.002065404 | 0.280922 | 0.33 | 0.573 | 1 |
| Rgs5_Contractile | Hook3 | 0.000566208 | 0.280769 | 0.132 | 0.255 | 1 |
| Intermediate | Ran | 4.69297E-11 | 0.280546 | 0.735 | 0.645 | 6.2149E-07 |
| Intermediate | Slc39a7 | 2.84569E-12 | 0.28023 | 0.482 | 0.341 | 3.76854E-08 |
| Intermediate | Cfl1 | 1.2361E-18 | 0.280168 | 0.913 | 0.843 | 1.63696E-14 |
| Intermediate | Gpr153 | 4.51056E-24 | 0.279892 | 0.214 | 0.072 | 5.97334E-20 |
| Contractile | Emc6 | 0.009805247 | 0.279675 | 0.391 | 0.371 | 1 |
| Intermediate | Prrx1 | 2.00248E-12 | 0.279655 | 0.48 | 0.329 | 2.65188E-08 |

| | | | | | | |
|---|---|---|---|---|---|---|
| Intermediate | Bin1 | 6.80835E-13 | 0.279523 | 0.372 | 0.235 | 9.01629E-09 |
| Contractile | Stom | 2.66505E-06 | 0.279495 | 0.206 | 0.12 | 0.0352932 |
| Contractile | Atl3 | 3.39528E-05 | 0.279481 | 0.182 | 0.112 | 0.449637464 |
| Terminal | Adgrg2 | 1.4034E-26 | 0.279412 | 0.202 | 0.06 | 1.85852E-22 |
| Intermediate | Nrp2 | 2.78569E-19 | 0.279249 | 0.22 | 0.089 | 3.68909E-15 |
| Contractile | Ndufb6 | 2.43052E-05 | 0.279248 | 0.659 | 0.651 | 0.321874333 |
| Contractile | Psma6 | 0.000713732 | 0.279103 | 0.544 | 0.529 | 1 |
| Contractile | Abi1 | 0.000483288 | 0.279093 | 0.256 | 0.192 | 1 |
| Intermediate | Aen | 9.79661E-13 | 0.279008 | 0.237 | 0.125 | 1.29737E-08 |
| Contractile | Pxmp4 | 0.000123849 | 0.279001 | 0.215 | 0.145 | 1 |
| Contractile | Trak2 | 3.38207E-06 | 0.278747 | 0.194 | 0.112 | 0.04478869 |
| Intermediate | Nav1 | 5.39278E-14 | 0.278712 | 0.346 | 0.204 | 7.14166E-10 |
| Intermediate | Adprh | 8.83329E-16 | 0.278585 | 0.403 | 0.242 | 1.16979E-11 |
| Intermediate | Mpv17 | 2.11053E-17 | 0.278444 | 0.384 | 0.217 | 2.79498E-13 |
| Contractile | Eif3g | 0.000829202 | 0.278372 | 0.529 | 0.51 | 1 |
| Contractile | Eif1b | 0.001894852 | 0.278302 | 0.509 | 0.499 | 1 |
| Intermediate | Ggct | 9.36895E-08 | 0.278262 | 0.236 | 0.151 | 0.001240729 |
| Contractile | Nsmaf | 3.48122E-05 | 0.278205 | 0.118 | 0.06 | 0.461018591 |
| Terminal | Nim1k | 4.84106E-27 | 0.278182 | 0.184 | 0.048 | 6.41101E-23 |
| Contractile | Ctdsp2 | 0.000318361 | 0.278046 | 0.406 | 0.347 | 1 |
| Intermediate | Tfpi | 3.07808E-23 | 0.277322 | 0.151 | 0.037 | 4.0763E-19 |
| Rgs5_Contractile | Kpna4 | 0.007012216 | 0.277317 | 0.153 | 0.261 | 1 |
| Intermediate | Ctps | 1.85535E-11 | 0.277182 | 0.227 | 0.121 | 2.45704E-07 |
| Contractile | Foxp1 | 8.68823E-05 | 0.277175 | 0.606 | 0.569 | 1 |
| Contractile | Gnb2 | 2.38683E-07 | 0.277117 | 0.803 | 0.762 | 0.003160878 |
| Intermediate | Mtpn | 3.85557E-15 | 0.277111 | 0.469 | 0.3 | 5.10593E-11 |
| Contractile | Mef2c | 4.94463E-06 | 0.276713 | 0.703 | 0.645 | 0.065481708 |
| Terminal | Cetn4 | 3.61434E-25 | 0.276703 | 0.261 | 0.098 | 4.78647E-21 |
| Contractile | Insr | 5.19676E-10 | 0.276338 | 0.153 | 0.06 | 6.88206E-06 |
| Rgs5_Contractile | Foxo1 | 0.006655459 | 0.276093 | 0.125 | 0.218 | 1 |
| Rgs5_Contractile | Bptf | 0.007215863 | 0.276013 | 0.163 | 0.276 | 1 |
| Contractile | 1500009L1 | 0.001426683 | 0.275952 | 0.188 | 0.132 | 1 |
| Contractile | Ntrk3 | 2.22059E-07 | 0.275905 | 0.306 | 0.189 | 0.002940723 |
| Contractile | Mark1 | 1.90416E-07 | 0.275748 | 0.209 | 0.112 | 0.00252168 |
| Contractile | Wscd2 | 6.59683E-08 | 0.275693 | 0.129 | 0.054 | 0.000873618 |
| Intermediate | Ndfip1 | 1.68776E-13 | 0.275578 | 0.311 | 0.18 | 2.2351E-09 |
| Contractile | Psmb4 | 0.00045623 | 0.275469 | 0.582 | 0.551 | 1 |
| Contractile | Samm50 | 0.005508317 | 0.275326 | 0.341 | 0.305 | 1 |
| Contractile | Hist1h2bc | 7.37332E-07 | 0.275097 | 0.812 | 0.723 | 0.009764492 |
| Intermediate | Psmb10 | 3.78837E-12 | 0.275075 | 0.252 | 0.141 | 5.01694E-08 |
| Contractile | Etfb | 0.00021656 | 0.275057 | 0.521 | 0.486 | 1 |
| Intermediate | Bola2 | 8.36583E-13 | 0.274965 | 0.597 | 0.448 | 1.10789E-08 |
| Rgs5_Contractile | Morf4l1 | 1.64651E-05 | 0.274914 | 0.872 | 0.949 | 0.218047895 |
| Intermediate | Metrnl | 2.18273E-10 | 0.274887 | 0.52 | 0.386 | 2.89059E-06 |
| Rgs5_Contractile | Polr2j | 0.002844311 | 0.274708 | 0.257 | 0.452 | 1 |
| Intermediate | Akr1a1 | 9.98215E-16 | 0.274623 | 0.861 | 0.806 | 1.32194E-11 |
| Intermediate | Sertad4 | 6.23179E-10 | 0.274571 | 0.284 | 0.179 | 8.25276E-06 |
| Intermediate | Arf1 | 2.37585E-15 | 0.274477 | 0.852 | 0.758 | 3.14634E-11 |
| Contractile | Zbtb38 | 0.000160708 | 0.274468 | 0.459 | 0.394 | 1 |
| Intermediate | Lhfp | 4.48664E-15 | 0.274435 | 0.879 | 0.76 | 5.94166E-11 |
| Terminal | Ostc | 1.26443E-15 | 0.273757 | 0.842 | 0.623 | 1.67449E-11 |
| Terminal | Pigp | 1.00131E-15 | 0.273723 | 0.729 | 0.522 | 1.32604E-11 |
| Contractile | Ubxn2a | 0.000203251 | 0.273688 | 0.224 | 0.154 | 1 |
| Intermediate | Rala | 2.64803E-13 | 0.273621 | 0.366 | 0.229 | 3.50679E-09 |
| Contractile | Zeb1 | 0.00026296 | 0.273598 | 0.353 | 0.284 | 1 |
| Intermediate | Ahr | 1.0127E-16 | 0.273597 | 0.179 | 0.07 | 1.34111E-12 |
| Contractile | Mrpl14 | 0.005466623 | 0.273493 | 0.459 | 0.448 | 1 |
| Rgs5_Contractile | Rabac1 | 0.006256951 | 0.273404 | 0.771 | 0.914 | 1 |
| Terminal | Chst12 | 4.49646E-19 | 0.273357 | 0.553 | 0.344 | 5.95466E-15 |

| | | | | | | |
|---|---|---|---|---|---|---|
| Contractile | Surf1 | 0.005254583 | 0.273003 | 0.356 | 0.324 | 1 |
| Intermediate | Man2b1 | 1.05071E-11 | 0.272994 | 0.31 | 0.186 | 1.39145E-07 |
| Intermediate | Eif3c | 2.31746E-12 | 0.272969 | 0.577 | 0.424 | 3.06901E-08 |
| Intermediate | Gjb5 | 1.08108E-26 | 0.272958 | 0.153 | 0.032 | 1.43167E-22 |
| Contractile | Gdi1 | 0.00523383 | 0.27292 | 0.291 | 0.246 | 1 |
| Terminal | Pla1a | 1.13981E-20 | 0.272826 | 0.304 | 0.141 | 1.50945E-16 |
| Contractile | Thoc7 | 0.001914009 | 0.272646 | 0.503 | 0.477 | 1 |
| Contractile | Fh1 | 0.006140017 | 0.272585 | 0.244 | 0.196 | 1 |
| Contractile | Stx1a | 8.93008E-10 | 0.27256 | 0.118 | 0.04 | 1.18261E-05 |
| Contractile | Gata6 | 0.000129522 | 0.272487 | 0.385 | 0.311 | 1 |
| Intermediate | Nars | 1.40344E-10 | 0.272297 | 0.494 | 0.354 | 1.85858E-06 |
| Contractile | Agl | 1.10712E-10 | 0.272287 | 0.191 | 0.082 | 1.46615E-06 |
| Contractile | Hsdl2 | 0.000647209 | 0.272242 | 0.171 | 0.114 | 1 |
| Contractile | Asah1 | 0.006864332 | 0.272222 | 0.453 | 0.417 | 1 |
| Contractile | Nek9 | 0.001037699 | 0.272102 | 0.209 | 0.152 | 1 |
| Contractile | Mef2d | 0.001259285 | 0.27209 | 0.221 | 0.165 | 1 |
| Contractile | Mfap1b | 0.005287583 | 0.27202 | 0.229 | 0.182 | 1 |
| Terminal | Dap | 1.09022E-16 | 0.27199 | 0.883 | 0.66 | 1.44378E-12 |
| Intermediate | Lrrc59 | 3.24233E-12 | 0.271919 | 0.461 | 0.326 | 4.29382E-08 |
| Intermediate | Mrps33 | 4.08597E-11 | 0.271883 | 0.602 | 0.47 | 5.41105E-07 |
| Rgs5_Contractile | Clpp | 0.006871149 | 0.271808 | 0.194 | 0.327 | 1 |
| Contractile | Fbxo9 | 0.00050237 | 0.271581 | 0.335 | 0.273 | 1 |
| Contractile | Klf7 | 0.000238707 | 0.271405 | 0.271 | 0.198 | 1 |
| Contractile | Fam110b | 4.3741E-07 | 0.271212 | 0.15 | 0.072 | 0.005792626 |
| Contractile | Picalm | 0.000121361 | 0.271183 | 0.465 | 0.405 | 1 |
| Contractile | Fam162a | 0.000864241 | 0.270819 | 0.591 | 0.585 | 1 |
| Contractile | Serf2 | 5.26341E-12 | 0.270734 | 0.991 | 0.978 | 6.97034E-08 |
| Contractile | Ndn | 1.65342E-14 | 0.270505 | 0.109 | 0.025 | 2.18963E-10 |
| Contractile | Phip | 0.000780862 | 0.270495 | 0.282 | 0.22 | 1 |
| Contractile | Clybl | 0.00125501 | 0.27048 | 0.179 | 0.125 | 1 |
| Terminal | Herpud1 | 5.72649E-17 | 0.270427 | 0.687 | 0.471 | 7.5836E-13 |
| Contractile | Tmtc1 | 3.92034E-07 | 0.270388 | 0.226 | 0.13 | 0.005191705 |
| Intermediate | Bid | 2.44592E-24 | 0.270356 | 0.198 | 0.062 | 3.23913E-20 |
| Contractile | Ankrd10 | 0.000119384 | 0.270205 | 0.212 | 0.14 | 1 |
| Terminal | Gale | 2.06058E-19 | 0.270117 | 0.196 | 0.074 | 2.72882E-15 |
| Intermediate | Gapdh | 2.21547E-13 | 0.270107 | 0.893 | 0.834 | 2.93395E-09 |
| Contractile | Tm9sf2 | 0.00471471 | 0.269932 | 0.518 | 0.498 | 1 |
| Terminal | Ildr2 | 3.84564E-18 | 0.269896 | 0.238 | 0.104 | 5.09278E-14 |
| Intermediate | Anpep | 1.15305E-13 | 0.269734 | 0.159 | 0.066 | 1.52698E-09 |
| Contractile | Acyp1 | 0.005880615 | 0.269727 | 0.2 | 0.155 | 1 |
| Terminal | Olfml3 | 1.89861E-18 | 0.26971 | 0.391 | 0.212 | 2.51433E-14 |
| Contractile | Ndrg1 | 0.000252326 | 0.269453 | 0.335 | 0.256 | 1 |
| Intermediate | Pisd | 1.31894E-20 | 0.269358 | 0.205 | 0.075 | 1.74667E-16 |
| Intermediate | Commd4 | 8.89581E-16 | 0.269343 | 0.426 | 0.261 | 1.17807E-11 |
| Terminal | Nebl | 2.96566E-28 | 0.269333 | 0.158 | 0.033 | 3.92742E-24 |
| Contractile | Slirp | 0.000549018 | 0.26915 | 0.576 | 0.562 | 1 |
| Intermediate | Tpst2 | 8.99708E-14 | 0.268929 | 0.334 | 0.198 | 1.19148E-09 |
| Contractile | Mrgprf | 0.005826544 | 0.268845 | 0.188 | 0.141 | 1 |
| Contractile | Nhsl2 | 1.66381E-22 | 0.268844 | 0.1 | 0.012 | 2.20338E-18 |
| Intermediate | Pdia4 | 1.00211E-07 | 0.268793 | 0.516 | 0.406 | 0.001327101 |
| Intermediate | Nipsnap3b | 2.98795E-19 | 0.268679 | 0.282 | 0.132 | 3.95694E-15 |
| Intermediate | Lrp10 | 4.0752E-12 | 0.268645 | 0.453 | 0.308 | 5.39678E-08 |
| Contractile | Vgll3 | 0.000314748 | 0.268274 | 0.144 | 0.086 | 1 |
| Contractile | Grhpr | 0.00073015 | 0.268101 | 0.232 | 0.172 | 1 |
| Intermediate | Nrip1 | 2.29612E-13 | 0.268088 | 0.32 | 0.188 | 3.04075E-09 |
| Contractile | Tmem141 | 7.46768E-05 | 0.268059 | 0.176 | 0.11 | 0.988945133 |
| Terminal | Cyp26b1 | 4.33224E-27 | 0.268054 | 0.147 | 0.029 | 5.73718E-23 |
| Rgs5_Contractile | Dek | 0.000401033 | 0.268052 | 0.292 | 0.519 | 1 |
| Contractile | Fam126b | 7.75578E-08 | 0.267893 | 0.174 | 0.084 | 0.001027097 |

| | | | | | | |
|---|---|---|---|---|---|---|
| Contractile | Casp12 | 0.000116669 | 0.267822 | 0.326 | 0.25 | 1 |
| Terminal | Clmp | 2.82751E-15 | 0.267737 | 0.385 | 0.227 | 3.74447E-11 |
| Contractile | Sft2d2 | 0.000248198 | 0.267704 | 0.185 | 0.122 | 1 |
| Intermediate | Elovl1 | 1.14031E-16 | 0.267458 | 0.475 | 0.296 | 1.51011E-12 |
| Intermediate | Twf1 | 1.98304E-13 | 0.267257 | 0.332 | 0.196 | 2.62615E-09 |
| Intermediate | Pros1 | 2.00404E-11 | 0.267248 | 0.473 | 0.333 | 2.65394E-07 |
| Rgs5_Contractile | Jund | 0.000550615 | 0.267175 | 0.74 | 0.81 | 1 |
| Contractile | Psmd7 | 0.000198246 | 0.26673 | 0.638 | 0.633 | 1 |
| Terminal | Os9 | 2.92869E-19 | 0.266721 | 0.609 | 0.387 | 3.87847E-15 |
| Rgs5_Contractile | Chtop | 0.002056812 | 0.266443 | 0.17 | 0.304 | 1 |
| Terminal | Crtap | 7.8961E-14 | 0.266223 | 0.616 | 0.425 | 1.04568E-09 |
| Intermediate | Mdh2 | 3.16256E-14 | 0.266179 | 0.732 | 0.585 | 4.18817E-10 |
| Contractile | Taldo1 | 0.008858064 | 0.266132 | 0.576 | 0.585 | 1 |
| Terminal | Kcnq1ot1 | 3.82195E-20 | 0.266121 | 0.39 | 0.206 | 5.06141E-16 |
| Contractile | Gsk3b | 0.003737095 | 0.266093 | 0.403 | 0.367 | 1 |
| Contractile | Osbpl1a | 0.001978622 | 0.265915 | 0.194 | 0.14 | 1 |
| Contractile | Ndufa8 | 0.004651738 | 0.265813 | 0.597 | 0.613 | 1 |
| Contractile | Rap1a | 0.00042525 | 0.265721 | 0.624 | 0.617 | 1 |
| Contractile | Rhoc | 2.21351E-07 | 0.265599 | 0.829 | 0.786 | 0.002931352 |
| Intermediate | Dnajc1 | 2.11138E-10 | 0.265566 | 0.499 | 0.36 | 2.79609E-06 |
| Intermediate | Kdelr3 | 1.43923E-08 | 0.265338 | 0.568 | 0.462 | 0.000190598 |
| Contractile | Actr3 | 5.14175E-05 | 0.265335 | 0.591 | 0.559 | 0.680921353 |
| Contractile | Tspan5 | 4.16593E-05 | 0.265263 | 0.197 | 0.122 | 0.551694547 |
| Intermediate | Ezr | 9.78392E-23 | 0.265135 | 0.133 | 0.029 | 1.29569E-18 |
| Rgs5_Contractile | Prkacb | 0.009586259 | 0.26501 | 0.194 | 0.32 | 1 |
| Contractile | Chchd4 | 7.3477E-05 | 0.264799 | 0.174 | 0.107 | 0.973055974 |
| Terminal | Uxs1 | 1.03762E-19 | 0.264713 | 0.413 | 0.226 | 1.37412E-15 |
| Contractile | Pde4b | 0.002206444 | 0.264489 | 0.259 | 0.201 | 1 |
| Contractile | Mrpl18 | 0.001922925 | 0.264423 | 0.474 | 0.438 | 1 |
| Contractile | Kat2b | 0.000908021 | 0.264397 | 0.171 | 0.115 | 1 |
| Contractile | Mrpl51 | 0.001140936 | 0.264306 | 0.468 | 0.438 | 1 |
| Intermediate | Plbd2 | 2.61517E-09 | 0.264272 | 0.497 | 0.38 | 3.46326E-05 |
| Contractile | Wac | 0.006079545 | 0.264067 | 0.294 | 0.255 | 1 |
| Contractile | Anp32a | 0.006534026 | 0.263982 | 0.444 | 0.427 | 1 |
| Intermediate | Gpx8 | 2.70553E-13 | 0.263936 | 0.775 | 0.651 | 3.58294E-09 |
| Intermediate | Susd6 | 2.19532E-11 | 0.263633 | 0.265 | 0.152 | 2.90726E-07 |
| Contractile | Grip2 | 5.11704E-21 | 0.263592 | 0.103 | 0.014 | 6.77649E-17 |
| Contractile | Ccdc50 | 0.009870287 | 0.263454 | 0.244 | 0.202 | 1 |
| Intermediate | Hmox1 | 1.97206E-10 | 0.263441 | 0.173 | 0.088 | 2.6116E-06 |
| Contractile | Sept4 | 4.94295E-08 | 0.263358 | 0.124 | 0.049 | 0.000654595 |
| Contractile | Anp32b | 0.002452719 | 0.263329 | 0.635 | 0.633 | 1 |
| Contractile | Nacad | 2.08421E-12 | 0.263189 | 0.103 | 0.026 | 2.76012E-08 |
| Contractile | 2610001J0 | 0.004581323 | 0.263172 | 0.521 | 0.519 | 1 |
| Rgs5_Contractile | Oard1 | 0.001890075 | 0.263115 | 0.118 | 0.222 | 1 |
| Rgs5_Contractile | Rragc | 0.008918183 | 0.263058 | 0.149 | 0.251 | 1 |
| Intermediate | Ephx2 | 3.19579E-14 | 0.263012 | 0.12 | 0.039 | 4.23219E-10 |
| Intermediate | B3galnt1 | 2.81051E-20 | 0.262955 | 0.214 | 0.081 | 3.72196E-16 |
| Intermediate | Ube2l3 | 2.25822E-15 | 0.262808 | 0.512 | 0.334 | 2.99055E-11 |
| Intermediate | Egfr | 5.68432E-12 | 0.262655 | 0.194 | 0.096 | 7.52774E-08 |
| Rgs5_Contractile | Tnrc6c | 0.005840233 | 0.262629 | 0.222 | 0.371 | 1 |
| Terminal | Hmgn2 | 1.45118E-18 | 0.262418 | 0.452 | 0.263 | 1.9218E-14 |
| Terminal | Sdc3 | 6.78673E-21 | 0.262412 | 0.564 | 0.332 | 8.98766E-17 |
| Intermediate | Gbp2 | 3.65673E-13 | 0.262346 | 0.158 | 0.065 | 4.84261E-09 |
| Rgs5_Contractile | Eif4h | 0.000815677 | 0.262278 | 0.295 | 0.527 | 1 |
| Intermediate | H6pd | 1.29062E-17 | 0.262239 | 0.219 | 0.092 | 1.70917E-13 |
| Rgs5_Contractile | Zranb2 | 0.00332944 | 0.262211 | 0.188 | 0.321 | 1 |
| Intermediate | Tubb4b | 3.02334E-10 | 0.262134 | 0.554 | 0.418 | 4.00381E-06 |
| Rgs5_Contractile | Prpsap1 | 0.00899454 | 0.262026 | 0.17 | 0.282 | 1 |
| Contractile | Crbn | 0.008975969 | 0.261937 | 0.156 | 0.114 | 1 |

| | | | | | | |
|---|---|---|---|---|---|---|
| Intermediate | Fscn1 | 5.02721E-27 | 0.261735 | 0.147 | 0.028 | 6.65754E-23 |
| Contractile | Scarb2 | 0.000737816 | 0.261669 | 0.429 | 0.374 | 1 |
| Intermediate | Nek6 | 4.65055E-18 | 0.26156 | 0.196 | 0.077 | 6.15872E-14 |
| Intermediate | Cfap20 | 3.66671E-12 | 0.261482 | 0.385 | 0.25 | 4.85582E-08 |
| Contractile | Fam26e | 0.008207719 | 0.261466 | 0.218 | 0.171 | 1 |
| Intermediate | Leprotl1 | 1.19411E-14 | 0.261322 | 0.408 | 0.258 | 1.58135E-10 |
| Contractile | Tns2 | 0.007211531 | 0.261255 | 0.309 | 0.264 | 1 |
| Rgs5_Contractile | Nfat5 | 0.002612605 | 0.261248 | 0.184 | 0.32 | 1 |
| Terminal | Slc20a1 | 5.69173E-19 | 0.2612 | 0.171 | 0.058 | 7.53755E-15 |
| Contractile | Ywhab | 0.000511025 | 0.26109 | 0.568 | 0.541 | 1 |
| Terminal | C1qtnf7 | 4.50185E-14 | 0.261036 | 0.239 | 0.118 | 5.96181E-10 |
| Intermediate | Lipa | 4.03785E-28 | 0.260933 | 0.195 | 0.051 | 5.34733E-24 |
| Contractile | Actg1 | 8.52945E-13 | 0.260924 | 0.988 | 0.964 | 1.12956E-08 |
| Intermediate | Pofut2 | 3.68807E-10 | 0.260812 | 0.628 | 0.504 | 4.88411E-06 |
| Contractile | Rnf146 | 0.00444599 | 0.260356 | 0.232 | 0.184 | 1 |
| Rgs5_Contractile | Sfxn3 | 0.001830867 | 0.26033 | 0.181 | 0.322 | 1 |
| Intermediate | Snrpd1 | 2.37595E-12 | 0.260314 | 0.506 | 0.356 | 3.14648E-08 |
| Rgs5_Contractile | Flii | 0.002628207 | 0.260257 | 0.132 | 0.238 | 1 |
| Contractile | Gem | 0.001182789 | 0.260252 | 0.406 | 0.347 | 1 |
| Intermediate | Sf3b6 | 4.52823E-11 | 0.260204 | 0.601 | 0.464 | 5.99673E-07 |
| Terminal | Slc43a3 | 5.41115E-20 | 0.260057 | 0.369 | 0.192 | 7.16599E-16 |
| Contractile | Rrp1 | 0.00730923 | 0.260047 | 0.421 | 0.401 | 1 |
| Intermediate | Itm2c | 2.99875E-12 | 0.260028 | 0.839 | 0.74 | 3.97124E-08 |
| Intermediate | Far1 | 2.17631E-11 | 0.260005 | 0.277 | 0.162 | 2.88209E-07 |
| Contractile | Zmat5 | 0.004442866 | 0.259936 | 0.344 | 0.303 | 1 |
| Contractile | Tceb2 | 1.71191E-09 | 0.25992 | 0.938 | 0.93 | 2.26708E-05 |
| Intermediate | Grina | 2.27246E-10 | 0.259889 | 0.479 | 0.342 | 3.00942E-06 |
| Intermediate | Csf1 | 3.20688E-15 | 0.259865 | 0.275 | 0.139 | 4.24688E-11 |
| Intermediate | Lmo4 | 3.27702E-11 | 0.259781 | 0.765 | 0.623 | 4.33975E-07 |
| Terminal | Hfe | 7.80052E-22 | 0.259761 | 0.218 | 0.08 | 1.03302E-17 |
| Contractile | Nampt | 0.000509523 | 0.259737 | 0.206 | 0.142 | 1 |
| Terminal | Ugdh | 6.04421E-16 | 0.259701 | 0.461 | 0.28 | 8.00435E-12 |
| Intermediate | Ppfibp1 | 1.58636E-12 | 0.259564 | 0.406 | 0.261 | 2.10082E-08 |
| Terminal | BC026585 | 3.96209E-21 | 0.259482 | 0.273 | 0.12 | 5.24699E-17 |
| Contractile | Rab14 | 0.001635063 | 0.259476 | 0.588 | 0.604 | 1 |
| Intermediate | Iqgap1 | 2.04575E-13 | 0.259433 | 0.624 | 0.469 | 2.70919E-09 |
| Intermediate | Idh1 | 9.05164E-13 | 0.259371 | 0.235 | 0.122 | 1.19871E-08 |
| Intermediate | Dpm2 | 1.44631E-15 | 0.259297 | 0.413 | 0.245 | 1.91535E-11 |
| Rgs5_Contractile | Nap1l4 | 0.004950442 | 0.259196 | 0.188 | 0.321 | 1 |
| Contractile | Vstm2b | 4.64751E-10 | 0.259144 | 0.1 | 0.029 | 6.1547E-06 |
| Contractile | Hoxb6 | 3.73405E-09 | 0.259017 | 0.109 | 0.037 | 4.945E-05 |
| Contractile | Trappc6b | 0.003342442 | 0.258988 | 0.465 | 0.438 | 1 |
| Rgs5_Contractile | Cdk2ap2 | 0.007442746 | 0.25894 | 0.323 | 0.536 | 1 |
| Terminal | Acsf2 | 3.9384E-27 | 0.258928 | 0.155 | 0.034 | 5.21562E-23 |
| Intermediate | Inf2 | 1.63876E-19 | 0.258808 | 0.195 | 0.07 | 2.1702E-15 |
| Contractile | Bckdha | 0.002911006 | 0.258742 | 0.259 | 0.205 | 1 |
| Terminal | Fkbp7 | 5.25924E-12 | 0.258608 | 0.804 | 0.621 | 6.96481E-08 |
| Intermediate | Timm13 | 7.12206E-13 | 0.25856 | 0.763 | 0.641 | 9.43174E-09 |
| Contractile | Wdr83os | 0.003538156 | 0.258557 | 0.482 | 0.472 | 1 |
| Terminal | Creg1 | 1.58873E-20 | 0.258485 | 0.547 | 0.323 | 2.10396E-16 |
| Intermediate | Tmed9 | 4.52385E-12 | 0.258443 | 0.874 | 0.781 | 5.99094E-08 |
| Intermediate | Cnih4 | 7.64764E-11 | 0.258322 | 0.488 | 0.354 | 1.01278E-06 |
| Contractile | Psma5 | 0.006057153 | 0.258314 | 0.524 | 0.522 | 1 |
| Contractile | Cbx7 | 4.97185E-09 | 0.25821 | 0.121 | 0.044 | 6.58423E-05 |
| Contractile | Gabbr1 | 8.20507E-05 | 0.258174 | 0.147 | 0.085 | 1 |
| Rgs5_Contractile | Dnajc5 | 0.008120982 | 0.258039 | 0.135 | 0.233 | 1 |
| Contractile | Lonp2 | 0.008229976 | 0.258011 | 0.188 | 0.143 | 1 |
| Contractile | Frg1 | 0.006746263 | 0.257917 | 0.3 | 0.256 | 1 |
| Intermediate | Gars | 2.21209E-14 | 0.257418 | 0.321 | 0.182 | 2.92947E-10 |

| | | | | | | |
|---|---|---|---|---|---|---|
| Rgs5_Contractile | St6galnac6 | 0.009186395 | 0.257416 | 0.163 | 0.272 | 1 |
| Intermediate | Ralbp1 | 9.87968E-10 | 0.257401 | 0.643 | 0.512 | 1.30837E-05 |
| Contractile | Gpx4 | 6.93261E-09 | 0.257394 | 0.932 | 0.882 | 9.18085E-05 |
| Contractile | Txnip | 0.000738744 | 0.25725 | 0.371 | 0.306 | 1 |
| Contractile | Specc1l | 0.001668084 | 0.257196 | 0.153 | 0.101 | 1 |
| Contractile | Tmx4 | 0.002551159 | 0.25714 | 0.226 | 0.173 | 1 |
| Contractile | Map6 | 5.88467E-05 | 0.257106 | 0.171 | 0.102 | 0.779306367 |
| Intermediate | Rhoq | 4.36577E-13 | 0.257032 | 0.339 | 0.201 | 5.78158E-09 |
| Intermediate | Sntb2 | 6.28249E-12 | 0.256821 | 0.329 | 0.204 | 8.31991E-08 |
| Contractile | Sult1a1 | 2.15803E-15 | 0.256747 | 0.118 | 0.027 | 2.85788E-11 |
| Rgs5_Contractile | Zbtb7a | 0.003847962 | 0.256475 | 0.194 | 0.333 | 1 |
| Contractile | Stra13 | 0.001955124 | 0.256402 | 0.406 | 0.367 | 1 |
| Contractile | Prps2 | 4.29744E-05 | 0.256352 | 0.171 | 0.102 | 0.569109397 |
| Intermediate | Apoe | 2.12889E-13 | 0.256312 | 0.159 | 0.066 | 2.81929E-09 |
| Intermediate | Nt5c | 9.2346E-16 | 0.25626 | 0.434 | 0.265 | 1.22294E-11 |
| Intermediate | Rab8a | 2.53034E-16 | 0.256237 | 0.331 | 0.182 | 3.35092E-12 |
| Intermediate | Mvp | 1.12024E-10 | 0.256187 | 0.374 | 0.25 | 1.48353E-06 |
| Intermediate | P4ha1 | 4.69437E-09 | 0.256131 | 0.473 | 0.352 | 6.21675E-05 |
| Contractile | Banf1 | 0.000644889 | 0.256092 | 0.535 | 0.503 | 1 |
| Intermediate | Gpatch11 | 1.76006E-09 | 0.256022 | 0.419 | 0.303 | 2.33085E-05 |
| Intermediate | Plpp2 | 7.87301E-15 | 0.256018 | 0.228 | 0.111 | 1.04262E-10 |
| Intermediate | Fam102b | 3.78961E-15 | 0.256006 | 0.252 | 0.125 | 5.01859E-11 |
| Terminal | Adk | 1.55765E-14 | 0.255975 | 0.622 | 0.436 | 2.06279E-10 |
| Intermediate | Nans | 3.81497E-09 | 0.255959 | 0.321 | 0.219 | 5.05217E-05 |
| Contractile | Fbxo3 | 0.000375699 | 0.255781 | 0.256 | 0.19 | 1 |
| Intermediate | Dpp7 | 7.08295E-18 | 0.255618 | 0.267 | 0.124 | 9.37996E-14 |
| Contractile | Fam168a | 0.003563222 | 0.25554 | 0.194 | 0.146 | 1 |
| Contractile | Smim1 | 0.000275068 | 0.255486 | 0.156 | 0.097 | 1 |
| Contractile | C2cd2 | 7.64891E-09 | 0.255477 | 0.115 | 0.042 | 0.000101295 |
| Intermediate | Adam10 | 1.64384E-18 | 0.25539 | 0.217 | 0.087 | 2.17694E-14 |
| Contractile | Nt5c3b | 0.006779726 | 0.255267 | 0.235 | 0.19 | 1 |
| Intermediate | Tmem14c | 2.31316E-12 | 0.255238 | 0.62 | 0.479 | 3.06332E-08 |
| Intermediate | Rbm3 | 2.03123E-10 | 0.254985 | 0.784 | 0.68 | 2.68996E-06 |
| Contractile | Camsap1 | 8.22317E-05 | 0.254923 | 0.126 | 0.069 | 1 |
| Intermediate | Chmp1a | 4.46255E-14 | 0.254744 | 0.384 | 0.231 | 5.90976E-10 |
| Contractile | Kidins220 | 0.000347668 | 0.254676 | 0.209 | 0.144 | 1 |
| Rgs5_Contractile | Gns | 0.002504041 | 0.254605 | 0.16 | 0.282 | 1 |
| Terminal | Jun | 1.10046E-07 | 0.254479 | 0.716 | 0.599 | 0.001457342 |
| Intermediate | Creb3 | 4.12126E-12 | 0.25438 | 0.475 | 0.326 | 5.45779E-08 |
| Contractile | Golim4 | 0.00560486 | 0.254375 | 0.471 | 0.46 | 1 |
| Intermediate | Antxr1 | 1.83506E-11 | 0.254357 | 0.46 | 0.316 | 2.43017E-07 |
| Terminal | Rrbp1 | 1.14384E-13 | 0.254182 | 0.946 | 0.8 | 1.51479E-09 |
| Intermediate | Psmd2 | 5.50341E-11 | 0.254024 | 0.489 | 0.356 | 7.28816E-07 |
| Contractile | Ctnnd1 | 0.001075214 | 0.254022 | 0.218 | 0.159 | 1 |
| Rgs5_Contractile | Banf1 | 0.005673238 | 0.253903 | 0.316 | 0.533 | 1 |
| Contractile | Tmem87b | 0.001775189 | 0.253896 | 0.221 | 0.166 | 1 |
| Terminal | Erf | 2.50095E-17 | 0.253657 | 0.296 | 0.149 | 3.312E-13 |
| Contractile | Hdac9 | 2.64125E-10 | 0.253483 | 0.103 | 0.03 | 3.49781E-06 |
| Contractile | Sdha | 0.00193023 | 0.253459 | 0.459 | 0.435 | 1 |
| Rgs5_Contractile | Aimp1 | 0.00252778 | 0.253393 | 0.222 | 0.387 | 1 |
| Rgs5_Contractile | Uqcrh | 0.004052725 | 0.253155 | 0.757 | 0.902 | 1 |
| Contractile | Cfdp1 | 0.003029599 | 0.253127 | 0.512 | 0.499 | 1 |
| Intermediate | Psmb2 | 1.31087E-12 | 0.25309 | 0.767 | 0.65 | 1.73599E-08 |
| Intermediate | Pkd1 | 3.31173E-11 | 0.253027 | 0.539 | 0.395 | 4.38572E-07 |
| Terminal | Il17d | 1.13793E-26 | 0.253021 | 0.145 | 0.028 | 1.50696E-22 |
| Contractile | Mecp2 | 0.006067972 | 0.253019 | 0.209 | 0.162 | 1 |
| Contractile | Inhba | 9.10906E-07 | 0.252994 | 0.359 | 0.25 | 0.012063133 |
| Intermediate | Spcs2 | 9.83715E-08 | 0.252955 | 0.748 | 0.642 | 0.001302734 |
| Contractile | Fdps | 0.000124329 | 0.25285 | 0.182 | 0.115 | 1 |

| | | | | | | |
|---|---|---|---|---|---|---|
| Rgs5_Contractile | Nr3c1 | 0.007569996 | 0.252843 | 0.24 | 0.398 | 1 |
| Terminal | Zc3h12a | 5.78642E-19 | 0.252805 | 0.204 | 0.08 | 7.66296E-15 |
| Intermediate | Bad | 1.07488E-11 | 0.252637 | 0.525 | 0.389 | 1.42347E-07 |
| Contractile | Txn1 | 4.64192E-07 | 0.252621 | 0.85 | 0.824 | 0.006147291 |
| Intermediate | Lsp1 | 3.0479E-10 | 0.252595 | 0.264 | 0.158 | 4.03633E-06 |
| Intermediate | Sec61a1 | 3.74523E-10 | 0.252512 | 0.507 | 0.375 | 4.95981E-06 |
| Intermediate | Pdcd6 | 4.92947E-09 | 0.252506 | 0.636 | 0.533 | 6.52809E-05 |
| Intermediate | Galk2 | 1.03822E-17 | 0.252491 | 0.204 | 0.083 | 1.37491E-13 |
| Intermediate | Svbp | 2.53857E-11 | 0.252433 | 0.564 | 0.421 | 3.36182E-07 |
| Intermediate | Psmd6 | 2.66573E-12 | 0.252395 | 0.494 | 0.343 | 3.53022E-08 |
| Contractile | Wbp5 | 4.62533E-05 | 0.252174 | 0.762 | 0.743 | 0.612531927 |
| Contractile | Loxl1 | 1.04113E-07 | 0.252169 | 0.55 | 0.406 | 0.001378764 |
| Intermediate | Maea | 1.80089E-14 | 0.252088 | 0.337 | 0.194 | 2.38492E-10 |
| Contractile | Nxt2 | 7.46494E-08 | 0.251894 | 0.147 | 0.066 | 0.000988582 |
| Contractile | Psmd8 | 0.00032134 | 0.251834 | 0.6 | 0.558 | 1 |
| Rgs5_Contractile | Cggbp1 | 0.005938301 | 0.251832 | 0.191 | 0.318 | 1 |
| Intermediate | Esd | 3.02805E-09 | 0.251715 | 0.701 | 0.586 | 4.01005E-05 |
| Contractile | Flot1 | 0.005544067 | 0.251628 | 0.409 | 0.38 | 1 |
| Contractile | Arl6ip1 | 0.001644378 | 0.251618 | 0.374 | 0.318 | 1 |
| Rgs5_Contractile | Eif4b | 0.008348495 | 0.251463 | 0.222 | 0.364 | 1 |
| Intermediate | Parva | 7.72055E-15 | 0.251335 | 0.393 | 0.231 | 1.02243E-10 |
| Intermediate | Snx18 | 1.88057E-12 | 0.251269 | 0.372 | 0.229 | 2.49044E-08 |
| Intermediate | Arpc1b | 7.47233E-17 | 0.251219 | 0.943 | 0.893 | 9.89561E-13 |
| Rgs5_Contractile | Ppp2ca | 0.006176235 | 0.251215 | 0.316 | 0.533 | 1 |
| Contractile | Cyb5r4 | 0.001464556 | 0.251154 | 0.188 | 0.135 | 1 |
| Contractile | Gipc2 | 1.69101E-08 | 0.25113 | 0.135 | 0.055 | 0.000223941 |
| Intermediate | Nhp2 | 5.44269E-11 | 0.250831 | 0.483 | 0.352 | 7.20776E-07 |
| Contractile | Ift27 | 0.004270306 | 0.250805 | 0.282 | 0.239 | 1 |
| Rgs5_Contractile | Mier1 | 0.002514562 | 0.250755 | 0.167 | 0.295 | 1 |
| Contractile | Aplp1 | 3.82511E-06 | 0.250748 | 0.144 | 0.072 | 0.050655962 |
| Contractile | Wdr45 | 0.006908925 | 0.250725 | 0.191 | 0.148 | 1 |
| Rgs5_Contractile | Matr3 | 0.006530175 | 0.250515 | 0.24 | 0.397 | 1 |
| Intermediate | H2-K1 | 0.001256075 | 0.250406 | 0.517 | 0.453 | 1 |
| Contractile | Hivep1 | 0.001413074 | 0.250293 | 0.156 | 0.104 | 1 |
| Intermediate | Dtnbp1 | 1.40451E-13 | 0.250191 | 0.419 | 0.271 | 1.85999E-09 |
| Contractile | Thumpd1 | 0.008718302 | 0.250104 | 0.138 | 0.098 | 1 |
| Intermediate | Cct3 | 1.19801E-12 | 0.250023 | 0.496 | 0.341 | 1.58652E-08 |
| Contractile | Ndufs4 | 0.009262382 | 0.250001 | 0.435 | 0.414 | 1 |

**Table S5. Marker genes for mesenchymal cell clusters in individual human coronary plaque scRNA-seq data. Corresponds to Fig. 1D.**
The listed genes have significantly higher expression in the indicated cluster than all other clusters combined.
Analysis was performed using the Wilcoxon test in Seurat v4.4.0 with P values adjusted for multiple testing by the Bonferroni method.

| cluster | gene | p_val | avg_log2F( | pct.1 | pct.2 | p_val_adj |
|---|---|---|---|---|---|---|
| Fibroblasts | APOD | 0 | 6.239827 | 0.956 | 0.24 | 0 |
| Fibroblasts | PLA2G2A | 6.8318E-290 | 6.109761 | 0.744 | 0.19 | 1.1221E-285 |
| Fibroblasts | CFD | 0 | 5.492189 | 0.881 | 0.366 | 0 |
| Fibroblasts | FBLN1 | 0 | 4.902937 | 0.998 | 0.16 | 0 |
| Fibroblasts | PTGDS | 0 | 4.497168 | 0.812 | 0.108 | 0 |
| Pericytes | CCL21 | 0 | 4.42203 | 0.656 | 0.045 | 0 |
| Fibroblasts | CXCL14 | 5.9949E-307 | 4.117714 | 0.55 | 0.029 | 9.846E-303 |
| Fibroblasts | C3 | 0 | 4.104663 | 0.933 | 0.055 | 0 |
| Fibroblasts | SERPINF1 | 0 | 3.982974 | 0.997 | 0.211 | 0 |
| RERGL_Cells | RERGL | 0 | 3.888293 | 0.889 | 0.091 | 0 |
| Fibroblasts | C7 | 0 | 3.867541 | 0.962 | 0.242 | 0 |
| Fibroblasts | DCN | 0 | 3.79242 | 1 | 0.48 | 0 |
| Fibroblasts | RARRES1 | 0 | 3.745849 | 0.773 | 0.08 | 0 |
| Pericytes | FABP4 | 6.1499E-279 | 3.722347 | 0.904 | 0.204 | 1.0101E-274 |
| Pericytes | CD36 | 0 | 3.614737 | 0.691 | 0.026 | 0 |
| Fibroblasts | MGST1 | 0 | 3.577756 | 0.91 | 0.036 | 0 |
| Fibroblasts | SFRP2 | 0 | 3.57037 | 0.615 | 0.018 | 0 |
| Pericytes | STEAP4 | 0 | 3.455872 | 0.922 | 0.19 | 0 |
| Fibroblasts | MYOC | 0 | 3.440352 | 0.644 | 0.025 | 0 |
| Fibroblasts | MFAP5 | 0 | 3.438744 | 0.721 | 0.032 | 0 |
| Fibroblasts | ITM2A | 0 | 3.233948 | 0.923 | 0.11 | 0 |
| RERGL_Cells | MUSTN1 | 4.2276E-292 | 3.23291 | 0.955 | 0.249 | 6.9434E-288 |
| Fibroblasts | HP | 1.8497E-167 | 3.22873 | 0.312 | 0.011 | 3.0379E-163 |
| Fibroblasts | FBLN2 | 0 | 3.174702 | 0.959 | 0.133 | 0 |
| Pericytes | RGS16 | 5.7686E-237 | 3.038748 | 0.758 | 0.154 | 9.4744E-233 |
| Fibroblasts | SERPINA3 | 0 | 2.975566 | 0.924 | 0.13 | 0 |
| Pericytes | FABP5 | 9.1774E-258 | 2.902851 | 0.705 | 0.102 | 1.5073E-253 |
| Intermediate | IGFBP2 | 1.1205E-285 | 2.781362 | 0.97 | 0.296 | 1.8403E-281 |
| Fibroblasts | GSN | 0 | 2.769489 | 1 | 0.957 | 0 |
| Fibroblasts | PI16 | 0 | 2.738845 | 0.6 | 0.027 | 0 |
| Pericytes | ID4 | 1.0491E-170 | 2.736637 | 0.893 | 0.508 | 1.723E-166 |
| RERGL_Cells | NET1 | 3.7153E-208 | 2.720686 | 0.784 | 0.205 | 6.102E-204 |
| Pericytes | TCIM | 1.1855E-238 | 2.70639 | 0.704 | 0.125 | 1.9471E-234 |
| Fibroblasts | PLIN2 | 7.0142E-157 | 2.669764 | 0.529 | 0.131 | 1.152E-152 |
| Terminal | COMP | 4.99538E-28 | 2.660841 | 0.146 | 0.033 | 8.20441E-24 |
| Intermediate | TNFRSF11I | 5.7642E-212 | 2.63085 | 0.654 | 0.106 | 9.4671E-208 |
| Fibroblasts | HTRA3 | 0 | 2.602822 | 0.652 | 0.031 | 0 |
| Pericytes | TM4SF1 | 1.4384E-153 | 2.592514 | 0.827 | 0.417 | 2.3624E-149 |
| Pericytes | CCL19 | 5.9073E-129 | 2.588044 | 0.464 | 0.084 | 9.7022E-125 |
| Intermediate | FN1 | 4.9528E-195 | 2.515592 | 0.964 | 0.539 | 8.1344E-191 |
| Fibroblasts | IGF1 | 0 | 2.485716 | 0.594 | 0.024 | 0 |
| Fibroblasts | LUM | 0 | 2.469418 | 0.988 | 0.281 | 0 |
| Intermediate | GAS6 | 2.1946E-157 | 2.464776 | 0.935 | 0.641 | 3.6044E-153 |
| Pericytes | ADAMTS1 | 2.4083E-164 | 2.462595 | 0.82 | 0.341 | 3.9553E-160 |
| Pericytes | ADAMTS4 | 1.4264E-177 | 2.461403 | 0.573 | 0.1 | 2.3428E-173 |
| Contractile | ACTG2 | 1.5636E-211 | 2.459003 | 0.717 | 0.128 | 2.5681E-207 |
| RERGL_Cells | PHLDA2 | 4.082E-227 | 2.436733 | 0.645 | 0.096 | 6.7042E-223 |
| Contractile | ACTC1 | 2.0785E-207 | 2.432782 | 0.427 | 0.025 | 3.4137E-203 |
| Fibroblasts | C1R | 0 | 2.40647 | 0.998 | 0.687 | 0 |
| Fibroblasts | CCDC80 | 1.0044E-303 | 2.396655 | 0.927 | 0.391 | 1.6497E-299 |
| Fibroblasts | PTN | 3.7134E-250 | 2.394432 | 0.694 | 0.149 | 6.099E-246 |

| | | | | | | |
|---|---|---|---|---|---|---|
| Fibroblasts | SRPX | 1.0739E-283 | 2.362803 | 0.773 | 0.196 | 1.7638E-279 |
| Pericytes | CCL8 | 4.8227E-155 | 2.325254 | 0.318 | 0.018 | 7.9207E-151 |
| Fibroblasts | C1S | 0 | 2.307748 | 0.999 | 0.61 | 0 |
| RERGL_Cells | BCAM | 4.5077E-225 | 2.305716 | 0.944 | 0.335 | 7.4035E-221 |
| Intermediate | VCAN | 2.9075E-215 | 2.288504 | 0.918 | 0.339 | 4.7753E-211 |
| Fibroblasts | MMP2 | 0 | 2.265734 | 0.897 | 0.233 | 0 |
| Fibroblasts | GPX3 | 1.2795E-209 | 2.235939 | 0.946 | 0.829 | 2.1015E-205 |
| Fibroblasts | DPT | 8.9157E-297 | 2.220396 | 0.806 | 0.173 | 1.4643E-292 |
| Fibroblasts | TAC1 | 1.09268E-94 | 2.206851 | 0.175 | 0.004 | 1.79463E-90 |
| Fibroblasts | PLTP | 1.2072E-299 | 2.18458 | 0.826 | 0.233 | 1.9827E-295 |
| RERGL_Cells | PTP4A3 | 1.9812E-194 | 2.182222 | 0.756 | 0.181 | 3.254E-190 |
| Pericytes | ATF3 | 6.2933E-185 | 2.17967 | 0.775 | 0.232 | 1.0336E-180 |
| Intermediate | CCN3 | 3.1899E-196 | 2.173054 | 0.905 | 0.294 | 5.2391E-192 |
| Intermediate | COL8A1 | 1.9669E-244 | 2.134417 | 0.845 | 0.2 | 3.2304E-240 |
| Pericytes | FGF7 | 1.043E-150 | 2.127136 | 0.878 | 0.387 | 1.713E-146 |
| Contractile | MYH11 | 4.801E-217 | 2.118697 | 0.986 | 0.411 | 7.8851E-213 |
| Pericytes | CITED2 | 9.38463E-65 | 2.104959 | 0.533 | 0.248 | 1.54133E-60 |
| Contractile | CNN1 | 2.1212E-184 | 2.101341 | 0.948 | 0.397 | 3.4838E-180 |
| Pericytes | ARID5A | 1.3262E-176 | 2.097842 | 0.825 | 0.341 | 2.1782E-172 |
| Contractile | PLN | 1.1273E-189 | 2.09385 | 0.986 | 0.482 | 1.8514E-185 |
| Contractile | TPM2 | 1.1241E-194 | 2.039194 | 0.998 | 0.832 | 1.8462E-190 |
| Pericytes | MYC | 5.8031E-136 | 2.036868 | 0.831 | 0.388 | 9.531E-132 |
| Intermediate | PRSS23 | 3.0003E-127 | 2.022729 | 0.746 | 0.287 | 4.9278E-123 |
| Fibroblasts | EMP1 | 0 | 2.004956 | 0.839 | 0.162 | 0 |
| Contractile | RAMP1 | 2.1847E-213 | 1.980926 | 0.94 | 0.293 | 3.5881E-209 |
| Pericytes | ARID5B | 5.524E-194 | 1.952508 | 0.962 | 0.659 | 9.0726E-190 |
| Pericytes | LGI4 | 1.0489E-193 | 1.937991 | 0.689 | 0.155 | 1.7227E-189 |
| Fibroblasts | GPC3 | 0 | 1.937946 | 0.648 | 0.063 | 0 |
| Pericytes | TGFBI | 1.789E-150 | 1.921622 | 0.669 | 0.192 | 2.9382E-146 |
| Fibroblasts | PODN | 0 | 1.914625 | 0.778 | 0.133 | 0 |
| Terminal | CRTAC1 | 7.88608E-75 | 1.914262 | 0.209 | 0.021 | 1.29521E-70 |
| Intermediate | SLC14A1 | 1.0308E-292 | 1.909481 | 0.656 | 0.055 | 1.693E-288 |
| Fibroblasts | FBLN5 | 0 | 1.909248 | 0.945 | 0.383 | 0 |
| Pericytes | LINC00924 | 4.3589E-217 | 1.8854 | 0.515 | 0.051 | 7.159E-213 |
| Fibroblasts | UAP1 | 2.1302E-207 | 1.884775 | 0.669 | 0.185 | 3.4986E-203 |
| Fibroblasts | PTX3 | 5.47866E-96 | 1.880024 | 0.207 | 0.013 | 8.99816E-92 |
| RERGL_Cells | LBH | 1.835E-110 | 1.868941 | 0.686 | 0.277 | 3.0138E-106 |
| Intermediate | KRT8 | 1.2614E-133 | 1.867455 | 0.578 | 0.137 | 2.0717E-129 |
| Fibroblasts | OLFML3 | 0 | 1.85803 | 0.708 | 0.088 | 0 |
| RERGL_Cells | IGFBP5 | 6.8843E-105 | 1.849323 | 0.846 | 0.444 | 1.1307E-100 |
| Fibroblasts | ABCA8 | 0 | 1.848546 | 0.66 | 0.053 | 0 |
| Fibroblasts | VIT | 0 | 1.842278 | 0.582 | 0.005 | 0 |
| Fibroblasts | SOD2 | 2.9546E-196 | 1.821922 | 0.738 | 0.244 | 4.8527E-192 |
| Fibroblasts | SCARA5 | 0 | 1.81648 | 0.575 | 0.02 | 0 |
| Fibroblasts | TNXB | 3.0518E-274 | 1.813643 | 0.653 | 0.092 | 5.0122E-270 |
| Intermediate | DKK3 | 2.7445E-186 | 1.807731 | 0.959 | 0.451 | 4.5075E-182 |
| Fibroblasts | PCOLCE | 6.9581E-279 | 1.803583 | 0.926 | 0.389 | 1.1428E-274 |
| Pericytes | GGT5 | 4.6723E-176 | 1.802873 | 0.742 | 0.215 | 7.6738E-172 |
| Intermediate | AQP1 | 3.5716E-201 | 1.786195 | 0.873 | 0.246 | 5.8659E-197 |
| Pericytes | CNN3 | 5.6983E-189 | 1.777714 | 0.964 | 0.816 | 9.3589E-185 |
| Contractile | C12orf75 | 7.496E-186 | 1.766059 | 0.95 | 0.372 | 1.2311E-181 |
| Fibroblasts | IGFBP4 | 1.0789E-218 | 1.764957 | 0.929 | 0.439 | 1.772E-214 |
| Fibroblasts | CES1 | 7.3703E-219 | 1.764417 | 0.648 | 0.144 | 1.2105E-214 |
| RERGL_Cells | COX4I2 | 4.1168E-166 | 1.759903 | 0.519 | 0.07 | 6.7615E-162 |
| RERGL_Cells | HES4 | 2.8482E-133 | 1.753253 | 0.75 | 0.284 | 4.6779E-129 |
| Intermediate | MFGE8 | 1.1551E-192 | 1.743279 | 0.998 | 0.839 | 1.8971E-188 |

| | | | | | | |
|---|---|---|---|---|---|---|
| Pericytes | PHLDA1 | 1.88431E-94 | 1.740644 | 0.485 | 0.141 | 3.09479E-90 |
| RERGL_Cells | ISYNA1 | 6.6802E-124 | 1.723499 | 0.778 | 0.323 | 1.0972E-119 |
| Contractile | PPP1R14A | 1.5526E-180 | 1.715831 | 0.992 | 0.573 | 2.55E-176 |
| Pericytes | FILIP1L | 1.5599E-104 | 1.712018 | 0.809 | 0.46 | 2.562E-100 |
| Fibroblasts | BICC1 | 2.7512E-297 | 1.696462 | 0.645 | 0.07 | 4.5186E-293 |
| Pericytes | TINAGL1 | 5.2425E-182 | 1.692632 | 0.989 | 0.545 | 8.6103E-178 |
| Pericytes | SOCS3 | 1.0365E-99 | 1.686843 | 0.804 | 0.418 | 1.70236E-95 |
| Fibroblasts | FMO2 | 8.3984E-203 | 1.686021 | 0.691 | 0.192 | 1.3794E-198 |
| Fibroblasts | ADH1B | 2.3551E-251 | 1.684434 | 0.935 | 0.427 | 3.868E-247 |
| RERGL_Cells | NRGN | 6.6323E-148 | 1.679743 | 0.573 | 0.117 | 1.0893E-143 |
| Pericytes | APOE | 6.0026E-177 | 1.677093 | 0.865 | 0.289 | 9.8586E-173 |
| Fibroblasts | IGFBP6 | 1.7174E-212 | 1.671019 | 0.975 | 0.731 | 2.8206E-208 |
| Contractile | MYL9 | 1.1276E-194 | 1.669172 | 1 | 0.916 | 1.852E-190 |
| Pericytes | TGFB3 | 1.8084E-107 | 1.654581 | 0.411 | 0.084 | 2.9701E-103 |
| Contractile | DSTN | 3.1812E-205 | 1.646522 | 1 | 0.951 | 5.2247E-201 |
| Intermediate | TGM2 | 9.1523E-157 | 1.640066 | 0.75 | 0.212 | 1.5032E-152 |
| RERGL_Cells | GUCY1A1 | 1.2365E-116 | 1.634478 | 0.709 | 0.264 | 2.0308E-112 |
| RERGL_Cells | OAZ2 | 1.3386E-104 | 1.631325 | 0.812 | 0.485 | 2.1984E-100 |
| Contractile | SORBS2 | 6.4195E-164 | 1.625449 | 0.848 | 0.271 | 1.0543E-159 |
| Pericytes | TMEM176 | 5.1797E-155 | 1.617298 | 0.736 | 0.207 | 8.5071E-151 |
| Fibroblasts | NNMT | 5.0864E-254 | 1.607233 | 0.981 | 0.679 | 8.3539E-250 |
| Intermediate | AEBP1 | 4.8094E-162 | 1.600439 | 0.983 | 0.763 | 7.8989E-158 |
| RERGL_Cells | KCNAB1 | 1.6199E-158 | 1.594859 | 0.526 | 0.083 | 2.6605E-154 |
| Pericytes | CD44 | 1.7396E-124 | 1.589863 | 0.793 | 0.411 | 2.8571E-120 |
| Terminal | BGN | 5.8511E-123 | 1.587 | 0.983 | 0.823 | 9.6098E-119 |
| Intermediate | BGN | 3.2682E-173 | 1.58633 | 0.994 | 0.817 | 5.3676E-169 |
| Contractile | ACTA2 | 1.4368E-142 | 1.581395 | 0.998 | 0.746 | 2.3597E-138 |
| Intermediate | ASPN | 4.3119E-123 | 1.574418 | 0.811 | 0.32 | 7.0819E-119 |
| RERGL_Cells | UBA2 | 5.7985E-120 | 1.564922 | 0.801 | 0.398 | 9.5234E-116 |
| RERGL_Cells | ARPC1A | 4.2717E-120 | 1.562929 | 0.808 | 0.443 | 7.0159E-116 |
| Fibroblasts | CYBRD1 | 1.108E-251 | 1.560589 | 0.868 | 0.313 | 1.8198E-247 |
| Contractile | DES | 9.61425E-58 | 1.560154 | 0.303 | 0.07 | 1.57904E-53 |
| Intermediate | SLPI | 1.2362E-07 | 1.559808 | 0.209 | 0.131 | 0.002030329 |
| Pericytes | SYTL2 | 5.4864E-151 | 1.555884 | 0.544 | 0.107 | 9.0108E-147 |
| Intermediate | CTHRC1 | 5.2982E-120 | 1.555569 | 0.578 | 0.147 | 8.7018E-116 |
| Pericytes | CTSC | 4.3461E-111 | 1.552153 | 0.573 | 0.177 | 7.138E-107 |
| Contractile | CSRP1 | 9.1857E-161 | 1.547546 | 0.99 | 0.701 | 1.5087E-156 |
| Pericytes | INPP4B | 1.6383E-166 | 1.546457 | 0.544 | 0.093 | 2.6908E-162 |
| Fibroblasts | ABCA6 | 8.1175E-267 | 1.539247 | 0.539 | 0.038 | 1.3332E-262 |
| Fibroblasts | MEG3 | 2.1315E-234 | 1.535904 | 0.617 | 0.101 | 3.5007E-230 |
| Fibroblasts | G0S2 | 1.13532E-77 | 1.532371 | 0.263 | 0.048 | 1.86465E-73 |
| Pericytes | PTGER1 | 5.6022E-225 | 1.527299 | 0.38 | 0.011 | 9.2011E-221 |
| Fibroblasts | TNFSF13B | 5.3603E-209 | 1.514313 | 0.463 | 0.043 | 8.8038E-205 |
| Fibroblasts | SELENOP | 1.7025E-219 | 1.511108 | 0.911 | 0.476 | 2.7961E-215 |
| Terminal | SFRP4 | 9.88362E-64 | 1.508118 | 0.564 | 0.219 | 1.62329E-59 |
| RERGL_Cells | SORBS2 | 9.71748E-69 | 1.504728 | 0.643 | 0.307 | 1.596E-64 |
| RERGL_Cells | SERPINI1 | 1.8231E-106 | 1.502261 | 0.588 | 0.175 | 2.9943E-102 |
| Fibroblasts | ADD3 | 1.7693E-261 | 1.497653 | 0.664 | 0.11 | 2.9059E-257 |
| Contractile | FLNA | 1.8243E-130 | 1.495853 | 0.94 | 0.56 | 2.9962E-126 |
| Pericytes | ESAM | 1.4995E-142 | 1.492737 | 0.836 | 0.342 | 2.4628E-138 |
| Fibroblasts | PDGFRL | 7.0619E-210 | 1.487378 | 0.525 | 0.074 | 1.1598E-205 |
| Fibroblasts | ZFP36L2 | 1.2256E-186 | 1.484075 | 0.86 | 0.432 | 2.013E-182 |
| Intermediate | S100A4 | 7.9256E-129 | 1.477706 | 0.974 | 0.731 | 1.3017E-124 |
| Contractile | HSPB1 | 2.2756E-149 | 1.475857 | 1 | 0.942 | 3.7374E-145 |
| Fibroblasts | LRP1 | 1.0504E-212 | 1.474893 | 0.79 | 0.261 | 1.7251E-208 |
| Pericytes | A2M | 3.3703E-125 | 1.473389 | 0.927 | 0.647 | 5.5354E-121 |

| | | | | | | |
|---|---|---|---|---|---|---|
| Pericytes | MT1A | 4.77322E-70 | 1.472374 | 0.653 | 0.303 | 7.83953E-66 |
| Intermediate | KRT18 | 1.03403E-70 | 1.470721 | 0.335 | 0.073 | 1.6983E-66 |
| Contractile | LDOC1 | 2.6733E-131 | 1.468848 | 0.651 | 0.182 | 4.3906E-127 |
| Pericytes | HES1 | 1.31539E-74 | 1.467932 | 0.653 | 0.31 | 2.1604E-70 |
| Intermediate | LTBP1 | 7.2573E-140 | 1.464707 | 0.892 | 0.414 | 1.1919E-135 |
| Pericytes | TMEM176 | 1.1745E-129 | 1.463004 | 0.693 | 0.2 | 1.9289E-125 |
| Fibroblasts | SFRP1 | 1.623E-264 | 1.462886 | 0.459 | 0.011 | 2.6656E-260 |
| Fibroblasts | NR4A1 | 1.6013E-151 | 1.461705 | 0.771 | 0.32 | 2.6299E-147 |
| Fibroblasts | ID2 | 1.231E-137 | 1.456588 | 0.829 | 0.458 | 2.0218E-133 |
| Contractile | HSPB7 | 8.3428E-134 | 1.455632 | 0.649 | 0.167 | 1.3702E-129 |
| Fibroblasts | CCL2 | 9.5031E-114 | 1.455454 | 0.805 | 0.412 | 1.5608E-109 |
| Intermediate | MGP | 3.1775E-116 | 1.454042 | 1 | 0.951 | 5.2188E-112 |
| Fibroblasts | DHRS3 | 2.9442E-170 | 1.453672 | 0.575 | 0.138 | 4.8355E-166 |
| Fibroblasts | AKR1C1 | 2.1824E-221 | 1.447908 | 0.666 | 0.134 | 3.5843E-217 |
| Intermediate | SUGCT | 4.6904E-183 | 1.442742 | 0.677 | 0.127 | 7.7034E-179 |
| Fibroblasts | CST3 | 0 | 1.440979 | 0.999 | 0.975 | 0 |
| Pericytes | CCN1 | 2.70166E-54 | 1.436587 | 0.891 | 0.756 | 4.43721E-50 |
| Fibroblasts | PCOLCE2 | 1.3891E-128 | 1.434004 | 0.666 | 0.255 | 2.2814E-124 |
| Fibroblasts | LEPR | 6.8016E-208 | 1.433789 | 0.431 | 0.031 | 1.1171E-203 |
| Pericytes | TPPP3 | 4.66929E-96 | 1.432797 | 0.569 | 0.178 | 7.66884E-92 |
| RERGL_Cells | PRPH | 3.5792E-173 | 1.43257 | 0.295 | 0.008 | 5.8785E-169 |
| RERGL_Cells | TOB1 | 1.17023E-72 | 1.432096 | 0.628 | 0.292 | 1.92199E-68 |
| Pericytes | SSTR2 | 2.9636E-170 | 1.428443 | 0.315 | 0.012 | 4.8675E-166 |
| Terminal | CLU | 3.6489E-47 | 1.42409 | 0.978 | 0.804 | 5.99295E-43 |
| Fibroblasts | CFH | 2.9254E-196 | 1.41955 | 0.97 | 0.499 | 4.8046E-192 |
| RERGL_Cells | CSRP2 | 9.8569E-101 | 1.417998 | 0.765 | 0.322 | 1.61889E-96 |
| Pericytes | ANGPT2 | 7.7391E-99 | 1.408031 | 0.433 | 0.101 | 1.27106E-94 |
| RERGL_Cells | NDUFA4L2 | 5.50593E-77 | 1.406964 | 0.812 | 0.469 | 9.04294E-73 |
| Contractile | PDLIM3 | 3.2519E-119 | 1.404657 | 0.93 | 0.575 | 5.3409E-115 |
| Intermediate | OGN | 1.7455E-109 | 1.404055 | 0.877 | 0.461 | 2.8668E-105 |
| Pericytes | SERPINE1 | 2.49352E-17 | 1.403062 | 0.273 | 0.142 | 4.09535E-13 |
| RERGL_Cells | RBPMS2 | 4.658E-124 | 1.398815 | 0.459 | 0.081 | 7.6503E-120 |
| Contractile | CSRP2 | 2.2969E-121 | 1.397221 | 0.816 | 0.31 | 3.7725E-117 |
| Contractile | SBSPON | 4.5934E-150 | 1.395622 | 0.627 | 0.129 | 7.5443E-146 |
| RERGL_Cells | SPARCL1 | 1.2151E-140 | 1.395514 | 0.994 | 0.95 | 1.9956E-136 |
| Intermediate | LTBP2 | 8.2024E-146 | 1.393989 | 0.733 | 0.193 | 1.3472E-141 |
| Intermediate | THBS2 | 2.49399E-83 | 1.390807 | 0.69 | 0.312 | 4.09613E-79 |
| RERGL_Cells | PDK4 | 2.96809E-50 | 1.387195 | 0.697 | 0.434 | 4.87478E-46 |
| RERGL_Cells | NUDT4 | 3.74445E-83 | 1.386493 | 0.756 | 0.423 | 6.14988E-79 |
| Fibroblasts | CYGB | 1.9299E-217 | 1.379917 | 0.623 | 0.108 | 3.1697E-213 |
| Pericytes | ARHGAP15 | 6.9104E-156 | 1.378485 | 0.427 | 0.053 | 1.135E-151 |
| Fibroblasts | CD34 | 2.4981E-212 | 1.36854 | 0.527 | 0.071 | 4.1029E-208 |
| Fibroblasts | NAMPT | 1.6474E-175 | 1.367204 | 0.72 | 0.235 | 2.7056E-171 |
| RERGL_Cells | CPM | 2.8481E-111 | 1.365712 | 0.316 | 0.035 | 4.6778E-107 |
| Pericytes | CXCL12 | 2.09775E-96 | 1.364456 | 0.873 | 0.549 | 3.44535E-92 |
| Fibroblasts | PMP22 | 2.4489E-212 | 1.363908 | 0.906 | 0.464 | 4.0221E-208 |
| RERGL_Cells | TBX2-AS1 | 1.28619E-89 | 1.362758 | 0.474 | 0.126 | 2.11244E-85 |
| Contractile | MYH10 | 4.533E-110 | 1.360811 | 0.782 | 0.305 | 7.445E-106 |
| Fibroblasts | SERPINE2 | 1.4588E-195 | 1.359017 | 0.556 | 0.09 | 2.396E-191 |
| Fibroblasts | S100A10 | 7.288E-202 | 1.355736 | 0.939 | 0.527 | 1.197E-197 |
| Fibroblasts | CP | 2.4214E-198 | 1.35483 | 0.556 | 0.089 | 3.9768E-194 |
| RERGL_Cells | TSC22D1 | 7.40992E-78 | 1.352917 | 0.88 | 0.602 | 1.21701E-73 |
| Pericytes | GJA4 | 3.78884E-65 | 1.349453 | 0.433 | 0.142 | 6.2228E-61 |
| Pericytes | TSPAN12 | 7.1238E-136 | 1.347409 | 0.38 | 0.046 | 1.17E-131 |
| Contractile | CCDC3 | 9.2294E-104 | 1.347313 | 0.862 | 0.406 | 1.5158E-99 |
| Fibroblasts | MT1X | 3.90838E-55 | 1.346973 | 0.772 | 0.599 | 6.41912E-51 |

| | | | | | | |
|---|---|---|---|---|---|---|
| Contractile | LMOD1 | 6.1651E-125 | 1.343931 | 0.884 | 0.399 | 1.0126E-120 |
| Contractile | PALLD | 1.0196E-113 | 1.340458 | 0.892 | 0.517 | 1.6745E-109 |
| Intermediate | RAMP1 | 1.4979E-128 | 1.339433 | 0.864 | 0.299 | 2.4602E-124 |
| Contractile | SLC25A4 | 6.9691E-100 | 1.336362 | 0.792 | 0.394 | 1.14461E-95 |
| Pericytes | REM1 | 9.9838E-130 | 1.3339 | 0.418 | 0.065 | 1.6397E-125 |
| RERGL_Cells | KLHL23 | 5.1605E-122 | 1.333056 | 0.4 | 0.057 | 8.4756E-118 |
| Pericytes | DNAJB4 | 2.33194E-69 | 1.332437 | 0.622 | 0.318 | 3.82998E-65 |
| RERGL_Cells | AC079822 | 2.671E-149 | 1.331878 | 0.31 | 0.017 | 4.3868E-145 |
| RERGL_Cells | BTG1 | 1.21005E-89 | 1.331312 | 0.885 | 0.625 | 1.98739E-85 |
| RERGL_Cells | MTHFD2 | 1.43554E-91 | 1.326925 | 0.78 | 0.398 | 2.35773E-87 |
| RERGL_Cells | MAP3K20 | 2.1481E-82 | 1.324174 | 0.699 | 0.343 | 3.52803E-78 |
| Contractile | WFDC1 | 1.11957E-98 | 1.321032 | 0.599 | 0.183 | 1.83879E-94 |
| RERGL_Cells | PPP1R12A | 8.0373E-102 | 1.319133 | 0.85 | 0.472 | 1.32004E-97 |
| Fibroblasts | ABCA9 | 3.3416E-222 | 1.317199 | 0.421 | 0.019 | 5.4882E-218 |
| RERGL_Cells | PGF | 9.85376E-61 | 1.314196 | 0.53 | 0.215 | 1.61838E-56 |
| Pericytes | ZFP36 | 2.20345E-84 | 1.312333 | 0.945 | 0.705 | 3.61895E-80 |
| Pericytes | DEGS1 | 1.2577E-86 | 1.308165 | 0.727 | 0.438 | 2.06564E-82 |
| Fibroblasts | F3 | 1.7494E-132 | 1.307331 | 0.304 | 0.026 | 2.8733E-128 |
| Pericytes | DEPP1 | 0.008792669 | 1.307115 | 0.431 | 0.42 | 1 |
| Pericytes | ECRG4 | 2.31438E-35 | 1.305481 | 0.771 | 0.585 | 3.80114E-31 |
| RERGL_Cells | NTN4 | 5.67926E-69 | 1.304817 | 0.611 | 0.28 | 9.32762E-65 |
| Contractile | CARMN | 9.2379E-111 | 1.3043 | 0.673 | 0.205 | 1.5172E-106 |
| RERGL_Cells | AKAP1 | 6.75461E-87 | 1.303119 | 0.476 | 0.132 | 1.10938E-82 |
| RERGL_Cells | EDNRA | 7.367E-117 | 1.302626 | 0.436 | 0.074 | 1.21E-112 |
| Contractile | FABP3 | 1.5513E-131 | 1.301313 | 0.493 | 0.087 | 2.5478E-127 |
| RERGL_Cells | MCAM | 1.1994E-93 | 1.299407 | 0.686 | 0.26 | 1.9699E-89 |
| Contractile | PPP1R12B | 2.9506E-99 | 1.296535 | 0.689 | 0.255 | 4.84614E-95 |
| RERGL_Cells | CASQ2 | 1.35714E-86 | 1.296332 | 0.568 | 0.187 | 2.22896E-82 |
| Contractile | NEXN | 1.4433E-101 | 1.295582 | 0.854 | 0.468 | 2.3705E-97 |
| Contractile | RGS5 | 1.067E-110 | 1.294384 | 0.89 | 0.4 | 1.7525E-106 |
| Contractile | ACTN1 | 5.5378E-113 | 1.287293 | 0.904 | 0.578 | 9.0953E-109 |
| Pericytes | THBS1 | 1.3285E-38 | 1.286049 | 0.549 | 0.318 | 2.18193E-34 |
| Pericytes | ADGRF5 | 1.5585E-212 | 1.285294 | 0.329 | 0.004 | 2.5597E-208 |
| RERGL_Cells | TINAGL1 | 5.3703E-121 | 1.27768 | 0.974 | 0.559 | 8.8203E-117 |
| Contractile | CCN3 | 2.7767E-111 | 1.276053 | 0.83 | 0.313 | 4.5605E-107 |
| Pericytes | FHL5 | 2.28604E-97 | 1.275454 | 0.447 | 0.105 | 3.75459E-93 |
| Contractile | ALDH1B1 | 3.0512E-102 | 1.272803 | 0.493 | 0.12 | 5.01131E-98 |
| RERGL_Cells | ADIRF | 1.2644E-139 | 1.267559 | 0.998 | 0.908 | 2.0767E-135 |
| RERGL_Cells | SNCG | 1.6468E-42 | 1.266002 | 0.6 | 0.348 | 2.70471E-38 |
| Fibroblasts | CEBPD | 6.7745E-186 | 1.264194 | 0.965 | 0.705 | 1.1127E-181 |
| Intermediate | TIMP1 | 5.60468E-44 | 1.26368 | 0.996 | 0.916 | 9.20513E-40 |
| Pericytes | FAM162B | 2.4096E-122 | 1.261101 | 0.387 | 0.055 | 3.9576E-118 |
| Intermediate | SGCA | 1.3468E-117 | 1.256325 | 0.794 | 0.308 | 2.212E-113 |
| Fibroblasts | KLF4 | 2.7173E-142 | 1.249762 | 0.511 | 0.12 | 4.4629E-138 |
| RERGL_Cells | EPAS1 | 3.73824E-77 | 1.247204 | 0.778 | 0.477 | 6.13968E-73 |
| Pericytes | H19 | 1.75013E-76 | 1.246808 | 0.32 | 0.064 | 2.87442E-72 |
| Fibroblasts | ADM | 4.20974E-90 | 1.245688 | 0.239 | 0.029 | 6.91408E-86 |
| Fibroblasts | SERPING1 | 1.4331E-263 | 1.240811 | 0.994 | 0.888 | 2.3538E-259 |
| Intermediate | POSTN | 1.12148E-60 | 1.240323 | 0.271 | 0.053 | 1.84193E-56 |
| Fibroblasts | MT2A | 1.21195E-62 | 1.239872 | 0.94 | 0.927 | 1.99051E-58 |
| Pericytes | NOTCH3 | 1.02379E-86 | 1.239204 | 0.598 | 0.22 | 1.68147E-82 |
| Fibroblasts | FOS | 3.7341E-145 | 1.233311 | 0.903 | 0.51 | 6.1329E-141 |
| Terminal | AGT | 3.60104E-34 | 1.23175 | 0.556 | 0.3 | 5.91434E-30 |
| Fibroblasts | MARCKSL1 | 3.8915E-123 | 1.228366 | 0.306 | 0.033 | 6.3913E-119 |
| Pericytes | CFHR1 | 8.48338E-63 | 1.227985 | 0.107 | 0.003 | 1.39331E-58 |
| Pericytes | GEM | 6.83643E-37 | 1.225795 | 0.664 | 0.458 | 1.12281E-32 |

| | | | | | | |
|---|---|---|---|---|---|---|
| Fibroblasts | LSP1 | 3.3926E-189 | 1.222946 | 0.449 | 0.048 | 5.572E-185 |
| Fibroblasts | RNASE4 | 1.531E-183 | 1.22168 | 0.905 | 0.534 | 2.5145E-179 |
| Pericytes | COLEC11 | 4.36299E-89 | 1.219544 | 0.28 | 0.039 | 7.16578E-85 |
| Fibroblasts | ZNF385A | 5.1938E-172 | 1.218963 | 0.453 | 0.061 | 8.5302E-168 |
| Fibroblasts | S100A13 | 2.7221E-168 | 1.218909 | 0.814 | 0.355 | 4.4707E-164 |
| Fibroblasts | ZFP36 | 1.0395E-145 | 1.215662 | 0.945 | 0.654 | 1.7073E-141 |
| RERGL_Cells | ID1 | 5.78856E-55 | 1.210756 | 0.541 | 0.234 | 9.50713E-51 |
| Pericytes | CCL2 | 1.61524E-73 | 1.210197 | 0.829 | 0.476 | 2.65287E-69 |
| RERGL_Cells | SEPTIN4 | 1.10473E-48 | 1.209984 | 0.515 | 0.245 | 1.8144E-44 |
| RERGL_Cells | MEF2B | 6.5135E-85 | 1.208791 | 0.237 | 0.025 | 1.06978E-80 |
| Intermediate | MYH10 | 7.1405E-106 | 1.201713 | 0.787 | 0.299 | 1.1728E-101 |
| Pericytes | AGTR1 | 8.48099E-52 | 1.199806 | 0.32 | 0.099 | 1.39292E-47 |
| Intermediate | KRT17 | 3.4468E-97 | 1.199537 | 0.234 | 0.019 | 5.66103E-93 |
| Fibroblasts | PIM1 | 1.1619E-135 | 1.199193 | 0.531 | 0.131 | 1.9082E-131 |
| Contractile | ITGA8 | 3.5974E-104 | 1.19826 | 0.739 | 0.264 | 5.9084E-100 |
| Terminal | PRELP | 5.18478E-28 | 1.195896 | 0.824 | 0.707 | 8.51548E-24 |
| Pericytes | PARM1 | 1.96786E-93 | 1.194513 | 0.478 | 0.129 | 3.23201E-89 |
| Pericytes | NABP1 | 2.32138E-46 | 1.188485 | 0.355 | 0.131 | 3.81264E-42 |
| Intermediate | SERPINE1 | 1.88317E-72 | 1.18818 | 0.428 | 0.116 | 3.09292E-68 |
| Contractile | MYL6 | 4.2447E-151 | 1.18642 | 1 | 0.974 | 6.9714E-147 |
| RERGL_Cells | ENTPD3 | 1.4198E-157 | 1.186278 | 0.246 | 0.003 | 2.3318E-153 |
| Contractile | SPINT2 | 7.16549E-94 | 1.18569 | 0.647 | 0.207 | 1.17686E-89 |
| Intermediate | FMOD | 1.0149E-98 | 1.185023 | 0.811 | 0.392 | 1.66687E-94 |
| Intermediate | ITGA10 | 1.8684E-121 | 1.183334 | 0.492 | 0.094 | 3.0686E-117 |
| RERGL_Cells | CHCHD10 | 1.99189E-64 | 1.183176 | 0.694 | 0.378 | 3.27148E-60 |
| Fibroblasts | QSOX1 | 5.9197E-131 | 1.179071 | 0.723 | 0.319 | 9.7225E-127 |
| Intermediate | FXYD5 | 2.8593E-107 | 1.178457 | 0.68 | 0.222 | 4.6961E-103 |
| Terminal | CD9 | 3.70223E-86 | 1.177986 | 0.976 | 0.799 | 6.08055E-82 |
| Contractile | CALD1 | 1.9654E-125 | 1.177374 | 0.994 | 0.96 | 3.2279E-121 |
| Pericytes | RASD1 | 4.33225E-42 | 1.17147 | 0.335 | 0.116 | 7.11528E-38 |
| Pericytes | EDNRB | 2.80909E-68 | 1.171179 | 0.344 | 0.085 | 4.61366E-64 |
| RERGL_Cells | TBX2 | 1.02242E-73 | 1.170926 | 0.35 | 0.078 | 1.67922E-69 |
| RERGL_Cells | MYH11 | 5.49712E-98 | 1.170283 | 0.912 | 0.427 | 9.02847E-94 |
| Fibroblasts | CHI3L2 | 1.5647E-115 | 1.167381 | 0.223 | 0.008 | 2.5698E-111 |
| Fibroblasts | NEGR1 | 2.0396E-197 | 1.16554 | 0.445 | 0.042 | 3.3498E-193 |
| RERGL_Cells | PTK2 | 4.98127E-58 | 1.163355 | 0.607 | 0.297 | 8.18124E-54 |
| Intermediate | ELN | 7.3373E-84 | 1.158678 | 0.755 | 0.346 | 1.20508E-79 |
| Contractile | SPEG | 1.3663E-108 | 1.158376 | 0.485 | 0.103 | 2.244E-104 |
| Fibroblasts | PTGFR | 1.5619E-201 | 1.156833 | 0.375 | 0.014 | 2.5653E-197 |
| Contractile | KCNMB1 | 8.08881E-92 | 1.156263 | 0.717 | 0.276 | 1.32851E-87 |
| Terminal | FXYD5 | 2.04942E-61 | 1.156113 | 0.584 | 0.247 | 3.36597E-57 |
| Intermediate | C5orf46 | 3.344E-121 | 1.155653 | 0.34 | 0.037 | 5.4921E-117 |
| Fibroblasts | CRISPLD2 | 5.4133E-132 | 1.154104 | 0.593 | 0.169 | 8.8908E-128 |
| RERGL_Cells | CDH6 | 1.97911E-83 | 1.153685 | 0.38 | 0.078 | 3.2505E-79 |
| Fibroblasts | CYP1B1 | 1.7046E-122 | 1.151947 | 0.319 | 0.038 | 2.7996E-118 |
| Pericytes | APOLD1 | 4.24743E-91 | 1.149118 | 0.338 | 0.058 | 6.97598E-87 |
| Pericytes | PROCR | 1.15024E-66 | 1.145811 | 0.436 | 0.145 | 1.88915E-62 |
| Fibroblasts | OAF | 3.5275E-188 | 1.143543 | 0.447 | 0.047 | 5.7935E-184 |
| Fibroblasts | EFEMP1 | 4.6031E-169 | 1.143211 | 0.938 | 0.548 | 7.5602E-165 |
| RERGL_Cells | FABP4 | 5.5454E-207 | 1.141842 | 0.929 | 0.218 | 9.1078E-203 |
| Intermediate | KRT7 | 3.0779E-145 | 1.141139 | 0.308 | 0.019 | 5.0551E-141 |
| Pericytes | HGF | 7.36792E-29 | 1.141006 | 0.265 | 0.108 | 1.21011E-24 |
| Pericytes | THY1 | 5.21811E-38 | 1.139339 | 0.304 | 0.11 | 8.57022E-34 |
| Terminal | FMOD | 1.72459E-43 | 1.138923 | 0.678 | 0.422 | 2.83247E-39 |
| RERGL_Cells | ID3 | 2.62518E-68 | 1.134014 | 0.929 | 0.737 | 4.3116E-64 |
| RERGL_Cells | MYL9 | 6.0491E-109 | 1.13226 | 1 | 0.917 | 9.9351E-105 |

| | | | | | | |
|---|---|---|---|---|---|---|
| Pericytes | AGT | 3.74448E-68 | 1.131227 | 0.635 | 0.278 | 6.14994E-64 |
| Fibroblasts | PTGIS | 2.7048E-112 | 1.128903 | 0.606 | 0.232 | 4.4424E-108 |
| Fibroblasts | SAT1 | 2.60239E-98 | 1.127451 | 0.786 | 0.46 | 4.27416E-94 |
| Terminal | DKK3 | 1.23447E-70 | 1.127383 | 0.843 | 0.48 | 2.0275E-66 |
| Intermediate | COL21A1 | 1.5149E-125 | 1.126743 | 0.518 | 0.101 | 2.488E-121 |
| Pericytes | C11orf96 | 5.76083E-56 | 1.125949 | 0.827 | 0.606 | 9.46158E-52 |
| Pericytes | STOM | 1.08181E-73 | 1.125568 | 0.753 | 0.484 | 1.77676E-69 |
| Pericytes | ACTN4 | 2.95977E-81 | 1.125046 | 0.827 | 0.522 | 4.86113E-77 |
| Pericytes | HLA-B | 1.18637E-81 | 1.125025 | 0.895 | 0.66 | 1.94849E-77 |
| RERGL_Cells | CAV2 | 6.19977E-81 | 1.123642 | 0.842 | 0.504 | 1.01825E-76 |
| Fibroblasts | MT1M | 8.51419E-24 | 1.122686 | 0.634 | 0.508 | 1.39837E-19 |
| RERGL_Cells | ROCK1 | 5.94738E-67 | 1.122562 | 0.679 | 0.368 | 9.76798E-63 |
| Contractile | MYLK | 2.49436E-93 | 1.121263 | 0.874 | 0.429 | 4.09673E-89 |
| Pericytes | FHL2 | 1.25183E-60 | 1.121004 | 0.607 | 0.304 | 2.056E-56 |
| Fibroblasts | PLPP3 | 4.2735E-129 | 1.120804 | 0.671 | 0.252 | 7.0188E-125 |
| Fibroblasts | LXN | 1.2548E-133 | 1.120541 | 0.388 | 0.061 | 2.0608E-129 |
| Fibroblasts | COL6A3 | 3.2323E-107 | 1.119905 | 0.496 | 0.152 | 5.3087E-103 |
| Intermediate | FRZB | 8.74267E-70 | 1.118544 | 0.877 | 0.577 | 1.4359E-65 |
| Pericytes | TPM4 | 1.36745E-94 | 1.118293 | 0.904 | 0.72 | 2.2459E-90 |
| Pericytes | CYTOR | 2.00155E-41 | 1.117432 | 0.496 | 0.265 | 3.28735E-37 |
| Fibroblasts | FBN1 | 2.01319E-97 | 1.117155 | 0.467 | 0.143 | 3.30647E-93 |
| Pericytes | IGFBP5 | 8.44636E-75 | 1.117083 | 0.798 | 0.442 | 1.38723E-70 |
| Fibroblasts | FCGRT | 2.9131E-145 | 1.115934 | 0.808 | 0.395 | 4.7845E-141 |
| Pericytes | EPS8 | 2.55018E-82 | 1.114802 | 0.793 | 0.488 | 4.18841E-78 |
| RERGL_Cells | NDRG2 | 4.25241E-61 | 1.114706 | 0.72 | 0.432 | 6.98416E-57 |
| Intermediate | ITGA8 | 3.3718E-111 | 1.11454 | 0.761 | 0.255 | 5.5379E-107 |
| Fibroblasts | SPON2 | 4.7151E-149 | 1.114152 | 0.356 | 0.035 | 7.7441E-145 |
| Pericytes | PTK2 | 3.53519E-74 | 1.113429 | 0.625 | 0.285 | 5.80619E-70 |
| Fibroblasts | DAB2 | 6.4956E-146 | 1.11311 | 0.544 | 0.136 | 1.0668E-141 |
| RERGL_Cells | NRIP2 | 2.0248E-130 | 1.112066 | 0.278 | 0.016 | 3.3255E-126 |
| Fibroblasts | PDGFRA | 3.1613E-165 | 1.109752 | 0.493 | 0.087 | 5.1921E-161 |
| RERGL_Cells | GJA4 | 2.91528E-72 | 1.108647 | 0.479 | 0.143 | 4.78805E-68 |
| Fibroblasts | RARRES2 | 8.1585E-106 | 1.108641 | 0.877 | 0.56 | 1.3399E-101 |
| RERGL_Cells | FILIP1L | 2.00732E-49 | 1.107893 | 0.726 | 0.482 | 3.29683E-45 |
| RERGL_Cells | CARMN | 1.93977E-66 | 1.107866 | 0.577 | 0.224 | 3.18588E-62 |
| Fibroblasts | ALDH2 | 8.1413E-128 | 1.107114 | 0.799 | 0.421 | 1.3371E-123 |
| Fibroblasts | C16orf89 | 2.3481E-163 | 1.106942 | 0.288 | 0.004 | 3.8565E-159 |
| Pericytes | C20orf27 | 2.51423E-63 | 1.101079 | 0.453 | 0.166 | 4.12937E-59 |
| Fibroblasts | TSHZ2 | 5.8179E-206 | 1.100589 | 0.397 | 0.018 | 9.5554E-202 |
| Fibroblasts | HSD11B1 | 1.6126E-152 | 1.097883 | 0.34 | 0.027 | 2.6485E-148 |
| Fibroblasts | DUSP1 | 1.24048E-91 | 1.097477 | 0.9 | 0.689 | 2.03736E-87 |
| RERGL_Cells | CCDC3 | 1.6389E-80 | 1.097406 | 0.833 | 0.415 | 2.69173E-76 |
| Terminal | TSC22D3 | 3.47549E-47 | 1.096527 | 0.81 | 0.54 | 5.70814E-43 |
| RERGL_Cells | SYNE2 | 1.68289E-61 | 1.095728 | 0.417 | 0.13 | 2.76398E-57 |
| Terminal | LTBP2 | 2.67692E-61 | 1.095233 | 0.573 | 0.23 | 4.39657E-57 |
| Contractile | TNS1 | 2.27756E-78 | 1.09358 | 0.613 | 0.231 | 3.74066E-74 |
| Fibroblasts | S1PR2 | 3.8908E-172 | 1.09326 | 0.399 | 0.037 | 6.3903E-168 |
| Intermediate | CD151 | 8.2911E-110 | 1.093254 | 0.981 | 0.829 | 1.3617E-105 |
| Pericytes | ACTB | 1.3135E-136 | 1.090875 | 1 | 1 | 2.1574E-132 |
| Terminal | PCOLCE2 | 2.35072E-20 | 1.09034 | 0.527 | 0.357 | 3.86082E-16 |
| Intermediate | EDIL3 | 1.22904E-85 | 1.090235 | 0.613 | 0.211 | 2.01857E-81 |
| Contractile | RASSF3 | 3.42656E-77 | 1.090039 | 0.363 | 0.079 | 5.62778E-73 |
| Pericytes | SYNPO2 | 8.86521E-60 | 1.089784 | 0.616 | 0.297 | 1.45602E-55 |
| Pericytes | SOX4 | 2.72497E-19 | 1.089685 | 0.34 | 0.194 | 4.47549E-15 |
| Fibroblasts | FSTL1 | 4.33E-118 | 1.089637 | 0.853 | 0.502 | 7.1116E-114 |
| Pericytes | SLC38A2 | 7.23545E-38 | 1.088783 | 0.584 | 0.374 | 1.18835E-33 |

| | | | | | | |
|---|---|---|---|---|---|---|
| Intermediate | SPINT2 | 2.40087E-90 | 1.088007 | 0.634 | 0.204 | 3.9432E-86 |
| Fibroblasts | NOVA1 | 2.5215E-159 | 1.086009 | 0.481 | 0.082 | 4.1414E-155 |
| Intermediate | CTSD | 6.93322E-89 | 1.082591 | 0.936 | 0.723 | 1.13871E-84 |
| Fibroblasts | TMEM176 | 7.1745E-108 | 1.081717 | 0.534 | 0.164 | 1.1783E-103 |
| RERGL_Cells | MGST3 | 2.6571E-101 | 1.081581 | 0.966 | 0.875 | 4.36396E-97 |
| Pericytes | EIF4A1 | 6.6022E-114 | 1.080731 | 0.976 | 0.895 | 1.0843E-109 |
| Intermediate | RILPL2 | 4.42643E-89 | 1.076554 | 0.57 | 0.188 | 7.26998E-85 |
| RERGL_Cells | GUCY1B1 | 3.14964E-64 | 1.075101 | 0.442 | 0.141 | 5.17297E-60 |
| Terminal | SPINT2 | 4.97312E-49 | 1.073368 | 0.54 | 0.228 | 8.16785E-45 |
| Intermediate | SPARC | 2.13497E-98 | 1.072687 | 0.957 | 0.793 | 3.50647E-94 |
| Fibroblasts | FOSB | 1.91321E-85 | 1.072259 | 0.47 | 0.158 | 3.14225E-81 |
| Pericytes | ECEL1 | 5.5003E-146 | 1.069854 | 0.22 | 0.001 | 9.0337E-142 |
| Fibroblasts | LAMA2 | 1.9633E-163 | 1.069669 | 0.425 | 0.055 | 3.2245E-159 |
| RERGL_Cells | HEXIM1 | 3.41797E-55 | 1.069589 | 0.662 | 0.378 | 5.61367E-51 |
| Fibroblasts | JUNB | 2.8544E-140 | 1.068599 | 0.981 | 0.748 | 4.6881E-136 |
| Pericytes | IL6 | 4.41055E-09 | 1.068262 | 0.167 | 0.088 | 7.24389E-05 |
| Pericytes | NR2F2 | 1.93604E-60 | 1.068075 | 0.602 | 0.3 | 3.17975E-56 |
| Fibroblasts | TGFBR2 | 1.0719E-112 | 1.067906 | 0.54 | 0.169 | 1.7604E-108 |
| RERGL_Cells | ITGA7 | 7.61588E-70 | 1.065305 | 0.545 | 0.197 | 1.25083E-65 |
| Pericytes | CRIM1 | 7.02001E-42 | 1.064092 | 0.524 | 0.279 | 1.15297E-37 |
| Fibroblasts | RND3 | 1.1914E-117 | 1.063715 | 0.474 | 0.123 | 1.9567E-113 |
| Fibroblasts | NFE2L2 | 1.0373E-149 | 1.062829 | 0.864 | 0.471 | 1.7036E-145 |
| Pericytes | MEOX2 | 5.39851E-59 | 1.06272 | 0.389 | 0.125 | 8.86651E-55 |
| Pericytes | LHFPL6 | 2.58919E-70 | 1.061875 | 0.88 | 0.716 | 4.25249E-66 |
| Intermediate | EFEMP1 | 4.38292E-56 | 1.061731 | 0.908 | 0.623 | 7.19852E-52 |
| Terminal | SERPINE2 | 5.10821E-07 | 1.060102 | 0.309 | 0.218 | 0.008389723 |
| Fibroblasts | EPHX1 | 1.1963E-150 | 1.060055 | 0.804 | 0.338 | 1.9648E-146 |
| Intermediate | NDUFA4L2 | 3.70115E-43 | 1.059328 | 0.723 | 0.477 | 6.07877E-39 |
| Contractile | TAGLN | 2.8232E-121 | 1.059159 | 1 | 0.969 | 4.6369E-117 |
| Fibroblasts | CTSK | 1.6467E-143 | 1.058437 | 0.478 | 0.099 | 2.7045E-139 |
| Contractile | HACD1 | 2.75773E-88 | 1.057973 | 0.507 | 0.137 | 4.5293E-84 |
| RERGL_Cells | GPRC5C | 5.63572E-49 | 1.055891 | 0.485 | 0.21 | 9.25611E-45 |
| Fibroblasts | EGR1 | 8.4646E-170 | 1.055598 | 0.933 | 0.411 | 1.3902E-165 |
| Fibroblasts | LTBP4 | 1.0616E-137 | 1.054603 | 0.926 | 0.603 | 1.7436E-133 |
| Pericytes | ACTG1 | 1.847E-109 | 1.053786 | 0.995 | 0.991 | 3.0336E-105 |
| Fibroblasts | ECM1 | 1.2724E-100 | 1.05227 | 0.448 | 0.128 | 2.08971E-96 |
| Contractile | NDUFA4 | 1.6406E-117 | 1.051994 | 0.99 | 0.897 | 2.6945E-113 |
| Terminal | VCAN | 8.31891E-64 | 1.051066 | 0.756 | 0.377 | 1.3663E-59 |
| Pericytes | IER2 | 2.01096E-55 | 1.050634 | 0.793 | 0.564 | 3.3028E-51 |
| RERGL_Cells | HRC | 1.76706E-82 | 1.049294 | 0.35 | 0.066 | 2.90222E-78 |
| Fibroblasts | CFI | 4.3512E-185 | 1.045324 | 0.423 | 0.037 | 7.1465E-181 |
| RERGL_Cells | PCSK7 | 5.15034E-52 | 1.043801 | 0.673 | 0.388 | 8.45891E-48 |
| Fibroblasts | TIMP3 | 9.6902E-169 | 1.039835 | 0.992 | 0.715 | 1.5915E-164 |
| Contractile | VIM | 9.2284E-147 | 1.038691 | 1 | 0.998 | 1.5157E-142 |
| Intermediate | CDH13 | 2.70633E-95 | 1.037996 | 0.482 | 0.118 | 4.44487E-91 |
| Fibroblasts | HLA-DRB1 | 3.27584E-84 | 1.036189 | 0.438 | 0.142 | 5.38024E-80 |
| RERGL_Cells | ZFHX3 | 2.53454E-45 | 1.034616 | 0.427 | 0.18 | 4.16273E-41 |
| Fibroblasts | CDH19 | 1.10608E-78 | 1.031303 | 0.328 | 0.082 | 1.81662E-74 |
| Terminal | NPC2 | 6.38333E-59 | 1.028786 | 0.863 | 0.615 | 1.0484E-54 |
| Pericytes | FXYD6 | 8.7146E-60 | 1.027869 | 0.467 | 0.177 | 1.43129E-55 |
| Fibroblasts | COL4A4 | 2.2919E-155 | 1.026395 | 0.325 | 0.02 | 3.7641E-151 |
| RERGL_Cells | NTRK2 | 7.37885E-58 | 1.025517 | 0.716 | 0.429 | 1.2119E-53 |
| RERGL_Cells | ESAM | 9.93464E-66 | 1.025111 | 0.737 | 0.37 | 1.63167E-61 |
| Contractile | PDLIM7 | 1.03651E-76 | 1.024924 | 0.886 | 0.595 | 1.70236E-72 |
| RERGL_Cells | FBXL13 | 7.99162E-86 | 1.024019 | 0.224 | 0.02 | 1.31254E-81 |
| Fibroblasts | PAMR1 | 2.7431E-170 | 1.021335 | 0.316 | 0.01 | 4.5053E-166 |

| | | | | | | |
|---|---|---|---|---|---|---|
| Fibroblasts | STEAP1 | 3.9431E-148 | 1.020764 | 0.381 | 0.046 | 6.4761E-144 |
| Contractile | TCEAL2 | 1.01532E-80 | 1.0184 | 0.393 | 0.088 | 1.66756E-76 |
| Fibroblasts | ARL4D | 3.6362E-109 | 1.01812 | 0.368 | 0.074 | 5.9721E-105 |
| Fibroblasts | SLC43A3 | 9.7485E-149 | 1.016902 | 0.515 | 0.109 | 1.6011E-144 |
| Contractile | SH3BGR | 1.15202E-88 | 1.016585 | 0.473 | 0.121 | 1.89208E-84 |
| Pericytes | ID3 | 7.37472E-21 | 1.015735 | 0.791 | 0.757 | 1.21122E-16 |
| Intermediate | CPXM2 | 8.84513E-66 | 1.01565 | 0.613 | 0.268 | 1.45272E-61 |
| Fibroblasts | MATN2 | 6.0186E-144 | 1.014647 | 0.511 | 0.112 | 9.885E-140 |
| Contractile | CRYAB | 3.9332E-61 | 1.014098 | 0.938 | 0.778 | 6.45989E-57 |
| RERGL_Cells | PACSIN2 | 6.35379E-40 | 1.013563 | 0.487 | 0.246 | 1.04355E-35 |
| RERGL_Cells | DSTN | 1.09369E-98 | 1.013338 | 0.998 | 0.952 | 1.79628E-94 |
| Fibroblasts | GPNMB | 7.4131E-112 | 1.012701 | 0.747 | 0.371 | 1.2175E-107 |
| Fibroblasts | NEAT1 | 1.2184E-100 | 1.012563 | 0.902 | 0.655 | 2.00117E-96 |
| Pericytes | MAP3K8 | 2.16249E-59 | 1.012438 | 0.413 | 0.146 | 3.55167E-55 |
| RERGL_Cells | FILIP1 | 5.43313E-42 | 1.011135 | 0.459 | 0.21 | 8.92338E-38 |
| Fibroblasts | LOX | 1.1657E-170 | 1.009807 | 0.403 | 0.041 | 1.9146E-166 |
| Pericytes | CAV1 | 4.01824E-77 | 1.009607 | 0.935 | 0.695 | 6.59957E-73 |
| Pericytes | MYO1B | 4.82725E-43 | 1.00909 | 0.34 | 0.127 | 7.92828E-39 |
| Pericytes | EGR1 | 1.78519E-49 | 1.007874 | 0.807 | 0.524 | 2.932E-45 |
| Pericytes | NR4A1 | 1.09028E-34 | 1.005804 | 0.662 | 0.418 | 1.79068E-30 |
| Fibroblasts | SLIT2 | 4.9615E-145 | 1.004653 | 0.314 | 0.022 | 8.1488E-141 |
| Fibroblasts | CHRDL1 | 4.1451E-160 | 1.000687 | 0.381 | 0.037 | 6.8079E-156 |
| Contractile | PGAM2 | 8.78823E-73 | 0.999657 | 0.305 | 0.055 | 1.44338E-68 |
| RERGL_Cells | DMPK | 1.47789E-49 | 0.998553 | 0.429 | 0.165 | 2.42728E-45 |
| Contractile | SLMAP | 1.23779E-71 | 0.997855 | 0.563 | 0.199 | 2.03295E-67 |
| RERGL_Cells | MYLK | 2.5211E-69 | 0.997663 | 0.812 | 0.442 | 4.14066E-65 |
| Pericytes | LBR | 1.09403E-63 | 0.997238 | 0.391 | 0.122 | 1.79684E-59 |
| RERGL_Cells | MRGPRF | 9.69277E-42 | 0.996695 | 0.568 | 0.324 | 1.59194E-37 |
| Fibroblasts | AOX1 | 2.9135E-160 | 0.996574 | 0.344 | 0.024 | 4.7851E-156 |
| RERGL_Cells | RCAN2 | 1.42356E-51 | 0.99653 | 0.622 | 0.315 | 2.33806E-47 |
| Fibroblasts | IL33 | 1.9759E-136 | 0.993708 | 0.547 | 0.14 | 3.2452E-132 |
| Pericytes | SLC25A5 | 4.55369E-57 | 0.992703 | 0.715 | 0.49 | 7.47899E-53 |
| Contractile | SMTN | 2.54613E-60 | 0.992678 | 0.671 | 0.334 | 4.18176E-56 |
| Fibroblasts | CD248 | 1.3295E-152 | 0.992672 | 0.335 | 0.024 | 2.1836E-148 |
| Terminal | AEBP1 | 3.03942E-48 | 0.990627 | 0.913 | 0.779 | 4.99195E-44 |
| Fibroblasts | FAIM2 | 1.0284E-176 | 0.987521 | 0.327 | 0.01 | 1.6891E-172 |
| Fibroblasts | FIBIN | 2.20792E-93 | 0.986791 | 0.698 | 0.325 | 3.62629E-89 |
| Pericytes | COX4I2 | 5.73314E-67 | 0.986413 | 0.36 | 0.087 | 9.41611E-63 |
| Pericytes | VASP | 2.67781E-53 | 0.985835 | 0.498 | 0.221 | 4.39803E-49 |
| Contractile | AOPEP | 8.91257E-67 | 0.985667 | 0.727 | 0.376 | 1.4638E-62 |
| Pericytes | MRGPRF | 2.316E-43 | 0.984423 | 0.56 | 0.319 | 3.80379E-39 |
| Fibroblasts | BTG2 | 9.20385E-85 | 0.984156 | 0.774 | 0.44 | 1.51164E-80 |
| Pericytes | CCDC102B | 5.43622E-63 | 0.984082 | 0.313 | 0.077 | 8.92845E-59 |
| Pericytes | JUNB | 4.38966E-68 | 0.983558 | 0.953 | 0.793 | 7.20958E-64 |
| Fibroblasts | HAS2 | 1.16367E-94 | 0.982616 | 0.213 | 0.016 | 1.91122E-90 |
| Contractile | SVIL | 4.51076E-68 | 0.982182 | 0.609 | 0.249 | 7.40848E-64 |
| Pericytes | TMEM200 | 1.7979E-99 | 0.981636 | 0.231 | 0.018 | 2.95285E-95 |
| Pericytes | KLF6 | 5.02598E-24 | 0.981193 | 0.591 | 0.456 | 8.25466E-20 |
| Contractile | ILK | 2.27695E-72 | 0.981104 | 0.87 | 0.633 | 3.73966E-68 |
| Contractile | MCAM | 1.41529E-73 | 0.979828 | 0.667 | 0.259 | 2.32448E-69 |
| Fibroblasts | GAS1 | 4.5796E-150 | 0.979707 | 0.33 | 0.025 | 7.5215E-146 |
| Intermediate | PPP1R14A | 4.24142E-85 | 0.977962 | 0.976 | 0.571 | 6.96611E-81 |
| Fibroblasts | IL6 | 3.27739E-56 | 0.976951 | 0.222 | 0.048 | 5.38278E-52 |
| Contractile | ADIRF | 7.29451E-80 | 0.974188 | 0.998 | 0.907 | 1.19805E-75 |
| Terminal | GAP43 | 1.37694E-47 | 0.973495 | 0.211 | 0.04 | 2.26149E-43 |
| Contractile | NIBAN1 | 1.36088E-65 | 0.972388 | 0.553 | 0.211 | 2.23511E-61 |

| | | | | | | |
|---|---|---|---|---|---|---|
| Pericytes | ITGA7 | 3.45964E-62 | 0.971832 | 0.502 | 0.195 | 5.68211E-58 |
| RERGL_Cells | PTMA | 8.2069E-126 | 0.971741 | 1 | 0.993 | 1.3479E-121 |
| Fibroblasts | CYP4B1 | 1.049E-133 | 0.971426 | 0.333 | 0.035 | 1.723E-129 |
| Intermediate | CDH11 | 7.00876E-78 | 0.970773 | 0.464 | 0.129 | 1.15112E-73 |
| RERGL_Cells | WTIP | 4.14618E-50 | 0.969663 | 0.308 | 0.085 | 6.80969E-46 |
| Pericytes | ERRFI1 | 1.53488E-15 | 0.968801 | 0.38 | 0.26 | 2.52089E-11 |
| Fibroblasts | ABI3BP | 1.0414E-133 | 0.967556 | 0.506 | 0.118 | 1.7105E-129 |
| Fibroblasts | COL1A2 | 2.2344E-136 | 0.965911 | 0.939 | 0.605 | 3.6697E-132 |
| RERGL_Cells | MAP3K7CI | 7.7225E-35 | 0.965877 | 0.404 | 0.176 | 1.26834E-30 |
| RERGL_Cells | ENPEP | 5.66666E-44 | 0.965021 | 0.312 | 0.096 | 9.30692E-40 |
| Contractile | LPP | 4.21122E-59 | 0.964946 | 0.715 | 0.382 | 6.91652E-55 |
| RERGL_Cells | RELL1 | 1.1625E-33 | 0.964678 | 0.37 | 0.165 | 1.90928E-29 |
| RERGL_Cells | MAFB | 2.63017E-31 | 0.964154 | 0.355 | 0.157 | 4.3198E-27 |
| Contractile | TPM1 | 2.77504E-81 | 0.963716 | 0.982 | 0.789 | 4.55773E-77 |
| RERGL_Cells | COX7A2 | 1.18833E-91 | 0.963711 | 0.957 | 0.864 | 1.95171E-87 |
| Contractile | MOB2 | 3.72019E-54 | 0.963566 | 0.547 | 0.244 | 6.11004E-50 |
| Contractile | CRIP2 | 5.25349E-78 | 0.962777 | 0.944 | 0.704 | 8.62833E-74 |
| Fibroblasts | CYBA | 7.7382E-117 | 0.962683 | 0.712 | 0.315 | 1.2709E-112 |
| RERGL_Cells | NOTCH3 | 2.3445E-52 | 0.962197 | 0.547 | 0.238 | 3.85061E-48 |
| RERGL_Cells | DES | 2.84566E-30 | 0.962168 | 0.25 | 0.08 | 4.67371E-26 |
| Intermediate | VCAM1 | 6.26861E-26 | 0.960151 | 0.335 | 0.149 | 1.02956E-21 |
| Contractile | CRIP1 | 1.04778E-67 | 0.955895 | 0.986 | 0.852 | 1.72087E-63 |
| Fibroblasts | GFPT2 | 5.5871E-141 | 0.955261 | 0.319 | 0.026 | 9.1762E-137 |
| RERGL_Cells | NR2F2 | 7.1694E-39 | 0.953886 | 0.564 | 0.313 | 1.1775E-34 |
| RERGL_Cells | C11orf96 | 1.10105E-42 | 0.953312 | 0.825 | 0.613 | 1.80837E-38 |
| Contractile | SELENOW | 5.16804E-92 | 0.952501 | 0.95 | 0.782 | 8.48799E-88 |
| Fibroblasts | IL6ST | 8.5748E-110 | 0.951664 | 0.618 | 0.23 | 1.4083E-105 |
| Contractile | GAPDH | 3.7299E-125 | 0.951178 | 1 | 0.975 | 6.126E-121 |
| Fibroblasts | LIMA1 | 7.4204E-111 | 0.94818 | 0.697 | 0.294 | 1.2187E-106 |
| Contractile | TSTD1 | 2.96759E-57 | 0.945938 | 0.535 | 0.225 | 4.87397E-53 |
| Pericytes | PPP1R15A | 7.28337E-41 | 0.943936 | 0.651 | 0.445 | 1.19622E-36 |
| Fibroblasts | CYB5A | 2.4498E-117 | 0.942927 | 0.662 | 0.256 | 4.0236E-113 |
| RERGL_Cells | RASL12 | 3.44385E-45 | 0.942683 | 0.442 | 0.181 | 5.65618E-41 |
| Terminal | LTBP1 | 1.04642E-53 | 0.942281 | 0.758 | 0.445 | 1.71865E-49 |
| Pericytes | LRRC17 | 3.59967E-39 | 0.940975 | 0.433 | 0.2 | 5.9121E-35 |
| Contractile | FBXO32 | 2.93992E-49 | 0.940001 | 0.483 | 0.191 | 4.82852E-45 |
| Fibroblasts | OMD | 1.86445E-84 | 0.939701 | 0.543 | 0.207 | 3.06216E-80 |
| Intermediate | RGS2 | 4.57171E-51 | 0.939045 | 0.501 | 0.201 | 7.50858E-47 |
| Pericytes | LAMA4 | 2.92845E-39 | 0.938287 | 0.315 | 0.117 | 4.80968E-35 |
| Contractile | RCAN2 | 7.66189E-65 | 0.936925 | 0.677 | 0.303 | 1.25839E-60 |
| RERGL_Cells | TPM1 | 8.41643E-75 | 0.936605 | 0.979 | 0.791 | 1.38231E-70 |
| Fibroblasts | MT1A | 9.57834E-61 | 0.936016 | 0.554 | 0.272 | 1.57315E-56 |
| RERGL_Cells | BST2 | 4.04051E-47 | 0.934114 | 0.491 | 0.206 | 6.63614E-43 |
| RERGL_Cells | CAV1 | 7.56616E-79 | 0.934074 | 0.966 | 0.697 | 1.24267E-74 |
| Contractile | SYNPO2 | 1.24425E-48 | 0.933828 | 0.629 | 0.3 | 2.04356E-44 |
| Pericytes | PRXL2A | 4.87873E-36 | 0.933238 | 0.34 | 0.141 | 8.01283E-32 |
| Intermediate | ALCAM | 5.69349E-95 | 0.93294 | 0.396 | 0.076 | 9.35099E-91 |
| Pericytes | CDKN1A | 3.50928E-29 | 0.932438 | 0.607 | 0.416 | 5.76365E-25 |
| Intermediate | TSC22D3 | 1.38053E-52 | 0.931844 | 0.832 | 0.529 | 2.26738E-48 |
| Pericytes | SLC1A5 | 2.61674E-31 | 0.930123 | 0.362 | 0.176 | 4.29773E-27 |
| Fibroblasts | SAMHD1 | 1.0539E-106 | 0.92948 | 0.551 | 0.184 | 1.7309E-102 |
| Pericytes | KCNJ8 | 2.17269E-46 | 0.92868 | 0.276 | 0.079 | 3.56843E-42 |
| Fibroblasts | TFPI | 1.1636E-114 | 0.925917 | 0.824 | 0.417 | 1.9112E-110 |
| RERGL_Cells | TMEM38B | 3.36215E-38 | 0.925531 | 0.321 | 0.116 | 5.522E-34 |
| Contractile | KCNA5 | 6.68339E-81 | 0.924615 | 0.521 | 0.148 | 1.09768E-76 |
| Contractile | FBLIM1 | 4.94229E-64 | 0.924284 | 0.729 | 0.362 | 8.11721E-60 |

| | | | | | | |
|---|---|---|---|---|---|---|
| Pericytes | MAT2A | 2.30008E-36 | 0.924279 | 0.451 | 0.233 | 3.77766E-32 |
| Pericytes | DDX5 | 2.8335E-110 | 0.924041 | 0.987 | 0.933 | 4.6537E-106 |
| Fibroblasts | SVEP1 | 2.2067E-141 | 0.922676 | 0.285 | 0.014 | 3.6242E-137 |
| Contractile | HAND2 | 3.58138E-72 | 0.922646 | 0.397 | 0.102 | 5.88205E-68 |
| Intermediate | SULF1 | 2.41561E-39 | 0.92244 | 0.355 | 0.136 | 3.96739E-35 |
| Pericytes | BATF3 | 2.1974E-104 | 0.922402 | 0.255 | 0.022 | 3.609E-100 |
| Intermediate | IGFBP7 | 1.3484E-133 | 0.918555 | 1 | 1 | 2.2146E-129 |
| Fibroblasts | CNTFR | 2.7371E-151 | 0.91804 | 0.373 | 0.038 | 4.4954E-147 |
| Pericytes | VASN | 1.13514E-26 | 0.916287 | 0.516 | 0.346 | 1.86436E-22 |
| Pericytes | BMP2 | 1.51126E-89 | 0.915341 | 0.216 | 0.018 | 2.48209E-85 |
| Fibroblasts | SERPINA5 | 4.5787E-145 | 0.914571 | 0.263 | 0.006 | 7.5201E-141 |
| Fibroblasts | GLUL | 1.57366E-76 | 0.914192 | 0.781 | 0.51 | 2.58458E-72 |
| Intermediate | CRIP2 | 1.62306E-78 | 0.914051 | 0.946 | 0.701 | 2.66571E-74 |
| Contractile | SOD3 | 2.22158E-73 | 0.913806 | 0.976 | 0.881 | 3.64873E-69 |
| RERGL_Cells | KANK2 | 1.55275E-38 | 0.912745 | 0.562 | 0.328 | 2.55024E-34 |
| RERGL_Cells | EHD2 | 2.2947E-38 | 0.912734 | 0.573 | 0.353 | 3.76881E-34 |
| Fibroblasts | PRCP | 1.887E-109 | 0.911435 | 0.587 | 0.208 | 3.0991E-105 |
| Fibroblasts | TMEM176 | 1.21325E-92 | 0.911325 | 0.538 | 0.181 | 1.99264E-88 |
| RERGL_Cells | COX17 | 2.37737E-44 | 0.911158 | 0.662 | 0.427 | 3.9046E-40 |
| Fibroblasts | KLF10 | 1.26663E-76 | 0.910742 | 0.517 | 0.209 | 2.08031E-72 |
| RERGL_Cells | PLCG2 | 1.11823E-17 | 0.910574 | 0.342 | 0.191 | 1.83658E-13 |
| Terminal | COL3A1 | 1.22779E-07 | 0.909684 | 0.438 | 0.337 | 0.002016517 |
| Fibroblasts | LY6E | 9.8502E-101 | 0.909236 | 0.759 | 0.388 | 1.6178E-96 |
| RERGL_Cells | CRIP1 | 1.78181E-74 | 0.907992 | 0.985 | 0.854 | 2.92644E-70 |
| RERGL_Cells | NDUFA4 | 9.8169E-84 | 0.907635 | 0.987 | 0.898 | 1.61233E-79 |
| Fibroblasts | CCN1 | 1.8156E-117 | 0.905929 | 0.954 | 0.7 | 2.9819E-113 |
| Pericytes | MT1E | 1.38741E-22 | 0.905296 | 0.764 | 0.667 | 2.27868E-18 |
| RERGL_Cells | RGS5 | 1.24523E-10 | 0.904095 | 0.568 | 0.453 | 2.04516E-06 |
| Contractile | MAOB | 2.40493E-62 | 0.904061 | 0.535 | 0.196 | 3.94985E-58 |
| Contractile | MSRB3 | 3.35964E-58 | 0.903336 | 0.667 | 0.337 | 5.51787E-54 |
| Pericytes | CEBPD | 2.86975E-42 | 0.902766 | 0.922 | 0.758 | 4.71327E-38 |
| Pericytes | CYCS | 3.11179E-42 | 0.902625 | 0.693 | 0.515 | 5.1108E-38 |
| Contractile | SNCG | 2.89573E-33 | 0.901095 | 0.597 | 0.346 | 4.75594E-29 |
| RERGL_Cells | PPP2R5A | 8.69523E-40 | 0.900255 | 0.321 | 0.113 | 1.4281E-35 |
| Pericytes | MT2A | 1.09805E-40 | 0.899714 | 0.962 | 0.925 | 1.80343E-36 |
| RERGL_Cells | MEF2C | 9.1589E-35 | 0.899668 | 0.556 | 0.317 | 1.50426E-30 |
| RERGL_Cells | SLC25A6 | 1.00551E-76 | 0.898845 | 0.959 | 0.882 | 1.65146E-72 |
| Pericytes | FBXL3 | 1.19272E-39 | 0.897916 | 0.589 | 0.385 | 1.95892E-35 |
| RERGL_Cells | SLMAP | 3.25231E-27 | 0.897276 | 0.421 | 0.224 | 5.3416E-23 |
| Fibroblasts | SPON1 | 3.6954E-153 | 0.897267 | 0.364 | 0.035 | 6.0693E-149 |
| Terminal | SPARC | 8.56999E-44 | 0.897108 | 0.906 | 0.804 | 1.40753E-39 |
| RERGL_Cells | GADD45B | 1.48225E-30 | 0.896719 | 0.782 | 0.657 | 2.43445E-26 |
| Pericytes | LINC01197 | 4.40547E-44 | 0.896516 | 0.364 | 0.137 | 7.23555E-40 |
| Pericytes | MEF2C | 2.96129E-27 | 0.896446 | 0.509 | 0.319 | 4.86363E-23 |
| Intermediate | CD9 | 1.38349E-82 | 0.896349 | 0.979 | 0.794 | 2.27225E-78 |
| Fibroblasts | CCN5 | 1.34951E-81 | 0.895167 | 0.622 | 0.286 | 2.21643E-77 |
| Contractile | MAP3K7CI | 9.57083E-36 | 0.894927 | 0.413 | 0.173 | 1.57191E-31 |
| Pericytes | ARHGAP1 | 3.17801E-39 | 0.892617 | 0.393 | 0.174 | 5.21956E-35 |
| Contractile | SH3BGRL | 2.5578E-76 | 0.892593 | 0.924 | 0.723 | 4.20093E-72 |
| Fibroblasts | CD74 | 7.7011E-49 | 0.892473 | 0.555 | 0.31 | 1.26483E-44 |
| Pericytes | MYLK | 3.42302E-33 | 0.891519 | 0.685 | 0.455 | 5.62198E-29 |
| Pericytes | LAPTM4B | 2.20032E-48 | 0.891073 | 0.409 | 0.165 | 3.61381E-44 |
| Intermediate | ANTXR1 | 3.29571E-67 | 0.890722 | 0.546 | 0.195 | 5.41288E-63 |
| Pericytes | RDH5 | 1.37268E-57 | 0.888918 | 0.285 | 0.069 | 2.25449E-53 |
| Pericytes | TFPI | 9.72165E-53 | 0.888326 | 0.756 | 0.5 | 1.59668E-48 |
| Fibroblasts | DPYSL2 | 1.4692E-111 | 0.88607 | 0.65 | 0.238 | 2.4129E-107 |

| | | | | | | |
|---|---|---|---|---|---|---|
| Terminal | S100A4 | 4.33906E-53 | 0.88547 | 0.935 | 0.742 | 7.12648E-49 |
| RERGL_Cells | CLMN | 3.41237E-62 | 0.884567 | 0.278 | 0.053 | 5.60447E-58 |
| Pericytes | PCSK7 | 4.68857E-25 | 0.882461 | 0.578 | 0.398 | 7.7005E-21 |
| Intermediate | FBXO32 | 1.01342E-71 | 0.881419 | 0.544 | 0.177 | 1.66443E-67 |
| Contractile | CAMK2G | 2.64385E-63 | 0.881149 | 0.441 | 0.138 | 4.34227E-59 |
| Intermediate | CRIP1 | 8.806E-61 | 0.880542 | 0.983 | 0.851 | 1.4463E-56 |
| Pericytes | TPM1 | 1.74416E-74 | 0.879609 | 0.969 | 0.788 | 2.86461E-70 |
| Fibroblasts | SOCS3 | 1.7057E-120 | 0.879582 | 0.79 | 0.342 | 2.8015E-116 |
| Pericytes | LASP1 | 2.13811E-37 | 0.877622 | 0.422 | 0.209 | 3.51163E-33 |
| Pericytes | ACTA2 | 7.57654E-66 | 0.877523 | 0.987 | 0.744 | 1.24437E-61 |
| Intermediate | EPDR1 | 9.54377E-84 | 0.877374 | 0.507 | 0.143 | 1.56747E-79 |
| Contractile | CASQ2 | 1.23354E-60 | 0.87706 | 0.529 | 0.19 | 2.02596E-56 |
| Intermediate | IFITM10 | 2.3509E-108 | 0.876561 | 0.361 | 0.051 | 3.8611E-104 |
| Intermediate | ITGB1 | 5.43043E-82 | 0.87649 | 0.976 | 0.878 | 8.91893E-78 |
| Pericytes | LMCD1 | 9.07088E-10 | 0.875572 | 0.367 | 0.283 | 1.4898E-05 |
| RERGL_Cells | TPM2 | 2.48272E-80 | 0.874581 | 0.998 | 0.833 | 4.07761E-76 |
| Fibroblasts | LGALS3 | 2.2793E-106 | 0.874295 | 0.906 | 0.646 | 3.7435E-102 |
| RERGL_Cells | SH3BGRL | 1.33056E-61 | 0.873928 | 0.895 | 0.729 | 2.18531E-57 |
| Fibroblasts | C1QTNF3 | 6.93928E-63 | 0.873637 | 0.15 | 0.012 | 1.13971E-58 |
| RERGL_Cells | COX5B | 2.83742E-65 | 0.872672 | 0.895 | 0.811 | 4.66017E-61 |
| Contractile | GMDS | 5.04314E-45 | 0.872604 | 0.467 | 0.197 | 8.28285E-41 |
| Contractile | RASL12 | 8.41924E-57 | 0.872456 | 0.481 | 0.172 | 1.38278E-52 |
| Contractile | SELENOM | 2.6697E-106 | 0.871408 | 0.998 | 0.949 | 4.3848E-102 |
| Contractile | CFL2 | 2.54242E-42 | 0.870936 | 0.669 | 0.408 | 4.17567E-38 |
| Fibroblasts | CTSH | 1.2449E-130 | 0.870871 | 0.336 | 0.039 | 2.0447E-126 |
| Intermediate | CCN5 | 3.18208E-17 | 0.870235 | 0.553 | 0.358 | 5.22624E-13 |
| Fibroblasts | MARCH3 | 5.8463E-140 | 0.869676 | 0.309 | 0.023 | 9.602E-136 |
| Fibroblasts | PLSCR1 | 5.38889E-94 | 0.869575 | 0.544 | 0.192 | 8.85072E-90 |
| Contractile | MRPS6 | 1.69361E-48 | 0.868206 | 0.543 | 0.25 | 2.78159E-44 |
| Contractile | CHURC1 | 3.75274E-60 | 0.867703 | 0.824 | 0.55 | 6.16351E-56 |
| RERGL_Cells | AC091182 | 3.67432E-77 | 0.867004 | 0.194 | 0.016 | 6.0347E-73 |
| Fibroblasts | ITM2B | 5.5188E-200 | 0.866077 | 0.996 | 0.97 | 9.064E-196 |
| Fibroblasts | ABCA10 | 2.1157E-145 | 0.865611 | 0.259 | 0.004 | 3.4747E-141 |
| RERGL_Cells | PLK2 | 2.31428E-28 | 0.863743 | 0.338 | 0.15 | 3.80097E-24 |
| RERGL_Cells | AOC3 | 1.50577E-37 | 0.861517 | 0.607 | 0.37 | 2.47307E-33 |
| Pericytes | AOC3 | 4.91221E-36 | 0.859714 | 0.584 | 0.368 | 8.06782E-32 |
| Intermediate | GMDS | 1.77496E-69 | 0.859451 | 0.538 | 0.181 | 2.9152E-65 |
| Fibroblasts | C6 | 4.9869E-156 | 0.859226 | 0.273 | 0.003 | 8.1904E-152 |
| Terminal | KCNK17 | 2.09161E-28 | 0.858987 | 0.285 | 0.108 | 3.43527E-24 |
| Fibroblasts | COL6A2 | 2.2859E-137 | 0.858189 | 0.978 | 0.818 | 3.7544E-133 |
| Contractile | TCEAL4 | 1.92466E-52 | 0.857884 | 0.862 | 0.667 | 3.16107E-48 |
| Fibroblasts | SPRY1 | 5.63368E-69 | 0.857249 | 0.569 | 0.26 | 9.25276E-65 |
| RERGL_Cells | FAM241A | 4.7419E-26 | 0.857198 | 0.291 | 0.12 | 7.7881E-22 |
| Pericytes | TAGLN | 4.47588E-73 | 0.855961 | 1 | 0.968 | 7.35119E-69 |
| Pericytes | DLC1 | 6.64066E-37 | 0.855066 | 0.393 | 0.178 | 1.09066E-32 |
| Intermediate | FGF1 | 1.71974E-97 | 0.854879 | 0.402 | 0.075 | 2.8245E-93 |
| RERGL_Cells | ACTA2 | 1.33974E-65 | 0.853604 | 0.996 | 0.749 | 2.2004E-61 |
| Intermediate | NPC2 | 1.41253E-57 | 0.853501 | 0.879 | 0.606 | 2.31993E-53 |
| Intermediate | LRRC32 | 8.54269E-70 | 0.853302 | 0.482 | 0.148 | 1.40305E-65 |
| RERGL_Cells | C1QTNF1 | 9.8813E-32 | 0.853082 | 0.592 | 0.383 | 1.6229E-27 |
| RERGL_Cells | AK1 | 1.54593E-38 | 0.852391 | 0.686 | 0.487 | 2.53904E-34 |
| RERGL_Cells | AC023157 | 2.11516E-41 | 0.852373 | 0.269 | 0.076 | 3.47395E-37 |
| Contractile | PPP1R3C | 2.99816E-49 | 0.849274 | 0.385 | 0.129 | 4.92419E-45 |
| Intermediate | PRELP | 5.37997E-54 | 0.849027 | 0.935 | 0.685 | 8.83606E-50 |
| RERGL_Cells | WFDC2 | 8.31382E-93 | 0.846872 | 0.173 | 0.006 | 1.36546E-88 |
| Fibroblasts | STEAP2 | 5.7566E-102 | 0.846208 | 0.276 | 0.035 | 9.45471E-98 |

| | | | | | | |
|---|---|---|---|---|---|---|
| Pericytes | ANO1 | 1.3746E-51 | 0.84546 | 0.365 | 0.12 | 2.25764E-47 |
| RERGL_Cells | TMEM51 | 1.51488E-68 | 0.845293 | 0.248 | 0.038 | 2.48804E-64 |
| Terminal | HAPLN1 | 1.34171E-59 | 0.844585 | 0.305 | 0.068 | 2.20362E-55 |
| Intermediate | MAOB | 5.12942E-60 | 0.844489 | 0.529 | 0.193 | 8.42457E-56 |
| RERGL_Cells | WDR1 | 1.45264E-44 | 0.844291 | 0.75 | 0.53 | 2.38582E-40 |
| Intermediate | TMEM204 | 1.16969E-47 | 0.844104 | 0.538 | 0.239 | 1.9211E-43 |
| RERGL_Cells | TBC1D1 | 1.92466E-43 | 0.843809 | 0.361 | 0.128 | 3.16107E-39 |
| Pericytes | HSPA2 | 2.45964E-36 | 0.843553 | 0.362 | 0.15 | 4.03972E-32 |
| Fibroblasts | GALNT15 | 1.1314E-109 | 0.843522 | 0.268 | 0.025 | 1.8582E-105 |
| Fibroblasts | BNIP3L | 8.22425E-93 | 0.841589 | 0.761 | 0.42 | 1.35075E-88 |
| Contractile | FRZB | 5.96029E-29 | 0.841152 | 0.776 | 0.596 | 9.78918E-25 |
| Pericytes | SLC2A3 | 1.84321E-37 | 0.840756 | 0.351 | 0.143 | 3.02729E-33 |
| Terminal | TMSB4X | 2.85766E-71 | 0.840002 | 1 | 0.997 | 4.69343E-67 |
| RERGL_Cells | APOLD1 | 1.66159E-30 | 0.839822 | 0.248 | 0.079 | 2.72899E-26 |
| Fibroblasts | GLRX | 4.00377E-65 | 0.83935 | 0.419 | 0.158 | 6.5758E-61 |
| RERGL_Cells | ANGPTL4 | 2.74382E-36 | 0.838796 | 0.566 | 0.303 | 4.50645E-32 |
| RERGL_Cells | ATP5F1B | 6.74656E-52 | 0.838591 | 0.865 | 0.756 | 1.10806E-47 |
| Fibroblasts | CLEC2B | 2.7259E-117 | 0.838546 | 0.365 | 0.059 | 4.477E-113 |
| Contractile | TCEAL3 | 8.40886E-42 | 0.837708 | 0.607 | 0.338 | 1.38107E-37 |
| Fibroblasts | ALDH1A1 | 3.13966E-90 | 0.837426 | 0.53 | 0.183 | 5.15657E-86 |
| Contractile | MFAP4 | 2.2029E-57 | 0.837186 | 0.956 | 0.782 | 3.61805E-53 |
| Contractile | SLC22A3 | 7.61424E-77 | 0.837044 | 0.353 | 0.072 | 1.25056E-72 |
| Pericytes | MYL12A | 2.20086E-79 | 0.834269 | 0.975 | 0.908 | 3.61468E-75 |
| RERGL_Cells | PLN | 3.34207E-75 | 0.833795 | 0.934 | 0.495 | 5.48901E-71 |
| RERGL_Cells | KCNA5 | 8.51863E-30 | 0.833764 | 0.38 | 0.173 | 1.3991E-25 |
| Fibroblasts | TSPAN4 | 2.3677E-102 | 0.833193 | 0.769 | 0.376 | 3.88871E-98 |
| Terminal | PLXDC2 | 2.82848E-41 | 0.832131 | 0.436 | 0.18 | 4.6455E-37 |
| Fibroblasts | UGDH | 7.64337E-75 | 0.832028 | 0.524 | 0.215 | 1.25535E-70 |
| RERGL_Cells | NTRK3 | 2.31395E-35 | 0.830616 | 0.329 | 0.124 | 3.80042E-31 |
| Terminal | TFPI2 | 3.9354E-41 | 0.829691 | 0.224 | 0.051 | 6.4635E-37 |
| Contractile | CSDC2 | 1.12224E-49 | 0.829423 | 0.389 | 0.128 | 1.84317E-45 |
| Contractile | CCDC107 | 1.33102E-51 | 0.82932 | 0.772 | 0.501 | 2.18607E-47 |
| Pericytes | HSPA5 | 1.42451E-21 | 0.828046 | 0.764 | 0.695 | 2.33962E-17 |
| RERGL_Cells | ENTPD1 | 1.78869E-52 | 0.827663 | 0.282 | 0.068 | 2.93774E-48 |
| Pericytes | ETS2 | 5.21111E-23 | 0.827519 | 0.405 | 0.239 | 8.55873E-19 |
| Pericytes | CDH6 | 5.13165E-55 | 0.826883 | 0.315 | 0.082 | 8.42822E-51 |
| Fibroblasts | LARP6 | 3.68933E-94 | 0.826748 | 0.575 | 0.221 | 6.05935E-90 |
| RERGL_Cells | HSDL2 | 6.37311E-34 | 0.826183 | 0.363 | 0.157 | 1.04672E-29 |
| Pericytes | THBD | 5.83477E-18 | 0.825806 | 0.238 | 0.115 | 9.58303E-14 |
| Fibroblasts | FGF7 | 2.8178E-172 | 0.825738 | 0.862 | 0.291 | 4.6279E-168 |
| Pericytes | DNAJB5 | 4.20271E-47 | 0.825436 | 0.307 | 0.095 | 6.90253E-43 |
| RERGL_Cells | DBI | 1.14984E-33 | 0.824657 | 0.688 | 0.539 | 1.8885E-29 |
| Fibroblasts | LRRN4CL | 1.2316E-159 | 0.824588 | 0.311 | 0.013 | 2.0227E-155 |
| Contractile | GADD45G | 1.31856E-31 | 0.822372 | 0.345 | 0.136 | 2.1656E-27 |
| Pericytes | ASAP1 | 1.01512E-60 | 0.821303 | 0.26 | 0.054 | 1.66724E-56 |
| Pericytes | COCH | 5.83751E-58 | 0.82082 | 0.224 | 0.04 | 9.58753E-54 |
| Pericytes | DDX21 | 2.65325E-21 | 0.81941 | 0.498 | 0.361 | 4.35769E-17 |
| Fibroblasts | CTSF | 5.79412E-78 | 0.819274 | 0.745 | 0.415 | 9.51627E-74 |
| Fibroblasts | CDC42EP2 | 7.4162E-108 | 0.817699 | 0.344 | 0.059 | 1.218E-103 |
| Pericytes | CNN1 | 2.40115E-34 | 0.817463 | 0.696 | 0.433 | 3.94365E-30 |
| Contractile | DYNLT1 | 7.68929E-34 | 0.817321 | 0.657 | 0.463 | 1.26289E-29 |
| Pericytes | TYMP | 6.37539E-31 | 0.816815 | 0.451 | 0.244 | 1.04709E-26 |
| Terminal | TGM2 | 6.46541E-34 | 0.816386 | 0.516 | 0.26 | 1.06188E-29 |
| Fibroblasts | ICAM1 | 6.80619E-81 | 0.816211 | 0.273 | 0.049 | 1.11785E-76 |
| Intermediate | PTTG1IP | 2.06357E-53 | 0.815145 | 0.656 | 0.339 | 3.3892E-49 |
| Fibroblasts | CILP | 1.6761E-108 | 0.814347 | 0.194 | 0.002 | 2.7528E-104 |

| | | | | | | |
|---|---|---|---|---|---|---|
| RERGL_Cells | MYL6 | 6.13828E-85 | 0.81395 | 1 | 0.974 | 1.00815E-80 |
| Intermediate | CST6 | 1.47965E-87 | 0.813051 | 0.219 | 0.019 | 2.43017E-83 |
| Pericytes | MYADM | 5.06481E-30 | 0.812945 | 0.625 | 0.451 | 8.31844E-26 |
| Pericytes | SUCNR1 | 1.22563E-79 | 0.811728 | 0.12 | 0.001 | 2.01298E-75 |
| Intermediate | WFDC1 | 5.60305E-63 | 0.810964 | 0.551 | 0.186 | 9.20244E-59 |
| Contractile | EFHD1 | 7.81747E-51 | 0.810362 | 0.703 | 0.363 | 1.28394E-46 |
| Terminal | ISLR | 4.04825E-18 | 0.809722 | 0.407 | 0.26 | 6.64884E-14 |
| Intermediate | COL4A2 | 4.03377E-42 | 0.809003 | 0.621 | 0.336 | 6.62507E-38 |
| Intermediate | P4HA3 | 7.5418E-114 | 0.807917 | 0.279 | 0.023 | 1.2387E-109 |
| Contractile | RNF115 | 1.73225E-38 | 0.807085 | 0.445 | 0.205 | 2.84504E-34 |
| Contractile | TGFB1I1 | 4.25292E-59 | 0.806684 | 0.862 | 0.585 | 6.98499E-55 |
| Fibroblasts | COLEC12 | 8.7458E-126 | 0.80572 | 0.294 | 0.026 | 1.4364E-121 |
| Terminal | LEFTY2 | 8.42106E-46 | 0.805692 | 0.205 | 0.039 | 1.38308E-41 |
| Pericytes | SLC2A4 | 4.8629E-59 | 0.805294 | 0.213 | 0.034 | 7.98683E-55 |
| RERGL_Cells | HNRNPA1 | 3.79326E-59 | 0.804032 | 0.966 | 0.87 | 6.23005E-55 |
| Fibroblasts | MGP | 5.3264E-134 | 0.802213 | 1 | 0.94 | 8.7481E-130 |
| Pericytes | AXL | 1.28079E-28 | 0.80219 | 0.627 | 0.452 | 2.10357E-24 |
| Pericytes | MAP7D3 | 2.3908E-29 | 0.801211 | 0.495 | 0.298 | 3.92665E-25 |
| Fibroblasts | TMEM45A | 7.99486E-73 | 0.799033 | 0.449 | 0.166 | 1.31308E-68 |
| RERGL_Cells | ETS2 | 4.23641E-26 | 0.79886 | 0.429 | 0.24 | 6.95789E-22 |
| RERGL_Cells | GPR20 | 2.8692E-47 | 0.798689 | 0.222 | 0.045 | 4.71237E-43 |
| Pericytes | FAM241A | 3.32088E-46 | 0.797866 | 0.329 | 0.108 | 5.45421E-42 |
| Pericytes | HEXIM1 | 2.35725E-21 | 0.797182 | 0.535 | 0.393 | 3.87155E-17 |
| RERGL_Cells | RRAD | 3.72375E-23 | 0.797123 | 0.248 | 0.099 | 6.11589E-19 |
| RERGL_Cells | FAM162B | 9.03883E-58 | 0.796293 | 0.318 | 0.074 | 1.48454E-53 |
| Pericytes | CREB5 | 2.3991E-26 | 0.795944 | 0.191 | 0.062 | 3.94029E-22 |
| Contractile | SNRPN | 3.747E-43 | 0.795485 | 0.725 | 0.475 | 6.15407E-39 |
| Pericytes | DNAJB1 | 1.01085E-24 | 0.794748 | 0.373 | 0.202 | 1.66021E-20 |
| Fibroblasts | SEMA3C | 4.20312E-83 | 0.794715 | 0.27 | 0.048 | 6.90321E-79 |
| Pericytes | CHCHD10 | 2.78682E-40 | 0.794432 | 0.629 | 0.381 | 4.57707E-36 |
| Contractile | MGST3 | 2.82896E-75 | 0.794333 | 0.962 | 0.875 | 4.64628E-71 |
| Intermediate | ADI1 | 4.6309E-59 | 0.79382 | 0.871 | 0.568 | 7.60579E-55 |
| RERGL_Cells | HIST1H4C | 6.23814E-22 | 0.792338 | 0.389 | 0.224 | 1.02455E-17 |
| Pericytes | SPRY2 | 4.57224E-33 | 0.792252 | 0.318 | 0.13 | 7.50944E-29 |
| Fibroblasts | GAS7 | 7.2874E-129 | 0.79183 | 0.288 | 0.022 | 1.1969E-124 |
| Pericytes | PCDH9 | 6.70851E-28 | 0.790936 | 0.345 | 0.163 | 1.10181E-23 |
| Pericytes | EPHX1 | 8.89998E-32 | 0.790261 | 0.665 | 0.444 | 1.46173E-27 |
| RERGL_Cells | ACTN4 | 1.22747E-40 | 0.789896 | 0.763 | 0.539 | 2.01599E-36 |
| Fibroblasts | CSRNP1 | 4.74615E-83 | 0.788925 | 0.492 | 0.166 | 7.79508E-79 |
| RERGL_Cells | PLA2G5 | 2.25173E-05 | 0.788461 | 0.288 | 0.238 | 0.369824942 |
| RERGL_Cells | DBNDD2 | 9.81024E-36 | 0.788121 | 0.752 | 0.563 | 1.61123E-31 |
| Fibroblasts | BOC | 1.3665E-130 | 0.788107 | 0.293 | 0.022 | 2.2444E-126 |
| Fibroblasts | CYP27A1 | 7.8904E-102 | 0.786758 | 0.472 | 0.135 | 1.29592E-97 |
| Pericytes | HNRNPDL | 5.0449E-56 | 0.785397 | 0.887 | 0.808 | 8.28574E-52 |
| Pericytes | HSPB6 | 8.80228E-23 | 0.784542 | 0.62 | 0.477 | 1.44569E-18 |
| Contractile | CD151 | 1.21829E-66 | 0.784385 | 0.958 | 0.834 | 2.00093E-62 |
| RERGL_Cells | SMTN | 5.30609E-35 | 0.784218 | 0.594 | 0.349 | 8.71472E-31 |
| RERGL_Cells | A2M | 1.87751E-49 | 0.784209 | 0.872 | 0.663 | 3.08362E-45 |
| Pericytes | SLC25A25 | 3.63945E-43 | 0.783442 | 0.302 | 0.099 | 5.97743E-39 |
| Pericytes | PDLIM1 | 9.66838E-36 | 0.782354 | 0.716 | 0.542 | 1.58793E-31 |
| RERGL_Cells | LGALSL | 6.68253E-37 | 0.781854 | 0.25 | 0.073 | 1.09754E-32 |
| Pericytes | PAWR | 1.18343E-36 | 0.781331 | 0.375 | 0.159 | 1.94366E-32 |
| RERGL_Cells | MYOZ1 | 1.16347E-35 | 0.780795 | 0.368 | 0.148 | 1.91088E-31 |
| Fibroblasts | CLDN11 | 9.9097E-104 | 0.780607 | 0.241 | 0.02 | 1.6276E-99 |
| Pericytes | SLC7A2 | 5.28227E-58 | 0.780274 | 0.231 | 0.042 | 8.6756E-54 |
| Fibroblasts | RBMS3 | 5.1964E-64 | 0.780131 | 0.584 | 0.272 | 8.53457E-60 |

| | | | | | | |
|---|---|---|---|---|---|---|
| Fibroblasts | RAMP2 | 1.1817E-118 | 0.779867 | 0.302 | 0.032 | 1.9408E-114 |
| Fibroblasts | IL32 | 6.06015E-85 | 0.77971 | 0.581 | 0.225 | 9.9532E-81 |
| Contractile | TSPAN2 | 2.07771E-62 | 0.778313 | 0.307 | 0.068 | 3.41243E-58 |
| Pericytes | STX12 | 5.1529E-31 | 0.778036 | 0.453 | 0.254 | 8.46313E-27 |
| Pericytes | KLF11 | 3.64209E-46 | 0.77743 | 0.295 | 0.087 | 5.98176E-42 |
| Intermediate | FAP | 1.27257E-74 | 0.77661 | 0.307 | 0.057 | 2.09007E-70 |
| Fibroblasts | TYMP | 1.5589E-83 | 0.776413 | 0.506 | 0.177 | 2.56034E-79 |
| Pericytes | TMEM119 | 2.40561E-66 | 0.776185 | 0.189 | 0.022 | 3.95097E-62 |
| Contractile | PLS3 | 1.9843E-50 | 0.775379 | 0.896 | 0.709 | 3.25902E-46 |
| Contractile | UQCRB | 2.19984E-64 | 0.774988 | 0.952 | 0.848 | 3.61302E-60 |
| RERGL_Cells | KCNMB1 | 4.62558E-34 | 0.774778 | 0.558 | 0.305 | 7.59705E-30 |
| Terminal | AQP1 | 2.98648E-37 | 0.774097 | 0.597 | 0.302 | 4.90499E-33 |
| Pericytes | HIF1A | 9.2193E-27 | 0.773911 | 0.48 | 0.29 | 1.51418E-22 |
| RERGL_Cells | MGLL | 5.16124E-22 | 0.77372 | 0.496 | 0.34 | 8.47682E-18 |
| Contractile | PFN1 | 3.45743E-68 | 0.772983 | 0.98 | 0.91 | 5.67849E-64 |
| Fibroblasts | CCNL1 | 4.80482E-67 | 0.772455 | 0.68 | 0.35 | 7.89143E-63 |
| Fibroblasts | MIR99AHG | 2.63932E-80 | 0.772401 | 0.446 | 0.146 | 4.33482E-76 |
| RERGL_Cells | TAGLN | 7.18369E-66 | 0.772257 | 1 | 0.969 | 1.17985E-61 |
| Fibroblasts | ARRDC3 | 1.52587E-66 | 0.772137 | 0.324 | 0.093 | 2.50609E-62 |
| Fibroblasts | THBS1 | 2.75095E-67 | 0.772046 | 0.58 | 0.256 | 4.51817E-63 |
| Pericytes | CNGA1 | 3.40893E-71 | 0.771825 | 0.147 | 0.008 | 5.59882E-67 |
| Pericytes | CD59 | 2.39863E-54 | 0.771406 | 0.904 | 0.851 | 3.93952E-50 |
| Fibroblasts | CDO1 | 4.1358E-108 | 0.769705 | 0.359 | 0.065 | 6.7927E-104 |
| Intermediate | TSPO | 9.62372E-59 | 0.769471 | 0.865 | 0.59 | 1.5806E-54 |
| Fibroblasts | GSTM5 | 1.304E-102 | 0.768688 | 0.344 | 0.064 | 2.14176E-98 |
| Contractile | CAMTA1 | 1.24963E-42 | 0.768223 | 0.705 | 0.451 | 2.0524E-38 |
| Pericytes | AVPR1A | 1.51547E-82 | 0.76817 | 0.193 | 0.014 | 2.48901E-78 |
| RERGL_Cells | CAVIN3 | 4.0227E-39 | 0.767813 | 0.857 | 0.723 | 6.60689E-35 |
| Pericytes | EAF1 | 5.47599E-41 | 0.767372 | 0.218 | 0.055 | 8.99376E-37 |
| Fibroblasts | NDRG1 | 8.71131E-92 | 0.767218 | 0.492 | 0.155 | 1.43075E-87 |
| Intermediate | LIMCH1 | 2.32751E-59 | 0.766675 | 0.473 | 0.16 | 3.82271E-55 |
| Fibroblasts | BZW1 | 1.04344E-71 | 0.766193 | 0.758 | 0.469 | 1.71374E-67 |
| Fibroblasts | MCL1 | 3.46136E-67 | 0.766178 | 0.74 | 0.448 | 5.68493E-63 |
| RERGL_Cells | CHMP4C | 1.75436E-61 | 0.766099 | 0.194 | 0.024 | 2.88136E-57 |
| Fibroblasts | ACKR3 | 1.75235E-66 | 0.766016 | 0.196 | 0.027 | 2.87806E-62 |
| Pericytes | UGCG | 2.966E-26 | 0.765884 | 0.324 | 0.155 | 4.87135E-22 |
| Fibroblasts | FSTL3 | 2.05048E-51 | 0.765195 | 0.516 | 0.258 | 3.36771E-47 |
| Terminal | RGS2 | 1.30565E-15 | 0.76511 | 0.381 | 0.226 | 2.1444E-11 |
| Fibroblasts | GASK1B | 5.22917E-94 | 0.761658 | 0.355 | 0.077 | 8.58839E-90 |
| Terminal | MYH10 | 2.1352E-31 | 0.761111 | 0.617 | 0.336 | 3.50685E-27 |
| Intermediate | MYH9 | 5.45198E-41 | 0.760841 | 0.658 | 0.358 | 8.95434E-37 |
| Fibroblasts | TNFAIP6 | 1.32766E-51 | 0.76072 | 0.29 | 0.09 | 2.18055E-47 |
| Fibroblasts | DBN1 | 1.90683E-63 | 0.760372 | 0.42 | 0.162 | 3.13178E-59 |
| Fibroblasts | SHISA3 | 3.2493E-65 | 0.759971 | 0.172 | 0.02 | 5.33665E-61 |
| Fibroblasts | MRC2 | 1.29787E-92 | 0.759969 | 0.37 | 0.089 | 2.13162E-88 |
| Pericytes | GUCY1B1 | 2.22269E-37 | 0.759151 | 0.365 | 0.147 | 3.65054E-33 |
| Fibroblasts | PLBD1 | 3.7399E-118 | 0.757997 | 0.302 | 0.033 | 6.1425E-114 |
| Fibroblasts | B4GALT1 | 1.88462E-62 | 0.75767 | 0.387 | 0.138 | 3.0953E-58 |
| Fibroblasts | SEMA3B | 2.12645E-89 | 0.757084 | 0.319 | 0.063 | 3.49248E-85 |
| Contractile | PKIG | 8.65245E-47 | 0.756549 | 0.749 | 0.467 | 1.42108E-42 |
| RERGL_Cells | ARPC2 | 6.18347E-45 | 0.755931 | 0.876 | 0.757 | 1.01557E-40 |
| RERGL_Cells | ITIH3 | 7.05547E-36 | 0.755275 | 0.207 | 0.052 | 1.15879E-31 |
| Intermediate | CCN2 | 7.75931E-46 | 0.754626 | 0.944 | 0.753 | 1.27439E-41 |
| Pericytes | IFITM1 | 8.16374E-31 | 0.754285 | 0.282 | 0.11 | 1.34081E-26 |
| Fibroblasts | ANGPTL5 | 1.5579E-114 | 0.753994 | 0.232 | 0.011 | 2.5586E-110 |
| Pericytes | SERPINH1 | 4.77209E-26 | 0.753844 | 0.605 | 0.443 | 7.83769E-22 |

| | | | | | | |
|---|---|---|---|---|---|---|
| RERGL_Cells | SYPL1 | 7.13449E-30 | 0.75343 | 0.682 | 0.541 | 1.17177E-25 |
| RERGL_Cells | MYLIP | 7.58728E-23 | 0.753178 | 0.417 | 0.237 | 1.24613E-18 |
| Pericytes | SFPQ | 2.85759E-25 | 0.752985 | 0.585 | 0.444 | 4.6933E-21 |
| RERGL_Cells | COX6C | 2.1971E-54 | 0.752225 | 0.915 | 0.838 | 3.60852E-50 |
| Pericytes | MT1M | 3.45191E-15 | 0.752202 | 0.633 | 0.53 | 5.66942E-11 |
| Contractile | SMIM10 | 3.29651E-44 | 0.752121 | 0.433 | 0.169 | 5.41419E-40 |
| Pericytes | DDIT3 | 2.16529E-20 | 0.750908 | 0.4 | 0.242 | 3.55627E-16 |
| RERGL_Cells | EIF4B | 2.76439E-28 | 0.749925 | 0.624 | 0.448 | 4.54024E-24 |
| Contractile | LIMS2 | 7.99144E-42 | 0.748361 | 0.399 | 0.153 | 1.31251E-37 |
| Fibroblasts | PLXDC1 | 2.319E-103 | 0.74794 | 0.233 | 0.017 | 3.8087E-99 |
| Pericytes | ARPC5 | 1.86971E-41 | 0.747739 | 0.795 | 0.646 | 3.07081E-37 |
| Pericytes | MT1F | 5.76707E-43 | 0.747106 | 0.224 | 0.055 | 9.47183E-39 |
| RERGL_Cells | INTS6 | 5.33486E-17 | 0.747065 | 0.385 | 0.241 | 8.76197E-13 |
| RERGL_Cells | NDUFB7 | 3.06837E-42 | 0.747012 | 0.801 | 0.648 | 5.03948E-38 |
| Pericytes | STK38L | 5.46641E-22 | 0.746879 | 0.327 | 0.169 | 8.97803E-18 |
| RERGL_Cells | ADAP2 | 2.54506E-75 | 0.746633 | 0.184 | 0.014 | 4.18E-71 |
| Contractile | MAP4 | 1.1362E-30 | 0.746551 | 0.527 | 0.302 | 1.86609E-26 |
| Pericytes | GNAI1 | 7.31814E-40 | 0.74632 | 0.293 | 0.097 | 1.20193E-35 |
| RERGL_Cells | NDUFA7 | 7.05828E-24 | 0.745753 | 0.44 | 0.268 | 1.15925E-19 |
| Terminal | ITGA10 | 5.94149E-35 | 0.745394 | 0.342 | 0.125 | 9.7583E-31 |
| Pericytes | ANGPTL1 | 7.56274E-20 | 0.745109 | 0.351 | 0.198 | 1.2421E-15 |
| Contractile | AC092164 | 6.0675E-44 | 0.743771 | 0.351 | 0.114 | 9.96526E-40 |
| Pericytes | PHLDB2 | 1.33529E-28 | 0.743633 | 0.335 | 0.153 | 2.19308E-24 |
| Contractile | CAVIN3 | 9.05948E-54 | 0.743212 | 0.9 | 0.715 | 1.48793E-49 |
| Fibroblasts | TNFSF10 | 6.9629E-106 | 0.743205 | 0.291 | 0.037 | 1.1436E-101 |
| Contractile | PPP1R12A | 1.10472E-44 | 0.742804 | 0.747 | 0.484 | 1.81438E-40 |
| RERGL_Cells | ECH1 | 1.62952E-26 | 0.742463 | 0.635 | 0.502 | 2.67633E-22 |
| Intermediate | GYPC | 7.66006E-47 | 0.742122 | 0.693 | 0.373 | 1.25809E-42 |
| Fibroblasts | SRPX2 | 1.2143E-107 | 0.741668 | 0.286 | 0.035 | 1.9944E-103 |
| Fibroblasts | ENG | 1.6676E-67 | 0.740382 | 0.754 | 0.442 | 2.73887E-63 |
| Intermediate | HTRA1 | 3.26668E-34 | 0.740378 | 0.725 | 0.473 | 5.3652E-30 |
| Fibroblasts | SKIL | 7.97677E-67 | 0.738821 | 0.506 | 0.208 | 1.3101E-62 |
| Pericytes | INTS6 | 2.24856E-21 | 0.738074 | 0.402 | 0.234 | 3.69303E-17 |
| Pericytes | FOSB | 6.099E-15 | 0.737441 | 0.375 | 0.23 | 1.0017E-10 |
| Intermediate | SFRP4 | 6.95146E-63 | 0.736801 | 0.559 | 0.212 | 1.14171E-58 |
| RERGL_Cells | MAPRE2 | 1.59678E-29 | 0.736702 | 0.254 | 0.09 | 2.62255E-25 |
| Terminal | MGP | 1.09647E-33 | 0.736508 | 0.996 | 0.952 | 1.80085E-29 |
| Fibroblasts | TPBG | 2.53521E-83 | 0.736456 | 0.362 | 0.093 | 4.16383E-79 |
| Terminal | NUCB2 | 7.1835E-24 | 0.736208 | 0.614 | 0.431 | 1.17982E-19 |
| Fibroblasts | TPST1 | 2.3129E-102 | 0.735646 | 0.353 | 0.069 | 3.79872E-98 |
| Pericytes | PLPPR4 | 8.3035E-111 | 0.7352 | 0.189 | 0.005 | 1.3638E-106 |
| Fibroblasts | SPOCK1 | 1.1794E-104 | 0.734981 | 0.277 | 0.033 | 1.9371E-100 |
| Contractile | EIF3L | 2.04065E-33 | 0.734782 | 0.764 | 0.573 | 3.35157E-29 |
| Pericytes | SDC4 | 1.46663E-14 | 0.734409 | 0.298 | 0.176 | 2.40879E-10 |
| RERGL_Cells | ATP5MC2 | 3.54594E-45 | 0.734261 | 0.87 | 0.742 | 5.82386E-41 |
| Pericytes | ZFP36L1 | 9.63534E-22 | 0.734096 | 0.72 | 0.588 | 1.58251E-17 |
| Fibroblasts | ACVRL1 | 5.0935E-103 | 0.733963 | 0.335 | 0.056 | 8.3656E-99 |
| Contractile | PCDH7 | 1.80461E-50 | 0.733771 | 0.269 | 0.063 | 2.9639E-46 |
| Intermediate | GADD45A | 1.45213E-49 | 0.733665 | 0.576 | 0.248 | 2.38497E-45 |
| Fibroblasts | METTL7A | 3.14618E-71 | 0.733541 | 0.599 | 0.269 | 5.16728E-67 |
| Pericytes | SEPTIN7 | 4.5096E-59 | 0.732385 | 0.958 | 0.905 | 7.40656E-55 |
| Fibroblasts | AKAP12 | 4.1473E-67 | 0.732369 | 0.605 | 0.292 | 6.81153E-63 |
| Fibroblasts | NR4A2 | 1.73716E-36 | 0.731914 | 0.264 | 0.1 | 2.85311E-32 |
| Intermediate | HK1 | 1.75765E-45 | 0.730531 | 0.583 | 0.289 | 2.88676E-41 |
| Terminal | MMP23B | 5.09827E-18 | 0.730464 | 0.364 | 0.204 | 8.3734E-14 |
| RERGL_Cells | ITIH5 | 3.62302E-32 | 0.729713 | 0.269 | 0.093 | 5.95045E-28 |

| | | | | | | |
|---|---|---|---|---|---|---|
| Contractile | CRNDE | 1.75139E-44 | 0.729622 | 0.399 | 0.145 | 2.87648E-40 |
| Contractile | TMEM159 | 7.07784E-30 | 0.729451 | 0.413 | 0.2 | 1.16246E-25 |
| Pericytes | CHRD | 4.24082E-34 | 0.72923 | 0.309 | 0.12 | 6.96513E-30 |
| Contractile | ARL6IP4 | 1.18456E-46 | 0.727851 | 0.852 | 0.679 | 1.94553E-42 |
| Fibroblasts | LINC02802 | 3.08756E-93 | 0.727471 | 0.35 | 0.073 | 5.07102E-89 |
| RERGL_Cells | JADE1 | 4.18213E-15 | 0.72743 | 0.363 | 0.234 | 6.86873E-11 |
| Intermediate | FBLIM1 | 6.1083E-50 | 0.726847 | 0.721 | 0.359 | 1.00323E-45 |
| RERGL_Cells | B3GNT2 | 3.70596E-27 | 0.726763 | 0.265 | 0.1 | 6.08667E-23 |
| Fibroblasts | TNFSF14 | 2.3452E-108 | 0.725829 | 0.195 | 0.003 | 3.8518E-104 |
| RERGL_Cells | TIMP4 | 2.87755E-24 | 0.725657 | 0.209 | 0.072 | 4.7261E-20 |
| Terminal | CRISPLD1 | 6.45396E-25 | 0.725629 | 0.39 | 0.191 | 1.06E-20 |
| RERGL_Cells | COX7A1 | 1.39472E-34 | 0.725315 | 0.784 | 0.639 | 2.29069E-30 |
| Contractile | COX7C | 6.25929E-69 | 0.725167 | 0.968 | 0.899 | 1.02803E-64 |
| Pericytes | JUN | 7.49899E-20 | 0.725051 | 0.784 | 0.667 | 1.23163E-15 |
| Intermediate | LINC01615 | 6.7967E-59 | 0.724651 | 0.381 | 0.107 | 1.11629E-54 |
| Fibroblasts | CPVL | 5.5093E-124 | 0.724129 | 0.268 | 0.017 | 9.0484E-120 |
| Pericytes | FOS | 1.08938E-26 | 0.723652 | 0.804 | 0.596 | 1.7892E-22 |
| Intermediate | ITGA1 | 2.77977E-43 | 0.723445 | 0.469 | 0.188 | 4.56549E-39 |
| Contractile | CALM3 | 1.56754E-41 | 0.721958 | 0.774 | 0.529 | 2.57453E-37 |
| Contractile | RBPMS | 3.22259E-41 | 0.721606 | 0.663 | 0.367 | 5.29278E-37 |
| Contractile | GADD45A | 4.9204E-32 | 0.721463 | 0.509 | 0.263 | 8.08126E-28 |
| RERGL_Cells | SEPTIN7 | 1.01811E-56 | 0.72107 | 0.959 | 0.906 | 1.67214E-52 |
| Fibroblasts | SCPEP1 | 3.04972E-74 | 0.720907 | 0.769 | 0.452 | 5.00886E-70 |
| RERGL_Cells | UQCRB | 1.9397E-56 | 0.720902 | 0.936 | 0.852 | 3.18576E-52 |
| Fibroblasts | CCDC71L | 1.34025E-75 | 0.720576 | 0.269 | 0.053 | 2.20122E-71 |
| Pericytes | RGMA | 2.54055E-25 | 0.719115 | 0.349 | 0.175 | 4.17261E-21 |
| Pericytes | MAP1B | 2.27673E-33 | 0.717682 | 0.816 | 0.655 | 3.7393E-29 |
| Pericytes | CRISPLD2 | 1.06844E-25 | 0.716949 | 0.469 | 0.265 | 1.7548E-21 |
| RERGL_Cells | PLEKHO1 | 4.00464E-20 | 0.71691 | 0.308 | 0.153 | 6.57722E-16 |
| Contractile | NINJ1 | 2.93171E-33 | 0.716062 | 0.497 | 0.258 | 4.81504E-29 |
| Intermediate | TGFB1 | 8.42782E-63 | 0.715621 | 0.363 | 0.093 | 1.38418E-58 |
| Terminal | COL8A1 | 3.09705E-34 | 0.715539 | 0.545 | 0.26 | 5.08659E-30 |
| Fibroblasts | CFB | 9.57799E-66 | 0.715151 | 0.438 | 0.162 | 1.57309E-61 |
| Fibroblasts | COL4A3 | 4.1726E-103 | 0.715052 | 0.232 | 0.017 | 6.8531E-99 |
| Intermediate | RHOD | 1.56025E-61 | 0.714524 | 0.378 | 0.104 | 2.56255E-57 |
| Fibroblasts | PDIA4 | 3.77763E-68 | 0.714416 | 0.588 | 0.272 | 6.20439E-64 |
| Contractile | MXI1 | 2.35936E-29 | 0.714384 | 0.377 | 0.176 | 3.87502E-25 |
| Fibroblasts | TNFAIP2 | 4.84247E-85 | 0.714068 | 0.219 | 0.023 | 7.95328E-81 |
| RERGL_Cells | JAG1 | 4.51989E-22 | 0.713431 | 0.355 | 0.181 | 7.42347E-18 |
| Fibroblasts | RRBP1 | 2.03826E-69 | 0.713387 | 0.564 | 0.251 | 3.34763E-65 |
| Fibroblasts | INMT | 2.85398E-47 | 0.713147 | 0.612 | 0.346 | 4.68737E-43 |
| Pericytes | PICALM | 6.64258E-26 | 0.712861 | 0.473 | 0.294 | 1.09098E-21 |
| Pericytes | BST2 | 6.11422E-30 | 0.70989 | 0.425 | 0.21 | 1.0042E-25 |
| Pericytes | NOP16 | 5.78713E-21 | 0.709636 | 0.298 | 0.151 | 9.50479E-17 |
| Intermediate | MICAL2 | 4.46984E-75 | 0.709562 | 0.331 | 0.066 | 7.34127E-71 |
| Contractile | TTLL7 | 6.61255E-36 | 0.709348 | 0.461 | 0.209 | 1.08605E-31 |
| Pericytes | MAFF | 1.37403E-21 | 0.708905 | 0.376 | 0.217 | 2.25671E-17 |
| RERGL_Cells | EPS8 | 5.89459E-30 | 0.708239 | 0.69 | 0.511 | 9.68128E-26 |
| Pericytes | C1orf198 | 5.2779E-22 | 0.708093 | 0.442 | 0.284 | 8.66842E-18 |
| Fibroblasts | IFI16 | 1.76171E-74 | 0.707109 | 0.703 | 0.363 | 2.89343E-70 |
| RERGL_Cells | PCDH1 | 2.36299E-42 | 0.70685 | 0.194 | 0.038 | 3.88098E-38 |
| Terminal | F2R | 5.79435E-24 | 0.706643 | 0.259 | 0.102 | 9.51664E-20 |
| Pericytes | MTHFD2 | 2.09175E-29 | 0.706558 | 0.624 | 0.416 | 3.43549E-25 |
| Terminal | COL1A1 | 9.26031E-11 | 0.706516 | 0.582 | 0.462 | 1.52091E-06 |
| Fibroblasts | TMEM98 | 6.3888E-83 | 0.70631 | 0.576 | 0.228 | 1.0493E-78 |
| Contractile | HCFC1R1 | 5.52943E-41 | 0.70624 | 0.778 | 0.502 | 9.08153E-37 |

| | | | | | | |
|---|---|---|---|---|---|---|
| Pericytes | SH3KBP1 | 3.08968E-26 | 0.704542 | 0.356 | 0.18 | 5.07449E-22 |
| RERGL_Cells | PPP1R12B | 4.39123E-28 | 0.704349 | 0.511 | 0.286 | 7.21215E-24 |
| Contractile | CALM1 | 7.36773E-46 | 0.704211 | 0.91 | 0.772 | 1.21008E-41 |
| Pericytes | TP53I11 | 1.22098E-46 | 0.703938 | 0.244 | 0.06 | 2.00534E-42 |
| RERGL_Cells | RASGRP2 | 1.48667E-26 | 0.70325 | 0.31 | 0.133 | 2.4417E-22 |
| Pericytes | OLFML2B | 3.05351E-13 | 0.702989 | 0.324 | 0.211 | 5.01509E-09 |
| RERGL_Cells | COX7B | 5.05468E-26 | 0.702813 | 0.746 | 0.65 | 8.30181E-22 |
| Intermediate | CYB5R3 | 4.47584E-54 | 0.702773 | 0.925 | 0.762 | 7.35112E-50 |
| Contractile | MYOZ1 | 4.16764E-42 | 0.70272 | 0.393 | 0.142 | 6.84493E-38 |
| Fibroblasts | XBP1 | 4.2059E-69 | 0.70244 | 0.796 | 0.505 | 6.90777E-65 |
| RERGL_Cells | H1F0 | 3.65582E-15 | 0.702419 | 0.338 | 0.205 | 6.00432E-11 |
| Fibroblasts | GFRA1 | 9.1216E-112 | 0.702271 | 0.251 | 0.018 | 1.4981E-107 |
| Pericytes | TCF21 | 3.47499E-18 | 0.70217 | 0.469 | 0.32 | 5.70733E-14 |
| RERGL_Cells | ARID5A | 5.57132E-38 | 0.701719 | 0.65 | 0.38 | 9.15034E-34 |
| Terminal | GPM6B | 1.56722E-26 | 0.701697 | 0.242 | 0.083 | 2.57401E-22 |
| RERGL_Cells | ANXA6 | 1.76066E-28 | 0.701243 | 0.692 | 0.563 | 2.8917E-24 |
| Intermediate | PLS3 | 3.22289E-51 | 0.701232 | 0.92 | 0.703 | 5.29328E-47 |
| RERGL_Cells | TGFBI | 5.08746E-35 | 0.701105 | 0.496 | 0.231 | 8.35565E-31 |
| Pericytes | GADD45B | 2.32607E-13 | 0.70063 | 0.689 | 0.671 | 3.82034E-09 |
| Contractile | PFN2 | 2.3815E-32 | 0.700081 | 0.629 | 0.388 | 3.91138E-28 |
| Pericytes | DDX3X | 6.30895E-27 | 0.700005 | 0.547 | 0.363 | 1.03618E-22 |
| Fibroblasts | FGFR1 | 9.95372E-78 | 0.699163 | 0.637 | 0.295 | 1.6348E-73 |
| RERGL_Cells | CCND1 | 5.50867E-18 | 0.698745 | 0.348 | 0.2 | 9.04744E-14 |
| Fibroblasts | PLAC9 | 5.48899E-73 | 0.69844 | 0.969 | 0.793 | 9.01512E-69 |
| RERGL_Cells | CAVIN1 | 1.64578E-47 | 0.698401 | 0.912 | 0.829 | 2.70303E-43 |
| Intermediate | CTSZ | 9.41039E-42 | 0.69824 | 0.441 | 0.18 | 1.54556E-37 |
| RERGL_Cells | ZFP36L1 | 2.63409E-22 | 0.69817 | 0.729 | 0.59 | 4.32623E-18 |
| Pericytes | PELO | 1.35013E-18 | 0.697626 | 0.315 | 0.174 | 2.21746E-14 |
| Pericytes | CLSTN2 | 4.37822E-69 | 0.697531 | 0.209 | 0.027 | 7.19078E-65 |
| RERGL_Cells | PABPC1 | 4.27272E-32 | 0.696987 | 0.825 | 0.726 | 7.01752E-28 |
| Pericytes | TUBA1A | 1.20819E-27 | 0.696852 | 0.824 | 0.758 | 1.98434E-23 |
| Contractile | MAP1B | 7.11058E-37 | 0.696432 | 0.84 | 0.654 | 1.16784E-32 |
| Fibroblasts | TGFBR3 | 1.50035E-87 | 0.696066 | 0.269 | 0.041 | 2.46417E-83 |
| Fibroblasts | SERTAD1 | 8.99013E-69 | 0.695515 | 0.531 | 0.227 | 1.47654E-64 |
| RERGL_Cells | COX6A1 | 5.87841E-47 | 0.695011 | 0.912 | 0.822 | 9.6547E-43 |
| Pericytes | RBPMS | 3.24354E-27 | 0.694871 | 0.573 | 0.378 | 5.32719E-23 |
| RERGL_Cells | SMOC2 | 1.73334E-32 | 0.694603 | 0.752 | 0.549 | 2.84684E-28 |
| Pericytes | CAV2 | 1.0724E-35 | 0.694114 | 0.74 | 0.514 | 1.76132E-31 |
| Terminal | CFH | 4.53253E-11 | 0.694057 | 0.725 | 0.629 | 7.44422E-07 |
| Terminal | EMP3 | 2.18111E-19 | 0.694028 | 0.693 | 0.559 | 3.58225E-15 |
| Pericytes | FAM13C | 1.00427E-18 | 0.69371 | 0.218 | 0.096 | 1.64942E-14 |
| Intermediate | ECM2 | 6.95868E-44 | 0.693419 | 0.579 | 0.274 | 1.14289E-39 |
| Fibroblasts | TRAM1 | 2.3703E-67 | 0.693369 | 0.726 | 0.42 | 3.89298E-63 |
| Contractile | ADI1 | 1.28133E-36 | 0.693128 | 0.806 | 0.582 | 2.10445E-32 |
| RERGL_Cells | GEM | 1.16511E-11 | 0.692941 | 0.562 | 0.479 | 1.91358E-07 |
| RERGL_Cells | SOD3 | 8.51108E-54 | 0.691927 | 0.968 | 0.883 | 1.39786E-49 |
| Terminal | POSTN | 3.85905E-15 | 0.690151 | 0.179 | 0.072 | 6.3381E-11 |
| Pericytes | SNHG15 | 5.09302E-28 | 0.688773 | 0.309 | 0.136 | 8.36478E-24 |
| RERGL_Cells | ARHGAP2 | 1.60791E-16 | 0.688392 | 0.37 | 0.229 | 2.64084E-12 |
| RERGL_Cells | CALM2 | 1.96196E-81 | 0.687446 | 1 | 0.983 | 3.22233E-77 |
| Pericytes | PRRX1 | 1.31881E-22 | 0.687046 | 0.365 | 0.203 | 2.16602E-18 |
| Terminal | OGN | 8.55326E-28 | 0.687001 | 0.745 | 0.491 | 1.40479E-23 |
| RERGL_Cells | C20orf27 | 6.60331E-17 | 0.68693 | 0.335 | 0.191 | 1.08453E-12 |
| Intermediate | ENO1 | 8.29743E-47 | 0.686298 | 0.916 | 0.753 | 1.36277E-42 |
| Fibroblasts | RHOBTB3 | 1.52008E-76 | 0.68611 | 0.366 | 0.101 | 2.49658E-72 |
| Pericytes | CCNL1 | 6.147E-24 | 0.685882 | 0.602 | 0.422 | 1.00958E-19 |

| | | | | | | |
|---|---|---|---|---|---|---|
| RERGL_Cells | YBX3 | 1.9166E-25 | 0.685776 | 0.756 | 0.648 | 3.14783E-21 |
| Intermediate | KCNK15 | 1.4843E-24 | 0.685549 | 0.305 | 0.136 | 2.43782E-20 |
| Contractile | NPM1 | 3.94582E-48 | 0.685074 | 0.952 | 0.867 | 6.48061E-44 |
| Contractile | STMN1 | 1.4973E-34 | 0.684884 | 0.323 | 0.117 | 2.45916E-30 |
| Pericytes | HMOX2 | 3.28748E-25 | 0.684692 | 0.367 | 0.194 | 5.39936E-21 |
| RERGL_Cells | NT5DC2 | 2.7247E-26 | 0.684522 | 0.239 | 0.086 | 4.47505E-22 |
| RERGL_Cells | LDB3 | 6.20472E-28 | 0.684242 | 0.265 | 0.095 | 1.01906E-23 |
| Intermediate | CDH2 | 1.00299E-70 | 0.683913 | 0.34 | 0.073 | 1.64731E-66 |
| Intermediate | UACA | 4.2071E-38 | 0.683398 | 0.471 | 0.21 | 6.90973E-34 |
| Pericytes | KRT18 | 2.19818E-25 | 0.683246 | 0.24 | 0.089 | 3.61029E-21 |
| Fibroblasts | PHGDH | 4.55705E-76 | 0.682863 | 0.471 | 0.165 | 7.48449E-72 |
| Intermediate | PLXDC2 | 2.70051E-50 | 0.682672 | 0.462 | 0.169 | 4.43533E-46 |
| Contractile | MFGE8 | 1.05336E-64 | 0.68146 | 0.986 | 0.843 | 1.73004E-60 |
| Pericytes | FADS3 | 1.18404E-29 | 0.681252 | 0.451 | 0.246 | 1.94466E-25 |
| RERGL_Cells | HSPB8 | 3.68692E-15 | 0.680984 | 0.361 | 0.224 | 6.05539E-11 |
| Contractile | COL18A1 | 3.17386E-30 | 0.680741 | 0.717 | 0.477 | 5.21275E-26 |
| Fibroblasts | BHLHE40 | 1.50655E-45 | 0.680659 | 0.41 | 0.178 | 2.47436E-41 |
| Fibroblasts | CCL13 | 2.09277E-54 | 0.678768 | 0.138 | 0.014 | 3.43717E-50 |
| Pericytes | INSIG2 | 3.23033E-26 | 0.67847 | 0.367 | 0.188 | 5.3055E-22 |
| Terminal | LUM | 5.27556E-10 | 0.678135 | 0.619 | 0.475 | 8.66458E-06 |
| RERGL_Cells | MSRB3 | 8.38333E-23 | 0.677977 | 0.534 | 0.361 | 1.37688E-18 |
| Pericytes | ENPEP | 7.53197E-28 | 0.677476 | 0.26 | 0.1 | 1.23705E-23 |
| Intermediate | SCG2 | 6.2103E-74 | 0.676991 | 0.219 | 0.025 | 1.01998E-69 |
| Fibroblasts | GPRC5A | 2.57039E-70 | 0.676905 | 0.243 | 0.045 | 4.22161E-66 |
| Intermediate | TMSB4X | 7.76984E-77 | 0.676648 | 1 | 0.997 | 1.27612E-72 |
| RERGL_Cells | HOXB-AS1 | 6.54981E-23 | 0.676047 | 0.271 | 0.117 | 1.07574E-18 |
| Terminal | FN1 | 1.2146E-46 | 0.675845 | 0.821 | 0.57 | 1.99486E-42 |
| Pericytes | IGFBP4 | 1.25828E-69 | 0.675783 | 0.924 | 0.525 | 2.06661E-65 |
| RERGL_Cells | MEOX2 | 3.98959E-21 | 0.674959 | 0.306 | 0.145 | 6.5525E-17 |
| Terminal | COL1A2 | 1.6719E-09 | 0.674125 | 0.739 | 0.701 | 2.74593E-05 |
| Intermediate | ECRG4 | 1.87682E-45 | 0.674061 | 0.862 | 0.57 | 3.0825E-41 |
| Contractile | ITIH4 | 4.79436E-39 | 0.67326 | 0.21 | 0.049 | 7.87425E-35 |
| RERGL_Cells | UQCR11 | 6.7587E-46 | 0.673204 | 0.906 | 0.802 | 1.11005E-41 |
| RERGL_Cells | CYSTM1 | 6.7905E-17 | 0.673081 | 0.603 | 0.518 | 1.11527E-12 |
| Fibroblasts | KLF9 | 2.40309E-61 | 0.672947 | 0.463 | 0.186 | 3.94684E-57 |
| Terminal | ELN | 9.68835E-22 | 0.672384 | 0.601 | 0.378 | 1.59121E-17 |
| Fibroblasts | LGALS3BP | 8.63519E-58 | 0.672283 | 0.806 | 0.546 | 1.41824E-53 |
| Fibroblasts | MGAT1 | 3.89186E-66 | 0.672223 | 0.597 | 0.289 | 6.39198E-62 |
| Terminal | TSPO | 8.43997E-35 | 0.671951 | 0.817 | 0.604 | 1.38618E-30 |
| RERGL_Cells | CRTC3 | 3.83081E-20 | 0.671896 | 0.269 | 0.126 | 6.29172E-16 |
| Contractile | MRPL51 | 1.76308E-31 | 0.671576 | 0.687 | 0.458 | 2.89569E-27 |
| Fibroblasts | VSIR | 1.76462E-62 | 0.671524 | 0.553 | 0.255 | 2.89821E-58 |
| Contractile | PPP1CB | 3.44172E-26 | 0.671463 | 0.814 | 0.635 | 5.65268E-22 |
| Intermediate | MRAP2 | 1.19341E-61 | 0.670518 | 0.342 | 0.082 | 1.96006E-57 |
| Fibroblasts | LAMP2 | 3.11266E-66 | 0.670384 | 0.555 | 0.25 | 5.11223E-62 |
| Contractile | FHL1 | 2.09792E-36 | 0.670372 | 0.93 | 0.816 | 3.44562E-32 |
| Pericytes | ANKRD10 | 2.06977E-19 | 0.669942 | 0.413 | 0.26 | 3.39938E-15 |
| Fibroblasts | PSAP | 1.8924E-109 | 0.669928 | 0.982 | 0.886 | 3.1081E-105 |
| Pericytes | ITGA1 | 3.29316E-23 | 0.669526 | 0.375 | 0.203 | 5.40869E-19 |
| Pericytes | AC092807 | 2.71487E-31 | 0.669493 | 0.207 | 0.062 | 4.45891E-27 |
| Pericytes | ARHGDIB | 3.91517E-21 | 0.669454 | 0.625 | 0.489 | 6.43027E-17 |
| RERGL_Cells | FABP5 | 1.77E-60 | 0.669255 | 0.485 | 0.151 | 2.90705E-56 |
| RERGL_Cells | SBSPON | 5.45407E-30 | 0.669162 | 0.389 | 0.17 | 8.95777E-26 |
| Pericytes | PPARG | 3.70603E-45 | 0.668565 | 0.196 | 0.039 | 6.08678E-41 |
| RERGL_Cells | NDUFB10 | 1.25976E-24 | 0.667752 | 0.699 | 0.579 | 2.06903E-20 |
| Contractile | HEY2 | 7.71201E-30 | 0.667536 | 0.303 | 0.116 | 1.26662E-25 |

| | | | | | | |
|---|---|---|---|---|---|---|
| Fibroblasts | CDC42EP4 | 2.50524E-61 | 0.667025 | 0.456 | 0.182 | 4.1146E-57 |
| Pericytes | FST | 2.95054E-10 | 0.666843 | 0.189 | 0.102 | 4.84597E-06 |
| RERGL_Cells | NIBAN1 | 9.55759E-20 | 0.666609 | 0.406 | 0.236 | 1.56974E-15 |
| Pericytes | C1orf56 | 3.38898E-12 | 0.666525 | 0.389 | 0.273 | 5.56606E-08 |
| Contractile | CCT8 | 2.77679E-27 | 0.666037 | 0.677 | 0.49 | 4.5606E-23 |
| Fibroblasts | TIMP1 | 2.8413E-134 | 0.664522 | 0.995 | 0.899 | 4.6665E-130 |
| Intermediate | FOXS1 | 8.63131E-51 | 0.664389 | 0.411 | 0.135 | 1.41761E-46 |
| Pericytes | IMMP2L | 4.09499E-24 | 0.664216 | 0.325 | 0.162 | 6.72561E-20 |
| RERGL_Cells | CBX6 | 6.66166E-17 | 0.663991 | 0.585 | 0.463 | 1.09411E-12 |
| RERGL_Cells | SYNPO2 | 9.61016E-28 | 0.663563 | 0.545 | 0.316 | 1.57837E-23 |
| RERGL_Cells | TUBA1C | 2.55075E-09 | 0.66316 | 0.378 | 0.287 | 4.18936E-05 |
| RERGL_Cells | ELOVL5 | 7.94633E-17 | 0.663094 | 0.382 | 0.239 | 1.3051E-12 |
| Contractile | SUSD5 | 9.81816E-39 | 0.66175 | 0.319 | 0.104 | 1.61253E-34 |
| Pericytes | ITM2C | 2.14428E-25 | 0.660308 | 0.309 | 0.141 | 3.52176E-21 |
| Fibroblasts | JUN | 1.97077E-64 | 0.659464 | 0.857 | 0.61 | 3.2368E-60 |
| Contractile | HINT1 | 3.31592E-49 | 0.658478 | 0.902 | 0.769 | 5.44607E-45 |
| Fibroblasts | ROBO3 | 5.3395E-99 | 0.658432 | 0.242 | 0.023 | 8.76957E-95 |
| Pericytes | ADRA2A | 7.53048E-64 | 0.657978 | 0.182 | 0.02 | 1.23681E-59 |
| Pericytes | CDC37L1 | 2.47716E-16 | 0.657889 | 0.325 | 0.193 | 4.06848E-12 |
| Pericytes | IFITM2 | 1.52231E-52 | 0.657568 | 0.973 | 0.894 | 2.50024E-48 |
| Contractile | ZFYVE21 | 1.20908E-29 | 0.657432 | 0.641 | 0.427 | 1.98579E-25 |
| Terminal | TNFAIP6 | 6.61559E-13 | 0.657069 | 0.255 | 0.135 | 1.08655E-08 |
| Intermediate | S100A16 | 9.38799E-38 | 0.656985 | 0.566 | 0.286 | 1.54188E-33 |
| RERGL_Cells | PLAC9 | 4.20165E-48 | 0.656951 | 0.981 | 0.825 | 6.90079E-44 |
| Terminal | OMD | 2.40646E-18 | 0.65667 | 0.466 | 0.284 | 3.95237E-14 |
| Fibroblasts | MPZL1 | 4.99226E-69 | 0.65643 | 0.457 | 0.17 | 8.19929E-65 |
| Pericytes | PIM1 | 8.60642E-16 | 0.655973 | 0.367 | 0.23 | 1.41352E-11 |
| RERGL_Cells | ANAPC16 | 8.00719E-31 | 0.655959 | 0.791 | 0.659 | 1.3151E-26 |
| Pericytes | B4GALT3 | 2.01807E-25 | 0.655661 | 0.285 | 0.126 | 3.31447E-21 |
| Contractile | SOD1 | 2.51291E-53 | 0.65509 | 0.936 | 0.845 | 4.1272E-49 |
| Contractile | NPTX2 | 1.31998E-63 | 0.65498 | 0.188 | 0.021 | 2.16793E-59 |
| RERGL_Cells | VAMP2 | 3.69794E-35 | 0.654825 | 0.825 | 0.704 | 6.0735E-31 |
| Contractile | TSR2 | 2.44615E-31 | 0.653977 | 0.493 | 0.26 | 4.01755E-27 |
| Fibroblasts | IFITM3 | 4.7698E-114 | 0.653613 | 0.999 | 0.988 | 7.8339E-110 |
| Contractile | HSPB8 | 3.6481E-29 | 0.653396 | 0.425 | 0.212 | 5.99165E-25 |
| RERGL_Cells | COX8A | 9.84154E-29 | 0.653343 | 0.767 | 0.667 | 1.61637E-24 |
| Fibroblasts | LAPTM4A | 2.4229E-135 | 0.652783 | 0.994 | 0.942 | 3.9794E-131 |
| Intermediate | PAM | 1.41813E-32 | 0.652562 | 0.615 | 0.349 | 2.32914E-28 |
| Pericytes | SRSF3 | 2.20172E-36 | 0.651707 | 0.867 | 0.756 | 3.6161E-32 |
| Contractile | TCEAL1 | 5.32876E-19 | 0.65136 | 0.481 | 0.312 | 8.75196E-15 |
| Intermediate | FKBP10 | 2.41493E-29 | 0.651117 | 0.593 | 0.36 | 3.96628E-25 |
| Intermediate | ANXA1 | 7.574E-56 | 0.650894 | 0.964 | 0.825 | 1.24395E-51 |
| Fibroblasts | EPB41L2 | 3.78659E-66 | 0.650664 | 0.541 | 0.229 | 6.21909E-62 |
| Pericytes | RERG | 3.34114E-25 | 0.650436 | 0.347 | 0.167 | 5.48749E-21 |
| Contractile | BEX3 | 3.58054E-33 | 0.650089 | 0.814 | 0.653 | 5.88067E-29 |
| Intermediate | HCFC1R1 | 4.70374E-36 | 0.650014 | 0.787 | 0.497 | 7.72543E-32 |
| RERGL_Cells | ARHGEF17 | 2.28905E-25 | 0.650012 | 0.226 | 0.079 | 3.75954E-21 |
| RERGL_Cells | NTF3 | 4.76623E-22 | 0.649419 | 0.237 | 0.095 | 7.82805E-18 |
| Fibroblasts | AC020916 | 1.87091E-38 | 0.648948 | 0.4 | 0.188 | 3.07278E-34 |
| Pericytes | TNFRSF1A | 2.68781E-22 | 0.648368 | 0.718 | 0.626 | 4.41446E-18 |
| Pericytes | TMEM173 | 1.0821E-21 | 0.647995 | 0.504 | 0.335 | 1.77724E-17 |
| Terminal | PHPT1 | 1.67509E-18 | 0.647924 | 0.71 | 0.581 | 2.75117E-14 |
| Pericytes | CBX6 | 1.72704E-15 | 0.647858 | 0.564 | 0.464 | 2.83649E-11 |
| RERGL_Cells | HACD1 | 6.97686E-20 | 0.647699 | 0.329 | 0.167 | 1.14588E-15 |
| Pericytes | ISYNA1 | 9.93241E-38 | 0.647538 | 0.616 | 0.34 | 1.6313E-33 |
| Fibroblasts | RCN3 | 3.41594E-61 | 0.647313 | 0.635 | 0.318 | 5.61034E-57 |

| | | | | | | |
|---|---|---|---|---|---|---|
| Intermediate | OCIAD2 | 6.7426E-67 | 0.646996 | 0.323 | 0.069 | 1.1074E-62 |
| Contractile | HADH | 7.59282E-29 | 0.646799 | 0.387 | 0.18 | 1.24704E-24 |
| Fibroblasts | C1orf21 | 3.134E-84 | 0.646738 | 0.349 | 0.082 | 5.14728E-80 |
| Terminal | CTHRC1 | 4.47294E-15 | 0.646652 | 0.342 | 0.192 | 7.34635E-11 |
| Fibroblasts | EGR2 | 1.09591E-71 | 0.646091 | 0.208 | 0.028 | 1.79992E-67 |
| Fibroblasts | TRIP10 | 4.58445E-60 | 0.645445 | 0.481 | 0.204 | 7.52951E-56 |
| Fibroblasts | DIO3OS | 7.7542E-113 | 0.644506 | 0.206 | 0.004 | 1.2735E-108 |
| Contractile | ELN | 5.49552E-20 | 0.644358 | 0.565 | 0.381 | 9.02584E-16 |
| RERGL_Cells | PARM1 | 1.25117E-25 | 0.644348 | 0.348 | 0.158 | 2.05492E-21 |
| Contractile | BCAM | 4.22659E-60 | 0.643687 | 0.79 | 0.354 | 6.94175E-56 |
| Fibroblasts | MAN1A1 | 4.62362E-74 | 0.642503 | 0.246 | 0.043 | 7.59383E-70 |
| Fibroblasts | MID1IP1 | 2.80986E-43 | 0.642285 | 0.379 | 0.165 | 4.61492E-39 |
| Fibroblasts | MEDAG | 1.96636E-46 | 0.641944 | 0.37 | 0.152 | 3.22954E-42 |
| Pericytes | FERMT2 | 4.79422E-16 | 0.641566 | 0.487 | 0.368 | 7.87402E-12 |
| Intermediate | EMC10 | 1.60761E-37 | 0.641451 | 0.596 | 0.311 | 2.64034E-33 |
| Intermediate | SNRPN | 1.13504E-35 | 0.641384 | 0.731 | 0.472 | 1.86419E-31 |
| Contractile | PYGB | 5.6699E-31 | 0.640771 | 0.351 | 0.148 | 9.31224E-27 |
| Pericytes | NECTIN2 | 9.28549E-22 | 0.640401 | 0.387 | 0.221 | 1.52505E-17 |
| Pericytes | NDRG2 | 4.83131E-24 | 0.640149 | 0.616 | 0.443 | 7.93495E-20 |
| Contractile | MBNL1-AS | 1.90932E-28 | 0.639045 | 0.303 | 0.119 | 3.13587E-24 |
| Contractile | FXYD1 | 4.41188E-41 | 0.638318 | 0.928 | 0.778 | 7.24608E-37 |
| Terminal | FRZB | 1.93553E-17 | 0.637714 | 0.778 | 0.598 | 3.17892E-13 |
| Pericytes | HLA-C | 3.47204E-52 | 0.637668 | 0.949 | 0.855 | 5.70247E-48 |
| RERGL_Cells | CAP2 | 8.98869E-25 | 0.637626 | 0.316 | 0.142 | 1.4763E-20 |
| Contractile | ZBTB16 | 1.19721E-28 | 0.637118 | 0.651 | 0.418 | 1.9663E-24 |
| RERGL_Cells | ZNF703 | 5.29709E-16 | 0.636687 | 0.361 | 0.218 | 8.69993E-12 |
| Contractile | YWHAH | 2.43735E-24 | 0.636377 | 0.721 | 0.558 | 4.00311E-20 |
| Contractile | DBNDD2 | 4.07333E-35 | 0.635589 | 0.79 | 0.555 | 6.69003E-31 |
| Fibroblasts | BAMBI | 7.84891E-54 | 0.634945 | 0.196 | 0.039 | 1.2891E-49 |
| Intermediate | PKM | 1.83686E-34 | 0.634059 | 0.867 | 0.726 | 3.01685E-30 |
| Intermediate | FHL1 | 6.60916E-37 | 0.63383 | 0.955 | 0.811 | 1.08549E-32 |
| RERGL_Cells | RERG | 4.49098E-22 | 0.633188 | 0.346 | 0.172 | 7.37599E-18 |
| RERGL_Cells | CABP1 | 7.59508E-45 | 0.63305 | 0.162 | 0.024 | 1.24742E-40 |
| Fibroblasts | RBMS1 | 3.68966E-58 | 0.632647 | 0.606 | 0.311 | 6.05991E-54 |
| RERGL_Cells | SEPTIN9 | 5.28869E-19 | 0.632139 | 0.269 | 0.129 | 8.68615E-15 |
| Terminal | CAPG | 3.81742E-17 | 0.632071 | 0.255 | 0.122 | 6.26973E-13 |
| Contractile | DMPK | 5.35928E-35 | 0.631519 | 0.405 | 0.167 | 8.80208E-31 |
| RERGL_Cells | ZBTB16 | 1.83287E-16 | 0.630328 | 0.571 | 0.433 | 3.01031E-12 |
| RERGL_Cells | COX4I1 | 3.12044E-37 | 0.629957 | 0.925 | 0.859 | 5.12502E-33 |
| Intermediate | PDGFD | 6.57592E-28 | 0.629914 | 0.383 | 0.179 | 1.08003E-23 |
| RERGL_Cells | NDUFB9 | 1.1339E-26 | 0.62987 | 0.754 | 0.649 | 1.86231E-22 |
| Fibroblasts | SOX9 | 7.95406E-79 | 0.629644 | 0.237 | 0.035 | 1.30637E-74 |
| Fibroblasts | FKBP11 | 1.11665E-57 | 0.629362 | 0.373 | 0.133 | 1.83398E-53 |
| Contractile | LINC02269 | 2.10998E-37 | 0.629232 | 0.214 | 0.052 | 3.46544E-33 |
| Pericytes | SNAI1 | 3.62565E-25 | 0.628788 | 0.187 | 0.061 | 5.95476E-21 |
| Pericytes | TMEM47 | 1.31117E-13 | 0.628752 | 0.489 | 0.375 | 2.15346E-09 |
| Contractile | EIF3H | 2.10449E-34 | 0.628685 | 0.792 | 0.592 | 3.45642E-30 |
| Contractile | PCBD1 | 3.57508E-27 | 0.628175 | 0.503 | 0.287 | 5.87171E-23 |
| Pericytes | FOXO3 | 3.36119E-18 | 0.627409 | 0.425 | 0.278 | 5.52042E-14 |
| Pericytes | PELI1 | 5.04655E-32 | 0.627191 | 0.211 | 0.062 | 8.28845E-28 |
| Intermediate | PDLIM3 | 1.11585E-47 | 0.626981 | 0.884 | 0.579 | 1.83267E-43 |
| Contractile | BEX2 | 3.66843E-53 | 0.626269 | 0.194 | 0.029 | 6.02503E-49 |
| RERGL_Cells | LPP | 1.01858E-20 | 0.626173 | 0.573 | 0.407 | 1.67291E-16 |
| Pericytes | GPR176 | 1.71822E-25 | 0.625766 | 0.213 | 0.076 | 2.82201E-21 |
| Contractile | PITPNB | 7.14201E-24 | 0.625601 | 0.451 | 0.253 | 1.173E-19 |
| Contractile | NACA | 3.64802E-56 | 0.624962 | 0.966 | 0.938 | 5.9915E-52 |

| | | | | | | |
|---|---|---|---|---|---|---|
| Contractile | SERTAD4-/ | 7.51571E-31 | 0.624818 | 0.333 | 0.136 | 1.23438E-26 |
| Contractile | NME4 | 6.85184E-25 | 0.624229 | 0.591 | 0.392 | 1.12535E-20 |
| Contractile | OGN | 3.4687E-42 | 0.62399 | 0.792 | 0.48 | 5.697E-38 |
| Pericytes | SOCS1 | 7.41274E-21 | 0.623577 | 0.138 | 0.041 | 1.21747E-16 |
| Contractile | TNNC1 | 1.00679E-22 | 0.623255 | 0.182 | 0.059 | 1.65356E-18 |
| RERGL_Cells | DYNLT1 | 1.43123E-18 | 0.622238 | 0.592 | 0.475 | 2.35065E-14 |
| Contractile | ASAH1 | 1.54267E-26 | 0.622006 | 0.705 | 0.53 | 2.53368E-22 |
| Contractile | PARK7 | 6.77228E-47 | 0.621701 | 0.918 | 0.794 | 1.11228E-42 |
| Terminal | TPD52L1 | 2.26567E-26 | 0.621551 | 0.275 | 0.105 | 3.72113E-22 |
| Pericytes | ZNF703 | 1.73761E-13 | 0.620556 | 0.34 | 0.218 | 2.85384E-09 |
| Terminal | TENT5A | 2.46139E-24 | 0.620505 | 0.512 | 0.295 | 4.04259E-20 |
| Fibroblasts | CASP4 | 1.14013E-70 | 0.62014 | 0.437 | 0.152 | 1.87255E-66 |
| RERGL_Cells | CABIN1 | 1.60108E-15 | 0.619432 | 0.276 | 0.149 | 2.62961E-11 |
| Contractile | RAB4A | 1.42662E-24 | 0.619306 | 0.461 | 0.263 | 2.34307E-20 |
| Fibroblasts | CADM3 | 3.88599E-58 | 0.61922 | 0.164 | 0.021 | 6.38236E-54 |
| RERGL_Cells | EFHD1 | 4.53099E-25 | 0.618584 | 0.6 | 0.382 | 7.44169E-21 |
| RERGL_Cells | GSTK1 | 6.03138E-13 | 0.617333 | 0.485 | 0.395 | 9.90595E-09 |
| RERGL_Cells | SLC2A4 | 8.31486E-35 | 0.617155 | 0.188 | 0.042 | 1.36563E-30 |
| Fibroblasts | NFIA | 1.21735E-55 | 0.616993 | 0.723 | 0.426 | 1.99937E-51 |
| Pericytes | EGFLAM | 2.81229E-25 | 0.616983 | 0.205 | 0.07 | 4.61891E-21 |
| Fibroblasts | LAMC1 | 8.05177E-49 | 0.616391 | 0.42 | 0.182 | 1.32242E-44 |
| Terminal | DPT | 2.00083E-16 | 0.616344 | 0.525 | 0.34 | 3.28616E-12 |
| RERGL_Cells | DNAJB4 | 8.71639E-14 | 0.616061 | 0.47 | 0.349 | 1.43158E-09 |
| Contractile | FAM207A | 1.35188E-23 | 0.616017 | 0.281 | 0.119 | 2.22032E-19 |
| Pericytes | H3F3B | 7.94017E-75 | 0.615881 | 0.998 | 0.99 | 1.30409E-70 |
| Pericytes | IMPDH2 | 9.92237E-18 | 0.615712 | 0.522 | 0.396 | 1.62965E-13 |
| Fibroblasts | MVP | 6.37017E-64 | 0.615693 | 0.623 | 0.303 | 1.04624E-59 |
| Contractile | CAPZA2 | 2.40297E-31 | 0.61565 | 0.782 | 0.605 | 3.94664E-27 |
| Pericytes | TIMP3 | 6.60403E-59 | 0.615648 | 0.976 | 0.766 | 1.08465E-54 |
| Fibroblasts | ADAMTSL2 | 3.45192E-92 | 0.614999 | 0.217 | 0.018 | 5.66943E-88 |
| Pericytes | ITGB1 | 7.54049E-46 | 0.614218 | 0.942 | 0.883 | 1.23845E-41 |
| RERGL_Cells | ATP5F1D | 1.63637E-20 | 0.613694 | 0.699 | 0.632 | 2.68758E-16 |
| Pericytes | VMP1 | 4.86998E-14 | 0.613579 | 0.529 | 0.415 | 7.99845E-10 |
| RERGL_Cells | PAWR | 8.52794E-19 | 0.612949 | 0.327 | 0.171 | 1.40063E-14 |
| RERGL_Cells | ANKRD29 | 9.48994E-27 | 0.612838 | 0.229 | 0.077 | 1.55863E-22 |
| Intermediate | INHBA | 8.73249E-45 | 0.612734 | 0.316 | 0.094 | 1.43422E-40 |
| Pericytes | OAZ2 | 5.91484E-25 | 0.612178 | 0.656 | 0.505 | 9.71454E-21 |
| Contractile | MYRFL | 9.00943E-53 | 0.611717 | 0.17 | 0.022 | 1.47971E-48 |
| RERGL_Cells | SH3KBP1 | 7.58628E-15 | 0.611708 | 0.321 | 0.19 | 1.24597E-10 |
| Contractile | DUSP26 | 8.25118E-47 | 0.611646 | 0.242 | 0.055 | 1.35517E-42 |
| RERGL_Cells | GLRX5 | 5.67386E-15 | 0.611583 | 0.323 | 0.197 | 9.31875E-11 |
| Fibroblasts | LINC0114( | 1.16512E-97 | 0.611517 | 0.25 | 0.025 | 1.91359E-93 |
| RERGL_Cells | HCFC1R1 | 7.13177E-23 | 0.611211 | 0.677 | 0.52 | 1.17132E-18 |
| RERGL_Cells | EBF1 | 9.72933E-15 | 0.61088 | 0.502 | 0.366 | 1.59794E-10 |
| RERGL_Cells | COX14 | 1.42171E-17 | 0.610509 | 0.549 | 0.423 | 2.33502E-13 |
| Fibroblasts | FTL | 6.978E-110 | 0.609774 | 1 | 0.992 | 1.1461E-105 |
| RERGL_Cells | GPBAR1 | 1.14039E-29 | 0.609688 | 0.115 | 0.019 | 1.87297E-25 |
| Contractile | ATOH8 | 3.3349E-34 | 0.609409 | 0.313 | 0.111 | 5.47724E-30 |
| Contractile | SBDS | 4.80567E-38 | 0.609244 | 0.888 | 0.776 | 7.89283E-34 |
| RERGL_Cells | ARHGAP1( | 1.02784E-16 | 0.608845 | 0.323 | 0.18 | 1.68813E-12 |
| Fibroblasts | IFITM2 | 9.98766E-91 | 0.608199 | 0.984 | 0.873 | 1.64037E-86 |
| Fibroblasts | USP53 | 8.13808E-59 | 0.60783 | 0.319 | 0.096 | 1.3366E-54 |
| Fibroblasts | CXCL12 | 1.46541E-43 | 0.607498 | 0.784 | 0.519 | 2.40678E-39 |
| Pericytes | TOB1 | 3.30068E-18 | 0.607241 | 0.473 | 0.311 | 5.42104E-14 |
| RERGL_Cells | ZFAND1 | 2.32903E-13 | 0.607134 | 0.432 | 0.317 | 3.8252E-09 |
| Contractile | BEX4 | 2.70624E-25 | 0.606948 | 0.489 | 0.278 | 4.44473E-21 |

| | | | | | | |
|---|---|---|---|---|---|---|
| Fibroblasts | MCUB | 8.63309E-89 | 0.606924 | 0.313 | 0.06 | 1.4179E-84 |
| RERGL_Cells | CYC1 | 4.90023E-15 | 0.60673 | 0.556 | 0.465 | 8.04814E-11 |
| RERGL_Cells | OR51E1 | 3.75801E-78 | 0.605601 | 0.143 | 0.005 | 6.17215E-74 |
| Fibroblasts | CD68 | 1.37623E-66 | 0.605586 | 0.301 | 0.078 | 2.26032E-62 |
| Contractile | GSTP1 | 1.89285E-59 | 0.605516 | 0.984 | 0.933 | 3.10882E-55 |
| RERGL_Cells | MICAL1 | 9.07229E-23 | 0.604662 | 0.222 | 0.082 | 1.49003E-18 |
| Contractile | TLN1 | 4.30934E-27 | 0.604174 | 0.579 | 0.337 | 7.07766E-23 |
| Fibroblasts | MBP | 9.20616E-97 | 0.603987 | 0.254 | 0.028 | 1.51202E-92 |
| RERGL_Cells | NIPSNAP2 | 1.43522E-15 | 0.603913 | 0.344 | 0.211 | 2.35721E-11 |
| Contractile | MRVI1 | 1.40843E-30 | 0.603606 | 0.361 | 0.15 | 2.3132E-26 |
| Contractile | ISCU | 6.02983E-35 | 0.60357 | 0.868 | 0.706 | 9.9034E-31 |
| Fibroblasts | DAAM1 | 1.15019E-40 | 0.603537 | 0.519 | 0.276 | 1.88907E-36 |
| RERGL_Cells | RHOB | 6.16819E-12 | 0.603121 | 0.626 | 0.545 | 1.01306E-07 |
| Pericytes | CSGALNAC | 3.25086E-23 | 0.60299 | 0.233 | 0.093 | 5.33921E-19 |
| RERGL_Cells | LINC02381 | 8.0598E-16 | 0.602496 | 0.197 | 0.085 | 1.32374E-11 |
| Fibroblasts | SDCBP | 8.58052E-49 | 0.60161 | 0.768 | 0.522 | 1.40927E-44 |
| Fibroblasts | RASD1 | 2.96334E-18 | 0.601447 | 0.23 | 0.115 | 4.86699E-14 |
| Contractile | HRCT1 | 2.79377E-28 | 0.600332 | 0.238 | 0.081 | 4.58849E-24 |
| Fibroblasts | TANK | 7.39049E-55 | 0.599841 | 0.498 | 0.225 | 1.21381E-50 |
| RERGL_Cells | COL18A1 | 1.13308E-22 | 0.599242 | 0.654 | 0.489 | 1.86097E-18 |
| RERGL_Cells | KCNE4 | 5.80058E-23 | 0.598153 | 0.154 | 0.043 | 9.52687E-19 |
| RERGL_Cells | HSPB2 | 4.04357E-13 | 0.597957 | 0.449 | 0.337 | 6.64116E-09 |
| Contractile | DANCR | 9.90496E-20 | 0.597448 | 0.403 | 0.226 | 1.62679E-15 |
| Fibroblasts | PLAT | 1.07062E-52 | 0.597165 | 0.26 | 0.072 | 1.75839E-48 |
| Contractile | KANK1 | 2.59558E-33 | 0.597121 | 0.259 | 0.084 | 4.26298E-29 |
| Fibroblasts | FLRT2 | 1.87209E-93 | 0.597096 | 0.196 | 0.01 | 3.07471E-89 |
| Pericytes | EIF5A | 6.48303E-18 | 0.596044 | 0.705 | 0.637 | 1.06477E-13 |
| Contractile | RSL24D1 | 8.56877E-30 | 0.595869 | 0.828 | 0.658 | 1.40733E-25 |
| RERGL_Cells | UTRN | 3.8188E-16 | 0.595548 | 0.284 | 0.15 | 6.27199E-12 |
| RERGL_Cells | NR2F2-AS1 | 8.16206E-18 | 0.595486 | 0.25 | 0.115 | 1.34054E-13 |
| RERGL_Cells | HES1 | 1.51969E-29 | 0.595441 | 0.577 | 0.33 | 2.49594E-25 |
| RERGL_Cells | LDHB | 1.3636E-23 | 0.595357 | 0.791 | 0.695 | 2.23957E-19 |
| Intermediate | LAMP1 | 3.05742E-28 | 0.595129 | 0.856 | 0.702 | 5.0215E-24 |
| Fibroblasts | PCSK5 | 1.48473E-68 | 0.595085 | 0.242 | 0.045 | 2.43853E-64 |
| Intermediate | COL4A1 | 1.73593E-25 | 0.594746 | 0.529 | 0.292 | 2.8511E-21 |
| RERGL_Cells | FAU | 1.20808E-61 | 0.594736 | 0.996 | 0.98 | 1.98414E-57 |
| RERGL_Cells | AGT | 4.24299E-32 | 0.594351 | 0.573 | 0.297 | 6.96869E-28 |
| Contractile | LGR6 | 9.18645E-36 | 0.594242 | 0.246 | 0.071 | 1.50878E-31 |
| Fibroblasts | IFI27 | 2.60677E-22 | 0.593628 | 0.177 | 0.07 | 4.28137E-18 |
| RERGL_Cells | KLF11 | 1.44393E-21 | 0.593308 | 0.246 | 0.1 | 2.37151E-17 |
| Fibroblasts | ATP1A1 | 1.53288E-54 | 0.592765 | 0.694 | 0.404 | 2.5176E-50 |
| Fibroblasts | HIC1 | 2.3002E-74 | 0.592503 | 0.233 | 0.036 | 3.77785E-70 |
| Pericytes | BX664615. | 6.8421E-16 | 0.5925 | 0.204 | 0.095 | 1.12375E-11 |
| Pericytes | HNRNPF | 2.77224E-21 | 0.591874 | 0.615 | 0.492 | 4.55313E-17 |
| Pericytes | ST5 | 7.38163E-22 | 0.59173 | 0.267 | 0.121 | 1.21236E-17 |
| RERGL_Cells | ATP1B2 | 3.88419E-41 | 0.591721 | 0.154 | 0.023 | 6.37939E-37 |
| Contractile | MRAP2 | 2.249E-31 | 0.591341 | 0.277 | 0.096 | 3.69377E-27 |
| RERGL_Cells | MRVI1 | 1.31137E-19 | 0.591135 | 0.318 | 0.158 | 2.15379E-15 |
| Pericytes | NTN4 | 1.64925E-14 | 0.59105 | 0.438 | 0.303 | 2.70872E-10 |
| Intermediate | ITGB5 | 6.34337E-36 | 0.590843 | 0.31 | 0.107 | 1.04183E-31 |
| Fibroblasts | CHEK2 | 1.5742E-108 | 0.590335 | 0.222 | 0.011 | 2.5854E-104 |
| RERGL_Cells | LMOD1 | 1.27807E-27 | 0.590207 | 0.694 | 0.432 | 2.0991E-23 |
| Fibroblasts | LINC01133 | 1.76403E-65 | 0.590069 | 0.17 | 0.018 | 2.89724E-61 |
| RERGL_Cells | TNFSF12 | 1.15853E-12 | 0.589929 | 0.335 | 0.219 | 1.90278E-08 |
| Intermediate | GAPDH | 1.11729E-65 | 0.589791 | 0.998 | 0.975 | 1.83503E-61 |
| Contractile | LDB3 | 2.72655E-38 | 0.589759 | 0.293 | 0.089 | 4.47809E-34 |

| | | | | | | |
|---|---|---|---|---|---|---|
| Terminal | LINC0161! | 3.90917E-12 | 0.58955 | 0.246 | 0.133 | 6.42042E-08 |
| Terminal | GABARAPI | 3.65939E-27 | 0.589405 | 0.828 | 0.727 | 6.01018E-23 |
| Intermediate | DAP | 6.26635E-31 | 0.58916 | 0.518 | 0.273 | 1.02918E-26 |
| RERGL_Cells | GRIP2 | 1.04514E-59 | 0.587643 | 0.145 | 0.011 | 1.71654E-55 |
| Intermediate | ATP5F1E | 5.45705E-54 | 0.587552 | 0.966 | 0.886 | 8.96265E-50 |
| RERGL_Cells | TXNIP | 2.30856E-14 | 0.587384 | 0.756 | 0.644 | 3.79157E-10 |
| Contractile | FAU | 5.65229E-64 | 0.586943 | 0.994 | 0.98 | 9.28332E-60 |
| Terminal | LIMCH1 | 8.94092E-25 | 0.586512 | 0.383 | 0.181 | 1.46846E-20 |
| Fibroblasts | NFIB | 3.53266E-44 | 0.585002 | 0.747 | 0.501 | 5.80204E-40 |
| RERGL_Cells | CAT | 4.01908E-14 | 0.584569 | 0.491 | 0.38 | 6.60094E-10 |
| Contractile | RAP1A | 2.55097E-20 | 0.584437 | 0.545 | 0.373 | 4.18971E-16 |
| Intermediate | HSPB1 | 9.76088E-65 | 0.584332 | 0.998 | 0.941 | 1.60313E-60 |
| Fibroblasts | APOL3 | 2.98032E-71 | 0.584268 | 0.323 | 0.084 | 4.89488E-67 |
| Contractile | GMPR | 7.47465E-28 | 0.584065 | 0.283 | 0.109 | 1.22764E-23 |
| RERGL_Cells | CBLN1 | 8.43515E-65 | 0.583571 | 0.13 | 0.006 | 1.38539E-60 |
| Contractile | TOMM7 | 2.19084E-44 | 0.582972 | 0.97 | 0.902 | 3.59823E-40 |
| Fibroblasts | TWIST2 | 2.08527E-71 | 0.582702 | 0.294 | 0.068 | 3.42485E-67 |
| Pericytes | NTMT1 | 2.05352E-15 | 0.582189 | 0.282 | 0.159 | 3.3727E-11 |
| Fibroblasts | PDIA3 | 4.06332E-63 | 0.582126 | 0.911 | 0.751 | 6.6736E-59 |
| Intermediate | TMED9 | 1.22418E-31 | 0.581235 | 0.811 | 0.613 | 2.0106E-27 |
| Contractile | SIVA1 | 9.75907E-26 | 0.580816 | 0.693 | 0.52 | 1.60283E-21 |
| Pericytes | SEMA5A | 2.08196E-47 | 0.580774 | 0.195 | 0.036 | 3.41941E-43 |
| Pericytes | ADGRE5 | 4.57368E-13 | 0.580384 | 0.347 | 0.228 | 7.51182E-09 |
| Pericytes | ACTR3 | 3.80619E-18 | 0.580239 | 0.595 | 0.495 | 6.25129E-14 |
| Contractile | WDR1 | 2.35093E-28 | 0.57984 | 0.729 | 0.531 | 3.86116E-24 |
| Fibroblasts | CSTA | 1.3582E-95 | 0.579282 | 0.177 | 0.004 | 2.23071E-91 |
| RERGL_Cells | RCSD1 | 2.547E-23 | 0.579097 | 0.197 | 0.066 | 4.18319E-19 |
| Terminal | DPP7 | 1.40407E-14 | 0.579082 | 0.534 | 0.408 | 2.30605E-10 |
| Fibroblasts | VMP1 | 4.1113E-48 | 0.578926 | 0.631 | 0.347 | 6.7524E-44 |
| Fibroblasts | TIMP2 | 1.59123E-37 | 0.57892 | 0.608 | 0.356 | 2.61343E-33 |
| Contractile | ROCK1 | 5.73226E-25 | 0.57888 | 0.597 | 0.378 | 9.41466E-21 |
| Fibroblasts | CPQ | 5.54966E-48 | 0.578462 | 0.659 | 0.389 | 9.11475E-44 |
| Contractile | LDHB | 4.8948E-34 | 0.578172 | 0.848 | 0.685 | 8.03921E-30 |
| Pericytes | SRSF2 | 4.20609E-19 | 0.578157 | 0.678 | 0.602 | 6.90809E-15 |
| Fibroblasts | SLC39A14 | 3.36655E-47 | 0.578 | 0.29 | 0.097 | 5.52922E-43 |
| Fibroblasts | RNF13 | 4.12724E-51 | 0.577719 | 0.558 | 0.278 | 6.77859E-47 |
| RERGL_Cells | CSDE1 | 1.54562E-25 | 0.577661 | 0.761 | 0.65 | 2.53853E-21 |
| Pericytes | LDLR | 3.85496E-29 | 0.577605 | 0.176 | 0.049 | 6.33138E-25 |
| Fibroblasts | SH3D19 | 4.33279E-70 | 0.577553 | 0.285 | 0.065 | 7.11618E-66 |
| Contractile | MRPL33 | 4.72888E-26 | 0.576585 | 0.733 | 0.542 | 7.76671E-22 |
| Fibroblasts | CBLB | 8.12364E-63 | 0.576254 | 0.277 | 0.067 | 1.33423E-58 |
| Intermediate | CLIC4 | 1.96216E-17 | 0.576191 | 0.718 | 0.563 | 3.22266E-13 |
| RERGL_Cells | PDGFA | 5.28436E-11 | 0.575878 | 0.263 | 0.153 | 8.67903E-07 |
| Pericytes | COL4A1 | 3.14764E-15 | 0.575711 | 0.436 | 0.308 | 5.16969E-11 |
| Fibroblasts | SLC25A37 | 3.04929E-50 | 0.575277 | 0.364 | 0.137 | 5.00815E-46 |
| Contractile | CAV1 | 8.94906E-41 | 0.574796 | 0.946 | 0.697 | 1.46979E-36 |
| Contractile | CAP1 | 6.15378E-32 | 0.57459 | 0.83 | 0.671 | 1.0107E-27 |
| Fibroblasts | PPP3CA | 3.53245E-50 | 0.574187 | 0.35 | 0.129 | 5.8017E-46 |
| RERGL_Cells | ZNF503 | 1.9362E-14 | 0.573746 | 0.274 | 0.149 | 3.18002E-10 |
| RERGL_Cells | TESC | 1.17245E-61 | 0.57355 | 0.111 | 0.004 | 1.92564E-57 |
| RERGL_Cells | KTN1 | 1.32165E-20 | 0.573503 | 0.671 | 0.554 | 2.17068E-16 |
| Fibroblasts | ABHD5 | 1.00539E-55 | 0.573449 | 0.29 | 0.084 | 1.65126E-51 |
| RERGL_Cells | AC092171 | 3.61943E-23 | 0.57332 | 0.158 | 0.045 | 5.94455E-19 |
| Contractile | EEF1A1 | 3.17583E-66 | 0.573226 | 1 | 0.997 | 5.21599E-62 |
| Fibroblasts | OSR1 | 1.8923E-46 | 0.572809 | 0.227 | 0.063 | 3.10791E-42 |
| RERGL_Cells | PIP5K1B | 5.48865E-33 | 0.572673 | 0.162 | 0.034 | 9.01456E-29 |

| | | | | | | |
|---|---|---|---|---|---|---|
| Intermediate | ITGA11 | 2.58715E-66 | 0.572491 | 0.26 | 0.045 | 4.24913E-62 |
| Intermediate | COL18A1 | 5.30753E-23 | 0.571419 | 0.716 | 0.474 | 8.71709E-19 |
| Pericytes | AAK1 | 4.06847E-16 | 0.571197 | 0.276 | 0.15 | 6.68206E-12 |
| Terminal | ASPN | 4.65845E-19 | 0.571185 | 0.58 | 0.366 | 7.65104E-15 |
| Fibroblasts | PRELP | 3.35814E-97 | 0.571168 | 0.944 | 0.626 | 5.51541E-93 |
| Fibroblasts | LTC4S | 3.2527E-48 | 0.570687 | 0.248 | 0.072 | 5.34223E-44 |
| RERGL_Cells | UQCRC2 | 4.91142E-17 | 0.57068 | 0.575 | 0.461 | 8.06651E-13 |
| Pericytes | SNRK | 1.09138E-23 | 0.569941 | 0.173 | 0.055 | 1.79249E-19 |
| Fibroblasts | SMIM14 | 3.78263E-50 | 0.569899 | 0.546 | 0.275 | 6.21259E-46 |
| Pericytes | NEURL1B | 4.28942E-32 | 0.569522 | 0.145 | 0.03 | 7.04494E-28 |
| RERGL_Cells | ITM2C | 7.38367E-11 | 0.568881 | 0.261 | 0.152 | 1.21269E-06 |
| Fibroblasts | ACKR4 | 9.87027E-65 | 0.568737 | 0.183 | 0.023 | 1.62109E-60 |
| Fibroblasts | DNAJB9 | 6.68835E-37 | 0.568559 | 0.486 | 0.265 | 1.0985E-32 |
| RERGL_Cells | ATP5MD | 1.57435E-25 | 0.568312 | 0.829 | 0.754 | 2.58571E-21 |
| Pericytes | ARHGAP6 | 1.14763E-21 | 0.567487 | 0.231 | 0.097 | 1.88487E-17 |
| Intermediate | GLRB | 5.69602E-61 | 0.567434 | 0.26 | 0.048 | 9.35514E-57 |
| Pericytes | SMC6 | 1.025E-19 | 0.567427 | 0.211 | 0.087 | 1.68347E-15 |
| Intermediate | LGALS1 | 2.42327E-56 | 0.567342 | 0.996 | 0.981 | 3.97997E-52 |
| Terminal | FIBIN | 1.12372E-13 | 0.567051 | 0.569 | 0.418 | 1.8456E-09 |
| RERGL_Cells | YWHAQ | 1.50929E-26 | 0.566347 | 0.799 | 0.699 | 2.47886E-22 |
| RERGL_Cells | NDUFA3 | 2.79652E-16 | 0.566121 | 0.6 | 0.498 | 4.593E-12 |
| Pericytes | SRSF7 | 1.17189E-18 | 0.566041 | 0.658 | 0.578 | 1.92471E-14 |
| Terminal | SNHG29 | 1.15918E-23 | 0.566008 | 0.898 | 0.831 | 1.90384E-19 |
| Contractile | HSPB6 | 1.15278E-17 | 0.565871 | 0.653 | 0.474 | 1.89333E-13 |
| Fibroblasts | WSB1 | 4.93613E-49 | 0.565744 | 0.814 | 0.54 | 8.1071E-45 |
| Pericytes | GYG1 | 5.71519E-17 | 0.56553 | 0.551 | 0.423 | 9.38663E-13 |
| Terminal | SUGCT | 1.66218E-12 | 0.565465 | 0.327 | 0.192 | 2.72996E-08 |
| Terminal | RGS3 | 1.10813E-16 | 0.565404 | 0.318 | 0.167 | 1.81999E-12 |
| Pericytes | SMTN | 1.67383E-15 | 0.56533 | 0.504 | 0.358 | 2.7491E-11 |
| Terminal | GAS6 | 9.13883E-26 | 0.565249 | 0.824 | 0.665 | 1.50096E-21 |
| RERGL_Cells | NFASC | 1.8966E-24 | 0.56494 | 0.212 | 0.071 | 3.11498E-20 |
| RERGL_Cells | AL021155. | 6.32289E-17 | 0.564902 | 0.135 | 0.043 | 1.03847E-12 |
| Intermediate | PFN1 | 4.9244E-42 | 0.56485 | 0.972 | 0.911 | 8.08783E-38 |
| RERGL_Cells | CAPZA2 | 4.69305E-18 | 0.564824 | 0.684 | 0.621 | 7.70786E-14 |
| RERGL_Cells | SELENOW | 1.049E-32 | 0.564544 | 0.897 | 0.791 | 1.72288E-28 |
| Terminal | ITGBL1 | 8.87961E-12 | 0.56453 | 0.316 | 0.191 | 1.45839E-07 |
| Intermediate | PPIC | 5.32511E-26 | 0.564316 | 0.645 | 0.415 | 8.74596E-22 |
| Intermediate | SH3BGRL3 | 2.17034E-21 | 0.564225 | 0.789 | 0.631 | 3.56457E-17 |
| Terminal | CTSZ | 1.54097E-18 | 0.564131 | 0.362 | 0.198 | 2.53089E-14 |
| Intermediate | COL1A1 | 1.12967E-13 | 0.564086 | 0.622 | 0.452 | 1.85537E-09 |
| Pericytes | SASH1 | 4.42238E-16 | 0.563647 | 0.22 | 0.105 | 7.26331E-12 |
| Intermediate | TMEM130 | 1.57077E-70 | 0.563563 | 0.2 | 0.022 | 2.57983E-66 |
| Pericytes | FGF10 | 3.53171E-42 | 0.563375 | 0.136 | 0.019 | 5.80049E-38 |
| Contractile | COPRS | 3.28115E-19 | 0.563175 | 0.525 | 0.354 | 5.38896E-15 |
| Pericytes | EPHA2 | 1.76884E-38 | 0.56277 | 0.142 | 0.023 | 2.90514E-34 |
| Terminal | EDIL3 | 3.3787E-24 | 0.562717 | 0.464 | 0.243 | 5.54918E-20 |
| Fibroblasts | ITGBL1 | 4.63445E-55 | 0.562549 | 0.378 | 0.134 | 7.61161E-51 |
| RERGL_Cells | VDAC2 | 6.73261E-20 | 0.562434 | 0.697 | 0.598 | 1.10576E-15 |
| Contractile | PBXIP1 | 8.60285E-16 | 0.562136 | 0.485 | 0.335 | 1.41293E-11 |
| Pericytes | TRA2B | 1.18537E-17 | 0.562119 | 0.576 | 0.464 | 1.94686E-13 |
| RERGL_Cells | JUN | 1.34122E-22 | 0.562045 | 0.816 | 0.665 | 2.20282E-18 |
| Terminal | PRSS23 | 3.1706E-30 | 0.561901 | 0.584 | 0.322 | 5.20739E-26 |
| Intermediate | GLIPR1 | 1.58694E-32 | 0.561663 | 0.579 | 0.309 | 2.6064E-28 |
| Fibroblasts | PLAUR | 2.46695E-57 | 0.561453 | 0.254 | 0.063 | 4.05173E-53 |
| Terminal | PPP1CB | 7.27063E-16 | 0.561306 | 0.754 | 0.646 | 1.19413E-11 |
| RERGL_Cells | NDUFB8 | 6.56748E-24 | 0.560888 | 0.778 | 0.692 | 1.07864E-19 |

| | | | | | | |
|---|---|---|---|---|---|---|
| Intermediate | PLP2 | 4.19947E-28 | 0.560583 | 0.811 | 0.6 | 6.8972E-24 |
| Terminal | HEYL | 3.99114E-11 | 0.560574 | 0.364 | 0.24 | 6.55505E-07 |
| RERGL_Cells | FADS3 | 2.12102E-11 | 0.560506 | 0.376 | 0.263 | 3.48356E-07 |
| Intermediate | CRISPLD1 | 1.85128E-33 | 0.560385 | 0.421 | 0.18 | 3.04055E-29 |
| Contractile | EIF4A2 | 1.98825E-37 | 0.559868 | 0.952 | 0.886 | 3.2655E-33 |
| Pericytes | ELOVL5 | 4.02637E-16 | 0.559434 | 0.376 | 0.236 | 6.61291E-12 |
| RERGL_Cells | CKMT2 | 1.52645E-25 | 0.559308 | 0.179 | 0.052 | 2.50704E-21 |
| RERGL_Cells | ZEB1 | 2.19467E-12 | 0.559055 | 0.278 | 0.164 | 3.60453E-08 |
| Contractile | MTHFD1L | 7.8515E-20 | 0.558715 | 0.265 | 0.121 | 1.28953E-15 |
| Contractile | PRR34-AS1 | 5.48393E-34 | 0.558442 | 0.273 | 0.088 | 9.0068E-30 |
| Fibroblasts | CD200 | 3.72274E-58 | 0.558326 | 0.326 | 0.102 | 6.11423E-54 |
| Fibroblasts | EMILIN1 | 3.54337E-37 | 0.558242 | 0.599 | 0.356 | 5.81963E-33 |
| Pericytes | SGIP1 | 5.85385E-27 | 0.557708 | 0.176 | 0.052 | 9.61437E-23 |
| Fibroblasts | PLEKHA5 | 3.69343E-66 | 0.557299 | 0.321 | 0.086 | 6.06608E-62 |
| RERGL_Cells | EIF1AY | 8.20196E-13 | 0.556633 | 0.288 | 0.169 | 1.34709E-08 |
| Intermediate | CRIM1 | 6.81037E-27 | 0.556605 | 0.529 | 0.279 | 1.11854E-22 |
| Contractile | SOST | 1.55469E-47 | 0.556574 | 0.144 | 0.017 | 2.55342E-43 |
| Contractile | NGRN | 6.39053E-24 | 0.556571 | 0.677 | 0.502 | 1.04958E-19 |
| Intermediate | MIR100HG | 5.68306E-31 | 0.55612 | 0.424 | 0.19 | 9.33385E-27 |
| Contractile | COX14 | 7.92296E-20 | 0.556104 | 0.577 | 0.417 | 1.30127E-15 |
| Contractile | CCNI | 1.07689E-35 | 0.555917 | 0.95 | 0.859 | 1.76868E-31 |
| Pericytes | PSME1 | 1.86029E-22 | 0.555815 | 0.718 | 0.639 | 3.05534E-18 |
| Fibroblasts | ABHD14A | 2.00281E-60 | 0.555673 | 0.357 | 0.117 | 3.28942E-56 |
| Contractile | DYNLRB1 | 1.51827E-24 | 0.555482 | 0.741 | 0.559 | 2.4936E-20 |
| Intermediate | ENPP5 | 1.11996E-58 | 0.555467 | 0.196 | 0.027 | 1.83943E-54 |
| RERGL_Cells | ELOB | 1.08473E-32 | 0.55525 | 0.87 | 0.812 | 1.78156E-28 |
| RERGL_Cells | CKB | 3.65255E-06 | 0.554918 | 0.425 | 0.356 | 0.059989425 |
| Fibroblasts | HELLPAR | 1.3842E-73 | 0.554497 | 0.179 | 0.016 | 2.27341E-69 |
| Pericytes | SRSF5 | 3.54186E-21 | 0.554444 | 0.82 | 0.756 | 5.81716E-17 |
| Fibroblasts | SELENOF | 1.15388E-55 | 0.554379 | 0.818 | 0.529 | 1.89513E-51 |
| RERGL_Cells | SMARCD3 | 7.1261E-13 | 0.554366 | 0.325 | 0.199 | 1.17039E-08 |
| RERGL_Cells | ATP5ME | 5.72181E-24 | 0.554011 | 0.797 | 0.76 | 9.3975E-20 |
| Pericytes | TBCC | 1.1455E-16 | 0.553886 | 0.262 | 0.138 | 1.88137E-12 |
| RERGL_Cells | ATP5F1A | 8.54941E-21 | 0.553425 | 0.724 | 0.631 | 1.40416E-16 |
| Contractile | VPS29 | 5.1058E-23 | 0.553371 | 0.661 | 0.478 | 8.38576E-19 |
| RERGL_Cells | ARHGEF25 | 2.39438E-16 | 0.552782 | 0.269 | 0.137 | 3.93254E-12 |
| Intermediate | GJA1 | 1.20146E-47 | 0.552672 | 0.249 | 0.058 | 1.97328E-43 |
| Contractile | ST13 | 9.16366E-28 | 0.552529 | 0.776 | 0.612 | 1.50504E-23 |
| Fibroblasts | IL1R1 | 2.93437E-77 | 0.551836 | 0.254 | 0.042 | 4.81942E-73 |
| Terminal | SERTAD4 | 2.40332E-23 | 0.551609 | 0.192 | 0.061 | 3.94721E-19 |
| Intermediate | MRPS6 | 6.81194E-32 | 0.551294 | 0.512 | 0.252 | 1.11879E-27 |
| Pericytes | PDE4B | 2.6841E-33 | 0.551283 | 0.144 | 0.029 | 4.40837E-29 |
| Fibroblasts | NUPR1 | 2.57225E-45 | 0.551133 | 0.918 | 0.806 | 4.22467E-41 |
| Fibroblasts | LAMB1 | 9.96052E-68 | 0.550794 | 0.26 | 0.054 | 1.63592E-63 |
| Pericytes | SRSF6 | 2.18336E-14 | 0.550398 | 0.467 | 0.342 | 3.58595E-10 |
| RERGL_Cells | ST6GALNA | 6.9897E-10 | 0.549982 | 0.462 | 0.389 | 1.14799E-05 |
| Intermediate | BSG | 5.13554E-38 | 0.549638 | 0.921 | 0.804 | 8.43461E-34 |
| Intermediate | TSPAN8 | 1.48486E-14 | 0.549462 | 0.34 | 0.196 | 2.43873E-10 |
| RERGL_Cells | YBX1 | 1.70251E-43 | 0.549239 | 0.966 | 0.923 | 2.79621E-39 |
| Intermediate | HAPLN3 | 2.54333E-47 | 0.54909 | 0.256 | 0.06 | 4.17716E-43 |
| Pericytes | PLAU | 1.65398E-18 | 0.549003 | 0.231 | 0.103 | 2.71649E-14 |
| RERGL_Cells | HIF1A | 5.61431E-13 | 0.548987 | 0.427 | 0.303 | 9.22094E-09 |
| Terminal | FTL | 1.60932E-37 | 0.548694 | 1 | 0.994 | 2.64314E-33 |
| RERGL_Cells | EEF1D | 6.73097E-31 | 0.548673 | 0.936 | 0.912 | 1.10549E-26 |
| Fibroblasts | KDELR2 | 8.63991E-48 | 0.548142 | 0.8 | 0.561 | 1.41902E-43 |
| Contractile | KIF5B | 1.7501E-19 | 0.547999 | 0.657 | 0.507 | 2.87437E-15 |

| | | | | | | |
|---|---|---|---|---|---|---|
| Contractile | CDH13 | 1.85434E-24 | 0.547992 | 0.333 | 0.146 | 3.04557E-20 |
| Fibroblasts | SREBF1 | 7.54185E-66 | 0.547992 | 0.231 | 0.043 | 1.23867E-61 |
| Pericytes | SLCO3A1 | 9.5372E-17 | 0.547958 | 0.247 | 0.123 | 1.56639E-12 |
| RERGL_Cells | RASGEF1B | 1.13491E-13 | 0.547768 | 0.128 | 0.046 | 1.86397E-09 |
| Terminal | ABI3BP | 1.85835E-07 | 0.547759 | 0.316 | 0.223 | 0.00305215 |
| Fibroblasts | ABLIM1 | 6.14511E-74 | 0.547265 | 0.219 | 0.031 | 1.00927E-69 |
| Intermediate | CD82 | 8.50311E-34 | 0.547169 | 0.323 | 0.117 | 1.39655E-29 |
| Contractile | PRKAG2 | 2.0038E-29 | 0.546955 | 0.297 | 0.113 | 3.29105E-25 |
| Pericytes | WDR1 | 5.46521E-20 | 0.546362 | 0.667 | 0.539 | 8.97606E-16 |
| Fibroblasts | C2 | 5.04709E-85 | 0.54604 | 0.216 | 0.022 | 8.28934E-81 |
| Fibroblasts | TSKU | 3.11157E-56 | 0.545916 | 0.249 | 0.062 | 5.11044E-52 |
| Contractile | CPED1 | 5.12722E-26 | 0.545895 | 0.333 | 0.144 | 8.42095E-22 |
| Fibroblasts | IER3 | 1.47913E-34 | 0.545446 | 0.328 | 0.144 | 2.42933E-30 |
| RERGL_Cells | NPNT | 3.92639E-17 | 0.545261 | 0.274 | 0.133 | 6.44871E-13 |
| Intermediate | CALM1 | 2.06081E-33 | 0.544017 | 0.899 | 0.772 | 3.38467E-29 |
| Pericytes | ANKRD29 | 9.27002E-29 | 0.543861 | 0.224 | 0.074 | 1.52251E-24 |
| Fibroblasts | PRDX4 | 8.09848E-46 | 0.543614 | 0.712 | 0.44 | 1.33009E-41 |
| Pericytes | NFIL3 | 4.53983E-17 | 0.542856 | 0.175 | 0.07 | 7.45622E-13 |
| Terminal | SGCA | 1.57077E-18 | 0.542769 | 0.558 | 0.354 | 2.57984E-14 |
| RERGL_Cells | NDUFA1 | 1.70805E-18 | 0.542569 | 0.746 | 0.702 | 2.80529E-14 |
| Pericytes | ARHGAP24 | 5.06546E-20 | 0.542388 | 0.191 | 0.075 | 8.31951E-16 |
| Contractile | ATP5MC2 | 7.01335E-32 | 0.542166 | 0.874 | 0.74 | 1.15187E-27 |
| Pericytes | RICTOR | 2.73909E-12 | 0.542047 | 0.271 | 0.16 | 4.49868E-08 |
| Pericytes | RHOB | 1.05007E-07 | 0.54155 | 0.589 | 0.55 | 0.001724639 |
| Contractile | ACTN4 | 2.00329E-33 | 0.541335 | 0.8 | 0.531 | 3.29021E-29 |
| Pericytes | NMD3 | 2.9558E-14 | 0.541324 | 0.367 | 0.243 | 4.8546E-10 |
| Terminal | CD99 | 6.78472E-27 | 0.541296 | 0.895 | 0.859 | 1.11432E-22 |
| Pericytes | CPE | 6.75287E-36 | 0.541291 | 0.958 | 0.887 | 1.10909E-31 |
| Fibroblasts | CSF1 | 2.21548E-59 | 0.541009 | 0.351 | 0.112 | 3.63871E-55 |
| Terminal | FAP | 6.10845E-25 | 0.540544 | 0.222 | 0.075 | 1.00325E-20 |
| Pericytes | SORBS3 | 5.40969E-12 | 0.540419 | 0.415 | 0.309 | 8.88487E-08 |
| Fibroblasts | FGL2 | 5.32555E-30 | 0.540351 | 0.288 | 0.13 | 8.74669E-26 |
| Fibroblasts | CHL1 | 1.05536E-91 | 0.540003 | 0.172 | 0.004 | 1.73332E-87 |
| RERGL_Cells | SYTL2 | 1.1292E-22 | 0.539824 | 0.331 | 0.15 | 1.8546E-18 |
| Fibroblasts | KLF2 | 1.00746E-44 | 0.539602 | 0.44 | 0.195 | 1.65466E-40 |
| Contractile | CYFIP2 | 1.9886E-27 | 0.539225 | 0.228 | 0.075 | 3.26608E-23 |
| Pericytes | TMEM50A | 3.69822E-20 | 0.5392 | 0.705 | 0.628 | 6.07396E-16 |
| Fibroblasts | JUND | 8.66445E-44 | 0.539063 | 0.46 | 0.21 | 1.42305E-39 |
| Terminal | FKBP10 | 8.77503E-14 | 0.538796 | 0.516 | 0.377 | 1.44121E-09 |
| Contractile | PGM1 | 2.78532E-19 | 0.538752 | 0.321 | 0.166 | 4.57461E-15 |
| Intermediate | CSDC2 | 2.70678E-37 | 0.538705 | 0.361 | 0.13 | 4.44561E-33 |
| RERGL_Cells | ATP6V1G1 | 7.6904E-22 | 0.538541 | 0.771 | 0.704 | 1.26307E-17 |
| Contractile | KRT8 | 1.37702E-35 | 0.537909 | 0.421 | 0.168 | 2.26162E-31 |
| RERGL_Cells | MOB2 | 1.29485E-12 | 0.537778 | 0.4 | 0.269 | 2.12665E-08 |
| Contractile | ZFAND1 | 6.93423E-18 | 0.537502 | 0.475 | 0.309 | 1.13888E-13 |
| Fibroblasts | PROS1 | 6.35274E-55 | 0.537304 | 0.551 | 0.255 | 1.04337E-50 |
| Pericytes | PFDN2 | 2.64289E-17 | 0.537181 | 0.584 | 0.472 | 4.34068E-13 |
| RERGL_Cells | TNS1 | 9.16499E-15 | 0.537063 | 0.41 | 0.265 | 1.50526E-10 |
| RERGL_Cells | ATP5PO | 3.05182E-23 | 0.537041 | 0.786 | 0.68 | 5.01231E-19 |
| Intermediate | LGR6 | 6.42035E-45 | 0.536816 | 0.264 | 0.066 | 1.05448E-40 |
| Contractile | SCRG1 | 1.87546E-17 | 0.536737 | 0.293 | 0.148 | 3.08025E-13 |
| Fibroblasts | DCLK1 | 7.16949E-83 | 0.53664 | 0.205 | 0.019 | 1.17752E-78 |
| Pericytes | VCL | 1.19467E-13 | 0.536169 | 0.382 | 0.256 | 1.96212E-09 |
| Contractile | SGCA | 2.17098E-28 | 0.536055 | 0.617 | 0.342 | 3.56561E-24 |
| Contractile | ANAPC16 | 6.69697E-26 | 0.535933 | 0.802 | 0.656 | 1.09991E-21 |
| RERGL_Cells | PELI1 | 7.14765E-19 | 0.53587 | 0.188 | 0.069 | 1.17393E-14 |

| | | | | | | |
|---|---|---|---|---|---|---|
| Pericytes | MYO1C | 4.3286E-16 | 0.535242 | 0.438 | 0.3 | 7.10929E-12 |
| RERGL_Cells | EBF2 | 3.40788E-20 | 0.535017 | 0.199 | 0.074 | 5.59711E-16 |
| Contractile | EIF5B | 2.60627E-18 | 0.534969 | 0.659 | 0.51 | 4.28054E-14 |
| Fibroblasts | PLD3 | 3.67019E-44 | 0.534797 | 0.753 | 0.494 | 6.02792E-40 |
| Contractile | CNBP | 9.11122E-32 | 0.534792 | 0.886 | 0.81 | 1.49643E-27 |
| RERGL_Cells | ATP1A2 | 1.65833E-35 | 0.534655 | 0.158 | 0.029 | 2.72363E-31 |
| Contractile | PDLIM5 | 5.80498E-20 | 0.534531 | 0.525 | 0.337 | 9.53409E-16 |
| RERGL_Cells | VASP | 3.71889E-14 | 0.534166 | 0.385 | 0.246 | 6.1079E-10 |
| Contractile | SYF2 | 9.14183E-19 | 0.533963 | 0.655 | 0.492 | 1.50145E-14 |
| Pericytes | RGS5 | 0.001112151 | 0.533838 | 0.547 | 0.454 | 1 |
| Terminal | MFGE8 | 4.82973E-31 | 0.532583 | 0.967 | 0.847 | 7.93235E-27 |
| Fibroblasts | NUCB2 | 1.54095E-47 | 0.532556 | 0.662 | 0.365 | 2.53086E-43 |
| Contractile | SUGCT | 3.94577E-21 | 0.532456 | 0.373 | 0.183 | 6.48053E-17 |
| Intermediate | SYNGR2 | 4.87602E-33 | 0.532096 | 0.508 | 0.236 | 8.00837E-29 |
| Fibroblasts | C1GALT1 | 8.68595E-63 | 0.53202 | 0.263 | 0.061 | 1.42658E-58 |
| Contractile | GUK1 | 3.73292E-34 | 0.532011 | 0.906 | 0.785 | 6.13094E-30 |
| Contractile | NDUFS5 | 4.10212E-32 | 0.531961 | 0.886 | 0.792 | 6.73731E-28 |
| Terminal | CDH19 | 2.24396E-13 | 0.531689 | 0.27 | 0.139 | 3.68548E-09 |
| Fibroblasts | ALPL | 5.21769E-67 | 0.531578 | 0.126 | 0.003 | 8.56954E-63 |
| Pericytes | CDC42SE1 | 2.94949E-07 | 0.53147 | 0.264 | 0.186 | 0.004844239 |
| RERGL_Cells | GBP2 | 1.13127E-07 | 0.531395 | 0.393 | 0.32 | 0.001858 |
| Pericytes | SMIM3 | 4.63964E-11 | 0.531282 | 0.307 | 0.205 | 7.62014E-07 |
| Fibroblasts | CLEC11A | 4.36725E-54 | 0.531238 | 0.356 | 0.121 | 7.17278E-50 |
| Intermediate | SMOC2 | 3.18573E-20 | 0.531162 | 0.763 | 0.543 | 5.23224E-16 |
| Pericytes | AKIRIN1 | 3.85958E-13 | 0.530972 | 0.344 | 0.228 | 6.33898E-09 |
| Pericytes | TRPC6 | 2.25199E-47 | 0.530828 | 0.125 | 0.012 | 3.69868E-43 |
| Fibroblasts | SESTD1 | 1.30948E-46 | 0.530589 | 0.376 | 0.151 | 2.15069E-42 |
| Terminal | SCRG1 | 5.58044E-10 | 0.530394 | 0.264 | 0.154 | 9.16532E-06 |
| Contractile | CRYZL1 | 4.65487E-21 | 0.529955 | 0.359 | 0.184 | 7.64516E-17 |
| Contractile | DAPK3 | 7.54536E-20 | 0.529935 | 0.427 | 0.24 | 1.23925E-15 |
| Fibroblasts | ZC3H12A | 1.45187E-63 | 0.529755 | 0.214 | 0.037 | 2.38454E-59 |
| Contractile | POLR2E | 1.18019E-21 | 0.52974 | 0.617 | 0.442 | 1.93834E-17 |
| RERGL_Cells | NDUFS2 | 2.94343E-10 | 0.529624 | 0.427 | 0.343 | 4.83428E-06 |
| Fibroblasts | WIPI1 | 1.69893E-56 | 0.529527 | 0.305 | 0.092 | 2.79033E-52 |
| Pericytes | HNRNPH2 | 8.90823E-15 | 0.529403 | 0.551 | 0.446 | 1.46309E-10 |
| Contractile | YBX1 | 4.60037E-41 | 0.529337 | 0.968 | 0.922 | 7.55565E-37 |
| RERGL_Cells | INF2 | 1.45213E-09 | 0.5292 | 0.207 | 0.119 | 2.38498E-05 |
| RERGL_Cells | AC100830 | 9.90183E-41 | 0.528892 | 0.132 | 0.017 | 1.62628E-36 |
| RERGL_Cells | TGFB1I1 | 1.0481E-19 | 0.528291 | 0.726 | 0.608 | 1.72139E-15 |
| Terminal | SH3BGRL3 | 1.23329E-14 | 0.528226 | 0.739 | 0.642 | 2.02556E-10 |
| Fibroblasts | BCL3 | 2.72586E-54 | 0.528117 | 0.33 | 0.107 | 4.47694E-50 |
| Fibroblasts | TMEM258 | 1.33518E-51 | 0.528093 | 0.881 | 0.676 | 2.19291E-47 |
| Contractile | SORBS1 | 3.58136E-26 | 0.527786 | 0.208 | 0.066 | 5.88203E-22 |
| Contractile | FILIP1 | 1.43324E-19 | 0.5275 | 0.401 | 0.217 | 2.35395E-15 |
| Fibroblasts | DHCR7 | 8.04941E-77 | 0.5275 | 0.216 | 0.027 | 1.32204E-72 |
| Fibroblasts | MDFI | 1.22067E-75 | 0.527433 | 0.185 | 0.017 | 2.00482E-71 |
| Terminal | COL15A1 | 6.7277E-17 | 0.527424 | 0.336 | 0.178 | 1.10496E-12 |
| Contractile | PLEKHO1 | 3.12465E-20 | 0.527155 | 0.315 | 0.15 | 5.13192E-16 |
| Intermediate | DMKN | 5.69253E-39 | 0.526315 | 0.275 | 0.08 | 9.3494E-35 |
| Fibroblasts | S100A11 | 2.22389E-64 | 0.526177 | 0.979 | 0.887 | 3.65252E-60 |
| Pericytes | TUBB6 | 1.07855E-12 | 0.525706 | 0.567 | 0.489 | 1.7714E-08 |
| RERGL_Cells | UQCRH | 6.19296E-15 | 0.525692 | 0.69 | 0.609 | 1.01713E-10 |
| Fibroblasts | ANXA2 | 2.53368E-52 | 0.525443 | 0.956 | 0.825 | 4.16132E-48 |
| Intermediate | HRCT1 | 3.97674E-38 | 0.525376 | 0.262 | 0.075 | 6.5314E-34 |
| Fibroblasts | HSP90B1 | 4.68093E-36 | 0.525346 | 0.891 | 0.75 | 7.68796E-32 |
| RERGL_Cells | MICU3 | 1.74289E-20 | 0.525317 | 0.184 | 0.063 | 2.86253E-16 |

| | | | | | | |
|---|---|---|---|---|---|---|
| Fibroblasts | MIDN | 4.75226E-55 | 0.525215 | 0.359 | 0.121 | 7.80511E-51 |
| RERGL_Cells | GADD45G | 3.86104E-08 | 0.525108 | 0.244 | 0.153 | 0.000634137 |
| Pericytes | BTG3 | 7.61936E-11 | 0.524984 | 0.449 | 0.356 | 1.2514E-06 |
| Intermediate | TNFRSF12/ | 4.53032E-19 | 0.524983 | 0.628 | 0.413 | 7.44059E-15 |
| Terminal | NENF | 3.3416E-21 | 0.524954 | 0.817 | 0.704 | 5.48824E-17 |
| RERGL_Cells | HMGN2 | 1.04807E-20 | 0.524653 | 0.823 | 0.767 | 1.72136E-16 |
| Pericytes | TNFRSF12/ | 0.00050796 | 0.52448 | 0.467 | 0.441 | 1 |
| Fibroblasts | IER2 | 2.12063E-58 | 0.5243 | 0.814 | 0.507 | 3.48292E-54 |
| Intermediate | S100A6 | 1.14708E-57 | 0.524035 | 0.998 | 0.994 | 1.88397E-53 |
| Fibroblasts | DDR2 | 4.00717E-45 | 0.523971 | 0.354 | 0.14 | 6.58138E-41 |
| Pericytes | ATP8B4 | 1.33639E-18 | 0.523894 | 0.171 | 0.064 | 2.19488E-14 |
| Contractile | TUBA4A | 2.14246E-23 | 0.523548 | 0.236 | 0.091 | 3.51877E-19 |
| Terminal | NINJ1 | 6.87511E-13 | 0.52343 | 0.418 | 0.273 | 1.12917E-08 |
| RERGL_Cells | MAP7D3 | 4.02821E-14 | 0.523124 | 0.442 | 0.311 | 6.61594E-10 |
| RERGL_Cells | F10 | 1.12198E-12 | 0.523097 | 0.335 | 0.209 | 1.84275E-08 |
| Fibroblasts | KCNQ1OT1 | 2.42132E-40 | 0.523004 | 0.279 | 0.1 | 3.97678E-36 |
| Contractile | AC147651 | 2.23114E-32 | 0.522961 | 0.208 | 0.056 | 3.66443E-28 |
| RERGL_Cells | HRH2 | 3.25183E-36 | 0.522895 | 0.132 | 0.02 | 5.3408E-32 |
| Pericytes | CAPN2 | 1.06677E-16 | 0.522885 | 0.598 | 0.504 | 1.75207E-12 |
| RERGL_Cells | TLE1 | 2.81079E-09 | 0.522783 | 0.235 | 0.142 | 4.61644E-05 |
| Fibroblasts | PLSCR4 | 6.30788E-45 | 0.52276 | 0.477 | 0.225 | 1.03601E-40 |
| Fibroblasts | SCN7A | 1.49345E-43 | 0.522715 | 0.171 | 0.036 | 2.45285E-39 |
| Contractile | UBXN1 | 4.92363E-17 | 0.522575 | 0.645 | 0.498 | 8.08657E-13 |
| Intermediate | PFN2 | 1.28958E-28 | 0.522516 | 0.647 | 0.382 | 2.11801E-24 |
| Contractile | CAP2 | 1.09965E-24 | 0.522448 | 0.323 | 0.139 | 1.80606E-20 |
| Contractile | PART1 | 8.10953E-71 | 0.521646 | 0.172 | 0.013 | 1.33191E-66 |
| Fibroblasts | PALM | 5.45349E-81 | 0.52151 | 0.212 | 0.023 | 8.95681E-77 |
| Pericytes | ZEB2 | 4.24694E-13 | 0.521098 | 0.315 | 0.195 | 6.97517E-09 |
| Contractile | LINC00844 | 4.87629E-38 | 0.521 | 0.142 | 0.022 | 8.00882E-34 |
| Pericytes | TNC | 6.05143E-28 | 0.520474 | 0.109 | 0.019 | 9.93886E-24 |
| Intermediate | TRAPPC1 | 8.15179E-28 | 0.520248 | 0.675 | 0.42 | 1.33885E-23 |
| RERGL_Cells | AC104083 | 1.39013E-26 | 0.520112 | 0.156 | 0.038 | 2.28315E-22 |
| Fibroblasts | NFKBIZ | 1.05359E-44 | 0.520066 | 0.347 | 0.133 | 1.73042E-40 |
| Terminal | SSR4 | 7.86456E-20 | 0.519995 | 0.828 | 0.738 | 1.29168E-15 |
| Pericytes | CEBPB | 2.80317E-11 | 0.51998 | 0.724 | 0.695 | 4.60393E-07 |
| RERGL_Cells | H1FX | 5.33184E-09 | 0.519897 | 0.269 | 0.176 | 8.75701E-05 |
| Fibroblasts | SPTSSA | 3.46852E-30 | 0.519743 | 0.547 | 0.337 | 5.6967E-26 |
| Pericytes | EHD1 | 1.36314E-17 | 0.519695 | 0.245 | 0.117 | 2.23882E-13 |
| Fibroblasts | ABCC9 | 7.68803E-62 | 0.519664 | 0.244 | 0.051 | 1.26268E-57 |
| Fibroblasts | APLNR | 3.7611E-52 | 0.519649 | 0.134 | 0.014 | 6.17723E-48 |
| RERGL_Cells | PICALM | 3.91169E-09 | 0.51917 | 0.397 | 0.31 | 6.42456E-05 |
| Pericytes | PEA15 | 1.48536E-13 | 0.518944 | 0.451 | 0.344 | 2.43956E-09 |
| Fibroblasts | TMSB10 | 1.61175E-56 | 0.518646 | 0.978 | 0.926 | 2.64715E-52 |
| Intermediate | C9orf16 | 1.06727E-23 | 0.518522 | 0.574 | 0.345 | 1.75288E-19 |
| RERGL_Cells | PRKG1 | 4.80079E-10 | 0.518447 | 0.357 | 0.25 | 7.88482E-06 |
| Pericytes | NRP1 | 7.60809E-14 | 0.518194 | 0.336 | 0.213 | 1.24955E-09 |
| RERGL_Cells | SEM1 | 3.49276E-14 | 0.517954 | 0.692 | 0.641 | 5.73651E-10 |
| Pericytes | TUBB4B | 8.85055E-16 | 0.517797 | 0.64 | 0.537 | 1.45361E-11 |
| Fibroblasts | SULT1A1 | 1.62472E-76 | 0.517269 | 0.248 | 0.04 | 2.66844E-72 |
| Fibroblasts | NFYB | 2.9666E-44 | 0.516965 | 0.321 | 0.119 | 4.87234E-40 |
| Intermediate | TSTD1 | 2.39996E-31 | 0.51684 | 0.486 | 0.23 | 3.9417E-27 |
| Terminal | RCN3 | 1.55053E-14 | 0.516725 | 0.542 | 0.395 | 2.54659E-10 |
| RERGL_Cells | SNTA1 | 5.40219E-21 | 0.516721 | 0.212 | 0.079 | 8.87255E-17 |
| Contractile | GULP1 | 5.98685E-20 | 0.516666 | 0.401 | 0.223 | 9.8328E-16 |
| Contractile | ANP32B | 8.88404E-21 | 0.516477 | 0.595 | 0.404 | 1.45911E-16 |
| Pericytes | PLPP1 | 3.40752E-11 | 0.516475 | 0.558 | 0.479 | 5.59652E-07 |

| | | | | | | |
|---|---|---|---|---|---|---|
| Intermediate | ISCU | 9.09013E-33 | 0.516325 | 0.875 | 0.703 | 1.49296E-28 |
| Contractile | HSP90AB1 | 1.12241E-34 | 0.516299 | 0.954 | 0.913 | 1.84345E-30 |
| Contractile | TRIR | 7.18344E-24 | 0.515158 | 0.762 | 0.608 | 1.17981E-19 |
| Contractile | C4orf3 | 3.37508E-25 | 0.515024 | 0.822 | 0.704 | 5.54323E-21 |
| RERGL_Cells | WFDC1 | 4.92321E-13 | 0.514377 | 0.368 | 0.222 | 8.08588E-09 |
| Fibroblasts | NPR1 | 1.50162E-61 | 0.514351 | 0.264 | 0.063 | 2.46626E-57 |
| RERGL_Cells | DICER1-AS | 9.33179E-20 | 0.514307 | 0.135 | 0.038 | 1.53265E-15 |
| Terminal | AHNAK | 5.85499E-14 | 0.514216 | 0.534 | 0.396 | 9.61623E-10 |
| Fibroblasts | ADAM33 | 3.33216E-51 | 0.514152 | 0.239 | 0.061 | 5.47274E-47 |
| Intermediate | MFAP2 | 8.61385E-24 | 0.513548 | 0.267 | 0.109 | 1.41474E-19 |
| Contractile | ATP5MG | 1.98272E-30 | 0.513379 | 0.888 | 0.802 | 3.25643E-26 |
| Intermediate | HAPLN1 | 2.90638E-38 | 0.513219 | 0.254 | 0.071 | 4.77344E-34 |
| Intermediate | CSPG4 | 7.48936E-52 | 0.513207 | 0.249 | 0.051 | 1.23005E-47 |
| RERGL_Cells | TSPAN14 | 3.77614E-16 | 0.513117 | 0.182 | 0.074 | 6.20193E-12 |
| Fibroblasts | RARA | 1.2051E-41 | 0.513 | 0.397 | 0.176 | 1.97926E-37 |
| Intermediate | ENG | 1.3051E-12 | 0.512834 | 0.665 | 0.513 | 2.1435E-08 |
| Fibroblasts | TNFRSF1A | 1.37952E-49 | 0.512782 | 0.814 | 0.566 | 2.26572E-45 |
| Intermediate | SMOC1 | 1.39457E-17 | 0.51265 | 0.185 | 0.073 | 2.29044E-13 |
| Terminal | FGL2 | 1.49073E-11 | 0.512558 | 0.283 | 0.162 | 2.44837E-07 |
| Contractile | CDH2 | 1.58744E-23 | 0.512418 | 0.242 | 0.092 | 2.60722E-19 |
| Contractile | S1PR3 | 9.36402E-16 | 0.512301 | 0.261 | 0.127 | 1.53795E-11 |
| RERGL_Cells | COX7C | 2.38911E-36 | 0.512245 | 0.962 | 0.901 | 3.92387E-32 |
| Intermediate | ARHGAP18 | 1.83583E-57 | 0.511372 | 0.232 | 0.041 | 3.01516E-53 |
| Fibroblasts | SLC20A1 | 6.73333E-39 | 0.511245 | 0.212 | 0.064 | 1.10588E-34 |
| Terminal | AKR1C2 | 1.15767E-11 | 0.511179 | 0.281 | 0.162 | 1.90135E-07 |
| Pericytes | GJC1 | 4.30914E-33 | 0.511101 | 0.136 | 0.025 | 7.07734E-29 |
| Pericytes | PTMA | 3.70285E-48 | 0.510871 | 1 | 0.993 | 6.08156E-44 |
| Fibroblasts | XPNPEP2 | 2.89364E-94 | 0.510544 | 0.175 | 0.004 | 4.75251E-90 |
| Fibroblasts | GOLIM4 | 7.65968E-33 | 0.510472 | 0.535 | 0.308 | 1.25803E-28 |
| Terminal | DDB2 | 1.26378E-11 | 0.510369 | 0.229 | 0.12 | 2.07563E-07 |
| Fibroblasts | SLC16A2 | 3.34045E-73 | 0.510248 | 0.195 | 0.022 | 5.48635E-69 |
| Fibroblasts | NCOA7 | 1.94143E-54 | 0.509993 | 0.323 | 0.104 | 3.1886E-50 |
| Intermediate | COL14A1 | 7.91896E-23 | 0.509846 | 0.632 | 0.384 | 1.30061E-18 |
| Intermediate | CTSB | 3.98433E-20 | 0.509437 | 0.574 | 0.376 | 6.54387E-16 |
| RERGL_Cells | MT1E | 1.99341E-09 | 0.509427 | 0.739 | 0.673 | 3.27397E-05 |
| Contractile | SMS | 5.46319E-19 | 0.509316 | 0.445 | 0.271 | 8.97275E-15 |
| RERGL_Cells | CSRP1 | 8.05462E-24 | 0.509253 | 0.872 | 0.721 | 1.32289E-19 |
| Contractile | AQP1 | 6.13875E-39 | 0.509224 | 0.627 | 0.293 | 1.00823E-34 |
| Pericytes | PPP1R12A | 2.07704E-27 | 0.509057 | 0.707 | 0.487 | 3.41133E-23 |
| Contractile | DMKN | 1.74169E-23 | 0.50902 | 0.236 | 0.089 | 2.86056E-19 |
| Contractile | TBCA | 3.84217E-21 | 0.508944 | 0.766 | 0.629 | 6.31038E-17 |
| Fibroblasts | TCF4 | 1.6583E-36 | 0.5088 | 0.557 | 0.315 | 2.7236E-32 |
| RERGL_Cells | RBM24 | 2.64159E-43 | 0.50847 | 0.13 | 0.015 | 4.33854E-39 |
| RERGL_Cells | CAMTA1 | 9.1454E-12 | 0.508315 | 0.562 | 0.476 | 1.50204E-07 |
| RERGL_Cells | PPP1R1A | 4.53834E-27 | 0.507851 | 0.135 | 0.028 | 7.45377E-23 |
| Pericytes | NCKAP5 | 6.58757E-56 | 0.507568 | 0.127 | 0.009 | 1.08194E-51 |
| Pericytes | ADAMTS5 | 1.65389E-28 | 0.507557 | 0.138 | 0.031 | 2.71635E-24 |
| Contractile | ASPN | 1.48774E-18 | 0.50739 | 0.577 | 0.364 | 2.44347E-14 |
| Intermediate | LRPAP1 | 5.21876E-24 | 0.507334 | 0.785 | 0.599 | 8.5713E-20 |
| Pericytes | INKA2 | 1.28057E-34 | 0.506722 | 0.129 | 0.021 | 2.10321E-30 |
| Fibroblasts | TC2N | 1.90277E-66 | 0.506628 | 0.213 | 0.034 | 3.1251E-62 |
| Fibroblasts | ADA | 1.34708E-69 | 0.506583 | 0.188 | 0.022 | 2.21244E-65 |
| Intermediate | KCNMB1 | 6.10671E-36 | 0.506515 | 0.615 | 0.289 | 1.00297E-31 |
| RERGL_Cells | PHACTR2 | 1.48547E-08 | 0.505732 | 0.314 | 0.226 | 0.000243973 |
| Contractile | PFDN5 | 7.74006E-34 | 0.505619 | 0.956 | 0.877 | 1.27123E-29 |
| Contractile | HSPA2 | 1.42214E-19 | 0.505605 | 0.327 | 0.159 | 2.33572E-15 |

| | | | | | | |
|---|---|---|---|---|---|---|
| Intermediate | THY1 | 1.13625E-23 | 0.505429 | 0.282 | 0.115 | 1.86618E-19 |
| Fibroblasts | RTN4 | 1.18659E-57 | 0.505322 | 0.931 | 0.825 | 1.94886E-53 |
| RERGL_Cells | MYOM1 | 2.48989E-31 | 0.504906 | 0.147 | 0.029 | 4.08939E-27 |
| Fibroblasts | TRIB1 | 3.46977E-46 | 0.504836 | 0.178 | 0.037 | 5.69875E-42 |
| RERGL_Cells | ARHGEF10 | 2.25211E-23 | 0.504713 | 0.167 | 0.049 | 3.69887E-19 |
| Terminal | GDF10 | 1.3131E-17 | 0.504317 | 0.168 | 0.059 | 2.15663E-13 |
| RERGL_Cells | STBD1 | 5.5518E-12 | 0.504279 | 0.224 | 0.12 | 9.11828E-08 |
| Fibroblasts | TCN2 | 1.8209E-63 | 0.504076 | 0.236 | 0.048 | 2.99064E-59 |
| Fibroblasts | CCN2 | 0.000878549 | 0.50376 | 0.817 | 0.767 | 1 |
| Contractile | PPP2R2B | 4.50755E-37 | 0.503716 | 0.196 | 0.044 | 7.4032E-33 |
| Contractile | COX20 | 1.73518E-17 | 0.503695 | 0.451 | 0.279 | 2.84985E-13 |
| Intermediate | ISLR | 9.45256E-22 | 0.503439 | 0.45 | 0.248 | 1.55249E-17 |
| Intermediate | SHISA4 | 9.0905E-33 | 0.502892 | 0.383 | 0.156 | 1.49302E-28 |
| RERGL_Cells | KIF1C | 1.79719E-11 | 0.502782 | 0.218 | 0.118 | 2.95171E-07 |
| Intermediate | HSPB7 | 2.42436E-28 | 0.502462 | 0.437 | 0.198 | 3.98177E-24 |
| Contractile | DYNC1LI2 | 4.62781E-16 | 0.502248 | 0.379 | 0.226 | 7.60071E-12 |
| RERGL_Cells | KIAA0040 | 1.07213E-12 | 0.502093 | 0.19 | 0.09 | 1.76087E-08 |
| Contractile | EID1 | 2.8903E-41 | 0.502012 | 0.964 | 0.905 | 4.74702E-37 |
| Fibroblasts | CLMP | 1.79482E-83 | 0.501978 | 0.19 | 0.014 | 2.94781E-79 |
| Pericytes | PSMB9 | 3.56629E-12 | 0.501974 | 0.238 | 0.137 | 5.85727E-08 |
| Contractile | KCNK6 | 5.65689E-26 | 0.501848 | 0.23 | 0.08 | 9.29087E-22 |
| Terminal | KDELR3 | 1.42985E-15 | 0.501463 | 0.32 | 0.175 | 2.34839E-11 |
| Intermediate | SVIL | 6.94363E-25 | 0.501346 | 0.507 | 0.263 | 1.14042E-20 |
| Intermediate | SYNPO | 3.7559E-33 | 0.501218 | 0.333 | 0.123 | 6.16869E-29 |
| Contractile | DYNLL1 | 1.28718E-25 | 0.500654 | 0.886 | 0.833 | 2.11407E-21 |
| Intermediate | SERTAD4-A | 1.00869E-29 | 0.500606 | 0.336 | 0.133 | 1.65668E-25 |
| Pericytes | IFITM3 | 5.15976E-49 | 0.500416 | 0.993 | 0.991 | 8.4744E-45 |
| Contractile | CKB | 0.000130328 | 0.500324 | 0.419 | 0.357 | 1 |
| RERGL_Cells | SPEG | 9.51733E-14 | 0.500184 | 0.267 | 0.139 | 1.56313E-09 |
| Terminal | TIMP1 | 1.51621E-25 | 0.500102 | 0.972 | 0.921 | 2.49022E-21 |
| Intermediate | IL34 | 2.3297E-36 | 0.499507 | 0.26 | 0.078 | 3.82629E-32 |
| RERGL_Cells | NOP53 | 7.4276E-18 | 0.499322 | 0.806 | 0.736 | 1.21991E-13 |
| Contractile | MSRB2 | 5.55352E-17 | 0.499176 | 0.459 | 0.291 | 9.12109E-13 |
| Pericytes | BLOC1S1 | 1.38735E-12 | 0.499057 | 0.58 | 0.517 | 2.27858E-08 |
| Contractile | ESD | 1.04153E-27 | 0.498786 | 0.858 | 0.741 | 1.7106E-23 |
| Intermediate | CDKN2A | 9.96014E-47 | 0.49876 | 0.135 | 0.015 | 1.63585E-42 |
| Pericytes | MRPL32 | 3.23851E-10 | 0.498529 | 0.458 | 0.376 | 5.31893E-06 |
| Fibroblasts | TMBIM4 | 3.21435E-38 | 0.498368 | 0.802 | 0.593 | 5.27925E-34 |
| Intermediate | TSC22D1 | 5.42505E-30 | 0.498084 | 0.86 | 0.599 | 8.9101E-26 |
| Contractile | MAF1 | 2.16951E-19 | 0.497803 | 0.673 | 0.51 | 3.5632E-15 |
| Pericytes | ADGRL3 | 6.44036E-56 | 0.497639 | 0.131 | 0.01 | 1.05777E-51 |
| RERGL_Cells | UQCR10 | 1.7008E-14 | 0.497359 | 0.665 | 0.594 | 2.7934E-10 |
| Intermediate | MYO1D | 1.0054E-34 | 0.497241 | 0.308 | 0.106 | 1.65127E-30 |
| RERGL_Cells | MRPL33 | 1.91964E-14 | 0.497055 | 0.641 | 0.557 | 3.15281E-10 |
| Fibroblasts | HDAC7 | 6.23334E-54 | 0.496968 | 0.343 | 0.115 | 1.02376E-49 |
| Intermediate | PGF | 8.07183E-26 | 0.496767 | 0.45 | 0.222 | 1.32572E-21 |
| RERGL_Cells | PHLDB2 | 1.02044E-10 | 0.496629 | 0.276 | 0.167 | 1.67598E-06 |
| Fibroblasts | CERCAM | 7.83098E-49 | 0.496502 | 0.436 | 0.187 | 1.28616E-44 |
| Fibroblasts | CD55 | 5.55534E-27 | 0.496207 | 0.381 | 0.199 | 9.1241E-23 |
| Contractile | SNHG5 | 4.29784E-17 | 0.496194 | 0.762 | 0.642 | 7.05877E-13 |
| RERGL_Cells | KLHDC8B | 3.2975E-13 | 0.496135 | 0.241 | 0.128 | 5.41581E-09 |
| Fibroblasts | SURF4 | 1.09209E-35 | 0.496025 | 0.538 | 0.3 | 1.79365E-31 |
| Fibroblasts | MXRA8 | 6.92753E-47 | 0.495671 | 0.693 | 0.389 | 1.13778E-42 |
| Pericytes | CSRP1 | 1.20104E-28 | 0.495279 | 0.864 | 0.719 | 1.97258E-24 |
| RERGL_Cells | FAM122A | 7.7188E-09 | 0.495027 | 0.224 | 0.139 | 0.000126774 |
| Pericytes | VCP | 2.66132E-13 | 0.494897 | 0.584 | 0.513 | 4.37096E-09 |

| | | | | | | |
|---|---|---|---|---|---|---|
| Pericytes | ZYX | 5.21691E-14 | 0.494786 | 0.549 | 0.446 | 8.56826E-10 |
| RERGL_Cells | SMIM12 | 1.08069E-10 | 0.494509 | 0.357 | 0.248 | 1.77492E-06 |
| Terminal | MRPL51 | 8.99323E-15 | 0.494244 | 0.623 | 0.47 | 1.47705E-10 |
| Contractile | SKP1 | 4.9143E-44 | 0.494077 | 0.968 | 0.944 | 8.07125E-40 |
| Intermediate | PODNL1 | 2.76124E-54 | 0.493818 | 0.222 | 0.04 | 4.53506E-50 |
| Contractile | DDT | 2.43198E-16 | 0.493755 | 0.569 | 0.412 | 3.99429E-12 |
| RERGL_Cells | PALLD | 1.38784E-17 | 0.493351 | 0.688 | 0.551 | 2.2794E-13 |
| RERGL_Cells | FKBP4 | 1.46298E-11 | 0.493304 | 0.321 | 0.206 | 2.4028E-07 |
| Fibroblasts | SGK1 | 1.11717E-18 | 0.493135 | 0.181 | 0.08 | 1.83485E-14 |
| Pericytes | HSPA1B | 2.29824E-05 | 0.493086 | 0.235 | 0.17 | 0.37746331 |
| Intermediate | MASP1 | 1.21885E-41 | 0.492837 | 0.178 | 0.034 | 2.00183E-37 |
| RERGL_Cells | MAP1B | 5.44007E-22 | 0.492376 | 0.806 | 0.661 | 8.93477E-18 |
| RERGL_Cells | DACT3 | 9.64845E-15 | 0.491767 | 0.165 | 0.066 | 1.58466E-10 |
| Terminal | UACA | 1.85731E-12 | 0.491723 | 0.37 | 0.231 | 3.05044E-08 |
| Pericytes | CAVIN2 | 6.09484E-14 | 0.491716 | 0.196 | 0.094 | 1.00102E-09 |
| Intermediate | TUSC3 | 3.37656E-55 | 0.491488 | 0.256 | 0.052 | 5.54566E-51 |
| Intermediate | TM4SF1 | 8.17369E-78 | 0.491366 | 0.865 | 0.413 | 1.34245E-73 |
| Contractile | CSDE1 | 2.35171E-19 | 0.491345 | 0.774 | 0.647 | 3.86246E-15 |
| RERGL_Cells | OLFML2A | 8.85534E-26 | 0.490942 | 0.109 | 0.02 | 1.4544E-21 |
| Terminal | S100A6 | 1.25521E-42 | 0.490846 | 1 | 0.994 | 2.06155E-38 |
| Terminal | PDLIM3 | 1.01851E-24 | 0.490732 | 0.821 | 0.595 | 1.67279E-20 |
| Terminal | TTC3 | 5.82164E-13 | 0.490533 | 0.519 | 0.376 | 9.56146E-09 |
| Pericytes | CCT2 | 3.71626E-10 | 0.490473 | 0.445 | 0.359 | 6.10358E-06 |
| RERGL_Cells | SPECC1 | 4.22806E-09 | 0.490415 | 0.229 | 0.138 | 6.94417E-05 |
| Contractile | MAP3K20 | 8.67739E-23 | 0.490352 | 0.573 | 0.359 | 1.42517E-18 |
| Pericytes | DTWD1 | 4.63887E-10 | 0.490169 | 0.327 | 0.227 | 7.61888E-06 |
| Fibroblasts | COL15A1 | 8.64939E-36 | 0.489671 | 0.329 | 0.142 | 1.42058E-31 |
| RERGL_Cells | ACADVL | 2.896E-11 | 0.489534 | 0.652 | 0.588 | 4.75638E-07 |
| Contractile | NTRK3 | 1.45463E-21 | 0.489366 | 0.293 | 0.127 | 2.38908E-17 |
| Contractile | ATP5PO | 4.52706E-24 | 0.489243 | 0.794 | 0.678 | 7.43524E-20 |
| Terminal | TPT1 | 1.6451E-26 | 0.488628 | 0.989 | 0.982 | 2.70191E-22 |
| Fibroblasts | ADAMTSL4 | 1.2065E-64 | 0.488495 | 0.197 | 0.029 | 1.98156E-60 |
| Pericytes | MYO10 | 5.05252E-45 | 0.488473 | 0.133 | 0.016 | 8.29826E-41 |
| Fibroblasts | MYC | 1.09898E-91 | 0.488374 | 0.754 | 0.328 | 1.80497E-87 |
| Pericytes | MAP2K1 | 3.92659E-12 | 0.488368 | 0.267 | 0.159 | 6.44904E-08 |
| Contractile | NAP1L1 | 8.17449E-21 | 0.488312 | 0.796 | 0.656 | 1.34258E-16 |
| RERGL_Cells | CACNA1A | 4.03256E-47 | 0.488296 | 0.115 | 0.009 | 6.62307E-43 |
| Pericytes | UBE2E1 | 4.76299E-11 | 0.488207 | 0.398 | 0.296 | 7.82273E-07 |
| Contractile | PRKG1 | 1.13138E-15 | 0.488157 | 0.403 | 0.241 | 1.85817E-11 |
| Intermediate | SLC16A3 | 1.73991E-56 | 0.487644 | 0.193 | 0.027 | 2.85763E-52 |
| Intermediate | SUSD5 | 2.09857E-28 | 0.487482 | 0.29 | 0.107 | 3.44669E-24 |
| Contractile | SYNGR2 | 6.39308E-22 | 0.487462 | 0.457 | 0.247 | 1.05E-17 |
| Terminal | TMEM204 | 2.81431E-07 | 0.48746 | 0.364 | 0.272 | 0.004622217 |
| Fibroblasts | FOSL2 | 2.02269E-38 | 0.487436 | 0.362 | 0.156 | 3.32206E-34 |
| Fibroblasts | APP | 3.38125E-39 | 0.487433 | 0.809 | 0.587 | 5.55337E-35 |
| RERGL_Cells | ATP5MC3 | 2.6568E-13 | 0.487295 | 0.707 | 0.682 | 4.36353E-09 |
| RERGL_Cells | C12orf75 | 0.004039365 | 0.487278 | 0.479 | 0.449 | 1 |
| Contractile | UFC1 | 2.94667E-15 | 0.487064 | 0.583 | 0.443 | 4.83962E-11 |
| Terminal | SVIL | 1.82387E-12 | 0.486706 | 0.429 | 0.28 | 2.99552E-08 |
| RERGL_Cells | HNRNPM | 1.90143E-10 | 0.486531 | 0.598 | 0.552 | 3.12291E-06 |
| RERGL_Cells | TWSG1 | 5.8629E-08 | 0.485335 | 0.37 | 0.289 | 0.000962923 |
| Contractile | GYPC | 2.31432E-22 | 0.485225 | 0.611 | 0.39 | 3.80104E-18 |
| Intermediate | LOXL1 | 1.54192E-23 | 0.485185 | 0.493 | 0.273 | 2.53245E-19 |
| Pericytes | MAP1LC3A | 1.07056E-07 | 0.48492 | 0.485 | 0.42 | 0.001758287 |
| RERGL_Cells | TCF21 | 2.15956E-08 | 0.484774 | 0.427 | 0.33 | 0.000354686 |
| RERGL_Cells | FCMR | 2.15135E-68 | 0.484415 | 0.113 | 0.002 | 3.53338E-64 |

| | | | | | | |
|---|---|---|---|---|---|---|
| Pericytes | THOC7 | 1.05546E-09 | 0.484217 | 0.395 | 0.305 | 1.73349E-05 |
| Pericytes | CCT5 | 4.04493E-10 | 0.484208 | 0.44 | 0.355 | 6.6434E-06 |
| Fibroblasts | RPN1 | 9.50705E-41 | 0.483961 | 0.494 | 0.249 | 1.56144E-36 |
| Intermediate | NINJ1 | 7.8196E-24 | 0.483738 | 0.484 | 0.258 | 1.28429E-19 |
| Fibroblasts | MYCBP2 | 8.434E-37 | 0.483721 | 0.374 | 0.169 | 1.3852E-32 |
| RERGL_Cells | MRPL32 | 7.46166E-10 | 0.48368 | 0.459 | 0.378 | 1.2255E-05 |
| Contractile | ACAT1 | 7.02258E-18 | 0.483421 | 0.489 | 0.312 | 1.15339E-13 |
| Pericytes | EHD2 | 1.70752E-12 | 0.483317 | 0.48 | 0.364 | 2.80442E-08 |
| RERGL_Cells | BAG1 | 3.84498E-10 | 0.483264 | 0.444 | 0.364 | 6.315E-06 |
| Terminal | PKM | 1.47197E-20 | 0.483139 | 0.852 | 0.731 | 2.41756E-16 |
| Fibroblasts | ISM1 | 6.6066E-89 | 0.483042 | 0.154 | 0 | 1.08507E-84 |
| Fibroblasts | ANG | 2.09937E-41 | 0.482862 | 0.441 | 0.205 | 3.448E-37 |
| Contractile | ATP5F1A | 6.74342E-23 | 0.482858 | 0.768 | 0.623 | 1.10754E-18 |
| Pericytes | PRXL2C | 3.42219E-12 | 0.482804 | 0.275 | 0.165 | 5.62061E-08 |
| Pericytes | CFL1 | 6.71159E-35 | 0.482628 | 0.964 | 0.939 | 1.10231E-30 |
| Contractile | PTMS | 2.95961E-23 | 0.482384 | 0.876 | 0.749 | 4.86087E-19 |
| Pericytes | JAK1 | 1.58747E-09 | 0.48233 | 0.373 | 0.278 | 2.60727E-05 |
| Fibroblasts | ACAA2 | 3.50012E-55 | 0.482192 | 0.292 | 0.084 | 5.74859E-51 |
| Contractile | PIN1 | 1.36432E-17 | 0.482157 | 0.579 | 0.413 | 2.24076E-13 |
| Terminal | SARAF | 1.3397E-15 | 0.481484 | 0.773 | 0.69 | 2.20033E-11 |
| Terminal | ST13 | 3.67343E-11 | 0.481222 | 0.717 | 0.622 | 6.03324E-07 |
| Intermediate | COPRS | 3.64026E-22 | 0.480907 | 0.563 | 0.346 | 5.97877E-18 |
| RERGL_Cells | PGAM1 | 2.70268E-15 | 0.480701 | 0.66 | 0.585 | 4.43888E-11 |
| Pericytes | MCL1 | 1.25526E-11 | 0.480602 | 0.616 | 0.521 | 2.06163E-07 |
| RERGL_Cells | REEP1 | 2.16309E-20 | 0.480555 | 0.15 | 0.045 | 3.55265E-16 |
| Intermediate | UNC5B | 5.9972E-44 | 0.480357 | 0.215 | 0.046 | 9.84981E-40 |
| Pericytes | PDK4 | 1.04348E-07 | 0.480343 | 0.535 | 0.456 | 0.00171381 |
| RERGL_Cells | ACTB | 4.90509E-58 | 0.480309 | 1 | 1 | 8.05613E-54 |
| Contractile | YPEL5 | 1.26152E-14 | 0.480141 | 0.501 | 0.348 | 2.07192E-10 |
| Contractile | SLC25A3 | 1.39447E-28 | 0.480089 | 0.882 | 0.801 | 2.29028E-24 |
| Pericytes | TLE2 | 3.64141E-19 | 0.479417 | 0.173 | 0.064 | 5.98065E-15 |
| Pericytes | CLMN | 4.15536E-20 | 0.479263 | 0.18 | 0.065 | 6.82476E-16 |
| Pericytes | CAT | 1.97861E-11 | 0.479178 | 0.48 | 0.379 | 3.24966E-07 |
| Pericytes | PSMA3 | 9.70262E-15 | 0.478882 | 0.536 | 0.418 | 1.59356E-10 |
| Contractile | NOL3 | 1.57565E-13 | 0.478863 | 0.329 | 0.192 | 2.58786E-09 |
| Pericytes | EIF4A3 | 8.9916E-10 | 0.478617 | 0.389 | 0.3 | 1.47678E-05 |
| RERGL_Cells | RALY | 4.31046E-10 | 0.478399 | 0.496 | 0.416 | 7.0795E-06 |
| Contractile | MPC1 | 2.87113E-15 | 0.478352 | 0.521 | 0.369 | 4.71554E-11 |
| RERGL_Cells | NDUFA12 | 2.79758E-11 | 0.47817 | 0.536 | 0.454 | 4.59474E-07 |
| Contractile | CCDC90B | 2.92021E-14 | 0.478084 | 0.357 | 0.221 | 4.79616E-10 |
| Fibroblasts | DUSP6 | 1.43468E-26 | 0.478049 | 0.221 | 0.091 | 2.35632E-22 |
| Contractile | COX6C | 2.12199E-31 | 0.477912 | 0.92 | 0.836 | 3.48516E-27 |
| Pericytes | BTG1 | 3.83625E-10 | 0.477906 | 0.711 | 0.649 | 6.30066E-06 |
| RERGL_Cells | ID4 | 2.88199E-42 | 0.477705 | 0.793 | 0.533 | 4.73338E-38 |
| Contractile | CHMP3 | 2.04951E-14 | 0.477603 | 0.511 | 0.369 | 3.36612E-10 |
| Terminal | GPX4 | 3.70502E-22 | 0.477416 | 0.865 | 0.806 | 6.08513E-18 |
| Fibroblasts | GLT8D1 | 1.08535E-40 | 0.477216 | 0.394 | 0.176 | 1.78258E-36 |
| Intermediate | DYNLL1 | 8.60638E-29 | 0.477071 | 0.921 | 0.826 | 1.41351E-24 |
| Contractile | BTF3 | 1.06781E-34 | 0.476852 | 0.95 | 0.921 | 1.75376E-30 |
| Fibroblasts | CLIC2 | 4.31889E-74 | 0.47685 | 0.194 | 0.021 | 7.09334E-70 |
| RERGL_Cells | CALM3 | 5.21279E-13 | 0.476623 | 0.637 | 0.552 | 8.56149E-09 |
| Fibroblasts | BST1 | 1.25754E-58 | 0.476583 | 0.288 | 0.077 | 2.06539E-54 |
| Contractile | FOXS1 | 6.40455E-14 | 0.47648 | 0.293 | 0.157 | 1.05188E-09 |
| RERGL_Cells | HADHB | 7.77936E-10 | 0.4764 | 0.455 | 0.37 | 1.27768E-05 |
| Pericytes | EPAS1 | 4.65358E-15 | 0.476292 | 0.631 | 0.496 | 7.64304E-11 |
| Contractile | CALM2 | 2.1643E-42 | 0.476264 | 0.992 | 0.984 | 3.55465E-38 |

| | | | | | | |
|---|---|---|---|---|---|---|
| Pericytes | PPP2CA | 2.28407E-07 | 0.476249 | 0.342 | 0.268 | 0.003751353 |
| Contractile | KCNMA1 | 5.82957E-25 | 0.476086 | 0.196 | 0.062 | 9.57449E-21 |
| Intermediate | GABARAPL | 2.89896E-28 | 0.475917 | 0.879 | 0.716 | 4.76125E-24 |
| Terminal | CRIP2 | 1.86279E-22 | 0.475823 | 0.9 | 0.714 | 3.05945E-18 |
| RERGL_Cells | CCDC107 | 3.19723E-12 | 0.475571 | 0.613 | 0.527 | 5.25114E-08 |
| Pericytes | GRK5 | 3.21404E-11 | 0.475342 | 0.278 | 0.174 | 5.27874E-07 |
| Pericytes | B2M | 1.51254E-50 | 0.475316 | 1 | 0.999 | 2.4842E-46 |
| Fibroblasts | HTR2B | 3.42597E-46 | 0.475164 | 0.153 | 0.025 | 5.62681E-42 |
| Contractile | CCNG1 | 3.43834E-11 | 0.475131 | 0.445 | 0.319 | 5.64713E-07 |
| RERGL_Cells | ABTB1 | 1.52741E-11 | 0.475105 | 0.348 | 0.231 | 2.50862E-07 |
| Terminal | SNRPN | 1.70974E-09 | 0.475084 | 0.601 | 0.497 | 2.80808E-05 |
| Pericytes | TRIB2 | 4.51457E-12 | 0.475051 | 0.189 | 0.095 | 7.41474E-08 |
| Contractile | TOMM20 | 4.23958E-12 | 0.474966 | 0.519 | 0.392 | 6.96309E-08 |
| Contractile | NPNT | 2.62233E-16 | 0.474876 | 0.273 | 0.132 | 4.30691E-12 |
| Pericytes | NAMPT | 6.95237E-08 | 0.474826 | 0.464 | 0.366 | 0.001141858 |
| RERGL_Cells | FRY | 5.76882E-11 | 0.474775 | 0.209 | 0.111 | 9.4747E-07 |
| Intermediate | AKR1C2 | 5.26948E-20 | 0.474775 | 0.318 | 0.153 | 8.65459E-16 |
| Contractile | GYG1 | 1.70961E-17 | 0.474505 | 0.589 | 0.419 | 2.80786E-13 |
| Pericytes | SNRPB | 3.37596E-10 | 0.474416 | 0.52 | 0.446 | 5.54468E-06 |
| Pericytes | RAN | 3.21945E-15 | 0.474172 | 0.735 | 0.721 | 5.28763E-11 |
| RERGL_Cells | RAB13 | 1.68682E-08 | 0.474065 | 0.444 | 0.363 | 0.000277044 |
| Intermediate | ITPR1 | 2.73616E-25 | 0.473968 | 0.273 | 0.105 | 4.49386E-21 |
| Terminal | FXYD1 | 4.09515E-17 | 0.473921 | 0.88 | 0.787 | 6.72587E-13 |
| Contractile | PRMT2 | 5.53693E-14 | 0.473866 | 0.457 | 0.307 | 9.09385E-10 |
| Intermediate | THUMPD2 | 1.42024E-30 | 0.473821 | 0.31 | 0.117 | 2.33261E-26 |
| Contractile | PSMB6 | 1.7609E-21 | 0.473495 | 0.745 | 0.606 | 2.8921E-17 |
| Pericytes | DPY19L2 | 1.78461E-51 | 0.473395 | 0.113 | 0.007 | 2.93104E-47 |
| RERGL_Cells | TNS2 | 2.82513E-08 | 0.47337 | 0.359 | 0.271 | 0.000463999 |
| Pericytes | TPI1 | 1.10528E-21 | 0.473246 | 0.844 | 0.808 | 1.81531E-17 |
| RERGL_Cells | PPP1R12C | 5.19548E-17 | 0.473086 | 0.171 | 0.064 | 8.53306E-13 |
| Contractile | MAGED2 | 3.54274E-22 | 0.472909 | 0.78 | 0.616 | 5.8186E-18 |
| Fibroblasts | PID1 | 2.61235E-52 | 0.472597 | 0.295 | 0.091 | 4.29053E-48 |
| Contractile | LINC01615 | 7.12722E-21 | 0.472478 | 0.285 | 0.125 | 1.17057E-16 |
| Contractile | CAVIN1 | 3.4334E-30 | 0.471998 | 0.93 | 0.825 | 5.63901E-26 |
| Fibroblasts | LMAN1 | 1.34488E-35 | 0.471866 | 0.45 | 0.227 | 2.20883E-31 |
| RERGL_Cells | CDC42 | 1.95445E-13 | 0.471697 | 0.737 | 0.662 | 3.20998E-09 |
| Pericytes | WNT6 | 1.0713E-59 | 0.47142 | 0.124 | 0.007 | 1.75951E-55 |
| Terminal | EIF3H | 4.80308E-10 | 0.471214 | 0.686 | 0.61 | 7.88858E-06 |
| Terminal | HK1 | 1.56318E-12 | 0.471125 | 0.466 | 0.313 | 2.56736E-08 |
| Pericytes | RASL11A | 4.70729E-22 | 0.470363 | 0.171 | 0.056 | 7.73125E-18 |
| Terminal | LINC02269 | 4.96024E-09 | 0.470358 | 0.139 | 0.065 | 8.1467E-05 |
| Intermediate | ARHGDIB | 1.20605E-21 | 0.470304 | 0.682 | 0.48 | 1.98081E-17 |
| Terminal | OCIAD2 | 4.23073E-13 | 0.469971 | 0.2 | 0.093 | 6.94856E-09 |
| Terminal | LRPAP1 | 1.38513E-14 | 0.469751 | 0.73 | 0.612 | 2.27494E-10 |
| RERGL_Cells | SUN1 | 1.32169E-07 | 0.469632 | 0.303 | 0.219 | 0.002170749 |
| Intermediate | TMSB10 | 9.53803E-30 | 0.469521 | 0.978 | 0.935 | 1.56653E-25 |
| Pericytes | PUM3 | 1.20025E-13 | 0.468836 | 0.225 | 0.118 | 1.97128E-09 |
| Fibroblasts | BDH2 | 7.5331E-37 | 0.46866 | 0.52 | 0.28 | 1.23724E-32 |
| Contractile | CAV2 | 3.91743E-22 | 0.468306 | 0.731 | 0.519 | 6.43399E-18 |
| Pericytes | VGLL3 | 9.69159E-17 | 0.468077 | 0.173 | 0.07 | 1.59175E-12 |
| Pericytes | MGLL | 1.92462E-10 | 0.468011 | 0.445 | 0.345 | 3.16099E-06 |
| Pericytes | TMEM30B | 4.27884E-14 | 0.467986 | 0.178 | 0.081 | 7.02757E-10 |
| Contractile | MZT2B | 1.89E-15 | 0.467941 | 0.727 | 0.608 | 3.10414E-11 |
| Fibroblasts | IFNGR1 | 1.30106E-35 | 0.467932 | 0.513 | 0.273 | 2.13687E-31 |
| RERGL_Cells | NOL3 | 4.15274E-11 | 0.467896 | 0.314 | 0.196 | 6.82046E-07 |
| RERGL_Cells | FXYD6 | 0.000182121 | 0.467538 | 0.271 | 0.214 | 1 |

| | | | | | | |
|---|---|---|---|---|---|---|
| RERGL_Cells | SUMO2 | 7.25836E-25 | 0.467512 | 0.915 | 0.86 | 1.19211E-20 |
| Contractile | CCDC115 | 1.61246E-15 | 0.466783 | 0.419 | 0.26 | 2.6483E-11 |
| Terminal | KCNMA1 | 2.28943E-19 | 0.466726 | 0.185 | 0.065 | 3.76016E-15 |
| Terminal | TGFB1 | 1.16692E-14 | 0.466671 | 0.244 | 0.117 | 1.91655E-10 |
| Intermediate | GPX8 | 2.41608E-28 | 0.466617 | 0.348 | 0.147 | 3.96817E-24 |
| Terminal | CCNI | 1.40896E-19 | 0.466499 | 0.904 | 0.867 | 2.31408E-15 |
| Intermediate | MFAP4 | 1.08587E-15 | 0.466432 | 0.933 | 0.784 | 1.78344E-11 |
| RERGL_Cells | ANP32B | 6.30283E-11 | 0.466129 | 0.515 | 0.418 | 1.03518E-06 |
| Intermediate | MSC | 1.54052E-21 | 0.466059 | 0.404 | 0.208 | 2.53014E-17 |
| Terminal | VIM | 3.98015E-38 | 0.465671 | 1 | 0.998 | 6.537E-34 |
| RERGL_Cells | EEF1G | 1.40487E-28 | 0.465159 | 0.983 | 0.949 | 2.30736E-24 |
| Terminal | EPDR1 | 5.54481E-10 | 0.465148 | 0.294 | 0.183 | 9.10679E-06 |
| Contractile | LINC02762 | 3.00068E-25 | 0.465097 | 0.22 | 0.075 | 4.92832E-21 |
| Pericytes | ASXL1 | 1.59014E-10 | 0.464866 | 0.205 | 0.115 | 2.61165E-06 |
| RERGL_Cells | HEY2 | 1.95622E-10 | 0.464847 | 0.229 | 0.129 | 3.2129E-06 |
| Terminal | TSC22D1 | 5.12662E-17 | 0.464812 | 0.802 | 0.614 | 8.41996E-13 |
| Pericytes | PDGFRB | 6.0878E-09 | 0.464525 | 0.455 | 0.367 | 9.9986E-05 |
| Intermediate | UBE2E2 | 6.03758E-21 | 0.464437 | 0.516 | 0.304 | 9.91612E-17 |
| Intermediate | ITGAV | 1.12129E-25 | 0.464322 | 0.29 | 0.116 | 1.84161E-21 |
| Pericytes | FAM133B | 1.0187E-06 | 0.464081 | 0.369 | 0.302 | 0.016731145 |
| Pericytes | MAPK1IP1 | 1.00695E-09 | 0.463939 | 0.365 | 0.275 | 1.65381E-05 |
| Fibroblasts | LSAMP | 1.03598E-79 | 0.463697 | 0.144 | 0.002 | 1.70149E-75 |
| Pericytes | CFL2 | 5.97983E-10 | 0.463598 | 0.513 | 0.431 | 9.82128E-06 |
| Intermediate | PDGFA | 2.02814E-15 | 0.463286 | 0.292 | 0.146 | 3.33101E-11 |
| Fibroblasts | AGTRAP | 2.04918E-46 | 0.463086 | 0.328 | 0.12 | 3.36558E-42 |
| Fibroblasts | DDX21 | 2.86241E-37 | 0.462774 | 0.554 | 0.308 | 4.70122E-33 |
| Fibroblasts | NAV1 | 1.45757E-54 | 0.462495 | 0.207 | 0.042 | 2.39392E-50 |
| RERGL_Cells | PDCL3 | 1.48634E-08 | 0.462397 | 0.248 | 0.16 | 0.000244116 |
| Fibroblasts | ARL6IP5 | 5.25962E-42 | 0.462245 | 0.886 | 0.678 | 8.63839E-38 |
| Fibroblasts | TAGLN2 | 2.23968E-32 | 0.462215 | 0.862 | 0.689 | 3.67845E-28 |
| Fibroblasts | MMP23B | 4.61147E-35 | 0.462186 | 0.365 | 0.164 | 7.57388E-31 |
| Pericytes | COX7B | 2.80598E-17 | 0.462089 | 0.735 | 0.649 | 4.60853E-13 |
| RERGL_Cells | SELENOM | 2.2739E-26 | 0.462064 | 0.957 | 0.955 | 3.73466E-22 |
| Terminal | MAOB | 1.62064E-11 | 0.461785 | 0.366 | 0.225 | 2.66174E-07 |
| Fibroblasts | CHPF | 6.52495E-34 | 0.461657 | 0.504 | 0.273 | 1.07166E-29 |
| Fibroblasts | SSR1 | 7.47349E-35 | 0.461626 | 0.508 | 0.272 | 1.22745E-30 |
| Terminal | GAPDH | 1.09316E-22 | 0.461535 | 0.989 | 0.977 | 1.7954E-18 |
| Contractile | COMMD6 | 7.85522E-25 | 0.4611 | 0.874 | 0.765 | 1.29014E-20 |
| RERGL_Cells | NES | 2.35833E-14 | 0.46086 | 0.158 | 0.062 | 3.87332E-10 |
| Terminal | SLC41A3 | 6.88277E-14 | 0.460784 | 0.338 | 0.194 | 1.13043E-09 |
| RERGL_Cells | CHURC1 | 9.76354E-13 | 0.460438 | 0.65 | 0.579 | 1.60356E-08 |
| Contractile | NDUFB8 | 5.47288E-20 | 0.460363 | 0.796 | 0.688 | 8.98866E-16 |
| Terminal | UBA52 | 3.7006E-29 | 0.460262 | 0.978 | 0.959 | 6.07787E-25 |
| Terminal | PFDN5 | 1.87261E-22 | 0.459872 | 0.948 | 0.879 | 3.07558E-18 |
| Pericytes | DAAM2 | 1.40201E-16 | 0.459751 | 0.173 | 0.069 | 2.30266E-12 |
| Terminal | FAM114A1 | 1.3279E-11 | 0.459037 | 0.407 | 0.277 | 2.18095E-07 |
| Contractile | MUSTN1 | 2.88149E-41 | 0.458837 | 0.637 | 0.293 | 4.73256E-37 |
| Contractile | CHMP4A | 1.01148E-14 | 0.458573 | 0.473 | 0.316 | 1.66126E-10 |
| Fibroblasts | SRPRB | 2.74073E-40 | 0.458559 | 0.362 | 0.154 | 4.50137E-36 |
| Fibroblasts | AC008760 | 1.30205E-72 | 0.458502 | 0.128 | 0.001 | 2.13849E-68 |
| Contractile | NDUFB10 | 1.77037E-16 | 0.458319 | 0.705 | 0.577 | 2.90765E-12 |
| Pericytes | NUDT4 | 2.95623E-13 | 0.458266 | 0.565 | 0.448 | 4.85532E-09 |
| RERGL_Cells | NMNAT2 | 8.44553E-55 | 0.458239 | 0.113 | 0.006 | 1.38709E-50 |
| Pericytes | MBNL2 | 1.59504E-09 | 0.458185 | 0.305 | 0.207 | 2.61969E-05 |
| Intermediate | CCND3 | 6.02406E-21 | 0.457824 | 0.342 | 0.167 | 9.89392E-17 |
| Contractile | ATP5PF | 8.14169E-20 | 0.457716 | 0.826 | 0.731 | 1.33719E-15 |

| | | | | | | |
|---|---|---|---|---|---|---|
| Contractile | SYNM | 4.8455E-18 | 0.457554 | 0.192 | 0.074 | 7.95825E-14 |
| RERGL_Cells | IPO9 | 8.22442E-11 | 0.457221 | 0.235 | 0.133 | 1.35078E-06 |
| RERGL_Cells | ARPC5 | 1.9745E-17 | 0.456663 | 0.754 | 0.656 | 3.24291E-13 |
| Contractile | NEDD8 | 5.13868E-20 | 0.456639 | 0.798 | 0.677 | 8.43976E-16 |
| Fibroblasts | PLAU | 1.99211E-09 | 0.456399 | 0.173 | 0.101 | 3.27185E-05 |
| Fibroblasts | TMEM35B | 1.3363E-37 | 0.456325 | 0.588 | 0.329 | 2.19474E-33 |
| Intermediate | LMO7 | 1.41135E-36 | 0.45628 | 0.243 | 0.067 | 2.318E-32 |
| Intermediate | PPP1R3C | 9.65003E-24 | 0.455776 | 0.32 | 0.137 | 1.58492E-19 |
| Fibroblasts | ANGPTL2 | 1.76451E-34 | 0.45575 | 0.301 | 0.125 | 2.89804E-30 |
| Contractile | MRPS24 | 1.05295E-13 | 0.455715 | 0.587 | 0.457 | 1.72937E-09 |
| Contractile | BANF1 | 2.37197E-18 | 0.455576 | 0.723 | 0.556 | 3.89572E-14 |
| Intermediate | SLC29A1 | 1.89913E-16 | 0.455471 | 0.464 | 0.279 | 3.11913E-12 |
| Pericytes | GFPT1 | 1.46126E-08 | 0.455375 | 0.253 | 0.168 | 0.000239997 |
| RERGL_Cells | EIF4G2 | 1.76647E-17 | 0.455282 | 0.759 | 0.722 | 2.90125E-13 |
| RERGL_Cells | SLIRP | 5.48075E-12 | 0.455224 | 0.641 | 0.597 | 9.00158E-08 |
| Fibroblasts | C4orf48 | 3.98827E-46 | 0.455212 | 0.195 | 0.046 | 6.55033E-42 |
| RERGL_Cells | ALKBH7 | 2.83325E-09 | 0.454919 | 0.395 | 0.305 | 4.65334E-05 |
| Intermediate | FLNA | 8.33107E-28 | 0.454772 | 0.843 | 0.573 | 1.36829E-23 |
| Contractile | EIF3F | 4.84945E-19 | 0.45447 | 0.774 | 0.654 | 7.96474E-15 |
| Contractile | HNRNPA1 | 1.74886E-24 | 0.454223 | 0.94 | 0.873 | 2.87233E-20 |
| RERGL_Cells | H3F3A | 2.95989E-30 | 0.454103 | 0.951 | 0.942 | 4.86133E-26 |
| Contractile | ZNF385D | 1.23189E-24 | 0.453862 | 0.251 | 0.092 | 2.02326E-20 |
| Intermediate | TBXA2R | 5.75538E-41 | 0.453705 | 0.222 | 0.052 | 9.45264E-37 |
| RERGL_Cells | LHFPL6 | 3.29994E-17 | 0.453472 | 0.803 | 0.731 | 5.41982E-13 |
| Contractile | HYI | 3.61439E-15 | 0.453374 | 0.403 | 0.251 | 5.93628E-11 |
| Terminal | HLA-C | 3.62235E-16 | 0.453276 | 0.891 | 0.866 | 5.94935E-12 |
| Fibroblasts | APOL4 | 1.28907E-72 | 0.453247 | 0.162 | 0.01 | 2.11717E-68 |
| RERGL_Cells | FBXL3 | 8.13755E-09 | 0.453139 | 0.489 | 0.405 | 0.000133651 |
| Terminal | MXRA8 | 7.48606E-09 | 0.452988 | 0.56 | 0.468 | 0.000122951 |
| RERGL_Cells | LGI4 | 3.84783E-21 | 0.452924 | 0.412 | 0.21 | 6.31968E-17 |
| Fibroblasts | CRELD2 | 2.53095E-27 | 0.452784 | 0.414 | 0.226 | 4.15683E-23 |
| Contractile | PRDM16-D | 1.66521E-34 | 0.452503 | 0.172 | 0.037 | 2.73495E-30 |
| RERGL_Cells | ILK | 1.43243E-17 | 0.452493 | 0.746 | 0.654 | 2.35263E-13 |
| Contractile | IK | 2.26023E-14 | 0.451954 | 0.519 | 0.364 | 3.7122E-10 |
| Contractile | MRFAP1 | 1.64575E-20 | 0.451924 | 0.828 | 0.723 | 2.70299E-16 |
| Intermediate | COL16A1 | 2.93301E-20 | 0.451833 | 0.333 | 0.161 | 4.81717E-16 |
| Terminal | PMEPA1 | 2.84667E-13 | 0.451732 | 0.218 | 0.103 | 4.67538E-09 |
| Contractile | ATF4 | 1.07524E-15 | 0.451644 | 0.695 | 0.572 | 1.76597E-11 |
| Contractile | SMDT1 | 1.6359E-14 | 0.451397 | 0.607 | 0.473 | 2.68679E-10 |
| Contractile | AC093390 | 2.26173E-40 | 0.451306 | 0.154 | 0.025 | 3.71466E-36 |
| Terminal | LAMC3 | 3.15002E-40 | 0.451168 | 0.107 | 0.01 | 5.1736E-36 |
| Terminal | S100A16 | 3.92442E-10 | 0.451123 | 0.438 | 0.312 | 6.44547E-06 |
| Intermediate | SMC4 | 7.0685E-25 | 0.45104 | 0.314 | 0.133 | 1.16093E-20 |
| RERGL_Cells | NDUFB2 | 2.47963E-14 | 0.450768 | 0.72 | 0.671 | 4.07254E-10 |
| RERGL_Cells | CHRD | 3.26661E-08 | 0.450745 | 0.224 | 0.137 | 0.000536508 |
| Contractile | RASGRP2 | 1.49516E-17 | 0.450647 | 0.287 | 0.135 | 2.45565E-13 |
| Intermediate | PKIG | 2.90392E-20 | 0.45035 | 0.707 | 0.471 | 4.76941E-16 |
| Contractile | PNKD | 5.94254E-17 | 0.450348 | 0.517 | 0.339 | 9.76004E-13 |
| Fibroblasts | IL11RA | 2.19788E-34 | 0.450029 | 0.277 | 0.111 | 3.60979E-30 |
| Fibroblasts | CD302 | 2.73464E-30 | 0.449996 | 0.542 | 0.325 | 4.49137E-26 |
| Fibroblasts | SLC38A10 | 1.08488E-38 | 0.449915 | 0.374 | 0.164 | 1.78181E-34 |
| Pericytes | ANXA6 | 8.87125E-11 | 0.449887 | 0.616 | 0.573 | 1.45701E-06 |
| Pericytes | SRSF10 | 1.09648E-09 | 0.44977 | 0.409 | 0.315 | 1.80086E-05 |
| Terminal | S100A10 | 7.23205E-11 | 0.449625 | 0.758 | 0.635 | 1.18779E-06 |
| Fibroblasts | FST | 4.43119E-36 | 0.449049 | 0.219 | 0.071 | 7.27779E-32 |
| Terminal | ZFYVE21 | 5.17237E-10 | 0.448776 | 0.566 | 0.441 | 8.4951E-06 |

| | | | | | | |
|---|---|---|---|---|---|---|
| Pericytes | CALD1 | 5.82536E-37 | 0.448534 | 0.996 | 0.959 | 9.56757E-33 |
| Terminal | ADI1 | 1.25367E-09 | 0.448189 | 0.695 | 0.602 | 2.05903E-05 |
| Terminal | AKR1C3 | 4.52312E-12 | 0.448145 | 0.455 | 0.312 | 7.42877E-08 |
| Pericytes | TRAF4 | 6.86187E-22 | 0.447896 | 0.155 | 0.048 | 1.12699E-17 |
| Intermediate | CHPF | 2.26276E-15 | 0.447746 | 0.486 | 0.317 | 3.71636E-11 |
| Contractile | PACSIN3 | 3.37451E-25 | 0.447717 | 0.216 | 0.072 | 5.5423E-21 |
| Fibroblasts | SIRPA | 3.28489E-65 | 0.447636 | 0.191 | 0.026 | 5.39511E-61 |
| RERGL_Cells | MSRB1 | 3.78877E-07 | 0.447587 | 0.218 | 0.142 | 0.006222673 |
| Fibroblasts | PLCXD3 | 9.2616E-51 | 0.447485 | 0.161 | 0.025 | 1.52113E-46 |
| Fibroblasts | ANK2 | 1.24001E-46 | 0.447118 | 0.2 | 0.047 | 2.03659E-42 |
| RERGL_Cells | SLCO3A1 | 1.42962E-06 | 0.446959 | 0.205 | 0.133 | 0.023480041 |
| Terminal | NACA | 2.09039E-17 | 0.446815 | 0.969 | 0.938 | 3.43325E-13 |
| RERGL_Cells | NDUFA5 | 2.04343E-09 | 0.446804 | 0.549 | 0.496 | 3.35613E-05 |
| Pericytes | PDHB | 1.08287E-10 | 0.446746 | 0.445 | 0.345 | 1.7785E-06 |
| RERGL_Cells | SMDT1 | 6.50959E-09 | 0.446604 | 0.536 | 0.485 | 0.000106914 |
| Intermediate | SUN2 | 5.73305E-17 | 0.446573 | 0.649 | 0.439 | 9.41597E-13 |
| RERGL_Cells | SLC25A5 | 2.30138E-08 | 0.446546 | 0.571 | 0.517 | 0.000377978 |
| RERGL_Cells | TLN1 | 1.02652E-07 | 0.446337 | 0.434 | 0.361 | 0.001685953 |
| Intermediate | BEX3 | 1.51761E-21 | 0.44631 | 0.832 | 0.648 | 2.49253E-17 |
| RERGL_Cells | AP2M1 | 5.34305E-11 | 0.446167 | 0.684 | 0.65 | 8.77543E-07 |
| Contractile | ACTB | 1.70191E-56 | 0.44597 | 1 | 1 | 2.79522E-52 |
| Fibroblasts | IL15RA | 7.43679E-55 | 0.445846 | 0.204 | 0.04 | 1.22142E-50 |
| Terminal | SYNPO | 3.32516E-12 | 0.445502 | 0.259 | 0.139 | 5.46125E-08 |
| Fibroblasts | ARSA | 6.83225E-53 | 0.445265 | 0.269 | 0.075 | 1.12213E-48 |
| RERGL_Cells | COX6B1 | 7.55253E-19 | 0.445048 | 0.814 | 0.767 | 1.24043E-14 |
| Terminal | IL32 | 2.07004E-05 | 0.444693 | 0.407 | 0.321 | 0.339984168 |
| Pericytes | HOPX | 4.43872E-21 | 0.444655 | 0.113 | 0.028 | 7.29016E-17 |
| Contractile | GAS5 | 4.28243E-14 | 0.444355 | 0.776 | 0.677 | 7.03346E-10 |
| Intermediate | PCDH7 | 2.06032E-29 | 0.444325 | 0.224 | 0.069 | 3.38387E-25 |
| Fibroblasts | MYDGF | 7.83167E-33 | 0.444321 | 0.736 | 0.514 | 1.28627E-28 |
| RERGL_Cells | OPTN | 1.33539E-06 | 0.444277 | 0.397 | 0.334 | 0.021932461 |
| RERGL_Cells | ARHGEF7 | 1.09256E-13 | 0.444251 | 0.165 | 0.068 | 1.79442E-09 |
| Fibroblasts | TRIOBP | 2.17816E-41 | 0.443753 | 0.279 | 0.098 | 3.5774E-37 |
| Contractile | COX7A2L | 2.12697E-15 | 0.44369 | 0.659 | 0.527 | 3.49333E-11 |
| Contractile | METTL5 | 2.67999E-12 | 0.443462 | 0.403 | 0.264 | 4.40162E-08 |
| Pericytes | IFI16 | 1.87162E-07 | 0.443438 | 0.515 | 0.456 | 0.003073956 |
| RERGL_Cells | PCNP | 4.54059E-11 | 0.443102 | 0.656 | 0.609 | 7.45746E-07 |
| Terminal | CTSD | 2.72339E-13 | 0.442843 | 0.837 | 0.742 | 4.4729E-09 |
| Intermediate | BANF1 | 1.03687E-17 | 0.4428 | 0.714 | 0.556 | 1.70296E-13 |
| Intermediate | LRRC17 | 1.53205E-17 | 0.442745 | 0.387 | 0.209 | 2.51624E-13 |
| Pericytes | RBM7 | 5.4036E-11 | 0.442428 | 0.382 | 0.279 | 8.87487E-07 |
| Contractile | BRK1 | 2.5633E-22 | 0.442232 | 0.844 | 0.741 | 4.20996E-18 |
| Contractile | TMEM106 | 7.92792E-14 | 0.441733 | 0.405 | 0.257 | 1.30208E-09 |
| RERGL_Cells | SDHA | 3.06103E-08 | 0.441707 | 0.393 | 0.315 | 0.000502744 |
| Fibroblasts | PXDC1 | 3.87178E-38 | 0.44157 | 0.353 | 0.152 | 6.35902E-34 |
| Fibroblasts | SLC19A2 | 3.29653E-65 | 0.441284 | 0.163 | 0.015 | 5.41422E-61 |
| RERGL_Cells | ATP5MG | 2.09145E-18 | 0.4412 | 0.874 | 0.805 | 3.435E-14 |
| Contractile | TSC22D3 | 2.27104E-22 | 0.441106 | 0.752 | 0.546 | 3.72996E-18 |
| RERGL_Cells | FHL5 | 2.49458E-12 | 0.441035 | 0.265 | 0.141 | 4.09709E-08 |
| Intermediate | PPP1CB | 6.8753E-15 | 0.440989 | 0.802 | 0.635 | 1.1292E-10 |
| Pericytes | PRMT1 | 2.68168E-08 | 0.440926 | 0.502 | 0.444 | 0.000440439 |
| Fibroblasts | IFITM1 | 1.55767E-50 | 0.440892 | 0.273 | 0.077 | 2.55832E-46 |
| Fibroblasts | HLA-E | 2.85652E-37 | 0.440625 | 0.821 | 0.583 | 4.69155E-33 |
| Intermediate | SYT12 | 1.66738E-66 | 0.440606 | 0.153 | 0.011 | 2.7385E-62 |
| Contractile | CRTAP | 1.06062E-12 | 0.44045 | 0.533 | 0.392 | 1.74196E-08 |
| Fibroblasts | PTGES | 1.55735E-68 | 0.440439 | 0.18 | 0.019 | 2.55779E-64 |

| | | | | | | |
|---|---|---|---|---|---|---|
| Terminal | MATN2 | 2.27365E-06 | 0.440388 | 0.309 | 0.221 | 0.037342496 |
| Intermediate | HSPA8 | 3.00839E-17 | 0.440185 | 0.856 | 0.762 | 4.94098E-13 |
| Fibroblasts | CEBPB | 1.64322E-15 | 0.440116 | 0.783 | 0.663 | 2.69883E-11 |
| Pericytes | IRF1 | 1.80063E-09 | 0.440078 | 0.271 | 0.173 | 2.95735E-05 |
| RERGL_Cells | HIGD2A | 3.85949E-05 | 0.439928 | 0.376 | 0.336 | 0.633882163 |
| Pericytes | SMIM12 | 2.77221E-09 | 0.4398 | 0.345 | 0.248 | 4.55309E-05 |
| Fibroblasts | PGD | 8.19954E-51 | 0.439588 | 0.248 | 0.067 | 1.34669E-46 |
| Pericytes | SSB | 7.55328E-10 | 0.439174 | 0.638 | 0.595 | 1.24055E-05 |
| Pericytes | SH3GLB1 | 3.65525E-06 | 0.439062 | 0.385 | 0.331 | 0.060033783 |
| RERGL_Cells | BTF3 | 7.50582E-29 | 0.438548 | 0.964 | 0.92 | 1.23276E-24 |
| Intermediate | OMD | 9.86746E-21 | 0.438488 | 0.493 | 0.275 | 1.62063E-16 |
| Contractile | CHCHD2 | 8.00731E-29 | 0.438382 | 0.934 | 0.879 | 1.31512E-24 |
| RERGL_Cells | FUS | 4.6942E-11 | 0.438293 | 0.667 | 0.626 | 7.70976E-07 |
| Terminal | GOLIM4 | 9.57713E-07 | 0.438062 | 0.449 | 0.366 | 0.015729477 |
| RERGL_Cells | PIP4K2A | 1.72306E-08 | 0.438037 | 0.241 | 0.152 | 0.000282995 |
| Fibroblasts | SELENOS | 1.67395E-31 | 0.438014 | 0.741 | 0.506 | 2.7493E-27 |
| Intermediate | MYL12B | 3.73202E-30 | 0.43791 | 0.938 | 0.864 | 6.12948E-26 |
| Terminal | CHPF | 8.39837E-10 | 0.437227 | 0.444 | 0.328 | 1.37935E-05 |
| Pericytes | LINC0098 | 2.77525E-13 | 0.437118 | 0.127 | 0.049 | 4.55807E-09 |
| Pericytes | NBEAL1 | 1.17493E-06 | 0.437094 | 0.275 | 0.202 | 0.019297111 |
| Fibroblasts | IGFBP3 | 2.73685E-10 | 0.436973 | 0.154 | 0.084 | 4.495E-06 |
| RERGL_Cells | CAMK2G | 8.71928E-09 | 0.436803 | 0.265 | 0.168 | 0.000143206 |
| Pericytes | FUS | 6.23505E-14 | 0.436533 | 0.684 | 0.622 | 1.02404E-09 |
| Pericytes | ARF6 | 2.15454E-09 | 0.436405 | 0.442 | 0.348 | 3.53862E-05 |
| Terminal | GADD45A | 6.53224E-06 | 0.436315 | 0.368 | 0.287 | 0.107285441 |
| Fibroblasts | HSD17B11 | 1.72571E-35 | 0.43607 | 0.502 | 0.264 | 2.83431E-31 |
| RERGL_Cells | IDH3G | 3.62823E-07 | 0.435698 | 0.4 | 0.332 | 0.005958998 |
| Contractile | PLP2 | 3.44338E-18 | 0.435393 | 0.762 | 0.61 | 5.65541E-14 |
| Contractile | MTFR1L | 3.89295E-14 | 0.435337 | 0.337 | 0.197 | 6.39377E-10 |
| Pericytes | NFASC | 6.94945E-16 | 0.435285 | 0.178 | 0.074 | 1.14138E-11 |
| Contractile | EPB41L4A- | 8.77854E-14 | 0.435197 | 0.371 | 0.222 | 1.44179E-09 |
| Pericytes | TGFB1I1 | 2.36335E-15 | 0.434993 | 0.705 | 0.609 | 3.88157E-11 |
| Contractile | PEBP1 | 8.85436E-25 | 0.434699 | 0.92 | 0.867 | 1.45424E-20 |
| RERGL_Cells | PDE3A | 6.25977E-14 | 0.43455 | 0.162 | 0.066 | 1.0281E-09 |
| Contractile | AKAP1 | 5.83349E-23 | 0.434398 | 0.339 | 0.15 | 9.58092E-19 |
| Intermediate | TPD52L1 | 5.02726E-29 | 0.43435 | 0.28 | 0.1 | 8.25677E-25 |
| RERGL_Cells | STAT3 | 9.05738E-10 | 0.434199 | 0.526 | 0.441 | 1.48758E-05 |
| Fibroblasts | DNAJC3 | 2.11782E-32 | 0.434079 | 0.6 | 0.354 | 3.4783E-28 |
| Terminal | BRK1 | 1.12049E-16 | 0.433894 | 0.854 | 0.741 | 1.84029E-12 |
| Pericytes | RBM8A | 3.69453E-09 | 0.433726 | 0.545 | 0.481 | 6.0679E-05 |
| Contractile | PEF1 | 5.04261E-12 | 0.4337 | 0.433 | 0.301 | 8.28198E-08 |
| Terminal | TCEA3 | 1.83407E-09 | 0.433666 | 0.255 | 0.154 | 3.01228E-05 |
| RERGL_Cells | MDH2 | 2.43262E-08 | 0.433477 | 0.538 | 0.497 | 0.000399534 |
| Terminal | COMMD6 | 6.46054E-14 | 0.433404 | 0.837 | 0.771 | 1.06108E-09 |
| Pericytes | RASSF1 | 2.8357E-07 | 0.433361 | 0.255 | 0.178 | 0.004657355 |
| Pericytes | DDX39A | 1.06614E-10 | 0.433169 | 0.233 | 0.136 | 1.75103E-06 |
| Fibroblasts | CELF2 | 3.50198E-40 | 0.433014 | 0.365 | 0.153 | 5.75165E-36 |
| Fibroblasts | CHPT1 | 1.94316E-35 | 0.432704 | 0.491 | 0.256 | 3.19144E-31 |
| Pericytes | TLE4 | 2.29499E-10 | 0.432695 | 0.202 | 0.111 | 3.7693E-06 |
| RERGL_Cells | PPARG | 6.2453E-13 | 0.432681 | 0.137 | 0.052 | 1.02573E-08 |
| Pericytes | TMEM256 | 1.91278E-07 | 0.432585 | 0.438 | 0.38 | 0.003141547 |
| Contractile | UNC45B | 3.89421E-45 | 0.431994 | 0.124 | 0.012 | 6.39585E-41 |
| RERGL_Cells | ATF3 | 3.52168E-15 | 0.431893 | 0.472 | 0.292 | 5.78401E-11 |
| RERGL_Cells | ARHGAP17 | 3.84744E-13 | 0.43184 | 0.145 | 0.057 | 6.31903E-09 |
| Intermediate | GPRC5C | 2.11006E-24 | 0.431791 | 0.441 | 0.211 | 3.46556E-20 |
| Contractile | PERP | 8.2817E-17 | 0.431699 | 0.357 | 0.195 | 1.36019E-12 |

| | | | | | | |
|---|---|---|---|---|---|---|
| Terminal | PLP2 | 1.21963E-11 | 0.431677 | 0.712 | 0.619 | 2.00312E-07 |
| Contractile | MRPL20 | 3.92794E-13 | 0.431615 | 0.625 | 0.5 | 6.45125E-09 |
| Fibroblasts | HEXB | 1.26596E-34 | 0.431535 | 0.631 | 0.377 | 2.07921E-30 |
| Pericytes | SEPTIN4 | 6.88421E-09 | 0.431518 | 0.367 | 0.265 | 0.000113066 |
| Contractile | CIRBP | 1.10097E-21 | 0.431226 | 0.95 | 0.895 | 1.80823E-17 |
| Fibroblasts | NGFR | 6.48438E-26 | 0.431204 | 0.22 | 0.088 | 1.06499E-21 |
| Contractile | PMVK | 1.64535E-14 | 0.430746 | 0.473 | 0.316 | 2.70232E-10 |
| Contractile | PDAP1 | 1.43418E-12 | 0.430725 | 0.563 | 0.433 | 2.35549E-08 |
| Pericytes | MYH9 | 1.98247E-08 | 0.430682 | 0.478 | 0.389 | 0.000325602 |
| Contractile | HSF1 | 4.12026E-12 | 0.430469 | 0.407 | 0.272 | 6.76712E-08 |
| Intermediate | PDLIM7 | 1.74953E-18 | 0.430314 | 0.785 | 0.609 | 2.87342E-14 |
| Intermediate | MEGF6 | 1.61324E-32 | 0.430281 | 0.243 | 0.074 | 2.64959E-28 |
| RERGL_Cells | EGFLAM | 6.99522E-14 | 0.429901 | 0.182 | 0.077 | 1.14889E-09 |
| RERGL_Cells | LIN7B | 4.87921E-15 | 0.429859 | 0.141 | 0.05 | 8.01361E-11 |
| Pericytes | SERPINI1 | 1.27365E-11 | 0.429847 | 0.333 | 0.209 | 2.09185E-07 |
| RERGL_Cells | NDUFB11 | 1.70907E-09 | 0.429563 | 0.605 | 0.566 | 2.80698E-05 |
| Intermediate | F2R | 4.31656E-17 | 0.429533 | 0.232 | 0.103 | 7.08951E-13 |
| Fibroblasts | ODF3B | 1.65233E-49 | 0.429339 | 0.244 | 0.065 | 2.71379E-45 |
| Terminal | ISCU | 1.18609E-12 | 0.429313 | 0.802 | 0.717 | 1.94803E-08 |
| Contractile | EIF4B | 3.98759E-10 | 0.429238 | 0.553 | 0.457 | 6.54922E-06 |
| RERGL_Cells | UBE2D2 | 5.02531E-08 | 0.429016 | 0.517 | 0.476 | 0.000825357 |
| Fibroblasts | TAPBPL | 6.42782E-49 | 0.428938 | 0.264 | 0.076 | 1.0557E-44 |
| RERGL_Cells | SIVA1 | 3.98983E-12 | 0.428733 | 0.624 | 0.532 | 6.5529E-08 |
| Contractile | COX8A | 8.70214E-19 | 0.428578 | 0.788 | 0.662 | 1.42924E-14 |
| RERGL_Cells | OSBPL1A | 2.0185E-06 | 0.428555 | 0.261 | 0.189 | 0.033151803 |
| Pericytes | CNTN4 | 1.54507E-28 | 0.428539 | 0.136 | 0.03 | 2.53763E-24 |
| Terminal | MIR100HG | 2.66522E-13 | 0.428526 | 0.346 | 0.207 | 4.37736E-09 |
| Contractile | TRAPPC3 | 8.97978E-14 | 0.428411 | 0.457 | 0.31 | 1.47484E-09 |
| Pericytes | ENAH | 3.62898E-06 | 0.428222 | 0.275 | 0.202 | 0.059602298 |
| Pericytes | PDE1A | 4.38068E-07 | 0.428155 | 0.304 | 0.223 | 0.007194821 |
| Contractile | ZCRB1 | 3.33524E-12 | 0.427935 | 0.461 | 0.33 | 5.47779E-08 |
| RERGL_Cells | ANXA11 | 8.3002E-11 | 0.427736 | 0.618 | 0.565 | 1.36323E-06 |
| Fibroblasts | H19 | 2.85443E-15 | 0.427703 | 0.167 | 0.076 | 4.68812E-11 |
| Pericytes | PDCD10 | 1.81932E-06 | 0.42755 | 0.311 | 0.238 | 0.029880528 |
| Fibroblasts | IFI35 | 2.70005E-43 | 0.427232 | 0.337 | 0.129 | 4.43456E-39 |
| Pericytes | CYP26B1 | 1.37212E-14 | 0.426951 | 0.1 | 0.03 | 2.25357E-10 |
| Pericytes | SON | 1.5436E-20 | 0.426433 | 0.867 | 0.796 | 2.53521E-16 |
| Intermediate | AKR1C3 | 7.4393E-17 | 0.426403 | 0.49 | 0.302 | 1.22183E-12 |
| Fibroblasts | GABARAPL | 9.13009E-30 | 0.426217 | 0.428 | 0.227 | 1.49953E-25 |
| Pericytes | SLC7A5 | 9.20987E-33 | 0.426179 | 0.102 | 0.013 | 1.51263E-28 |
| RERGL_Cells | TTLL7 | 3.18883E-06 | 0.425264 | 0.308 | 0.234 | 0.052373292 |
| Intermediate | SLC9A3R2 | 1.09944E-21 | 0.425247 | 0.348 | 0.164 | 1.80571E-17 |
| Fibroblasts | ADAM17 | 8.14721E-38 | 0.425137 | 0.297 | 0.114 | 1.3381E-33 |
| Pericytes | LIN7A | 1.47685E-43 | 0.42511 | 0.109 | 0.01 | 2.42557E-39 |
| Pericytes | ATP5F1B | 2.84154E-22 | 0.425085 | 0.822 | 0.761 | 4.66695E-18 |
| Intermediate | C7orf50 | 3.24052E-21 | 0.424986 | 0.604 | 0.368 | 5.32222E-17 |
| RERGL_Cells | UQCRQ | 2.95197E-12 | 0.424937 | 0.756 | 0.736 | 4.84831E-08 |
| Terminal | SDC2 | 2.12647E-07 | 0.424846 | 0.702 | 0.624 | 0.003492518 |
| RERGL_Cells | THRA | 1.95111E-05 | 0.424788 | 0.246 | 0.18 | 0.320450777 |
| RERGL_Cells | NDUFB1 | 7.54854E-10 | 0.424694 | 0.66 | 0.645 | 1.23977E-05 |
| Fibroblasts | NTM | 1.21158E-62 | 0.42468 | 0.134 | 0.008 | 1.9899E-58 |
| Pericytes | GCA | 2.62456E-10 | 0.424544 | 0.204 | 0.114 | 4.31057E-06 |
| RERGL_Cells | PNMA1 | 7.78461E-06 | 0.424365 | 0.293 | 0.228 | 0.127854363 |
| Intermediate | XIST | 3.25497E-29 | 0.424202 | 0.221 | 0.066 | 5.34596E-25 |
| RERGL_Cells | OTUD6B-A | 2.14857E-05 | 0.424172 | 0.382 | 0.335 | 0.352881699 |
| Pericytes | PDE1B | 2.4909E-16 | 0.424089 | 0.14 | 0.049 | 4.09106E-12 |

| | | | | | | |
|---|---|---|---|---|---|---|
| Pericytes | WDR43 | 4.84182E-07 | 0.423993 | 0.216 | 0.144 | 0.007952204 |
| Intermediate | GPC1 | 3.43363E-21 | 0.423895 | 0.286 | 0.125 | 5.6394E-17 |
| RERGL_Cells | KLF6 | 3.59887E-07 | 0.423691 | 0.538 | 0.468 | 0.005910781 |
| RERGL_Cells | BRD2 | 0.000172702 | 0.423596 | 0.348 | 0.31 | 1 |
| Pericytes | GNL3 | 1.23504E-07 | 0.423589 | 0.327 | 0.248 | 0.002028432 |
| Pericytes | BCAR1 | 6.09403E-07 | 0.423388 | 0.255 | 0.18 | 0.01000884 |
| Intermediate | KCNK6 | 2.01664E-28 | 0.423137 | 0.236 | 0.077 | 3.31213E-24 |
| Fibroblasts | TCF21 | 3.41516E-31 | 0.423117 | 0.502 | 0.275 | 5.60906E-27 |
| Pericytes | NME1 | 6.4692E-06 | 0.423039 | 0.329 | 0.268 | 0.106250169 |
| Intermediate | PPDPF | 1.45737E-19 | 0.423035 | 0.961 | 0.887 | 2.39358E-15 |
| RERGL_Cells | TPPP3 | 1.22914E-08 | 0.422981 | 0.335 | 0.223 | 0.000201874 |
| Pericytes | HSPA9 | 1.19883E-07 | 0.422794 | 0.475 | 0.413 | 0.001968965 |
| Fibroblasts | LDHA | 8.00515E-36 | 0.422767 | 0.939 | 0.867 | 1.31477E-31 |
| Pericytes | LINC0140 | 1.75142E-18 | 0.422719 | 0.124 | 0.037 | 2.87653E-14 |
| RERGL_Cells | MZT2B | 1.12414E-12 | 0.422217 | 0.677 | 0.617 | 1.84629E-08 |
| Pericytes | CCDC126 | 7.8074E-12 | 0.421948 | 0.169 | 0.082 | 1.28229E-07 |
| RERGL_Cells | PPA2 | 5.79426E-05 | 0.42182 | 0.288 | 0.235 | 0.951648475 |
| RERGL_Cells | ACYP1 | 1.02822E-07 | 0.421819 | 0.154 | 0.084 | 0.001688749 |
| RERGL_Cells | MYOCD | 4.48469E-20 | 0.42179 | 0.143 | 0.041 | 7.36566E-16 |
| Pericytes | SNRPD1 | 4.26015E-08 | 0.421671 | 0.444 | 0.375 | 0.000699687 |
| Fibroblasts | IGSF10 | 7.40488E-58 | 0.421372 | 0.119 | 0.005 | 1.21618E-53 |
| Intermediate | COL13A1 | 1.76125E-50 | 0.421286 | 0.172 | 0.024 | 2.89268E-46 |
| Contractile | MRPL49 | 4.25273E-13 | 0.421159 | 0.275 | 0.15 | 6.98469E-09 |
| Terminal | C9orf16 | 5.68968E-11 | 0.421142 | 0.49 | 0.363 | 9.34473E-07 |
| Fibroblasts | AKR1B1 | 2.81654E-39 | 0.421056 | 0.429 | 0.198 | 4.62589E-35 |
| RERGL_Cells | ZYX | 5.63668E-08 | 0.420942 | 0.513 | 0.454 | 0.000925768 |
| Intermediate | MAP3K7CI | 6.62703E-28 | 0.420697 | 0.396 | 0.173 | 1.08842E-23 |
| Pericytes | PPP6C | 1.10843E-07 | 0.420582 | 0.315 | 0.234 | 0.001820487 |
| Terminal | STMN1 | 3.06275E-07 | 0.420496 | 0.22 | 0.135 | 0.005030264 |
| Contractile | RBPMS2 | 8.54945E-24 | 0.420493 | 0.277 | 0.107 | 1.40416E-19 |
| Pericytes | HRC | 4.9192E-17 | 0.420319 | 0.204 | 0.085 | 8.07929E-13 |
| Fibroblasts | F10 | 2.47731E-23 | 0.420075 | 0.34 | 0.177 | 4.06873E-19 |
| RERGL_Cells | NBEAL1 | 3.9287E-05 | 0.419996 | 0.267 | 0.205 | 0.645250478 |
| Terminal | RBP1 | 2.58557E-07 | 0.419932 | 0.224 | 0.14 | 0.004246534 |
| Fibroblasts | SPTBN1 | 6.59692E-27 | 0.419882 | 0.506 | 0.296 | 1.08348E-22 |
| RERGL_Cells | AMN1 | 8.38734E-10 | 0.419756 | 0.167 | 0.084 | 1.37754E-05 |
| RERGL_Cells | EIF3K | 4.98107E-14 | 0.419694 | 0.776 | 0.722 | 8.18092E-10 |
| RERGL_Cells | LYSMD2 | 1.15413E-06 | 0.419446 | 0.194 | 0.125 | 0.018955469 |
| Terminal | EEF1A1 | 8.03869E-24 | 0.41938 | 0.998 | 0.998 | 1.32028E-19 |
| Fibroblasts | TRIM8 | 1.74887E-34 | 0.419319 | 0.309 | 0.127 | 2.87234E-30 |
| Fibroblasts | SLC2A3 | 9.99838E-40 | 0.41912 | 0.31 | 0.116 | 1.64213E-35 |
| RERGL_Cells | PNRC1 | 1.18709E-16 | 0.418947 | 0.882 | 0.826 | 1.94968E-12 |
| Fibroblasts | NID1 | 8.84489E-39 | 0.418834 | 0.187 | 0.049 | 1.45269E-34 |
| RERGL_Cells | DEPP1 | 0.000330833 | 0.418798 | 0.476 | 0.413 | 1 |
| Pericytes | SMOC2 | 2.91745E-16 | 0.41819 | 0.698 | 0.553 | 4.79161E-12 |
| Terminal | XIST | 1.9285E-12 | 0.41797 | 0.174 | 0.077 | 3.16737E-08 |
| RERGL_Cells | CSKMT | 5.61452E-09 | 0.417818 | 0.188 | 0.104 | 9.2213E-05 |
| Contractile | NDUFB7 | 5.18063E-18 | 0.417701 | 0.786 | 0.649 | 8.50866E-14 |
| Contractile | UQCRH | 7.59398E-14 | 0.417611 | 0.709 | 0.605 | 1.24724E-09 |
| Terminal | GUK1 | 9.27387E-15 | 0.417474 | 0.867 | 0.792 | 1.52314E-10 |
| Contractile | ALDH3A2 | 1.11376E-06 | 0.417415 | 0.214 | 0.14 | 0.01829242 |
| Contractile | EIF3D | 2.60934E-14 | 0.417105 | 0.631 | 0.483 | 4.28558E-10 |
| Pericytes | RCAN1 | 2.06691E-06 | 0.417091 | 0.151 | 0.091 | 0.033946917 |
| Pericytes | DMD | 5.38566E-11 | 0.417073 | 0.211 | 0.115 | 8.84541E-07 |
| Contractile | EEF1B2 | 2.63127E-15 | 0.417043 | 0.862 | 0.788 | 4.3216E-11 |
| Contractile | AC068888 | 4.6222E-12 | 0.416958 | 0.357 | 0.224 | 7.59149E-08 |

| | | | | | | |
|---|---|---|---|---|---|---|
| Intermediate | CCND1 | 1.83323E-21 | 0.416726 | 0.385 | 0.191 | 3.01089E-17 |
| Terminal | TAGLN2 | 2.88148E-10 | 0.416705 | 0.784 | 0.734 | 4.73254E-06 |
| Fibroblasts | TMED10 | 5.74763E-30 | 0.416663 | 0.731 | 0.509 | 9.43991E-26 |
| RERGL_Cells | AURKAIP1 | 6.78689E-07 | 0.416364 | 0.502 | 0.463 | 0.011146794 |
| Terminal | GYPC | 1.6004E-08 | 0.41633 | 0.521 | 0.406 | 0.00026285 |
| Fibroblasts | LMAN2 | 1.21387E-32 | 0.416048 | 0.659 | 0.41 | 1.99366E-28 |
| Contractile | NSRP1 | 7.50835E-10 | 0.416046 | 0.491 | 0.368 | 1.23317E-05 |
| Fibroblasts | OSMR | 3.21843E-40 | 0.415987 | 0.269 | 0.092 | 5.28594E-36 |
| Pericytes | STX11 | 6.97386E-22 | 0.41568 | 0.125 | 0.033 | 1.14539E-17 |
| Contractile | UQCRC1 | 2.44303E-12 | 0.41558 | 0.485 | 0.35 | 4.01244E-08 |
| Pericytes | PNMA1 | 2.17236E-09 | 0.41553 | 0.322 | 0.221 | 3.56788E-05 |
| RERGL_Cells | SSBP2 | 3.42834E-06 | 0.41549 | 0.293 | 0.225 | 0.056307093 |
| Pericytes | PFKL | 1.0748E-07 | 0.415377 | 0.331 | 0.248 | 0.001765245 |
| RERGL_Cells | HDAC5 | 1.13414E-07 | 0.415243 | 0.276 | 0.19 | 0.001862704 |
| Pericytes | NAA50 | 2.28562E-08 | 0.415214 | 0.304 | 0.217 | 0.00037539 |
| Terminal | UNC5B | 2.24827E-16 | 0.415205 | 0.161 | 0.058 | 3.69256E-12 |
| Fibroblasts | AHI1 | 8.44603E-32 | 0.415127 | 0.347 | 0.162 | 1.38718E-27 |
| RERGL_Cells | NT5C3B | 6.53608E-05 | 0.415001 | 0.314 | 0.264 | 1 |
| Terminal | MSC | 3.71407E-07 | 0.414567 | 0.322 | 0.225 | 0.006099982 |
| Fibroblasts | SLIT3 | 3.13144E-30 | 0.414325 | 0.409 | 0.209 | 5.14307E-26 |
| Intermediate | MYL12A | 4.15113E-28 | 0.414297 | 0.966 | 0.91 | 6.81782E-24 |
| Fibroblasts | PBX1 | 5.82418E-35 | 0.414232 | 0.309 | 0.126 | 9.56564E-31 |
| Contractile | SNHG14 | 5.19585E-12 | 0.414199 | 0.242 | 0.129 | 8.53367E-08 |
| Fibroblasts | PDGFD | 3.35131E-37 | 0.414165 | 0.348 | 0.149 | 5.50419E-33 |
| Terminal | TSTD1 | 4.194E-10 | 0.413955 | 0.377 | 0.252 | 6.88823E-06 |
| RERGL_Cells | UBXN1 | 4.40498E-10 | 0.413828 | 0.583 | 0.509 | 7.23474E-06 |
| Pericytes | TCP1 | 7.16879E-09 | 0.41367 | 0.5 | 0.428 | 0.00011774 |
| Contractile | CALCOCO1 | 2.95625E-09 | 0.413657 | 0.261 | 0.159 | 4.85535E-05 |
| RERGL_Cells | RASSF3 | 1.86802E-08 | 0.413359 | 0.192 | 0.107 | 0.000306804 |
| RERGL_Cells | SYNGR2 | 1.47968E-08 | 0.413131 | 0.365 | 0.263 | 0.000243023 |
| Terminal | SCG2 | 1.34314E-09 | 0.413034 | 0.113 | 0.045 | 2.20598E-05 |
| Pericytes | BCL3 | 1.22157E-07 | 0.413 | 0.242 | 0.161 | 0.002006312 |
| RERGL_Cells | TCP1 | 1.06188E-08 | 0.412904 | 0.494 | 0.431 | 0.000174403 |
| RERGL_Cells | TBCA | 1.64887E-11 | 0.412886 | 0.69 | 0.641 | 2.70811E-07 |
| Contractile | FAM241B | 6.82956E-36 | 0.412813 | 0.142 | 0.024 | 1.12169E-31 |
| Fibroblasts | CHIC2 | 4.72203E-38 | 0.412802 | 0.278 | 0.103 | 7.75546E-34 |
| Intermediate | PCDH10 | 2.57348E-37 | 0.412779 | 0.189 | 0.041 | 4.22669E-33 |
| RERGL_Cells | CARHSP1 | 1.71977E-05 | 0.41264 | 0.397 | 0.347 | 0.282455426 |
| Contractile | KANK2 | 2.79982E-13 | 0.412454 | 0.497 | 0.336 | 4.59842E-09 |
| Contractile | CKMT2 | 1.50603E-25 | 0.412453 | 0.178 | 0.051 | 2.47351E-21 |
| Contractile | KALRN | 7.41328E-20 | 0.412236 | 0.19 | 0.069 | 1.21756E-15 |
| Intermediate | COL6A1 | 2.44193E-14 | 0.412026 | 0.86 | 0.722 | 4.01063E-10 |
| Contractile | PSMD4 | 8.19332E-12 | 0.411765 | 0.575 | 0.445 | 1.34567E-07 |
| Intermediate | PDGFC | 4.64214E-32 | 0.411685 | 0.23 | 0.066 | 7.62425E-28 |
| Pericytes | NOP56 | 2.65644E-08 | 0.411675 | 0.396 | 0.311 | 0.000436294 |
| Pericytes | BRD2 | 2.64791E-07 | 0.411343 | 0.382 | 0.303 | 0.004348925 |
| Fibroblasts | EMP3 | 1.07008E-21 | 0.411236 | 0.714 | 0.517 | 1.7575E-17 |
| Fibroblasts | RUNX1 | 2.98683E-42 | 0.411207 | 0.191 | 0.047 | 4.90557E-38 |
| Intermediate | BCAP29 | 2.57501E-16 | 0.411177 | 0.589 | 0.391 | 4.2292E-12 |
| Fibroblasts | CHID1 | 1.32239E-33 | 0.411082 | 0.51 | 0.274 | 2.1719E-29 |
| Terminal | ANXA2 | 1.11955E-17 | 0.410971 | 0.926 | 0.855 | 1.83874E-13 |
| Intermediate | HMGN3 | 1.60619E-14 | 0.410593 | 0.671 | 0.487 | 2.63801E-10 |
| RERGL_Cells | PYGM | 7.57148E-15 | 0.410476 | 0.139 | 0.049 | 1.24354E-10 |
| RERGL_Cells | MICOS10 | 3.845E-12 | 0.410338 | 0.69 | 0.653 | 6.31503E-08 |
| Fibroblasts | MEST | 2.77652E-30 | 0.410034 | 0.118 | 0.024 | 4.56016E-26 |
| Intermediate | VKORC1 | 9.37172E-18 | 0.409977 | 0.748 | 0.582 | 1.53921E-13 |

| | | | | | | |
|---|---|---|---|---|---|---|
| Intermediate | C12orf75 | 2.40304E-46 | 0.409898 | 0.815 | 0.389 | 3.94676E-42 |
| RERGL_Cells | N4BP2L1 | 6.28095E-07 | 0.409895 | 0.188 | 0.117 | 0.010315838 |
| RERGL_Cells | EEF1A1 | 4.69E-41 | 0.409688 | 1 | 0.997 | 7.70286E-37 |
| Contractile | DUSP3 | 3.05814E-13 | 0.409606 | 0.255 | 0.133 | 5.02269E-09 |
| RERGL_Cells | MOCS1 | 1.23688E-12 | 0.409564 | 0.145 | 0.059 | 2.03145E-08 |
| Pericytes | UBA2 | 5.5527E-15 | 0.409562 | 0.573 | 0.429 | 9.11975E-11 |
| Contractile | H2AFZ | 1.27796E-13 | 0.409295 | 0.689 | 0.564 | 2.09892E-09 |
| Contractile | UQCR11 | 2.57654E-20 | 0.409065 | 0.904 | 0.802 | 4.23171E-16 |
| RERGL_Cells | ACO2 | 3.64433E-05 | 0.409059 | 0.303 | 0.246 | 0.598544858 |
| Fibroblasts | CD81 | 2.02303E-19 | 0.408956 | 0.461 | 0.284 | 3.32262E-15 |
| Terminal | UQCRH | 2.91804E-11 | 0.408293 | 0.71 | 0.606 | 4.79259E-07 |
| RERGL_Cells | RAB20 | 9.58879E-09 | 0.408177 | 0.143 | 0.071 | 0.000157486 |
| Contractile | ATP1A2 | 5.26387E-19 | 0.407711 | 0.124 | 0.034 | 8.64539E-15 |
| Fibroblasts | DSE | 1.73649E-40 | 0.407272 | 0.288 | 0.102 | 2.85201E-36 |
| Pericytes | PSME2 | 1.40169E-07 | 0.407265 | 0.462 | 0.394 | 0.002302131 |
| Terminal | FTH1 | 7.3926E-25 | 0.40722 | 1 | 1 | 1.21416E-20 |
| Intermediate | SNAP23 | 1.26559E-15 | 0.407038 | 0.409 | 0.238 | 2.0786E-11 |
| Pericytes | SF1 | 2.09365E-06 | 0.406601 | 0.436 | 0.377 | 0.034386172 |
| Contractile | NDUFC1 | 3.2439E-15 | 0.406592 | 0.693 | 0.541 | 5.32778E-11 |
| Intermediate | RGS5 | 1.68457E-52 | 0.406589 | 0.832 | 0.404 | 2.76674E-48 |
| Fibroblasts | FZD4 | 6.96617E-39 | 0.406377 | 0.187 | 0.049 | 1.14412E-34 |
| Pericytes | SNRPB2 | 2.40897E-07 | 0.406284 | 0.578 | 0.549 | 0.003956488 |
| Contractile | SPRYD3 | 4.80756E-15 | 0.406106 | 0.295 | 0.157 | 7.89594E-11 |
| Contractile | VDAC2 | 1.53265E-18 | 0.405933 | 0.733 | 0.591 | 2.51722E-14 |
| Intermediate | ZNF385D | 6.29497E-22 | 0.405854 | 0.239 | 0.093 | 1.03389E-17 |
| Pericytes | EDNRA | 2.41003E-15 | 0.405545 | 0.225 | 0.103 | 3.95823E-11 |
| Intermediate | RCN2 | 5.39091E-18 | 0.405544 | 0.639 | 0.432 | 8.85403E-14 |
| Fibroblasts | MID1 | 7.58144E-54 | 0.405473 | 0.174 | 0.028 | 1.24518E-49 |
| Fibroblasts | EBF1 | 1.60752E-33 | 0.405411 | 0.556 | 0.31 | 2.64019E-29 |
| Fibroblasts | HM13 | 2.76464E-29 | 0.405322 | 0.562 | 0.336 | 4.54064E-25 |
| Intermediate | APLP2 | 9.46658E-18 | 0.405246 | 0.826 | 0.656 | 1.55479E-13 |
| Contractile | TPT1 | 2.33452E-38 | 0.405227 | 0.994 | 0.981 | 3.83421E-34 |
| RERGL_Cells | TRAF5 | 1.93599E-11 | 0.404946 | 0.162 | 0.074 | 3.17967E-07 |
| RERGL_Cells | CDS2 | 9.68871E-07 | 0.404896 | 0.256 | 0.177 | 0.015912743 |
| RERGL_Cells | EML2 | 2.46025E-09 | 0.404864 | 0.171 | 0.089 | 4.04072E-05 |
| Terminal | SCARA3 | 9.09521E-09 | 0.404751 | 0.233 | 0.138 | 0.00014938 |
| Contractile | ACTN2 | 3.29404E-46 | 0.404708 | 0.126 | 0.012 | 5.41013E-42 |
| RERGL_Cells | THOC2 | 3.06649E-06 | 0.404651 | 0.261 | 0.188 | 0.050364093 |
| Intermediate | SLC41A3 | 3.6986E-19 | 0.404609 | 0.363 | 0.186 | 6.07458E-15 |
| RERGL_Cells | CIRBP | 9.68185E-16 | 0.404296 | 0.934 | 0.898 | 1.59015E-11 |
| Contractile | EIF4H | 7.24639E-13 | 0.40388 | 0.629 | 0.483 | 1.19015E-08 |
| Intermediate | CUTA | 2.64138E-18 | 0.403756 | 0.751 | 0.568 | 4.3382E-14 |
| Pericytes | CSNK1D | 3.66287E-06 | 0.403644 | 0.305 | 0.238 | 0.060159056 |
| Contractile | C7orf50 | 9.02482E-12 | 0.40337 | 0.521 | 0.384 | 1.48224E-07 |
| Intermediate | RGCC | 1.02005E-08 | 0.403362 | 0.252 | 0.152 | 0.000167533 |
| Contractile | SRP14 | 2.37739E-33 | 0.40323 | 0.964 | 0.952 | 3.90463E-29 |
| Pericytes | BAZ1A | 2.37792E-08 | 0.403141 | 0.193 | 0.115 | 0.00039055 |
| Intermediate | S100A11 | 1.1259E-23 | 0.40311 | 0.966 | 0.906 | 1.84918E-19 |
| Terminal | PARK7 | 4.14545E-12 | 0.403021 | 0.871 | 0.802 | 6.80849E-08 |
| Terminal | SAT1 | 0.000103798 | 0.402868 | 0.612 | 0.55 | 1 |
| Contractile | PRPSAP1 | 7.70222E-10 | 0.402847 | 0.259 | 0.156 | 1.26501E-05 |
| Pericytes | NXT1 | 8.83725E-06 | 0.402756 | 0.271 | 0.208 | 0.145142989 |
| Pericytes | NOLC1 | 1.35685E-08 | 0.402742 | 0.224 | 0.14 | 0.000222849 |
| Terminal | CCPG1 | 2.59561E-07 | 0.402314 | 0.481 | 0.401 | 0.004263027 |
| Contractile | TRAPPC1 | 3.92013E-13 | 0.402298 | 0.581 | 0.438 | 6.43842E-09 |
| Fibroblasts | CTSL | 3.01944E-24 | 0.402203 | 0.658 | 0.459 | 4.95912E-20 |

| | | | | | | |
|---|---|---|---|---|---|---|
| RERGL_Cells | LINC00702 | 7.08296E-07 | 0.402187 | 0.179 | 0.108 | 0.011633059 |
| Fibroblasts | PNRC1 | 1.35974E-21 | 0.401603 | 0.9 | 0.805 | 2.23324E-17 |
| Pericytes | NBL1 | 5.92948E-10 | 0.40149 | 0.622 | 0.549 | 9.73858E-06 |
| Intermediate | HMCN1 | 1.93133E-40 | 0.401435 | 0.176 | 0.033 | 3.17201E-36 |
| Contractile | TXNRD1 | 1.77626E-06 | 0.401242 | 0.325 | 0.24 | 0.029173357 |
| Fibroblasts | SNX9 | 5.19111E-33 | 0.401231 | 0.433 | 0.223 | 8.52587E-29 |
| Contractile | MRPL21 | 3.2918E-10 | 0.401171 | 0.393 | 0.268 | 5.40646E-06 |
| RERGL_Cells | MTRNR2L1 | 3.05252E-06 | 0.401031 | 0.237 | 0.163 | 0.050134565 |
| Fibroblasts | FYN | 1.07187E-60 | 0.400674 | 0.195 | 0.031 | 1.76044E-56 |
| Fibroblasts | EFEMP2 | 1.35135E-28 | 0.40045 | 0.674 | 0.434 | 2.21946E-24 |
| Fibroblasts | PTP4A2 | 3.61639E-32 | 0.400432 | 0.487 | 0.262 | 5.93956E-28 |
| Contractile | AMIGO2 | 8.1106E-14 | 0.400369 | 0.178 | 0.077 | 1.33208E-09 |
| Contractile | PDLIM1 | 2.21926E-07 | 0.400218 | 0.631 | 0.558 | 0.003644911 |
| Intermediate | GMPR | 7.21696E-24 | 0.400115 | 0.273 | 0.108 | 1.18531E-19 |
| Fibroblasts | AKR1C3 | 1.70677E-26 | 0.400104 | 0.471 | 0.27 | 2.8032E-22 |
| Terminal | PLEKHA1 | 1.83037E-08 | 0.40009 | 0.22 | 0.13 | 0.00030062 |
| Fibroblasts | RAB3IL1 | 5.84055E-56 | 0.399991 | 0.171 | 0.025 | 9.59252E-52 |
| Intermediate | KALRN | 5.09582E-30 | 0.399965 | 0.215 | 0.063 | 8.36938E-26 |
| Intermediate | KCNS3 | 1.38898E-31 | 0.399811 | 0.178 | 0.044 | 2.28126E-27 |
| Intermediate | P4HB | 1.67148E-15 | 0.399807 | 0.834 | 0.713 | 2.74524E-11 |
| Contractile | LITAF | 3.86421E-12 | 0.399757 | 0.677 | 0.562 | 6.34658E-08 |
| Pericytes | NOP58 | 3.72249E-07 | 0.399692 | 0.305 | 0.226 | 0.006113818 |
| Contractile | PSMC3 | 1.82342E-12 | 0.399615 | 0.603 | 0.471 | 2.99479E-08 |
| RERGL_Cells | CHCHD2 | 6.07889E-21 | 0.399554 | 0.912 | 0.883 | 9.98396E-17 |
| Contractile | RNF7 | 4.40875E-12 | 0.399522 | 0.581 | 0.442 | 7.24093E-08 |
| Terminal | DDIT4 | 2.57734E-05 | 0.399446 | 0.346 | 0.262 | 0.423302972 |
| RERGL_Cells | ZNF22 | 3.73355E-05 | 0.399388 | 0.269 | 0.209 | 0.613198575 |
| RERGL_Cells | PSME1 | 8.56177E-13 | 0.399359 | 0.716 | 0.641 | 1.40619E-08 |
| Pericytes | DACT1 | 0.001052058 | 0.399248 | 0.151 | 0.111 | 1 |
| Pericytes | LRRC32 | 4.28409E-09 | 0.399162 | 0.284 | 0.183 | 7.03619E-05 |
| Terminal | ANP32B | 9.05305E-07 | 0.398987 | 0.503 | 0.42 | 0.014868734 |
| RERGL_Cells | SMARCB1 | 5.09363E-05 | 0.398822 | 0.306 | 0.25 | 0.836577867 |
| Contractile | EIF2S3 | 1.39441E-10 | 0.398769 | 0.469 | 0.341 | 2.29018E-06 |
| Intermediate | PDLIM2 | 4.22898E-15 | 0.398606 | 0.606 | 0.436 | 6.94568E-11 |
| Intermediate | SORT1 | 3.24255E-28 | 0.398456 | 0.305 | 0.113 | 5.32556E-24 |
| Terminal | ANXA4 | 9.01171E-08 | 0.398393 | 0.505 | 0.425 | 0.001480083 |
| Fibroblasts | TMEM9B | 3.041E-33 | 0.398184 | 0.433 | 0.218 | 4.99454E-29 |
| Fibroblasts | SMAD3 | 3.92612E-33 | 0.398164 | 0.181 | 0.054 | 6.44827E-29 |
| Contractile | C1orf198 | 4.19005E-13 | 0.398139 | 0.441 | 0.287 | 6.88174E-09 |
| Intermediate | VIM | 2.65329E-49 | 0.398112 | 1 | 0.998 | 4.35776E-45 |
| RERGL_Cells | HMGB1 | 4.86391E-21 | 0.398057 | 0.942 | 0.917 | 7.98848E-17 |
| Contractile | CTPS1 | 4.11853E-15 | 0.397847 | 0.265 | 0.131 | 6.76427E-11 |
| Contractile | HSPA1A | 7.2901E-15 | 0.397814 | 0.465 | 0.307 | 1.19733E-10 |
| Pericytes | CALM2 | 6.27719E-33 | 0.397797 | 0.995 | 0.983 | 1.03096E-28 |
| Contractile | GLIPR1 | 1.83945E-08 | 0.397505 | 0.441 | 0.334 | 0.000302111 |
| Fibroblasts | FEZ1 | 1.63817E-24 | 0.397421 | 0.481 | 0.284 | 2.69053E-20 |
| Contractile | PPP1R16A | 9.29949E-13 | 0.397374 | 0.202 | 0.098 | 1.52735E-08 |
| Terminal | ARL6IP5 | 5.14786E-13 | 0.397348 | 0.824 | 0.728 | 8.45484E-09 |
| Pericytes | ARHGDIA | 3.19373E-09 | 0.397288 | 0.589 | 0.536 | 5.24538E-05 |
| RERGL_Cells | CBX7 | 7.80075E-06 | 0.396907 | 0.214 | 0.147 | 0.128119585 |
| RERGL_Cells | EEF2 | 8.80639E-20 | 0.396783 | 0.953 | 0.945 | 1.44636E-15 |
| RERGL_Cells | FOXC2 | 1.16644E-06 | 0.396687 | 0.194 | 0.123 | 0.019157601 |
| RERGL_Cells | CACNA1H | 1.08335E-20 | 0.396664 | 0.111 | 0.025 | 1.7793E-16 |
| Terminal | PPP1R3C | 4.85181E-09 | 0.396606 | 0.257 | 0.151 | 7.96861E-05 |
| Fibroblasts | OLFML1 | 4.41351E-42 | 0.396585 | 0.258 | 0.083 | 7.24875E-38 |
| Terminal | IGFBP6 | 3.72956E-11 | 0.396462 | 0.898 | 0.791 | 6.12543E-07 |

| | | | | | | |
|---|---|---|---|---|---|---|
| Pericytes | GLIPR2 | 4.30197E-09 | 0.396353 | 0.273 | 0.179 | 7.06556E-05 |
| Contractile | RTF2 | 1.47194E-11 | 0.396326 | 0.541 | 0.416 | 2.41752E-07 |
| RERGL_Cells | GNAI1 | 5.85217E-08 | 0.396275 | 0.199 | 0.116 | 0.000961161 |
| Contractile | STUB1 | 4.72695E-12 | 0.396142 | 0.615 | 0.472 | 7.76354E-08 |
| Pericytes | PPIA | 3.82965E-18 | 0.396138 | 0.898 | 0.867 | 6.28981E-14 |
| Contractile | RRAD | 2.31959E-17 | 0.396021 | 0.236 | 0.1 | 3.8097E-13 |
| Contractile | SNHG29 | 9.55899E-18 | 0.395911 | 0.904 | 0.829 | 1.56997E-13 |
| RERGL_Cells | PRXL2A | 1.43575E-08 | 0.395848 | 0.254 | 0.159 | 0.000235808 |
| RERGL_Cells | SNRPB | 3.88075E-07 | 0.39579 | 0.496 | 0.451 | 0.006373736 |
| Contractile | SARAF | 1.05937E-14 | 0.395729 | 0.792 | 0.686 | 1.7399E-10 |
| RERGL_Cells | PSMD4 | 6.65215E-06 | 0.395714 | 0.487 | 0.46 | 0.109254874 |
| Contractile | MPP7 | 1.07563E-35 | 0.395703 | 0.122 | 0.017 | 1.76661E-31 |
| Fibroblasts | TXNRD1 | 1.33798E-26 | 0.395386 | 0.377 | 0.198 | 2.1975E-22 |
| Pericytes | ZFAND5 | 2.7188E-06 | 0.395326 | 0.433 | 0.376 | 0.044653639 |
| Intermediate | CLEC18A | 1.73711E-47 | 0.395305 | 0.15 | 0.019 | 2.85302E-43 |
| Pericytes | MOB4 | 6.74924E-09 | 0.395291 | 0.409 | 0.321 | 0.00011085 |
| Terminal | CIB1 | 3.30478E-07 | 0.395262 | 0.505 | 0.426 | 0.005427764 |
| Pericytes | SPECC1 | 1.16523E-10 | 0.395165 | 0.235 | 0.134 | 1.91377E-06 |
| Contractile | COPS9 | 4.49478E-15 | 0.395055 | 0.689 | 0.568 | 7.38223E-11 |
| Fibroblasts | NPDC1 | 8.88346E-29 | 0.39497 | 0.56 | 0.337 | 1.45902E-24 |
| Contractile | FMC1 | 1.64615E-08 | 0.39494 | 0.355 | 0.251 | 0.000270364 |
| Contractile | ZMAT2 | 1.0676E-10 | 0.394915 | 0.399 | 0.274 | 1.75343E-06 |
| Fibroblasts | ECHDC2 | 4.69483E-28 | 0.394812 | 0.378 | 0.193 | 7.71079E-24 |
| Fibroblasts | SERPINB1 | 3.75385E-34 | 0.394687 | 0.438 | 0.217 | 6.16532E-30 |
| Intermediate | ITGBL1 | 4.34626E-08 | 0.394651 | 0.292 | 0.192 | 0.00071383 |
| Intermediate | TPM4 | 1.98323E-19 | 0.394148 | 0.869 | 0.727 | 3.25725E-15 |
| RERGL_Cells | C1orf54 | 1.28324E-05 | 0.394059 | 0.333 | 0.272 | 0.210759169 |
| Pericytes | DDAH1 | 2.00368E-12 | 0.393847 | 0.18 | 0.087 | 3.29085E-08 |
| RERGL_Cells | WBP11 | 5.95891E-07 | 0.393776 | 0.318 | 0.239 | 0.009786909 |
| Contractile | FIS1 | 1.91052E-14 | 0.393742 | 0.747 | 0.617 | 3.13784E-10 |
| Pericytes | UBE2N | 1.53386E-08 | 0.393727 | 0.513 | 0.451 | 0.000251922 |
| Terminal | NUCKS1 | 5.54898E-11 | 0.393671 | 0.786 | 0.696 | 9.11364E-07 |
| RERGL_Cells | FRMD4A | 7.17391E-11 | 0.393657 | 0.154 | 0.069 | 1.17824E-06 |
| RERGL_Cells | KLF2 | 0.002251502 | 0.393273 | 0.31 | 0.263 | 1 |
| Contractile | THRA | 5.36258E-12 | 0.393064 | 0.297 | 0.171 | 8.80749E-08 |
| Fibroblasts | DRAM1 | 1.66086E-49 | 0.392719 | 0.18 | 0.034 | 2.72779E-45 |
| Fibroblasts | CALU | 1.67717E-24 | 0.392582 | 0.677 | 0.472 | 2.75458E-20 |
| Pericytes | TEAD3 | 3.85329E-08 | 0.392438 | 0.218 | 0.137 | 0.000632865 |
| RERGL_Cells | RBPMS | 1.06115E-08 | 0.392391 | 0.496 | 0.395 | 0.000174283 |
| Pericytes | HNRNPU | 1.6932E-10 | 0.392234 | 0.644 | 0.592 | 2.78091E-06 |
| Fibroblasts | HLA-DPA1 | 6.02057E-25 | 0.391497 | 0.331 | 0.168 | 9.88818E-21 |
| Pericytes | TSPAN14 | 1.4079E-15 | 0.390993 | 0.175 | 0.072 | 2.31233E-11 |
| Intermediate | HEYL | 6.06044E-23 | 0.390939 | 0.445 | 0.223 | 9.95366E-19 |
| Fibroblasts | GUSB | 2.65693E-36 | 0.390753 | 0.356 | 0.154 | 4.36375E-32 |
| RERGL_Cells | NDUFV3 | 1.43028E-05 | 0.390748 | 0.301 | 0.241 | 0.234908501 |
| Contractile | NDUFAB1 | 3.43305E-12 | 0.390643 | 0.593 | 0.48 | 5.63844E-08 |
| Contractile | ADH5 | 1.13569E-12 | 0.390258 | 0.663 | 0.534 | 1.86526E-08 |
| Intermediate | HLA-A | 2.16929E-22 | 0.390252 | 0.946 | 0.865 | 3.56285E-18 |
| Intermediate | SEC11C | 1.89869E-18 | 0.390231 | 0.385 | 0.202 | 3.11841E-14 |
| Contractile | FN3KRP | 9.07451E-13 | 0.390048 | 0.265 | 0.147 | 1.4904E-08 |
| Terminal | HSP90AB1 | 9.39523E-16 | 0.389686 | 0.954 | 0.913 | 1.54307E-11 |
| RERGL_Cells | ADGRA2 | 2.02687E-10 | 0.389566 | 0.109 | 0.042 | 3.32893E-06 |
| Pericytes | DIMT1 | 9.62277E-07 | 0.389305 | 0.275 | 0.203 | 0.015804434 |
| Terminal | GMDS | 5.42351E-10 | 0.389266 | 0.342 | 0.219 | 8.90757E-06 |
| Intermediate | TMEM14A | 2.27194E-15 | 0.389173 | 0.486 | 0.302 | 3.73144E-11 |
| Terminal | DPYSL3 | 1.18612E-10 | 0.389151 | 0.292 | 0.174 | 1.94809E-06 |

| | | | | | | |
|---|---|---|---|---|---|---|
| Contractile | BAG2 | 3.21209E-12 | 0.388966 | 0.345 | 0.209 | 5.27554E-08 |
| Contractile | ERP29 | 4.11346E-08 | 0.388963 | 0.509 | 0.418 | 0.000675595 |
| Intermediate | CD99 | 4.54185E-24 | 0.388923 | 0.938 | 0.85 | 7.45953E-20 |
| RERGL_Cells | EIF3G | 2.14763E-07 | 0.388695 | 0.615 | 0.597 | 0.003527271 |
| RERGL_Cells | TRA2B | 9.92071E-07 | 0.388497 | 0.521 | 0.475 | 0.016293767 |
| Terminal | TMEM106 | 2.69986E-07 | 0.388402 | 0.362 | 0.265 | 0.004434258 |
| Contractile | MSRB1 | 1.75066E-10 | 0.388194 | 0.242 | 0.138 | 2.87528E-06 |
| RERGL_Cells | TPR | 5.17825E-05 | 0.388052 | 0.4 | 0.355 | 0.850475289 |
| Terminal | SULF1 | 3.23516E-09 | 0.387958 | 0.261 | 0.155 | 5.31343E-05 |
| RERGL_Cells | MRPL24 | 0.000134188 | 0.387662 | 0.359 | 0.321 | 1 |
| Fibroblasts | P4HA1 | 1.33525E-36 | 0.387609 | 0.256 | 0.092 | 2.19302E-32 |
| Terminal | OS9 | 1.4972E-09 | 0.387578 | 0.641 | 0.548 | 2.459E-05 |
| Contractile | AURKAIP1 | 2.05887E-13 | 0.387147 | 0.593 | 0.448 | 3.38149E-09 |
| RERGL_Cells | GHITM | 2.86341E-09 | 0.386944 | 0.632 | 0.6 | 4.70287E-05 |
| Intermediate | CIB1 | 1.48899E-16 | 0.38689 | 0.6 | 0.408 | 2.44551E-12 |
| Fibroblasts | C1RL | 1.34037E-36 | 0.386797 | 0.278 | 0.103 | 2.20143E-32 |
| Terminal | ANXA1 | 1.86899E-14 | 0.386725 | 0.906 | 0.837 | 3.06963E-10 |
| Pericytes | NGFR | 2.18884E-08 | 0.386569 | 0.196 | 0.115 | 0.000359495 |
| Fibroblasts | FAXDC2 | 2.65419E-35 | 0.386534 | 0.217 | 0.07 | 4.35924E-31 |
| Fibroblasts | GSTM3 | 2.94374E-23 | 0.386473 | 0.547 | 0.349 | 4.8348E-19 |
| RERGL_Cells | DEK | 3.12025E-06 | 0.38634 | 0.481 | 0.448 | 0.051247041 |
| Fibroblasts | ZNF331 | 2.3565E-14 | 0.386274 | 0.205 | 0.105 | 3.87032E-10 |
| Terminal | LIMS1 | 1.45333E-06 | 0.386255 | 0.355 | 0.274 | 0.023869436 |
| Contractile | TRDN | 2.92608E-30 | 0.386123 | 0.104 | 0.015 | 4.80579E-26 |
| Intermediate | GUK1 | 2.79514E-23 | 0.386051 | 0.897 | 0.785 | 4.59074E-19 |
| RERGL_Cells | EIF2S3 | 0.000236253 | 0.385937 | 0.393 | 0.354 | 1 |
| RERGL_Cells | CORO1B | 4.62519E-05 | 0.385877 | 0.323 | 0.272 | 0.759641519 |
| Pericytes | RARRES2 | 8.00144E-25 | 0.385839 | 0.836 | 0.622 | 1.31416E-20 |
| Terminal | HSPB1 | 8.87408E-21 | 0.385615 | 0.987 | 0.944 | 1.45748E-16 |
| RERGL_Cells | HSPB11 | 1.93481E-05 | 0.385611 | 0.261 | 0.198 | 0.317772902 |
| Intermediate | RCN3 | 2.83203E-15 | 0.385334 | 0.578 | 0.385 | 4.65133E-11 |
| Pericytes | HES4 | 3.18984E-17 | 0.385318 | 0.496 | 0.317 | 5.23899E-13 |
| Contractile | NUDC | 1.83464E-13 | 0.385141 | 0.677 | 0.546 | 3.01322E-09 |
| Fibroblasts | CCDC69 | 8.83354E-52 | 0.384977 | 0.205 | 0.042 | 1.45082E-47 |
| Intermediate | PPIB | 1.8267E-16 | 0.384864 | 0.884 | 0.742 | 3.00017E-12 |
| Pericytes | CNN2 | 1.11092E-06 | 0.384827 | 0.331 | 0.255 | 0.018245729 |
| Fibroblasts | SCARB2 | 2.45691E-29 | 0.384733 | 0.502 | 0.281 | 4.03522E-25 |
| Terminal | ZBTB16 | 9.69634E-06 | 0.384708 | 0.512 | 0.442 | 0.159252607 |
| Pericytes | SUSD2 | 1.16663E-08 | 0.38462 | 0.156 | 0.082 | 0.000191607 |
| Contractile | KCTD10 | 2.66825E-10 | 0.384495 | 0.275 | 0.166 | 4.38234E-06 |
| Pericytes | GRINA | 1.57555E-05 | 0.384173 | 0.318 | 0.257 | 0.258769004 |
| Fibroblasts | PPIB | 8.27506E-30 | 0.38409 | 0.875 | 0.716 | 1.3591E-25 |
| RERGL_Cells | FAM107B | 1.21739E-06 | 0.383995 | 0.177 | 0.108 | 0.019994495 |
| Terminal | RAB13 | 2.04977E-09 | 0.383963 | 0.477 | 0.359 | 3.36654E-05 |
| Intermediate | ERG | 8.8309E-37 | 0.383814 | 0.17 | 0.034 | 1.45039E-32 |
| Pericytes | EBF2 | 1.06351E-11 | 0.383758 | 0.164 | 0.077 | 1.74672E-07 |
| Pericytes | CCDC50 | 2.3342E-06 | 0.38373 | 0.291 | 0.221 | 0.038336916 |
| Pericytes | SNAI2 | 8.69131E-05 | 0.383699 | 0.185 | 0.129 | 1 |
| RERGL_Cells | HIP1 | 3.08773E-11 | 0.383595 | 0.145 | 0.063 | 5.0713E-07 |
| Contractile | EIF3E | 1.43176E-15 | 0.383518 | 0.878 | 0.814 | 2.35152E-11 |
| Terminal | FDXR | 2.40196E-10 | 0.383415 | 0.111 | 0.043 | 3.94498E-06 |
| RERGL_Cells | UBE2N | 5.64194E-06 | 0.383396 | 0.489 | 0.457 | 0.092663198 |
| Fibroblasts | SSPN | 7.81685E-29 | 0.383301 | 0.653 | 0.411 | 1.28384E-24 |
| Intermediate | MPG | 2.60926E-13 | 0.383143 | 0.596 | 0.43 | 4.28545E-09 |
| Pericytes | TDG | 4.14051E-06 | 0.38306 | 0.224 | 0.156 | 0.06800373 |
| Contractile | ARL2 | 1.20615E-13 | 0.383059 | 0.695 | 0.566 | 1.98098E-09 |

| | | | | | | |
|---|---|---|---|---|---|---|
| Contractile | KLHDC2 | 7.58331E-09 | 0.383004 | 0.467 | 0.351 | 0.000124548 |
| Terminal | METTL26 | 4.26147E-07 | 0.382931 | 0.423 | 0.323 | 0.006999044 |
| Terminal | CRYAB | 3.74169E-13 | 0.382835 | 0.895 | 0.787 | 6.14535E-09 |
| Terminal | PDGFD | 2.3083E-07 | 0.382831 | 0.294 | 0.197 | 0.003791159 |
| Terminal | SERTAD4-/ | 1.33263E-11 | 0.382809 | 0.268 | 0.148 | 2.18871E-07 |
| Contractile | NDUFB9 | 3.66102E-15 | 0.382647 | 0.756 | 0.648 | 6.01286E-11 |
| RERGL_Cells | SUSD2 | 1.1659E-07 | 0.382473 | 0.156 | 0.084 | 0.001914879 |
| RERGL_Cells | ARL6IP4 | 7.95026E-11 | 0.3823 | 0.731 | 0.698 | 1.30575E-06 |
| Pericytes | RNF139 | 1.09792E-07 | 0.382292 | 0.304 | 0.221 | 0.001803221 |
| Pericytes | TUBB2A | 8.71008E-08 | 0.382266 | 0.284 | 0.199 | 0.001430543 |
| Contractile | PRUNE2 | 6.63558E-17 | 0.382102 | 0.158 | 0.057 | 1.08983E-12 |
| Fibroblasts | PPIC | 5.92833E-28 | 0.381804 | 0.613 | 0.378 | 9.7367E-24 |
| Pericytes | IL32 | 2.10209E-07 | 0.38176 | 0.415 | 0.317 | 0.00345247 |
| Fibroblasts | PDIA6 | 1.07658E-20 | 0.381725 | 0.743 | 0.575 | 1.76818E-16 |
| Fibroblasts | NAV3 | 6.02372E-57 | 0.381686 | 0.113 | 0.004 | 9.89336E-53 |
| RERGL_Cells | NDUFA13 | 9.87306E-13 | 0.381676 | 0.827 | 0.807 | 1.62155E-08 |
| RERGL_Cells | SPOP | 0.000209121 | 0.381667 | 0.359 | 0.331 | 1 |
| Intermediate | NANS | 1.11303E-14 | 0.381554 | 0.35 | 0.197 | 1.82804E-10 |
| Contractile | COL14A1 | 9.30208E-14 | 0.38134 | 0.571 | 0.397 | 1.52777E-09 |
| Pericytes | SEC24D | 2.17293E-07 | 0.381077 | 0.264 | 0.185 | 0.003568826 |
| Contractile | RSU1 | 1.00357E-11 | 0.381037 | 0.549 | 0.415 | 1.64826E-07 |
| RERGL_Cells | MAT2A | 4.26552E-06 | 0.380914 | 0.329 | 0.257 | 0.070056872 |
| Pericytes | TMX1 | 0.000174668 | 0.380716 | 0.285 | 0.237 | 1 |
| Fibroblasts | PXK | 1.42713E-40 | 0.380595 | 0.263 | 0.088 | 2.34393E-36 |
| Contractile | SMIM4 | 5.76226E-14 | 0.380582 | 0.283 | 0.151 | 9.46393E-10 |
| Pericytes | B3GNT2 | 4.63836E-08 | 0.380525 | 0.187 | 0.11 | 0.000761805 |
| Terminal | ANXA5 | 4.04234E-12 | 0.380504 | 0.852 | 0.799 | 6.63914E-08 |
| Contractile | CRELD1 | 6.48569E-14 | 0.380458 | 0.269 | 0.141 | 1.06521E-09 |
| Terminal | C12orf57 | 6.49242E-13 | 0.380396 | 0.854 | 0.793 | 1.06632E-08 |
| Fibroblasts | NTRK2 | 3.59297E-28 | 0.380279 | 0.628 | 0.397 | 5.9011E-24 |
| Intermediate | ATP6AP2 | 6.09219E-16 | 0.380271 | 0.836 | 0.698 | 1.00058E-11 |
| RERGL_Cells | ANO1 | 4.09775E-05 | 0.38006 | 0.212 | 0.149 | 0.67301455 |
| Terminal | GLT8D2 | 6.24104E-07 | 0.380025 | 0.336 | 0.244 | 0.010250281 |
| Pericytes | MIR4435-2 | 5.56853E-08 | 0.379921 | 0.164 | 0.093 | 0.000914576 |
| Intermediate | GALNT18 | 1.7539E-38 | 0.379578 | 0.25 | 0.067 | 2.8806E-34 |
| Terminal | ANKH | 7.50675E-07 | 0.379532 | 0.272 | 0.185 | 0.012329083 |
| Terminal | PFN2 | 2.12622E-06 | 0.379368 | 0.503 | 0.409 | 0.034920989 |
| Fibroblasts | NPY1R | 5.51395E-22 | 0.379271 | 0.149 | 0.052 | 9.05611E-18 |
| Pericytes | CSRNP1 | 2.39716E-09 | 0.379258 | 0.351 | 0.248 | 3.93709E-05 |
| Pericytes | CCN2 | 6.48334E-09 | 0.379082 | 0.833 | 0.772 | 0.000106482 |
| Pericytes | NIFK | 1.2906E-05 | 0.379049 | 0.284 | 0.22 | 0.211968157 |
| Intermediate | SLC25A34- | 3.46687E-39 | 0.378875 | 0.151 | 0.025 | 5.69399E-35 |
| Contractile | PDHA1 | 1.36109E-10 | 0.378452 | 0.311 | 0.192 | 2.23545E-06 |
| Contractile | ARHGEF25 | 3.26965E-10 | 0.378384 | 0.246 | 0.139 | 5.37008E-06 |
| Intermediate | FKBP5 | 5.59737E-15 | 0.378241 | 0.344 | 0.189 | 9.19312E-11 |
| Contractile | SHC4 | 3.74761E-15 | 0.377529 | 0.168 | 0.067 | 6.15508E-11 |
| Intermediate | GNG5 | 3.20519E-14 | 0.377473 | 0.707 | 0.532 | 5.2642E-10 |
| Contractile | ENTPD1 | 3.25638E-20 | 0.377294 | 0.21 | 0.077 | 5.34828E-16 |
| Fibroblasts | C17orf58 | 1.48797E-30 | 0.377099 | 0.161 | 0.046 | 2.44384E-26 |
| Fibroblasts | DEPTOR | 6.2662E-65 | 0.376873 | 0.142 | 0.008 | 1.02916E-60 |
| RERGL_Cells | CHMP4A | 1.29134E-05 | 0.376858 | 0.385 | 0.33 | 0.212089492 |
| RERGL_Cells | ITPK1 | 3.47962E-09 | 0.376822 | 0.137 | 0.064 | 5.71492E-05 |
| Contractile | RWDD1 | 6.38405E-11 | 0.376773 | 0.657 | 0.55 | 1.04852E-06 |
| Contractile | C19orf53 | 3.38212E-12 | 0.376652 | 0.707 | 0.599 | 5.55479E-08 |
| Fibroblasts | SRSF6 | 8.7586E-27 | 0.376611 | 0.516 | 0.295 | 1.43851E-22 |
| RERGL_Cells | ZNF580 | 7.41306E-05 | 0.37652 | 0.265 | 0.207 | 1 |

| | | | | | | |
|---|---|---|---|---|---|---|
| Intermediate | COL15A1 | 4.77289E-10 | 0.376426 | 0.295 | 0.181 | 7.839E-06 |
| Fibroblasts | PRDX5 | 4.45E-26 | 0.376342 | 0.756 | 0.568 | 7.30868E-22 |
| Pericytes | SYPL1 | 1.49623E-10 | 0.376249 | 0.611 | 0.549 | 2.45741E-06 |
| Intermediate | ADGRE5 | 2.4099E-20 | 0.376216 | 0.421 | 0.216 | 3.95803E-16 |
| Contractile | UNC45A | 3.74791E-09 | 0.376161 | 0.303 | 0.197 | 6.15557E-05 |
| Terminal | FKBP5 | 3.75153E-05 | 0.376138 | 0.275 | 0.203 | 0.616151511 |
| Pericytes | RAD21 | 1.23102E-05 | 0.375871 | 0.324 | 0.259 | 0.202182913 |
| Terminal | RHOD | 1.45234E-09 | 0.375853 | 0.233 | 0.132 | 2.38532E-05 |
| Pericytes | PCDH1 | 1.55973E-21 | 0.375696 | 0.147 | 0.043 | 2.5617E-17 |
| Pericytes | MFSD14A | 9.29941E-05 | 0.375648 | 0.287 | 0.232 | 1 |
| Contractile | KCNAB1 | 1.39128E-19 | 0.375595 | 0.279 | 0.118 | 2.28504E-15 |
| Contractile | TSC22D1 | 8.49237E-23 | 0.375426 | 0.834 | 0.607 | 1.39479E-18 |
| Fibroblasts | TENT5A | 7.42384E-14 | 0.375415 | 0.42 | 0.281 | 1.21929E-09 |
| Fibroblasts | BACE2 | 3.67365E-30 | 0.375178 | 0.372 | 0.182 | 6.0336E-26 |
| Terminal | PERP | 0.000238214 | 0.375126 | 0.27 | 0.209 | 1 |
| Pericytes | ADAM19 | 2.61894E-17 | 0.375103 | 0.105 | 0.029 | 4.30135E-13 |
| Intermediate | PIK3R1 | 3.07508E-16 | 0.37497 | 0.265 | 0.126 | 5.0505E-12 |
| RERGL_Cells | GNAS | 1.48114E-08 | 0.374924 | 0.583 | 0.526 | 0.000243263 |
| Intermediate | TMEM47 | 9.56919E-14 | 0.374768 | 0.553 | 0.364 | 1.57164E-09 |
| Contractile | COX7A2 | 7.45924E-23 | 0.374755 | 0.918 | 0.869 | 1.22511E-18 |
| Intermediate | COL3A1 | 0.003394092 | 0.374535 | 0.402 | 0.341 | 1 |
| Pericytes | C14orf28 | 3.03585E-10 | 0.374465 | 0.153 | 0.075 | 4.98608E-06 |
| Pericytes | ZNF207 | 6.82894E-06 | 0.374361 | 0.4 | 0.348 | 0.112158443 |
| Terminal | HLA-A | 6.81746E-11 | 0.374299 | 0.891 | 0.875 | 1.1197E-06 |
| Contractile | MTRNR2L8 | 3.53959E-07 | 0.374191 | 0.226 | 0.144 | 0.005813415 |
| Pericytes | RBMX | 4.88263E-08 | 0.374072 | 0.593 | 0.546 | 0.000801922 |
| Contractile | OPTN | 6.87784E-10 | 0.373974 | 0.447 | 0.326 | 1.12962E-05 |
| RERGL_Cells | POMP | 8.36078E-13 | 0.373828 | 0.729 | 0.694 | 1.37318E-08 |
| RERGL_Cells | VCAM1 | 8.59201E-06 | 0.373653 | 0.239 | 0.167 | 0.141115132 |
| Terminal | LGALS1 | 3.32314E-23 | 0.373557 | 1 | 0.981 | 5.45793E-19 |
| Pericytes | MPRIP | 3.91868E-11 | 0.373526 | 0.218 | 0.118 | 6.43605E-07 |
| Contractile | TRMT112 | 1.17729E-12 | 0.373371 | 0.804 | 0.733 | 1.93358E-08 |
| Pericytes | LIMA1 | 4.24946E-09 | 0.373294 | 0.5 | 0.4 | 6.97932E-05 |
| Pericytes | TSPAN4 | 1.75736E-08 | 0.373177 | 0.571 | 0.48 | 0.000288629 |
| Terminal | PPIG | 1.52712E-08 | 0.373101 | 0.566 | 0.46 | 0.000250815 |
| Fibroblasts | MARCKS | 1.95019E-28 | 0.373065 | 0.363 | 0.178 | 3.20298E-24 |
| Intermediate | GALNT2 | 8.46879E-22 | 0.372984 | 0.284 | 0.122 | 1.39091E-17 |
| Contractile | AC018521 | 1.39181E-12 | 0.37285 | 0.198 | 0.095 | 2.28591E-08 |
| Terminal | MFAP2 | 6.4299E-09 | 0.372708 | 0.214 | 0.12 | 0.000105605 |
| Contractile | PDE5A | 1.14077E-12 | 0.372441 | 0.238 | 0.12 | 1.8736E-08 |
| Intermediate | CCPG1 | 1.34469E-12 | 0.372414 | 0.55 | 0.387 | 2.20852E-08 |
| Pericytes | EFHD1 | 2.996E-11 | 0.372223 | 0.529 | 0.389 | 4.92063E-07 |
| Contractile | PAM | 5.43826E-09 | 0.372138 | 0.487 | 0.373 | 8.93181E-05 |
| Terminal | MTHFD1L | 1.06063E-09 | 0.371954 | 0.229 | 0.128 | 1.74198E-05 |
| Fibroblasts | THBS3 | 7.94665E-33 | 0.371734 | 0.315 | 0.135 | 1.30516E-28 |
| RERGL_Cells | GUK1 | 2.17263E-15 | 0.37118 | 0.846 | 0.795 | 3.56832E-11 |
| Intermediate | CTNNAL1 | 2.29348E-17 | 0.371165 | 0.316 | 0.156 | 3.76682E-13 |
| Contractile | HSP90AA1 | 3.6038E-17 | 0.370485 | 0.92 | 0.871 | 5.91887E-13 |
| Pericytes | TSPAN3 | 1.75639E-06 | 0.370469 | 0.342 | 0.27 | 0.028846945 |
| Contractile | FBXL22 | 1.90384E-36 | 0.370284 | 0.118 | 0.015 | 3.12686E-32 |
| Fibroblasts | FAM107A | 7.9428E-46 | 0.370151 | 0.138 | 0.02 | 1.30453E-41 |
| RERGL_Cells | MAPK10 | 5.98802E-06 | 0.370015 | 0.184 | 0.119 | 0.098347233 |
| Intermediate | LINC01088 | 1.26868E-15 | 0.369918 | 0.148 | 0.053 | 2.08367E-11 |
| Pericytes | SLIT3 | 6.51776E-06 | 0.369806 | 0.327 | 0.259 | 0.107047649 |
| Terminal | TOMM7 | 5.25204E-14 | 0.369763 | 0.93 | 0.909 | 8.62596E-10 |
| Contractile | DMAC1 | 1.07268E-09 | 0.369675 | 0.485 | 0.367 | 1.76177E-05 |

| | | | | | | |
|---|---|---|---|---|---|---|
| RERGL_Cells | DLC1 | 1.15573E-05 | 0.369667 | 0.271 | 0.201 | 0.18981703 |
| Terminal | MPG | 1.53304E-05 | 0.369363 | 0.51 | 0.447 | 0.251786486 |
| Fibroblasts | TNFSF13 | 2.51538E-34 | 0.369293 | 0.223 | 0.075 | 4.13126E-30 |
| Fibroblasts | DDOST | 1.35401E-27 | 0.369217 | 0.49 | 0.28 | 2.22383E-23 |
| Fibroblasts | CREB3L1 | 2.25681E-41 | 0.369143 | 0.148 | 0.028 | 3.70658E-37 |
| Contractile | COPS6 | 2.00736E-13 | 0.36907 | 0.647 | 0.503 | 3.29689E-09 |
| Pericytes | BAG2 | 0.00162411 | 0.369057 | 0.262 | 0.222 | 1 |
| Contractile | DTNBP1 | 3.91967E-12 | 0.369052 | 0.232 | 0.121 | 6.43766E-08 |
| Terminal | NFIX | 8.57837E-09 | 0.368887 | 0.248 | 0.15 | 0.000140891 |
| Pericytes | SDC2 | 1.84426E-05 | 0.368876 | 0.625 | 0.636 | 0.302900609 |
| Terminal | NGEF | 8.65677E-29 | 0.368762 | 0.102 | 0.015 | 1.42179E-24 |
| Contractile | LGALS1 | 1.05138E-29 | 0.368685 | 0.996 | 0.981 | 1.72678E-25 |
| RERGL_Cells | UBE2H | 6.33996E-06 | 0.368628 | 0.188 | 0.124 | 0.104127535 |
| Contractile | CABIN1 | 3.1149E-12 | 0.368571 | 0.273 | 0.148 | 5.11591E-08 |
| Fibroblasts | MSX2 | 1.90597E-59 | 0.368387 | 0.13 | 0.008 | 3.13036E-55 |
| Pericytes | LMOD1 | 7.77092E-16 | 0.368278 | 0.631 | 0.437 | 1.2763E-11 |
| Terminal | GAMT | 2.13414E-06 | 0.368266 | 0.275 | 0.189 | 0.035051059 |
| Fibroblasts | CDKN1C | 3.67285E-33 | 0.367998 | 0.199 | 0.063 | 6.03229E-29 |
| Pericytes | GPI | 1.22671E-05 | 0.367952 | 0.287 | 0.224 | 0.201474911 |
| Fibroblasts | RECQL | 2.10497E-28 | 0.367829 | 0.355 | 0.173 | 3.4572E-24 |
| Contractile | SAP18 | 5.32757E-15 | 0.367769 | 0.836 | 0.738 | 8.75001E-11 |
| RERGL_Cells | YWHAZ | 2.49365E-05 | 0.36773 | 0.421 | 0.375 | 0.40955735 |
| RERGL_Cells | TUFM | 9.41436E-07 | 0.367621 | 0.534 | 0.51 | 0.015462143 |
| Terminal | CALM3 | 3.63694E-07 | 0.367514 | 0.625 | 0.554 | 0.005973307 |
| Intermediate | AC092164 | 3.72389E-18 | 0.36736 | 0.277 | 0.124 | 6.11612E-14 |
| Contractile | RALY | 2.56054E-11 | 0.367333 | 0.533 | 0.409 | 4.20543E-07 |
| RERGL_Cells | PLEKHA4 | 0.000181595 | 0.367112 | 0.237 | 0.182 | 1 |
| Terminal | MEDAG | 1.36855E-05 | 0.367023 | 0.285 | 0.208 | 0.224770811 |
| Pericytes | TOMM40 | 4.57405E-06 | 0.366811 | 0.236 | 0.169 | 0.075124134 |
| RERGL_Cells | PNRC2 | 1.57401E-05 | 0.36681 | 0.423 | 0.371 | 0.258516104 |
| RERGL_Cells | RAB11B | 0.000641462 | 0.366563 | 0.449 | 0.442 | 1 |
| Pericytes | POLR2H | 9.86485E-07 | 0.366425 | 0.242 | 0.169 | 0.016202037 |
| Pericytes | PARVA | 8.88826E-06 | 0.366271 | 0.329 | 0.269 | 0.145980778 |
| Terminal | AC245297 | 1.19443E-06 | 0.366236 | 0.294 | 0.206 | 0.019617333 |
| Pericytes | PTGER4 | 2.26149E-06 | 0.366187 | 0.122 | 0.068 | 0.037142658 |
| RERGL_Cells | CAP1 | 6.63339E-11 | 0.366136 | 0.752 | 0.684 | 1.08947E-06 |
| Contractile | KIF1C | 1.10677E-10 | 0.366121 | 0.218 | 0.116 | 1.81776E-06 |
| Intermediate | ZFYVE21 | 2.43611E-16 | 0.366087 | 0.622 | 0.428 | 4.00106E-12 |
| RERGL_Cells | SORT1 | 4.7895E-07 | 0.366071 | 0.209 | 0.131 | 0.007866271 |
| Contractile | PPIE | 3.5971E-10 | 0.366056 | 0.327 | 0.212 | 5.90788E-06 |
| Contractile | RBFOX2 | 1.83803E-10 | 0.365993 | 0.297 | 0.181 | 3.01877E-06 |
| Fibroblasts | EFNA5 | 4.21908E-44 | 0.365681 | 0.177 | 0.038 | 6.92942E-40 |
| Terminal | CLEC18A | 7.96641E-16 | 0.365652 | 0.105 | 0.028 | 1.3084E-11 |
| Fibroblasts | H6PD | 2.52664E-33 | 0.365628 | 0.194 | 0.06 | 4.14975E-29 |
| RERGL_Cells | TADA3 | 0.000383619 | 0.365561 | 0.295 | 0.248 | 1 |
| Intermediate | GSTO1 | 2.61258E-15 | 0.36556 | 0.708 | 0.523 | 4.29089E-11 |
| RERGL_Cells | ATP5F1C | 1.63794E-07 | 0.365231 | 0.575 | 0.536 | 0.002690149 |
| Fibroblasts | NGF | 2.31847E-31 | 0.365208 | 0.2 | 0.066 | 3.80785E-27 |
| RERGL_Cells | CDKN1B | 0.000335887 | 0.36517 | 0.233 | 0.181 | 1 |
| Fibroblasts | PDPN | 1.55147E-57 | 0.365168 | 0.129 | 0.008 | 2.54813E-53 |
| Fibroblasts | ANXA1 | 9.35681E-31 | 0.365158 | 0.94 | 0.805 | 1.53676E-26 |
| Contractile | GABARAP | 2.69856E-22 | 0.364867 | 0.938 | 0.888 | 4.43211E-18 |
| Intermediate | TSPAN2 | 5.0745E-25 | 0.364559 | 0.23 | 0.079 | 8.33437E-21 |
| Intermediate | CLU | 1.86373E-19 | 0.36454 | 0.968 | 0.801 | 3.06099E-15 |
| Pericytes | ETV6 | 1.41018E-13 | 0.364465 | 0.127 | 0.048 | 2.31607E-09 |
| Contractile | ATP5PD | 3.97497E-10 | 0.364405 | 0.657 | 0.548 | 6.5285E-06 |

| | | | | | | |
|---|---|---|---|---|---|---|
| Intermediate | PBXIP1 | 3.96694E-14 | 0.3644 | 0.518 | 0.328 | 6.51531E-10 |
| RERGL_Cells | C3orf70 | 3.96995E-09 | 0.364245 | 0.122 | 0.054 | 6.52024E-05 |
| RERGL_Cells | MRPL23 | 0.000965439 | 0.364226 | 0.306 | 0.269 | 1 |
| Fibroblasts | SLC35G2 | 2.50956E-50 | 0.364023 | 0.168 | 0.028 | 4.1217E-46 |
| RERGL_Cells | CITED4 | 1.8409E-09 | 0.363992 | 0.122 | 0.053 | 3.0235E-05 |
| Contractile | MICOS10 | 3.83854E-14 | 0.363972 | 0.78 | 0.638 | 6.30442E-10 |
| RERGL_Cells | KNOP1 | 0.00031776 | 0.363958 | 0.169 | 0.119 | 1 |
| RERGL_Cells | RASL11A | 1.39922E-11 | 0.363911 | 0.147 | 0.062 | 2.29808E-07 |
| Contractile | HSPB3 | 2.74838E-42 | 0.363855 | 0.108 | 0.009 | 4.51394E-38 |
| Intermediate | WIPF3 | 8.56408E-27 | 0.363678 | 0.135 | 0.03 | 1.40656E-22 |
| Fibroblasts | WBP1L | 6.27045E-27 | 0.363644 | 0.312 | 0.148 | 1.02986E-22 |
| Fibroblasts | KRT222 | 1.56741E-67 | 0.363464 | 0.118 | 0 | 2.57431E-63 |
| Contractile | PDE4DIP | 9.79529E-11 | 0.363235 | 0.22 | 0.118 | 1.60878E-06 |
| Pericytes | MARCH7 | 1.18359E-05 | 0.363206 | 0.293 | 0.227 | 0.194393632 |
| Intermediate | MGST2 | 7.33578E-20 | 0.363153 | 0.368 | 0.182 | 1.20483E-15 |
| RERGL_Cells | AL035071. | 4.83138E-09 | 0.363131 | 0.165 | 0.084 | 7.93505E-05 |
| RERGL_Cells | EEF1B2 | 1.03211E-11 | 0.362864 | 0.842 | 0.792 | 1.69514E-07 |
| Contractile | REEP1 | 7.4361E-27 | 0.362791 | 0.164 | 0.041 | 1.2213E-22 |
| Fibroblasts | RNF24 | 2.47395E-26 | 0.362741 | 0.38 | 0.197 | 4.06322E-22 |
| Fibroblasts | DYNC2LI1 | 1.42921E-35 | 0.362638 | 0.276 | 0.104 | 2.34733E-31 |
| Contractile | PTGES3 | 5.04812E-15 | 0.362576 | 0.83 | 0.76 | 8.29104E-11 |
| Pericytes | TUBA1B | 4.73012E-21 | 0.362376 | 0.949 | 0.917 | 7.76874E-17 |
| Pericytes | TAF1D | 0.000303969 | 0.362321 | 0.335 | 0.293 | 1 |
| RERGL_Cells | RNF14 | 1.31734E-05 | 0.362148 | 0.179 | 0.118 | 0.216360234 |
| RERGL_Cells | CZIB | 3.49804E-05 | 0.361767 | 0.393 | 0.345 | 0.5745185 |
| RERGL_Cells | NR3C1 | 4.79583E-05 | 0.361703 | 0.346 | 0.291 | 0.78766755 |
| Pericytes | C6orf62 | 5.73326E-07 | 0.36166 | 0.365 | 0.285 | 0.009416312 |
| Terminal | SEC11A | 8.59174E-10 | 0.361459 | 0.621 | 0.532 | 1.41111E-05 |
| Pericytes | PTBP1 | 9.28657E-06 | 0.361368 | 0.315 | 0.251 | 0.15252259 |
| Contractile | ANAPC15 | 4.76318E-16 | 0.361303 | 0.253 | 0.118 | 7.82304E-12 |
| Terminal | EMC10 | 2.57757E-07 | 0.361154 | 0.442 | 0.34 | 0.004233406 |
| RERGL_Cells | LAMA5 | 4.66322E-09 | 0.360955 | 0.132 | 0.061 | 7.65887E-05 |
| Pericytes | ATG101 | 0.000289206 | 0.360925 | 0.325 | 0.283 | 1 |
| Pericytes | LPCAT3 | 2.07387E-07 | 0.360851 | 0.165 | 0.098 | 0.003406118 |
| Fibroblasts | MAP1A | 1.41622E-25 | 0.360816 | 0.314 | 0.152 | 2.326E-21 |
| Pericytes | MAPRE1 | 9.79709E-07 | 0.360761 | 0.393 | 0.32 | 0.016090743 |
| Pericytes | FHOD1 | 6.87303E-14 | 0.360722 | 0.115 | 0.039 | 1.12883E-09 |
| Contractile | DTNA | 7.31293E-21 | 0.360676 | 0.152 | 0.045 | 1.20108E-16 |
| Pericytes | PDGFA | 2.73539E-06 | 0.360597 | 0.233 | 0.156 | 0.044926059 |
| Fibroblasts | C1orf56 | 6.07702E-27 | 0.360504 | 0.431 | 0.231 | 9.9809E-23 |
| Fibroblasts | PTGFRN | 4.52248E-50 | 0.360209 | 0.14 | 0.017 | 7.42773E-46 |
| RERGL_Cells | VEZF1 | 1.30514E-05 | 0.360177 | 0.203 | 0.138 | 0.21435605 |
| Contractile | TPD52L2 | 3.75178E-08 | 0.360067 | 0.421 | 0.319 | 0.000616192 |
| Fibroblasts | SFT2D1 | 2.85507E-30 | 0.360048 | 0.485 | 0.262 | 4.68917E-26 |
| Terminal | GNG11 | 4.08085E-07 | 0.360038 | 0.588 | 0.504 | 0.006702396 |
| Fibroblasts | SAR1A | 9.54473E-27 | 0.359999 | 0.566 | 0.341 | 1.56763E-22 |
| Contractile | SMARCD3 | 1.37177E-10 | 0.359948 | 0.325 | 0.198 | 2.25299E-06 |
| Fibroblasts | PTGIR | 2.64967E-19 | 0.359841 | 0.394 | 0.235 | 4.35182E-15 |
| Terminal | SNHG32 | 3.23396E-08 | 0.359813 | 0.662 | 0.583 | 0.000531146 |
| Contractile | SNHG6 | 1.24246E-08 | 0.359806 | 0.623 | 0.529 | 0.000204061 |
| RERGL_Cells | TIMM17B | 0.000259187 | 0.359799 | 0.259 | 0.207 | 1 |
| Pericytes | HSPD1 | 0.002523086 | 0.359689 | 0.465 | 0.458 | 1 |
| Pericytes | PABPC1 | 8.28269E-12 | 0.359596 | 0.78 | 0.731 | 1.36035E-07 |
| Fibroblasts | GRN | 2.12342E-26 | 0.359327 | 0.579 | 0.357 | 3.48751E-22 |
| RERGL_Cells | NDUFV1 | 0.000538771 | 0.359312 | 0.387 | 0.361 | 1 |
| Contractile | PRDM6 | 1.23836E-15 | 0.359279 | 0.154 | 0.057 | 2.03388E-11 |

| | | | | | | |
|---|---|---|---|---|---|---|
| Contractile | ALDOC | 5.57736E-14 | 0.359164 | 0.18 | 0.077 | 9.16026E-10 |
| Contractile | SAE1 | 1.08285E-09 | 0.359137 | 0.311 | 0.201 | 1.77847E-05 |
| Intermediate | EMP3 | 6.72958E-18 | 0.358981 | 0.753 | 0.545 | 1.10527E-13 |
| RERGL_Cells | SLC25A33 | 1.00782E-08 | 0.358965 | 0.135 | 0.064 | 0.000165525 |
| Intermediate | ARPC2 | 4.58311E-16 | 0.358722 | 0.845 | 0.76 | 7.5273E-12 |
| Terminal | TXNDC12 | 2.32803E-08 | 0.358708 | 0.357 | 0.249 | 0.000382356 |
| Intermediate | FRMD4B | 6.14335E-30 | 0.358593 | 0.217 | 0.064 | 1.00898E-25 |
| Contractile | RNF5 | 5.11099E-10 | 0.358397 | 0.419 | 0.297 | 8.39429E-06 |
| Fibroblasts | BNC2 | 3.66566E-64 | 0.358331 | 0.119 | 0.002 | 6.02047E-60 |
| Pericytes | CSKMT | 1.06212E-05 | 0.358189 | 0.164 | 0.106 | 0.174441821 |
| Pericytes | PTPRA | 5.50444E-05 | 0.358167 | 0.338 | 0.288 | 0.904048754 |
| Contractile | TP53TG1 | 1.32491E-10 | 0.358133 | 0.261 | 0.15 | 2.17603E-06 |
| Pericytes | NARS | 6.88801E-08 | 0.357908 | 0.415 | 0.334 | 0.001131286 |
| Intermediate | MAGED2 | 4.17604E-15 | 0.357859 | 0.783 | 0.613 | 6.85874E-11 |
| Fibroblasts | CNIH1 | 2.4037E-24 | 0.357768 | 0.563 | 0.356 | 3.94784E-20 |
| RERGL_Cells | HVCN1 | 7.50369E-11 | 0.357718 | 0.122 | 0.049 | 1.23241E-06 |
| Terminal | ARRDC2 | 1.40156E-07 | 0.357663 | 0.166 | 0.09 | 0.002301925 |
| RERGL_Cells | MEA1 | 0.000674457 | 0.357649 | 0.331 | 0.302 | 1 |
| Fibroblasts | HEXA | 1.91704E-26 | 0.357634 | 0.487 | 0.282 | 3.14855E-22 |
| Fibroblasts | MDK | 1.49203E-27 | 0.357514 | 0.295 | 0.133 | 2.45051E-23 |
| Fibroblasts | PRNP | 2.55561E-22 | 0.357411 | 0.708 | 0.503 | 4.19734E-18 |
| Contractile | TTC1 | 3.13221E-09 | 0.35734 | 0.371 | 0.257 | 5.14434E-05 |
| Intermediate | NAP1L1 | 4.7987E-16 | 0.357281 | 0.807 | 0.652 | 7.88138E-12 |
| RERGL_Cells | GRK5 | 1.00254E-05 | 0.357276 | 0.25 | 0.181 | 0.164657434 |
| Fibroblasts | HDLBP | 1.21313E-25 | 0.357256 | 0.502 | 0.296 | 1.99244E-21 |
| Terminal | GADD45G | 0.005637637 | 0.357059 | 0.205 | 0.159 | 1 |
| RERGL_Cells | SEPTIN2 | 7.21638E-08 | 0.356991 | 0.609 | 0.571 | 0.001185219 |
| Pericytes | SFT2D1 | 4.48406E-05 | 0.35689 | 0.373 | 0.321 | 0.736462596 |
| RERGL_Cells | COPS6 | 5.55178E-07 | 0.356852 | 0.547 | 0.519 | 0.009118243 |
| Contractile | TAF15 | 1.22112E-09 | 0.356759 | 0.277 | 0.172 | 2.00557E-05 |
| RERGL_Cells | INAFM1 | 3.57699E-05 | 0.3567 | 0.291 | 0.228 | 0.58748543 |
| RERGL_Cells | FGD4 | 6.54207E-08 | 0.356553 | 0.115 | 0.053 | 0.001074469 |
| Pericytes | CPSF4 | 1.10944E-06 | 0.356529 | 0.165 | 0.101 | 0.018221421 |
| Pericytes | DNAJA1 | 2.87767E-05 | 0.356496 | 0.544 | 0.515 | 0.472628557 |
| Terminal | IFI27L2 | 1.18425E-05 | 0.356429 | 0.414 | 0.336 | 0.194502025 |
| Pericytes | DHX15 | 4.75881E-06 | 0.35642 | 0.262 | 0.194 | 0.078158771 |
| RERGL_Cells | CCNI | 8.06506E-18 | 0.356358 | 0.925 | 0.864 | 1.32461E-13 |
| Intermediate | STMN1 | 2.38244E-19 | 0.356352 | 0.28 | 0.122 | 3.91293E-15 |
| RERGL_Cells | TEAD2 | 4.88314E-06 | 0.356275 | 0.19 | 0.123 | 0.080200673 |
| Intermediate | MMP23B | 3.9188E-11 | 0.355987 | 0.333 | 0.205 | 6.43623E-07 |
| Contractile | ZNHIT3 | 1.12287E-08 | 0.355923 | 0.301 | 0.198 | 0.00018442 |
| Contractile | NSA2 | 2.1484E-06 | 0.355863 | 0.341 | 0.258 | 0.03528529 |
| Fibroblasts | SAMD11 | 4.07966E-47 | 0.355629 | 0.135 | 0.017 | 6.70043E-43 |
| RERGL_Cells | HNRNPU | 9.39821E-09 | 0.355496 | 0.632 | 0.595 | 0.000154356 |
| Fibroblasts | TMBIM6 | 1.30198E-31 | 0.355377 | 0.901 | 0.738 | 2.13838E-27 |
| RERGL_Cells | HNRNPD | 8.31172E-05 | 0.355238 | 0.297 | 0.242 | 1 |
| Fibroblasts | TMED5 | 3.88634E-32 | 0.355215 | 0.356 | 0.161 | 6.38293E-28 |
| Fibroblasts | FBXO2 | 4.652E-47 | 0.355195 | 0.141 | 0.02 | 7.64044E-43 |
| Intermediate | MYDGF | 1.28081E-12 | 0.355172 | 0.725 | 0.555 | 2.10359E-08 |
| Contractile | NDUFB4 | 1.53547E-11 | 0.355047 | 0.768 | 0.678 | 2.52185E-07 |
| Fibroblasts | STAT3 | 9.27658E-27 | 0.354721 | 0.612 | 0.383 | 1.52359E-22 |
| Intermediate | CRELD1 | 1.8328E-17 | 0.354696 | 0.288 | 0.136 | 3.01019E-13 |
| Fibroblasts | RPN2 | 4.88044E-24 | 0.354618 | 0.681 | 0.483 | 8.01563E-20 |
| Contractile | SSBP1 | 6.89586E-09 | 0.354532 | 0.609 | 0.514 | 0.000113258 |
| RERGL_Cells | LSM2 | 0.000613355 | 0.354506 | 0.308 | 0.266 | 1 |
| Contractile | PXMP2 | 9.8759E-13 | 0.354504 | 0.202 | 0.096 | 1.62202E-08 |

| | | | | | | |
|---|---|---|---|---|---|---|
| Pericytes | STK3 | 4.49582E-07 | 0.354386 | 0.2 | 0.128 | 0.007383932 |
| Contractile | XIST | 8.9857E-14 | 0.354324 | 0.178 | 0.075 | 1.47581E-09 |
| Terminal | ZMAT3 | 1.92793E-08 | 0.354218 | 0.218 | 0.127 | 0.000316644 |
| RERGL_Cells | RRAS | 0.000254285 | 0.354138 | 0.333 | 0.291 | 1 |
| Contractile | UBE2V2 | 6.96528E-10 | 0.354005 | 0.491 | 0.372 | 1.14398E-05 |
| Pericytes | ARL5A | 0.000316416 | 0.353995 | 0.453 | 0.427 | 1 |
| RERGL_Cells | CNN1 | 5.39729E-29 | 0.353827 | 0.748 | 0.432 | 8.86451E-25 |
| RERGL_Cells | EIF3D | 2.82606E-05 | 0.353788 | 0.53 | 0.5 | 0.46415172 |
| Fibroblasts | ITGA5 | 4.40682E-24 | 0.353576 | 0.206 | 0.083 | 7.23777E-20 |
| Terminal | CUTA | 8.97675E-10 | 0.353557 | 0.691 | 0.582 | 1.47434E-05 |
| Pericytes | EIF2S2 | 9.1658E-08 | 0.35348 | 0.578 | 0.536 | 0.001505391 |
| Contractile | MGST2 | 1.04105E-08 | 0.353423 | 0.301 | 0.195 | 0.000170982 |
| Terminal | FAM3C | 3.60612E-06 | 0.353264 | 0.536 | 0.462 | 0.059226913 |
| Fibroblasts | SEC22B | 1.38781E-25 | 0.353229 | 0.454 | 0.255 | 2.27934E-21 |
| Pericytes | PSMD7 | 1.25203E-06 | 0.353154 | 0.435 | 0.37 | 0.020563331 |
| Fibroblasts | MAPK3 | 1.66012E-31 | 0.353093 | 0.336 | 0.15 | 2.72658E-27 |
| Pericytes | HMG20B | 3.34474E-07 | 0.352975 | 0.32 | 0.236 | 0.005493405 |
| Fibroblasts | TJP2 | 5.39642E-26 | 0.35295 | 0.277 | 0.126 | 8.86308E-22 |
| Fibroblasts | ATG101 | 8.08822E-25 | 0.352923 | 0.423 | 0.232 | 1.32841E-20 |
| Pericytes | CEBPZ | 0.000185784 | 0.352636 | 0.311 | 0.265 | 1 |
| RERGL_Cells | NDUFS5 | 7.55241E-12 | 0.352631 | 0.827 | 0.802 | 1.24041E-07 |
| Terminal | DAP | 1.04761E-07 | 0.352614 | 0.399 | 0.297 | 0.0017206 |
| Intermediate | ACTN1 | 1.2412E-24 | 0.352525 | 0.864 | 0.581 | 2.03855E-20 |
| Pericytes | PAG1 | 1.45687E-13 | 0.352249 | 0.102 | 0.033 | 2.39276E-09 |
| Contractile | RIMKLB | 1.31555E-10 | 0.352137 | 0.23 | 0.125 | 2.16067E-06 |
| RERGL_Cells | ARL3 | 0.000334399 | 0.352064 | 0.382 | 0.347 | 1 |
| RERGL_Cells | DPM3 | 0.003646781 | 0.352043 | 0.325 | 0.299 | 1 |
| Fibroblasts | SEC61A1 | 1.62979E-20 | 0.352008 | 0.372 | 0.215 | 2.67677E-16 |
| Pericytes | ABCE1 | 1.57167E-05 | 0.351719 | 0.256 | 0.194 | 0.258130413 |
| Contractile | NDUFS6 | 1.55474E-11 | 0.35167 | 0.639 | 0.506 | 2.55351E-07 |
| RERGL_Cells | PPP2R5E | 2.79176E-05 | 0.35155 | 0.205 | 0.143 | 0.458519473 |
| Intermediate | SSR4 | 3.02824E-18 | 0.351528 | 0.86 | 0.73 | 4.97357E-14 |
| RERGL_Cells | RGS16 | 2.06592E-51 | 0.351482 | 0.538 | 0.203 | 3.39307E-47 |
| Contractile | ATP5F1E | 6.67535E-20 | 0.351384 | 0.944 | 0.89 | 1.09636E-15 |
| RERGL_Cells | HNRNPA3 | 2.51383E-08 | 0.351017 | 0.701 | 0.667 | 0.000412871 |
| Pericytes | SNHG16 | 0.000107601 | 0.350973 | 0.355 | 0.303 | 1 |
| Pericytes | NES | 2.55933E-09 | 0.350936 | 0.135 | 0.064 | 4.20345E-05 |
| RERGL_Cells | PRPSAP1 | 1.76841E-05 | 0.35091 | 0.226 | 0.162 | 0.290443061 |
| Pericytes | ANGPTL4 | 2.01431E-06 | 0.350896 | 0.413 | 0.323 | 0.033082958 |
| Terminal | CCND3 | 1.48266E-06 | 0.350895 | 0.266 | 0.182 | 0.02435125 |
| RERGL_Cells | CBWD1 | 0.000165681 | 0.350891 | 0.212 | 0.158 | 1 |
| RERGL_Cells | RAD23A | 2.60581E-08 | 0.350717 | 0.618 | 0.575 | 0.000427978 |
| Contractile | S100A4 | 6.3564E-07 | 0.350706 | 0.784 | 0.764 | 0.010439749 |
| Intermediate | ERP29 | 6.49289E-12 | 0.350648 | 0.578 | 0.404 | 1.06639E-07 |
| Terminal | SLC14A1 | 1.73745E-08 | 0.350553 | 0.231 | 0.132 | 0.00028536 |
| Contractile | RTL8C | 1.01477E-10 | 0.350551 | 0.637 | 0.52 | 1.66666E-06 |
| Intermediate | TMED4 | 9.14363E-14 | 0.350476 | 0.512 | 0.325 | 1.50175E-09 |
| Pericytes | NIP7 | 2.3152E-07 | 0.350351 | 0.195 | 0.122 | 0.003802482 |
| RERGL_Cells | PDGFRB | 0.000613224 | 0.350249 | 0.412 | 0.375 | 1 |
| Contractile | TBXA2R | 1.25214E-15 | 0.349949 | 0.164 | 0.063 | 2.05651E-11 |
| Contractile | PLPBP | 4.47624E-08 | 0.349815 | 0.323 | 0.221 | 0.000735177 |
| RERGL_Cells | CAMK1 | 0.000122176 | 0.349801 | 0.274 | 0.216 | 1 |
| Pericytes | RTN4 | 2.13419E-15 | 0.349543 | 0.893 | 0.85 | 3.50519E-11 |
| Fibroblasts | PER2 | 3.14942E-27 | 0.349472 | 0.237 | 0.096 | 5.1726E-23 |
| RERGL_Cells | DTX3 | 0.001029856 | 0.34934 | 0.207 | 0.16 | 1 |
| Contractile | ECHS1 | 1.87223E-08 | 0.349338 | 0.475 | 0.373 | 0.000307494 |

| | | | | | | |
|---|---|---|---|---|---|---|
| Fibroblasts | LINC01697 | 4.03986E-56 | 0.349311 | 0.103 | 0.002 | 6.63506E-52 |
| Contractile | SMIM26 | 2.16157E-08 | 0.349274 | 0.561 | 0.472 | 0.000355015 |
| Fibroblasts | LGR4 | 1.08371E-40 | 0.349201 | 0.131 | 0.021 | 1.77988E-36 |
| RERGL_Cells | TPI1 | 1.69042E-11 | 0.349055 | 0.825 | 0.812 | 2.77635E-07 |
| Contractile | ODC1 | 2.22767E-08 | 0.348975 | 0.317 | 0.216 | 0.000365872 |
| Fibroblasts | GLG1 | 1.88466E-24 | 0.348956 | 0.478 | 0.277 | 3.09536E-20 |
| Pericytes | PTEN | 1.10958E-06 | 0.34888 | 0.256 | 0.181 | 0.018223669 |
| Pericytes | MRPL42 | 5.76696E-05 | 0.348829 | 0.291 | 0.234 | 0.947165809 |
| Contractile | MOAP1 | 1.14227E-09 | 0.34882 | 0.279 | 0.171 | 1.87606E-05 |
| Terminal | P4HA2 | 1.9504E-06 | 0.348796 | 0.283 | 0.201 | 0.032033435 |
| Contractile | ST6GAL2 | 5.55364E-35 | 0.348771 | 0.114 | 0.015 | 9.1213E-31 |
| Contractile | CCDC8 | 4.51778E-06 | 0.348703 | 0.194 | 0.127 | 0.074199951 |
| Pericytes | RPF2 | 4.37739E-06 | 0.34864 | 0.291 | 0.223 | 0.071894317 |
| Contractile | ACTR3 | 1.56217E-08 | 0.348631 | 0.589 | 0.497 | 0.000256571 |
| Pericytes | RBM39 | 3.44001E-09 | 0.34863 | 0.678 | 0.624 | 5.64987E-05 |
| Terminal | ZCRB1 | 3.67254E-07 | 0.34857 | 0.434 | 0.335 | 0.006031776 |
| Fibroblasts | GFM1 | 3.54875E-11 | 0.348559 | 0.194 | 0.11 | 5.82846E-07 |
| Pericytes | TECR | 0.00010655 | 0.348509 | 0.413 | 0.376 | 1 |
| Pericytes | TUBB3 | 1.45386E-25 | 0.348495 | 0.102 | 0.018 | 2.38782E-21 |
| Contractile | GALNT18 | 3.85234E-12 | 0.348404 | 0.176 | 0.081 | 6.32708E-08 |
| RERGL_Cells | SUSD5 | 3.9924E-08 | 0.348403 | 0.209 | 0.123 | 0.000655712 |
| Intermediate | PHPT1 | 3.18646E-17 | 0.348261 | 0.744 | 0.572 | 5.23344E-13 |
| Pericytes | KCTD11 | 4.77439E-08 | 0.348251 | 0.144 | 0.076 | 0.000784145 |
| Contractile | C22orf39 | 1.94683E-09 | 0.347984 | 0.277 | 0.169 | 3.19747E-05 |
| RERGL_Cells | RAP1A | 5.79624E-06 | 0.347953 | 0.442 | 0.39 | 0.095197518 |
| Pericytes | RSU1 | 3.2245E-06 | 0.347928 | 0.487 | 0.424 | 0.052959106 |
| Contractile | MRAS | 6.61205E-11 | 0.347631 | 0.204 | 0.106 | 1.08596E-06 |
| Contractile | ENO2 | 4.92692E-13 | 0.347614 | 0.212 | 0.101 | 8.09198E-09 |
| Contractile | PGM5 | 1.40467E-12 | 0.347585 | 0.184 | 0.084 | 2.30704E-08 |
| RERGL_Cells | TP53INP2 | 1.17214E-14 | 0.347584 | 0.109 | 0.033 | 1.92512E-10 |
| Contractile | SPARCL1 | 5.145E-25 | 0.347573 | 0.998 | 0.949 | 8.45015E-21 |
| Terminal | COPZ2 | 6.31787E-06 | 0.347368 | 0.235 | 0.159 | 0.10376475 |
| RERGL_Cells | DTNA | 3.5125E-10 | 0.347272 | 0.122 | 0.05 | 5.76894E-06 |
| Intermediate | ARL6IP5 | 1.5162E-17 | 0.34727 | 0.888 | 0.715 | 2.49021E-13 |
| Terminal | FAM177A1 | 4.44771E-06 | 0.347211 | 0.436 | 0.356 | 0.073049177 |
| Contractile | MYOCD | 7.73075E-18 | 0.347197 | 0.136 | 0.041 | 1.2697E-13 |
| Fibroblasts | OS9 | 7.01454E-24 | 0.347049 | 0.708 | 0.496 | 1.15207E-19 |
| Pericytes | ITSN1 | 3.57371E-06 | 0.34701 | 0.191 | 0.127 | 0.058694671 |
| Intermediate | CCDC3 | 9.32388E-29 | 0.346962 | 0.763 | 0.418 | 1.53135E-24 |
| Contractile | ADCY5 | 1.60414E-21 | 0.346675 | 0.13 | 0.033 | 2.63465E-17 |
| Contractile | CORO1C | 3.40332E-08 | 0.346641 | 0.265 | 0.169 | 0.000558962 |
| Intermediate | IGFBP3 | 8.18309E-06 | 0.346528 | 0.159 | 0.095 | 0.134398991 |
| Fibroblasts | MFSD1 | 1.56199E-31 | 0.346479 | 0.307 | 0.131 | 2.56542E-27 |
| Pericytes | PIM3 | 6.73511E-06 | 0.346414 | 0.191 | 0.126 | 0.110617424 |
| Contractile | TAX1BP1 | 1.64441E-08 | 0.346325 | 0.593 | 0.495 | 0.000270078 |
| Pericytes | MAP1S | 1.25464E-08 | 0.346317 | 0.164 | 0.089 | 0.000206062 |
| Terminal | IGBP1 | 1.69997E-05 | 0.34623 | 0.338 | 0.263 | 0.279203102 |
| RERGL_Cells | ACTG2 | 7.18896E-24 | 0.346225 | 0.387 | 0.183 | 1.18071E-19 |
| Contractile | MRPL10 | 1.87884E-09 | 0.346097 | 0.313 | 0.201 | 3.08581E-05 |
| RERGL_Cells | ATP5PB | 5.11802E-05 | 0.346045 | 0.521 | 0.508 | 0.840583622 |
| Pericytes | LYPLA1 | 1.0967E-06 | 0.345937 | 0.231 | 0.16 | 0.018012252 |
| Terminal | MAMDC2 | 1.28088E-07 | 0.345862 | 0.205 | 0.118 | 0.00210372 |
| Contractile | COX5B | 5.40022E-20 | 0.345834 | 0.914 | 0.807 | 8.86931E-16 |
| Intermediate | ARPC3 | 6.67973E-14 | 0.345808 | 0.738 | 0.629 | 1.09708E-09 |
| RERGL_Cells | SELENOK | 1.65616E-06 | 0.345749 | 0.575 | 0.559 | 0.027200724 |
| Contractile | TRIM36 | 6.18739E-25 | 0.345581 | 0.114 | 0.023 | 1.01622E-20 |

| | | | | | | |
|---|---|---|---|---|---|---|
| Intermediate | NUCB1 | 3.17989E-13 | 0.345504 | 0.74 | 0.597 | 5.22266E-09 |
| Fibroblasts | LTBP3 | 1.49983E-19 | 0.345303 | 0.346 | 0.192 | 2.46332E-15 |
| Pericytes | NFKBIZ | 0.000367769 | 0.345188 | 0.244 | 0.189 | 1 |
| RERGL_Cells | PSMB6 | 1.51573E-07 | 0.345123 | 0.637 | 0.623 | 0.002489431 |
| Intermediate | AL390755. | 3.97199E-33 | 0.345015 | 0.191 | 0.046 | 6.52359E-29 |
| Fibroblasts | FAM180B | 5.43956E-58 | 0.344969 | 0.103 | 0.001 | 8.93394E-54 |
| Contractile | UQCRQ | 6.15984E-12 | 0.344871 | 0.808 | 0.728 | 1.01169E-07 |
| Pericytes | TCEAL9 | 1.70836E-06 | 0.344868 | 0.64 | 0.623 | 0.02805803 |
| Terminal | GLRB | 4.37445E-09 | 0.344816 | 0.146 | 0.07 | 7.18459E-05 |
| Pericytes | GALNT16 | 5.2753E-09 | 0.344766 | 0.153 | 0.079 | 8.66416E-05 |
| RERGL_Cells | NUTF2 | 0.00100251 | 0.344762 | 0.318 | 0.283 | 1 |
| Contractile | CAPZB | 5.60218E-14 | 0.344562 | 0.784 | 0.68 | 9.20103E-10 |
| Contractile | COA3 | 1.17327E-07 | 0.344446 | 0.401 | 0.303 | 0.00192698 |
| RERGL_Cells | NME4 | 5.67251E-07 | 0.344372 | 0.481 | 0.411 | 0.009316529 |
| RERGL_Cells | TACC1 | 0.000124189 | 0.344356 | 0.447 | 0.42 | 1 |
| Pericytes | PNPLA8 | 5.07523E-05 | 0.344323 | 0.267 | 0.208 | 0.833556138 |
| RERGL_Cells | R3HCC1 | 4.53543E-05 | 0.344297 | 0.209 | 0.15 | 0.74489889 |
| Contractile | SLIRP | 5.56529E-11 | 0.344182 | 0.693 | 0.588 | 9.14043E-07 |
| Intermediate | DSTN | 1.18936E-34 | 0.344144 | 1 | 0.951 | 1.9534E-30 |
| RERGL_Cells | SET | 1.3093E-05 | 0.344013 | 0.485 | 0.436 | 0.215039095 |
| Fibroblasts | AC097662 | 4.04559E-42 | 0.343891 | 0.101 | 0.008 | 6.64448E-38 |
| Contractile | MANBAL | 4.48842E-08 | 0.343866 | 0.399 | 0.298 | 0.000737179 |
| RERGL_Cells | HADHA | 3.72023E-05 | 0.343788 | 0.47 | 0.437 | 0.611010267 |
| Pericytes | ILF3 | 0.000181647 | 0.343659 | 0.296 | 0.246 | 1 |
| RERGL_Cells | MYO1C | 0.001085457 | 0.343613 | 0.35 | 0.317 | 1 |
| Terminal | SLC9A3R2 | 0.001188881 | 0.343514 | 0.24 | 0.184 | 1 |
| Terminal | RNASET2 | 1.43794E-05 | 0.343461 | 0.22 | 0.151 | 0.236166845 |
| Terminal | KCNK15 | 3.42401E-07 | 0.343461 | 0.237 | 0.15 | 0.005623602 |
| Contractile | MRPS18C | 2.28231E-08 | 0.343316 | 0.441 | 0.331 | 0.000374847 |
| Fibroblasts | PLK3 | 1.13232E-23 | 0.34326 | 0.319 | 0.162 | 1.85973E-19 |
| Intermediate | PLEC | 1.41681E-16 | 0.343156 | 0.28 | 0.134 | 2.32697E-12 |
| Intermediate | CYFIP2 | 1.42416E-20 | 0.343104 | 0.209 | 0.077 | 2.33904E-16 |
| Fibroblasts | HSPG2 | 7.15687E-16 | 0.343024 | 0.313 | 0.184 | 1.17544E-11 |
| Fibroblasts | CERS2 | 7.26913E-25 | 0.343018 | 0.413 | 0.225 | 1.19388E-20 |
| Contractile | RYR2 | 4.62411E-27 | 0.342943 | 0.102 | 0.016 | 7.59464E-23 |
| Intermediate | PEBP1 | 1.24791E-18 | 0.342936 | 0.942 | 0.863 | 2.04957E-14 |
| Intermediate | CRNDE | 1.17604E-21 | 0.342761 | 0.338 | 0.152 | 1.93154E-17 |
| Intermediate | PDLIM1 | 2.88219E-16 | 0.342699 | 0.729 | 0.54 | 4.73371E-12 |
| Pericytes | TXNIP | 0.006731148 | 0.342628 | 0.655 | 0.659 | 1 |
| Terminal | SRP14 | 8.3082E-16 | 0.342529 | 0.989 | 0.949 | 1.36454E-11 |
| Contractile | MRPS23 | 7.58586E-10 | 0.342393 | 0.313 | 0.2 | 1.2459E-05 |
| Fibroblasts | FAM49A | 1.23747E-53 | 0.342299 | 0.133 | 0.012 | 2.03242E-49 |
| RERGL_Cells | HNRNPUL1 | 5.54302E-05 | 0.342293 | 0.31 | 0.254 | 0.910384976 |
| RERGL_Cells | PRDX3 | 7.72624E-06 | 0.342283 | 0.457 | 0.418 | 0.126895692 |
| Contractile | ADARB1 | 5.83655E-19 | 0.342253 | 0.152 | 0.049 | 9.58596E-15 |
| Contractile | MRPL55 | 7.60924E-10 | 0.342202 | 0.457 | 0.332 | 1.24974E-05 |
| Intermediate | SLC25A4 | 9.33444E-21 | 0.342053 | 0.664 | 0.411 | 1.53309E-16 |
| Terminal | ERGIC3 | 2.96253E-06 | 0.34201 | 0.532 | 0.459 | 0.048656541 |
| Pericytes | STRN3 | 5.14707E-05 | 0.341988 | 0.205 | 0.147 | 0.845355126 |
| Contractile | PSMC5 | 3.86885E-10 | 0.341955 | 0.629 | 0.504 | 6.3542E-06 |
| Contractile | NDUFB1 | 1.95823E-08 | 0.341817 | 0.703 | 0.637 | 0.000321619 |
| Pericytes | MOCS1 | 4.37428E-13 | 0.341792 | 0.142 | 0.057 | 7.18432E-09 |
| Pericytes | RELA | 0.000369279 | 0.341756 | 0.264 | 0.217 | 1 |
| Terminal | COL14A1 | 0.000808147 | 0.341672 | 0.471 | 0.414 | 1 |
| Fibroblasts | ANKRD35 | 4.67465E-33 | 0.341604 | 0.261 | 0.1 | 7.67764E-29 |
| RERGL_Cells | SSBP4 | 0.00123232 | 0.341596 | 0.244 | 0.2 | 1 |

| | | | | | | |
|---|---|---|---|---|---|---|
| Intermediate | SPSB1 | 3.02852E-15 | 0.341548 | 0.363 | 0.204 | 4.97404E-11 |
| Fibroblasts | RBP1 | 4.07337E-35 | 0.341516 | 0.27 | 0.1 | 6.6901E-31 |
| Contractile | ANP32A | 1.96486E-07 | 0.341415 | 0.335 | 0.242 | 0.003227089 |
| Pericytes | CALHM5 | 2.59073E-07 | 0.34125 | 0.116 | 0.059 | 0.004255011 |
| Contractile | SORT1 | 1.82978E-13 | 0.341229 | 0.248 | 0.124 | 3.00523E-09 |
| Intermediate | TPT1 | 8.62004E-18 | 0.341196 | 0.989 | 0.982 | 1.41576E-13 |
| Pericytes | COQ10B | 3.2599E-06 | 0.341189 | 0.347 | 0.28 | 0.053540599 |
| Contractile | AC093673 | 3.63151E-16 | 0.341172 | 0.16 | 0.059 | 5.96439E-12 |
| Fibroblasts | NID2 | 3.49584E-47 | 0.341166 | 0.135 | 0.017 | 5.74158E-43 |
| Fibroblasts | DPYD | 2.88851E-42 | 0.341094 | 0.209 | 0.055 | 4.74409E-38 |
| Terminal | EEF1G | 4.82529E-14 | 0.34109 | 0.976 | 0.951 | 7.92505E-10 |
| Contractile | PDK2 | 7.64176E-11 | 0.341085 | 0.242 | 0.134 | 1.25508E-06 |
| Fibroblasts | CAB39L | 1.52239E-28 | 0.341035 | 0.27 | 0.114 | 2.50038E-24 |
| Contractile | JPH2 | 5.51215E-25 | 0.340986 | 0.132 | 0.03 | 9.05315E-21 |
| Intermediate | CNPY2 | 1.46397E-12 | 0.340823 | 0.624 | 0.449 | 2.40442E-08 |
| RERGL_Cells | AP001160 | 2.14547E-08 | 0.340819 | 0.107 | 0.046 | 0.000352371 |
| Contractile | AVPI1 | 1.21445E-12 | 0.340798 | 0.22 | 0.11 | 1.99462E-08 |
| Contractile | POLR2C | 2.27209E-09 | 0.340702 | 0.477 | 0.362 | 3.73168E-05 |
| Terminal | CDH11 | 3.14474E-05 | 0.34067 | 0.24 | 0.17 | 0.516492681 |
| Fibroblasts | GCHFR | 1.62373E-41 | 0.340646 | 0.162 | 0.033 | 2.66682E-37 |
| Fibroblasts | SIL1 | 3.61389E-27 | 0.340574 | 0.37 | 0.186 | 5.93544E-23 |
| Terminal | IGLC2 | 0.005506197 | 0.340367 | 0.209 | 0.16 | 1 |
| Pericytes | HNRNPH1 | 0.005387601 | 0.340238 | 0.431 | 0.411 | 1 |
| Pericytes | CWC25 | 1.61411E-06 | 0.34014 | 0.255 | 0.181 | 0.026510133 |
| Fibroblasts | ODF2L | 1.42246E-34 | 0.339963 | 0.207 | 0.066 | 2.33625E-30 |
| Pericytes | UBE2A | 0.000125229 | 0.339888 | 0.347 | 0.301 | 1 |
| Contractile | CLTA | 3.39977E-11 | 0.339756 | 0.705 | 0.593 | 5.58378E-07 |
| Contractile | KTN1 | 6.61849E-09 | 0.339743 | 0.661 | 0.555 | 0.000108702 |
| Pericytes | CAMK2D | 2.13577E-07 | 0.339633 | 0.178 | 0.106 | 0.003507788 |
| Pericytes | CHN1 | 2.57283E-13 | 0.339584 | 0.111 | 0.038 | 4.22561E-09 |
| Pericytes | CSNK1A1 | 2.011E-06 | 0.339433 | 0.591 | 0.559 | 0.033028607 |
| Terminal | SUN2 | 1.12426E-06 | 0.339386 | 0.553 | 0.458 | 0.018464894 |
| Fibroblasts | TMEM150 | 6.9098E-58 | 0.339332 | 0.113 | 0.004 | 1.13486E-53 |
| Pericytes | CYP7B1 | 3.02362E-10 | 0.339324 | 0.149 | 0.071 | 4.96599E-06 |
| Fibroblasts | SLC16A4 | 2.83377E-35 | 0.339276 | 0.208 | 0.065 | 4.65419E-31 |
| RERGL_Cells | NDUFB4 | 3.65053E-10 | 0.339197 | 0.716 | 0.687 | 5.99562E-06 |
| RERGL_Cells | NDUFV2 | 5.1374E-07 | 0.339186 | 0.551 | 0.515 | 0.008437659 |
| Fibroblasts | C5orf15 | 5.84417E-26 | 0.339127 | 0.445 | 0.245 | 9.59846E-22 |
| Fibroblasts | COLGALT2 | 5.07612E-33 | 0.339031 | 0.102 | 0.015 | 8.33702E-29 |
| Intermediate | SMOX | 9.04235E-22 | 0.338934 | 0.144 | 0.041 | 1.48512E-17 |
| Contractile | UBL5 | 9.64742E-13 | 0.338844 | 0.86 | 0.799 | 1.58449E-08 |
| Contractile | SRP9 | 4.84032E-10 | 0.338402 | 0.681 | 0.599 | 7.94974E-06 |
| Terminal | PLD3 | 4.93913E-08 | 0.338333 | 0.647 | 0.561 | 0.000811203 |
| Contractile | ATP1B1 | 6.58325E-07 | 0.338157 | 0.238 | 0.155 | 0.010812322 |
| RERGL_Cells | SELENBP1 | 0.002434593 | 0.338112 | 0.297 | 0.263 | 1 |
| Terminal | RILPL2 | 5.00163E-06 | 0.33801 | 0.322 | 0.234 | 0.082146701 |
| Fibroblasts | ARF4 | 2.2671E-20 | 0.337969 | 0.799 | 0.616 | 3.72349E-16 |
| RERGL_Cells | LSM14A | 0.000405067 | 0.337956 | 0.353 | 0.316 | 1 |
| RERGL_Cells | IGLC1 | 0.00279313 | 0.337953 | 0.184 | 0.138 | 1 |
| Fibroblasts | PHLDB1 | 2.26198E-21 | 0.337791 | 0.264 | 0.128 | 3.71508E-17 |
| Contractile | PSMA2 | 1.30611E-08 | 0.337767 | 0.655 | 0.571 | 0.000214515 |
| RERGL_Cells | PSMC5 | 4.01924E-05 | 0.33776 | 0.53 | 0.52 | 0.660119229 |
| Fibroblasts | HACD3 | 5.39236E-27 | 0.337647 | 0.358 | 0.179 | 8.8564E-23 |
| Contractile | S100A6 | 2.85005E-32 | 0.337579 | 1 | 0.994 | 4.68092E-28 |
| Fibroblasts | RAB8B | 1.6915E-31 | 0.337493 | 0.194 | 0.063 | 2.77812E-27 |
| Terminal | MYO1D | 4.80728E-07 | 0.337419 | 0.207 | 0.126 | 0.007895479 |

| | | | | | | |
|---|---|---|---|---|---|---|
| Pericytes | CDC42EP1 | 0.000369656 | 0.33735 | 0.407 | 0.367 | 1 |
| Contractile | NDUFA12 | 1.46398E-10 | 0.337338 | 0.575 | 0.447 | 2.40444E-06 |
| Intermediate | MYOZ1 | 1.02916E-20 | 0.337092 | 0.333 | 0.15 | 1.69029E-16 |
| Contractile | LSM3 | 8.62727E-10 | 0.336853 | 0.561 | 0.438 | 1.41694E-05 |
| RERGL_Cells | POLR2J3 | 0.002694052 | 0.336838 | 0.207 | 0.167 | 1 |
| Pericytes | ETF1 | 4.23221E-06 | 0.336739 | 0.289 | 0.219 | 0.069509846 |
| Pericytes | FKBP4 | 2.29814E-05 | 0.336674 | 0.276 | 0.211 | 0.377447208 |
| Pericytes | CMTM7 | 2.37025E-12 | 0.336563 | 0.116 | 0.044 | 3.8929E-08 |
| Terminal | AIG1 | 2.36921E-06 | 0.336547 | 0.349 | 0.255 | 0.038911958 |
| Intermediate | DAD1 | 1.17011E-13 | 0.336494 | 0.815 | 0.698 | 1.92178E-09 |
| Fibroblasts | TLNRD1 | 1.33518E-29 | 0.336486 | 0.241 | 0.093 | 2.1929E-25 |
| Contractile | SYNPO | 8.97724E-10 | 0.336396 | 0.244 | 0.14 | 1.47442E-05 |
| Pericytes | POLR2M | 0.000204943 | 0.336369 | 0.196 | 0.145 | 1 |
| RERGL_Cells | C12orf57 | 1.59566E-09 | 0.336169 | 0.808 | 0.8 | 2.62071E-05 |
| Intermediate | ADIRF | 6.67979E-27 | 0.336109 | 0.998 | 0.906 | 1.09709E-22 |
| Fibroblasts | NRP1 | 1.73687E-21 | 0.336018 | 0.346 | 0.183 | 2.85264E-17 |
| Fibroblasts | CFLAR | 3.49938E-23 | 0.335866 | 0.352 | 0.186 | 5.74738E-19 |
| Contractile | DGCR6L | 4.31491E-07 | 0.335821 | 0.329 | 0.236 | 0.0070868 |
| Contractile | TPD52L1 | 2.40802E-09 | 0.335775 | 0.206 | 0.114 | 3.95493E-05 |
| Intermediate | SH3BGR | 1.14827E-17 | 0.33572 | 0.308 | 0.146 | 1.88591E-13 |
| Fibroblasts | RASSF8-AS | 2.11377E-35 | 0.335592 | 0.17 | 0.045 | 3.47166E-31 |
| Terminal | RHOC | 5.61904E-09 | 0.335557 | 0.782 | 0.727 | 9.22871E-05 |
| Terminal | PRPS1 | 4.11641E-07 | 0.33551 | 0.218 | 0.137 | 0.006760795 |
| Pericytes | RBM3 | 2.2381E-08 | 0.335392 | 0.729 | 0.712 | 0.000367586 |
| Terminal | CFAP36 | 2.53212E-05 | 0.33538 | 0.318 | 0.24 | 0.415875309 |
| Pericytes | H2AFX | 4.48289E-05 | 0.335239 | 0.127 | 0.079 | 0.736269642 |
| Contractile | DEXI | 5.13179E-07 | 0.335213 | 0.345 | 0.253 | 0.008428446 |
| Contractile | EEF2 | 5.86153E-16 | 0.335209 | 0.972 | 0.942 | 9.62698E-12 |
| Pericytes | AEBP2 | 2.03695E-08 | 0.3352 | 0.167 | 0.092 | 0.000334549 |
| Contractile | ARHGEF9 | 3.70122E-12 | 0.335087 | 0.19 | 0.09 | 6.07888E-08 |
| Intermediate | KCNA5 | 3.99105E-17 | 0.335063 | 0.35 | 0.173 | 6.55489E-13 |
| Contractile | SURF1 | 8.12481E-09 | 0.334967 | 0.339 | 0.228 | 0.000133442 |
| RERGL_Cells | PARL | 0.002956399 | 0.334802 | 0.274 | 0.237 | 1 |
| Pericytes | ABCC9 | 1.62854E-05 | 0.334699 | 0.156 | 0.1 | 0.267470624 |
| Contractile | CCT3 | 6.47844E-10 | 0.334616 | 0.571 | 0.449 | 1.06402E-05 |
| Fibroblasts | ST3GAL4 | 2.6103E-21 | 0.334612 | 0.337 | 0.182 | 4.28715E-17 |
| Contractile | DNAJB6 | 2.63691E-10 | 0.334529 | 0.661 | 0.551 | 4.33086E-06 |
| RERGL_Cells | PSMC3 | 4.13565E-05 | 0.334437 | 0.504 | 0.487 | 0.679239703 |
| Fibroblasts | TGIF1 | 5.38644E-23 | 0.334425 | 0.281 | 0.135 | 8.84669E-19 |
| Pericytes | NEXN | 1.12032E-09 | 0.334246 | 0.629 | 0.502 | 1.84002E-05 |
| Intermediate | PLD3 | 1.25989E-13 | 0.334036 | 0.738 | 0.543 | 2.06925E-09 |
| Fibroblasts | PHYKPL | 4.0534E-30 | 0.334032 | 0.27 | 0.11 | 6.6573E-26 |
| Fibroblasts | CREG1 | 1.63779E-23 | 0.334023 | 0.509 | 0.307 | 2.68991E-19 |
| Terminal | EDF1 | 6.66665E-11 | 0.334016 | 0.861 | 0.805 | 1.09493E-06 |
| Fibroblasts | SLC39A7 | 6.42423E-20 | 0.333796 | 0.465 | 0.29 | 1.05512E-15 |
| RERGL_Cells | GRAMD2B | 3.84647E-06 | 0.333689 | 0.13 | 0.072 | 0.063174365 |
| Terminal | C7orf50 | 1.25986E-05 | 0.333574 | 0.471 | 0.393 | 0.206920021 |
| Contractile | NUPR1 | 9.06106E-15 | 0.333566 | 0.916 | 0.827 | 1.48819E-10 |
| Intermediate | SP100 | 5.26858E-15 | 0.33356 | 0.471 | 0.281 | 8.65312E-11 |
| RERGL_Cells | RBBP7 | 1.22866E-05 | 0.333508 | 0.427 | 0.378 | 0.201794619 |
| Fibroblasts | OSTC | 9.85924E-21 | 0.333472 | 0.669 | 0.464 | 1.61928E-16 |
| Fibroblasts | CNOT8 | 1.23718E-26 | 0.333294 | 0.269 | 0.118 | 2.03195E-22 |
| Fibroblasts | SCP2 | 2.34959E-22 | 0.333258 | 0.753 | 0.542 | 3.85897E-18 |
| Fibroblasts | GPCPD1 | 8.26994E-34 | 0.333228 | 0.181 | 0.052 | 1.35825E-29 |
| Terminal | MAGED2 | 1.18174E-08 | 0.333178 | 0.715 | 0.627 | 0.00019409 |
| Fibroblasts | ELMSAN1 | 1.27266E-24 | 0.333176 | 0.231 | 0.097 | 2.09022E-20 |

| | | | | | | |
|---|---|---|---|---|---|---|
| Contractile | LETMD1 | 4.24111E-09 | 0.333037 | 0.259 | 0.158 | 6.96559E-05 |
| Fibroblasts | TNS2 | 1.26126E-22 | 0.332875 | 0.408 | 0.228 | 2.0715E-18 |
| Terminal | CTSB | 6.96229E-07 | 0.332857 | 0.486 | 0.393 | 0.011434867 |
| RERGL_Cells | SYF2 | 1.22899E-06 | 0.332841 | 0.56 | 0.508 | 0.020184954 |
| Terminal | WDR83OS | 5.18199E-07 | 0.332835 | 0.73 | 0.662 | 0.008510896 |
| Pericytes | MORF4L2 | 6.69177E-07 | 0.332785 | 0.735 | 0.731 | 0.010990565 |
| Pericytes | IFRD1 | 0.00341353 | 0.33269 | 0.32 | 0.286 | 1 |
| Pericytes | RELL1 | 3.68696E-07 | 0.332646 | 0.262 | 0.179 | 0.006055466 |
| Fibroblasts | SH3BGRL3 | 2.87658E-16 | 0.332624 | 0.768 | 0.606 | 4.72449E-12 |
| Fibroblasts | SOAT1 | 3.12269E-35 | 0.33245 | 0.202 | 0.061 | 5.12871E-31 |
| Fibroblasts | ZCCHC24 | 1.68451E-34 | 0.332405 | 0.185 | 0.054 | 2.76663E-30 |
| Pericytes | FLOT1 | 1.25304E-05 | 0.332256 | 0.455 | 0.406 | 0.205799446 |
| Contractile | MLF2 | 2.59219E-09 | 0.332218 | 0.575 | 0.473 | 4.25742E-05 |
| Terminal | DYNLL1 | 1.87347E-11 | 0.332186 | 0.917 | 0.829 | 3.07698E-07 |
| Pericytes | AMD1 | 0.000127283 | 0.332073 | 0.375 | 0.324 | 1 |
| Terminal | ERP29 | 2.00836E-08 | 0.332065 | 0.521 | 0.417 | 0.000329853 |
| Fibroblasts | COL6A1 | 6.67044E-24 | 0.332013 | 0.861 | 0.692 | 1.09555E-19 |
| Contractile | YWHAQ | 2.89542E-13 | 0.331982 | 0.812 | 0.696 | 4.75543E-09 |
| RERGL_Cells | SLC25A3 | 8.27823E-11 | 0.331967 | 0.84 | 0.808 | 1.35962E-06 |
| Intermediate | CETN2 | 1.14186E-12 | 0.331798 | 0.473 | 0.304 | 1.87538E-08 |
| Fibroblasts | TPM3 | 3.32866E-25 | 0.33178 | 0.401 | 0.216 | 5.46698E-21 |
| Contractile | MRPS34 | 4.12935E-07 | 0.331771 | 0.341 | 0.244 | 0.006782051 |
| RERGL_Cells | STRN3 | 8.32391E-05 | 0.33175 | 0.209 | 0.148 | 1 |
| RERGL_Cells | UQCRC1 | 0.000251778 | 0.331738 | 0.4 | 0.364 | 1 |
| Contractile | PHGDH | 1.40739E-06 | 0.331733 | 0.339 | 0.244 | 0.023115045 |
| Intermediate | NEXN | 7.23425E-15 | 0.331669 | 0.703 | 0.49 | 1.18815E-10 |
| Fibroblasts | RARG | 2.62584E-32 | 0.331629 | 0.19 | 0.059 | 4.31268E-28 |
| Contractile | EEF1G | 2.27422E-20 | 0.331629 | 0.984 | 0.949 | 3.73517E-16 |
| Contractile | OCIAD2 | 6.21346E-13 | 0.331553 | 0.198 | 0.092 | 1.0205E-08 |
| RERGL_Cells | BTN2A1 | 1.12634E-06 | 0.331494 | 0.16 | 0.093 | 0.018498968 |
| Intermediate | PSAP | 5.27617E-18 | 0.331482 | 0.966 | 0.906 | 8.66559E-14 |
| Contractile | PHB | 2.06639E-07 | 0.331339 | 0.487 | 0.397 | 0.003393842 |
| Contractile | SSB | 2.72233E-11 | 0.331286 | 0.695 | 0.586 | 4.47115E-07 |
| RERGL_Cells | SLC8B1 | 8.65373E-08 | 0.331223 | 0.135 | 0.068 | 0.001421289 |
| RERGL_Cells | FMC1 | 0.000932969 | 0.330985 | 0.301 | 0.26 | 1 |
| Fibroblasts | NINJ2 | 2.26771E-46 | 0.330849 | 0.124 | 0.014 | 3.72449E-42 |
| Fibroblasts | RGMA | 6.59779E-33 | 0.330749 | 0.336 | 0.143 | 1.08362E-28 |
| Contractile | RBBP7 | 2.7235E-07 | 0.330715 | 0.463 | 0.371 | 0.004473076 |
| Fibroblasts | CHD1 | 1.1925E-25 | 0.330536 | 0.274 | 0.122 | 1.95856E-21 |
| Pericytes | RCN1 | 5.5395E-07 | 0.330499 | 0.56 | 0.52 | 0.009098074 |
| Intermediate | TFPI2 | 1.78939E-15 | 0.330394 | 0.157 | 0.059 | 2.9389E-11 |
| Fibroblasts | IFNGR2 | 6.65271E-26 | 0.330365 | 0.35 | 0.175 | 1.09264E-21 |
| Contractile | HMGN3 | 1.30667E-09 | 0.330354 | 0.617 | 0.498 | 2.14607E-05 |
| Terminal | MRPL49 | 2.43347E-05 | 0.330308 | 0.229 | 0.158 | 0.399673585 |
| Terminal | SSPN | 6.14244E-06 | 0.330233 | 0.547 | 0.474 | 0.100883486 |
| Intermediate | ANKH | 3.51295E-11 | 0.330076 | 0.303 | 0.177 | 5.76967E-07 |
| Contractile | RTL8A | 4.5098E-07 | 0.329967 | 0.401 | 0.306 | 0.007406892 |
| Contractile | COX7A1 | 2.65564E-11 | 0.329926 | 0.747 | 0.644 | 4.36162E-07 |
| Terminal | YIPF3 | 4.5657E-06 | 0.32989 | 0.444 | 0.371 | 0.074987068 |
| Terminal | NCOR1 | 0.002373333 | 0.329809 | 0.394 | 0.354 | 1 |
| RERGL_Cells | NDUFA2 | 3.67101E-05 | 0.329421 | 0.474 | 0.449 | 0.602926887 |
| RERGL_Cells | GAMT | 0.003784408 | 0.329199 | 0.233 | 0.195 | 1 |
| Contractile | RTN1 | 8.40595E-22 | 0.329199 | 0.118 | 0.027 | 1.38059E-17 |
| Contractile | C1QA | 1.32159E-09 | 0.329193 | 0.192 | 0.101 | 2.17058E-05 |
| Pericytes | SLC25A24 | 8.85932E-07 | 0.329179 | 0.196 | 0.126 | 0.014550551 |
| Intermediate | ARPC1B | 5.30173E-08 | 0.329106 | 0.639 | 0.523 | 0.000870756 |

| | | | | | | |
|---|---|---|---|---|---|---|
| Pericytes | OSR2 | 0.000443731 | 0.329058 | 0.105 | 0.065 | 1 |
| Terminal | ENO1 | 1.22425E-11 | 0.329041 | 0.847 | 0.767 | 2.01071E-07 |
| Terminal | HSPB7 | 0.001834743 | 0.328935 | 0.283 | 0.227 | 1 |
| Fibroblasts | APOL1 | 3.69669E-29 | 0.328909 | 0.196 | 0.068 | 6.07144E-25 |
| Intermediate | ABHD2 | 8.74414E-16 | 0.328855 | 0.213 | 0.092 | 1.43614E-11 |
| Fibroblasts | FAM114A1 | 2.41923E-21 | 0.328844 | 0.414 | 0.241 | 3.97335E-17 |
| Intermediate | KDELR3 | 7.93845E-12 | 0.328792 | 0.301 | 0.175 | 1.30381E-07 |
| Fibroblasts | AZI2 | 2.30634E-22 | 0.328705 | 0.391 | 0.215 | 3.78794E-18 |
| Pericytes | RHOC | 1.96726E-09 | 0.32844 | 0.738 | 0.733 | 3.23103E-05 |
| Fibroblasts | GNS | 1.15725E-28 | 0.328433 | 0.208 | 0.076 | 1.90067E-24 |
| Terminal | TMEM219 | 2.05204E-06 | 0.328426 | 0.492 | 0.407 | 0.033702776 |
| RERGL_Cells | NDUFC1 | 5.54526E-06 | 0.328387 | 0.577 | 0.56 | 0.091075339 |
| Contractile | ARPC2 | 9.15598E-15 | 0.328377 | 0.82 | 0.765 | 1.50378E-10 |
| Contractile | TIMM8B | 1.22199E-07 | 0.328321 | 0.437 | 0.334 | 0.002006989 |
| Pericytes | SERPING1 | 3.7454E-21 | 0.328197 | 0.967 | 0.912 | 6.15145E-17 |
| Pericytes | ACVRL1 | 1.85695E-05 | 0.328149 | 0.191 | 0.131 | 0.304985906 |
| Terminal | ATP6V1F | 8.18216E-07 | 0.328143 | 0.523 | 0.439 | 0.013438382 |
| Pericytes | MPDU1 | 0.000103775 | 0.328071 | 0.169 | 0.119 | 1 |
| RERGL_Cells | ATP5IF1 | 3.96313E-06 | 0.328066 | 0.583 | 0.556 | 0.06509048 |
| Terminal | NAP1L1 | 0.000100248 | 0.328065 | 0.715 | 0.669 | 1 |
| Intermediate | PARK7 | 6.48504E-18 | 0.32803 | 0.888 | 0.797 | 1.0651E-13 |
| Contractile | COX5A | 1.12716E-07 | 0.327961 | 0.481 | 0.39 | 0.001851241 |
| Fibroblasts | BCHE | 2.28471E-58 | 0.327891 | 0.108 | 0.002 | 3.75241E-54 |
| Fibroblasts | SNED1 | 9.69936E-37 | 0.327833 | 0.132 | 0.025 | 1.59302E-32 |
| Pericytes | PFKP | 1.61705E-06 | 0.327791 | 0.198 | 0.128 | 0.026558488 |
| Contractile | HMGN5 | 7.74754E-08 | 0.327734 | 0.172 | 0.095 | 0.001272455 |
| Terminal | RNF7 | 2.78936E-05 | 0.327705 | 0.516 | 0.454 | 0.458124487 |
| Pericytes | SNHG3 | 1.18631E-05 | 0.327696 | 0.158 | 0.101 | 0.19483993 |
| Terminal | STX8 | 0.000618775 | 0.327624 | 0.29 | 0.23 | 1 |
| Fibroblasts | CHST12 | 1.0133E-26 | 0.327476 | 0.329 | 0.158 | 1.66425E-22 |
| RERGL_Cells | PSMF1 | 0.000531967 | 0.327451 | 0.391 | 0.36 | 1 |
| Terminal | DUSP26 | 7.10502E-06 | 0.327417 | 0.131 | 0.073 | 0.116692899 |
| Intermediate | NDUFC1 | 6.36041E-13 | 0.327382 | 0.71 | 0.537 | 1.04463E-08 |
| Pericytes | DDX18 | 3.85341E-06 | 0.327249 | 0.505 | 0.453 | 0.063288382 |
| Intermediate | LNP1 | 3.15878E-15 | 0.327013 | 0.23 | 0.105 | 5.18799E-11 |
| Intermediate | PLIN3 | 6.95924E-13 | 0.327007 | 0.404 | 0.247 | 1.14299E-08 |
| Pericytes | TAF7 | 8.37002E-06 | 0.32685 | 0.551 | 0.519 | 0.137469274 |
| Pericytes | CLTB | 1.53506E-07 | 0.326463 | 0.596 | 0.566 | 0.00252118 |
| Terminal | IRS2 | 4.07382E-07 | 0.326438 | 0.159 | 0.088 | 0.006690849 |
| RERGL_Cells | SLC8A1 | 9.05565E-10 | 0.326398 | 0.115 | 0.048 | 1.4873E-05 |
| Contractile | ANXA7 | 8.79512E-08 | 0.326376 | 0.665 | 0.578 | 0.00144451 |
| Contractile | CDKN2C | 1.0134E-05 | 0.326344 | 0.242 | 0.168 | 0.166440316 |
| Terminal | ANTXR1 | 1.3963E-06 | 0.326093 | 0.329 | 0.235 | 0.022932838 |
| RERGL_Cells | PPP2R2B | 2.66414E-11 | 0.326056 | 0.135 | 0.055 | 4.37558E-07 |
| RERGL_Cells | HIPK2 | 7.71866E-08 | 0.326021 | 0.105 | 0.047 | 0.001267714 |
| RERGL_Cells | MDH1 | 0.000737638 | 0.325942 | 0.453 | 0.437 | 1 |
| Fibroblasts | ZBTB20 | 6.51475E-26 | 0.325931 | 0.485 | 0.271 | 1.06998E-21 |
| Pericytes | PELI2 | 2.92237E-16 | 0.325826 | 0.105 | 0.03 | 4.7997E-12 |
| RERGL_Cells | NDN | 0.000300914 | 0.325823 | 0.534 | 0.532 | 1 |
| Intermediate | MTRNR2L8 | 4.78967E-09 | 0.325793 | 0.241 | 0.14 | 7.86655E-05 |
| Intermediate | SNHG14 | 3.05494E-13 | 0.325682 | 0.249 | 0.127 | 5.01743E-09 |
| Contractile | SLURP2 | 3.97523E-22 | 0.325673 | 0.132 | 0.033 | 6.52892E-18 |
| Intermediate | PMEPA1 | 1.47521E-21 | 0.325637 | 0.247 | 0.096 | 2.42288E-17 |
| Pericytes | HMGN4 | 6.34412E-06 | 0.325609 | 0.376 | 0.31 | 0.104195798 |
| Contractile | TWSG1 | 3.10749E-07 | 0.325571 | 0.383 | 0.286 | 0.005103748 |
| Intermediate | SOD1 | 8.26089E-15 | 0.325511 | 0.925 | 0.846 | 1.35677E-10 |

| | | | | | | |
|---|---|---|---|---|---|---|
| Contractile | MEAF6 | 6.29624E-09 | 0.325451 | 0.521 | 0.404 | 0.000103409 |
| Pericytes | SYDE1 | 0.000488973 | 0.325428 | 0.196 | 0.148 | 1 |
| RERGL_Cells | RBBP4 | 0.005444108 | 0.3254 | 0.286 | 0.258 | 1 |
| RERGL_Cells | IQCK | 4.62597E-07 | 0.325396 | 0.113 | 0.056 | 0.007597693 |
| Fibroblasts | TRPS1 | 1.35749E-27 | 0.325381 | 0.189 | 0.066 | 2.22954E-23 |
| Fibroblasts | CTSO | 6.58874E-23 | 0.325333 | 0.355 | 0.191 | 1.08213E-18 |
| Terminal | FOXS1 | 4.85824E-09 | 0.325321 | 0.275 | 0.162 | 7.97917E-05 |
| Intermediate | MYLIP | 1.10513E-13 | 0.325278 | 0.406 | 0.235 | 1.81507E-09 |
| Fibroblasts | CLU | 3.67528E-51 | 0.325145 | 0.956 | 0.77 | 6.03629E-47 |
| Contractile | ACTR2 | 1.30322E-07 | 0.325136 | 0.395 | 0.292 | 0.002140401 |
| Terminal | NFIC | 2.09545E-05 | 0.325012 | 0.545 | 0.493 | 0.344155987 |
| RERGL_Cells | PARP1 | 0.000162392 | 0.324974 | 0.229 | 0.175 | 1 |
| Contractile | EVA1C | 7.13301E-10 | 0.32478 | 0.216 | 0.118 | 1.17153E-05 |
| Pericytes | TSPYL1 | 0.002161256 | 0.324733 | 0.402 | 0.375 | 1 |
| Contractile | ZEB1 | 2.91947E-08 | 0.324599 | 0.261 | 0.166 | 0.000479494 |
| Contractile | MSC | 9.11004E-08 | 0.324437 | 0.327 | 0.223 | 0.001496233 |
| RERGL_Cells | UBE2S | 4.02513E-08 | 0.324425 | 0.137 | 0.068 | 0.000661087 |
| Intermediate | ASAH1 | 5.94783E-10 | 0.324358 | 0.684 | 0.532 | 9.76871E-06 |
| Pericytes | EBF1 | 5.83598E-05 | 0.324354 | 0.431 | 0.375 | 0.958500591 |
| RERGL_Cells | TAX1BP1 | 1.1095E-05 | 0.324119 | 0.534 | 0.505 | 0.182224671 |
| RERGL_Cells | HAGH | 0.001017663 | 0.324079 | 0.318 | 0.277 | 1 |
| Contractile | SDHB | 2.33851E-08 | 0.324072 | 0.465 | 0.356 | 0.000384076 |
| Terminal | SPSB1 | 1.11597E-06 | 0.324046 | 0.307 | 0.216 | 0.018328731 |
| RERGL_Cells | POLR2I | 0.00013133 | 0.323917 | 0.455 | 0.428 | 1 |
| Intermediate | TOMM7 | 4.95506E-19 | 0.323787 | 0.946 | 0.906 | 8.13819E-15 |
| Pericytes | TACC1 | 0.000199707 | 0.323686 | 0.453 | 0.418 | 1 |
| Contractile | LMO7 | 4.11866E-09 | 0.32354 | 0.162 | 0.082 | 6.76448E-05 |
| Pericytes | SMPDL3A | 3.28462E-07 | 0.323536 | 0.138 | 0.077 | 0.005394659 |
| Contractile | MTHFD1 | 7.38715E-09 | 0.323401 | 0.21 | 0.12 | 0.000121327 |
| Intermediate | FUCA1 | 3.65715E-16 | 0.323281 | 0.321 | 0.165 | 6.0065E-12 |
| Fibroblasts | IRF1 | 5.85249E-20 | 0.323212 | 0.286 | 0.146 | 9.61212E-16 |
| Pericytes | MMP14 | 4.83991E-07 | 0.322796 | 0.344 | 0.263 | 0.007949075 |
| Contractile | AK1 | 5.98945E-06 | 0.322711 | 0.573 | 0.503 | 0.098370788 |
| Contractile | USO1 | 4.35558E-05 | 0.322677 | 0.369 | 0.298 | 0.715360723 |
| Pericytes | RHEB | 2.77704E-07 | 0.322627 | 0.658 | 0.649 | 0.004561018 |
| Pericytes | FUBP1 | 2.22067E-06 | 0.322615 | 0.236 | 0.163 | 0.0364723 |
| RERGL_Cells | PET100 | 6.18973E-07 | 0.322571 | 0.526 | 0.481 | 0.010166011 |
| RERGL_Cells | AC092164 | 0.000475814 | 0.322322 | 0.194 | 0.139 | 1 |
| RERGL_Cells | SYNM | 2.55131E-08 | 0.322247 | 0.156 | 0.08 | 0.000419027 |
| RERGL_Cells | PFKM | 0.00292694 | 0.321788 | 0.201 | 0.16 | 1 |
| RERGL_Cells | IRF1 | 8.01807E-05 | 0.321729 | 0.244 | 0.18 | 1 |
| Fibroblasts | ZFP36L1 | 2.89616E-31 | 0.321632 | 0.775 | 0.537 | 4.75665E-27 |
| Fibroblasts | ATRAID | 1.63954E-18 | 0.321608 | 0.752 | 0.569 | 2.69278E-14 |
| Contractile | LIMCH1 | 6.62638E-07 | 0.321603 | 0.285 | 0.194 | 0.010883161 |
| Contractile | PSMA6 | 8.38931E-08 | 0.321598 | 0.513 | 0.413 | 0.00137786 |
| Contractile | PSMB4 | 1.1051E-07 | 0.321111 | 0.433 | 0.33 | 0.001815015 |
| Terminal | MPC2 | 9.89442E-06 | 0.321088 | 0.438 | 0.355 | 0.162505949 |
| Terminal | PRAF2 | 1.98887E-05 | 0.321075 | 0.333 | 0.259 | 0.32665188 |
| RERGL_Cells | SRSF4 | 0.005565663 | 0.321034 | 0.359 | 0.345 | 1 |
| Contractile | TCEA2 | 2.83943E-06 | 0.321014 | 0.323 | 0.238 | 0.046634861 |
| Contractile | PPM1B | 5.05923E-08 | 0.321009 | 0.283 | 0.186 | 0.000830928 |
| Pericytes | FNBP4 | 0.000377108 | 0.320985 | 0.216 | 0.166 | 1 |
| RERGL_Cells | ANAPC15 | 0.001982253 | 0.320696 | 0.173 | 0.131 | 1 |
| Contractile | MACF1 | 9.85521E-07 | 0.320678 | 0.267 | 0.182 | 0.016186201 |
| Contractile | RAB13 | 2.00125E-09 | 0.320668 | 0.489 | 0.355 | 3.28685E-05 |
| RERGL_Cells | RBM23 | 0.000731723 | 0.320554 | 0.201 | 0.154 | 1 |

| | | | | | | |
|---|---|---|---|---|---|---|
| Contractile | MDH1 | 7.75254E-08 | 0.320509 | 0.523 | 0.426 | 0.001273278 |
| RERGL_Cells | JAK1 | 0.00200912 | 0.320398 | 0.323 | 0.288 | 1 |
| Pericytes | PAIP2 | 5.86516E-07 | 0.320394 | 0.596 | 0.547 | 0.009632942 |
| RERGL_Cells | SBDS | 8.39864E-09 | 0.320321 | 0.797 | 0.791 | 0.000137939 |
| RERGL_Cells | YPEL5 | 4.12376E-05 | 0.320278 | 0.41 | 0.363 | 0.677285905 |
| Fibroblasts | LIFR | 1.03747E-24 | 0.320174 | 0.139 | 0.042 | 1.70394E-20 |
| Contractile | PSMB7 | 1.90608E-08 | 0.320113 | 0.601 | 0.5 | 0.000313055 |
| Terminal | FKBP7 | 1.37878E-05 | 0.320089 | 0.325 | 0.244 | 0.22645146 |
| RERGL_Cells | SVBP | 0.001749724 | 0.320003 | 0.233 | 0.195 | 1 |
| Terminal | CERCAM | 4.58617E-05 | 0.320002 | 0.325 | 0.253 | 0.753233229 |
| RERGL_Cells | ENAH | 0.001362948 | 0.319992 | 0.252 | 0.207 | 1 |
| Intermediate | SARAF | 6.16315E-13 | 0.319866 | 0.813 | 0.681 | 1.01224E-08 |
| Terminal | CRABP2 | 2.81664E-11 | 0.319796 | 0.12 | 0.045 | 4.62605E-07 |
| Intermediate | PLAAT3 | 8.18712E-12 | 0.319589 | 0.469 | 0.311 | 1.34465E-07 |
| Pericytes | LAP3 | 0.004656797 | 0.319169 | 0.307 | 0.275 | 1 |
| Terminal | TCEAL2 | 1.0167E-08 | 0.319156 | 0.214 | 0.118 | 0.000166982 |
| Fibroblasts | ACSL4 | 1.01795E-23 | 0.318787 | 0.131 | 0.039 | 1.67187E-19 |
| RERGL_Cells | NDUFA6 | 2.39886E-05 | 0.318671 | 0.485 | 0.455 | 0.393989207 |
| Terminal | CALU | 1.46056E-05 | 0.318438 | 0.58 | 0.527 | 0.239881809 |
| Fibroblasts | PARVB | 2.09225E-41 | 0.318364 | 0.153 | 0.029 | 3.43631E-37 |
| Pericytes | EMG1 | 4.42444E-05 | 0.318357 | 0.244 | 0.183 | 0.726670758 |
| Fibroblasts | TSPAN8 | 3.0393E-17 | 0.318355 | 0.314 | 0.176 | 4.99174E-13 |
| Contractile | POLR2I | 2.02889E-08 | 0.318293 | 0.533 | 0.415 | 0.000333225 |
| Contractile | ENAH | 9.35524E-09 | 0.318245 | 0.309 | 0.198 | 0.00015365 |
| Fibroblasts | RORA | 3.1343E-19 | 0.318159 | 0.169 | 0.071 | 5.14777E-15 |
| Pericytes | SSBP4 | 7.24246E-05 | 0.318157 | 0.255 | 0.197 | 1 |
| RERGL_Cells | EHD1 | 0.000122146 | 0.318132 | 0.186 | 0.13 | 1 |
| RERGL_Cells | CISD3 | 0.00663464 | 0.318051 | 0.246 | 0.211 | 1 |
| RERGL_Cells | COX20 | 0.000490126 | 0.318046 | 0.344 | 0.297 | 1 |
| Terminal | MOCS2 | 8.12408E-05 | 0.318039 | 0.314 | 0.24 | 1 |
| Intermediate | MRPL51 | 3.75848E-13 | 0.317894 | 0.649 | 0.462 | 6.17293E-09 |
| Terminal | CMTM3 | 2.31886E-05 | 0.317832 | 0.244 | 0.172 | 0.380849621 |
| Contractile | NOP53 | 1.26334E-09 | 0.317756 | 0.806 | 0.735 | 2.07491E-05 |
| Contractile | MIR100HG | 1.61266E-05 | 0.317679 | 0.291 | 0.214 | 0.264863879 |
| Pericytes | DNM1L | 2.48379E-05 | 0.317651 | 0.189 | 0.13 | 0.407937037 |
| Pericytes | MVP | 9.00672E-05 | 0.317562 | 0.435 | 0.393 | 1 |
| Contractile | TBCB | 4.77008E-09 | 0.317527 | 0.599 | 0.486 | 7.83438E-05 |
| Fibroblasts | TAPBP | 2.22664E-23 | 0.317215 | 0.419 | 0.235 | 3.65703E-19 |
| RERGL_Cells | AL135925. | 1.30371E-07 | 0.317165 | 0.1 | 0.045 | 0.002141217 |
| Terminal | PRDX4 | 2.13444E-06 | 0.317163 | 0.582 | 0.513 | 0.035056028 |
| Contractile | PPP1R13L | 3.09855E-09 | 0.317031 | 0.168 | 0.086 | 5.08906E-05 |
| RERGL_Cells | HNRNPK | 2.41044E-12 | 0.317024 | 0.878 | 0.848 | 3.9589E-08 |
| RERGL_Cells | YTHDC1 | 0.000182365 | 0.317004 | 0.241 | 0.186 | 1 |
| RERGL_Cells | SMIM10 | 4.99094E-05 | 0.316923 | 0.265 | 0.197 | 0.819712215 |
| Contractile | FKBP5 | 5.52022E-07 | 0.316843 | 0.293 | 0.199 | 0.009066402 |
| Terminal | ARF1 | 6.29235E-09 | 0.316773 | 0.791 | 0.751 | 0.000103346 |
| RERGL_Cells | GPAT2 | 1.54709E-07 | 0.316713 | 0.103 | 0.046 | 0.002540938 |
| RERGL_Cells | GNB4 | 0.000373965 | 0.316632 | 0.175 | 0.125 | 1 |
| RERGL_Cells | PM20D2 | 4.4266E-07 | 0.316502 | 0.141 | 0.075 | 0.007270246 |
| Contractile | WFS1 | 1.74513E-08 | 0.316382 | 0.265 | 0.167 | 0.000286619 |
| Terminal | HTRA1 | 1.61256E-06 | 0.316371 | 0.575 | 0.501 | 0.026484657 |
| Fibroblasts | AHNAK2 | 7.10957E-30 | 0.316261 | 0.148 | 0.04 | 1.16768E-25 |
| Pericytes | LARP1B | 0.001229627 | 0.316161 | 0.147 | 0.106 | 1 |
| RERGL_Cells | PDLIM5 | 0.000159088 | 0.316131 | 0.402 | 0.358 | 1 |
| Terminal | PSMD8 | 1.96865E-06 | 0.316116 | 0.654 | 0.596 | 0.032333135 |
| RERGL_Cells | AMT | 2.10917E-06 | 0.315881 | 0.118 | 0.061 | 0.03464099 |

| | | | | | | |
|---|---|---|---|---|---|---|
| RERGL_Cells | NAXE | 0.004540036 | 0.315764 | 0.31 | 0.283 | 1 |
| Intermediate | PIK3IP1 | 1.12446E-10 | 0.315685 | 0.267 | 0.152 | 1.84681E-06 |
| RERGL_Cells | PBXIP1 | 9.83111E-05 | 0.315571 | 0.4 | 0.349 | 1 |
| RERGL_Cells | PPP1CB | 3.13997E-09 | 0.315548 | 0.714 | 0.652 | 5.15709E-05 |
| Terminal | IFT22 | 6.52258E-06 | 0.315429 | 0.224 | 0.15 | 0.10712688 |
| Intermediate | PIGT | 1.79201E-10 | 0.31527 | 0.619 | 0.469 | 2.94319E-06 |
| Fibroblasts | SLC35F6 | 3.50521E-29 | 0.315244 | 0.294 | 0.127 | 5.75696E-25 |
| Terminal | PEBP1 | 4.11792E-10 | 0.315212 | 0.893 | 0.872 | 6.76327E-06 |
| Intermediate | HEBP1 | 1.38685E-12 | 0.315009 | 0.471 | 0.302 | 2.27777E-08 |
| Pericytes | HMGCS1 | 6.33207E-06 | 0.315007 | 0.127 | 0.073 | 0.10399792 |
| Terminal | TSR2 | 0.000369216 | 0.314989 | 0.342 | 0.285 | 1 |
| Fibroblasts | ANGPTL4 | 5.00253E-20 | 0.314527 | 0.467 | 0.281 | 8.21616E-16 |
| Pericytes | CDC42 | 7.37063E-09 | 0.314495 | 0.705 | 0.666 | 0.000121055 |
| Contractile | SELENOK | 2.6709E-10 | 0.314477 | 0.669 | 0.544 | 4.38668E-06 |
| Terminal | SAP18 | 2.59411E-07 | 0.314457 | 0.793 | 0.745 | 0.00426056 |
| Intermediate | CDK2AP2 | 4.84981E-10 | 0.314452 | 0.381 | 0.243 | 7.96533E-06 |
| Fibroblasts | SMPDL3A | 1.82176E-37 | 0.314449 | 0.18 | 0.046 | 2.99206E-33 |
| Fibroblasts | TNNT3 | 1.3105E-46 | 0.314326 | 0.102 | 0.006 | 2.15237E-42 |
| RERGL_Cells | PCBP2 | 8.43366E-06 | 0.314271 | 0.637 | 0.628 | 0.138514433 |
| Terminal | ECRG4 | 1.50797E-12 | 0.314227 | 0.749 | 0.593 | 2.47669E-08 |
| Contractile | PSMA7 | 3.61852E-14 | 0.314224 | 0.898 | 0.844 | 5.94305E-10 |
| Contractile | NSFL1C | 3.90522E-09 | 0.314158 | 0.369 | 0.251 | 6.41394E-05 |
| Pericytes | UBE2D1 | 0.000884195 | 0.314128 | 0.258 | 0.212 | 1 |
| Contractile | MYEF2 | 4.19487E-17 | 0.314062 | 0.13 | 0.04 | 6.88966E-13 |
| Intermediate | AC147651 | 8.65719E-19 | 0.313838 | 0.174 | 0.061 | 1.42186E-14 |
| Fibroblasts | GNPDA2 | 1.45782E-29 | 0.313798 | 0.265 | 0.108 | 2.39432E-25 |
| Pericytes | POR | 0.000183525 | 0.313559 | 0.273 | 0.217 | 1 |
| Contractile | MTMR11 | 4.76919E-11 | 0.313553 | 0.188 | 0.093 | 7.83292E-07 |
| Terminal | GSTP1 | 1.20358E-08 | 0.313459 | 0.963 | 0.937 | 0.000197676 |
| RERGL_Cells | PIN1 | 0.003950534 | 0.313435 | 0.438 | 0.436 | 1 |
| RERGL_Cells | RASAL2 | 6.90573E-06 | 0.313026 | 0.158 | 0.095 | 0.113419751 |
| Terminal | NDUFS4 | 4.67809E-05 | 0.313023 | 0.547 | 0.491 | 0.768329966 |
| Pericytes | PDE5A | 2.20931E-06 | 0.31298 | 0.196 | 0.126 | 0.036285778 |
| Fibroblasts | FMO3 | 1.35855E-33 | 0.312886 | 0.14 | 0.031 | 2.23128E-29 |
| Pericytes | GTPBP4 | 0.001576753 | 0.312882 | 0.211 | 0.169 | 1 |
| Pericytes | SMARCA1 | 0.000239775 | 0.312765 | 0.209 | 0.156 | 1 |
| RERGL_Cells | ARIH2 | 0.004795657 | 0.312628 | 0.19 | 0.153 | 1 |
| Contractile | PFDN6 | 3.96786E-06 | 0.312603 | 0.295 | 0.216 | 0.065168162 |
| Pericytes | CISD1 | 0.000429036 | 0.312497 | 0.331 | 0.286 | 1 |
| Pericytes | ODF3B | 0.001110269 | 0.312473 | 0.153 | 0.112 | 1 |
| Terminal | TMBIM1 | 2.54335E-06 | 0.312317 | 0.562 | 0.488 | 0.041771963 |
| Fibroblasts | RASSF4 | 2.36351E-26 | 0.312308 | 0.184 | 0.065 | 3.88184E-22 |
| RERGL_Cells | RNF5 | 0.004987474 | 0.312231 | 0.331 | 0.311 | 1 |
| Pericytes | REV3L | 0.001981672 | 0.312168 | 0.187 | 0.145 | 1 |
| RERGL_Cells | GP1BB | 5.34562E-09 | 0.311995 | 0.115 | 0.05 | 8.77965E-05 |
| Pericytes | INAFM1 | 0.00213503 | 0.311975 | 0.271 | 0.23 | 1 |
| Fibroblasts | PTGER4 | 1.37446E-36 | 0.311879 | 0.163 | 0.039 | 2.25741E-32 |
| RERGL_Cells | SPATS2L | 0.000795255 | 0.311871 | 0.278 | 0.233 | 1 |
| Contractile | C1QTNF1 | 4.12304E-07 | 0.311778 | 0.507 | 0.394 | 0.006771677 |
| Contractile | HAAO | 6.68873E-08 | 0.31165 | 0.244 | 0.153 | 0.001098557 |
| RERGL_Cells | DRAP1 | 0.000100146 | 0.311569 | 0.543 | 0.533 | 1 |
| Fibroblasts | ITPKC | 5.93871E-33 | 0.31152 | 0.127 | 0.026 | 9.75373E-29 |
| Fibroblasts | MAN1C1 | 9.27632E-43 | 0.311466 | 0.144 | 0.024 | 1.52354E-38 |
| RERGL_Cells | CALD1 | 1.65359E-27 | 0.311463 | 0.991 | 0.961 | 2.71586E-23 |
| Contractile | HNRNPA3 | 3.89115E-09 | 0.31145 | 0.752 | 0.659 | 6.39083E-05 |
| Intermediate | TMEM30A | 2.68773E-07 | 0.311381 | 0.4 | 0.286 | 0.004414326 |

| | | | | | | |
|---|---|---|---|---|---|---|
| Pericytes | ARVCF | 3.42184E-06 | 0.311331 | 0.145 | 0.086 | 0.056200265 |
| Intermediate | EFEMP2 | 8.97936E-09 | 0.311303 | 0.611 | 0.488 | 0.000147477 |
| Fibroblasts | LPAR1 | 8.77136E-44 | 0.311275 | 0.131 | 0.018 | 1.44061E-39 |
| Contractile | MCRIP1 | 3.31838E-07 | 0.311056 | 0.483 | 0.382 | 0.005450111 |
| Fibroblasts | SLC16A7 | 7.83036E-25 | 0.310984 | 0.169 | 0.059 | 1.28606E-20 |
| Pericytes | PSMC4 | 0.000326912 | 0.310826 | 0.362 | 0.32 | 1 |
| RERGL_Cells | EIF3H | 2.86359E-08 | 0.310759 | 0.667 | 0.612 | 0.000470316 |
| Terminal | SCN7A | 0.000649708 | 0.310745 | 0.115 | 0.071 | 1 |
| Intermediate | IRS2 | 1.32067E-13 | 0.310705 | 0.185 | 0.081 | 2.16907E-09 |
| Intermediate | HEPH | 2.76182E-17 | 0.310542 | 0.221 | 0.091 | 4.53602E-13 |
| Fibroblasts | DDX5 | 5.54243E-37 | 0.310191 | 0.981 | 0.925 | 9.10289E-33 |
| Fibroblasts | HNRNPF | 7.96354E-21 | 0.31016 | 0.656 | 0.449 | 1.30793E-16 |
| Fibroblasts | NABP1 | 9.79371E-45 | 0.310104 | 0.307 | 0.104 | 1.60852E-40 |
| Intermediate | GULP1 | 5.06612E-09 | 0.310023 | 0.35 | 0.229 | 8.32059E-05 |
| Terminal | DNAJC1 | 0.000364155 | 0.30978 | 0.272 | 0.211 | 1 |
| Pericytes | TWISTNB | 0.000119227 | 0.309758 | 0.269 | 0.212 | 1 |
| Terminal | SEC11C | 4.9432E-05 | 0.309724 | 0.29 | 0.22 | 0.811871552 |
| RERGL_Cells | GNB5 | 0.000104097 | 0.309653 | 0.147 | 0.096 | 1 |
| Terminal | PTMS | 1.49649E-07 | 0.309424 | 0.804 | 0.761 | 0.00245783 |
| RERGL_Cells | NDUFS8 | 0.000232687 | 0.309285 | 0.479 | 0.454 | 1 |
| Contractile | RHOD | 2.37324E-10 | 0.309271 | 0.234 | 0.13 | 3.89782E-06 |
| Fibroblasts | JAM2 | 2.75771E-34 | 0.309255 | 0.137 | 0.029 | 4.52927E-30 |
| Contractile | PDCL3 | 2.84696E-11 | 0.309159 | 0.275 | 0.155 | 4.67585E-07 |
| Terminal | CTSF | 1.89661E-07 | 0.309151 | 0.593 | 0.503 | 0.00311499 |
| Terminal | CALM1 | 7.94418E-10 | 0.309139 | 0.847 | 0.783 | 1.30475E-05 |
| Intermediate | LINC02269 | 2.22482E-18 | 0.309089 | 0.168 | 0.058 | 3.65404E-14 |
| Pericytes | STAT3 | 0.000132386 | 0.308763 | 0.485 | 0.446 | 1 |
| RERGL_Cells | PKIG | 8.97395E-06 | 0.308719 | 0.553 | 0.499 | 0.147388212 |
| Contractile | GHITM | 1.81797E-08 | 0.30845 | 0.673 | 0.593 | 0.000298583 |
| Fibroblasts | NUMBL | 2.58935E-34 | 0.308335 | 0.169 | 0.045 | 4.25276E-30 |
| Intermediate | PPP1R13L | 1.60137E-18 | 0.308261 | 0.204 | 0.079 | 2.63009E-14 |
| Fibroblasts | SNHG9 | 9.28409E-22 | 0.308131 | 0.306 | 0.153 | 1.52482E-17 |
| Terminal | CRTAP | 1.78819E-05 | 0.307963 | 0.479 | 0.402 | 0.293691966 |
| Intermediate | NOL3 | 2.66108E-16 | 0.307927 | 0.359 | 0.185 | 4.37056E-12 |
| Fibroblasts | CD38 | 3.81712E-52 | 0.307918 | 0.107 | 0.005 | 6.26924E-48 |
| RERGL_Cells | HNRNPR | 0.000393301 | 0.307913 | 0.455 | 0.443 | 1 |
| Pericytes | ECHDC3 | 5.68528E-13 | 0.307882 | 0.144 | 0.059 | 9.33751E-09 |
| Pericytes | RDX | 0.001846626 | 0.307631 | 0.373 | 0.343 | 1 |
| RERGL_Cells | UQCRFS1 | 0.00010209 | 0.307598 | 0.474 | 0.448 | 1 |
| Pericytes | EIF1 | 2.94933E-29 | 0.307502 | 1 | 0.997 | 4.84397E-25 |
| Terminal | HAND2 | 5.09736E-06 | 0.307319 | 0.209 | 0.133 | 0.083719026 |
| Fibroblasts | CSGALNAC | 2.43868E-27 | 0.307214 | 0.206 | 0.076 | 4.00528E-23 |
| Intermediate | COL1A2 | 0.000125852 | 0.30712 | 0.796 | 0.689 | 1 |
| Contractile | ATP6V1E1 | 8.5213E-08 | 0.307117 | 0.555 | 0.454 | 0.001399538 |
| Contractile | OLA1 | 7.16705E-08 | 0.307063 | 0.333 | 0.23 | 0.001177116 |
| Terminal | DYNC1LI2 | 1.33902E-05 | 0.306987 | 0.32 | 0.237 | 0.219920421 |
| Pericytes | RSRC2 | 9.12103E-05 | 0.306882 | 0.471 | 0.43 | 1 |
| Terminal | ALCAM | 1.10511E-06 | 0.306834 | 0.192 | 0.114 | 0.018150335 |
| Fibroblasts | FKBP9 | 3.95112E-16 | 0.306817 | 0.356 | 0.214 | 6.48932E-12 |
| Intermediate | DNASE2 | 9.78759E-13 | 0.306778 | 0.335 | 0.191 | 1.60751E-08 |
| Fibroblasts | RDH10 | 3.37358E-40 | 0.306641 | 0.101 | 0.01 | 5.54077E-36 |
| Terminal | MAP4 | 7.22825E-07 | 0.306528 | 0.423 | 0.321 | 0.011871677 |
| Contractile | SRP68 | 7.1947E-07 | 0.306505 | 0.317 | 0.226 | 0.011816573 |
| Fibroblasts | BLVRB | 1.18484E-19 | 0.306442 | 0.594 | 0.397 | 1.94598E-15 |
| Fibroblasts | ABCA1 | 3.05224E-28 | 0.306409 | 0.148 | 0.042 | 5.01301E-24 |
| RERGL_Cells | ZEB2 | 0.00252218 | 0.306379 | 0.25 | 0.208 | 1 |

| | | | | | | |
|---|---|---|---|---|---|---|
| Contractile | ATP5MD | 8.0935E-11 | 0.30633 | 0.832 | 0.753 | 1.32928E-06 |
| RERGL_Cells | MARCH2 | 0.002497587 | 0.305886 | 0.192 | 0.151 | 1 |
| Terminal | NME2 | 4.43522E-10 | 0.305841 | 0.893 | 0.875 | 7.28441E-06 |
| Fibroblasts | SLC9A9 | 5.76621E-42 | 0.305833 | 0.117 | 0.014 | 9.47042E-38 |
| Contractile | MIF4GD | 1.9846E-07 | 0.30583 | 0.234 | 0.148 | 0.003259501 |
| Contractile | NDUFS8 | 4.38991E-06 | 0.305802 | 0.515 | 0.448 | 0.072099838 |
| Pericytes | MCMBP | 7.6724E-05 | 0.305778 | 0.225 | 0.168 | 1 |
| Fibroblasts | SYNGR1 | 2.83055E-26 | 0.30576 | 0.313 | 0.147 | 4.6489E-22 |
| Contractile | NDUFA7 | 3.33774E-09 | 0.305755 | 0.401 | 0.273 | 5.4819E-05 |
| Pericytes | DDX39B | 5.22546E-05 | 0.305666 | 0.507 | 0.474 | 0.8582289 |
| Terminal | CHID1 | 2.73601E-05 | 0.305626 | 0.41 | 0.336 | 0.449362698 |
| Fibroblasts | EGR3 | 3.82344E-30 | 0.305474 | 0.121 | 0.026 | 6.27962E-26 |
| RERGL_Cells | UNC45A | 0.00443194 | 0.305463 | 0.241 | 0.207 | 1 |
| RERGL_Cells | HNRNPL | 0.000647521 | 0.305163 | 0.192 | 0.144 | 1 |
| Fibroblasts | KPNA2 | 6.54041E-21 | 0.305122 | 0.231 | 0.106 | 1.0742E-16 |
| Fibroblasts | RBPJ | 4.44971E-24 | 0.305065 | 0.533 | 0.322 | 7.3082E-20 |
| Contractile | EDF1 | 9.14453E-14 | 0.305035 | 0.884 | 0.8 | 1.5019E-09 |
| Terminal | NUDC | 5.60775E-05 | 0.305005 | 0.601 | 0.559 | 0.921016296 |
| Intermediate | PPP1R14B | 7.87606E-09 | 0.304943 | 0.258 | 0.155 | 0.000129356 |
| Intermediate | HINT1 | 6.76822E-16 | 0.304777 | 0.886 | 0.77 | 1.11161E-11 |
| Pericytes | MBNL1-AS | 0.004232063 | 0.304657 | 0.18 | 0.138 | 1 |
| Intermediate | CAVIN3 | 3.15031E-14 | 0.304619 | 0.875 | 0.717 | 5.17407E-10 |
| Fibroblasts | UCHL1 | 1.34567E-19 | 0.3044 | 0.261 | 0.131 | 2.21013E-15 |
| Terminal | BACE2 | 0.001154803 | 0.30417 | 0.283 | 0.233 | 1 |
| RERGL_Cells | LUC7L2 | 0.001048183 | 0.304102 | 0.397 | 0.372 | 1 |
| Intermediate | NT5DC1 | 7.34937E-10 | 0.304101 | 0.312 | 0.192 | 1.20706E-05 |
| Terminal | HINT1 | 1.20463E-05 | 0.303883 | 0.813 | 0.784 | 0.19784876 |
| RERGL_Cells | MICOS13 | 2.26463E-05 | 0.303844 | 0.551 | 0.522 | 0.371942202 |
| Contractile | ITGA3 | 1.12691E-09 | 0.303673 | 0.134 | 0.06 | 1.85084E-05 |
| Pericytes | NR2F2-AS1 | 1.09523E-05 | 0.303649 | 0.187 | 0.122 | 0.179880681 |
| Intermediate | OS9 | 1.02896E-08 | 0.303605 | 0.669 | 0.541 | 0.000168997 |
| Pericytes | MICALL2 | 8.51076E-08 | 0.303596 | 0.136 | 0.072 | 0.001397807 |
| Pericytes | NUAK1 | 1.7382E-11 | 0.303539 | 0.109 | 0.041 | 2.85483E-07 |
| Terminal | P4HB | 2.85475E-07 | 0.303311 | 0.756 | 0.728 | 0.004688646 |
| RERGL_Cells | ADCY3 | 7.8096E-06 | 0.303277 | 0.162 | 0.1 | 0.128264833 |
| Terminal | VKORC1 | 4.58439E-07 | 0.303254 | 0.678 | 0.597 | 0.007529406 |
| Intermediate | PRUNE2 | 1.30972E-18 | 0.303242 | 0.163 | 0.055 | 2.15108E-14 |
| Intermediate | SHC4 | 5.46351E-24 | 0.303161 | 0.194 | 0.062 | 8.97327E-20 |
| Intermediate | CASK | 1.90945E-22 | 0.303147 | 0.123 | 0.03 | 3.13609E-18 |
| Intermediate | TMEM219 | 6.12088E-12 | 0.303146 | 0.57 | 0.391 | 1.00529E-07 |
| Fibroblasts | HLA-A | 2.25532E-19 | 0.303119 | 0.944 | 0.848 | 3.70413E-15 |
| Pericytes | WASHC4 | 0.005789243 | 0.302962 | 0.196 | 0.161 | 1 |
| Terminal | TMED9 | 7.17209E-07 | 0.302959 | 0.712 | 0.632 | 0.011779446 |
| Terminal | MRPS34 | 0.000786994 | 0.302884 | 0.309 | 0.25 | 1 |
| Contractile | AK5 | 7.78023E-26 | 0.302802 | 0.118 | 0.023 | 1.27782E-21 |
| Pericytes | DAZAP1 | 0.000167215 | 0.302789 | 0.213 | 0.159 | 1 |
| Terminal | TMED4 | 1.82045E-05 | 0.302754 | 0.423 | 0.343 | 0.298991503 |
| Pericytes | TRIM41 | 0.000294671 | 0.302703 | 0.131 | 0.087 | 1 |
| Fibroblasts | GDF10 | 5.46971E-30 | 0.302551 | 0.149 | 0.04 | 8.98345E-26 |
| Intermediate | SEC11A | 8.55231E-10 | 0.302528 | 0.658 | 0.524 | 1.40463E-05 |
| Intermediate | HAND2 | 7.38461E-18 | 0.302527 | 0.271 | 0.12 | 1.21285E-13 |
| Intermediate | FXYD1 | 3.71179E-13 | 0.302473 | 0.901 | 0.781 | 6.09624E-09 |
| Terminal | ERG | 5.43501E-12 | 0.302429 | 0.122 | 0.045 | 8.92646E-08 |
| Contractile | LAYN | 2.13944E-15 | 0.302284 | 0.128 | 0.042 | 3.51382E-11 |
| Contractile | AC245297 | 3.31945E-06 | 0.302276 | 0.289 | 0.206 | 0.054518585 |
| RERGL_Cells | DCTN3 | 0.000362578 | 0.302251 | 0.494 | 0.471 | 1 |

| | | | | | | |
|---|---|---|---|---|---|---|
| Intermediate | FGL2 | 1.44738E-08 | 0.302248 | 0.271 | 0.161 | 0.000237717 |
| Fibroblasts | CISD1 | 2.01652E-19 | 0.302247 | 0.411 | 0.242 | 3.31194E-15 |
| Terminal | GPAA1 | 9.0494E-05 | 0.302216 | 0.46 | 0.401 | 1 |
| Pericytes | NCL | 0.001071964 | 0.302179 | 0.553 | 0.546 | 1 |
| Fibroblasts | ST3GAL1 | 4.56928E-23 | 0.302147 | 0.256 | 0.116 | 7.50459E-19 |
| Contractile | ATP5MPL | 9.77557E-11 | 0.30198 | 0.822 | 0.746 | 1.60554E-06 |
| Terminal | MRPL34 | 0.000310729 | 0.30191 | 0.392 | 0.331 | 1 |
| Intermediate | TMEM25 | 3.64943E-17 | 0.301768 | 0.172 | 0.063 | 5.99383E-13 |
| RERGL_Cells | HINT2 | 0.006731414 | 0.301649 | 0.28 | 0.249 | 1 |
| Pericytes | HNRNPAB | 0.000596286 | 0.301637 | 0.291 | 0.248 | 1 |
| Terminal | CREB3L2 | 3.54394E-08 | 0.301621 | 0.19 | 0.104 | 0.000582056 |
| Contractile | NDUFA13 | 1.10078E-12 | 0.301611 | 0.87 | 0.799 | 1.80792E-08 |
| Fibroblasts | CSTF2T | 4.60587E-25 | 0.301593 | 0.222 | 0.092 | 7.56468E-21 |
| Contractile | ITIH3 | 3.29876E-14 | 0.301589 | 0.154 | 0.059 | 5.41788E-10 |
| Pericytes | HNRNPA0 | 2.41777E-07 | 0.301546 | 0.733 | 0.73 | 0.003970953 |
| Contractile | MRPL57 | 2.61205E-06 | 0.301529 | 0.441 | 0.352 | 0.042900337 |
| RERGL_Cells | LAMTOR1 | 0.000648168 | 0.301408 | 0.513 | 0.512 | 1 |
| Fibroblasts | ACADVL | 5.65765E-21 | 0.301397 | 0.737 | 0.536 | 9.29213E-17 |
| Intermediate | CD55 | 2.29138E-09 | 0.301348 | 0.357 | 0.235 | 3.76336E-05 |
| Terminal | SERGEF | 5.96541E-06 | 0.301348 | 0.24 | 0.159 | 0.097975883 |
| Pericytes | RABGGTB | 0.005935129 | 0.301129 | 0.231 | 0.194 | 1 |
| Terminal | MFAP4 | 9.5244E-07 | 0.301055 | 0.9 | 0.792 | 0.015642877 |
| RERGL_Cells | NAB1 | 6.83655E-07 | 0.301028 | 0.143 | 0.078 | 0.011228347 |
| Pericytes | SLC20A2 | 4.98649E-06 | 0.301022 | 0.156 | 0.096 | 0.081898069 |
| Pericytes | RCSD1 | 2.88292E-10 | 0.300684 | 0.151 | 0.07 | 4.73491E-06 |
| Intermediate | SNX18 | 3.8888E-14 | 0.30052 | 0.256 | 0.126 | 6.38696E-10 |
| Terminal | ZC3H13 | 3.62048E-05 | 0.300488 | 0.351 | 0.273 | 0.594628261 |
| Contractile | EMC4 | 3.64466E-07 | 0.300471 | 0.551 | 0.472 | 0.005985998 |
| Contractile | BMERB1 | 3.49898E-06 | 0.300448 | 0.299 | 0.212 | 0.057467272 |
| Terminal | COL21A1 | 5.81786E-08 | 0.300367 | 0.248 | 0.151 | 0.000955525 |
| Pericytes | GBP1 | 5.77968E-06 | 0.300209 | 0.16 | 0.101 | 0.094925395 |
| Fibroblasts | TXNIP | 6.59058E-19 | 0.300106 | 0.767 | 0.612 | 1.08244E-14 |
| RERGL_Cells | MAGEF1 | 0.00047443 | 0.299924 | 0.261 | 0.211 | 1 |
| Terminal | TALDO1 | 8.61977E-05 | 0.299915 | 0.49 | 0.426 | 1 |
| Contractile | SNRPE | 1.50895E-07 | 0.299833 | 0.577 | 0.487 | 0.002478305 |
| Pericytes | RNPEP | 0.001344954 | 0.299742 | 0.218 | 0.175 | 1 |
| Contractile | PSMB1 | 5.65102E-11 | 0.299566 | 0.822 | 0.771 | 9.28123E-07 |
| Pericytes | PAK1IP1 | 0.001834238 | 0.29955 | 0.207 | 0.167 | 1 |
| Pericytes | IMP4 | 0.002812011 | 0.299433 | 0.265 | 0.227 | 1 |
| Intermediate | SLC22A17 | 4.4341E-13 | 0.299427 | 0.318 | 0.176 | 7.28257E-09 |
| Terminal | RRBP1 | 1.87212E-05 | 0.29936 | 0.418 | 0.335 | 0.307477443 |
| RERGL_Cells | NCOA4 | 0.004332023 | 0.299351 | 0.31 | 0.279 | 1 |
| Pericytes | HLA-F | 6.86556E-06 | 0.299306 | 0.189 | 0.125 | 0.112759977 |
| Intermediate | MIR497HG | 1.64523E-11 | 0.299291 | 0.303 | 0.172 | 2.70213E-07 |
| Contractile | RPP21 | 2.32014E-07 | 0.29926 | 0.405 | 0.299 | 0.003810604 |
| Pericytes | EIF4E | 0.00076309 | 0.299242 | 0.364 | 0.326 | 1 |
| RERGL_Cells | CPEB4 | 0.000113784 | 0.299242 | 0.126 | 0.077 | 1 |
| Terminal | EEF1D | 1.43871E-10 | 0.29907 | 0.932 | 0.913 | 2.36294E-06 |
| Contractile | HMG20B | 4.27698E-07 | 0.299 | 0.331 | 0.236 | 0.007024509 |
| RERGL_Cells | RB1CC1 | 0.007794845 | 0.298939 | 0.237 | 0.205 | 1 |
| Fibroblasts | PIGT | 1.10351E-17 | 0.298926 | 0.626 | 0.433 | 1.81241E-13 |
| Pericytes | RNF138 | 4.70801E-07 | 0.298817 | 0.116 | 0.061 | 0.007732428 |
| Contractile | ROR1 | 1.63997E-12 | 0.298745 | 0.104 | 0.035 | 2.69348E-08 |
| RERGL_Cells | STARD7 | 0.000397318 | 0.298538 | 0.194 | 0.143 | 1 |
| Pericytes | ADAMTS2 | 1.53368E-07 | 0.298477 | 0.136 | 0.073 | 0.002518912 |
| Terminal | GSTO1 | 1.42814E-05 | 0.298472 | 0.601 | 0.543 | 0.234557001 |

| | | | | | | |
|---|---|---|---|---|---|---|
| Fibroblasts | SLPI | 1.86271E-63 | 0.29838 | 0.296 | 0.077 | 3.05932E-59 |
| Terminal | EBPL | 0.00117808 | 0.298307 | 0.277 | 0.224 | 1 |
| Pericytes | OSMR | 0.001201328 | 0.298096 | 0.18 | 0.138 | 1 |
| Terminal | HMGN1 | 7.96151E-05 | 0.298023 | 0.481 | 0.421 | 1 |
| Intermediate | NRP2 | 6.95752E-23 | 0.298022 | 0.114 | 0.025 | 1.1427E-18 |
| Intermediate | OST4 | 6.02405E-15 | 0.297997 | 0.92 | 0.847 | 9.89391E-11 |
| RERGL_Cells | MSRB2 | 0.00027988 | 0.297957 | 0.353 | 0.309 | 1 |
| Terminal | HEXB | 0.001077963 | 0.29785 | 0.486 | 0.449 | 1 |
| Terminal | POLR2C | 0.001759502 | 0.297661 | 0.423 | 0.371 | 1 |
| Contractile | HACD4 | 5.56512E-14 | 0.297622 | 0.132 | 0.047 | 9.14016E-10 |
| Terminal | AC092164 | 5.47914E-05 | 0.297618 | 0.205 | 0.138 | 0.899894124 |
| RERGL_Cells | MTMR2 | 0.002211171 | 0.297571 | 0.15 | 0.11 | 1 |
| Terminal | GNAI2 | 8.4545E-07 | 0.297541 | 0.314 | 0.219 | 0.01388567 |
| Intermediate | GPC4 | 2.53991E-12 | 0.297464 | 0.204 | 0.098 | 4.17154E-08 |
| RERGL_Cells | SART1 | 0.002397748 | 0.297455 | 0.201 | 0.159 | 1 |
| Pericytes | TRPC4AP | 1.7472E-05 | 0.29745 | 0.227 | 0.164 | 0.286960792 |
| Contractile | MZT2A | 2.26022E-05 | 0.297415 | 0.353 | 0.279 | 0.371218614 |
| Pericytes | STRAP | 2.00107E-05 | 0.297173 | 0.509 | 0.47 | 0.328655185 |
| Intermediate | PTMS | 1.2745E-12 | 0.297112 | 0.882 | 0.746 | 2.09325E-08 |
| Terminal | SDF4 | 9.8893E-07 | 0.297108 | 0.545 | 0.458 | 0.016242187 |
| Fibroblasts | BMP1 | 4.83191E-26 | 0.297061 | 0.164 | 0.054 | 7.93592E-22 |
| Terminal | ITSN2 | 9.6417E-06 | 0.296999 | 0.237 | 0.161 | 0.158355353 |
| Fibroblasts | RFTN1 | 1.05658E-31 | 0.296951 | 0.2 | 0.065 | 1.73532E-27 |
| Terminal | PDAP1 | 0.000713085 | 0.296945 | 0.492 | 0.445 | 1 |
| Contractile | BTC | 1.26186E-23 | 0.29694 | 0.114 | 0.024 | 2.07248E-19 |
| Terminal | YIPF2 | 6.08515E-06 | 0.296938 | 0.34 | 0.249 | 0.09994247 |
| Contractile | GSTO1 | 1.94659E-07 | 0.296912 | 0.629 | 0.538 | 0.003197087 |
| Terminal | CRNDE | 2.27475E-06 | 0.296872 | 0.259 | 0.168 | 0.037360513 |
| Pericytes | HSPE1 | 0.001185885 | 0.296769 | 0.524 | 0.54 | 1 |
| Terminal | ECM2 | 0.00098418 | 0.296713 | 0.375 | 0.311 | 1 |
| Terminal | MPHOSPH | 2.47777E-05 | 0.296668 | 0.514 | 0.44 | 0.406948836 |
| Intermediate | MRPL49 | 1.09411E-15 | 0.296613 | 0.295 | 0.145 | 1.79697E-11 |
| RERGL_Cells | ANGPTL1 | 0.003359014 | 0.296608 | 0.261 | 0.216 | 1 |
| Pericytes | RPP25L | 0.001716927 | 0.296548 | 0.147 | 0.109 | 1 |
| Fibroblasts | AKIRIN2 | 7.27574E-21 | 0.296342 | 0.337 | 0.182 | 1.19497E-16 |
| Contractile | LRRFIP2 | 3.7813E-08 | 0.296341 | 0.283 | 0.182 | 0.00062104 |
| Pericytes | LINC02381 | 1.37277E-05 | 0.296321 | 0.147 | 0.091 | 0.22546368 |
| Pericytes | NEDD9 | 0.000374584 | 0.296249 | 0.125 | 0.081 | 1 |
| RERGL_Cells | PHB2 | 0.005340304 | 0.296244 | 0.382 | 0.366 | 1 |
| RERGL_Cells | ANAPC11 | 1.24358E-05 | 0.29624 | 0.611 | 0.601 | 0.204246087 |
| RERGL_Cells | SRRM1 | 0.000200167 | 0.296074 | 0.496 | 0.474 | 1 |
| Contractile | ZYX | 8.39494E-08 | 0.295862 | 0.559 | 0.446 | 0.001378786 |
| Pericytes | E2F4 | 0.001421026 | 0.295693 | 0.195 | 0.152 | 1 |
| RERGL_Cells | SMIM30 | 0.004638404 | 0.295614 | 0.359 | 0.337 | 1 |
| Pericytes | MRVI1 | 1.22623E-08 | 0.295576 | 0.264 | 0.164 | 0.000201396 |
| Intermediate | TXN | 1.07985E-12 | 0.295534 | 0.821 | 0.67 | 1.77354E-08 |
| Contractile | SNAPIN | 4.78731E-07 | 0.295418 | 0.339 | 0.24 | 0.007862683 |
| Contractile | CNOT7 | 1.20354E-06 | 0.295358 | 0.307 | 0.218 | 0.019766998 |
| Pericytes | DCTN2 | 2.51777E-06 | 0.295213 | 0.529 | 0.492 | 0.041351935 |
| Contractile | FRY | 4.34198E-09 | 0.295013 | 0.204 | 0.111 | 7.13127E-05 |
| Contractile | COX6A1 | 1.04203E-16 | 0.294868 | 0.902 | 0.822 | 1.71142E-12 |
| Terminal | EEF1B2 | 1.0368E-06 | 0.29482 | 0.832 | 0.793 | 0.017028397 |
| Intermediate | ADH1B | 1.64999E-06 | 0.294758 | 0.74 | 0.551 | 0.027099425 |
| Fibroblasts | SRGAP1 | 5.0494E-36 | 0.294691 | 0.149 | 0.033 | 8.29314E-32 |
| Intermediate | TIMP2 | 6.98268E-11 | 0.29468 | 0.572 | 0.408 | 1.14684E-06 |
| RERGL_Cells | CCT7 | 0.000374244 | 0.29463 | 0.506 | 0.497 | 1 |

| | | | | | | |
|---|---|---|---|---|---|---|
| RERGL_Cells | NPM1 | 3.85212E-12 | 0.294573 | 0.917 | 0.873 | 6.32673E-08 |
| RERGL_Cells | RNF152 | 0.000316127 | 0.294567 | 0.132 | 0.085 | 1 |
| Contractile | RAB23 | 2.26003E-05 | 0.294504 | 0.226 | 0.158 | 0.371187431 |
| Fibroblasts | LOXL1 | 0.000189943 | 0.294416 | 0.35 | 0.286 | 1 |
| Contractile | VCL | 2.97842E-07 | 0.294374 | 0.365 | 0.261 | 0.004891759 |
| RERGL_Cells | CCT8 | 0.000330522 | 0.294346 | 0.526 | 0.515 | 1 |
| Terminal | DHRS7 | 0.000200828 | 0.294248 | 0.473 | 0.418 | 1 |
| Intermediate | EIF4A2 | 1.94572E-12 | 0.294195 | 0.94 | 0.887 | 3.19565E-08 |
| Contractile | TMOD1 | 2.78207E-07 | 0.294155 | 0.279 | 0.18 | 0.004569265 |
| Terminal | PHLDA3 | 0.001304718 | 0.294103 | 0.401 | 0.345 | 1 |
| Contractile | FOXC1 | 5.74207E-09 | 0.294079 | 0.148 | 0.073 | 9.43077E-05 |
| Terminal | DMKN | 1.59223E-05 | 0.293971 | 0.166 | 0.101 | 0.261508614 |
| Terminal | CPQ | 5.91503E-05 | 0.293918 | 0.547 | 0.459 | 0.971484844 |
| RERGL_Cells | ATP5PF | 5.63455E-07 | 0.293759 | 0.741 | 0.744 | 0.009254181 |
| Intermediate | MAGED1 | 1.75314E-10 | 0.29369 | 0.35 | 0.219 | 2.87936E-06 |
| RERGL_Cells | USP16 | 0.000765103 | 0.293672 | 0.37 | 0.33 | 1 |
| RERGL_Cells | POLR2J | 0.007650148 | 0.29363 | 0.38 | 0.377 | 1 |
| Fibroblasts | STMP1 | 2.84035E-20 | 0.293414 | 0.462 | 0.278 | 4.66499E-16 |
| Pericytes | TWF1 | 0.00045431 | 0.293367 | 0.202 | 0.153 | 1 |
| RERGL_Cells | SKP1 | 3.76366E-16 | 0.293276 | 0.955 | 0.946 | 6.18143E-12 |
| Terminal | SPTBN1 | 1.88311E-05 | 0.293269 | 0.429 | 0.349 | 0.309281829 |
| RERGL_Cells | LAMTOR4 | 0.000134876 | 0.293219 | 0.564 | 0.573 | 1 |
| Terminal | BAX | 0.000119448 | 0.293129 | 0.355 | 0.285 | 1 |
| Fibroblasts | AKR1C2 | 1.65719E-22 | 0.293118 | 0.278 | 0.134 | 2.72178E-18 |
| Intermediate | CTPS1 | 2.4447E-12 | 0.293046 | 0.254 | 0.131 | 4.01518E-08 |
| Fibroblasts | APOL6 | 1.3082E-24 | 0.292992 | 0.148 | 0.047 | 2.14858E-20 |
| Contractile | MMADHC | 5.73889E-06 | 0.292965 | 0.511 | 0.43 | 0.094255569 |
| Fibroblasts | CAST | 2.9385E-17 | 0.292943 | 0.615 | 0.438 | 4.82619E-13 |
| Contractile | POLR1D | 5.07773E-05 | 0.292863 | 0.343 | 0.27 | 0.833965696 |
| Contractile | HS1BP3 | 1.86639E-10 | 0.292631 | 0.152 | 0.07 | 3.06536E-06 |
| Fibroblasts | ARHGAP21 | 9.81582E-27 | 0.292611 | 0.162 | 0.052 | 1.61215E-22 |
| Terminal | ARHGDIB | 0.000953013 | 0.292609 | 0.549 | 0.504 | 1 |
| RERGL_Cells | C5orf24 | 0.003075199 | 0.292606 | 0.25 | 0.214 | 1 |
| Contractile | ATP5ME | 5.62976E-09 | 0.292364 | 0.842 | 0.752 | 9.24631E-05 |
| Fibroblasts | MESD | 1.61825E-20 | 0.292089 | 0.441 | 0.26 | 2.65782E-16 |
| Contractile | PFKP | 1.78154E-08 | 0.292061 | 0.216 | 0.126 | 0.0002926 |
| Intermediate | SDC4 | 6.00829E-14 | 0.29188 | 0.321 | 0.172 | 9.86802E-10 |
| Pericytes | ADCY3 | 2.47737E-07 | 0.291829 | 0.167 | 0.097 | 0.004068835 |
| Intermediate | TMEM109 | 7.82364E-10 | 0.291684 | 0.654 | 0.507 | 1.28496E-05 |
| Pericytes | AL118516. | 0.000946932 | 0.291679 | 0.111 | 0.073 | 1 |
| Fibroblasts | SPATA6 | 1.60437E-30 | 0.291574 | 0.148 | 0.039 | 2.63501E-26 |
| Pericytes | LMNA | 2.43571E-10 | 0.291523 | 0.922 | 0.897 | 4.00042E-06 |
| Intermediate | ENDOD1 | 1.42176E-15 | 0.29151 | 0.234 | 0.106 | 2.33511E-11 |
| Terminal | GABARAP | 1.53138E-10 | 0.291494 | 0.93 | 0.89 | 2.51514E-06 |
| Intermediate | RNF7 | 1.74465E-11 | 0.291486 | 0.609 | 0.436 | 2.86541E-07 |
| RERGL_Cells | LINC01197 | 3.91797E-05 | 0.291453 | 0.231 | 0.163 | 0.643487675 |
| RERGL_Cells | NDUFC2 | 8.76082E-06 | 0.291395 | 0.652 | 0.652 | 0.143887656 |
| Terminal | AKR1A1 | 4.85661E-05 | 0.291393 | 0.42 | 0.346 | 0.797649268 |
| Fibroblasts | TMEM167 | 2.90617E-20 | 0.291317 | 0.378 | 0.211 | 4.77309E-16 |
| Contractile | CHMP4B | 1.29495E-05 | 0.291172 | 0.505 | 0.43 | 0.212682994 |
| Contractile | NDUFV2 | 4.65432E-07 | 0.291154 | 0.589 | 0.509 | 0.007644251 |
| RERGL_Cells | CNBP | 8.43172E-11 | 0.291116 | 0.868 | 0.813 | 1.38483E-06 |
| Fibroblasts | ARHGAP12 | 1.01943E-31 | 0.29108 | 0.151 | 0.039 | 1.67431E-27 |
| Intermediate | DSEL | 1.4948E-10 | 0.290984 | 0.249 | 0.138 | 2.45506E-06 |
| RERGL_Cells | CYCS | 1.85122E-06 | 0.290858 | 0.573 | 0.538 | 0.03040438 |
| Intermediate | CARD19 | 5.5331E-10 | 0.290843 | 0.353 | 0.22 | 9.08757E-06 |

| | | | | | | |
|---|---|---|---|---|---|---|
| Terminal | SERF2 | 7.49866E-13 | 0.290791 | 0.985 | 0.973 | 1.23158E-08 |
| RERGL_Cells | GALNT16 | 0.001052024 | 0.290636 | 0.126 | 0.085 | 1 |
| Fibroblasts | ANKH | 8.10303E-19 | 0.290523 | 0.29 | 0.156 | 1.33084E-14 |
| RERGL_Cells | MAL | 4.57609E-08 | 0.29048 | 0.1 | 0.043 | 0.000751577 |
| Contractile | DACT3 | 7.04859E-12 | 0.290274 | 0.156 | 0.066 | 1.15766E-07 |
| Contractile | EI24 | 2.12908E-08 | 0.290252 | 0.497 | 0.384 | 0.00034968 |
| RERGL_Cells | C9orf78 | 0.009312278 | 0.290245 | 0.314 | 0.292 | 1 |
| Fibroblasts | SLFN5 | 1.48331E-27 | 0.290237 | 0.157 | 0.047 | 2.43618E-23 |
| RERGL_Cells | MYL6B | 0.009281146 | 0.290099 | 0.239 | 0.21 | 1 |
| RERGL_Cells | NDUFS6 | 0.000207801 | 0.290086 | 0.532 | 0.523 | 1 |
| Terminal | RAMP1 | 1.95945E-05 | 0.290069 | 0.484 | 0.368 | 0.321820221 |
| Intermediate | TBCB | 1.64819E-11 | 0.28998 | 0.645 | 0.477 | 2.70699E-07 |
| Contractile | TBC1D1 | 7.37721E-08 | 0.289955 | 0.24 | 0.145 | 0.001211634 |
| Intermediate | NDUFA4 | 2.208E-20 | 0.289946 | 0.968 | 0.899 | 3.62642E-16 |
| RERGL_Cells | GLYR1 | 0.00413618 | 0.289945 | 0.16 | 0.123 | 1 |
| Terminal | RHBDD2 | 3.23177E-05 | 0.289917 | 0.307 | 0.231 | 0.530785296 |
| Terminal | DCXR | 0.001869055 | 0.289906 | 0.312 | 0.259 | 1 |
| Contractile | CRISPLD1 | 5.95122E-05 | 0.289804 | 0.279 | 0.206 | 0.977428496 |
| Terminal | RGN | 1.40097E-06 | 0.289778 | 0.144 | 0.079 | 0.023009573 |
| Pericytes | PSMA4 | 4.61568E-05 | 0.289729 | 0.573 | 0.565 | 0.758078681 |
| Contractile | IGBP1 | 4.09741E-07 | 0.289484 | 0.355 | 0.259 | 0.006729589 |
| RERGL_Cells | MAGED2 | 9.51739E-05 | 0.289447 | 0.641 | 0.638 | 1 |
| Contractile | ENY2 | 3.80687E-06 | 0.289437 | 0.531 | 0.446 | 0.062524065 |
| Contractile | RRAS | 5.28648E-05 | 0.289418 | 0.361 | 0.287 | 0.868250831 |
| Terminal | PIGT | 1.85482E-05 | 0.289259 | 0.547 | 0.483 | 0.304636349 |
| Pericytes | MITF | 1.06749E-05 | 0.289215 | 0.12 | 0.068 | 0.175325336 |
| RERGL_Cells | ST5 | 0.004020443 | 0.289115 | 0.177 | 0.138 | 1 |
| RERGL_Cells | DCTN1 | 0.0015044 | 0.289096 | 0.192 | 0.147 | 1 |
| Intermediate | ZMAT3 | 2.00096E-10 | 0.28905 | 0.228 | 0.123 | 3.28637E-06 |
| Pericytes | HOXB-AS1 | 0.001997734 | 0.289027 | 0.173 | 0.13 | 1 |
| Fibroblasts | PDCD4 | 1.85522E-22 | 0.288966 | 0.313 | 0.157 | 3.04701E-18 |
| Contractile | KIAA1191 | 8.2983E-08 | 0.288927 | 0.22 | 0.131 | 0.001362913 |
| Intermediate | DNASE1L1 | 2.74535E-22 | 0.288911 | 0.157 | 0.046 | 4.50897E-18 |
| Pericytes | RBM14 | 0.001191217 | 0.288891 | 0.165 | 0.123 | 1 |
| RERGL_Cells | C18orf32 | 6.98234E-05 | 0.28887 | 0.498 | 0.472 | 1 |
| Intermediate | PERP | 3.14546E-12 | 0.288838 | 0.338 | 0.196 | 5.1661E-08 |
| Intermediate | REEP5 | 4.55436E-09 | 0.288811 | 0.708 | 0.562 | 7.48008E-05 |
| Pericytes | ZNF22 | 0.002199101 | 0.288784 | 0.255 | 0.21 | 1 |
| Pericytes | MTHFR | 0.000450783 | 0.288757 | 0.144 | 0.099 | 1 |
| RERGL_Cells | C6orf47 | 0.003358226 | 0.288715 | 0.175 | 0.136 | 1 |
| Pericytes | MED21 | 0.00355473 | 0.288706 | 0.311 | 0.28 | 1 |
| Intermediate | SPCS1 | 2.50347E-11 | 0.288698 | 0.789 | 0.678 | 4.1117E-07 |
| Contractile | HEBP1 | 6.93411E-06 | 0.288569 | 0.397 | 0.316 | 0.113885887 |
| RERGL_Cells | CAMLG | 0.001060825 | 0.288407 | 0.451 | 0.428 | 1 |
| Contractile | TRAPPC2L | 1.06136E-06 | 0.288309 | 0.411 | 0.315 | 0.017431819 |
| Contractile | RNF11 | 1.44347E-05 | 0.288292 | 0.467 | 0.389 | 0.23707602 |
| Intermediate | HIGD1A | 1.24909E-07 | 0.288238 | 0.533 | 0.401 | 0.002051502 |
| Terminal | CNPY2 | 0.000138118 | 0.288232 | 0.521 | 0.469 | 1 |
| Pericytes | EPN2 | 3.80394E-05 | 0.288182 | 0.175 | 0.118 | 0.624758638 |
| Contractile | PCNP | 2.5739E-07 | 0.288174 | 0.675 | 0.605 | 0.004227374 |
| Pericytes | ZCCHC10 | 0.002907896 | 0.288154 | 0.202 | 0.162 | 1 |
| Pericytes | ASPH | 0.000954445 | 0.288117 | 0.425 | 0.394 | 1 |
| RERGL_Cells | TNNC1 | 7.52514E-07 | 0.288038 | 0.13 | 0.068 | 0.012359283 |
| Pericytes | CCT6A | 0.003917093 | 0.287971 | 0.409 | 0.392 | 1 |
| RERGL_Cells | THAP11 | 0.001846755 | 0.287962 | 0.113 | 0.077 | 1 |
| Pericytes | VPS16 | 0.000137154 | 0.287949 | 0.129 | 0.083 | 1 |

| | | | | | | |
|---|---|---|---|---|---|---|
| Contractile | ADSSL1 | 1.11411E-17 | 0.287893 | 0.1 | 0.025 | 1.82982E-13 |
| Fibroblasts | SETD7 | 4.5242E-26 | 0.287867 | 0.17 | 0.057 | 7.43054E-22 |
| Pericytes | NXF1 | 0.000561729 | 0.287819 | 0.224 | 0.173 | 1 |
| Pericytes | PAPOLA | 6.61699E-06 | 0.287765 | 0.433 | 0.376 | 0.108677376 |
| Contractile | NR1H2 | 1.98217E-06 | 0.287747 | 0.467 | 0.382 | 0.03255518 |
| Terminal | MTPN | 0.000404812 | 0.287734 | 0.266 | 0.208 | 1 |
| Intermediate | UPP1 | 3.07937E-21 | 0.287532 | 0.179 | 0.059 | 5.05755E-17 |
| Contractile | RTRAF | 1.17069E-06 | 0.287523 | 0.627 | 0.557 | 0.019227492 |
| Intermediate | LRRFIP1 | 1.03286E-13 | 0.287467 | 0.492 | 0.303 | 1.69638E-09 |
| Contractile | SERBP1 | 3.33823E-08 | 0.287436 | 0.679 | 0.588 | 0.000548271 |
| Contractile | AIF1L | 5.46524E-14 | 0.287421 | 0.112 | 0.036 | 8.97611E-10 |
| Contractile | CCNB1IP1 | 2.80788E-07 | 0.287411 | 0.22 | 0.137 | 0.004611658 |
| RERGL_Cells | SNRPF | 0.005396749 | 0.287391 | 0.421 | 0.418 | 1 |
| Intermediate | WDR83OS | 3.27332E-09 | 0.287357 | 0.753 | 0.656 | 5.3761E-05 |
| Intermediate | DUSP26 | 1.66251E-22 | 0.2872 | 0.191 | 0.062 | 2.73051E-18 |
| RERGL_Cells | GADD45GI | 6.17513E-05 | 0.287187 | 0.575 | 0.569 | 1 |
| Terminal | RABAC1 | 8.27895E-08 | 0.287155 | 0.865 | 0.822 | 0.001359735 |
| Contractile | GNAS | 1.93575E-08 | 0.287145 | 0.629 | 0.518 | 0.000317928 |
| Terminal | C1QTNF2 | 7.77607E-07 | 0.287136 | 0.246 | 0.157 | 0.01277141 |
| Contractile | HAX1 | 8.63185E-06 | 0.287082 | 0.465 | 0.386 | 0.14176953 |
| Contractile | CUTA | 9.07004E-08 | 0.28704 | 0.671 | 0.583 | 0.001489664 |
| RERGL_Cells | A1BG | 0.001409752 | 0.287032 | 0.19 | 0.143 | 1 |
| Fibroblasts | SMARCA2 | 2.76044E-25 | 0.287022 | 0.246 | 0.105 | 4.53375E-21 |
| RERGL_Cells | ECI1 | 0.001202591 | 0.286986 | 0.13 | 0.09 | 1 |
| RERGL_Cells | CD27-AS1 | 0.001834415 | 0.286959 | 0.254 | 0.211 | 1 |
| Terminal | C1QB | 1.9551E-05 | 0.28691 | 0.157 | 0.095 | 0.321105229 |
| Terminal | BCAP31 | 5.04239E-05 | 0.286871 | 0.512 | 0.444 | 0.828161603 |
| RERGL_Cells | TAF3 | 1.95901E-05 | 0.286845 | 0.152 | 0.093 | 0.321747081 |
| RERGL_Cells | MAF1 | 0.000201391 | 0.28681 | 0.545 | 0.531 | 1 |
| Pericytes | MRPS25 | 1.49734E-05 | 0.28676 | 0.164 | 0.105 | 0.245923419 |
| Pericytes | CRY1 | 8.62257E-05 | 0.28674 | 0.156 | 0.105 | 1 |
| Fibroblasts | PGM3 | 1.27256E-26 | 0.286695 | 0.231 | 0.094 | 2.09004E-22 |
| Contractile | MPHOSPH | 2.46574E-08 | 0.286678 | 0.539 | 0.435 | 0.000404973 |
| Fibroblasts | MMP19 | 7.53472E-26 | 0.286651 | 0.144 | 0.043 | 1.2375E-21 |
| Fibroblasts | SPSB1 | 5.40193E-11 | 0.286633 | 0.302 | 0.195 | 8.87213E-07 |
| Pericytes | FNBP1 | 2.25808E-06 | 0.286568 | 0.195 | 0.125 | 0.037086753 |
| Pericytes | MLF1 | 6.15601E-05 | 0.286426 | 0.155 | 0.102 | 1 |
| Contractile | KCTD9 | 1.05628E-07 | 0.286349 | 0.212 | 0.127 | 0.001734829 |
| Contractile | IDH2 | 1.24694E-07 | 0.286152 | 0.246 | 0.156 | 0.002047978 |
| RERGL_Cells | ATP5PD | 4.08308E-05 | 0.28614 | 0.581 | 0.561 | 0.670605765 |
| RERGL_Cells | TCF25 | 0.00015024 | 0.285949 | 0.536 | 0.523 | 1 |
| Fibroblasts | TXNDC5 | 2.36663E-25 | 0.285791 | 0.188 | 0.069 | 3.88696E-21 |
| Pericytes | PIP5K1A | 1.30099E-06 | 0.285786 | 0.14 | 0.08 | 0.021367429 |
| Terminal | MTCH1 | 1.18423E-05 | 0.285567 | 0.656 | 0.604 | 0.194497755 |
| Intermediate | CALM2 | 9.94384E-22 | 0.285552 | 1 | 0.982 | 1.63318E-17 |
| Pericytes | ADSS | 0.000150281 | 0.285492 | 0.207 | 0.152 | 1 |
| Terminal | PSAP | 2.23719E-13 | 0.285477 | 0.946 | 0.911 | 3.67435E-09 |
| Pericytes | ARHGAP17 | 4.99819E-08 | 0.285471 | 0.122 | 0.059 | 0.000820903 |
| Fibroblasts | APCDD1 | 3.99042E-28 | 0.285465 | 0.109 | 0.023 | 6.55386E-24 |
| Intermediate | SRGN | 6.76691E-05 | 0.285384 | 0.28 | 0.202 | 1 |
| Pericytes | TMEM123 | 0.000381089 | 0.285374 | 0.362 | 0.316 | 1 |
| RERGL_Cells | CCDC102A | 0.001057762 | 0.285356 | 0.143 | 0.1 | 1 |
| Terminal | CTBS | 6.3433E-06 | 0.285253 | 0.233 | 0.155 | 0.10418233 |
| Fibroblasts | NFIX | 1.4687E-22 | 0.285183 | 0.26 | 0.121 | 2.41219E-18 |
| Contractile | DYNLT3 | 2.36868E-05 | 0.285127 | 0.481 | 0.398 | 0.389031465 |
| Fibroblasts | CSTB | 1.41083E-17 | 0.284963 | 0.744 | 0.583 | 2.31714E-13 |

| | | | | | | |
|---|---|---|---|---|---|---|
| Intermediate | NPDC1 | 2.42809E-08 | 0.284903 | 0.521 | 0.384 | 0.00039879 |
| RERGL_Cells | PDHB | 0.007021627 | 0.284849 | 0.374 | 0.359 | 1 |
| Contractile | TRIM47 | 3.07216E-06 | 0.284827 | 0.345 | 0.259 | 0.050457182 |
| Terminal | ALKBH7 | 0.000848665 | 0.284721 | 0.368 | 0.309 | 1 |
| Terminal | IFNGR2 | 2.23544E-05 | 0.284665 | 0.294 | 0.218 | 0.367148298 |
| Pericytes | DTX3 | 0.00045384 | 0.284653 | 0.209 | 0.159 | 1 |
| Fibroblasts | PYGL | 8.62193E-31 | 0.284644 | 0.183 | 0.057 | 1.41607E-26 |
| Fibroblasts | SPTLC1 | 1.59432E-26 | 0.28456 | 0.392 | 0.197 | 2.61851E-22 |
| Terminal | TM7SF3 | 0.001139767 | 0.284482 | 0.248 | 0.195 | 1 |
| RERGL_Cells | SWAP70 | 0.006554708 | 0.284415 | 0.177 | 0.141 | 1 |
| Pericytes | HMGXB4 | 0.000929625 | 0.284388 | 0.211 | 0.162 | 1 |
| Pericytes | AP2A1 | 0.000187386 | 0.284361 | 0.233 | 0.176 | 1 |
| Contractile | SLC25A34- | 4.32493E-12 | 0.284355 | 0.102 | 0.034 | 7.10327E-08 |
| Pericytes | MAOA | 1.05858E-05 | 0.284165 | 0.231 | 0.161 | 0.173861341 |
| Fibroblasts | CMTM3 | 3.16452E-23 | 0.284043 | 0.285 | 0.136 | 5.19741E-19 |
| Terminal | PDE1A | 3.83429E-05 | 0.283961 | 0.309 | 0.225 | 0.629743974 |
| Pericytes | EIF4G2 | 5.73799E-08 | 0.283954 | 0.749 | 0.723 | 0.000942408 |
| Fibroblasts | ADAMTSL3 | 2.49389E-33 | 0.283836 | 0.117 | 0.021 | 4.09597E-29 |
| Pericytes | HNRNPR | 0.000395174 | 0.283834 | 0.469 | 0.44 | 1 |
| Pericytes | IFT43 | 0.001333863 | 0.283817 | 0.26 | 0.217 | 1 |
| RERGL_Cells | CPNE2 | 4.22805E-07 | 0.283812 | 0.124 | 0.062 | 0.006944152 |
| Contractile | UBE2D2 | 4.65757E-07 | 0.283793 | 0.563 | 0.469 | 0.007649587 |
| Pericytes | ATF4 | 2.17966E-06 | 0.2836 | 0.62 | 0.584 | 0.035798787 |
| Terminal | RGS10 | 1.72984E-06 | 0.283537 | 0.163 | 0.095 | 0.028410941 |
| Fibroblasts | GNL3 | 2.89325E-21 | 0.283463 | 0.381 | 0.208 | 4.75188E-17 |
| RERGL_Cells | ADAMTS1 | 1.09614E-07 | 0.283381 | 0.515 | 0.399 | 0.001800292 |
| Pericytes | MAFB | 0.001459697 | 0.283364 | 0.224 | 0.175 | 1 |
| Terminal | SESN3 | 6.76953E-07 | 0.283185 | 0.122 | 0.061 | 0.01111827 |
| Fibroblasts | SDF4 | 6.1693E-17 | 0.283167 | 0.605 | 0.411 | 1.01325E-12 |
| RERGL_Cells | SNRK | 1.59433E-07 | 0.283092 | 0.13 | 0.064 | 0.002618534 |
| Intermediate | ICAM3 | 1.30391E-25 | 0.283051 | 0.142 | 0.034 | 2.14154E-21 |
| Pericytes | ZNF331 | 0.000342653 | 0.283005 | 0.176 | 0.128 | 1 |
| Terminal | WWP2 | 0.000245322 | 0.282981 | 0.148 | 0.097 | 1 |
| Terminal | S100A13 | 2.92869E-06 | 0.282938 | 0.582 | 0.48 | 0.048100767 |
| Terminal | ZFAS1 | 0.000828013 | 0.282822 | 0.606 | 0.576 | 1 |
| Fibroblasts | TM9SF1 | 1.08699E-19 | 0.282607 | 0.384 | 0.219 | 1.78528E-15 |
| Terminal | THBS2 | 5.54362E-07 | 0.282589 | 0.462 | 0.354 | 0.009104845 |
| Fibroblasts | P3H3 | 5.27581E-21 | 0.282455 | 0.245 | 0.116 | 8.66499E-17 |
| RERGL_Cells | IGHG4 | 7.57818E-05 | 0.282455 | 0.139 | 0.086 | 1 |
| RERGL_Cells | UBL5 | 7.85501E-09 | 0.282343 | 0.827 | 0.804 | 0.000129011 |
| Pericytes | C1QTNF1 | 3.9749E-05 | 0.28234 | 0.465 | 0.4 | 0.652837958 |
| Pericytes | TSPYL2 | 0.006528211 | 0.282175 | 0.258 | 0.225 | 1 |
| Terminal | TP53I13 | 0.00045051 | 0.282096 | 0.207 | 0.15 | 1 |
| RERGL_Cells | NSMCE1 | 0.003125909 | 0.282093 | 0.344 | 0.319 | 1 |
| Contractile | PRKAR1A | 4.30295E-06 | 0.282014 | 0.677 | 0.609 | 0.070671685 |
| RERGL_Cells | NUCKS1 | 8.76611E-07 | 0.281937 | 0.729 | 0.705 | 0.014397453 |
| Contractile | STN1 | 4.71937E-06 | 0.281925 | 0.142 | 0.082 | 0.077510905 |
| Terminal | EMC4 | 0.000503783 | 0.281865 | 0.516 | 0.478 | 1 |
| Pericytes | FBXO3 | 0.003163952 | 0.281858 | 0.202 | 0.164 | 1 |
| Fibroblasts | SPHK1 | 3.51644E-21 | 0.281785 | 0.133 | 0.045 | 5.7754E-17 |
| Contractile | POLR3GL | 4.95183E-05 | 0.281717 | 0.309 | 0.236 | 0.813287912 |
| Terminal | EFEMP2 | 0.00010321 | 0.281676 | 0.564 | 0.498 | 1 |
| Terminal | COL6A1 | 1.53431E-06 | 0.281595 | 0.776 | 0.738 | 0.025199555 |
| Contractile | HNRNPH3 | 2.47872E-07 | 0.281517 | 0.523 | 0.429 | 0.004071047 |
| Terminal | TIMP3 | 1.39148E-11 | 0.281476 | 0.619 | 0.824 | 2.28536E-07 |
| Contractile | NUTF2 | 1.9842E-05 | 0.281452 | 0.359 | 0.276 | 0.325885288 |

| | | | | | | |
|---|---|---|---|---|---|---|
| Pericytes | TRPV2 | 3.27461E-05 | 0.281433 | 0.173 | 0.114 | 0.537821201 |
| RERGL_Cells | ZNF512 | 0.000639447 | 0.281418 | 0.115 | 0.074 | 1 |
| Terminal | YIPF5 | 0.000188885 | 0.281252 | 0.281 | 0.214 | 1 |
| Terminal | UBB | 8.0761E-09 | 0.281211 | 0.985 | 0.949 | 0.000132642 |
| Intermediate | ANXA3 | 8.38607E-21 | 0.281175 | 0.116 | 0.028 | 1.37733E-16 |
| Fibroblasts | DNAJC10 | 2.19561E-18 | 0.281092 | 0.323 | 0.179 | 3.60606E-14 |
| Fibroblasts | ASAP3 | 1.27484E-40 | 0.280879 | 0.114 | 0.014 | 2.0938E-36 |
| Fibroblasts | GPX8 | 9.45537E-20 | 0.280836 | 0.269 | 0.137 | 1.55295E-15 |
| Intermediate | IFI27L2 | 1.02393E-09 | 0.280794 | 0.473 | 0.323 | 1.6817E-05 |
| Contractile | GMPR2 | 2.3398E-07 | 0.280781 | 0.24 | 0.152 | 0.003842886 |
| RERGL_Cells | DHPS | 0.004178072 | 0.280762 | 0.293 | 0.259 | 1 |
| Pericytes | THOC6 | 0.000105767 | 0.280758 | 0.16 | 0.109 | 1 |
| Intermediate | SLC22A3 | 1.23134E-10 | 0.280752 | 0.194 | 0.096 | 2.02235E-06 |
| Fibroblasts | TMEM91 | 1.553E-23 | 0.280653 | 0.136 | 0.043 | 2.55065E-19 |
| RERGL_Cells | HDAC1 | 0.004205059 | 0.280575 | 0.205 | 0.166 | 1 |
| Contractile | NT5C3B | 4.6105E-06 | 0.280505 | 0.345 | 0.258 | 0.075722775 |
| RERGL_Cells | EMD | 0.0037259 | 0.280495 | 0.318 | 0.287 | 1 |
| RERGL_Cells | PBRM1 | 0.006391381 | 0.280475 | 0.162 | 0.126 | 1 |
| Intermediate | WWP2 | 4.60878E-17 | 0.280394 | 0.209 | 0.085 | 7.56946E-13 |
| Intermediate | COPS9 | 1.11331E-08 | 0.280228 | 0.688 | 0.566 | 0.00018285 |
| Contractile | PDZRN3 | 4.26864E-10 | 0.280169 | 0.114 | 0.046 | 7.01082E-06 |
| Contractile | SNRNP25 | 3.48581E-09 | 0.280059 | 0.168 | 0.085 | 5.7251E-05 |
| RERGL_Cells | SEPTIN1 | 0.008586708 | 0.280001 | 0.126 | 0.093 | 1 |
| RERGL_Cells | STARD3NL | 0.000997331 | 0.279962 | 0.184 | 0.138 | 1 |
| Pericytes | MYOF | 4.5183E-06 | 0.27993 | 0.144 | 0.085 | 0.074208621 |
| Fibroblasts | FKBP2 | 1.54393E-22 | 0.279736 | 0.345 | 0.177 | 2.53575E-18 |
| Fibroblasts | SPCS2 | 2.57332E-16 | 0.279705 | 0.801 | 0.676 | 4.22643E-12 |
| RERGL_Cells | ANKRD12 | 0.001535025 | 0.279613 | 0.41 | 0.382 | 1 |
| Terminal | TP53TG1 | 0.006882149 | 0.279568 | 0.2 | 0.16 | 1 |
| Pericytes | GRWD1 | 0.002076132 | 0.279532 | 0.173 | 0.132 | 1 |
| Contractile | ENSA | 1.3215E-05 | 0.279452 | 0.529 | 0.446 | 0.217043133 |
| Contractile | HNRNPM | 5.59804E-09 | 0.279426 | 0.645 | 0.544 | 9.19421E-05 |
| Contractile | PTP4A3 | 4.90877E-21 | 0.27934 | 0.447 | 0.225 | 8.06216E-17 |
| Terminal | BBX | 0.00013754 | 0.279327 | 0.475 | 0.403 | 1 |
| Contractile | CAV3 | 6.83159E-19 | 0.279321 | 0.1 | 0.023 | 1.12202E-14 |
| Contractile | TENT5B | 2.87599E-15 | 0.279241 | 0.124 | 0.04 | 4.72353E-11 |
| Terminal | MRAP2 | 9.35168E-05 | 0.279189 | 0.174 | 0.113 | 1 |
| Pericytes | HNRNPM | 3.32462E-05 | 0.278972 | 0.582 | 0.553 | 0.546034887 |
| Intermediate | NBDY | 5.31479E-12 | 0.278943 | 0.535 | 0.362 | 8.729E-08 |
| Intermediate | AGA | 2.332E-17 | 0.278939 | 0.206 | 0.082 | 3.83008E-13 |
| Intermediate | CALCRL | 7.63899E-12 | 0.278833 | 0.211 | 0.104 | 1.25463E-07 |
| Fibroblasts | FAM20C | 6.45563E-27 | 0.278803 | 0.136 | 0.037 | 1.06027E-22 |
| Intermediate | HAAO | 2.27101E-14 | 0.278696 | 0.288 | 0.144 | 3.72991E-10 |
| Pericytes | COTL1 | 0.003196052 | 0.278684 | 0.136 | 0.1 | 1 |
| Contractile | RILPL2 | 8.07965E-09 | 0.278671 | 0.351 | 0.228 | 0.0001327 |
| Intermediate | MGAT4C | 1.16973E-16 | 0.278666 | 0.112 | 0.032 | 1.92117E-12 |
| Pericytes | PRKAA1 | 0.001000931 | 0.278659 | 0.185 | 0.139 | 1 |
| Fibroblasts | HNMT | 1.50411E-17 | 0.278582 | 0.456 | 0.289 | 2.47035E-13 |
| Terminal | GNG5 | 1.80602E-05 | 0.27858 | 0.617 | 0.549 | 0.296620961 |
| Fibroblasts | PRRX1 | 3.96208E-23 | 0.278496 | 0.346 | 0.176 | 6.50732E-19 |
| Fibroblasts | FOXP2 | 2.02716E-45 | 0.278492 | 0.105 | 0.008 | 3.3294E-41 |
| Intermediate | ANK3 | 7.64923E-20 | 0.278468 | 0.153 | 0.047 | 1.25631E-15 |
| RERGL_Cells | DNAJC9 | 0.001069505 | 0.278464 | 0.152 | 0.107 | 1 |
| Contractile | R3HCC1 | 3.33735E-09 | 0.278436 | 0.246 | 0.143 | 5.48126E-05 |
| Contractile | PAAF1 | 1.09177E-09 | 0.278409 | 0.158 | 0.077 | 1.79313E-05 |
| Terminal | EIF3L | 1.66712E-05 | 0.278388 | 0.647 | 0.593 | 0.273808595 |

| | | | | | | |
|---|---|---|---|---|---|---|
| Pericytes | SLC25A6 | 2.40308E-12 | 0.278384 | 0.911 | 0.889 | 3.94682E-08 |
| Terminal | UFC1 | 0.000112109 | 0.27833 | 0.516 | 0.454 | 1 |
| Pericytes | HDAC5 | 0.000937882 | 0.278199 | 0.242 | 0.194 | 1 |
| Pericytes | SMARCD3 | 1.13368E-06 | 0.278188 | 0.289 | 0.202 | 0.018619487 |
| Terminal | EID1 | 3.72936E-09 | 0.278071 | 0.952 | 0.907 | 6.12511E-05 |
| RERGL_Cells | PFDN5 | 3.48742E-11 | 0.277767 | 0.912 | 0.884 | 5.72775E-07 |
| RERGL_Cells | TAF9B | 1.15692E-06 | 0.277735 | 0.165 | 0.094 | 0.019001222 |
| Pericytes | LONRF1 | 2.6058E-06 | 0.277701 | 0.136 | 0.079 | 0.042797686 |
| Terminal | TOB2 | 0.006123603 | 0.277593 | 0.29 | 0.247 | 1 |
| Pericytes | IMMT | 0.001378497 | 0.277527 | 0.233 | 0.188 | 1 |
| Intermediate | DYNLT3 | 5.82246E-10 | 0.277507 | 0.548 | 0.385 | 9.56281E-06 |
| Terminal | FBXO32 | 0.000550246 | 0.277461 | 0.29 | 0.223 | 1 |
| Pericytes | MTREX | 0.001389178 | 0.277356 | 0.173 | 0.131 | 1 |
| RERGL_Cells | NDUFS3 | 0.003815852 | 0.277347 | 0.333 | 0.31 | 1 |
| Intermediate | METTL5 | 8.89502E-10 | 0.277284 | 0.402 | 0.263 | 1.46092E-05 |
| Contractile | COX4I1 | 5.82629E-12 | 0.277067 | 0.92 | 0.859 | 9.5691E-08 |
| Pericytes | DAPK3 | 0.000573791 | 0.27706 | 0.307 | 0.258 | 1 |
| Terminal | DCTN3 | 0.000341006 | 0.277025 | 0.519 | 0.467 | 1 |
| Fibroblasts | RAB32 | 3.73917E-21 | 0.277004 | 0.346 | 0.188 | 6.1412E-17 |
| Pericytes | CYP51A1 | 0.000264911 | 0.276916 | 0.138 | 0.093 | 1 |
| Fibroblasts | ARSG | 4.15689E-26 | 0.276785 | 0.191 | 0.069 | 6.82727E-22 |
| Contractile | AIP | 3.14533E-05 | 0.276753 | 0.375 | 0.294 | 0.516588473 |
| RERGL_Cells | PDCD10 | 0.005435583 | 0.276707 | 0.278 | 0.245 | 1 |
| Fibroblasts | TXNDC15 | 2.05803E-18 | 0.276664 | 0.415 | 0.247 | 3.3801E-14 |
| Terminal | ELOF1 | 0.002685089 | 0.276599 | 0.305 | 0.262 | 1 |
| RERGL_Cells | IPO5 | 0.004661602 | 0.276581 | 0.212 | 0.175 | 1 |
| RERGL_Cells | DCTN2 | 0.003086251 | 0.276504 | 0.491 | 0.498 | 1 |
| Pericytes | DYNC1I2 | 1.79643E-06 | 0.276327 | 0.593 | 0.551 | 0.029504543 |
| Fibroblasts | RECK | 4.68364E-26 | 0.276177 | 0.181 | 0.063 | 7.69241E-22 |
| Pericytes | OSTM1 | 0.002249546 | 0.276151 | 0.249 | 0.21 | 1 |
| Fibroblasts | GOLGA2 | 1.39575E-19 | 0.276106 | 0.354 | 0.197 | 2.29237E-15 |
| Terminal | PKIG | 9.51571E-05 | 0.275995 | 0.564 | 0.497 | 1 |
| Terminal | ANXA7 | 1.39582E-05 | 0.275891 | 0.625 | 0.585 | 0.229248824 |
| Pericytes | GJA5 | 0.000613364 | 0.275884 | 0.102 | 0.064 | 1 |
| Fibroblasts | NRBF2 | 7.76436E-25 | 0.27587 | 0.254 | 0.11 | 1.27522E-20 |
| RERGL_Cells | CPED1 | 0.001730915 | 0.275851 | 0.212 | 0.164 | 1 |
| Intermediate | RCAN2 | 1.04499E-14 | 0.275808 | 0.548 | 0.322 | 1.71629E-10 |
| RERGL_Cells | AAMP | 0.006322452 | 0.275731 | 0.41 | 0.408 | 1 |
| RERGL_Cells | RABGAP1 | 0.000225422 | 0.27567 | 0.135 | 0.086 | 1 |
| Contractile | PRDX1 | 5.29885E-12 | 0.275661 | 0.908 | 0.859 | 8.70283E-08 |
| Contractile | LAMA5 | 4.29251E-07 | 0.275653 | 0.124 | 0.062 | 0.007050014 |
| Intermediate | MIF4GD | 3.89314E-13 | 0.27557 | 0.275 | 0.14 | 6.3941E-09 |
| Pericytes | FAM110B | 1.35137E-06 | 0.275543 | 0.127 | 0.069 | 0.022194912 |
| RERGL_Cells | TP53I11 | 3.22285E-05 | 0.275534 | 0.137 | 0.081 | 0.529320602 |
| Pericytes | MDK | 0.008681091 | 0.275424 | 0.215 | 0.176 | 1 |
| Terminal | SP100 | 0.00027561 | 0.275387 | 0.368 | 0.301 | 1 |
| Intermediate | COL12A1 | 8.10307E-08 | 0.275251 | 0.157 | 0.084 | 0.001330849 |
| RERGL_Cells | VPS28 | 0.001928835 | 0.27521 | 0.547 | 0.565 | 1 |
| Contractile | FKBP8 | 2.24082E-06 | 0.275187 | 0.699 | 0.63 | 0.036803258 |
| Intermediate | TNS3 | 4.16332E-22 | 0.274972 | 0.138 | 0.037 | 6.83784E-18 |
| Terminal | FAM32A | 0.000763632 | 0.274967 | 0.357 | 0.301 | 1 |
| Fibroblasts | DNM1 | 6.07175E-41 | 0.27479 | 0.111 | 0.013 | 9.97224E-37 |
| Contractile | EMC10 | 2.0708E-05 | 0.27474 | 0.423 | 0.342 | 0.340107725 |
| RERGL_Cells | MRPL2 | 0.008967074 | 0.274616 | 0.286 | 0.256 | 1 |
| Pericytes | ENPP4 | 3.58712E-05 | 0.274577 | 0.116 | 0.068 | 0.589148208 |
| RERGL_Cells | NDUFAB1 | 0.000159671 | 0.274563 | 0.517 | 0.492 | 1 |

| | | | | | | |
|---|---|---|---|---|---|---|
| RERGL_Cells | EGLN1 | 0.000453947 | 0.274502 | 0.162 | 0.113 | 1 |
| RERGL_Cells | C1QTNF2 | 0.002973857 | 0.274476 | 0.205 | 0.163 | 1 |
| Contractile | NUDT16 | 8.07074E-06 | 0.274353 | 0.206 | 0.136 | 0.132553814 |
| Pericytes | USP47 | 0.002652056 | 0.274316 | 0.202 | 0.162 | 1 |
| Pericytes | M6PR | 0.000895767 | 0.274299 | 0.282 | 0.235 | 1 |
| RERGL_Cells | PLEKHH3 | 1.50981E-06 | 0.274162 | 0.113 | 0.057 | 0.024797191 |
| RERGL_Cells | SMARCE1 | 0.002096231 | 0.274095 | 0.382 | 0.358 | 1 |
| Contractile | EXOC7 | 6.29813E-06 | 0.274016 | 0.305 | 0.22 | 0.103440501 |
| Contractile | HINT2 | 4.50001E-06 | 0.274001 | 0.327 | 0.241 | 0.073908152 |
| Intermediate | HSPG2 | 2.43361E-08 | 0.273986 | 0.32 | 0.206 | 0.000399697 |
| RERGL_Cells | HTATSF1 | 0.00450937 | 0.273972 | 0.265 | 0.229 | 1 |
| Contractile | DDB2 | 1.5497E-05 | 0.273889 | 0.192 | 0.124 | 0.25452308 |
| Pericytes | MICAL1 | 7.63217E-07 | 0.273869 | 0.156 | 0.09 | 0.012535071 |
| Pericytes | PTRH2 | 0.003180586 | 0.273859 | 0.109 | 0.076 | 1 |
| Contractile | RSF1 | 3.84876E-05 | 0.27383 | 0.371 | 0.294 | 0.632120371 |
| Contractile | C9orf16 | 6.87875E-05 | 0.2738 | 0.443 | 0.369 | 1 |
| RERGL_Cells | ZNF37A | 0.004365248 | 0.273751 | 0.141 | 0.104 | 1 |
| Terminal | MRFAP1 | 3.50898E-07 | 0.273704 | 0.789 | 0.731 | 0.005763156 |
| Contractile | UBA52 | 4.71996E-16 | 0.273636 | 0.974 | 0.96 | 7.75206E-12 |
| RERGL_Cells | PDE1B | 2.57395E-06 | 0.273599 | 0.111 | 0.056 | 0.04227461 |
| RERGL_Cells | RNASEK | 3.94867E-07 | 0.273549 | 0.795 | 0.78 | 0.006485294 |
| RERGL_Cells | CYP7B1 | 0.001567593 | 0.273487 | 0.118 | 0.078 | 1 |
| RERGL_Cells | EIF3L | 1.57E-05 | 0.273368 | 0.618 | 0.597 | 0.257857599 |
| Pericytes | RAB10 | 0.002357855 | 0.273187 | 0.367 | 0.338 | 1 |
| RERGL_Cells | H2AFZ | 0.000171506 | 0.273167 | 0.585 | 0.581 | 1 |
| Contractile | VTI1B | 1.04948E-06 | 0.273009 | 0.345 | 0.248 | 0.017236734 |
| Contractile | WDR13 | 6.59118E-06 | 0.272995 | 0.387 | 0.301 | 0.108253548 |
| Terminal | SNHG6 | 0.000597199 | 0.272989 | 0.584 | 0.536 | 1 |
| Intermediate | HEXA | 9.3714E-09 | 0.272921 | 0.46 | 0.323 | 0.000153916 |
| RERGL_Cells | DAPK3 | 0.001706852 | 0.272782 | 0.306 | 0.26 | 1 |
| Intermediate | PGM2L1 | 4.82461E-16 | 0.272777 | 0.148 | 0.052 | 7.92394E-12 |
| Intermediate | YIPF2 | 2.29009E-11 | 0.272655 | 0.387 | 0.239 | 3.76124E-07 |
| RERGL_Cells | ID2 | 3.50663E-10 | 0.272651 | 0.679 | 0.553 | 5.7593E-06 |
| RERGL_Cells | HSD17B10 | 0.000239672 | 0.27259 | 0.436 | 0.394 | 1 |
| Fibroblasts | SSR4 | 6.50965E-18 | 0.272588 | 0.866 | 0.699 | 1.06914E-13 |
| Intermediate | CD58 | 1.03582E-13 | 0.272477 | 0.338 | 0.187 | 1.70123E-09 |
| Pericytes | ACSS1 | 8.37194E-06 | 0.272472 | 0.14 | 0.083 | 0.13750067 |
| Contractile | HRAS | 4.22664E-06 | 0.272392 | 0.202 | 0.13 | 0.069418328 |
| Contractile | PTMA | 5.04857E-18 | 0.272322 | 0.998 | 0.994 | 8.29177E-14 |
| Terminal | ZNF385D | 0.000702041 | 0.272047 | 0.159 | 0.108 | 1 |
| Contractile | CACYBP | 6.25494E-05 | 0.272029 | 0.373 | 0.301 | 1 |
| Fibroblasts | ALG2 | 1.97769E-26 | 0.272018 | 0.231 | 0.093 | 3.24816E-22 |
| RERGL_Cells | CD82 | 0.002881264 | 0.272013 | 0.186 | 0.142 | 1 |
| RERGL_Cells | MRPS10 | 0.004957552 | 0.271928 | 0.188 | 0.149 | 1 |
| Intermediate | TCEAL4 | 6.31548E-10 | 0.271926 | 0.802 | 0.675 | 1.03726E-05 |
| Contractile | TXN | 2.6115E-07 | 0.271876 | 0.752 | 0.682 | 0.004289122 |
| Pericytes | SMARCB1 | 0.000377348 | 0.271754 | 0.3 | 0.25 | 1 |
| Contractile | NAP1L3 | 5.7181E-10 | 0.271682 | 0.166 | 0.081 | 9.3914E-06 |
| Fibroblasts | TLE5 | 1.56074E-17 | 0.271528 | 0.378 | 0.221 | 2.56336E-13 |
| Intermediate | TMCO3 | 5.33854E-12 | 0.271391 | 0.303 | 0.168 | 8.76801E-08 |
| RERGL_Cells | NSMCE4A | 0.003947121 | 0.271388 | 0.194 | 0.156 | 1 |
| Terminal | VAMP5 | 0.000169332 | 0.271364 | 0.471 | 0.416 | 1 |
| Terminal | B2M | 9.98775E-13 | 0.271154 | 1 | 0.999 | 1.64039E-08 |
| RERGL_Cells | TNFAIP8 | 7.20481E-07 | 0.271107 | 0.132 | 0.069 | 0.011833184 |
| Terminal | FAU | 5.65536E-12 | 0.271059 | 0.996 | 0.98 | 9.28836E-08 |
| Pericytes | TEAD2 | 0.003061755 | 0.270997 | 0.165 | 0.126 | 1 |

| | | | | | | |
|---|---|---|---|---|---|---|
| RERGL_Cells | DAAM2 | 0.000105156 | 0.270907 | 0.128 | 0.078 | 1 |
| Contractile | LAS1L | 7.2262E-06 | 0.270891 | 0.172 | 0.108 | 0.118683156 |
| RERGL_Cells | TCEA1 | 0.001339005 | 0.270876 | 0.47 | 0.455 | 1 |
| Terminal | MAN1B1 | 0.000218408 | 0.270872 | 0.222 | 0.161 | 1 |
| Fibroblasts | SMOC1 | 5.45412E-23 | 0.27076 | 0.163 | 0.058 | 8.95784E-19 |
| Fibroblasts | FAM3C | 5.22948E-18 | 0.270704 | 0.609 | 0.412 | 8.5889E-14 |
| Terminal | CD74 | 0.002972245 | 0.270649 | 0.427 | 0.377 | 1 |
| Terminal | LRRFIP1 | 2.19771E-05 | 0.270548 | 0.407 | 0.32 | 0.360952498 |
| Pericytes | SRI | 2.02962E-05 | 0.270481 | 0.633 | 0.605 | 0.333345528 |
| RERGL_Cells | KMT5B | 0.00821183 | 0.270384 | 0.126 | 0.093 | 1 |
| Contractile | GADD45GI | 2.20146E-08 | 0.270355 | 0.665 | 0.554 | 0.000361568 |
| Fibroblasts | BNIP3 | 8.64922E-19 | 0.270305 | 0.306 | 0.164 | 1.42055E-14 |
| Fibroblasts | POLR2L | 1.05646E-18 | 0.270223 | 0.883 | 0.756 | 1.73512E-14 |
| Contractile | PMPCB | 7.94426E-05 | 0.270202 | 0.309 | 0.239 | 1 |
| Pericytes | TMEM165 | 0.000542386 | 0.270165 | 0.48 | 0.451 | 1 |
| Contractile | RNASEH2C | 2.07082E-06 | 0.270101 | 0.531 | 0.44 | 0.034011142 |
| Terminal | SNHG5 | 0.002361951 | 0.270028 | 0.669 | 0.657 | 1 |
| Intermediate | AC245297 | 1.13414E-09 | 0.269964 | 0.323 | 0.199 | 1.86271E-05 |
| Intermediate | DPP7 | 1.0877E-07 | 0.269794 | 0.536 | 0.404 | 0.001786433 |
| Fibroblasts | PRKCSH | 1.24636E-17 | 0.269755 | 0.44 | 0.268 | 2.04702E-13 |
| Pericytes | PSMB2 | 2.91281E-06 | 0.269703 | 0.491 | 0.434 | 0.047839997 |
| Pericytes | SNHG17 | 8.96131E-05 | 0.269654 | 0.176 | 0.122 | 1 |
| Pericytes | TMA16 | 0.003952533 | 0.269581 | 0.229 | 0.194 | 1 |
| Pericytes | ELP5 | 0.000617415 | 0.269555 | 0.2 | 0.151 | 1 |
| Intermediate | CRTAP | 5.68677E-09 | 0.269516 | 0.54 | 0.389 | 9.33996E-05 |
| Terminal | PDCD6 | 0.000135743 | 0.269465 | 0.431 | 0.367 | 1 |
| Intermediate | CCDC107 | 1.34466E-10 | 0.269264 | 0.692 | 0.512 | 2.20847E-06 |
| Pericytes | NFATC4 | 4.28962E-07 | 0.269219 | 0.127 | 0.067 | 0.007045277 |
| Fibroblasts | GMPPB | 2.04903E-23 | 0.26901 | 0.213 | 0.088 | 3.36533E-19 |
| Contractile | GAMT | 3.29654E-07 | 0.268993 | 0.283 | 0.187 | 0.005414245 |
| Fibroblasts | HCG11 | 1.52187E-29 | 0.26894 | 0.146 | 0.039 | 2.49951E-25 |
| Fibroblasts | STK17B | 2.21721E-25 | 0.268915 | 0.156 | 0.051 | 3.64155E-21 |
| Fibroblasts | NFIC | 1.12533E-18 | 0.268907 | 0.642 | 0.438 | 1.84824E-14 |
| Fibroblasts | LMO4 | 4.47256E-17 | 0.268798 | 0.439 | 0.271 | 7.34574E-13 |
| Fibroblasts | CKAP4 | 3.28739E-16 | 0.26877 | 0.288 | 0.161 | 5.39921E-12 |
| Intermediate | CPPED1 | 6.69016E-17 | 0.268767 | 0.14 | 0.046 | 1.09879E-12 |
| Intermediate | DUSP6 | 9.141E-14 | 0.268763 | 0.236 | 0.112 | 1.50132E-09 |
| Pericytes | U2AF2 | 0.006417261 | 0.268665 | 0.207 | 0.172 | 1 |
| RERGL_Cells | C1QBP | 0.00257064 | 0.268604 | 0.442 | 0.443 | 1 |
| Fibroblasts | PLAGL1 | 8.43849E-23 | 0.268538 | 0.29 | 0.139 | 1.38594E-18 |
| RERGL_Cells | FIS1 | 1.09382E-05 | 0.268529 | 0.645 | 0.633 | 0.17964979 |
| Terminal | DGCR6L | 0.000577879 | 0.2685 | 0.301 | 0.242 | 1 |
| Intermediate | COL5A2 | 4.75127E-14 | 0.268469 | 0.204 | 0.091 | 7.80349E-10 |
| Fibroblasts | SRPRA | 1.92227E-21 | 0.26843 | 0.37 | 0.201 | 3.15714E-17 |
| Contractile | MRPS26 | 5.95265E-05 | 0.268425 | 0.355 | 0.281 | 0.97766317 |
| Terminal | DAAM1 | 0.00119188 | 0.268389 | 0.401 | 0.341 | 1 |
| Contractile | FAM210B | 3.03974E-05 | 0.268315 | 0.226 | 0.158 | 0.49924668 |
| Terminal | MLLT11 | 1.73566E-08 | 0.268293 | 0.126 | 0.057 | 0.000285064 |
| Fibroblasts | KRT10 | 4.58996E-14 | 0.268221 | 0.625 | 0.454 | 7.53854E-10 |
| RERGL_Cells | EDNRB | 2.1783E-05 | 0.268217 | 0.182 | 0.116 | 0.3577634 |
| Intermediate | TMEM106 | 5.10632E-10 | 0.268169 | 0.396 | 0.257 | 8.38662E-06 |
| Fibroblasts | ATP6V0E1 | 1.65879E-15 | 0.26815 | 0.811 | 0.653 | 2.72439E-11 |
| RERGL_Cells | LIMS2 | 9.4604E-05 | 0.268108 | 0.244 | 0.179 | 1 |
| Terminal | PODNL1 | 6.22821E-11 | 0.268049 | 0.137 | 0.057 | 1.02292E-06 |
| Terminal | BEX3 | 0.000682947 | 0.268001 | 0.708 | 0.67 | 1 |
| Contractile | ZFAND2B | 6.63105E-06 | 0.267933 | 0.194 | 0.123 | 0.108908348 |

| | | | | | | |
|---|---|---|---|---|---|---|
| Contractile | PIP5K1B | 8.2536E-15 | 0.267899 | 0.122 | 0.039 | 1.35557E-10 |
| Intermediate | DYRK4 | 1.76243E-12 | 0.267866 | 0.234 | 0.116 | 2.89462E-08 |
| RERGL_Cells | UBE2W | 0.003542482 | 0.267783 | 0.201 | 0.159 | 1 |
| Fibroblasts | CD63 | 6.95281E-31 | 0.267687 | 0.997 | 0.992 | 1.14193E-26 |
| Pericytes | KLHL23 | 1.1852E-10 | 0.267651 | 0.18 | 0.087 | 1.94657E-06 |
| Intermediate | CAP1 | 8.74411E-12 | 0.267567 | 0.813 | 0.672 | 1.43613E-07 |
| Fibroblasts | CBR3 | 8.27923E-16 | 0.267518 | 0.206 | 0.102 | 1.35978E-11 |
| RERGL_Cells | RTN2 | 0.000206853 | 0.267511 | 0.107 | 0.064 | 1 |
| RERGL_Cells | FAM76A | 0.000798208 | 0.267456 | 0.156 | 0.11 | 1 |
| Fibroblasts | SKAP2 | 3.08486E-27 | 0.267388 | 0.122 | 0.03 | 5.06657E-23 |
| Contractile | TMEM14A | 0.000177767 | 0.267331 | 0.389 | 0.32 | 1 |
| Terminal | PAXX | 0.000605637 | 0.267306 | 0.246 | 0.188 | 1 |
| Terminal | FUCA1 | 0.00362311 | 0.267244 | 0.229 | 0.182 | 1 |
| Contractile | CCT4 | 4.09496E-06 | 0.267184 | 0.569 | 0.477 | 0.067255581 |
| Contractile | DYNLL2 | 1.19363E-09 | 0.267002 | 0.144 | 0.067 | 1.96041E-05 |
| Contractile | NUDT2 | 5.14164E-07 | 0.266872 | 0.214 | 0.132 | 0.008444627 |
| RERGL_Cells | HNRNPH2 | 7.90651E-05 | 0.266739 | 0.483 | 0.459 | 1 |
| Terminal | TCF4 | 5.84702E-05 | 0.266707 | 0.46 | 0.377 | 0.960314847 |
| Fibroblasts | RICTOR | 3.33058E-19 | 0.26667 | 0.271 | 0.137 | 5.47015E-15 |
| Terminal | NGRN | 0.000162576 | 0.266493 | 0.575 | 0.519 | 1 |
| RERGL_Cells | PTEN | 0.006121008 | 0.26641 | 0.226 | 0.187 | 1 |
| Intermediate | CRYAB | 1.02405E-20 | 0.266331 | 0.912 | 0.781 | 1.6819E-16 |
| Terminal | TMEM9 | 0.004441694 | 0.266324 | 0.298 | 0.253 | 1 |
| Terminal | TNFRSF11I | 1.81604E-10 | 0.266311 | 0.301 | 0.171 | 2.98267E-06 |
| Fibroblasts | UBXN4 | 9.04832E-18 | 0.266292 | 0.596 | 0.393 | 1.4861E-13 |
| Terminal | BSG | 7.09199E-08 | 0.266276 | 0.856 | 0.817 | 0.001164788 |
| Fibroblasts | RALGDS | 1.6971E-30 | 0.266229 | 0.108 | 0.02 | 2.78731E-26 |
| Pericytes | PLRG1 | 0.00308107 | 0.266225 | 0.236 | 0.196 | 1 |
| Fibroblasts | SLC39A8 | 6.20097E-29 | 0.266076 | 0.113 | 0.023 | 1.01845E-24 |
| Intermediate | CD46 | 1.64377E-07 | 0.266048 | 0.445 | 0.32 | 0.002699724 |
| Pericytes | KPNA1 | 0.004029205 | 0.265883 | 0.144 | 0.108 | 1 |
| Terminal | TMEM14C | 0.004287871 | 0.265752 | 0.58 | 0.553 | 1 |
| Fibroblasts | TMEM39A | 6.71425E-27 | 0.265735 | 0.226 | 0.088 | 1.10275E-22 |
| Contractile | EPDR1 | 2.0265E-08 | 0.265733 | 0.291 | 0.182 | 0.000332832 |
| Fibroblasts | RNF149 | 5.84596E-21 | 0.265684 | 0.294 | 0.147 | 9.60141E-17 |
| Contractile | CMAS | 1.12355E-06 | 0.265674 | 0.299 | 0.207 | 0.018453157 |
| Contractile | MAPRE1 | 1.64831E-06 | 0.265639 | 0.415 | 0.317 | 0.027071848 |
| Fibroblasts | IL10RB | 3.35877E-23 | 0.26559 | 0.253 | 0.114 | 5.51645E-19 |
| Pericytes | CCNL2 | 0.009530514 | 0.26554 | 0.256 | 0.227 | 1 |
| Terminal | ZNF667-AS | 6.62219E-05 | 0.265447 | 0.281 | 0.206 | 1 |
| Fibroblasts | TM9SF3 | 6.89468E-22 | 0.265331 | 0.338 | 0.176 | 1.13238E-17 |
| Terminal | CLTA | 1.70785E-05 | 0.26531 | 0.643 | 0.604 | 0.280497514 |
| Contractile | NONO | 0.000109611 | 0.265296 | 0.353 | 0.285 | 1 |
| Fibroblasts | KDELR3 | 1.56454E-18 | 0.265217 | 0.289 | 0.153 | 2.5696E-14 |
| Contractile | SMIM30 | 1.77815E-05 | 0.265215 | 0.413 | 0.328 | 0.292043496 |
| Fibroblasts | TCTN1 | 3.80356E-21 | 0.265212 | 0.269 | 0.13 | 6.24697E-17 |
| Terminal | TRIM56 | 0.003909965 | 0.265208 | 0.203 | 0.157 | 1 |
| Contractile | ARHGEF10 | 2.94663E-09 | 0.265137 | 0.124 | 0.055 | 4.83955E-05 |
| Fibroblasts | SYNE3 | 1.19428E-25 | 0.264957 | 0.112 | 0.027 | 1.96149E-21 |
| Intermediate | ACYP2 | 6.60972E-15 | 0.264929 | 0.254 | 0.119 | 1.08558E-10 |
| RERGL_Cells | DCAF7 | 0.002749538 | 0.264903 | 0.154 | 0.113 | 1 |
| Pericytes | ITPA | 0.003656065 | 0.264891 | 0.236 | 0.197 | 1 |
| Fibroblasts | BCL6 | 1.1581E-20 | 0.264871 | 0.177 | 0.07 | 1.90206E-16 |
| Contractile | SMIM10L1 | 2.85945E-07 | 0.264763 | 0.226 | 0.142 | 0.004696362 |
| Contractile | NPRL3 | 0.000298885 | 0.264589 | 0.182 | 0.127 | 1 |
| Pericytes | LPP | 6.43539E-05 | 0.264575 | 0.476 | 0.42 | 1 |

| | | | | | | |
|---|---|---|---|---|---|---|
| Contractile | VPS51 | 2.24429E-05 | 0.264532 | 0.339 | 0.257 | 0.368601986 |
| Terminal | RSF1 | 0.001720541 | 0.26451 | 0.346 | 0.298 | 1 |
| Terminal | PGAM2 | 0.000100974 | 0.264401 | 0.137 | 0.083 | 1 |
| Terminal | NPDC1 | 0.000538595 | 0.26434 | 0.455 | 0.397 | 1 |
| Fibroblasts | RNF146 | 2.51199E-19 | 0.264336 | 0.357 | 0.198 | 4.12569E-15 |
| Pericytes | ACTL6A | 0.002918469 | 0.264306 | 0.218 | 0.178 | 1 |
| Fibroblasts | AKAP13 | 1.31101E-18 | 0.26428 | 0.402 | 0.235 | 2.1532E-14 |
| Intermediate | TMEM120 | 1.46869E-10 | 0.264245 | 0.325 | 0.194 | 2.41218E-06 |
| RERGL_Cells | NGRN | 0.000492961 | 0.26415 | 0.534 | 0.525 | 1 |
| Fibroblasts | ARL4A | 1.01103E-23 | 0.264139 | 0.196 | 0.076 | 1.66051E-19 |
| Contractile | NIPSNAP2 | 3.86857E-06 | 0.264058 | 0.305 | 0.216 | 0.063537389 |
| Fibroblasts | NUDT5 | 7.97876E-18 | 0.264 | 0.256 | 0.133 | 1.31043E-13 |
| Pericytes | FBXL5 | 0.00185642 | 0.263896 | 0.382 | 0.354 | 1 |
| Contractile | PPP2R1A | 1.58522E-06 | 0.263823 | 0.587 | 0.506 | 0.026035707 |
| Fibroblasts | CPXM2 | 1.82508E-16 | 0.263727 | 0.433 | 0.27 | 2.99751E-12 |
| Pericytes | RRN3 | 0.001126272 | 0.26366 | 0.135 | 0.094 | 1 |
| Pericytes | C1QBP | 0.000186326 | 0.263622 | 0.471 | 0.438 | 1 |
| Contractile | AAMDC | 1.02964E-06 | 0.263528 | 0.429 | 0.332 | 0.016910794 |
| Contractile | SF3B6 | 5.98811E-06 | 0.263495 | 0.631 | 0.555 | 0.09834876 |
| Fibroblasts | ZBTB44 | 1.55533E-17 | 0.263363 | 0.123 | 0.044 | 2.55448E-13 |
| Terminal | MAGED1 | 0.002995586 | 0.263328 | 0.283 | 0.232 | 1 |
| RERGL_Cells | RAC1 | 1.40799E-09 | 0.263217 | 0.895 | 0.878 | 2.31248E-05 |
| RERGL_Cells | UBE2M | 0.000293588 | 0.262976 | 0.122 | 0.077 | 1 |
| Pericytes | LEPROT | 0.001460229 | 0.262925 | 0.611 | 0.61 | 1 |
| Intermediate | UCHL1 | 2.09891E-06 | 0.262874 | 0.241 | 0.157 | 0.034472467 |
| Contractile | RTF1 | 7.20898E-05 | 0.26278 | 0.483 | 0.427 | 1 |
| Pericytes | PPP2R1A | 0.000167336 | 0.262766 | 0.538 | 0.514 | 1 |
| Terminal | ARPC1B | 0.001598779 | 0.262764 | 0.571 | 0.536 | 1 |
| Intermediate | TUBA4A | 7.97956E-11 | 0.262714 | 0.196 | 0.097 | 1.31056E-06 |
| RERGL_Cells | PYGB | 0.007589765 | 0.262651 | 0.209 | 0.172 | 1 |
| Fibroblasts | TMEM205 | 5.07336E-16 | 0.262586 | 0.473 | 0.303 | 8.33249E-12 |
| Fibroblasts | EGFR | 1.82187E-28 | 0.262584 | 0.19 | 0.063 | 2.99223E-24 |
| Contractile | EIF3G | 1.93461E-06 | 0.262402 | 0.661 | 0.59 | 0.031774054 |
| Fibroblasts | SLC43A1 | 1.27374E-33 | 0.262364 | 0.143 | 0.032 | 2.09199E-29 |
| RERGL_Cells | RSU1 | 0.001003112 | 0.262316 | 0.449 | 0.431 | 1 |
| Intermediate | SDF4 | 2.58015E-07 | 0.262253 | 0.576 | 0.451 | 0.004237633 |
| Contractile | NDUFS3 | 4.83172E-05 | 0.262195 | 0.379 | 0.303 | 0.793561564 |
| Contractile | PARL | 1.56975E-05 | 0.262171 | 0.311 | 0.231 | 0.257816014 |
| Contractile | ANXA11 | 6.21304E-06 | 0.262154 | 0.643 | 0.561 | 0.102043049 |
| Fibroblasts | SPCS3 | 3.6368E-16 | 0.262085 | 0.36 | 0.215 | 5.97308E-12 |
| RERGL_Cells | CNOT7 | 0.007438748 | 0.261939 | 0.261 | 0.226 | 1 |
| Pericytes | NUS1 | 0.000383553 | 0.261908 | 0.178 | 0.13 | 1 |
| Terminal | SIVA1 | 0.000512244 | 0.261875 | 0.584 | 0.538 | 1 |
| Pericytes | APMAP | 0.002202363 | 0.261851 | 0.271 | 0.231 | 1 |
| Intermediate | TPD52L2 | 9.04499E-12 | 0.261811 | 0.484 | 0.307 | 1.48555E-07 |
| Fibroblasts | LMF2 | 8.01379E-18 | 0.26178 | 0.347 | 0.198 | 1.31618E-13 |
| Terminal | MRPL41 | 0.000139177 | 0.261653 | 0.429 | 0.353 | 1 |
| Terminal | CDKN2C | 0.000254246 | 0.261652 | 0.233 | 0.17 | 1 |
| Intermediate | VOPP1 | 1.97658E-14 | 0.261624 | 0.234 | 0.109 | 3.24633E-10 |
| Contractile | MRPL54 | 3.64544E-06 | 0.261528 | 0.419 | 0.331 | 0.059872678 |
| Terminal | GPC1 | 0.000441525 | 0.261443 | 0.198 | 0.142 | 1 |
| Terminal | ATRAID | 2.3437E-05 | 0.261438 | 0.658 | 0.619 | 0.38492886 |
| Fibroblasts | VAT1 | 8.23684E-19 | 0.26142 | 0.295 | 0.156 | 1.35282E-14 |
| Pericytes | PLEKHB2 | 7.45666E-05 | 0.261377 | 0.153 | 0.101 | 1 |
| Intermediate | PLPP5 | 4.21436E-11 | 0.261369 | 0.256 | 0.139 | 6.92166E-07 |
| Intermediate | STN1 | 1.82929E-16 | 0.261352 | 0.187 | 0.074 | 3.00442E-12 |

| | | | | | | |
|---|---|---|---|---|---|---|
| Contractile | SHISA4 | 0.000343692 | 0.261228 | 0.238 | 0.182 | 1 |
| Fibroblasts | FNDC3B | 4.67341E-16 | 0.261059 | 0.202 | 0.098 | 7.67561E-12 |
| Contractile | TCEA3 | 3.38695E-05 | 0.261055 | 0.226 | 0.157 | 0.556273373 |
| RERGL_Cells | MUS81 | 0.002339617 | 0.260942 | 0.173 | 0.13 | 1 |
| Pericytes | STBD1 | 0.007990787 | 0.260935 | 0.164 | 0.128 | 1 |
| Fibroblasts | NUFIP2 | 8.85138E-20 | 0.260668 | 0.156 | 0.059 | 1.45375E-15 |
| Intermediate | B2M | 1.21409E-18 | 0.260659 | 1 | 0.999 | 1.99402E-14 |
| Intermediate | JMJD8 | 1.05998E-09 | 0.260637 | 0.284 | 0.167 | 1.74092E-05 |
| Contractile | PQBP1 | 1.81636E-05 | 0.260545 | 0.331 | 0.251 | 0.298318349 |
| RERGL_Cells | MEF2D | 0.002383085 | 0.260487 | 0.103 | 0.067 | 1 |
| Pericytes | TEX41 | 2.55722E-07 | 0.260477 | 0.107 | 0.052 | 0.004199974 |
| Contractile | CCNQ | 1.33136E-07 | 0.260443 | 0.17 | 0.095 | 0.00218662 |
| Fibroblasts | MAGI2-AS | 7.83041E-18 | 0.260422 | 0.366 | 0.209 | 1.28607E-13 |
| Pericytes | MAPK10 | 0.003676002 | 0.260407 | 0.16 | 0.122 | 1 |
| RERGL_Cells | POLR2K | 0.000402202 | 0.26028 | 0.509 | 0.495 | 1 |
| Intermediate | CBR1 | 5.0878E-07 | 0.260246 | 0.452 | 0.327 | 0.008356198 |
| Fibroblasts | ZDHHC9 | 5.8023E-31 | 0.260082 | 0.117 | 0.023 | 9.5297E-27 |
| Terminal | KCNS3 | 3.83097E-06 | 0.260074 | 0.111 | 0.057 | 0.062919855 |
| Fibroblasts | BTG3 | 1.3226E-19 | 0.260003 | 0.508 | 0.311 | 2.17224E-15 |
| Terminal | PXMP2 | 0.0095289 | 0.259992 | 0.142 | 0.107 | 1 |
| Terminal | PPP1R13L | 5.66865E-07 | 0.259897 | 0.161 | 0.088 | 0.009310184 |
| Pericytes | SLC35A4 | 0.000155341 | 0.259776 | 0.153 | 0.102 | 1 |
| RERGL_Cells | FKBP8 | 2.10735E-06 | 0.259768 | 0.658 | 0.637 | 0.034611137 |
| Terminal | GMPR | 2.26103E-05 | 0.259746 | 0.192 | 0.124 | 0.371351124 |
| RERGL_Cells | CHMP2A | 0.000500008 | 0.259583 | 0.564 | 0.573 | 1 |
| Intermediate | SPRY1 | 1.16967E-09 | 0.259578 | 0.484 | 0.33 | 1.92107E-05 |
| Intermediate | ATP6V0B | 1.35281E-07 | 0.259483 | 0.589 | 0.445 | 0.002221863 |
| Terminal | DBNDD2 | 1.88927E-07 | 0.259333 | 0.699 | 0.571 | 0.003102937 |
| Contractile | WDR83OS | 5.77093E-08 | 0.259216 | 0.735 | 0.66 | 0.000947817 |
| Contractile | HEBP2 | 0.000176283 | 0.259208 | 0.279 | 0.212 | 1 |
| Contractile | USP16 | 0.001067162 | 0.259169 | 0.389 | 0.326 | 1 |
| Terminal | FBXW5 | 0.000703455 | 0.259108 | 0.333 | 0.269 | 1 |
| RERGL_Cells | ZNF511 | 0.004137463 | 0.25906 | 0.212 | 0.169 | 1 |
| Fibroblasts | A4GALT | 3.26397E-17 | 0.259021 | 0.383 | 0.226 | 5.36075E-13 |
| RERGL_Cells | CPSF7 | 0.002643811 | 0.258979 | 0.179 | 0.137 | 1 |
| Contractile | DCXR | 0.000233312 | 0.258931 | 0.323 | 0.257 | 1 |
| Terminal | KRTCAP2 | 5.19048E-05 | 0.258906 | 0.736 | 0.707 | 0.852484749 |
| Fibroblasts | PER1 | 1.66799E-17 | 0.25883 | 0.262 | 0.134 | 2.7395E-13 |
| Contractile | PHPT1 | 0.000119224 | 0.258811 | 0.661 | 0.587 | 1 |
| Pericytes | EDEM2 | 0.000735766 | 0.258806 | 0.144 | 0.101 | 1 |
| Terminal | PIH1D1 | 0.002590589 | 0.258749 | 0.292 | 0.243 | 1 |
| Terminal | HTRA2 | 0.000649551 | 0.258737 | 0.237 | 0.178 | 1 |
| Intermediate | CCDC90B | 5.95827E-09 | 0.258699 | 0.346 | 0.221 | 9.78586E-05 |
| Intermediate | REEP3 | 1.24327E-09 | 0.258537 | 0.346 | 0.215 | 2.04195E-05 |
| RERGL_Cells | OLA1 | 0.003190094 | 0.258509 | 0.276 | 0.24 | 1 |
| Terminal | BLOC1S2 | 4.21871E-05 | 0.258487 | 0.251 | 0.178 | 0.692880737 |
| RERGL_Cells | FBXO3 | 0.005827503 | 0.258465 | 0.203 | 0.165 | 1 |
| Terminal | NMRK1 | 0.000138909 | 0.258437 | 0.194 | 0.133 | 1 |
| Pericytes | JMJD6 | 0.000504099 | 0.258408 | 0.224 | 0.169 | 1 |
| Terminal | GTF3A | 0.006158482 | 0.258308 | 0.349 | 0.306 | 1 |
| Fibroblasts | GALNS | 1.8734E-26 | 0.258212 | 0.149 | 0.045 | 3.07687E-22 |
| Pericytes | CCT4 | 2.59724E-05 | 0.258122 | 0.522 | 0.484 | 0.426570989 |
| Contractile | CENPX | 0.000816404 | 0.258113 | 0.303 | 0.245 | 1 |
| Fibroblasts | CALHM2 | 1.50772E-19 | 0.257974 | 0.421 | 0.245 | 2.47627E-15 |
| Fibroblasts | DTWD1 | 5.89206E-20 | 0.257887 | 0.356 | 0.193 | 9.67712E-16 |
| Fibroblasts | ELF1 | 1.64625E-20 | 0.257758 | 0.26 | 0.126 | 2.7038E-16 |

| | | | | | | |
|---|---|---|---|---|---|---|
| Fibroblasts | RAP1B | 2.48084E-13 | 0.257735 | 0.694 | 0.535 | 4.07454E-09 |
| RERGL_Cells | CD164 | 0.000864477 | 0.257729 | 0.624 | 0.652 | 1 |
| Pericytes | PLK3 | 0.008212189 | 0.257716 | 0.24 | 0.203 | 1 |
| Fibroblasts | UBL3 | 3.87459E-20 | 0.257591 | 0.316 | 0.164 | 6.36363E-16 |
| Contractile | ZDHHC4 | 1.05022E-06 | 0.257414 | 0.259 | 0.173 | 0.017248753 |
| Intermediate | GLT8D2 | 1.18688E-10 | 0.257408 | 0.378 | 0.235 | 1.94934E-06 |
| Contractile | CTNNAL1 | 5.98558E-07 | 0.257405 | 0.257 | 0.168 | 0.009830709 |
| Contractile | SUB1 | 9.29244E-06 | 0.257388 | 0.695 | 0.664 | 0.152619083 |
| Intermediate | CTDSPL | 2.16475E-14 | 0.257338 | 0.153 | 0.059 | 3.55539E-10 |
| Contractile | SNF8 | 1.53286E-05 | 0.257335 | 0.427 | 0.344 | 0.251756392 |
| Pericytes | RAB11A | 0.008339043 | 0.257218 | 0.364 | 0.344 | 1 |
| Pericytes | ARHGAP10 | 0.000713121 | 0.257123 | 0.24 | 0.191 | 1 |
| Contractile | SNRNP35 | 1.40946E-06 | 0.256981 | 0.224 | 0.143 | 0.023149047 |
| Fibroblasts | FAM92A | 1.65591E-22 | 0.256768 | 0.255 | 0.117 | 2.71967E-18 |
| Fibroblasts | GANAB | 5.98598E-22 | 0.256722 | 0.284 | 0.137 | 9.83137E-18 |
| Terminal | CMPK1 | 0.000478247 | 0.256715 | 0.573 | 0.529 | 1 |
| Pericytes | AL021155. | 1.38899E-08 | 0.256474 | 0.105 | 0.046 | 0.000228128 |
| Contractile | FAM153CP | 2.2134E-10 | 0.256429 | 0.106 | 0.04 | 3.63528E-06 |
| Terminal | POLR2I | 0.002205448 | 0.256428 | 0.466 | 0.427 | 1 |
| Contractile | MYL12A | 2.40816E-10 | 0.256404 | 0.932 | 0.916 | 3.95517E-06 |
| Contractile | EEF1D | 2.07728E-11 | 0.256317 | 0.956 | 0.908 | 3.41173E-07 |
| Terminal | AOPEP | 0.000164669 | 0.256175 | 0.499 | 0.414 | 1 |
| Pericytes | YWHAG | 0.002159796 | 0.255987 | 0.227 | 0.184 | 1 |
| Intermediate | MXRA8 | 7.21796E-05 | 0.25594 | 0.563 | 0.466 | 1 |
| Pericytes | YRDC | 0.000243368 | 0.255913 | 0.127 | 0.083 | 1 |
| Intermediate | NME2 | 1.34375E-11 | 0.255905 | 0.925 | 0.869 | 2.20698E-07 |
| Contractile | PSMC1 | 4.1895E-05 | 0.255871 | 0.557 | 0.475 | 0.688084101 |
| Pericytes | SEPTIN9 | 0.000177582 | 0.255807 | 0.195 | 0.138 | 1 |
| Contractile | BCL2L1 | 1.46567E-06 | 0.25578 | 0.24 | 0.157 | 0.024072171 |
| Intermediate | PRKAR1B | 6.39109E-24 | 0.255728 | 0.146 | 0.037 | 1.04967E-19 |
| Terminal | SEC61B | 4.05138E-07 | 0.25569 | 0.782 | 0.715 | 0.00665398 |
| Contractile | SEC11C | 0.000152043 | 0.255586 | 0.289 | 0.219 | 1 |
| Terminal | C5orf46 | 1.78065E-05 | 0.255586 | 0.133 | 0.075 | 0.292453877 |
| Terminal | TMEM160 | 0.000803632 | 0.255564 | 0.316 | 0.257 | 1 |
| Fibroblasts | HLA-DRA | 0.000103786 | 0.255451 | 0.311 | 0.245 | 1 |
| Intermediate | ATP1B1 | 6.72003E-08 | 0.255433 | 0.25 | 0.152 | 0.001103698 |
| Contractile | MRPL11 | 1.70322E-05 | 0.255231 | 0.325 | 0.24 | 0.279737249 |
| Contractile | TBL1X | 2.78345E-12 | 0.255229 | 0.126 | 0.047 | 4.57154E-08 |
| Contractile | SQSTM1 | 2.75254E-08 | 0.255192 | 0.828 | 0.759 | 0.000452077 |
| Contractile | RHOB | 0.000118129 | 0.255185 | 0.633 | 0.543 | 1 |
| Contractile | CWC27 | 7.51245E-06 | 0.255175 | 0.182 | 0.115 | 0.123384448 |
| Contractile | RTCB | 5.88596E-05 | 0.255066 | 0.331 | 0.26 | 0.966709609 |
| Terminal | HAX1 | 0.002032493 | 0.255015 | 0.436 | 0.391 | 1 |
| Pericytes | PSMA5 | 0.008485302 | 0.255009 | 0.322 | 0.295 | 1 |
| Fibroblasts | FRMD6 | 2.88679E-25 | 0.25499 | 0.146 | 0.045 | 4.74127E-21 |
| Contractile | C1orf43 | 2.61492E-05 | 0.254984 | 0.519 | 0.437 | 0.42947476 |
| Pericytes | RBBP6 | 0.0067508 | 0.254957 | 0.205 | 0.168 | 1 |
| Intermediate | DOCK5 | 2.15566E-17 | 0.254752 | 0.146 | 0.048 | 3.54045E-13 |
| Terminal | PTGIR | 0.009831516 | 0.254716 | 0.322 | 0.277 | 1 |
| RERGL_Cells | SORBS1 | 5.46613E-06 | 0.254646 | 0.139 | 0.078 | 0.089775734 |
| Contractile | SERPINB6 | 3.28122E-05 | 0.254623 | 0.513 | 0.442 | 0.538907768 |
| Pericytes | SEPTIN8 | 0.006002606 | 0.254622 | 0.196 | 0.159 | 1 |
| Contractile | EMP2 | 4.67472E-06 | 0.254535 | 0.655 | 0.581 | 0.076777589 |
| Terminal | KDELR2 | 0.000231509 | 0.254478 | 0.667 | 0.628 | 1 |
| Terminal | COPRS | 0.002592856 | 0.254457 | 0.42 | 0.372 | 1 |
| Fibroblasts | TMEM71 | 5.49216E-41 | 0.254301 | 0.102 | 0.009 | 9.02033E-37 |

| | | | | | | |
|---|---|---|---|---|---|---|
| Terminal | DENND10 | 0.000174158 | 0.254284 | 0.179 | 0.121 | 1 |
| Pericytes | BYSL | 7.00789E-07 | 0.25425 | 0.105 | 0.052 | 0.011509765 |
| Pericytes | CPEB4 | 2.23496E-06 | 0.254207 | 0.133 | 0.075 | 0.036706902 |
| Intermediate | UBB | 1.42526E-13 | 0.254157 | 0.974 | 0.95 | 2.34084E-09 |
| Contractile | JADE1 | 0.000634548 | 0.254092 | 0.307 | 0.242 | 1 |
| Pericytes | TBL3 | 6.78924E-05 | 0.254036 | 0.149 | 0.097 | 1 |
| Pericytes | NUP35 | 0.000612815 | 0.253798 | 0.1 | 0.063 | 1 |
| Pericytes | CCT3 | 0.001185031 | 0.253761 | 0.487 | 0.462 | 1 |
| Pericytes | TUBB | 4.15386E-05 | 0.253585 | 0.809 | 0.823 | 0.682229937 |
| Fibroblasts | XAF1 | 2.43966E-26 | 0.253497 | 0.113 | 0.026 | 4.00689E-22 |
| Fibroblasts | CAMK2N1 | 6.93604E-25 | 0.253436 | 0.144 | 0.044 | 1.13917E-20 |
| Contractile | GNG10 | 1.66812E-07 | 0.253421 | 0.541 | 0.433 | 0.002739717 |
| Fibroblasts | TMED3 | 1.92347E-16 | 0.253411 | 0.511 | 0.338 | 3.15911E-12 |
| Contractile | NBDY | 1.73489E-05 | 0.253399 | 0.467 | 0.375 | 0.284938002 |
| Intermediate | KDELR1 | 1.49138E-07 | 0.253388 | 0.604 | 0.466 | 0.002449435 |
| RERGL_Cells | EHBP1L1 | 0.00058581 | 0.253326 | 0.107 | 0.066 | 1 |
| Terminal | FDX2 | 0.001261218 | 0.25332 | 0.172 | 0.125 | 1 |
| Fibroblasts | SRSF7 | 1.31233E-16 | 0.253254 | 0.723 | 0.533 | 2.15538E-12 |
| Contractile | SSNA1 | 1.2959E-06 | 0.253217 | 0.365 | 0.266 | 0.021283904 |
| Pericytes | BRIX1 | 0.005336074 | 0.253159 | 0.273 | 0.24 | 1 |
| RERGL_Cells | LINC01088 | 4.17388E-06 | 0.253128 | 0.115 | 0.06 | 0.068551842 |
| Pericytes | CASP3 | 8.065E-05 | 0.253028 | 0.107 | 0.063 | 1 |
| Pericytes | RUVBL1 | 0.000184034 | 0.252981 | 0.153 | 0.104 | 1 |
| Terminal | SNRPD2 | 0.000117152 | 0.252965 | 0.763 | 0.738 | 1 |
| Pericytes | PAFAH1B1 | 0.005412523 | 0.252937 | 0.262 | 0.226 | 1 |
| Intermediate | ENPP2 | 9.57973E-22 | 0.252858 | 0.103 | 0.022 | 1.57338E-17 |
| Contractile | ZC3H13 | 1.38926E-05 | 0.252788 | 0.357 | 0.271 | 0.228171516 |
| Contractile | TMEM120 | 0.000224654 | 0.252599 | 0.269 | 0.205 | 1 |
| Fibroblasts | LPCAT1 | 1.18407E-28 | 0.252595 | 0.152 | 0.043 | 1.94472E-24 |
| Terminal | RACK1 | 3.4853E-06 | 0.25255 | 0.972 | 0.971 | 0.057242599 |
| Intermediate | LIMS2 | 3.78487E-14 | 0.252536 | 0.32 | 0.164 | 6.21628E-10 |
| Intermediate | PRKG1 | 2.24646E-14 | 0.252489 | 0.421 | 0.236 | 3.68959E-10 |
| Fibroblasts | BMP4 | 9.02568E-18 | 0.252425 | 0.19 | 0.086 | 1.48238E-13 |
| Contractile | CISD3 | 3.05134E-05 | 0.252393 | 0.281 | 0.205 | 0.501152254 |
| Contractile | LRRFIP1 | 5.28494E-06 | 0.252384 | 0.413 | 0.318 | 0.086799869 |
| RERGL_Cells | THAP5 | 0.005211778 | 0.252264 | 0.165 | 0.127 | 1 |
| Fibroblasts | DNAJA1 | 5.84739E-16 | 0.252254 | 0.651 | 0.463 | 9.60375E-12 |
| Fibroblasts | SLC9B2 | 3.42324E-31 | 0.252089 | 0.131 | 0.029 | 5.62232E-27 |
| RERGL_Cells | DDT | 0.007631511 | 0.252064 | 0.434 | 0.434 | 1 |
| Pericytes | TRIP6 | 0.000158209 | 0.252052 | 0.433 | 0.384 | 1 |
| Fibroblasts | ECHDC1 | 7.30722E-19 | 0.252051 | 0.261 | 0.131 | 1.20014E-14 |
| Pericytes | PALM2-AK | 0.007143034 | 0.251975 | 0.175 | 0.14 | 1 |
| Terminal | CKS1B | 0.001846654 | 0.251914 | 0.325 | 0.269 | 1 |
| Pericytes | HNRNPK | 1.53779E-07 | 0.251907 | 0.851 | 0.852 | 0.002525669 |
| Contractile | PIN4 | 0.002013214 | 0.251708 | 0.367 | 0.315 | 1 |
| Pericytes | VPS37A | 0.001115317 | 0.251667 | 0.173 | 0.128 | 1 |
| RERGL_Cells | ATG4D | 0.00044367 | 0.251665 | 0.1 | 0.061 | 1 |
| Contractile | NDUFA2 | 7.04938E-06 | 0.251589 | 0.525 | 0.44 | 0.11577897 |
| Terminal | STUB1 | 0.000244485 | 0.251544 | 0.538 | 0.485 | 1 |
| Intermediate | PDLIM5 | 5.60845E-10 | 0.251525 | 0.51 | 0.338 | 9.21132E-06 |
| Fibroblasts | ATP8B4 | 1.12916E-25 | 0.251495 | 0.156 | 0.048 | 1.85452E-21 |
| Terminal | FSCN1 | 4.36805E-07 | 0.251436 | 0.109 | 0.051 | 0.007174086 |
| Pericytes | RMND1 | 5.52111E-06 | 0.251391 | 0.118 | 0.065 | 0.090678689 |
| Intermediate | CDK5RAP2 | 4.20842E-09 | 0.251356 | 0.159 | 0.079 | 6.9119E-05 |
| Fibroblasts | CYBC1 | 3.01825E-21 | 0.251343 | 0.255 | 0.119 | 4.95717E-17 |
| Fibroblasts | NQO1 | 8.33818E-16 | 0.251295 | 0.206 | 0.102 | 1.36946E-11 |

| | | | | | | |
|---|---|---|---|---|---|---|
| Fibroblasts | STXBP1 | 6.13087E-24 | 0.251284 | 0.183 | 0.068 | 1.00693E-19 |
| Pericytes | TCEAL1 | 3.65507E-05 | 0.251211 | 0.396 | 0.325 | 0.600308537 |
| Terminal | PLPP5 | 0.000770363 | 0.251186 | 0.205 | 0.149 | 1 |
| Pericytes | FAIM | 0.000297019 | 0.251024 | 0.151 | 0.104 | 1 |
| Contractile | XPA | 5.62175E-06 | 0.25094 | 0.259 | 0.178 | 0.092331634 |
| Contractile | RACK1 | 2.06525E-13 | 0.250939 | 0.982 | 0.969 | 3.39196E-09 |
| Contractile | SCOC | 6.76715E-05 | 0.250807 | 0.419 | 0.341 | 1 |
| Intermediate | BAD | 2.01004E-08 | 0.250723 | 0.486 | 0.346 | 0.000330129 |
| Terminal | CD82 | 0.003323134 | 0.250612 | 0.19 | 0.142 | 1 |
| Intermediate | CCDC85B | 1.15616E-07 | 0.25061 | 0.699 | 0.565 | 0.001898885 |
| Contractile | IDH3A | 0.00038201 | 0.250597 | 0.232 | 0.175 | 1 |
| Contractile | NDRG3 | 1.27344E-06 | 0.250573 | 0.184 | 0.11 | 0.020915012 |
| Terminal | C12orf10 | 0.000791893 | 0.250566 | 0.275 | 0.218 | 1 |
| Contractile | SNAPC3 | 3.06569E-05 | 0.250463 | 0.186 | 0.123 | 0.5035084 |
| Intermediate | ANXA2 | 9.27242E-13 | 0.250438 | 0.942 | 0.85 | 1.5229E-08 |
| Contractile | CRIM1-DT | 2.57702E-07 | 0.250353 | 0.118 | 0.058 | 0.004232505 |
| Fibroblasts | MILR1 | 3.10659E-36 | 0.250311 | 0.106 | 0.014 | 5.10226E-32 |
| Contractile | VPS28 | 1.31408E-07 | 0.250296 | 0.649 | 0.548 | 0.002158246 |
| Fibroblasts | SEC63 | 7.67973E-14 | 0.250244 | 0.578 | 0.405 | 1.26132E-09 |
| Fibroblasts | CTTN | 1.98132E-13 | 0.250176 | 0.553 | 0.388 | 3.25411E-09 |
| Intermediate | SESN3 | 2.03956E-14 | 0.250042 | 0.148 | 0.055 | 3.34977E-10 |

**Table S6. Marker genes for mesenchymal cell clusters in pig coronary plaque scRNA-seq data. Corresponds to Fig. 1E.**
The listed genes have significantly higher expression in the indicated cluster than all other clusters combined.
Analysis was performed using the Wilcoxon test in Seurat v4.4.0 with P values adjusted for multiple testing by the Bonferroni method.

| cluster | gene | p_val | avg_log2F( | pct.1 | pct.2 | p_val_adj |
|---|---|---|---|---|---|---|
| Fibroblasts | APOD | 9.84723E-17 | 4.749427 | 0.289 | 0.126 | 1.91775E-12 |
| VEGFA_Terminal | SCGB2A2 | 0 | 4.725957 | 0.482 | 0.021 | 0 |
| Pericytes | CXCL2 | 5.58036E-39 | 4.567386 | 0.68 | 0.125 | 1.08678E-34 |
| IFN_Intermediate | ISG15 | 4.8233E-158 | 4.449024 | 0.944 | 0.126 | 9.3933E-154 |
| Fibroblasts | CFD | 0 | 4.383033 | 0.752 | 0.033 | 0 |
| IFN_Intermediate | CXCL10 | 1.00998E-83 | 4.063385 | 0.159 | 0.004 | 1.96693E-79 |
| VEGFA_Terminal | IGFBP2 | 2.6416E-297 | 3.81891 | 0.857 | 0.172 | 5.1445E-293 |
| Contractile | TNNC1 | 0 | 3.769554 | 0.838 | 0.019 | 0 |
| Contractile | MYH11 | 0 | 3.756324 | 1 | 0.347 | 0 |
| Contractile | PPP1R14A | 0 | 3.746697 | 0.988 | 0.2 | 0 |
| Contractile | CNN1 | 0 | 3.640167 | 0.986 | 0.184 | 0 |
| Pericytes | RGS5 | 4.36576E-52 | 3.636207 | 0.84 | 0.136 | 8.50233E-48 |
| VEGFA_Terminal | IGFBP3 | 0 | 3.518879 | 0.37 | 0.013 | 0 |
| Fibroblasts | MFAP5 | 0 | 3.509947 | 0.647 | 0.03 | 0 |
| Contractile | ACTG2 | 0 | 3.459011 | 0.744 | 0.022 | 0 |
| Pericytes | CCL2 | 1.0894E-06 | 3.437804 | 0.48 | 0.262 | 0.02121598 |
| Pericytes | CCL19 | 2.8087E-24 | 3.350367 | 0.68 | 0.188 | 5.46995E-20 |
| Pericytes | ENSSSCG0( | 1.68213E-24 | 3.265759 | 0.6 | 0.144 | 3.27594E-20 |
| Pericytes | GJA4 | 0 | 3.250557 | 0.66 | 0.005 | 0 |
| Pericytes | RGS16 | 4.90691E-88 | 3.172734 | 0.58 | 0.036 | 9.5562E-84 |
| Contractile | MYL9 | 0 | 3.143199 | 0.998 | 0.576 | 0 |
| Contractile | CSRP1 | 0 | 3.085564 | 0.996 | 0.726 | 0 |
| Pericytes | ADAMTS4 | 4.69917E-70 | 3.057452 | 0.48 | 0.031 | 9.15164E-66 |
| Pericytes | ZFP36 | 2.93978E-18 | 3.018624 | 0.8 | 0.423 | 5.72522E-14 |
| Contractile | SMTN | 0 | 3.008522 | 0.968 | 0.266 | 0 |
| Contractile | TPM2 | 0 | 2.987528 | 0.998 | 0.819 | 0 |
| Fibroblasts | C3 | 1.2399E-224 | 2.959179 | 0.394 | 0.018 | 2.4147E-220 |
| Fibroblasts | PI16 | 7.6353E-143 | 2.944801 | 0.835 | 0.2 | 1.487E-138 |
| Contractile | ACTA2 | 0 | 2.94437 | 1 | 0.621 | 0 |
| Fibroblasts | FMO2 | 0 | 2.904087 | 0.523 | 0.017 | 0 |
| Pericytes | BTG2 | 5.53138E-16 | 2.797862 | 0.72 | 0.351 | 1.07724E-11 |
| Intermediate | IL1RL1 | 2.4073E-238 | 2.796494 | 0.637 | 0.139 | 4.6883E-234 |
| IFN_Intermediate | GBP1 | 1.914E-129 | 2.785771 | 0.523 | 0.038 | 3.7276E-125 |
| Fibroblasts | IGFBP5 | 2.04309E-37 | 2.762968 | 0.592 | 0.264 | 3.97892E-33 |
| Contractile | DES | 0 | 2.760461 | 0.52 | 0.017 | 0 |
| Contractile | LMOD1 | 0 | 2.721013 | 0.978 | 0.351 | 0 |
| Pericytes | ENSSSCG0( | 5.43866E-64 | 2.719094 | 0.52 | 0.04 | 1.05918E-59 |
| VEGFA_Terminal | VEGFA | 0 | 2.701279 | 0.814 | 0.132 | 0 |
| IFN_Intermediate | MX1 | 9.28639E-87 | 2.68124 | 0.869 | 0.207 | 1.80853E-82 |
| Pericytes | NFKBIA | 2.47055E-12 | 2.677076 | 0.72 | 0.45 | 4.8114E-08 |
| IFN_Intermediate | ENSSSCG0( | 3.50933E-73 | 2.672873 | 0.963 | 0.35 | 6.83442E-69 |
| Contractile | FLNA | 0 | 2.663736 | 0.998 | 0.749 | 0 |
| Fibroblasts | TXNIP | 6.29415E-90 | 2.654284 | 0.908 | 0.492 | 1.22579E-85 |
| VEGFA_Terminal | IGFBP5 | 2.5261E-179 | 2.648895 | 0.806 | 0.237 | 4.9196E-175 |
| VEGFA_Terminal | LOXL2 | 5.7842E-284 | 2.627632 | 0.906 | 0.223 | 1.1265E-279 |
| VEGFA_Terminal | ECM1 | 4.2666E-186 | 2.618229 | 0.978 | 0.64 | 8.3091E-182 |
| Contractile | RGS5 | 0 | 2.605438 | 0.618 | 0.05 | 0 |
| Pericytes | FABP4 | 5.5707E-40 | 2.591282 | 0.72 | 0.134 | 1.08489E-35 |
| Contractile | DSTN | 0 | 2.551904 | 0.996 | 0.829 | 0 |
| Contractile | MYLK | 0 | 2.549861 | 0.992 | 0.531 | 0 |
| Fibroblasts | MYOC | 0 | 2.544565 | 0.289 | 0.001 | 0 |
| Terminal | SAA3 | 3.62347E-45 | 2.531125 | 0.118 | 0.029 | 7.05671E-41 |
| Pericytes | FOSB | 5.78977E-18 | 2.530772 | 0.68 | 0.254 | 1.12756E-13 |
| Contractile | SORBS1 | 0 | 2.528008 | 0.891 | 0.216 | 0 |

| | | | | | | |
|---|---|---|---|---|---|---|
| Pericytes | SOCS3 | 2.09915E-16 | 2.509845 | 0.64 | 0.236 | 4.08809E-12 |
| Pericytes | JUNB | 5.45452E-14 | 2.507813 | 0.8 | 0.552 | 1.06227E-09 |
| IFN_Intermediate | IFIT1 | 9.6108E-177 | 2.503086 | 0.757 | 0.061 | 1.8717E-172 |
| Pericytes | C7 | 6.70995E-27 | 2.488914 | 0.74 | 0.195 | 1.30676E-22 |
| VEGFA_Terminal | PRSS23 | 2.9844E-203 | 2.473191 | 0.821 | 0.244 | 5.812E-199 |
| Pericytes | FOS | 6.43635E-08 | 2.459532 | 0.76 | 0.585 | 0.00125348 |
| Pericytes | ITIH4 | 1.3665E-19 | 2.449327 | 0.9 | 0.447 | 2.66125E-15 |
| VEGFA_Terminal | BNIP3 | 6.0888E-173 | 2.443855 | 0.942 | 0.473 | 1.1858E-168 |
| Pericytes | HIGD1B | 0 | 2.441256 | 0.56 | 0.006 | 0 |
| Contractile | ITGA8 | 0 | 2.422144 | 0.979 | 0.445 | 0 |
| Pericytes | TINAGL1 | 1.6684E-58 | 2.41603 | 0.68 | 0.081 | 3.24922E-54 |
| Pericytes | RASD1 | 4.13869E-43 | 2.402777 | 0.46 | 0.047 | 8.06009E-39 |
| Fibroblasts | SELENBP1 | 9.5085E-124 | 2.397334 | 0.807 | 0.213 | 1.8518E-119 |
| Fibroblasts | C7 | 4.27061E-33 | 2.367876 | 0.472 | 0.189 | 8.31702E-29 |
| Contractile | C6H19ORF | 0 | 2.365919 | 0.884 | 0.354 | 0 |
| IFN_Intermediate | MX2 | 1.8361E-284 | 2.361415 | 0.664 | 0.024 | 3.5759E-280 |
| Contractile | SYNPO2 | 0 | 2.348626 | 0.959 | 0.407 | 0 |
| Contractile | ITIH4 | 8.2555E-224 | 2.348423 | 0.788 | 0.385 | 1.6078E-219 |
| Pericytes | GADD45B | 1.54468E-13 | 2.340811 | 0.72 | 0.356 | 3.00826E-09 |
| Contractile | MYOZ2 | 0 | 2.335457 | 0.843 | 0.257 | 0 |
| Pericytes | NOTCH3 | 1.60831E-24 | 2.335379 | 0.66 | 0.178 | 3.13218E-20 |
| Terminal | COMP | 1.5099E-117 | 2.334128 | 0.177 | 0.019 | 2.9406E-113 |
| Fibroblasts | GPX3 | 2.8218E-92 | 2.329869 | 0.72 | 0.217 | 5.49546E-88 |
| Pericytes | MYC | 3.70655E-46 | 2.325971 | 0.56 | 0.066 | 7.21851E-42 |
| Fibroblasts | ABCA9 | 5.7336E-217 | 2.306669 | 0.619 | 0.059 | 1.1166E-212 |
| Pericytes | FAM162B | 0 | 2.30659 | 0.64 | 0.003 | 0 |
| Pericytes | LOC10052 | 8.91925E-55 | 2.292994 | 0.28 | 0.013 | 1.73702E-50 |
| Pericytes | EDNRB | 0 | 2.289823 | 0.52 | 0.001 | 0 |
| IFN_Intermediate | RSAD2 | 0 | 2.284355 | 0.486 | 0.008 | 0 |
| Fibroblasts | NID1 | 0 | 2.277897 | 0.573 | 0.022 | 0 |
| Fibroblasts | ENSSSCG0 | 4.488E-277 | 2.266913 | 0.509 | 0.025 | 8.7403E-273 |
| Fibroblasts | LAMA2 | 1.643E-131 | 2.264073 | 0.642 | 0.112 | 3.1998E-127 |
| IFN_Intermediate | LOC11025 | 4.6303E-57 | 2.251142 | 0.477 | 0.073 | 9.01751E-53 |
| Pericytes | ATF3 | 1.22034E-27 | 2.250731 | 0.66 | 0.149 | 2.37661E-23 |
| Pericytes | CYCS | 4.68504E-07 | 2.226359 | 0.6 | 0.386 | 0.009124112 |
| Pericytes | RRAD | 2.10736E-15 | 2.221681 | 0.34 | 0.068 | 4.10408E-11 |
| Fibroblasts | IGF2 | 1.5412E-112 | 2.196549 | 0.665 | 0.132 | 3.0014E-108 |
| IFN_Intermediate | ENSSSCG0 | 1.21468E-93 | 2.190701 | 0.748 | 0.124 | 2.36558E-89 |
| Contractile | SPARCL1 | 0 | 2.189287 | 0.994 | 0.651 | 0 |
| Pericytes | IL6 | 1.05505E-09 | 2.146648 | 0.18 | 0.031 | 2.0547E-05 |
| Pericytes | IER3 | 8.92082E-23 | 2.140056 | 0.56 | 0.129 | 1.73733E-18 |
| Fibroblasts | CRISPLD2 | 4.63164E-95 | 2.121879 | 0.716 | 0.214 | 9.02013E-91 |
| Fibroblasts | SLC6A4 | 0 | 2.109973 | 0.436 | 0.004 | 0 |
| Contractile | FBXO32 | 0 | 2.103539 | 0.874 | 0.336 | 0 |
| Pericytes | NR4A1 | 2.00781E-14 | 2.101561 | 0.6 | 0.218 | 3.91022E-10 |
| Contractile | ADIRF | 0 | 2.093203 | 0.986 | 0.648 | 0 |
| Fibroblasts | OGN | 2.42766E-95 | 2.062094 | 0.821 | 0.301 | 4.72787E-91 |
| IFN_Intermediate | ISG12(A) | 3.14325E-44 | 2.060837 | 0.972 | 0.652 | 6.12148E-40 |
| Pericytes | NR2F2 | 4.75141E-24 | 2.057888 | 0.8 | 0.29 | 9.25338E-20 |
| Pericytes | IFRD1 | 1.6103E-16 | 2.04448 | 0.58 | 0.19 | 3.13605E-12 |
| Pericytes | PPP1R15A | 7.09713E-16 | 2.043182 | 0.72 | 0.308 | 1.38217E-11 |
| Contractile | SORBS2 | 0 | 2.042765 | 0.816 | 0.194 | 0 |
| Contractile | TAGLN | 0 | 2.040148 | 1 | 0.956 | 0 |
| IFN_Intermediate | EIF2AK2 | 1.08102E-57 | 2.034917 | 0.841 | 0.273 | 2.1053E-53 |
| Contractile | PPP1R12B | 0 | 2.018169 | 0.656 | 0.047 | 0 |
| Fibroblasts | ABCA6 | 0 | 2.011674 | 0.5 | 0.003 | 0 |
| Contractile | PDLIM3 | 0 | 2.010221 | 0.817 | 0.215 | 0 |
| IFN_Intermediate | RIGI | 3.4582E-132 | 2.005972 | 0.617 | 0.054 | 6.7349E-128 |
| Pericytes | ADAMTS1 | 3.3464E-13 | 2.005734 | 0.58 | 0.206 | 6.51712E-09 |

| | | | | | | |
|---|---|---|---|---|---|---|
| Contractile | CALD1 | 0 | 1.999967 | 1 | 0.962 | 0 |
| Terminal | NDUFA4L2 | 0 | 1.995777 | 0.735 | 0.209 | 0 |
| Pericytes | IRF1 | 2.26384E-26 | 1.991705 | 0.48 | 0.083 | 4.40884E-22 |
| Pericytes | JUN | 7.06302E-11 | 1.987952 | 0.84 | 0.739 | 1.37552E-06 |
| Contractile | PAWR | 0 | 1.983551 | 0.97 | 0.546 | 0 |
| Fibroblasts | ENSSSCG0( | 1.4944E-146 | 1.978134 | 0.656 | 0.104 | 2.9103E-142 |
| Pericytes | ITGA7 | 6.92625E-32 | 1.956848 | 0.74 | 0.168 | 1.34889E-27 |
| Contractile | PGM5 | 0 | 1.952684 | 0.692 | 0.059 | 0 |
| Fibroblasts | MARCKS | 1.6856E-129 | 1.949267 | 0.615 | 0.105 | 3.2827E-125 |
| Terminal | ANXA8 | 0 | 1.946732 | 0.842 | 0.279 | 0 |
| Pericytes | ITIH5 | 5.28154E-27 | 1.934633 | 0.68 | 0.168 | 1.02858E-22 |
| Pericytes | CEBPB | 2.2507E-14 | 1.931736 | 0.8 | 0.501 | 4.38323E-10 |
| VEGFA_Terminal | CA4 | 2.6776E-195 | 1.927598 | 0.799 | 0.202 | 5.2147E-191 |
| Contractile | CPE | 0 | 1.926429 | 0.991 | 0.471 | 0 |
| Fibroblasts | CLEC3B | 9.35126E-43 | 1.914379 | 0.514 | 0.186 | 1.82116E-38 |
| CD74_Intermediat | SLA-DMA | 2.20874E-71 | 0.328573 | 0.112 | 0.012 | 4.30152E-67 |
| Fibroblasts | GPC3 | 1.468E-205 | 1.90664 | 0.417 | 0.023 | 2.8589E-201 |
| Contractile | MYL6 | 0 | 1.902123 | 0.997 | 0.965 | 0 |
| Intermediate | CCN3 | 1.29E-164 | 1.901006 | 0.962 | 0.658 | 2.5124E-160 |
| Fibroblasts | DCN | 3.42513E-96 | 1.895882 | 1 | 0.874 | 6.67045E-92 |
| Pericytes | EGR1 | 5.49385E-09 | 1.889503 | 0.7 | 0.468 | 0.000106993 |
| VEGFA_Terminal | EGLN3 | 2.4515E-184 | 1.878608 | 0.77 | 0.212 | 4.7742E-180 |
| IFN_Intermediate | OAS2 | 1.5295E-108 | 1.878471 | 0.617 | 0.066 | 2.9787E-104 |
| VEGFA_Terminal | NMB | 2.6702E-156 | 1.87674 | 0.55 | 0.102 | 5.2002E-152 |
| Pericytes | GUCY1B1 | 7.97981E-25 | 1.868702 | 0.68 | 0.191 | 1.55407E-20 |
| Contractile | KCNMB1 | 0 | 1.868186 | 0.667 | 0.016 | 0 |
| IFN_Intermediate | VCAM1 | 3.93557E-18 | 1.867775 | 0.645 | 0.293 | 7.66452E-14 |
| Pericytes | PKIG | 9.49738E-15 | 1.855213 | 0.62 | 0.255 | 1.84962E-10 |
| Pericytes | SEPTIN4 | 2.17525E-75 | 1.855012 | 0.64 | 0.053 | 4.2363E-71 |
| Contractile | CKM | 0 | 1.853987 | 0.402 | 0 | 0 |
| VEGFA_Terminal | SERPINE1 | 1.64295E-95 | 1.839793 | 0.901 | 0.598 | 3.19965E-91 |
| Contractile | RASSF3 | 0 | 1.837036 | 0.793 | 0.216 | 0 |
| Pericytes | ZFP36L1 | 2.36559E-10 | 1.835655 | 0.86 | 0.677 | 4.607E-06 |
| Contractile | PLN | 0 | 1.834225 | 0.675 | 0.076 | 0 |
| Contractile | MCAM | 0 | 1.826303 | 0.662 | 0.046 | 0 |
| IFN_Intermediate | EPSTI1 | 3.76046E-88 | 1.82392 | 0.551 | 0.066 | 7.32349E-84 |
| Contractile | CAMK2G | 0 | 1.819407 | 0.777 | 0.202 | 0 |
| Contractile | PDLIM7 | 0 | 1.810289 | 0.916 | 0.427 | 0 |
| Contractile | IGF2 | 1.0996E-256 | 1.804231 | 0.505 | 0.082 | 2.1414E-252 |
| Pericytes | ID4 | 3.36726E-14 | 1.803076 | 0.76 | 0.4 | 6.55773E-10 |
| Terminal | S100A4 | 0 | 1.803018 | 0.97 | 0.531 | 0 |
| Fibroblasts | SPTBN1 | 1.68502E-81 | 1.79379 | 0.968 | 0.764 | 3.28158E-77 |
| Pericytes | FRMD3 | 1.4913E-154 | 1.792741 | 0.44 | 0.01 | 2.9043E-150 |
| IFN_Intermediate | ENSSSCG0( | 1.24105E-35 | 1.789861 | 0.888 | 0.519 | 2.41695E-31 |
| Contractile | VCL | 0 | 1.785515 | 0.985 | 0.664 | 0 |
| Pericytes | ISYNA1 | 7.31669E-44 | 1.783827 | 0.52 | 0.06 | 1.42492E-39 |
| IFN_Intermediate | GHSR | 4.08679E-26 | 1.773282 | 0.542 | 0.17 | 7.95903E-22 |
| VEGFA_Terminal | DDIT4 | 2.20831E-45 | 1.766533 | 0.625 | 0.338 | 4.30069E-41 |
| Pericytes | GUCY1A1 | 1.09676E-32 | 1.762887 | 0.72 | 0.161 | 2.13594E-28 |
| Pericytes | KLHL23 | 5.07888E-35 | 1.762477 | 0.46 | 0.059 | 9.89112E-31 |
| Pericytes | COL4A1 | 8.63458E-11 | 1.755361 | 0.88 | 0.692 | 1.68158E-06 |
| IFN_Intermediate | STAT1 | 9.4309E-56 | 1.75479 | 0.813 | 0.242 | 1.83667E-51 |
| Pericytes | GJC1 | 6.80722E-20 | 1.753845 | 0.62 | 0.174 | 1.32571E-15 |
| Contractile | SLC22A3 | 0 | 1.747311 | 0.575 | 0.016 | 0 |
| Pericytes | KLF2 | 3.59165E-09 | 1.746671 | 0.68 | 0.374 | 6.99474E-05 |
| Intermediate | IFI6 | 3.1674E-127 | 1.745076 | 0.939 | 0.672 | 6.1684E-123 |
| Contractile | TNS1 | 0 | 1.741473 | 0.804 | 0.269 | 0 |
| Fibroblasts | THBS4 | 6.89572E-52 | 1.738372 | 0.138 | 0.011 | 1.34294E-47 |
| VEGFA_Terminal | ERO1A | 1.4164E-101 | 1.734504 | 0.758 | 0.327 | 2.75838E-97 |

| | | | | | | |
|---|---|---|---|---|---|---|
| Fibroblasts | SOCS3 | 4.66245E-20 | 1.732398 | 0.445 | 0.232 | 9.08013E-16 |
| Pericytes | LOC10012 | 2.01174E-44 | 1.730276 | 0.52 | 0.058 | 3.91786E-40 |
| Pericytes | SLC2A3 | 4.19929E-11 | 1.724474 | 0.6 | 0.279 | 8.17812E-07 |
| CD74_Intermediat | ENSSSCG0 | 1.32897E-58 | 0.30577 | 0.107 | 0.016 | 2.58817E-54 |
| Pericytes | PEAR1 | 1.10178E-30 | 1.711916 | 0.54 | 0.091 | 2.14571E-26 |
| Terminal | IGFBP6 | 1.3902E-286 | 1.711309 | 0.859 | 0.502 | 2.7075E-282 |
| IFN_Intermediate | IFI6 | 3.1879E-28 | 1.708463 | 0.972 | 0.695 | 6.20843E-24 |
| Fibroblasts | CXCL12 | 1.14875E-64 | 1.706242 | 0.619 | 0.192 | 2.23719E-60 |
| Fibroblasts | GHSR | 2.61358E-39 | 1.701332 | 0.468 | 0.166 | 5.08995E-35 |
| VEGFA_Terminal | GAPDH | 1.8547E-160 | 1.699724 | 0.998 | 0.928 | 3.612E-156 |
| IFN_Intermediate | PARP14 | 5.513E-135 | 1.699007 | 0.57 | 0.043 | 1.0736E-130 |
| Contractile | LOC10051 | 1.2477E-248 | 1.697518 | 0.684 | 0.207 | 2.4298E-244 |
| Pericytes | IL13RA1 | 2.15469E-20 | 1.697158 | 0.8 | 0.33 | 4.19626E-16 |
| Fibroblasts | HSD11B1 | 9.49936E-66 | 1.691206 | 0.844 | 0.432 | 1.85E-61 |
| IFN_Intermediate | PLAC8 | 9.10049E-97 | 1.688145 | 0.636 | 0.079 | 1.77232E-92 |
| Contractile | KANK1 | 0 | 1.679578 | 0.825 | 0.326 | 0 |
| Contractile | NIBAN1 | 0 | 1.677527 | 0.973 | 0.732 | 0 |
| Fibroblasts | HTRA3 | 5.1848E-177 | 1.667536 | 0.372 | 0.022 | 1.0097E-172 |
| Contractile | EFHD1 | 0 | 1.662543 | 0.587 | 0.037 | 0 |
| Pericytes | ARPC1A | 6.29361E-21 | 1.649125 | 0.74 | 0.26 | 1.22568E-16 |
| Pericytes | TXNIP | 2.79265E-07 | 1.648187 | 0.74 | 0.504 | 0.005438683 |
| Pericytes | GADD45A | 2.43367E-08 | 1.647815 | 0.6 | 0.326 | 0.000473956 |
| Contractile | RBPMS | 0 | 1.644322 | 0.819 | 0.294 | 0 |
| Contractile | FBLIM1 | 0 | 1.64387 | 0.875 | 0.377 | 0 |
| VEGFA_Terminal | NDUFA4L2 | 1.734E-101 | 1.643028 | 0.818 | 0.347 | 3.37698E-97 |
| VEGFA_Terminal | SLC16A3 | 7.4519E-227 | 1.641434 | 0.678 | 0.115 | 1.4513E-222 |
| Fibroblasts | ANGPTL4 | 6.5897E-59 | 1.627005 | 0.505 | 0.138 | 1.28334E-54 |
| Pericytes | ANGPTL4 | 0.00189412 | 1.613655 | 0.28 | 0.149 | 1 |
| Contractile | PPP1R12A | 0 | 1.612815 | 0.954 | 0.632 | 0 |
| Pericytes | RND3 | 1.36204E-12 | 1.612737 | 0.62 | 0.254 | 2.65258E-08 |
| Fibroblasts | CHRDL1 | 0 | 1.607954 | 0.422 | 0.009 | 0 |
| VEGFA_Terminal | PTGES | 2.5842E-112 | 1.600905 | 0.831 | 0.396 | 5.0327E-108 |
| Fibroblasts | PCOLCE2 | 1.33965E-51 | 1.599949 | 0.642 | 0.253 | 2.60898E-47 |
| Pericytes | NEURL1B | 4.7894E-244 | 1.597372 | 0.48 | 0.006 | 9.3273E-240 |
| Terminal | FNDC1 | 0 | 1.59716 | 0.803 | 0.322 | 0 |
| Contractile | NEXN | 1.2698E-288 | 1.596603 | 0.951 | 0.611 | 2.4728E-284 |
| Pericytes | UTRN | 8.18842E-14 | 1.591789 | 0.72 | 0.38 | 1.59469E-09 |
| VEGFA_Terminal | ADM | 1.29543E-62 | 1.590692 | 0.385 | 0.11 | 2.52285E-58 |
| Pericytes | MYO1B | 5.9138E-16 | 1.586709 | 0.46 | 0.117 | 1.15171E-11 |
| Contractile | LMCD1 | 3.7229E-264 | 1.586006 | 0.725 | 0.231 | 7.2504E-260 |
| Pericytes | ITIH3 | 4.9416E-133 | 1.581992 | 0.3 | 0.005 | 9.6237E-129 |
| VEGFA_Terminal | PLAT | 1.00638E-30 | 1.578976 | 0.521 | 0.281 | 1.95992E-26 |
| Pericytes | COL5A3 | 7.39829E-08 | 1.577611 | 0.44 | 0.189 | 0.001440817 |
| CD74_Intermediat | GPNMB | 3.7646E-298 | 1.57367 | 0.77 | 0.33 | 7.3315E-294 |
| Fibroblasts | FMO1 | 2.9686E-99 | 1.57203 | 0.33 | 0.034 | 5.78125E-95 |
| Pericytes | CYGB | 1.63119E-53 | 1.571195 | 0.32 | 0.017 | 3.17675E-49 |
| Contractile | TPM1 | 0 | 1.56712 | 0.994 | 0.789 | 0 |
| Pericytes | RASL11A | 5.6998E-11 | 1.564539 | 0.6 | 0.273 | 1.11004E-06 |
| VEGFA_Terminal | SPON2 | 9.28027E-82 | 1.555492 | 0.596 | 0.209 | 1.80733E-77 |
| Pericytes | COL18A1 | 1.7324E-14 | 1.552713 | 0.62 | 0.225 | 3.37385E-10 |
| Contractile | RASL12 | 0 | 1.54667 | 0.562 | 0.04 | 0 |
| Fibroblasts | CXCL14 | 1.67977E-21 | 1.545565 | 0.106 | 0.016 | 3.27136E-17 |
| Pericytes | ICAM1 | 1.67874E-08 | 1.537291 | 0.44 | 0.178 | 0.000326935 |
| Contractile | CALML4 | 5.777E-210 | 1.535901 | 0.506 | 0.122 | 1.1251E-205 |
| Pericytes | CCNL1 | 9.21746E-10 | 1.533556 | 0.54 | 0.246 | 1.7951E-05 |
| VEGFA_Terminal | TSC22D1 | 2.18889E-89 | 1.52962 | 0.714 | 0.304 | 4.26286E-85 |
| Pericytes | IER2 | 1.00353E-06 | 1.529073 | 0.52 | 0.294 | 0.019543747 |
| Pericytes | TCF21 | 5.72063E-47 | 1.527845 | 0.36 | 0.026 | 1.11409E-42 |
| Pericytes | IL17B | 1.08798E-13 | 1.522339 | 0.58 | 0.213 | 2.11885E-09 |

| | | | | | | |
|---|---|---|---|---|---|---|
| VEGFA_Terminal | ENO1 | 1.0065E-152 | 1.519589 | 1 | 0.916 | 1.9601E-148 |
| Contractile | RHOB | 3.3797E-227 | 1.518932 | 0.846 | 0.438 | 6.5819E-223 |
| Contractile | SVIL | 6.6285E-244 | 1.515684 | 0.812 | 0.372 | 1.2909E-239 |
| IFN_Intermediate | HERC6 | 6.59788E-60 | 1.514893 | 0.645 | 0.131 | 1.28494E-55 |
| Pericytes | IER5 | 1.86341E-14 | 1.512477 | 0.48 | 0.137 | 3.629E-10 |
| Pericytes | LOC10052 | 9.71855E-18 | 1.509556 | 0.96 | 0.927 | 1.89269E-13 |
| VEGFA_Terminal | NDRG1 | 6.93593E-95 | 1.508786 | 0.741 | 0.319 | 1.35077E-90 |
| Fibroblasts | PLSCR4 | 2.48053E-60 | 1.508099 | 0.789 | 0.382 | 4.83082E-56 |
| Contractile | MGLL | 8.0221E-253 | 1.505058 | 0.819 | 0.372 | 1.5623E-248 |
| CD74_Intermediat | ENSSSCG0( | 1.9251E-220 | 1.503722 | 0.884 | 0.641 | 3.7491E-216 |
| Contractile | ADAMTS1 | 3.8906E-109 | 1.50032 | 0.456 | 0.161 | 7.5769E-105 |
| Fibroblasts | APOE | 3.63958E-10 | 1.498931 | 0.495 | 0.336 | 7.08809E-06 |
| VEGFA_Terminal | LDHA | 2.7508E-146 | 1.493083 | 0.985 | 0.685 | 5.3572E-142 |
| IFN_Intermediate | UBE2L6 | 2.16497E-56 | 1.491646 | 0.626 | 0.134 | 4.21628E-52 |
| Pericytes | ABCC9 | 8.60988E-10 | 1.491614 | 0.24 | 0.052 | 1.67677E-05 |
| Pericytes | ANO3 | 1.0669E-153 | 1.490836 | 0.34 | 0.005 | 2.0778E-149 |
| Pericytes | NRP1 | 6.68159E-10 | 1.489558 | 0.68 | 0.371 | 1.30124E-05 |
| Fibroblasts | FOS | 9.10499E-09 | 1.48585 | 0.651 | 0.584 | 0.00017732 |
| Contractile | S100A2 | 1.9649E-180 | 1.474736 | 0.511 | 0.136 | 3.8266E-176 |
| Fibroblasts | ZFP36 | 1.64003E-08 | 1.472077 | 0.528 | 0.422 | 0.000319396 |
| Contractile | LPP | 2.219E-296 | 1.468455 | 0.79 | 0.28 | 4.3215E-292 |
| Pericytes | ENSSSCG0( | 7.61949E-08 | 1.458954 | 0.56 | 0.293 | 0.001483895 |
| Fibroblasts | ADGRD1 | 7.25848E-45 | 1.456482 | 0.624 | 0.27 | 1.41359E-40 |
| IFN_Intermediate | LGALS3BP | 4.39418E-35 | 1.453268 | 0.813 | 0.322 | 8.55766E-31 |
| Fibroblasts | FBLN2 | 3.35017E-37 | 1.447021 | 0.711 | 0.423 | 6.52445E-33 |
| Fibroblasts | FBLN1 | 1.63783E-43 | 1.437986 | 0.894 | 0.688 | 3.18967E-39 |
| Pericytes | ENSSSCG0( | 1.05085E-06 | 1.43779 | 0.4 | 0.177 | 0.020465239 |
| Contractile | NPNT | 3.1336E-235 | 1.427866 | 0.683 | 0.216 | 6.1027E-231 |
| Fibroblasts | OMD | 1.66543E-40 | 1.421346 | 0.665 | 0.322 | 3.24343E-36 |
| Terminal | POSTN | 4.8677E-257 | 1.418707 | 0.942 | 0.634 | 9.4799E-253 |
| Fibroblasts | LTBP4 | 6.7621E-62 | 1.417592 | 0.95 | 0.731 | 1.31692E-57 |
| Intermediate | EFEMP1 | 1.589E-122 | 1.409676 | 0.906 | 0.639 | 3.0945E-118 |
| CD74_Intermediat | A2M | 1.1971E-109 | 1.409058 | 0.738 | 0.542 | 2.3313E-105 |
| VEGFA_Terminal | PHLDA2 | 7.61923E-74 | 1.407888 | 0.838 | 0.491 | 1.48385E-69 |
| VEGFA_Terminal | CCL2 | 4.34214E-28 | 1.406188 | 0.475 | 0.249 | 8.45633E-24 |
| Pericytes | EBF1 | 8.14963E-15 | 1.401375 | 0.68 | 0.271 | 1.58714E-10 |
| Pericytes | KCNA5 | 6.04761E-13 | 1.39669 | 0.32 | 0.07 | 1.17777E-08 |
| VEGFA_Terminal | P4HA1 | 5.6408E-111 | 1.386984 | 0.893 | 0.482 | 1.0986E-106 |
| Pericytes | SRSF7 | 1.1734E-07 | 1.385684 | 0.7 | 0.475 | 0.002285191 |
| VEGFA_Terminal | CSRP2 | 1.96211E-70 | 1.382336 | 0.77 | 0.417 | 3.82121E-66 |
| Contractile | KCNAB1 | 2.1763E-272 | 1.378976 | 0.667 | 0.175 | 4.2384E-268 |
| Fibroblasts | DPT | 7.71697E-40 | 1.377662 | 0.688 | 0.374 | 1.50288E-35 |
| Fibroblasts | ITGBL1 | 1.6758E-105 | 1.37691 | 0.252 | 0.017 | 3.2637E-101 |
| Fibroblasts | RNASE4 | 4.46537E-62 | 1.376246 | 0.927 | 0.763 | 8.69632E-58 |
| Contractile | FERMT2 | 1.3998E-290 | 1.375932 | 0.953 | 0.688 | 2.726E-286 |
| Fibroblasts | PDGFRA | 2.04146E-54 | 1.375825 | 0.683 | 0.303 | 3.97573E-50 |
| Pericytes | CPM | 6.97751E-23 | 1.374966 | 0.12 | 0.006 | 1.35887E-18 |
| Pericytes | OLFML2A | 2.84742E-15 | 1.374687 | 0.32 | 0.06 | 5.54535E-11 |
| Fibroblasts | MATN2 | 5.50438E-56 | 1.374622 | 0.344 | 0.068 | 1.07198E-51 |
| Terminal | COL8A1 | 1.9146E-237 | 1.368527 | 0.644 | 0.245 | 3.7287E-233 |
| Pericytes | FHL5 | 0 | 1.367439 | 0.34 | 0 | 0 |
| Fibroblasts | F3 | 1.87086E-55 | 1.363999 | 0.252 | 0.035 | 3.6435E-51 |
| IFN_Intermediate | LOC10051! | 2.54893E-35 | 1.363837 | 0.533 | 0.139 | 4.96405E-31 |
| VEGFA_Terminal | POSTN | 3.31544E-80 | 1.363804 | 0.952 | 0.717 | 6.45683E-76 |
| Fibroblasts | ADD3 | 6.25075E-42 | 1.361896 | 0.729 | 0.416 | 1.21733E-37 |
| Contractile | FLNC | 0 | 1.361895 | 0.476 | 0.037 | 0 |
| Fibroblasts | PTN | 6.91701E-96 | 1.358629 | 0.229 | 0.015 | 1.34709E-91 |
| Pericytes | ENSSSCG0( | 9.3187E-157 | 1.357329 | 0.16 | 0 | 1.8148E-152 |
| Contractile | PDZRN4 | 0 | 1.357275 | 0.452 | 0.016 | 0 |

| | | | | | | |
|---|---|---|---|---|---|---|
| Pericytes | CD4 | 1.2179E-266 | 1.356424 | 0.4 | 0.003 | 2.3719E-262 |
| Terminal | SERPINE2 | 2.5927E-191 | 1.348655 | 0.803 | 0.464 | 5.0492E-187 |
| Terminal | COL3A1 | 8.0894E-167 | 1.342821 | 0.976 | 0.933 | 1.5754E-162 |
| Fibroblasts | SVEP1 | 9.37909E-70 | 1.339956 | 0.317 | 0.046 | 1.82658E-65 |
| Fibroblasts | CD302 | 3.38179E-43 | 1.339308 | 0.775 | 0.476 | 6.58604E-39 |
| IFN_Intermediate | ENSSSCG0( | 5.58051E-77 | 1.334223 | 0.514 | 0.062 | 1.0868E-72 |
| Contractile | NET1 | 2.5644E-196 | 1.334135 | 0.716 | 0.283 | 4.9942E-192 |
| IFN_Intermediate | ZNFX1 | 6.76392E-42 | 1.33289 | 0.617 | 0.155 | 1.31727E-37 |
| Intermediate | MYH10 | 1.20828E-82 | 1.33129 | 0.829 | 0.585 | 2.35313E-78 |
| Pericytes | ARL5B | 5.50628E-10 | 1.330219 | 0.4 | 0.13 | 1.07235E-05 |
| Pericytes | RCAN2 | 2.52149E-09 | 1.329621 | 0.52 | 0.209 | 4.91061E-05 |
| IFN_Intermediate | IFITM1 | 1.061E-53 | 1.321048 | 0.458 | 0.069 | 2.0663E-49 |
| Fibroblasts | CYBRD1 | 7.6914E-40 | 1.319032 | 0.78 | 0.522 | 1.4979E-35 |
| VEGFA_Terminal | UPP1 | 1.4672E-247 | 1.318684 | 0.433 | 0.033 | 2.8573E-243 |
| Contractile | TGM2 | 1.106E-167 | 1.315778 | 0.839 | 0.476 | 2.154E-163 |
| Pericytes | ENSSSCG0( | 8.3545E-126 | 1.313365 | 0.32 | 0.006 | 1.627E-121 |
| Pericytes | EPAS1 | 1.78964E-09 | 1.312605 | 0.8 | 0.572 | 3.48531E-05 |
| Pericytes | RHOB | 8.01117E-06 | 1.312227 | 0.68 | 0.502 | 0.156017592 |
| Pericytes | STUM | 1.06835E-92 | 1.312067 | 0.38 | 0.014 | 2.08062E-88 |
| Contractile | TGFB1I1 | 4.2261E-210 | 1.305309 | 0.736 | 0.319 | 8.2303E-206 |
| Terminal | PRELP | 1.1701E-167 | 1.304893 | 0.898 | 0.794 | 2.2788E-163 |
| Contractile | SPINT2 | 1.7573E-210 | 1.303549 | 0.802 | 0.374 | 3.4224E-206 |
| Pericytes | SEPTIN11 | 2.15104E-12 | 1.302915 | 0.78 | 0.466 | 4.18915E-08 |
| Contractile | IRAG1 | 3.7151E-233 | 1.302862 | 0.679 | 0.217 | 7.2351E-229 |
| Pericytes | EPS8 | 1.44943E-11 | 1.300888 | 0.56 | 0.239 | 2.82276E-07 |
| Intermediate | A2M | 2.5531E-115 | 1.300821 | 0.877 | 0.564 | 4.9722E-111 |
| Contractile | FHL1 | 8.7033E-242 | 1.300083 | 0.959 | 0.718 | 1.695E-237 |
| Contractile | RAMP1 | 1.3228E-266 | 1.297609 | 0.538 | 0.096 | 2.5761E-262 |
| Contractile | TLN1 | 2.7226E-230 | 1.297319 | 0.899 | 0.564 | 5.3022E-226 |
| CD74_Intermediat | FGL2 | 1.2431E-304 | 1.294703 | 0.956 | 0.599 | 2.4209E-300 |
| IFN_Intermediate | HERC5 | 1.156E-124 | 1.292851 | 0.439 | 0.026 | 2.2514E-120 |
| IFN_Intermediate | PSMB9 | 7.05896E-34 | 1.285812 | 0.364 | 0.068 | 1.37473E-29 |
| Contractile | ENSSSCG0( | 7.6366E-287 | 1.28434 | 0.54 | 0.095 | 1.4872E-282 |
| Pericytes | EDNRA | 1.26064E-18 | 1.284168 | 0.38 | 0.071 | 2.4551E-14 |
| Contractile | CAP1 | 3.6236E-217 | 1.280409 | 0.855 | 0.512 | 7.0571E-213 |
| Contractile | GYG1 | 3.985E-176 | 1.279432 | 0.768 | 0.425 | 7.7608E-172 |
| Fibroblasts | JUNB | 1.19758E-08 | 1.275351 | 0.619 | 0.551 | 0.000233229 |
| Pericytes | TRA2B | 1.66317E-05 | 1.274039 | 0.66 | 0.539 | 0.323902006 |
| Fibroblasts | GFPT2 | 6.955E-131 | 1.271592 | 0.298 | 0.019 | 1.3545E-126 |
| Pericytes | P3H2 | 2.89374E-41 | 1.271263 | 0.36 | 0.029 | 5.63555E-37 |
| CD74_Intermediat | ENSSSCG0( | 1.3718E-218 | 1.269637 | 0.934 | 0.756 | 2.6715E-214 |
| Pericytes | FMO2 | 1.80341E-24 | 1.267812 | 0.3 | 0.033 | 3.51213E-20 |
| Intermediate | FRZB | 1.0918E-124 | 1.267461 | 0.57 | 0.19 | 2.1263E-120 |
| VEGFA_Terminal | TPI1 | 2.0685E-106 | 1.266481 | 0.942 | 0.73 | 4.0283E-102 |
| Pericytes | APOD | 1.96833E-23 | 1.260991 | 0.58 | 0.128 | 3.83332E-19 |
| Pericytes | MCAM | 3.30193E-16 | 1.259808 | 0.54 | 0.142 | 6.4305E-12 |
| Fibroblasts | EGR1 | 6.60293E-16 | 1.259769 | 0.651 | 0.463 | 1.28592E-11 |
| Pericytes | SMARCA5 | 7.62888E-11 | 1.259723 | 0.78 | 0.495 | 1.48572E-06 |
| IFN_Intermediate | PSMB8 | 1.46609E-19 | 1.252833 | 0.505 | 0.193 | 2.85521E-15 |
| Contractile | RCAN2 | 2.4764E-185 | 1.252601 | 0.542 | 0.148 | 4.8227E-181 |
| Contractile | HSPB7 | 7.9708E-230 | 1.251539 | 0.541 | 0.119 | 1.5523E-225 |
| Fibroblasts | ENSSSCG0( | 4.6076E-66 | 1.25093 | 0.312 | 0.047 | 8.9733E-62 |
| Pericytes | PTGER4 | 5.0664E-39 | 1.250432 | 0.32 | 0.024 | 9.86681E-35 |
| VEGFA_Terminal | IER3 | 8.54727E-70 | 1.247909 | 0.409 | 0.113 | 1.66458E-65 |
| Intermediate | ACTA2 | 7.39727E-89 | 1.247671 | 0.911 | 0.656 | 1.44062E-84 |
| Intermediate | ITIH4 | 1.65457E-75 | 1.244906 | 0.725 | 0.418 | 3.22227E-71 |
| Pericytes | GTF2E2 | 6.5907E-09 | 1.241319 | 0.7 | 0.453 | 0.000128354 |
| Contractile | SLC25A4 | 1.0963E-213 | 1.240567 | 0.693 | 0.263 | 2.1351E-209 |
| Pericytes | PDGFA | 3.45042E-11 | 1.238805 | 0.54 | 0.214 | 6.7197E-07 |

| | | | | | | |
|---|---|---|---|---|---|---|
| Contractile | SH3BGR | 1.2593E-202 | 1.236238 | 0.652 | 0.236 | 2.4524E-198 |
| Pericytes | LBH | 2.97802E-09 | 1.235482 | 0.38 | 0.128 | 5.7997E-05 |
| Fibroblasts | BTG2 | 2.97473E-15 | 1.233043 | 0.537 | 0.347 | 5.79328E-11 |
| Pericytes | STOM | 2.21445E-13 | 1.230882 | 0.86 | 0.524 | 4.31264E-09 |
| Contractile | ITGA5 | 4.7526E-134 | 1.217882 | 0.639 | 0.296 | 9.2557E-130 |
| Fibroblasts | TGFBR3 | 1.50036E-36 | 1.215374 | 0.564 | 0.252 | 2.92196E-32 |
| Fibroblasts | ACKR4 | 1.02434E-41 | 1.214898 | 0.211 | 0.033 | 1.99489E-37 |
| Pericytes | CTSC | 0.001956776 | 1.210729 | 0.48 | 0.333 | 1 |
| Fibroblasts | C4B | 4.49629E-06 | 1.209592 | 0.353 | 0.255 | 0.087565313 |
| Fibroblasts | ALDH1A1 | 1.15366E-44 | 1.203311 | 0.486 | 0.158 | 2.24674E-40 |
| Pericytes | EBF2 | 5.72031E-46 | 1.198903 | 0.26 | 0.013 | 1.11403E-41 |
| Pericytes | SLC12A2 | 0.004464807 | 1.196962 | 0.18 | 0.081 | 1 |
| VEGFA_Terminal | COL6A3 | 9.05973E-78 | 1.196622 | 0.961 | 0.714 | 1.76438E-73 |
| Pericytes | ENPEP | 1.4422E-167 | 1.1947 | 0.24 | 0.002 | 2.8087E-163 |
| Fibroblasts | C1S | 3.89712E-33 | 1.194108 | 0.927 | 0.751 | 7.58965E-29 |
| VEGFA_Terminal | SLC38A1 | 5.1474E-180 | 1.193942 | 0.494 | 0.07 | 1.0025E-175 |
| Contractile | LRPAP1 | 5.6141E-188 | 1.192498 | 0.903 | 0.706 | 1.0933E-183 |
| Contractile | LIMCH1 | 6.0988E-171 | 1.191839 | 0.775 | 0.407 | 1.1877E-166 |
| Fibroblasts | TNXB | 1.11592E-35 | 1.186039 | 0.858 | 0.538 | 2.17326E-31 |
| IFN_Intermediate | IFIH1 | 1.17689E-40 | 1.18453 | 0.561 | 0.134 | 2.29199E-36 |
| Contractile | ENSSSCG0( | 6.5559E-125 | 1.18364 | 0.365 | 0.093 | 1.2768E-120 |
| IFN_Intermediate | B2M.1 | 1.23316E-23 | 1.183118 | 0.991 | 0.873 | 2.40157E-19 |
| CD74_Intermediat | COL6A6 | 1.98438E-74 | 0.458 | 0.134 | 0.019 | 3.86458E-70 |
| Contractile | PALLD | 1.3866E-185 | 1.178426 | 0.697 | 0.288 | 2.7005E-181 |
| VEGFA_Terminal | SLC2A3 | 3.12119E-95 | 1.17755 | 0.697 | 0.252 | 6.07851E-91 |
| Fibroblasts | ITIH5 | 2.37442E-44 | 1.177155 | 0.495 | 0.16 | 4.62418E-40 |
| Intermediate | C7 | 6.80966E-91 | 1.172442 | 0.496 | 0.164 | 1.32618E-86 |
| Pericytes | SIK1 | 1.27269E-19 | 1.171513 | 0.36 | 0.059 | 2.47857E-15 |
| Pericytes | RAMP1 | 1.81229E-10 | 1.170225 | 0.48 | 0.165 | 3.52944E-06 |
| Contractile | LRRFIP1 | 7.0172E-204 | 1.169016 | 0.902 | 0.64 | 1.3666E-199 |
| Terminal | CRYAB | 6.8642E-205 | 1.167495 | 0.953 | 0.812 | 1.3368E-200 |
| Intermediate | CP | 8.10551E-85 | 1.165268 | 0.576 | 0.252 | 1.57855E-80 |
| Pericytes | DUSP5 | 1.0853E-24 | 1.165028 | 0.28 | 0.03 | 2.11362E-20 |
| Pericytes | PDGFRB | 0.001481462 | 1.164818 | 0.54 | 0.451 | 1 |
| IFN_Intermediate | SAMHD1 | 2.97069E-16 | 1.163371 | 0.411 | 0.149 | 5.78542E-12 |
| Contractile | ITGA7 | 2.2266E-244 | 1.162964 | 0.528 | 0.104 | 4.3363E-240 |
| Contractile | PDE4DIP | 1.1695E-183 | 1.160913 | 0.56 | 0.173 | 2.2776E-179 |
| Pericytes | LRRC10B | 2.123E-150 | 1.160878 | 0.34 | 0.006 | 4.1346E-146 |
| Fibroblasts | PDPN | 6.6415E-108 | 1.160356 | 0.349 | 0.035 | 1.2934E-103 |
| Contractile | CCDC3 | 5.0493E-303 | 1.158821 | 0.471 | 0.055 | 9.8335E-299 |
| Fibroblasts | CPXM2 | 3.65472E-09 | 1.158389 | 0.578 | 0.525 | 7.11757E-05 |
| Terminal | FMOD | 1.1172E-167 | 1.157441 | 0.892 | 0.737 | 2.1757E-163 |
| Fibroblasts | MTR | 7.51079E-54 | 1.156923 | 0.312 | 0.058 | 1.46273E-49 |
| Pericytes | PDE5A | 9.81591E-06 | 1.15315 | 0.44 | 0.225 | 0.191164892 |
| Pericytes | NR4A3 | 6.16576E-21 | 1.151687 | 0.34 | 0.049 | 1.20078E-16 |
| Contractile | FILIP1L | 1.3764E-125 | 1.150833 | 0.802 | 0.537 | 2.6806E-121 |
| Pericytes | ENSSSCG0( | 3.61477E-11 | 1.149643 | 0.5 | 0.187 | 7.03976E-07 |
| Fibroblasts | SPON2 | 1.19391E-44 | 1.149026 | 0.601 | 0.222 | 2.32514E-40 |
| Contractile | MSRB3 | 2.7329E-151 | 1.147229 | 0.744 | 0.427 | 5.3223E-147 |
| Contractile | MXI1 | 5.5644E-152 | 1.146825 | 0.67 | 0.306 | 1.0837E-147 |
| CD74_Intermediat | ENSSSCG0( | 2.0535E-49 | 0.289906 | 0.105 | 0.02 | 3.99919E-45 |
| Contractile | DMD | 2.9528E-186 | 1.142967 | 0.534 | 0.15 | 5.7506E-182 |
| Pericytes | SYTL2 | 2.03988E-09 | 1.141647 | 0.4 | 0.136 | 3.97267E-05 |
| Pericytes | ITGA1 | 2.9115E-07 | 1.139928 | 0.8 | 0.614 | 0.005670151 |
| IFN_Intermediate | IFI44L | 6.16773E-35 | 1.13432 | 0.617 | 0.177 | 1.20117E-30 |
| IFN_Intermediate | TAP1 | 7.75126E-40 | 1.133245 | 0.43 | 0.081 | 1.50956E-35 |
| Pericytes | PLCB1 | 1.75757E-05 | 1.13236 | 0.38 | 0.181 | 0.342285795 |
| IFN_Intermediate | ENSSSCG0( | 7.9957E-29 | 1.132265 | 0.579 | 0.188 | 1.55716E-24 |
| VEGFA_Terminal | SLC20A1 | 1.4112E-47 | 1.130603 | 0.55 | 0.25 | 2.74832E-43 |

| | | | | | | |
|---|---|---|---|---|---|---|
| Pericytes | LGALSL | 6.9641E-07 | 1.129233 | 0.56 | 0.345 | 0.013562583 |
| Pericytes | PALM2AKA | 1.65925E-11 | 1.124476 | 0.76 | 0.434 | 3.23139E-07 |
| VEGFA_Terminal | TNC | 5.03327E-42 | 1.122623 | 0.458 | 0.187 | 9.80229E-38 |
| VEGFA_Terminal | LGALS3 | 6.39079E-62 | 1.121887 | 0.971 | 0.774 | 1.24461E-57 |
| Terminal | CDO1 | 1.2141E-139 | 1.121516 | 0.872 | 0.601 | 2.3645E-135 |
| VEGFA_Terminal | CSTB | 1.94849E-90 | 1.117925 | 0.966 | 0.809 | 3.79469E-86 |
| Pericytes | DES | 1.01684E-07 | 1.115873 | 0.32 | 0.097 | 0.001980304 |
| VEGFA_Terminal | ENO2 | 7.5278E-112 | 1.115479 | 0.431 | 0.085 | 1.466E-107 |
| Pericytes | SLC8A1 | 1.64892E-08 | 1.114348 | 0.4 | 0.149 | 0.000321126 |
| Terminal | SOD2 | 1.2913E-103 | 1.113336 | 0.601 | 0.338 | 2.5149E-99 |
| IFN_Intermediate | RTP4 | 1.11128E-45 | 1.111949 | 0.439 | 0.073 | 2.16422E-41 |
| Fibroblasts | MEDAG | 8.3438E-32 | 1.10978 | 0.422 | 0.158 | 1.62495E-27 |
| Pericytes | KCNAB1 | 1.54148E-05 | 1.109634 | 0.46 | 0.253 | 0.300202433 |
| IFN_Intermediate | SLA-DQB1 | 2.96343E-07 | 1.108028 | 0.318 | 0.15 | 0.005771276 |
| Contractile | LIMS2 | 2.6479E-154 | 1.104837 | 0.794 | 0.463 | 5.1568E-150 |
| VEGFA_Terminal | SOD2 | 1.10045E-22 | 1.103823 | 0.61 | 0.409 | 2.14313E-18 |
| Fibroblasts | EBF1 | 6.80407E-31 | 1.103805 | 0.546 | 0.265 | 1.32509E-26 |
| Contractile | IL17B | 1.5407E-147 | 1.102829 | 0.504 | 0.161 | 3.0006E-143 |
| Contractile | PCDH7 | 1.3381E-152 | 1.099443 | 0.729 | 0.365 | 2.6059E-148 |
| Pericytes | PDE1C | 1.88734E-25 | 1.098133 | 0.24 | 0.021 | 3.6756E-21 |
| Fibroblasts | PID1 | 2.4634E-60 | 1.096198 | 0.417 | 0.09 | 4.79747E-56 |
| Pericytes | CXCL14 | 2.74975E-05 | 1.095528 | 0.1 | 0.019 | 0.535514515 |
| Pericytes | CSRNP1 | 5.74162E-17 | 1.094834 | 0.38 | 0.076 | 1.11818E-12 |
| Pericytes | SPARCL1 | 1.9525E-07 | 1.092727 | 0.84 | 0.706 | 0.003802491 |
| IFN_Intermediate | ENSSSCG0( | 3.70621E-09 | 1.092555 | 0.757 | 0.623 | 7.21785E-05 |
| VEGFA_Terminal | PTPRR | 1.11459E-85 | 1.090749 | 0.574 | 0.188 | 2.17067E-81 |
| Contractile | SBSPON | 0 | 1.090004 | 0.395 | 0.019 | 0 |
| CD74_Intermediat | GSN | 1.9778E-292 | 1.089494 | 1 | 0.988 | 3.8518E-288 |
| Contractile | SMARCD3 | 6.8689E-174 | 1.08889 | 0.567 | 0.186 | 1.3377E-169 |
| Intermediate | VCAN | 1.59241E-59 | 1.088469 | 0.896 | 0.764 | 3.10122E-55 |
| IFN_Intermediate | WARS1 | 3.08919E-09 | 1.087854 | 0.28 | 0.109 | 6.01621E-05 |
| IFN_Intermediate | LGALS9 | 9.40808E-58 | 1.087018 | 0.374 | 0.043 | 1.83222E-53 |
| VEGFA_Terminal | ANPEP | 6.91099E-53 | 1.085817 | 0.809 | 0.498 | 1.34592E-48 |
| Contractile | DMPK | 5.1406E-206 | 1.084972 | 0.501 | 0.115 | 1.0011E-201 |
| Pericytes | SLC7A2 | 4.18294E-07 | 1.084244 | 0.48 | 0.23 | 0.008146267 |
| Pericytes | TGFB3 | 0.009115137 | 1.083819 | 0.34 | 0.223 | 1 |
| IFN_Intermediate | SETX | 4.74039E-25 | 1.083616 | 0.533 | 0.174 | 9.2319E-21 |
| Fibroblasts | SERPING1 | 9.00191E-44 | 1.080974 | 0.954 | 0.817 | 1.75312E-39 |
| Contractile | PLCB4 | 0 | 1.079565 | 0.372 | 0.023 | 0 |
| Pericytes | ATP1B2 | 8.90632E-18 | 1.078261 | 0.2 | 0.021 | 1.73451E-13 |
| VEGFA_Terminal | PKM | 7.1102E-110 | 1.07752 | 0.993 | 0.869 | 1.3847E-105 |
| Pericytes | PLN | 2.77539E-08 | 1.07727 | 0.46 | 0.171 | 0.000540508 |
| Pericytes | PTK2 | 6.07719E-08 | 1.076615 | 0.44 | 0.187 | 0.001183533 |
| VEGFA_Terminal | ALDOA | 2.0206E-102 | 1.07634 | 0.978 | 0.824 | 3.9352E-98 |
| Fibroblasts | TGFBR2 | 3.40516E-29 | 1.076011 | 0.688 | 0.469 | 6.63154E-25 |
| VEGFA_Terminal | TIMP1 | 2.44266E-30 | 1.075228 | 0.949 | 0.899 | 4.75708E-26 |
| Fibroblasts | ENSSSCG0( | 8.28796E-29 | 1.074958 | 0.225 | 0.051 | 1.61408E-24 |
| Contractile | MAP3K20 | 3.3878E-186 | 1.074524 | 0.577 | 0.178 | 6.5977E-182 |
| Contractile | CLIC4 | 6.6697E-233 | 1.072792 | 0.978 | 0.871 | 1.2989E-228 |
| IFN_Intermediate | STAT2 | 3.41697E-28 | 1.071948 | 0.477 | 0.13 | 6.65455E-24 |
| Terminal | EDIL3 | 7.2602E-226 | 1.070045 | 0.693 | 0.271 | 1.4139E-221 |
| VEGFA_Terminal | FAM13A | 2.76135E-52 | 1.070028 | 0.538 | 0.231 | 5.37773E-48 |
| Contractile | PBXIP1 | 6.9155E-144 | 1.069902 | 0.667 | 0.319 | 1.3468E-139 |
| Pericytes | AHI1 | 6.04767E-05 | 1.069311 | 0.56 | 0.384 | 1 |
| Pericytes | DNAJA1 | 1.34547E-05 | 1.06918 | 0.68 | 0.512 | 0.262030856 |
| Fibroblasts | MATN4 | 5.01501E-47 | 1.068513 | 0.353 | 0.081 | 9.76674E-43 |
| Fibroblasts | UACA | 7.50629E-16 | 1.068375 | 0.596 | 0.437 | 1.46185E-11 |
| Intermediate | PI16 | 1.88224E-40 | 1.068012 | 0.416 | 0.2 | 3.66566E-36 |
| Fibroblasts | MEOX2 | 1.58165E-13 | 1.067759 | 0.33 | 0.169 | 3.08027E-09 |

| | | | | | | |
|---|---|---|---|---|---|---|
| Fibroblasts | CDKN1C | 6.53884E-27 | 1.067608 | 0.317 | 0.101 | 1.27344E-22 |
| CD74_Intermediat | ENSSSCG0( | 2.7586E-164 | 1.066413 | 0.823 | 0.547 | 5.3724E-160 |
| VEGFA_Terminal | PGAM1 | 1.74218E-83 | 1.066133 | 0.879 | 0.521 | 3.39289E-79 |
| Contractile | CFL2 | 6.9307E-152 | 1.065871 | 0.695 | 0.338 | 1.3498E-147 |
| IFN_Intermediate | SAMD9 | 4.58893E-33 | 1.065354 | 0.383 | 0.075 | 8.93693E-29 |
| Pericytes | MEF2C | 2.31935E-07 | 1.062629 | 0.66 | 0.427 | 0.004516939 |
| Pericytes | PACSIN2 | 1.04248E-06 | 1.062056 | 0.48 | 0.238 | 0.020302289 |
| Pericytes | PRKAR2B | 5.55781E-05 | 1.062025 | 0.4 | 0.217 | 1 |
| Fibroblasts | IGFBP6 | 2.02955E-09 | 1.061896 | 0.702 | 0.613 | 3.95255E-05 |
| CD74_Intermediat | SPON1 | 1.7435E-203 | 1.06187 | 0.702 | 0.332 | 3.3955E-199 |
| CD74_Intermediat | PPL | 8.43176E-34 | 0.276079 | 0.101 | 0.028 | 1.64209E-29 |
| Pericytes | REL | 8.98456E-05 | 1.057314 | 0.3 | 0.132 | 1 |
| VEGFA_Terminal | EMP1 | 4.42592E-65 | 1.056898 | 0.927 | 0.627 | 8.61948E-61 |
| Contractile | ACTN4 | 8.291E-128 | 1.055885 | 0.738 | 0.428 | 1.6147E-123 |
| Contractile | PPP1R3B | 6.175E-193 | 1.050841 | 0.424 | 0.084 | 1.2026E-188 |
| Pericytes | PELI1 | 1.3222E-05 | 1.050005 | 0.32 | 0.134 | 0.257498792 |
| Contractile | CIRBP | 2.3077E-124 | 1.049121 | 0.658 | 0.34 | 4.4942E-120 |
| Terminal | ASPN | 6.69088E-97 | 1.048479 | 0.524 | 0.281 | 1.30305E-92 |
| Pericytes | PDE4B | 0.000608788 | 1.046147 | 0.32 | 0.166 | 1 |
| Pericytes | ACTN4 | 2.44767E-08 | 1.045624 | 0.7 | 0.477 | 0.000476684 |
| VEGFA_Terminal | METRNL | 3.25343E-48 | 1.04457 | 0.591 | 0.282 | 6.33605E-44 |
| Contractile | ZYX | 9.5047E-155 | 1.044439 | 0.8 | 0.458 | 1.851E-150 |
| CD74_Intermediat | B2M.1 | 5.3127E-214 | 1.044379 | 0.961 | 0.841 | 1.0347E-209 |
| IFN_Intermediate | SP140 | 2.16001E-19 | 1.041745 | 0.551 | 0.221 | 4.20662E-15 |
| Terminal | S100A10 | 3.7623E-209 | 1.040125 | 0.957 | 0.72 | 7.3271E-205 |
| Fibroblasts | F2RL2 | 7.88022E-66 | 1.039128 | 0.39 | 0.075 | 1.53467E-61 |
| CD74_Intermediat | TIMP3 | 3.0837E-211 | 1.037055 | 0.993 | 0.919 | 6.0056E-207 |
| Pericytes | DDIT3 | 0.001664531 | 1.036741 | 0.38 | 0.238 | 1 |
| CD74_Intermediat | SPRY1 | 8.2885E-157 | 1.036101 | 0.754 | 0.449 | 1.6142E-152 |
| Pericytes | CRIM1 | 2.97615E-06 | 1.035145 | 0.58 | 0.362 | 0.057960583 |
| Fibroblasts | NR4A1 | 2.01279E-08 | 1.034724 | 0.349 | 0.217 | 0.000391991 |
| Terminal | SH3BGRL3 | 3.5287E-228 | 1.034459 | 0.895 | 0.67 | 6.8721E-224 |
| Contractile | NDUFB1 | 3.7277E-122 | 1.032785 | 0.781 | 0.535 | 7.2597E-118 |
| Pericytes | SPRY2 | 3.26917E-07 | 1.032243 | 0.34 | 0.124 | 0.006366702 |
| VEGFA_Terminal | ATF4 | 4.20177E-41 | 1.031908 | 0.901 | 0.736 | 8.18295E-37 |
| Pericytes | CDC42EP4 | 7.42774E-20 | 1.031153 | 0.4 | 0.072 | 1.44655E-15 |
| Pericytes | EIF4A1 | 0.000121987 | 1.030351 | 0.6 | 0.498 | 1 |
| Intermediate | THBS2 | 1.02023E-70 | 1.028584 | 0.756 | 0.506 | 1.9869E-66 |
| Fibroblasts | TCF21 | 1.0025E-116 | 1.02767 | 0.28 | 0.019 | 1.9524E-112 |
| Terminal | SMPDL3B | 2.6758E-214 | 1.024493 | 0.496 | 0.136 | 5.2111E-210 |
| Pericytes | SCN1B | 2.8076E-09 | 1.022173 | 0.34 | 0.102 | 5.46781E-05 |
| Pericytes | LRRC32 | 4.36913E-05 | 1.021721 | 0.34 | 0.159 | 0.850887189 |
| Contractile | ITPR1 | 1.0807E-136 | 1.021146 | 0.536 | 0.198 | 2.1047E-132 |
| Pericytes | KLHL15 | 9.24713E-09 | 1.017877 | 0.36 | 0.116 | 0.000180088 |
| VEGFA_Terminal | FGFR2 | 6.19647E-73 | 1.016485 | 0.511 | 0.164 | 1.20676E-68 |
| IFN_Intermediate | ENSSSCG0( | 1.67676E-11 | 1.016485 | 0.944 | 0.804 | 3.26548E-07 |
| Fibroblasts | FBLN5 | 6.85472E-18 | 1.015781 | 0.734 | 0.587 | 1.33496E-13 |
| Intermediate | NEXN | 1.87426E-72 | 1.014813 | 0.869 | 0.642 | 3.65012E-68 |
| Pericytes | PLEKHA2 | 1.35359E-06 | 1.013376 | 0.34 | 0.132 | 0.026361226 |
| Fibroblasts | ENSSSCG0( | 2.84971E-24 | 1.012424 | 0.835 | 0.665 | 5.54981E-20 |
| Terminal | PRSS35 | 5.6354E-136 | 1.012242 | 0.445 | 0.161 | 1.0975E-131 |
| CD74_Intermediat | C1S | 7.5706E-251 | 1.011621 | 0.949 | 0.68 | 1.4744E-246 |
| Pericytes | LOC10015: | 6.03033E-11 | 1.011223 | 0.22 | 0.04 | 1.17441E-06 |
| Fibroblasts | ABLIM1 | 1.21055E-19 | 1.010978 | 0.56 | 0.361 | 2.35754E-15 |
| Intermediate | SLIT3 | 2.93997E-98 | 1.006403 | 0.585 | 0.227 | 5.72559E-94 |
| IFN_Intermediate | PPA1 | 2.3523E-13 | 1.005641 | 0.411 | 0.168 | 4.58111E-09 |
| Fibroblasts | CAT | 1.23776E-28 | 1.003009 | 0.606 | 0.337 | 2.41054E-24 |
| VEGFA_Terminal | SULF1 | 2.07729E-54 | 1.002623 | 0.714 | 0.385 | 4.04551E-50 |
| VEGFA_Terminal | PGK1 | 4.16667E-69 | 1.002621 | 0.814 | 0.465 | 8.11459E-65 |

| | | | | | | |
|---|---|---|---|---|---|---|
| Pericytes | TM4SF1 | 0.000454089 | 0.999768 | 0.76 | 0.663 | 1 |
| Fibroblasts | OLFML2A | 1.01792E-37 | 0.999024 | 0.261 | 0.055 | 1.9824E-33 |
| IFN_Intermediate | IFIT5 | 3.66177E-40 | 0.998934 | 0.495 | 0.103 | 7.1313E-36 |
| VEGFA_Terminal | KIT | 4.9789E-116 | 0.998736 | 0.479 | 0.098 | 9.6963E-112 |
| Contractile | ATP1A2 | 1.8315E-291 | 0.998681 | 0.322 | 0.014 | 3.5668E-287 |
| VEGFA_Terminal | TXN | 3.78884E-81 | 0.99852 | 0.985 | 0.936 | 7.37877E-77 |
| Pericytes | SNRK | 1.99283E-08 | 0.99809 | 0.38 | 0.131 | 0.000388104 |
| Contractile | COL18A1 | 3.3864E-104 | 0.997031 | 0.481 | 0.179 | 6.5949E-100 |
| Contractile | FILIP1 | 2.7163E-122 | 0.996866 | 0.589 | 0.25 | 5.29E-118 |
| Contractile | FHOD3 | 5.3415E-200 | 0.994217 | 0.381 | 0.06 | 1.0403E-195 |
| Pericytes | FILIP1 | 5.68808E-06 | 0.994192 | 0.54 | 0.303 | 0.110775408 |
| VEGFA_Terminal | ENSSSCG0( | 1.6677E-104 | 0.993294 | 0.344 | 0.055 | 3.2478E-100 |
| Pericytes | ETS2 | 1.82062E-06 | 0.99275 | 0.4 | 0.173 | 0.035456647 |
| Fibroblasts | CPED1 | 2.58826E-25 | 0.992116 | 0.656 | 0.403 | 5.04064E-21 |
| Fibroblasts | JUN | 4.16145E-11 | 0.991586 | 0.817 | 0.737 | 8.10441E-07 |
| Contractile | ITGA1 | 1.3744E-134 | 0.989342 | 0.842 | 0.572 | 2.6766E-130 |
| Pericytes | SERTAD1 | 1.48475E-05 | 0.989325 | 0.48 | 0.275 | 0.289155963 |
| Contractile | CALM1 | 2.5745E-219 | 0.987582 | 0.991 | 0.95 | 5.0138E-215 |
| Contractile | KCNE1 | 4.2991E-135 | 0.987372 | 0.503 | 0.167 | 8.3725E-131 |
| Contractile | DTNA | 8.4033E-229 | 0.986277 | 0.373 | 0.046 | 1.6365E-224 |
| Contractile | SCHIP1 | 9.4097E-121 | 0.985583 | 0.49 | 0.184 | 1.8325E-116 |
| Contractile | ARHGEF17 | 4.4204E-135 | 0.985123 | 0.531 | 0.196 | 8.6087E-131 |
| IFN_Intermediate | CD74 | 3.03442E-07 | 0.984655 | 0.477 | 0.267 | 0.005909538 |
| Fibroblasts | KLF2 | 3.31821E-09 | 0.983514 | 0.509 | 0.372 | 6.46221E-05 |
| Pericytes | ENSSSCG0( | 1.68545E-07 | 0.983322 | 0.58 | 0.31 | 0.00328242 |
| Terminal | COL1A1 | 2.3533E-157 | 0.980942 | 0.978 | 0.921 | 4.583E-153 |
| Contractile | ENSSSCG0( | 2.0927E-143 | 0.980838 | 0.52 | 0.177 | 4.0756E-139 |
| Pericytes | IGFBP5 | 7.87964E-05 | 0.9808 | 0.48 | 0.274 | 1 |
| Contractile | TACC1 | 1.3797E-119 | 0.980556 | 0.805 | 0.595 | 2.6869E-115 |
| Contractile | ARHGEF25 | 1.5477E-128 | 0.980377 | 0.673 | 0.333 | 3.0142E-124 |
| Pericytes | HIPK3 | 5.71016E-08 | 0.980292 | 0.52 | 0.252 | 0.001112053 |
| Terminal | LUM | 8.7782E-223 | 0.979396 | 0.997 | 0.901 | 1.7096E-218 |
| CD74_Intermediat | ENSSSCG0( | 6.15992E-58 | 0.332419 | 0.138 | 0.03 | 1.19965E-53 |
| VEGFA_Terminal | HK2 | 1.1974E-173 | 0.978546 | 0.278 | 0.017 | 2.332E-169 |
| Pericytes | CALD1 | 3.09369E-14 | 0.978235 | 1 | 0.968 | 6.02495E-10 |
| CD74_Intermediat | TMEM245 | 2.7999E-142 | 0.978123 | 0.73 | 0.452 | 5.4528E-138 |
| Pericytes | ENSSSCG0( | 0.001984565 | 0.97763 | 0.8 | 0.903 | 1 |
| IFN_Intermediate | LOC10015! | 4.2959E-200 | 0.977366 | 0.28 | 0.004 | 8.3662E-196 |
| Pericytes | GNG2 | 6.63499E-08 | 0.977352 | 0.14 | 0.024 | 0.001292165 |
| VEGFA_Terminal | C4ORF3 | 7.3904E-66 | 0.975512 | 0.908 | 0.694 | 1.43928E-61 |
| Fibroblasts | PLPP3 | 1.25443E-23 | 0.972897 | 0.931 | 0.903 | 2.443E-19 |
| Terminal | SPARC | 3.3889E-196 | 0.972576 | 0.978 | 0.903 | 6.5998E-192 |
| Pericytes | COL4A2 | 0.000597482 | 0.972495 | 0.7 | 0.585 | 1 |
| VEGFA_Terminal | VMP1 | 6.05659E-59 | 0.9724 | 0.92 | 0.659 | 1.17952E-54 |
| IFN_Intermediate | C1S | 5.92484E-14 | 0.97133 | 0.935 | 0.754 | 1.15386E-09 |
| Pericytes | ANGPT4 | 6.83562E-10 | 0.97119 | 0.2 | 0.037 | 1.33124E-05 |
| Contractile | IL11RA | 1.9061E-144 | 0.971081 | 0.545 | 0.193 | 3.7121E-140 |
| Pericytes | LOC10051( | 0.002677601 | 0.97054 | 0.44 | 0.283 | 1 |
| Intermediate | MEGF6 | 2.24236E-94 | 0.969959 | 0.335 | 0.08 | 4.36699E-90 |
| Contractile | SMOC2 | 7.3914E-184 | 0.9696 | 0.44 | 0.091 | 1.4395E-179 |
| Intermediate | CCN1 | 3.98794E-26 | 0.969359 | 0.601 | 0.421 | 7.76651E-22 |
| Pericytes | ESAM | 4.32801E-21 | 0.969337 | 0.32 | 0.044 | 8.42881E-17 |
| Fibroblasts | CD55 | 1.508E-17 | 0.968626 | 0.257 | 0.095 | 2.93684E-13 |
| Pericytes | MAP3K8 | 2.06383E-07 | 0.968532 | 0.36 | 0.134 | 0.004019302 |
| Pericytes | SLC25A25 | 5.08258E-23 | 0.968158 | 0.26 | 0.028 | 9.89833E-19 |
| Pericytes | PHLDA2 | 0.002792397 | 0.96808 | 0.62 | 0.513 | 1 |
| Pericytes | SOCS1 | 3.90062E-12 | 0.966746 | 0.26 | 0.049 | 7.59646E-08 |
| VEGFA_Terminal | HK1 | 4.77657E-63 | 0.965607 | 0.893 | 0.644 | 9.30237E-59 |
| Fibroblasts | PDK4 | 3.44362E-22 | 0.965458 | 0.312 | 0.107 | 6.70646E-18 |

| | | | | | | |
|---|---|---|---|---|---|---|
| Pericytes | PGF | 1.23256E-13 | 0.96405 | 0.3 | 0.058 | 2.40041E-09 |
| Pericytes | TAGLN | 9.86904E-10 | 0.963378 | 1 | 0.963 | 1.92199E-05 |
| Contractile | KANK2 | 1.8155E-124 | 0.962368 | 0.778 | 0.496 | 3.5357E-120 |
| Contractile | CLDN10 | 0 | 0.961412 | 0.271 | 0.002 | 0 |
| Pericytes | TPM1 | 6.38413E-07 | 0.960746 | 0.9 | 0.821 | 0.012433087 |
| Pericytes | COQ10B | 4.63085E-06 | 0.960362 | 0.42 | 0.194 | 0.090185808 |
| Contractile | ENSSSCG0( | 9.7601E-132 | 0.959542 | 0.827 | 0.589 | 1.9008E-127 |
| Terminal | S100A1 | 6.5429E-51 | 0.957449 | 0.247 | 0.103 | 1.27423E-46 |
| Pericytes | KITLG | 1.32035E-05 | 0.957428 | 0.32 | 0.128 | 0.257138713 |
| Pericytes | NAMPT | 0.000357308 | 0.957111 | 0.44 | 0.269 | 1 |
| Pericytes | TNFSF12 | 7.16878E-06 | 0.956661 | 0.5 | 0.279 | 0.139611942 |
| Pericytes | TIPARP | 3.03609E-05 | 0.955269 | 0.24 | 0.085 | 0.591279055 |
| Pericytes | COX17 | 2.14595E-05 | 0.955116 | 0.56 | 0.374 | 0.417923921 |
| Pericytes | FXYD6 | 9.47397E-07 | 0.954801 | 0.54 | 0.287 | 0.018450552 |
| VEGFA_Terminal | ID2 | 1.27852E-28 | 0.954121 | 0.877 | 0.755 | 2.48993E-24 |
| VEGFA_Terminal | RGS4 | 2.65612E-50 | 0.953435 | 0.329 | 0.095 | 5.1728E-46 |
| Pericytes | FRMD4A | 4.30364E-13 | 0.951839 | 0.32 | 0.068 | 8.38135E-09 |
| Pericytes | DDX5 | 4.38983E-09 | 0.951546 | 0.92 | 0.845 | 8.54919E-05 |
| Pericytes | NPY1R | 2.92673E-06 | 0.950127 | 0.2 | 0.054 | 0.056998072 |
| Contractile | TMEM47 | 9.36E-119 | 0.950011 | 0.8 | 0.575 | 1.8229E-114 |
| Contractile | MAST4 | 2.6902E-117 | 0.948184 | 0.473 | 0.172 | 5.2391E-113 |
| Pericytes | GALR1 | 3.1577E-218 | 0.947948 | 0.18 | 0 | 6.1497E-214 |
| Pericytes | OLFML2B | 1.90287E-06 | 0.947499 | 0.28 | 0.096 | 0.037058414 |
| Contractile | ATP2A2 | 4.1793E-106 | 0.94618 | 0.602 | 0.303 | 8.1391E-102 |
| Pericytes | ADGRA2 | 7.07773E-10 | 0.945602 | 0.26 | 0.059 | 1.37839E-05 |
| Pericytes | FAM13C | 0.000147269 | 0.943843 | 0.44 | 0.255 | 1 |
| Pericytes | IFITM1 | 2.5618E-05 | 0.93852 | 0.22 | 0.074 | 0.498911093 |
| Pericytes | ADAP2 | 2.75623E-37 | 0.938192 | 0.14 | 0.005 | 5.36776E-33 |
| Fibroblasts | SESN3 | 1.33122E-19 | 0.934431 | 0.518 | 0.3 | 2.59256E-15 |
| Fibroblasts | EPHA3 | 2.65448E-69 | 0.934023 | 0.266 | 0.032 | 5.1696E-65 |
| Pericytes | ETS1 | 1.3398E-07 | 0.93312 | 0.54 | 0.268 | 0.00260926 |
| Pericytes | ART4 | 7.03944E-46 | 0.932315 | 0.24 | 0.011 | 1.37093E-41 |
| Contractile | TTLL7 | 6.0897E-111 | 0.931767 | 0.704 | 0.399 | 1.186E-106 |
| Pericytes | ELL2 | 0.000190742 | 0.931412 | 0.36 | 0.195 | 1 |
| VEGFA_Terminal | PLOD1 | 3.28091E-74 | 0.929765 | 0.564 | 0.197 | 6.38958E-70 |
| Intermediate | PENK | 9.95296E-30 | 0.928239 | 0.233 | 0.094 | 1.93834E-25 |
| Contractile | NUF2 | 5.5283E-171 | 0.927415 | 0.334 | 0.053 | 1.0766E-166 |
| Contractile | ENSSSCG0( | 2.2606E-101 | 0.92734 | 0.694 | 0.41 | 4.40258E-97 |
| Pericytes | CTDSPL | 0.000189564 | 0.92661 | 0.34 | 0.181 | 1 |
| Pericytes | RASGEF1B | 5.75184E-08 | 0.926254 | 0.32 | 0.104 | 0.00112017 |
| IFN_Intermediate | ENSSSCG0( | 2.07457E-45 | 0.924874 | 0.364 | 0.051 | 4.04023E-41 |
| Pericytes | KLF6 | 0.000781099 | 0.923869 | 0.66 | 0.617 | 1 |
| VEGFA_Terminal | SOX9 | 1.45205E-91 | 0.923118 | 0.349 | 0.065 | 2.82786E-87 |
| VEGFA_Terminal | P4HA2 | 3.53156E-52 | 0.923026 | 0.789 | 0.486 | 6.87772E-48 |
| Intermediate | MFAP4 | 3.40506E-95 | 0.922428 | 0.971 | 0.844 | 6.63135E-91 |
| Pericytes | C2CD2 | 1.83195E-07 | 0.922367 | 0.32 | 0.107 | 0.003567726 |
| Pericytes | TBX2 | 1.91621E-18 | 0.922069 | 0.28 | 0.039 | 3.73181E-14 |
| VEGFA_Terminal | GDF15 | 7.56324E-63 | 0.921866 | 0.186 | 0.026 | 1.47294E-58 |
| Fibroblasts | TMEM150( | 3.08223E-25 | 0.919175 | 0.445 | 0.192 | 6.00264E-21 |
| VEGFA_Terminal | PKIB | 1.5759E-173 | 0.916466 | 0.269 | 0.016 | 3.069E-169 |
| Fibroblasts | TCF4 | 1.06585E-22 | 0.915452 | 0.711 | 0.522 | 2.07574E-18 |
| Contractile | ATL2 | 5.3282E-127 | 0.915088 | 0.426 | 0.129 | 1.0377E-122 |
| Pericytes | KCNJ8 | 6.81542E-23 | 0.91505 | 0.22 | 0.02 | 1.3273E-18 |
| VEGFA_Terminal | PRDX6 | 2.56177E-72 | 0.914754 | 0.552 | 0.194 | 4.98905E-68 |
| Pericytes | RELL1 | 3.6993E-07 | 0.911956 | 0.3 | 0.1 | 0.007204385 |
| IFN_Intermediate | HLA-DRA | 2.33662E-07 | 0.910303 | 0.224 | 0.085 | 0.00455057 |
| Contractile | BCAM | 4.9674E-160 | 0.907242 | 0.439 | 0.108 | 9.6739E-156 |
| Pericytes | ENSSSCG0( | 1.32382E-14 | 0.907078 | 0.22 | 0.031 | 2.57813E-10 |
| IFN_Intermediate | IRF7 | 7.48347E-61 | 0.90515 | 0.383 | 0.042 | 1.45741E-56 |

| | | | | | | |
|---|---|---|---|---|---|---|
| Pericytes | RALGPS2 | 4.41518E-12 | 0.904977 | 0.28 | 0.057 | 8.59856E-08 |
| Fibroblasts | ZMIZ1 | 9.68384E-24 | 0.904364 | 0.537 | 0.291 | 1.88593E-19 |
| Pericytes | APOLD1 | 1.94708E-29 | 0.90386 | 0.1 | 0.003 | 3.79195E-25 |
| Pericytes | LOC10052( | 3.37022E-05 | 0.903805 | 0.42 | 0.213 | 0.656350288 |
| Pericytes | RERGL | 3.7425E-139 | 0.900182 | 0.12 | 0 | 7.2884E-135 |
| IFN_Intermediate | CMPK2 | 1.8172E-64 | 0.900065 | 0.299 | 0.024 | 3.539E-60 |
| Pericytes | BCAM | 8.37566E-09 | 0.899892 | 0.44 | 0.16 | 0.000163116 |
| IFN_Intermediate | DDX60 | 3.3593E-114 | 0.899253 | 0.355 | 0.018 | 6.5422E-110 |
| Contractile | ENSSSCG0( | 0 | 0.899125 | 0.297 | 0.005 | 0 |
| Fibroblasts | ENSSSCG0( | 1.46582E-58 | 0.899011 | 0.261 | 0.036 | 2.85469E-54 |
| Contractile | PYGB | 1.9911E-124 | 0.898829 | 0.463 | 0.155 | 3.8777E-120 |
| Fibroblasts | HPGD | 2.6653E-157 | 0.89705 | 0.183 | 0.003 | 5.1907E-153 |
| Contractile | SUN2 | 7.503E-101 | 0.896811 | 0.597 | 0.295 | 1.4612E-96 |
| Pericytes | CCDC3 | 5.25575E-07 | 0.896192 | 0.34 | 0.121 | 0.010235577 |
| Terminal | MYBL1 | 7.66344E-95 | 0.895214 | 0.397 | 0.17 | 1.49245E-90 |
| IFN_Intermediate | B2M | 6.20581E-11 | 0.895184 | 0.953 | 0.86 | 1.20858E-06 |
| Terminal | LCN2 | 1.6645E-100 | 0.894812 | 0.327 | 0.107 | 3.24155E-96 |
| VEGFA_Terminal | ACKR3 | 5.30634E-51 | 0.894647 | 0.322 | 0.091 | 1.03341E-46 |
| Pericytes | SEMA5A | 3.54488E-63 | 0.894631 | 0.24 | 0.008 | 6.90365E-59 |
| Contractile | TES | 9.5253E-122 | 0.894586 | 0.442 | 0.144 | 1.855E-117 |
| Pericytes | ENSSSCG0( | 8.16208E-10 | 0.894396 | 0.22 | 0.044 | 1.58957E-05 |
| Fibroblasts | KCTD12 | 1.04142E-25 | 0.894371 | 0.381 | 0.146 | 2.02817E-21 |
| Contractile | RAP1A | 1.9003E-109 | 0.893362 | 0.711 | 0.431 | 3.7008E-105 |
| Pericytes | MAP3K5 | 5.06014E-15 | 0.892827 | 0.2 | 0.025 | 9.85462E-11 |
| CD74_Intermediat | FBLN1 | 6.6458E-211 | 0.89199 | 0.933 | 0.601 | 1.2943E-206 |
| CD74_Intermediat | C1R | 2.9952E-236 | 0.891705 | 0.986 | 0.84 | 5.8331E-232 |
| Fibroblasts | THBD | 6.76933E-21 | 0.891111 | 0.22 | 0.064 | 1.31833E-16 |
| Pericytes | RCSD1 | 3.05825E-09 | 0.890119 | 0.34 | 0.103 | 5.95593E-05 |
| Fibroblasts | COLEC12 | 8.1229E-176 | 0.887428 | 0.229 | 0.006 | 1.5819E-171 |
| Pericytes | ENSSSCG0( | 6.17866E-05 | 0.886836 | 0.5 | 0.318 | 1 |
| Contractile | FAM13C | 1.5594E-109 | 0.886496 | 0.516 | 0.207 | 3.037E-105 |
| CD74_Intermediat | FBLN2 | 1.1925E-162 | 0.884619 | 0.68 | 0.335 | 2.3223E-158 |
| Pericytes | CTNNA3 | 7.4474E-51 | 0.884051 | 0.22 | 0.008 | 1.45038E-46 |
| VEGFA_Terminal | LCN2 | 3.5478E-20 | 0.883959 | 0.341 | 0.166 | 6.90934E-16 |
| Terminal | FN1 | 4.7724E-203 | 0.882707 | 1 | 0.988 | 9.2942E-199 |
| Fibroblasts | FOXO1 | 1.48759E-26 | 0.882591 | 0.45 | 0.197 | 2.89709E-22 |
| Terminal | TNC | 1.9475E-130 | 0.880849 | 0.385 | 0.119 | 3.7927E-126 |
| CD74_Intermediat | SERPINF1 | 1.909E-212 | 0.880646 | 0.982 | 0.676 | 3.7177E-208 |
| CD74_Intermediat | CTSH | 6.7892E-191 | 0.880004 | 0.953 | 0.717 | 1.3222E-186 |
| Pericytes | SH2B3 | 2.83783E-09 | 0.879294 | 0.22 | 0.047 | 5.52668E-05 |
| Intermediate | RARRES2 | 5.38083E-90 | 0.878619 | 0.98 | 0.902 | 1.04792E-85 |
| Fibroblasts | FLRT2 | 2.53079E-30 | 0.875843 | 0.312 | 0.09 | 4.92872E-26 |
| Contractile | MICAL3 | 1.1259E-111 | 0.875293 | 0.382 | 0.113 | 2.1927E-107 |
| Terminal | SEMA3C | 1.15705E-86 | 0.875258 | 0.486 | 0.253 | 2.25336E-82 |
| CD74_Intermediat | B2M | 3.2287E-167 | 0.875092 | 0.938 | 0.831 | 6.2878E-163 |
| Contractile | UNC45A | 8.0873E-118 | 0.874767 | 0.444 | 0.149 | 1.575E-113 |
| Terminal | NBL1 | 2.1588E-127 | 0.874658 | 0.802 | 0.576 | 4.2042E-123 |
| Pericytes | PLEKHH3 | 2.62564E-06 | 0.873751 | 0.28 | 0.099 | 0.051134328 |
| Terminal | CD9 | 9.3696E-222 | 0.873691 | 0.987 | 0.924 | 1.8247E-217 |
| Pericytes | INPP4B | 6.62276E-20 | 0.873569 | 0.18 | 0.015 | 1.28978E-15 |
| CD74_Intermediat | ENSSSCG0( | 1.9983E-161 | 0.824321 | 0.255 | 0.032 | 3.8916E-157 |
| Fibroblasts | KLF5 | 4.90647E-12 | 0.872767 | 0.193 | 0.073 | 9.55536E-08 |
| Pericytes | ARHGAP29 | 1.2564E-06 | 0.87246 | 0.5 | 0.245 | 0.024468409 |
| Fibroblasts | EIF4EBP2 | 1.08479E-21 | 0.872235 | 0.615 | 0.399 | 2.11262E-17 |
| Fibroblasts | FBP1 | 6.79686E-13 | 0.872059 | 0.312 | 0.149 | 1.32369E-08 |
| Fibroblasts | DHRS3 | 1.24218E-18 | 0.87185 | 0.564 | 0.338 | 2.41915E-14 |
| Fibroblasts | LAMC1 | 1.94336E-21 | 0.871724 | 0.794 | 0.665 | 3.78469E-17 |
| Terminal | CLU | 5.4728E-231 | 0.871432 | 0.999 | 0.945 | 1.0658E-226 |
| Intermediate | FABP4 | 2.42941E-44 | 0.871261 | 0.31 | 0.119 | 4.73128E-40 |

| | | | | | | |
|---|---|---|---|---|---|---|
| Pericytes | NSG1 | 8.5842E-20 | 0.871217 | 0.24 | 0.027 | 1.67177E-15 |
| VEGFA_Terminal | MIF | 2.99561E-72 | 0.870975 | 0.976 | 0.856 | 5.83395E-68 |
| Fibroblasts | ADH1A | 7.3347E-150 | 0.87094 | 0.138 | 0.001 | 1.4284E-145 |
| Fibroblasts | ANG | 1.32879E-20 | 0.868822 | 0.619 | 0.418 | 2.58782E-16 |
| Pericytes | GPER1 | 8.57195E-06 | 0.866838 | 0.28 | 0.103 | 0.166938773 |
| Pericytes | LIFR | 0.000187241 | 0.865556 | 0.22 | 0.081 | 1 |
| VEGFA_Terminal | SMPDL3B | 1.26129E-44 | 0.865534 | 0.533 | 0.232 | 2.45636E-40 |
| VEGFA_Terminal | CTSH | 7.66294E-37 | 0.864641 | 0.927 | 0.774 | 1.49236E-32 |
| Contractile | LDHB | 1.843E-113 | 0.864192 | 0.853 | 0.682 | 3.5892E-109 |
| Pericytes | CNN3 | 1.18469E-08 | 0.863862 | 0.84 | 0.661 | 0.000230718 |
| Pericytes | PLAU | 7.59456E-06 | 0.863764 | 0.16 | 0.039 | 0.147904137 |
| CD74_Intermediat | PLAC9 | 7.4338E-162 | 0.863755 | 0.862 | 0.614 | 1.4477E-157 |
| Terminal | GALNT15 | 2.04393E-76 | 0.862574 | 0.491 | 0.274 | 3.98055E-72 |
| Fibroblasts | CD34 | 8.46317E-17 | 0.860591 | 0.454 | 0.25 | 1.6482E-12 |
| Pericytes | PRXL2A | 1.16756E-11 | 0.860268 | 0.38 | 0.1 | 2.27382E-07 |
| VEGFA_Terminal | ENSSSCG0( | 5.97812E-58 | 0.859966 | 0.702 | 0.344 | 1.16424E-53 |
| Pericytes | CAMK1 | 9.31887E-10 | 0.859682 | 0.46 | 0.163 | 1.81485E-05 |
| VEGFA_Terminal | BHLHE40 | 2.00032E-38 | 0.858039 | 0.821 | 0.559 | 3.89563E-34 |
| Intermediate | RCAN2 | 2.40353E-52 | 0.85784 | 0.432 | 0.186 | 4.68087E-48 |
| Contractile | FIBIN | 3.2574E-105 | 0.857613 | 0.299 | 0.071 | 6.3438E-101 |
| Contractile | HSPB1 | 1.0502E-117 | 0.85688 | 0.962 | 0.902 | 2.0453E-113 |
| IFN_Intermediate | FUNDC1 | 7.80763E-10 | 0.856668 | 0.495 | 0.262 | 1.52054E-05 |
| VEGFA_Terminal | KITLG | 6.94794E-72 | 0.85648 | 0.409 | 0.109 | 1.35311E-67 |
| Pericytes | PHLDB2 | 6.02072E-06 | 0.856187 | 0.5 | 0.266 | 0.11725344 |
| Contractile | HEY2 | 9.5214E-92 | 0.855976 | 0.597 | 0.312 | 1.85429E-87 |
| Contractile | ENSSSCG0( | 1.37534E-86 | 0.855421 | 0.607 | 0.344 | 2.67848E-82 |
| Contractile | HSPB6 | 1.1572E-118 | 0.855323 | 0.878 | 0.686 | 2.2537E-114 |
| VEGFA_Terminal | VDAC1 | 7.04881E-46 | 0.854208 | 0.794 | 0.541 | 1.37276E-41 |
| VEGFA_Terminal | ATF3 | 1.35283E-16 | 0.853408 | 0.293 | 0.143 | 2.63464E-12 |
| IFN_Intermediate | PSME1 | 1.20544E-11 | 0.852929 | 0.757 | 0.548 | 2.34759E-07 |
| Contractile | CCL19 | 8.1318E-150 | 0.851534 | 0.484 | 0.135 | 1.5837E-145 |
| Pericytes | PPP1R12A | 1.09469E-08 | 0.85074 | 0.86 | 0.683 | 0.00021319 |
| Contractile | ENSSSCG0( | 4.865E-266 | 0.850513 | 1 | 0.997 | 9.4747E-262 |
| Pericytes | CEBPD | 0.002038106 | 0.850298 | 0.78 | 0.829 | 1 |
| Contractile | ATF3 | 7.04586E-09 | 0.850247 | 0.209 | 0.142 | 0.000137218 |
| Contractile | YAP1 | 7.47439E-89 | 0.849105 | 0.687 | 0.42 | 1.45564E-84 |
| VEGFA_Terminal | PPP1R3C | 3.49629E-23 | 0.844657 | 0.455 | 0.243 | 6.80902E-19 |
| Contractile | PLIN4 | 2.1626E-303 | 0.844291 | 0.277 | 0.003 | 4.2116E-299 |
| CD74_Intermediat | ABCA1 | 2.382E-118 | 0.843765 | 0.76 | 0.513 | 4.639E-114 |
| VEGFA_Terminal | FAM162A | 8.59085E-27 | 0.842659 | 0.656 | 0.431 | 1.67307E-22 |
| Intermediate | C4B | 5.18358E-39 | 0.84226 | 0.45 | 0.236 | 1.0095E-34 |
| Pericytes | CAV2 | 4.57929E-06 | 0.842018 | 0.6 | 0.403 | 0.089181738 |
| Contractile | SPC24 | 2.7263E-118 | 0.841071 | 0.305 | 0.066 | 5.3094E-114 |
| Fibroblasts | PI4K2B | 7.96208E-18 | 0.840241 | 0.509 | 0.319 | 1.55061E-13 |
| CD74_Intermediat | FBLN5 | 1.1009E-137 | 0.839818 | 0.802 | 0.509 | 2.144E-133 |
| Terminal | HTRA1 | 4.569E-203 | 0.839532 | 0.976 | 0.858 | 8.8982E-199 |
| Contractile | ROCK1 | 1.38132E-87 | 0.838245 | 0.787 | 0.571 | 2.69013E-83 |
| Pericytes | WTIP | 5.78698E-07 | 0.837999 | 0.26 | 0.08 | 0.011270136 |
| Fibroblasts | GALNT15 | 4.78338E-13 | 0.837422 | 0.523 | 0.337 | 9.31563E-09 |
| Contractile | ENSSSCG0( | 3.50592E-94 | 0.836676 | 0.682 | 0.393 | 6.82777E-90 |
| VEGFA_Terminal | PRDX4 | 3.94998E-49 | 0.835059 | 0.843 | 0.638 | 7.69259E-45 |
| IFN_Intermediate | XAF1 | 3.37736E-69 | 0.832847 | 0.346 | 0.03 | 6.57741E-65 |
| Pericytes | FAM107B | 4.47041E-05 | 0.832437 | 0.34 | 0.159 | 0.870611626 |
| Contractile | ANO6 | 4.43711E-88 | 0.830802 | 0.607 | 0.332 | 8.64128E-84 |
| Pericytes | PPP1R14A | 5.40536E-18 | 0.829121 | 0.96 | 0.323 | 1.05269E-13 |
| Pericytes | SEPTIN8 | 0.00634579 | 0.828596 | 0.44 | 0.314 | 1 |
| Pericytes | KCNS3 | 1.67091E-26 | 0.828382 | 0.18 | 0.011 | 3.2541E-22 |
| IFN_Intermediate | ZC3HAV1 | 7.26221E-18 | 0.828108 | 0.43 | 0.143 | 1.41432E-13 |
| Pericytes | AJUBA | 1.16034E-25 | 0.827813 | 0.3 | 0.033 | 2.25977E-21 |

| | | | | | | |
|---|---|---|---|---|---|---|
| Fibroblasts | MTARC2 | 1.66329E-46 | 0.826568 | 0.252 | 0.043 | 3.23925E-42 |
| IFN_Intermediate | LAP3 | 5.93456E-07 | 0.82589 | 0.607 | 0.461 | 0.011557556 |
| Terminal | FAP | 2.9157E-117 | 0.825669 | 0.614 | 0.339 | 5.6784E-113 |
| Contractile | ADCY5 | 1.0122E-189 | 0.825402 | 0.327 | 0.042 | 1.9712E-185 |
| Pericytes | ENSSSCG0( | 3.23289E-05 | 0.82459 | 0.86 | 0.709 | 0.629605095 |
| Pericytes | SEPTIN7 | 3.81134E-10 | 0.824419 | 1 | 0.934 | 7.42258E-06 |
| CD74_Intermediat | ENSSSCG0( | 4.55952E-73 | 0.447022 | 0.161 | 0.032 | 8.87967E-69 |
| Fibroblasts | IL33 | 8.76536E-42 | 0.824191 | 0.206 | 0.031 | 1.70705E-37 |
| Contractile | VASP | 6.21941E-98 | 0.823885 | 0.484 | 0.204 | 1.21123E-93 |
| VEGFA_Terminal | CRLF1 | 1.35903E-24 | 0.823227 | 0.596 | 0.368 | 2.64671E-20 |
| Fibroblasts | IL6ST | 2.26652E-23 | 0.822503 | 0.817 | 0.663 | 4.41405E-19 |
| Contractile | TOB2 | 8.52749E-83 | 0.821207 | 0.759 | 0.53 | 1.66073E-78 |
| Pericytes | ENSSSCG0( | 0.000101661 | 0.820883 | 0.24 | 0.088 | 1 |
| Pericytes | GPC3 | 5.45453E-12 | 0.820622 | 0.22 | 0.036 | 1.06227E-07 |
| Contractile | ILK | 1.6149E-104 | 0.820044 | 0.8 | 0.543 | 3.145E-100 |
| Pericytes | TRPV2 | 1.11134E-36 | 0.818665 | 0.2 | 0.01 | 2.16434E-32 |
| Pericytes | CHN1 | 1.44867E-20 | 0.81862 | 0.2 | 0.018 | 2.82128E-16 |
| VEGFA_Terminal | GPI | 1.4229E-44 | 0.818525 | 0.634 | 0.323 | 2.77109E-40 |
| Fibroblasts | EBF2 | 2.23639E-71 | 0.8169 | 0.161 | 0.01 | 4.35537E-67 |
| Terminal | VMP1 | 1.458E-153 | 0.816738 | 0.86 | 0.589 | 2.8394E-149 |
| Pericytes | PPA1 | 0.000179518 | 0.815867 | 0.34 | 0.17 | 1 |
| Pericytes | SLC38A2 | 0.000150416 | 0.815428 | 0.64 | 0.468 | 1 |
| VEGFA_Terminal | BHLHE41 | 1.09864E-35 | 0.814858 | 0.639 | 0.358 | 2.1396E-31 |
| VEGFA_Terminal | S100A10 | 7.52284E-39 | 0.814621 | 0.913 | 0.788 | 1.46507E-34 |
| IFN_Intermediate | SNX10 | 1.07449E-12 | 0.813506 | 0.383 | 0.15 | 2.09256E-08 |
| IFN_Intermediate | LOC10051 | 2.88827E-15 | 0.81258 | 0.336 | 0.105 | 5.6249E-11 |
| Contractile | LYNX1 | 2.17877E-89 | 0.812511 | 0.569 | 0.283 | 4.24316E-85 |
| Pericytes | EPHX3 | 1.7837E-34 | 0.81095 | 0.16 | 0.007 | 3.47375E-30 |
| IFN_Intermediate | IFI44 | 8.99263E-34 | 0.809744 | 0.299 | 0.046 | 1.75131E-29 |
| VEGFA_Terminal | TCEA3 | 3.65666E-28 | 0.809615 | 0.666 | 0.416 | 7.12134E-24 |
| IFN_Intermediate | TENT5A | 1.90819E-09 | 0.806092 | 0.738 | 0.493 | 3.71619E-05 |
| Pericytes | EIF1 | 2.05812E-07 | 0.806016 | 0.96 | 0.986 | 0.004008187 |
| Contractile | INKA2 | 3.6008E-197 | 0.805906 | 0.304 | 0.032 | 7.0126E-193 |
| Pericytes | ENSSSCG0( | 0.007405835 | 0.805354 | 0.44 | 0.327 | 1 |
| Intermediate | LOC10051 | 6.74495E-41 | 0.803749 | 0.492 | 0.259 | 1.31358E-36 |
| Fibroblasts | ENSSSCG0( | 2.55273E-64 | 0.803505 | 0.239 | 0.027 | 4.97144E-60 |
| CD74_Intermediat | IFI30 | 1.9658E-133 | 0.803284 | 0.64 | 0.331 | 3.8284E-129 |
| VEGFA_Terminal | TMEM45A | 1.73727E-61 | 0.802948 | 0.579 | 0.226 | 3.38334E-57 |
| Fibroblasts | FBN1 | 2.45927E-08 | 0.802531 | 0.688 | 0.64 | 0.000478942 |
| Terminal | COL1A2 | 5.4845E-126 | 0.801755 | 0.981 | 0.929 | 1.0681E-121 |
| Contractile | DACT3 | 2.3392E-106 | 0.801601 | 0.436 | 0.154 | 4.5555E-102 |
| Pericytes | PDE3A | 0.00024768 | 0.801331 | 0.52 | 0.342 | 1 |
| Pericytes | LGI4 | 1.53687E-05 | 0.801203 | 0.2 | 0.061 | 0.299304609 |
| Pericytes | BTG3 | 6.73355E-05 | 0.801145 | 0.46 | 0.256 | 1 |
| IFN_Intermediate | SLFN11 | 4.26893E-57 | 0.801072 | 0.271 | 0.022 | 8.31374E-53 |
| VEGFA_Terminal | THBS1 | 1.03497E-24 | 0.800884 | 0.755 | 0.536 | 2.0156E-20 |
| Fibroblasts | MMP2 | 6.94014E-18 | 0.800733 | 0.835 | 0.772 | 1.35159E-13 |
| Fibroblasts | ZFP36L2 | 9.21446E-16 | 0.800131 | 0.917 | 0.884 | 1.79452E-11 |
| Pericytes | ARHGAP28 | 5.75773E-80 | 0.79967 | 0.18 | 0.003 | 1.12132E-75 |
| Terminal | RGCC | 3.52171E-80 | 0.799523 | 0.426 | 0.204 | 6.85853E-76 |
| Pericytes | NR2F1 | 9.74945E-27 | 0.799375 | 0.24 | 0.02 | 1.89871E-22 |
| Contractile | PARVA | 6.68728E-75 | 0.799265 | 0.603 | 0.35 | 1.30235E-70 |
| Pericytes | UBB | 4.13916E-09 | 0.799117 | 0.98 | 0.979 | 8.06102E-05 |
| Contractile | THRB | 4.84229E-74 | 0.798687 | 0.523 | 0.274 | 9.43036E-70 |
| Terminal | ENSSSCG0( | 8.7105E-143 | 0.797874 | 0.589 | 0.264 | 1.6964E-138 |
| Contractile | BEND5 | 9.4231E-207 | 0.797689 | 0.273 | 0.02 | 1.8352E-202 |
| Fibroblasts | UAP1 | 3.45653E-07 | 0.79704 | 0.417 | 0.316 | 0.006731584 |
| Fibroblasts | CCNL1 | 4.73148E-16 | 0.796792 | 0.436 | 0.242 | 9.21457E-12 |
| VEGFA_Terminal | VDAC3 | 4.72769E-44 | 0.796029 | 0.668 | 0.368 | 9.20717E-40 |

| | | | | | | |
|---|---|---|---|---|---|---|
| VEGFA_Terminal | ADAMTSL4 | 4.11695E-63 | 0.795291 | 0.55 | 0.205 | 8.01776E-59 |
| Pericytes | ENSSSCG0( | 2.98141E-08 | 0.794998 | 0.24 | 0.059 | 0.000580629 |
| Pericytes | TMEM200 | 3.87956E-08 | 0.79477 | 0.18 | 0.036 | 0.000755545 |
| Terminal | DCN | 2.58732E-61 | 0.794524 | 0.913 | 0.862 | 5.0388E-57 |
| Contractile | MRGPRF | 2.3998E-91 | 0.794237 | 0.469 | 0.196 | 4.67361E-87 |
| VEGFA_Terminal | GALNT1 | 7.0117E-34 | 0.793942 | 0.622 | 0.37 | 1.36553E-29 |
| IFN_Intermediate | SOD2 | 1.07385E-06 | 0.792022 | 0.598 | 0.419 | 0.020913136 |
| Contractile | BEX1 | 1.1323E-153 | 0.79094 | 0.271 | 0.035 | 2.2052E-149 |
| VEGFA_Terminal | NDFIP1 | 1.05876E-46 | 0.790916 | 0.891 | 0.679 | 2.06193E-42 |
| Intermediate | FILIP1L | 1.14451E-32 | 0.790582 | 0.733 | 0.562 | 2.22893E-28 |
| VEGFA_Terminal | SLC22A18 | 8.28495E-77 | 0.790312 | 0.32 | 0.065 | 1.61349E-72 |
| Pericytes | OSMR | 0.002024441 | 0.790296 | 0.42 | 0.283 | 1 |
| Terminal | CST3 | 1.1752E-252 | 0.790267 | 0.998 | 0.993 | 2.2887E-248 |
| Fibroblasts | MPZL1 | 1.33952E-17 | 0.790257 | 0.647 | 0.471 | 2.60871E-13 |
| Contractile | PIAS1 | 3.99129E-76 | 0.790166 | 0.477 | 0.225 | 7.77303E-72 |
| Pericytes | TSC22D1 | 4.39402E-05 | 0.789847 | 0.54 | 0.329 | 0.855735244 |
| Fibroblasts | CYP27A1 | 4.87676E-21 | 0.788771 | 0.307 | 0.11 | 9.49749E-17 |
| Intermediate | RSPO3 | 2.01187E-44 | 0.788255 | 0.358 | 0.147 | 3.91812E-40 |
| Pericytes | EHD1 | 2.50875E-05 | 0.788156 | 0.38 | 0.181 | 0.488578548 |
| Fibroblasts | DEPTOR | 6.29929E-40 | 0.787388 | 0.252 | 0.049 | 1.22679E-35 |
| Fibroblasts | ZFP36L1 | 4.27524E-22 | 0.787128 | 0.83 | 0.673 | 8.32602E-18 |
| Fibroblasts | SPARCL1 | 3.04406E-13 | 0.786992 | 0.839 | 0.702 | 5.92831E-09 |
| Contractile | CCNG1 | 9.48312E-76 | 0.78698 | 0.576 | 0.322 | 1.84684E-71 |
| VEGFA_Terminal | PLOD2 | 3.32586E-36 | 0.786775 | 0.785 | 0.548 | 6.47712E-32 |
| Contractile | CNBP | 3.3048E-137 | 0.786764 | 0.929 | 0.811 | 6.4361E-133 |
| VEGFA_Terminal | COL6A1 | 2.72643E-48 | 0.785775 | 0.973 | 0.875 | 5.30972E-44 |
| VEGFA_Terminal | NFKBIA | 3.99359E-11 | 0.785655 | 0.571 | 0.443 | 7.77753E-07 |
| Intermediate | CHI3L1 | 5.091E-18 | 0.785059 | 0.676 | 0.528 | 9.91473E-14 |
| CD74_Intermediat | HLA-DRA | 3.9808E-129 | 1.058962 | 0.224 | 0.033 | 7.7527E-125 |
| Intermediate | CCN2 | 1.36142E-22 | 0.783764 | 0.791 | 0.679 | 2.65137E-18 |
| Contractile | ZBTB16 | 1.5562E-96 | 0.783641 | 0.556 | 0.25 | 3.0307E-92 |
| Pericytes | IRAG1 | 4.84234E-06 | 0.783165 | 0.54 | 0.29 | 0.094304652 |
| Contractile | EIF5B | 3.6146E-80 | 0.782727 | 0.773 | 0.564 | 7.03943E-76 |
| Intermediate | THBS1 | 2.94051E-39 | 0.782399 | 0.754 | 0.527 | 5.72665E-35 |
| Fibroblasts | SAT1 | 1.28787E-14 | 0.782162 | 0.807 | 0.677 | 2.50812E-10 |
| Pericytes | BAALC | 1.59656E-07 | 0.781811 | 0.16 | 0.031 | 0.00310931 |
| Pericytes | ENSSSCG0( | 0.00022546 | 0.780494 | 0.2 | 0.071 | 1 |
| Fibroblasts | VEGFD | 1.21496E-74 | 0.780464 | 0.202 | 0.016 | 2.36614E-70 |
| Pericytes | CDKN1A | 8.76148E-07 | 0.780062 | 0.34 | 0.122 | 0.017062975 |
| Intermediate | TPM1 | 1.85145E-56 | 0.780052 | 0.945 | 0.807 | 3.60571E-52 |
| Pericytes | CISH | 5.43758E-24 | 0.77977 | 0.2 | 0.015 | 1.05897E-19 |
| VEGFA_Terminal | F5 | 7.34073E-39 | 0.779707 | 0.613 | 0.323 | 1.42961E-34 |
| Fibroblasts | CD248 | 3.29814E-13 | 0.777154 | 0.284 | 0.13 | 6.42312E-09 |
| Pericytes | DAAM2 | 7.46286E-12 | 0.77689 | 0.24 | 0.044 | 1.45339E-07 |
| Terminal | TRPS1 | 5.46095E-95 | 0.776242 | 0.637 | 0.399 | 1.06352E-90 |
| Contractile | ENSSSCG0( | 1.1482E-115 | 0.776206 | 0.378 | 0.106 | 2.2362E-111 |
| Contractile | STOM | 1.84422E-77 | 0.775497 | 0.724 | 0.488 | 3.59161E-73 |
| VEGFA_Terminal | ENSSSCG0( | 2.17191E-52 | 0.774911 | 0.971 | 0.891 | 4.22979E-48 |
| Fibroblasts | GABARAPL | 4.69012E-18 | 0.774427 | 0.624 | 0.436 | 9.134E-14 |
| IFN_Intermediate | FAM111A | 5.45741E-11 | 0.773585 | 0.804 | 0.549 | 1.06283E-06 |
| VEGFA_Terminal | TNFRSF12A | 4.95843E-29 | 0.773136 | 0.7 | 0.46 | 9.65655E-25 |
| Fibroblasts | RND3 | 4.69627E-12 | 0.772453 | 0.417 | 0.251 | 9.14598E-08 |
| Fibroblasts | LRRC17 | 1.03503E-08 | 0.770728 | 0.528 | 0.41 | 0.000201572 |
| Contractile | CCND2 | 1.33526E-81 | 0.770593 | 0.749 | 0.502 | 2.60041E-77 |
| Fibroblasts | LARP6 | 7.62443E-16 | 0.769377 | 0.358 | 0.176 | 1.48486E-11 |
| Intermediate | LTBP2 | 1.08231E-25 | 0.769282 | 0.773 | 0.725 | 2.10781E-21 |
| Pericytes | CRISPLD2 | 5.52951E-06 | 0.768967 | 0.48 | 0.229 | 0.107687112 |
| Fibroblasts | CDC42EP2 | 1.15337E-35 | 0.768579 | 0.239 | 0.049 | 2.24619E-31 |
| CD74_Intermediat | NUDT4 | 2.73513E-93 | 0.768534 | 0.641 | 0.413 | 5.32667E-89 |

| | | | | | | |
|---|---|---|---|---|---|---|
| Pericytes | AXL | 0.000917849 | 0.768481 | 0.5 | 0.352 | 1 |
| Intermediate | TGM2 | 4.27422E-36 | 0.76813 | 0.708 | 0.515 | 8.32405E-32 |
| Pericytes | ADGRF5 | 1.61258E-56 | 0.767598 | 0.1 | 0.001 | 3.14051E-52 |
| Fibroblasts | ENSSSCG0( | 1.06291E-54 | 0.767172 | 0.128 | 0.009 | 2.07002E-50 |
| IFN_Intermediate | SAT1 | 1.05861E-06 | 0.763962 | 0.832 | 0.679 | 0.020616442 |
| Contractile | ANGPTL7 | 3.6876E-120 | 0.763379 | 0.275 | 0.051 | 7.1816E-116 |
| Fibroblasts | KLF4 | 6.0145E-07 | 0.763066 | 0.399 | 0.287 | 0.011713231 |
| Pericytes | ENSSSCG0( | 1.90219E-08 | 0.76304 | 0.24 | 0.056 | 0.000370452 |
| Contractile | CRIM1 | 2.63655E-48 | 0.762721 | 0.53 | 0.332 | 5.13467E-44 |
| Pericytes | MOB3B | 8.26037E-10 | 0.762475 | 0.2 | 0.037 | 1.60871E-05 |
| Fibroblasts | FAM180B | 1.79803E-36 | 0.76176 | 0.206 | 0.035 | 3.50166E-32 |
| VEGFA_Terminal | PGF | 1.11958E-37 | 0.761669 | 0.203 | 0.05 | 2.18039E-33 |
| Fibroblasts | OLFML3 | 1.4654E-09 | 0.760696 | 0.523 | 0.4 | 2.85386E-05 |
| Pericytes | LOC10051! | 0.000278425 | 0.760459 | 0.3 | 0.144 | 1 |
| Contractile | SEPTIN9 | 6.07026E-69 | 0.760071 | 0.575 | 0.33 | 1.18218E-64 |
| Fibroblasts | ZCCHC24 | 5.0237E-12 | 0.760013 | 0.615 | 0.527 | 9.78365E-08 |
| Terminal | WIF1 | 4.56488E-27 | 0.75998 | 0.488 | 0.381 | 8.89011E-23 |
| Fibroblasts | PLAC9 | 3.29485E-18 | 0.759559 | 0.803 | 0.68 | 6.41672E-14 |
| Fibroblasts | TGFBI | 9.53958E-07 | 0.759098 | 0.454 | 0.347 | 0.018578336 |
| IFN_Intermediate | NUB1 | 1.11911E-06 | 0.75848 | 0.57 | 0.373 | 0.021794578 |
| Fibroblasts | PDGFRL | 1.74003E-16 | 0.758156 | 0.665 | 0.49 | 3.38871E-12 |
| Fibroblasts | ADAMTS5 | 3.41408E-74 | 0.758097 | 0.151 | 0.008 | 6.64892E-70 |
| Terminal | MT1X | 2.39116E-18 | 0.758087 | 0.178 | 0.1 | 4.65678E-14 |
| Pericytes | ENSSSCG0( | 5.68059E-09 | 0.758043 | 0.26 | 0.064 | 0.000110629 |
| Terminal | STK38L | 2.77583E-91 | 0.757846 | 0.63 | 0.394 | 5.40592E-87 |
| Contractile | ALCAM | 5.59549E-70 | 0.757646 | 0.723 | 0.495 | 1.08972E-65 |
| Contractile | DNAJB4 | 5.33419E-71 | 0.757315 | 0.721 | 0.492 | 1.03883E-66 |
| VEGFA_Terminal | SARS1 | 1.44552E-31 | 0.757159 | 0.644 | 0.405 | 2.81514E-27 |
| Fibroblasts | P2RY1 | 6.57434E-11 | 0.757095 | 0.257 | 0.124 | 1.28035E-06 |
| Fibroblasts | RBMS3 | 1.88177E-12 | 0.756335 | 0.491 | 0.326 | 3.66476E-08 |
| Pericytes | PIM3 | 1.20215E-05 | 0.755667 | 0.26 | 0.092 | 0.234119474 |
| Pericytes | SYNPO2 | 4.40012E-05 | 0.75541 | 0.7 | 0.495 | 0.856922447 |
| VEGFA_Terminal | CAPG | 4.80094E-39 | 0.755327 | 0.867 | 0.673 | 9.34983E-35 |
| Contractile | MFAP4 | 1.3719E-108 | 0.7548 | 0.965 | 0.837 | 2.6719E-104 |
| Intermediate | NET1 | 1.55205E-37 | 0.754774 | 0.556 | 0.33 | 3.02261E-33 |
| IFN_Intermediate | CASP5 | 2.71614E-09 | 0.754511 | 0.449 | 0.228 | 5.28969E-05 |
| Fibroblasts | FOSB | 0.000210219 | 0.754291 | 0.349 | 0.254 | 1 |
| Contractile | GASK1B | 1.37824E-67 | 0.753579 | 0.675 | 0.433 | 2.68411E-63 |
| Contractile | GPR20 | 3.9843E-220 | 0.753164 | 0.249 | 0.011 | 7.7595E-216 |
| Pericytes | HNRNPAB | 3.93423E-05 | 0.752837 | 0.56 | 0.375 | 0.766192126 |
| Contractile | MAP7D1 | 1.65285E-79 | 0.752809 | 0.712 | 0.472 | 3.21892E-75 |
| Fibroblasts | SLC25A36 | 1.19702E-13 | 0.751451 | 0.528 | 0.358 | 2.3312E-09 |
| Fibroblasts | NOVA1 | 1.25096E-38 | 0.751314 | 0.22 | 0.039 | 2.43625E-34 |
| Fibroblasts | PTGER4 | 3.25967E-52 | 0.7505 | 0.188 | 0.021 | 6.34821E-48 |
| VEGFA_Terminal | COL4A1 | 1.39872E-12 | 0.750451 | 0.78 | 0.687 | 2.724E-08 |
| Contractile | RASSF2 | 3.62406E-87 | 0.748648 | 0.343 | 0.11 | 7.05786E-83 |
| Intermediate | PLK2 | 2.51084E-34 | 0.748495 | 0.292 | 0.124 | 4.88986E-30 |
| Fibroblasts | AKAP12 | 4.43836E-10 | 0.748442 | 0.436 | 0.296 | 8.6437E-06 |
| Fibroblasts | VGLL3 | 9.47218E-11 | 0.748409 | 0.289 | 0.15 | 1.84471E-06 |
| Pericytes | JAG1 | 0.000297864 | 0.747296 | 0.62 | 0.454 | 1 |
| CD74_Intermediat | PLXDC1 | 1.029E-106 | 0.746352 | 0.824 | 0.616 | 2.004E-102 |
| Fibroblasts | ENSSSCG0( | 3.67125E-15 | 0.745998 | 0.693 | 0.544 | 7.14976E-11 |
| VEGFA_Terminal | CD44 | 1.65889E-38 | 0.745868 | 0.661 | 0.356 | 3.23068E-34 |
| Fibroblasts | LAMB1 | 1.06282E-12 | 0.745384 | 0.739 | 0.646 | 2.06984E-08 |
| Pericytes | SELENOW | 0.00056374 | 0.745153 | 0.64 | 0.549 | 1 |
| Contractile | RNF150 | 9.0282E-188 | 0.744928 | 0.264 | 0.023 | 1.7582E-183 |
| Pericytes | SPRY4 | 7.61166E-05 | 0.744853 | 0.3 | 0.13 | 1 |
| Fibroblasts | FRMD6 | 6.48298E-14 | 0.744432 | 0.417 | 0.233 | 1.26256E-09 |
| VEGFA_Terminal | FCHSD2 | 2.15822E-47 | 0.744406 | 0.416 | 0.15 | 4.20314E-43 |

| | | | | | | |
|---|---|---|---|---|---|---|
| Pericytes | NEK4 | 2.21958E-06 | 0.742979 | 0.2 | 0.053 | 0.043226306 |
| Contractile | THRA | 3.34759E-83 | 0.742777 | 0.52 | 0.248 | 6.51942E-79 |
| IFN_Intermediate | TRIM25 | 2.29436E-13 | 0.741459 | 0.402 | 0.149 | 4.46827E-09 |
| CD74_Intermediat | ENC1 | 1.88215E-30 | 0.339584 | 0.105 | 0.033 | 3.66548E-26 |
| Pericytes | DLC1 | 0.002287489 | 0.740811 | 0.36 | 0.227 | 1 |
| CD74_Intermediat | BHMT2 | 4.60443E-67 | 0.412478 | 0.176 | 0.043 | 8.96713E-63 |
| Contractile | CSPG4 | 4.42947E-73 | 0.740387 | 0.511 | 0.253 | 8.6264E-69 |
| Pericytes | CH25H | 9.20715E-09 | 0.74023 | 0.24 | 0.056 | 0.000179309 |
| Fibroblasts | ENSSSCG0( | 3.30307E-24 | 0.739918 | 0.17 | 0.035 | 6.43274E-20 |
| Contractile | ACTN1 | 8.6245E-115 | 0.739536 | 0.974 | 0.847 | 1.6796E-110 |
| Pericytes | PNRC1 | 0.001395416 | 0.73751 | 0.72 | 0.639 | 1 |
| Contractile | SH3BGRL | 2.29907E-84 | 0.736726 | 0.772 | 0.559 | 4.47744E-80 |
| VEGFA_Terminal | MTURN | 1.02263E-46 | 0.736194 | 0.557 | 0.242 | 1.99158E-42 |
| Fibroblasts | PTGFR | 4.6493E-40 | 0.736041 | 0.216 | 0.036 | 9.05452E-36 |
| Fibroblasts | KLF9 | 1.02986E-08 | 0.735142 | 0.445 | 0.318 | 0.000200566 |
| VEGFA_Terminal | HSBP1 | 1.39846E-50 | 0.734847 | 0.935 | 0.77 | 2.7235E-46 |
| Terminal | MEOX1 | 3.74266E-90 | 0.734846 | 0.231 | 0.059 | 7.28882E-86 |
| Pericytes | MAFF | 0.001127927 | 0.734553 | 0.42 | 0.276 | 1 |
| Pericytes | NOSTRIN | 0.001847059 | 0.734351 | 0.3 | 0.163 | 1 |
| IFN_Intermediate | TAP2 | 1.46861E-42 | 0.734346 | 0.346 | 0.048 | 2.86011E-38 |
| Pericytes | AMOTL2 | 8.88388E-05 | 0.734195 | 0.32 | 0.143 | 1 |
| VEGFA_Terminal | LTC4S | 2.69194E-41 | 0.733928 | 0.901 | 0.687 | 5.24255E-37 |
| Intermediate | SOD3 | 8.23713E-54 | 0.733792 | 0.911 | 0.795 | 1.60418E-49 |
| Pericytes | CACNA2D1 | 0.000587298 | 0.733692 | 0.46 | 0.296 | 1 |
| Pericytes | ENSSSCG0( | 0.000966942 | 0.733436 | 0.34 | 0.188 | 1 |
| Pericytes | PTH1R | 1.19297E-06 | 0.733223 | 0.26 | 0.08 | 0.023233027 |
| IFN_Intermediate | CNP | 1.7623E-17 | 0.733191 | 0.327 | 0.09 | 3.43209E-13 |
| Pericytes | SFPQ | 1.89743E-05 | 0.732967 | 0.66 | 0.492 | 0.369523948 |
| Contractile | PHLDB2 | 2.66023E-69 | 0.732501 | 0.472 | 0.228 | 5.18079E-65 |
| Fibroblasts | LAMA4 | 5.63089E-09 | 0.732354 | 0.541 | 0.428 | 0.000109662 |
| Intermediate | ENSSSCG0( | 3.82767E-64 | 0.731484 | 0.959 | 0.895 | 7.45439E-60 |
| CD74_Intermediat | KLF5 | 1.59524E-28 | 0.273445 | 0.136 | 0.053 | 3.10673E-24 |
| IFN_Intermediate | GBP2 | 1.17863E-28 | 0.728817 | 0.187 | 0.022 | 2.29539E-24 |
| Contractile | GLDN | 8.1442E-198 | 0.728615 | 0.229 | 0.011 | 1.5861E-193 |
| Fibroblasts | ALDH1A3 | 1.76156E-10 | 0.728356 | 0.307 | 0.162 | 3.43065E-06 |
| Fibroblasts | SPRY1 | 1.03147E-09 | 0.727963 | 0.647 | 0.532 | 2.00879E-05 |
| Terminal | TPPP3 | 5.81377E-44 | 0.726517 | 0.345 | 0.191 | 1.13223E-39 |
| CD74_Intermediat | CTSB | 2.2399E-103 | 0.725696 | 0.882 | 0.728 | 4.3622E-99 |
| Terminal | CCND3 | 9.5335E-108 | 0.725621 | 0.671 | 0.425 | 1.8566E-103 |
| Terminal | NT5E | 9.828E-131 | 0.725299 | 0.398 | 0.129 | 1.914E-126 |
| Pericytes | PVR | 3.45365E-06 | 0.724829 | 0.18 | 0.045 | 0.067259855 |
| VEGFA_Terminal | GARS1 | 4.63041E-32 | 0.724691 | 0.538 | 0.279 | 9.01771E-28 |
| Pericytes | STX11 | 0.000220992 | 0.724612 | 0.24 | 0.095 | 1 |
| Contractile | INPP5A | 3.0254E-101 | 0.724252 | 0.337 | 0.095 | 5.892E-97 |
| VEGFA_Terminal | IGSF23 | 1.5428E-100 | 0.722896 | 0.288 | 0.04 | 3.00455E-96 |
| Contractile | KCNA5 | 1.1367E-162 | 0.722389 | 0.276 | 0.033 | 2.2138E-158 |
| Pericytes | CCDC9B | 5.40291E-05 | 0.722354 | 0.22 | 0.075 | 1 |
| Terminal | COL5A2 | 5.3862E-112 | 0.7218 | 0.875 | 0.706 | 1.049E-107 |
| CD74_Intermediat | ETV1 | 3.3011E-107 | 0.721469 | 0.671 | 0.397 | 6.4289E-103 |
| Contractile | RRAGD | 1.3798E-134 | 0.720473 | 0.289 | 0.05 | 2.6871E-130 |
| Contractile | HACD1 | 1.69257E-79 | 0.72017 | 0.433 | 0.183 | 3.29627E-75 |
| Fibroblasts | DPYSL3 | 3.70769E-11 | 0.719753 | 0.468 | 0.3 | 7.22073E-07 |
| Contractile | SGCA | 2.92026E-87 | 0.719665 | 0.38 | 0.133 | 5.6872E-83 |
| VEGFA_Terminal | EIF4EBP1 | 8.65319E-41 | 0.718854 | 0.484 | 0.209 | 1.68521E-36 |
| Intermediate | IGFBP7 | 1.58313E-95 | 0.718309 | 1 | 0.999 | 3.08315E-91 |
| Contractile | KIAA2026 | 5.05598E-74 | 0.71823 | 0.531 | 0.274 | 9.84652E-70 |
| Terminal | EMP3 | 3.1181E-135 | 0.717907 | 0.922 | 0.786 | 6.0726E-131 |
| VEGFA_Terminal | IL18 | 9.8528E-44 | 0.717832 | 0.465 | 0.185 | 1.91883E-39 |
| Contractile | FRZB | 2.35083E-98 | 0.71691 | 0.487 | 0.18 | 4.57825E-94 |

| | | | | | | |
|---|---|---|---|---|---|---|
| Contractile | EIF4A2 | 1.20944E-92 | 0.716854 | 0.916 | 0.78 | 2.35538E-88 |
| Pericytes | PAPLN | 0.001491844 | 0.716844 | 0.22 | 0.097 | 1 |
| Pericytes | PEAK1 | 0.008569611 | 0.716783 | 0.32 | 0.215 | 1 |
| CD74_Intermediat | PLPP3 | 1.7937E-112 | 0.715954 | 0.971 | 0.877 | 3.4933E-108 |
| Fibroblasts | PROCR | 5.4323E-07 | 0.715787 | 0.202 | 0.104 | 0.010579408 |
| Fibroblasts | NR2F2 | 3.90248E-05 | 0.714748 | 0.376 | 0.291 | 0.76000789 |
| Contractile | DYNLL2 | 2.5526E-68 | 0.714384 | 0.573 | 0.336 | 4.97118E-64 |
| VEGFA_Terminal | ENSSSCG0( | 2.83123E-37 | 0.714373 | 0.562 | 0.277 | 5.51383E-33 |
| Contractile | ATOH8 | 1.5217E-113 | 0.714114 | 0.299 | 0.066 | 2.9634E-109 |
| Pericytes | RNF149 | 5.67594E-05 | 0.714098 | 0.4 | 0.203 | 1 |
| Pericytes | TNFSF13 | 0.000241772 | 0.713506 | 0.2 | 0.072 | 1 |
| VEGFA_Terminal | PDGFA | 1.87661E-48 | 0.712748 | 0.499 | 0.196 | 3.65469E-44 |
| Pericytes | PABPC1 | 0.001187889 | 0.71226 | 0.76 | 0.765 | 1 |
| VEGFA_Terminal | VEGFC | 3.85814E-82 | 0.711682 | 0.363 | 0.076 | 7.51374E-78 |
| Pericytes | FERMT2 | 4.15661E-06 | 0.710267 | 0.86 | 0.73 | 0.080949895 |
| Pericytes | NAALAD2 | 5.56677E-33 | 0.709702 | 0.18 | 0.009 | 1.08413E-28 |
| Contractile | UBC | 1.9885E-111 | 0.709523 | 0.982 | 0.95 | 3.8727E-107 |
| Fibroblasts | ARGLU1 | 5.96035E-18 | 0.709244 | 0.748 | 0.583 | 1.16078E-13 |
| Terminal | FOLR3 | 3.0719E-110 | 0.709092 | 0.463 | 0.199 | 5.9825E-106 |
| Pericytes | ANGPT2 | 0.000104958 | 0.709076 | 0.3 | 0.129 | 1 |
| Pericytes | PCP4L1 | 6.89165E-08 | 0.708283 | 0.18 | 0.037 | 0.001342149 |
| IFN_Intermediate | C1R | 2.97102E-12 | 0.707929 | 0.963 | 0.88 | 5.78606E-08 |
| Fibroblasts | GLUL | 2.82313E-06 | 0.707557 | 0.468 | 0.387 | 0.054980482 |
| Pericytes | PLCL1 | 0.000685328 | 0.706618 | 0.2 | 0.078 | 1 |
| Terminal | EZR | 4.45219E-82 | 0.706365 | 0.568 | 0.339 | 8.67064E-78 |
| Contractile | ADAMTSL5 | 2.04116E-68 | 0.706243 | 0.486 | 0.24 | 3.97516E-64 |
| IFN_Intermediate | IFI30 | 4.27628E-05 | 0.706006 | 0.57 | 0.416 | 0.832805587 |
| Pericytes | NID2 | 0.005277142 | 0.705121 | 0.34 | 0.21 | 1 |
| VEGFA_Terminal | ALPK3 | 4.9158E-71 | 0.705082 | 0.339 | 0.077 | 9.57353E-67 |
| Contractile | REV1 | 2.19141E-75 | 0.704992 | 0.363 | 0.136 | 4.26778E-71 |
| Terminal | COL5A1 | 2.28439E-81 | 0.704662 | 0.694 | 0.48 | 4.44885E-77 |
| Intermediate | ITGA8 | 1.62958E-64 | 0.704065 | 0.806 | 0.499 | 3.1736E-60 |
| Intermediate | TAGLN | 2.00369E-51 | 0.703631 | 0.997 | 0.959 | 3.90218E-47 |
| CD74_Intermediat | EPHA7 | 2.91502E-44 | 0.333329 | 0.165 | 0.055 | 5.67699E-40 |
| Contractile | ADGRL1 | 1.5789E-161 | 0.701523 | 0.263 | 0.03 | 3.0749E-157 |
| Contractile | PRKG1 | 4.59469E-62 | 0.701204 | 0.655 | 0.421 | 8.94817E-58 |
| Contractile | VIM | 2.8222E-184 | 0.700595 | 1 | 1 | 5.4963E-180 |
| Intermediate | SOST | 3.02673E-37 | 0.700493 | 0.203 | 0.064 | 5.89456E-33 |
| VEGFA_Terminal | EIF1 | 1.61633E-54 | 0.699334 | 0.998 | 0.985 | 3.14781E-50 |
| Contractile | IL1RL1 | 2.48572E-49 | 0.699068 | 0.363 | 0.158 | 4.84093E-45 |
| Contractile | CDS2 | 1.47346E-86 | 0.698932 | 0.303 | 0.087 | 2.86957E-82 |
| Contractile | COX6C | 7.2077E-113 | 0.698219 | 0.933 | 0.865 | 1.4037E-108 |
| Contractile | PRKACA | 1.61887E-70 | 0.697544 | 0.45 | 0.213 | 3.15275E-66 |
| Fibroblasts | ACER3 | 1.23832E-15 | 0.697472 | 0.275 | 0.113 | 2.41163E-11 |
| Pericytes | PPP1R10 | 8.73725E-05 | 0.69712 | 0.34 | 0.16 | 1 |
| VEGFA_Terminal | PLP2 | 2.86409E-41 | 0.696667 | 0.91 | 0.734 | 5.57781E-37 |
| CD74_Intermediat | SAT1 | 2.6944E-109 | 0.696375 | 0.846 | 0.617 | 5.2473E-105 |
| Pericytes | P2RY14 | 1.48293E-29 | 0.695366 | 0.14 | 0.006 | 2.88801E-25 |
| Fibroblasts | DKK2 | 2.0529E-152 | 0.695335 | 0.128 | 0.001 | 3.9979E-148 |
| Contractile | SLC2A13 | 6.82736E-50 | 0.695254 | 0.468 | 0.269 | 1.32963E-45 |
| Contractile | DIAPH2 | 8.2725E-57 | 0.695191 | 0.613 | 0.394 | 1.61107E-52 |
| Contractile | CDC42EP3 | 1.4481E-65 | 0.694855 | 0.639 | 0.392 | 2.82017E-61 |
| Terminal | PALMD | 1.4804E-62 | 0.6948 | 0.483 | 0.28 | 2.88307E-58 |
| Pericytes | HNRNPUL1 | 0.001055563 | 0.694717 | 0.48 | 0.328 | 1 |
| Pericytes | TPM4 | 1.36125E-05 | 0.694182 | 0.94 | 0.954 | 0.26510272 |
| Fibroblasts | RNF217 | 7.52218E-11 | 0.693608 | 0.317 | 0.172 | 1.46495E-06 |
| Intermediate | PDLIM7 | 2.82273E-44 | 0.693248 | 0.713 | 0.482 | 5.49726E-40 |
| Contractile | PI15 | 1.22152E-76 | 0.692758 | 0.272 | 0.076 | 2.37891E-72 |
| Contractile | ENTPD3 | 6.0371E-224 | 0.692295 | 0.223 | 0.005 | 1.1757E-219 |

| | | | | | | |
|---|---|---|---|---|---|---|
| CD74_Intermediat | SFRP1 | 2.1992E-119 | 0.692133 | 0.943 | 0.772 | 4.2829E-115 |
| Fibroblasts | GLT8D2 | 6.81279E-13 | 0.691235 | 0.514 | 0.35 | 1.32679E-08 |
| Pericytes | ENSSSCG0( | 6.11191E-10 | 0.691149 | 0.2 | 0.035 | 1.19029E-05 |
| Contractile | TEAD1 | 1.52108E-56 | 0.691136 | 0.474 | 0.261 | 2.96231E-52 |
| IFN_Intermediate | LOC10651( | 6.19528E-28 | 0.690371 | 0.346 | 0.07 | 1.20653E-23 |
| Intermediate | RASL11A | 7.14215E-31 | 0.690227 | 0.445 | 0.256 | 1.39093E-26 |
| Contractile | C5ORF24 | 1.17981E-61 | 0.689469 | 0.667 | 0.469 | 2.29768E-57 |
| Contractile | PTK2B | 1.11087E-71 | 0.688809 | 0.364 | 0.14 | 2.16343E-67 |
| Contractile | CCDC107 | 1.11726E-82 | 0.688668 | 0.353 | 0.123 | 2.17586E-78 |
| Fibroblasts | MBTPS1 | 3.55897E-17 | 0.688644 | 0.61 | 0.413 | 6.93109E-13 |
| Pericytes | SGIP1 | 1.52549E-16 | 0.687143 | 0.22 | 0.027 | 2.97088E-12 |
| Pericytes | ITM2B | 5.98366E-07 | 0.687057 | 0.98 | 0.973 | 0.011653172 |
| Contractile | ENSSSCG0( | 2.10047E-65 | 0.686737 | 0.69 | 0.472 | 4.09066E-61 |
| Pericytes | MGST1 | 0.005591306 | 0.686338 | 0.38 | 0.242 | 1 |
| Fibroblasts | SRSF7 | 3.27855E-09 | 0.685556 | 0.587 | 0.473 | 6.38498E-05 |
| VEGFA_Terminal | FN1 | 7.25937E-40 | 0.685262 | 0.998 | 0.991 | 1.41376E-35 |
| VEGFA_Terminal | COX7C | 7.84145E-47 | 0.684981 | 0.947 | 0.844 | 1.52712E-42 |
| VEGFA_Terminal | ANKRD37 | 8.2168E-33 | 0.68456 | 0.337 | 0.128 | 1.60022E-28 |
| Terminal | CTHRC1 | 2.88588E-62 | 0.684203 | 0.536 | 0.323 | 5.62025E-58 |
| Contractile | RASL11A | 8.1476E-76 | 0.683706 | 0.503 | 0.232 | 1.58674E-71 |
| Pericytes | TNFAIP3 | 6.28708E-07 | 0.683196 | 0.24 | 0.067 | 0.012244081 |
| VEGFA_Terminal | PRRX2 | 4.31865E-37 | 0.682918 | 0.91 | 0.759 | 8.41058E-33 |
| Fibroblasts | TMEM100 | 0.001427357 | 0.682287 | 0.202 | 0.14 | 1 |
| Pericytes | MARCKS | 0.006386665 | 0.681845 | 0.24 | 0.122 | 1 |
| Terminal | RFLNB | 1.4796E-121 | 0.68158 | 0.295 | 0.073 | 2.8816E-117 |
| IFN_Intermediate | PARP12 | 2.7083E-26 | 0.681554 | 0.234 | 0.035 | 5.27442E-22 |
| Contractile | ENO3 | 5.3063E-175 | 0.681386 | 0.241 | 0.019 | 1.0334E-170 |
| Contractile | SLMAP | 2.80622E-59 | 0.681347 | 0.442 | 0.225 | 5.46512E-55 |
| VEGFA_Terminal | WSB1 | 3.1377E-18 | 0.680258 | 0.731 | 0.554 | 6.11066E-14 |
| Fibroblasts | COL15A1 | 3.12863E-09 | 0.680098 | 0.697 | 0.628 | 6.09302E-05 |
| Fibroblasts | CYP1B1 | 1.38076E-09 | 0.679987 | 0.275 | 0.141 | 2.68903E-05 |
| Fibroblasts | ENSSSCG0( | 1.39525E-10 | 0.679477 | 0.445 | 0.301 | 2.71726E-06 |
| Fibroblasts | ATP2B1 | 1.59277E-09 | 0.679054 | 0.394 | 0.255 | 3.10193E-05 |
| Fibroblasts | RBP1 | 1.04377E-92 | 0.679054 | 0.183 | 0.01 | 2.03274E-88 |
| Contractile | BMPR1A | 2.39338E-56 | 0.679035 | 0.534 | 0.317 | 4.6611E-52 |
| Terminal | ANXA2 | 1.2149E-120 | 0.678934 | 0.964 | 0.907 | 2.3661E-116 |
| Fibroblasts | NFIA | 4.7643E-13 | 0.67875 | 0.674 | 0.549 | 9.27848E-09 |
| CD74_Intermediat | PDGFD | 1.69671E-97 | 0.678499 | 0.67 | 0.394 | 3.30434E-93 |
| Pericytes | TNFRSF21 | 5.35819E-06 | 0.67837 | 0.2 | 0.056 | 0.104350694 |
| Intermediate | GAS6 | 8.99089E-40 | 0.678219 | 0.73 | 0.536 | 1.75098E-35 |
| Contractile | MBNL1 | 6.43896E-69 | 0.677933 | 0.852 | 0.718 | 1.25399E-64 |
| Contractile | ZNF503 | 2.38742E-58 | 0.677799 | 0.477 | 0.255 | 4.6495E-54 |
| Pericytes | FAM241A | 5.63065E-10 | 0.677371 | 0.24 | 0.051 | 1.09657E-05 |
| Contractile | WDR1 | 5.72408E-69 | 0.676795 | 0.669 | 0.439 | 1.11476E-64 |
| Fibroblasts | ARRDC3 | 1.17657E-07 | 0.675953 | 0.257 | 0.144 | 0.002291376 |
| Contractile | OAT | 5.25987E-83 | 0.675545 | 0.851 | 0.693 | 1.02436E-78 |
| Contractile | DENND2B | 4.97517E-70 | 0.675439 | 0.379 | 0.157 | 9.68915E-66 |
| Fibroblasts | NR3C1 | 4.04183E-15 | 0.675051 | 0.619 | 0.442 | 7.87146E-11 |
| Contractile | ENSSSCG0( | 1.42713E-35 | 0.674688 | 0.488 | 0.325 | 2.77934E-31 |
| Contractile | ROCK2 | 8.98233E-59 | 0.673854 | 0.696 | 0.503 | 1.74931E-54 |
| Terminal | CLEC11A | 1.52838E-90 | 0.673346 | 0.571 | 0.311 | 2.97651E-86 |
| CD74_Intermediat | COL14A1 | 2.8267E-130 | 0.673339 | 0.986 | 0.742 | 5.505E-126 |
| Intermediate | PCDH7 | 2.73277E-34 | 0.673162 | 0.618 | 0.401 | 5.32206E-30 |
| Contractile | SETD3 | 8.61801E-58 | 0.672974 | 0.524 | 0.303 | 1.67836E-53 |
| Pericytes | TMEM150 | 0.008037816 | 0.672962 | 0.32 | 0.2 | 1 |
| Pericytes | PARM1 | 1.33272E-59 | 0.672894 | 0.18 | 0.004 | 2.59548E-55 |
| Pericytes | CAV1 | 0.000245186 | 0.672633 | 0.34 | 0.162 | 1 |
| IFN_Intermediate | PSME2 | 1.95438E-10 | 0.672258 | 0.495 | 0.246 | 3.80616E-06 |
| CD74_Intermediat | PITPNC1 | 1.75626E-32 | 0.294748 | 0.147 | 0.056 | 3.42031E-28 |

| | | | | | | |
|---|---|---|---|---|---|---|
| Contractile | SOST | 4.8235E-102 | 0.670509 | 0.246 | 0.046 | 9.39371E-98 |
| VEGFA_Terminal | STBD1 | 4.30781E-33 | 0.670378 | 0.489 | 0.24 | 8.38946E-29 |
| Contractile | DAAM1 | 1.88887E-57 | 0.670217 | 0.725 | 0.521 | 3.67857E-53 |
| Pericytes | RASL12 | 4.36725E-09 | 0.669478 | 0.4 | 0.122 | 8.50522E-05 |
| Contractile | AKAP6 | 3.7489E-125 | 0.669282 | 0.264 | 0.044 | 7.3011E-121 |
| Contractile | MACF1 | 2.58338E-49 | 0.668957 | 0.56 | 0.352 | 5.03113E-45 |
| Fibroblasts | ENSSSCG0( | 7.12151E-09 | 0.668596 | 0.491 | 0.362 | 0.000138691 |
| Intermediate | ISG12(A) | 7.13528E-39 | 0.668417 | 0.831 | 0.637 | 1.3896E-34 |
| Contractile | ENSSSCG0( | 9.20479E-65 | 0.667652 | 0.345 | 0.138 | 1.79263E-60 |
| Contractile | CAV2 | 1.65026E-57 | 0.66734 | 0.586 | 0.369 | 3.21388E-53 |
| Pericytes | RND1 | 2.42468E-09 | 0.666312 | 0.2 | 0.037 | 4.72206E-05 |
| Pericytes | APBB2 | 0.000532917 | 0.666273 | 0.28 | 0.132 | 1 |
| Pericytes | ENSSSCG0( | 7.80913E-09 | 0.666041 | 0.98 | 0.998 | 0.000152083 |
| Fibroblasts | S1PR2 | 3.49163E-22 | 0.665859 | 0.206 | 0.055 | 6.79995E-18 |
| Contractile | HSPG2 | 4.33804E-61 | 0.665101 | 0.759 | 0.57 | 8.44833E-57 |
| Terminal | C1QTNF5 | 7.79549E-93 | 0.664551 | 0.75 | 0.522 | 1.51817E-88 |
| Contractile | ENSSSCG0( | 1.5601E-109 | 0.664393 | 0.271 | 0.055 | 3.0383E-105 |
| Pericytes | TRPC6 | 4.53116E-13 | 0.664047 | 0.26 | 0.046 | 8.82444E-09 |
| Pericytes | ENSSSCG0( | 1.04563E-19 | 0.66292 | 0.1 | 0.005 | 2.03636E-15 |
| Fibroblasts | FGL2 | 1.67547E-17 | 0.662836 | 0.89 | 0.694 | 3.26298E-13 |
| CD74_Intermediat | CCN3 | 3.97596E-62 | 0.662115 | 0.82 | 0.638 | 7.74317E-58 |
| IFN_Intermediate | C2 | 1.74642E-11 | 0.661298 | 0.393 | 0.16 | 3.40116E-07 |
| Intermediate | C1S | 1.92677E-56 | 0.660932 | 0.916 | 0.738 | 3.75239E-52 |
| IFN_Intermediate | IFIT3 | 8.37644E-63 | 0.660405 | 0.178 | 0.008 | 1.63131E-58 |
| CD74_Intermediat | TNXB | 1.2553E-100 | 0.660188 | 0.762 | 0.465 | 2.44467E-96 |
| Fibroblasts | RAP2A | 4.13047E-10 | 0.660034 | 0.339 | 0.2 | 8.0441E-06 |
| Fibroblasts | GOLIM4 | 8.7554E-11 | 0.659648 | 0.716 | 0.633 | 1.70511E-06 |
| Terminal | ID1 | 6.21328E-37 | 0.658414 | 0.625 | 0.484 | 1.21004E-32 |
| Intermediate | TTLL7 | 8.44125E-28 | 0.658113 | 0.605 | 0.43 | 1.64393E-23 |
| Pericytes | PTP4A3 | 1.99463E-18 | 0.656128 | 0.28 | 0.038 | 3.88453E-14 |
| Terminal | LGALS3 | 1.7927E-140 | 0.656123 | 0.961 | 0.705 | 3.4913E-136 |
| Fibroblasts | AXL | 4.0279E-10 | 0.655946 | 0.472 | 0.348 | 7.84434E-06 |
| VEGFA_Terminal | CTSC | 5.15081E-16 | 0.65528 | 0.504 | 0.322 | 1.00312E-11 |
| Pericytes | C1ORF54 | 0.000168987 | 0.654242 | 0.4 | 0.214 | 1 |
| Contractile | CAPZA2 | 1.83919E-79 | 0.654069 | 0.858 | 0.714 | 3.58182E-75 |
| Intermediate | TNFRSF11E | 2.0985E-15 | 0.653815 | 0.386 | 0.253 | 4.08683E-11 |
| Contractile | GRSF1 | 4.24316E-54 | 0.653776 | 0.548 | 0.338 | 8.26355E-50 |
| VEGFA_Terminal | VLDLR | 2.95736E-35 | 0.653565 | 0.751 | 0.463 | 5.75946E-31 |
| Fibroblasts | ECE1 | 1.77808E-20 | 0.653111 | 0.197 | 0.053 | 3.46281E-16 |
| Fibroblasts | IER2 | 7.09757E-07 | 0.652654 | 0.404 | 0.292 | 0.01382252 |
| Contractile | PLCL1 | 1.3677E-125 | 0.652382 | 0.263 | 0.043 | 2.6636E-121 |
| Terminal | ENSSSCG0( | 2.1539E-125 | 0.652345 | 0.953 | 0.87 | 4.1947E-121 |
| VEGFA_Terminal | INHBA | 5.28045E-45 | 0.652313 | 0.392 | 0.136 | 1.02837E-40 |
| VEGFA_Terminal | ITGA5 | 5.37264E-33 | 0.652248 | 0.62 | 0.332 | 1.04632E-28 |
| Contractile | FOXO3 | 2.09953E-52 | 0.651425 | 0.694 | 0.497 | 4.08883E-48 |
| Contractile | CSDE1 | 8.31922E-67 | 0.650169 | 0.807 | 0.651 | 1.62017E-62 |
| Contractile | RRAS | 5.16828E-79 | 0.650122 | 0.871 | 0.697 | 1.00652E-74 |
| Pericytes | CRTC3 | 0.001666181 | 0.649606 | 0.36 | 0.218 | 1 |
| Fibroblasts | NFIB | 1.06203E-15 | 0.648604 | 0.739 | 0.574 | 2.06831E-11 |
| VEGFA_Terminal | FAM111A | 4.41188E-31 | 0.648072 | 0.792 | 0.536 | 8.59213E-27 |
| VEGFA_Terminal | WLS | 6.84955E-33 | 0.646831 | 0.412 | 0.176 | 1.33395E-28 |
| Fibroblasts | FCGRT | 1.80196E-13 | 0.646731 | 0.528 | 0.355 | 3.50932E-09 |
| Fibroblasts | GGT5 | 1.38145E-14 | 0.646643 | 0.193 | 0.063 | 2.69037E-10 |
| Intermediate | CALD1 | 1.91975E-54 | 0.645422 | 0.994 | 0.965 | 3.7387E-50 |
| Contractile | ZFAND5 | 1.77834E-46 | 0.645314 | 0.577 | 0.383 | 3.46331E-42 |
| Pericytes | ADAMTS12 | 0.009408074 | 0.644772 | 0.22 | 0.113 | 1 |
| Terminal | THBS1 | 4.22882E-50 | 0.644699 | 0.687 | 0.486 | 8.23563E-46 |
| VEGFA_Terminal | RHOD | 1.02451E-40 | 0.644489 | 0.581 | 0.281 | 1.99523E-36 |
| VEGFA_Terminal | TAGLN2 | 2.97546E-38 | 0.644354 | 0.886 | 0.753 | 5.79472E-34 |

| | | | | | | |
|---|---|---|---|---|---|---|
| Intermediate | ARID5B | 6.58496E-31 | 0.644112 | 0.601 | 0.413 | 1.28242E-26 |
| VEGFA_Terminal | TMSB10 | 3.96853E-43 | 0.643996 | 1 | 0.984 | 7.72872E-39 |
| Terminal | SDCBP | 7.4497E-101 | 0.643903 | 0.811 | 0.623 | 1.45082E-96 |
| Contractile | LHFPL6 | 1.08053E-72 | 0.643867 | 0.863 | 0.704 | 2.10433E-68 |
| CD74_Intermediat | ISLR | 2.40532E-86 | 0.643402 | 0.705 | 0.452 | 4.68436E-82 |
| Pericytes | NFKBIB | 0.000255136 | 0.643396 | 0.26 | 0.111 | 1 |
| Contractile | NDUFA4 | 4.74032E-87 | 0.643084 | 0.926 | 0.871 | 9.23177E-83 |
| Pericytes | PTGER1 | 4.49424E-42 | 0.642694 | 0.12 | 0.003 | 8.75254E-38 |
| Pericytes | TMEM182 | 3.6279E-101 | 0.642129 | 0.16 | 0.001 | 7.06533E-97 |
| Pericytes | HNRNPA2B | 1.85479E-06 | 0.641993 | 0.94 | 0.898 | 0.036122118 |
| Terminal | PCOLCE2 | 3.15363E-75 | 0.641077 | 0.415 | 0.197 | 6.1417E-71 |
| Pericytes | TP53I11 | 3.35963E-05 | 0.640836 | 0.12 | 0.026 | 0.654287023 |
| Pericytes | CFAP210 | 1.46445E-05 | 0.640813 | 0.18 | 0.048 | 0.285201956 |
| Contractile | CAB39 | 4.58576E-58 | 0.640694 | 0.412 | 0.199 | 8.93077E-54 |
| Pericytes | HIC1 | 6.52558E-05 | 0.640589 | 0.2 | 0.065 | 1 |
| Contractile | MGST2 | 1.52124E-75 | 0.640036 | 0.342 | 0.123 | 2.96262E-71 |
| Intermediate | OSR1 | 3.05725E-22 | 0.639763 | 0.505 | 0.35 | 5.95399E-18 |
| Fibroblasts | BMP4 | 2.65226E-10 | 0.63959 | 0.275 | 0.138 | 5.16528E-06 |
| VEGFA_Terminal | LGMN | 7.11498E-24 | 0.639392 | 0.564 | 0.341 | 1.38564E-19 |
| Fibroblasts | NDRG2 | 9.14199E-10 | 0.639346 | 0.468 | 0.332 | 1.7804E-05 |
| Fibroblasts | ENSSSCG0( | 1.21349E-11 | 0.639222 | 0.583 | 0.434 | 2.36327E-07 |
| Contractile | LSM6 | 1.61954E-46 | 0.639011 | 0.479 | 0.289 | 3.15406E-42 |
| Intermediate | CCL19 | 1.35462E-35 | 0.638847 | 0.369 | 0.171 | 2.63812E-31 |
| Contractile | ATP8B1 | 4.79768E-89 | 0.63864 | 0.317 | 0.09 | 9.34348E-85 |
| Intermediate | SYTL2 | 2.38807E-40 | 0.638129 | 0.301 | 0.119 | 4.65077E-36 |
| Terminal | PCSK1N | 4.12432E-71 | 0.637993 | 0.803 | 0.67 | 8.03212E-67 |
| VEGFA_Terminal | SNX9 | 9.61503E-25 | 0.637559 | 0.758 | 0.532 | 1.87253E-20 |
| Fibroblasts | CRYBG3 | 1.40303E-08 | 0.63733 | 0.22 | 0.106 | 0.00027324 |
| Fibroblasts | IGF1 | 2.98612E-60 | 0.637325 | 0.119 | 0.006 | 5.81546E-56 |
| Fibroblasts | ENSSSCG0( | 3.47898E-16 | 0.637241 | 0.853 | 0.71 | 6.77531E-12 |
| CD74_Intermediat | SLPI | 6.66218E-50 | 0.702839 | 0.188 | 0.064 | 1.29746E-45 |
| Pericytes | ARHGAP1 | 0.00036145 | 0.636859 | 0.44 | 0.255 | 1 |
| VEGFA_Terminal | AP3S1 | 1.38822E-38 | 0.636747 | 0.738 | 0.433 | 2.70356E-34 |
| Intermediate | PTGS1 | 9.09798E-44 | 0.636591 | 0.303 | 0.113 | 1.77183E-39 |
| VEGFA_Terminal | ANXA5 | 1.79699E-56 | 0.636486 | 0.993 | 0.942 | 3.49964E-52 |
| Pericytes | ENSSSCG0( | 7.14301E-05 | 0.635732 | 0.14 | 0.036 | 1 |
| Contractile | PPP1CB | 1.70052E-56 | 0.635684 | 0.689 | 0.493 | 3.31176E-52 |
| Pericytes | BAZ1A | 0.000127717 | 0.635617 | 0.22 | 0.08 | 1 |
| VEGFA_Terminal | PSAT1 | 9.66342E-53 | 0.635048 | 0.23 | 0.047 | 1.88195E-48 |
| Contractile | FBXO30 | 2.04114E-49 | 0.634932 | 0.494 | 0.292 | 3.97511E-45 |
| Intermediate | PRKG1 | 1.35026E-33 | 0.634759 | 0.647 | 0.437 | 2.62964E-29 |
| Pericytes | TDG | 0.000132872 | 0.634532 | 0.26 | 0.106 | 1 |
| VEGFA_Terminal | SSR3 | 3.68386E-31 | 0.634523 | 0.797 | 0.583 | 7.17432E-27 |
| Pericytes | OSTF1 | 0.007499684 | 0.634089 | 0.56 | 0.484 | 1 |
| Pericytes | LOC10062( | 0.0028052 | 0.633869 | 0.14 | 0.05 | 1 |
| Contractile | VWA1 | 5.1623E-105 | 0.632489 | 0.265 | 0.055 | 1.0054E-100 |
| CD74_Intermediat | MAMDC2 | 3.24392E-76 | 0.632045 | 0.657 | 0.432 | 6.31754E-72 |
| VEGFA_Terminal | DNASE1L1 | 2.55972E-31 | 0.631932 | 0.613 | 0.351 | 4.98506E-27 |
| VEGFA_Terminal | ZNF395 | 2.42585E-18 | 0.631902 | 0.472 | 0.281 | 4.72435E-14 |
| Intermediate | ACTN1 | 8.40718E-49 | 0.631716 | 0.939 | 0.859 | 1.6373E-44 |
| Contractile | TGFBR3 | 1.14065E-49 | 0.631538 | 0.434 | 0.23 | 2.22142E-45 |
| Fibroblasts | WNT5A | 2.15215E-58 | 0.631358 | 0.183 | 0.017 | 4.19131E-54 |
| Fibroblasts | EPB41L2 | 1.0835E-09 | 0.631282 | 0.596 | 0.48 | 2.11011E-05 |
| Intermediate | ENSSSCG0( | 1.80842E-24 | 0.630995 | 0.51 | 0.332 | 3.5219E-20 |
| VEGFA_Terminal | MGAT4A | 7.32405E-49 | 0.6308 | 0.354 | 0.107 | 1.42636E-44 |
| Contractile | SCAPER | 4.37301E-58 | 0.629279 | 0.372 | 0.168 | 8.51643E-54 |
| Fibroblasts | ABCA8 | 1.7542E-106 | 0.629042 | 0.147 | 0.004 | 3.4162E-102 |
| Pericytes | UQCRC1 | 0.009849463 | 0.628749 | 0.44 | 0.333 | 1 |
| VEGFA_Terminal | SYPL1 | 1.02429E-27 | 0.628617 | 0.676 | 0.422 | 1.99481E-23 |

| | | | | | | |
|---|---|---|---|---|---|---|
| Contractile | APP | 1.37605E-87 | 0.628515 | 0.951 | 0.876 | 2.67985E-83 |
| Contractile | MAP4 | 1.36225E-62 | 0.628204 | 0.759 | 0.569 | 2.65298E-58 |
| VEGFA_Terminal | TMBIM6 | 1.60837E-27 | 0.62816 | 0.881 | 0.744 | 3.13229E-23 |
| Pericytes | LMO3 | 2.06163E-48 | 0.628032 | 0.14 | 0.003 | 4.01503E-44 |
| Terminal | ENSSSCG0( | 2.65035E-74 | 0.62799 | 0.586 | 0.364 | 5.16155E-70 |
| Terminal | RRBP1 | 4.99331E-89 | 0.627724 | 0.865 | 0.706 | 9.72448E-85 |
| Intermediate | ENSSSCG0( | 1.87542E-23 | 0.627141 | 0.143 | 0.047 | 3.65238E-19 |
| Contractile | MAF1 | 6.47572E-55 | 0.626618 | 0.474 | 0.261 | 1.26115E-50 |
| Pericytes | ENSSSCG0( | 0.001021144 | 0.626259 | 0.48 | 0.313 | 1 |
| Fibroblasts | PIK3R1 | 1.82914E-06 | 0.625956 | 0.518 | 0.433 | 0.035622548 |
| IFN_Intermediate | USP18 | 4.41269E-36 | 0.625949 | 0.234 | 0.026 | 8.59371E-32 |
| Contractile | ATP2B4 | 2.98485E-57 | 0.625881 | 0.786 | 0.624 | 5.81299E-53 |
| VEGFA_Terminal | FXYD6 | 9.61613E-32 | 0.625759 | 0.54 | 0.271 | 1.87274E-27 |
| Contractile | MAP1B | 9.19286E-77 | 0.625406 | 0.932 | 0.809 | 1.79031E-72 |
| Terminal | ACAN | 1.45919E-85 | 0.624792 | 0.11 | 0.006 | 2.84177E-81 |
| VEGFA_Terminal | ARPC2 | 6.57608E-42 | 0.624426 | 0.927 | 0.797 | 1.28069E-37 |
| Fibroblasts | RBPJ | 2.79983E-07 | 0.624298 | 0.628 | 0.58 | 0.005452673 |
| Contractile | RIMS3 | 6.2663E-175 | 0.624105 | 0.227 | 0.015 | 1.2204E-170 |
| Pericytes | DDX3X | 0.00748882 | 0.623993 | 0.54 | 0.441 | 1 |
| Fibroblasts | HLF | 6.52019E-11 | 0.623869 | 0.339 | 0.191 | 1.26981E-06 |
| Pericytes | ENSSSCG0( | 5.55563E-11 | 0.623842 | 0.18 | 0.028 | 1.08196E-06 |
| VEGFA_Terminal | KDELR3 | 5.42879E-29 | 0.623693 | 0.702 | 0.447 | 1.05726E-24 |
| Pericytes | ENSSSCG0( | 6.08206E-07 | 0.623337 | 0.96 | 0.976 | 0.011844821 |
| Pericytes | PTBP1 | 0.007606439 | 0.623141 | 0.46 | 0.354 | 1 |
| Pericytes | LYAR | 0.000497897 | 0.623083 | 0.22 | 0.086 | 1 |
| VEGFA_Terminal | APLN | 1.7439E-141 | 0.623066 | 0.249 | 0.018 | 3.3962E-137 |
| VEGFA_Terminal | TUBB | 2.34279E-25 | 0.623063 | 0.729 | 0.505 | 4.56257E-21 |
| Pericytes | PPP2R5A | 0.008914456 | 0.622914 | 0.36 | 0.233 | 1 |
| Contractile | KCNMA1 | 1.87101E-49 | 0.622488 | 0.524 | 0.317 | 3.64379E-45 |
| Fibroblasts | TIMP2 | 5.13382E-18 | 0.621863 | 0.927 | 0.924 | 9.99811E-14 |
| Terminal | SELENOM | 2.63101E-89 | 0.621375 | 0.854 | 0.682 | 5.12389E-85 |
| Fibroblasts | CELF2 | 3.19758E-15 | 0.620953 | 0.344 | 0.161 | 6.22729E-11 |
| Intermediate | ENSSSCG0( | 4.52141E-48 | 0.620939 | 0.868 | 0.691 | 8.80545E-44 |
| Pericytes | ENSSSCG0( | 5.62087E-05 | 0.620379 | 0.9 | 0.965 | 1 |
| Contractile | JAZF1 | 7.60068E-68 | 0.619858 | 0.34 | 0.13 | 1.48023E-63 |
| Intermediate | MYH9 | 1.76086E-30 | 0.619215 | 0.876 | 0.801 | 3.42927E-26 |
| Pericytes | ENSSSCG0( | 4.66532E-06 | 0.619188 | 0.96 | 0.962 | 0.090857048 |
| Contractile | MRPS6 | 4.91262E-63 | 0.619101 | 0.364 | 0.151 | 9.56733E-59 |
| VEGFA_Terminal | ENSSSCG0( | 0.000173699 | 0.618479 | 0.208 | 0.143 | 1 |
| Pericytes | CIRBP | 0.000431756 | 0.61756 | 0.56 | 0.39 | 1 |
| Contractile | WWP2 | 2.07059E-49 | 0.617399 | 0.401 | 0.204 | 4.03247E-45 |
| Contractile | IGF1R | 1.53627E-51 | 0.616908 | 0.346 | 0.156 | 2.99189E-47 |
| VEGFA_Terminal | APOA1 | 5.14039E-30 | 0.616762 | 0.31 | 0.114 | 1.00109E-25 |
| VEGFA_Terminal | CXCL2 | 2.90088E-14 | 0.616468 | 0.252 | 0.121 | 5.64947E-10 |
| Contractile | MEF2C | 3.74821E-47 | 0.616299 | 0.597 | 0.397 | 7.29963E-43 |
| VEGFA_Terminal | MAFF | 3.74759E-26 | 0.616224 | 0.494 | 0.261 | 7.29843E-22 |
| Terminal | TXN | 1.8217E-119 | 0.616157 | 0.977 | 0.921 | 3.5478E-115 |
| Fibroblasts | C1QTNF4 | 9.93202E-13 | 0.615129 | 0.174 | 0.059 | 1.93426E-08 |
| Intermediate | SERPING1 | 1.34895E-51 | 0.615049 | 0.946 | 0.807 | 2.62708E-47 |
| Pericytes | REM1 | 1.11749E-10 | 0.614402 | 0.22 | 0.04 | 2.17632E-06 |
| Pericytes | CDS2 | 0.001585303 | 0.613604 | 0.26 | 0.121 | 1 |
| CD74_Intermediat | ADD3 | 2.2155E-86 | 0.613337 | 0.608 | 0.355 | 4.31469E-82 |
| Fibroblasts | BACH1 | 1.10278E-05 | 0.61296 | 0.372 | 0.285 | 0.214765703 |
| Fibroblasts | COL5A3 | 4.25755E-10 | 0.612425 | 0.33 | 0.186 | 8.29157E-06 |
| IFN_Intermediate | FNDC1 | 1.63614E-05 | 0.612399 | 0.673 | 0.473 | 0.318638117 |
| Pericytes | GUCY1A2 | 3.81074E-30 | 0.61221 | 0.14 | 0.006 | 7.42142E-26 |
| VEGFA_Terminal | S100A4 | 1.94557E-24 | 0.612046 | 0.906 | 0.656 | 3.78899E-20 |
| Contractile | NACA | 2.2686E-119 | 0.611961 | 0.989 | 0.982 | 4.4181E-115 |
| Contractile | PJA2 | 4.10109E-48 | 0.611445 | 0.617 | 0.435 | 7.98687E-44 |

| | | | | | | |
|---|---|---|---|---|---|---|
| Intermediate | PLCB1 | 7.19542E-23 | 0.611117 | 0.315 | 0.167 | 1.40131E-18 |
| Pericytes | BRD2 | 0.002992674 | 0.610878 | 0.54 | 0.379 | 1 |
| Contractile | PKIA | 2.64772E-55 | 0.610793 | 0.391 | 0.189 | 5.15643E-51 |
| Contractile | ENSSSCG00 | 1.7334E-198 | 0.610442 | 0.209 | 0.007 | 3.3757E-194 |
| Pericytes | NAPA | 9.52662E-06 | 0.610199 | 0.3 | 0.113 | 0.185530871 |
| VEGFA_Terminal | TJP2 | 8.52965E-35 | 0.609842 | 0.45 | 0.199 | 1.66115E-30 |
| Terminal | IVNS1ABP | 8.10442E-57 | 0.60903 | 0.449 | 0.261 | 1.57834E-52 |
| Intermediate | COL18A1 | 9.59659E-19 | 0.608647 | 0.355 | 0.213 | 1.86894E-14 |
| VEGFA_Terminal | DUSP1 | 8.25763E-11 | 0.608492 | 0.603 | 0.466 | 1.60817E-06 |
| CD74_Intermediat | FLRT2 | 7.12989E-41 | 0.343466 | 0.178 | 0.066 | 1.38855E-36 |
| IFN_Intermediate | IL34 | 8.14024E-07 | 0.607968 | 0.374 | 0.19 | 0.015853116 |
| Fibroblasts | MYC | 7.1074E-09 | 0.607677 | 0.165 | 0.067 | 0.000138417 |
| Terminal | MFGE8 | 2.17291E-85 | 0.607519 | 0.885 | 0.765 | 4.23173E-81 |
| Pericytes | LAMB2 | 0.002963646 | 0.606875 | 0.64 | 0.523 | 1 |
| Fibroblasts | CEBPD | 5.84498E-08 | 0.606587 | 0.867 | 0.828 | 0.001138309 |
| Pericytes | ENSSSCG00 | 0.000235025 | 0.606583 | 0.24 | 0.092 | 1 |
| Fibroblasts | TMEFF2 | 4.25646E-98 | 0.606515 | 0.151 | 0.005 | 8.28946E-94 |
| Fibroblasts | TENM3 | 5.21412E-18 | 0.606064 | 0.234 | 0.078 | 1.01545E-13 |
| Intermediate | TPM4 | 8.20359E-45 | 0.605222 | 0.985 | 0.951 | 1.59765E-40 |
| Pericytes | ENSSSCG00 | 0.000975372 | 0.604759 | 0.14 | 0.045 | 1 |
| VEGFA_Terminal | FARP1 | 3.0252E-31 | 0.604596 | 0.525 | 0.264 | 5.89157E-27 |
| Contractile | CRABP1 | 6.6875E-120 | 0.604587 | 0.202 | 0.023 | 1.3024E-115 |
| Pericytes | G3BP2 | 0.003073358 | 0.603951 | 0.42 | 0.276 | 1 |
| VEGFA_Terminal | UGP2 | 7.57118E-17 | 0.603592 | 0.61 | 0.409 | 1.47449E-12 |
| CD74_Intermediat | TMTC1 | 1.91827E-52 | 0.379411 | 0.196 | 0.067 | 3.73583E-48 |
| Fibroblasts | CAVIN2 | 1.49369E-06 | 0.603284 | 0.344 | 0.236 | 0.029089542 |
| Contractile | DYNLT3 | 1.17212E-57 | 0.603173 | 0.739 | 0.568 | 2.28271E-53 |
| Fibroblasts | HES1 | 1.54351E-07 | 0.603121 | 0.317 | 0.19 | 0.003005989 |
| Pericytes | GLDN | 0.00971056 | 0.603086 | 0.12 | 0.046 | 1 |
| Pericytes | BMP2 | 1.09791E-08 | 0.602216 | 0.16 | 0.027 | 0.000213817 |
| Fibroblasts | TSPAN11 | 1.95117E-69 | 0.601538 | 0.161 | 0.01 | 3.79991E-65 |
| Contractile | RNF11 | 4.75725E-50 | 0.601345 | 0.594 | 0.394 | 9.26474E-46 |
| VEGFA_Terminal | IFNGR2 | 1.75681E-28 | 0.601118 | 0.584 | 0.326 | 3.42139E-24 |
| Fibroblasts | QKI | 2.64501E-10 | 0.600968 | 0.606 | 0.491 | 5.15116E-06 |
| Pericytes | ARHGEF17 | 0.001533782 | 0.599894 | 0.42 | 0.249 | 1 |
| VEGFA_Terminal | FKBP8 | 2.33526E-24 | 0.5996 | 0.637 | 0.402 | 4.54792E-20 |
| Contractile | EIF3A | 3.20501E-53 | 0.599437 | 0.741 | 0.583 | 6.24177E-49 |
| Pericytes | ADCY3 | 1.81868E-05 | 0.599196 | 0.2 | 0.06 | 0.354188095 |
| Pericytes | MYOM1 | 2.29524E-07 | 0.598739 | 0.2 | 0.047 | 0.004469974 |
| Contractile | GPRC5B | 1.16304E-74 | 0.598389 | 0.294 | 0.093 | 2.26502E-70 |
| IFN_Intermediate | ETV1 | 1.12607E-07 | 0.597732 | 0.692 | 0.471 | 0.002193012 |
| Pericytes | SNTB1 | 7.73709E-46 | 0.597413 | 0.16 | 0.005 | 1.5068E-41 |
| Fibroblasts | SLC1A5 | 1.08333E-06 | 0.597336 | 0.417 | 0.321 | 0.021097898 |
| Pericytes | WNT6 | 5.64243E-57 | 0.597152 | 0.14 | 0.003 | 1.09886E-52 |
| Contractile | STK17A | 8.91719E-60 | 0.597116 | 0.448 | 0.218 | 1.73662E-55 |
| Contractile | ID4 | 1.63405E-48 | 0.597047 | 0.588 | 0.367 | 3.18231E-44 |
| Terminal | PCOLCE | 1.27397E-89 | 0.596845 | 0.851 | 0.626 | 2.48105E-85 |
| Intermediate | KCNE1 | 3.24424E-30 | 0.596799 | 0.387 | 0.202 | 6.31816E-26 |
| Contractile | SCOC | 1.74013E-50 | 0.59637 | 0.41 | 0.213 | 3.3889E-46 |
| Pericytes | RACK1 | 1.01513E-05 | 0.596352 | 0.94 | 0.935 | 0.197696673 |
| Fibroblasts | C1R | 1.16889E-15 | 0.59633 | 0.968 | 0.878 | 2.27641E-11 |
| Pericytes | CCM2 | 0.000446636 | 0.596304 | 0.3 | 0.14 | 1 |
| VEGFA_Terminal | PAPSS1 | 2.63054E-27 | 0.595941 | 0.671 | 0.416 | 5.12297E-23 |
| VEGFA_Terminal | BASP1 | 9.39913E-33 | 0.595546 | 0.559 | 0.286 | 1.83048E-28 |
| Fibroblasts | HDAC7 | 1.08463E-08 | 0.594831 | 0.394 | 0.262 | 0.000211232 |
| Pericytes | PSPC1 | 0.007722466 | 0.594153 | 0.36 | 0.232 | 1 |
| VEGFA_Terminal | DDIT3 | 1.07341E-14 | 0.593971 | 0.39 | 0.229 | 2.09046E-10 |
| Pericytes | PTMS | 1.04324E-08 | 0.593469 | 0.92 | 0.957 | 0.00020317 |
| Fibroblasts | CDK14 | 2.7884E-11 | 0.593414 | 0.252 | 0.118 | 5.43042E-07 |

| | | | | | | |
|---|---|---|---|---|---|---|
| Pericytes | INPP5A | 0.003843905 | 0.593147 | 0.26 | 0.133 | 1 |
| Terminal | OMD | 3.5884E-51 | 0.593103 | 0.462 | 0.274 | 6.98841E-47 |
| Contractile | FABP4 | 2.8235E-113 | 0.592923 | 0.369 | 0.094 | 5.4989E-109 |
| Terminal | PROCR | 2.37102E-36 | 0.592523 | 0.178 | 0.075 | 4.61755E-32 |
| Contractile | CAVIN3 | 4.92266E-91 | 0.592077 | 0.989 | 0.953 | 9.58688E-87 |
| Contractile | KCTD10 | 2.59851E-42 | 0.59124 | 0.485 | 0.3 | 5.0606E-38 |
| Contractile | ENSSSCG0( | 4.27633E-51 | 0.590982 | 0.722 | 0.516 | 8.32816E-47 |
| Terminal | S100A6 | 1.2407E-157 | 0.590349 | 0.999 | 0.99 | 2.4163E-153 |
| Pericytes | SINHCAF | 0.000994964 | 0.589173 | 0.22 | 0.09 | 1 |
| Pericytes | SNRPB | 0.008712397 | 0.58812 | 0.46 | 0.332 | 1 |
| VEGFA_Terminal | TBX3 | 1.48036E-37 | 0.588057 | 0.349 | 0.123 | 2.88301E-33 |
| Contractile | PDE3A | 3.31544E-42 | 0.587832 | 0.51 | 0.311 | 6.45681E-38 |
| Contractile | NREP | 3.66482E-49 | 0.587731 | 0.308 | 0.131 | 7.13724E-45 |
| CD74_Intermediat | SMOC1 | 1.712E-67 | 0.587633 | 0.687 | 0.478 | 3.33412E-63 |
| Pericytes | NRIP2 | 1.04985E-59 | 0.587623 | 0.14 | 0.002 | 2.04457E-55 |
| Contractile | RAB27B | 3.69426E-51 | 0.587612 | 0.338 | 0.15 | 7.19457E-47 |
| Contractile | ARHGAP1 | 4.47313E-48 | 0.58749 | 0.424 | 0.224 | 8.71142E-44 |
| Pericytes | CD36 | 1.90521E-20 | 0.587471 | 0.1 | 0.005 | 3.7104E-16 |
| Contractile | CAP2 | 2.9618E-122 | 0.587313 | 0.21 | 0.025 | 5.7682E-118 |
| VEGFA_Terminal | SLC39A14 | 2.43909E-18 | 0.587111 | 0.4 | 0.221 | 4.75013E-14 |
| Terminal | TSPO | 6.4575E-131 | 0.586991 | 0.96 | 0.912 | 1.2576E-126 |
| Pericytes | TP53INP2 | 8.0979E-08 | 0.586966 | 0.16 | 0.03 | 0.001577066 |
| VEGFA_Terminal | TSPAN15 | 3.77449E-44 | 0.586053 | 0.341 | 0.109 | 7.35082E-40 |
| Fibroblasts | ASB9 | 4.98934E-40 | 0.585865 | 0.161 | 0.02 | 9.71675E-36 |
| CD74_Intermediat | CTSK | 5.4455E-101 | 0.585682 | 0.948 | 0.853 | 1.06051E-96 |
| IFN_Intermediate | SHISA5 | 3.51064E-07 | 0.585537 | 0.551 | 0.349 | 0.006836962 |
| VEGFA_Terminal | HECTD2 | 1.74836E-32 | 0.584854 | 0.426 | 0.189 | 3.40494E-28 |
| Pericytes | ENSSSCG0( | 1.61784E-07 | 0.58477 | 0.16 | 0.032 | 0.003150752 |
| Pericytes | ENSSSCG0( | 0.000426664 | 0.584217 | 0.14 | 0.042 | 1 |
| Contractile | PID1 | 1.85093E-97 | 0.58419 | 0.285 | 0.066 | 3.60469E-93 |
| VEGFA_Terminal | CYP7A1 | 1.45116E-07 | 0.584175 | 0.131 | 0.065 | 0.002826135 |
| VEGFA_Terminal | SLC2A5 | 1.42151E-50 | 0.584119 | 0.153 | 0.022 | 2.76839E-46 |
| Intermediate | LAMB1 | 1.69872E-22 | 0.583809 | 0.753 | 0.637 | 3.30825E-18 |
| Intermediate | HGF | 1.01342E-17 | 0.583718 | 0.386 | 0.25 | 1.97364E-13 |
| VEGFA_Terminal | ARRDC3 | 4.28523E-25 | 0.583618 | 0.32 | 0.136 | 8.34549E-21 |
| Terminal | KDELR3 | 1.73598E-75 | 0.582911 | 0.617 | 0.392 | 3.38081E-71 |
| Contractile | RNF149 | 1.16081E-48 | 0.582619 | 0.365 | 0.174 | 2.26067E-44 |
| VEGFA_Terminal | CIDEC | 7.7633E-36 | 0.582452 | 0.194 | 0.048 | 1.5119E-31 |
| Fibroblasts | TNPO1 | 5.87914E-09 | 0.581644 | 0.417 | 0.282 | 0.000114496 |
| Fibroblasts | CYGB | 3.4698E-43 | 0.580961 | 0.147 | 0.015 | 6.75744E-39 |
| Intermediate | B2M | 6.87419E-45 | 0.580288 | 0.931 | 0.853 | 1.33875E-40 |
| Intermediate | ZYX | 9.96195E-28 | 0.580039 | 0.657 | 0.496 | 1.94009E-23 |
| VEGFA_Terminal | HOPX | 3.46652E-25 | 0.579928 | 0.588 | 0.34 | 6.75104E-21 |
| Pericytes | ENSSSCG0( | 0.000750818 | 0.579872 | 0.52 | 0.359 | 1 |
| Pericytes | PAWR | 1.93216E-05 | 0.579829 | 0.8 | 0.613 | 0.37628739 |
| Pericytes | STAT1 | 0.00376419 | 0.579493 | 0.4 | 0.251 | 1 |
| CD74_Intermediat | ENSSSCG0( | 9.41033E-85 | 0.57943 | 0.837 | 0.66 | 1.83266E-80 |
| VEGFA_Terminal | RNH1 | 3.04809E-28 | 0.578596 | 0.84 | 0.654 | 5.93616E-24 |
| Intermediate | ENSSSCG0( | 2.62723E-32 | 0.578445 | 0.344 | 0.159 | 5.11653E-28 |
| Contractile | NFIA | 2.91762E-49 | 0.578428 | 0.708 | 0.523 | 5.68207E-45 |
| Contractile | SLIT3 | 6.52204E-59 | 0.578317 | 0.468 | 0.225 | 1.27017E-54 |
| Fibroblasts | TTC14 | 0.000212086 | 0.577911 | 0.39 | 0.323 | 1 |
| Terminal | EMP1 | 2.05719E-73 | 0.577689 | 0.792 | 0.578 | 4.00638E-69 |
| Pericytes | ANGPTL1 | 8.01467E-09 | 0.576897 | 0.14 | 0.021 | 0.000156086 |
| Contractile | PTGES3L | 8.6498E-115 | 0.57645 | 0.227 | 0.034 | 1.6845E-110 |
| Pericytes | CLTB | 0.007591251 | 0.576173 | 0.5 | 0.433 | 1 |
| Pericytes | ARHGAP6 | 0.003246198 | 0.575986 | 0.22 | 0.104 | 1 |
| VEGFA_Terminal | FKBP1A | 5.0088E-21 | 0.575645 | 0.758 | 0.567 | 9.75464E-17 |
| Terminal | HOPX | 5.8748E-75 | 0.575518 | 0.515 | 0.282 | 1.14412E-70 |

| | | | | | | |
|---|---|---|---|---|---|---|
| Intermediate | FBLN5 | 6.4375E-36 | 0.575465 | 0.776 | 0.571 | 1.2537E-31 |
| Pericytes | BCL3 | 0.009001697 | 0.57534 | 0.2 | 0.098 | 1 |
| VEGFA_Terminal | ENSSSCG0( | 1.14381E-53 | 0.574689 | 1 | 0.997 | 2.22758E-49 |
| Pericytes | MYH9 | 0.003386394 | 0.574587 | 0.84 | 0.809 | 1 |
| Pericytes | ENSSSCG0( | 0.000794741 | 0.57444 | 0.26 | 0.116 | 1 |
| Contractile | CRTC3 | 6.4562E-54 | 0.573596 | 0.392 | 0.186 | 1.25734E-49 |
| Contractile | TMEM70 | 1.91791E-41 | 0.57352 | 0.419 | 0.242 | 3.73514E-37 |
| Intermediate | VCL | 6.22473E-36 | 0.573475 | 0.845 | 0.701 | 1.21227E-31 |
| Pericytes | RBM24 | 3.55303E-15 | 0.573422 | 0.16 | 0.016 | 6.91953E-11 |
| Contractile | CIB4 | 2.0788E-168 | 0.573393 | 0.186 | 0.008 | 4.0485E-164 |
| Intermediate | CDC42EP3 | 2.86632E-30 | 0.572737 | 0.613 | 0.41 | 5.58216E-26 |
| CD74_Intermediat | CFB | 4.09076E-55 | 0.405109 | 0.206 | 0.07 | 7.96676E-51 |
| Contractile | VAMP2 | 7.69657E-44 | 0.570373 | 0.55 | 0.362 | 1.49891E-39 |
| Fibroblasts | MDM4 | 3.36522E-06 | 0.570332 | 0.312 | 0.216 | 0.065537689 |
| IFN_Intermediate | PML | 2.53857E-12 | 0.570267 | 0.318 | 0.107 | 4.94387E-08 |
| Terminal | ENPP1 | 1.52078E-41 | 0.5702 | 0.319 | 0.173 | 2.96171E-37 |
| Fibroblasts | PCOLCE | 2.90005E-08 | 0.570182 | 0.798 | 0.695 | 0.000564785 |
| Pericytes | ENSSSCG0( | 0.005469113 | 0.569994 | 0.48 | 0.352 | 1 |
| VEGFA_Terminal | SEC61B | 8.44147E-28 | 0.569581 | 0.901 | 0.774 | 1.64398E-23 |
| Fibroblasts | TRIOBP | 4.10044E-14 | 0.569456 | 0.294 | 0.133 | 7.98561E-10 |
| Contractile | KLF9 | 7.20155E-35 | 0.569414 | 0.466 | 0.294 | 1.4025E-30 |
| Fibroblasts | SESTD1 | 1.67361E-07 | 0.568907 | 0.482 | 0.377 | 0.003259351 |
| VEGFA_Terminal | PPIA | 4.58239E-40 | 0.568781 | 0.954 | 0.883 | 8.9242E-36 |
| Contractile | NR4A3 | 1.08758E-29 | 0.568661 | 0.123 | 0.038 | 2.11806E-25 |
| CD74_Intermediat | PAPLN | 4.35763E-32 | 0.252503 | 0.17 | 0.07 | 8.48648E-28 |
| CD74_Intermediat | NTN4 | 8.03147E-64 | 0.567805 | 0.601 | 0.392 | 1.56413E-59 |
| IFN_Intermediate | SHFL | 2.29456E-12 | 0.566688 | 0.336 | 0.118 | 4.46866E-08 |
| Intermediate | MYL9 | 4.71427E-55 | 0.566474 | 0.863 | 0.619 | 9.18104E-51 |
| Contractile | LRRK2 | 7.2241E-39 | 0.56612 | 0.452 | 0.274 | 1.40689E-34 |
| Fibroblasts | CGNL1 | 2.66351E-09 | 0.566105 | 0.427 | 0.281 | 5.18719E-05 |
| CD74_Intermediat | B4GALNT1 | 1.26484E-36 | 0.29428 | 0.177 | 0.071 | 2.46327E-32 |
| VEGFA_Terminal | PRSS35 | 3.9668E-22 | 0.565037 | 0.448 | 0.238 | 7.72533E-18 |
| Terminal | COX4I2 | 1.94309E-86 | 0.564964 | 0.895 | 0.78 | 3.78417E-82 |
| Pericytes | GNAI3 | 0.00842647 | 0.564908 | 0.48 | 0.368 | 1 |
| VEGFA_Terminal | PFKL | 3.4825E-43 | 0.564872 | 0.429 | 0.159 | 6.78216E-39 |
| Fibroblasts | CDON | 1.87731E-07 | 0.564588 | 0.216 | 0.112 | 0.003656053 |
| Terminal | TMSB10 | 1.0819E-117 | 0.564401 | 0.997 | 0.98 | 2.107E-113 |
| IFN_Intermediate | COL14A1 | 1.60746E-07 | 0.564104 | 0.944 | 0.809 | 0.003130536 |
| Pericytes | UBE2H | 0.002318267 | 0.563438 | 0.58 | 0.455 | 1 |
| Intermediate | C1R | 2.14247E-51 | 0.562814 | 0.969 | 0.871 | 4.17245E-47 |
| CD74_Intermediat | SERPING1 | 1.6844E-105 | 0.56242 | 0.949 | 0.771 | 3.2803E-101 |
| Contractile | PDLIM5 | 2.90727E-43 | 0.562181 | 0.507 | 0.313 | 5.6619E-39 |
| Terminal | SHISA4 | 6.97419E-59 | 0.562071 | 0.611 | 0.428 | 1.35822E-54 |
| Contractile | ASB2 | 1.7867E-169 | 0.56195 | 0.199 | 0.01 | 3.4797E-165 |
| IFN_Intermediate | IFIT2 | 3.75597E-94 | 0.561562 | 0.112 | 0.001 | 7.31476E-90 |
| Fibroblasts | PEAR1 | 4.67985E-14 | 0.560798 | 0.234 | 0.09 | 9.114E-10 |
| VEGFA_Terminal | FAM107B | 3.82524E-34 | 0.560744 | 0.373 | 0.145 | 7.44965E-30 |
| Fibroblasts | LOC10052: | 2.62266E-07 | 0.560658 | 0.913 | 0.928 | 0.00510764 |
| Fibroblasts | MARCHF6 | 7.20123E-09 | 0.560552 | 0.472 | 0.347 | 0.000140244 |
| Intermediate | NOX4 | 1.14004E-23 | 0.560382 | 0.473 | 0.305 | 2.22023E-19 |
| VEGFA_Terminal | ELK3 | 8.8646E-33 | 0.560045 | 0.358 | 0.141 | 1.72638E-28 |
| Pericytes | ARID5A | 1.87326E-05 | 0.559754 | 0.28 | 0.103 | 0.36481755 |
| Contractile | ADGRL2 | 8.05459E-38 | 0.559585 | 0.475 | 0.297 | 1.56863E-33 |
| Pericytes | RBM14 | 0.002397442 | 0.559473 | 0.22 | 0.1 | 1 |
| VEGFA_Terminal | ID4 | 9.658E-13 | 0.55937 | 0.554 | 0.392 | 1.8809E-08 |
| Pericytes | JADE3 | 0.000246713 | 0.559163 | 0.14 | 0.04 | 1 |
| Fibroblasts | CLK1 | 1.34489E-06 | 0.558766 | 0.564 | 0.505 | 0.026191683 |
| VEGFA_Terminal | TPM3 | 1.97123E-28 | 0.558746 | 0.77 | 0.507 | 3.83897E-24 |
| CD74_Intermediat | ENSSSCG0( | 7.79434E-32 | 0.262022 | 0.169 | 0.071 | 1.51795E-27 |

| | | | | | | |
|---|---|---|---|---|---|---|
| Contractile | ANTXR2 | 1.12733E-36 | 0.557848 | 0.444 | 0.274 | 2.19547E-32 |
| Fibroblasts | EMP1 | 5.63357E-12 | 0.557838 | 0.794 | 0.642 | 1.09714E-07 |
| Contractile | DNAJB6 | 1.32474E-44 | 0.55782 | 0.62 | 0.447 | 2.57994E-40 |
| Fibroblasts | CAMKK2 | 6.8212E-08 | 0.557324 | 0.243 | 0.132 | 0.00132843 |
| Contractile | ACTB | 1.40814E-56 | 0.557156 | 0.829 | 0.686 | 2.74236E-52 |
| Contractile | ENSSSCG0( | 1.15043E-87 | 0.556206 | 0.205 | 0.038 | 2.24047E-83 |
| Terminal | COL6A3 | 9.44996E-61 | 0.55586 | 0.854 | 0.672 | 1.84038E-56 |
| Contractile | SMIM13 | 6.78615E-48 | 0.555857 | 0.372 | 0.183 | 1.3216E-43 |
| Contractile | DNAJB5 | 6.8503E-133 | 0.555562 | 0.206 | 0.021 | 1.3341E-128 |
| Contractile | ZNF395 | 5.71408E-44 | 0.555528 | 0.457 | 0.262 | 1.11282E-39 |
| VEGFA_Terminal | ACTG1 | 2.63467E-28 | 0.555513 | 0.981 | 0.986 | 5.13102E-24 |
| Pericytes | CSNK1E | 0.005889938 | 0.554895 | 0.36 | 0.23 | 1 |
| IFN_Intermediate | SP110 | 8.23009E-12 | 0.55388 | 0.364 | 0.137 | 1.60281E-07 |
| VEGFA_Terminal | RFTN1 | 6.79447E-69 | 0.553727 | 0.249 | 0.043 | 1.32322E-64 |
| Pericytes | SLC16A12 | 0.000791789 | 0.552494 | 0.14 | 0.044 | 1 |
| IFN_Intermediate | PHF11 | 6.40611E-08 | 0.552135 | 0.383 | 0.184 | 0.00124759 |
| Pericytes | KDM5B | 0.002544689 | 0.552115 | 0.3 | 0.165 | 1 |
| Pericytes | SSR4 | 0.004872557 | 0.552078 | 0.78 | 0.736 | 1 |
| Intermediate | LMCD1 | 4.58869E-22 | 0.551903 | 0.475 | 0.292 | 8.93648E-18 |
| Contractile | SEMA3E | 3.12596E-45 | 0.551833 | 0.232 | 0.084 | 6.0878E-41 |
| Intermediate | PCGF5 | 7.00095E-16 | 0.5518 | 0.651 | 0.557 | 1.36343E-11 |
| VEGFA_Terminal | PYCR1 | 1.87257E-33 | 0.551566 | 0.424 | 0.179 | 3.64682E-29 |
| IFN_Intermediate | PLEKHA4 | 2.09424E-16 | 0.551216 | 0.271 | 0.066 | 4.07854E-12 |
| VEGFA_Terminal | PDK1 | 5.01184E-26 | 0.551122 | 0.523 | 0.274 | 9.76056E-22 |
| CD74_Intermediat | PDCD4 | 3.56508E-48 | 0.550823 | 0.628 | 0.453 | 6.94299E-44 |
| Contractile | BLTP3B | 2.53044E-34 | 0.550797 | 0.485 | 0.325 | 4.92804E-30 |
| Contractile | ENSSSCG0( | 6.0709E-101 | 0.550637 | 0.22 | 0.037 | 1.18231E-96 |
| Fibroblasts | FZD7 | 1.36121E-06 | 0.550353 | 0.275 | 0.171 | 0.026509568 |
| CD74_Intermediat | CPXM2 | 2.70855E-74 | 0.550236 | 0.691 | 0.462 | 5.27489E-70 |
| Contractile | COL4A2 | 4.90394E-57 | 0.549697 | 0.765 | 0.551 | 9.55042E-53 |
| Intermediate | CPED1 | 5.69904E-30 | 0.549572 | 0.587 | 0.392 | 1.10989E-25 |
| Contractile | BOC | 2.6149E-52 | 0.54939 | 0.308 | 0.126 | 5.09251E-48 |
| Fibroblasts | NR1D2 | 1.10859E-07 | 0.548636 | 0.436 | 0.322 | 0.002158982 |
| Contractile | NDRG4 | 1.98386E-82 | 0.54829 | 0.181 | 0.031 | 3.86358E-78 |
| Intermediate | KCNAB1 | 2.01299E-30 | 0.547857 | 0.433 | 0.233 | 3.9203E-26 |
| Pericytes | MMD | 0.002306832 | 0.547855 | 0.12 | 0.039 | 1 |
| VEGFA_Terminal | TXNDC17 | 6.0823E-28 | 0.547744 | 0.777 | 0.556 | 1.18453E-23 |
| Fibroblasts | TPT1 | 4.50761E-19 | 0.547129 | 0.982 | 0.981 | 8.77858E-15 |
| Fibroblasts | ITM2B | 2.09412E-24 | 0.547108 | 0.982 | 0.973 | 4.0783E-20 |
| IFN_Intermediate | TLR3 | 7.40076E-16 | 0.546959 | 0.262 | 0.065 | 1.4413E-11 |
| Fibroblasts | LAPTM4A | 1.29667E-17 | 0.546738 | 0.89 | 0.831 | 2.52527E-13 |
| Pericytes | ENSSSCG0( | 6.76539E-06 | 0.546607 | 0.1 | 0.017 | 0.131755946 |
| CD74_Intermediat | PROS1 | 4.98852E-72 | 0.546413 | 0.715 | 0.517 | 9.71515E-68 |
| VEGFA_Terminal | EIF4EBP3 | 1.04286E-26 | 0.545994 | 0.322 | 0.134 | 2.03097E-22 |
| CD74_Intermediat | MMP2 | 3.03011E-94 | 0.545545 | 0.889 | 0.729 | 5.90115E-90 |
| Intermediate | PDE3A | 8.64932E-25 | 0.545463 | 0.501 | 0.325 | 1.68445E-20 |
| Fibroblasts | PPL | 2.84803E-24 | 0.545248 | 0.193 | 0.043 | 5.54654E-20 |
| CD74_Intermediat | LTC4S | 1.02012E-86 | 0.545193 | 0.877 | 0.631 | 1.98668E-82 |
| VEGFA_Terminal | ST6GALNA | 7.5373E-26 | 0.544904 | 0.443 | 0.224 | 1.46789E-21 |
| Terminal | COL6A1 | 8.35917E-68 | 0.544858 | 0.94 | 0.853 | 1.62795E-63 |
| Contractile | PPP1R14C | 1.6585E-150 | 0.54457 | 0.196 | 0.013 | 3.2299E-146 |
| VEGFA_Terminal | ARRDC4 | 7.54612E-24 | 0.544232 | 0.278 | 0.112 | 1.46961E-19 |
| Pericytes | SAG | 1.01165E-09 | 0.543834 | 0.1 | 0.01 | 1.97018E-05 |
| Intermediate | HMG20B | 2.01915E-19 | 0.543834 | 0.422 | 0.283 | 3.93229E-15 |
| Contractile | MPRIP | 8.93352E-43 | 0.543644 | 0.378 | 0.197 | 1.7398E-38 |
| Fibroblasts | SPON1 | 1.64743E-06 | 0.543208 | 0.537 | 0.433 | 0.032083643 |
| Contractile | NCOR1 | 2.03511E-42 | 0.543105 | 0.538 | 0.341 | 3.96337E-38 |
| Fibroblasts | LVRN | 4.69072E-52 | 0.543061 | 0.124 | 0.008 | 9.13519E-48 |
| Fibroblasts | CFLAR | 2.93876E-06 | 0.542916 | 0.321 | 0.22 | 0.057232264 |

| | | | | | | |
|---|---|---|---|---|---|---|
| Intermediate | RAMP1 | 2.23986E-18 | 0.542583 | 0.284 | 0.154 | 4.36213E-14 |
| Terminal | CAPG | 3.25738E-66 | 0.54196 | 0.797 | 0.633 | 6.34374E-62 |
| Fibroblasts | GASK1A | 1.74461E-22 | 0.541312 | 0.174 | 0.04 | 3.39763E-18 |
| Contractile | ENSSSCG0( | 6.00715E-61 | 0.541105 | 0.332 | 0.128 | 1.16989E-56 |
| Intermediate | CTSS | 8.03686E-26 | 0.540898 | 0.469 | 0.284 | 1.56518E-21 |
| Contractile | CD47 | 9.99495E-40 | 0.540768 | 0.709 | 0.561 | 1.94652E-35 |
| Contractile | EIF3L | 1.9195E-41 | 0.540758 | 0.659 | 0.493 | 3.73822E-37 |
| Intermediate | MYH11 | 1.14741E-76 | 0.540297 | 0.81 | 0.41 | 2.23459E-72 |
| Terminal | CGREF1 | 4.46459E-76 | 0.540275 | 0.278 | 0.098 | 8.69478E-72 |
| VEGFA_Terminal | ENSSSCG0( | 1.1585E-71 | 0.540054 | 0.201 | 0.026 | 2.25617E-67 |
| IFN_Intermediate | DTX3L | 1.19053E-25 | 0.539943 | 0.243 | 0.038 | 2.31856E-21 |
| IFN_Intermediate | CLU | 8.54018E-11 | 0.539847 | 0.991 | 0.962 | 1.6632E-06 |
| Contractile | LIN7C | 5.99929E-38 | 0.53961 | 0.519 | 0.343 | 1.16836E-33 |
| Pericytes | MTA2 | 0.002725742 | 0.539597 | 0.22 | 0.103 | 1 |
| Fibroblasts | RAMP2 | 3.95819E-07 | 0.539245 | 0.495 | 0.363 | 0.007708569 |
| Intermediate | SPINT2 | 8.59082E-34 | 0.539159 | 0.648 | 0.419 | 1.67306E-29 |
| Fibroblasts | CTSV | 4.19849E-15 | 0.539032 | 0.885 | 0.849 | 8.17657E-11 |
| Terminal | DYNLL1 | 1.63617E-69 | 0.539017 | 0.844 | 0.712 | 3.18644E-65 |
| Fibroblasts | MAP1A | 4.13148E-06 | 0.538998 | 0.56 | 0.486 | 0.080460478 |
| Fibroblasts | TFPI2 | 8.32435E-11 | 0.538608 | 0.872 | 0.812 | 1.62117E-06 |
| Pericytes | KANK2 | 0.005426118 | 0.538513 | 0.64 | 0.541 | 1 |
| Fibroblasts | ATP1B1 | 2.23271E-05 | 0.538509 | 0.417 | 0.322 | 0.434819938 |
| VEGFA_Terminal | C9H1ORF2 | 4.99148E-25 | 0.538416 | 0.673 | 0.414 | 9.7209E-21 |
| VEGFA_Terminal | LMAN1 | 1.92596E-25 | 0.538401 | 0.661 | 0.426 | 3.75081E-21 |
| CD74_Intermediat | FOS | 1.47571E-27 | 0.538334 | 0.672 | 0.553 | 2.87394E-23 |
| Contractile | PRDM6 | 1.667E-63 | 0.538218 | 0.267 | 0.085 | 3.24649E-59 |
| Fibroblasts | PCSK6 | 1.65824E-21 | 0.538144 | 0.161 | 0.035 | 3.22941E-17 |
| Pericytes | TCF7 | 0.002407825 | 0.538142 | 0.16 | 0.061 | 1 |
| Intermediate | LTBP4 | 2.18603E-19 | 0.538114 | 0.806 | 0.731 | 4.25728E-15 |
| VEGFA_Terminal | P4HB | 2.17742E-22 | 0.538086 | 0.678 | 0.444 | 4.24053E-18 |
| Contractile | NUCKS1 | 4.28674E-81 | 0.537778 | 0.965 | 0.906 | 8.34842E-77 |
| Fibroblasts | FNDC3A | 3.74322E-05 | 0.537705 | 0.537 | 0.473 | 0.728991544 |
| CD74_Intermediat | SLA-DQB1 | 1.3107E-165 | 1.17845 | 0.348 | 0.075 | 2.5526E-161 |
| Pericytes | PTBP2 | 0.002669817 | 0.537222 | 0.24 | 0.114 | 1 |
| Pericytes | ENSSSCG0( | 6.29751E-11 | 0.536645 | 0.12 | 0.013 | 1.22644E-06 |
| CD74_Intermediat | PLSCR4 | 2.56735E-71 | 0.535728 | 0.564 | 0.33 | 4.99992E-67 |
| Pericytes | SHOX2 | 4.01871E-15 | 0.535474 | 0.12 | 0.009 | 7.82643E-11 |
| Intermediate | CXCL16 | 3.39614E-22 | 0.535239 | 0.524 | 0.356 | 6.61398E-18 |
| IFN_Intermediate | TNXB | 4.17155E-07 | 0.535138 | 0.766 | 0.546 | 0.008124095 |
| Terminal | NDFIP1 | 1.76511E-79 | 0.534956 | 0.806 | 0.64 | 3.43756E-75 |
| IFN_Intermediate | SDCBP | 0.00010872 | 0.534807 | 0.785 | 0.682 | 1 |
| Fibroblasts | PLEKHA5 | 1.76653E-09 | 0.534685 | 0.239 | 0.115 | 3.44032E-05 |
| Fibroblasts | FGF7 | 1.81459E-07 | 0.534182 | 0.307 | 0.178 | 0.003533909 |
| Pericytes | STARD8 | 1.36793E-13 | 0.534157 | 0.16 | 0.018 | 2.66404E-09 |
| Terminal | CD44 | 1.2214E-59 | 0.534099 | 0.513 | 0.312 | 2.37868E-55 |
| Intermediate | MYL6 | 1.46807E-33 | 0.533852 | 0.978 | 0.969 | 2.85906E-29 |
| Contractile | ARID4B | 2.50848E-31 | 0.533479 | 0.575 | 0.429 | 4.88527E-27 |
| Contractile | NPY1R | 6.4561E-101 | 0.532988 | 0.196 | 0.028 | 1.25733E-96 |
| VEGFA_Terminal | RHOC | 1.86415E-18 | 0.532914 | 0.789 | 0.634 | 3.63042E-14 |
| Pericytes | AIF1 | 4.85115E-13 | 0.532869 | 0.12 | 0.011 | 9.44762E-09 |
| VEGFA_Terminal | TPM4 | 4.47902E-30 | 0.532762 | 0.981 | 0.952 | 8.7229E-26 |
| Intermediate | BASP1 | 9.04694E-22 | 0.532697 | 0.447 | 0.287 | 1.76189E-17 |
| Pericytes | MIDN | 0.000715606 | 0.531842 | 0.26 | 0.118 | 1 |
| Intermediate | C1QTNF1 | 3.61698E-21 | 0.531727 | 0.677 | 0.552 | 7.04407E-17 |
| Pericytes | SRSF3 | 0.007287202 | 0.531485 | 0.62 | 0.565 | 1 |
| VEGFA_Terminal | HOMER3 | 1.00289E-29 | 0.531452 | 0.494 | 0.24 | 1.95313E-25 |
| CD74_Intermediat | RARRES1 | 7.84579E-73 | 0.531379 | 0.858 | 0.683 | 1.52797E-68 |
| IFN_Intermediate | GPNMB | 4.09844E-07 | 0.530984 | 0.692 | 0.451 | 0.007981715 |
| Contractile | ENSSSCG0( | 9.29229E-47 | 0.530835 | 0.285 | 0.119 | 1.80967E-42 |

| | | | | | | |
|---|---|---|---|---|---|---|
| Terminal | CRLF1 | 6.33412E-41 | 0.529657 | 0.503 | 0.326 | 1.23357E-36 |
| Intermediate | SFRP4 | 3.89053E-27 | 0.529584 | 0.384 | 0.207 | 7.57682E-23 |
| Fibroblasts | ENSSSCG0( | 3.51053E-13 | 0.529551 | 0.165 | 0.053 | 6.83675E-09 |
| Terminal | FGF7 | 6.35899E-40 | 0.529252 | 0.276 | 0.138 | 1.23841E-35 |
| Intermediate | CXCL2 | 5.29024E-09 | 0.528679 | 0.2 | 0.121 | 0.000103027 |
| Contractile | ESD | 2.93307E-38 | 0.52865 | 0.653 | 0.503 | 5.71216E-34 |
| Fibroblasts | GFRA1 | 1.71595E-21 | 0.528587 | 0.151 | 0.032 | 3.34182E-17 |
| Pericytes | HEYL | 0.000554494 | 0.528425 | 0.42 | 0.234 | 1 |
| Contractile | KIF1C | 9.35023E-42 | 0.528326 | 0.422 | 0.238 | 1.82096E-37 |
| Contractile | RSBN1L | 2.34238E-39 | 0.528279 | 0.607 | 0.426 | 4.56179E-35 |
| Fibroblasts | NOX4 | 3.01326E-05 | 0.52772 | 0.399 | 0.32 | 0.586833177 |
| Intermediate | TINAGL1 | 5.0571E-45 | 0.527512 | 0.23 | 0.069 | 9.8487E-41 |
| Terminal | CHI3L1 | 1.35326E-11 | 0.527192 | 0.599 | 0.517 | 2.63546E-07 |
| Pericytes | GPR4 | 4.95138E-46 | 0.52685 | 0.12 | 0.002 | 9.64281E-42 |
| Contractile | AKT3 | 8.04962E-34 | 0.526811 | 0.491 | 0.328 | 1.56766E-29 |
| VEGFA_Terminal | SDCBP | 1.20887E-21 | 0.526802 | 0.85 | 0.672 | 2.35428E-17 |
| Contractile | ARHGAP1( | 7.48518E-35 | 0.526755 | 0.5 | 0.334 | 1.45774E-30 |
| Contractile | FOXN3 | 2.16786E-33 | 0.526671 | 0.484 | 0.321 | 4.22191E-29 |
| Contractile | NES | 1.09638E-53 | 0.526462 | 0.273 | 0.099 | 2.13521E-49 |
| Pericytes | ENSSSCG0( | 0.000936318 | 0.5264 | 0.1 | 0.026 | 1 |
| Pericytes | PPP1R12C | 6.32407E-06 | 0.525974 | 0.24 | 0.075 | 0.123161241 |
| Pericytes | CYP7B1 | 6.01571E-05 | 0.525908 | 0.18 | 0.055 | 1 |
| Fibroblasts | ENSSSCG0( | 3.11025E-15 | 0.52574 | 0.128 | 0.031 | 6.05721E-11 |
| Contractile | ITGA3 | 3.26571E-71 | 0.525255 | 0.21 | 0.05 | 6.35997E-67 |
| IFN_Intermediate | OAS1 | 2.69847E-57 | 0.525092 | 0.14 | 0.005 | 5.25526E-53 |
| Contractile | PPDPF | 6.59648E-65 | 0.52497 | 0.923 | 0.868 | 1.28466E-60 |
| Contractile | LOC10052( | 6.34795E-48 | 0.524388 | 0.377 | 0.184 | 1.23626E-43 |
| VEGFA_Terminal | AK4 | 1.45045E-34 | 0.524387 | 0.419 | 0.17 | 2.82476E-30 |
| Fibroblasts | KLF11 | 5.26082E-26 | 0.523877 | 0.179 | 0.036 | 1.02454E-21 |
| Contractile | ZFP36L2 | 1.15051E-32 | 0.523642 | 0.926 | 0.877 | 2.24062E-28 |
| Intermediate | SMOC2 | 1.17013E-29 | 0.523469 | 0.293 | 0.13 | 2.27883E-25 |
| Contractile | ENSSSCG0( | 4.68949E-80 | 0.523439 | 0.225 | 0.05 | 9.13279E-76 |
| Fibroblasts | FZD2 | 1.20933E-25 | 0.5234 | 0.151 | 0.027 | 2.35518E-21 |
| Contractile | ENSSSCG0( | 3.93726E-48 | 0.523311 | 0.264 | 0.101 | 7.66782E-44 |
| Contractile | MYO1C | 3.4993E-30 | 0.522788 | 0.528 | 0.375 | 6.81488E-26 |
| Fibroblasts | PTCH1 | 7.66589E-11 | 0.522785 | 0.128 | 0.041 | 1.49293E-06 |
| Pericytes | TSPAN12 | 0.004175826 | 0.522777 | 0.12 | 0.041 | 1 |
| Fibroblasts | TSC22D3 | 6.40726E-05 | 0.522536 | 0.486 | 0.418 | 1 |
| Contractile | ADARB1 | 7.4462E-36 | 0.522521 | 0.454 | 0.275 | 1.45015E-31 |
| Fibroblasts | PITPNC1 | 2.81694E-11 | 0.522446 | 0.197 | 0.077 | 5.48599E-07 |
| Fibroblasts | APOA1 | 0.001730408 | 0.522426 | 0.188 | 0.125 | 1 |
| Contractile | RMND5A | 8.70591E-54 | 0.522387 | 0.313 | 0.128 | 1.69548E-49 |
| Terminal | ENO1 | 1.8294E-126 | 0.52196 | 0.973 | 0.898 | 3.5628E-122 |
| IFN_Intermediate | EDIL3 | 4.40488E-05 | 0.521862 | 0.589 | 0.404 | 0.857850293 |
| CD74_Intermediat | ENSSSCG0( | 1.21192E-73 | 0.521731 | 0.833 | 0.646 | 2.36022E-69 |
| Pericytes | RBPMS2 | 0.001061929 | 0.521695 | 0.12 | 0.035 | 1 |
| VEGFA_Terminal | LAT2 | 1.53092E-26 | 0.521354 | 0.416 | 0.196 | 2.98146E-22 |
| Pericytes | ARID2 | 0.002882801 | 0.521352 | 0.26 | 0.131 | 1 |
| Contractile | CCDC146 | 1.85356E-51 | 0.521308 | 0.255 | 0.091 | 3.60981E-47 |
| Pericytes | NUDT15 | 0.000285687 | 0.521118 | 0.16 | 0.05 | 1 |
| Pericytes | BCL9L | 0.003075484 | 0.521059 | 0.24 | 0.117 | 1 |
| Fibroblasts | EEF1B2 | 4.71592E-12 | 0.520513 | 0.927 | 0.938 | 9.18426E-08 |
| VEGFA_Terminal | ENSSSCG0( | 1.45998E-22 | 0.520411 | 0.54 | 0.313 | 2.84331E-18 |
| VEGFA_Terminal | GJC1 | 2.27194E-22 | 0.520355 | 0.354 | 0.165 | 4.42461E-18 |
| Contractile | PLEKHO1 | 2.07544E-40 | 0.519927 | 0.284 | 0.126 | 4.04192E-36 |
| Pericytes | EBNA1BP2 | 0.005289429 | 0.519867 | 0.28 | 0.156 | 1 |
| Fibroblasts | GALNT12 | 5.10371E-49 | 0.519821 | 0.17 | 0.018 | 9.93947E-45 |
| Fibroblasts | MTUS1 | 1.86461E-07 | 0.519694 | 0.344 | 0.225 | 0.003631324 |
| Fibroblasts | NDEL1 | 2.19622E-08 | 0.519595 | 0.261 | 0.141 | 0.000427714 |

| | | | | | | |
|---|---|---|---|---|---|---|
| Fibroblasts | ENSSSCG0( | 4.29704E-10 | 0.51912 | 0.156 | 0.057 | 8.36848E-06 |
| Contractile | EMC6 | 8.47216E-39 | 0.519055 | 0.631 | 0.456 | 1.64995E-34 |
| Pericytes | KCTD20 | 0.006767459 | 0.519026 | 0.3 | 0.177 | 1 |
| CD74_Intermediat | ENSSSCG0( | 3.2323E-72 | 0.443413 | 0.245 | 0.075 | 6.2949E-68 |
| Pericytes | NGF | 3.6706E-11 | 0.518713 | 0.18 | 0.027 | 7.14849E-07 |
| Pericytes | PDE4A | 0.000178332 | 0.518689 | 0.18 | 0.059 | 1 |
| Contractile | CD93 | 2.3017E-123 | 0.51841 | 0.187 | 0.018 | 4.4825E-119 |
| Terminal | RNH1 | 5.93412E-66 | 0.518399 | 0.782 | 0.611 | 1.15567E-61 |
| Contractile | PDK1 | 6.10976E-38 | 0.518367 | 0.441 | 0.261 | 1.18988E-33 |
| Pericytes | NFIL3 | 0.002898472 | 0.518352 | 0.24 | 0.115 | 1 |
| VEGFA_Terminal | YBX3 | 7.45263E-34 | 0.518352 | 0.969 | 0.869 | 1.4514E-29 |
| Terminal | IL18 | 5.56285E-63 | 0.518319 | 0.328 | 0.145 | 1.08336E-58 |
| Contractile | TCEAL4 | 1.34333E-39 | 0.518097 | 0.628 | 0.453 | 2.61614E-35 |
| Terminal | CYP1B1 | 4.20643E-34 | 0.517953 | 0.222 | 0.109 | 8.19201E-30 |
| Pericytes | PPP1R1A | 2.55408E-11 | 0.517908 | 0.12 | 0.012 | 4.97406E-07 |
| Pericytes | TLE4 | 0.001166519 | 0.517727 | 0.18 | 0.069 | 1 |
| Fibroblasts | CLMP | 7.86837E-08 | 0.517029 | 0.261 | 0.146 | 0.001532364 |
| Fibroblasts | ENSSSCG0( | 3.57586E-06 | 0.516972 | 0.271 | 0.166 | 0.069639841 |
| Fibroblasts | TSPAN4 | 1.65242E-05 | 0.516642 | 0.243 | 0.154 | 0.321809498 |
| IFN_Intermediate | ENSSSCG0( | 0.001392227 | 0.51649 | 0.776 | 0.709 | 1 |
| VEGFA_Terminal | TPST1 | 1.35795E-34 | 0.51615 | 0.368 | 0.14 | 2.64461E-30 |
| VEGFA_Terminal | SDC2 | 2.41469E-25 | 0.515944 | 0.937 | 0.785 | 4.70261E-21 |
| Pericytes | NCK2 | 0.000803263 | 0.515603 | 0.26 | 0.116 | 1 |
| Pericytes | VASP | 0.002177557 | 0.515435 | 0.4 | 0.248 | 1 |
| Fibroblasts | GNG2 | 2.16203E-42 | 0.515365 | 0.165 | 0.02 | 4.21055E-38 |
| Pericytes | CIDEA | 0.000134333 | 0.515241 | 0.1 | 0.022 | 1 |
| Contractile | ANKMY2 | 2.95201E-43 | 0.514419 | 0.502 | 0.303 | 5.74904E-39 |
| Pericytes | ATAD5 | 2.12684E-06 | 0.514235 | 0.1 | 0.015 | 0.041420255 |
| Pericytes | HS6ST1 | 3.43275E-05 | 0.514122 | 0.22 | 0.071 | 0.668528291 |
| Fibroblasts | MTCH1 | 2.2818E-08 | 0.513489 | 0.661 | 0.596 | 0.000444381 |
| Contractile | SLC25A3 | 7.45899E-62 | 0.513328 | 0.935 | 0.872 | 1.45264E-57 |
| Pericytes | ARHGEF7 | 7.62872E-05 | 0.513267 | 0.24 | 0.088 | 1 |
| Contractile | SLC48A1 | 8.40507E-41 | 0.513058 | 0.296 | 0.137 | 1.63689E-36 |
| Fibroblasts | SP3 | 0.000506826 | 0.512963 | 0.358 | 0.3 | 1 |
| Contractile | CCDC85B | 1.93206E-36 | 0.512959 | 0.497 | 0.325 | 3.76269E-32 |
| Contractile | TOMM20 | 4.80739E-35 | 0.512739 | 0.55 | 0.387 | 9.36239E-31 |
| Terminal | FOLR2 | 5.94963E-53 | 0.512703 | 0.577 | 0.395 | 1.15869E-48 |
| Pericytes | ENSSSCG0( | 9.78062E-05 | 0.512584 | 0.12 | 0.028 | 1 |
| VEGFA_Terminal | ACTR3 | 1.98008E-16 | 0.512524 | 0.746 | 0.557 | 3.8562E-12 |
| Contractile | C17ORF58 | 4.55437E-91 | 0.512469 | 0.192 | 0.031 | 8.86963E-87 |
| Contractile | NFIB | 1.27773E-35 | 0.511685 | 0.705 | 0.556 | 2.48838E-31 |
| Fibroblasts | TAMALIN | 5.95656E-23 | 0.511389 | 0.133 | 0.024 | 1.16004E-18 |
| Pericytes | EIF3D | 0.00791283 | 0.510515 | 0.56 | 0.493 | 1 |
| Fibroblasts | CPZ | 2.51969E-73 | 0.510288 | 0.147 | 0.008 | 4.9071E-69 |
| IFN_Intermediate | SLA-DMA | 1.78763E-16 | 0.509781 | 0.196 | 0.038 | 3.48141E-12 |
| Fibroblasts | TMEM98 | 2.64377E-09 | 0.509773 | 0.303 | 0.169 | 5.14875E-05 |
| Contractile | CELF1 | 5.28515E-33 | 0.509723 | 0.459 | 0.297 | 1.02928E-28 |
| Intermediate | PDLIM5 | 4.08286E-17 | 0.509485 | 0.464 | 0.33 | 7.95137E-13 |
| CD74_Intermediat | CDO1 | 6.5872E-101 | 0.509369 | 0.891 | 0.607 | 1.28285E-96 |
| Pericytes | EPDR1 | 0.001268401 | 0.509123 | 0.32 | 0.165 | 1 |
| Pericytes | TAF1D | 0.00696428 | 0.50888 | 0.62 | 0.517 | 1 |
| Pericytes | CAP2 | 0.005551125 | 0.508736 | 0.14 | 0.054 | 1 |
| Contractile | DIPK2A | 1.04108E-35 | 0.50859 | 0.357 | 0.192 | 2.0275E-31 |
| IFN_Intermediate | PSMB10 | 8.22624E-06 | 0.508155 | 0.346 | 0.191 | 0.160205936 |
| Terminal | DAB2 | 2.25537E-53 | 0.507904 | 0.64 | 0.438 | 4.39234E-49 |
| Intermediate | SEPTIN11 | 1.10346E-22 | 0.507807 | 0.601 | 0.453 | 2.14898E-18 |
| Fibroblasts | ENSSSCG0( | 7.53646E-07 | 0.50775 | 0.546 | 0.472 | 0.014677246 |
| VEGFA_Terminal | RASA3 | 8.25989E-28 | 0.507636 | 0.458 | 0.218 | 1.60861E-23 |
| CD74_Intermediat | GPM6B | 5.22888E-38 | 0.299412 | 0.191 | 0.077 | 1.01832E-33 |

| | | | | | | |
|---|---|---|---|---|---|---|
| CD74_Intermediat | CCN5 | 2.34533E-60 | 0.507391 | 0.615 | 0.393 | 4.56754E-56 |
| Intermediate | WASF3 | 8.08324E-20 | 0.507029 | 0.416 | 0.263 | 1.57421E-15 |
| Pericytes | GPCPD1 | 0.000134667 | 0.506825 | 0.26 | 0.101 | 1 |
| VEGFA_Terminal | P2RY6 | 7.8382E-124 | 0.506581 | 0.169 | 0.008 | 1.5265E-119 |
| CD74_Intermediat | LAMA2 | 1.92076E-69 | 0.428132 | 0.252 | 0.082 | 3.74067E-65 |
| Contractile | PDE5A | 1.07566E-37 | 0.506371 | 0.371 | 0.199 | 2.09485E-33 |
| Contractile | ACSS3 | 7.70319E-47 | 0.506165 | 0.291 | 0.12 | 1.5002E-42 |
| Fibroblasts | GINM1 | 6.30576E-08 | 0.505945 | 0.679 | 0.622 | 0.001228047 |
| Intermediate | GJA5 | 7.95977E-22 | 0.505935 | 0.3 | 0.158 | 1.55017E-17 |
| Fibroblasts | HYCC1 | 4.81872E-07 | 0.505615 | 0.353 | 0.239 | 0.009384461 |
| IFN_Intermediate | CXCL16 | 8.35447E-08 | 0.505585 | 0.598 | 0.37 | 0.001627033 |
| Fibroblasts | ECRG4 | 9.33665E-10 | 0.505533 | 0.284 | 0.138 | 1.81831E-05 |
| CD74_Intermediat | NRP1 | 1.60288E-54 | 0.505498 | 0.515 | 0.318 | 3.1216E-50 |
| Fibroblasts | ENSSSCG0( | 1.09611E-05 | 0.504879 | 0.372 | 0.269 | 0.213468252 |
| VEGFA_Terminal | HSPA5 | 4.35488E-05 | 0.504789 | 0.746 | 0.652 | 0.84811198 |
| Pericytes | FAIM | 0.000225792 | 0.504782 | 0.14 | 0.039 | 1 |
| CD74_Intermediat | ENSSSCG0( | 1.99231E-33 | 0.295286 | 0.19 | 0.083 | 3.88003E-29 |
| Pericytes | GATAD2A | 0.000475119 | 0.504689 | 0.22 | 0.087 | 1 |
| Contractile | SREK1IP1 | 1.50118E-30 | 0.504516 | 0.476 | 0.325 | 2.92354E-26 |
| Fibroblasts | CSRNP1 | 0.001802146 | 0.504389 | 0.128 | 0.077 | 1 |
| Intermediate | COL16A1 | 1.27621E-23 | 0.504305 | 0.573 | 0.403 | 2.48543E-19 |
| VEGFA_Terminal | COL5A1 | 2.33548E-17 | 0.50428 | 0.743 | 0.535 | 4.54835E-13 |
| Contractile | CPED1 | 2.0134E-42 | 0.503864 | 0.591 | 0.378 | 3.9211E-38 |
| Contractile | TEAD3 | 8.76223E-47 | 0.503574 | 0.271 | 0.109 | 1.70644E-42 |
| Contractile | IDH2 | 8.90986E-36 | 0.503327 | 0.39 | 0.225 | 1.73519E-31 |
| Pericytes | YWHAG | 0.007665395 | 0.503177 | 0.52 | 0.414 | 1 |
| VEGFA_Terminal | ENSSSCG0( | 1.24158E-31 | 0.502976 | 0.448 | 0.199 | 2.41797E-27 |
| CD74_Intermediat | CD302 | 1.6355E-60 | 0.502938 | 0.636 | 0.428 | 3.18514E-56 |
| VEGFA_Terminal | IARS1 | 6.72788E-19 | 0.502221 | 0.426 | 0.24 | 1.31025E-14 |
| IFN_Intermediate | COL6A3 | 1.6291E-05 | 0.501801 | 0.907 | 0.727 | 0.317267887 |
| Contractile | CHD5 | 8.759E-143 | 0.501617 | 0.153 | 0.005 | 1.7058E-138 |
| Terminal | PAPSS1 | 3.37521E-53 | 0.500947 | 0.56 | 0.373 | 6.57322E-49 |
| Contractile | ENSSSCG0( | 2.93292E-30 | 0.500823 | 0.633 | 0.5 | 5.71187E-26 |
| Terminal | DDIT4 | 4.2556E-41 | 0.500731 | 0.469 | 0.304 | 8.28779E-37 |
| Contractile | HSD17B10 | 1.15675E-31 | 0.500618 | 0.445 | 0.289 | 2.25277E-27 |
| Fibroblasts | FOXO3 | 1.61608E-07 | 0.500617 | 0.615 | 0.526 | 0.00314731 |
| Contractile | YWHAG | 1.16016E-33 | 0.500316 | 0.552 | 0.388 | 2.25941E-29 |
| Contractile | HMG20B | 1.84431E-30 | 0.500312 | 0.426 | 0.272 | 3.59179E-26 |
| Contractile | ACTR2 | 1.17301E-34 | 0.5003 | 0.644 | 0.493 | 2.28443E-30 |
| Contractile | DZIP1 | 7.4334E-30 | 0.500189 | 0.456 | 0.309 | 1.44765E-25 |
| Contractile | BCAP29 | 8.55906E-37 | 0.500186 | 0.657 | 0.509 | 1.66688E-32 |
| Fibroblasts | CREB5 | 2.44225E-06 | 0.499628 | 0.349 | 0.244 | 0.047562847 |
| VEGFA_Terminal | AARS1 | 1.53634E-18 | 0.499441 | 0.332 | 0.165 | 2.99202E-14 |
| Intermediate | COL4A2 | 5.79173E-19 | 0.499361 | 0.691 | 0.573 | 1.12794E-14 |
| IFN_Intermediate | SERPINF1 | 2.07955E-07 | 0.499348 | 0.953 | 0.76 | 0.004049932 |
| VEGFA_Terminal | CLEC14A | 1.42827E-16 | 0.499283 | 0.659 | 0.457 | 2.78156E-12 |
| Contractile | DAP | 1.41047E-34 | 0.499216 | 0.671 | 0.529 | 2.74688E-30 |
| Intermediate | FHL1 | 3.6802E-28 | 0.499177 | 0.862 | 0.745 | 7.1672E-24 |
| VEGFA_Terminal | SEC61G | 1.16027E-22 | 0.499059 | 0.879 | 0.778 | 2.25962E-18 |
| CD74_Intermediat | ISG12(A) | 2.07252E-85 | 0.498718 | 0.836 | 0.586 | 4.03624E-81 |
| VEGFA_Terminal | IMPA2 | 7.78534E-41 | 0.498382 | 0.283 | 0.082 | 1.51619E-36 |
| IFN_Intermediate | ICAM1 | 0.000309614 | 0.498082 | 0.308 | 0.178 | 1 |
| IFN_Intermediate | DHX58 | 7.65605E-34 | 0.498076 | 0.178 | 0.016 | 1.49102E-29 |
| Pericytes | MGAT2 | 0.008066329 | 0.497751 | 0.3 | 0.177 | 1 |
| CD74_Intermediat | IFI6 | 4.5951E-113 | 0.497737 | 0.88 | 0.629 | 8.949E-109 |
| Contractile | AIMP1 | 9.91092E-40 | 0.49726 | 0.752 | 0.617 | 1.93015E-35 |
| IFN_Intermediate | ENSSSCG0( | 3.35472E-11 | 0.497073 | 0.355 | 0.135 | 6.53331E-07 |
| Contractile | NHSL1 | 1.51455E-86 | 0.496616 | 0.212 | 0.04 | 2.94959E-82 |
| Contractile | CABIN1 | 1.46454E-43 | 0.496505 | 0.274 | 0.115 | 2.85219E-39 |

| | | | | | | |
|---|---|---|---|---|---|---|
| Pericytes | MAGI1 | 4.0543E-06 | 0.496108 | 0.14 | 0.03 | 0.078957477 |
| Contractile | TNFRSF21 | 2.6213E-74 | 0.496053 | 0.179 | 0.034 | 5.10499E-70 |
| Fibroblasts | ENSSSCG0( | 7.76364E-05 | 0.495838 | 0.564 | 0.532 | 1 |
| Contractile | ARPC1A | 4.7086E-43 | 0.495743 | 0.423 | 0.233 | 9.17E-39 |
| VEGFA_Terminal | NME2 | 5.32872E-31 | 0.495605 | 0.959 | 0.908 | 1.03777E-26 |
| Contractile | DDT | 1.21947E-29 | 0.495408 | 0.453 | 0.303 | 2.37492E-25 |
| Intermediate | SMAD6 | 9.60724E-16 | 0.495404 | 0.429 | 0.3 | 1.87101E-11 |
| Contractile | SGK1 | 2.21555E-39 | 0.495381 | 0.183 | 0.061 | 4.31478E-35 |
| IFN_Intermediate | FN1 | 1.15104E-06 | 0.495147 | 1 | 0.992 | 0.022416422 |
| VEGFA_Terminal | DARS1 | 1.74197E-21 | 0.494772 | 0.487 | 0.274 | 3.39248E-17 |
| IFN_Intermediate | RSPO3 | 3.30554E-05 | 0.494643 | 0.318 | 0.167 | 0.643753305 |
| VEGFA_Terminal | ETS1 | 5.21418E-18 | 0.494268 | 0.455 | 0.257 | 1.01546E-13 |
| Contractile | CDKN1B | 3.39855E-30 | 0.494236 | 0.485 | 0.328 | 6.61867E-26 |
| IFN_Intermediate | HES4 | 2.56621E-20 | 0.493844 | 0.187 | 0.029 | 4.9977E-16 |
| Terminal | TUBB2A | 6.74916E-44 | 0.493697 | 0.417 | 0.254 | 1.3144E-39 |
| Pericytes | ENSSSCG0( | 4.05913E-05 | 0.493695 | 1 | 0.994 | 0.790516187 |
| Pericytes | EEF1G | 0.001236414 | 0.493206 | 0.88 | 0.838 | 1 |
| Pericytes | MARCHF5 | 0.002620083 | 0.493017 | 0.34 | 0.194 | 1 |
| Contractile | MCL1 | 3.19685E-29 | 0.493012 | 0.674 | 0.546 | 6.22587E-25 |
| Contractile | PPP2R5A | 2.16361E-37 | 0.492463 | 0.374 | 0.207 | 4.21362E-33 |
| Intermediate | MCFD2 | 2.20071E-19 | 0.492397 | 0.535 | 0.395 | 4.28589E-15 |
| Contractile | RAB11FIP2 | 1.37526E-33 | 0.492076 | 0.349 | 0.191 | 2.67833E-29 |
| Terminal | COL12A1 | 5.6552E-35 | 0.492008 | 0.475 | 0.317 | 1.10135E-30 |
| VEGFA_Terminal | DNAJB9 | 6.2343E-13 | 0.491826 | 0.446 | 0.278 | 1.21413E-08 |
| Pericytes | DBNDD2 | 0.0042528 | 0.491604 | 0.42 | 0.266 | 1 |
| Fibroblasts | BRD4 | 3.86871E-06 | 0.491383 | 0.472 | 0.39 | 0.07534313 |
| Intermediate | TMSB4X | 6.12769E-41 | 0.491201 | 1 | 0.999 | 1.19337E-36 |
| Intermediate | COL4A1 | 5.55049E-23 | 0.491135 | 0.8 | 0.681 | 1.08096E-18 |
| Contractile | WASL | 6.36919E-33 | 0.490855 | 0.594 | 0.447 | 1.2404E-28 |
| Fibroblasts | USP47 | 1.17386E-05 | 0.490638 | 0.367 | 0.274 | 0.228609462 |
| Contractile | RGS4 | 2.5151E-46 | 0.490279 | 0.24 | 0.085 | 4.89816E-42 |
| Terminal | ENSSSCG0( | 8.1084E-112 | 0.489788 | 0.997 | 0.985 | 1.5791E-107 |
| Pericytes | AIMP1 | 0.008214136 | 0.489201 | 0.74 | 0.638 | 1 |
| Contractile | ZNRF3 | 1.31211E-73 | 0.489173 | 0.203 | 0.045 | 2.55533E-69 |
| Contractile | RBPMS2 | 4.3351E-132 | 0.488468 | 0.168 | 0.011 | 8.4426E-128 |
| Contractile | KIF5B | 2.14663E-40 | 0.488413 | 0.784 | 0.649 | 4.18055E-36 |
| IFN_Intermediate | EHD4 | 1.30116E-09 | 0.488089 | 0.29 | 0.107 | 2.53401E-05 |
| Fibroblasts | SEMA6A | 3.35044E-80 | 0.488074 | 0.142 | 0.006 | 6.52498E-76 |
| Intermediate | HSPB1 | 2.77206E-12 | 0.48796 | 0.923 | 0.911 | 5.39858E-08 |
| Intermediate | ENSSSCG0( | 1.50336E-14 | 0.487372 | 0.247 | 0.136 | 2.92779E-10 |
| Contractile | ANP32E | 2.5656E-32 | 0.487295 | 0.392 | 0.234 | 4.9965E-28 |
| Pericytes | PPP1R12B | 0.001736312 | 0.487168 | 0.3 | 0.145 | 1 |
| VEGFA_Terminal | VAPA | 1.17315E-18 | 0.486631 | 0.746 | 0.568 | 2.28471E-14 |
| Contractile | CALCOCO1 | 8.96619E-41 | 0.486173 | 0.335 | 0.164 | 1.74617E-36 |
| Contractile | MSRB1 | 5.90829E-34 | 0.48589 | 0.28 | 0.138 | 1.15064E-29 |
| Contractile | ATP1B3 | 2.70307E-35 | 0.485763 | 0.848 | 0.752 | 5.26424E-31 |
| Fibroblasts | NAV1 | 5.10267E-08 | 0.485527 | 0.216 | 0.109 | 0.000993745 |
| Contractile | PDCD5 | 1.61882E-39 | 0.485229 | 0.752 | 0.614 | 3.15265E-35 |
| Fibroblasts | EHBP1 | 2.34585E-08 | 0.485073 | 0.266 | 0.144 | 0.000456853 |
| Intermediate | MAP1B | 7.0621E-20 | 0.484965 | 0.889 | 0.822 | 1.37534E-15 |
| CD74_Intermediat | ENSSSCG0( | 2.23435E-72 | 0.484869 | 0.878 | 0.712 | 4.3514E-68 |
| Contractile | PFKFB3 | 1.80699E-48 | 0.484777 | 0.225 | 0.076 | 3.51911E-44 |
| Contractile | A4GALT | 6.056E-27 | 0.484642 | 0.505 | 0.37 | 1.17941E-22 |
| VEGFA_Terminal | MMP14 | 1.80517E-12 | 0.484287 | 0.55 | 0.366 | 3.51558E-08 |
| Contractile | CPT1B | 2.6306E-119 | 0.484088 | 0.162 | 0.012 | 5.1232E-115 |
| Intermediate | QSOX1 | 4.09906E-16 | 0.484015 | 0.551 | 0.424 | 7.98292E-12 |
| Pericytes | SLC25A3 | 0.000585999 | 0.483818 | 0.9 | 0.882 | 1 |
| IFN_Intermediate | LIMA1 | 1.37951E-05 | 0.483626 | 0.664 | 0.488 | 0.268659681 |
| Contractile | PRMT1 | 1.21812E-31 | 0.483238 | 0.516 | 0.361 | 2.37229E-27 |

| | | | | | | |
|---|---|---|---|---|---|---|
| Pericytes | TPM2 | 1.0773E-06 | 0.483053 | 0.96 | 0.847 | 0.020980503 |
| Pericytes | ENSSSCG0( | 1.35154E-38 | 0.483025 | 0.12 | 0.003 | 2.63213E-34 |
| Fibroblasts | PCDHGA4 | 0.003300353 | 0.482687 | 0.408 | 0.382 | 1 |
| VEGFA_Terminal | ZMIZ1 | 9.3172E-14 | 0.482686 | 0.458 | 0.288 | 1.81452E-09 |
| VEGFA_Terminal | VDAC2 | 1.44114E-15 | 0.482683 | 0.685 | 0.531 | 2.80663E-11 |
| Contractile | ACYP2 | 2.23137E-53 | 0.482664 | 0.239 | 0.08 | 4.3456E-49 |
| VEGFA_Terminal | SELENOM | 1.14999E-20 | 0.482396 | 0.879 | 0.727 | 2.2396E-16 |
| CD74_Intermediat | UACA | 1.2746E-43 | 0.482293 | 0.569 | 0.392 | 2.48229E-39 |
| Fibroblasts | SNRNP70 | 0.000593271 | 0.482274 | 0.376 | 0.315 | 1 |
| Pericytes | TNS3 | 1.18312E-05 | 0.482114 | 0.16 | 0.04 | 0.230412025 |
| Pericytes | MKNK2 | 0.006844539 | 0.481761 | 0.26 | 0.14 | 1 |
| Pericytes | HRC | 1.13663E-22 | 0.4817 | 0.12 | 0.006 | 2.21358E-18 |
| Intermediate | ID4 | 1.00123E-07 | 0.481622 | 0.47 | 0.395 | 0.001949891 |
| Fibroblasts | SPRED1 | 0.000174559 | 0.481613 | 0.294 | 0.219 | 1 |
| VEGFA_Terminal | CEMIP | 5.33333E-26 | 0.481101 | 0.177 | 0.05 | 1.03867E-21 |
| Fibroblasts | RAB3IL1 | 1.8495E-14 | 0.480738 | 0.17 | 0.053 | 3.6019E-10 |
| Fibroblasts | STAG1 | 3.83132E-06 | 0.480626 | 0.468 | 0.371 | 0.074615037 |
| Fibroblasts | SPRY2 | 7.59975E-06 | 0.480494 | 0.211 | 0.123 | 0.148005225 |
| VEGFA_Terminal | PTPN2 | 2.89022E-23 | 0.48021 | 0.402 | 0.197 | 5.6287E-19 |
| VEGFA_Terminal | PNP | 1.59271E-27 | 0.479912 | 0.186 | 0.054 | 3.1018E-23 |
| Contractile | NOL7 | 1.17861E-42 | 0.479826 | 0.791 | 0.657 | 2.29535E-38 |
| Contractile | ENAH | 4.67346E-29 | 0.479732 | 0.406 | 0.254 | 9.10157E-25 |
| CD74_Intermediat | CLEC14A | 6.28112E-54 | 0.479416 | 0.614 | 0.413 | 1.22325E-49 |
| Pericytes | ECE1 | 0.000139777 | 0.479154 | 0.18 | 0.057 | 1 |
| VEGFA_Terminal | TPD52L2 | 4.73965E-18 | 0.479 | 0.586 | 0.38 | 9.23046E-14 |
| Fibroblasts | IER5L | 0.000153402 | 0.478629 | 0.289 | 0.207 | 1 |
| Fibroblasts | ALDH6A1 | 1.36933E-11 | 0.478593 | 0.202 | 0.08 | 2.66677E-07 |
| Fibroblasts | TMX4 | 3.99761E-06 | 0.478578 | 0.477 | 0.39 | 0.077853389 |
| Contractile | YPEL2 | 4.06889E-31 | 0.478474 | 0.402 | 0.244 | 7.92415E-27 |
| VEGFA_Terminal | PXDC1 | 5.467E-12 | 0.478306 | 0.571 | 0.403 | 1.0647E-07 |
| Contractile | EIF4B | 1.36336E-33 | 0.477549 | 0.64 | 0.477 | 2.65513E-29 |
| Intermediate | AEBP1 | 8.94611E-33 | 0.477127 | 0.949 | 0.881 | 1.74225E-28 |
| CD74_Intermediat | LRRC17 | 1.04548E-56 | 0.476853 | 0.565 | 0.354 | 2.03607E-52 |
| Intermediate | NEDD9 | 5.37472E-15 | 0.476779 | 0.353 | 0.226 | 1.04673E-10 |
| Fibroblasts | ORAI1 | 2.7729E-08 | 0.476147 | 0.22 | 0.11 | 0.000540022 |
| CD74_Intermediat | INMT | 4.79077E-16 | 0.298717 | 0.151 | 0.083 | 9.33002E-12 |
| VEGFA_Terminal | FTH1 | 1.64704E-30 | 0.476041 | 0.998 | 0.998 | 3.2076E-26 |
| Contractile | BEX3 | 3.57357E-31 | 0.475172 | 0.337 | 0.184 | 6.95953E-27 |
| Contractile | LIX1L | 2.31419E-27 | 0.474986 | 0.373 | 0.236 | 4.50688E-23 |
| Contractile | RTN4 | 1.02671E-41 | 0.474944 | 0.856 | 0.764 | 1.99953E-37 |
| VEGFA_Terminal | PPP1R14B | 2.60334E-15 | 0.474221 | 0.656 | 0.469 | 5.07E-11 |
| CD74_Intermediat | LAMB1 | 5.5924E-66 | 0.47391 | 0.797 | 0.591 | 1.08912E-61 |
| Contractile | DEK | 3.65309E-54 | 0.473699 | 0.915 | 0.858 | 7.11439E-50 |
| Pericytes | GVQW3 | 0.000737027 | 0.473632 | 0.16 | 0.054 | 1 |
| Contractile | TET1 | 3.37032E-38 | 0.47331 | 0.28 | 0.127 | 6.56369E-34 |
| Fibroblasts | ENSSSCG0( | 3.5857E-43 | 0.473193 | 0.101 | 0.007 | 6.98314E-39 |
| CD74_Intermediat | SRGN | 3.7634E-77 | 0.603324 | 0.264 | 0.085 | 7.32923E-73 |
| VEGFA_Terminal | NAMPT | 2.35678E-19 | 0.47273 | 0.46 | 0.256 | 4.58984E-15 |
| Contractile | CUL1 | 6.8016E-29 | 0.472425 | 0.41 | 0.257 | 1.32461E-24 |
| Pericytes | NOB1 | 0.000108696 | 0.472322 | 0.18 | 0.056 | 1 |
| Contractile | SH3PXD2A | 1.52017E-33 | 0.472097 | 0.308 | 0.159 | 2.96052E-29 |
| Terminal | PERP | 6.0775E-89 | 0.471682 | 0.162 | 0.025 | 1.18359E-84 |
| Contractile | DSG2 | 2.4665E-112 | 0.471524 | 0.158 | 0.013 | 4.8035E-108 |
| Terminal | PRDX1 | 5.14372E-46 | 0.471298 | 0.702 | 0.55 | 1.00174E-41 |
| Contractile | GPC6 | 8.98936E-38 | 0.471151 | 0.278 | 0.125 | 1.75068E-33 |
| VEGFA_Terminal | COX7A2L | 3.21139E-20 | 0.470604 | 0.736 | 0.522 | 6.25419E-16 |
| Terminal | PLOD2 | 6.02073E-43 | 0.470567 | 0.668 | 0.515 | 1.17254E-38 |
| Intermediate | MACF1 | 1.57828E-11 | 0.470263 | 0.473 | 0.375 | 3.07371E-07 |
| Contractile | CALM2 | 7.90356E-71 | 0.47011 | 0.969 | 0.94 | 1.53922E-66 |

| | | | | | | |
|---|---|---|---|---|---|---|
| VEGFA_Terminal | IDH2 | 7.18343E-23 | 0.469505 | 0.46 | 0.236 | 1.39897E-18 |
| Fibroblasts | LIFR | 2.85952E-07 | 0.469385 | 0.17 | 0.079 | 0.005568913 |
| Fibroblasts | ARHGAP29 | 2.24632E-06 | 0.469303 | 0.353 | 0.243 | 0.043747116 |
| Contractile | ENSSSCG0( | 6.96421E-44 | 0.469298 | 0.321 | 0.146 | 1.35628E-39 |
| CD74_Intermediat | CFI | 3.61254E-34 | 0.329342 | 0.198 | 0.088 | 7.03542E-30 |
| Pericytes | ZNF618 | 9.78062E-05 | 0.469055 | 0.12 | 0.028 | 1 |
| Terminal | LDHA | 1.61366E-75 | 0.468997 | 0.83 | 0.646 | 3.14259E-71 |
| VEGFA_Terminal | RAI14 | 2.07486E-15 | 0.468879 | 0.542 | 0.355 | 4.04079E-11 |
| VEGFA_Terminal | SMAP2 | 3.61453E-24 | 0.468838 | 0.404 | 0.195 | 7.0393E-20 |
| VEGFA_Terminal | SOCS5 | 2.0911E-18 | 0.46881 | 0.431 | 0.24 | 4.07242E-14 |
| Fibroblasts | ZBTB44 | 1.16031E-08 | 0.468495 | 0.339 | 0.201 | 0.00022597 |
| CD74_Intermediat | DPT | 1.3305E-44 | 0.468126 | 0.521 | 0.331 | 2.59114E-40 |
| IFN_Intermediate | USP15 | 0.000148428 | 0.467944 | 0.383 | 0.233 | 1 |
| Pericytes | TTPAL | 0.000222735 | 0.467921 | 0.1 | 0.023 | 1 |
| Pericytes | ENSSSCG0( | 0.000866354 | 0.467906 | 0.12 | 0.034 | 1 |
| Contractile | EIF4EBP2 | 2.08124E-31 | 0.467898 | 0.546 | 0.379 | 4.05321E-27 |
| Contractile | LGR4 | 3.23459E-75 | 0.467523 | 0.213 | 0.047 | 6.29937E-71 |
| Terminal | SDC2 | 1.88812E-57 | 0.467382 | 0.88 | 0.755 | 3.67711E-53 |
| Pericytes | ENSSSCG0( | 3.71502E-07 | 0.467251 | 0.16 | 0.032 | 0.007235002 |
| Contractile | ETS2 | 4.19996E-40 | 0.467181 | 0.314 | 0.148 | 8.17942E-36 |
| VEGFA_Terminal | TRPC4 | 7.55789E-31 | 0.467179 | 0.324 | 0.121 | 1.4719E-26 |
| Contractile | OCIAD1 | 6.75562E-33 | 0.466463 | 0.68 | 0.545 | 1.31566E-28 |
| Terminal | MIF | 3.0568E-77 | 0.465835 | 0.931 | 0.832 | 5.95313E-73 |
| Fibroblasts | COL14A1 | 1.16135E-11 | 0.465663 | 0.908 | 0.808 | 2.26173E-07 |
| Fibroblasts | ENSSSCG0( | 1.02641E-12 | 0.465429 | 0.248 | 0.103 | 1.99893E-08 |
| Contractile | RSF1 | 4.1158E-26 | 0.465335 | 0.525 | 0.385 | 8.01552E-22 |
| Fibroblasts | MAFB | 0.001424993 | 0.465313 | 0.216 | 0.149 | 1 |
| Terminal | CDA | 6.02619E-98 | 0.465152 | 0.134 | 0.01 | 1.1736E-93 |
| Intermediate | NPNT | 2.58828E-27 | 0.464199 | 0.473 | 0.27 | 5.04068E-23 |
| Pericytes | NOLC1 | 0.008767204 | 0.463659 | 0.32 | 0.193 | 1 |
| Contractile | CAT | 4.37562E-28 | 0.463606 | 0.477 | 0.321 | 8.52151E-24 |
| Contractile | CREBRF | 2.1186E-28 | 0.463263 | 0.425 | 0.274 | 4.12598E-24 |
| Intermediate | REXO2 | 3.87075E-27 | 0.462848 | 0.886 | 0.794 | 7.53829E-23 |
| Intermediate | NR2F2 | 2.8261E-15 | 0.462602 | 0.41 | 0.281 | 5.50384E-11 |
| VEGFA_Terminal | COL5A2 | 7.38461E-19 | 0.462547 | 0.867 | 0.753 | 1.43815E-14 |
| Fibroblasts | PLBD2 | 1.26114E-07 | 0.462433 | 0.422 | 0.296 | 0.002456071 |
| VEGFA_Terminal | PDLIM4 | 3.9697E-31 | 0.462396 | 0.303 | 0.109 | 7.731E-27 |
| Pericytes | LDHB | 0.007956406 | 0.462124 | 0.72 | 0.709 | 1 |
| Fibroblasts | BICC1 | 9.41493E-06 | 0.461412 | 0.312 | 0.213 | 0.183355852 |
| Intermediate | MMP23B | 9.54687E-21 | 0.460781 | 0.648 | 0.5 | 1.85925E-16 |
| Contractile | CACNA2D1 | 3.84235E-26 | 0.460746 | 0.418 | 0.274 | 7.48299E-22 |
| Pericytes | PFDN2 | 0.007540347 | 0.460737 | 0.62 | 0.518 | 1 |
| CD74_Intermediat | ACKR2 | 5.75051E-30 | 0.276935 | 0.193 | 0.089 | 1.11991E-25 |
| Fibroblasts | PAPLN | 2.71622E-06 | 0.460242 | 0.183 | 0.095 | 0.052898316 |
| Pericytes | DLX5 | 1.04474E-11 | 0.459722 | 0.14 | 0.016 | 2.03464E-07 |
| Contractile | PRKAR1A | 6.72276E-32 | 0.459685 | 0.681 | 0.559 | 1.30926E-27 |
| Contractile | MARCHF6 | 6.40874E-26 | 0.459558 | 0.471 | 0.328 | 1.2481E-21 |
| VEGFA_Terminal | OPTN | 1.67018E-11 | 0.45955 | 0.746 | 0.639 | 3.25268E-07 |
| Fibroblasts | PELI1 | 0.000223691 | 0.45925 | 0.206 | 0.133 | 1 |
| Fibroblasts | HS6ST1 | 2.89567E-12 | 0.459081 | 0.188 | 0.068 | 5.63932E-08 |
| VEGFA_Terminal | NDUFV2 | 3.52908E-19 | 0.45905 | 0.671 | 0.469 | 6.87288E-15 |
| CD74_Intermediat | FNDC3A | 3.34507E-44 | 0.458971 | 0.594 | 0.428 | 6.51452E-40 |
| VEGFA_Terminal | PAPSS2 | 6.6999E-25 | 0.458926 | 0.324 | 0.138 | 1.30481E-20 |
| CD74_Intermediat | C2 | 2.1238E-135 | 0.730432 | 0.349 | 0.091 | 4.1362E-131 |
| Fibroblasts | NEO1 | 4.2385E-05 | 0.457846 | 0.193 | 0.112 | 0.825447385 |
| Fibroblasts | DHX9 | 0.000161949 | 0.457559 | 0.45 | 0.386 | 1 |
| VEGFA_Terminal | PPP1R15A | 2.03199E-26 | 0.457557 | 0.562 | 0.294 | 3.9573E-22 |
| Contractile | LTA4H | 4.77362E-28 | 0.457514 | 0.392 | 0.245 | 9.29662E-24 |
| Pericytes | PHAF1 | 0.008425592 | 0.457442 | 0.14 | 0.056 | 1 |

| | | | | | | |
|---|---|---|---|---|---|---|
| VEGFA_Terminal | EDIL3 | 1.04911E-19 | 0.457373 | 0.632 | 0.391 | 2.04314E-15 |
| Terminal | BHLHE41 | 8.73135E-33 | 0.456907 | 0.478 | 0.329 | 1.70043E-28 |
| Fibroblasts | NTRK2 | 3.9187E-08 | 0.456803 | 0.216 | 0.107 | 0.000763167 |
| Fibroblasts | ENSSSCG00 | 0.000129231 | 0.456172 | 0.183 | 0.111 | 1 |
| Fibroblasts | ENSSSCG00 | 5.756E-07 | 0.456091 | 0.587 | 0.5 | 0.011209816 |
| IFN_Intermediate | ENPP4 | 6.60522E-12 | 0.455968 | 0.15 | 0.031 | 1.28637E-07 |
| Terminal | ANXA5 | 5.32978E-80 | 0.455677 | 0.983 | 0.927 | 1.03797E-75 |
| Fibroblasts | ARAP1 | 2.60253E-05 | 0.45544 | 0.211 | 0.127 | 0.506842546 |
| Terminal | CTSD | 9.81988E-57 | 0.455072 | 0.944 | 0.873 | 1.91242E-52 |
| Fibroblasts | CPT1A | 7.4977E-07 | 0.455043 | 0.225 | 0.124 | 0.014601764 |
| CD74_Intermediat | ENSSSCG00 | 4.88225E-47 | 0.45477 | 0.777 | 0.629 | 9.50819E-43 |
| Intermediate | STON1 | 1.06139E-15 | 0.453785 | 0.444 | 0.311 | 2.06706E-11 |
| CD74_Intermediat | CTTNBP2 | 7.33295E-33 | 0.348616 | 0.204 | 0.094 | 1.42809E-28 |
| IFN_Intermediate | NBN | 3.01914E-06 | 0.453455 | 0.355 | 0.183 | 0.058797735 |
| CD74_Intermediat | ECRG4 | 1.3638E-71 | 1.143935 | 0.265 | 0.095 | 2.65601E-67 |
| Pericytes | ENSSSCG00 | 0.004664447 | 0.452833 | 0.12 | 0.041 | 1 |
| Contractile | PATJ | 1.05194E-41 | 0.452676 | 0.227 | 0.085 | 2.04866E-37 |
| IFN_Intermediate | MFSD12 | 1.35027E-11 | 0.452598 | 0.14 | 0.028 | 2.62965E-07 |
| Fibroblasts | DNM1 | 3.90463E-20 | 0.452563 | 0.124 | 0.023 | 7.60427E-16 |
| Fibroblasts | CPD | 4.41482E-05 | 0.452185 | 0.358 | 0.269 | 0.859786262 |
| Fibroblasts | ENSSSCG00 | 0.002934585 | 0.452148 | 0.248 | 0.176 | 1 |
| Terminal | ANXA1 | 3.00663E-49 | 0.452115 | 0.835 | 0.703 | 5.8554E-45 |
| Contractile | UBE2H | 2.6947E-25 | 0.451958 | 0.568 | 0.435 | 5.24792E-21 |
| CD74_Intermediat | ASAH1 | 5.80054E-54 | 0.451867 | 0.526 | 0.323 | 1.12965E-49 |
| Fibroblasts | BCLAF1 | 0.000105652 | 0.451856 | 0.5 | 0.439 | 1 |
| Fibroblasts | EIF4A2 | 1.53906E-09 | 0.4518 | 0.807 | 0.802 | 2.99733E-05 |
| Intermediate | GJA1 | 2.64853E-07 | 0.451774 | 0.608 | 0.57 | 0.005158013 |
| Fibroblasts | RGMB | 1.15261E-07 | 0.451719 | 0.142 | 0.059 | 0.002244716 |
| Contractile | PTGER3 | 3.7716E-108 | 0.451501 | 0.124 | 0.006 | 7.3452E-104 |
| Fibroblasts | DENND4C | 0.000861048 | 0.451212 | 0.28 | 0.212 | 1 |
| Fibroblasts | PGRMC1 | 9.08993E-06 | 0.451157 | 0.454 | 0.373 | 0.177026466 |
| Fibroblasts | HIC1 | 4.53632E-11 | 0.451007 | 0.17 | 0.063 | 8.83448E-07 |
| IFN_Intermediate | STAT3 | 0.000171233 | 0.450433 | 0.57 | 0.392 | 1 |
| Terminal | TMEM45A | 3.54909E-54 | 0.450198 | 0.373 | 0.191 | 6.91186E-50 |
| VEGFA_Terminal | CFDP1 | 5.86191E-17 | 0.450039 | 0.797 | 0.61 | 1.14161E-12 |
| CD74_Intermediat | ENSSSCG00 | 8.74677E-67 | 0.460485 | 0.263 | 0.095 | 1.70343E-62 |
| Contractile | CCND1 | 5.94625E-25 | 0.44967 | 0.598 | 0.459 | 1.15803E-20 |
| Fibroblasts | MEIS1 | 0.000112433 | 0.449488 | 0.28 | 0.199 | 1 |
| VEGFA_Terminal | SSR4 | 9.54041E-14 | 0.449418 | 0.826 | 0.73 | 1.858E-09 |
| Fibroblasts | DNAI7 | 3.89952E-08 | 0.449281 | 0.202 | 0.097 | 0.000759431 |
| Fibroblasts | PLAU | 5.23586E-09 | 0.44893 | 0.115 | 0.037 | 0.000101968 |
| Contractile | STAU2 | 1.0758E-33 | 0.448748 | 0.29 | 0.144 | 2.09511E-29 |
| Contractile | MMP28 | 1.65647E-54 | 0.448584 | 0.217 | 0.066 | 3.22598E-50 |
| VEGFA_Terminal | PLEKHO1 | 1.67266E-14 | 0.448551 | 0.281 | 0.143 | 3.25751E-10 |
| Fibroblasts | RUNX1T1 | 2.40045E-05 | 0.448055 | 0.261 | 0.167 | 0.467487774 |
| Fibroblasts | GAS7 | 2.63971E-14 | 0.447943 | 0.188 | 0.062 | 5.14084E-10 |
| Pericytes | GZF1 | 4.21182E-07 | 0.447509 | 0.14 | 0.026 | 0.008202516 |
| Contractile | FZD7 | 4.27941E-32 | 0.447143 | 0.297 | 0.151 | 8.33416E-28 |
| CD74_Intermediat | CYP27A1 | 6.58216E-18 | 0.271112 | 0.173 | 0.095 | 1.28188E-13 |
| Fibroblasts | SCARB2 | 3.01862E-06 | 0.446251 | 0.656 | 0.616 | 0.058787536 |
| Intermediate | LRRC17 | 1.09299E-16 | 0.446019 | 0.547 | 0.398 | 2.1286E-12 |
| IFN_Intermediate | IRF1 | 0.00035382 | 0.445601 | 0.178 | 0.084 | 1 |
| Intermediate | FILIP1 | 6.20135E-12 | 0.44545 | 0.409 | 0.292 | 1.20771E-07 |
| Intermediate | STK17A | 6.14169E-20 | 0.445403 | 0.393 | 0.239 | 1.19609E-15 |
| VEGFA_Terminal | YWHAE | 6.15308E-18 | 0.445158 | 0.891 | 0.766 | 1.19831E-13 |
| Terminal | PKM | 1.77975E-85 | 0.445105 | 0.943 | 0.846 | 3.46606E-81 |
| Pericytes | SPHK1 | 1.13114E-18 | 0.444947 | 0.12 | 0.007 | 2.2029E-14 |
| Terminal | C16ORF74 | 8.10671E-49 | 0.444921 | 0.31 | 0.151 | 1.57878E-44 |
| CD74_Intermediat | EPAS1 | 7.75944E-50 | 0.444606 | 0.703 | 0.522 | 1.51115E-45 |

| | | | | | | |
|---|---|---|---|---|---|---|
| Contractile | MINDY2 | 1.81562E-39 | 0.444379 | 0.274 | 0.12 | 3.53592E-35 |
| Terminal | ALDH2 | 1.32491E-39 | 0.444092 | 0.665 | 0.497 | 2.58026E-35 |
| Fibroblasts | UBE3A | 0.000563862 | 0.444086 | 0.353 | 0.291 | 1 |
| Pericytes | TUBA1B | 0.009856538 | 0.443787 | 0.76 | 0.712 | 1 |
| Pericytes | BCL7A | 0.00482508 | 0.443578 | 0.16 | 0.063 | 1 |
| CD74_Intermediat | KCTD12 | 1.42436E-75 | 0.504764 | 0.289 | 0.1 | 2.77395E-71 |
| Intermediate | PROS1 | 1.17242E-19 | 0.443374 | 0.696 | 0.559 | 2.28328E-15 |
| Pericytes | FGF13 | 2.89498E-50 | 0.443306 | 0.14 | 0.003 | 5.63797E-46 |
| Contractile | PROM1 | 3.9646E-128 | 0.443212 | 0.152 | 0.008 | 7.7211E-124 |
| Pericytes | ADCY6 | 0.001440758 | 0.443114 | 0.2 | 0.081 | 1 |
| Fibroblasts | EDA | 2.79579E-73 | 0.442999 | 0.106 | 0.003 | 5.44481E-69 |
| Contractile | SEZ6L | 8.28913E-72 | 0.442973 | 0.177 | 0.033 | 1.61431E-67 |
| Pericytes | THSD7A | 5.47681E-05 | 0.442968 | 0.12 | 0.027 | 1 |
| Fibroblasts | JAK1 | 4.20003E-08 | 0.442946 | 0.748 | 0.714 | 0.000817956 |
| Pericytes | ANGPT1 | 2.30098E-06 | 0.442723 | 0.2 | 0.052 | 0.044811674 |
| Intermediate | GEM | 1.34186E-18 | 0.442609 | 0.75 | 0.616 | 2.61327E-14 |
| IFN_Intermediate | SLA-8 | 0.000363179 | 0.442604 | 0.449 | 0.299 | 1 |
| Pericytes | COTL1 | 0.000106136 | 0.442511 | 0.14 | 0.036 | 1 |
| Terminal | COL14A1 | 4.88658E-46 | 0.442345 | 0.931 | 0.755 | 9.51662E-42 |
| Contractile | ENSSSCG0( | 1.0143E-119 | 0.442299 | 0.165 | 0.013 | 1.9753E-115 |
| Fibroblasts | PTGFRN | 1.10825E-05 | 0.442189 | 0.248 | 0.155 | 0.215832646 |
| Fibroblasts | FUS | 4.29844E-05 | 0.442051 | 0.349 | 0.257 | 0.837120846 |
| Contractile | WAC | 1.9535E-23 | 0.441685 | 0.435 | 0.303 | 3.80444E-19 |
| Terminal | AP3S1 | 6.72603E-50 | 0.441306 | 0.576 | 0.395 | 1.30989E-45 |
| Fibroblasts | ACSF2 | 1.98046E-05 | 0.441176 | 0.317 | 0.221 | 0.385693617 |
| Contractile | TSPAN7 | 2.7186E-141 | 0.441097 | 0.146 | 0.004 | 5.2945E-137 |
| Contractile | SEMA3D | 1.36012E-37 | 0.441086 | 0.195 | 0.07 | 2.64884E-33 |
| Pericytes | EIF4A2 | 0.00103062 | 0.441031 | 0.82 | 0.802 | 1 |
| VEGFA_Terminal | PHGDH | 1.95194E-40 | 0.441023 | 0.184 | 0.038 | 3.8014E-36 |
| Fibroblasts | PTPN9 | 0.000826085 | 0.441001 | 0.216 | 0.152 | 1 |
| IFN_Intermediate | SCARB2 | 5.31944E-06 | 0.440981 | 0.766 | 0.615 | 0.10359603 |
| Contractile | HIP1 | 5.75626E-24 | 0.440876 | 0.328 | 0.198 | 1.12103E-19 |
| VEGFA_Terminal | RUNX2 | 2.6042E-30 | 0.44053 | 0.235 | 0.072 | 5.07169E-26 |
| VEGFA_Terminal | RCN3 | 4.21289E-19 | 0.44052 | 0.683 | 0.459 | 8.20461E-15 |
| VEGFA_Terminal | NRP1 | 3.31705E-11 | 0.440502 | 0.528 | 0.363 | 6.45995E-07 |
| Fibroblasts | CHD2 | 3.90149E-07 | 0.440306 | 0.298 | 0.179 | 0.007598157 |
| Intermediate | SPARCL1 | 3.12481E-24 | 0.440226 | 0.843 | 0.691 | 6.08557E-20 |
| Intermediate | SMOC1 | 7.94601E-13 | 0.44019 | 0.621 | 0.527 | 1.54749E-08 |
| Intermediate | APOA1 | 8.10328E-24 | 0.440147 | 0.249 | 0.113 | 1.57811E-19 |
| Contractile | ERBIN | 5.24121E-26 | 0.440083 | 0.579 | 0.44 | 1.02073E-21 |
| Contractile | ASH1L | 6.28701E-26 | 0.439892 | 0.676 | 0.554 | 1.22439E-21 |
| VEGFA_Terminal | BCL2A1 | 2.13131E-35 | 0.43988 | 0.228 | 0.062 | 4.15073E-31 |
| Terminal | BGN | 1.70125E-81 | 0.439358 | 0.998 | 0.986 | 3.31318E-77 |
| Fibroblasts | BIN1 | 0.002296599 | 0.439342 | 0.261 | 0.201 | 1 |
| Fibroblasts | ENSSSCG0( | 0.000159843 | 0.439229 | 0.427 | 0.35 | 1 |
| Terminal | P4HA2 | 1.85512E-46 | 0.439216 | 0.623 | 0.451 | 3.61284E-42 |
| Intermediate | LIMCH1 | 2.1699E-17 | 0.439115 | 0.601 | 0.451 | 4.22588E-13 |
| Terminal | FAM180A | 2.47185E-34 | 0.439034 | 0.281 | 0.152 | 4.81393E-30 |
| VEGFA_Terminal | WNT5B | 1.68754E-28 | 0.438939 | 0.341 | 0.135 | 3.28649E-24 |
| Fibroblasts | CCNG2 | 8.52317E-06 | 0.438923 | 0.225 | 0.133 | 0.165988755 |
| Fibroblasts | ADA | 9.11745E-08 | 0.438718 | 0.188 | 0.089 | 0.001775624 |
| CD74_Intermediat | GXYLT2 | 3.57678E-42 | 0.438509 | 0.56 | 0.381 | 6.96577E-38 |
| Pericytes | KLHL21 | 0.006786183 | 0.438254 | 0.2 | 0.093 | 1 |
| VEGFA_Terminal | OSTC | 2.771E-18 | 0.438072 | 0.598 | 0.381 | 5.39652E-14 |
| Fibroblasts | GXYLT1 | 1.81549E-05 | 0.437563 | 0.156 | 0.081 | 0.353567455 |
| Pericytes | ENSSSCG0( | 0.003569773 | 0.437493 | 1 | 0.997 | 1 |
| Pericytes | ENSSSCG0( | 7.83631E-06 | 0.437446 | 0.12 | 0.023 | 0.152612083 |
| Intermediate | C1ORF54 | 9.65037E-21 | 0.437236 | 0.349 | 0.2 | 1.87941E-16 |
| Contractile | NDUFB9 | 5.92582E-26 | 0.437029 | 0.681 | 0.578 | 1.15405E-21 |

| | | | | | | |
|---|---|---|---|---|---|---|
| Pericytes | TULP3 | 0.001291765 | 0.436811 | 0.26 | 0.122 | 1 |
| IFN_Intermediate | ABI3BP | 0.000220596 | 0.436802 | 0.636 | 0.506 | 1 |
| Contractile | FAM168B | 9.6578E-30 | 0.436083 | 0.347 | 0.197 | 1.88086E-25 |
| Terminal | SERTAD4 | 4.7242E-46 | 0.435785 | 0.272 | 0.128 | 9.20037E-42 |
| Terminal | CD59 | 3.32338E-40 | 0.435765 | 0.716 | 0.586 | 6.47229E-36 |
| Contractile | CBX6 | 2.032E-33 | 0.435664 | 0.273 | 0.128 | 3.95733E-29 |
| Contractile | ITGB1 | 3.22553E-60 | 0.435641 | 0.995 | 0.977 | 6.28173E-56 |
| IFN_Intermediate | UGT1A6 | 0.000527786 | 0.435614 | 0.168 | 0.08 | 1 |
| VEGFA_Terminal | PLAUR | 3.30129E-21 | 0.435443 | 0.242 | 0.096 | 6.42926E-17 |
| Fibroblasts | UST | 1.02051E-16 | 0.435376 | 0.17 | 0.047 | 1.98744E-12 |
| CD74_Intermediat | CHST7 | 2.12657E-21 | 0.259272 | 0.192 | 0.104 | 4.1415E-17 |
| Fibroblasts | ITPR2 | 9.59876E-06 | 0.435312 | 0.248 | 0.153 | 0.186935912 |
| CD74_Intermediat | PTGS1 | 3.96265E-24 | 0.261649 | 0.203 | 0.105 | 7.71725E-20 |
| Contractile | WIPF1 | 5.94018E-24 | 0.435171 | 0.426 | 0.289 | 1.15685E-19 |
| Contractile | ST6GAL2 | 2.8274E-114 | 0.435099 | 0.164 | 0.014 | 5.5063E-110 |
| Contractile | HIP1R | 2.24239E-57 | 0.435007 | 0.17 | 0.04 | 4.36705E-53 |
| Contractile | EIF3H | 5.42407E-37 | 0.434998 | 0.851 | 0.773 | 1.05634E-32 |
| Contractile | CACNA1H | 5.3472E-128 | 0.434996 | 0.134 | 0.004 | 1.0414E-123 |
| Pericytes | FRY | 0.00019489 | 0.434954 | 0.18 | 0.058 | 1 |
| CD74_Intermediat | LAMP2 | 2.75041E-56 | 0.434773 | 0.827 | 0.695 | 5.35642E-52 |
| Intermediate | ENSSSCG0( | 3.50664E-26 | 0.43457 | 0.756 | 0.61 | 6.82918E-22 |
| Contractile | PEA15 | 5.58152E-27 | 0.434562 | 0.54 | 0.393 | 1.087E-22 |
| Contractile | NPM1 | 4.87094E-38 | 0.434408 | 0.877 | 0.819 | 9.48615E-34 |
| Contractile | ARHGAP29 | 2.32727E-30 | 0.434327 | 0.378 | 0.222 | 4.53236E-26 |
| CD74_Intermediat | MYLIP | 5.34253E-36 | 0.434256 | 0.544 | 0.388 | 1.04046E-31 |
| Terminal | CYP7A1 | 1.21603E-46 | 0.434093 | 0.136 | 0.038 | 2.36822E-42 |
| Fibroblasts | LDLRAD3 | 2.09946E-10 | 0.433435 | 0.165 | 0.061 | 4.0887E-06 |
| Contractile | SRSF3 | 1.18906E-24 | 0.433329 | 0.665 | 0.546 | 2.3157E-20 |
| Fibroblasts | RASSF8 | 0.003344943 | 0.433244 | 0.298 | 0.244 | 1 |
| Fibroblasts | ENSSSCG0( | 1.41705E-05 | 0.432685 | 0.771 | 0.759 | 0.275970655 |
| Fibroblasts | GXYLT2 | 0.001315615 | 0.432684 | 0.472 | 0.431 | 1 |
| Pericytes | ENSSSCG0( | 0.004539973 | 0.432416 | 0.12 | 0.041 | 1 |
| Contractile | UBE2D2 | 2.95239E-26 | 0.432404 | 0.597 | 0.471 | 5.74978E-22 |
| VEGFA_Terminal | EMP3 | 1.22223E-18 | 0.432169 | 0.913 | 0.824 | 2.3803E-14 |
| Fibroblasts | PDE3B | 3.39023E-07 | 0.431985 | 0.188 | 0.092 | 0.00660247 |
| VEGFA_Terminal | KLF4 | 3.76016E-13 | 0.431959 | 0.45 | 0.279 | 7.32292E-09 |
| VEGFA_Terminal | ALDH1A3 | 1.07424E-05 | 0.431941 | 0.242 | 0.162 | 0.20920839 |
| Fibroblasts | SLIT2 | 9.39218E-11 | 0.431571 | 0.128 | 0.04 | 1.82913E-06 |
| Contractile | PLPPR4 | 8.5136E-138 | 0.431468 | 0.134 | 0.003 | 1.658E-133 |
| IFN_Intermediate | SLC7A2 | 0.001199066 | 0.431467 | 0.355 | 0.23 | 1 |
| CD74_Intermediat | ABI3BP | 3.05786E-49 | 0.431353 | 0.658 | 0.448 | 5.95519E-45 |
| Contractile | AEBP1 | 1.17914E-46 | 0.431281 | 0.961 | 0.875 | 2.29638E-42 |
| Contractile | PFKP | 3.10582E-25 | 0.431262 | 0.507 | 0.366 | 6.04858E-21 |
| IFN_Intermediate | ENSSSCG0( | 7.03838E-07 | 0.431079 | 0.252 | 0.105 | 0.013707239 |
| Contractile | ENSSSCG0( | 9.51282E-76 | 0.430889 | 0.991 | 0.991 | 1.85262E-71 |
| CD74_Intermediat | OLFM2 | 7.56368E-37 | 0.342121 | 0.23 | 0.107 | 1.47303E-32 |
| Fibroblasts | DDX5 | 1.59535E-09 | 0.430636 | 0.858 | 0.845 | 3.10694E-05 |
| Contractile | ENSSSCG0( | 1.52456E-23 | 0.430609 | 0.358 | 0.228 | 2.96908E-19 |
| Contractile | PITHD1 | 9.32589E-27 | 0.430551 | 0.353 | 0.213 | 1.81622E-22 |
| Terminal | CCDC80 | 1.24178E-45 | 0.430483 | 0.92 | 0.817 | 2.41836E-41 |
| VEGFA_Terminal | TNIP1 | 3.92533E-13 | 0.430287 | 0.375 | 0.222 | 7.64457E-09 |
| Intermediate | AGRN | 6.06144E-18 | 0.430132 | 0.284 | 0.158 | 1.18047E-13 |
| Fibroblasts | RGMA | 4.90067E-12 | 0.430011 | 0.147 | 0.046 | 9.54406E-08 |
| Intermediate | LPP | 3.56798E-22 | 0.429992 | 0.524 | 0.344 | 6.94863E-18 |
| CD74_Intermediat | RARRES2 | 1.55332E-71 | 0.429982 | 0.963 | 0.889 | 3.02509E-67 |
| Fibroblasts | PTGS1 | 2.88173E-06 | 0.429946 | 0.229 | 0.129 | 0.056121783 |
| VEGFA_Terminal | TSPO | 2.42464E-19 | 0.429929 | 0.981 | 0.923 | 4.72199E-15 |
| CD74_Intermediat | FGF2 | 1.44597E-32 | 0.429436 | 0.629 | 0.493 | 2.81603E-28 |
| Fibroblasts | RC3H1 | 0.001414864 | 0.42913 | 0.275 | 0.211 | 1 |

| | | | | | | |
|---|---|---|---|---|---|---|
| Fibroblasts | LPAR1 | 3.92375E-08 | 0.429129 | 0.211 | 0.103 | 0.000764151 |
| Fibroblasts | EZH1 | 0.000259904 | 0.429026 | 0.261 | 0.187 | 1 |
| CD74_Intermediat | HOMER1 | 8.87661E-44 | 0.350473 | 0.252 | 0.111 | 1.72872E-39 |
| Fibroblasts | SMOC2 | 0.000548666 | 0.428175 | 0.22 | 0.144 | 1 |
| Fibroblasts | TMOD2 | 0.000144023 | 0.428145 | 0.197 | 0.121 | 1 |
| CD74_Intermediat | FAM43A | 2.28194E-45 | 0.428232 | 0.276 | 0.128 | 4.44408E-41 |
| Fibroblasts | ST3GAL5 | 2.53897E-14 | 0.428071 | 0.119 | 0.029 | 4.94464E-10 |
| Pericytes | GPR20 | 0.000241927 | 0.428025 | 0.16 | 0.049 | 1 |
| IFN_Intermediate | ENSSSCG0( | 3.47741E-05 | 0.427975 | 0.234 | 0.111 | 0.677225801 |
| Fibroblasts | ENSSSCG0( | 9.15169E-06 | 0.427956 | 0.284 | 0.184 | 0.178229165 |
| VEGFA_Terminal | FGF7 | 3.13198E-21 | 0.427689 | 0.363 | 0.169 | 6.09953E-17 |
| Pericytes | EML2 | 0.001034941 | 0.427681 | 0.2 | 0.079 | 1 |
| Fibroblasts | ENSSSCG0( | 6.65499E-05 | 0.427491 | 0.321 | 0.231 | 1 |
| Intermediate | SERPINE1 | 1.25006E-22 | 0.427407 | 0.759 | 0.602 | 2.43449E-18 |
| VEGFA_Terminal | MAP2K1 | 8.64253E-24 | 0.427286 | 0.286 | 0.113 | 1.68313E-19 |
| VEGFA_Terminal | RPUSD3 | 2.42162E-26 | 0.427267 | 0.257 | 0.092 | 4.7161E-22 |
| Pericytes | HMGB1 | 2.67443E-05 | 0.427238 | 1 | 0.965 | 0.520845487 |
| VEGFA_Terminal | NEDD9 | 4.68293E-13 | 0.42712 | 0.387 | 0.229 | 9.12001E-09 |
| Contractile | LYRM1 | 3.78801E-30 | 0.427026 | 0.235 | 0.109 | 7.37716E-26 |
| Pericytes | GPM6B | 0.007173067 | 0.427011 | 0.22 | 0.108 | 1 |
| Pericytes | ENSSSCG0( | 0.001019529 | 0.426044 | 0.98 | 0.997 | 1 |
| CD74_Intermediat | CDH11 | 7.62608E-38 | 0.425999 | 0.527 | 0.362 | 1.48518E-33 |
| Fibroblasts | OLFM1 | 5.78174E-12 | 0.425811 | 0.138 | 0.042 | 1.12599E-07 |
| Fibroblasts | VGLL4 | 0.005697969 | 0.425772 | 0.211 | 0.155 | 1 |
| IFN_Intermediate | PSMA2 | 0.000129011 | 0.425469 | 0.664 | 0.53 | 1 |
| Contractile | HLF | 1.73305E-29 | 0.425233 | 0.318 | 0.173 | 3.37511E-25 |
| IFN_Intermediate | LTBP2 | 9.76993E-05 | 0.425008 | 0.85 | 0.728 | 1 |
| Contractile | PDK4 | 5.34312E-30 | 0.424896 | 0.22 | 0.094 | 1.04057E-25 |
| VEGFA_Terminal | DOCK7 | 7.80829E-16 | 0.424767 | 0.356 | 0.19 | 1.52066E-11 |
| Contractile | JUN | 2.38479E-43 | 0.424723 | 0.88 | 0.713 | 4.64437E-39 |
| Pericytes | H1-0 | 0.004882776 | 0.424622 | 0.18 | 0.078 | 1 |
| Intermediate | SAMD11 | 3.54179E-18 | 0.424585 | 0.246 | 0.126 | 6.89764E-14 |
| CD74_Intermediat | TPT1 | 1.799E-74 | 0.424576 | 0.99 | 0.978 | 3.50355E-70 |
| Contractile | DSC3 | 9.55191E-93 | 0.424462 | 0.139 | 0.013 | 1.86023E-88 |
| VEGFA_Terminal | FTL | 2.06725E-14 | 0.42443 | 0.99 | 0.994 | 4.02598E-10 |
| Terminal | C4ORF48 | 7.15589E-49 | 0.424374 | 0.443 | 0.259 | 1.39361E-44 |
| Pericytes | DUSP19 | 0.004391786 | 0.424308 | 0.18 | 0.078 | 1 |
| Terminal | RHOD | 9.63707E-43 | 0.424056 | 0.412 | 0.248 | 1.87682E-38 |
| IFN_Intermediate | JAK2 | 6.22878E-05 | 0.423792 | 0.383 | 0.231 | 1 |
| Terminal | CDH13 | 1.47942E-44 | 0.423001 | 0.755 | 0.597 | 2.88118E-40 |
| VEGFA_Terminal | LOC10062 | 1.11059E-20 | 0.422986 | 0.257 | 0.105 | 2.16287E-16 |
| Fibroblasts | MITF | 8.68239E-05 | 0.422548 | 0.266 | 0.18 | 1 |
| Pericytes | CLDN10 | 0.008576053 | 0.422498 | 0.12 | 0.045 | 1 |
| Pericytes | ENSSSCG0( | 0.001800618 | 0.422405 | 0.76 | 0.66 | 1 |
| Contractile | HMGN1 | 1.00622E-23 | 0.422166 | 0.638 | 0.52 | 1.95962E-19 |
| Fibroblasts | SHISA5 | 0.007833788 | 0.422136 | 0.376 | 0.351 | 1 |
| VEGFA_Terminal | CLIC1.1 | 2.87441E-14 | 0.422025 | 0.826 | 0.683 | 5.59792E-10 |
| Contractile | SMAD9 | 6.93107E-26 | 0.421999 | 0.529 | 0.382 | 1.34983E-21 |
| IFN_Intermediate | ERAP1 | 2.09231E-08 | 0.421812 | 0.196 | 0.063 | 0.000407477 |
| Fibroblasts | ACKR2 | 0.000100523 | 0.421647 | 0.193 | 0.116 | 1 |
| Contractile | PABPC1 | 2.34384E-38 | 0.421433 | 0.864 | 0.745 | 4.56463E-34 |
| Contractile | WBP11 | 7.39455E-27 | 0.420907 | 0.321 | 0.186 | 1.44009E-22 |
| Terminal | GABARAPL | 3.21201E-44 | 0.420885 | 0.751 | 0.613 | 6.25539E-40 |
| VEGFA_Terminal | RASSF1 | 7.95464E-19 | 0.420764 | 0.295 | 0.137 | 1.54917E-14 |
| Contractile | SYDE2 | 9.18107E-62 | 0.42066 | 0.161 | 0.033 | 1.78801E-57 |
| Fibroblasts | WSB1 | 8.30811E-08 | 0.420556 | 0.651 | 0.562 | 0.001618005 |
| Pericytes | ADAMTS9 | 0.002715387 | 0.420473 | 0.12 | 0.039 | 1 |
| Intermediate | TIMP1 | 1.30346E-33 | 0.420472 | 0.957 | 0.896 | 2.53848E-29 |
| IFN_Intermediate | SLA-7 | 1.72104E-07 | 0.420444 | 0.374 | 0.181 | 0.003351726 |

| | | | | | | |
|---|---|---|---|---|---|---|
| Intermediate | PMEPA1 | 2.89059E-17 | 0.420314 | 0.851 | 0.805 | 5.62943E-13 |
| Contractile | ENSSSCG0( | 2.4566E-51 | 0.420154 | 0.194 | 0.056 | 4.78423E-47 |
| Contractile | CDHR1 | 1.46866E-34 | 0.419976 | 0.222 | 0.091 | 2.86021E-30 |
| VEGFA_Terminal | MRPL17 | 1.65074E-16 | 0.419713 | 0.797 | 0.608 | 3.21482E-12 |
| VEGFA_Terminal | PVR | 9.80448E-34 | 0.419578 | 0.167 | 0.038 | 1.90942E-29 |
| VEGFA_Terminal | RHOA | 6.46242E-19 | 0.419433 | 0.86 | 0.704 | 1.25856E-14 |
| Contractile | SLC8A1 | 9.03832E-33 | 0.419425 | 0.27 | 0.128 | 1.76021E-28 |
| VEGFA_Terminal | CMC1 | 1.54344E-17 | 0.419405 | 0.53 | 0.324 | 3.00584E-13 |
| Contractile | TNS2 | 3.59148E-25 | 0.418895 | 0.443 | 0.297 | 6.9944E-21 |
| VEGFA_Terminal | CD9 | 1.60129E-24 | 0.41874 | 0.985 | 0.941 | 3.11851E-20 |
| Terminal | RHOA | 1.0951E-49 | 0.418696 | 0.8 | 0.673 | 2.13271E-45 |
| Fibroblasts | PCM1 | 0.001794772 | 0.418666 | 0.541 | 0.526 | 1 |
| Intermediate | SYNPO2 | 9.38261E-33 | 0.418511 | 0.707 | 0.472 | 1.82726E-28 |
| Contractile | ADAMTSL3 | 5.14218E-17 | 0.418207 | 0.389 | 0.278 | 1.00144E-12 |
| Fibroblasts | TIPARP | 1.71092E-06 | 0.418159 | 0.17 | 0.083 | 0.033320185 |
| VEGFA_Terminal | NT5E | 2.40261E-13 | 0.417791 | 0.358 | 0.206 | 4.67909E-09 |
| CD74_Intermediat | ENSSSCG0( | 1.00223E-34 | 0.334547 | 0.259 | 0.131 | 1.95183E-30 |
| Contractile | DYRK2 | 5.57757E-28 | 0.417293 | 0.289 | 0.156 | 1.08623E-23 |
| Contractile | BBX | 1.60212E-23 | 0.417237 | 0.519 | 0.386 | 3.12012E-19 |
| Fibroblasts | SERPINF1 | 1.319E-10 | 0.417232 | 0.959 | 0.756 | 2.56875E-06 |
| Contractile | RALY | 6.74707E-25 | 0.416923 | 0.536 | 0.399 | 1.31399E-20 |
| Fibroblasts | DBN1 | 0.00036001 | 0.416827 | 0.229 | 0.157 | 1 |
| IFN_Intermediate | SYNE2 | 0.005716737 | 0.416575 | 0.505 | 0.378 | 1 |
| VEGFA_Terminal | ENSSSCG0( | 6.66189E-06 | 0.416196 | 0.409 | 0.308 | 0.12974035 |
| Contractile | ZMYND11 | 4.46214E-19 | 0.415921 | 0.551 | 0.44 | 8.69002E-15 |
| IFN_Intermediate | CXCL12 | 0.000973558 | 0.415881 | 0.336 | 0.205 | 1 |
| Intermediate | DYRK3 | 3.28398E-16 | 0.41571 | 0.295 | 0.172 | 6.39556E-12 |
| Fibroblasts | CTTNBP2 | 6.5847E-07 | 0.41566 | 0.229 | 0.122 | 0.012823704 |
| Contractile | ALDOC | 2.22019E-21 | 0.415522 | 0.248 | 0.137 | 4.32382E-17 |
| Intermediate | RALGPS2 | 1.47179E-36 | 0.415459 | 0.167 | 0.046 | 2.86632E-32 |
| VEGFA_Terminal | MCUB | 3.92082E-19 | 0.415056 | 0.438 | 0.237 | 7.63579E-15 |
| VEGFA_Terminal | SYCP2 | 1.46021E-17 | 0.414923 | 0.373 | 0.194 | 2.84375E-13 |
| Contractile | ADGRE5 | 7.45939E-20 | 0.414663 | 0.403 | 0.281 | 1.45272E-15 |
| VEGFA_Terminal | ELL2 | 1.98754E-17 | 0.414647 | 0.356 | 0.184 | 3.87073E-13 |
| VEGFA_Terminal | H2AZ2 | 1.69616E-14 | 0.414537 | 0.816 | 0.656 | 3.30326E-10 |
| Contractile | GABPA | 2.23556E-21 | 0.414514 | 0.324 | 0.201 | 4.35376E-17 |
| Contractile | PTPRK | 4.10196E-22 | 0.41445 | 0.459 | 0.317 | 7.98858E-18 |
| VEGFA_Terminal | CMIP | 2.25306E-39 | 0.414431 | 0.228 | 0.058 | 4.38784E-35 |
| Contractile | ATP5MC2 | 9.47973E-29 | 0.414072 | 0.695 | 0.564 | 1.84618E-24 |
| Fibroblasts | RHOJ | 0.000105694 | 0.413758 | 0.193 | 0.117 | 1 |
| Fibroblasts | ARL4A | 0.001533105 | 0.413699 | 0.252 | 0.185 | 1 |
| Terminal | UGP2 | 1.02562E-24 | 0.413675 | 0.507 | 0.382 | 1.9974E-20 |
| Pericytes | ENSSSCG0( | 0.000496707 | 0.413662 | 0.98 | 0.977 | 1 |
| IFN_Intermediate | PLSCR1 | 9.15214E-06 | 0.41347 | 0.29 | 0.143 | 0.178237852 |
| Fibroblasts | UTRN | 0.000676301 | 0.41332 | 0.436 | 0.381 | 1 |
| Contractile | PSMG3 | 8.82E-31 | 0.413189 | 0.282 | 0.143 | 1.7177E-26 |
| Contractile | CS | 7.94775E-27 | 0.413044 | 0.365 | 0.22 | 1.54782E-22 |
| IFN_Intermediate | DDX24 | 2.75036E-06 | 0.412995 | 0.551 | 0.35 | 0.053563213 |
| CD74_Intermediat | PCOLCE | 1.69845E-57 | 0.41266 | 0.857 | 0.636 | 3.30774E-53 |
| Pericytes | ENSSSCG0( | 0.001334507 | 0.412658 | 0.78 | 0.624 | 1 |
| CD74_Intermediat | MDK | 1.3807E-32 | 0.358112 | 0.253 | 0.131 | 2.68891E-28 |
| Contractile | TUBB6 | 1.06455E-22 | 0.412465 | 0.429 | 0.294 | 2.07321E-18 |
| Fibroblasts | TRA2B | 0.000136085 | 0.412317 | 0.564 | 0.539 | 1 |
| Fibroblasts | TOP1 | 0.000353036 | 0.412287 | 0.555 | 0.526 | 1 |
| Fibroblasts | TLE1 | 0.000114166 | 0.412284 | 0.339 | 0.256 | 1 |
| Fibroblasts | COQ8A | 6.23636E-09 | 0.41215 | 0.165 | 0.067 | 0.000121453 |
| IFN_Intermediate | AKAP7 | 0.006426885 | 0.412104 | 0.178 | 0.102 | 1 |
| CD74_Intermediat | SPTBN1 | 2.31436E-56 | 0.412069 | 0.869 | 0.733 | 4.50722E-52 |
| Fibroblasts | PSPC1 | 0.000109404 | 0.411193 | 0.312 | 0.231 | 1 |

| | | | | | | |
|---|---|---|---|---|---|---|
| VEGFA_Terminal | CHCHD3 | 3.13096E-24 | 0.411068 | 0.387 | 0.178 | 6.09754E-20 |
| Terminal | CA4 | 1.18857E-66 | 0.410849 | 0.387 | 0.173 | 2.31475E-62 |
| VEGFA_Terminal | SLC38A2 | 6.41781E-11 | 0.410559 | 0.615 | 0.459 | 1.24987E-06 |
| Fibroblasts | MT1X | 2.35724E-08 | 0.410411 | 0.243 | 0.121 | 0.000459072 |
| Fibroblasts | BEX3 | 0.000133999 | 0.410304 | 0.294 | 0.206 | 1 |
| Contractile | ZC3H6 | 7.47895E-24 | 0.410296 | 0.279 | 0.156 | 1.45653E-19 |
| Contractile | CCDC68 | 2.78317E-78 | 0.410241 | 0.156 | 0.023 | 5.42023E-74 |
| Fibroblasts | IGFBP4 | 2.95656E-05 | 0.410172 | 0.165 | 0.089 | 0.575790221 |
| VEGFA_Terminal | PACSIN2 | 5.95957E-15 | 0.4101 | 0.4 | 0.229 | 1.16063E-10 |
| Contractile | BLOC1S1 | 8.07918E-25 | 0.409945 | 0.615 | 0.49 | 1.57342E-20 |
| CD74_Intermediat | LTBP1 | 1.67505E-51 | 0.409904 | 0.894 | 0.78 | 3.26217E-47 |
| Intermediate | GADD45B | 1.02433E-20 | 0.409896 | 0.512 | 0.342 | 1.99488E-16 |
| Contractile | PAIP2 | 4.06778E-27 | 0.409646 | 0.601 | 0.459 | 7.92201E-23 |
| Fibroblasts | CDC25B | 5.68641E-18 | 0.409615 | 0.138 | 0.031 | 1.10743E-13 |
| Fibroblasts | NUFIP2 | 0.000178221 | 0.40947 | 0.509 | 0.454 | 1 |
| VEGFA_Terminal | SPI1 | 1.95119E-38 | 0.409415 | 0.206 | 0.049 | 3.79994E-34 |
| Intermediate | SLPI | 3.22741E-12 | 0.409246 | 0.177 | 0.09 | 6.28539E-08 |
| Contractile | DCTN6 | 2.74635E-21 | 0.409235 | 0.451 | 0.324 | 5.34852E-17 |
| Fibroblasts | TMEM59 | 2.52485E-07 | 0.408944 | 0.798 | 0.772 | 0.004917154 |
| Pericytes | CD247 | 5.6437E-14 | 0.408942 | 0.12 | 0.01 | 1.09911E-09 |
| IFN_Intermediate | ENSSSCG0( | 0.008142116 | 0.408412 | 0.168 | 0.094 | 1 |
| Terminal | PRRX2 | 1.61152E-45 | 0.408287 | 0.847 | 0.732 | 3.13844E-41 |
| Fibroblasts | SRSF5 | 0.001164023 | 0.408276 | 0.307 | 0.242 | 1 |
| Contractile | ENSSSCG0( | 3.99143E-66 | 0.40827 | 0.186 | 0.041 | 7.77332E-62 |
| Fibroblasts | FOLH1 | 2.39686E-09 | 0.40815 | 0.225 | 0.103 | 4.66789E-05 |
| Contractile | NUMA1 | 1.0548E-22 | 0.408138 | 0.428 | 0.291 | 2.05423E-18 |
| Fibroblasts | THOC2 | 0.001844222 | 0.407885 | 0.381 | 0.329 | 1 |
| VEGFA_Terminal | ZNHIT1 | 8.20582E-15 | 0.407884 | 0.678 | 0.484 | 1.59808E-10 |
| Contractile | PUM2 | 4.09016E-20 | 0.407631 | 0.424 | 0.301 | 7.96559E-16 |
| Fibroblasts | DPYSL2 | 0.000103221 | 0.407522 | 0.353 | 0.275 | 1 |
| Pericytes | SOCS4 | 0.005858838 | 0.407379 | 0.3 | 0.166 | 1 |
| VEGFA_Terminal | RFNG | 1.56414E-19 | 0.407319 | 0.54 | 0.316 | 3.04617E-15 |
| Fibroblasts | RAB3GAP2 | 0.000509065 | 0.407292 | 0.326 | 0.251 | 1 |
| Contractile | TNRC18 | 2.70993E-36 | 0.407209 | 0.219 | 0.087 | 5.27758E-32 |
| Intermediate | ANTXR1 | 2.17049E-17 | 0.406983 | 0.76 | 0.658 | 4.22702E-13 |
| Pericytes | PPFIBP2 | 0.002540389 | 0.40692 | 0.18 | 0.073 | 1 |
| Intermediate | DAAM1 | 2.21429E-14 | 0.406862 | 0.661 | 0.542 | 4.31233E-10 |
| CD74_Intermediat | MFAP2 | 2.7539E-43 | 0.406452 | 0.566 | 0.375 | 5.36322E-39 |
| Fibroblasts | TNS2 | 0.002889966 | 0.406378 | 0.367 | 0.319 | 1 |
| Pericytes | EHD3 | 0.000129206 | 0.406289 | 0.16 | 0.047 | 1 |
| Fibroblasts | PALM | 1.21828E-07 | 0.406244 | 0.179 | 0.083 | 0.002372605 |
| VEGFA_Terminal | RCAN1 | 9.49542E-13 | 0.406122 | 0.281 | 0.151 | 1.84923E-08 |
| CD74_Intermediat | PDGFRL | 1.70002E-35 | 0.406077 | 0.615 | 0.449 | 3.31079E-31 |
| Contractile | LOC10015: | 6.04625E-81 | 0.405945 | 0.15 | 0.02 | 1.17751E-76 |
| VEGFA_Terminal | HSPE1 | 3.26062E-15 | 0.405904 | 0.809 | 0.644 | 6.35005E-11 |
| CD74_Intermediat | HSD11B1 | 1.46659E-42 | 0.405853 | 0.579 | 0.394 | 2.85618E-38 |
| IFN_Intermediate | ANKRD12 | 0.000591113 | 0.405772 | 0.832 | 0.716 | 1 |
| Fibroblasts | TPM3 | 8.24197E-05 | 0.405687 | 0.583 | 0.522 | 1 |
| Terminal | SELENOS | 1.47761E-37 | 0.405678 | 0.644 | 0.494 | 2.87765E-33 |
| Contractile | PPP2CB | 1.20707E-18 | 0.405632 | 0.436 | 0.323 | 2.35078E-14 |
| Contractile | ZNF185 | 3.67374E-77 | 0.405626 | 0.159 | 0.025 | 7.15462E-73 |
| Fibroblasts | CREG1 | 0.000599411 | 0.405577 | 0.472 | 0.424 | 1 |
| Contractile | RTF1 | 1.84744E-23 | 0.405557 | 0.668 | 0.565 | 3.59789E-19 |
| CD74_Intermediat | ALDH2 | 1.16179E-48 | 0.405453 | 0.688 | 0.497 | 2.26259E-44 |
| VEGFA_Terminal | EGLN1 | 3.98416E-14 | 0.405319 | 0.433 | 0.259 | 7.75916E-10 |
| CD74_Intermediat | MASP1 | 3.56917E-30 | 0.307458 | 0.253 | 0.134 | 6.95095E-26 |
| Contractile | ANKRD12 | 7.056E-28 | 0.404832 | 0.793 | 0.703 | 1.37416E-23 |
| Contractile | PHF19 | 4.33466E-43 | 0.404607 | 0.178 | 0.055 | 8.44175E-39 |
| Intermediate | CNN1 | 2.88508E-76 | 0.404406 | 0.676 | 0.271 | 5.61869E-72 |

| | | | | | | |
|---|---|---|---|---|---|---|
| Fibroblasts | GSN | 4.78659E-08 | 0.404387 | 1 | 0.991 | 0.000932188 |
| Contractile | ITGAV | 1.32467E-18 | 0.404337 | 0.697 | 0.594 | 2.57979E-14 |
| Contractile | UBALD1 | 3.11891E-30 | 0.404306 | 0.249 | 0.119 | 6.07408E-26 |
| Contractile | CAV3 | 2.3284E-108 | 0.404245 | 0.137 | 0.008 | 4.5346E-104 |
| Contractile | HNRNPDL | 7.56326E-29 | 0.404044 | 0.73 | 0.609 | 1.47294E-24 |
| IFN_Intermediate | PTPN9 | 4.44367E-06 | 0.403816 | 0.308 | 0.151 | 0.086540515 |
| IFN_Intermediate | GDF10 | 3.28173E-06 | 0.403681 | 0.187 | 0.071 | 0.063911763 |
| Contractile | PDZRN3 | 5.55049E-25 | 0.403408 | 0.25 | 0.13 | 1.08096E-20 |
| Terminal | TSPAN6 | 1.85032E-37 | 0.403264 | 0.518 | 0.363 | 3.60351E-33 |
| Fibroblasts | ENSSSCG0( | 3.82181E-17 | 0.403245 | 1 | 0.997 | 7.44298E-13 |
| Fibroblasts | SON | 2.78073E-11 | 0.403177 | 0.894 | 0.87 | 5.41548E-07 |
| Pericytes | RAB12 | 0.006392126 | 0.403021 | 0.34 | 0.203 | 1 |
| Contractile | USP14 | 2.15364E-22 | 0.402942 | 0.422 | 0.29 | 4.19422E-18 |
| Contractile | AKAP11 | 1.41187E-22 | 0.402759 | 0.307 | 0.183 | 2.74962E-18 |
| CD74_Intermediat | CCDC80 | 4.64799E-54 | 0.402716 | 0.927 | 0.82 | 9.05197E-50 |
| Intermediate | FMOD | 1.88571E-31 | 0.402623 | 0.905 | 0.773 | 3.67243E-27 |
| IFN_Intermediate | IRF9 | 6.49237E-08 | 0.40262 | 0.449 | 0.223 | 0.001264389 |
| Terminal | PRDX4 | 3.80046E-45 | 0.402463 | 0.746 | 0.607 | 7.40139E-41 |
| Contractile | PPFIBP1 | 4.59403E-20 | 0.402418 | 0.622 | 0.511 | 8.94688E-16 |
| Contractile | PPP1R12C | 3.41748E-60 | 0.402181 | 0.201 | 0.052 | 6.65554E-56 |
| Contractile | GJA1 | 1.68712E-20 | 0.402136 | 0.692 | 0.551 | 3.28566E-16 |
| Contractile | ATRX | 2.01641E-23 | 0.402109 | 0.667 | 0.558 | 3.92696E-19 |
| Intermediate | TPM2 | 1.42054E-39 | 0.401844 | 0.949 | 0.836 | 2.7665E-35 |
| IFN_Intermediate | SLA-6 | 0.007224833 | 0.401573 | 0.514 | 0.387 | 1 |
| Fibroblasts | PIP5K1A | 0.000162019 | 0.401531 | 0.239 | 0.157 | 1 |
| Contractile | H1-10 | 1.92351E-25 | 0.401521 | 0.745 | 0.655 | 3.74604E-21 |
| Pericytes | PAG1 | 5.62539E-08 | 0.401285 | 0.14 | 0.023 | 0.001095544 |
| VEGFA_Terminal | ENSSSCG0( | 4.87157E-09 | 0.401136 | 0.102 | 0.04 | 9.48739E-05 |
| VEGFA_Terminal | COL1A1 | 2.75034E-07 | 0.401043 | 0.942 | 0.939 | 0.005356287 |
| IFN_Intermediate | PARP9 | 1.84806E-16 | 0.400967 | 0.206 | 0.041 | 3.59911E-12 |
| Pericytes | PAK2 | 0.007026967 | 0.400801 | 0.62 | 0.5 | 1 |
| Contractile | PARP1 | 2.72936E-20 | 0.40075 | 0.294 | 0.18 | 5.31542E-16 |
| Intermediate | ITGA1 | 5.92329E-15 | 0.400683 | 0.705 | 0.605 | 1.15356E-10 |
| VEGFA_Terminal | RAC1 | 1.40426E-17 | 0.400481 | 0.84 | 0.712 | 2.7348E-13 |
| CD74_Intermediat | PDGFRB | 1.9898E-40 | 0.399932 | 0.572 | 0.404 | 3.87514E-36 |
| VEGFA_Terminal | GLS | 5.81394E-10 | 0.399893 | 0.554 | 0.398 | 1.13226E-05 |
| Contractile | SNU13 | 1.60747E-18 | 0.399786 | 0.533 | 0.436 | 3.13055E-14 |
| Contractile | HNRNPD | 1.19026E-22 | 0.399257 | 0.574 | 0.451 | 2.31804E-18 |
| Contractile | LAMB2 | 5.73083E-24 | 0.399231 | 0.632 | 0.503 | 1.11608E-19 |
| Intermediate | AKAP12 | 7.01023E-16 | 0.399169 | 0.429 | 0.286 | 1.36524E-11 |
| Fibroblasts | ANTXR2 | 0.001647885 | 0.399165 | 0.353 | 0.3 | 1 |
| Contractile | CTNNA1 | 9.68983E-18 | 0.399055 | 0.59 | 0.503 | 1.88709E-13 |
| Contractile | FADS3 | 1.36929E-21 | 0.399028 | 0.41 | 0.282 | 2.6667E-17 |
| Pericytes | JARID2 | 0.002790967 | 0.398924 | 0.14 | 0.05 | 1 |
| VEGFA_Terminal | ATP5MK | 1.58637E-16 | 0.398898 | 0.843 | 0.724 | 3.08946E-12 |
| Contractile | PYGM | 2.4741E-107 | 0.398823 | 0.126 | 0.006 | 4.8182E-103 |
| CD74_Intermediat | EFEMP1 | 2.20177E-65 | 0.398812 | 0.828 | 0.603 | 4.28794E-61 |
| VEGFA_Terminal | CTNNBIP1 | 1.32188E-21 | 0.3988 | 0.349 | 0.16 | 2.57437E-17 |
| Fibroblasts | MRC2 | 0.003201471 | 0.398769 | 0.482 | 0.437 | 1 |
| Pericytes | NFKBID | 2.90561E-06 | 0.398706 | 0.12 | 0.022 | 0.056586823 |
| Contractile | FHL3 | 2.03788E-33 | 0.398527 | 0.212 | 0.088 | 3.96877E-29 |
| Intermediate | NIBAN1 | 7.06951E-24 | 0.398494 | 0.859 | 0.76 | 1.37679E-19 |
| Terminal | AQP3 | 1.42661E-27 | 0.398453 | 0.527 | 0.38 | 2.77832E-23 |
| Fibroblasts | EIF4A1 | 0.000328895 | 0.398349 | 0.528 | 0.497 | 1 |
| Contractile | NPTN | 7.77205E-21 | 0.398074 | 0.603 | 0.499 | 1.51361E-16 |
| Pericytes | CCDC86 | 1.21088E-07 | 0.398026 | 0.14 | 0.024 | 0.00235818 |
| Pericytes | ENSSSCG0( | 6.64143E-13 | 0.397926 | 0.14 | 0.014 | 1.29342E-08 |
| Contractile | HNRNPA1 | 3.96458E-28 | 0.397848 | 0.704 | 0.585 | 7.72102E-24 |
| Contractile | ATP5PF | 3.13454E-37 | 0.397816 | 0.888 | 0.833 | 6.10452E-33 |

| | | | | | | |
|---|---|---|---|---|---|---|
| Intermediate | PKDCC | 7.66789E-13 | 0.396736 | 0.544 | 0.432 | 1.49332E-08 |
| Contractile | RSU1 | 2.00736E-25 | 0.396678 | 0.636 | 0.513 | 3.90933E-21 |
| Contractile | C4B | 2.28192E-38 | 0.39659 | 0.428 | 0.226 | 4.44404E-34 |
| Fibroblasts | IL6R | 1.75074E-11 | 0.396561 | 0.165 | 0.057 | 3.40957E-07 |
| IFN_Intermediate | DNASE1L1 | 0.001953393 | 0.396439 | 0.505 | 0.366 | 1 |
| Terminal | ECM1 | 4.95752E-82 | 0.396434 | 0.803 | 0.596 | 9.65476E-78 |
| VEGFA_Terminal | FMNL3 | 2.62565E-34 | 0.396428 | 0.24 | 0.07 | 5.11346E-30 |
| Contractile | ABLIM1 | 5.36585E-23 | 0.396372 | 0.492 | 0.344 | 1.045E-18 |
| VEGFA_Terminal | MOXD1 | 5.89868E-32 | 0.396299 | 0.223 | 0.064 | 1.14877E-27 |
| Fibroblasts | RPIA | 0.000370491 | 0.396249 | 0.257 | 0.181 | 1 |
| Fibroblasts | KLHL15 | 0.000878815 | 0.396249 | 0.179 | 0.116 | 1 |
| Pericytes | STAC | 2.93342E-05 | 0.396193 | 0.18 | 0.051 | 0.57128339 |
| Fibroblasts | TMED4 | 0.000229943 | 0.396104 | 0.546 | 0.504 | 1 |
| Terminal | SLC7A2 | 4.44391E-24 | 0.39607 | 0.308 | 0.197 | 8.65451E-20 |
| VEGFA_Terminal | EPAS1 | 1.4251E-07 | 0.395954 | 0.7 | 0.565 | 0.002775385 |
| VEGFA_Terminal | CTSK | 2.66854E-10 | 0.395735 | 0.93 | 0.877 | 5.19697E-06 |
| Pericytes | ZNF697 | 0.008319161 | 0.395614 | 0.16 | 0.07 | 1 |
| Pericytes | KSR1 | 0.00638107 | 0.395392 | 0.22 | 0.106 | 1 |
| Contractile | ARID4A | 5.56838E-21 | 0.395301 | 0.416 | 0.285 | 1.08444E-16 |
| Fibroblasts | SASH1 | 0.001914633 | 0.39514 | 0.532 | 0.503 | 1 |
| Fibroblasts | ENSSSCG0( | 0.007220895 | 0.39512 | 0.248 | 0.197 | 1 |
| IFN_Intermediate | ENSSSCG0( | 0.00037177 | 0.394787 | 0.131 | 0.052 | 1 |
| Intermediate | ADAMTS1 | 0.000612363 | 0.394787 | 0.25 | 0.204 | 1 |
| VEGFA_Terminal | RCN1 | 2.70642E-11 | 0.394631 | 0.608 | 0.443 | 5.27076E-07 |
| IFN_Intermediate | PIK3C2A | 0.00118567 | 0.394071 | 0.346 | 0.223 | 1 |
| Intermediate | PRKAR2B | 1.24251E-15 | 0.393727 | 0.332 | 0.205 | 2.41978E-11 |
| VEGFA_Terminal | SLC3A2 | 3.6617E-17 | 0.393617 | 0.329 | 0.162 | 7.13115E-13 |
| IFN_Intermediate | MORC3 | 3.6862E-06 | 0.393567 | 0.29 | 0.136 | 0.071788772 |
| Terminal | COPZ2 | 1.12887E-31 | 0.393499 | 0.621 | 0.49 | 2.19847E-27 |
| Contractile | NR1H2 | 4.44004E-19 | 0.393451 | 0.35 | 0.235 | 8.64698E-15 |
| VEGFA_Terminal | BRI3 | 1.29537E-13 | 0.393356 | 0.772 | 0.628 | 2.52274E-09 |
| VEGFA_Terminal | SQSTM1 | 2.52538E-11 | 0.393274 | 0.814 | 0.67 | 4.91818E-07 |
| Contractile | YWHAH | 1.55489E-20 | 0.3932 | 0.488 | 0.363 | 3.02815E-16 |
| Intermediate | TENT5A | 2.48721E-15 | 0.393161 | 0.619 | 0.482 | 4.84384E-11 |
| Contractile | RNF38 | 7.83119E-23 | 0.392627 | 0.315 | 0.188 | 1.52512E-18 |
| Fibroblasts | JMJD1C | 0.002397802 | 0.392543 | 0.528 | 0.488 | 1 |
| Terminal | TNFAIP6 | 2.37E-33 | 0.392541 | 0.208 | 0.097 | 4.61558E-29 |
| Pericytes | COBLL1 | 0.006661484 | 0.392468 | 0.2 | 0.093 | 1 |
| Contractile | GYG2 | 2.11282E-28 | 0.392441 | 0.251 | 0.124 | 4.11472E-24 |
| Intermediate | HMCN1 | 3.29207E-07 | 0.392248 | 0.425 | 0.355 | 0.006411308 |
| Intermediate | ELN | 3.44677E-15 | 0.391917 | 0.161 | 0.074 | 6.71258E-11 |
| Fibroblasts | PRNP | 0.000447119 | 0.391345 | 0.44 | 0.389 | 1 |
| Contractile | BOK | 1.53771E-20 | 0.391156 | 0.374 | 0.251 | 2.9947E-16 |
| Contractile | LRCH1 | 9.70543E-34 | 0.391119 | 0.243 | 0.107 | 1.89013E-29 |
| Pericytes | ENTPD1 | 0.003201726 | 0.391086 | 0.22 | 0.097 | 1 |
| Fibroblasts | AK3 | 3.25567E-05 | 0.390966 | 0.312 | 0.218 | 0.634041776 |
| Contractile | COL4A5 | 6.32331E-25 | 0.390903 | 0.298 | 0.168 | 1.23146E-20 |
| Fibroblasts | SERINC1 | 1.44204E-05 | 0.390638 | 0.619 | 0.565 | 0.280836912 |
| Pericytes | ENSSSCG0( | 0.004627923 | 0.39059 | 0.12 | 0.041 | 1 |
| Intermediate | PDLIM1 | 1.09088E-12 | 0.390552 | 0.465 | 0.347 | 2.12448E-08 |
| Terminal | ABI3BP | 7.88037E-22 | 0.390154 | 0.595 | 0.467 | 1.5347E-17 |
| Fibroblasts | PTPRS | 2.36531E-06 | 0.390151 | 0.188 | 0.098 | 0.046064422 |
| IFN_Intermediate | GSN | 8.25478E-05 | 0.390126 | 0.991 | 0.992 | 1 |
| Fibroblasts | EMILIN2 | 1.17785E-07 | 0.390125 | 0.174 | 0.08 | 0.002293855 |
| VEGFA_Terminal | MFF | 1.24056E-11 | 0.390069 | 0.545 | 0.378 | 2.41599E-07 |
| Contractile | PFDN1 | 3.21425E-19 | 0.389901 | 0.533 | 0.423 | 6.25974E-15 |
| VEGFA_Terminal | UNC5B | 7.15829E-38 | 0.389771 | 0.206 | 0.049 | 1.39408E-33 |
| Contractile | HSPB8 | 6.21936E-24 | 0.389657 | 0.36 | 0.22 | 1.21122E-19 |
| IFN_Intermediate | CD274 | 5.22435E-20 | 0.389326 | 0.131 | 0.015 | 1.01744E-15 |

| | | | | | | |
|---|---|---|---|---|---|---|
| Contractile | C7ORF50 | 1.54566E-22 | 0.389278 | 0.449 | 0.312 | 3.01017E-18 |
| Pericytes | FUT2A | 3.92014E-07 | 0.389254 | 0.1 | 0.014 | 0.007634473 |
| Intermediate | CAVIN3 | 3.70638E-19 | 0.389212 | 0.972 | 0.957 | 7.21817E-15 |
| Fibroblasts | SGCE | 0.003230202 | 0.389191 | 0.381 | 0.338 | 1 |
| VEGFA_Terminal | NANS | 6.14912E-16 | 0.389154 | 0.339 | 0.177 | 1.19754E-11 |
| Terminal | NPC2 | 7.06313E-36 | 0.389131 | 0.762 | 0.633 | 1.37555E-31 |
| Contractile | ITGB1BP2 | 2.305E-118 | 0.388796 | 0.137 | 0.006 | 4.489E-114 |
| VEGFA_Terminal | LXN | 8.58351E-12 | 0.388741 | 0.441 | 0.28 | 1.67164E-07 |
| Contractile | NBR1 | 2.60908E-18 | 0.388669 | 0.378 | 0.261 | 5.08119E-14 |
| Contractile | SBDS | 1.09231E-23 | 0.388649 | 0.65 | 0.535 | 2.12727E-19 |
| Contractile | NFIC | 1.33952E-24 | 0.388471 | 0.724 | 0.626 | 2.60872E-20 |
| Contractile | DUSP3 | 1.25808E-22 | 0.388374 | 0.375 | 0.245 | 2.4501E-18 |
| Fibroblasts | TIMP3 | 3.17521E-06 | 0.388264 | 0.95 | 0.94 | 0.061837273 |
| Fibroblasts | RBM26 | 0.002629329 | 0.388223 | 0.335 | 0.278 | 1 |
| Terminal | GPX1 | 2.13768E-34 | 0.387591 | 0.952 | 0.925 | 4.16313E-30 |
| Fibroblasts | NFIX | 9.28129E-05 | 0.387468 | 0.633 | 0.574 | 1 |
| IFN_Intermediate | PTHLH | 0.000141405 | 0.387349 | 0.178 | 0.079 | 1 |
| Contractile | UQCR10 | 5.66531E-27 | 0.387347 | 0.773 | 0.666 | 1.10332E-22 |
| Contractile | ZBTB47 | 2.85044E-28 | 0.387336 | 0.28 | 0.145 | 5.55124E-24 |
| Intermediate | CLEC14A | 2.15634E-08 | 0.387191 | 0.547 | 0.461 | 0.000419947 |
| Pericytes | ADAMTS1! | 1.13375E-05 | 0.387097 | 0.14 | 0.032 | 0.220797294 |
| Contractile | ABCC9 | 5.39005E-56 | 0.387016 | 0.156 | 0.034 | 1.04971E-51 |
| Fibroblasts | ACKR3 | 2.55637E-06 | 0.386766 | 0.197 | 0.103 | 0.049785222 |
| Fibroblasts | RAB8B | 0.006819127 | 0.386654 | 0.28 | 0.232 | 1 |
| Fibroblasts | DENND2A | 7.92828E-07 | 0.386175 | 0.156 | 0.07 | 0.015440326 |
| Contractile | PPM1L | 2.95411E-54 | 0.385928 | 0.162 | 0.038 | 5.75313E-50 |
| Contractile | TAF7 | 8.10508E-22 | 0.385857 | 0.561 | 0.428 | 1.57846E-17 |
| Fibroblasts | NCOR2 | 0.001206373 | 0.385811 | 0.312 | 0.248 | 1 |
| IFN_Intermediate | ITPR2 | 0.000631824 | 0.385677 | 0.271 | 0.154 | 1 |
| Contractile | SSBP3 | 3.19866E-29 | 0.385668 | 0.219 | 0.098 | 6.22939E-25 |
| Contractile | MEF2D | 9.28698E-19 | 0.385118 | 0.324 | 0.211 | 1.80864E-14 |
| Fibroblasts | EPHA7 | 7.69359E-07 | 0.384911 | 0.174 | 0.083 | 0.014983272 |
| Fibroblasts | IFNAR1 | 0.003150643 | 0.384679 | 0.28 | 0.224 | 1 |
| Contractile | RUSC2 | 2.16255E-31 | 0.384522 | 0.264 | 0.126 | 4.21156E-27 |
| Contractile | GATA4 | 2.83119E-20 | 0.384394 | 0.284 | 0.172 | 5.51375E-16 |
| Contractile | INPP4A | 5.54235E-36 | 0.383804 | 0.182 | 0.064 | 1.07937E-31 |
| Contractile | ENSSSCG0( | 9.33344E-55 | 0.383763 | 0.148 | 0.031 | 1.81769E-50 |
| Fibroblasts | FRMD3 | 1.3429E-43 | 0.383528 | 0.119 | 0.01 | 2.6153E-39 |
| CD74_Intermediat | ALDH1A3 | 1.15614E-29 | 0.383524 | 0.252 | 0.134 | 2.25159E-25 |
| Fibroblasts | TST | 0.004126179 | 0.38346 | 0.335 | 0.285 | 1 |
| Contractile | NPTX1 | 3.52382E-97 | 0.383332 | 0.106 | 0.004 | 6.86264E-93 |
| Contractile | IPO8 | 5.70925E-27 | 0.383187 | 0.225 | 0.108 | 1.11188E-22 |
| Intermediate | FGF7 | 1.51555E-06 | 0.383101 | 0.246 | 0.175 | 0.029515301 |
| Intermediate | FSTL3 | 1.35549E-13 | 0.383006 | 0.488 | 0.367 | 2.63982E-09 |
| Pericytes | ENSSSCG0( | 5.82877E-05 | 0.382797 | 0.16 | 0.044 | 1 |
| Contractile | RNF19A | 2.27019E-26 | 0.382793 | 0.752 | 0.63 | 4.42119E-22 |
| Contractile | PNPLA2 | 5.68497E-18 | 0.382609 | 0.339 | 0.229 | 1.10715E-13 |
| Terminal | SEC62 | 8.83107E-57 | 0.382466 | 0.944 | 0.9 | 1.71985E-52 |
| CD74_Intermediat | ALDH1A1 | 3.42857E-30 | 0.349377 | 0.255 | 0.136 | 6.67714E-26 |
| VEGFA_Terminal | EEF1G | 1.74292E-15 | 0.382124 | 0.896 | 0.835 | 3.39434E-11 |
| Fibroblasts | VEZF1 | 0.000661665 | 0.382094 | 0.344 | 0.275 | 1 |
| Contractile | KCTD15 | 3.31318E-47 | 0.382072 | 0.187 | 0.056 | 6.45242E-43 |
| Contractile | MAP2K7 | 2.25623E-24 | 0.382026 | 0.245 | 0.127 | 4.39401E-20 |
| Contractile | CYC1 | 6.87149E-22 | 0.381827 | 0.447 | 0.315 | 1.33822E-17 |
| Terminal | ID2 | 3.50288E-37 | 0.381779 | 0.831 | 0.731 | 6.82185E-33 |
| Contractile | HIPK3 | 2.18843E-17 | 0.381738 | 0.348 | 0.235 | 4.26198E-13 |
| CD74_Intermediat | C7 | 5.34011E-76 | 0.608319 | 0.352 | 0.138 | 1.03999E-71 |
| Fibroblasts | NOTCH2 | 0.000111513 | 0.381662 | 0.486 | 0.427 | 1 |
| Contractile | ITPRID2 | 3.15993E-20 | 0.381656 | 0.52 | 0.389 | 6.15396E-16 |

| | | | | | | |
|---|---|---|---|---|---|---|
| VEGFA_Terminal | EFCAB7 | 1.74788E-16 | 0.381591 | 0.484 | 0.285 | 3.404E-12 |
| CD74_Intermediat | CELF2 | 5.8531E-20 | 0.27106 | 0.235 | 0.141 | 1.13989E-15 |
| Contractile | ATP5MK | 1.45493E-25 | 0.381427 | 0.81 | 0.717 | 2.83348E-21 |
| Fibroblasts | ITGB3 | 0.000577711 | 0.381419 | 0.179 | 0.112 | 1 |
| Intermediate | CFL2 | 9.62025E-13 | 0.381403 | 0.505 | 0.383 | 1.87354E-08 |
| Fibroblasts | QSER1 | 0.004269094 | 0.381337 | 0.289 | 0.236 | 1 |
| Pericytes | NOCT | 2.48593E-05 | 0.381306 | 0.16 | 0.041 | 0.484135045 |
| VEGFA_Terminal | WARS1 | 2.63486E-18 | 0.381277 | 0.242 | 0.103 | 5.1314E-14 |
| Contractile | FKBP4 | 5.96765E-20 | 0.380982 | 0.421 | 0.295 | 1.1622E-15 |
| Intermediate | SYCP2 | 9.36854E-09 | 0.380849 | 0.281 | 0.197 | 0.000182452 |
| CD74_Intermediat | TMEM150 | 2.38035E-74 | 0.571886 | 0.346 | 0.144 | 4.63574E-70 |
| Fibroblasts | TRPS1 | 3.49469E-06 | 0.380449 | 0.569 | 0.472 | 0.068059139 |
| Pericytes | COX4I2 | 0.006948784 | 0.380359 | 0.86 | 0.817 | 1 |
| Pericytes | AKAP6 | 0.001226218 | 0.380176 | 0.2 | 0.079 | 1 |
| IFN_Intermediate | FMNL2 | 0.002254533 | 0.380172 | 0.243 | 0.138 | 1 |
| Fibroblasts | DDX46 | 0.002202211 | 0.380153 | 0.491 | 0.451 | 1 |
| Terminal | PHLDA2 | 2.39309E-21 | 0.379789 | 0.593 | 0.476 | 4.66055E-17 |
| CD74_Intermediat | ICAM1 | 8.16911E-34 | 0.369545 | 0.272 | 0.144 | 1.59093E-29 |
| Contractile | DIP2C | 4.41349E-23 | 0.379686 | 0.265 | 0.146 | 8.59527E-19 |
| CD74_Intermediat | RSPO3 | 2.22729E-16 | 0.469216 | 0.23 | 0.145 | 4.33764E-12 |
| Fibroblasts | PCDH18 | 0.000243 | 0.379236 | 0.179 | 0.107 | 1 |
| Contractile | GNAI2 | 3.89874E-27 | 0.378742 | 0.813 | 0.738 | 7.5928E-23 |
| Intermediate | SH3BGR | 7.33642E-15 | 0.378686 | 0.425 | 0.289 | 1.42877E-10 |
| Fibroblasts | GSTK1 | 0.007669402 | 0.378558 | 0.161 | 0.111 | 1 |
| VEGFA_Terminal | TBCA | 1.88144E-13 | 0.378375 | 0.775 | 0.615 | 3.6641E-09 |
| Contractile | ENSSSCG0 | 1.83142E-22 | 0.378372 | 0.359 | 0.229 | 3.5667E-18 |
| Contractile | SRFBP1 | 3.7251E-20 | 0.378042 | 0.32 | 0.203 | 7.25464E-16 |
| Contractile | PGLYRP1 | 5.11338E-28 | 0.378023 | 0.25 | 0.121 | 9.9583E-24 |
| Fibroblasts | SAMHD1 | 8.39563E-05 | 0.377936 | 0.234 | 0.151 | 1 |
| IFN_Intermediate | SLC25A28 | 5.65249E-05 | 0.377556 | 0.28 | 0.146 | 1 |
| VEGFA_Terminal | RIOK3 | 5.37859E-12 | 0.377484 | 0.695 | 0.514 | 1.04748E-07 |
| Fibroblasts | ANKFY1 | 2.20772E-07 | 0.377188 | 0.188 | 0.092 | 0.004299544 |
| Contractile | BPTF | 1.62216E-17 | 0.37707 | 0.532 | 0.428 | 3.15916E-13 |
| Contractile | FAM210B | 4.43905E-20 | 0.377015 | 0.396 | 0.272 | 8.64506E-16 |
| VEGFA_Terminal | ENSSSCG0 | 1.2542E-21 | 0.376432 | 0.983 | 0.975 | 2.44255E-17 |
| Contractile | XPR1 | 1.15794E-52 | 0.376323 | 0.164 | 0.04 | 2.25509E-48 |
| VEGFA_Terminal | UAP1 | 9.4212E-12 | 0.376315 | 0.479 | 0.308 | 1.83478E-07 |
| VEGFA_Terminal | ARFGAP1 | 1.39451E-14 | 0.375997 | 0.496 | 0.3 | 2.71582E-10 |
| CD74_Intermediat | GPX3 | 1.0275E-151 | 0.87315 | 0.46 | 0.146 | 2.0011E-147 |
| VEGFA_Terminal | ARHGDIA | 2.80675E-15 | 0.375907 | 0.755 | 0.573 | 5.46614E-11 |
| VEGFA_Terminal | ARHGDIB | 2.49561E-11 | 0.375778 | 0.533 | 0.362 | 4.86021E-07 |
| IFN_Intermediate | H2AC4 | 0.000779744 | 0.375646 | 0.402 | 0.263 | 1 |
| Contractile | SNRNP25 | 2.33026E-19 | 0.375548 | 0.407 | 0.285 | 4.53819E-15 |
| IFN_Intermediate | ENSSSCG0 | 5.84995E-06 | 0.375442 | 0.206 | 0.082 | 0.11392773 |
| Fibroblasts | CD81 | 6.91122E-13 | 0.375352 | 0.995 | 0.979 | 1.34596E-08 |
| IFN_Intermediate | LPAR1 | 0.000377819 | 0.375286 | 0.206 | 0.105 | 1 |
| Contractile | BCL2 | 1.385E-20 | 0.375125 | 0.375 | 0.247 | 2.69728E-16 |
| VEGFA_Terminal | CRYAB | 1.02579E-24 | 0.375067 | 0.978 | 0.848 | 1.99773E-20 |
| Contractile | CNTFR | 1.80744E-98 | 0.374976 | 0.109 | 0.004 | 3.51999E-94 |
| Contractile | ECHS1 | 6.63549E-29 | 0.374623 | 0.8 | 0.707 | 1.29226E-24 |
| Contractile | NSMCE1 | 2.18782E-19 | 0.374459 | 0.474 | 0.357 | 4.26077E-15 |
| Fibroblasts | COL6A6 | 5.43461E-05 | 0.374299 | 0.11 | 0.05 | 1 |
| Fibroblasts | ANGPT4 | 6.31025E-07 | 0.374218 | 0.101 | 0.036 | 0.012289212 |
| IFN_Intermediate | PLA2G4A | 0.000915817 | 0.374216 | 0.308 | 0.184 | 1 |
| Fibroblasts | TNKS | 0.000200741 | 0.374211 | 0.252 | 0.173 | 1 |
| IFN_Intermediate | BACH1 | 9.44469E-05 | 0.374201 | 0.449 | 0.286 | 1 |
| Fibroblasts | DDX39B | 0.002451133 | 0.374184 | 0.326 | 0.262 | 1 |
| Terminal | PGAM1 | 2.16603E-33 | 0.37417 | 0.647 | 0.496 | 4.21834E-29 |
| Contractile | SYNC | 2.05339E-34 | 0.373675 | 0.208 | 0.083 | 3.99898E-30 |

| | | | | | | |
|---|---|---|---|---|---|---|
| Pericytes | ENSSSCG0( | 0.001262963 | 0.373577 | 0.16 | 0.057 | 1 |
| Pericytes | ENSSSCG0( | 0.009455566 | 0.373563 | 0.12 | 0.046 | 1 |
| Pericytes | IFT172 | 0.004510626 | 0.373496 | 0.16 | 0.064 | 1 |
| Pericytes | ALG14 | 0.000987907 | 0.373438 | 0.12 | 0.035 | 1 |
| Contractile | COL4A1 | 5.44844E-44 | 0.373333 | 0.837 | 0.665 | 1.06108E-39 |
| IFN_Intermediate | PCYOX1 | 0.008290649 | 0.373284 | 0.505 | 0.381 | 1 |
| VEGFA_Terminal | ENSSSCG0( | 2.05022E-28 | 0.373167 | 0.998 | 0.995 | 3.9928E-24 |
| VEGFA_Terminal | SHISA4 | 1.40569E-13 | 0.373072 | 0.659 | 0.475 | 2.73759E-09 |
| Fibroblasts | ANKRD46 | 0.000393337 | 0.373043 | 0.271 | 0.195 | 1 |
| Contractile | CCT7 | 4.95564E-21 | 0.372929 | 0.557 | 0.434 | 9.65112E-17 |
| Contractile | PWWP3A | 1.27454E-19 | 0.37271 | 0.328 | 0.209 | 2.48216E-15 |
| Intermediate | HSPB7 | 2.19327E-12 | 0.372674 | 0.287 | 0.176 | 4.2714E-08 |
| Fibroblasts | HSPA12B | 4.16682E-65 | 0.372629 | 0.101 | 0.004 | 8.11488E-61 |
| Contractile | JCAD | 7.42509E-20 | 0.372585 | 0.278 | 0.167 | 1.44604E-15 |
| IFN_Intermediate | LUM | 2.13316E-06 | 0.372582 | 0.991 | 0.931 | 0.041543299 |
| IFN_Intermediate | TRIM14 | 4.004E-16 | 0.372545 | 0.15 | 0.024 | 7.7978E-12 |
| Pericytes | ENSSSCG0( | 0.00940219 | 0.372473 | 0.1 | 0.034 | 1 |
| Contractile | SUCLG2 | 2.99293E-16 | 0.37239 | 0.394 | 0.288 | 5.82873E-12 |
| VEGFA_Terminal | ASAP2 | 2.55244E-15 | 0.372233 | 0.424 | 0.246 | 4.97087E-11 |
| IFN_Intermediate | TAPBP | 0.00017775 | 0.372081 | 0.308 | 0.171 | 1 |
| Intermediate | PCSK5 | 8.22993E-14 | 0.372006 | 0.339 | 0.218 | 1.60278E-09 |
| Intermediate | AMOTL2 | 3.99147E-15 | 0.371949 | 0.241 | 0.133 | 7.7734E-11 |
| Pericytes | FAM110B | 0.000622173 | 0.371885 | 0.16 | 0.054 | 1 |
| Fibroblasts | ENSSSCG0( | 0.007123069 | 0.371775 | 0.261 | 0.212 | 1 |
| Contractile | STON1 | 4.01033E-18 | 0.371609 | 0.428 | 0.305 | 7.81012E-14 |
| Intermediate | FLNA | 6.17123E-46 | 0.371262 | 0.925 | 0.774 | 1.20185E-41 |
| IFN_Intermediate | TTF2 | 8.93886E-07 | 0.371228 | 0.131 | 0.038 | 0.017408428 |
| Intermediate | IRAG1 | 4.38923E-12 | 0.371103 | 0.399 | 0.279 | 8.54803E-08 |
| IFN_Intermediate | RHBDD1 | 5.03241E-06 | 0.371026 | 0.168 | 0.061 | 0.098006143 |
| VEGFA_Terminal | ANXA2 | 2.10202E-14 | 0.370759 | 0.956 | 0.923 | 4.09367E-10 |
| Fibroblasts | PAIP2 | 0.000786725 | 0.370349 | 0.518 | 0.48 | 1 |
| CD74_Intermediat | WFDC1 | 5.64884E-38 | 0.370241 | 0.842 | 0.72 | 1.10011E-33 |
| Terminal | HSD3B7 | 2.29859E-37 | 0.370215 | 0.359 | 0.209 | 4.4765E-33 |
| VEGFA_Terminal | MRPL14 | 8.79025E-13 | 0.370147 | 0.613 | 0.422 | 1.7119E-08 |
| Contractile | EVI5 | 1.72281E-20 | 0.37008 | 0.315 | 0.198 | 3.35518E-16 |
| Terminal | TUSC3 | 2.7969E-37 | 0.370063 | 0.33 | 0.186 | 5.44695E-33 |
| VEGFA_Terminal | CMPK1 | 5.99288E-14 | 0.370048 | 0.663 | 0.478 | 1.16711E-09 |
| Contractile | RAB18 | 3.70479E-21 | 0.369787 | 0.611 | 0.499 | 7.21507E-17 |
| CD74_Intermediat | HES1 | 3.92154E-36 | 0.430683 | 0.294 | 0.155 | 7.63721E-32 |
| Contractile | PRR13 | 1.48906E-21 | 0.369452 | 0.464 | 0.331 | 2.89994E-17 |
| Fibroblasts | CORO2B | 0.001130446 | 0.369341 | 0.174 | 0.113 | 1 |
| Fibroblasts | LONRF1 | 1.7244E-05 | 0.369161 | 0.161 | 0.084 | 0.335826167 |
| Pericytes | SLC2A4 | 2.32994E-05 | 0.369025 | 0.1 | 0.019 | 0.453756571 |
| VEGFA_Terminal | EVA1A | 1.39041E-59 | 0.368999 | 0.169 | 0.022 | 2.70783E-55 |
| Contractile | EP300 | 3.39169E-18 | 0.368823 | 0.385 | 0.27 | 6.60532E-14 |
| Terminal | SCX | 7.88375E-75 | 0.368607 | 0.106 | 0.009 | 1.53536E-70 |
| Terminal | LAT2 | 3.44494E-40 | 0.368596 | 0.31 | 0.163 | 6.70902E-36 |
| Contractile | ENSSSCG0( | 3.29705E-55 | 0.368591 | 0.999 | 0.996 | 6.421E-51 |
| CD74_Intermediat | IL6ST | 3.20987E-32 | 0.368403 | 0.75 | 0.637 | 6.25123E-28 |
| Intermediate | EFEMP2 | 2.07379E-13 | 0.368341 | 0.676 | 0.582 | 4.03871E-09 |
| Contractile | RUNX1T1 | 9.60332E-22 | 0.368266 | 0.269 | 0.151 | 1.87025E-17 |
| CD74_Intermediat | JUNB | 9.40666E-18 | 0.368262 | 0.616 | 0.529 | 1.83195E-13 |
| Contractile | ZCCHC17 | 7.40272E-18 | 0.367999 | 0.372 | 0.259 | 1.44168E-13 |
| Pericytes | B2M | 0.002790466 | 0.367783 | 0.9 | 0.861 | 1 |
| Contractile | SAMD4B | 2.72642E-23 | 0.367779 | 0.226 | 0.114 | 5.3097E-19 |
| Pericytes | NACA | 0.00036076 | 0.367556 | 1 | 0.983 | 1 |
| Terminal | F5 | 4.96126E-26 | 0.36748 | 0.436 | 0.298 | 9.66206E-22 |
| VEGFA_Terminal | RER1 | 1.30256E-14 | 0.367413 | 0.622 | 0.426 | 2.53674E-10 |
| Contractile | MRPL49 | 1.29748E-23 | 0.367394 | 0.275 | 0.154 | 2.52684E-19 |

| | | | | | | |
|---|---|---|---|---|---|---|
| Terminal | NGEF | 7.3088E-78 | 0.367363 | 0.164 | 0.031 | 1.42339E-73 |
| CD74_Intermediat | CXCL12 | 1.28885E-49 | 0.566099 | 0.329 | 0.158 | 2.51004E-45 |
| Contractile | SPOPL | 4.11E-20 | 0.366982 | 0.245 | 0.138 | 8.00422E-16 |
| VEGFA_Terminal | SERPINH1 | 1.54169E-09 | 0.366811 | 0.743 | 0.622 | 3.00244E-05 |
| Fibroblasts | ARL5A | 0.000181366 | 0.36638 | 0.394 | 0.32 | 1 |
| CD74_Intermediat | TCF4 | 2.16803E-29 | 0.36618 | 0.628 | 0.489 | 4.22224E-25 |
| Contractile | SERBP1 | 8.27833E-28 | 0.366057 | 0.823 | 0.749 | 1.6122E-23 |
| Contractile | HACD2 | 2.09765E-16 | 0.366051 | 0.388 | 0.28 | 4.08518E-12 |
| Contractile | TMEM51 | 1.08356E-54 | 0.366018 | 0.149 | 0.032 | 2.11024E-50 |
| Fibroblasts | EFNB1 | 4.38229E-05 | 0.365983 | 0.188 | 0.109 | 0.85345177 |
| Intermediate | ENSSSCG0( | 1.74449E-12 | 0.365953 | 0.736 | 0.652 | 3.3974E-08 |
| VEGFA_Terminal | RNF7 | 2.31195E-13 | 0.365804 | 0.63 | 0.454 | 4.50253E-09 |
| Contractile | LNPEP | 2.78481E-20 | 0.365718 | 0.359 | 0.236 | 5.42342E-16 |
| Contractile | JAG1 | 1.65381E-21 | 0.365484 | 0.573 | 0.432 | 3.22079E-17 |
| Contractile | SMC2 | 1.56882E-18 | 0.365456 | 0.219 | 0.12 | 3.05528E-14 |
| Contractile | LZTS2 | 5.59903E-18 | 0.365423 | 0.338 | 0.226 | 1.09041E-13 |
| Contractile | AAMDC | 1.65093E-16 | 0.36524 | 0.511 | 0.411 | 3.2152E-12 |
| Contractile | PYGO1 | 4.46926E-25 | 0.36495 | 0.229 | 0.113 | 8.70388E-21 |
| Fibroblasts | LOC10051: | 0.001507685 | 0.364945 | 0.248 | 0.183 | 1 |
| IFN_Intermediate | IFI35 | 9.23553E-05 | 0.364884 | 0.234 | 0.115 | 1 |
| Pericytes | EFHD2 | 0.006104894 | 0.364878 | 0.24 | 0.121 | 1 |
| Contractile | FKBP8 | 9.00782E-18 | 0.364876 | 0.509 | 0.399 | 1.75427E-13 |
| Terminal | NUCB2 | 4.167E-29 | 0.36487 | 0.517 | 0.377 | 8.11523E-25 |
| Contractile | ST13 | 1.08865E-33 | 0.364867 | 0.925 | 0.866 | 2.12015E-29 |
| VEGFA_Terminal | MYH9 | 8.10464E-08 | 0.364512 | 0.891 | 0.803 | 0.001578379 |
| Contractile | CLPX | 3.03697E-20 | 0.364211 | 0.261 | 0.151 | 5.91451E-16 |
| Contractile | PABPN1 | 6.86095E-20 | 0.364116 | 0.565 | 0.454 | 1.33617E-15 |
| Fibroblasts | SLC46A3 | 8.21151E-18 | 0.36401 | 0.142 | 0.032 | 1.59919E-13 |
| Terminal | ALDOA | 1.96807E-44 | 0.363871 | 0.902 | 0.803 | 3.83281E-40 |
| VEGFA_Terminal | LMNA | 4.58911E-12 | 0.363855 | 0.91 | 0.855 | 8.93729E-08 |
| Fibroblasts | GPR173 | 0.000949809 | 0.363739 | 0.11 | 0.06 | 1 |
| Pericytes | FMNL3 | 0.001314917 | 0.363706 | 0.2 | 0.08 | 1 |
| Fibroblasts | SEC63 | 0.00042226 | 0.363371 | 0.509 | 0.465 | 1 |
| Terminal | PTPRR | 2.25702E-47 | 0.363368 | 0.328 | 0.16 | 4.39554E-43 |
| CD74_Intermediat | CXCL16 | 5.68156E-31 | 0.363276 | 0.478 | 0.332 | 1.10648E-26 |
| VEGFA_Terminal | MGST3 | 3.17357E-14 | 0.363262 | 0.801 | 0.631 | 6.18052E-10 |
| Contractile | FKBP1A | 2.01372E-19 | 0.363023 | 0.663 | 0.563 | 3.92171E-15 |
| Fibroblasts | SDF4 | 6.06679E-07 | 0.362962 | 0.78 | 0.732 | 0.011815069 |
| Contractile | PCMTD1 | 6.82646E-16 | 0.362799 | 0.564 | 0.47 | 1.32945E-11 |
| Fibroblasts | SLC4A4 | 5.27643E-13 | 0.362635 | 0.128 | 0.035 | 1.02758E-08 |
| Intermediate | GDF6 | 7.41327E-46 | 0.362598 | 0.138 | 0.026 | 1.44373E-41 |
| Fibroblasts | KAT2B | 0.000222417 | 0.362578 | 0.193 | 0.121 | 1 |
| Contractile | NONO | 8.36051E-14 | 0.362476 | 0.418 | 0.326 | 1.62821E-09 |
| VEGFA_Terminal | HUS1 | 3.57964E-21 | 0.362314 | 0.305 | 0.135 | 6.97135E-17 |
| Fibroblasts | H6PD | 0.000589603 | 0.362246 | 0.174 | 0.109 | 1 |
| Fibroblasts | AMOTL2 | 0.001976913 | 0.362179 | 0.206 | 0.142 | 1 |
| Contractile | DCAF11 | 5.3598E-22 | 0.362088 | 0.22 | 0.113 | 1.04382E-17 |
| Fibroblasts | GATM | 3.07477E-06 | 0.362082 | 0.138 | 0.062 | 0.059881067 |
| Fibroblasts | DACT1 | 0.000370422 | 0.362059 | 0.206 | 0.134 | 1 |
| Fibroblasts | PHF3 | 0.004498149 | 0.361847 | 0.427 | 0.383 | 1 |
| Contractile | SON | 2.10151E-29 | 0.361622 | 0.908 | 0.864 | 4.09269E-25 |
| Terminal | C1QTNF2 | 4.08426E-27 | 0.361618 | 0.543 | 0.416 | 7.95409E-23 |
| Contractile | TMED5 | 1.63664E-16 | 0.361612 | 0.399 | 0.29 | 3.18735E-12 |
| Contractile | SELENOW | 5.67623E-19 | 0.361451 | 0.644 | 0.532 | 1.10545E-14 |
| Contractile | ENSSSCG0( | 2.08861E-19 | 0.361387 | 0.422 | 0.295 | 4.06756E-15 |
| VEGFA_Terminal | ENSSSCG0( | 1.67618E-18 | 0.361274 | 0.961 | 0.962 | 3.26436E-14 |
| Intermediate | ENSSSCG0( | 4.09669E-17 | 0.361187 | 0.445 | 0.29 | 7.97831E-13 |
| VEGFA_Terminal | ATP5MGL | 3.73602E-17 | 0.361126 | 0.884 | 0.771 | 7.2759E-13 |
| Contractile | CEP70 | 3.85302E-37 | 0.361023 | 0.19 | 0.067 | 7.50375E-33 |

| | | | | | | |
|---|---|---|---|---|---|---|
| VEGFA_Terminal | RALA | 6.78336E-14 | 0.361017 | 0.433 | 0.262 | 1.32106E-09 |
| CD74_Intermediat | ENSSSCG0( | 6.42836E-38 | 0.360992 | 0.774 | 0.616 | 1.25192E-33 |
| Contractile | PKN2 | 3.25626E-18 | 0.360834 | 0.475 | 0.353 | 6.34157E-14 |
| Contractile | NTF3 | 4.6028E-115 | 0.360769 | 0.111 | 0.002 | 8.964E-111 |
| Fibroblasts | PROS1 | 0.00012349 | 0.360734 | 0.624 | 0.572 | 1 |
| VEGFA_Terminal | PDXK | 2.124E-25 | 0.360674 | 0.199 | 0.062 | 4.13649E-21 |
| VEGFA_Terminal | ADAMTS12 | 4.65438E-24 | 0.360294 | 0.269 | 0.102 | 9.0644E-20 |
| VEGFA_Terminal | NFKBIZ | 2.75006E-11 | 0.360226 | 0.671 | 0.49 | 5.35575E-07 |
| VEGFA_Terminal | LAMA5 | 1.7894E-11 | 0.360195 | 0.642 | 0.455 | 3.48485E-07 |
| Terminal | ANPEP | 4.7391E-27 | 0.360135 | 0.627 | 0.468 | 9.2294E-23 |
| Contractile | MTUS2 | 3.27517E-68 | 0.36012 | 0.139 | 0.021 | 6.3784E-64 |
| IFN_Intermediate | EPS15 | 0.008242253 | 0.360055 | 0.495 | 0.396 | 1 |
| CD74_Intermediat | ENSSSCG0( | 9.33537E-32 | 0.359849 | 0.618 | 0.458 | 1.81806E-27 |
| Terminal | ENSSSCG0( | 7.24268E-17 | 0.35978 | 0.43 | 0.328 | 1.41051E-12 |
| Contractile | GADD45A | 4.51965E-21 | 0.35966 | 0.45 | 0.305 | 8.80201E-17 |
| Contractile | MPC1 | 1.45858E-19 | 0.359522 | 0.681 | 0.586 | 2.84058E-15 |
| Terminal | P4HA1 | 3.56326E-38 | 0.359443 | 0.628 | 0.453 | 6.93944E-34 |
| IFN_Intermediate | NMI | 9.57783E-05 | 0.359314 | 0.159 | 0.066 | 1 |
| VEGFA_Terminal | RRAGC | 5.02955E-11 | 0.35931 | 0.499 | 0.332 | 9.79506E-07 |
| Terminal | COL15A1 | 2.36989E-19 | 0.359305 | 0.689 | 0.603 | 4.61536E-15 |
| Pericytes | YY1 | 0.007501418 | 0.35924 | 0.66 | 0.523 | 1 |
| Terminal | MRPL17 | 4.94659E-35 | 0.359192 | 0.711 | 0.578 | 9.63348E-31 |
| Contractile | EFCAB7 | 1.93314E-12 | 0.359039 | 0.365 | 0.285 | 3.76478E-08 |
| Fibroblasts | GPR137B | 8.54217E-05 | 0.358562 | 0.211 | 0.13 | 1 |
| Contractile | TRIM24 | 4.6999E-26 | 0.358451 | 0.211 | 0.098 | 9.15306E-22 |
| IFN_Intermediate | OPTN | 0.003260289 | 0.358428 | 0.729 | 0.644 | 1 |
| Intermediate | WFDC1 | 1.1643E-10 | 0.358361 | 0.814 | 0.747 | 2.26747E-06 |
| Pericytes | ACTA2 | 5.45134E-06 | 0.358219 | 0.94 | 0.68 | 0.106164789 |
| Fibroblasts | RNASET2 | 0.00069338 | 0.358152 | 0.335 | 0.263 | 1 |
| CD74_Intermediat | CD74 | 2.349E-235 | 1.911289 | 0.55 | 0.159 | 4.5747E-231 |
| Fibroblasts | PMP22 | 0.000158562 | 0.358076 | 0.72 | 0.7 | 1 |
| Pericytes | ENSSSCG0( | 4.25555E-22 | 0.357953 | 0.12 | 0.006 | 8.28768E-18 |
| Contractile | NDUFA7 | 1.22807E-20 | 0.357952 | 0.459 | 0.329 | 2.39166E-16 |
| Pericytes | YBX1 | 0.007888283 | 0.357846 | 0.96 | 0.936 | 1 |
| Fibroblasts | VPS13A | 0.000987409 | 0.357781 | 0.211 | 0.143 | 1 |
| Contractile | NDRG2 | 1.25194E-21 | 0.357781 | 0.452 | 0.314 | 2.43815E-17 |
| Fibroblasts | HEG1 | 0.005670292 | 0.357694 | 0.344 | 0.292 | 1 |
| Contractile | PPIG | 5.46836E-18 | 0.357563 | 0.632 | 0.536 | 1.06496E-13 |
| Terminal | FAM167B | 5.26245E-30 | 0.357495 | 0.245 | 0.131 | 1.02486E-25 |
| Fibroblasts | C21ORF91 | 0.002589896 | 0.357473 | 0.33 | 0.269 | 1 |
| VEGFA_Terminal | DHRS1 | 9.08952E-16 | 0.357464 | 0.351 | 0.183 | 1.77018E-11 |
| Pericytes | SIK3 | 0.004054128 | 0.357458 | 0.18 | 0.074 | 1 |
| Contractile | PNRC1 | 1.75105E-22 | 0.357291 | 0.726 | 0.623 | 3.41018E-18 |
| VEGFA_Terminal | CREB3L1 | 8.68716E-13 | 0.357267 | 0.344 | 0.195 | 1.69182E-08 |
| Contractile | AGAP3 | 3.18767E-19 | 0.357234 | 0.209 | 0.112 | 6.20798E-15 |
| Fibroblasts | ETNK1 | 0.000389097 | 0.357224 | 0.243 | 0.165 | 1 |
| IFN_Intermediate | GNG11 | 0.00035592 | 0.357034 | 0.991 | 0.929 | 1 |
| VEGFA_Terminal | ENSSSCG0( | 2.05372E-10 | 0.356921 | 0.581 | 0.424 | 3.99963E-06 |
| Contractile | DAG1 | 3.56896E-15 | 0.356884 | 0.478 | 0.381 | 6.95054E-11 |
| VEGFA_Terminal | GBE1 | 3.7949E-13 | 0.356655 | 0.361 | 0.205 | 7.39057E-09 |
| IFN_Intermediate | MPZL1 | 0.003754013 | 0.356541 | 0.598 | 0.475 | 1 |
| Contractile | TINAGL1 | 4.16873E-43 | 0.356101 | 0.198 | 0.064 | 8.11861E-39 |
| Fibroblasts | KAZALD1 | 1.2746E-09 | 0.3557 | 0.133 | 0.045 | 2.48228E-05 |
| VEGFA_Terminal | RGS16 | 6.77651E-23 | 0.355696 | 0.133 | 0.034 | 1.31972E-18 |
| Terminal | PYCR1 | 7.64473E-31 | 0.355666 | 0.277 | 0.157 | 1.48881E-26 |
| IFN_Intermediate | EZR | 0.005784466 | 0.355556 | 0.523 | 0.411 | 1 |
| Fibroblasts | RBMS1 | 0.007504711 | 0.35544 | 0.486 | 0.469 | 1 |
| Fibroblasts | ZBTB20 | 0.001083973 | 0.355371 | 0.468 | 0.408 | 1 |
| CD74_Intermediat | PSMB8 | 3.20861E-33 | 0.355356 | 0.291 | 0.161 | 6.24877E-29 |

| | | | | | | |
|---|---|---|---|---|---|---|
| VEGFA_Terminal | MYDGF | 1.19597E-13 | 0.355283 | 0.584 | 0.39 | 2.32915E-09 |
| Intermediate | DACT1 | 1.20539E-13 | 0.355103 | 0.229 | 0.126 | 2.3475E-09 |
| Intermediate | MAP3K8 | 1.59174E-14 | 0.355018 | 0.229 | 0.125 | 3.09991E-10 |
| Fibroblasts | APBB2 | 0.003166841 | 0.355003 | 0.188 | 0.131 | 1 |
| IFN_Intermediate | ACKR2 | 0.001052968 | 0.354838 | 0.215 | 0.117 | 1 |
| Contractile | KHDRBS1 | 1.19612E-16 | 0.354825 | 0.487 | 0.378 | 2.32944E-12 |
| Pericytes | ENSSSCG0( | 0.008127757 | 0.354462 | 0.14 | 0.056 | 1 |
| Contractile | NTRK3 | 1.79834E-44 | 0.354374 | 0.158 | 0.043 | 3.50226E-40 |
| Contractile | HNRNPH3 | 2.46584E-21 | 0.354343 | 0.672 | 0.579 | 4.80222E-17 |
| Pericytes | NCKAP5 | 2.11299E-48 | 0.354179 | 0.1 | 0.001 | 4.11505E-44 |
| Contractile | RYR2 | 9.20374E-33 | 0.354026 | 0.168 | 0.06 | 1.79243E-28 |
| Intermediate | ATP1B3 | 8.98963E-12 | 0.354023 | 0.816 | 0.762 | 1.75073E-07 |
| Terminal | POGLUT3 | 1.30116E-29 | 0.353978 | 0.36 | 0.225 | 2.53401E-25 |
| Fibroblasts | CASD1 | 2.15639E-06 | 0.353401 | 0.151 | 0.071 | 0.041995714 |
| Fibroblasts | GDF10 | 8.34238E-05 | 0.35336 | 0.138 | 0.071 | 1 |
| IFN_Intermediate | DNAJC7 | 0.000736561 | 0.353305 | 0.411 | 0.259 | 1 |
| Intermediate | SULF2 | 1.07997E-08 | 0.352645 | 0.335 | 0.248 | 0.000210324 |
| Fibroblasts | N4BP2 | 2.07913E-05 | 0.352565 | 0.161 | 0.084 | 0.40491034 |
| Terminal | SLC39A14 | 1.49846E-23 | 0.35252 | 0.309 | 0.197 | 2.91825E-19 |
| Fibroblasts | SNX33 | 0.000602743 | 0.352517 | 0.197 | 0.128 | 1 |
| VEGFA_Terminal | TGFB1 | 4.69017E-16 | 0.35243 | 0.438 | 0.241 | 9.13411E-12 |
| Intermediate | MAP2 | 1.19918E-08 | 0.352351 | 0.369 | 0.277 | 0.000233541 |
| Fibroblasts | ENSSSCG0( | 8.65507E-07 | 0.351875 | 0.78 | 0.775 | 0.016855749 |
| Intermediate | FGF2 | 1.04841E-07 | 0.351838 | 0.599 | 0.524 | 0.002041774 |
| Terminal | CD109 | 2.45801E-22 | 0.351761 | 0.578 | 0.465 | 4.78697E-18 |
| Fibroblasts | TAPT1 | 0.000606084 | 0.351668 | 0.174 | 0.108 | 1 |
| Fibroblasts | CHIC2 | 0.009720375 | 0.351578 | 0.358 | 0.326 | 1 |
| VEGFA_Terminal | MYBL1 | 9.40792E-10 | 0.351509 | 0.373 | 0.234 | 1.83219E-05 |
| VEGFA_Terminal | SLC9A3R2 | 3.85435E-27 | 0.351437 | 0.264 | 0.093 | 7.50634E-23 |
| Contractile | AMIGO2 | 5.24044E-29 | 0.351409 | 0.168 | 0.064 | 1.02058E-24 |
| Contractile | VEZF1 | 5.64346E-15 | 0.351368 | 0.366 | 0.261 | 1.09906E-10 |
| Contractile | TRIP6 | 3.16015E-27 | 0.35136 | 0.294 | 0.159 | 6.1544E-23 |
| Fibroblasts | SH3D19 | 5.15413E-05 | 0.351336 | 0.463 | 0.378 | 1 |
| VEGFA_Terminal | MTSS1 | 1.98425E-36 | 0.351287 | 0.177 | 0.039 | 3.86433E-32 |
| Contractile | SLC2A12 | 1.67916E-63 | 0.351261 | 0.122 | 0.017 | 3.27016E-59 |
| VEGFA_Terminal | ENSSSCG0( | 5.06263E-20 | 0.351179 | 0.15 | 0.046 | 9.85948E-16 |
| Fibroblasts | LGR4 | 7.36929E-08 | 0.351125 | 0.165 | 0.071 | 0.001435168 |
| VEGFA_Terminal | EDEM1 | 1.5441E-17 | 0.351119 | 0.31 | 0.145 | 3.00714E-13 |
| VEGFA_Terminal | CGREF1 | 2.75887E-07 | 0.351114 | 0.242 | 0.15 | 0.005372901 |
| Fibroblasts | TM9SF3 | 0.000297407 | 0.351003 | 0.624 | 0.605 | 1 |
| CD74_Intermediat | AEBP1 | 3.03798E-46 | 0.350922 | 0.948 | 0.865 | 5.91647E-42 |
| Contractile | YTHDC1 | 7.46044E-16 | 0.350876 | 0.497 | 0.393 | 1.45292E-11 |
| IFN_Intermediate | TRIM38 | 0.000139577 | 0.350782 | 0.206 | 0.098 | 1 |
| CD74_Intermediat | ARSK | 1.49952E-38 | 0.367016 | 0.306 | 0.162 | 2.92032E-34 |
| Terminal | AP1S2 | 5.5297E-37 | 0.350389 | 0.4 | 0.241 | 1.07691E-32 |
| Fibroblasts | IRF2BP2 | 0.003032979 | 0.350357 | 0.472 | 0.438 | 1 |
| Contractile | PTBP2 | 6.05747E-26 | 0.350292 | 0.21 | 0.097 | 1.17969E-21 |
| VEGFA_Terminal | ENSSSCG0( | 5.99916E-14 | 0.350288 | 0.511 | 0.318 | 1.16834E-09 |
| Terminal | ITM2C | 7.75216E-38 | 0.350198 | 0.787 | 0.662 | 1.50973E-33 |
| Terminal | PDGFC | 2.73241E-40 | 0.350197 | 0.247 | 0.115 | 5.32136E-36 |
| Terminal | ITGA10 | 3.76678E-35 | 0.350186 | 0.28 | 0.148 | 7.3358E-31 |
| VEGFA_Terminal | SNRPG | 1.54583E-10 | 0.350119 | 0.743 | 0.59 | 3.0105E-06 |
| Contractile | SMAD4 | 6.70468E-15 | 0.350105 | 0.368 | 0.268 | 1.30574E-10 |
| Pericytes | PLK3 | 0.000668119 | 0.349894 | 0.16 | 0.054 | 1 |
| Intermediate | EPDR1 | 2.67567E-12 | 0.349611 | 0.255 | 0.156 | 5.21088E-08 |
| Contractile | CNST | 3.08253E-23 | 0.349567 | 0.27 | 0.149 | 6.00323E-19 |
| IFN_Intermediate | MAD2L2 | 0.001749962 | 0.349446 | 0.243 | 0.139 | 1 |
| IFN_Intermediate | ADAR | 1.95831E-08 | 0.349416 | 0.178 | 0.053 | 0.00038138 |
| CD74_Intermediat | CLEC3B | 1.21134E-27 | 0.476071 | 0.284 | 0.162 | 2.35908E-23 |

| | | | | | | |
|---|---|---|---|---|---|---|
| Terminal | RCN2 | 5.38324E-32 | 0.349357 | 0.699 | 0.573 | 1.04839E-27 |
| VEGFA_Terminal | PDIA6 | 2.13269E-09 | 0.349291 | 0.668 | 0.523 | 4.15342E-05 |
| Contractile | RXRA | 8.69658E-24 | 0.349265 | 0.207 | 0.1 | 1.69366E-19 |
| VEGFA_Terminal | ANXA1 | 6.67652E-10 | 0.349028 | 0.85 | 0.738 | 1.30025E-05 |
| Terminal | BBLN | 2.9273E-24 | 0.349024 | 0.417 | 0.292 | 5.70091E-20 |
| Fibroblasts | WBP1L | 0.000535734 | 0.349022 | 0.211 | 0.137 | 1 |
| Terminal | HSPA8 | 2.14776E-28 | 0.348858 | 0.772 | 0.69 | 4.18276E-24 |
| Terminal | TPM3 | 2.16572E-35 | 0.348837 | 0.634 | 0.472 | 4.21774E-31 |
| Contractile | TMOD1 | 1.16716E-24 | 0.348804 | 0.223 | 0.109 | 2.27305E-20 |
| Fibroblasts | IMP3 | 0.000711219 | 0.34876 | 0.284 | 0.213 | 1 |
| IFN_Intermediate | MYCBP2 | 0.009325902 | 0.348753 | 0.467 | 0.344 | 1 |
| Fibroblasts | MKNK2 | 0.000752024 | 0.34872 | 0.211 | 0.138 | 1 |
| CD74_Intermediat | COL5A3 | 8.53008E-22 | 0.250053 | 0.265 | 0.162 | 1.66123E-17 |
| VEGFA_Terminal | P4HA3 | 2.46568E-26 | 0.348594 | 0.194 | 0.058 | 4.80191E-22 |
| IFN_Intermediate | TRIM21 | 4.77943E-08 | 0.348589 | 0.15 | 0.042 | 0.000930795 |
| IFN_Intermediate | BATF2 | 6.97675E-59 | 0.348589 | 0.131 | 0.004 | 1.35872E-54 |
| IFN_Intermediate | GLUL | 0.006204101 | 0.348343 | 0.495 | 0.388 | 1 |
| Contractile | CDV3 | 2.23696E-20 | 0.348298 | 0.671 | 0.571 | 4.35649E-16 |
| Pericytes | FMO3 | 8.45051E-97 | 0.348262 | 0.1 | 0 | 1.64574E-92 |
| Fibroblasts | SSR2 | 0.002612493 | 0.348108 | 0.518 | 0.492 | 1 |
| VEGFA_Terminal | YBX1 | 9.54814E-19 | 0.348078 | 0.976 | 0.933 | 1.8595E-14 |
| Fibroblasts | CDKN1B | 0.002033353 | 0.348019 | 0.404 | 0.352 | 1 |
| Contractile | ENSSSCG0( | 1.35505E-13 | 0.347983 | 0.322 | 0.23 | 2.63895E-09 |
| VEGFA_Terminal | C4ORF48 | 2.3611E-16 | 0.347911 | 0.518 | 0.303 | 4.59824E-12 |
| Pericytes | SGTB | 6.4209E-07 | 0.347576 | 0.12 | 0.02 | 0.012504707 |
| VEGFA_Terminal | APC | 7.22425E-10 | 0.347568 | 0.416 | 0.272 | 1.40692E-05 |
| IFN_Intermediate | TNFAIP8 | 0.000860769 | 0.347528 | 0.402 | 0.261 | 1 |
| Fibroblasts | TMEM248 | 0.002732928 | 0.347468 | 0.362 | 0.311 | 1 |
| VEGFA_Terminal | ENSSSCG0( | 2.11744E-56 | 0.347371 | 0.143 | 0.017 | 4.12372E-52 |
| CD74_Intermediat | ITGB5 | 2.59667E-29 | 0.347317 | 0.669 | 0.541 | 5.05701E-25 |
| Contractile | PHACTR1 | 9.16693E-93 | 0.347298 | 0.115 | 0.007 | 1.78526E-88 |
| Contractile | CCDC186 | 5.25942E-17 | 0.347076 | 0.545 | 0.439 | 1.02427E-12 |
| Contractile | TM4SF1 | 2.2604E-12 | 0.346888 | 0.711 | 0.654 | 4.40213E-08 |
| Pericytes | KCNE4 | 4.04913E-25 | 0.346816 | 0.12 | 0.005 | 7.88568E-21 |
| Fibroblasts | KLF12 | 0.009305552 | 0.346571 | 0.216 | 0.164 | 1 |
| Fibroblasts | DCLK1 | 6.59947E-07 | 0.346407 | 0.133 | 0.056 | 0.012852475 |
| VEGFA_Terminal | STMP1 | 9.14181E-13 | 0.346382 | 0.455 | 0.285 | 1.78037E-08 |
| Fibroblasts | LBR | 0.000208198 | 0.346039 | 0.179 | 0.107 | 1 |
| Fibroblasts | LENG8 | 0.000532902 | 0.345948 | 0.28 | 0.206 | 1 |
| Pericytes | MBP | 3.30468E-09 | 0.345939 | 0.12 | 0.015 | 6.43587E-05 |
| VEGFA_Terminal | PGM3 | 4.36699E-17 | 0.345692 | 0.346 | 0.175 | 8.50472E-13 |
| Pericytes | PPP1R3E | 0.003875257 | 0.345582 | 0.18 | 0.075 | 1 |
| Fibroblasts | UGCG | 4.85424E-05 | 0.345581 | 0.17 | 0.094 | 0.94536262 |
| Pericytes | ENSSSCG0( | 0.004767788 | 0.34552 | 0.12 | 0.042 | 1 |
| Intermediate | NDUFB1 | 1.13404E-05 | 0.345044 | 0.601 | 0.572 | 0.220853572 |
| Fibroblasts | GNG11 | 4.73506E-07 | 0.344966 | 0.963 | 0.929 | 0.009221537 |
| VEGFA_Terminal | CD63 | 1.1874E-22 | 0.344869 | 0.981 | 0.974 | 2.31247E-18 |
| VEGFA_Terminal | CLEC11A | 1.96701E-11 | 0.344669 | 0.564 | 0.382 | 3.83075E-07 |
| CD74_Intermediat | OSR1 | 4.40684E-24 | 0.34464 | 0.461 | 0.328 | 8.58233E-20 |
| Contractile | PSMC5 | 2.01981E-16 | 0.344627 | 0.585 | 0.496 | 3.93358E-12 |
| Contractile | PHACTR2 | 1.26039E-10 | 0.344474 | 0.355 | 0.277 | 2.45461E-06 |
| Intermediate | ITGAV | 1.03203E-12 | 0.344464 | 0.69 | 0.601 | 2.00988E-08 |
| Contractile | PDLIM2 | 1.25315E-18 | 0.344441 | 0.303 | 0.189 | 2.44051E-14 |
| VEGFA_Terminal | TWF2 | 1.07549E-14 | 0.344083 | 0.429 | 0.246 | 2.09452E-10 |
| IFN_Intermediate | EMP1 | 0.00316619 | 0.344071 | 0.785 | 0.645 | 1 |
| VEGFA_Terminal | HRAS | 2.8859E-12 | 0.34386 | 0.443 | 0.274 | 5.62028E-08 |
| VEGFA_Terminal | THOC7 | 2.40303E-12 | 0.343714 | 0.666 | 0.487 | 4.6799E-08 |
| Contractile | TMEM144 | 2.55937E-39 | 0.343642 | 0.147 | 0.041 | 4.98437E-35 |
| CD74_Intermediat | RENBP | 6.21314E-18 | 0.252275 | 0.256 | 0.162 | 1.21001E-13 |

| | | | | | | |
|---|---|---|---|---|---|---|
| Contractile | KLHL23 | 5.08227E-44 | 0.34342 | 0.159 | 0.043 | 9.89771E-40 |
| Contractile | TTC39B | 1.55596E-23 | 0.343404 | 0.203 | 0.097 | 3.03023E-19 |
| Intermediate | NFKBIZ | 1.40433E-07 | 0.343214 | 0.576 | 0.494 | 0.002734926 |
| Contractile | MICALL1 | 3.64411E-54 | 0.342934 | 0.124 | 0.022 | 7.09691E-50 |
| Fibroblasts | TPP2 | 0.001284795 | 0.342932 | 0.183 | 0.122 | 1 |
| VEGFA_Terminal | HSD3B7 | 7.02745E-11 | 0.342874 | 0.392 | 0.248 | 1.3686E-06 |
| VEGFA_Terminal | STC2 | 6.03647E-33 | 0.34283 | 0.131 | 0.024 | 1.1756E-28 |
| Contractile | YPEL5 | 2.48478E-15 | 0.342809 | 0.459 | 0.361 | 4.83911E-11 |
| Contractile | COX7A1 | 7.52627E-13 | 0.342739 | 0.529 | 0.449 | 1.46574E-08 |
| VEGFA_Terminal | AVPI1 | 6.25358E-16 | 0.342725 | 0.237 | 0.105 | 1.21789E-11 |
| Pericytes | B2M.1 | 0.00215953 | 0.34271 | 0.94 | 0.875 | 1 |
| Terminal | MCUB | 2.03858E-37 | 0.342687 | 0.354 | 0.201 | 3.97014E-33 |
| Contractile | CCT2 | 1.27671E-15 | 0.342554 | 0.598 | 0.515 | 2.48639E-11 |
| Contractile | NEK9 | 7.13539E-20 | 0.34242 | 0.312 | 0.195 | 1.38962E-15 |
| Intermediate | ENSSSCG0( | 1.54873E-24 | 0.342334 | 0.886 | 0.797 | 3.01615E-20 |
| Contractile | ENSSSCG0( | 1.42334E-13 | 0.342319 | 0.257 | 0.164 | 2.77196E-09 |
| CD74_Intermediat | ENSSSCG0( | 9.81828E-14 | 0.288571 | 0.239 | 0.162 | 1.91211E-09 |
| Pericytes | ANKRD24 | 9.02353E-34 | 0.341834 | 0.1 | 0.002 | 1.75733E-29 |
| Intermediate | DENND2A | 5.80758E-21 | 0.341665 | 0.163 | 0.063 | 1.13103E-16 |
| Contractile | GNG12 | 4.71202E-16 | 0.34162 | 0.662 | 0.588 | 9.17665E-12 |
| Contractile | NDUFS6 | 6.33738E-16 | 0.341515 | 0.576 | 0.491 | 1.23421E-11 |
| Fibroblasts | ENSSSCG0( | 1.62703E-13 | 0.341075 | 1 | 0.996 | 3.16863E-09 |
| Fibroblasts | SMG1 | 0.003756646 | 0.341034 | 0.353 | 0.306 | 1 |
| IFN_Intermediate | MOV10 | 2.32961E-05 | 0.340778 | 0.121 | 0.04 | 0.45369085 |
| CD74_Intermediat | AQP3 | 1.1801E-21 | 0.340729 | 0.52 | 0.39 | 2.29824E-17 |
| VEGFA_Terminal | LIMS1 | 2.11205E-09 | 0.340668 | 0.663 | 0.512 | 4.11322E-05 |
| Contractile | MAP1LC3E | 2.94714E-19 | 0.340646 | 0.714 | 0.638 | 5.73956E-15 |
| Contractile | PDCL3 | 2.77275E-21 | 0.340643 | 0.197 | 0.098 | 5.39994E-17 |
| Contractile | CAVIN2 | 2.11009E-13 | 0.340604 | 0.319 | 0.225 | 4.10939E-09 |
| Terminal | MGP | 3.37737E-10 | 0.340598 | 0.999 | 0.998 | 6.57743E-06 |
| Contractile | ARMCX2 | 1.73029E-45 | 0.340534 | 0.155 | 0.04 | 3.36974E-41 |
| Contractile | STBD1 | 1.08364E-18 | 0.340532 | 0.356 | 0.238 | 2.11039E-14 |
| Terminal | RCAN1 | 4.82214E-18 | 0.340497 | 0.217 | 0.132 | 9.39111E-14 |
| Fibroblasts | CIRBP | 0.001325996 | 0.340351 | 0.445 | 0.39 | 1 |
| Terminal | NMB | 3.84492E-23 | 0.340143 | 0.194 | 0.102 | 7.48799E-19 |
| Contractile | LOC10051 | 1.20812E-33 | 0.340024 | 0.176 | 0.064 | 2.35281E-29 |
| Fibroblasts | MGP | 0.000367505 | 0.339935 | 0.991 | 0.999 | 1 |
| Fibroblasts | ZHX3 | 0.002491702 | 0.339894 | 0.147 | 0.092 | 1 |
| IFN_Intermediate | CD47 | 0.002064181 | 0.339714 | 0.701 | 0.583 | 1 |
| Contractile | MARCHF2 | 6.13782E-16 | 0.339628 | 0.335 | 0.231 | 1.19534E-11 |
| CD74_Intermediat | RAP2A | 1.72925E-40 | 0.379705 | 0.31 | 0.163 | 3.36771E-36 |
| Fibroblasts | LRRK1 | 0.000367307 | 0.33909 | 0.183 | 0.112 | 1 |
| Terminal | C4ORF3 | 3.01659E-29 | 0.338948 | 0.772 | 0.678 | 5.87482E-25 |
| Fibroblasts | PDCD4 | 0.000309943 | 0.338892 | 0.56 | 0.501 | 1 |
| VEGFA_Terminal | NME1 | 1.16609E-17 | 0.338883 | 0.402 | 0.21 | 2.27097E-13 |
| Contractile | MYOM1 | 1.17366E-57 | 0.338681 | 0.147 | 0.029 | 2.2857E-53 |
| VEGFA_Terminal | MRPL28 | 2.69597E-08 | 0.33857 | 0.644 | 0.501 | 0.00052504 |
| Pericytes | NME4 | 0.002223388 | 0.338468 | 0.1 | 0.028 | 1 |
| Contractile | MAP4K3 | 1.06423E-16 | 0.338222 | 0.255 | 0.156 | 2.07259E-12 |
| Fibroblasts | KLHL9 | 0.001555321 | 0.338179 | 0.252 | 0.183 | 1 |
| Contractile | SCAF11 | 1.18776E-15 | 0.338007 | 0.501 | 0.4 | 2.31317E-11 |
| Terminal | CEMIP | 1.01962E-54 | 0.337842 | 0.126 | 0.027 | 1.98572E-50 |
| Fibroblasts | RECK | 0.001111218 | 0.33779 | 0.183 | 0.12 | 1 |
| Contractile | KMT2E | 4.72052E-15 | 0.337743 | 0.674 | 0.585 | 9.19321E-11 |
| VEGFA_Terminal | RACK1 | 1.80741E-14 | 0.337553 | 0.966 | 0.933 | 3.51993E-10 |
| Fibroblasts | SMARCD2 | 7.01223E-05 | 0.337496 | 0.17 | 0.096 | 1 |
| Fibroblasts | PTPRG | 0.003491914 | 0.337482 | 0.257 | 0.197 | 1 |
| Terminal | LOXL3 | 3.30532E-34 | 0.337453 | 0.188 | 0.083 | 6.4371E-30 |
| Contractile | HIF1A | 2.47476E-19 | 0.337341 | 0.711 | 0.624 | 4.8196E-15 |

| | | | | | | |
|---|---|---|---|---|---|---|
| Contractile | ATOSA | 3.98672E-19 | 0.337326 | 0.213 | 0.114 | 7.76414E-15 |
| Pericytes | RASGRP2 | 7.12577E-09 | 0.337093 | 0.1 | 0.011 | 0.000138774 |
| Intermediate | SERPINB9 | 1.00492E-18 | 0.336913 | 0.184 | 0.081 | 1.95709E-14 |
| CD74_Intermediat | CLU | 7.88173E-60 | 0.336812 | 0.997 | 0.948 | 1.53497E-55 |
| Contractile | CBLB | 1.0413E-17 | 0.336799 | 0.38 | 0.261 | 2.02793E-13 |
| Contractile | NUDT7 | 4.66617E-22 | 0.336778 | 0.241 | 0.128 | 9.08737E-18 |
| Fibroblasts | BRD3 | 0.00554889 | 0.336634 | 0.349 | 0.299 | 1 |
| Fibroblasts | PLSCR1 | 3.28857E-06 | 0.336611 | 0.243 | 0.141 | 0.064044864 |
| Contractile | ITGA9 | 5.6745E-16 | 0.336538 | 0.22 | 0.127 | 1.10511E-11 |
| Contractile | ADCY6 | 3.06348E-28 | 0.336435 | 0.168 | 0.066 | 5.96613E-24 |
| Contractile | FOXP2 | 2.13322E-30 | 0.336428 | 0.174 | 0.066 | 4.15445E-26 |
| VEGFA_Terminal | MVP | 2.50083E-09 | 0.336289 | 0.409 | 0.27 | 4.87037E-05 |
| VEGFA_Terminal | ATP13A3 | 1.24328E-16 | 0.336253 | 0.298 | 0.143 | 2.42128E-12 |
| IFN_Intermediate | AP2A2 | 0.000909082 | 0.336227 | 0.346 | 0.224 | 1 |
| Fibroblasts | RAD21 | 0.004953869 | 0.33622 | 0.422 | 0.374 | 1 |
| Intermediate | MYLK | 3.42094E-35 | 0.335793 | 0.793 | 0.583 | 6.66228E-31 |
| IFN_Intermediate | PLAT | 0.000101099 | 0.335768 | 0.467 | 0.294 | 1 |
| Intermediate | CFI | 8.13522E-21 | 0.335675 | 0.232 | 0.106 | 1.58433E-16 |
| Terminal | TMED3 | 3.98573E-25 | 0.335553 | 0.384 | 0.261 | 7.76221E-21 |
| Fibroblasts | PNN | 0.001336033 | 0.335435 | 0.615 | 0.612 | 1 |
| Intermediate | NPR3 | 5.26361E-44 | 0.335427 | 0.112 | 0.018 | 1.02509E-39 |
| Contractile | MRPS26 | 3.00678E-18 | 0.33534 | 0.581 | 0.468 | 5.8557E-14 |
| Contractile | PARP8 | 4.81569E-24 | 0.335316 | 0.225 | 0.11 | 9.37855E-20 |
| Contractile | CCNI | 1.36112E-16 | 0.335188 | 0.68 | 0.584 | 2.65077E-12 |
| VEGFA_Terminal | SSR2 | 9.7361E-13 | 0.33516 | 0.654 | 0.481 | 1.89611E-08 |
| Intermediate | IL17B | 2.90147E-07 | 0.335134 | 0.289 | 0.208 | 0.00565061 |
| Contractile | NAB1 | 4.88136E-17 | 0.334855 | 0.366 | 0.253 | 9.50646E-13 |
| Contractile | ANKRD46 | 1.83202E-17 | 0.334827 | 0.288 | 0.181 | 3.56786E-13 |
| Contractile | DHX40 | 1.11534E-14 | 0.334813 | 0.38 | 0.279 | 2.17213E-10 |
| Fibroblasts | TNFAIP6 | 0.000128768 | 0.334811 | 0.211 | 0.13 | 1 |
| Pericytes | ENSSSCG0( | 0.00445105 | 0.334712 | 0.14 | 0.052 | 1 |
| Contractile | PALM2AKA | 3.21188E-18 | 0.334584 | 0.533 | 0.419 | 6.25513E-14 |
| CD74_Intermediat | ARL4A | 3.44011E-16 | 0.281404 | 0.251 | 0.163 | 6.69962E-12 |
| VEGFA_Terminal | BACE2 | 2.85222E-15 | 0.334516 | 0.409 | 0.229 | 5.5547E-11 |
| VEGFA_Terminal | CPNE8 | 5.58716E-14 | 0.334449 | 0.22 | 0.102 | 1.0881E-09 |
| Fibroblasts | ALDH2 | 0.005791085 | 0.334263 | 0.601 | 0.549 | 1 |
| Contractile | DAPK3 | 2.16068E-14 | 0.334232 | 0.218 | 0.133 | 4.20792E-10 |
| VEGFA_Terminal | VKORC1 | 1.49197E-12 | 0.334203 | 0.792 | 0.636 | 2.90562E-08 |
| Intermediate | BGN | 1.56624E-26 | 0.334057 | 0.998 | 0.989 | 3.05024E-22 |
| Intermediate | LITAF | 4.0166E-10 | 0.333991 | 0.553 | 0.451 | 7.82232E-06 |
| Pericytes | ENSSSCG0( | 4.22322E-28 | 0.333958 | 0.1 | 0.003 | 8.22471E-24 |
| VEGFA_Terminal | TOLLIP | 5.66528E-12 | 0.333887 | 0.47 | 0.303 | 1.10331E-07 |
| VEGFA_Terminal | ANO1 | 5.04032E-09 | 0.333829 | 0.438 | 0.298 | 9.81602E-05 |
| IFN_Intermediate | PDCD4 | 0.000344442 | 0.333765 | 0.682 | 0.5 | 1 |
| Fibroblasts | DPYD | 0.003282113 | 0.333574 | 0.197 | 0.139 | 1 |
| Fibroblasts | KCTD9 | 0.000552287 | 0.333493 | 0.197 | 0.128 | 1 |
| CD74_Intermediat | SELENBP1 | 3.55582E-90 | 0.671088 | 0.407 | 0.165 | 6.92496E-86 |
| Terminal | FSTL1 | 2.53703E-26 | 0.333199 | 0.873 | 0.837 | 4.94087E-22 |
| Contractile | EVA1C | 1.60071E-24 | 0.333055 | 0.199 | 0.091 | 3.11738E-20 |
| Contractile | PA2G4 | 3.13738E-18 | 0.333011 | 0.659 | 0.561 | 6.11005E-14 |
| Contractile | CD151 | 5.28528E-18 | 0.332949 | 0.767 | 0.686 | 1.02931E-13 |
| Fibroblasts | EDEM2 | 1.04629E-05 | 0.332932 | 0.161 | 0.082 | 0.203764304 |
| Pericytes | DCUN1D3 | 8.53596E-06 | 0.332845 | 0.12 | 0.024 | 0.166237732 |
| Contractile | LARP4 | 4.2531E-16 | 0.33283 | 0.262 | 0.163 | 8.28291E-12 |
| CD74_Intermediat | IL34 | 1.65993E-17 | 0.254398 | 0.259 | 0.167 | 3.23271E-13 |
| Fibroblasts | GMFB | 1.6031E-06 | 0.332353 | 0.165 | 0.08 | 0.031220295 |
| Contractile | DGKG | 2.73886E-78 | 0.332283 | 0.11 | 0.009 | 5.33393E-74 |
| Terminal | AGFG1 | 2.0478E-28 | 0.332273 | 0.334 | 0.208 | 3.98809E-24 |
| Intermediate | ENSSSCG0( | 7.52931E-20 | 0.332267 | 0.134 | 0.047 | 1.46633E-15 |

| | | | | | | |
|---|---|---|---|---|---|---|
| VEGFA_Terminal | CALR | 1.2305E-06 | 0.332174 | 0.799 | 0.686 | 0.023963914 |
| Fibroblasts | GPC1 | 5.53469E-08 | 0.332122 | 0.142 | 0.056 | 0.00107788 |
| VEGFA_Terminal | CAPZA1 | 8.69871E-14 | 0.332099 | 0.504 | 0.312 | 1.69407E-09 |
| VEGFA_Terminal | IL18BP | 2.76983E-19 | 0.332047 | 0.16 | 0.052 | 5.39424E-15 |
| VEGFA_Terminal | ARHGEF2 | 3.42498E-08 | 0.332032 | 0.387 | 0.257 | 0.000667015 |
| IFN_Intermediate | LGALS8 | 4.9101E-05 | 0.331883 | 0.252 | 0.123 | 0.956242029 |
| Contractile | ARHGEF10 | 4.53607E-14 | 0.331835 | 0.261 | 0.169 | 8.834E-10 |
| Fibroblasts | SOCS1 | 6.84799E-09 | 0.331782 | 0.133 | 0.048 | 0.000133365 |
| Fibroblasts | ENSSSCG0( | 0.000597781 | 0.331719 | 0.665 | 0.626 | 1 |
| IFN_Intermediate | TMEM100 | 0.00011949 | 0.331713 | 0.271 | 0.14 | 1 |
| Fibroblasts | NRF1 | 0.00019262 | 0.331684 | 0.156 | 0.088 | 1 |
| Fibroblasts | PRPF4B | 0.000267749 | 0.331661 | 0.5 | 0.449 | 1 |
| Fibroblasts | TWSG1 | 0.003175719 | 0.331409 | 0.358 | 0.303 | 1 |
| Terminal | SEC61B | 1.35287E-32 | 0.331277 | 0.843 | 0.754 | 2.63471E-28 |
| Contractile | RIMKLB | 5.69129E-21 | 0.331025 | 0.256 | 0.141 | 1.10838E-16 |
| Fibroblasts | ATXN10 | 0.003667696 | 0.33087 | 0.28 | 0.221 | 1 |
| Pericytes | CMTM4 | 0.006193431 | 0.330812 | 0.16 | 0.065 | 1 |
| Terminal | ATP5MC1 | 2.34683E-25 | 0.330743 | 0.645 | 0.53 | 4.57044E-21 |
| VEGFA_Terminal | LBH | 4.41053E-14 | 0.330519 | 0.254 | 0.122 | 8.5895E-10 |
| Contractile | CFL1 | 7.73882E-28 | 0.330412 | 0.897 | 0.834 | 1.50714E-23 |
| Fibroblasts | HNRNPA2E | 1.30455E-05 | 0.33033 | 0.885 | 0.899 | 0.2540613 |
| VEGFA_Terminal | NIBAN2 | 7.38779E-17 | 0.330248 | 0.31 | 0.15 | 1.43877E-12 |
| Intermediate | SMARCD3 | 2.70137E-11 | 0.330189 | 0.349 | 0.236 | 5.26091E-07 |
| Contractile | ANXA6 | 1.41135E-14 | 0.330174 | 0.431 | 0.328 | 2.7486E-10 |
| Contractile | CROT | 7.05692E-19 | 0.329937 | 0.202 | 0.106 | 1.37434E-14 |
| IFN_Intermediate | CIITA | 3.07968E-38 | 0.329756 | 0.121 | 0.006 | 5.99768E-34 |
| IFN_Intermediate | ENSSSCG0( | 4.19664E-07 | 0.329702 | 0.159 | 0.05 | 0.008172959 |
| Contractile | GJC1 | 4.25201E-19 | 0.329667 | 0.269 | 0.16 | 8.2808E-15 |
| VEGFA_Terminal | MAP4K4 | 2.11404E-11 | 0.329634 | 0.477 | 0.309 | 4.1171E-07 |
| Intermediate | PODN | 6.80867E-14 | 0.329627 | 0.189 | 0.097 | 1.32599E-09 |
| Contractile | CASKIN2 | 1.05215E-33 | 0.3296 | 0.153 | 0.05 | 2.04906E-29 |
| CD74_Intermediat | SDC2 | 2.06246E-37 | 0.32946 | 0.874 | 0.764 | 4.01665E-33 |
| Terminal | CCDC141 | 4.17089E-33 | 0.329431 | 0.252 | 0.131 | 8.12281E-29 |
| Contractile | NKTR | 2.48824E-13 | 0.329386 | 0.475 | 0.373 | 4.84585E-09 |
| CD74_Intermediat | COL21A1 | 7.68842E-61 | 0.506526 | 0.373 | 0.178 | 1.49732E-56 |
| Contractile | RBFOX2 | 2.03229E-16 | 0.329302 | 0.455 | 0.343 | 3.95788E-12 |
| Contractile | ZEB1 | 3.06463E-16 | 0.329229 | 0.343 | 0.232 | 5.96837E-12 |
| Fibroblasts | POFUT1 | 3.62861E-05 | 0.32908 | 0.202 | 0.117 | 0.706672043 |
| Terminal | VKORC1 | 4.60451E-26 | 0.328942 | 0.719 | 0.612 | 8.96728E-22 |
| Terminal | PGK1 | 2.92086E-30 | 0.328689 | 0.59 | 0.44 | 5.68837E-26 |
| CD74_Intermediat | BICC1 | 9.41688E-26 | 0.278275 | 0.303 | 0.182 | 1.83394E-21 |
| CD74_Intermediat | ARPC1B | 1.03854E-27 | 0.328513 | 0.707 | 0.587 | 2.02255E-23 |
| Contractile | EIF3D | 6.7582E-16 | 0.328277 | 0.58 | 0.477 | 1.31616E-11 |
| Contractile | VPS25 | 1.0969E-14 | 0.328102 | 0.431 | 0.331 | 2.13621E-10 |
| VEGFA_Terminal | UBE2E1 | 7.11505E-11 | 0.32804 | 0.455 | 0.298 | 1.38566E-06 |
| Contractile | PHTF2 | 3.62571E-11 | 0.327992 | 0.378 | 0.292 | 7.06108E-07 |
| IFN_Intermediate | ADD3 | 0.000132379 | 0.32788 | 0.607 | 0.424 | 1 |
| Intermediate | SAMD4A | 1.2494E-06 | 0.327844 | 0.551 | 0.481 | 0.02433209 |
| Fibroblasts | TOPBP1 | 0.004685211 | 0.327214 | 0.17 | 0.117 | 1 |
| VEGFA_Terminal | BTG1 | 8.75227E-14 | 0.32717 | 0.249 | 0.12 | 1.7045E-09 |
| Intermediate | CLIP1 | 1.66863E-10 | 0.327127 | 0.664 | 0.564 | 3.24965E-06 |
| Fibroblasts | IRS2 | 0.001045094 | 0.326995 | 0.211 | 0.142 | 1 |
| Contractile | MPDZ | 1.74095E-15 | 0.326887 | 0.423 | 0.311 | 3.3905E-11 |
| Intermediate | DNAJB4 | 1.57893E-09 | 0.326807 | 0.599 | 0.521 | 3.07496E-05 |
| Terminal | TAGLN2 | 2.13297E-26 | 0.326776 | 0.827 | 0.731 | 4.15396E-22 |
| Fibroblasts | ENSSSCG0( | 0.000940365 | 0.326758 | 0.229 | 0.159 | 1 |
| Fibroblasts | SERINC3 | 0.003001111 | 0.326693 | 0.555 | 0.55 | 1 |
| Contractile | KRR1 | 2.05593E-16 | 0.326568 | 0.295 | 0.191 | 4.00391E-12 |
| Contractile | CTBP2 | 3.19933E-13 | 0.326545 | 0.26 | 0.173 | 6.23069E-09 |

| | | | | | | |
|---|---|---|---|---|---|---|
| VEGFA_Terminal | MANF | 5.34546E-08 | 0.326274 | 0.591 | 0.45 | 0.001041029 |
| Contractile | EIF4H | 4.66073E-14 | 0.326239 | 0.59 | 0.507 | 9.07678E-10 |
| VEGFA_Terminal | TUBB2A | 9.0521E-08 | 0.326198 | 0.421 | 0.298 | 0.001762896 |
| Contractile | PRDX3 | 5.71348E-18 | 0.326109 | 0.329 | 0.217 | 1.1127E-13 |
| Contractile | MED28 | 3.04359E-19 | 0.325992 | 0.213 | 0.115 | 5.92739E-15 |
| IFN_Intermediate | UBA7 | 4.30629E-11 | 0.32598 | 0.112 | 0.02 | 8.38649E-07 |
| Fibroblasts | SKAP2 | 0.007912532 | 0.325972 | 0.193 | 0.141 | 1 |
| VEGFA_Terminal | ARL8B | 2.04338E-09 | 0.325961 | 0.581 | 0.429 | 3.97949E-05 |
| Contractile | DYSF | 1.61278E-62 | 0.325829 | 0.113 | 0.014 | 3.14089E-58 |
| Terminal | NDUFB2 | 8.20431E-22 | 0.325819 | 0.597 | 0.484 | 1.59779E-17 |
| Contractile | KMT5A | 8.21929E-11 | 0.325804 | 0.28 | 0.202 | 1.60071E-06 |
| Contractile | MIGA1 | 8.10614E-16 | 0.325707 | 0.28 | 0.179 | 1.57867E-11 |
| Terminal | OLFML3 | 2.78746E-29 | 0.325659 | 0.505 | 0.357 | 5.42857E-25 |
| VEGFA_Terminal | SEC11A | 2.21119E-11 | 0.325526 | 0.695 | 0.543 | 4.3063E-07 |
| Fibroblasts | CDC42EP4 | 3.43848E-05 | 0.325467 | 0.142 | 0.072 | 0.669643535 |
| IFN_Intermediate | ENTPD4 | 0.004606924 | 0.325318 | 0.196 | 0.113 | 1 |
| Fibroblasts | CDC14B | 5.63436E-05 | 0.325316 | 0.188 | 0.109 | 1 |
| Contractile | ZNF644 | 4.97384E-14 | 0.325204 | 0.301 | 0.202 | 9.68655E-10 |
| Fibroblasts | EP400 | 0.000335011 | 0.32514 | 0.193 | 0.12 | 1 |
| CD74_Intermediat | LUM | 1.12645E-69 | 0.325121 | 0.999 | 0.906 | 2.19376E-65 |
| VEGFA_Terminal | ARHGAP18 | 1.82267E-52 | 0.324882 | 0.138 | 0.017 | 3.54966E-48 |
| CD74_Intermediat | GABARAPL | 6.66456E-28 | 0.32483 | 0.542 | 0.403 | 1.29792E-23 |
| CD74_Intermediat | NID2 | 1.71183E-20 | 0.290357 | 0.283 | 0.182 | 3.33379E-16 |
| Fibroblasts | ENSSSCG0( | 0.001191234 | 0.324603 | 0.22 | 0.153 | 1 |
| Intermediate | FAM107B | 7.86998E-09 | 0.324519 | 0.235 | 0.151 | 0.000153268 |
| IFN_Intermediate | ENSSSCG0( | 0.000923444 | 0.324255 | 0.121 | 0.051 | 1 |
| Contractile | TADA3 | 3.74741E-22 | 0.324198 | 0.223 | 0.115 | 7.29808E-18 |
| Fibroblasts | VAMP2 | 0.002386271 | 0.324156 | 0.436 | 0.391 | 1 |
| Fibroblasts | NISCH | 0.003891968 | 0.32412 | 0.17 | 0.115 | 1 |
| Contractile | MANBAL.1 | 3.25024E-15 | 0.323884 | 0.23 | 0.141 | 6.32984E-11 |
| IFN_Intermediate | ENSSSCG0( | 0.000570576 | 0.323862 | 0.308 | 0.184 | 1 |
| Terminal | ACKR3 | 2.48551E-44 | 0.323814 | 0.186 | 0.069 | 4.84053E-40 |
| Fibroblasts | ENSSSCG0( | 0.001599089 | 0.323814 | 0.133 | 0.079 | 1 |
| Terminal | YBX3 | 1.10107E-36 | 0.323765 | 0.912 | 0.858 | 2.14433E-32 |
| Contractile | SGCD | 8.85566E-16 | 0.323656 | 0.328 | 0.223 | 1.72464E-11 |
| Intermediate | ENSSSCG0( | 2.99341E-14 | 0.323647 | 0.273 | 0.158 | 5.82967E-10 |
| Intermediate | RBPMS | 3.98315E-15 | 0.323397 | 0.522 | 0.362 | 7.75719E-11 |
| Contractile | CTDSPL | 1.93789E-14 | 0.323267 | 0.26 | 0.167 | 3.77404E-10 |
| Contractile | LRRC1 | 3.15073E-51 | 0.323173 | 0.122 | 0.022 | 6.13605E-47 |
| Terminal | ROPN1L | 2.04983E-34 | 0.323144 | 0.202 | 0.09 | 3.99204E-30 |
| Fibroblasts | PRRC2C | 0.002451661 | 0.323134 | 0.734 | 0.744 | 1 |
| VEGFA_Terminal | RTN4 | 1.17197E-10 | 0.322985 | 0.867 | 0.772 | 2.28241E-06 |
| Contractile | RAN | 1.08271E-17 | 0.322968 | 0.709 | 0.643 | 2.10857E-13 |
| IFN_Intermediate | DPT | 0.007336165 | 0.32271 | 0.486 | 0.383 | 1 |
| Contractile | ENSSSCG0( | 7.51485E-27 | 0.322696 | 0.18 | 0.075 | 1.46352E-22 |
| Pericytes | MYL6 | 0.002910929 | 0.322685 | 0.94 | 0.97 | 1 |
| VEGFA_Terminal | CASP3 | 2.65938E-10 | 0.322662 | 0.344 | 0.208 | 5.17914E-06 |
| IFN_Intermediate | GM2A | 4.59021E-05 | 0.32265 | 0.252 | 0.123 | 0.893942568 |
| VEGFA_Terminal | ENSSSCG0( | 2.74828E-26 | 0.322645 | 0.14 | 0.033 | 5.35227E-22 |
| VEGFA_Terminal | MED31 | 1.10188E-11 | 0.32258 | 0.516 | 0.343 | 2.14591E-07 |
| VEGFA_Terminal | CIAO2B | 2.7939E-13 | 0.322454 | 0.448 | 0.267 | 5.44112E-09 |
| Contractile | FNTA | 3.85327E-13 | 0.322388 | 0.361 | 0.269 | 7.50424E-09 |
| Contractile | PREX2 | 4.30084E-30 | 0.32228 | 0.159 | 0.057 | 8.3759E-26 |
| Fibroblasts | CEP170 | 0.004364154 | 0.322254 | 0.339 | 0.278 | 1 |
| VEGFA_Terminal | SAMD4A | 9.47511E-09 | 0.322182 | 0.637 | 0.478 | 0.000184528 |
| Contractile | AFF4 | 1.04187E-12 | 0.32212 | 0.417 | 0.321 | 2.02903E-08 |
| Contractile | ENSSSCG0( | 1.95668E-40 | 0.321866 | 0.138 | 0.036 | 3.81064E-36 |
| IFN_Intermediate | SERPING1 | 0.001036905 | 0.321859 | 0.869 | 0.821 | 1 |
| Intermediate | RSU1 | 3.09029E-08 | 0.321709 | 0.605 | 0.524 | 0.000601833 |

| | | | | | | |
|---|---|---|---|---|---|---|
| VEGFA_Terminal | ARFGAP3 | 2.04326E-11 | 0.321415 | 0.692 | 0.515 | 3.97924E-07 |
| Intermediate | BTG2 | 0.004621664 | 0.321385 | 0.392 | 0.35 | 1 |
| VEGFA_Terminal | ENSSSCG0( | 1.1001E-11 | 0.321347 | 0.385 | 0.228 | 2.14245E-07 |
| IFN_Intermediate | S100A1 | 0.006716179 | 0.321345 | 0.243 | 0.147 | 1 |
| Fibroblasts | CCDC170 | 5.27521E-06 | 0.321255 | 0.11 | 0.046 | 0.102734764 |
| Contractile | ACTR3 | 4.86605E-17 | 0.321247 | 0.651 | 0.554 | 9.47664E-13 |
| IFN_Intermediate | TDRD7 | 6.99438E-06 | 0.320984 | 0.234 | 0.1 | 0.136215496 |
| VEGFA_Terminal | ENSSSCG0( | 4.15557E-08 | 0.320983 | 0.571 | 0.426 | 0.000809297 |
| Contractile | PPCDC | 5.34271E-54 | 0.320794 | 0.119 | 0.02 | 1.04049E-49 |
| Contractile | CPEB4 | 3.52415E-14 | 0.320745 | 0.367 | 0.271 | 6.86328E-10 |
| Terminal | OSTC | 5.53224E-23 | 0.320685 | 0.476 | 0.357 | 1.0774E-18 |
| Fibroblasts | KLHL24 | 0.006662762 | 0.320379 | 0.284 | 0.227 | 1 |
| VEGFA_Terminal | TNFAIP3 | 2.01972E-15 | 0.320369 | 0.165 | 0.061 | 3.93341E-11 |
| Fibroblasts | ENSSSCG0( | 8.87274E-07 | 0.320295 | 0.138 | 0.059 | 0.017279657 |
| Intermediate | ID3 | 3.24365E-15 | 0.320172 | 0.899 | 0.813 | 6.31701E-11 |
| CD74_Intermediat | ANPEP | 4.54605E-36 | 0.320072 | 0.645 | 0.469 | 8.85344E-32 |
| Intermediate | MATN4 | 7.33093E-13 | 0.319959 | 0.166 | 0.082 | 1.4277E-08 |
| Terminal | SPPL2A | 7.1975E-24 | 0.319592 | 0.556 | 0.425 | 1.40171E-19 |
| Fibroblasts | DPY19L3 | 0.000146679 | 0.319573 | 0.151 | 0.083 | 1 |
| VEGFA_Terminal | ADAM12 | 2.65972E-17 | 0.319571 | 0.235 | 0.099 | 5.17981E-13 |
| Contractile | EIF1AX | 3.68997E-15 | 0.319472 | 0.669 | 0.586 | 7.18621E-11 |
| Terminal | ITGA11 | 4.50341E-32 | 0.31935 | 0.284 | 0.156 | 8.77038E-28 |
| VEGFA_Terminal | FSCN1 | 1.49282E-28 | 0.319282 | 0.199 | 0.057 | 2.90726E-24 |
| Intermediate | CCN5 | 9.53374E-08 | 0.319002 | 0.53 | 0.447 | 0.001856695 |
| Fibroblasts | GALNT11 | 0.002042687 | 0.318883 | 0.261 | 0.198 | 1 |
| Intermediate | DUSP10 | 2.84531E-15 | 0.318835 | 0.121 | 0.048 | 5.54124E-11 |
| Contractile | HUWE1 | 4.88713E-11 | 0.318708 | 0.394 | 0.307 | 9.51768E-07 |
| Fibroblasts | GIT2 | 0.004770252 | 0.318522 | 0.183 | 0.128 | 1 |
| Fibroblasts | TMBIM6 | 8.975E-05 | 0.31844 | 0.775 | 0.753 | 1 |
| IFN_Intermediate | SLA-DMB | 3.83647E-10 | 0.318336 | 0.131 | 0.028 | 7.47153E-06 |
| Terminal | ENSSSCG0( | 1.06438E-31 | 0.318301 | 0.398 | 0.247 | 2.07288E-27 |
| VEGFA_Terminal | CHMP5 | 3.8569E-12 | 0.318279 | 0.588 | 0.397 | 7.51132E-08 |
| CD74_Intermediat | FTL | 3.41114E-67 | 0.318097 | 0.998 | 0.992 | 6.6432E-63 |
| Intermediate | BOK | 5.75249E-13 | 0.318024 | 0.382 | 0.258 | 1.1203E-08 |
| Fibroblasts | ACVRL1 | 3.22868E-07 | 0.317763 | 0.151 | 0.065 | 0.006287846 |
| VEGFA_Terminal | TSPAN6 | 7.22235E-11 | 0.31774 | 0.576 | 0.401 | 1.40655E-06 |
| Terminal | BHLHE40 | 2.90215E-37 | 0.317553 | 0.698 | 0.519 | 5.65193E-33 |
| Contractile | MSX1 | 2.82062E-40 | 0.317518 | 0.115 | 0.026 | 5.49316E-36 |
| Contractile | NCS1 | 2.19768E-24 | 0.31743 | 0.157 | 0.065 | 4.27997E-20 |
| Fibroblasts | LSP1 | 3.63315E-10 | 0.317274 | 0.128 | 0.041 | 7.07556E-06 |
| Contractile | STK40 | 1.10971E-19 | 0.317225 | 0.261 | 0.151 | 2.16115E-15 |
| Contractile | EEF2 | 7.74382E-28 | 0.317132 | 0.958 | 0.911 | 1.50811E-23 |
| VEGFA_Terminal | RNF13 | 2.25404E-12 | 0.317124 | 0.501 | 0.32 | 4.38975E-08 |
| VEGFA_Terminal | RNFT1 | 2.09802E-12 | 0.317114 | 0.3 | 0.165 | 4.08589E-08 |
| CD74_Intermediat | LGALS3 | 1.54408E-53 | 0.317095 | 0.939 | 0.727 | 3.00709E-49 |
| Contractile | PBX2 | 4.14717E-16 | 0.31707 | 0.193 | 0.107 | 8.0766E-12 |
| Intermediate | DMD | 2.13213E-16 | 0.316943 | 0.338 | 0.197 | 4.15233E-12 |
| IFN_Intermediate | REEP5 | 0.00023788 | 0.316935 | 0.832 | 0.748 | 1 |
| Fibroblasts | SPSB1 | 5.5813E-05 | 0.316917 | 0.124 | 0.06 | 1 |
| Terminal | ANKH | 9.08188E-13 | 0.316893 | 0.513 | 0.429 | 1.7687E-08 |
| Fibroblasts | ENSSSCG0( | 0.000372504 | 0.316743 | 0.679 | 0.66 | 1 |
| Terminal | TPI1 | 5.49437E-32 | 0.316723 | 0.819 | 0.709 | 1.07003E-27 |
| VEGFA_Terminal | FAM114A1 | 2.44793E-09 | 0.31662 | 0.671 | 0.526 | 4.76734E-05 |
| Contractile | SWAP70 | 1.92184E-13 | 0.316618 | 0.475 | 0.379 | 3.74279E-09 |
| Fibroblasts | ZNF662 | 0.003390199 | 0.316529 | 0.161 | 0.104 | 1 |
| Contractile | HDAC9 | 5.55618E-23 | 0.316528 | 0.23 | 0.116 | 1.08207E-18 |
| Intermediate | CXXC5 | 5.89439E-08 | 0.316504 | 0.435 | 0.35 | 0.001147933 |
| Contractile | ADAMTS19 | 6.4179E-108 | 0.31636 | 0.104 | 0.002 | 1.2499E-103 |
| CD74_Intermediat | VPS13C | 1.40676E-20 | 0.31635 | 0.547 | 0.434 | 2.73966E-16 |

| | | | | | | |
|---|---|---|---|---|---|---|
| Intermediate | STX12 | 1.56253E-09 | 0.316126 | 0.398 | 0.298 | 3.04303E-05 |
| VEGFA_Terminal | GNG5 | 5.74117E-13 | 0.316035 | 0.516 | 0.329 | 1.11809E-08 |
| VEGFA_Terminal | EIF2S3B | 1.14079E-11 | 0.315839 | 0.656 | 0.479 | 2.22169E-07 |
| VEGFA_Terminal | KDELR1 | 2.1558E-12 | 0.31568 | 0.792 | 0.633 | 4.19842E-08 |
| Fibroblasts | EXOC6B | 0.003506761 | 0.3156 | 0.216 | 0.156 | 1 |
| Terminal | SCARA3 | 5.53096E-28 | 0.315596 | 0.564 | 0.411 | 1.07715E-23 |
| IFN_Intermediate | TRIM56 | 0.000186851 | 0.315378 | 0.178 | 0.078 | 1 |
| Contractile | PPM1K | 3.9376E-16 | 0.31537 | 0.216 | 0.123 | 7.66848E-12 |
| IFN_Intermediate | TRIM34 | 5.52087E-25 | 0.315198 | 0.112 | 0.009 | 1.07519E-20 |
| CD74_Intermediat | MGST1 | 2.58996E-67 | 0.51877 | 0.391 | 0.184 | 5.04395E-63 |
| Terminal | COX5A | 5.84811E-25 | 0.314932 | 0.603 | 0.48 | 1.13892E-20 |
| VEGFA_Terminal | SLC5A3 | 5.4934E-20 | 0.314896 | 0.109 | 0.027 | 1.06984E-15 |
| VEGFA_Terminal | TMEM100 | 6.09205E-12 | 0.314882 | 0.259 | 0.133 | 1.18643E-07 |
| Contractile | EML2 | 9.81965E-33 | 0.314869 | 0.173 | 0.062 | 1.91238E-28 |
| Contractile | DRAP1 | 1.32097E-17 | 0.314832 | 0.653 | 0.565 | 2.5726E-13 |
| Contractile | AK4 | 3.24918E-14 | 0.314658 | 0.264 | 0.172 | 6.32777E-10 |
| VEGFA_Terminal | BTG3 | 1.55322E-11 | 0.314631 | 0.404 | 0.247 | 3.0249E-07 |
| Contractile | PACSIN3 | 2.62487E-14 | 0.314518 | 0.306 | 0.208 | 5.11193E-10 |
| Contractile | STX12 | 1.36651E-15 | 0.314495 | 0.402 | 0.291 | 2.66128E-11 |
| Contractile | UTP11 | 5.02602E-19 | 0.314395 | 0.282 | 0.17 | 9.78817E-15 |
| Terminal | TSPAN2 | 3.84463E-15 | 0.314206 | 0.187 | 0.116 | 7.48741E-11 |
| Fibroblasts | IQSEC1 | 0.001142837 | 0.314169 | 0.211 | 0.143 | 1 |
| Contractile | DCAF7 | 4.93503E-13 | 0.314109 | 0.302 | 0.212 | 9.61098E-09 |
| Intermediate | BCAM | 5.13888E-12 | 0.314024 | 0.252 | 0.151 | 1.0008E-07 |
| Contractile | NRGN | 2.89865E-51 | 0.313984 | 0.123 | 0.023 | 5.64511E-47 |
| Fibroblasts | ENSSSCG0( | 4.38217E-10 | 0.313872 | 0.119 | 0.037 | 8.53427E-06 |
| Contractile | FGL2 | 1.30061E-45 | 0.313717 | 0.919 | 0.659 | 2.53295E-41 |
| Contractile | ARID5B | 3.12724E-12 | 0.313696 | 0.508 | 0.418 | 6.0903E-08 |
| Contractile | U2AF1 | 1.60651E-17 | 0.313602 | 0.62 | 0.506 | 3.12868E-13 |
| VEGFA_Terminal | TTLL3 | 2.49709E-11 | 0.313585 | 0.772 | 0.624 | 4.86309E-07 |
| VEGFA_Terminal | SRPX2 | 4.25623E-28 | 0.313565 | 0.254 | 0.085 | 8.28901E-24 |
| Contractile | CAV1 | 1.00344E-14 | 0.313495 | 0.242 | 0.149 | 1.95419E-10 |
| Fibroblasts | NAALADL2 | 3.57362E-05 | 0.313437 | 0.165 | 0.088 | 0.695963016 |
| Contractile | ENSSSCG0( | 2.95603E-19 | 0.313407 | 0.767 | 0.695 | 5.75687E-15 |
| Contractile | TBL1XR1 | 1.95769E-16 | 0.313396 | 0.667 | 0.561 | 3.8126E-12 |
| Terminal | TNFRSF11I | 3.57157E-09 | 0.313295 | 0.312 | 0.245 | 6.95563E-05 |
| Contractile | HRH1 | 1.81521E-43 | 0.313218 | 0.126 | 0.028 | 3.53512E-39 |
| Pericytes | BEX1 | 0.003614456 | 0.313075 | 0.18 | 0.072 | 1 |
| VEGFA_Terminal | SMIM1 | 1.36532E-19 | 0.312925 | 0.138 | 0.04 | 2.65896E-15 |
| CD74_Intermediat | PSAP | 4.6526E-39 | 0.312913 | 0.96 | 0.903 | 9.06094E-35 |
| Contractile | ARHGAP5 | 2.11765E-12 | 0.312781 | 0.55 | 0.454 | 4.12413E-08 |
| Fibroblasts | ENSSSCG0( | 0.00044374 | 0.312701 | 0.142 | 0.081 | 1 |
| Contractile | ENSSSCG0( | 2.18672E-15 | 0.312559 | 0.464 | 0.348 | 4.25864E-11 |
| Contractile | HSP90AB1 | 2.24656E-24 | 0.312557 | 0.9 | 0.841 | 4.37517E-20 |
| Pericytes | PGM2 | 0.00601718 | 0.312411 | 0.14 | 0.054 | 1 |
| Contractile | MRPS33 | 2.64626E-11 | 0.312281 | 0.417 | 0.331 | 5.15358E-07 |
| Fibroblasts | SYNE3 | 5.45027E-05 | 0.312276 | 0.119 | 0.057 | 1 |
| CD74_Intermediat | SAMD12 | 1.29505E-19 | 0.28307 | 0.287 | 0.184 | 2.52211E-15 |
| IFN_Intermediate | ENSSSCG0( | 0.002413652 | 0.312109 | 0.439 | 0.306 | 1 |
| CD74_Intermediat | CRISPLD2 | 1.31009E-34 | 0.265045 | 0.337 | 0.19 | 2.5514E-30 |
| Contractile | RNF146 | 7.2377E-17 | 0.312014 | 0.354 | 0.241 | 1.40954E-12 |
| Fibroblasts | NPEPPS | 0.009144803 | 0.311933 | 0.317 | 0.269 | 1 |
| VEGFA_Terminal | NARS1 | 2.46536E-07 | 0.311736 | 0.576 | 0.44 | 0.004801285 |
| Fibroblasts | NOL9 | 4.46325E-05 | 0.31171 | 0.156 | 0.083 | 0.869218336 |
| Contractile | RAD21 | 4.84522E-10 | 0.311625 | 0.438 | 0.364 | 9.43606E-06 |
| Terminal | METRNL | 5.6514E-30 | 0.311558 | 0.399 | 0.257 | 1.10061E-25 |
| Pericytes | SOX5 | 0.004647141 | 0.311335 | 0.1 | 0.031 | 1 |
| IFN_Intermediate | IL15 | 0.000261822 | 0.31127 | 0.103 | 0.036 | 1 |
| Fibroblasts | LYSMD3 | 0.008441045 | 0.311083 | 0.275 | 0.225 | 1 |

| | | | | | | |
|---|---|---|---|---|---|---|
| Contractile | CHCHD2 | 3.14024E-17 | 0.311031 | 0.722 | 0.632 | 6.11562E-13 |
| Contractile | MRPL33 | 1.15131E-13 | 0.310944 | 0.473 | 0.377 | 2.24218E-09 |
| Contractile | CHD6 | 2.13003E-16 | 0.310936 | 0.305 | 0.197 | 4.14823E-12 |
| Contractile | MYL12A | 5.92932E-21 | 0.310676 | 0.898 | 0.859 | 1.15474E-16 |
| Intermediate | ITGA7 | 1.60621E-13 | 0.310663 | 0.275 | 0.16 | 3.12809E-09 |
| Contractile | ENSSSCG0( | 2.63391E-24 | 0.310549 | 0.888 | 0.849 | 5.12954E-20 |
| VEGFA_Terminal | HILPDA | 2.83653E-15 | 0.31018 | 0.264 | 0.123 | 5.52414E-11 |
| Contractile | CCDC149 | 3.13291E-17 | 0.310057 | 0.2 | 0.108 | 6.10134E-13 |
| VEGFA_Terminal | NPC2 | 6.89724E-11 | 0.309889 | 0.806 | 0.665 | 1.34324E-06 |
| Fibroblasts | JUP | 0.004452798 | 0.309863 | 0.307 | 0.25 | 1 |
| VEGFA_Terminal | PLOD3 | 1.08588E-17 | 0.309788 | 0.332 | 0.158 | 2.11475E-13 |
| Fibroblasts | PDZD8 | 0.002140929 | 0.309614 | 0.197 | 0.134 | 1 |
| Intermediate | RCN1 | 6.92717E-11 | 0.309611 | 0.558 | 0.442 | 1.34907E-06 |
| Intermediate | TUBA1B | 4.59806E-13 | 0.309608 | 0.791 | 0.703 | 8.95473E-09 |
| IFN_Intermediate | PHLDA1 | 0.000632912 | 0.309459 | 0.28 | 0.158 | 1 |
| Fibroblasts | CDC27 | 0.006939803 | 0.309391 | 0.188 | 0.136 | 1 |
| Fibroblasts | GPC4 | 1.12755E-09 | 0.309231 | 0.11 | 0.033 | 2.1959E-05 |
| Contractile | NSA2 | 1.39105E-16 | 0.309224 | 0.78 | 0.702 | 2.70908E-12 |
| Contractile | CYLD | 6.24375E-17 | 0.309201 | 0.279 | 0.173 | 1.21597E-12 |
| Pericytes | ENSSSCG0( | 0.008741174 | 0.30917 | 0.12 | 0.044 | 1 |
| VEGFA_Terminal | FYN | 6.15148E-16 | 0.309118 | 0.298 | 0.143 | 1.198E-11 |
| Fibroblasts | ENSSSCG0( | 0.000473592 | 0.308868 | 0.138 | 0.077 | 1 |
| Fibroblasts | MCMBP | 0.006874095 | 0.308737 | 0.206 | 0.155 | 1 |
| Contractile | ARMCX3 | 1.61605E-13 | 0.308676 | 0.307 | 0.211 | 3.14726E-09 |
| Fibroblasts | TFDP2 | 0.004134927 | 0.308669 | 0.28 | 0.22 | 1 |
| Fibroblasts | GLI2 | 0.00016632 | 0.308638 | 0.119 | 0.06 | 1 |
| Contractile | TST | 1.43553E-13 | 0.308186 | 0.372 | 0.27 | 2.7957E-09 |
| Fibroblasts | TSPYL2 | 0.000665275 | 0.308166 | 0.193 | 0.124 | 1 |
| Contractile | GNB5 | 5.50744E-17 | 0.308152 | 0.262 | 0.16 | 1.07257E-12 |
| VEGFA_Terminal | IKBIP | 7.47416E-11 | 0.308064 | 0.663 | 0.496 | 1.45559E-06 |
| Contractile | COX17 | 1.25214E-14 | 0.308047 | 0.465 | 0.359 | 2.43854E-10 |
| Fibroblasts | CPXM1 | 9.12871E-13 | 0.307912 | 0.101 | 0.023 | 1.77782E-08 |
| VEGFA_Terminal | SLC39A13 | 1.28026E-15 | 0.307866 | 0.247 | 0.112 | 2.49331E-11 |
| Intermediate | ESAM | 2.53926E-24 | 0.30783 | 0.126 | 0.037 | 4.94521E-20 |
| VEGFA_Terminal | CDC42EP5 | 1.61197E-10 | 0.30766 | 0.545 | 0.377 | 3.13932E-06 |
| Contractile | TAX1BP1 | 6.0881E-20 | 0.307518 | 0.774 | 0.707 | 1.18566E-15 |
| Intermediate | PLXDC2 | 1.42737E-06 | 0.307472 | 0.768 | 0.739 | 0.027797981 |
| CD74_Intermediat | HEYL | 5.40011E-34 | 0.417628 | 0.337 | 0.195 | 1.05167E-29 |
| Contractile | KLHL2 | 1.86287E-23 | 0.307325 | 0.165 | 0.071 | 3.62793E-19 |
| Contractile | WBP2 | 1.79144E-15 | 0.30732 | 0.576 | 0.478 | 3.48884E-11 |
| VEGFA_Terminal | AMDHD2 | 3.38944E-11 | 0.307312 | 0.712 | 0.529 | 6.60093E-07 |
| Contractile | CRBN | 9.38396E-13 | 0.307222 | 0.315 | 0.225 | 1.82753E-08 |
| Contractile | KLHL21 | 4.0136E-21 | 0.307173 | 0.172 | 0.079 | 7.81649E-17 |
| Fibroblasts | TOP3A | 0.000627256 | 0.307154 | 0.11 | 0.057 | 1 |
| Contractile | CCT4 | 1.24036E-12 | 0.307128 | 0.495 | 0.411 | 2.4156E-08 |
| Terminal | MFAP2 | 3.97769E-23 | 0.30703 | 0.52 | 0.386 | 7.74655E-19 |
| Contractile | ZNF358 | 1.54397E-20 | 0.307019 | 0.237 | 0.129 | 3.00688E-16 |
| VEGFA_Terminal | FJX1 | 5.37402E-16 | 0.306958 | 0.295 | 0.142 | 1.04659E-11 |
| Terminal | RCN3 | 5.58196E-21 | 0.306908 | 0.551 | 0.438 | 1.08709E-16 |
| IFN_Intermediate | DNAJC13 | 0.000954681 | 0.306598 | 0.299 | 0.178 | 1 |
| Terminal | TNFAIP8L3 | 1.44496E-67 | 0.306592 | 0.145 | 0.028 | 2.81406E-63 |
| VEGFA_Terminal | EMD | 2.54108E-12 | 0.306341 | 0.598 | 0.403 | 4.94875E-08 |
| Contractile | CYP27A1 | 4.31213E-20 | 0.306338 | 0.203 | 0.1 | 8.39787E-16 |
| Fibroblasts | ENSSSCG0( | 0.000193484 | 0.306029 | 0.119 | 0.06 | 1 |
| VEGFA_Terminal | PPP4R1 | 4.07207E-16 | 0.30601 | 0.24 | 0.106 | 7.93035E-12 |
| Intermediate | RIMKLB | 5.87663E-12 | 0.306007 | 0.249 | 0.149 | 1.14447E-07 |
| Contractile | ACTG1 | 2.82075E-53 | 0.305999 | 0.999 | 0.983 | 5.49341E-49 |
| VEGFA_Terminal | GNAS | 8.43717E-20 | 0.305902 | 0.99 | 0.979 | 1.64314E-15 |
| Pericytes | ENSSSCG0( | 3.74529E-17 | 0.30586 | 0.1 | 0.006 | 7.29395E-13 |

| | | | | | | |
|---|---|---|---|---|---|---|
| CD74_Intermediat | NR4A1 | 3.85403E-16 | 0.315124 | 0.286 | 0.196 | 7.50572E-12 |
| Terminal | ST8SIA1 | 1.61071E-55 | 0.305764 | 0.152 | 0.04 | 3.13685E-51 |
| VEGFA_Terminal | PDE1C | 1.35745E-40 | 0.305757 | 0.119 | 0.016 | 2.64363E-36 |
| Pericytes | HEY1 | 0.002536589 | 0.305732 | 0.1 | 0.029 | 1 |
| Contractile | ANKRD13A | 6.22082E-12 | 0.305663 | 0.37 | 0.283 | 1.2115E-07 |
| Intermediate | STX6 | 1.32137E-08 | 0.305641 | 0.349 | 0.256 | 0.000257337 |
| Fibroblasts | OGT | 0.007286779 | 0.305636 | 0.225 | 0.168 | 1 |
| VEGFA_Terminal | PPP2R3A | 2.00263E-13 | 0.305629 | 0.315 | 0.168 | 3.90013E-09 |
| Contractile | DDX42 | 4.34959E-11 | 0.305606 | 0.39 | 0.306 | 8.47083E-07 |
| CD74_Intermediat | FAM111A | 7.46711E-31 | 0.305543 | 0.674 | 0.506 | 1.45422E-26 |
| Contractile | ATF7IP | 4.33121E-14 | 0.305299 | 0.345 | 0.24 | 8.43504E-10 |
| Contractile | PCGF5 | 6.05024E-19 | 0.305293 | 0.674 | 0.546 | 1.17828E-14 |
| Terminal | SRPX2 | 1.98535E-41 | 0.305194 | 0.169 | 0.062 | 3.86647E-37 |
| Intermediate | TEAD3 | 6.49921E-15 | 0.305088 | 0.233 | 0.124 | 1.26572E-10 |
| Terminal | FAM114A1 | 3.74068E-23 | 0.304924 | 0.617 | 0.497 | 7.28497E-19 |
| VEGFA_Terminal | TFG | 2.0954E-09 | 0.304856 | 0.622 | 0.463 | 4.08079E-05 |
| VEGFA_Terminal | KDELR2 | 4.0611E-11 | 0.304819 | 0.649 | 0.469 | 7.90899E-07 |
| Fibroblasts | CRY2 | 0.008784284 | 0.304748 | 0.133 | 0.089 | 1 |
| VEGFA_Terminal | CHMP4B | 2.94115E-06 | 0.304625 | 0.702 | 0.603 | 0.057278983 |
| Fibroblasts | HNRNPU | 0.007858172 | 0.304623 | 0.619 | 0.637 | 1 |
| Fibroblasts | TSPAN9 | 1.52742E-07 | 0.304615 | 0.133 | 0.053 | 0.002974657 |
| IFN_Intermediate | CCDC80 | 0.009357828 | 0.30455 | 0.925 | 0.849 | 1 |
| Fibroblasts | FSTL1 | 0.00053118 | 0.304544 | 0.89 | 0.847 | 1 |
| VEGFA_Terminal | MDFIC | 2.55792E-13 | 0.304332 | 0.334 | 0.182 | 4.98154E-09 |
| VEGFA_Terminal | GPX8 | 2.28616E-10 | 0.304275 | 0.567 | 0.392 | 4.45229E-06 |
| IFN_Intermediate | CGNL1 | 0.000193043 | 0.304063 | 0.467 | 0.283 | 1 |
| Pericytes | ZCCHC3 | 0.000366044 | 0.304051 | 0.1 | 0.024 | 1 |
| Contractile | CNOT7 | 2.43885E-14 | 0.304037 | 0.309 | 0.211 | 4.74965E-10 |
| Contractile | USP8 | 8.6235E-12 | 0.304007 | 0.456 | 0.367 | 1.67943E-07 |
| Contractile | MEIS2 | 5.97631E-18 | 0.303923 | 0.288 | 0.175 | 1.16389E-13 |
| Contractile | NEDD9 | 3.63927E-11 | 0.303897 | 0.312 | 0.225 | 7.08748E-07 |
| Contractile | SYT11 | 1.2144E-13 | 0.303855 | 0.524 | 0.434 | 2.36505E-09 |
| Contractile | EIF4G2 | 1.45415E-25 | 0.303386 | 0.928 | 0.882 | 2.83196E-21 |
| Intermediate | ENSSSCG00 | 2.05358E-09 | 0.3031 | 0.793 | 0.755 | 3.99935E-05 |
| Intermediate | SEPTIN9 | 1.36584E-08 | 0.302977 | 0.449 | 0.36 | 0.000265997 |
| Contractile | RBBP7 | 7.02993E-14 | 0.302757 | 0.408 | 0.307 | 1.36908E-09 |
| VEGFA_Terminal | H2AC6 | 1.26573E-07 | 0.302723 | 0.179 | 0.098 | 0.002465006 |
| VEGFA_Terminal | ENSSSCG00 | 0.000543263 | 0.302619 | 0.455 | 0.354 | 1 |
| Contractile | DCAF5 | 4.8047E-14 | 0.302539 | 0.321 | 0.224 | 9.35715E-10 |
| VEGFA_Terminal | UGDH | 2.86483E-07 | 0.302343 | 0.387 | 0.271 | 0.005579247 |
| Contractile | RABGAP1 | 9.094E-18 | 0.302315 | 0.222 | 0.124 | 1.77106E-13 |
| Intermediate | PDLIM3 | 8.0477E-10 | 0.302266 | 0.415 | 0.3 | 1.56729E-05 |
| Contractile | HNRNPUL2 | 3.88662E-12 | 0.302145 | 0.477 | 0.391 | 7.56919E-08 |
| Fibroblasts | RETREG1 | 0.003981357 | 0.302141 | 0.197 | 0.139 | 1 |
| Contractile | TESK1 | 3.2577E-15 | 0.301948 | 0.216 | 0.127 | 6.34437E-11 |
| Terminal | VEGFC | 2.79246E-33 | 0.301791 | 0.16 | 0.065 | 5.43831E-29 |
| Terminal | HHIPL1 | 1.37854E-38 | 0.301743 | 0.217 | 0.096 | 2.68471E-34 |
| VEGFA_Terminal | EMC6 | 2.10801E-11 | 0.301318 | 0.659 | 0.472 | 4.10535E-07 |
| Terminal | CA5B | 9.38374E-35 | 0.301316 | 0.197 | 0.088 | 1.82748E-30 |
| CD74_Intermediat | ENSSSCG00 | 5.52452E-21 | 0.301179 | 0.519 | 0.399 | 1.0759E-16 |
| IFN_Intermediate | NCBP1 | 0.003913962 | 0.301064 | 0.168 | 0.09 | 1 |
| Contractile | COPS2 | 1.08773E-13 | 0.300961 | 0.584 | 0.494 | 2.11836E-09 |
| Terminal | MXRA8 | 7.89901E-15 | 0.300836 | 0.712 | 0.652 | 1.53833E-10 |
| IFN_Intermediate | ALDH2 | 0.004650461 | 0.300809 | 0.664 | 0.549 | 1 |
| Fibroblasts | SNRK | 0.00022616 | 0.300738 | 0.206 | 0.13 | 1 |
| VEGFA_Terminal | UBA52 | 1.08089E-13 | 0.300685 | 0.971 | 0.948 | 2.10503E-09 |
| Intermediate | PEA15 | 2.27084E-06 | 0.300512 | 0.478 | 0.41 | 0.044224676 |
| Contractile | ARPP19 | 8.82227E-13 | 0.3003 | 0.584 | 0.504 | 1.71814E-08 |
| VEGFA_Terminal | RGS2 | 1.85153E-06 | 0.300017 | 0.184 | 0.107 | 0.036058606 |

| | | | | | | |
|---|---|---|---|---|---|---|
| VEGFA_Terminal | TAX1BP3 | 5.58929E-10 | 0.299998 | 0.453 | 0.295 | 1.08852E-05 |
| Contractile | RND3 | 5.66549E-08 | 0.299966 | 0.316 | 0.246 | 0.001103354 |
| Intermediate | RARRES1 | 2.94416E-13 | 0.299888 | 0.811 | 0.723 | 5.73375E-09 |
| Intermediate | PPP1R14A | 1.67104E-56 | 0.299831 | 0.651 | 0.29 | 3.25434E-52 |
| IFN_Intermediate | DST | 0.008972116 | 0.299808 | 0.72 | 0.63 | 1 |
| Intermediate | TNFAIP3 | 1.25061E-10 | 0.299726 | 0.127 | 0.061 | 2.43555E-06 |
| Intermediate | TMEM70 | 1.20097E-07 | 0.299631 | 0.346 | 0.262 | 0.002338893 |
| Contractile | NEMF | 5.07162E-13 | 0.299597 | 0.36 | 0.265 | 9.87698E-09 |
| CD74_Intermediat | HMCN1 | 1.51727E-25 | 0.299591 | 0.464 | 0.322 | 2.95488E-21 |
| Contractile | ITIH5 | 3.22097E-23 | 0.299502 | 0.286 | 0.15 | 6.27283E-19 |
| Contractile | ARMC1 | 1.63168E-12 | 0.299445 | 0.299 | 0.211 | 3.17769E-08 |
| IFN_Intermediate | CTSH | 0.000157215 | 0.299424 | 0.916 | 0.782 | 1 |
| CD74_Intermediat | CD34 | 6.57945E-55 | 0.449728 | 0.396 | 0.203 | 1.28135E-50 |
| Fibroblasts | CERS4 | 0.000399049 | 0.299353 | 0.11 | 0.056 | 1 |
| Pericytes | DUSP16 | 0.005987789 | 0.299322 | 0.14 | 0.054 | 1 |
| Pericytes | ENSSSCG0( | 0.002996662 | 0.299201 | 0.12 | 0.039 | 1 |
| IFN_Intermediate | UNC45B | 2.92127E-28 | 0.299038 | 0.112 | 0.008 | 5.68917E-24 |
| Intermediate | PEAR1 | 6.55364E-18 | 0.29902 | 0.189 | 0.084 | 1.27632E-13 |
| IFN_Intermediate | IRAK1BP1 | 0.000587859 | 0.29896 | 0.234 | 0.125 | 1 |
| Contractile | PPP2R1A | 7.56985E-13 | 0.298837 | 0.364 | 0.268 | 1.47423E-08 |
| Contractile | TMEM43 | 4.70534E-11 | 0.298812 | 0.468 | 0.38 | 9.16364E-07 |
| Fibroblasts | VPS36 | 0.002522178 | 0.298754 | 0.482 | 0.447 | 1 |
| Intermediate | CPQ | 1.89392E-09 | 0.29872 | 0.45 | 0.347 | 3.68842E-05 |
| CD74_Intermediat | FRMD6 | 3.58979E-30 | 0.375942 | 0.332 | 0.203 | 6.99111E-26 |
| Terminal | TSKU | 1.3956E-22 | 0.298709 | 0.378 | 0.255 | 2.71793E-18 |
| Fibroblasts | ACOX1 | 0.001497071 | 0.298549 | 0.344 | 0.276 | 1 |
| Fibroblasts | IGDCC4 | 3.06007E-13 | 0.298497 | 0.101 | 0.023 | 5.95949E-09 |
| Contractile | SRC | 2.7386E-28 | 0.298423 | 0.139 | 0.048 | 5.33343E-24 |
| Contractile | PRKAG2 | 4.88684E-18 | 0.298343 | 0.223 | 0.124 | 9.51712E-14 |
| VEGFA_Terminal | MOB2 | 1.17341E-11 | 0.298281 | 0.438 | 0.275 | 2.28522E-07 |
| Terminal | UCHL1 | 4.16388E-29 | 0.298245 | 0.126 | 0.048 | 8.10916E-25 |
| Contractile | GTF2I | 2.01183E-11 | 0.298242 | 0.34 | 0.254 | 3.91804E-07 |
| Fibroblasts | AKAP9 | 0.006382599 | 0.297962 | 0.661 | 0.669 | 1 |
| VEGFA_Terminal | MKNK2 | 2.39533E-09 | 0.297921 | 0.242 | 0.134 | 4.66491E-05 |
| Intermediate | LMO4 | 2.55421E-08 | 0.297826 | 0.842 | 0.828 | 0.000497433 |
| Terminal | TGFBI | 4.28754E-18 | 0.29781 | 0.422 | 0.317 | 8.34999E-14 |
| Contractile | SEM1 | 4.70121E-20 | 0.297742 | 0.812 | 0.759 | 9.15561E-16 |
| Intermediate | TLN1 | 2.90674E-09 | 0.297613 | 0.676 | 0.612 | 5.66087E-05 |
| Contractile | FYTTD1 | 3.63878E-11 | 0.297547 | 0.447 | 0.364 | 7.08653E-07 |
| Contractile | ENSSSCG0( | 1.83705E-20 | 0.297523 | 0.81 | 0.739 | 3.57765E-16 |
| VEGFA_Terminal | RAB5IF | 1.67087E-10 | 0.297511 | 0.62 | 0.454 | 3.25402E-06 |
| Contractile | SPAG8 | 9.33846E-11 | 0.297338 | 0.419 | 0.331 | 1.81867E-06 |
| VEGFA_Terminal | TMEM258 | 7.27701E-09 | 0.297271 | 0.748 | 0.603 | 0.00014172 |
| Fibroblasts | TGS1 | 0.00145839 | 0.296882 | 0.289 | 0.216 | 1 |
| VEGFA_Terminal | LAMTOR5 | 6.28049E-10 | 0.296875 | 0.707 | 0.551 | 1.22313E-05 |
| Contractile | PKP4 | 1.16973E-22 | 0.296845 | 0.196 | 0.093 | 2.27805E-18 |
| Intermediate | ADARB1 | 1.91105E-08 | 0.296684 | 0.389 | 0.294 | 0.000372177 |
| Contractile | EMC2 | 6.91948E-11 | 0.296667 | 0.391 | 0.312 | 1.34757E-06 |
| Fibroblasts | BPTF | 0.00658129 | 0.296584 | 0.477 | 0.444 | 1 |
| Pericytes | LOC11025! | 1.2299E-121 | 0.296584 | 0.12 | 0 | 2.3952E-117 |
| Contractile | TJP1 | 4.57642E-12 | 0.296519 | 0.491 | 0.403 | 8.91259E-08 |
| CD74_Intermediat | ENSSSCG0( | 8.71773E-17 | 0.296481 | 0.438 | 0.34 | 1.69778E-12 |
| Contractile | KLF6 | 1.37294E-08 | 0.296406 | 0.664 | 0.609 | 0.000267381 |
| CD74_Intermediat | SULF2 | 4.67349E-37 | 0.381687 | 0.368 | 0.213 | 9.10161E-33 |
| VEGFA_Terminal | ENSSSCG0( | 1.32688E-15 | 0.29623 | 0.281 | 0.133 | 2.58411E-11 |
| Contractile | NMT1 | 3.1678E-12 | 0.296225 | 0.377 | 0.286 | 6.1693E-08 |
| Pericytes | TRABD2B | 0.000689434 | 0.296185 | 0.1 | 0.025 | 1 |
| VEGFA_Terminal | IGF2R | 1.85561E-08 | 0.296171 | 0.358 | 0.233 | 0.00036138 |
| IFN_Intermediate | PNPT1 | 0.000500664 | 0.295819 | 0.131 | 0.054 | 1 |

| | | | | | | |
|---|---|---|---|---|---|---|
| VEGFA_Terminal | CANX | 8.24602E-08 | 0.295795 | 0.857 | 0.754 | 0.001605912 |
| Contractile | DNAJC21 | 8.6457E-14 | 0.295751 | 0.291 | 0.197 | 1.68375E-09 |
| CD74_Intermediat | RBMS3 | 3.63639E-19 | 0.295733 | 0.411 | 0.3 | 7.08187E-15 |
| Contractile | ZNF385D | 1.71319E-10 | 0.295611 | 0.237 | 0.161 | 3.33644E-06 |
| CD74_Intermediat | XG | 3.07632E-48 | 0.453283 | 0.391 | 0.217 | 5.99114E-44 |
| CD74_Intermediat | GADD45A | 1.77196E-15 | 0.295466 | 0.399 | 0.3 | 3.45089E-11 |
| Terminal | P4HB | 8.92843E-23 | 0.295296 | 0.548 | 0.418 | 1.73881E-18 |
| CD74_Intermediat | LMO7 | 6.16338E-24 | 0.295515 | 0.339 | 0.218 | 1.20032E-19 |
| IFN_Intermediate | ENSSSCG0( | 0.000752635 | 0.295273 | 0.131 | 0.056 | 1 |
| Contractile | NAA50 | 2.84759E-13 | 0.295252 | 0.379 | 0.281 | 5.54567E-09 |
| Terminal | ADAMTS6 | 2.35789E-48 | 0.2952 | 0.118 | 0.026 | 4.59199E-44 |
| Fibroblasts | BEX1 | 3.68153E-06 | 0.294934 | 0.151 | 0.07 | 0.071697787 |
| Fibroblasts | OGA | 0.005469385 | 0.29493 | 0.289 | 0.232 | 1 |
| VEGFA_Terminal | TMEM176 | 9.58311E-13 | 0.29492 | 0.312 | 0.169 | 1.86631E-08 |
| CD74_Intermediat | LOXL1 | 5.35419E-23 | 0.294894 | 0.647 | 0.515 | 1.04273E-18 |
| Contractile | DDX6 | 8.25957E-13 | 0.29484 | 0.412 | 0.316 | 1.60855E-08 |
| CD74_Intermediat | FOSB | 4.48729E-31 | 0.740437 | 0.352 | 0.22 | 8.73899E-27 |
| Fibroblasts | WDFY3 | 0.00609943 | 0.294706 | 0.206 | 0.151 | 1 |
| CD74_Intermediat | CPQ | 9.94525E-24 | 0.294703 | 0.449 | 0.322 | 1.93684E-19 |
| Fibroblasts | LFNG | 0.002219436 | 0.29467 | 0.101 | 0.055 | 1 |
| Intermediate | S100A11 | 7.59197E-15 | 0.294638 | 0.951 | 0.924 | 1.47854E-10 |
| Contractile | ZBTB38 | 4.62036E-12 | 0.294576 | 0.579 | 0.5 | 8.99815E-08 |
| CD74_Intermediat | PSME1 | 5.11025E-21 | 0.294567 | 0.634 | 0.519 | 9.95221E-17 |
| VEGFA_Terminal | NDUFB5 | 7.19293E-09 | 0.294521 | 0.596 | 0.453 | 0.000140082 |
| Contractile | SNX27 | 2.36016E-18 | 0.294458 | 0.254 | 0.147 | 4.59642E-14 |
| Contractile | CDC5L | 4.40441E-13 | 0.294415 | 0.535 | 0.438 | 8.57759E-09 |
| Contractile | NUP88 | 6.65062E-17 | 0.294398 | 0.197 | 0.107 | 1.29521E-12 |
| Contractile | ZC3H13 | 1.09194E-13 | 0.294318 | 0.4 | 0.299 | 2.12655E-09 |
| Contractile | ABHD16A | 1.70501E-14 | 0.294281 | 0.25 | 0.158 | 3.32051E-10 |
| CD74_Intermediat | STAT1 | 1.7554E-17 | 0.269913 | 0.324 | 0.223 | 3.41865E-13 |
| Contractile | HYPK | 2.16564E-16 | 0.294182 | 0.649 | 0.547 | 4.21759E-12 |
| Terminal | YKT6 | 7.54364E-20 | 0.294029 | 0.424 | 0.312 | 1.46912E-15 |
| Contractile | ENSSSCG0( | 2.5379E-31 | 0.293809 | 0.13 | 0.04 | 4.94256E-27 |
| Pericytes | ENSSSCG0( | 1.25862E-30 | 0.29373 | 0.1 | 0.003 | 2.45115E-26 |
| Contractile | ALAD | 3.92165E-27 | 0.293725 | 0.173 | 0.07 | 7.63742E-23 |
| Fibroblasts | IFI44L | 0.006189555 | 0.293707 | 0.239 | 0.182 | 1 |
| Fibroblasts | CTSC | 0.003016975 | 0.293697 | 0.394 | 0.331 | 1 |
| Fibroblasts | ELMO2 | 0.000203889 | 0.293669 | 0.128 | 0.067 | 1 |
| IFN_Intermediate | ENSSSCG0( | 0.001732242 | 0.293604 | 0.187 | 0.097 | 1 |
| Terminal | GPX8 | 2.11201E-24 | 0.293516 | 0.495 | 0.36 | 4.11314E-20 |
| Fibroblasts | SLC35E2B | 7.57738E-06 | 0.29351 | 0.161 | 0.08 | 0.147569543 |
| Contractile | ACADL | 2.14507E-15 | 0.293508 | 0.322 | 0.216 | 4.17752E-11 |
| Intermediate | SFRP1 | 2.30082E-07 | 0.293503 | 0.842 | 0.818 | 0.004480839 |
| VEGFA_Terminal | ESD | 1.83279E-09 | 0.293424 | 0.688 | 0.516 | 3.56936E-05 |
| Contractile | TPR | 5.2968E-14 | 0.293347 | 0.629 | 0.547 | 1.03155E-09 |
| IFN_Intermediate | KYNU | 9.31043E-08 | 0.29314 | 0.178 | 0.055 | 0.001813207 |
| VEGFA_Terminal | LDLRAD4 | 7.43089E-11 | 0.293106 | 0.252 | 0.134 | 1.44717E-06 |
| Terminal | MEDAG | 1.97215E-31 | 0.292942 | 0.248 | 0.128 | 3.84076E-27 |
| Contractile | C1ORF35 | 6.8988E-12 | 0.292902 | 0.435 | 0.341 | 1.34354E-07 |
| Fibroblasts | C1GALT1 | 0.005963801 | 0.292754 | 0.211 | 0.155 | 1 |
| Terminal | NDUFC2 | 2.50291E-23 | 0.292746 | 0.715 | 0.616 | 4.87442E-19 |
| CD74_Intermediat | TMSB4X | 4.50788E-55 | 0.29274 | 0.999 | 0.999 | 8.7791E-51 |
| Intermediate | GPC6 | 3.81866E-09 | 0.292727 | 0.224 | 0.141 | 7.43683E-05 |
| Fibroblasts | ST3GAL4 | 0.003713187 | 0.292594 | 0.211 | 0.151 | 1 |
| Intermediate | APOE | 1.01105E-17 | 0.292534 | 0.498 | 0.323 | 1.96903E-13 |
| Contractile | HNRNPR | 1.94777E-11 | 0.292503 | 0.54 | 0.465 | 3.79329E-07 |
| Terminal | C9H1ORF2 | 6.69902E-25 | 0.29236 | 0.525 | 0.387 | 1.30463E-20 |
| Intermediate | KANK1 | 3.0495E-13 | 0.292352 | 0.535 | 0.392 | 5.93889E-09 |
| Intermediate | MPRIP | 6.82406E-09 | 0.292313 | 0.307 | 0.216 | 0.000132899 |

| | | | | | | |
|---|---|---|---|---|---|---|
| Fibroblasts | C6ORF62 | 0.009803602 | 0.292254 | 0.39 | 0.347 | 1 |
| Fibroblasts | PABPC1 | 0.002354245 | 0.292147 | 0.711 | 0.766 | 1 |
| VEGFA_Terminal | ERI1 | 5.25362E-23 | 0.292069 | 0.138 | 0.036 | 1.02314E-18 |
| Terminal | TFPI2 | 2.84769E-16 | 0.292005 | 0.859 | 0.792 | 5.54587E-12 |
| Contractile | AKT1S1 | 8.75674E-13 | 0.29193 | 0.254 | 0.169 | 1.70537E-08 |
| Contractile | PPP4R3B | 6.21936E-12 | 0.291892 | 0.456 | 0.361 | 1.21122E-07 |
| Contractile | COL4A6 | 7.73039E-23 | 0.291861 | 0.195 | 0.091 | 1.50549E-18 |
| Intermediate | DST | 1.00442E-06 | 0.291852 | 0.677 | 0.626 | 0.019561138 |
| CD74_Intermediat | ZFP36 | 6.19508E-17 | 0.291741 | 0.5 | 0.396 | 1.20649E-12 |
| Terminal | P3H4 | 3.04845E-20 | 0.291715 | 0.363 | 0.252 | 5.93685E-16 |
| VEGFA_Terminal | ATP6V0E1 | 1.80007E-09 | 0.29168 | 0.763 | 0.617 | 3.50563E-05 |
| CD74_Intermediat | SCARA3 | 1.13062E-16 | 0.291658 | 0.536 | 0.43 | 2.20189E-12 |
| VEGFA_Terminal | XPC | 2.57634E-08 | 0.291617 | 0.293 | 0.179 | 0.000501742 |
| Intermediate | EPAS1 | 4.50254E-10 | 0.29153 | 0.659 | 0.564 | 8.7687E-06 |
| VEGFA_Terminal | PTPRB | 2.4241E-16 | 0.291435 | 0.228 | 0.097 | 4.72093E-12 |
| Contractile | TFEB | 2.82758E-19 | 0.291418 | 0.191 | 0.096 | 5.50671E-15 |
| Contractile | NDUFV1 | 1.67701E-12 | 0.291394 | 0.37 | 0.278 | 3.26598E-08 |
| Contractile | MARF1 | 1.44096E-14 | 0.29132 | 0.286 | 0.188 | 2.80627E-10 |
| Intermediate | CSRP1 | 3.16462E-34 | 0.291273 | 0.886 | 0.756 | 6.1631E-30 |
| Contractile | AHSA1 | 9.70982E-14 | 0.291214 | 0.381 | 0.282 | 1.89099E-09 |
| Contractile | ENSSSCG00 | 1.77228E-21 | 0.291211 | 0.89 | 0.859 | 3.45152E-17 |
| Contractile | AEBP2 | 4.70784E-11 | 0.291177 | 0.293 | 0.21 | 9.16852E-07 |
| Fibroblasts | APH1B | 0.00604606 | 0.291077 | 0.161 | 0.11 | 1 |
| Pericytes | CACNA1H | 0.000599354 | 0.291042 | 0.1 | 0.024 | 1 |
| Fibroblasts | LAMP2 | 0.001277665 | 0.291033 | 0.734 | 0.732 | 1 |
| CD74_Intermediat | CMKLR1 | 3.26229E-18 | 0.296337 | 0.329 | 0.225 | 6.35332E-14 |
| Fibroblasts | MAPK12 | 0.002184977 | 0.290891 | 0.165 | 0.108 | 1 |
| Contractile | CHCHD10 | 2.19461E-12 | 0.290713 | 0.686 | 0.607 | 4.274E-08 |
| Intermediate | ENSSSCG00 | 5.4245E-07 | 0.290696 | 0.541 | 0.472 | 0.010564211 |
| Fibroblasts | ENSSSCG00 | 1.92641E-05 | 0.290584 | 0.11 | 0.048 | 0.375168918 |
| Contractile | ENSSSCG00 | 1.89103E-30 | 0.290532 | 0.979 | 0.976 | 3.68277E-26 |
| Intermediate | CREB3L2 | 2.33838E-05 | 0.29048 | 0.545 | 0.501 | 0.455398882 |
| VEGFA_Terminal | COPE | 2.53635E-09 | 0.290476 | 0.768 | 0.634 | 4.93955E-05 |
| Fibroblasts | F2R | 0.001203799 | 0.290427 | 0.128 | 0.073 | 1 |
| Contractile | CACNB3 | 1.52804E-20 | 0.290378 | 0.156 | 0.07 | 2.97586E-16 |
| CD74_Intermediat | APOE | 9.011E-213 | 1.712587 | 0.617 | 0.232 | 1.7549E-208 |
| Fibroblasts | C3ORF70 | 7.79319E-07 | 0.290351 | 0.138 | 0.058 | 0.015177243 |
| CD74_Intermediat | CLDND1 | 1.70841E-18 | 0.290183 | 0.478 | 0.37 | 3.32712E-14 |
| Fibroblasts | CALCRL | 5.89581E-05 | 0.290166 | 0.115 | 0.054 | 1 |
| Contractile | NAPEPLD | 1.29076E-12 | 0.290127 | 0.22 | 0.138 | 2.51375E-08 |
| Intermediate | ENSSSCG00 | 5.82679E-07 | 0.290103 | 0.39 | 0.303 | 0.011347669 |
| VEGFA_Terminal | ASNS | 1.09638E-10 | 0.290094 | 0.196 | 0.096 | 2.13521E-06 |
| Contractile | TAF11 | 2.8236E-12 | 0.290044 | 0.325 | 0.236 | 5.49897E-08 |
| CD74_Intermediat | TNFAIP8 | 3.09707E-20 | 0.290891 | 0.341 | 0.232 | 6.03154E-16 |
| Intermediate | PNRC2 | 1.89914E-08 | 0.289867 | 0.677 | 0.595 | 0.000369858 |
| VEGFA_Terminal | PDE8A | 1.35883E-10 | 0.289781 | 0.215 | 0.11 | 2.64632E-06 |
| Contractile | MRPL11 | 9.18018E-14 | 0.289724 | 0.291 | 0.197 | 1.78784E-09 |
| Contractile | EPB41 | 7.11055E-25 | 0.289652 | 0.138 | 0.051 | 1.38478E-20 |
| Intermediate | LOXL1 | 5.57166E-09 | 0.289618 | 0.63 | 0.544 | 0.000108508 |
| CD74_Intermediat | HGF | 1.17939E-18 | 0.272373 | 0.341 | 0.234 | 2.29687E-14 |
| Contractile | NAP1L1 | 1.41397E-20 | 0.289316 | 0.909 | 0.881 | 2.7537E-16 |
| VEGFA_Terminal | UBE2D3 | 5.76436E-09 | 0.28922 | 0.668 | 0.528 | 0.000112261 |
| Contractile | ENSSSCG00 | 3.02735E-28 | 0.289096 | 0.163 | 0.062 | 5.89577E-24 |
| Intermediate | MICAL3 | 4.83278E-10 | 0.289058 | 0.238 | 0.147 | 9.41183E-06 |
| Contractile | PTP4A3 | 8.94988E-34 | 0.289058 | 0.108 | 0.027 | 1.74299E-29 |
| IFN_Intermediate | ELAVL1 | 0.0001569 | 0.289048 | 0.477 | 0.305 | 1 |
| Fibroblasts | ENPP1 | 0.000749846 | 0.288968 | 0.298 | 0.217 | 1 |
| VEGFA_Terminal | SCAMP1 | 3.99932E-10 | 0.288911 | 0.414 | 0.259 | 7.78868E-06 |
| Terminal | TXNDC5 | 1.08648E-16 | 0.288858 | 0.574 | 0.467 | 2.11593E-12 |

| | | | | | | |
|---|---|---|---|---|---|---|
| Terminal | SLC40A1 | 6.81041E-22 | 0.288833 | 0.26 | 0.157 | 1.32633E-17 |
| Contractile | ZBTB37 | 3.37466E-16 | 0.288808 | 0.178 | 0.093 | 6.57216E-12 |
| CD74_Intermediat | ENG | 5.22198E-24 | 0.288674 | 0.473 | 0.34 | 1.01698E-19 |
| Contractile | HECTD1 | 1.92441E-11 | 0.288657 | 0.56 | 0.482 | 3.74778E-07 |
| Terminal | FAT4 | 2.31201E-15 | 0.288653 | 0.475 | 0.378 | 4.50264E-11 |
| VEGFA_Terminal | HTRA1 | 9.49851E-10 | 0.288633 | 0.935 | 0.893 | 1.84983E-05 |
| Intermediate | PDE5A | 1.14462E-08 | 0.288603 | 0.31 | 0.217 | 0.000222915 |
| Contractile | AFF1 | 1.36786E-08 | 0.288603 | 0.347 | 0.276 | 0.000266391 |
| Intermediate | LOX | 3.49933E-07 | 0.288575 | 0.843 | 0.818 | 0.006814954 |
| CD74_Intermediat | VCAM1 | 6.04852E-71 | 0.537256 | 0.461 | 0.235 | 1.17795E-66 |
| CD74_Intermediat | CYBRD1 | 2.08606E-28 | 0.288448 | 0.631 | 0.491 | 4.0626E-24 |
| Contractile | LOC10015: | 1.29421E-11 | 0.288418 | 0.486 | 0.409 | 2.52047E-07 |
| IFN_Intermediate | PALMD | 0.001436342 | 0.288274 | 0.477 | 0.343 | 1 |
| Fibroblasts | SCAMP2 | 2.22295E-05 | 0.288083 | 0.174 | 0.095 | 0.432919274 |
| Contractile | ABL1 | 5.10211E-13 | 0.287974 | 0.28 | 0.188 | 9.93637E-09 |
| VEGFA_Terminal | MAFG | 2.37929E-11 | 0.287892 | 0.228 | 0.114 | 4.63368E-07 |
| Terminal | BASP1 | 3.00349E-23 | 0.287826 | 0.395 | 0.261 | 5.84929E-19 |
| Terminal | CSRP2 | 8.20447E-17 | 0.287781 | 0.509 | 0.408 | 1.59782E-12 |
| Contractile | ENSSSCG0( | 4.10779E-13 | 0.287778 | 0.226 | 0.144 | 7.99992E-09 |
| Intermediate | LMOD1 | 1.28775E-32 | 0.287759 | 0.677 | 0.426 | 2.50789E-28 |
| Fibroblasts | CEP135 | 0.00134384 | 0.287648 | 0.101 | 0.053 | 1 |
| Fibroblasts | ENSSSCG0( | 0.00979621 | 0.287613 | 0.124 | 0.08 | 1 |
| Contractile | ENSSSCG0( | 3.55325E-11 | 0.287519 | 0.406 | 0.32 | 6.91996E-07 |
| Contractile | ENSSSCG0( | 1.4485E-11 | 0.287498 | 0.355 | 0.263 | 2.82096E-07 |
| IFN_Intermediate | CSF1 | 0.006566843 | 0.287353 | 0.393 | 0.285 | 1 |
| Contractile | CTDSP1 | 5.03051E-12 | 0.287337 | 0.331 | 0.242 | 9.79692E-08 |
| Terminal | XYLT1 | 2.17604E-22 | 0.287273 | 0.248 | 0.148 | 4.23784E-18 |
| Intermediate | IGFBP4 | 1.7109E-11 | 0.287031 | 0.161 | 0.084 | 3.33198E-07 |
| VEGFA_Terminal | CHCHD5 | 5.10872E-11 | 0.286975 | 0.305 | 0.172 | 9.94924E-07 |
| Contractile | RAB22A | 5.74358E-10 | 0.286948 | 0.28 | 0.205 | 1.11856E-05 |
| Contractile | CHMP4B | 3.80564E-13 | 0.286928 | 0.658 | 0.601 | 7.41148E-09 |
| VEGFA_Terminal | SLC2A1 | 1.74063E-39 | 0.286907 | 0.131 | 0.02 | 3.38988E-35 |
| Contractile | ENSSSCG0( | 1.30898E-30 | 0.286815 | 0.135 | 0.043 | 2.54924E-26 |
| IFN_Intermediate | ENSSSCG0( | 0.000642703 | 0.286804 | 1 | 0.997 | 1 |
| CD74_Intermediat | PHYH | 4.63532E-19 | 0.286778 | 0.582 | 0.477 | 9.02729E-15 |
| Contractile | ACO2 | 2.1334E-11 | 0.286676 | 0.305 | 0.22 | 4.15479E-07 |
| CD74_Intermediat | LAMC1 | 1.58172E-26 | 0.286651 | 0.756 | 0.636 | 3.0804E-22 |
| VEGFA_Terminal | PTS | 6.36308E-10 | 0.286072 | 0.76 | 0.618 | 1.23921E-05 |
| Terminal | TRIM47 | 1.1591E-18 | 0.28596 | 0.482 | 0.376 | 2.25734E-14 |
| Contractile | RAD23A | 4.35929E-12 | 0.285852 | 0.299 | 0.211 | 8.48973E-08 |
| Terminal | SPON2 | 2.53554E-19 | 0.285685 | 0.308 | 0.201 | 4.93796E-15 |
| Contractile | DCTN4 | 2.42296E-12 | 0.285534 | 0.29 | 0.201 | 4.71872E-08 |
| CD74_Intermediat | VPS36 | 9.40996E-22 | 0.285447 | 0.537 | 0.412 | 1.83259E-17 |
| Terminal | FOXS1 | 5.61964E-20 | 0.285245 | 0.511 | 0.394 | 1.09442E-15 |
| VEGFA_Terminal | SRP72 | 8.31193E-10 | 0.285222 | 0.426 | 0.277 | 1.61875E-05 |
| VEGFA_Terminal | TAF1D | 1.06424E-08 | 0.285213 | 0.673 | 0.507 | 0.000207262 |
| IFN_Intermediate | APLP2 | 0.006263211 | 0.285191 | 0.738 | 0.602 | 1 |
| VEGFA_Terminal | KLF6 | 6.94881E-09 | 0.285188 | 0.743 | 0.609 | 0.000135328 |
| VEGFA_Terminal | BNIP3L | 2.79734E-09 | 0.285154 | 0.63 | 0.468 | 5.44782E-05 |
| Contractile | UBE2M | 7.32336E-09 | 0.285082 | 0.5 | 0.444 | 0.000142622 |
| Contractile | LRP5 | 7.69057E-20 | 0.2847 | 0.167 | 0.078 | 1.49774E-15 |
| Terminal | PLP2 | 1.67293E-25 | 0.284662 | 0.803 | 0.719 | 3.25803E-21 |
| Contractile | SSBP2 | 5.66173E-18 | 0.284573 | 0.201 | 0.107 | 1.10262E-13 |
| Contractile | SIN3B | 1.12402E-15 | 0.284559 | 0.173 | 0.091 | 2.18902E-11 |
| Fibroblasts | TPBG | 3.84138E-06 | 0.28455 | 0.124 | 0.053 | 0.07481081 |
| CD74_Intermediat | EGR1 | 1.24027E-19 | 0.284373 | 0.559 | 0.434 | 2.41543E-15 |
| CD74_Intermediat | ZBTB20 | 1.29611E-20 | 0.284353 | 0.497 | 0.376 | 2.52417E-16 |
| CD74_Intermediat | ANTXR1 | 2.38792E-25 | 0.284352 | 0.759 | 0.633 | 4.65047E-21 |
| Fibroblasts | DPY19L4 | 0.002322641 | 0.284276 | 0.133 | 0.081 | 1 |

| | | | | | | |
|---|---|---|---|---|---|---|
| Intermediate | CUX1 | 2.22078E-05 | 0.284276 | 0.602 | 0.562 | 0.432496099 |
| VEGFA_Terminal | ENSSSCG0( | 1.38258E-22 | 0.284275 | 0.102 | 0.021 | 2.69257E-18 |
| Fibroblasts | RGL1 | 7.88675E-07 | 0.284215 | 0.142 | 0.061 | 0.015359453 |
| Terminal | MRC2 | 1.04692E-23 | 0.284185 | 0.533 | 0.394 | 2.03888E-19 |
| VEGFA_Terminal | BACH1 | 4.26314E-10 | 0.284172 | 0.438 | 0.278 | 8.30246E-06 |
| VEGFA_Terminal | ZNRF2 | 1.82951E-18 | 0.284161 | 0.155 | 0.051 | 3.56296E-14 |
| Pericytes | MARK1 | 2.1245E-10 | 0.284124 | 0.12 | 0.013 | 4.13747E-06 |
| Fibroblasts | RNF144A | 3.97058E-09 | 0.28409 | 0.115 | 0.037 | 7.7327E-05 |
| VEGFA_Terminal | PFDN2 | 1.99452E-09 | 0.283958 | 0.673 | 0.507 | 3.88432E-05 |
| VEGFA_Terminal | ZDHHC12 | 3.32326E-15 | 0.283664 | 0.247 | 0.112 | 6.47205E-11 |
| Intermediate | FZD7 | 1.07874E-08 | 0.283659 | 0.253 | 0.166 | 0.000210085 |
| Pericytes | COX4I1 | 0.008561489 | 0.283643 | 0.96 | 0.957 | 1 |
| Fibroblasts | NR2C1 | 3.11227E-05 | 0.283532 | 0.133 | 0.065 | 0.606114148 |
| CD74_Intermediat | SDF4 | 1.46239E-28 | 0.283404 | 0.814 | 0.702 | 2.848E-24 |
| VEGFA_Terminal | RGCC | 7.3376E-06 | 0.283401 | 0.373 | 0.269 | 0.142899822 |
| IFN_Intermediate | FZD6 | 0.000248711 | 0.283248 | 0.196 | 0.092 | 1 |
| Contractile | HDGFL3 | 4.02748E-11 | 0.28312 | 0.398 | 0.317 | 7.84352E-07 |
| Terminal | P4HA3 | 3.99872E-46 | 0.283073 | 0.133 | 0.036 | 7.78751E-42 |
| Contractile | HNRNPUL1 | 6.31171E-13 | 0.283071 | 0.412 | 0.313 | 1.22921E-08 |
| CD74_Intermediat | PLAT | 3.23996E-60 | 0.31205 | 0.449 | 0.236 | 6.30982E-56 |
| IFN_Intermediate | IPO9 | 0.000839084 | 0.282984 | 0.215 | 0.112 | 1 |
| Fibroblasts | ENSSSCG0( | 0.001183856 | 0.282832 | 0.353 | 0.281 | 1 |
| IFN_Intermediate | NDFIP1 | 0.003232685 | 0.282715 | 0.757 | 0.692 | 1 |
| VEGFA_Terminal | TRIB1 | 8.52721E-16 | 0.282623 | 0.191 | 0.076 | 1.66067E-11 |
| IFN_Intermediate | ANXA1 | 0.003984161 | 0.282548 | 0.832 | 0.744 | 1 |
| CD74_Intermediat | BTG2 | 5.28514E-09 | 0.282381 | 0.405 | 0.334 | 0.000102928 |
| VEGFA_Terminal | HYPK | 1.87034E-08 | 0.282356 | 0.69 | 0.554 | 0.000364249 |
| Contractile | UBE4B | 1.01017E-13 | 0.282347 | 0.19 | 0.109 | 1.96731E-09 |
| Contractile | BCKDHB | 1.171E-21 | 0.282252 | 0.154 | 0.066 | 2.28052E-17 |
| Fibroblasts | PAICS | 0.001528533 | 0.282011 | 0.174 | 0.114 | 1 |
| VEGFA_Terminal | VAT1 | 9.45295E-05 | 0.282007 | 0.467 | 0.38 | 1 |
| Terminal | TMEM208 | 8.5819E-21 | 0.28187 | 0.472 | 0.357 | 1.67133E-16 |
| Terminal | CHPF | 4.8769E-20 | 0.281814 | 0.454 | 0.339 | 9.49776E-16 |
| VEGFA_Terminal | COL12A1 | 4.6739E-08 | 0.281651 | 0.501 | 0.358 | 0.000910242 |
| Terminal | SSR3 | 6.61407E-25 | 0.281647 | 0.68 | 0.557 | 1.28809E-20 |
| VEGFA_Terminal | CEBPG | 1.56419E-09 | 0.281608 | 0.24 | 0.13 | 3.04627E-05 |
| CD74_Intermediat | CGNL1 | 1.09887E-45 | 0.435186 | 0.412 | 0.236 | 2.14006E-41 |
| Terminal | SLC29A1 | 1.69678E-28 | 0.2816 | 0.192 | 0.093 | 3.30448E-24 |
| Intermediate | ACTN4 | 3.37957E-05 | 0.281496 | 0.525 | 0.473 | 0.658171377 |
| Contractile | NR2F6 | 3.4394E-11 | 0.281473 | 0.233 | 0.156 | 6.69823E-07 |
| CD74_Intermediat | LAMC3 | 4.42715E-67 | 0.637001 | 0.452 | 0.239 | 8.62187E-63 |
| VEGFA_Terminal | BOLA2 | 1.00444E-09 | 0.281353 | 0.639 | 0.472 | 1.95615E-05 |
| VEGFA_Terminal | IL13RA1 | 1.60354E-09 | 0.2813 | 0.477 | 0.324 | 3.12289E-05 |
| Contractile | PLAAT1 | 4.44325E-53 | 0.280945 | 0.121 | 0.021 | 8.65323E-49 |
| Terminal | MYOF | 4.95678E-16 | 0.280863 | 0.67 | 0.566 | 9.65333E-12 |
| Terminal | TCEA3 | 1.28416E-19 | 0.280806 | 0.511 | 0.396 | 2.5009E-15 |
| VEGFA_Terminal | ATXN3 | 2.33255E-07 | 0.280803 | 0.242 | 0.145 | 0.004542641 |
| Contractile | PREPL | 4.9963E-11 | 0.280737 | 0.293 | 0.21 | 9.7303E-07 |
| Contractile | EHBP1L1 | 3.58399E-11 | 0.280691 | 0.303 | 0.219 | 6.97983E-07 |
| CD74_Intermediat | CTSV | 6.70994E-31 | 0.280551 | 0.907 | 0.828 | 1.30676E-26 |
| Contractile | POLDIP3 | 9.25266E-11 | 0.280481 | 0.317 | 0.236 | 1.80196E-06 |
| IFN_Intermediate | HEG1 | 0.000304696 | 0.280459 | 0.458 | 0.291 | 1 |
| VEGFA_Terminal | ENSSSCG0( | 2.20507E-17 | 0.280455 | 0.99 | 0.989 | 4.29436E-13 |
| Terminal | SERPINF1 | 7.41596E-28 | 0.280422 | 0.895 | 0.701 | 1.44426E-23 |
| VEGFA_Terminal | ENSSSCG0( | 9.04114E-07 | 0.280416 | 0.632 | 0.495 | 0.017607623 |
| Fibroblasts | GGCX | 0.004965832 | 0.280291 | 0.211 | 0.155 | 1 |
| Intermediate | PHLDB2 | 2.9393E-09 | 0.280255 | 0.363 | 0.256 | 5.72428E-05 |
| Fibroblasts | COMMD1( | 0.002757321 | 0.280182 | 0.202 | 0.14 | 1 |
| VEGFA_Terminal | DBP | 4.0297E-11 | 0.280124 | 0.249 | 0.131 | 7.84784E-07 |

| | | | | | | |
|---|---|---|---|---|---|---|
| Pericytes | PWP2 | 0.001341337 | 0.279981 | 0.12 | 0.035 | 1 |
| Contractile | SNRNP200 | 5.022E-10 | 0.279947 | 0.365 | 0.288 | 9.78035E-06 |
| CD74_Intermediat | ADGRD1 | 1.01013E-32 | 0.312187 | 0.392 | 0.239 | 1.96723E-28 |
| Contractile | NDUFB7 | 6.48919E-15 | 0.279912 | 0.619 | 0.538 | 1.26377E-10 |
| Terminal | CDK2AP1 | 1.39945E-20 | 0.279872 | 0.441 | 0.326 | 2.72542E-16 |
| Intermediate | SMAD9 | 4.06152E-06 | 0.27984 | 0.464 | 0.399 | 0.079098147 |
| Fibroblasts | FAM8A1 | 0.003126703 | 0.279839 | 0.261 | 0.197 | 1 |
| Fibroblasts | KIAA0408 | 6.21009E-05 | 0.279687 | 0.128 | 0.063 | 1 |
| Contractile | BLOC1S6 | 3.97709E-13 | 0.27954 | 0.352 | 0.258 | 7.74538E-09 |
| Contractile | BDP1 | 1.53021E-09 | 0.279516 | 0.334 | 0.256 | 2.98009E-05 |
| VEGFA_Terminal | ACVR1 | 2.54638E-10 | 0.279344 | 0.329 | 0.195 | 4.95908E-06 |
| Contractile | PNRC2 | 6.1418E-16 | 0.279326 | 0.684 | 0.589 | 1.19612E-11 |
| VEGFA_Terminal | FAU | 2.01227E-15 | 0.279285 | 0.995 | 0.992 | 3.91889E-11 |
| Contractile | EHD1 | 1.84961E-11 | 0.279115 | 0.251 | 0.17 | 3.60212E-07 |
| VEGFA_Terminal | ENSSSCG0( | 3.74734E-16 | 0.279103 | 0.167 | 0.061 | 7.29795E-12 |
| Contractile | PLEKHJ1 | 6.89237E-10 | 0.278984 | 0.319 | 0.246 | 1.34229E-05 |
| CD74_Intermediat | ENSSSCG0( | 1.13386E-53 | 0.278977 | 0.999 | 0.997 | 2.20819E-49 |
| Contractile | PSMC3 | 4.43849E-11 | 0.278963 | 0.655 | 0.584 | 8.64396E-07 |
| Terminal | LTBP3 | 7.77487E-22 | 0.278926 | 0.784 | 0.686 | 1.51416E-17 |
| VEGFA_Terminal | CAMK1 | 2.15805E-11 | 0.278904 | 0.288 | 0.157 | 4.20281E-07 |
| Fibroblasts | ZDHHC9 | 0.000447739 | 0.278781 | 0.142 | 0.082 | 1 |
| CD74_Intermediat | TFPI2 | 1.54408E-30 | 0.278716 | 0.883 | 0.786 | 3.0071E-26 |
| Fibroblasts | CANX | 0.002329691 | 0.278687 | 0.766 | 0.761 | 1 |
| Contractile | SLC25A28 | 3.86679E-14 | 0.278608 | 0.223 | 0.134 | 7.53058E-10 |
| Terminal | SOD1 | 2.32647E-22 | 0.278514 | 0.798 | 0.726 | 4.53079E-18 |
| VEGFA_Terminal | SELENOF | 1.93406E-10 | 0.278514 | 0.787 | 0.649 | 3.76658E-06 |
| Contractile | HNRNPU | 2.25358E-13 | 0.278482 | 0.686 | 0.627 | 4.38884E-09 |
| VEGFA_Terminal | BET1 | 7.40675E-10 | 0.278394 | 0.419 | 0.267 | 1.44246E-05 |
| CD74_Intermediat | AIG1 | 1.23106E-36 | 0.382455 | 0.401 | 0.244 | 2.39748E-32 |
| Fibroblasts | IL17RD | 3.66878E-08 | 0.278128 | 0.119 | 0.043 | 0.000714495 |
| Contractile | PEX5 | 1.90736E-30 | 0.278059 | 0.132 | 0.042 | 3.71458E-26 |
| CD74_Intermediat | CTSD | 4.19668E-31 | 0.278047 | 0.954 | 0.873 | 8.17303E-27 |
| Terminal | BCAP31 | 2.74291E-18 | 0.278024 | 0.471 | 0.362 | 5.34181E-14 |
| Terminal | GAPDH | 7.94334E-65 | 0.277854 | 0.972 | 0.914 | 1.54697E-60 |
| IFN_Intermediate | NFKBIA | 0.003682855 | 0.277818 | 0.589 | 0.45 | 1 |
| VEGFA_Terminal | THBS2 | 0.004320309 | 0.27775 | 0.605 | 0.527 | 1 |
| VEGFA_Terminal | TARS1 | 2.81823E-09 | 0.277745 | 0.358 | 0.225 | 5.48849E-05 |
| Contractile | RNF115 | 4.75987E-10 | 0.277719 | 0.251 | 0.178 | 9.26985E-06 |
| Contractile | CAMKK2 | 4.11458E-14 | 0.27763 | 0.208 | 0.122 | 8.01315E-10 |
| Intermediate | ENG | 1.72623E-05 | 0.277481 | 0.438 | 0.371 | 0.336182339 |
| IFN_Intermediate | LTBP3 | 0.006647431 | 0.277394 | 0.822 | 0.715 | 1 |
| Terminal | ARL6IP5 | 1.63596E-17 | 0.277261 | 0.716 | 0.646 | 3.18603E-13 |
| Fibroblasts | CNN3 | 0.006675045 | 0.277256 | 0.642 | 0.663 | 1 |
| Fibroblasts | SIRPA | 0.000107815 | 0.27724 | 0.174 | 0.098 | 1 |
| Contractile | ENSSSCG0( | 1.5259E-09 | 0.277157 | 0.38 | 0.302 | 2.9717E-05 |
| Terminal | TMEM258 | 1.29315E-17 | 0.277137 | 0.676 | 0.582 | 2.5184E-13 |
| Contractile | PRRT1 | 1.62724E-17 | 0.277078 | 0.177 | 0.09 | 3.16904E-13 |
| Contractile | PCYT1A | 1.03434E-12 | 0.277014 | 0.263 | 0.176 | 2.01437E-08 |
| Pericytes | NTF3 | 3.80446E-05 | 0.276984 | 0.1 | 0.019 | 0.740918087 |
| Terminal | DHRS7 | 4.96684E-20 | 0.276936 | 0.454 | 0.337 | 9.67292E-16 |
| CD74_Intermediat | CTSS | 2.8568E-61 | 0.567933 | 0.449 | 0.245 | 5.56363E-57 |
| VEGFA_Terminal | SLC16A1.1 | 1.20401E-13 | 0.276894 | 0.288 | 0.145 | 2.34481E-09 |
| VEGFA_Terminal | MTHFD2 | 3.14882E-19 | 0.276827 | 0.22 | 0.084 | 6.13233E-15 |
| VEGFA_Terminal | CHRAC1 | 9.77482E-12 | 0.276798 | 0.545 | 0.357 | 1.90365E-07 |
| Fibroblasts | MTDH | 0.000934001 | 0.276574 | 0.734 | 0.715 | 1 |
| CD74_Intermediat | C1QTNF1 | 1.85072E-25 | 0.276538 | 0.665 | 0.525 | 3.60428E-21 |
| Contractile | CXXC5 | 4.86218E-10 | 0.276375 | 0.428 | 0.346 | 9.4691E-06 |
| VEGFA_Terminal | TVP23B | 2.56181E-08 | 0.276315 | 0.617 | 0.47 | 0.000498912 |
| CD74_Intermediat | ALCAM | 1.44044E-23 | 0.276314 | 0.635 | 0.491 | 2.80526E-19 |

| | | | | | | |
|---|---|---|---|---|---|---|
| Intermediate | C12ORF75 | 9.56017E-12 | 0.276188 | 0.233 | 0.136 | 1.86184E-07 |
| Terminal | GEM | 1.64851E-20 | 0.276099 | 0.699 | 0.597 | 3.21048E-16 |
| IFN_Intermediate | WSB2 | 0.009295494 | 0.276093 | 0.346 | 0.24 | 1 |
| CD74_Intermediat | LGALS3BP | 4.2601E-106 | 0.74098 | 0.526 | 0.253 | 8.2966E-102 |
| Intermediate | HDAC9 | 9.47826E-08 | 0.275833 | 0.198 | 0.127 | 0.001845892 |
| Contractile | TRAPPC3 | 1.16529E-10 | 0.275741 | 0.405 | 0.324 | 2.26941E-06 |
| Contractile | GUCY1A1 | 9.57119E-19 | 0.275554 | 0.259 | 0.148 | 1.86399E-14 |
| VEGFA_Terminal | POLR1D | 4.50581E-09 | 0.275497 | 0.816 | 0.712 | 8.77507E-05 |
| Contractile | TRAF5 | 1.27816E-34 | 0.275401 | 0.114 | 0.029 | 2.48922E-30 |
| Fibroblasts | PRPSAP2 | 2.12732E-05 | 0.275356 | 0.156 | 0.081 | 0.414295159 |
| VEGFA_Terminal | LGALS1 | 1.60004E-07 | 0.275344 | 0.971 | 0.975 | 0.003116075 |
| Contractile | PDE1A | 1.86072E-11 | 0.275339 | 0.303 | 0.215 | 3.62376E-07 |
| IFN_Intermediate | COL1A2 | 0.002186987 | 0.275271 | 0.972 | 0.945 | 1 |
| Contractile | AIP | 2.97808E-11 | 0.275255 | 0.363 | 0.274 | 5.79981E-07 |
| Contractile | ARFGEF2 | 1.29961E-17 | 0.275237 | 0.202 | 0.108 | 2.53099E-13 |
| Contractile | AMFR | 6.1334E-14 | 0.275186 | 0.25 | 0.158 | 1.19448E-09 |
| Contractile | ARL6IP4 | 9.75598E-12 | 0.275178 | 0.287 | 0.2 | 1.89998E-07 |
| Fibroblasts | ZNF609 | 0.000586251 | 0.275069 | 0.174 | 0.107 | 1 |
| Contractile | KLF15 | 4.54163E-15 | 0.275046 | 0.204 | 0.116 | 8.84483E-11 |
| Contractile | PHIP | 1.53224E-10 | 0.274879 | 0.272 | 0.191 | 2.98403E-06 |
| Contractile | ATP10A | 3.44495E-16 | 0.27476 | 0.185 | 0.099 | 6.70903E-12 |
| Contractile | SNF8 | 5.73939E-12 | 0.274705 | 0.64 | 0.563 | 1.11775E-07 |
| IFN_Intermediate | SMCHD1 | 0.000568048 | 0.274632 | 0.421 | 0.263 | 1 |
| Contractile | ATP8A1 | 3.78355E-47 | 0.274497 | 0.106 | 0.018 | 7.36846E-43 |
| Contractile | PTMS | 1.1143E-30 | 0.274451 | 0.971 | 0.954 | 2.1701E-26 |
| VEGFA_Terminal | PRDX1 | 2.05238E-09 | 0.274362 | 0.719 | 0.59 | 3.99701E-05 |
| IFN_Intermediate | PLOD3 | 0.006300511 | 0.27433 | 0.262 | 0.168 | 1 |
| Contractile | ENSSSCG0( | 9.3659E-12 | 0.274328 | 0.597 | 0.526 | 1.82401E-07 |
| Contractile | TRIM28 | 1.10961E-10 | 0.274309 | 0.345 | 0.262 | 2.16096E-06 |
| Contractile | GPC1 | 4.10612E-27 | 0.274296 | 0.132 | 0.045 | 7.99666E-23 |
| Terminal | ENSSSCG0( | 1.44869E-16 | 0.274257 | 0.524 | 0.421 | 2.82132E-12 |
| Fibroblasts | ENSSSCG0( | 0.000456722 | 0.274 | 0.133 | 0.073 | 1 |
| Contractile | SPSB3 | 5.65443E-11 | 0.273914 | 0.255 | 0.177 | 1.1012E-06 |
| Contractile | OLFM1 | 8.88091E-38 | 0.273906 | 0.122 | 0.03 | 1.72956E-33 |
| VEGFA_Terminal | BCL2L1 | 2.64841E-17 | 0.273883 | 0.186 | 0.07 | 5.15777E-13 |
| Contractile | PRRC2A | 2.1308E-12 | 0.273825 | 0.288 | 0.198 | 4.14973E-08 |
| Fibroblasts | ENSSSCG0( | 3.49274E-08 | 0.273802 | 0.101 | 0.032 | 0.00068021 |
| Contractile | PTGR3 | 9.15691E-13 | 0.273713 | 0.215 | 0.134 | 1.78331E-08 |
| Contractile | PPP2CA | 1.03651E-12 | 0.273609 | 0.33 | 0.237 | 2.01859E-08 |
| Terminal | VASN | 1.12556E-17 | 0.273575 | 0.475 | 0.363 | 2.19203E-13 |
| Contractile | PCBP4 | 2.02523E-12 | 0.273476 | 0.228 | 0.146 | 3.94414E-08 |
| CD74_Intermediat | SESN3 | 3.63002E-49 | 0.472767 | 0.442 | 0.254 | 7.06946E-45 |
| CD74_Intermediat | RCN1 | 8.19456E-23 | 0.273401 | 0.552 | 0.415 | 1.59589E-18 |
| Contractile | SSRP1 | 8.88656E-11 | 0.27339 | 0.329 | 0.244 | 1.73066E-06 |
| Contractile | KPNA4 | 5.38338E-09 | 0.273261 | 0.42 | 0.346 | 0.000104841 |
| Contractile | TXNDC9 | 1.83366E-09 | 0.273193 | 0.356 | 0.283 | 3.57106E-05 |
| Contractile | METAP2 | 2.3444E-12 | 0.27311 | 0.631 | 0.557 | 4.56573E-08 |
| Contractile | DCAF8 | 1.15117E-10 | 0.273031 | 0.309 | 0.227 | 2.2419E-06 |
| Terminal | LTC4S | 1.31141E-21 | 0.27302 | 0.795 | 0.657 | 2.55397E-17 |
| Contractile | CLIP1 | 1.12574E-11 | 0.272963 | 0.641 | 0.561 | 2.19237E-07 |
| Intermediate | BAMBI | 2.72036E-08 | 0.272953 | 0.257 | 0.173 | 0.00052979 |
| Fibroblasts | EIF3A | 0.004689734 | 0.272919 | 0.606 | 0.608 | 1 |
| Contractile | PLEKHB2 | 3.03678E-11 | 0.272827 | 0.25 | 0.17 | 5.91414E-07 |
| Terminal | SEC61G | 1.79027E-19 | 0.272795 | 0.827 | 0.766 | 3.48655E-15 |
| Terminal | C1QTNF7 | 6.79148E-11 | 0.272792 | 0.323 | 0.245 | 1.32264E-06 |
| CD74_Intermediat | C12ORF57 | 3.46322E-19 | 0.27269 | 0.601 | 0.489 | 6.74463E-15 |
| Terminal | TGFB2 | 1.78705E-22 | 0.272685 | 0.347 | 0.226 | 3.48027E-18 |
| IFN_Intermediate | PDIA3 | 0.006464575 | 0.272661 | 0.879 | 0.776 | 1 |
| Contractile | DDAH2 | 1.44382E-11 | 0.272623 | 0.717 | 0.663 | 2.81184E-07 |

| | | | | | | |
|---|---|---|---|---|---|---|
| VEGFA_Terminal | ATP6V0B | 2.38792E-08 | 0.272447 | 0.588 | 0.43 | 0.000465048 |
| Intermediate | KCNA5 | 1.6283E-09 | 0.272445 | 0.129 | 0.066 | 3.17111E-05 |
| VEGFA_Terminal | ISCA1 | 1.63826E-06 | 0.272424 | 0.639 | 0.501 | 0.031905129 |
| Terminal | PLPP2 | 4.02959E-30 | 0.272373 | 0.199 | 0.095 | 7.84764E-26 |
| CD74_Intermediat | IL1R1 | 8.81314E-24 | 0.324783 | 0.382 | 0.258 | 1.71636E-19 |
| Fibroblasts | ENSSSCG0( | 0.004930606 | 0.272365 | 0.183 | 0.125 | 1 |
| Intermediate | ENSSSCG0( | 3.7615E-06 | 0.272311 | 0.571 | 0.516 | 0.073255165 |
| CD74_Intermediat | MAT2A | 6.1478E-14 | 0.272253 | 0.572 | 0.481 | 1.19728E-09 |
| IFN_Intermediate | PLAA | 1.60977E-05 | 0.272231 | 0.215 | 0.092 | 0.313502951 |
| Fibroblasts | EFS | 1.28301E-05 | 0.272152 | 0.124 | 0.056 | 0.249866171 |
| Terminal | UBL5 | 5.4182E-29 | 0.272152 | 0.881 | 0.809 | 1.05519E-24 |
| Contractile | MINDY3 | 3.62582E-19 | 0.272075 | 0.174 | 0.084 | 7.06128E-15 |
| Fibroblasts | WNT11 | 1.54471E-05 | 0.272073 | 0.119 | 0.054 | 0.300832293 |
| Contractile | HNRNPK | 1.34894E-19 | 0.271921 | 0.817 | 0.746 | 2.62706E-15 |
| Contractile | PLS3 | 1.47473E-10 | 0.271618 | 0.768 | 0.718 | 2.87204E-06 |
| Contractile | RBM3 | 8.80931E-11 | 0.271549 | 0.561 | 0.483 | 1.71561E-06 |
| Fibroblasts | GPD1L | 5.22447E-05 | 0.271415 | 0.106 | 0.047 | 1 |
| Contractile | ARL4D | 8.94653E-37 | 0.27139 | 0.123 | 0.031 | 1.74234E-32 |
| Contractile | CD200 | 2.99149E-09 | 0.271373 | 0.436 | 0.353 | 5.82592E-05 |
| CD74_Intermediat | FOXS1 | 5.6995E-19 | 0.271249 | 0.523 | 0.395 | 1.10998E-14 |
| VEGFA_Terminal | CSDC2 | 1.71088E-10 | 0.271152 | 0.23 | 0.121 | 3.33195E-06 |
| Contractile | CDC42 | 1.07837E-12 | 0.271143 | 0.7 | 0.625 | 2.10013E-08 |
| CD74_Intermediat | PDGFRA | 4.30608E-52 | 0.453547 | 0.46 | 0.259 | 8.3861E-48 |
| Contractile | UBP1 | 1.58769E-09 | 0.271107 | 0.257 | 0.187 | 3.09202E-05 |
| CD74_Intermediat | ADAMTS2 | 1.45999E-53 | 0.507587 | 0.448 | 0.259 | 2.84333E-49 |
| IFN_Intermediate | PIK3IP1 | 0.00844987 | 0.271042 | 0.327 | 0.225 | 1 |
| Terminal | TMEM167/ | 4.9424E-19 | 0.271008 | 0.471 | 0.356 | 9.62532E-15 |
| Contractile | SETD1B | 1.11879E-17 | 0.270949 | 0.16 | 0.077 | 2.17884E-13 |
| VEGFA_Terminal | SLC35A3 | 3.13601E-12 | 0.270889 | 0.242 | 0.121 | 6.10738E-08 |
| Contractile | SNTA1 | 1.82791E-16 | 0.270774 | 0.206 | 0.115 | 3.55985E-12 |
| Contractile | ENSSSCG0( | 8.38151E-11 | 0.270701 | 0.265 | 0.183 | 1.6323E-06 |
| Fibroblasts | AKR7A2 | 0.001939429 | 0.270541 | 0.124 | 0.073 | 1 |
| Contractile | ARHGEF26 | 3.10093E-20 | 0.270539 | 0.161 | 0.073 | 6.03905E-16 |
| Fibroblasts | SRRM1 | 0.003142227 | 0.270536 | 0.583 | 0.574 | 1 |
| Contractile | AKT2 | 1.76729E-17 | 0.270498 | 0.183 | 0.095 | 3.4418E-13 |
| Contractile | PRKAB2 | 1.41503E-18 | 0.270463 | 0.173 | 0.085 | 2.75578E-14 |
| Terminal | ENSSSCG0( | 1.50174E-18 | 0.270134 | 0.662 | 0.55 | 2.92464E-14 |
| CD74_Intermediat | TIMP2 | 4.68049E-39 | 0.27011 | 0.965 | 0.907 | 9.11525E-35 |
| VEGFA_Terminal | PSPH | 1.8165E-15 | 0.270079 | 0.182 | 0.072 | 3.53764E-11 |
| Contractile | ZFR | 2.00189E-09 | 0.27007 | 0.378 | 0.298 | 3.89868E-05 |
| Contractile | CCDC124 | 4.79759E-13 | 0.269924 | 0.378 | 0.282 | 9.3433E-09 |
| CD74_Intermediat | LRIG1 | 1.69857E-62 | 0.558554 | 0.467 | 0.26 | 3.30796E-58 |
| Fibroblasts | ATP8B1 | 0.004284934 | 0.269897 | 0.183 | 0.125 | 1 |
| VEGFA_Terminal | DLGAP4 | 7.22689E-11 | 0.269733 | 0.433 | 0.271 | 1.40744E-06 |
| Contractile | AKAP1 | 3.17923E-22 | 0.269587 | 0.145 | 0.06 | 6.19154E-18 |
| Contractile | PLCE1 | 4.70292E-12 | 0.269545 | 0.237 | 0.153 | 9.15894E-08 |
| Contractile | DYNLRB1 | 4.71026E-15 | 0.269398 | 0.781 | 0.701 | 9.17324E-11 |
| Fibroblasts | RBM25 | 0.004444421 | 0.269386 | 0.491 | 0.447 | 1 |
| Fibroblasts | LLGL1 | 0.006487021 | 0.269379 | 0.151 | 0.101 | 1 |
| Contractile | NIPBL | 7.57564E-11 | 0.269324 | 0.444 | 0.354 | 1.47536E-06 |
| Contractile | PRKAR2A | 1.04145E-12 | 0.269219 | 0.24 | 0.155 | 2.02823E-08 |
| Contractile | RANBP9 | 1.15863E-11 | 0.269218 | 0.208 | 0.132 | 2.25644E-07 |
| Contractile | ENSSSCG0( | 7.50682E-12 | 0.269144 | 0.634 | 0.564 | 1.46195E-07 |
| VEGFA_Terminal | COX7A2 | 1.79536E-08 | 0.268938 | 0.85 | 0.759 | 0.000349647 |
| Contractile | SIK2 | 9.94796E-18 | 0.268932 | 0.152 | 0.071 | 1.93737E-13 |
| Contractile | VPS26A | 3.78859E-11 | 0.268671 | 0.495 | 0.408 | 7.37828E-07 |
| Contractile | RBSN | 1.39759E-13 | 0.268651 | 0.223 | 0.136 | 2.7218E-09 |
| VEGFA_Terminal | SLC12A2 | 2.26393E-15 | 0.26861 | 0.186 | 0.075 | 4.40901E-11 |
| Intermediate | GGT5 | 1.86873E-07 | 0.268599 | 0.115 | 0.062 | 0.003639343 |

| | | | | | | |
|---|---|---|---|---|---|---|
| Contractile | MRAP2 | 3.44742E-16 | 0.268573 | 0.237 | 0.138 | 6.71385E-12 |
| Terminal | BZW1 | 2.18683E-19 | 0.268481 | 0.665 | 0.565 | 4.25886E-15 |
| Terminal | KCNN4 | 2.04725E-56 | 0.268462 | 0.117 | 0.021 | 3.98702E-52 |
| Contractile | FLOT2 | 4.28358E-12 | 0.26844 | 0.284 | 0.196 | 8.34226E-08 |
| Contractile | TERF2IP | 2.25602E-11 | 0.26828 | 0.432 | 0.347 | 4.3936E-07 |
| Contractile | MTMR9 | 3.61589E-13 | 0.26823 | 0.15 | 0.081 | 7.04195E-09 |
| Terminal | ARF4 | 5.91015E-17 | 0.268224 | 0.681 | 0.602 | 1.151E-12 |
| Contractile | TRIM65 | 4.21184E-28 | 0.268182 | 0.135 | 0.045 | 8.20256E-24 |
| Terminal | KLF2 | 1.35757E-28 | 0.268081 | 0.486 | 0.325 | 2.64386E-24 |
| VEGFA_Terminal | KCMF1 | 4.00706E-11 | 0.268075 | 0.375 | 0.223 | 7.80375E-07 |
| Terminal | ENSSSCG0( | 4.99868E-34 | 0.268057 | 0.217 | 0.101 | 9.73492E-30 |
| Contractile | TMEM94 | 2.28219E-23 | 0.268056 | 0.117 | 0.041 | 4.44457E-19 |
| Contractile | KANSL1L | 1.05544E-15 | 0.267981 | 0.2 | 0.111 | 2.05548E-11 |
| VEGFA_Terminal | LOC10051 | 1.37178E-07 | 0.26791 | 0.663 | 0.534 | 0.002671539 |
| Terminal | NME2 | 3.82765E-30 | 0.267861 | 0.94 | 0.897 | 7.45435E-26 |
| CD74_Intermediat | RNF19A | 4.48077E-16 | 0.267808 | 0.709 | 0.626 | 8.7263E-12 |
| Contractile | CRY2 | 1.02371E-16 | 0.267732 | 0.157 | 0.077 | 1.99368E-12 |
| Fibroblasts | ALAD | 0.000195631 | 0.267648 | 0.151 | 0.084 | 1 |
| Intermediate | SRPX | 1.49588E-05 | 0.26756 | 0.442 | 0.372 | 0.291321829 |
| VEGFA_Terminal | PLK3 | 2.22423E-14 | 0.267538 | 0.138 | 0.049 | 4.33169E-10 |
| VEGFA_Terminal | THYN1 | 3.38073E-07 | 0.267511 | 0.472 | 0.335 | 0.006583964 |
| Terminal | COPE | 1.67581E-20 | 0.267427 | 0.709 | 0.612 | 3.26363E-16 |
| VEGFA_Terminal | BTF3 | 2.16458E-09 | 0.267383 | 0.935 | 0.891 | 4.21553E-05 |
| VEGFA_Terminal | ARHGAP22 | 1.56067E-11 | 0.267294 | 0.295 | 0.161 | 3.0394E-07 |
| Contractile | NEDD4 | 1.76964E-11 | 0.266893 | 0.257 | 0.173 | 3.44637E-07 |
| CD74_Intermediat | ECM2 | 1.3107E-23 | 0.266715 | 0.785 | 0.671 | 2.5526E-19 |
| Terminal | HOMER3 | 6.3692E-19 | 0.266711 | 0.328 | 0.224 | 1.2404E-14 |
| Fibroblasts | TBC1D10A | 0.002317373 | 0.266625 | 0.138 | 0.084 | 1 |
| Contractile | ZNF664 | 2.38519E-10 | 0.266531 | 0.266 | 0.189 | 4.64515E-06 |
| VEGFA_Terminal | DUSP7 | 2.02841E-13 | 0.266511 | 0.194 | 0.084 | 3.95033E-09 |
| VEGFA_Terminal | ATP1B3 | 6.80154E-06 | 0.266462 | 0.821 | 0.764 | 0.132459993 |
| Contractile | EPHX1 | 8.9704E-11 | 0.266319 | 0.307 | 0.222 | 1.74699E-06 |
| Contractile | SENP6 | 6.22083E-09 | 0.266278 | 0.41 | 0.334 | 0.000121151 |
| VEGFA_Terminal | PRELID1 | 1.38853E-07 | 0.266264 | 0.506 | 0.377 | 0.002704162 |
| Contractile | LUC7L3 | 3.13316E-09 | 0.266214 | 0.528 | 0.458 | 6.10184E-05 |
| VEGFA_Terminal | MRPL23 | 9.98844E-08 | 0.266163 | 0.712 | 0.573 | 0.001945249 |
| CD74_Intermediat | RNASE4 | 3.25128E-29 | 0.266031 | 0.85 | 0.736 | 6.33187E-25 |
| Intermediate | MYL12A | 1.95175E-07 | 0.26602 | 0.865 | 0.865 | 0.003801027 |
| Terminal | LBH | 2.07172E-27 | 0.265939 | 0.198 | 0.098 | 4.03467E-23 |
| CD74_Intermediat | OXCT1 | 6.49025E-18 | 0.265922 | 0.516 | 0.405 | 1.26398E-13 |
| Fibroblasts | TMEM107 | 0.000429995 | 0.265732 | 0.151 | 0.088 | 1 |
| VEGFA_Terminal | ATP1A1 | 7.75017E-07 | 0.265725 | 0.586 | 0.458 | 0.015093459 |
| Terminal | REXO2 | 1.79099E-24 | 0.265721 | 0.852 | 0.781 | 3.48795E-20 |
| VEGFA_Terminal | S100A6 | 7.68194E-08 | 0.265565 | 0.985 | 0.994 | 0.001496057 |
| Terminal | SULF1 | 3.94616E-18 | 0.26544 | 0.492 | 0.366 | 7.68514E-14 |
| Contractile | HSDL2 | 4.62337E-11 | 0.26542 | 0.219 | 0.143 | 9.00402E-07 |
| Contractile | POLDIP2 | 2.04913E-11 | 0.265383 | 0.228 | 0.149 | 3.99067E-07 |
| Contractile | SNRPC | 1.66506E-10 | 0.265359 | 0.369 | 0.285 | 3.24271E-06 |
| Contractile | NCAM1 | 1.31576E-11 | 0.265237 | 0.457 | 0.352 | 2.56243E-07 |
| Contractile | LOC10062( | 8.02726E-13 | 0.265162 | 0.234 | 0.148 | 1.56331E-08 |
| Contractile | ZBTB10 | 9.95462E-12 | 0.265159 | 0.263 | 0.179 | 1.93866E-07 |
| Contractile | METTL24 | 1.17698E-19 | 0.265107 | 0.159 | 0.072 | 2.29217E-15 |
| VEGFA_Terminal | UQCC2 | 1.10984E-08 | 0.265089 | 0.571 | 0.412 | 0.000216141 |
| VEGFA_Terminal | PDCD5 | 9.45498E-08 | 0.265073 | 0.741 | 0.629 | 0.001841357 |
| Contractile | BANF1 | 2.02383E-11 | 0.265059 | 0.771 | 0.733 | 3.9414E-07 |
| VEGFA_Terminal | PERP | 3.12351E-13 | 0.265053 | 0.157 | 0.062 | 6.08303E-09 |
| CD74_Intermediat | CHRD | 5.381E-16 | 0.279924 | 0.357 | 0.261 | 1.04795E-11 |
| Contractile | CSNK1A1 | 1.7405E-11 | 0.264999 | 0.703 | 0.623 | 3.38963E-07 |
| VEGFA_Terminal | PTPA | 1.85971E-07 | 0.264854 | 0.579 | 0.44 | 0.003621776 |

| | | | | | | |
|---|---|---|---|---|---|---|
| Contractile | CHD4 | 5.64266E-12 | 0.264751 | 0.612 | 0.532 | 1.09891E-07 |
| Intermediate | FAM13C | 5.61152E-08 | 0.264691 | 0.343 | 0.247 | 0.001092843 |
| Contractile | YWHAQ | 1.99403E-13 | 0.264617 | 0.69 | 0.61 | 3.88337E-09 |
| CD74_Intermediat | PAM | 3.16566E-23 | 0.264575 | 0.814 | 0.716 | 6.16511E-19 |
| VEGFA_Terminal | LIMD1 | 3.63283E-08 | 0.264557 | 0.213 | 0.119 | 0.000707493 |
| VEGFA_Terminal | KLF10 | 0.003649358 | 0.264524 | 0.356 | 0.28 | 1 |
| VEGFA_Terminal | TMEM223 | 4.02897E-07 | 0.264484 | 0.7 | 0.584 | 0.007846426 |
| Terminal | RETREG1 | 7.75453E-26 | 0.264426 | 0.21 | 0.108 | 1.5102E-21 |
| CD74_Intermediat | TPP1 | 2.91708E-17 | 0.264423 | 0.584 | 0.477 | 5.68101E-13 |
| Intermediate | SAMD12 | 7.90659E-10 | 0.264332 | 0.304 | 0.203 | 1.53981E-05 |
| IFN_Intermediate | TM9SF2 | 0.005025422 | 0.264252 | 0.617 | 0.482 | 1 |
| Contractile | BTF3 | 3.81132E-20 | 0.264071 | 0.904 | 0.892 | 7.42255E-16 |
| Fibroblasts | ADGRA2 | 0.000396131 | 0.26395 | 0.115 | 0.059 | 1 |
| Terminal | GALNT1 | 5.12574E-14 | 0.263728 | 0.453 | 0.355 | 9.98239E-10 |
| IFN_Intermediate | FKBP11 | 0.002402018 | 0.263598 | 0.308 | 0.193 | 1 |
| Contractile | XPO1 | 1.38455E-08 | 0.263548 | 0.294 | 0.223 | 0.00026964 |
| VEGFA_Terminal | ENSSSCG0( | 1.58722E-10 | 0.263535 | 0.308 | 0.175 | 3.09111E-06 |
| IFN_Intermediate | CTSD | 0.00147115 | 0.263437 | 0.963 | 0.895 | 1 |
| Fibroblasts | HERC4 | 0.006286073 | 0.263342 | 0.138 | 0.089 | 1 |
| Contractile | KAT6B | 1.91478E-08 | 0.263318 | 0.384 | 0.308 | 0.000372903 |
| Terminal | CARD19 | 6.20767E-18 | 0.263187 | 0.338 | 0.239 | 1.20894E-13 |
| Intermediate | WDR1 | 1.40298E-05 | 0.263049 | 0.528 | 0.47 | 0.273231115 |
| Contractile | PDK2 | 6.06389E-19 | 0.262985 | 0.142 | 0.063 | 1.18094E-14 |
| Contractile | SUCO | 6.86765E-12 | 0.262974 | 0.273 | 0.188 | 1.33747E-07 |
| Contractile | SLAIN2 | 8.85094E-11 | 0.26294 | 0.304 | 0.219 | 1.72372E-06 |
| VEGFA_Terminal | SLC1A4 | 6.77699E-16 | 0.26291 | 0.169 | 0.063 | 1.31982E-11 |
| Contractile | PANK3 | 2.00886E-12 | 0.262823 | 0.174 | 0.101 | 3.91225E-08 |
| IFN_Intermediate | TSPAN6 | 0.00874541 | 0.262798 | 0.533 | 0.41 | 1 |
| VEGFA_Terminal | BCL3 | 9.68201E-12 | 0.262736 | 0.199 | 0.092 | 1.88557E-07 |
| IFN_Intermediate | TOR3A | 2.00112E-05 | 0.262591 | 0.15 | 0.054 | 0.389718699 |
| IFN_Intermediate | SCARA3 | 0.004706859 | 0.262495 | 0.589 | 0.458 | 1 |
| CD74_Intermediat | GRN | 2.70902E-15 | 0.262369 | 0.563 | 0.465 | 5.27581E-11 |
| VEGFA_Terminal | TTC7A | 1.93045E-11 | 0.262204 | 0.237 | 0.12 | 3.75955E-07 |
| CD74_Intermediat | CHI3L1 | 4.27888E-61 | 0.262154 | 0.732 | 0.469 | 8.33313E-57 |
| VEGFA_Terminal | NDUFS4 | 1.56854E-08 | 0.26208 | 0.688 | 0.543 | 0.000305473 |
| CD74_Intermediat | S100A14 | 7.7955E-39 | 0.435338 | 0.419 | 0.262 | 1.51817E-34 |
| IFN_Intermediate | ACSL4 | 0.005244938 | 0.261884 | 0.327 | 0.22 | 1 |
| Contractile | SVBP | 1.86705E-12 | 0.261871 | 0.39 | 0.29 | 3.63607E-08 |
| Contractile | KHSRP | 3.30863E-11 | 0.261856 | 0.253 | 0.17 | 6.44356E-07 |
| Contractile | ZNF106 | 2.59618E-10 | 0.261758 | 0.294 | 0.214 | 5.05606E-06 |
| VEGFA_Terminal | HNRNPF | 5.1122E-07 | 0.261725 | 0.751 | 0.602 | 0.009956002 |
| VEGFA_Terminal | PCDH17 | 3.70971E-12 | 0.261696 | 0.196 | 0.09 | 7.22466E-08 |
| Terminal | FUT8 | 1.99564E-25 | 0.261685 | 0.27 | 0.158 | 3.8865E-21 |
| CD74_Intermediat | ENSSSCG0( | 2.014E-31 | 0.381443 | 0.41 | 0.265 | 3.92227E-27 |
| Contractile | PNKD.1 | 2.83705E-08 | 0.261617 | 0.38 | 0.313 | 0.000552515 |
| VEGFA_Terminal | RAP1B | 1.40957E-08 | 0.26154 | 0.63 | 0.459 | 0.000274514 |
| Intermediate | JUNB | 3.39216E-05 | 0.261421 | 0.61 | 0.547 | 0.660623242 |
| Contractile | ARL14EP | 5.34228E-10 | 0.261376 | 0.429 | 0.343 | 1.04041E-05 |
| Contractile | PCBD1 | 2.51905E-11 | 0.261116 | 0.344 | 0.253 | 4.90585E-07 |
| VEGFA_Terminal | BAZ1A | 1.73257E-14 | 0.26106 | 0.182 | 0.074 | 3.37418E-10 |
| Contractile | EMILIN1 | 7.85245E-08 | 0.260938 | 0.375 | 0.307 | 0.001529264 |
| Contractile | PAK1 | 4.75321E-39 | 0.260895 | 0.118 | 0.028 | 9.25687E-35 |
| Contractile | PAFAH1B1 | 2.79272E-10 | 0.260888 | 0.661 | 0.616 | 5.43882E-06 |
| Intermediate | DCBLD2 | 8.80173E-07 | 0.26088 | 0.346 | 0.266 | 0.017141372 |
| Contractile | ANKRD39 | 5.64293E-11 | 0.26076 | 0.305 | 0.223 | 1.09896E-06 |
| CD74_Intermediat | CTNNB1 | 6.38481E-14 | 0.260747 | 0.661 | 0.588 | 1.24344E-09 |
| Fibroblasts | YES1 | 0.006291174 | 0.260713 | 0.161 | 0.108 | 1 |
| VEGFA_Terminal | TMEM50A | 4.45795E-08 | 0.260674 | 0.755 | 0.618 | 0.000868187 |
| Contractile | FMC1 | 2.27698E-10 | 0.260529 | 0.281 | 0.203 | 4.43442E-06 |

| | | | | | | |
|---|---|---|---|---|---|---|
| Fibroblasts | EFNA5 | 0.000198157 | 0.260528 | 0.147 | 0.08 | 1 |
| Contractile | MYH10 | 3.77434E-12 | 0.260501 | 0.689 | 0.596 | 7.35054E-08 |
| Terminal | FARP1 | 8.40831E-22 | 0.260484 | 0.363 | 0.244 | 1.63752E-17 |
| Intermediate | FOXP1 | 0.00030032 | 0.260402 | 0.53 | 0.491 | 1 |
| Fibroblasts | BCOR | 0.004407016 | 0.260386 | 0.124 | 0.076 | 1 |
| Terminal | DSE | 5.58724E-23 | 0.260382 | 0.264 | 0.157 | 1.08812E-18 |
| CD74_Intermediat | PLXDC2 | 9.19828E-30 | 0.260291 | 0.839 | 0.703 | 1.79137E-25 |
| Contractile | SECISBP2L | 6.62706E-08 | 0.260226 | 0.424 | 0.358 | 0.001290619 |
| Contractile | MAGEF1 | 1.9129E-11 | 0.260205 | 0.327 | 0.237 | 3.72538E-07 |
| VEGFA_Terminal | BZW1 | 3.75783E-08 | 0.260201 | 0.746 | 0.587 | 0.000731837 |
| Intermediate | MSRB3 | 2.06017E-08 | 0.260097 | 0.561 | 0.469 | 0.000401219 |
| IFN_Intermediate | CALCOCO2 | 0.004760946 | 0.260027 | 0.785 | 0.667 | 1 |
| Terminal | RUNX1 | 1.66525E-20 | 0.25973 | 0.197 | 0.111 | 3.24307E-16 |
| VEGFA_Terminal | PSME2 | 1.6465E-06 | 0.259497 | 0.351 | 0.244 | 0.03206559 |
| Contractile | PRPF8 | 6.2457E-09 | 0.259481 | 0.286 | 0.212 | 0.000121635 |
| Contractile | THRAP3 | 4.40684E-09 | 0.259464 | 0.595 | 0.538 | 8.58231E-05 |
| Contractile | PRKAA2 | 1.83729E-36 | 0.259342 | 0.115 | 0.028 | 3.57812E-32 |
| CD74_Intermediat | RAMP2 | 1.0055E-168 | 0.979048 | 0.622 | 0.267 | 1.9583E-164 |
| VEGFA_Terminal | SPRYD7 | 3.366E-08 | 0.259264 | 0.358 | 0.233 | 0.000655529 |
| Fibroblasts | UBA52 | 9.94502E-05 | 0.259235 | 0.968 | 0.948 | 1 |
| Contractile | ENSSSCG00 | 6.43772E-10 | 0.259103 | 0.813 | 0.768 | 1.25375E-05 |
| Intermediate | CREB5 | 0.003959221 | 0.259098 | 0.28 | 0.244 | 1 |
| VEGFA_Terminal | EIF2S2 | 2.06629E-08 | 0.258987 | 0.748 | 0.617 | 0.000402411 |
| Contractile | INMT | 5.19275E-17 | 0.258789 | 0.176 | 0.088 | 1.01129E-12 |
| IFN_Intermediate | DCN | 0.000497995 | 0.258736 | 0.916 | 0.878 | 1 |
| VEGFA_Terminal | CD46 | 4.86801E-06 | 0.258702 | 0.673 | 0.54 | 0.09480454 |
| Contractile | TMEM106 | 1.3196E-09 | 0.258682 | 0.579 | 0.519 | 2.56993E-05 |
| Contractile | ZNF330 | 9.24316E-12 | 0.258488 | 0.227 | 0.146 | 1.80011E-07 |
| IFN_Intermediate | TIMP3 | 0.000480485 | 0.258459 | 0.953 | 0.94 | 1 |
| Contractile | PPP3CB | 7.46349E-10 | 0.25845 | 0.4 | 0.315 | 1.45352E-05 |
| Terminal | PXDC1 | 1.57212E-18 | 0.258388 | 0.491 | 0.378 | 3.06171E-14 |
| CD74_Intermediat | UBA52 | 3.57127E-34 | 0.258344 | 0.965 | 0.943 | 6.95505E-30 |
| CD74_Intermediat | LHFPL6 | 2.02335E-22 | 0.258273 | 0.809 | 0.698 | 3.94047E-18 |
| VEGFA_Terminal | FKBP10 | 2.07841E-09 | 0.258258 | 0.462 | 0.31 | 4.0477E-05 |
| Intermediate | KCTD10 | 6.4993E-08 | 0.258074 | 0.419 | 0.319 | 0.001265739 |
| Intermediate | BHMT2 | 1.34232E-08 | 0.258016 | 0.138 | 0.074 | 0.000261417 |
| Contractile | WWTR1 | 1.05891E-09 | 0.257918 | 0.579 | 0.516 | 2.06223E-05 |
| Contractile | LSM7 | 3.48478E-09 | 0.257857 | 0.376 | 0.301 | 6.78661E-05 |
| Contractile | IL6R | 1.41757E-20 | 0.257844 | 0.125 | 0.049 | 2.76072E-16 |
| Contractile | ZFP91 | 4.68535E-11 | 0.257797 | 0.636 | 0.564 | 9.12472E-07 |
| Contractile | TENT5C | 6.21877E-14 | 0.25774 | 0.187 | 0.104 | 1.21111E-09 |
| Contractile | YY1 | 7.20073E-11 | 0.257728 | 0.595 | 0.51 | 1.40234E-06 |
| VEGFA_Terminal | BIRC3 | 3.5243E-12 | 0.257429 | 0.363 | 0.205 | 6.86357E-08 |
| VEGFA_Terminal | SPCS1 | 1.68258E-08 | 0.257419 | 0.792 | 0.657 | 0.000327682 |
| VEGFA_Terminal | LOC10052 | 7.86254E-09 | 0.257387 | 0.964 | 0.925 | 0.000153123 |
| Contractile | DLD | 9.05627E-10 | 0.257103 | 0.335 | 0.256 | 1.76371E-05 |
| Contractile | DYRK3 | 2.62214E-11 | 0.256651 | 0.255 | 0.172 | 5.10662E-07 |
| VEGFA_Terminal | HSPA9 | 1.59694E-06 | 0.256438 | 0.397 | 0.281 | 0.031100502 |
| Fibroblasts | PDE5A | 0.001250472 | 0.256419 | 0.307 | 0.224 | 1 |
| Contractile | CD2AP | 6.8093E-08 | 0.25627 | 0.402 | 0.334 | 0.001326112 |
| Contractile | BRD2 | 3.05587E-09 | 0.256226 | 0.447 | 0.368 | 5.9513E-05 |
| Contractile | IRF2BP2 | 3.34491E-09 | 0.256225 | 0.504 | 0.427 | 6.51421E-05 |
| VEGFA_Terminal | FOSB | 1.0123E-08 | 0.256217 | 0.385 | 0.248 | 0.000197145 |
| Contractile | FN3K | 1.93259E-16 | 0.256204 | 0.195 | 0.105 | 3.76371E-12 |
| Contractile | SAMD11 | 1.91338E-15 | 0.256083 | 0.217 | 0.124 | 3.7263E-11 |
| Intermediate | VGLL3 | 1.88719E-09 | 0.256036 | 0.233 | 0.145 | 3.67531E-05 |
| Contractile | WDR82 | 3.91767E-08 | 0.256001 | 0.478 | 0.413 | 0.000762966 |
| Contractile | ATF5 | 7.15833E-25 | 0.255933 | 0.149 | 0.057 | 1.39408E-20 |
| CD74_Intermediat | CAT | 8.98914E-24 | 0.25588 | 0.448 | 0.306 | 1.75064E-19 |

| | | | | | | |
|---|---|---|---|---|---|---|
| Intermediate | ADGRL2 | 7.85842E-07 | 0.255735 | 0.402 | 0.316 | 0.015304269 |
| Intermediate | PALM2AKA | 0.003300522 | 0.255581 | 0.469 | 0.433 | 1 |
| VEGFA_Terminal | CTTN | 1.24721E-06 | 0.255487 | 0.632 | 0.496 | 0.024289401 |
| VEGFA_Terminal | ENSSSCG00 | 1.50918E-13 | 0.255399 | 0.99 | 0.994 | 2.93914E-09 |
| IFN_Intermediate | ASB13 | 0.00333407 | 0.255274 | 0.168 | 0.086 | 1 |
| CD74_Intermediat | EPB41L2 | 4.42398E-20 | 0.255243 | 0.574 | 0.449 | 8.61569E-16 |
| VEGFA_Terminal | ENSSSCG00 | 7.48014E-11 | 0.255221 | 0.262 | 0.14 | 1.45676E-06 |
| Contractile | KDSR | 1.60141E-07 | 0.255164 | 0.45 | 0.389 | 0.003118737 |
| Fibroblasts | ENSSSCG00 | 0.000139503 | 0.254899 | 0.986 | 0.961 | 1 |
| VEGFA_Terminal | ORMDL2 | 1.64678E-08 | 0.25488 | 0.274 | 0.16 | 0.000320711 |
| Contractile | ARMC10 | 1.30185E-09 | 0.254864 | 0.567 | 0.5 | 2.53536E-05 |
| Contractile | PAIP2B | 8.44059E-21 | 0.254712 | 0.13 | 0.051 | 1.64381E-16 |
| VEGFA_Terminal | COL4A2 | 0.001675828 | 0.254668 | 0.663 | 0.58 | 1 |
| Terminal | TNXB | 7.76666E-13 | 0.254637 | 0.623 | 0.515 | 1.51256E-08 |
| Terminal | TJP2 | 7.09496E-21 | 0.254472 | 0.287 | 0.182 | 1.38174E-16 |
| Terminal | TNFRSF12A | 1.16805E-08 | 0.254461 | 0.517 | 0.457 | 0.000227478 |
| Contractile | CDK13 | 5.37303E-10 | 0.254458 | 0.286 | 0.208 | 1.0464E-05 |
| VEGFA_Terminal | GOLM1 | 2.3439E-11 | 0.25443 | 0.409 | 0.247 | 4.56475E-07 |
| CD74_Intermediat | SLA-8 | 1.4278E-23 | 0.281606 | 0.391 | 0.267 | 2.78064E-19 |
| Fibroblasts | ATG2B | 0.000324052 | 0.254395 | 0.133 | 0.071 | 1 |
| CD74_Intermediat | ITM2B | 6.29093E-38 | 0.254293 | 0.984 | 0.969 | 1.22516E-33 |
| Intermediate | KIF5B | 5.43533E-06 | 0.254291 | 0.714 | 0.666 | 0.105853121 |
| IFN_Intermediate | SFRP1 | 0.000425604 | 0.254217 | 0.981 | 0.818 | 1 |
| Intermediate | PAM | 1.50769E-07 | 0.254181 | 0.782 | 0.739 | 0.002936228 |
| VEGFA_Terminal | MYADM | 3.23345E-08 | 0.254105 | 0.642 | 0.489 | 0.000629715 |
| Contractile | FUS | 2.58286E-12 | 0.25391 | 0.341 | 0.245 | 5.03013E-08 |
| Contractile | CEP128 | 1.0127E-22 | 0.253893 | 0.12 | 0.043 | 1.97224E-18 |
| Intermediate | LAMA4 | 2.08193E-06 | 0.253877 | 0.499 | 0.425 | 0.040545648 |
| Contractile | PCED1A | 3.60125E-19 | 0.253829 | 0.171 | 0.082 | 7.01344E-15 |
| Contractile | TRMT112 | 6.09366E-13 | 0.25352 | 0.739 | 0.692 | 1.18674E-08 |
| Contractile | COX7B | 9.76334E-14 | 0.253355 | 0.811 | 0.766 | 1.90141E-09 |
| VEGFA_Terminal | YARS1 | 2.00022E-17 | 0.253345 | 0.167 | 0.059 | 3.89543E-13 |
| VEGFA_Terminal | FOS | 1.4082E-06 | 0.253282 | 0.671 | 0.581 | 0.027424735 |
| VEGFA_Terminal | CDKN1A | 1.10637E-13 | 0.25327 | 0.247 | 0.115 | 2.15466E-09 |
| VEGFA_Terminal | ATF5 | 0.000387236 | 0.253173 | 0.114 | 0.069 | 1 |
| Fibroblasts | L2HGDH | 0.001689122 | 0.253146 | 0.115 | 0.064 | 1 |
| VEGFA_Terminal | COX5B | 2.58537E-08 | 0.252941 | 0.85 | 0.74 | 0.000503502 |
| IFN_Intermediate | ENSSSCG00 | 1.05453E-05 | 0.252925 | 0.187 | 0.072 | 0.205369201 |
| Contractile | ENSSSCG00 | 1.45756E-14 | 0.252924 | 0.798 | 0.693 | 2.8386E-10 |
| VEGFA_Terminal | IER3IP1 | 1.02219E-08 | 0.252875 | 0.499 | 0.344 | 0.000199071 |
| Terminal | H2AZ2 | 2.39608E-18 | 0.252857 | 0.727 | 0.638 | 4.66636E-14 |
| Contractile | UBQLN1 | 8.84951E-11 | 0.252828 | 0.333 | 0.248 | 1.72344E-06 |
| Contractile | CSNK2B | 6.07883E-11 | 0.25281 | 0.519 | 0.429 | 1.18385E-06 |
| IFN_Intermediate | ATAD1 | 6.98196E-05 | 0.252745 | 0.224 | 0.104 | 1 |
| VEGFA_Terminal | SRD5A1 | 8.05517E-32 | 0.252738 | 0.131 | 0.025 | 1.56874E-27 |
| Contractile | FBXO21 | 1.2138E-08 | 0.252633 | 0.223 | 0.156 | 0.000236387 |
| CD74_Intermediat | SMAD6 | 3.63387E-16 | 0.289318 | 0.384 | 0.285 | 7.07697E-12 |
| VEGFA_Terminal | ENSSSCG00 | 1.82046E-05 | 0.252494 | 0.816 | 0.708 | 0.354534648 |
| VEGFA_Terminal | CD109 | 1.5335E-09 | 0.252375 | 0.663 | 0.49 | 2.98649E-05 |
| CD74_Intermediat | ALDH1A2 | 5.4211E-120 | 0.784742 | 0.593 | 0.291 | 1.0558E-115 |
| Contractile | ZNF277 | 2.19159E-11 | 0.252249 | 0.489 | 0.396 | 4.26812E-07 |
| IFN_Intermediate | LMO7 | 0.007937205 | 0.25219 | 0.355 | 0.25 | 1 |
| Fibroblasts | NDE1 | 0.008831043 | 0.252066 | 0.133 | 0.087 | 1 |
| Fibroblasts | USP36 | 0.000327338 | 0.252063 | 0.106 | 0.052 | 1 |
| Terminal | PMEPA1 | 6.59173E-19 | 0.252045 | 0.848 | 0.791 | 1.28374E-14 |
| CD74_Intermediat | SASH1 | 1.43648E-14 | 0.252034 | 0.574 | 0.476 | 2.79755E-10 |
| Intermediate | TNFRSF1B | 2.80669E-06 | 0.251918 | 0.461 | 0.386 | 0.0546603 |
| Contractile | LTBP4 | 1.56868E-21 | 0.251905 | 0.831 | 0.721 | 3.05501E-17 |
| Contractile | MBD3 | 1.25715E-08 | 0.25183 | 0.198 | 0.136 | 0.00024483 |

| | | | | | | |
|---|---|---|---|---|---|---|
| IFN_Intermediate | PXK | 0.005507622 | 0.251724 | 0.271 | 0.169 | 1 |
| Contractile | EXTL3 | 1.09649E-19 | 0.251589 | 0.127 | 0.052 | 2.13541E-15 |
| Terminal | ATOX1 | 3.63503E-17 | 0.251565 | 0.643 | 0.54 | 7.07921E-13 |
| Contractile | CCDC6 | 4.75396E-10 | 0.251551 | 0.288 | 0.209 | 9.25834E-06 |
| IFN_Intermediate | KHNYN | 0.004647821 | 0.251531 | 0.168 | 0.092 | 1 |
| VEGFA_Terminal | SGMS2 | 6.46545E-12 | 0.251518 | 0.184 | 0.084 | 1.25915E-07 |
| Contractile | PRXL2B | 8.40347E-17 | 0.251518 | 0.143 | 0.068 | 1.63658E-12 |
| VEGFA_Terminal | EDNRA | 9.94741E-16 | 0.251413 | 0.174 | 0.066 | 1.93726E-11 |
| Terminal | SERPINH1 | 6.3937E-15 | 0.251395 | 0.679 | 0.607 | 1.24517E-10 |
| Intermediate | PDGFD | 3.13407E-07 | 0.251325 | 0.556 | 0.463 | 0.006103603 |
| VEGFA_Terminal | BMPR2 | 1.23275E-06 | 0.251025 | 0.45 | 0.321 | 0.024007826 |
| IFN_Intermediate | ACAP2 | 0.005969859 | 0.251007 | 0.579 | 0.431 | 1 |
| Contractile | MCCC2 | 8.37585E-16 | 0.250992 | 0.198 | 0.109 | 1.6312E-11 |
| Fibroblasts | ARSA | 0.002421669 | 0.250984 | 0.128 | 0.076 | 1 |
| VEGFA_Terminal | ENSSSCG0( | 8.08424E-29 | 0.250928 | 0.143 | 0.032 | 1.57441E-24 |
| CD74_Intermediat | ENSSSCG0( | 1.53846E-17 | 0.250868 | 0.625 | 0.519 | 2.99615E-13 |
| VEGFA_Terminal | AGFG1 | 1.5387E-09 | 0.250799 | 0.385 | 0.239 | 2.99661E-05 |
| Contractile | ANP32B | 2.78178E-11 | 0.250723 | 0.75 | 0.706 | 5.41751E-07 |
| Contractile | C12ORF75 | 1.36493E-10 | 0.250695 | 0.21 | 0.134 | 2.6582E-06 |
| Contractile | DDX21 | 9.11851E-08 | 0.250578 | 0.492 | 0.435 | 0.00177583 |
| Intermediate | C1ORF198 | 4.95229E-05 | 0.250532 | 0.313 | 0.25 | 0.964458392 |
| Contractile | HNRNPC | 9.12666E-09 | 0.250435 | 0.491 | 0.424 | 0.000177742 |
| Contractile | ENSSSCG0( | 7.71833E-13 | 0.250381 | 0.164 | 0.091 | 1.50314E-08 |
| Contractile | PBRM1 | 1.11369E-08 | 0.250326 | 0.386 | 0.312 | 0.000216892 |
| VEGFA_Terminal | POLR3H | 1.8631E-19 | 0.250241 | 0.186 | 0.065 | 3.62838E-15 |
| Contractile | DHX36 | 4.72875E-09 | 0.250234 | 0.427 | 0.353 | 9.20924E-05 |
| Contractile | USP7 | 4.21757E-08 | 0.250229 | 0.367 | 0.296 | 0.000821372 |
| Terminal | ARPC2 | 1.05433E-21 | 0.250194 | 0.845 | 0.787 | 2.0533E-17 |
| IFN_Intermediate | IGFBP6 | 0.006377935 | 0.250157 | 0.738 | 0.614 | 1 |
| Contractile | SZRD1 | 2.11194E-09 | 0.250154 | 0.271 | 0.198 | 4.11301E-05 |
| Contractile | ANGPT4 | 1.06159E-39 | 0.250129 | 0.112 | 0.024 | 2.06745E-35 |
| Contractile | PINK1 | 1.16602E-08 | 0.250121 | 0.27 | 0.202 | 0.000227083 |
| CD74_Intermediat | CTSC | 2.33279E-29 | 0.380762 | 0.435 | 0.293 | 4.54311E-25 |
| IFN_Intermediate | CNDP2 | 0.007223186 | 0.250004 | 0.28 | 0.182 | 1 |

**Table S7. Marker genes for shared and mesenchymal cell types based on consensus analysis. Corresponds to Figs. 3B and 6C.**

Marker genes were identified in merged scRNA-seq datasets of mouse, pig, and human elastic artery atherosclerotic plaques, using cell labels based on consensus among integrations. Markers with log2FC ≥ 0.6 are shown, sorted by log2FC values with highest values in the top row.

Analysis was performed using the Wilcoxon test in Seurat v4.4.0 with P values adjusted for multiple testing by the Bonferroni method.

| Shared Contractile | | | | | Shared Intermediate | | | | | Shared Terminal | | | | | Shared Proliferating | | | Shared Fibroblasts | | Shared IFN_Intermedi | | Contractile Human-Pig-sp | | Pericytes | | | | DLX5_Cells | VEGFA_Ter | CD74_Inte | RERGL_Cel | Chondromyocytes |
|---|---|---|---|---|---|---|---|---|---|---|---|---|---|---|---|---|---|---|---|---|---|---|---|---|---|---|---|---|---|---|---|---|
| Elastic | | | Coronary | | Elastic | | | Coronary | | Elastic | | | Coronary | | Elastic | | | Coronary | | Coronary | | Elastic | | Elastic | | Coronary | | Carotid | Coronary | Coronary | Coronary | Elastic |
| Mouse | Pig | Human | Pig | Human | Mouse | Pig | Human | Pig | Human | Mouse | Pig | Human | Pig | Human | Mouse | Pig | Human | Pig | Human | Pig | Human | Pig | Human | Pig | Human | Pig | Human | Human | Pig | Pig | Human | Mouse |
| Eln | IL1RL1 | KRT17 | TNNC1 | ACTG2 | Chil1 | CHI3L1 | CCL19 | IL1RL1 | IGFBP2 | Slurp1 | DCN | LUM | SAA3 | COMP | Stmn1 | H2AC4 | IBSP | APOD | PLA2G2A | ISG15 | ISG15 | RGS5 | PLN | CXCL2 | COX4I2 | CXCL2 | CCL21 | DLX6-AS1 | SCGB2A2 | CD74 | RERGL | 3110079O15Rik |
| Igfbp2 | MYH11 | ASPN | MYH11 | ACTC1 | Gpx3 | CA4 | ECRG4 | CCN3 | CCN3 | Ly6a | SAA3 | DCN | NDUFA4L2 | CRTAC1 | Eln | DUT | ELN | CFD | APOD | CXCL10 | CXCL10 | PPP1R14A | MYH11 | RASD1 | THY1 | RGS5 | CD36 | SUCNR1 | IGFBP3 | APOE | MUSTN1 | Ibsp |
| Rgs5 | MYH10 | CCN3 | PPP1R14A | MYH11 | Spp1 | | CCL21 | ITIH4 | TNFRSF11B | Timp1 | ANXA8 | COL3A1 | COMP | PCOLCE2 | Cenpa | ENSSSCG0 | MMP11 | MFAP5 | CFD | GBP1 | ICAM1 | CNN1 | C11orf96 | CCL19 | MUSTN1 | CCL2 | STEAP4 | SOST | IGFBP2 | GPNMB | NET1 | Chad |
| Thbs1 | CCN3 | KRT18 | CNN1 | CNN1 | Saa3 | | SELENOP | MYH10 | VCAN | Ly6c1 | SERPINF1 | APOE | ANXA8 | SERPINE2 | H2afz | ANLN | STMN1 | PI16 | PTGDS | MX1 | MX1 | SPARCL1 | CNN1 | GJA4 | CCDC102B | CCL19 | FABP4 | PTN | IGFBP5 | ENSSSCG0 | PHLDA2 | Alpl |
| Ankrd1 | A2M | FHL1 | ACTG2 | PLN | Bpifb1 | | CXCL12 | IFI6 | FN1 | Col3a1 | COMP | CFH | S100A4 | BGN | Birc5 | RRM2 | S100A10 | C3 | FBLN1 | ENSSSCG0 | RSAD2 | ACTG2 | CSRP2 | ATF3 | RGS16 | GJA4 | FABP5 | LMO2 | VEGFA | A2M | BCAM | Pcolce2 |
| Rgs5 | ACTA2 | TNFRSF12A | MYL9 | TPM2 | Lcn2 | | STEAP4 | ACTA2 | GAS6 | S100a10 | IGFBP6 | PTGDS | IGFBP6 | TNFAIP6 | Cks2 | NUSAP1 | PTN | FMO2 | CXCL14 | IFIT1 | IFI44L | MYL9 | C12orf75 | C7 | STEAP4 | ENSSSCG0 | TCIM | DLX5 | LOXL2 | FGL2 | PTP4A3 | Mia |
| Col18a1 | EFEMP1 | PPP1R14A | CSRP1 | RAMP1 | Mmp3 | | CP | EFEMP1 | COL8A1 | Col1a2 | COL14A1 | COL3A1 | FNDC1 | POSTN | Smc2 | HMGB2 | TK1 | C7 | C3 | MX2 | IFI6 | MYH11 | LMOD1 | BTG2 | NDUFA4L2 | RGS16 | RGS16 | FRZB | ECM1 | ENSSSCG0 | LBH | Col2a1 |
| Mfap5 | FLNA | MYOZ2 | SMTN | C12orf75 | Saa1 | | FGF7 | A2M | AQP1 | Col6a3 | SERPINE2 | SLPI | POSTN | CLU | 2700094K | PCNA.1 | COL1A1 | TXNIP | C7 | RSAD2 | CX3CL1 | ACTA2 | TPM2 | HIGD1B | ABCC9 | ADAMTS4 | PTGER1 | ATP1B1 | PRSS23 | SLA-DQB1 | IGFBP5 | Pth1r |
| Fbln5 | PCGF5 | OGN | TPM2 | PPP1R14A | Mmp13 | | ZFP36 | FRZB | SLC14A1 | Pcolce | COL6A3 | POSTN | COL8A1 | LUM | Top2a | S100A10 | PCNA | IGFBP5 | SERPINF1 | LOC11025 | CCL5 | CSRP1 | FILIP1L | COL5A3 | IGFBP4 | ZFP36 | LINC00924 | PDE5A | BNIP3 | ECRG4 | COX4I2 | Omd |
| Postn | ITGA8 | KRT7 | ACTA2 | SORBS2 | Hp | | CCL2 | NEXN | ASPN | Vcam1 | NDUFA4L2 | COL1A1 | SERPINE2 | COL3A1 | Lockd | PCLAF | EMP3 | MYOC | RARRES1 | ENSSSCG0 | IFI27 | ITIH4 | FLNA | FABP4 | CD36 | BTG2 | ADGRF5 | MECOM | SERPINE1 | GSN | HES4 | Bglap |
| Ctgf | NEXN | CRYAB | DES | MYL9 | Saa2 | | IGKC | CCN1 | MFGE8 | Fmod | TIMP3 | TIMP1 | SOD2 | FXYD5 | Gpx7 | H2AZ2 | TYMS | SELENBP1 | DCN | ISG12(A) | IFIT2 | TNNC1 | PPP1R12B | RGS16 | PGF | ENSSSCG0 | ARID5B | ENPP2 | PLAT | SPON1 | ISYNA1 | Lect1 |
| C4b | VCAN | BDNF | LMOD1 | DSTN | Clu | | IGFBP5 | VCAN | RAMP1 | Dpt | EDIL3 | KDELR3 | COL3A1 | SFRP4 | Col4a2 | ATAD2 | CYTL1 | GPX3 | MGST1 | RIGI | TNFAIP2 | DSTN | MCAM | GJC1 | APOE | NFKBIA | LGI4 | ACTC1 | NMB | SPRY1 | NRGN | Col10a1 |
| Mfap4 | MYLK | ACTA2 | FLNA | DES | Penk | | HSPA1A | THBS1 | KRT8 | Col6a1 | CTHRC1 | ITM2B | S100A10 | TIMP1 | Hmgn2 | CKS2 | COL1A2 | LAMA2 | SFRP2 | EIF2AK2 | IFIT3 | ADIRF | SORBS2 | PGF | PTP4A3 | FABP4 | CNN3 | LMO7 | EGLN3 | ENSSSCG0 | OAZ2 | Col11a1 |
| Col4a1 | SLIT3 | CYFIP2 | RGS5 | ACTA2 | Slpi | | C7 | CP | DKK3 | Col1a1 | S100A10 | ECM1 | PRELP | FN1 | Ccnb2 | ENSSSCG0 | PTTG1 | ABCA9 | HP | OAS2 | XAF1 | TAGLN | KCNMA1 | TINAGL1 | MT2A | FOSB | ATF3 | FGL2 | CA4 | HLA-DRA | GUCY1A1 | Cst3 |
| Nov | SMTN | CCN2 | DSTN | CSRP1 | Cp | | FMO2 | PENK | OGN | Col2a1 | LGALS3 | CRTAC1 | SMPDL3B | CTHRC1 | Cenpf | TPX2 | TUBA1B | NID1 | MYOC | VCAM1 | LY6E | TPM2 | MYL9 | FHL5 | HIGD1B | JUNB | TINAGL1 | PLK2 | PTGES | TIMP3 | KCNAB1 | Hapln1 |
| Itih4 | SYTL2 | INHBA | MYLK | FLNA | Steap4 | | CFH | MFAP4 | MYH10 | Cthrc1 | ECM1 | RCN3 | PRSS35 | GAP43 | Col4a1 | PTTG1 | NUDT1 | ENSSSCG0 | MFAP5 | ENSSSCG0 | IFIT1 | GLDN | MYLK | RGS5 | HLA-B | SOCS3 | ADAMTS4 | NBL1 | ERO1A | B2M.1 | ARPC1A | Comp |
| Prss23 | VCL | SERPINE1 | SORBS1 | LDOC1 | Comp | | FOS | TPM1 | TGM2 | Mmp2 | LUM | EMP1 | LCN2 | CD9 | Rrm2 | PRC1 | SH3BGRL3 | CRISPLD2 | ITM2A | EPSTI1 | MX2 | NEXN | ACTC1 | CYGB | SEPTIN4 | C7 | APOE | ID1 | TSC22D1 | C1S | UBA2 | Cytl1 |
| Loxl1 | MYL9 | EFEMP1 | ITGA8 | HSPB1 | Prg4 | | NR2F2 | FABP4 | PRSS23 | Col5a2 | SMPDL3B | PLAC9 | CRYAB | PRELP | Racgap1 | CENPE | HELLS | IGF2 | FBLN2 | GHSR | OASL | FHL1 | RERGL | ADAMTS1 | IGKC | FOS | GGT5 | TSPAN13 | SLC16A3 | RAMP2 | SORBS2 | Scrg1 |
| Fgl2 | CNN1 | NEXN | C6H19ORF | SBSPON | Srgn | | VCAM1 | RCAN2 | AEBP1 | Tgfbi | S100A1 | PLTP | EDIL3 | VCAN | Hmgb2 | SMC4 | COL15A1 | SLC6A4 | SERPINA3 | STAT1 | IGHM | DES | LDOC1 | FRMD3 | PHLDA1 | HIGD1B | ARID5A | CAVIN2 | GAPDH | TMEM245 | SERPINI1 | Bglap2 |
| 2210407C | MAP1B | ACKR4 | SYNPO2 | HSPB7 | C3 | | C1R | THBS2 | SPINT2 | Cst6 | ALDH2 | TYMP | SH3BGRL3 | NPC2 | 2810417H | TUBB | COL6A1 | OGN | PLIN2 | IFI6 | PLSCR1 | HSPB7 | FILIP1 | GUCY1B1 | TBX2 | ITIH4 | ID4 | AK1 | SPON2 | CTSH | TOB1 | Col9a3 |
| Crispld2 | PPP1R14A | CRIM1 | MYOZ2 | PPP1R12B | Wfdc21 | | FABP4 | SLIT3 | S100A4 | Ndufa4l2 | ABI3BP | PPIB | ASPN | PRSS23 | Mki67 | CCL2 | ARPC1B | DCN | PI16 | PARP14 | OAS1 | SMTN | LPP | ZFP36 | LHFPL6 | TINAGL1 | SSTR2 | SAMD11 | DDIT4 | FBLN1 | PRPH | Rarres1 |
| Ltbp1 | MYH9 | WFDC1 | ITIH4 | CSRP2 | Tsc22d1 | | HLA-B | CCN2 | SUGCT | Npc2 | POSTN | COL1A2 | FMOD | COL1A1 | Tubb5 | ZNF367 | LGALS1 | ENSSSCG0 | GSN | PLAC8 | IRF1 | MYLK | SUSD5 | ITIH5 | RGS5 | RASD1 | INPP4B | THSD4 | CCL2 | FBLN2 | NDUFA4L2 | Sparc |
| Sncg | PALLD | CCDC144N | SPARCL1 | PALLD | Fth1 | | SFRP2 | LOC10051 | ELN | Serpinf1 | PCOLCE | CLEC11A | COL1A1 | EMP3 | Cks1b | ANXA2 | PCLAF | ABCA6 | HTRA3 | UBE2L6 | ZC3HAV1 | LMOD1 | SYNPO2 | SOCS3 | GJA4 | NOTCH3 | ADAMTS1 | HMCN1 | CSRP2 | GPX3 | CSRP2 | Smpd3 |
| Fstl1 | LMOD1 | TNFRSF11B | FBXO32 | PDLIM3 | Dcn | | IGLC2 | COL18A1 | FRZB | Prg4 | FNDC1 | CRABP2 | TNC | COL1A2 | Col5a1 | TK1 | PLAU | MARCKS | C1R | HERC6 | APOL1 | FILIP1L | DSTN | FAM162B | ZFP36 | MYC | ARHGAP15 | BST2 | IER3 | C1R | RBPMS2 | Col27a1 |
| Wisp2 | ACTN4 | FSTL3 | ADIRF | CCDC3 | Lrg1 | | IFITM2 | ITGA8 | BGN | Col6a2 | PI16 | MXRA5 | CDO1 | DKK3 | Tmsb10 | TOP2A | PHLDA2 | HSD11B1 | SRPX | LGALS3BP | EPSTI1 | EFHD1 | MFAP4 | SEPTIN4 | ARHGAP29 | FAM162B | TMEM176 | GREM2 | ENO1 | PLAC9 | SPARCL1 | Ccdc80 |
| Mgst1 | PDLIM7 | FBLN5 | SORBS2 | SLC25A4 | Mt2 | | NPC2 | MEGF6 | LTBP1 | Nbl1 | PCOLCE2 | COL6A2 | CD9 | LTBP2 | Col1a1 | SPC24 | SLC25A5 | CLEC3B | GPX3 | LOC10051 | USP18 | MYL6 | TNS1 | MYO1B | SOCS3 | GADD45B | CCL8 | PRDM6 | UPP1 | SERPINF1 | PDK4 | Dhx58os |
| Lox | ACTN1 | CREB5 | TAGLN | FABP3 | Cxcl14 | | HLA-A | TNFRSF11B | LTBP2 | Sfrp4 | TFPI2 | PMP22 | MYBL1 | SH3BGRL3 | Ccnb1 | COL6A3 | MCM7 | SPTBN1 | PTN | IFITM1 | GBP4 | FBXO32 | ITGA8 | PTP4A3 | GUCY1B1 | LOC10052 | TM4SF1 | RSPO2 | POSTN | B2M | NUDT4 | Crispld1 |
| Col4a2 | COL15A1 | PXDC1 | PDLIM3 | LMOD1 | Serpine2 | | C1S | TAGLN | PPP1R14A | Fn1 | SEMA3C | UNC5B | VMP1 | FMOD | Cenpe | STMN1 | ITGA5 | GPC3 | C1S | ZNFX1 | CSF1 | LMCD1 | MTRNR2L12 | APOD | FABP5 | EDNRB | FGF7 | RAMP1 | ADM | FBLN5 | TBX2-AS1 | Ucma |
| Actg2 | PRKG1 | SUGCT | PPP1R12B | MYH10 | Lum | | NNMT | FILIP1L | ITGA8 | Ctsk | PRSS35 | TNFSF13B | FN1 | GAS6 | Ube2c | H2AC16 | H2AFV | GHSR | IGF1 | ENSSSCG0 | APOL3 | SYNPO2 | SLMAP | FOS | JUN | ATF3 | TGFBI | DLX6 | SLC38A1 | ABCA1 | TSC22D1 | A230065H16Rik |
| Lmod1 | LOC10051 | FSTL1 | PAWR | ALDH1B1 | Ier2 | | TFPI | IGFBP7 | SGCA | Anxa8 | CLU | TFPI | ENSSSCG0 | MGP | Col3a1 | ENSSSCG0 | KITLG | FBLN1 | PLTP | HERC5 | IRF7 | FLNA | HSPB7 | NR4A1 | PLXDC1 | CYCS | SYTL2 | CDO1 | LDHA | ENSSSCG0 | CPM | Cdkn1c |
| Crlf1 | CALD1 | ELN | CALD1 | CARMN | Cst3 | | CALCRL | SOD3 | THBS2 | Ltbp2 | SOD2 | FKBP11 | SPARC | LTBP1 | Tubb6 | SMC2 | CENPK | SOCS3 | TAC1 | PSMB9 | TFPI2 | IGF2 | GRIA2 | NR2F2 | NOTCH3 | RRAD | ECEL1 | IGFBP3 | NDUFA4L2 | IFI30 | KLHL23 | Wif1 |
| Serpinb6b | PKDCC | MYH10 | PGM5 | WFDC1 | Gadd45a | | MARCKS | RASL11A | COL21A1 | Igfbp4 | COL11A1 | CYP1B1 | ECM1 | SPARC | Col6a3 | HMMR | GINS2 | CXCL12 | LUM | PSMB8 | TNFSF13B | FIBIN | CSRP1 | JUN | CPM | IL6 | ESAM | OGN | PHLDA2 | ALDH1A2 | BTG1 | Acan |
| Itga8 | MGLL | CCN1 | CKM | NEXN | Junb | | SOCS3 | MYH9 | FMOD | Fxyd5 | SFRP2 | SMOC1 | MT1X | S100A10 | Prc1 | CAPG | RFC3 | PLSCR4 | CCDC80 | B2M.1 | STAT1 | FRZB | ACTG2 | ARPC1A | OR51E1 | NR4A1 | MYC | GPC3 | NDRG1 | NUDT4 | AC079822 | Col11a2 |
| Cald1 | PCDH7 | MFAP4 | MYL6 | ACTN1 | Trf | | LY6E | PDLIM7 | FBXO32 | Sdc1 | OMD | HLA-B | LUM | TNFRSF11B | Lgals1 | MCM4 | COL3A1 | HTRA3 | FBLN5 | IFIH1 | IFI35 | LOC10051 | PPP1R12A | FOSB | HES4 | NR2F2 | TSPAN12 | PTGIS | TPI1 | FOSB | MTHFD2 | Col9a1 |
| C1s1 | TNC | TGFBR1 | KCNMB1 | RGS5 | Serpina3n | | CCN1 | CALD1 | CRIP2 | Lum | ASPN | STEAP1 | DDIT4 | ISLR | Lmna | TYMS | SMC4 | PCOLCE2 | MMP2 | SAMHD1 | REL | SEMA3E | PALLD | CCL2 | SLCO1C1 | IER3 | ACTB | RGS7BP | SLC2A3 | PLXDC1 | PPP1R12A | Myoc |
| Fblim1 | ITIH4 | TCEAL4 | CPE | CCN3 | Gas1 | | ANGPT2 | RARRES2 | GMDS | Capg | LOXL2 | RRBP1 | SEMA3C | DPT | Tuba1b | LOC11025 | H2AFZ | APOE | GPC3 | IFI44L | ADPRHL2 | C6H19ORF | FIBIN | AVPR1A | TPPP3 | PPP1R15A | TMEM176 | CCDC68 | FGFR2 | LGALS3BP | MAP3K20 | Frzb |
| Tpm4 | MACF1 | FN1 | PLN | ITGA8 | Lbp | | RARRES2 | PLK2 | CD151 | Igfbp6 | COL6A1 | LTC4S | NBL1 | FGL2 | Hmmr | ECM1 | ZWINT | ANGPTL4 | OLFML3 | TAP1 | HLA-A | PDLIM7 | SMTN | EPS8 | LINGO1 | IFRD1 | REM1 | GEM | P4HA1 | C2 | NTN4 | Chil1 |
| Igfbp7 | CD74 | RAB32 | RASSF3 | CALD1 | Scara3 | | JUNB | PRKG1 | KRT7 | 1500015O | NBL1 | CD9 | GALNT15 | HAPLN1 | Cdkn3 | CENPF | PLP2 | FOS | DPT | ENSSSCG0 | IL15RA | SORBS1 | CARMN | SLC2A3 | NEURL1B | ADAMTS1 | CCL19 | LINC02544 | ATF4 | CTSB | PGF | Col9a2 |
| F2r | CSRP1 | ACTN4 | CAMK2G | KCNMB1 | Phlda1 | | EGR1 | TTLL7 | EDIL3 | Anxa2 | IGFBP5 | NUCB2 | HTRA1 | TSC22D3 | Cdca3 | LMNA | COL5A1 | LTBP4 | PODN | RTP4 | VSIR | NET1 | SBSPON | NRP1 | ENPEP | IRF1 | A2M | IER2 | PKIB | SLPI | AKAP1 | Hp |
| Htra1 | ADGRE5 | CNN1 | MCAM | SPINT2 | 1500015O10Rik | | SPRY1 | SYTL2 | MAOB | Mmp14 | MT1X | FKBP10 | RGCC | S100A4 | Kif22 | CENPU | RPA3 | DPT | EMP1 | ENSSSCG0 | UBE2L6 | CALD1 | SVIL | ADAMTS4 | ADAP2 | JUN | CD44 | ID3 | CSTB | ETV1 | MCAM | Fmod |
| Angpt2 | NIBAN1 | PRSS23 | IGF2 | MYL6 | Mafb | | CEBPD | SOST | C5orf46 | Nupr1 | BHLHE41 | HSP90B1 | COL1A2 | F2R | Ccl9 | TUBB2A | SLC14A1 | CHRDL1 | PTX3 | LGALS9 | SOD2 | MCAM | PRUNE2 | BTG3 | EDNRB | ITGA7 | FAM162B | MSX2 | ENO2 | PLPP3 | EDNRA | Panx3 |
| Tgfb2 | SMOC2 | ACTG2 | PDLIM7 | MYLK | Klf4 | | BST2 | VCL | CRIP1 | Sparc | FN1 | B2M | S100A1 | FAP | Adamts2 | PLP2 | S100A11 | FMO1 | VIT | SLA-DQB1 | DAB2 | RASSF3 | SCRG1 | ITIH3 | NCKAP5 | CEBPB | EIF4A1 | CDKN1C | ENSSSCG0 | SAT1 | CASQ2 | Fzd9 |
| Cyr61 | HMCN1 | COL8A1 | VCL | TNS1 | Gdf10 | | NTRK2 | TGM2 | TSC22D3 | Timp3 | APOD | PPIC | FAP | COL8A1 | Knstrn | TNFRSF12A | PSMB2 | FBLN2 | ABCA8 | SETX | PLAAT4 | PLN | RTN1 | LOC10052 | FABP4 | ITIH5 | CTSC | INMT | NFKBIA | SELENBP1 | TINAGL1 | Mepe |
| Fbn1 | DPYSL3 | PDLIM7 | SLC22A3 | RASSF3 | Cdkn1a | | PLAAT4 | MYL9 | CDH13 | Gm17455 | LCN2 | ISLR | LGALS3 | PLXDC2 | Clic1 | SERPINE2 | CD59 | C1S | UAP1 | WARS1 | CMPK2 | LPP | SORBS1 | GUCY1A1 | C20orf27 | EGR1 | DDX5 | OMD | COL6A3 | SFRP1 | ADIRF | Srpx |
| Cnn2 | ITGA7 | GEM | TNS1 | SPEG | Fosb | | FOSB | HSPB1 | RILPL2 | Ppic | COL1A2 | CD59 | CLU | S100A16 | Fstl1 | UBE2T | SLC16A3 | ENSSSCG0 | PCOLCE | STAT2 | MARCKS | RHOB | CALD1 | NOTCH3 | PLAC9 | GUCY1B1 | PLPPR4 | DAPL1 | LGALS3 | PDGFD | SNCG | Fap |
| Cdh13 | VLDLR | FLNA | LOC10051 | HACD1 | | | ANGPT1 | ENSSSCG0 | ITGA10 | S100a4 | CDO1 | NNMT | NT5E | SULF1 | Spc24 | CKAP2 | GPX7 | ENSSSCG0 | SOD2 | SAMD9 | PARP9 | KANK1 | PCDH7 | KLHL23 | AC079822.1 | SEPTIN4 | ACTG1 | KLF2 | KIT | TNXB | EPAS1 | Pltp |
| Fat1 | COL4A2 | VCAN | NIBAN1 | TAGLN | | | JUN | MYH11 | ADI1 | Lgals3 | NT5E | LRRC17 | EMP3 | RGS2 | Cdc20 | UBALD2 | PSME2 | F3 | FMO2 | SP140 | EIF2AK2 | ITGA7 | TAGLN | ENSSSCG0 | CRISPLD2 | PKIG | TGFB3 | IGFBP5 | ENSSSCG0 | COL14A1 | SEPTIN4 | Mfge8 |
| Lamb2 | L1CAM | WTIP | KANK1 | SH3BGR | | | CTSC | PCDH7 | CTSD | Prelp | SPP1 | PTHLH | TRPS1 | IL32 | Ckap4 | KIF23 | CSF1 | RNASE4 | ADH1B | ENSSSCG0 | JCHAIN | ISG12(A) | NEXN | APOLD1 | GUCY1A2 | ZFP36L1 | BATF3 | DPT | FAM13A | CCN3 | MEF2B | Sox9 |
| Cd74 | COL18A1 | CRLF1 | EFHD1 | SLMAP | | | CD74 | CHI3L1 | WFDC1 | Col15a1 | ITGBL1 | HLA-C | MEOX1 | NUCB2 | Rangap1 | ENSSSCG0 | HMGB3 | ZFP36 | SCARA5 | PPA1 | DNAJA1 | CPE | DES | MYC | AC100803.4 | ID4 | FILIP1L | ECRG4 | PTPRR | ISLR | ID1 | Cd24a |
| Cxcl12 | PALM2AKA | TPM2 | FBLIM1 | VIM | | | SLC38A2 | TPM4 | MGP | Col14a1 | MMP2 | SPARC | COL6A3 | CFH | Tpx2 | TPM3 | THY1 | ADGRD1 | CES1 | IFIT5 | TNFSF10 | SH3BGR | RAMP1 | PDGFRB | TESC | ISYNA1 | SOCS3 | TPD52L1 | RGS4 | MAMDC2 | ENTPD3 | Itm2b |
| Ace | KANK1 | ADH1B | RBPMS | SMTN | | | IER2 | ENSSSCG0 | CTHRC1 | Timp2 | CD9 | TNXB | EMP1 | AEBP1 | Anln | KIF20A | NES | CD302 | TNXB | CD74 | EFEMP1 | SORBS2 | TCEAL2 | ENSSSCG0 | GMFG | FRMD3 | CXCL12 | OSR1 | TIMP1 | LAMC3 | CHCHD10 | Pdzrn4 |
| Aebp1 | ADARB1 | DNAJB4 | PPP1R12A | NDUFA4 | | | HLA-DRA | NET1 | EPDR1 | Emp3 | SFRP1 | SSR4 | ENO1 | IGFBP6 | Ckap2 | ENSSSCG0 | CLSPN | C4B | BICC1 | LOC10051 | ANGPTL7 | LIMCH1 | TTLL7 | CYCS | EPHX1 | KLHL23 | TMEM200 | LMO3 | ANPEP | C7 | TBX2 | Tspan13 |
| CR974586 | GJA5 | NDUFA4 | LMCD1 | PDLIM7 | | | RGS16 | CCL19 | KRT18 | Fbln2 | EMP1 | HLA-A | ANXA2 | SERPINE1 | Ran | TAGLN2 | GGH | CYBRD1 | IGFBP4 | C1S | PSMB9 | FILIP1 | MYOCD | RHOB | JUNB | GUCY1A1 | ANGPT2 | MBNL1-AS | GDF15 | ADD3 | MYH11 | Ppa1 |
| Pde3a | COL4A1 | MBNL1-AS | RASL12 | NIBAN1 | | | HSPA1B | C4B | PDLIM3 | Dcn | HAPLN1 | COL5A2 | TXN | VCAM1 | Notch3 | PBK | MMP14 | OMD | G0S2 | ENSSSCG0 | OLFML3 | TTLL7 | MAP3K20 | ENSSSCG0 | ADGRF5 | GJC1 | TPPP3 | LMCD1 | SULF1 | | PTK2 | Serpinh1 |
| Flna | DMD | MMP23B | CALML4 | SYNPO2 | | | DEPP1 | MYL6 | HSPB7 | Cd34 | VMP1 | TAGLN2 | COL5A2 | KDELR3 | Anxa2 | DLGAP5 | CENPH | ADD3 | NNMT | IRF7 | TLK1 | TGM2 | NPNT | ENSSSCG0 | PHLDA2 | COL4A1 | PHLDA1 | FOS | METRNL | | CDH6 | Chadl |
| Myh10 | AEBP1 | MYL12A | TPM1 | TPM1 | | | | COL4A1 | SNRPN | Oaf | CYP1B1 | PHLDA2 | EZR | ABI3BP | Cenpw | MAL | CKAP2 | ITGBL1 | IGFBP6 | B2M | IFI44 | ADGRE5 | RGS5 | EBF1 | CDH6 | LOC10012 | FHL5 | IL11RA | CRLF1 | | FABP4 | Haus8 |
| Tmsb10 | LPP | KRT8 | NEXN | TCEAL2 | | | | COL4A2 | IGFBP7 | Emp1 | PDGFRA | NDUFA4L2 | COL5A1 | COL15A1 | Lamb2 | H1-3 | ANLN | PDGFRA | TNFSF13B | DDX60 | SHMT2 | S100A2 | ACTA2 | PDGFA | NRARP | PEAR1 | TPM4 | BEX2 | CXCL2 | | MYL9 | Phlda1 |
| Tmsb4x | TPM2 | TAGLN | RHOB | LPP | | | | PCGF5 | LINC01615 | Sdc2 | BNIP3 | MMP2 | CTHRC1 | ITGA10 | Ptprs | KIF11 | TMEM106 | JUNB | ABCA6 | HLA-DRA | PRCP | RCAN2 | HSPB6 | NFKBIA | LPL | SLC2A3 | PARM1 | NDNF | SOX9 | | CAV2 | Pdgfrl |
| Ctsl | MCAM | FILIP1L | SVIL | ILK | | | | ARID5B | KRT17 | Ecm1 | CRLF1 | AKR1C2 | BNIP3 | RCN3 | Cdk1 | MAD2L1 | DEK | SVEP1 | CYBRD1 | CMPK2 | ODF2L | PDZRN4 | ADRA2C | ADAP2 | FAM162B | IL13RA1 | APOLD1 | RGS17 | SLC20A1 | | ID3 | Gpx3 |
| Rarres2 | TTLL7 | BAG2 | MGLL | MCAM | | | | GAS6 | SPARC | Gda | HSD11B1 | CCL19 | CST3 | CDH19 | Calm3 | TUBA1B | DTYMK | MATN2 | ID2 | FUNDC1 | PSME2 | KCNAB1 | TLN1 | CAMK2N1 | LGI4 | KLF2 | COLEC11 | NRGN | PGAM1 | | ROCK1 | Srpx2 |
| Psap | SYCP2 | TPH1 | ADAMTS1 | MAP3K7CL | | | | ACTN1 | CDH11 | Aqp1 | TRPS1 | LAMB1 | ENSSSCG0 | CAPG | Ube2s | NUDC | TAGLN2 | TNXB | PDGFRL | PSME1 | PARP12 | CFL2 | DACT3 | EDNRB | NR2F2 | GADD45A | BMP2 | CXXC5 | CTSH | | NDRG2 | Dnajb1 |
| Fbln2 | SYNPO2 | | S100A2 | SELENOW | | | | ID4 | MFAP4 | Acan | RFLNB | MMP23B | TPPP3 | FKBP10 | Col12a1 | CLIC1.1 | E2F1 | SERPING1 | SELENOP | XAF1 | SAMD9L | CAVIN3 | PART1 | DEPP1 | BTG1 | ARPC1A | NOTCH3 | PHYHIPL | TNC | | NRIP2 | Trpv4 |
| Vcl | HSPG2 | | LPP | SVIL | | | | CDC42EP3 | EFEMP1 | Ppp1r14b | PRELP | PROCR | FOLR3 | OMD | Hs6st2 | ASPM | CCPG1 | MATN4 | ADD3 | IFI44 | DDX58 | LIMS2 | NET1 | SLC7A2 | FXYD6 | TXNIP | DEGS1 | BTG2 | HK1 | | FILIP1L | Fam104a |
| Itgb1 | NOTCH3 | | NPNT | PGAM2 | | | | LMCD1 | HSPB1 | Col11a2 | OGN | IL34 | CLEC11A | AGT | Pgf | ITGA6 | PPIH | PLPP3 | CFH | ZC3HAV1 | SAMD9 | PAWR | SPEG | SPRY2 | GJC1 | ANGPTL4 | ZFP36 | PRRX2 | ID2 | | GJA4 | Sgms2 |
| Col8a1 | ITGA1 | | FERMT2 | AOPEP | | | | | FBLIM1 | AA467197 | CAPG | CDH11 | SELENOM | CRABP2 | Cenpm | EMILIN2 | DYNLT1 | ALDH1A1 | SFRP1 | LAP3 | IL6ST | LRRFIP1 | PPP1R14A | PHLDA2 | TDO2 | RND3 | AVPR1A | ARID5B | HK2 | | CARMN | Tubb2a |
| Ltbp3 | CRIM1 | | KCNAB1 | CRYAB | | | | | IFITM10 | Anxa1 | SELENOM | CFB | RFLNB | ANXA2 | Cdca8 | SLBP | TMEM50A | GFPT2 | DHRS3 | TENT5A | TLR2 | FABP4 | MYH10 | GTF2E2 | RCSD1 | NEURL1B | HLA-B | ENG | ALDOA | | SYNE2 | Car8 |
| Nexn | PAWR | | PDZRN4 | FBLIM1 | | | | | CPXM2 | Serping1 | RGCC | KDELR2 | LDHA | ANKH | Dctpp1 | RNH1 | COLGALT1 | ACKR4 | MT1X | LOC10051 | AGTRAP | IL17B | FHL5 | RCSD1 | ATP1B2 | MYO1B | EPS8 | NPTX2 | PKM | | CCDC3 | Cox4i2 |
| Hspg2 | LAMB1 | | FLNC | MOB2 | | | | | ANTXR1 | Nme1 | ANXA2 | LAMA2 | DCN | CST6 | Cd44 | GAPDH | TPST2 | PTN | ZFP36L2 | SNX10 | NNMT | IRAG1 | SCX | ANO3 | CLMN | ITIH3 | SUCNR1 | MT1X | ATF3 | | MGST3 | 1500015O10Rik |
| Pbxip1 | ITGB1 | | NET1 | GAPDH | | | | | LIMCH1 | Matn2 | MAL | HTRA1 | KDELR3 | ITGBL1 | Lox | DNMT1 | TOP2A | BTG2 | LRP1 | SLFN11 | ZNF207 | NDUFB1 | ALDH1B1 | IFRD1 | PTMA | UTRN | MYL12A | CCN2 | EMP1 | | GUCY1B1 | Znhit6 |
| Pcdh7 | PDLIM5 | | RAMP1 | CASQ2 | | | | | MYH9 | Aspn | SOX9 | DPP7 | SDCBP | CHPF | Banf1 | TUBB6 | UQCC2 | CPXM2 | LEPR | SOD2 | CFI | VCL | MTRNR2L8 | JUNB | ANKRD29 | CYGB | ACTN4 | EFEMP1 | TXN | | HEXIM1 | Cgref1 |
| Pi15 | WIF1 | | FHL1 | RASL12 | | | | | FOXS1 | Lman1 | CCL2 | ARPC1B | HOPX | TAGLN2 | Kif20a | CDKN3 | PGP | ENSSSCG0 | NR4A1 | FAM111A | IRF3 | ID4 | CPED1 | EPHX3 | KCNE4 | COL5A3 | IGFBP5 | SNCG | C4ORF3 | | ITGA7 | AW112010 |
| Sulf1 | EFHD1 | | ENSSSCG0 | CFL2 | | | | | CCN5 | Crabp2 | IL18 | EMP3 | NMB | FIBIN | Ranbp1 | PPP1R14B | SLC29A1 | CEBPD | CP | CASP5 | ABHD5 | TPM1 | MXI1 | ENSSSCG0 | APOLD1 | COL18A1 | CAV1 | JUNB | PGF | | HRC | Scin |
| Mustn1 | TLN1 | | SPINT2 | ADIRF | | | | | RHOD | Ftl1 | CLEC11A | HLA-E | PALMD | TSPO | Fbn1 | SMPDL3B | PPDPF | TGFBR3 | CCL2 | SAT1 | RAD50 | TNS1 | NCS1 | RELL1 | IFITM3 | CCNL1 | H19 | GDF10 | P4HA2 | | PCSK7 | Sdf4 |
| Adam9 | GAS6 | | TGM2 | SNCG | | | | | C12orf75 | Gsn | PTGES | CTSZ | CA4 | NDUFA4L2 | Pbk | DIAPH3 | CBX5 | EGR1 | CYGB | NUB1 | TSPAN4 | RBPMS | SLC25A4 | ABCC9 | CAMK2N1 | ICAM1 | HES1 | KAZALD1 | CTSC | | ZFHX3 | Maged2 |
| Gpc6 | AKAP12 | | TLN1 | KCNA5 | | | | | ITGB1 | | GALNT15 | FAM114A1 | CAPG | TMSB10 | S100a10 | MYBL1 | TXNDC17 | ITIH5 | CST3 | TRIM25 | RSPO3 | FAM13C | CAMK2G | STOM | TMEM176B | TCF21 | H3F3B | SYNGR1 | ENSSSCG0000002549 | | GPRC5C | Slc38a3 |
| H2-K1 | IRAG1 | | IRAG1 | MSRB3 | | | | | SFRP4 | | CTSK | FBN1 | STK38L | CD55 | Kif20b | POLR2L | GAPDH | SLPI | CD34 | TAP2 | TICAM2 | TMEM70 | RASSF3 | TRA2B | HLA-C | IER2 | STOM | KRT19 | PGK1 | | NTRK2 | Ogfrl1 |
| Mcam | SORBS1 | | GYG1 | HAND2 | | | | | FGF1 | | ENSSSCG0 | CERCAM | C1QTNF5 | TFPI2 | Cdca5 | LMNB1 | PON2 | PDPN | NAMPT | CNP | GLDN | FGL2 | HSPB8 | CXCL14 | AGT | LOC10052 | CCL2 | BEX3 | PRDX6 | | ESAM | Ssr4 |
| Spint2 | COL13A1 | | TGFB1I1 | RCAN2 | | | | | ALCAM | | CTSH | COL5A1 | TSPO | TGM2 | Fbln2 | BHLHE41 | PRC1 | FBP1 | MEG3 | GBP2 | HIC1 | PHLDB2 | ACTN1 | DES | H3F3B | NRP1 | TPM1 | MAB21L2 | VMP1 | | FILIP1 | Rgcc |
| Sox4 | AIF1L | | CAP1 | CRIP2 | | | | | GYPC | | TSPO | THBS2 | CCND3 | IGFBP2 | Anp32e | MCM7 | FNDC1 | UACA | AKR1C1 | PSME2 | COL10A1 | ZYX | ADSSL1 | GADD45A | SLC7A2 | RASL11A | CNGA1 | SLIT3 | PLOD1 | | PACSIN2 | Serinc5 |
| Aoc3 | FILIP1 | | RCAN2 | MAOB | | | | | HCFC1R1 | | CCDC80 | CALU | TMSB10 | MEG3 | Smc4 | ALYREF | PDCD1LG2 | FBLN5 | ABCA9 | C1R | CMTM3 | RASL11A | FOXN3 | PPP1R15A | ARHGAP42 | ABCC9 | PTK2 | TAGLN2 | F5 | | DSTN | Vkorc1 |
| Sulf2 | C1ORF198 | | SLC25A4 | TSTD1 | | | | | TSTD1 | | ANKH | FGF7 | S100A6 | TMSB4X | Fndc1 | PSMA7 | LASP1 | MTR | ADM | IFI30 | C6orf62 | ITGA8 | NIBAN1 | GNG2 | IFITM2 | ANO3 | TAGLN | KCNK17 | VDAC1 | | RCAN2 | Susd5 |
| Lama5 | MSRB3 | | SH3BGR | SELENOM | | | | | ITGA1 | | GAPDH | MRC2 | ENSSSCG0 | ECM1 | Ass1 | CENPH | RAD51AP1 | TGFBR2 | F3 | PARP12 | PML | KCNMB1 | RNF115 | ZFP36L1 | ADGRA2 | IER5 | MT1A | S100A13 | MTURN | | MYLK | Enho |
| Actn1 | CD93 | | ITGA5 | CAMK2G | | | | | ECM2 | | ACKR3 | PDIA3 | BHLHE41 | AKR1C2 | Megf8 | TXN | BIRC5 | CDKN1C | MT2A | LOC10651 | PLEKHA4 | MSRB3 | PRKG1 | EDNRA | CD248 | IL17B | IGFBP4 | HAND2 | KITLG | | DMPK | Clec11a |
| Rhob | HSPB1 | | HSPB7 | SLC22A3 | | | | | PLS3 | | S100A4 | CXCL12 | CGREF1 | S100A13 | Khdrbs1 | HTRA3 | LIG1 | EBF1 | CEBPD | IFIT3 | IFIH1 | TNC | CPE | KCNE4 | TBX2-AS1 | ENSSSCG0 | LHFPL6 | SOD3 | COL4A1 | | MRGPRF | Tsc22d3 |
| Sod3 | ENSSSCG0000000523 | | LRPAP1 | FBXO32 | | | | | SVIL | | ENO1 | PRDX4 | CD44 | KRT17 | Loxl2 | LDHA | ATP1B3 | CPED1 | PMP22 | C2 | STEAP1 | PALLD | DMD | CDKN1C | IFITM1 | ENSSSCG0 | JUNB | CLIC3 | SARS1 | | FBXL13 | Bambi |
| Lpp | LIMCH1 | | PDE4DIP | SH3BGRL | | | | | FXYD5 | | GALNT1 | TSPAN4 | RNH1 | DPYSL3 | Hn1 | PPIA | ACAN | MEDAG | SERPING1 | USP18 | MMP28 | MYOM1 | PDLIM5 | RASL11A | ANGPT2 | SRSF7 | CLSTN2 | KCTD12 | TMEM45A | | WTIP | Steap4 |
| Cnn1 | FRZB | | ENSSSCG0 | MRPS6 | | | | | PTTG1IP | | KDELR3 | TPM3 | IL18 | PKM | Nucks1 | MCM6 | AP1S2 | PDK4 | MARCKSL1 | | SNX33 | SPINT2 | FBXO32 | ITIH4 | ARPC5 | OLFML2A | EDNRB | CXCL12 | COL6A1 | | PTMA | Azgp1 |
| Fbln1 | FBLIM1 | | LIMCH1 | RNF115 | | | | | GADD45A | | COL1A1 | FAM3C | RRBP1 | MMP2 | Tyms | UBE2C | TP53I11 | CAT | RNASE4 | | CTSK | | HIST1H4C | PEAR1 | FRMD3 | CPM | DNAJB4 | LGALS3 | GARS1 | | ENPEP | Fgfr3 |
| Myh11 | LYNX1 | | PALLD | TCEAL4 | | | | | COL4A2 | | FBLN7 | OMD | PHLDA2 | RGS3 | Plp2 | C1QB | SMC2 | F2RL2 | PIM1 | | ITSN1 | | ANGPTL1 | STUM | EDNRA | KCNA5 | AGT | GMDS | PRDX4 | | MAFB | Cdkn1a |
| Gas6 | LSAMP | | MXI1 | FRZB | | | | | FHL1 | | SLC20A1 | SERPINH1 | NDFIP1 | HTRA1 | Mmp14 | CENPW | CNIH1 | TCF4 | CHI3L2 | | COL1A2 | | SPINT2 | LRRC10B | AC100793.2 | EBF1 | COX4I2 | LNP1 | ADAMTSL4 | | DES | Ppp1r1b |
| Ltbp4 | TGFB3 | | ITGA7 | CCDC107 | | | | | MRPS6 | | NPC2 | MDK | SHISA4 | MXRA8 | Ccdc34 | BNIP3 | SERPINE1 | TCF21 | MT1M | | GBP2 | | CFL2 | EGR1 | SCIN | FHL5 | PROCR | NFKBIA | THBS1 | | NOTCH3 | Vit |
| Cbr2 | TNS1 | | LRRFIP1 | CRIP1 | | | | | TMSB4X | | SULF1 | MYDGF | ID1 | MMP23B | Sdc1 | HOMER3 | DPP7 | DHRS3 | QSOX1 | | SNHG32 | | MRPS6 | OLFML2A | TINAGL1 | ENSSSCG0 | TMEM119 | CDKN2C | STBD1 | | MAP3K7CL | Sp7 |
| Pdgfrb | CAP1 | | MSRB3 | CHURC1 | | | | | MRAP2 | | RCAN1 | PDGFRA | PCOLCE2 | AQP1 | Timp1 | EZR | CLIC1 | ANG | ZFP36 | | SLC16A7 | | ST6GAL2 | P3H2 | TYMP | ARL5B | GJA4 | | INHBA | | COX7A2 | Bglap3 |
| Prss12 | COL8A1 | | DMD | GADD45G | | | | | SYNGR2 | | CGREF1 | LGALS3BP | | ANXA5 | Cthrc1 | LOC11026 | MICU2 | SPON2 | PLPP3 | | OGFR | | MYL6 | IER2 | SLC2A3 | RCAN2 | ADRA2A | | IGSF23 | | RELL1 | Mef2c |
| Filip1l | ENSSSCG0000006199 | | CCDC3 | DYNLT1 | | | | | RGS2 | | GJA1 | CHPF | | | Mlec | PNP | SH3GL1 | MEOX2 | FCGRT | | ZC3H12A | | AKAP1 | PRXL2A | PSME1 | RHOB | CITED2 | | SOD2 | | C11orf96 | Il17d |
| Cx3cl1 | LTBP1 | | FILIP1L | CSDC2 | | | | | KCNMB1 | | HOPX | IFI27L2 | | | Nid1 | DEK | PDPN | PID1 | S100A10 | | TRIM22 | | ZBTB16 | NEURL1B | SERPINF1 | EPAS1 | ACTA2 | | SLC22A18 | | NR2F2 | Ncmap |
| Ecm1 | FAM13C | | SBSPON | SOD3 | | | | | COL18A1 | | P4HA1 | DIO2 | | | C330027C | CYCS | TMEM209 | LRRC17 | PDGFRA | | GBP3 | | MRAP2 | ENSSSCG0 | UGCG | ENSSSCG0 | CFHR1 | | NDFIP1 | | TPM1 | Rnaset2a |
| Tagln2 | GSN | | PLCB4 | MFAP4 | | | | | PFN2 | | COL5A1 | RGS2 | | | H2afv | ANXA1 | RFC4 | CD34 | EFEMP1 | | AC074032.1 | | HACD1 | PLAU | IGHA1 | STUM | LBR | | GALNT1 | | BST2 | Id1 |
| Itgb5 | PEA15 | | MAP3K20 | TGFB1I1 | | | | | GAPDH | | DAB2 | KLF2 | | | Rbm3 | RFC2 | MTRF1L | GALNT15 | NEGR1 | | SDF2 | | ZNF385D | IFITM1 | CYGB | EPS8 | CCDC102B | | FCHSD2 | | CAV1 | Adk |
| Mical2 | ISLR | | LIMS2 | EFHD1 | | | | | CDH2 | | GNG11 | RGS16 | | | Ehd2 | ENO1 | BRCA2 | EPHA3 | TGFBR2 | | SLC2A3 | | SLC22A3 | RALGPS2 | ISYNA1 | SEPTIN11 | C20orf27 | | MAFF | | RASL12 | Ddit4l |
| Acta2 | DST | | IL17B | PPP1R12A | | | | | SLC25A4 | | LOXL3 | COL6A1 | | | Myof | TIMP1 | PSMD1 | JUN | FOS | | MYOM2 | | BTC | GPR4 | TGFBI | EDNRA | NR2F2 | | VDAC3 | | TMEM38B | Golph3 |
| Arid5b | IGFBP7 | | DMPK | PPP1R3C | | | | | SYNPO | | CILP2 | S100A13 | | | Cycs | RAN | SEPTIN9 | OLFML2A | C16orf89 | | ISG20 | | MRVI1 | CD59 | CEBPD | TRA2B | ITGA7 | | EIF4EBP1 | | EHD2 | Fgfr1 |
| Thbs2 | | | SMARCD3 | MGST3 | | | | | FLNA | | FAM180A | ANKH | | | Tipin | H2AC20 | CNPY3 | GGT5 | LSP1 | | AHI1 | | RBPMS | ITGA7 | PCOLCE | FMO2 | FXYD6 | | ACKR3 | | COX17 | Dcn |
| Map1b | | | PCDH7 | CAMTA1 | | | | | RGS5 | | CP | MXRA8 | | | Dtymk | PEG10 | TRIM59 | ABLIM1 | S100A13 | | BIRC3 | | SMOC2 | GPER1 | ATP5MC2 | CD4 | ASAP1 | | APOA1 | | PLCG2 | Cpxm2 |
| C1ra | | | PPP1R3B | CD151 | | | | | SCG2 | | EMP3 | S100A16 | | | Trip12 | IL33 | UBE2T | SAT1 | RARRES2 | | RARB | | RAP1A | MCAM | ETS1 | MCAM | MEOX2 | | GPI | | NDUFA4 | Lgals3 |
| Dclk1 | | | PBXIP1 | GMDS | | | | | PPP1R3C | | SH3BGRL3 | LRRC32 | | | Cct2 | LYAR | C1QL1 | ZMIZ1 | TMEM176A | | OAS2 | | FABP3 | EGFLAM | FAM13C | SMARCA5 | SYNPO2 | | TMBIM6 | | KANK2 | Atp2b1 |
| Lgals1 | | | CLIC4 | GADD45A | | | | | CD9 | | PDGFRL | SERPINE1 | | | Dnajc9 | YWHAQ | P3H4 | FLRT2 | ALDH2 | | GTPBP1 | | GASK1B | LOC10051 | SNRK | LBH | TIMP3 | | WLS | | RGS5 | Lxn |
| Scarf2 | | | CIRBP | TSPAN2 | | | | | PLXDC2 | | SDCBP | TM4SF1 | | | Mad2l1 | HINT1 | CTSL | NR4A1 | LAMA2 | | SLC11A2 | | GSTP1 | RBM24 | CHCHD10 | P3H2 | SLC2A4 | | IFNGR2 | | CRIP1 | Ldlrad3 |
| Adamts2 | | | CFL2 | TCEAL3 | | | | | BEX3 | | USP53 | CCDC80 | | | Snrpe | HNRNPAB | WDR76 | KCTD12 | CYP1B1 | | FGL2 | | MAP1B | SPP1 | IFI16 | GTF2E2 | FHL2 | | CAPG | | PPP2R5A | Gm42792 |
| Scube3 | | | ITPR1 | RBPMS | | | | | HK1 | | | PDGFRL | | | Fbxw8 | PARP1 | CENPW | C1R | HSD11B1 | | NOC4L | | | RASL12 | EPAS1 | PDGFA | MAP3K8 | | BHLHE40 | | GADD45B | Ppib |
| Tm4sf1 | | | ZYX | EIF3L | | | | | ISCU | | | MEG3 | | | Acot7 | VDAC1 | ORC6 | CYP27A1 | OAF | | GALNS | | | SNX10 | EIF4A1 | APOD | SEPTIN7 | | SMPDL3B | | MEF2C | |
| Ccl19 | | | ACTN4 | MXI1 | | | | | | | | SOX4 | | | Tpm4 | EMP3 | NCAPH2 | LAMB1 | LXN | | EBF3 | | | PCP4L1 | B3GNT2 | STOM | SLC25A5 | | ITGA5 | | SLC25A6 | |
| Pmepa1 | | | ATP1A2 | SMIM10 | | | | | | | | ABI3BP | | | Ddx21 | RCN1 | ACTR1A | FGF7 | PTGIS | | CEP162 | | | SOX18 | STOM | CTSC | SLC7A2 | | PRRX2 | | SLMAP | |
| Pde5a | | | SCHIP1 | PKIG | | | | | | | | NBL1 | | | Chpf | ANXA8 | ZDHHC16 | PDGFRL | PTGFR | | LRRC28 | | | MARCKS | PNRC1 | EBF2 | COCH | | ZNF395 | | CLMN | |
| Gxylt2 | | | NDUFB1 | CAVIN3 | | | | | | | | EFEMP1 | | | Brip1os | NUF2 | KDELR3 | TFPI2 | PCOLCE2 | | HLA-E | | | DLX5 | SLC25A5 | SLC12A2 | HNRNPDL | | IL18 | | SH3BGRL | |
| Picalm | | | DTNA | PLS3 | | | | | | | | DDIT4 | | | Hmgb1 | LSM4 | B4GALT2 | CD55 | ZNF385A | | COL1A1 | | | CHN1 | PTK2 | ENPEP | RDH5 | | LGMN | | TPM2 | |
| Kif26b | | | ARHGEF25 | AC092164.1 | | | | | | | | IER3 | | | Calu | DHFR | ADNP | KLF2 | CTSK | | REV3L | | | SYNE2 | GGT5 | PTGER4 | IER2 | | TSPAN15 | | COX5B | |
| Gm26771 | | | FHOD3 | HCFC1R1 | | | | | | | | PLPP1 | | | Tk1 | PHGDH | GLRB | OLFML3 | DAB2 | | PSORS1C1 | | | SEPTIN11 | GUCY1A1 | PDGFRB | CDH6 | | MIF | | AC091182.1 | |
| Csrp1 | | | IL11RA | SNRPN | | | | | | | | TGM2 | | | Crip1 | NDC80 | RACGAP1 | THBD | KLF4 | | ANKRD17 | | | VCAM1 | SLCO3A1 | DUSP5 | C11orf96 | | TBX3 | | PLK2 | |
| Col14a1 | | | CALM1 | PCDH7 | | | | | | | | IGFBP4 | | | Plaur | RANBP1 | ALYREF | PTGS1 | FOSB | | CSTF1 | | | ANGPT1 | SLC12A2 | LRRC10B | CD59 | | HSBP1 | | AOC3 | |
| Myl12a | | | FILIP1 | PFN1 | | | | | | | | HLA-DRB5 | | | Erh | C4BPA | TSPAN15 | FOXO1 | CFI | | FAM126A | | | ENSSSCG0 | ARHGDIB | PDE5A | IFITM2 | | APLN | | FAM241A | |
| Ctps | | | KCNE1 | UQCRB | | | | | | | | FGL2 | | | Mthfd2 | SRSF3 | POMGNT1 | SPRY1 | DUSP1 | | OSBPL8 | | | NRIP2 | ZEB2 | NR4A3 | CCN1 | | SNX9 | | C1QTNF1 | |
| Epas1 | | | ENSSSCG0 | TTLL7 | | | | | | | | CP | | | Txndc5 | CD44 | RECQL | EIF4EBP2 | TSHZ2 | | TCN2 | | | HGF | DDX5 | ENSSSCG0 | TFPI | | TCEA3 | | ACTA2 | |
| Pdlim1 | | | TACC1 | MAP1B | | | | | | | | CCL21 | | | Nudcd2 | PTGES | AGTRAP | TMEM150 | HLA-DRB1 | | GNB4 | | | PARM1 | DLC1 | SIK1 | AGTR1 | | FXYD6 | | WFDC2 | |
| Dst | | | CLDN10 | MAP4 | | | | | | | | CD74 | | | Rtn4 | CLSPN | ZBTB1 | SESN3 | EPHX1 | | BLVRA | | | ANGPT2 | RERGL | PALM2AKA | HLA-C | | CD44 | | AK1 | |
| Hspb6 | | | ARHGEF17 | CCT8 | | | | | | | | TNFRSF11B | | | Cfl1 | CENPS | CTNNAL1 | MPZL1 | NFE2L2 | | C5orf56 | | | ISYNA1 | BST2 | SYTL2 | ANO1 | | FKBP8 | | WDR1 | |
| Mrvi1 | | | ITGA1 | CALM1 | | | | | | | | SOD2 | | | H1fx | JPT1 | TVP23B | GABARAPL | CRISPLD2 | | AC092807.3 | | | PDE1C | CYSTM1 | LGALSL | VASP | | PSAT1 | | ANGPTL4 | |
| Slco2a1 | | | ENSSSCG0 | COX7C | | | | | | | | TIMP3 | | | S100a11 | SPCS3 | COL11A1 | HPGD | COL4A4 | | ECM1 | | | CEBPB | NDRG2 | ITGA1 | EGR1 | | DNASE1L1 | | AC023157.3 | |
| Slc6a6 | | | MAST4 | CRNDE | | | | | | | | CLU | | | Lmf2 | SGO2 | SEZ6L2 | GADD45B | NOVA1 | | NEO1 | | | LOC10012 | HOXB-AS1 | DES | LAPTM4B | | FN1 | | TMEM51 | |
| Ncam1 | | | KANK2 | TMEM159 | | | | | | | | CCL2 | | | Diaph3 | PRDX1 | FURIN | COLEC12 | SAT1 | | PTAR1 | | | ENSSSCG0 | CHN1 | RAMP1 | NABP1 | | ELK3 | | ATP5F1B | |
| Fhl1 | | | ATP2A2 | ARL6IP4 | | | | | | | | PCOLCE2 | | | Cct5 | SNRPD1 | SCAMP3 | CCNL1 | AOX1 | | TRAF7 | | | DLC1 | REM1 | PDE1C | KCNJ8 | | MGAT4A | | APOLD1 | |
| Col5a1 | | | TMEM47 | LIMS2 | | | | | | | | SFRP4 | | | Rc3h2 | BBLN | SLC39A3 | TIMP3 | EGR1 | | BAZ1A | | | ETS1 | MAP3K7CL | CXCL14 | DNAJB5 | | ENSSSCG0000001094 | | TBC1D1 | |
| | | | COL18A1 | NINJ1 | | | | | | | | SERPINF1 | | | Cenpl | KIF20B | EIF5A2 | ENSSSCG0 | SPON2 | | STXBP3 | | | ENSSSCG0 | MARCKSL1 | CSRNP1 | FAM241A | | EIF1 | | NTRK3 | |
| | | | ENSSSCG0 | CALM3 | | | | | | | | COL6A3 | | | Col1a2 | UBE2S | ORMDL3 | PI4K2B | PAMR1 | | TCF4 | | | ETS2 | INPP4B | KCNAB1 | TP53I11 | | FAM107B | | PLN | |
| | | | NUF2 | FHL1 | | | | | | | | GSN | | | Trim47 | RCAN1 | C17orf75 | | STEAP1 | | FAM102B | | | AIG1 | HLA-F | SLC7A2 | KLF11 | | FAM162A | | DBI | |
| | | | ATL2 | SUSD5 | | | | | | | | TFPI2 | | | Por | LTC4S | IRX3 | | GPNMB | | SCARB2 | | | PLEKHA5 | GPRC5C | SLC8A1 | ITGB1 | | PAPSS1 | | KCNA5 | |
| | | | SMOC2 | MRVI1 | | | | | | | | COL15A1 | | | Rabggtb | NUCKS1 | HMMR | | ARL4D | | SLC20A1 | | | ADAMTS1 | MIR4435-2HG | PLCB1 | PPARG | | BHLHE41 | | HSDL2 | |
| | | | BCAM | PFN2 | | | | | | | | MGP | | | Iqgap1 | TMSB10 | PPFIA1 | | JUNB | | DPM1 | | | COL4A1 | MEF2B | PTK2 | CEBPD | | VLDLR | | ENTPD1 | |
| | | | TTLL7 | BEX2 | | | | | | | | C1R | | | Dazap1 | PKM | ITGA6 | | CYP4B1 | | WASHC1 | | | PRPF4B | SH3KBP1 | ATP1B2 | MRGPRF | | COX7C | | MYL6 | |
| | | | TES | PYGB | | | | | | | | SERPINA3 | | | Cmc2 | IGFBP6 | TUBB2B | | SLIT2 | | ZNF232 | | | PIM3 | ARHGEF17 | DNAJA1 | LINC01197 | | ALPK3 | | HNRNPA1 | |
| | | | SUN2 | ADI1 | | | | | | | | S100A10 | | | Hnrnph1 | CDK2AP1 | KRT14 | | LOX | | TMEM140 | | | SHOX2 | EGR1 | AHI1 | RASD1 | | VEGFC | | ETS2 | |
| | | | ENSSSCG0 | BCAM | | | | | | | | IFI27 | | | Fkbp2 | MCUB | ACOT7 | | S1PR2 | | ADAM17 | | | RERGL | CSRP2 | PLN | MT1F | | PLOD2 | | FAM162B | |
| | | | ATF3 | TSR2 | | | | | | | | VCAM1 | | | Fbln5 | CLEC11A | ACTL6A | | SLC43A3 | | GPC4 | | | COX4I2 | JAK1 | PACSIN2 | MYO1B | | LTC4S | | HIST1H4C | |
| | | | MICAL3 | STMN1 | | | | | | | | PCOLCE | | | Pcolce | ARHGDIB | NUSAP1 | | COL6A3 | | CTDSPL2 | | | ENSSSCG0 | PPP1R15A | MEF2C | ARPC5 | | TNFRSF12A | | RRAD | |
| | | | PYGB | TNNC1 | | | | | | | | TNFSF10 | | | Medag | USP1 | MRM3 | | IL33 | | SEZ6 | | | SIK1 | NRP1 | SPARCL1 | SLC25A25 | | ARPC2 | | GPR20 | |
| | | | RAP1A | ZFYVE21 | | | | | | | | FN1 | | | Fgfr1op | COX17 | LRP11 | | CHRDL1 | | IER3 | | | TRPC6 | SYTL2 | EIF4A1 | MT2A | | GJC1 | | PLA2G5 | |
| | | | UNC45A | ASAH1 | | | | | | | | TNC | | | Rrm1 | CALM3 | TLK2 | | CYBA | | SLC44A3 | | | JAG1 | HGF | SPRY2 | EAF1 | | DDIT3 | | ACTN4 | |
| | | | FAM13C | PARK7 | | | | | | | | MARCKS | | | B4galt5 | TUBB3 | MYCBP | | RND3 | | TATDN1 | | | LDHB | MT1A | DDIT3 | CYCS | | ENSSSCG0000005946 | | SMTN | |
| | | | PLIN4 | HSPB8 | | | | | | | | PRSS23 | | | Cby1 | HMGN1 | MCM10 | | CD248 | | MIR22HG | | | EPAS1 | HOPX | REL | CRIM1 | | WNT5B | | LGALSL | |
| | | | LDHB | ELN | | | | | | | | DPT | | | Bzw1 | CDKN2C | GPR137B | | CYB5A | | MGP | | | SLC8A1 | CYBA | SCN1B | CHCHD10 | | S100A10 | | DBNDD2 | |
| | | | ENSSSCG0 | NME4 | | | | | | | | IGFBP6 | | | Fkbp10 | OSTC | CFL1 | | MATN2 | | OS9 | | | CCNL1 | PLEKHH3 | CRIM1 | PPP1R15A | | HECTD2 | | A2M | |
| | | | ENSSSCG0 | HEY2 | | | | | | | | FBLN2 | | | Mfap5 | MRPS25 | ARF5 | | NEAT1 | | LSM5 | | | TNFRSF21 | LGALSL | PELI1 | CYTOR | | ST6GALNAC4 | | KCNMB1 | |
| | | | ENSSSCG0 | MRPL51 | | | | | | | | CTSK | | | Snrpd1 | RACGAP1 | RPP25 | | SAMHD1 | | CCL4 | | | CISD1 | KRT18 | PDE4B | FBXL3 | | TPST1 | | MYOZ1 | |
| | | | YAP1 | ITIH4 | | | | | | | | FBLN1 | | | Eif3a | GMNN | CENPM | | SERPINE2 | | AKR1B1 | | | NCKAP5 | CCNL1 | KLHL15 | GNAI1 | | MFF | | MGLL | |
| | | | HSPB6 | BEX3 | | | | | | | | AGT | | | Angptl2 | SNRPG | PSMD14 | | GAS1 | | PEX7 | | | LDLR | UBA2 | ACTN4 | ARHGAP1 | | TJP2 | | TAGLN | |
| | | | HEY2 | MFGE8 | | | | | | | | C7 | | | Tpd52l2 | YBX1 | POLE3 | | BTG2 | | SYNC | | | LRRC8A | COL4A2 | CDC42EP4 | LRRC17 | | PLP2 | | CAVIN3 | |
| | | | SPC24 | DMPK | | | | | | | | VCAN | | | Aurka | SSR3 | POC1A | | FAIM2 | | MPEG1 | | | TBX2 | IER5L | PRKAR2B | THBS1 | | RHOD | | CHMP4C | |
| | | | ANO6 | MYOZ1 | | | | | | | | LTBP2 | | | 1810055G | FNDC1 | NXPH4 | | CD74 | | C11orf80 | | | LAMA4 | INAFM1 | LOC10015 | LAMA4 | | ANKRD37 | | ARPC2 | |
| | | | VASP | YWHAH | | | | | | | | FAP | | | Actg1 | SPDL1 | CTDSP2 | | PRCP | | IRF9 | | | IGFBP4 | FOXS1 | LRRC32 | SLC38A2 | | TAGLN2 | | ITIH3 | |
| | | | TOB2 | NPM1 | | | | | | | | APOC1 | | | Ddx39 | NSD2 | HRH1 | | LTBP4 | | HELZ2 | | | ARHGAP29 | ACYP1 | TM4SF1 | THY1 | | DUSP1 | | SYPL1 | |

ROCK1
FIBIN
INKA2
DACT3
CCL19
ADCY5
ILK
THRB
EIF5B
BEND5
PARVA
HSPB1
BEX1
LYNX1
MRGPRF
CCNG1
PIAS1
CNBP
ZBTB16
STOM
DNAJB4
ENSSSCG0(
SEPTIN9
CCND2
ALCAM
RASSF2
RNF150
GPR20
CSPG4
GASK1B
THRA
INPP5A
ATOH8
MAP7D1
CRIM1
KIAA2026
ANGPTL7
SH3BGRL
ACTN1
PI15
MFAP4
CDS2
RRAGD
GLDN
ADGRL1
PHLDB2
EIF4A2
DYNLL2
REV1
ENTPD3
KCNA5
VIM
HACD1
SLC2A13
PRKACA
UBC
ZNF503
SGCA
ENSSSCG00000013600
C5ORF24
CCDC107
TEAD1
COX6C
PTK2B
SLMAP
CDC42EP3
MACF1
FRZB
ENSSSCG00000061997
DAAM1
DIAPH2
ENSSSCG00000061867
MBNL1
HSPG2
BMPR1A
CAV2
WDR1
SOST
OAT
SETD3
GRSF1
ENO3
RASL11A
ENSSSCG00000059682
ROCK2
PLCL1
ADAMTSL5
CAPZA2
FBXO30
NDUFA4
AKAP6
DENND2B
SCAPER
FOXO3
ZFAND5
PRKG1
CSDE1
PPP1CB
CAB39
LSM6
MGST2
TGFBR3
RIMS3
LHFPL6
ATP8B1
APP
RRAS
VWA1
MAP4
JAZF1
ATP2B4
PKIA
MAF1
MRPS6
ENSSSCG00000032522
SCOC
NACA
MEF2C
KCNMA1
PJA2
IL1RL1
EIF3A
MAP1B
WWP2
RNF11
IGF1R
CRABP1
STK17A
CRTC3

TCEAL1
PITPNB
PPP1CB
ACTN4
BEX4
SOD1
LGR6
ATOH8
ZBTB16
HINT1
CAPZA2
PCBD1
LDB3
FXYD1
ISCU
SERTAD4-AS1
EIF3H
NACA
RAB4A
HADH
SBDS
DBNDD2
HRCT1
DUSP26
MYRFL
CAP1
PRKAG2
NGRN
COL18A1
SCRG1
WDR1

CYTL1
GAP43
SERPINE2
CCL5
THY1
IGFBP3
BGN
SH3BGRL3
SULF1
TNFAIP6
CTSC
AEBP1
SPRY1
F2R
MT2A
TMSB10
GGT5
CST3
FXYD5
LY6E
HLA-DRB1
C1S
IL32
G0S2
NPC2

C1qtnf6
Eif5a
Snx7
Polr2a
Ssr3
Kcnn4
Loxl3
Wsb2
Ckap2l
Ptms
Ezh2
Carhsp1
Nrarp
Tpm3
Arpc4
Rfc2
Col5a3
Cx3cl1
Mcm5
Vdac1
Dek
Snx3
Bok
Tagln2
4930427A
Lman1
Rad21
Ptn
Surf4
Cars
Samm50
Ttc28
P4hb
Gpc6
Mettl9
Rpn1
Esyt1
Nudt21
Neurl1b
Cinp
Fen1
Hmgn5
Acsl5
Tpi1
Nusap1
Purb
Alyref
Col5a2
Pabpc1
Nfic
Fam102b
Rfc5
Atad2
Manf
Nudt5
Mtch1
Eif4ebp1
Serpinh1
Mgat5
Snrpc
Slc25a24
Ak2
Kazald1
Cnot6
Vim
Adam10
Col15a1
Edem1
Gmnn
Kdelr2
Cpt2
Exosc8
Dbf4
Hint1
Srsf7
Nsmce1
Ppp1ca
Pgam1
Calm2
Ptma
Ccnd1
Mfn2
Cox19
Vps16
Erp29
Ttll12
Sar1a
Naa50
Slc37a3
Tmem167
Vcam1
Psmc3
Lmnb2
Txndc12
Mzt1
Ptgr1
Dnajc10
Pdxdc1
Gins2
Ccdc8
Notch1
Marf1
Hp1bp3
Timm13
Eny2
Tbc1d1
Pfn1
Slmo2
Cep295
Crabp2
Ipo5
Ccna2
Thbs2
Msn
Ap1s1
Atp5o
Phc2
Shcbp1
Dtl
Cenph
Kdelr3
Mterf3
Lsm5
Stac2
Ccdc6
Actr3
Ect2
Stom
Cdk2
Fam129b
Mob1b
Ednra
Polr2g
Rnps1
Setd5
Rnf5
4930579G
Hist1h2ap
Clca3a2
Pcyt1a
Bscl2
Utp3
Rnf10
P4ha3
DHRSX
Necap2
Hsp90b1
Rpp30
Mri1
Dyrk1a
Loxl1
Ncapd2
Mrpl51
Ubl4a
Baiap2
Psma4
Fam96a
Gemin5
Rangrf
Ccdc61
Rac1
Ywhah
Psat1
Rgl2
Prickle3
Scpep1
Eri1

PSIP1
H1-10
LOXL2
SNRPF
POC1A
HSPD1
ARPC5L
DRAP1
PSMD2
TOPBP1
TCEA3
SSR4
LGALS3
ATP5MK
RFC4
BIRC5
HMGB1
HSPA8
MCM2
UHRF1
ENSSSCG0(
LOC10052
HNRNPA2I
LSM5
SKA2
TMEM258
HAT1
KPNA2
LGALS1
ENSSSCG0(
HMGN3
ATOX1
CCDC167
CALM2
UAP1
BUB1B
P4HA3
GINS2
RRM1
MAZ
CALR
MIF
CENPK
TSPO
AVPI1
PSMG4
GARS1
RCN3
UBE2L3
TEX30
NME1
AP3S1
KIFC1
EIF4EBP1
FOSL1
ENSSSCG00000012879
KDELR2
PA2G4
APRT
MRPL20
CNIH1
RPA2
EZH2
RHOC
TSPAN5
CKS1B
CISD2
HOPX
TUBA1A
LRRC59
SPTSSA
YWHAZ
RPA3
COL6A1
UBE2I
ESM1
KCNK12
ARHGDIA
RCC2
CCT3
HSPA5
MKI67
TPI1
NCAM1
IPO7
ENSSSCG00000037274
NOP56
PLIN2
ARF2
ATP5MGL
SNRPE
NDUFA4L2
RALY
AQP1
IMPDH2
CCNYL1
NID1
EIF4E
NUCB2
NCAPD3
ORC6
VIM
BUB3
PCOLCE2
EMP1
DNASE1L1
TAF10
ENSSSCG00000003612
NPM3
SERPINH1
CMC2
ANXA5
PTGES3
PGP
E2F1
BZW1
TSKU
DNAJA1
METTL26
UQCC2
DDIT3
IER3
UBE2M
LNPK
MANF
NDUFV3
PLAUR
CDC20
ANP32E
B3GLCT
LIG1
NCAPG
CALM1
NME2
PRDX4
ALDOA
CHCHD2
TRA2B
GGCT
ACTR3
POSTN
ABL2
CCDC34
NOLC1
RAP1B
RFC3
GINS1
CCNB3
PRKCSH
CCDC85B
IL18
MTHFD1
EDIL3
COL5A1
KPNB1
ADAM12
PPA1
ERH
LSM8
MBD3
RBM3
IDH2
MRPL17
SET
NDUFA11
PRIM1

BX470102.2
FAM131A
GOLT1B
KIAA0408
CDK1
SAC3D1
FUCA2
FAM104B
CBWD2
MFAP5
CALML4
C1orf52
CENPB
ADAMTS2
E2F6
TMEM160
ADRM1
TMEM54
SLC50A1
HPRT1
SNHG30
SUZ12
AKR1B1
ALG5
AC099343.3
TXNDC16
ARPP21
MAD2L2
C1orf131
PARVB
TRIM16L
CREB3L1
FANCI
MRPL12
KIF20B
ARSB
FAM229B
TFDP1
RGS19
GUF1
SCO1
HIPK1
ARHGEF10L
CD276
COX19
CENPF
CD68
FAM83G

CDH19
TMEM176B
FBN1
IL6
SVEP1
MGP
GFPT2
IL6ST
OMD
MT1A
FIBIN
GLUL
SERPINA5
TFPI
FSTL1
SPON1
CCN5
SPRY1
CNTFR
ICAM1
CTSH
HAS2
PLSCR1
COL1A2
CCN1
GALNT15
BNIP3L
ALDH1A1
MARCH3
C6
LIMA1
CLEC2B
ABCA10
SOCS3
LY6E
COL6A2
ITM2B
ABI3BP
LRRN4CL
TIMP3
DPYSL2
CTSF
KLF10
TSPAN4
GLRX
STEAP2
LGALS3
TMEM45A
LARP6
FGF7
COLEC12
CLDN11
CILP
MRC2
ACKR3
ECM1
CCNL1
CSRNP1
CYP27A1
METTL7A
BOC
MCL1
UGDH
SHISA3
RAMP2
SCPEP1
SEMA3B
SPOCK1
CCDC71L
CDO1
NDRG1
CPVL
GASK1B
PLBD1
CDC42EP2
PDIA4
GSTM5
ENG
BZW1
MIR99AHG
GFRA1
ACVRL1
CFB
TGFBR3
CCL13
TNFSF14
INMT
B4GALT1
C1QTNF3
TPST1
SEMA3C
PHGDH
AKAP12
TNFSF10
COL4A3
PSAP
LGALS3BP
SRPX2
TYMP
FGFR1
FSTL3
IFI16
CXCL12
FTL
CDC42EP4
IFITM3
TNFAIP2
RHOBTB3
ADAMTSL2
KLF9
TIMP1
ANGPTL5
RASD1
PLXDC1
C1orf21
TMEM98
TPBG
MEDAG
TRAM1
PCSK5
MID1IP1
ARRDC3
THBS1
ROBO3
GAS7
NFIA
RRBP1
RBMS3
LAPTM4A
DIO3OS
MPZL1
MAN1A1
XBP1
BHLHE40
RCN3
EPB41L2
LAMP2
NR4A2
IFITM2
RBMS1

HAPLN3
GLT8D2
SSH1
LRRC59
ACKR4
MFHAS1
PIGS
CYREN
NEIL1
MXRA7
GLIS2
POLG2
DDX19A

KCNJ8
FARP1
CDC42EP4
NAALAD2
DBNDD2
KCNA5
ECE1
TOP1
CRISPLD2
BAALC
TAGLN2
WTIP
TPPP3
REM1
OAZ2
HEYL
BCAM
REV3L
ITM2B
ENSSSCG0(
ENSSSCG0(
NR4A3
ENSSSCG0(
PLEKHA2
AJUBA
SEPTIN8
TULP3
HNRNPA2I
ARHGAP1
ANKRD24
RFTN1
CSNK1E
ENSSSCG0(
NREP
RCAN2
ARHGDIB
PLCL1
IL13RA1
TMEM150
ENSSSCG0(
MAFB
FILIP1
IER5
COTL1
SOCS1
LOC10051!
CAMK1
ATP2B1
PPP1R12A
CHD4
RND3
EIF1B
COL4A2
SPTBN1
LBH
ARL5B
MSX2
ENSSSCG00000021343
PLEKHH3
UTRN
INPP4B
GLUL
SLC22A18
ARHGEF7
EIF4A1
BZW2
B2M.1
RBPMS
BRD3
TLR5
TM4SF1
ENSSSCG00000052979
ATP5MC2
RAP2A
EFHD2
SMARCA5
MXI1
ITPK1
SEMA6D
BAMBI
NEK4
NFATC4
TBC1D1
MAN1A1
HSP90AB1
ARGLU1
RASA3
PPA1
PELI1
MAT2A
FGF7
ENSSSCG00000037565
ETFB
GGTA1
LRRC32
PSME1
TBX3
ESAM
MDK
BTBD10
F3
ODC1
HSBP1
TGFBR2
DOCK11
ENSSSCG00000058351
ID3
ARHGEF17
ADGRF5
DOCK6
AIF1
ARHGAP42
LAMC1
SH2B3
SPC24
CNN3
ENPEP
PDE3A
MAP3K20
DEPTOR
IFNAR2
CCDC9B
KANK2
TP53I11
MEF2C
TXNIP
TNFRSF1A
TESC
REL
UACA
ENSSSCG00000055977
NDRG2
ENSSSCG00000062137
CTSC
SIX1
PDE1A
JAK1
PNISR
BTG1
PSPC1
MSI2
SEMA5A
OSMR
ENSSSCG00000059774
ENSSSCG00000047783
SEPTIN7
PTK2
CACNA2D1
ENSSSCG00000042114
AHI1
LOC100520329
CD4
VPS36
TTYH2
PLK2
ADCY3
MAP3K7CL
HOXA9
NCK2
NFE2L3
ENSSSCG00000055139
ENSSSCG00000061649
NAB1
ZFHX3
EBF2
CD248
FILIP1L

ATF3
IER2
FARP1
JAG1
EVA1B
RASL11A
PPP1R10
ABTB1
SMARCB1
PRXL2A
SOX4
ADAMTS1
HES1
LINC01197
PMEPA1
PSMB8
THBS1
BTG2
NR4A1
CPE
BCAM
HIGD2A
SERTAD1
MT1M
DEPP1
PDGFRB
COL18A1
MYC
CCL19
HLA-A
FOS
KLHL23
CYTOR
TFPI
UACA
MYO1B
OLFML2A
OAZ2
B2M
IGLC2
COL4A1
EPS8
MT1E
ZFP36L1
LAMA4
SERPINI1
NT5DC2
KCNJ8
LBH
FOSB
IFI27
SPARCL1
KIAA0040
SEPTIN7
MGLL
GADD45B

IGFBP5
HIPK3
SNRK
PLEKHA2
SERTAD1
ETS2
FILIP1
ENSSSCG0(
GNG2
COL4A2
ANGPT4
ESAM
LOC10051!
SLC25A25
SOCS1
PGF
FXYD6
KITLG
CALD1
TIPARP
FRMD4A
NPY1R
NAMPT
OLFML2B
COQ10B
GALR1
ADGRA2
DDX5
ENSSSCG0(
COX17
ADAP2
PHLDA2
ELL2
TNFSF12
KLF6
ART4
CTDSPL
RASGEF1B
ETS1
TBX2
TAGLN
RELL1
FAM13C
LOC10052!
ENSSSCG0(
MAP3K8
C2CD2
TPM1
APOLD1
IFITM1
CCDC3
RERGL
ENSSSCG0(
SEMA5A
ENSSSCG0(
KCNJ8
MAP3K5
RCSD1
BCAM
RALGPS2
CTNNA3
SH2B3
PLEKHH3
INPP4B
NSG1
GPER1
LIFR
ARHGAP2!
CAMK1
PHLDB2
CEBPD
PLAU
WTIP
FAM107B
CNN3
PPP1R12A
KCNS3
AJUBA
SLC38A2
PRXL2A
ENSSSCG0(
GPC3
SEPTIN8
PPA1
CHN1
TRPV2
CAV2
SEPTIN7
EPHX3
EIF1
LGI4
PDE3A
ENSSSCG0(
TMEM200
OSMR
ENSSSCG0(
BAALC
UBB
IRAG1
DAAM2
BTG3
AXL
CRISPLD2
NR2F1
EHD1
ARHGAP2!
PPP1R14A
ADGRF5
CDKN1A
ENSSSCG0(
TSC22D1
ENSSSCG0(
LOC10051!
MOB3B
SYNPO2
ENSSSCG0(
CISH
SPRY4
HNRNPAB
JAG1
DLC1
NEK4
PNRC1
CH25H
MAFF
CACNA2D1
ENSSSCG00000057146
NOSTRIN
AMOTL2
STX11
PEAK1
PAPLN
TNFSF13
PABPC1
PTH1R
SFPQ
ANGPT2
NID2
NAALAD2
PCP4L1
PLCL1
FERMT2
PVR
HNRNPUL1
RNF149
CCDC9B
PPP1R10
P2RY14
MGST1
ENSSSCG00000056370
SGIP1
MARCKS
SELENOW
TNFRSF21
TMEM150C
FAM241A
TPM4
PIM3
PARM1
ITM2B
APBB2
RND1
ENSSSCG00000037274
TRPC6
ENSSSCG00000009399
CAV1
PTP4A3

SLC2A3
GEM
LASP1
GUCY1B1
PAWR
HSPA2
ISYNA1
AOC3
PRXL2A
DLC1
CAV2
SRSF3
MAT2A
ECRG4
NR4A1
PDLIM1
CHRD
SPRY2
MYLK
CNN1
EPHX1
MAP1B
AC092807.3
PELI1
IFITM1
TYMP
SLC1A5
MYADM
STX12
FADS3
HGF
AXL
BST2
MTHFD2
PHLDB2
CDKN1A
MAP7D3
PCDH9
ENPEP
SNHG15
FOS
MEF2C
SH3KBP1
HIF1A
VASN
TUBA1A
UGCG
INSIG2
DDX3X
CRISPLD2
RBPMS
SERPINH1
ITM2C
RGMA
PICALM
EGFLAM
PCSK7
CREB5
OAZ2
HMOX2
RERG
SNAI1
NDRG2
B4GALT3
SFPQ
GPR176
CCNL1
KRT18
IMMP2L
DNAJB1
MT1E
ETS2
CSGALNACT1
ITGA1
KLF6
HSPB6
PRRX1
TNFRSF1A
HSPA5
TMEM173
NECTIN2
ZFP36L1
INTS6
ID3
HEXIM1
DDX21
SOCS1
ARHGDIB
MAFF
NOP16
C1orf198
DDIT3
JUN
STK38L
ANGPTL1
ANKRD10
FAM13C
TOB1
SOX4
TCF21
IMPDH2
FOXO3
PELO
THBD
PIM1
SERPINE1
FERMT2
CBX6
MT1M
FOSB
ERRFI1
CDC37L1
SDC4
VMP1
TMEM47
ZNF703
GADD45B
OLFML2B
C1orf56
FST
IL6
LMCD1
DEPP1

SYPL1
FAM111A
PDGFA
FARP1
ENSSSCG00000003599
WSB1
ARRDC4
PTPN2
CYP7A1
VAPA
AP3S1
FKBP1A
NRP1
TNIP1
SOCS5
AK4
P2RY6
C9H1ORF21

COX6C
INTS6
NDUFB7
EIF4B
MYLIP
ADAP2
NDUFA7
ECH1
MAPRE2
ITIH5
COX7A1
TIMP4
ATP5MC2
B3GNT2
SEPTIN7
JADE1
UQCRB
JAG1
PCDH1
PLEKHO1
RASGRP2
ANXA6
PPP1R12B
COX7B
TGFBI
ZFP36L1
EPS8
CAVIN1
CCND1
GEM
SMOC2
ARID5A
COX6A1
H1F0
SOD3
ARHGAP29
PABPC1
LDB3
C20orf27
CALM2
HSPB8
YBX3
MEOX2
CRTC3
HOXB-AS1
FABP5
CYSTM1
MSRB3
UQCR11
NDUFB10
CBX6
ELOVL5
SYNPO2
NIBAN1
PLAC9
NT5DC2
TUBA1C
ANAPC16
ARHGEF17
NTF3
VAMP2
COX8A
HACD1
SBSPON
CAP2
PARM1
SEPTIN9
CABP1
RERG
ZNF703
NDUFB9
COX4I1
DYNLT1
CABIN1
LPP
ZBTB16
SLC2A4
ATP5F1D
PAWR
ANKRD29
SH3KBP1
GLRX5
ARHGAP10
GSTK1
EBF1
COX14
ZFAND1
EFHD1
DNAJB4
NIPSNAP2
OR51E1
MICAL1
HCFC1R1
CYC1
LINC02381
COL18A1
GPBAR1
RHOB
KCNE4
AGT

| | |
|---|---|
| Map4 | F10 |
| Lsm8 | CCNB1 |
| Psma3 | RNASEH2B |
| Ctsc | HNRNPA3 |
| Trnau1ap | PGLS |
| Lama5 | HSPE1 |
| Lsm2 | TMOD3 |
| Usp5 | EIF2S2 |
| Pop7 | HNRNPR |
| Pcna | PGAM1 |
| Ankrd13a | MAPK7 |
| Trim28 | UQCRC1 |
| Thy1 | SYNCRIP |
| Sept9 | CDK16 |
| Ssna1 | CDT1 |
| Zfp367 | FKBP1A |
| Slc25a51 | NME3 |
| Nup107 | CHAF1A |
| Kif11 | ENSSSCG00000043175 |
| Pdia5 | CHCHD3 |
| Adamtsl3 | ACTG1 |
| Gm10709 | CFL1 |
| Kpnb1 | SMG6 |
| Ilf2 | TCF19 |
| Rnpepl1 | ADAMTS6 |
| Ssr2 | RNFT1 |
| Dclk1 | PHLDA2 |
| Olfml2b | ARPC1B |
| Ipo9 | GOLT1B |
| Ptrh2 | MAGOHB |
| Coq9 | DERA |
| Cdc16 | YWHAE |
| Cers5 | PLOD1 |
| Tubb4b | GSPT1 |
| Vps36 | UPP1 |
| Arf5 | ZMAT5 |
| Orc6 | PSMA4 |
| Mrc2 | EEF1G |
| Far1 | CKAP2L |
| Mapre1 | ENSSSCG00000025499 |
| Cbx3 | SEC61A1 |
| Sept2 | PSMD14 |
| Gna13 | TFDP1 |
| Chac2 | MAD2L2 |
| Anapc15 | VDAC2 |
| Txnl1 | CDK4 |
| Lrfn4 | CCT6A |
| Igf2r | TPD52 |
| Pofut2 | PMF1 |
| Leo1 | C1QC |
| Mmgt1 | ENSSSCG00000033926 |
| Lbr | S100A4 |
| Tcf3 | COX7A2 |
| Swt1 | FUOM |
| Ptgfrn | YKT6 |
| Phb | BTG3 |
| Pfdn6 | CDK1 |
| Adamts4 | SARNP |
| Ppih | SELENOF |
| Pycard | CDA |
| Usp9x | PRTFDC1 |
| Gas2l1 | |

| | |
|---|---|
| PEMT | RASL12 |
| PDE4B | CRTC3 |
| IRAG1 | HIC1 |
| BCL3 | PTGER1 |
| JARID2 | TMEM182 |
| H3-3A | TP53I11 |
| PHLDB2 | TNFAIP3 |
| ENSSSCG00000041449 | TDG |
| RAP1A | ENSSSCG00000055139 |
| PSMB8 | LOC100624590 |
| SUB1 | PTPN1 |
| ABCC4 | PTBP1 |
| CCND1 | HNRNPA2B1 |
| PRMT1 | LMO3 |
| THSD7A | CIRBP |
| PRR13 | ENSSSCG00000058952 |
| ATP8B1 | NFKBIB |
| ENSSSCG00000058085 | ENSSSCG00000046461 |
| ENSSSCG00000062670 | BRD2 |
| CHD3 | ENSSSCG00000024974 |
| FAM76A | CFAP210 |
| ARPC1B | CDS2 |
| DGKZ | ENSSSCG00000025928 |
| DDX5 | ENSSSCG00000052671 |
| FBL | BAZ1A |
| SMARCC1 | NAPA |
| LDB2 | GUCY1A2 |
| SNTB1 | G3BP2 |
| PTPN1 | ARHGAP1 |
| GSTO1.1 | ENSSSCG00000058351 |
| ENSSSCG00000032436 | C1ORF54 |
| ARRDC2 | LAMB2 |
| ENSSSCG00000060262 | ENSSSCG00000060262 |
| CREM | GLDN |
| UBB | BMP2 |
| ENSSSCG00000061837 | LYAR |
| FXYD6 | ADCY3 |
| UGCG | MYOM1 |
| ENSSSCG00000056123 | RACK1 |
| MYLIP | SNTB1 |
| CPM | WNT6 |
| GALNT16 | ARHGEF17 |
| TMEM131L | REM1 |
| KLHL5 | |
| ARHGAP6 | |
| PRDX2 | |
| CEP85L | |
| ENSSSCG00000055827 | |
| S1PR5 | |
| H1-3 | |
| PPP2R2A | |
| AGTR1 | |
| PDCD4 | |
| B2M | |
| ENSSSCG00000010440 | |
| TRPV2 | |
| AIMP1 | |
| CDH6 | |
| ENSSSCG00000060237 | |
| MED13 | |
| SOX13 | |
| MOB3B | |
| EHD2 | |
| TIPARP | |
| ENSSSCG00000055499 | |
| ROCK1 | |
| ADAMTS9 | |
| ACYP1 | |
| CTNND1 | |
| ENSSSCG00000057146 | |
| RBPMS2 | |
| PNRC1 | |
| SLC12A2 | |
| AFAP1L2 | |
| MARCHF8 | |
| PABPC1 | |
| PAIP2 | |
| ENSSSCG00000048556 | |
| UBE2E3 | |
| C5ORF24 | |
| STAT3 | |
| MAPRE2 | |
| DYNC1LI1 | |
| MEOX2 | |
| BCAS3 | |
| SPRY1 | |
| LIFR | |
| PNPLA2 | |
| TAF1D | |
| CACNA1H | |
| CNN2 | |
| HNRNPM | |
| NR1H3 | |
| ENSSSCG00000038912 | |

**Table S8. Conserved markers for mesenchymal cell clusters across species. Corresponds to Figure 8.**
This table lists conserved gene markers for mesenchymal cell clusters identified in mouse, pig, and human elastic and coronary artery atherosclerotic plaques.
The markers were derived by overlapping the gene lists from Table S7 across species, highlighting those consistently expressed across all species

| Contractile | | | Intermediate | | | Terminal | | | Fibroblasts | IFN_SMC | Proliferating | Pericytes | | |
|---|---|---|---|---|---|---|---|---|---|---|---|---|---|---|
| Shared Elastic | Shared Coronary | Shared between Vasc Beds | Shared Elastic | Shared Coronary | Shared between Vasc Beds | Shared Elastic | Shared Coronary | Shared between Vasc Beds | Shared Coronary | Shared Coronary | Shared Elastic | Shared Elastic | Shared Coronary | Shared between Vasc Beds |
| FLNA | TNNC1 | CNN1 | | CCN3 | | S100A10 | NDUFA4L2 | NDUFA4L2 | APOD | ISG15 | STMN1 | CCL19 | CCL2 | CCL19 |
| MYH10 | MYH11 | ACTA2 | | MYH10 | | COL1A2 | COMP | IGFBP6 | CFD | CXCL10 | BIRC5 | GJA4 | CCL19 | GJA4 |
| COL8A1 | PPP1R14A | FLNA | | EFEMP1 | | COL6A3 | S100A4 | S100A10 | MFAP5 | MX1 | SMC2 | ATF3 | GJA4 | ATF3 |
| NEXN | CNN1 | NEXN | | FRZB | | PCOLCE | IGFBP6 | COL1A1 | PI16 | IFIT1 | TOP2A | BTG2 | RGS16 | FABP4 |
| CNN1 | ACTG2 | | | VCAN | | COL6A1 | POSTN | FN1 | C3 | MX2 | CENPF | HIGD1B | ADAMTS4 | RGS16 |
| ACTA2 | MYL9 | | | MFAP4 | | COL1A1 | COL8A1 | ECM1 | FMO2 | RSAD2 | RACGAP1 | FABP4 | ZFP36 | TINAGL1 |
| | CSRP1 | | | THBS2 | | MMP2 | SERPINE2 | LUM | C7 | EIF2AK2 | COL5A1 | RGS16 | FABP4 | ADAMTS1 |
| | SMTN | | | COL18A1 | | NDUFA4L2 | COL3A1 | COL1A2 | MYOC | OAS2 | PRC1 | GJC1 | FOSB | GUCY1B1 |
| | TPM2 | | | ITGA8 | | NPC2 | S100A10 | EMP3 | GPX3 | EPSTI1 | LGALS1 | PGF | JUNB | ZFP36 |
| | ACTA2 | | | TNFRSF11B | | SERPINF1 | PRELP | | LAMA2 | STAT1 | TUBA1B | TINAGL1 | SOCS3 | SOCS3 |
| | DES | | | IGFBP7 | | NBL1 | SH3BGRL3 | | ABCA9 | IFI6 | HMMR | RGS5 | FOS | FAM162B |
| | LMOD1 | | | MYH9 | | FN1 | FMOD | | CRISPLD2 | UBE2L6 | ANLN | CYGB | TINAGL1 | MYO1B |
| | FLNA | | | TGM2 | | CTSK | COL1A1 | | DCN | PSMB9 | CKAP2 | ADAMTS1 | RASD1 | FOS |
| | RGS5 | | | HSPB1 | | LUM | CD9 | | ABCA6 | IFIH1 | CENPW | FRMD3 | NOTCH3 | NR4A1 |
| | DSTN | | | COL4A2 | | IGFBP6 | FN1 | | HSD11B1 | IFI44L | CDK1 | GUCY1B1 | MYC | NR2F2 |
| | MYLK | | | GAS6 | | TIMP3 | SPARC | | GPC3 | SAMD9 | S100A10 | ZFP36 | FAM162B | JUN |
| | ITGA8 | | | | | EMP3 | ECM1 | | FBLN1 | IRF7 | KIF20B | SOCS3 | GADD45B | FOSB |
| | SYNPO2 | | | | | DCN | LUM | | SOCS3 | CMPK2 | SMC4 | FAM162B | EDNRB | EPS8 |
| | ITIH4 | | | | | EMP1 | HTRA1 | | CXCL12 | XAF1 | FNDC1 | SEPTIN4 | ATF3 | SLC2A3 |
| | FBXO32 | | | | | ECM1 | COL1A2 | | HTRA3 | IFI44 | TYMS | MYO1B | CYCS | NOTCH3 |
| | ADIRF | | | | | | FAP | | PCOLCE2 | ZC3HAV1 | PLP2 | PTP4A3 | IL6 | APOLD1 |
| | SORBS2 | | | | | | CLU | | FOS | SOD2 | TK1 | FOS | NR4A1 | MYC |
| | TAGLN | | | | | | EMP3 | | LTBP4 | GBP2 | CFL1 | NR4A1 | NR2F2 | EDNRB |
| | PDLIM3 | | | | | | ANXA2 | | DPT | PSME2 | DEK | NR2F2 | PPP1R15A | SLC7A2 |
| | PPP1R12B | | | | | | CTHRC1 | | CHRDL1 | PARP12 | TAGLN2 | JUN | ADAMTS1 | JUNB |
| | CALD1 | | | | | | KDELR3 | | FBLN2 | IFIT3 | NUSAP1 | FOSB | JUN | STOM |
| | MYL6 | | | | | | CAPG | | C1S | USP18 | ALYREF | EPS8 | ITGA7 | PPP1R15A |
| | KCNMB1 | | | | | | TSPO | | F3 | | GINS2 | SLC2A3 | EGR1 | ZFP36L1 |
| | PLN | | | | | | TMSB10 | | RNASE4 | | CENPH | NRP1 | GUCY1B1 | EGR1 |
| | RASSF3 | | | | | | PCOLCE2 | | ZFP36 | | ORC6 | GUCY1A1 | ZFP36L1 | IER2 |
| | CAMK2G | | | | | | | | CYBRD1 | | | NOTCH3 | ID4 | PRXL2A |
| | MCAM | | | | | | | | OMD | | | KLHL23 | ISYNA1 | IFITM1 |
| | PDLIM7 | | | | | | | | ADD3 | | | APOLD1 | SLC2A3 | ANGPT2 |
| | SLC22A3 | | | | | | | | PDGFRA | | | MYC | MYO1B | ISYNA1 |
| | TNS1 | | | | | | | | JUNB | | | PDGFRB | CCNL1 | DLC1 |
| | NIBAN1 | | | | | | | | SVEP1 | | | ADAP2 | TCF21 | CCNL1 |
| | EFHD1 | | | | | | | | MATN2 | | | CAMK2N1 | IER2 | KCNJ8 |
| | FBLIM1 | | | | | | | | TNXB | | | EDNRB | FHL5 | CRISPLD2 |
| | RBPMS | | | | | | | | SERPING1 | | | DEPP1 | EPS8 | REM1 |
| | PPP1R12A | | | | | | | | PLPP3 | | | SLC7A2 | STOM | INPP4B |
| | RASL12 | | | | | | | | ALDH1A1 | | | PHLDA2 | CTSC | EIF4A1 |
| | TPM1 | | | | | | | | GFPT2 | | | RCSD1 | ENPEP | ADGRF5 |
| | NEXN | | | | | | | | PTN | | | JUNB | SYTL2 | ENPEP |
| | SVIL | | | | | | | | BTG2 | | | ABCC9 | ITGA1 | SEPTIN7 |
| | LPP | | | | | | | | CEBPD | | | STOM | SLC7A2 | PTK2 |
| | RAMP1 | | | | | | | | TGFBR3 | | | PPP1R15A | PTK2 | DDX5 |
| | FHL1 | | | | | | | | EGR1 | | | ZFP36L1 | MEF2C | FXYD6 |
| | SPINT2 | | | | | | | | FBLN5 | | | EDNRA | EIF4A1 | |
| | TGFB1I1 | | | | | | | | TGFBR2 | | | KCNE4 | SPRY2 | |
| | CAP1 | | | | | | | | MEDAG | | | RASL11A | DDIT3 | |
| | RCAN2 | | | | | | | | DHRS3 | | | EGR1 | CRIM1 | |
| | SLC25A4 | | | | | | | | SPON2 | | | OLFML2A | PELI1 | |
| | SH3BGR | | | | | | | | CD34 | | | IER2 | ACTN4 | |
| | HSPB7 | | | | | | | | GALNT15 | | | PRXL2A | TM4SF1 | |
| | PALLD | | | | | | | | SAT1 | | | NEURL1B | IGFBP5 | |
| | MXI1 | | | | | | | | NR4A1 | | | IFITM1 | ETS2 | |
| | MSRB3 | | | | | | | | C1R | | | CHN1 | ESAM | |
| | CCDC3 | | | | | | | | CYP27A1 | | | HGF | SLC25A25 | |
| | SBSPON | | | | | | | | FGF7 | | | ANGPT2 | SOCS1 | |
| | LIMS2 | | | | | | | | PDGFRL | | | ISYNA1 | FXYD6 | |
| | DMPK | | | | | | | | OLFML3 | | | DLC1 | OLFML2B | |
| | PCDH7 | | | | | | | | SPRY1 | | | ETS1 | DDX5 | |
| | CFL2 | | | | | | | | MPZL1 | | | COL4A1 | KLF6 | |
| | ACTN4 | | | | | | | | COLEC12 | | | RERGL | TAGLN | |
| | CALM1 | | | | | | | | CCNL1 | | | COX4I2 | FAM13C | |
| | COL18A1 | | | | | | | | TIMP3 | | | JAG1 | MAP3K8 | |
| | BCAM | | | | | | | | | | | EPAS1 | TPM1 | |
| | TTLL7 | | | | | | | | | | | CCNL1 | APOLD1 | |
| | PYGB | | | | | | | | | | | NCKAP5 | IFITM1 | |

| | | |
|---|---|---|
| HEY2 | TBX2 | KCNJ8 |
| ILK | LAMA4 | INPP4B |
| HSPB1 | IGFBP4 | PHLDB2 |
| ZBTB16 | ARHGAP29 | CEBPD |
| ATOH8 | KCNJ8 | CNN3 |
| SH3BGRL | FARP1 | SLC38A2 |
| ACTN1 | CRISPLD2 | PRXL2A |
| MFAP4 | TPPP3 | CAV2 |
| KCNA5 | REM1 | SEPTIN7 |
| VIM | OAZ2 | LGI4 |
| HACD1 | BCAM | AXL |
| CCDC107 | ARHGDIB | CRISPLD2 |
| SLMAP | COL4A2 | ADGRF5 |
| FRZB | LBH | CDKN1A |
| WDR1 | PLEKHH3 | SYNPO2 |
| CAPZA2 | INPP4B | DLC1 |
| NDUFA4 | EIF4A1 | MAFF |
| PPP1CB | ATP5MC2 | SFPQ |
| MAP4 | PSME1 | ANGPT2 |
| MRPS6 | ARHGEF17 | FERMT2 |
| NACA | ADGRF5 | FAM241A |
| MAP1B | ARHGAP42 | TPM4 |
| | ENPEP | PARM1 |
| | TESC | CAV1 |
| | UACA | PTGER1 |
| | NDRG2 | TP53I11 |
| | JAK1 | ARHGAP1 |
| | BTG1 | BMP2 |
| | SEPTIN7 | REM1 |
| | PTK2 | |
| | MAP3K7CL | |
| | CD248 | |
| | PSMB8 | |
| | DDX5 | |
| | FXYD6 | |
| | UGCG | |
| | CPM | |
| | B2M | |
| | CDH6 | |
| | ACYP1 | |
| | PNRC1 | |
| | SLC12A2 | |

**Table S9. Published nomenclature for mesenchymal cells subtypes in scRNA-seq studies of human and mouse atherosclerosis. (Corresponds to Figure 8)**
The table lists cell subtype names and the markers used to identify them from the indicated references

| SMC name | Species | Vascular bed | Markers | Ref |
|---|---|---|---|---|
| Contractile | Human | Carotid/Coronary | TPM2, MYH11, MYL9, ACTA2, CNN1, TAGLN | Mosquera et al, 2023 (ref. 11) |
| Intermediate | Human | Carotid/Coronary | BGN, IGFBP2, TNFRSF11B, CTGF, VCAM1 | Mosquera et al, 2023 (ref. 11) |
| Fibromyocytes | Human | Carotid/Coronary | COL6A2, COL4A2, LTBP1, VCAN, FN1 | Mosquera et al, 2023 (ref. 11) |
| Fibrochondrocytes | Human | Carotid/Coronary | CRTAC1, CLU, COL3A1, COMP | Mosquera et al, 2023 (ref. 11) |
| Contractile | Mouse | Ascending aorta/BCT | Ppp1r14a, Acta2, Myl9, Tagln, Cald1, Myh11 | Sharma et al, Pan et al, Carramolino et al, Wirka et al (ref. 12, 13, 22, 23) |
| Intermediate/Pioneer/SEM | Mouse | Ascending aorta/BCT | Ly6a, Vcam1, Lgals3 | Sharma et al, Pan et al, Carramolino et al, Wirka et al (ref. 12, 13, 22, 23) |
| Macrophage-like | Mouse | Ascending aorta/BCT | Lgals3, Cd68, Mac3, Cd36, CD11b | Pan et al 2020, Chen et al, 2023 (ref 13, 66) |
| Fibromyocytes | Mouse | Ascending aorta/BCT | Vcam1 , Tnfrsf11b, Lum, Fbln5 | Sharma et al, Pan et al, Carramolino et al, Wirka et al (ref. 12, 13, 22, 23) |
| Chondromyocytes | Mouse | Ascending aorta/BCT | Col2a1, Acan, Spp1, Snorc | Sharma et al, Pan et al, Carramolino et al, Wirka et al (ref. 12, 13, 22, 23) |
| Fibromyocytes | Human | Coronary | IGFBP2, BGN, FN1, GAS6, MFGE8, LTBP1 | Wirka et al (ref. 12) |
